# Force-dependent Reorganization and Mechanostability of the Izumo1:Juno Complex Involved in Human Fertilization

**DOI:** 10.1101/2024.10.17.618658

**Authors:** Sean Boult, Paulina Pacak, Byeongseon Yang, Haipei Liu, Viola Vogel, Michael A. Nash

## Abstract

Izumo1:Juno-mediated adhesion between sperm and egg cells is essential for mammalian sexual reproduction. However, conventional biophysical and structural approaches have provided only limited functional insights. Using atomic force microscopy-based single-molecule force spectroscopy and all-atom steered molecular dynamic simulations, we explored the role of mechanical forces in regulating the human Izumo1:Juno complex. Our findings reveal a multi-state catch bond capable of withstanding forces up to 600 pN- mechanostability rarely observed among eukaryotic protein complexes. We further found that this mechanostability was impaired in the infertility-associated mutant, JunoH177Q. Detailed steered molecular dynamics simulations show how force-dependent reorganization of the Izumo1:Juno complex engages new binding conformations to achieve this state of high mechanostability. Overall, this study significantly enhances our understanding of the mechanical underpinnings that regulate human fertilization.

## Introduction

In mammals, gamete cell fusion is contingent on the remarkably specific cellular adhesion mechanisms that exist between sperm and egg cells (*1–3*). Where several fusion-indispensable genes have now been identified (*3–6*), sperm-egg adhesion is still primarily defined by the interaction between the sperm transmembrane protein Izumo1 (*1*) and the egg peripheral membrane protein Juno (also known as Folr4 or Izumo1r) (*2*) (Fig. 1A). Only for Izumo1:Juno have both *in vitro* and *in vivo* studies provided clear and convincing evidence for a role in maintaining the close membrane-to-membrane contact necessary for gamete cell fusion (*1*, *2*, *7*, *8*). Yet, to date, only limited insights into the molecular mechanisms underpinning Izumo1:Juno-mediated adhesion have been obtained from conventional structural and biophysical approaches (*2*, *9–11*). Further impeding our understanding of human infertility has been the absence of any biophysical characterization of infertility-associated Izumo1 or Juno variants.

**Figure 1:**
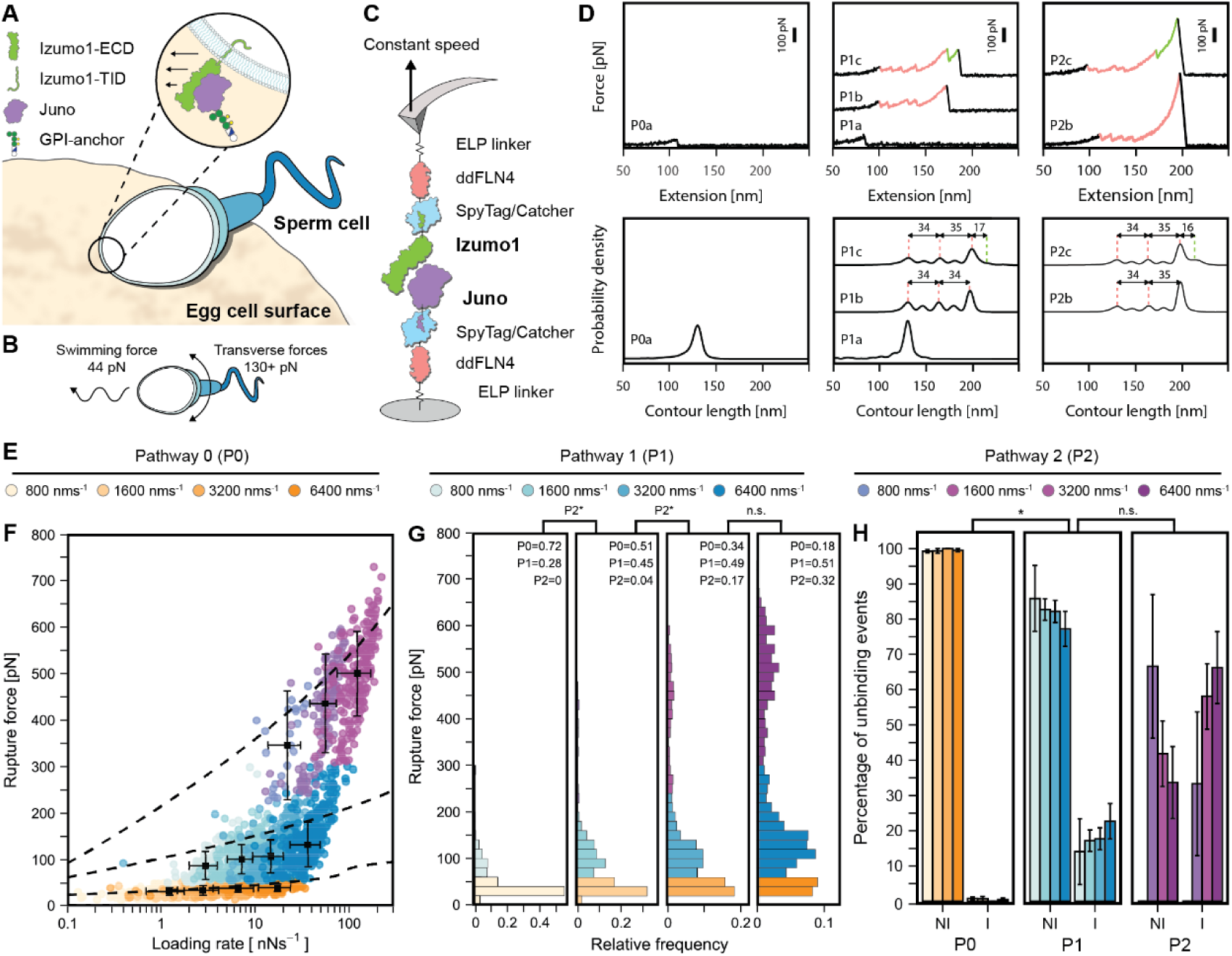
Constant speed AFM-SMFS of Izumo1:Juno reveals catch bonding and multiple unbinding pathways. (A) Sperm:egg adhesion through the Izumo1:Juno complex. (B) Forward swimming and transverse motive forces for healthy sperm. (C) Schematic showing protein constructs and surface immobilization for constant speed AFM-SMFS. (D) Above: representative P0, P1, and P2 force vs. extension curves and their subtypes (a, b, and c). Bottom: combined force vs. contour length histograms pooled from four independent experiments for P0a (n = 116), P1a (n = 22), P1b (n = 243), P1c (n = 62), P2b (n = 66), and P2c (n = 142) at 6400 nm·s^-1^. (E) Legend for P0, P1, and P2 unbinding events at each constant pulling speed. (F) Izumo1:Juno rupture force vs. loading rates (median ± MAD). Dashed lines show the DHS model fits (Eq. (6), Eq. (7), table S2). (G) Izumo1:Juno rupture force histograms (bins = 20 pN). (H) Direct Izumo1:Juno unbinding (NI, force vs. extension curves: P0a, P1a, P1b, and P2b) compared to Izumo1:Juno unbinding through intermediate folded states (I, force vs. extension curves: P1c and P2c) (mean ± SEM). Data in F, G, and H represent four pooled independent experiments (n = 433-693) with significance (*p < 0.05) determined using Kurskal-Wallis tests followed by pairwise Wilcoxon rank-sum tests.

From a mechanobiology perspective, it is evident that Izumo1:Juno must be sufficiently mechanostable to resist the powerful undulating forces generated by the sperm flagellum (Fig. 1B). Regularly exceeding 100 pN in primates (*12*), these forces are sufficient to rupture the vast majority of eukaryotic protein:protein interactions studied to date (*13*). The selective pressure for highly mechanostable protein interactions is manifested in several interesting ways (*14–18*). Of particular interest in the field of cellular adhesion is the catch bond (*19–21*). Canonically, protein interactions are observed as slip bonds; when force is applied to a slip bond, the bond lifetime decreases exponentially. Conversely, in protein interactions exhibiting catch bond behavior, the application of force increases the bond lifetime (*22*). Previous examples of molecular mechanisms that govern catch bond behavior include force-induced allosteric changes (*23*, *24*), alterations to hydrogen bonding networks and contact surface areas (*25–27*), changes to force propagation pathways (*28*), the presence of multiple binding modes (*29*), and the availability of alternate unbinding pathways (*23*, *30*, *31*). It is through these subtle molecular mechanisms that catch bonds facilitate the emergence of unique force-dependent behaviors at the supramolecular and cellular levels (*32*). In considering scenarios for gamete cell fusion, it becomes clear that catch bonding might not only stabilize sperm-egg adhesion, but in doing so directly influence the selection of motile sperm (*33*).

Here, by combining atomic force microscopy-based single-molecule force spectroscopy (AFM-SMFS) with kinetic Monte Carlo modeling and all-atom steered molecular dynamics (SMD) simulations, we show that the unbinding behavior of Izumo1:Juno is indeed consistent with that of a remarkably stable multi-state catch bond (*23*, *31*, *34*). In doing so we provide molecular insights into a previously unexplained infertility-associated mutation (JunoH177Q) (*35*), and identify a novel, non-equilibrium, binding interface for consideration in drug development.

## Results and Discussion

### Constant speed AFM-SMFS of the Izumo1:Juno complex

To study how the human Izumo1:Juno complex responds to force, we first expressed the glycosylated ectodomains (ECD) of Izumo1 and Juno using Sf9 cells. The presence of C-terminal SpyTags were used to confirm the successful expression and purification of Izumo1 and Juno. These C-terminal SpyTags were subsequently used in the covalent attachment of the polyproteins required for AFM-SMFS, termed AFM-handles. For the AFM-SMFS experiment described in Fig. 1, the AFM-handles were expressed in *E. coli* and consisted of SpyCatcher, a ddFLN4 fingerprint domain, an elastin-like polypeptide (ELP) linker, and a ybbR tag for site-specific surface immobilization. A flow cytometry-based bead-binding assay was first used to confirm the equilibrium binding affinity (*K_D_*) and validate recombinant protein quality for all purified Izumo1 and Juno variants in the presence of the AFM-handle and following our surface immobilization procedure (fig. S1). For our recombinantly expressed wild-type Izumo1:Juno complex, we measured a *K_D_* of 45 ± 5 nM (± SE), which was in excellent agreement with previously published surface plasmon resonance data (*9*).

We next measured the loading-rate dependent rupture force distributions of Izumo1:Juno using constant-speed AFM-SMFS. Here, cantilevers functionalized with Izumo1 were briefly brought into contact with cover glasses functionalized with Juno (Fig. 1C). Following this brief contact (0.2 s), the cantilever was retracted at a constant speed (800, 1600, 3200, or 6400 nm·s^-1^). Both Izumo1 and Juno were site-specifically and covalently linked to their respective surfaces through their C-termini, resulting in a pulling geometry that precisely mimicked their natively anchored orientation at the cell surface (*i.e*. attachment through the transmembrane and intracellular domains (TID) of Izumo1 and the glycosylphosphatidylinositol (GPI) anchor of Juno). When Izumo1:Juno complexes formed, retraction of the cantilever resulted in stretching and a build-up of tension that was recorded in force *vs.* extension curves. Sawtooth-like peaks in the force *vs.* extension curves indicated the sudden release of tension through unfolding of the ddFLN4 fingerprint domains, conformational changes within Izumo1 or Juno, or the eventual rupture of the Izumo1:Juno complex (Fig. 1D: top panels). By repeating these measurements tens of thousands of times, we could filter for single Izumo1:Juno complex formation and rupture events. Further explanation of the force *vs.* extension curve selection procedure and representative examples (fig. S2) are provided in the Supplementary Materials and Supplementary Text.

### Wild type Izumo1:Juno unbinds through three distinct pathways

At constant pulling speeds, Izumo1:Juno predominantly unbinds along three distinct pathways, observed in Figs. 1F and 1G as a trimodal rupture force distribution. We refer to these characteristic unbinding behaviors as pathway 0 (P0), pathway 1 (P1), or pathway 2 (P2). P1 and P2, in particular, are also associated with intermediate unfolding events within the Izumo1:Juno complex (see below). The discrete pathway switching of Izumo1:Juno is a phenomenon separate from the classical loading-rate dependency of rupture forces described by the Bell-Evans (BE) model and its extensions (*36–39*). Nonetheless, within each unbinding pathway we still observe rupture forces that increase with loading rate. P0, P1, and P2 exhibit median rupture forces of 32.3 to 39.9 pN for loading rates of 1.2 to 16.9 nN·s^-1^, 87.7 to 132.5 pN for loading rates of 3.0 to 36.5 nN·s^-1^, and 346.2 to 499.8 pN for loading rates of 22.1 to 121.5 nN·s^-1^, respectively (Fig. 1F). Across the trimodal rupture force distribution, the proportion of Izumo1:Juno complexes unbinding through either P0, P1, or P2 displayed a non-linear dependence on pulling speed. Separately fitting each of the four independent experiments using a gaussian mixed model (GMM), we observed that as pulling speed increased from 800 to 6400 nm·s^-1^, the pathway dominance of P0 in Izumo1:Juno decreased from 68.7 ± 8.3 to 23.1 ± 8.8% (± SEM). In parallel, unbinding through P1 increased from 31.3 ± 8.4 to 47.9 ± 5.8% (± SEM), and unbinding though P2 increased from 0 ± 0 to 28.9 ± 3.8% (± SEM). Based on these experimental P0, P1, and P2 distributions, we propose that P0 is representative of an initial near equilibrium unbinding configuration, whilst P1 and P2 result from sequential force-induced changes to the Izumo1:Juno complex.

### Higher order unbinding pathways desensitize Izumo1:Juno to mechanical stress

We next used the Dudko–Hummer–Szabo (DHS) and BE models to describe the underlying energy landscapes governing P0, P1, and P2 (*36–39*). For both DHS and BE fitting of P0, P1, and P2 rupture events, we observed a several fold decrease in Δ*x* when comparing P1 to P0 and then again when comparing P2 to P1 (table S1). These changes to Δ*x* indicate an increasing rigidity to the Izumo1:Juno interaction along P1 and P2, implying greater cooperativity during unbinding. In this way, successive pathway-dependent changes in Δ*x* are consistent with loading-rate dependent catch bond behavior (*34*).

During Izumo1:Juno unbinding, P1 and P2 exist only as force-activated pathways. In this scenario, extrapolation of the DHS and BE off-rate to loading rates of zero (*k_0_*) is not particularly meaningful. Instead, we compared the DHS and BE-derived force-dependent off-rates (*k(F)*) for P0, P1, and P2. We observed that despite a lower *k_0_*, the P0 *k(F)* overtakes the P1 *k(F)* at forces of ∼5–20 pN (fig. S3). Similarly, differences in the DHS and BE-derived *k(F)* for P1 and P2 become apparent above ∼20 pN, with P2 showing greater stability at higher forces (fig. S3). This theoretical analysis supports the interpretation that P0 represents a near-equilibrium conformation and that P1 and P2 are representative of conformational changes that enhance the stability of the Izumo1:Juno complex over biologically relevant force regimes.

### Higher order Izumo1:Juno unbinding pathways are linked to large conformational changes

Across the constant speed AFM-SMFS Izumo1:Juno datasets, we also identified the presence of several intermediate Izumo1:Juno folded states. To further explore these observations and their relationship to P0, P1, and P2, we subdivided the P0, P1, and P2 force *vs.* extension curves based on the presence of these intermediates (Fig. 1D). For each constant pulling speed, the resulting proportions of Izumo1:Juno complexes unbinding directly (no intermediate (NI). Curve subtypes: P0a, P1a, P1b, and P2b) and those unbinding through an intermediate state (intermediate (I). Curve subtypes: P1c and P2c) are shown in Fig. 1H. In P1, we observed transition through intermediate folded states increasing from 14.1 ± 9.4 to 22 ± 5% (± SEM) as the pulling speed increased from 800 to 6400 nm·s^-1^. Whilst in P2, we observe transition through intermediate folded states increasing from 33.3 ± 20.4 to 66.3 ± 10.2% (± SEM) over the same range of pulling speeds. In P1 and P2, these intermediate folded states appeared at median loading rates between 5.1 to 33.9 nN·s^-1^ and 6.4 to 45.6 nN·s^-1^, respectively. Whereas for P0, despite reaching median loading rates of 16.9 nN·s^-1^, we typically observed direct unbinding without the appearance of any intermediate folded states (see Supplementary Text). At pulling speeds of 3200 nm·s^-1^, the pathway-induced intermediate unfolding events have median contour length intervals of 14.8 ± 6.7 and 15.1 ± 8.1 nm (± MAD) for P1 and P2, respectively. At 6400 nm·s^-1^, the median contour length intervals increased to 15.1 ± 6.9 and 19.2 ± 9.1 (± MAD) for P1 and P2, respectively. Due to the presence of multiple disulfide bonds, it was not possible to assign these intermediates to discrete structures within the domains of Izumo1 or Juno. For reference, the distribution of the contour length increments associated with the Izumo1:Juno intermediates at all pulling speeds and a topology diagram of the disulfide networks of Izumo1 and Juno are provided in fig. S4.

### Force-stabilization and dissipation mechanism(s) observed by all-atom SMD

To gain high-resolution structural insights, all-atom explicit solution SMD simulations (*40*, *41*) of the Izumo1:Juno complex were performed at constant pulling velocities of 0.001 Å·ps^-1^. To best mimic the experiments, the C-terminal residue of Juno (S228) was fixed and the C-terminal residue of Izumo1 acted as the pulling point (P254) (Fig. 2A). Simulations were then run in 50 independent replicas for 90 ns. For a summary of all simulations performed see table S2. The resulting SMD simulations describing force-dependent Izumo1:Juno unbinding were then analyzed with several computational tools (see below). Approximately 20% of trajectories were classified as two-state unbinding (Fig. 2B, fig. S5, fig. S6; Supplementary Movie 1), while ∼80% were classified as three-state unbinding (Fig. 2, C and D; fig. S7; fig. S8; Supplementary Movie 2). Each state is defined by the interaction of three different clusters of amino acids on Izumo1:Juno (table S3). In State 1, (*i.e.* the equilibrium structure), Clusters 1 and 2 are bound; in State 2, only Cluster 2 is bound; in State 3, only Cluster 3 is bound. The time dependent change for contacts between Izumo1 and Juno is shown in fig. S9.

**Figure 2:**
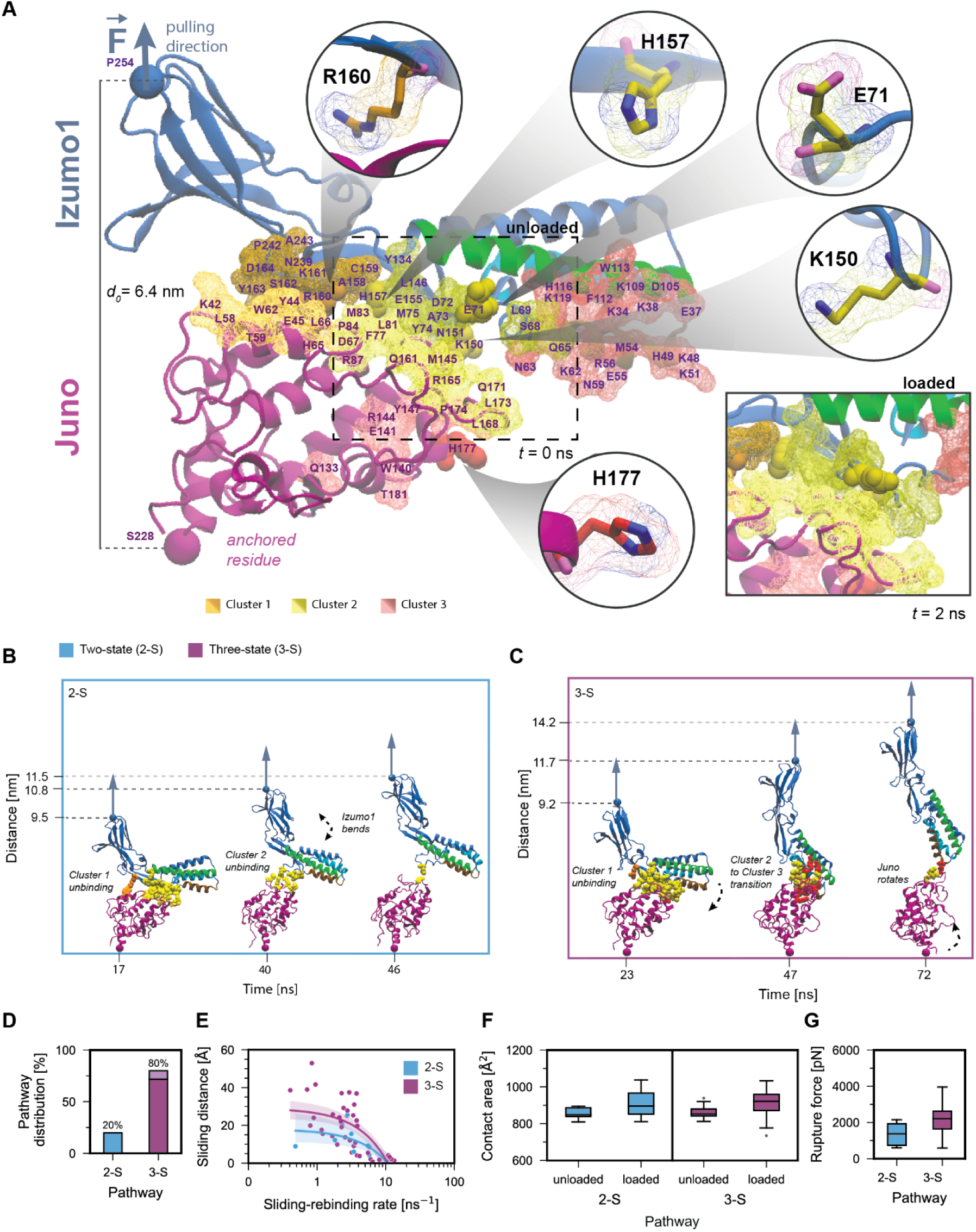
SMD simulations of Izumo1:Juno unbinding. (A) Known Izumo1:Juno binding interface shown in orange (Cluster 1) and yellow (Cluster 2), with the new binding interface shown in red (Cluster 3) (table S3). Circles highlight specific Izumo1 and Juno residues used in mutational studies. (B) Example of Izumo1:Juno unbinding along the known binding interface as part of the two-state (2-S) pathway. (C) Example of Izumo1:Juno forming new Cluster 3 interactions during unbinding as part of the three-state (3-S) pathway. In B and C, the starting distance between C terminal residues of the equilibrated Izumo1:Juno complex is 6.8 nm. (D) The two-state and three-state unbinding pathway distribution across 50 SMD trajectories. (E) Sliding-rebinding distance vs. sliding-rebinding rate for the force-dependent rearrangement of Izumo1:Juno complexes in the two-state and three-state unbinding pathways. (F) Observed increases in contact area between Izumo1 and Juno in unloaded (t = 0 ns) and loaded (t = 2 ns, inlet panel A) complexes for the two-state and three-state unbinding pathways. (G) Peak forces extracted from two-state and three-state SMD simulations of Izumo1:Juno complexes (median ± MAD).

Despite only minor secondary structural alterations to Izumo1:Juno in the two-and three-state unbinding trajectories, the distance between the C-termini of Izumo1 and Juno show marked increases at several time points. Firstly, as Cluster 1 unbound, the Immunoglobulin-like domain of Izumo1 was pulled away from the binding interface (Fig. 2, B and C). Secondly, as Cluster 2 unbound, the hinge-domain of Izumo1 was also pulled away from the binding interface (Fig. 2, B and C). Thirdly, as Izumo1 straightened, it underwent progressive sliding and rebinding across the surface of Juno (Fig. 2F). This sliding-rebinding process was greatly prolonged by the gradual binding of Cluster 3 in the three-state unbinding trajectories, further increasing the distance between the C-termini of Izumo1 and Juno.

Across the 50 SMD replicas, the median distance change between the C-termini of Izumo1 and Juno was 5.7 ± 1.6 and 6.4 ± 0.9 nm (± MAD), for the two-and three-state unbinding trajectories, respectively. Additionally, in 4% of Izumo1:Juno SMD simulations we observed partial unfolding of the Juno C-terminal α-helix (fig. S4F, fig. S10). Unfolding of this Juno α-helix until the proximal disulfide bond would contribute a maximum contour length increment of 7.7 nm (residues 207 to 228, 21 aa × 0.365 nm). Consequently, the combined distance intervals for Izumo1:Juno rearrangement and unfolding of the Juno C-terminal α-helix observed in SMD are consistent with the intermediate unfolding events observed experimentally (fig. S4).

Prior to Cluster 1 unbinding, the contact area between Izumo1 and Juno increased as the Izumo1:Juno complex transitioned from the unloaded-state (*t* = 0 ns) to the loaded-state (*t* = 2 ns). For the two-state unbinding trajectories, the median contact surface increased from 848.7 ± 25 to 894.9 ± 62.9 Å^2^ (± MAD) (Fig. 2F), while for the three-state unbinding trajectories, the median contact surface area increased from 883.4 ± 24.7 to 921.9 ± 58.5 Å^2^ (± MAD) (Fig. 2F). Subsequent unbinding of Izumo1:Juno from State 2 resulted in median peak forces of 1372 ± 910.1 pN (± MAD) (Fig. 2G). With the gradual binding of Cluster 3 and transition to State 3 (Fig. 2C), the protracted period of sliding and rebinding resulted in a 61.1% increase in the median peak forces for the three-state unbinding trajectories, to 2210 ± 739.5 pN (± MAD) (Fig. 2G). In considering all above analyses, we propose that the two-state pathway is representative of P1 and that the three-state pathway is representative of P2.

Finally, we note that the three-state unbinding trajectories can involve divergent residues on Izumo1 and Juno. We identified a subtype comprising ∼10% of the three-state unbinding trajectories (Fig. 2D, fig. S11, fig. S12, Supplementary Movie 3). In this sub-pathway, the partially buried Izumo1 residues implicated in its membrane fusion function (F28, W88, and W113) (*11*) underwent a corkscrew-like motion, aligning towards the same interaction plane with increased solvent exposed surface area (fig. S12, Supplementary Movie 4). This finding, alongside other force-dependent structural rearrangements (see Supplementary Text, fig. S13), could help explain the unexpectedly low fusion rates for Izumo1 expressing cells under equilibrium conditions (*11*). Thus, mechanical force should be considered as a relevant parameter for future Izumo1 membrane fusion studies.

### Functional redundancy of higher order Izumo1:Juno unbinding pathways

To pinpoint crucial residues, we asked how Izumo1 point mutations might influence P0, P1, and P2. We introduced single alanine substitutions into well conserved residues within Clusters 1 (Izumo1 R160A) or 2 (Izumo1 H157A) (*9*), as well as two divergent residues within Cluster 2 (Izumo1 E71A and Izumo1 K150A) (*9*) (Fig. 2A). Using our bead-based flow cytometry method, we observed increases in *K_D_* for Izumo1 E71A:Juno, Izumo1 H157A:Juno, and Izumo1 R160A:Juno, relative to the wildtype, that were consistent with prior work (fig. S1) (*9*). For the previously uncharacterized interaction, Izumo1 K150A:Juno, we measured a *K_D_* of 110 ± 27 nM (SE).

Overall, using the same constant speed AFM-SMFS setup described in Fig. 1C, the unbinding pathways of the selected Izumo1 mutants did not deviate in terms of the median rupture forces or Δ*x* trends described for wild-type Izumo1:Juno (fig. S14-17, table S1). Thus, it becomes clear that while alanine substitutions of single Izumo1 residues are sufficient to disrupt the equilibrium binding behavior of Izumo1:Juno (fig. S1) (*9*), the mechanical behavior and force-dependent unbinding pathways offer a greater level of functional redundancy. Where disruption of the mechanical behavior is observed, it is largely confined to the unbinding pathway distributions (fig. S14-17), thereby implicating the invisible transition barriers separating P0, P1, and P2 (see Supplementary Text).

### Monte Carlo simulations support the multi-state catch bond mechanism

Multi-state kinetic Monte Carlo simulations (*14*, *29*) were next used to describe the mechanically induced dissociation of Izumo1:Juno and evaluate competing catch bond models. Based on the experimental constant speed AFM-SMFS datasets showing three pathways and the SMD simulations showing up to distinct three-Izumo1:Juno states, we considered two models each with three rupture pathways and three states (Fig. 3A: native (N), intermediate 1 (I_1_) and intermediate 2 (I_2_)). One model simulated irreversible transitions and the other simulated reversible transitions from N to I_1_ and from I_1_ to I_2_ (fig. S18). We used experimentally derived BE parameter sets (table S1) to describe the unbinding (*i.e.* rupture) events pertaining to P0, P1 and P2, and we fit new BE parameter sets for each state transition (table S4, table S5). Indeed, the model with reversible transitions was able to more closely recapitulate the experimental rupture force histograms (see Supplementary Text). This simplified model supports the interpretation that P1 and P2 arise through force-dependent processes (table S5) and that the state transitions are reversible.

**Figure 3:**
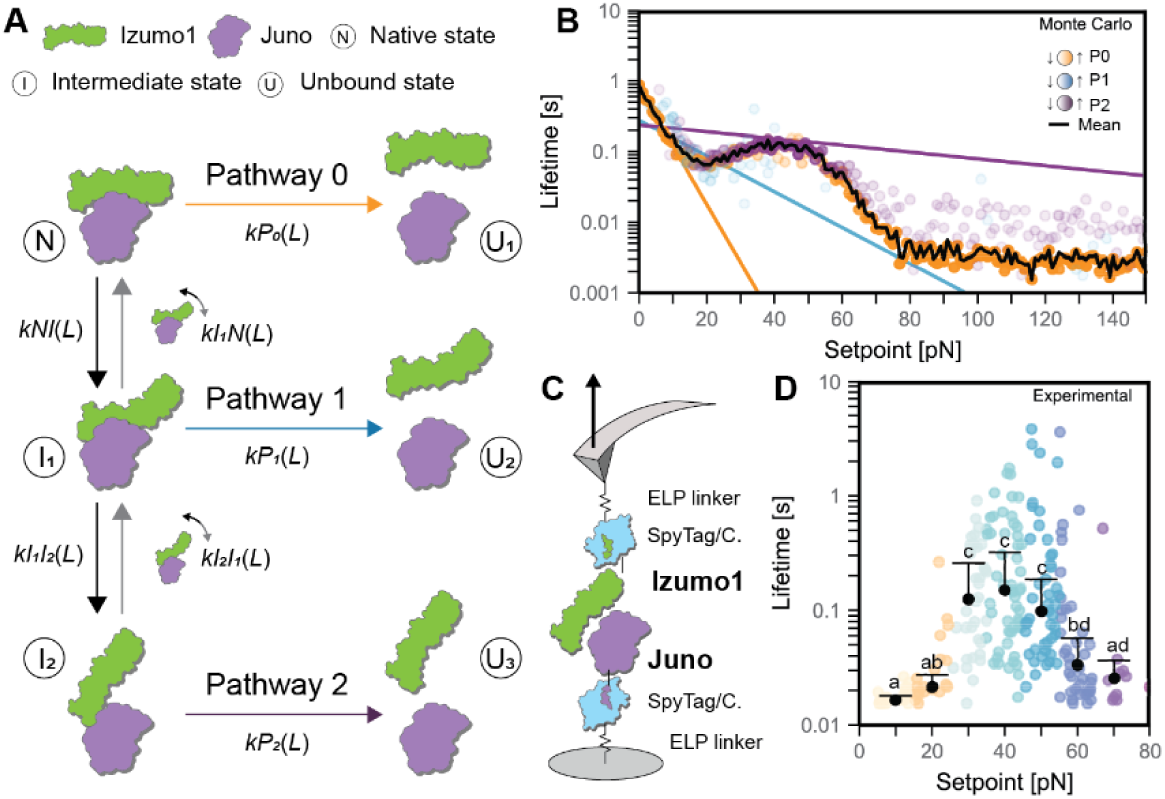
Modeling and validation of Izumo1:Juno forming a catch bond. (A) Proposed three-state kinetic model for unbinding of Izumo1:Juno with reversible transitions between multiple structural states. (B) Monte Carlo simulated force clamp AFM-SMFS for the Izumo1:Juno complex. Colored dots represent mean bond lifetime for each unbinding pathway at each setpoint with intensity representing the relative abundance of an unbinding pathway. Colored lines represent lifetimes calculated directly from BE-derived kinetic parameters (table S1, Eq. (11)). Black line represents the mean bond lifetime across all three pathways. (C) Schematic showing protein constructs and surface immobilization for force clamp AFM-SMFS. (D) Experimental bond lifetimes for Izumo1:Juno from force clamp AFM-SMFS. Colored dots represent individual bond lifetime measurements. Black dots represent median bond lifetime (± MAD) across 10 pN bins. Shared letters indicate the absence of statistically significant differences (p > 0.05) from Kruskal–Wallis tests followed by Dunn’s multiple-comparison tests.

The Monte Carlo model was then used to simulate Izumo1:Juno unbinding under constant force conditions (Fig. 3B). Our model predicted that the Izumo1:Juno slip-to-catch transition sets in at ∼20 pN, and is overtaken by a subsequent catch-to-slip transition at ∼40 pN. This predicted transition occurs close to the previously reported free swimming force of spermatocytes of 44 pN (*42*). The simulations show that from 20-40 pN, P2 was the most prominent unbinding pathway. While, at very low (<20 pN) and very high forces (>80 pN), P0 was found to dominate. We simulated clamping forces up to 150 pN to encompass the highest transverse forces recorded for hyperactivated sperm cells (*12*), such as would be present for sperm cells bound to egg cells. Interestingly, our simulation predicts a broad period of stability influenced by sequestering in I_2_ before eventual rupture through P0. Overall, these simulations highlight the potential interplay of all three unbinding pathways in stabilizing Izumo1:Juno across different biologically relevant force-regimes.

### Force clamp AFM-SMFS observation of Izumo1:Juno catch bond behavior

Force clamp AFM-SMFS was then used to experimentally validate our force clamp Monte Carlo simulations. Following a brief contact with the surface (0.2 s), the cantilever was retracted at 4000 nm·s^- 1^ until the target force setpoint was reached. The force was then maintained using a feedback loop that adjusted the height of the cantilever as required. The ddFLN4 fingerprint domains were omitted from the force clamp AFM-SMFS protein constructs to avoid biasing bond lifetime measurements at low forces (Fig. 3C). A detailed description of the curve selection procedure is provided (Supplementary Methods), which included replicates of Izumo1:blank controls (fig. S19). Representative force clamp curves are shown in fig. S20.

In agreement with the Monte Carlo simulations, the force clamp experiments show statistically significant increases in bond lifetime beginning at ∼20 pN and peaking at 40–50 pN (Fig. 3D, fig. S19). Over the slip-to-catch transition, we observed a 7.5-fold increase in median bond lifetime from 0.02 ± 0.003 to 0.15 ± 0.02 s (mean ± SEM) (Fig. 3D, fig. S19). At clamping forces from 80-160 pN, we observed approximately the same bond-clamping yield with no statistically significant difference between the bond lifetimes (representative examples: fig. S19, E and F). These results provide direct agreement with the force clamp Monte Carlo predictions regarding the emergence of the catch bond state at forces paralleling the free swimming force of spermatocytes (*42*). Further, they provide support for the predicted broad period of stability at the transverse forces typical of hyperactivated sperm cells on the egg cell membrane (*12*).

### Loss of mechanostability in the infertility-associated Juno mutant, JunoH177Q

To explore the role of mechanostability in human fertility, we characterized the infertility-associated mutant JunoH177Q. This particular variant was found in 9% of patients across a female idiopathic infertility cohort (*35*). Since the initial publication, the prevalence of the JunoH177Q variant has been measured at 0.15–0.18% in populations where greater than 150 Juno alleles have been sequenced (dbSNP database: rs76779571). Notably, Juno H177 is located distal to the equilibrium Izumo1:Juno binding interface and our flow cytometry-bead based binding assay demonstrates a high binding affinity (*K_D_* = 64 ± 14 nM (± SE)) for Izumo1:JunoH177Q. Thus, equilibrium-based analysis and structural data of the Izumo1:Juno complex (*9*, *43–45*) does not explain any deleterious effects of this infertility-associated mutation.

Within our SMD classification framework, Juno H177 is a Cluster 3 amino acid that interacts with Izumo1 as part of the three-state unbinding pathway (Fig. 2, A and C). In the protracted sliding-rebinding mechanism that defines State 3, Juno H177 forms both early and terminal interactions, first with Izumo1 N59&R56 (Fig. 4A) and later Izumo1 K48 (Fig. 4B). Across 50 SMD simulations of Izumo1:JunoH177Q unbinding, the stable interactions between Izumo1 N59&R56-Juno H177 and Izumo1K48-JunoH177 are either replaced with transient interactions or lost completely (Fig. 4, C and D; fig. S21). While the peak forces during the three-state unbinding of Izumo1:JunoH177Q are similar to wild-type Izumo1:Juno, we observed a 5.9% decrease in the median contact surface area when comparing the unloaded (*t* = 0 ns) to the loaded states (*t* = 2 ns), from 918.6 ± 81.8 to 864.1 ± 95.7 Å^2^ (± MAD) (Fig. 4E). This coincided with a 13.3% reduction in median peak force for the two-state unbinding trajectories (Fig. 4F), from 1372.4 ± 630.1 pN to 1190.1 ± 687.6 pN (± MAD) and a decrease in the prevalence of three-state unbinding trajectories, from 80% to 58% (Fig. 4G). Thus, we conclude that the stabilization of State 2 through force-dependent increases in contact surface area around Juno H177 facilitates Cluster 3 formation and catch bond activation in wild-type Izumo1:Juno.

**Figure 4:**
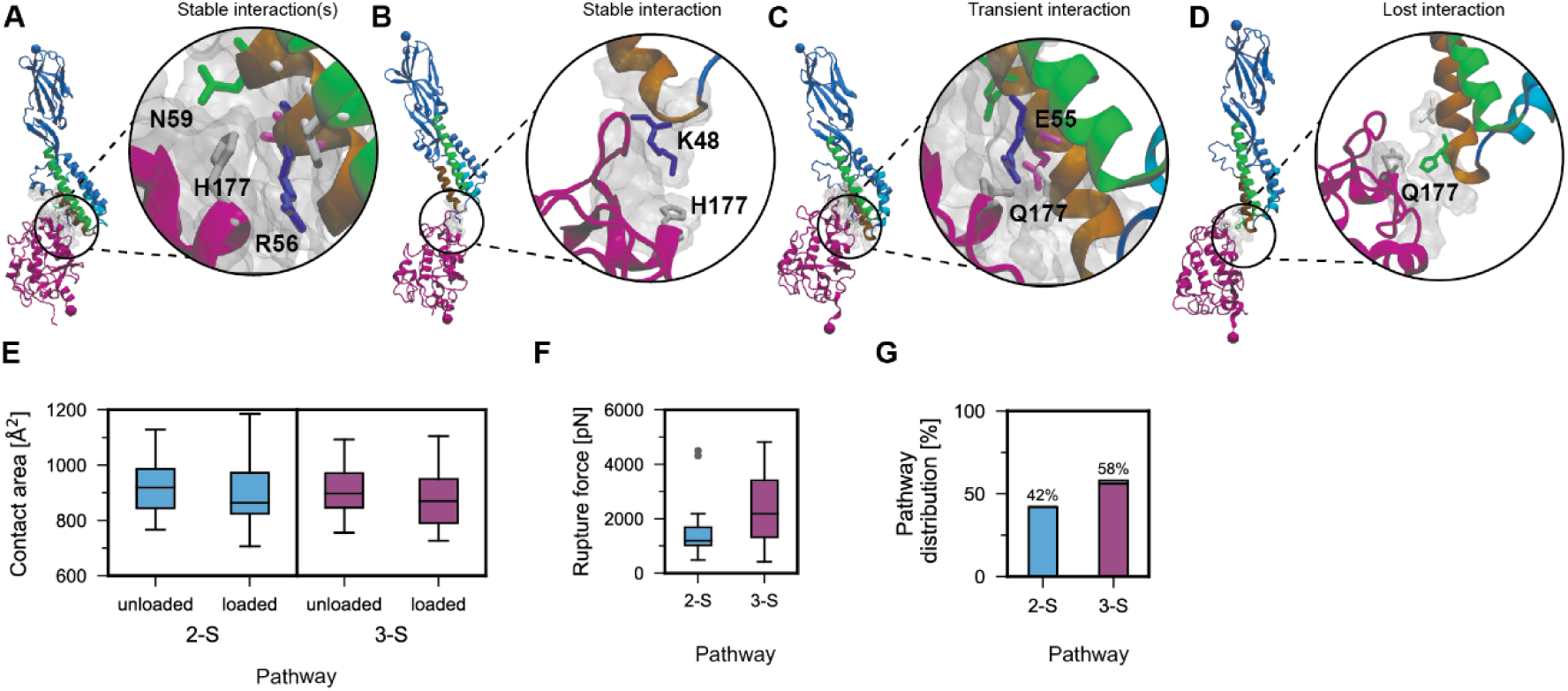
Juno H177 mediates the transition to higher order unbinding pathways in Izumo1:Juno. (A) Exemplary snapshot of stable long-lasting Cluster 3 interactions mediated by Juno H177. (B) Exemplary snapshot of terminal complex-prolonging interactions mediated by Izumo1 K48 and Juno H177. (C) Exemplary snapshot of transient Cluster 3 interactions formed in the Izumo1:JunoH177Q infertility mutant. (D) Exemplary snapshot of terminal interactions that are absent in Izumo1:JunoH177Q. (E) Observed decreases in contact area between Izumo1 and JunoH177Q in unloaded (t = 0 ns) and loaded (t = 2 ns) complexes for the two-state (2-S) and three-state (3-S) unbinding pathways. (F) Peak forces extracted from two-state and three-state SMD simulations of Izumo1:JunoH177Q complexes (median ± MAD). (G) Two-state and three-state unbinding pathway distribution for Izumo1:H177Q across 50 SMD trajectories.

Noteworthy, in SMD simulations where Juno H177 was mutated to a negatively charged glutamic acid residue (JunoH177E) the median peak-force of the two-state unbinding trajectories increased by 55%, from 1372.4 ± 630.1 pN to 2129.9 ± 654.5 pN (± MAD) (fig. S22A). Additionally, for Izumo1:JunoH177E we note the absence of the three-state unbinding subpathway (fig. S22B) linked to the solvent-exposure of membrane fusion-associated Izumo1 residues (fig. S12) (*11*). With contact surface area increases between the unloaded (*t* = 0 ns) to the loaded states (*t* = 2 ns) of Izumo1:JunoH177E (fig. S22C) similar to wild-type Izumo1:Juno, the increased mechanostability of JunoH177E in State 2 and the absence of the three-state unbinding subpathway appears to be charge related. Finally, similar to JunoH177Q, for JunoH177E we observed an increase in unfolding rate for the C-terminal Juno α-helix (residues 207 to 228, fig. S4F), from 4% to 14% (fig. S10C). This increase in Juno unfolding corresponded to a slightly increased median distance between the C-termini of Izumo1:JunoH177Q and Izumo1:JunoH177E at rupture, from 12.9 ± 7.2 to 13.4 ± 7.5 and 12.4 ± 7.3 to 13.9 ± 7.4 nm (± MAD), respectively.

Experimentally, constant speed AFM-SMFS shows that whilst three unbinding pathways can be classified for Izumo1:JunoH177Q using the GMM, the median rupture forces for P0, P1, and P2 have all decreased. Here, P0, P1, and P2 exhibit median rupture forces of 26.1 to 33.1 pN for loading rates of 1 to 12.3 nN·s^-1^, 48.5 to 59 pN for loading rates of 2.1 to 18.9 nN·s^-1^, and 114.2 to 115.9 pN for loading rates of 7.4 to 42.2 nN·s^-1^, respectively (Fig. 5, A and B). The DHS and BE fitted Δ*x* values for P1 and P2 are several-fold higher than in wild-type Izumo1:Juno (table S1), implicating increased conformational flexibility in the loss of mechanostability of these pathways. The disruption of temporary sequestration of complexes within mechanostable intermediate states (Fig. 3A) could explain secondary effects on the mechanostability of P0 in the absence of large changes to Δ*x* (*34*). In support of this hypothesis, we observed a lower prevalence of transition from P0 to P1 and P1 to P2 at all pulling speeds. We also observed an increased contour length change for P1 and P2 intermediate states that could be linked to an increased frequency of unfolding for the C-terminal Juno helix (fig. S10C, fig. S23B). The resulting median contour length changes amount to 19.9 ± 10 and 20 ± 9.8 nm (± MAD) for P1 and P2 at pulling speeds of 3200 nm·s^-1^, respectively (fig. S23, C and D). Overall, constant speed AFM-SMFS results showing lower mechanostability, greater conformational flexibility, decreased transition through higher order unbinding pathways, and an increased contour length change for the intermediate states, all support the SMD results implicating Juno H177 in unbinding pathway transition and the overall mechanostability of Izumo1:Juno.

**Figure 5:**
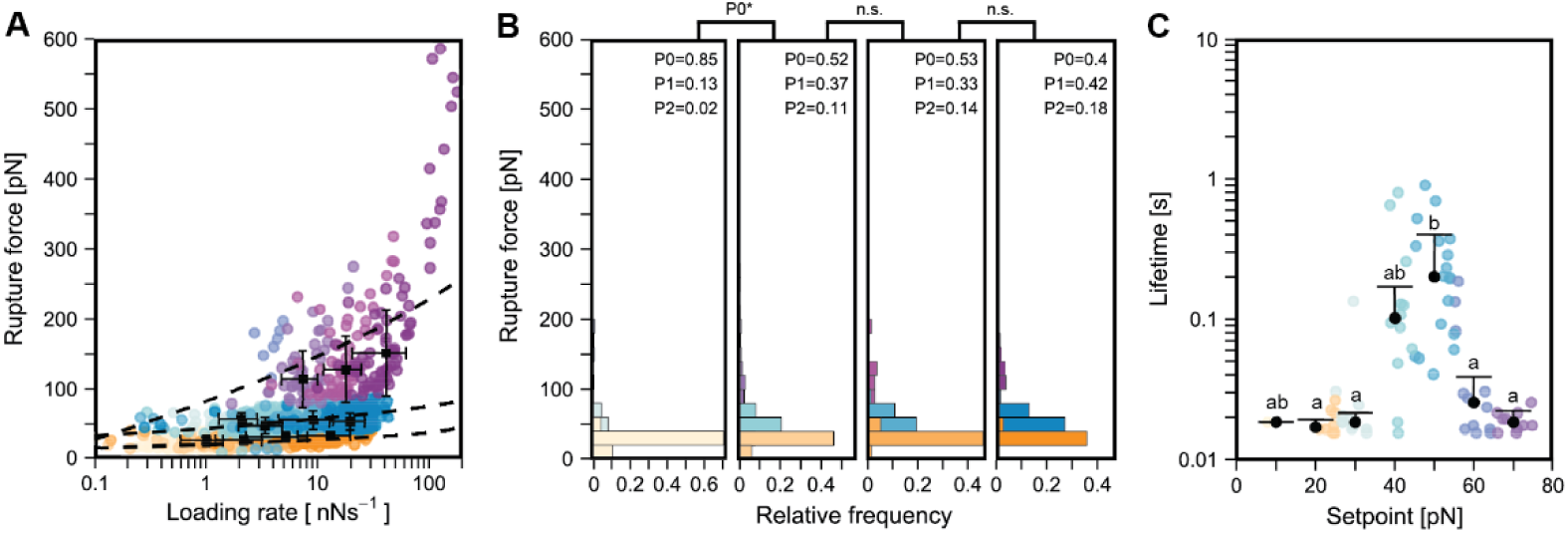
Constant speed and force clamp AFM-SMFS reveals the loss of mechanostability in Izumo1:JunoH177Q. (A) Rupture force vs. loading rate plot for Izumo1:JunoH177Q (median ± MAD). Dashed lines show the DHS model fits (Eq. (6), Eq. (7), table S1). (B) Izumo1:JunoH177Q rupture force histograms (bins = 20 pN). Experimental data in A and B was pooled from four independent experiments (n = 367-533) with significance (*p < 0.05) determined using Kurskal-Wallis tests followed by pairwise Wilcoxon rank-sum tests, see supplementary for specific details. (C) Bond lifetimes for Izumo1:JunoH177Q from force clamp AFM-SMFS. Colored dots represent individual bond lifetime measurements. Black dots represent median bond lifetime (± MAD) across 10 pN bins. Shared letters indicate the absence of statistically significant differences (p > 0.05) using Kruskal–Wallis tests followed by Dunn’s multiple-comparison tests.

Monte Carlo simulations at constant forces predict the preservation of catch bond behavior in Izumo1:JunoH177Q (fig. S23E). However, I_2_-dependent stabilization of mean bond lifetimes at 70-150 pN, as observed for wildtype Izumo1:Juno, is lost (Fig. 3B, fig. S23E). With the majority of the predicted bond lifetimes for Izumo1:JunoH177Q below the detection threshold of our force clamp AFM-SMFS setup (0.02 s), we expected a lower bond-clamping yield and an over sampling of longer lifetime events. In force clamp AFM-SMFS of Izumo1:JunoH177Q, we observed catch bond behavior with an average median peak bond lifetime of 0.17 ± 0.04 s (± SEM) at 40-50 pN (Fig. 5C; fig. S23, F and G). We were also unable to directly measure the loss of I_2_-dependent stabilization at 80-160 pN. However, in line with our Monte Carlo predictions, cantilevers confirmed to be active showed fewer bond clamping events in the 80–160 pN force range for Izumo1:JunoH177Q (representative example: fig. S23H) compared to wild-type Izumo1:Juno (representative examples: fig. S19, E and F). Together, the SMFS experiments and simulations for Izumo1:JunoH177Q strongly support the hypothesis that this infertility mutation inhibits the effect of sequestering complexes in I_2_, thereby degrading the overall mechanostability.

## Summary and conclusion

In summary, we used constant speed and force clamp AFM-SMFS, combined with kinetic Monte Carlo modeling and all-atom SMD to discover and quantify three unbinding pathways of the Izumo1:Juno complex involved in human fertilization. We showed how loading rate-dependent transitions between these three conformational states gave rise to a multi-state catch bond that generates mechanostability across a broad force range. Unbinding at forces up to 600 pN in constant speed AFM-SMFS experiments, Izumo1:Juno represents one of the most mechanostable eukaryotic protein:protein interactions studied to date. Comparable examples of mechanostability having previously only been observed in muscle protein complexes involved in force transduction (*13*) and bacterial receptors involved in matrix adhesion (*25*, *26*) and cellulose degradation (*29*).

The mechanical behavior of Izumo1:Juno is consistent with natural selection under the influence of two-different sperm-egg binding scenarios. In the first-binding scenario, marked by the initial adhesion of freely swimming sperm to egg cells, healthy human sperm cells have a swimming force of approximately 44 pN (*42*). This previously reported swimming force is remarkably consistent with the force at which the Izumo1:Juno catch bond state is most stable, experimentally and in our kinetic model. The second-binding scenario is characterized by the stable long-term adhesion between sperm and egg, directly preceding membrane fusion. Here, transverse forces generated by the flagellum of hyperactivated sperm cells can be on the order of 130 pN in primates (*12*). In this force range, our kinetic model predicts that despite a return to slip bond behavior, the bond lifetime of intact Izumo1:Juno complexes is stabilized by temporary sequestering in highly mechanostable states (*i.e.* I_1_ and I_2_). The overall implication of our kinetic model is that any loss of sperm motility impairing the transition of Izumo1:Juno to I_1_ or I_2_ would inhibit both sperm-egg binding scenarios. Importantly, we also find that the infertility-associated mutant, JunoH177Q, is implicated in the proposed high-force stabilization mechanism. This mechanism involves a novel Izumo1:Juno binding interface, not observed in the equilibrium crystal structure, that could represent a valid drug target for the development of both non-hormonal birth control and fertility treatments, which so far have had only limited success (*46*).

Our work therefore demonstrates a highly mechanostable multi-state catch bond mechanism in Izumo1:Juno, identifies a new non-equilibrium binding site for potential drug development, and provides new biophysical and computational avenues for understanding human fertility.

## Author Contributions

S.B. prepared biological samples, performed experiments and analyzed experimental data, with assistance from B.Y. S.B. performed the Monte Carlo simulations with assistance from H.L. P.P. performed and analyzed SMD simulations and prepared SMD visualizations. S.B. prepared the manuscript with contributions from all authors.

M.A.N. and V.V. conceived of and supervised the project.

## Acknowledgements

We would like to express our sincere gratitude to Dr. Carleen Kluger whose insightful preliminary discussions significantly shaped the direction of this project. We thank Dr. Maria Grazia Giuffreda and her team at Swiss National Supercomputing Center for their technical support and assistance in managing computational resources for this project.

## Supplementary Materials

Materials and Methods

Supplementary Text

Figures S1 to S23

Tables S1 to S5

Movies S1-S5

Supplementary PDB Files F1-F3

Supplementary References (*47–66*)

---

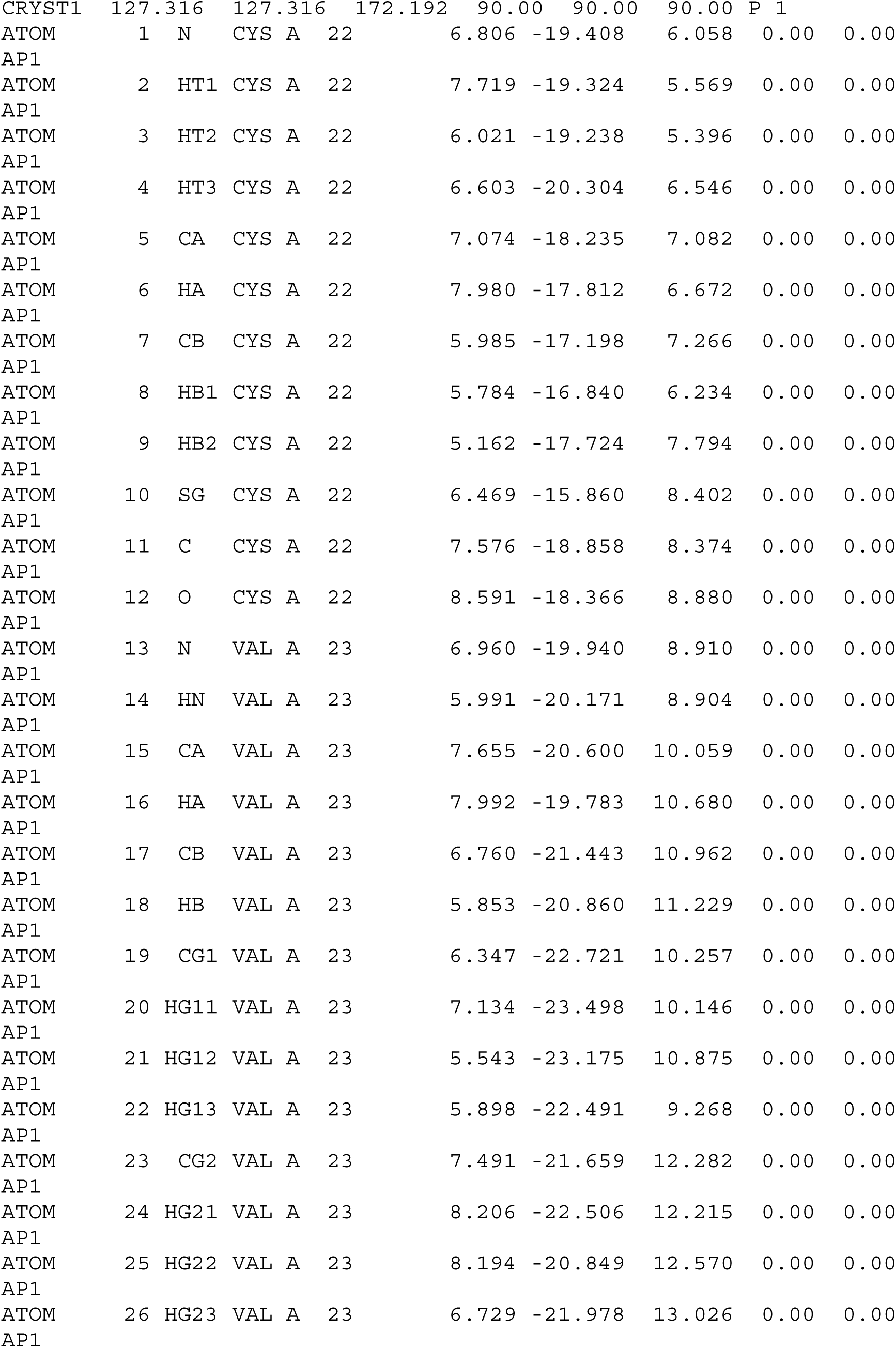

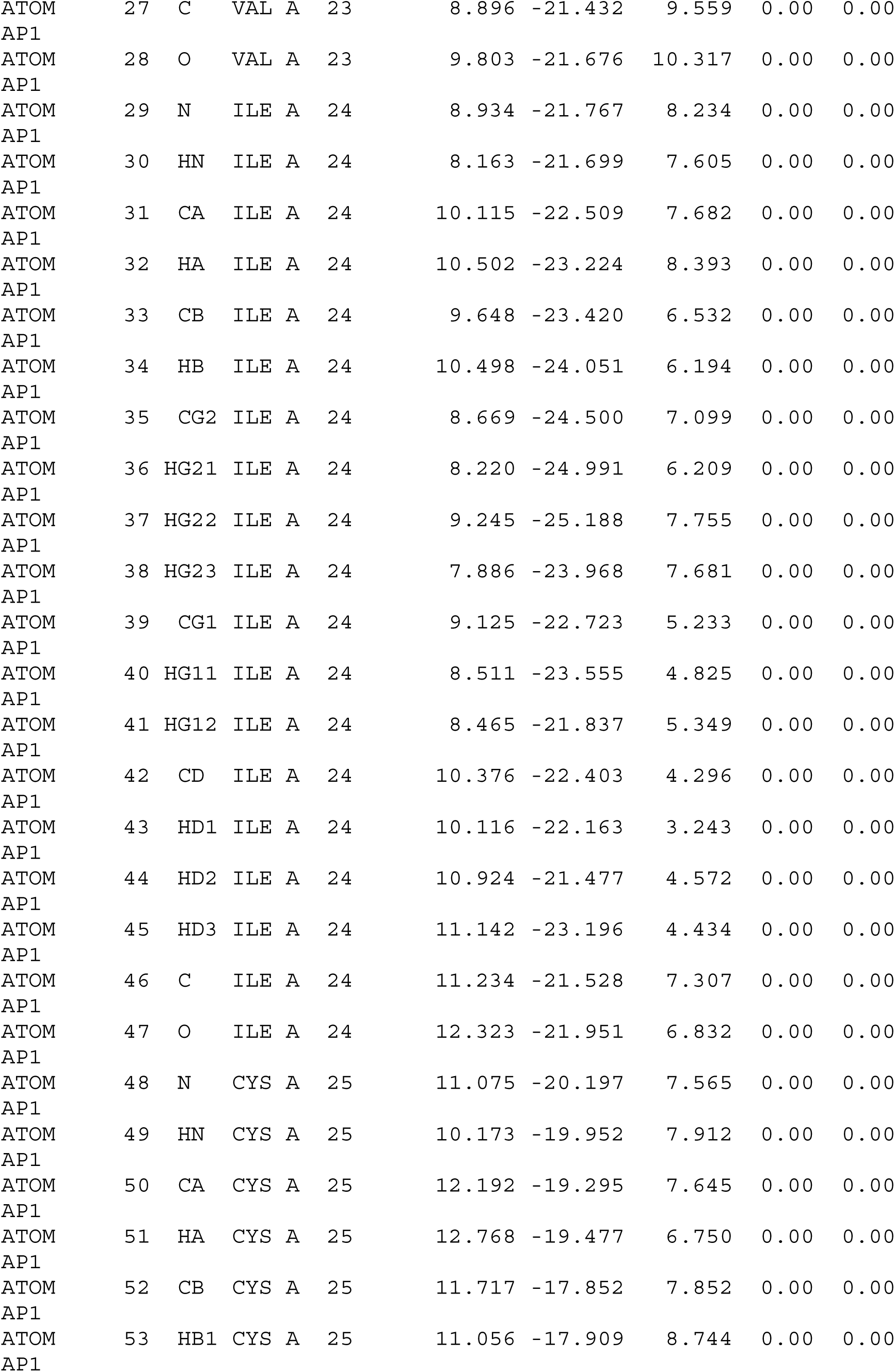

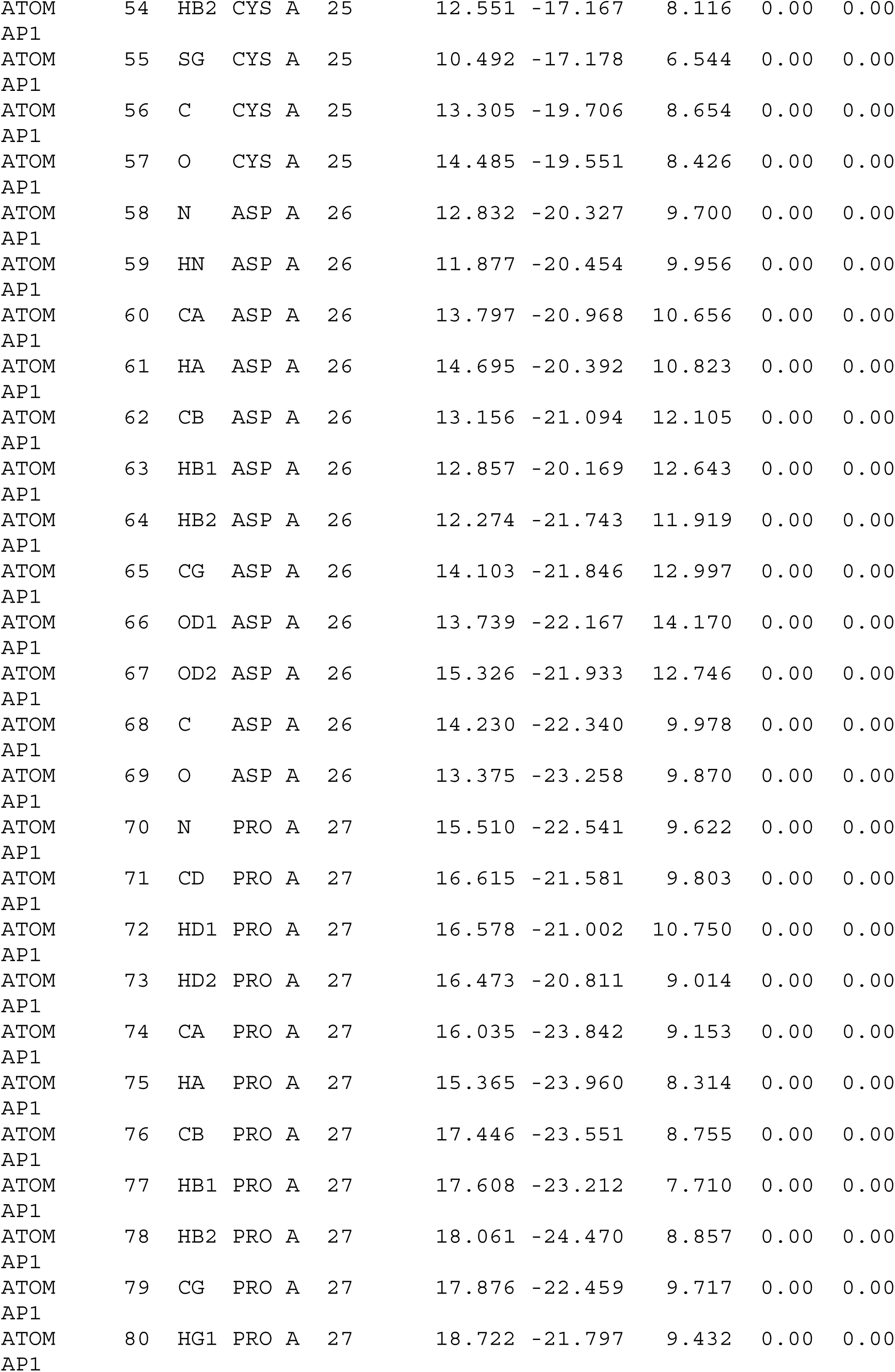

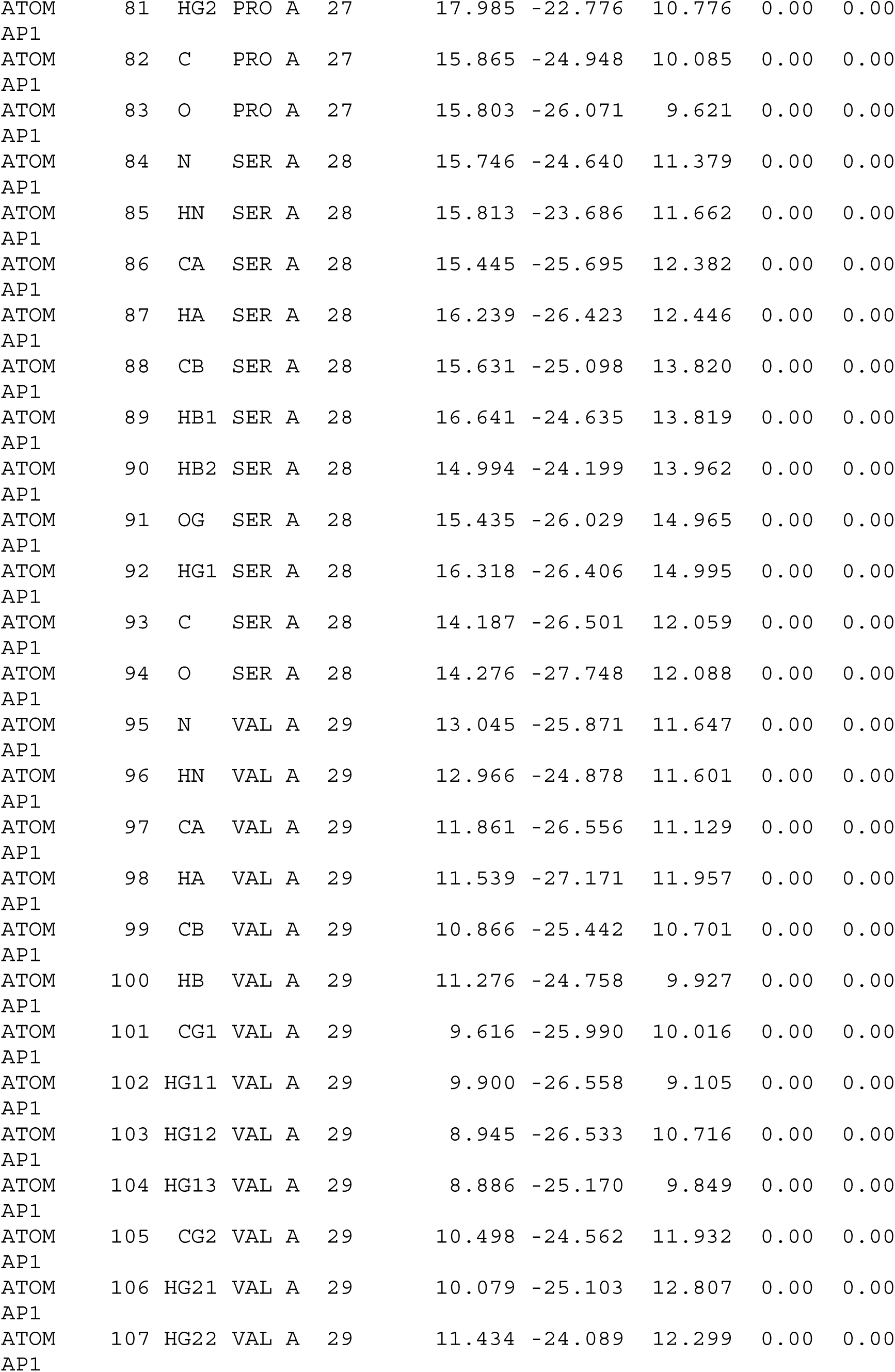

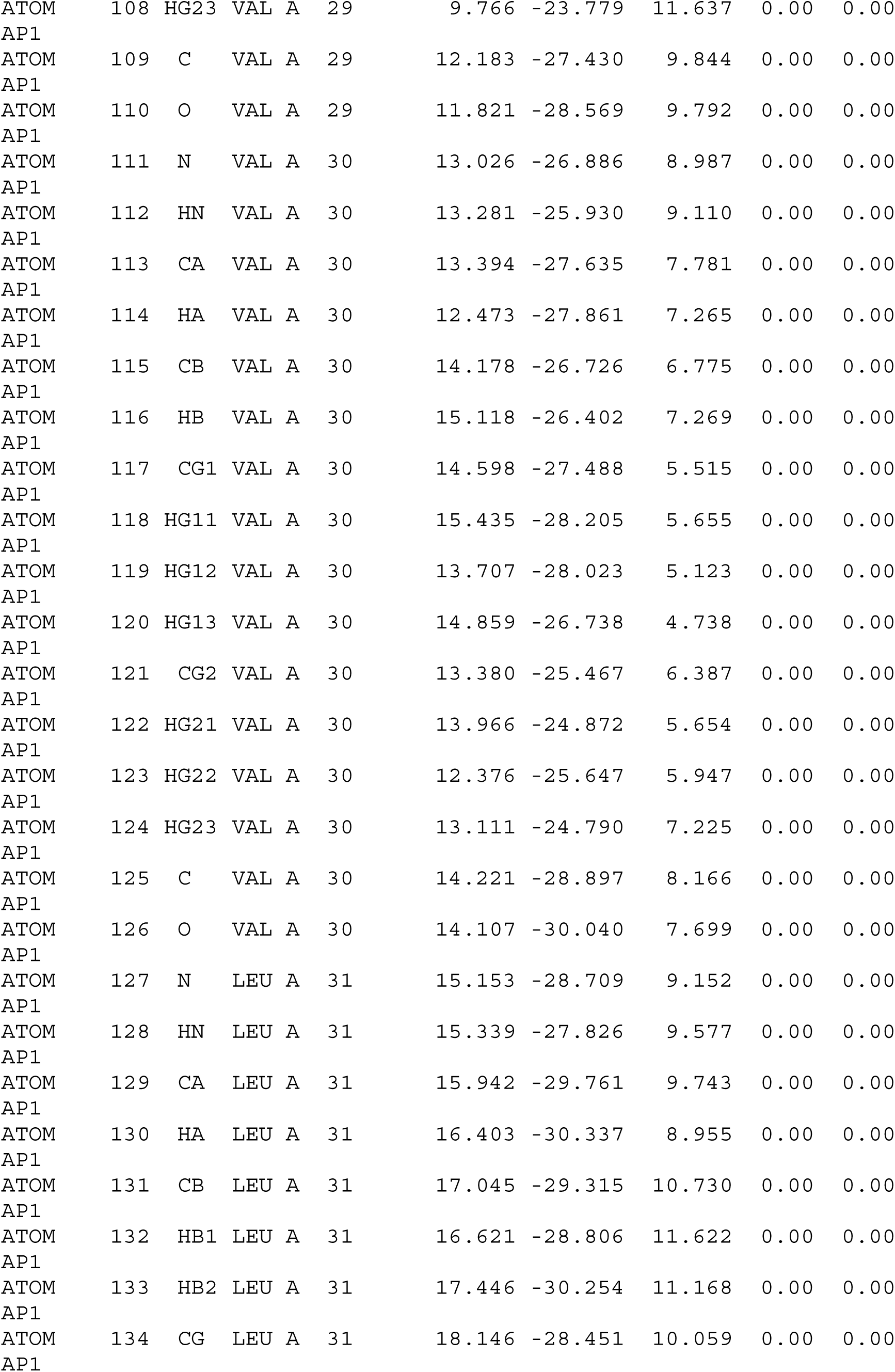

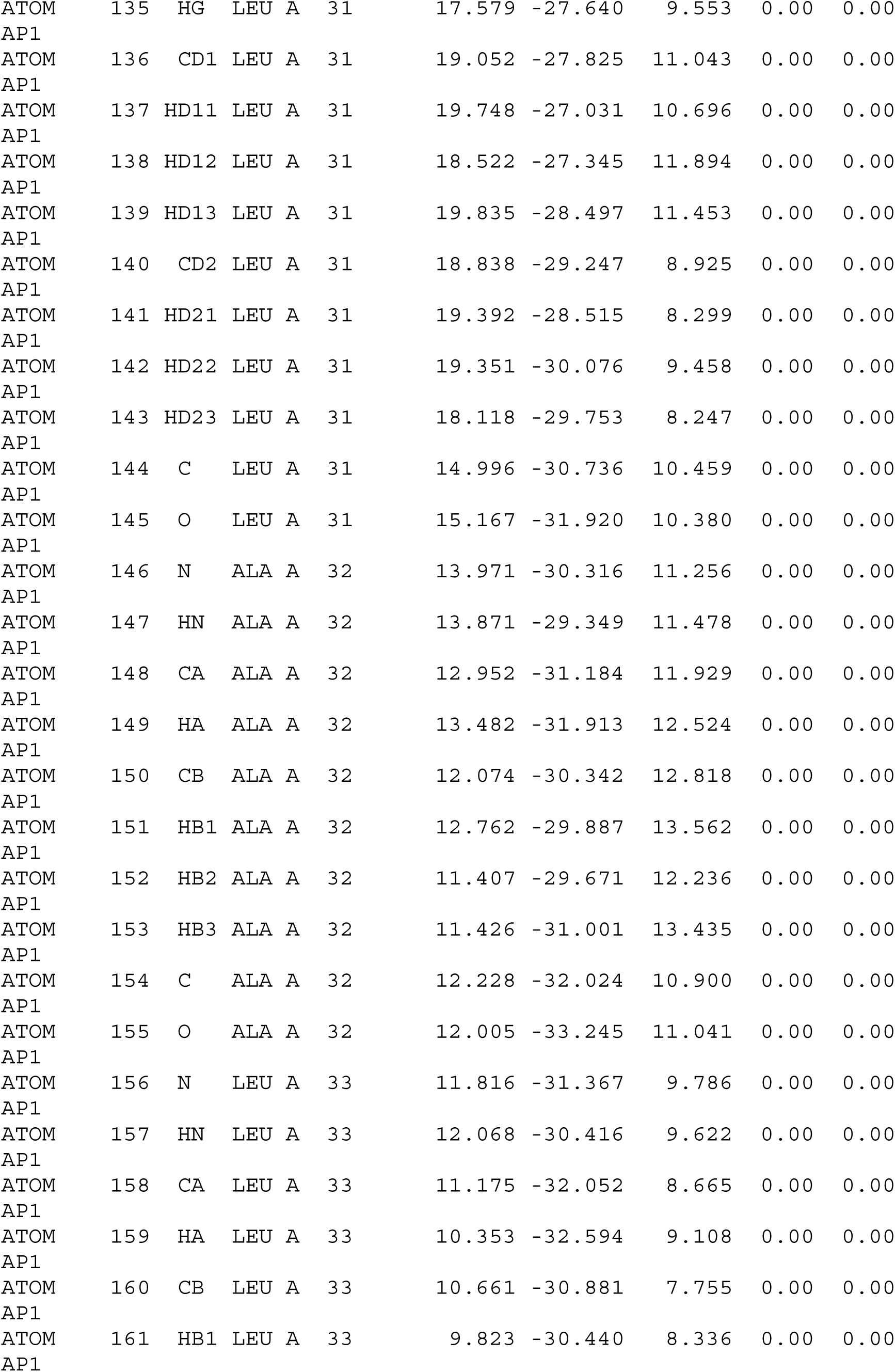

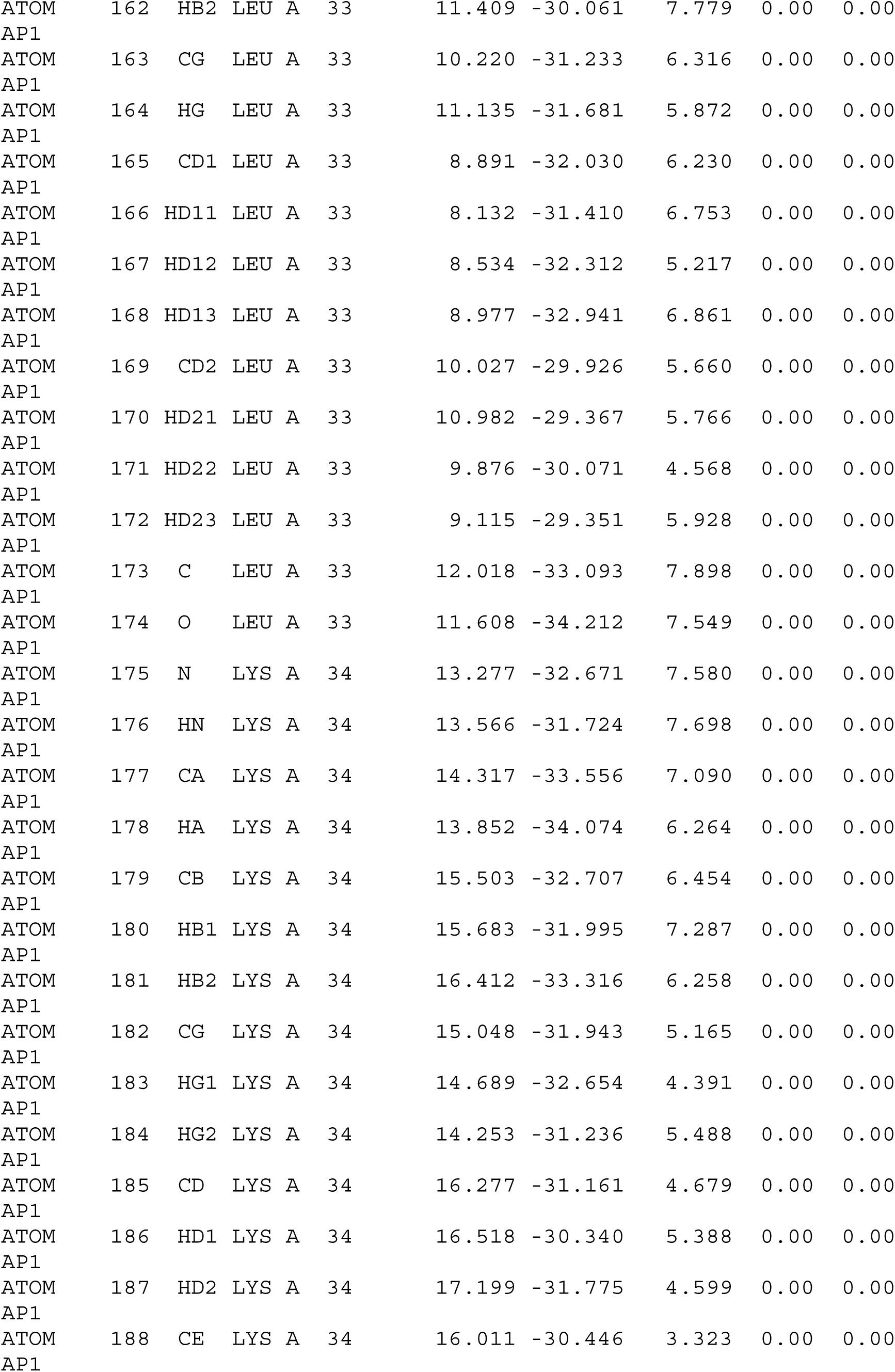

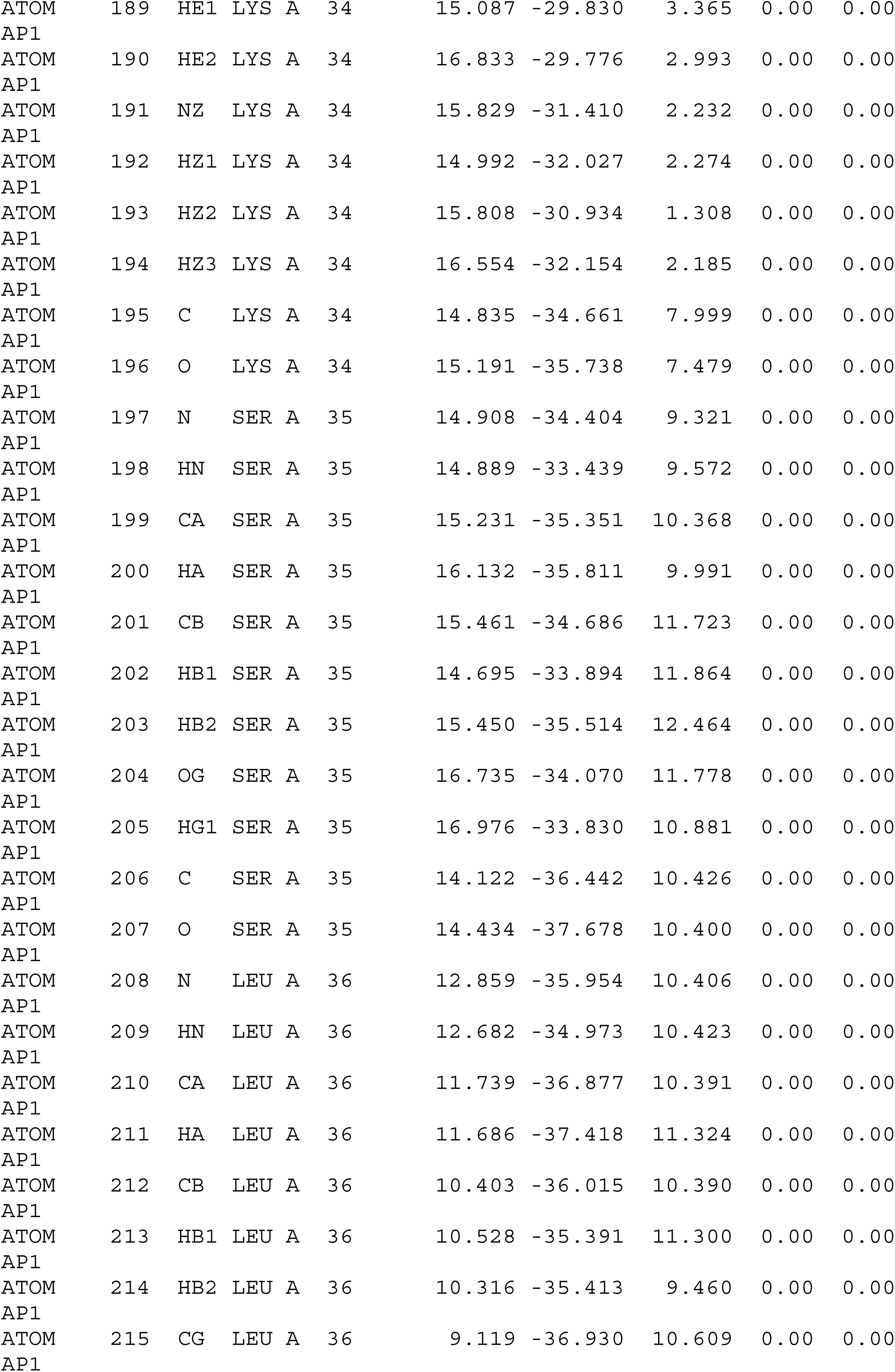

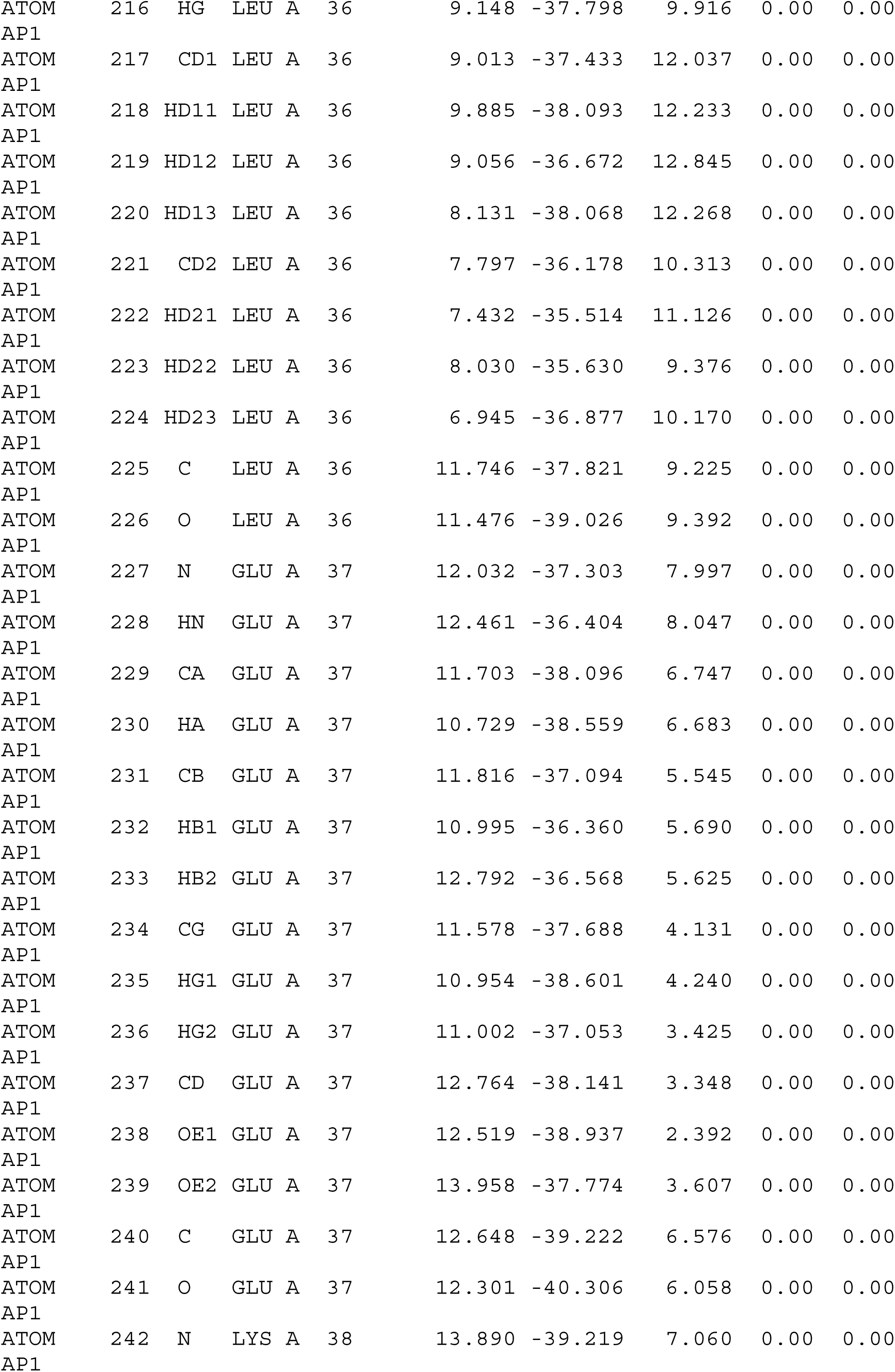

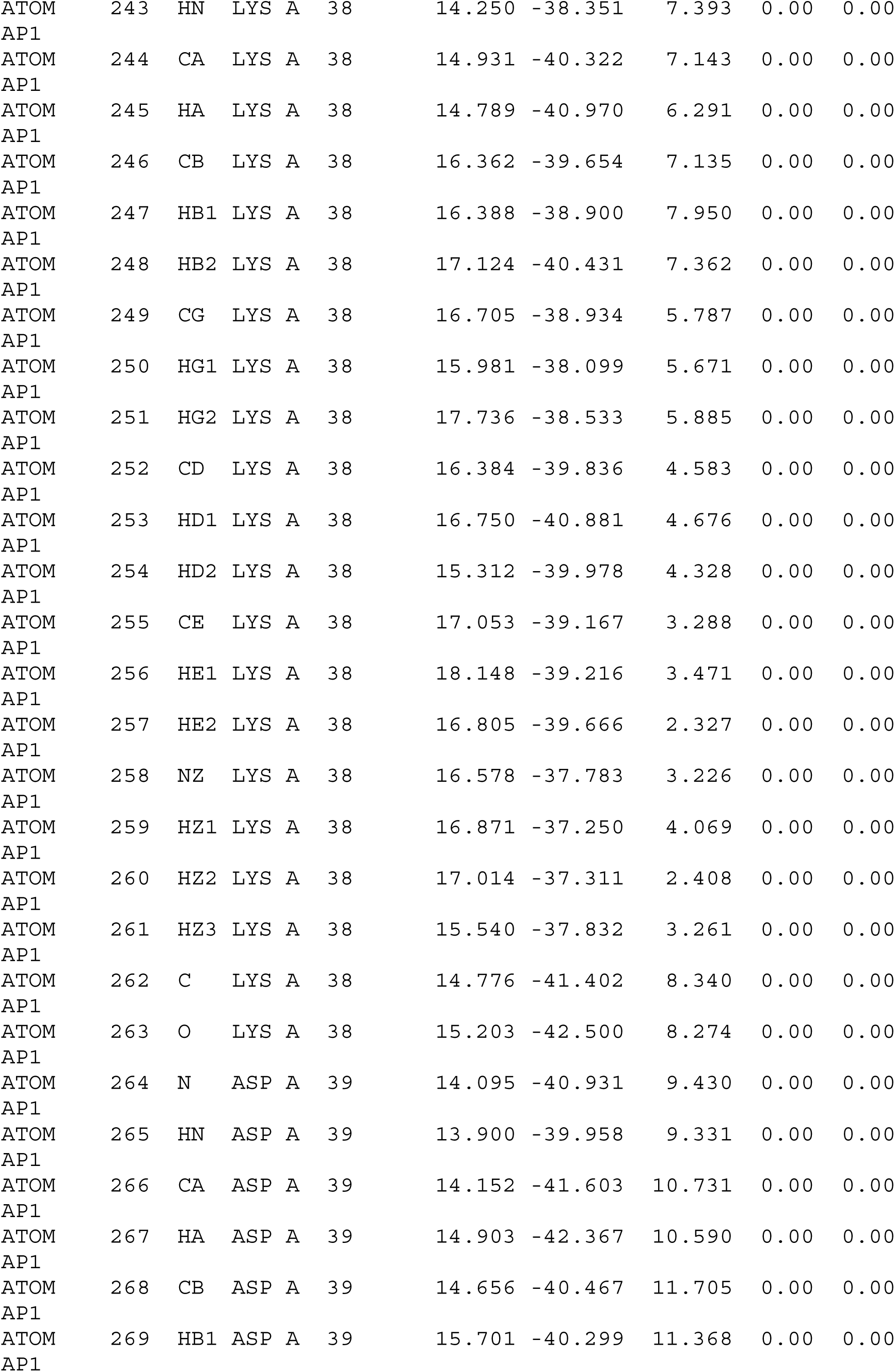

|  |  |  |  |  |  |  |  |  |  |  |  |
| --- | --- | --- | --- | --- | --- | --- | --- | --- | --- | --- | --- |
| ATOM<br>AP1 | 270 | HB2 | ASP | A | 39 |  | 14.087 | -39.557 | 11.416 | 0.00 | 0.00 |
| ATOM<br>AP1 | 271 | CG | ASP | A | 39 |  | 14.610 | -40.778 | 13.212 | 0.00 | 0.00 |
| ATOM<br>AP1 | 272 | OD1 | ASP | A | 39 |  | 14.611 | -39.829 | 14.012 | 0.00 | 0.00 |
| ATOM<br>AP1 | 273 | OD2 | ASP | A | 39 |  | 14.790 | -41.987 | 13.539 | 0.00 | 0.00 |
| ATOM<br>AP1 | 274 | C | ASP | A | 39 |  | 12.811 | -42.319 | 11.014 | 0.00 | 0.00 |
| ATOM<br>AP1 | 275 | O | ASP | A | 39 |  | 12.810 | -43.583 | 11.285 | 0.00 | 0.00 |
| ATOM<br>AP1 | 276 | N | TYR | A | 40 |  | 11.690 | -41.583 | 11.128 | 0.00 | 0.00 |
| ATOM<br>AP1 | 277 | HN | TYR | A | 40 |  | 11.670 | -40.633 | 10.824 | 0.00 | 0.00 |
| ATOM<br>AP1 | 278 | CA | TYR | A | 40 |  | 10.430 | -42.049 | 11.667 | 0.00 | 0.00 |
| ATOM<br>AP1 | 279 | HA | TYR | A | 40 |  | 10.521 | -42.939 | 12.273 | 0.00 | 0.00 |
| ATOM<br>AP1 | 280 | CB | TYR | A | 40 |  | 9.661 | -40.952 | 12.487 | 0.00 | 0.00 |
| ATOM<br>AP1 | 281 | HB1 | TYR | A | 40 |  | 10.478 | -40.617 | 13.161 | 0.00 | 0.00 |
| ATOM<br>AP1 | 282 | HB2 | TYR | A | 40 |  | 9.248 | -40.110 | 11.891 | 0.00 | 0.00 |
| ATOM<br>AP1 | 283 | CG | TYR | A | 40 |  | 8.456 | -41.456 | 13.274 | 0.00 | 0.00 |
| ATOM<br>AP1 | 284 | CD1 | TYR | A | 40 |  | 8.604 | -41.600 | 14.693 | 0.00 | 0.00 |
| ATOM<br>AP1 | 285 | HD1 | TYR | A | 40 |  | 9.619 | -41.488 | 15.046 | 0.00 | 0.00 |
| ATOM<br>AP1 | 286 | CE1 | TYR | A | 40 |  | 7.495 | -41.992 | 15.526 | 0.00 | 0.00 |
| ATOM<br>AP1 | 287 | HE1 | TYR | A | 40 |  | 7.571 | -42.141 | 16.593 | 0.00 | 0.00 |
| ATOM<br>AP1 | 288 | CZ | TYR | A | 40 |  | 6.232 | -42.077 | 14.970 | 0.00 | 0.00 |
| ATOM<br>AP1 | 289 | OH | TYR | A | 40 |  | 5.210 | -42.526 | 15.757 | 0.00 | 0.00 |
| ATOM<br>AP1 | 290 | HH | TYR | A | 40 |  | 4.651 | -43.091 | 15.218 | 0.00 | 0.00 |
| ATOM<br>AP1 | 291 | CD2 | TYR | A | 40 |  | 7.179 | -41.448 | 12.714 | 0.00 | 0.00 |
| ATOM<br>AP1 | 292 | HD2 | TYR | A | 40 |  | 7.039 | -41.318 | 11.651 | 0.00 | 0.00 |
| ATOM<br>AP1 | 293 | CE2 | TYR | A | 40 |  | 6.052 | -41.749 | 13.588 | 0.00 | 0.00 |
| ATOM<br>AP1 | 294 | HE2 | TYR | A | 40 |  | 5.039 | -41.797 | 13.217 | 0.00 | 0.00 |
| ATOM<br>AP1 | 295 | C | TYR | A | 40 |  | 9.513 | -42.552 | 10.532 | 0.00 | 0.00 |
| ATOM<br>AP1 | 296 | O | TYR | A | 40 |  | 8.788 | -43.557 | 10.697 | 0.00 | 0.00 |

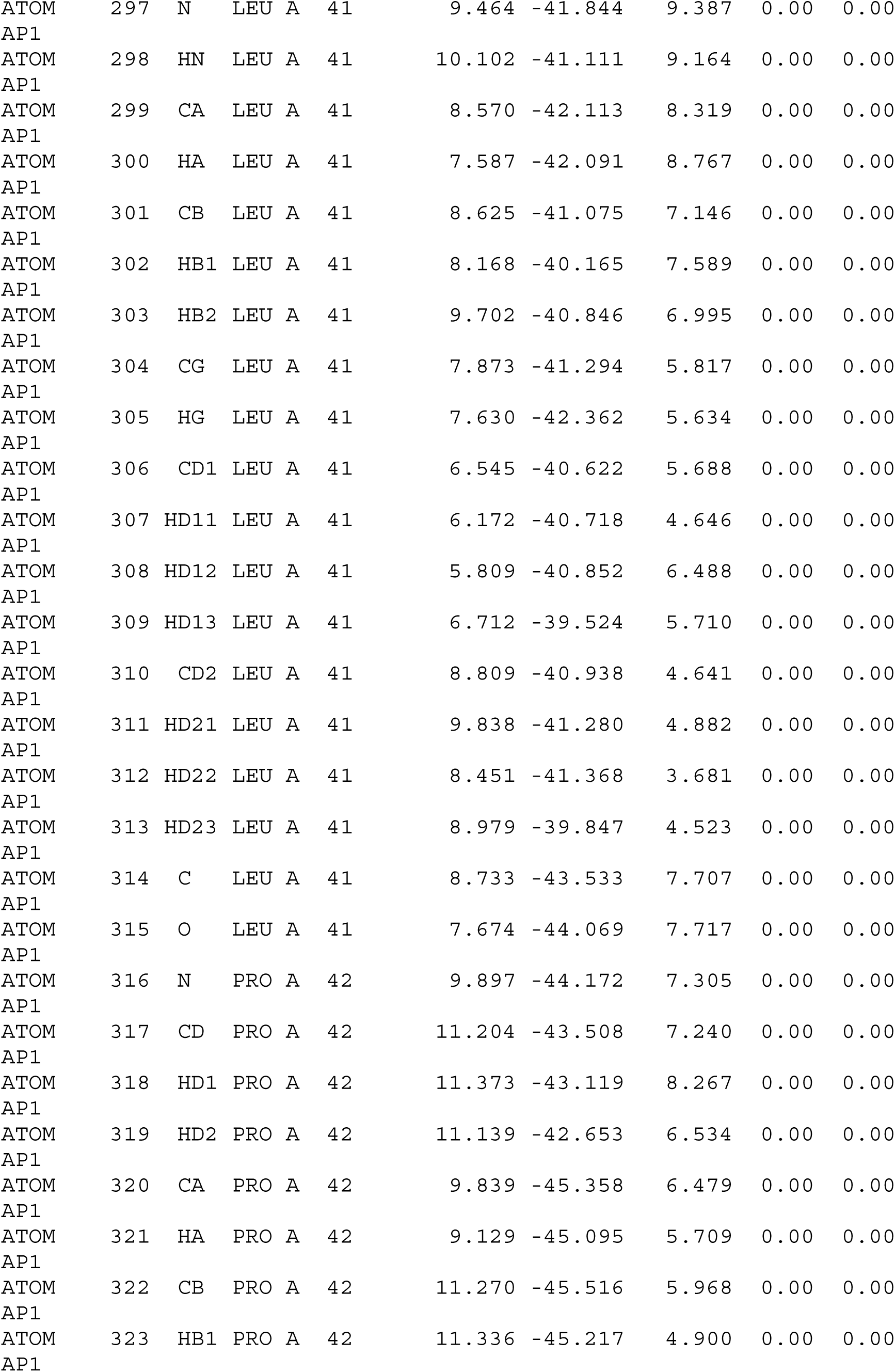

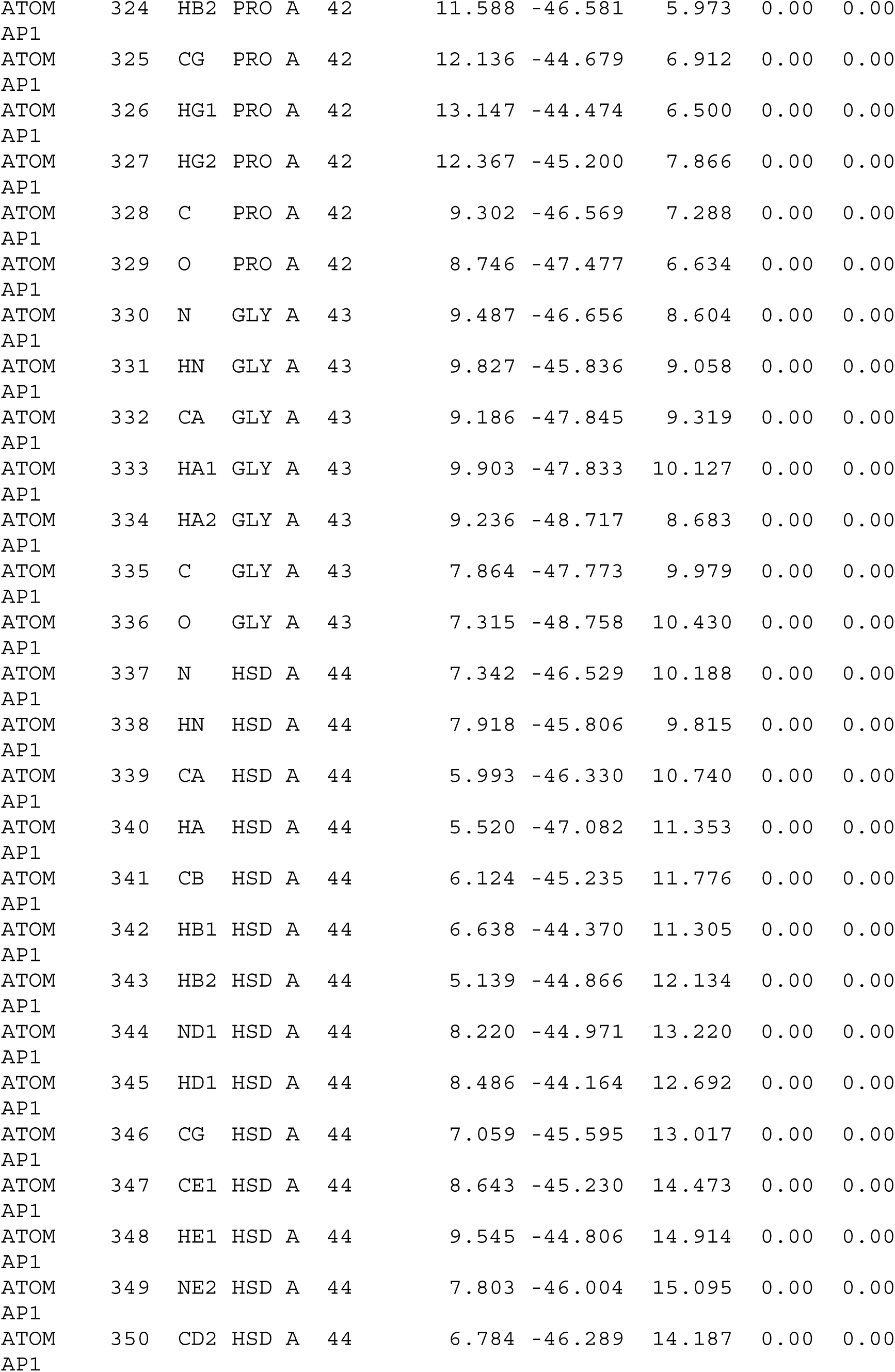

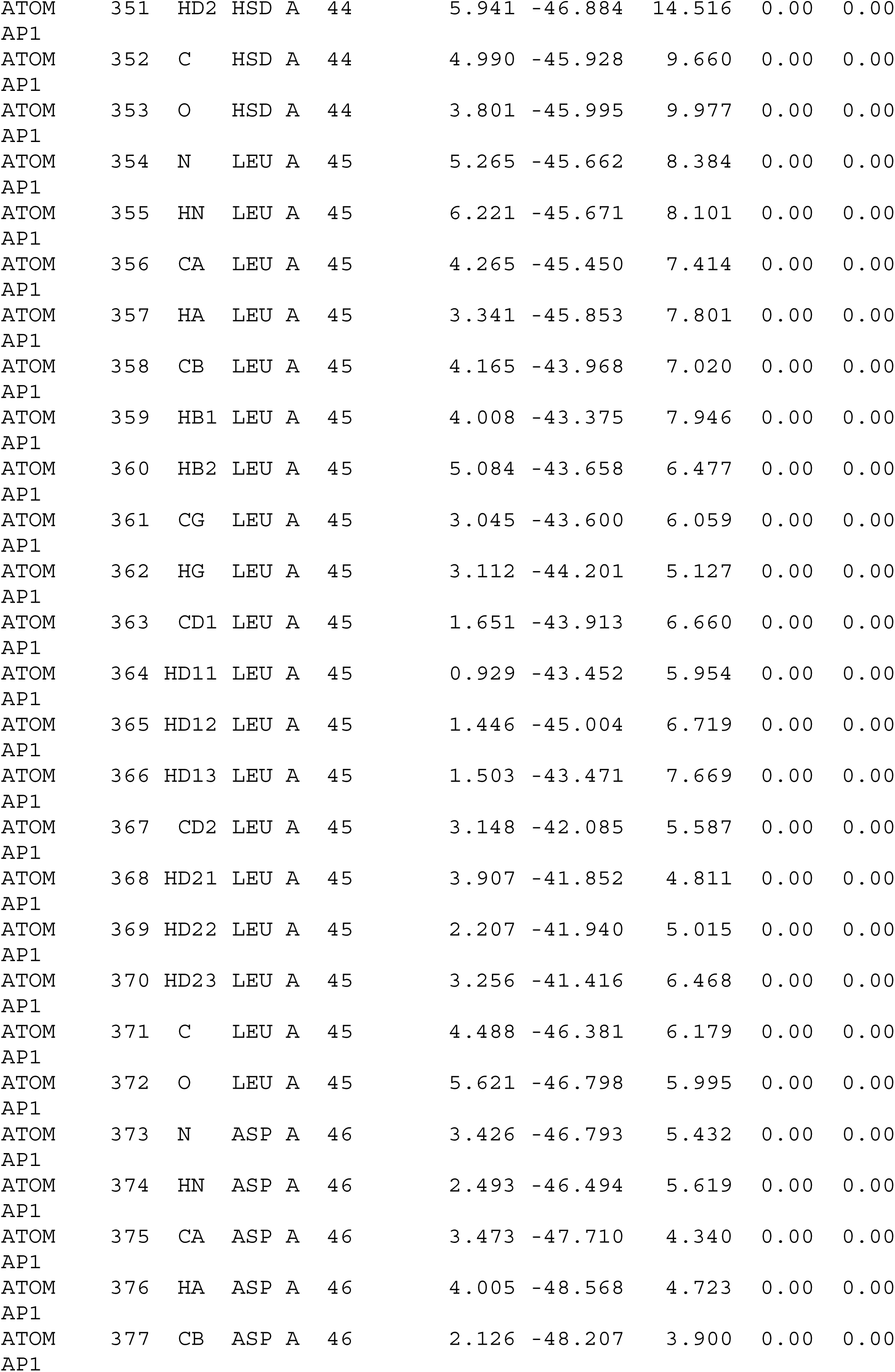

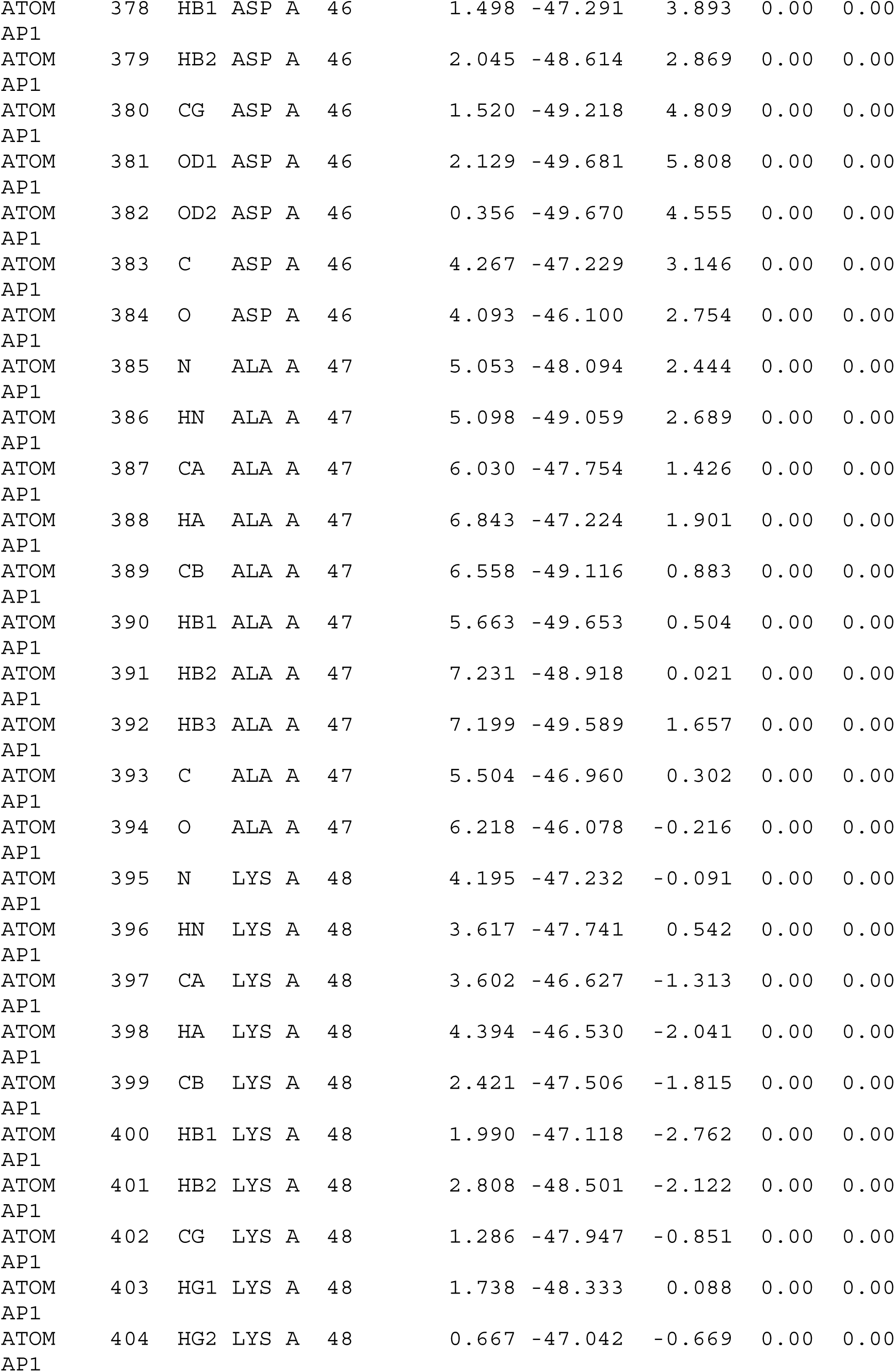

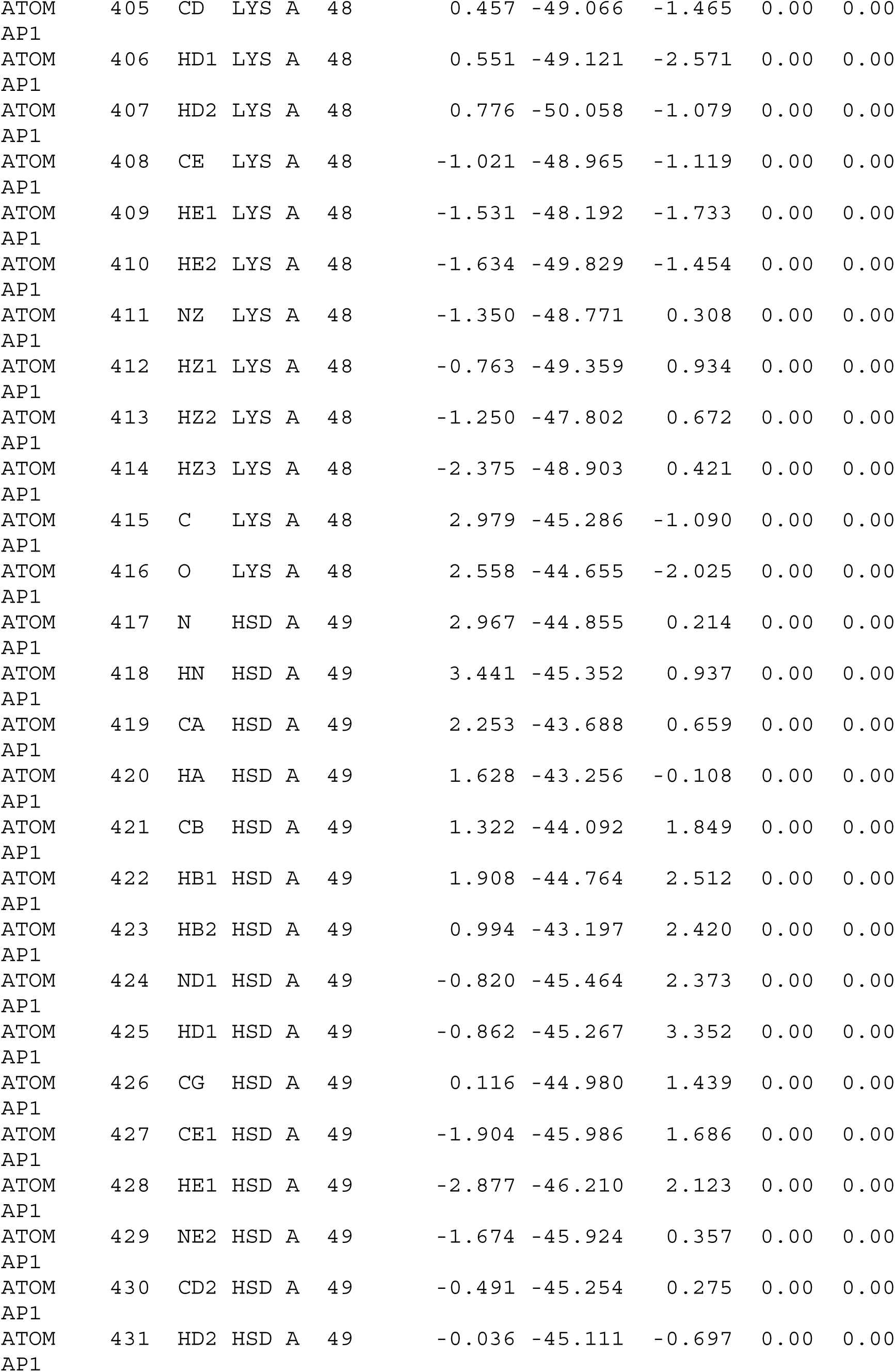

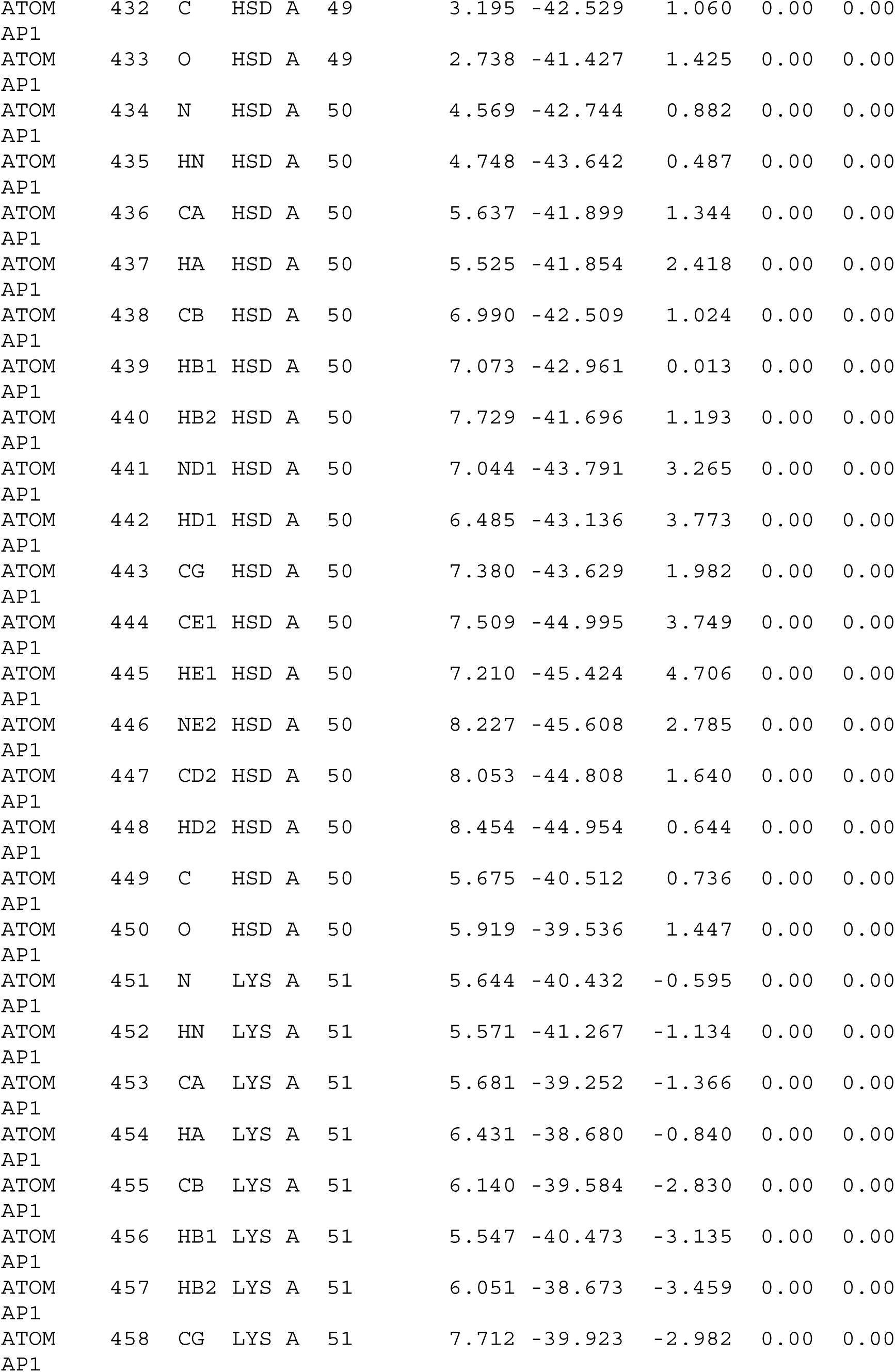

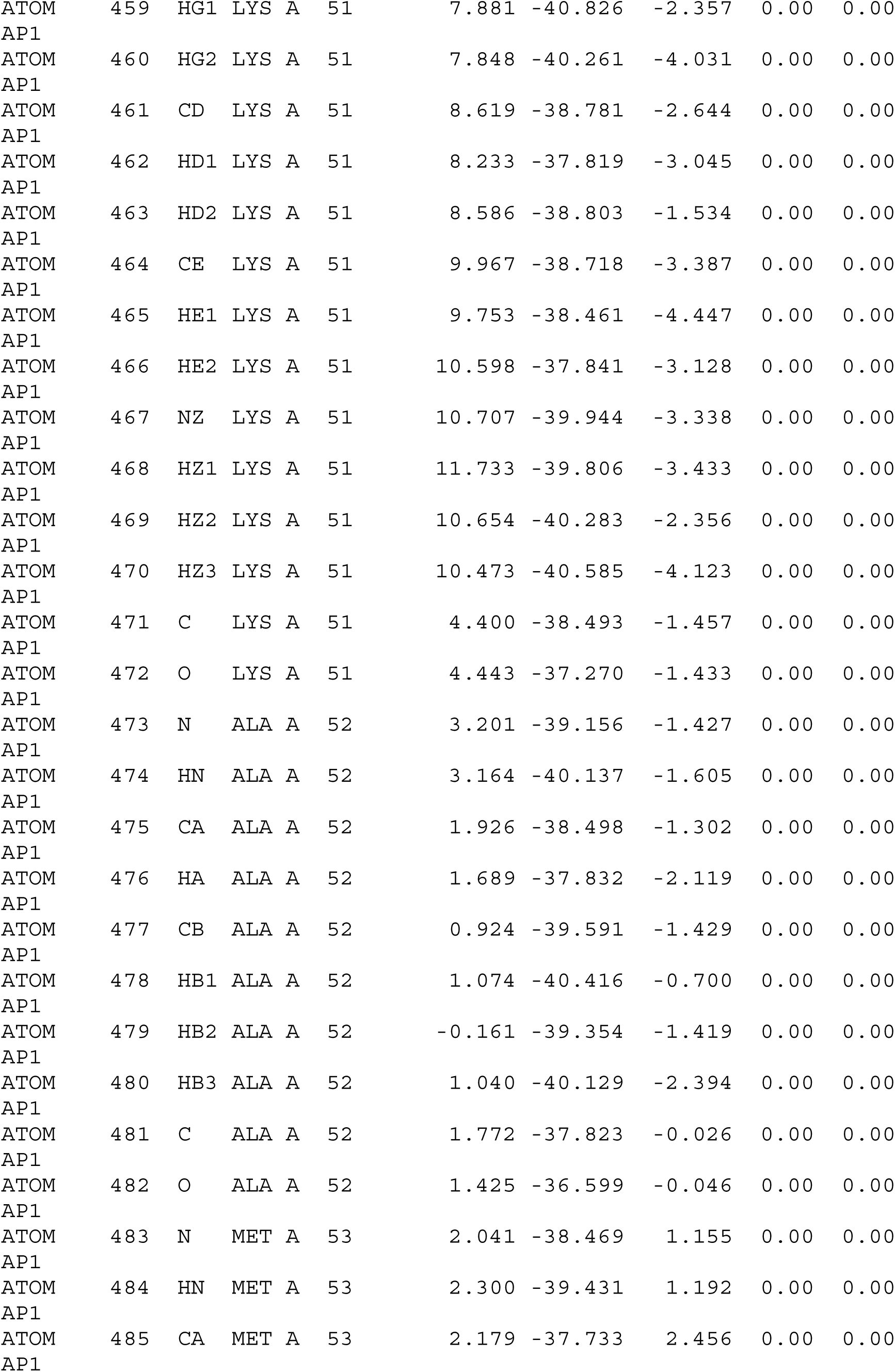

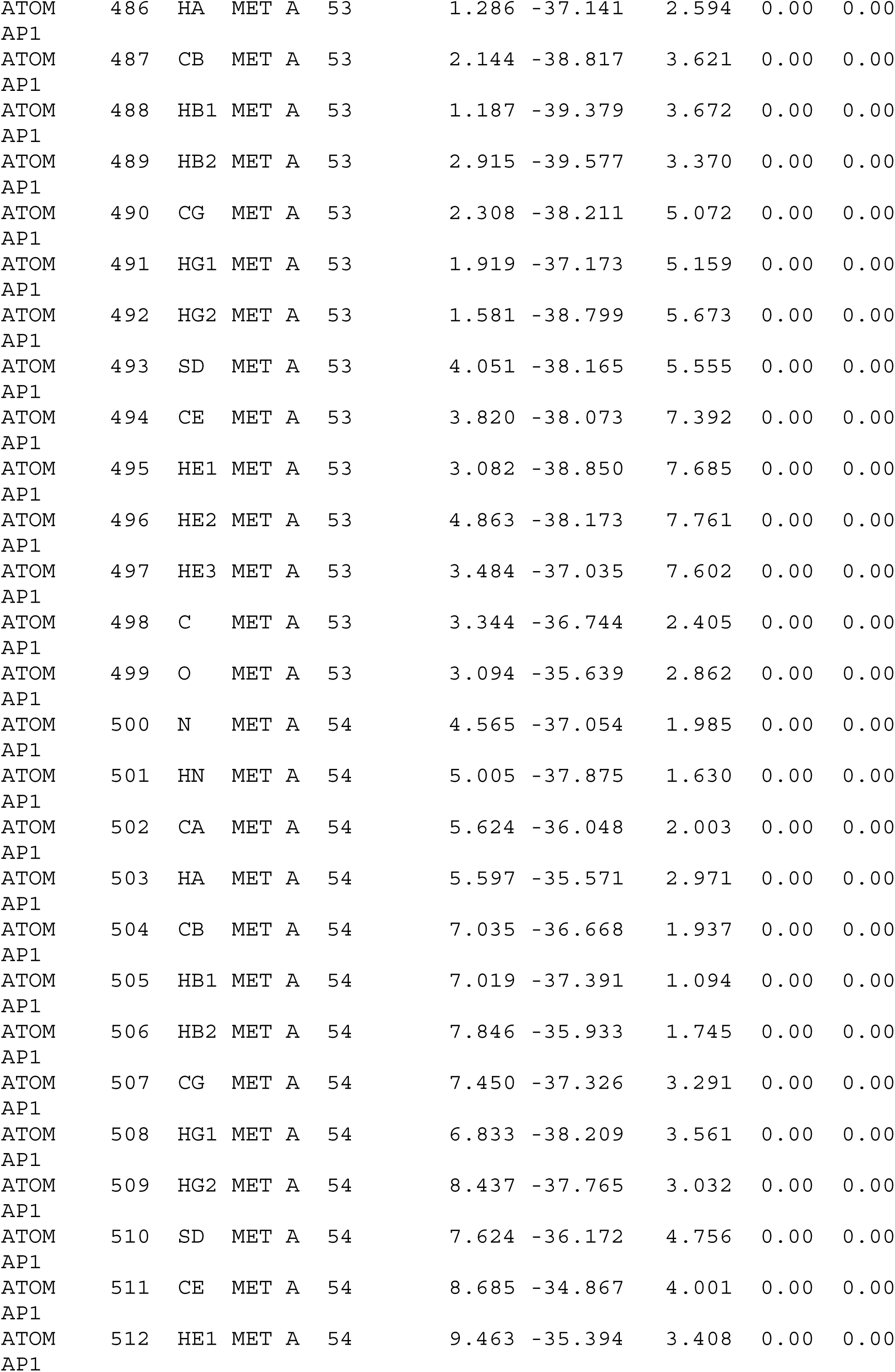

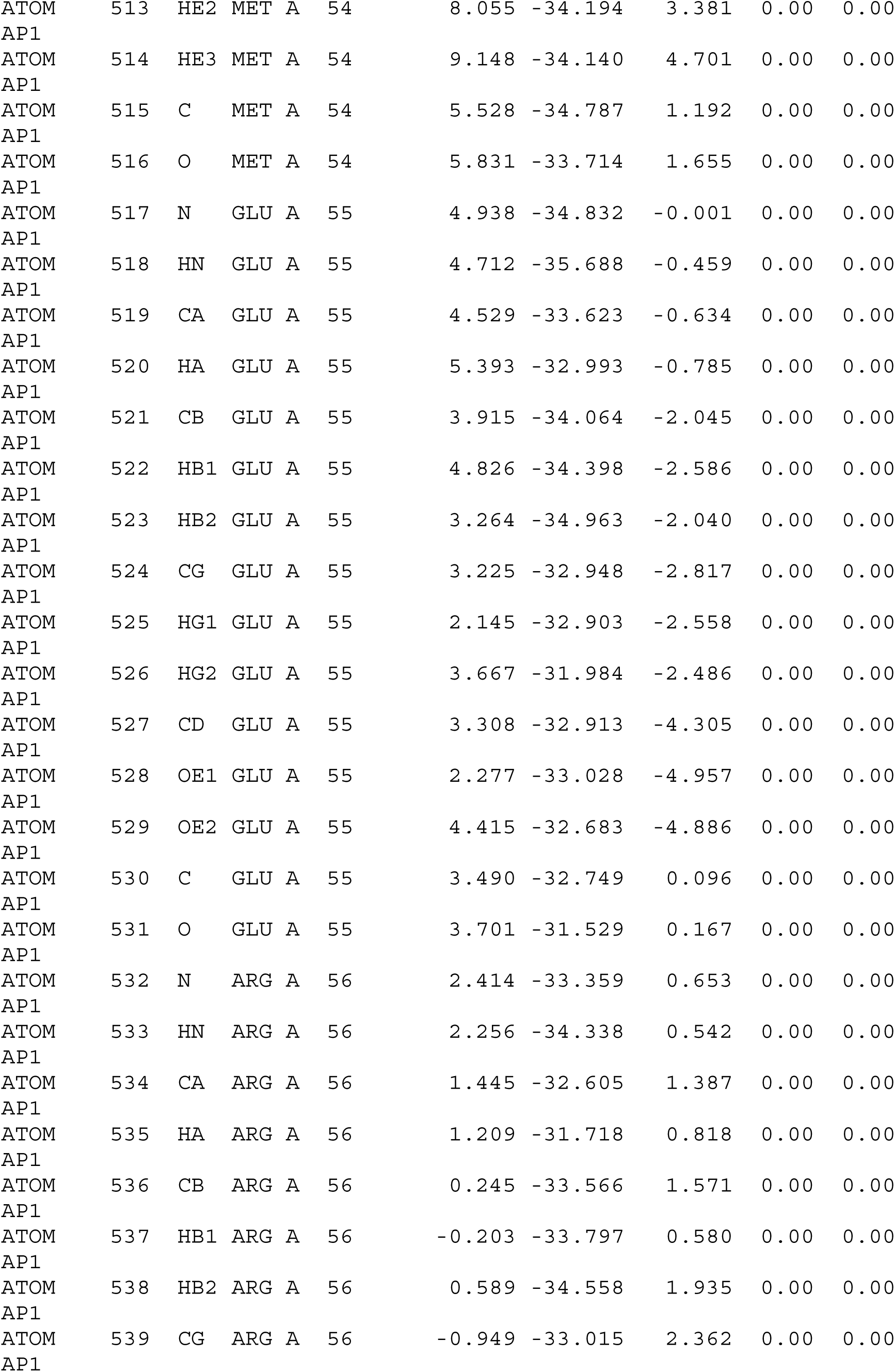

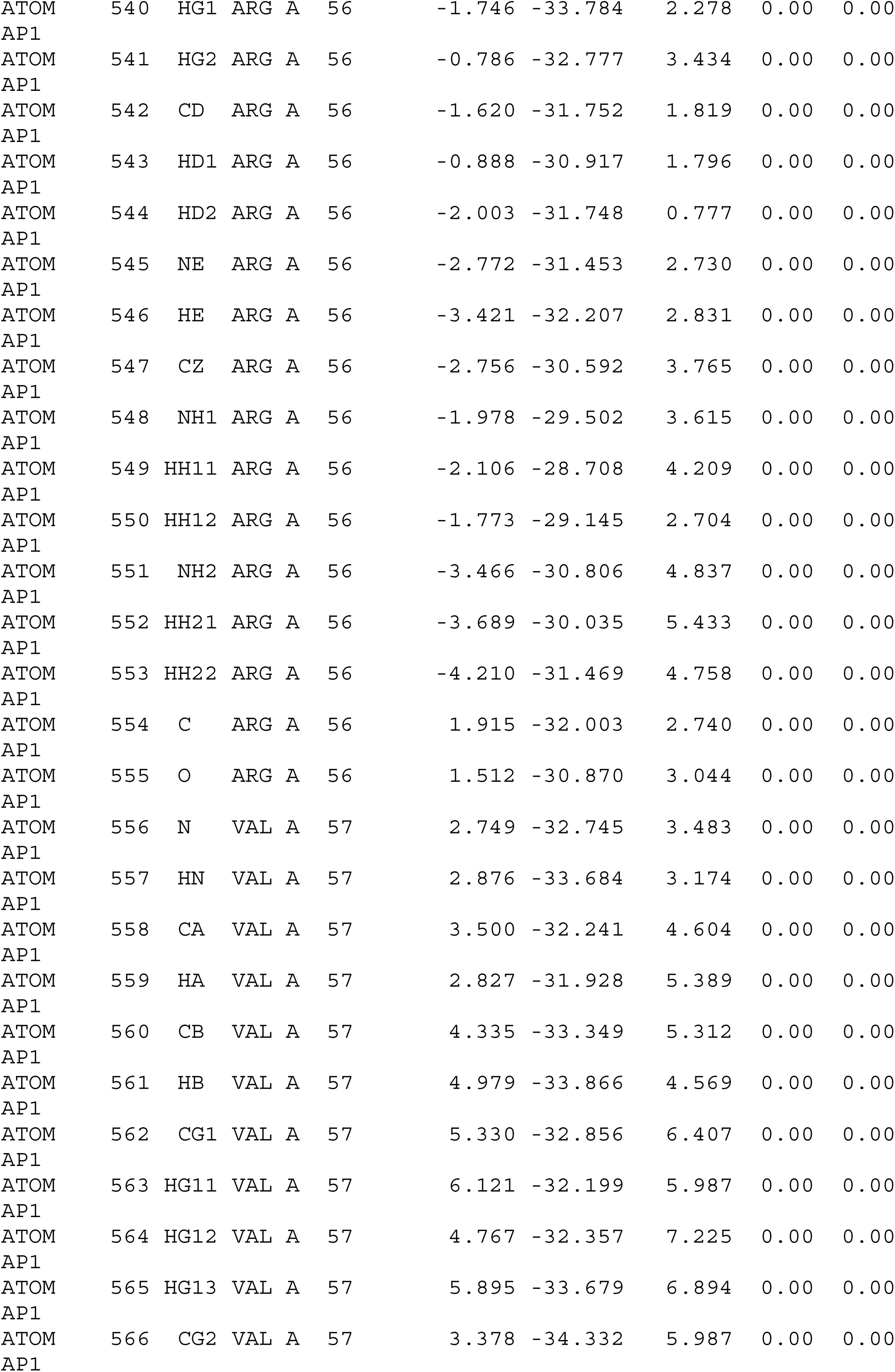

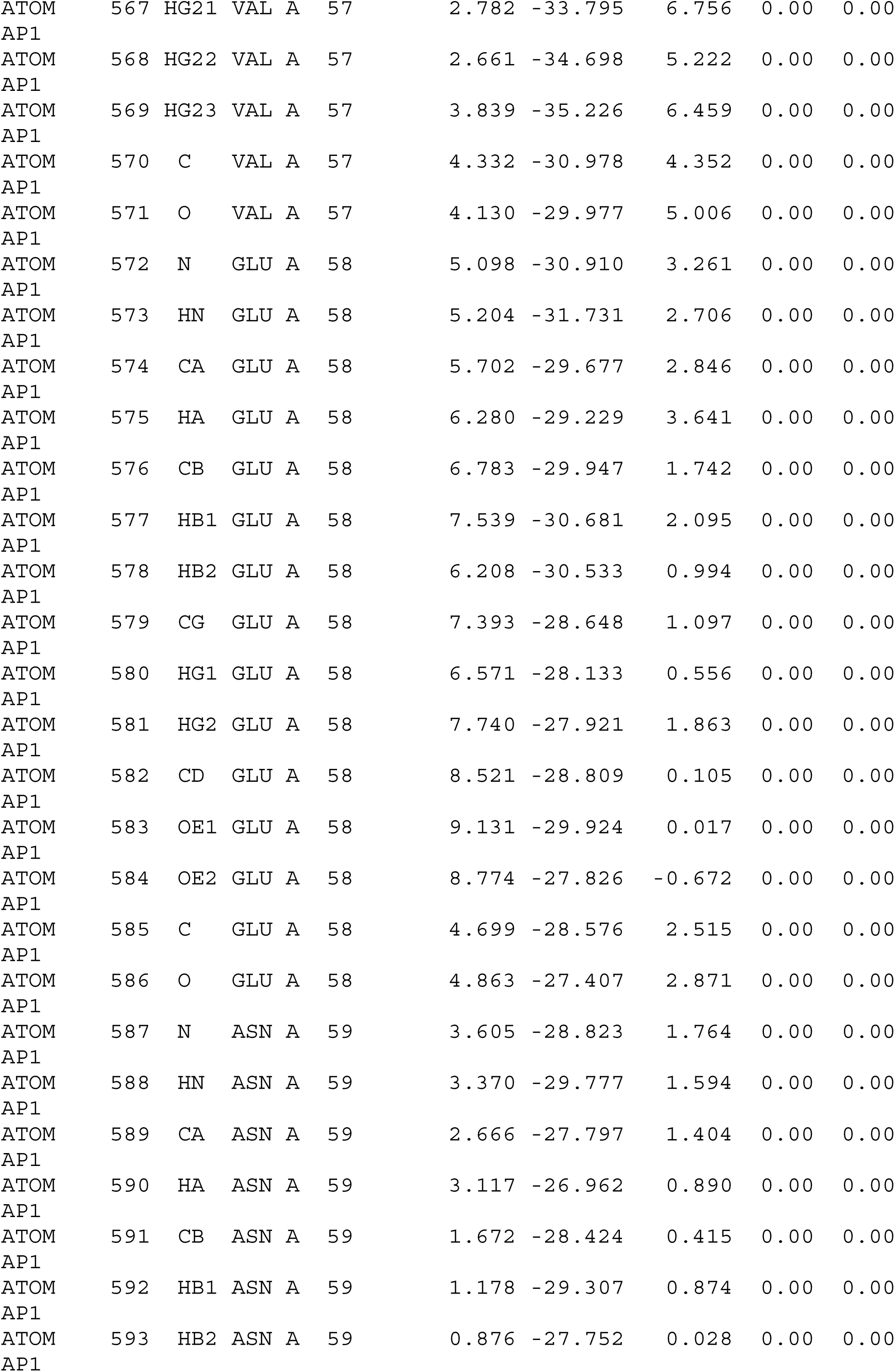

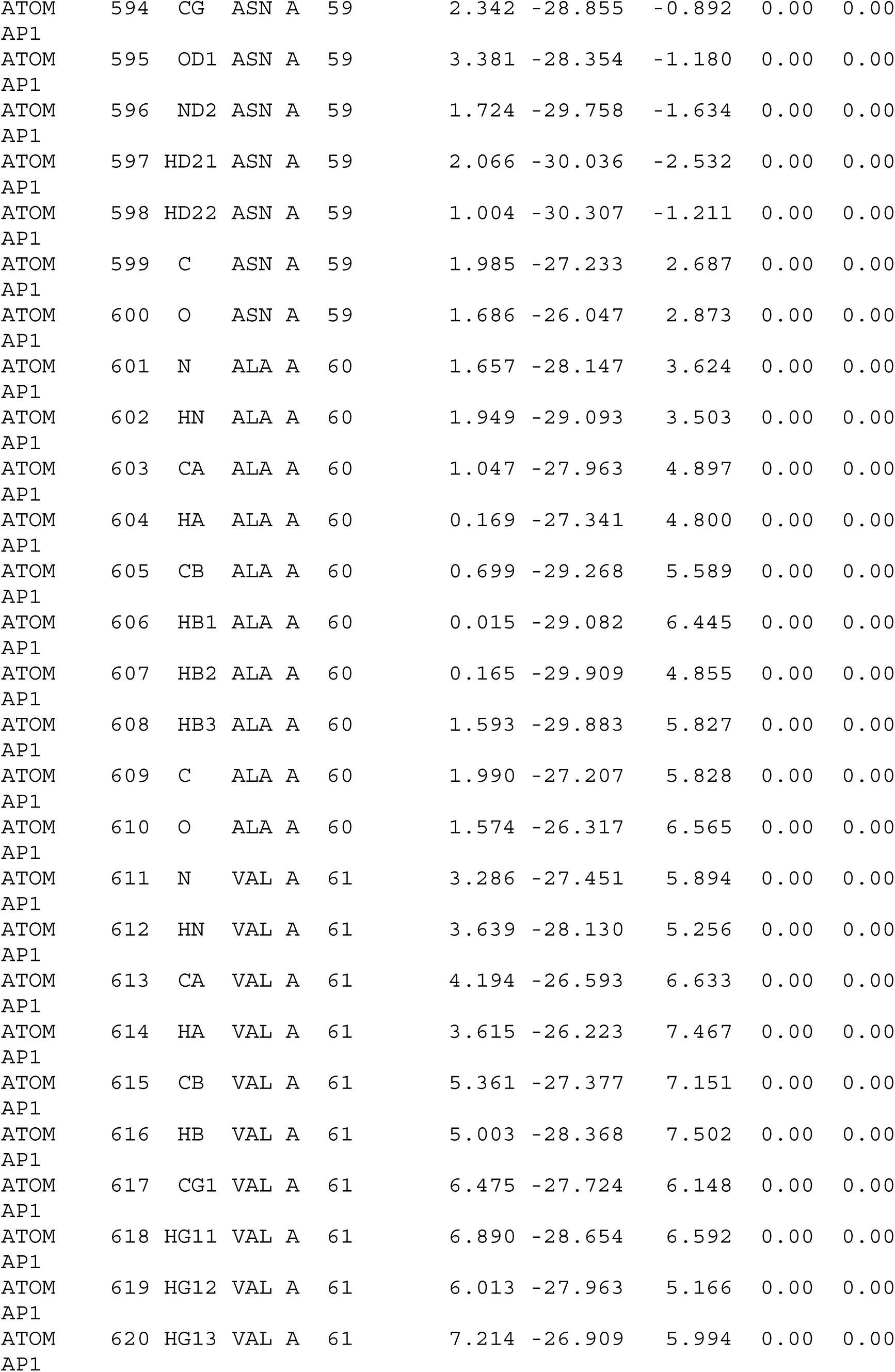

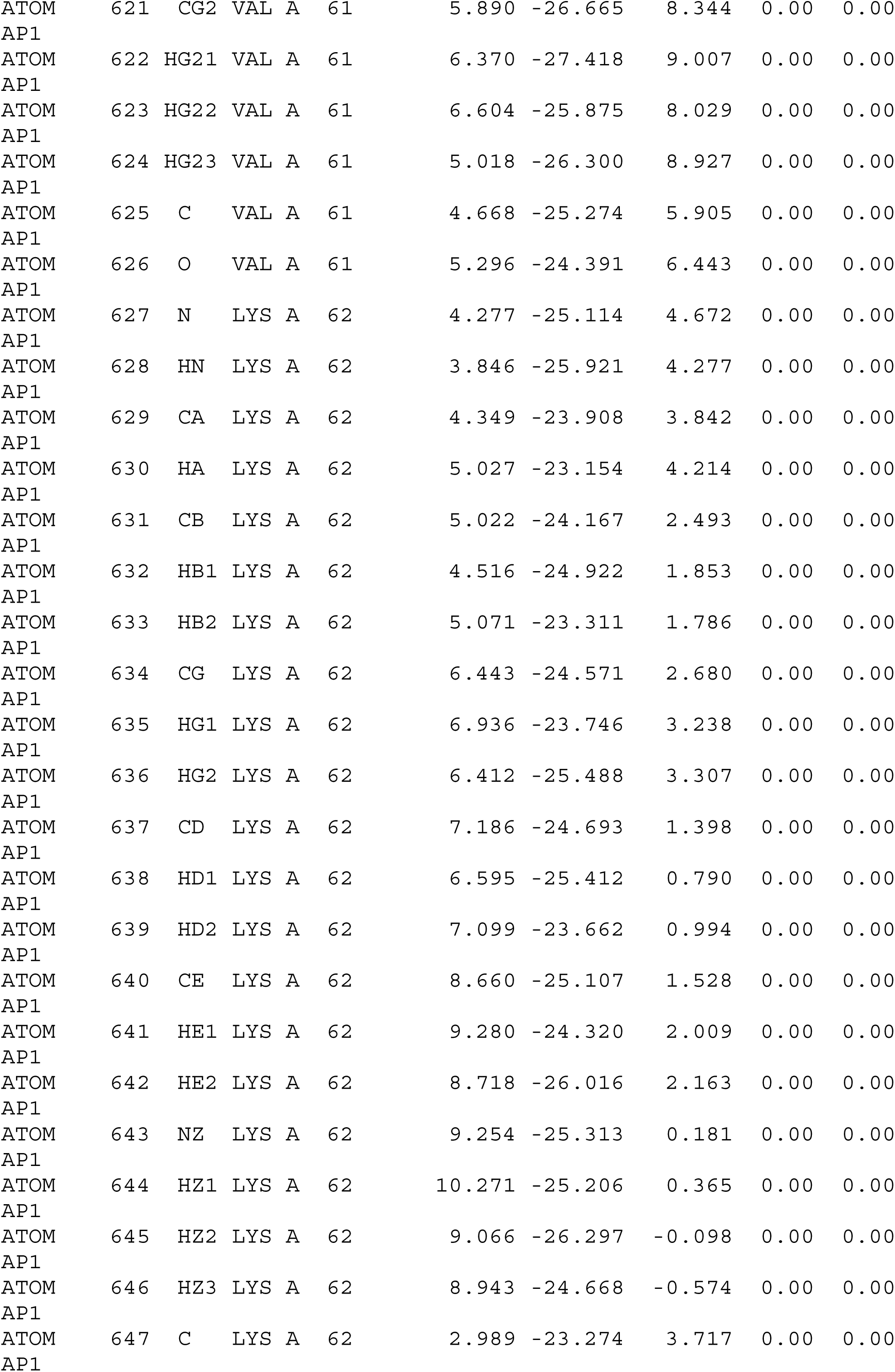

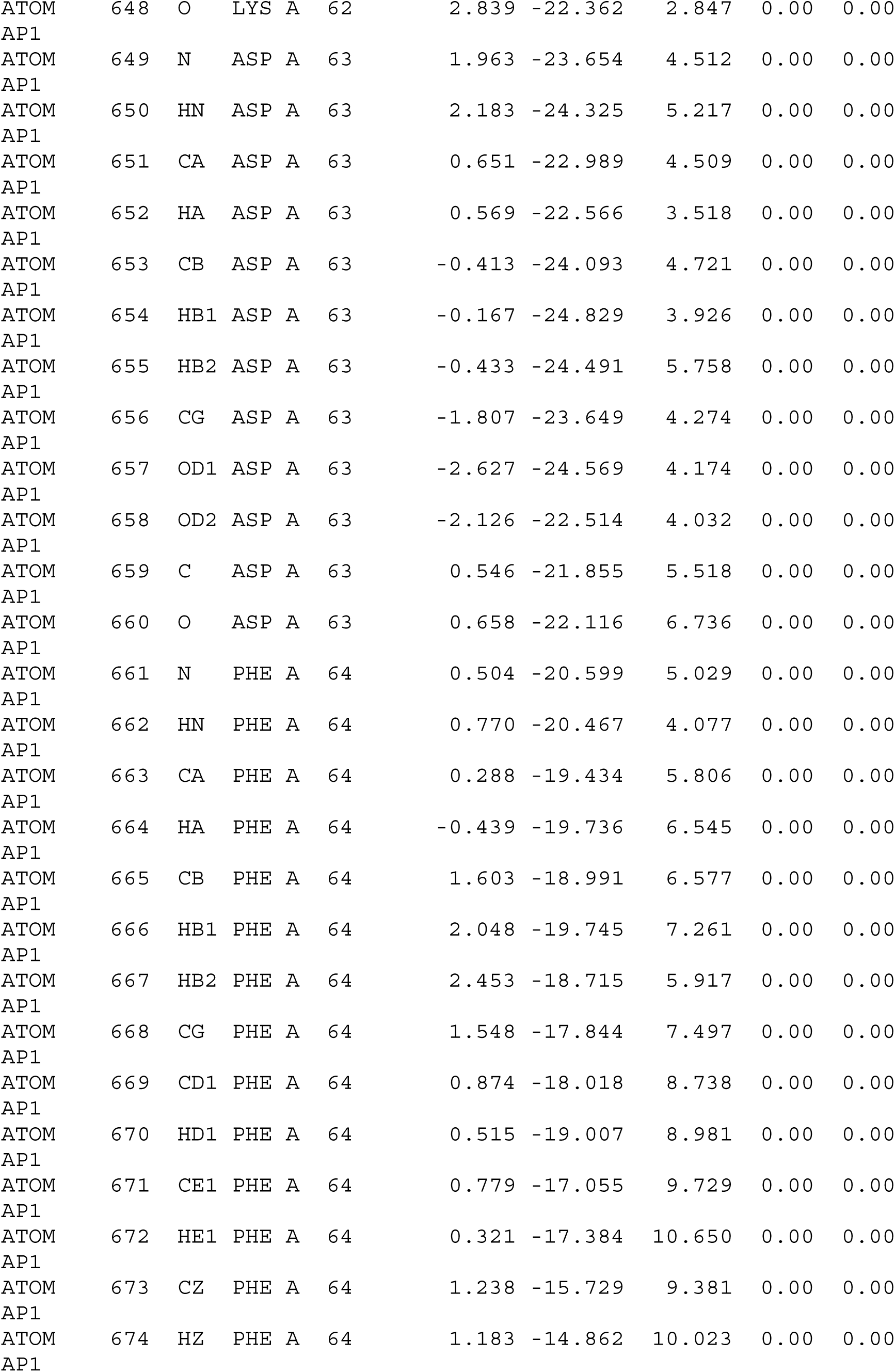

|  |  |  |  |  |  |  |  |  |  |  |  |
| --- | --- | --- | --- | --- | --- | --- | --- | --- | --- | --- | --- |
| ATOM<br>AP1 | 675 | CD2 | PHE | A | 64 |  | 2.084 | -16.538 | 7.176 | 0.00 | 0.00 |
| ATOM<br>AP1 | 676 | HD2 | PHE | A | 64 |  | 2.695 | -16.405 | 6.296 | 0.00 | 0.00 |
| ATOM<br>AP1 | 677 | CE2 | PHE | A | 64 |  | 1.869 | -15.526 | 8.116 | 0.00 | 0.00 |
| ATOM<br>AP1 | 678 | HE2 | PHE | A | 64 |  | 2.103 | -14.502 | 7.863 | 0.00 | 0.00 |
| ATOM<br>AP1 | 679 | C | PHE | A | 64 |  | -0.461 | -18.236 | 5.081 | 0.00 | 0.00 |
| ATOM<br>AP1 | 680 | O | PHE | A | 64 |  | 0.111 | -17.257 | 4.606 | 0.00 | 0.00 |
| ATOM<br>AP1 | 681 | N | GLN | A | 65 |  | -1.774 | -18.262 | 4.918 | 0.00 | 0.00 |
| ATOM<br>AP1 | 682 | HN | GLN | A | 65 |  | -2.134 | -19.178 | 5.079 | 0.00 | 0.00 |
| ATOM<br>AP1 | 683 | CA | GLN | A | 65 |  | -2.576 | -17.258 | 4.258 | 0.00 | 0.00 |
| ATOM<br>AP1 | 684 | HA | GLN | A | 65 |  | -2.061 | -16.312 | 4.185 | 0.00 | 0.00 |
| ATOM<br>AP1 | 685 | CB | GLN | A | 65 |  | -3.089 | -17.731 | 2.821 | 0.00 | 0.00 |
| ATOM<br>AP1 | 686 | HB1 | GLN | A | 65 |  | -3.224 | -18.831 | 2.907 | 0.00 | 0.00 |
| ATOM<br>AP1 | 687 | HB2 | GLN | A | 65 |  | -3.980 | -17.141 | 2.520 | 0.00 | 0.00 |
| ATOM<br>AP1 | 688 | CG | GLN | A | 65 |  | -2.016 | -17.698 | 1.712 | 0.00 | 0.00 |
| ATOM<br>AP1 | 689 | HG1 | GLN | A | 65 |  | -1.300 | -18.502 | 1.987 | 0.00 | 0.00 |
| ATOM<br>AP1 | 690 | HG2 | GLN | A | 65 |  | -2.432 | -17.892 | 0.700 | 0.00 | 0.00 |
| ATOM<br>AP1 | 691 | CD | GLN | A | 65 |  | -1.348 | -16.332 | 1.539 | 0.00 | 0.00 |
| ATOM<br>AP1 | 692 | OE1 | GLN | A | 65 |  | -0.147 | -16.198 | 1.905 | 0.00 | 0.00 |
| ATOM<br>AP1 | 693 | NE2 | GLN | A | 65 |  | -2.033 | -15.309 | 1.091 | 0.00 | 0.00 |
| ATOM<br>AP1 | 694 | HE21 | GLN | A | 65 |  | -1.518 | -14.462 | 1.228 | 0.00 | 0.00 |
| ATOM<br>AP1 | 695 | HE22 | GLN | A | 65 |  | -2.914 | -15.518 | 0.667 | 0.00 | 0.00 |
| ATOM<br>AP1 | 696 | C | GLN | A | 65 |  | -3.753 | -16.803 | 5.093 | 0.00 | 0.00 |
| ATOM<br>AP1 | 697 | O | GLN | A | 65 |  | -4.587 | -16.042 | 4.637 | 0.00 | 0.00 |
| ATOM<br>AP1 | 698 | N | GLU | A | 66 |  | -3.733 | -17.238 | 6.399 | 0.00 | 0.00 |
| ATOM<br>AP1 | 699 | HN | GLU | A | 66 |  | -2.897 | -17.628 | 6.777 | 0.00 | 0.00 |
| ATOM<br>AP1 | 700 | CA | GLU | A | 66 |  | -4.663 | -16.811 | 7.466 | 0.00 | 0.00 |
| ATOM<br>AP1 | 701 | HA | GLU | A | 66 |  | -5.580 | -17.342 | 7.257 | 0.00 | 0.00 |

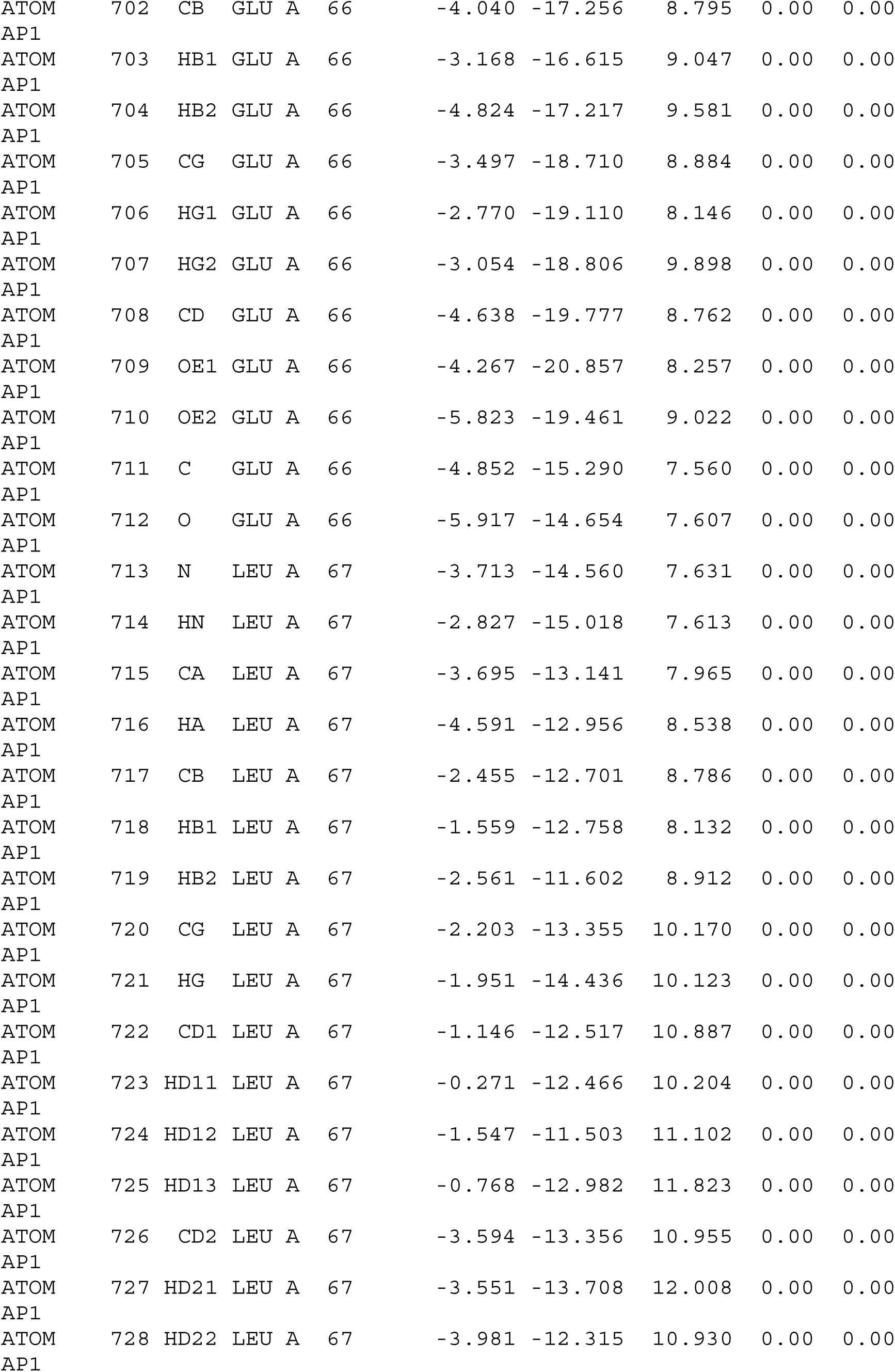

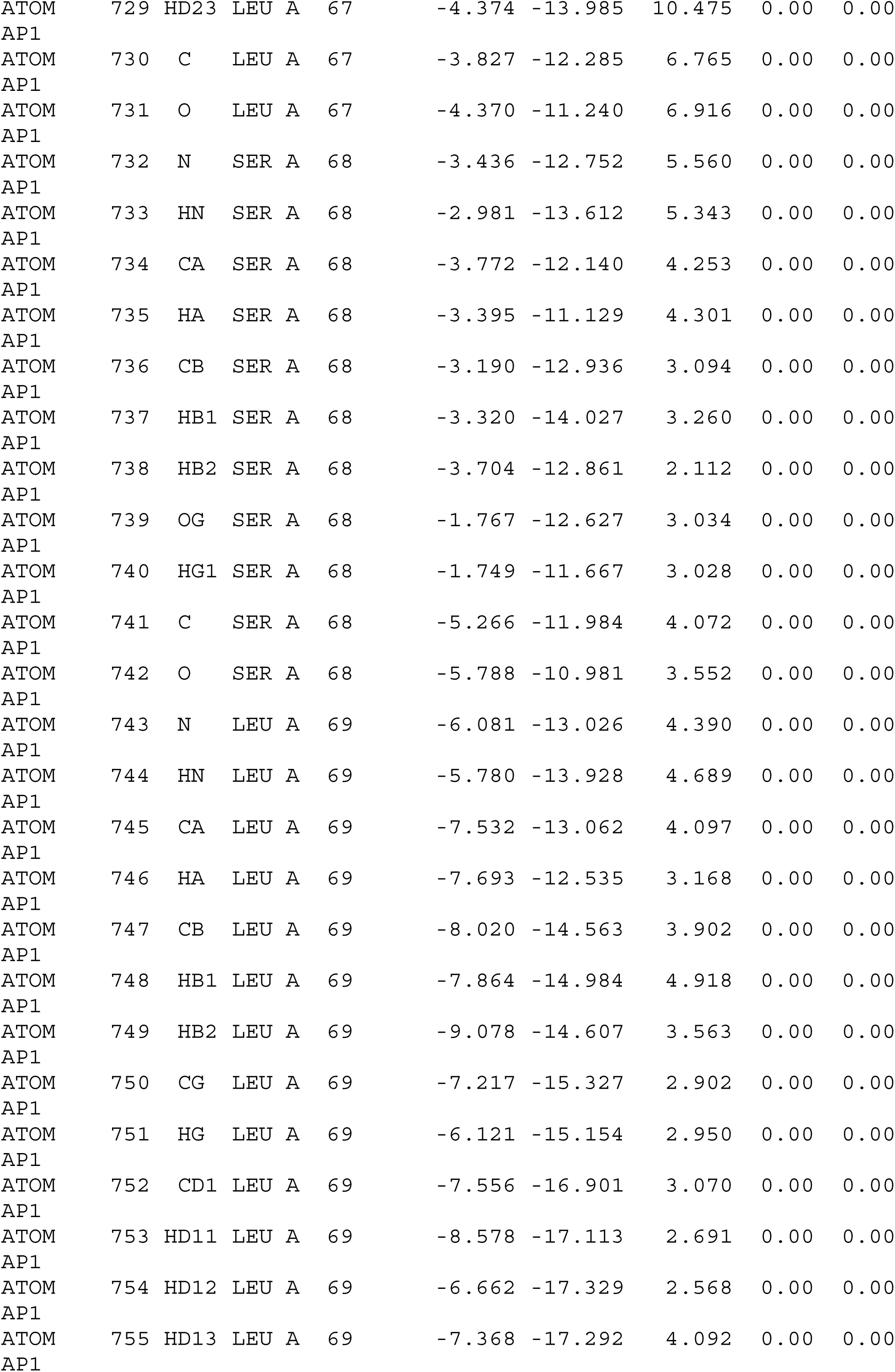

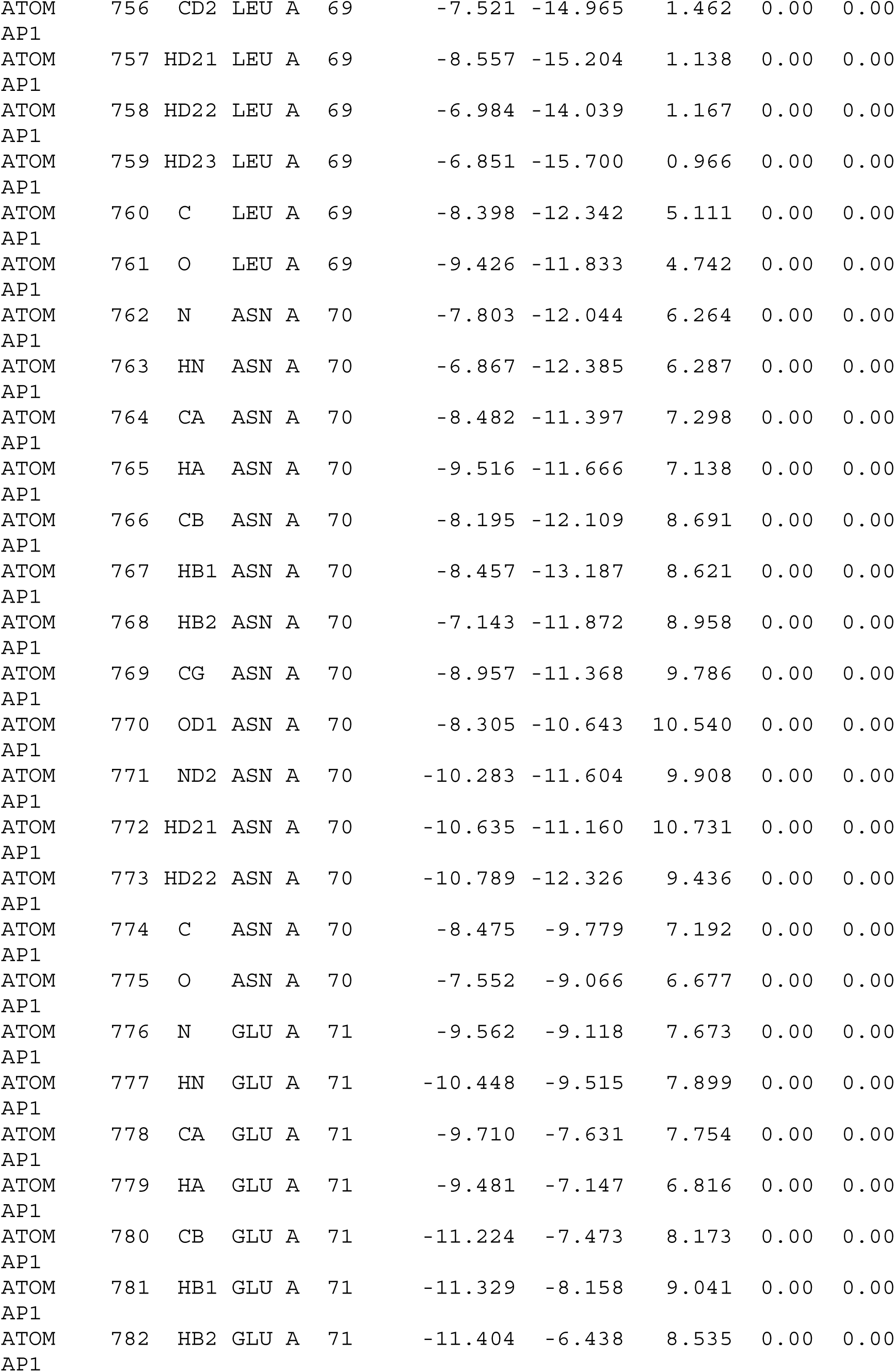

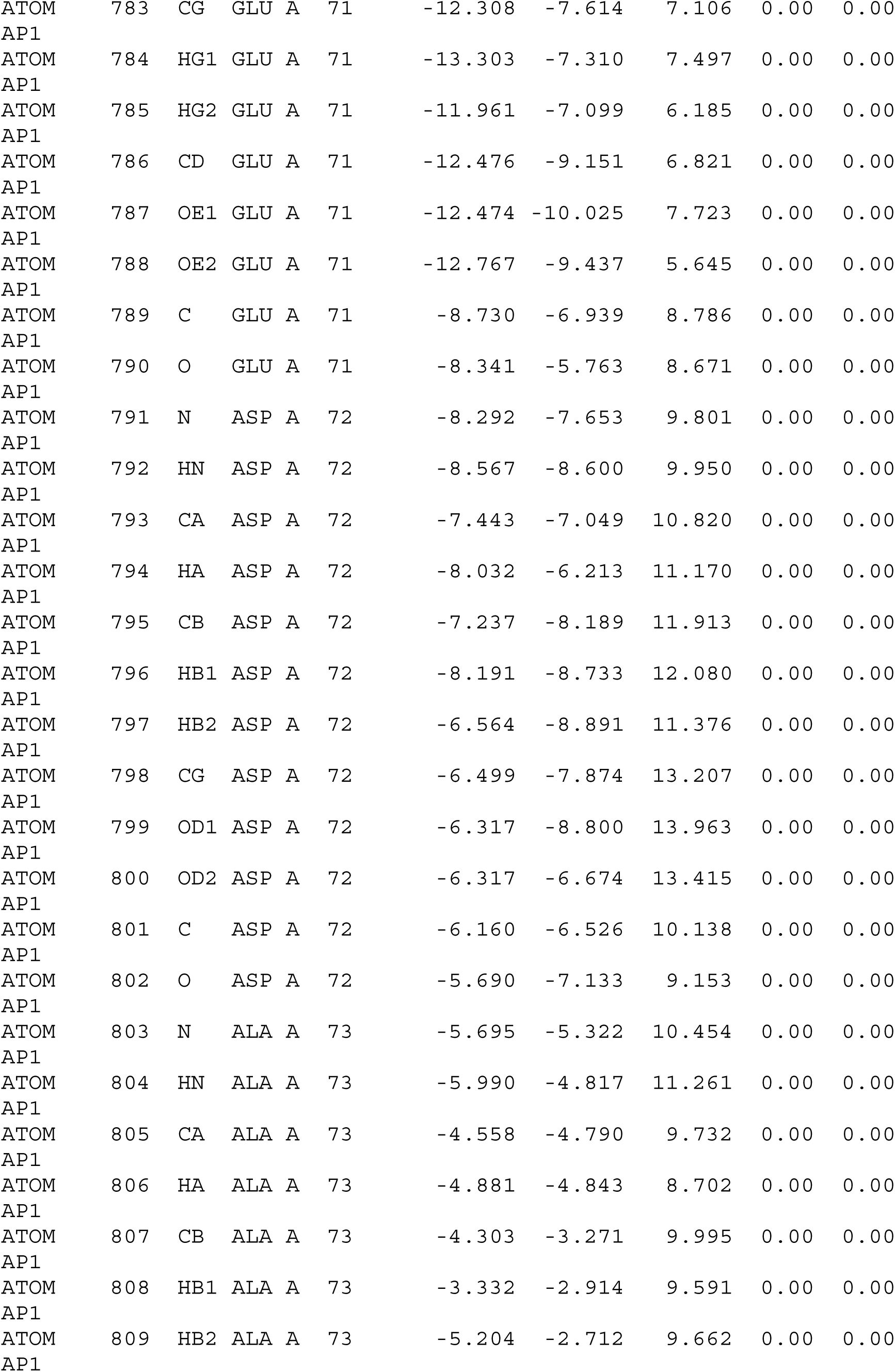

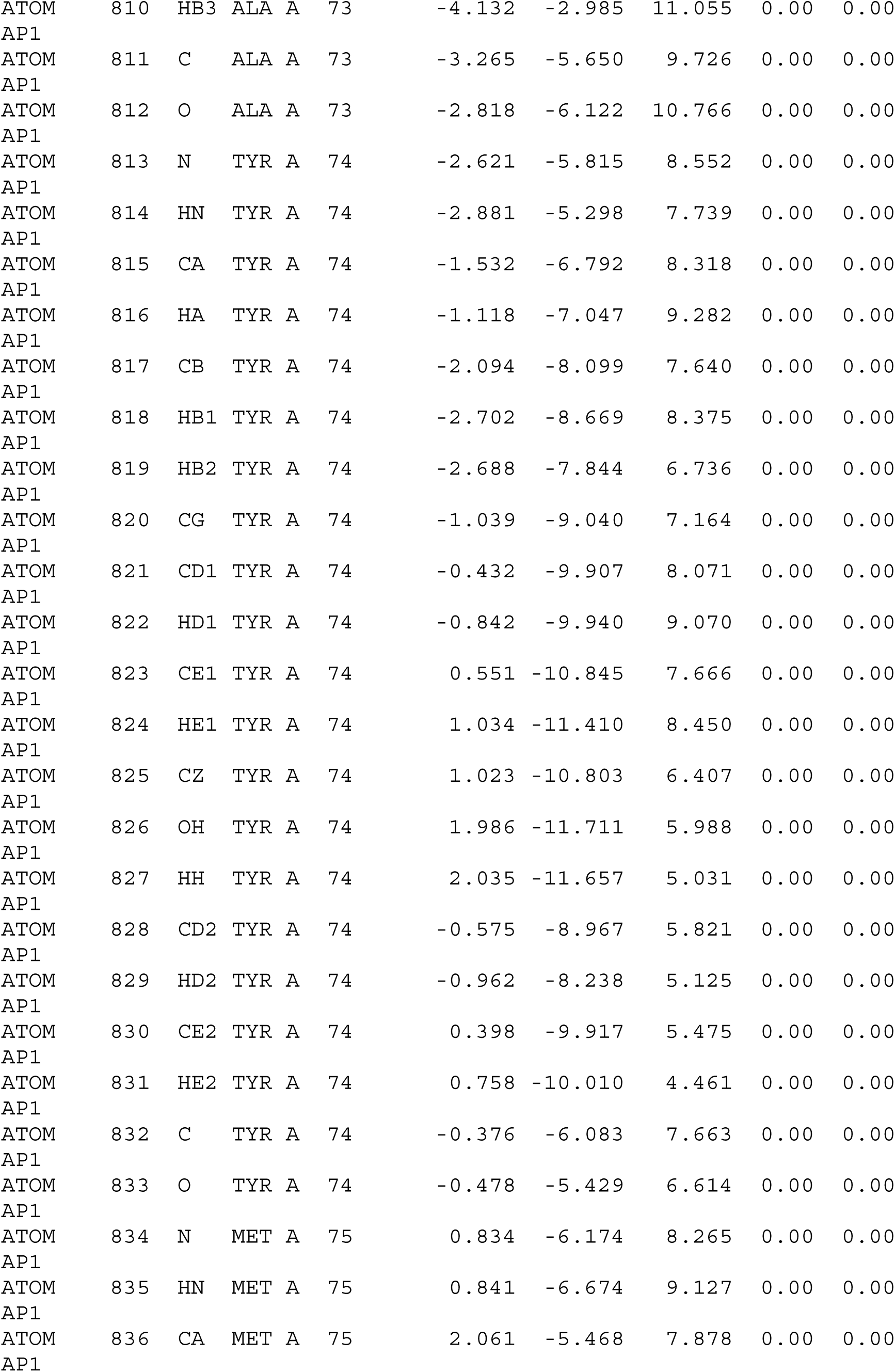

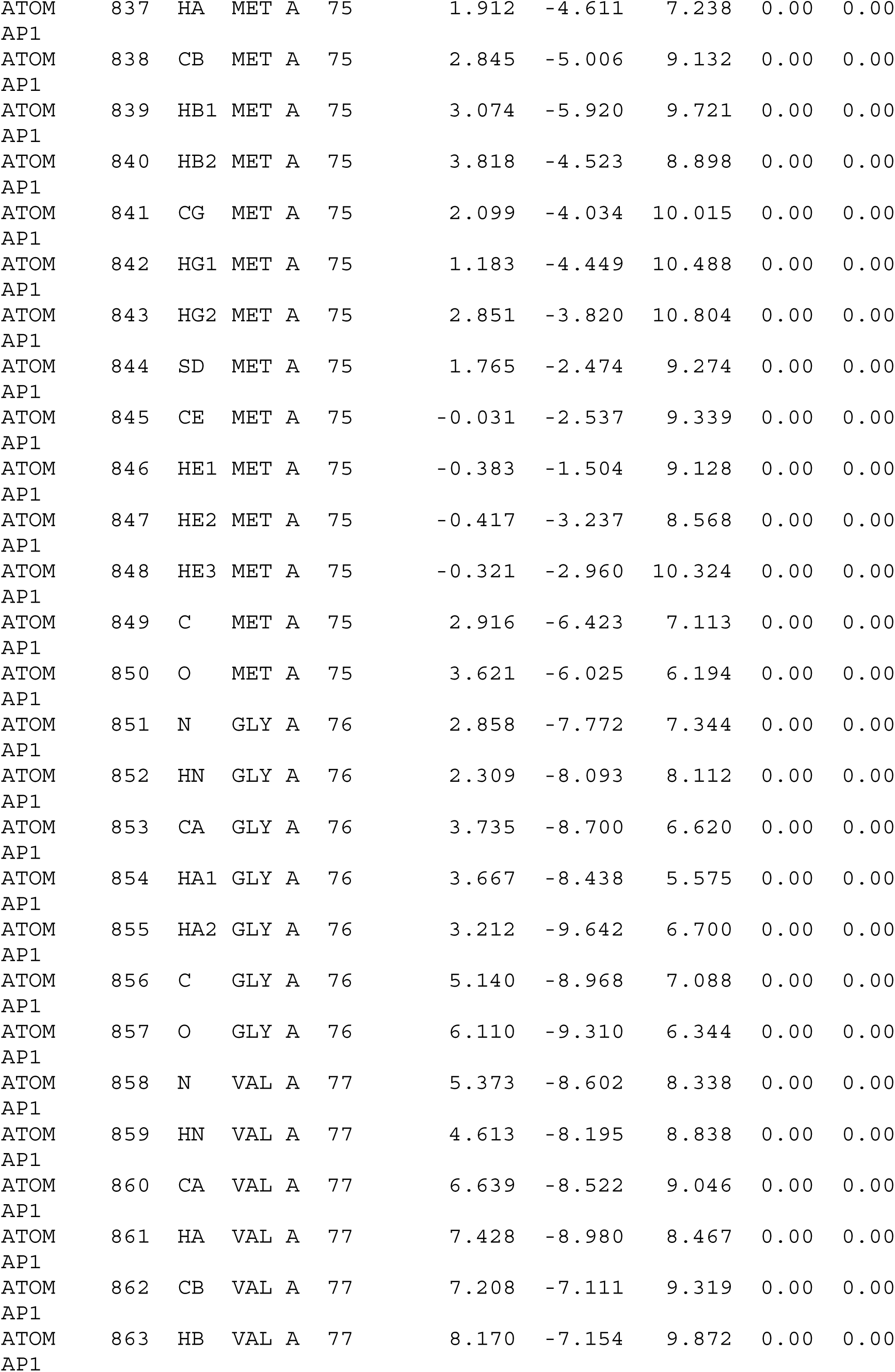

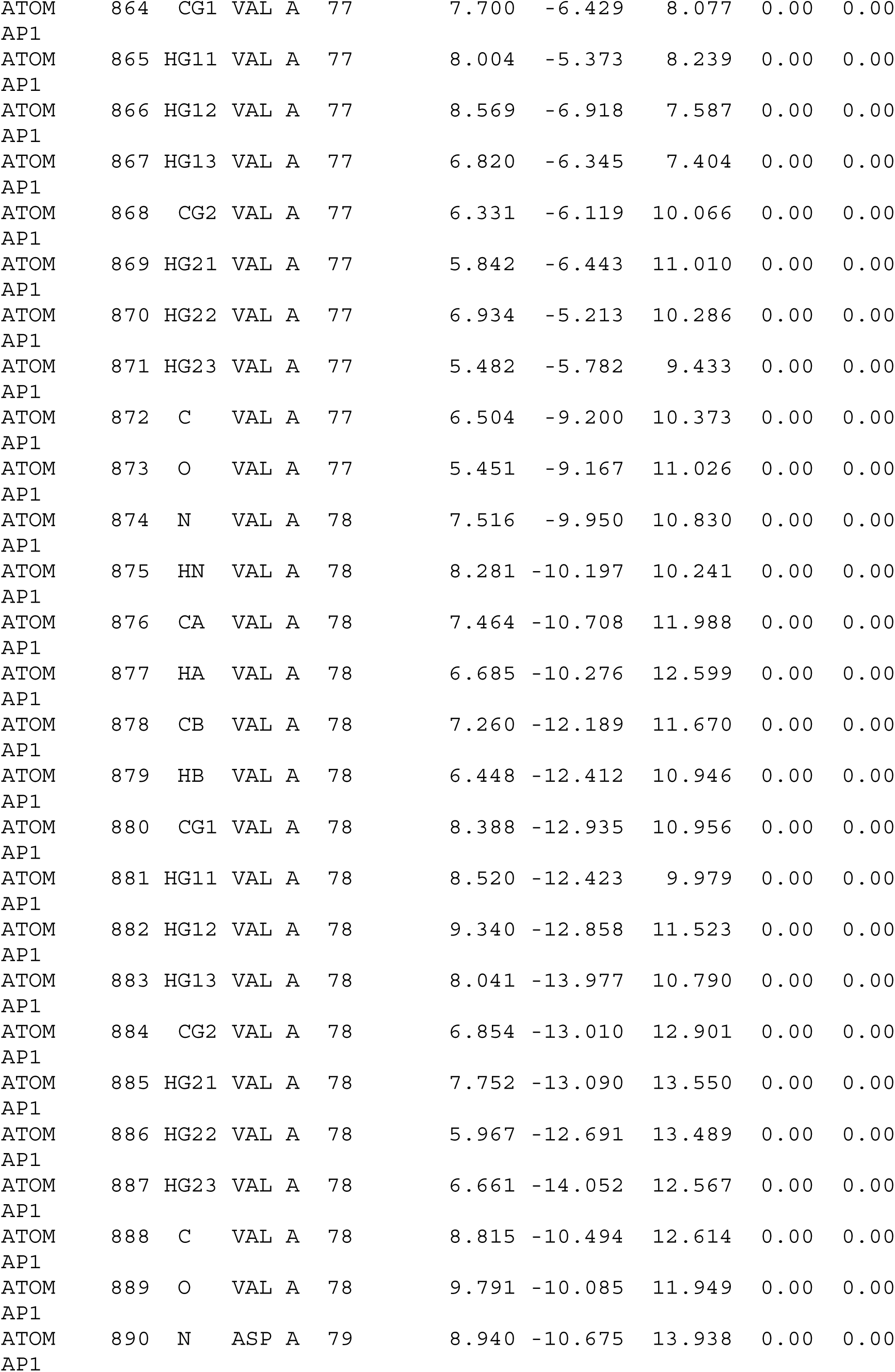

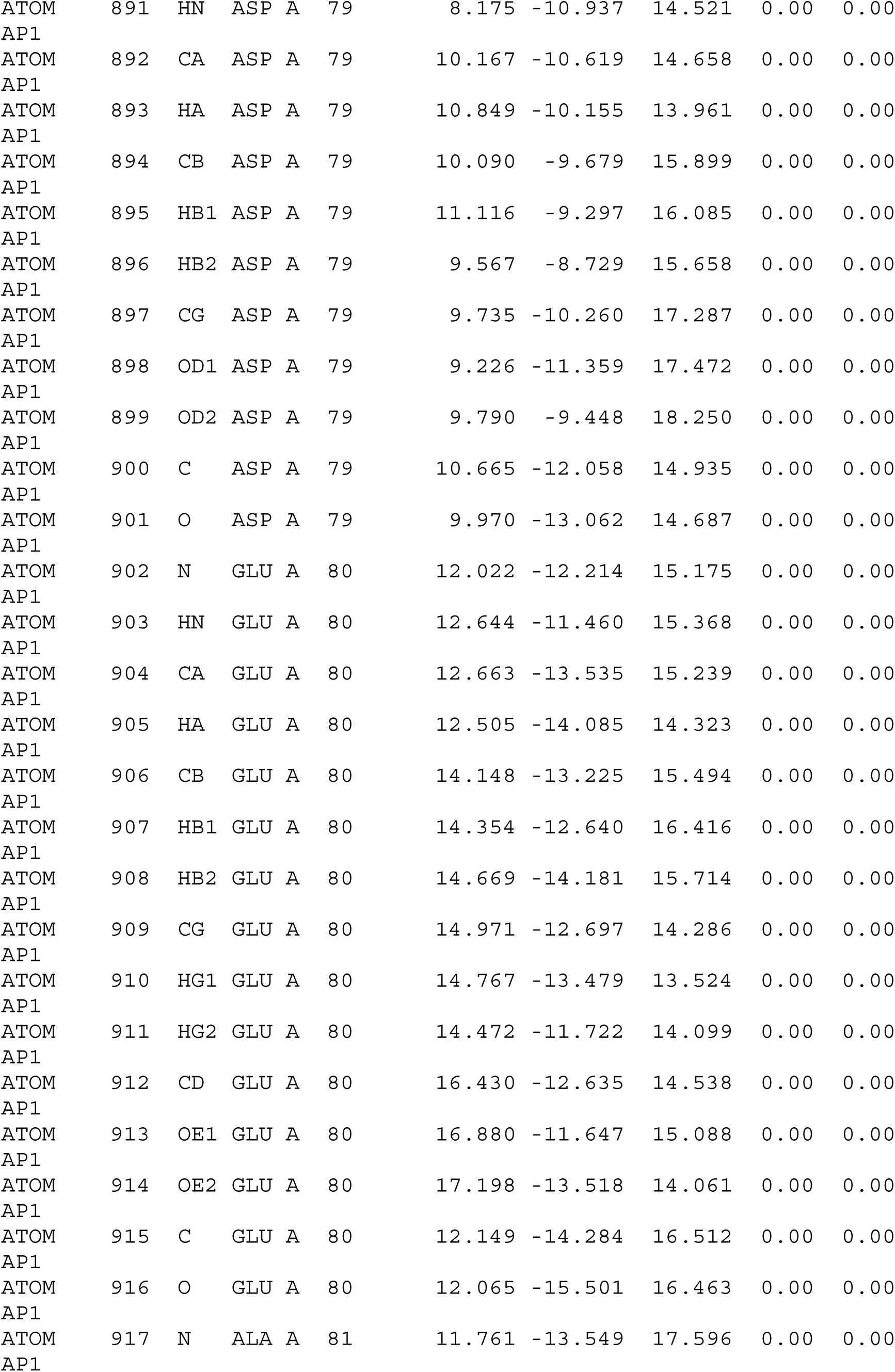

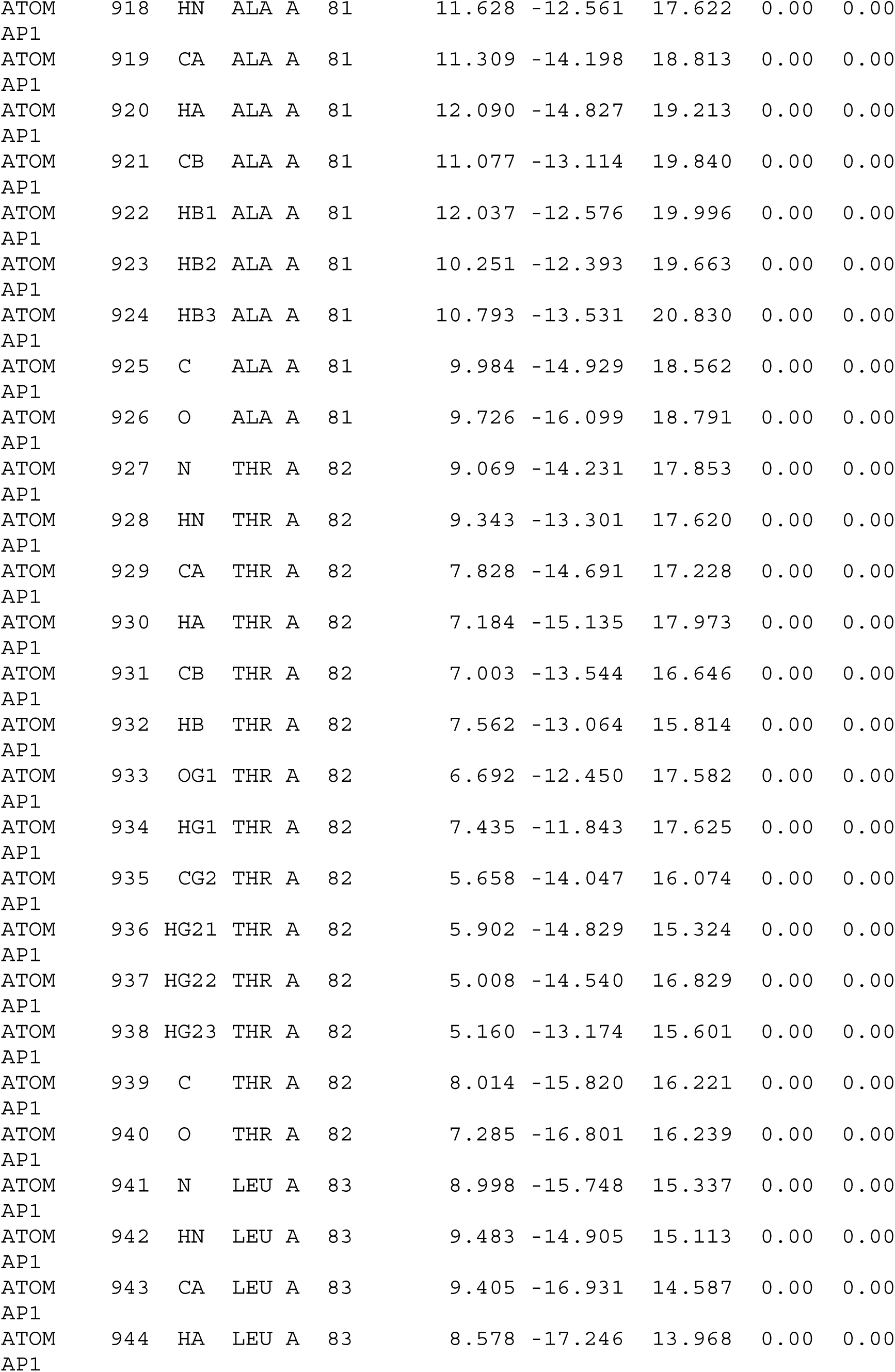

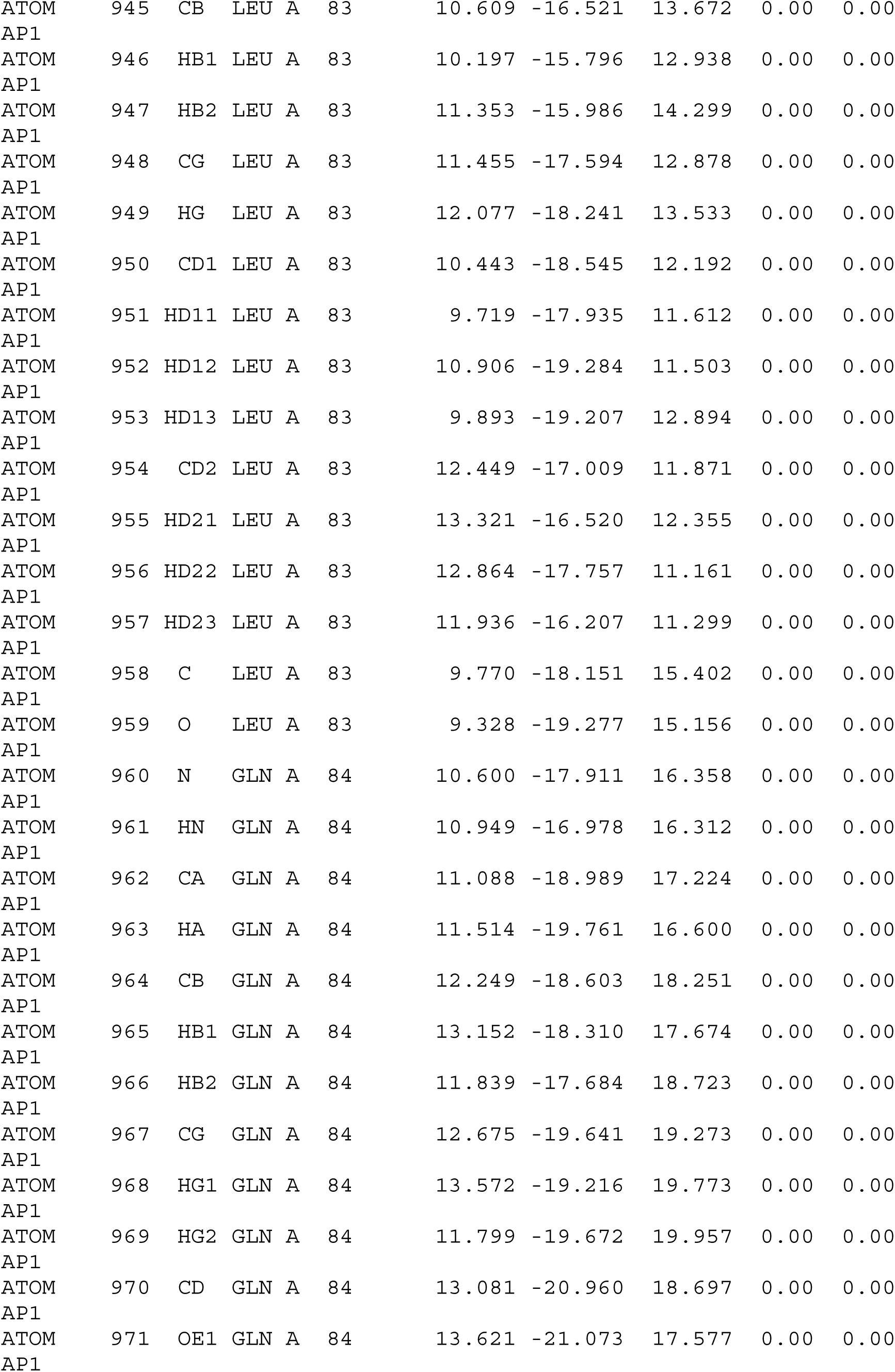

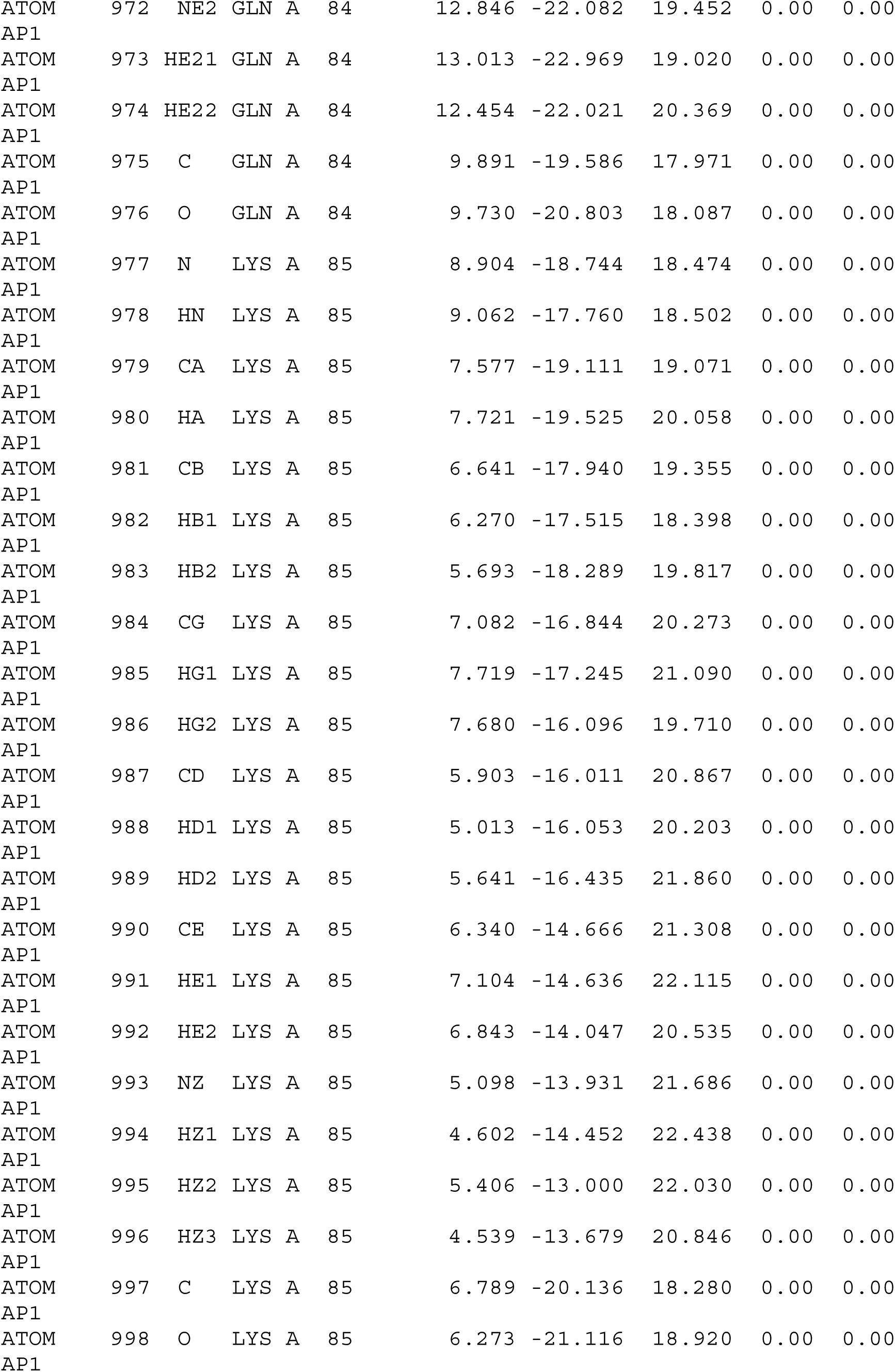

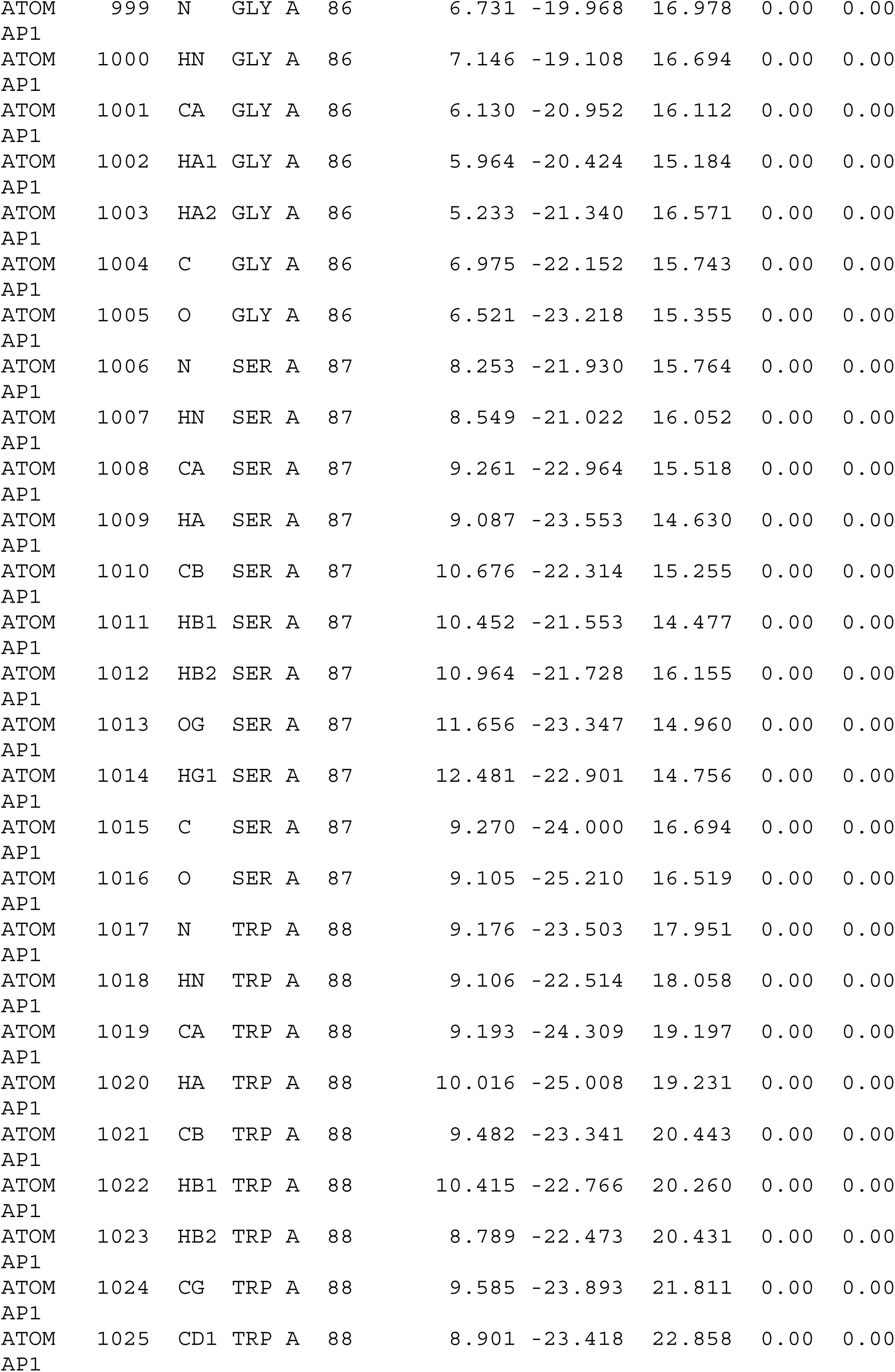

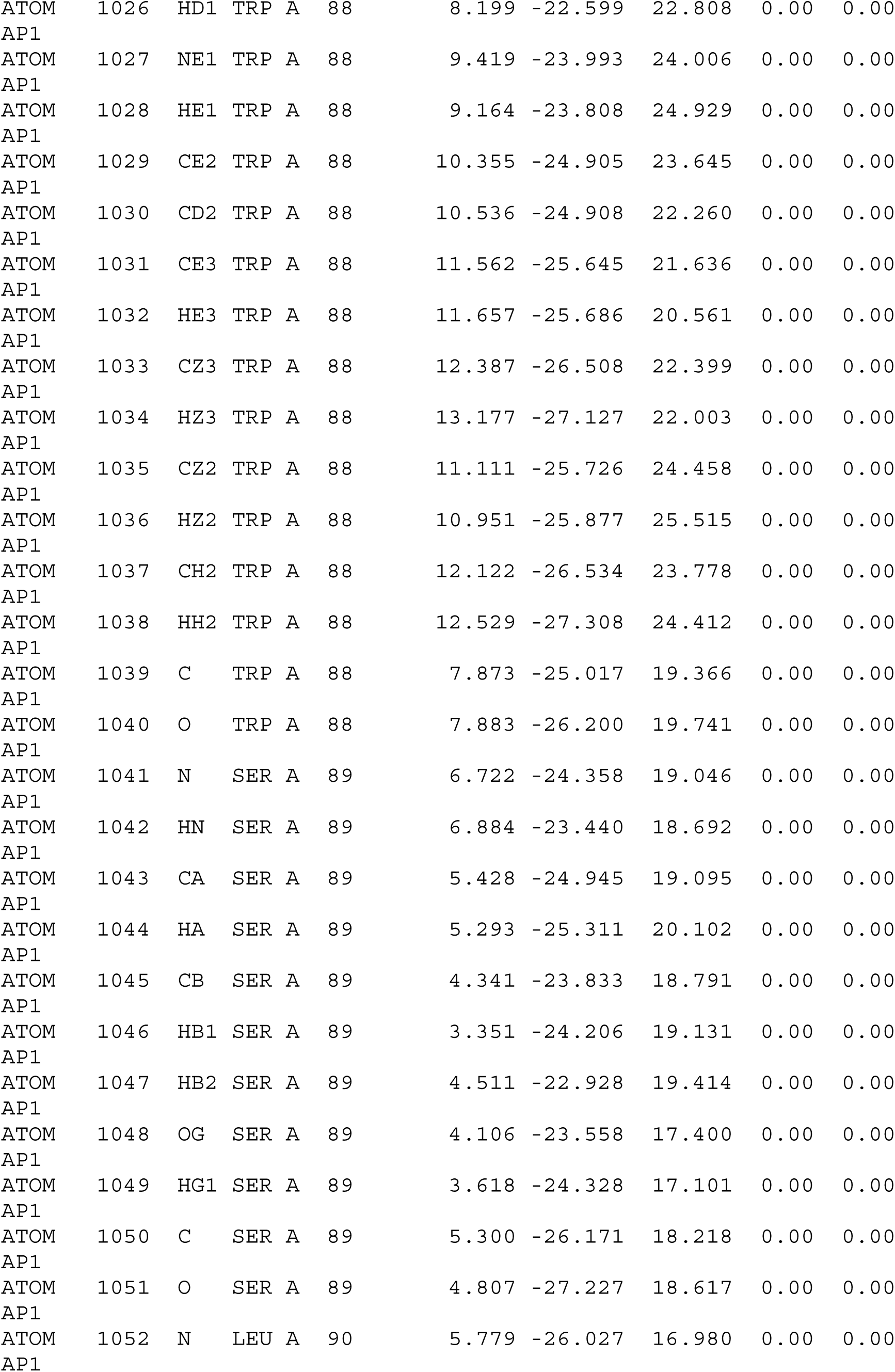

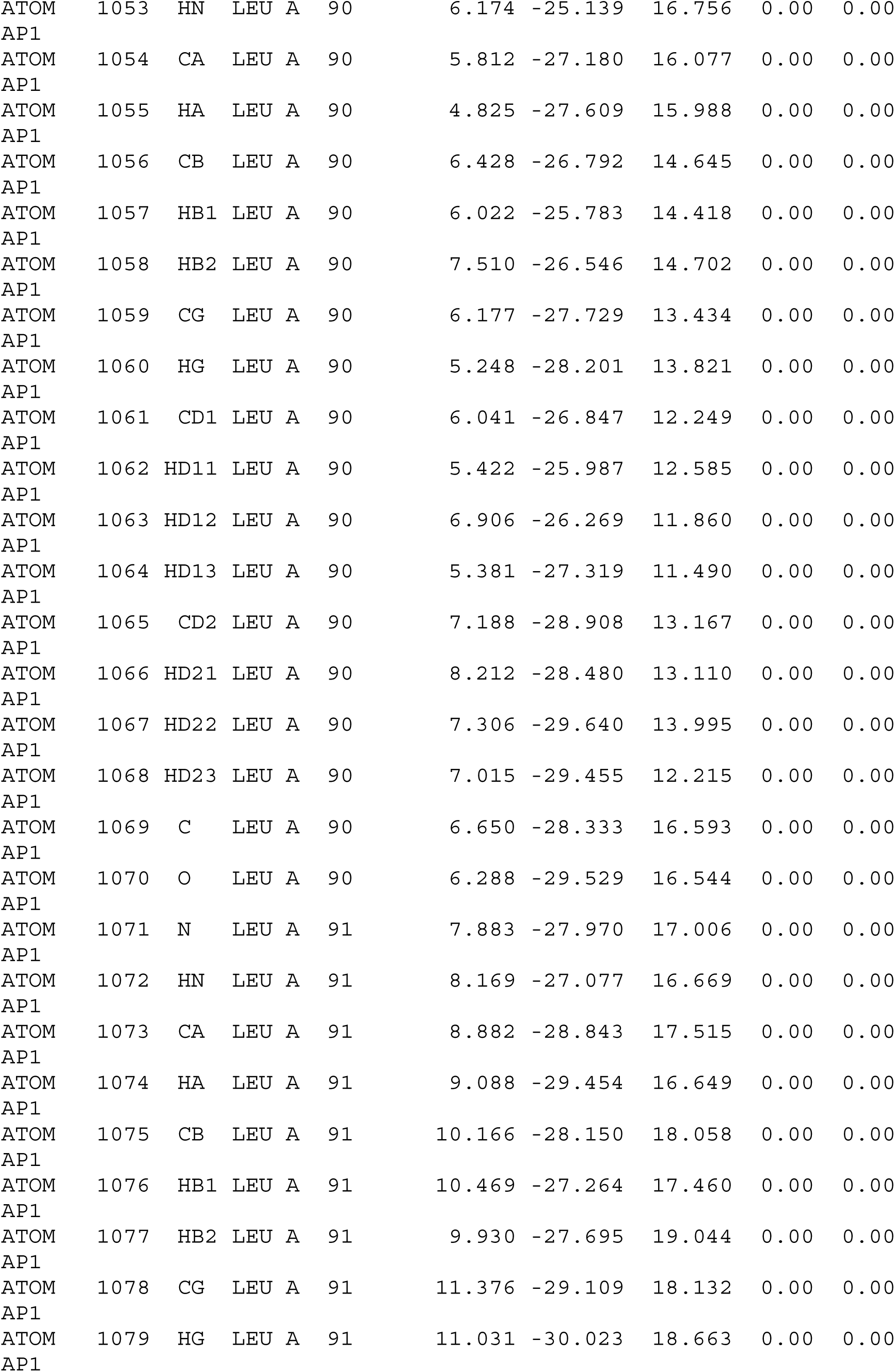

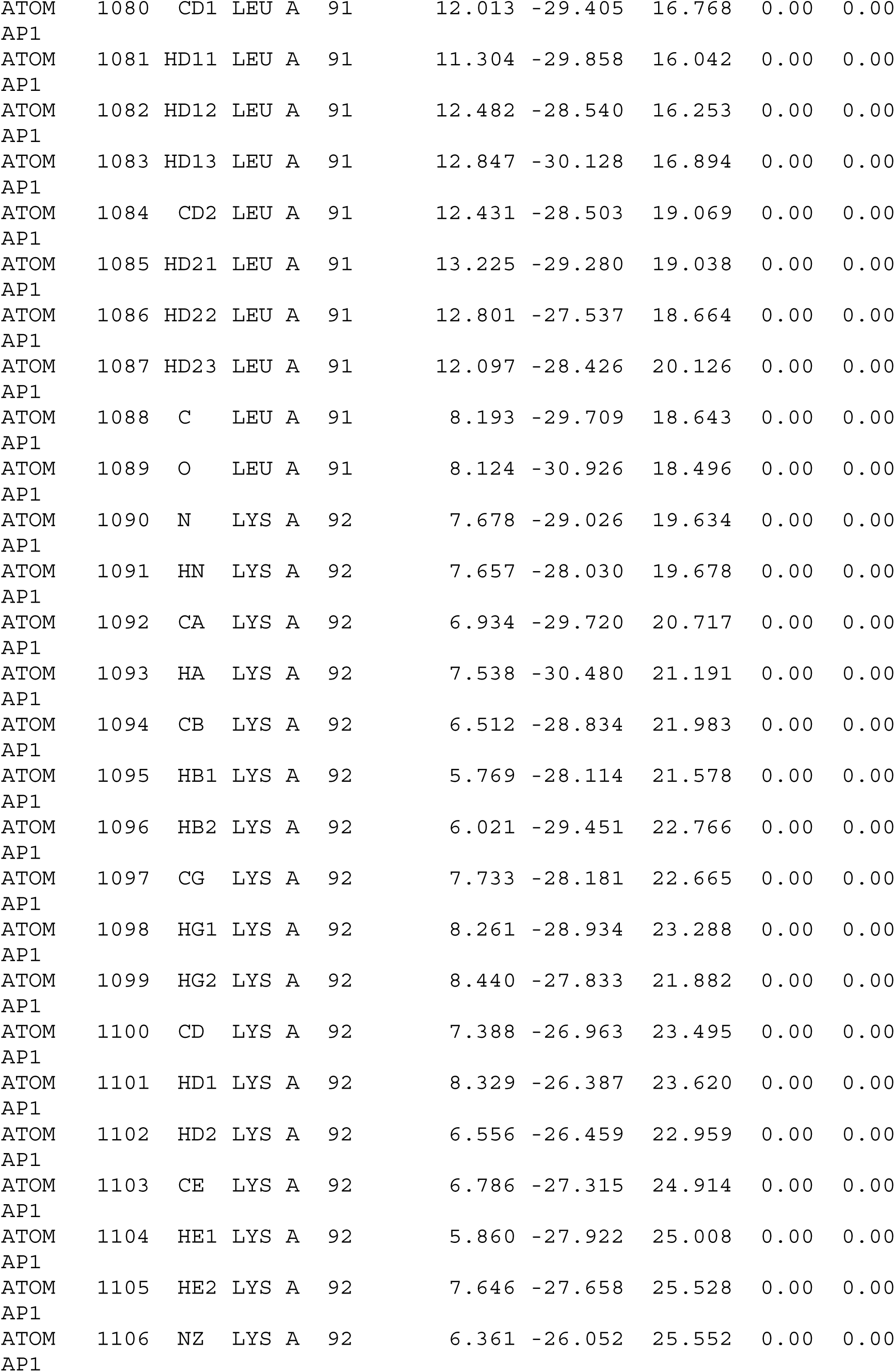

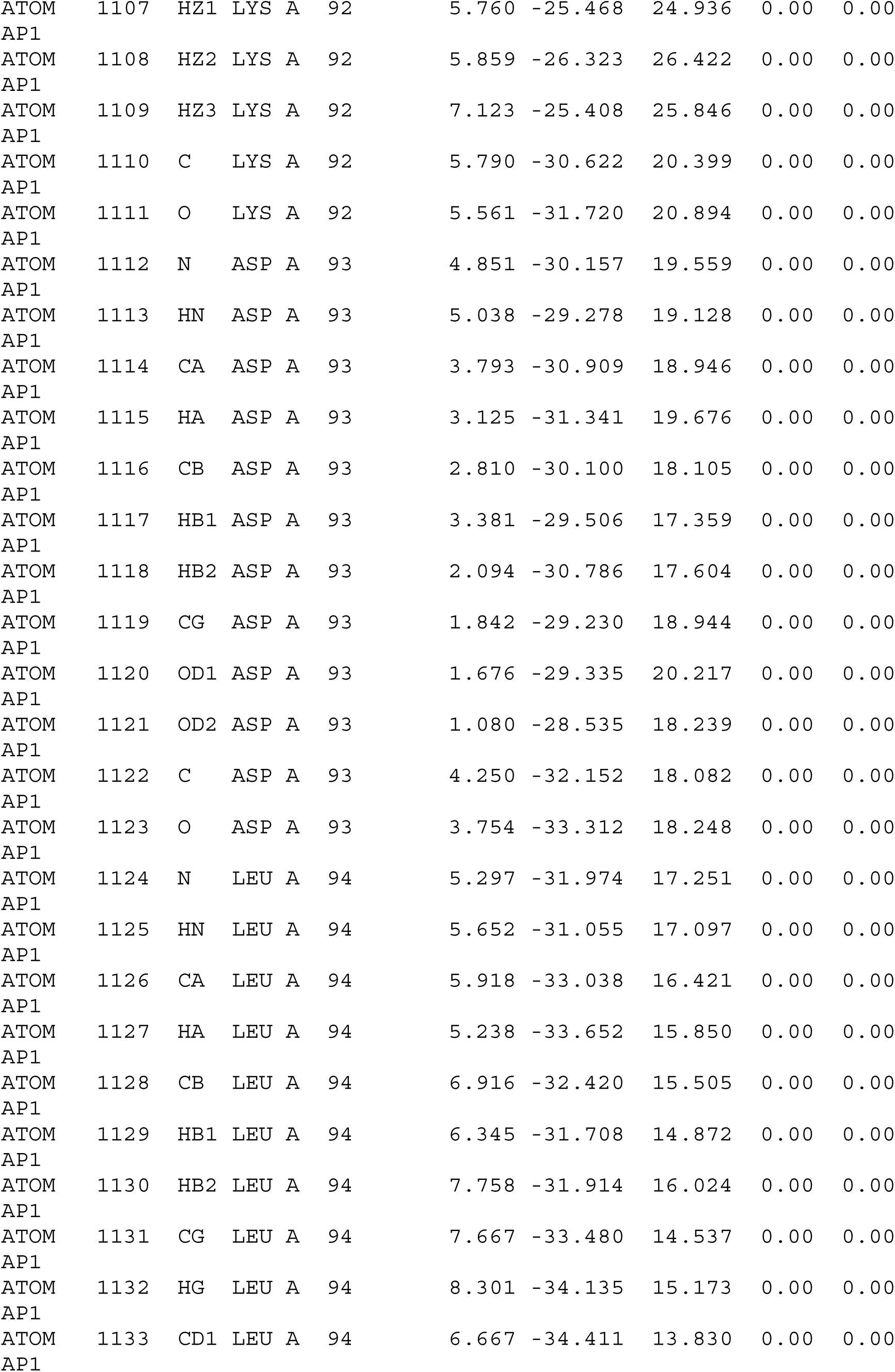

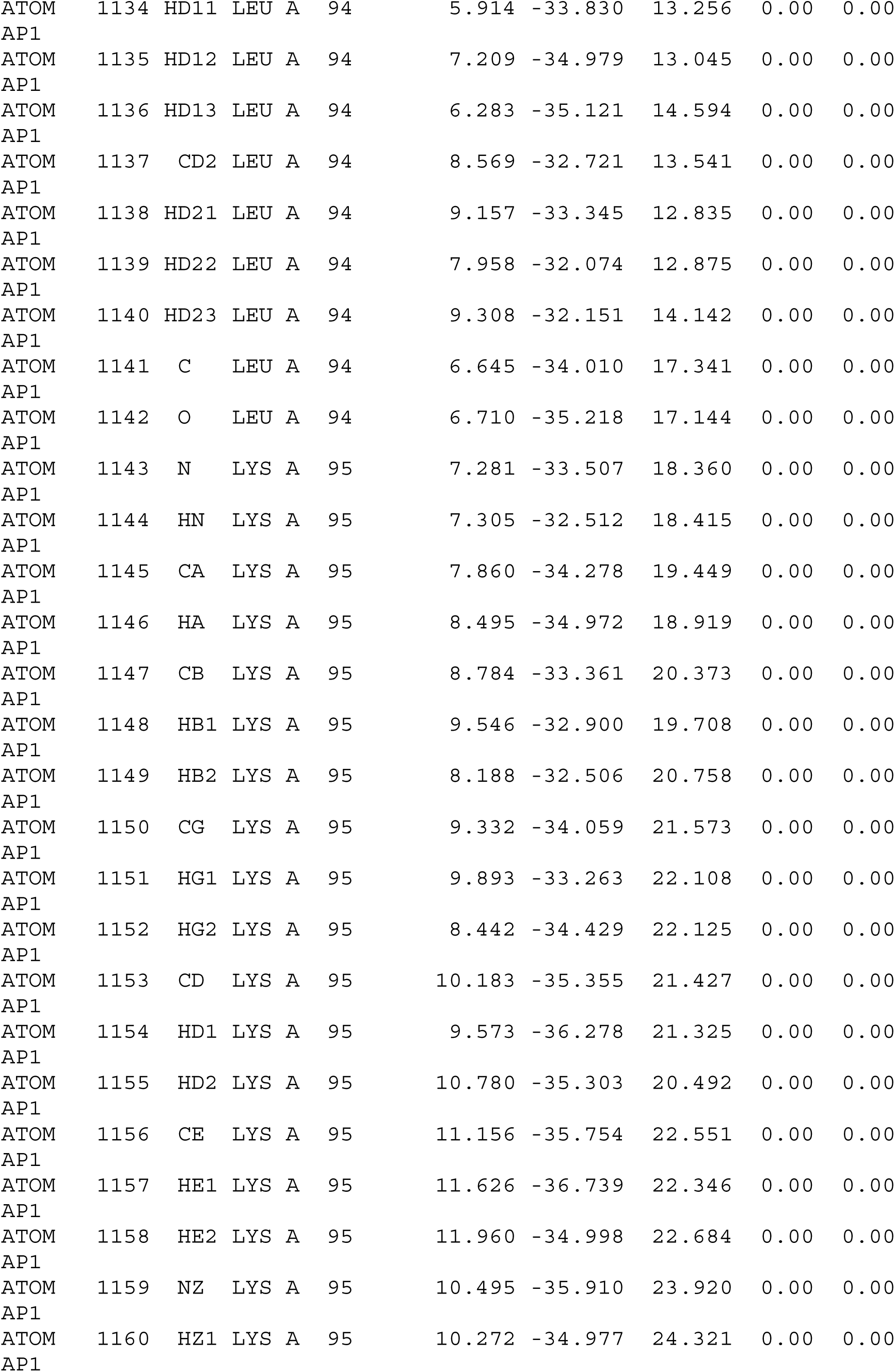

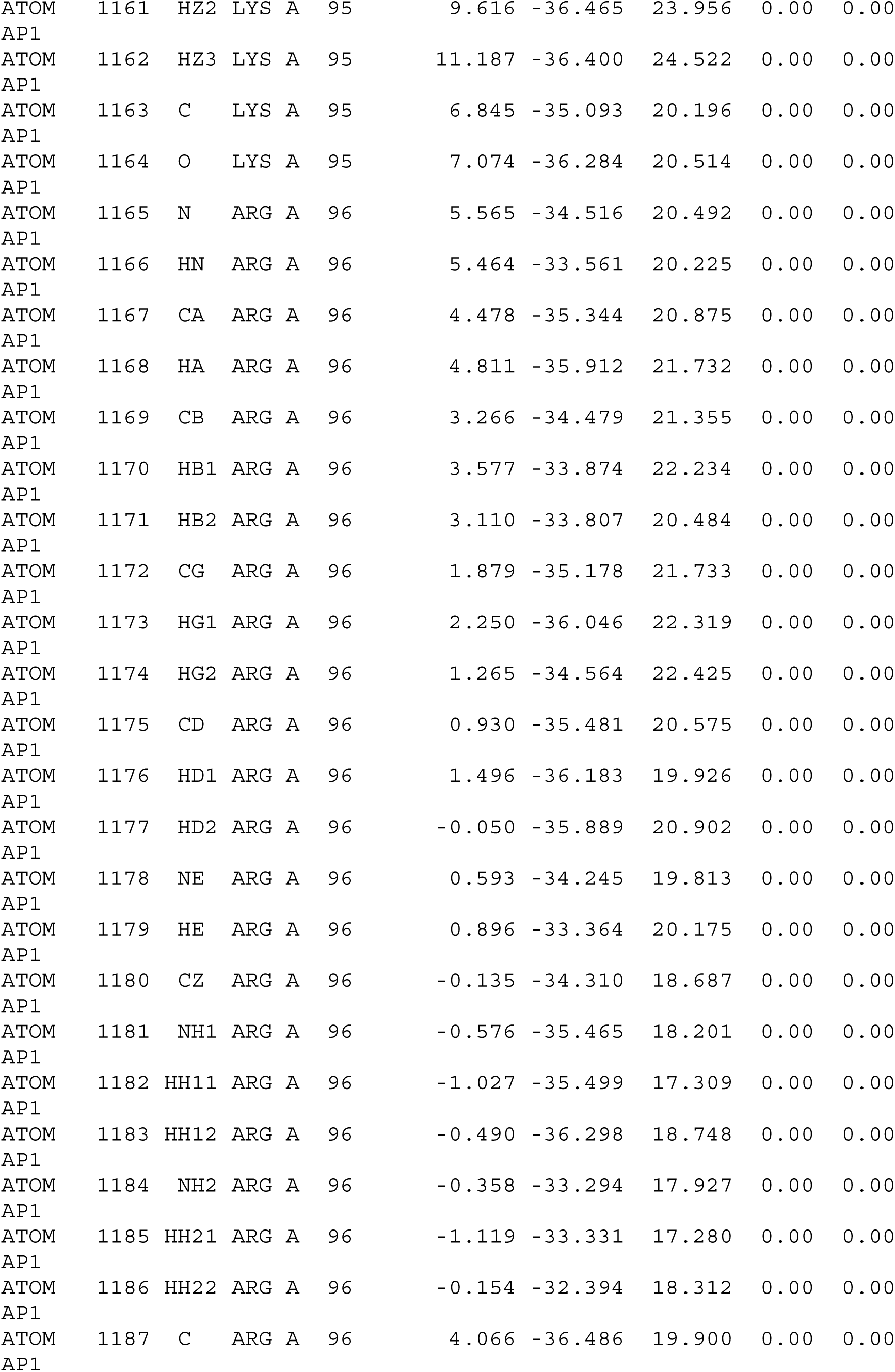

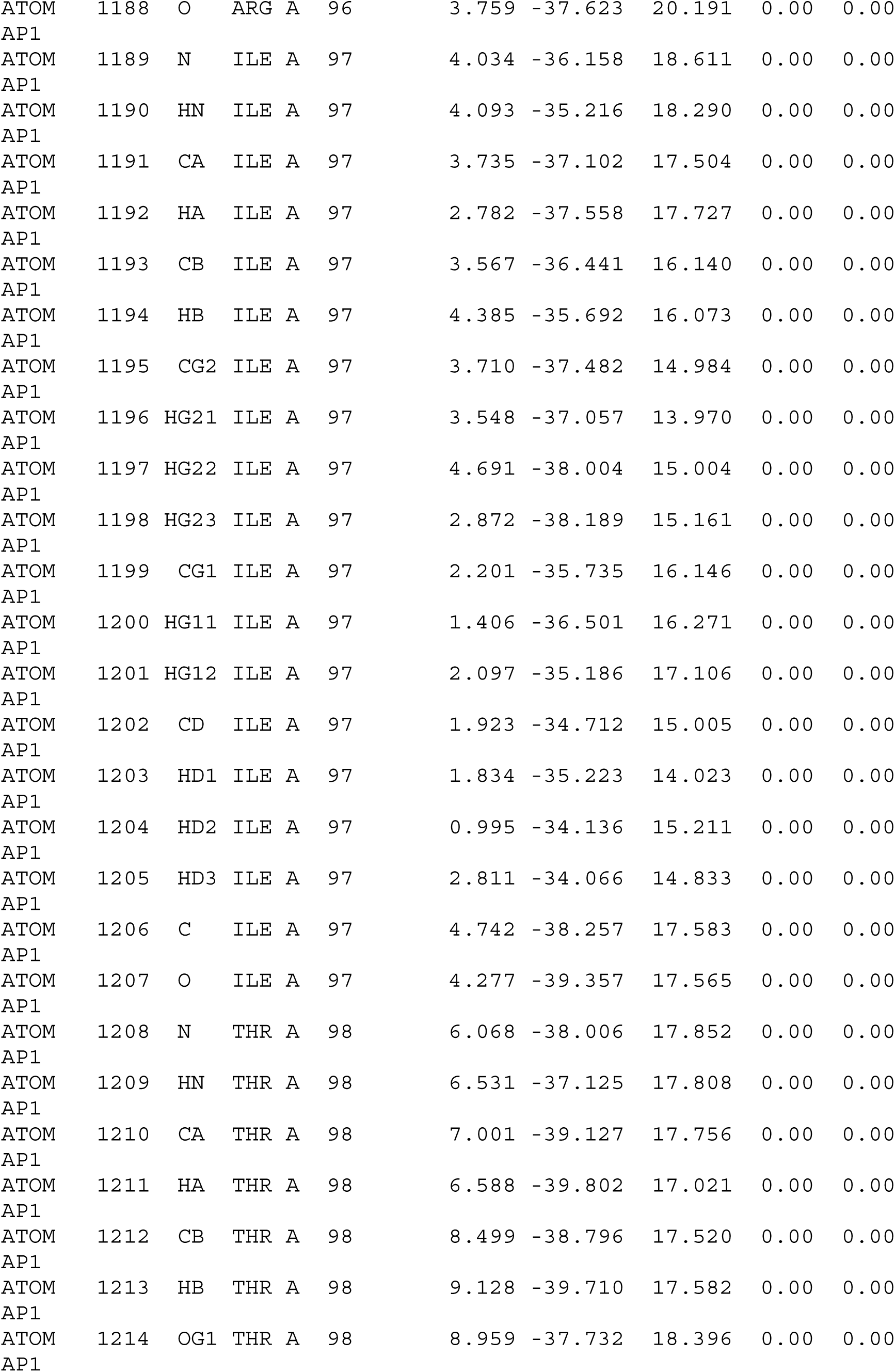

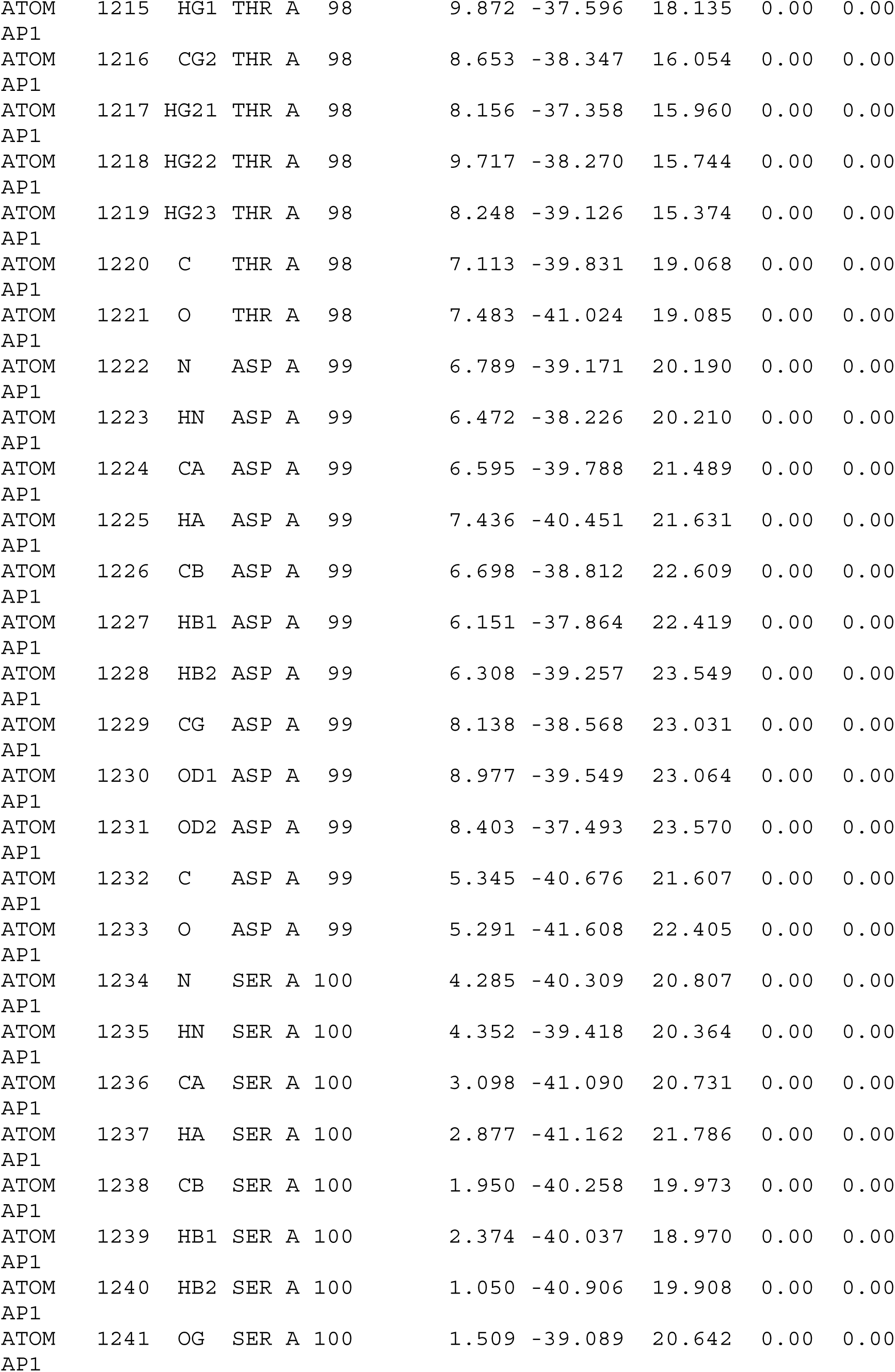

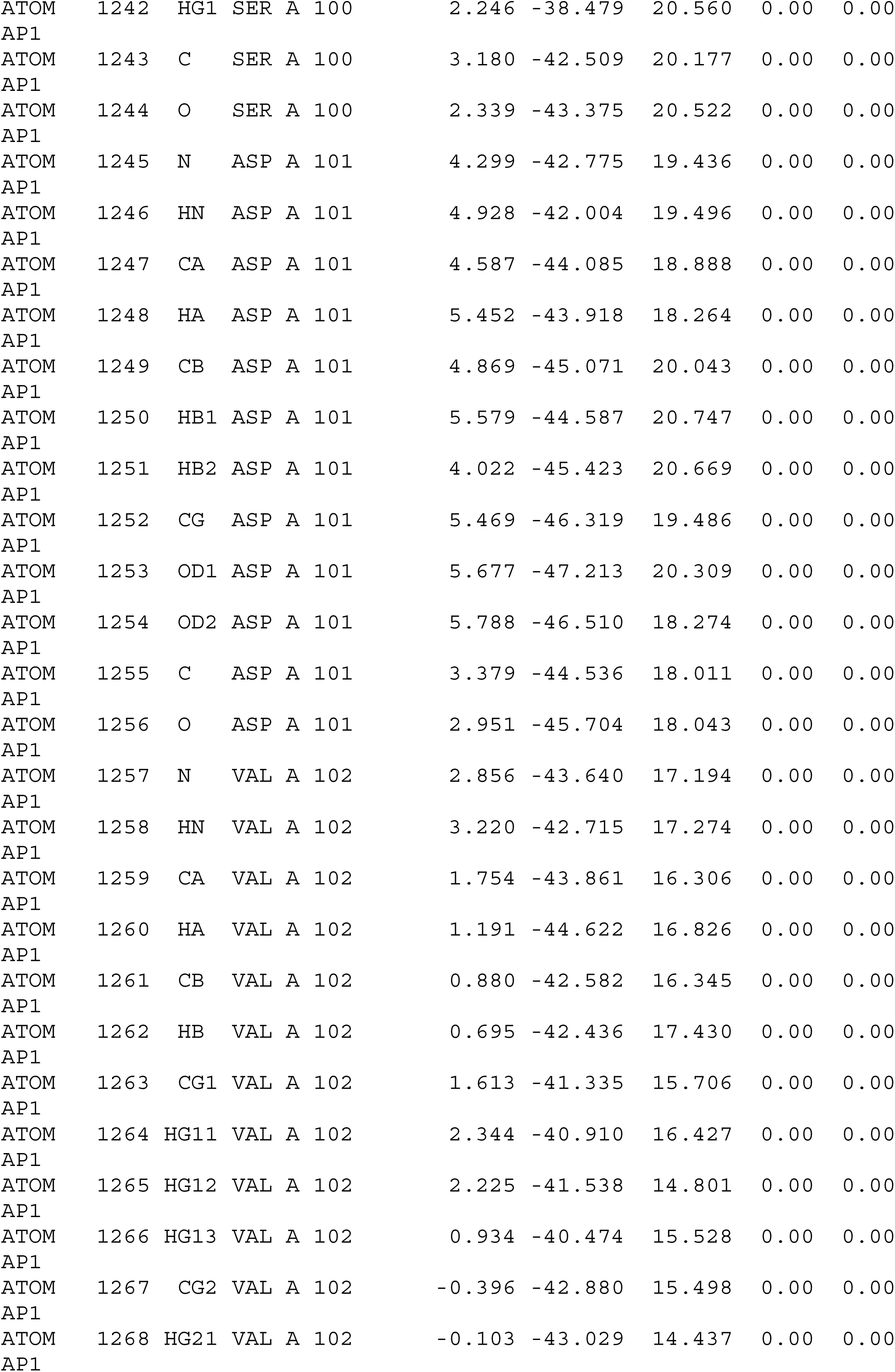

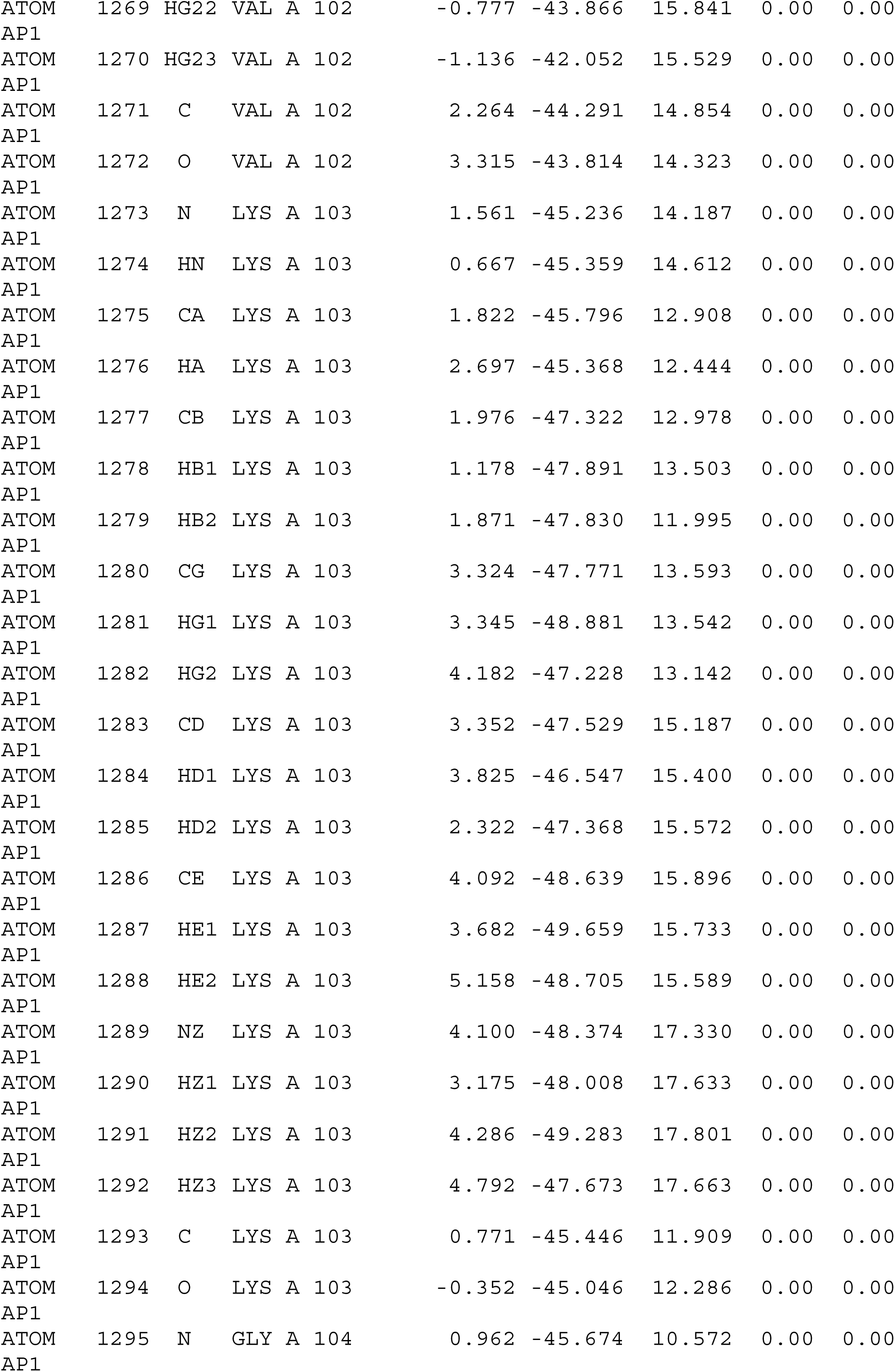

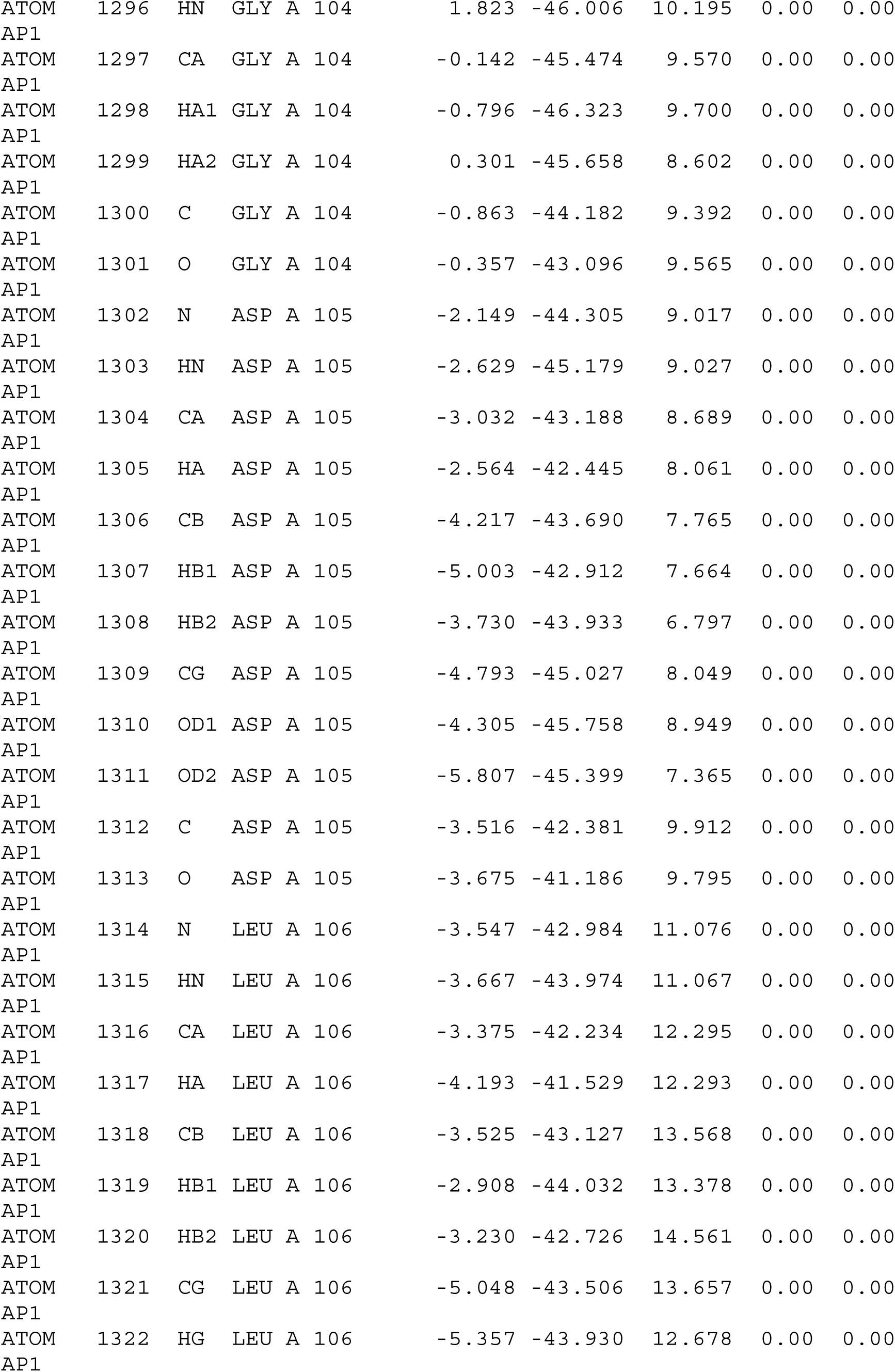

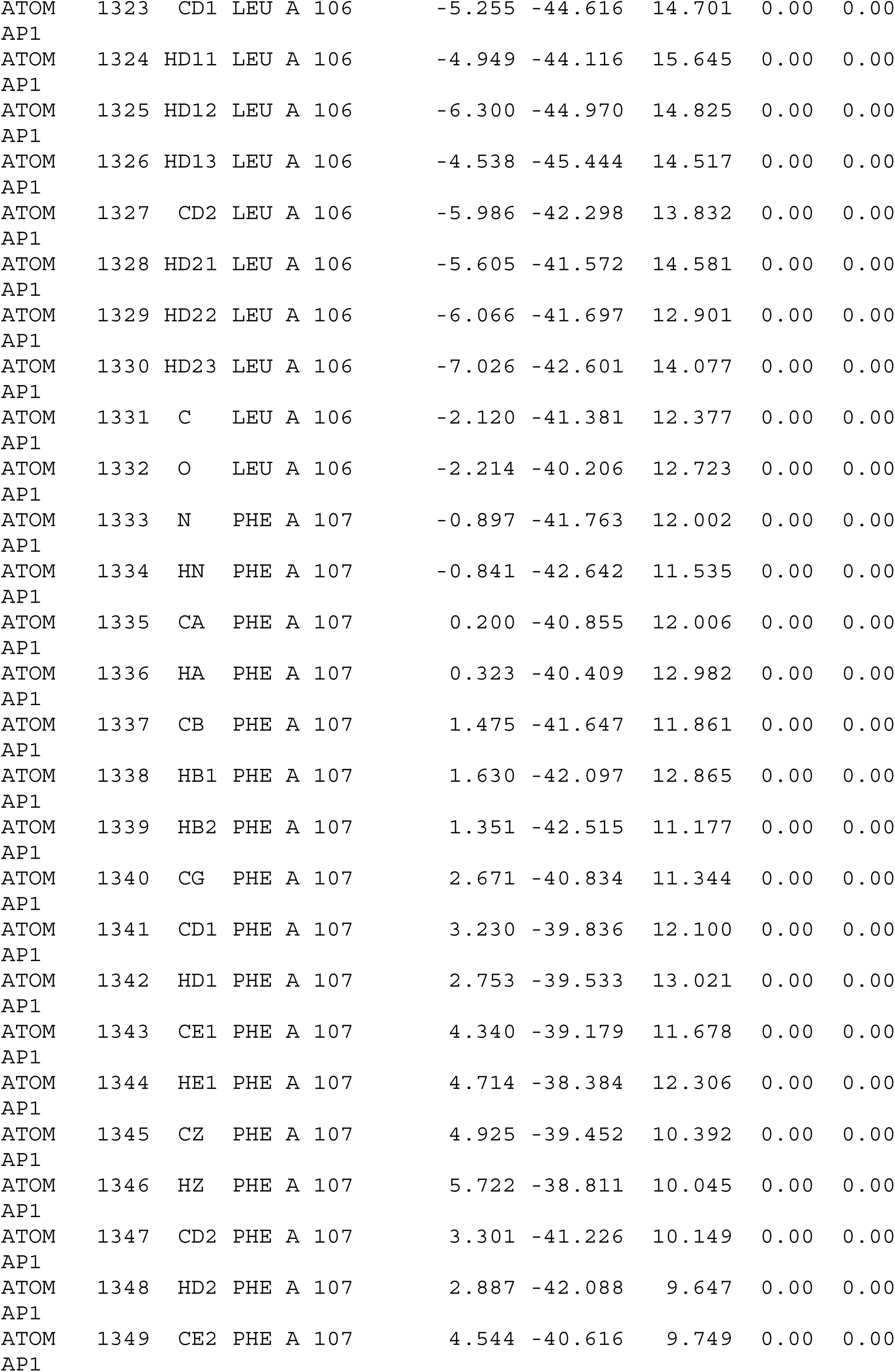

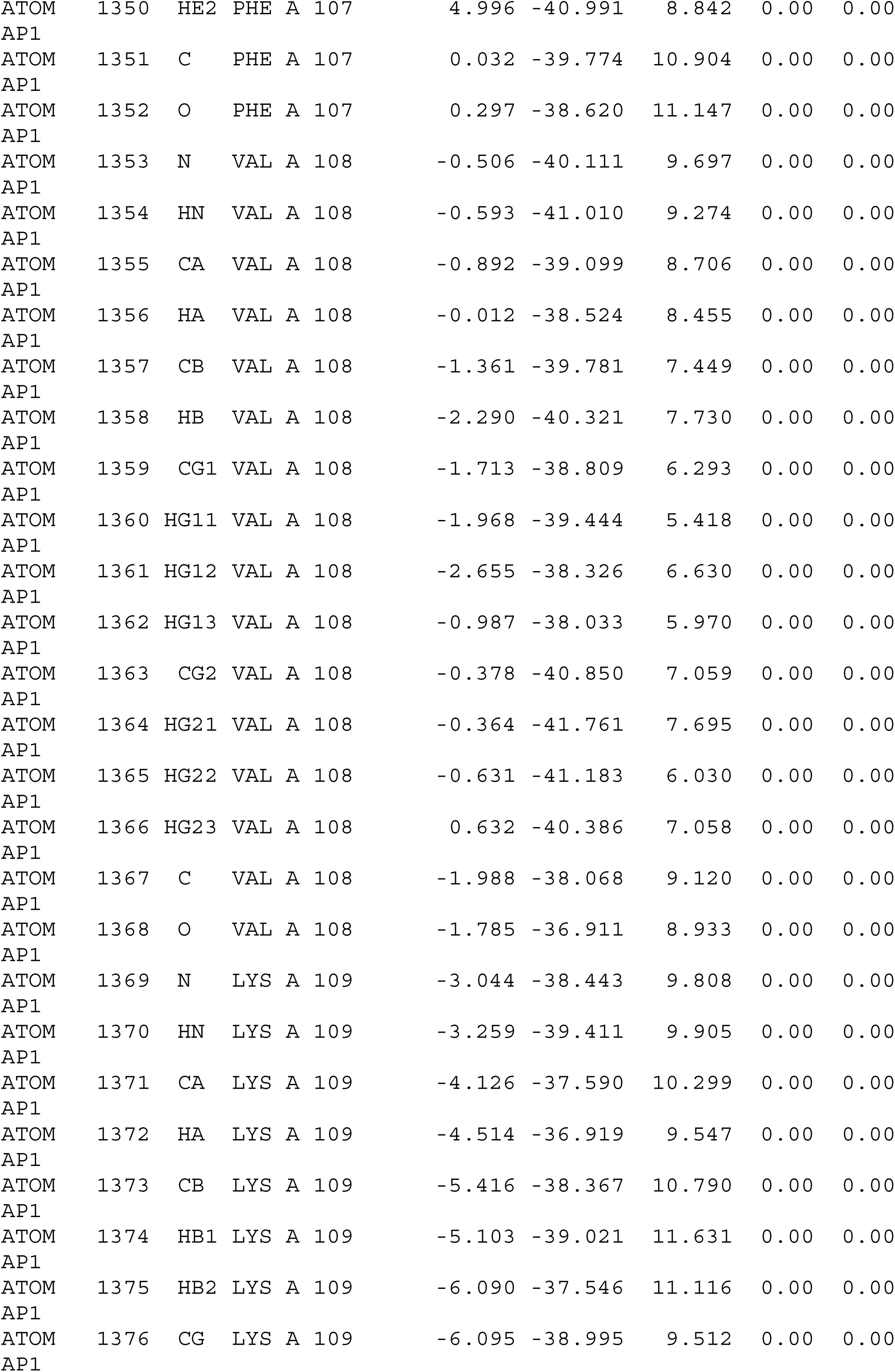

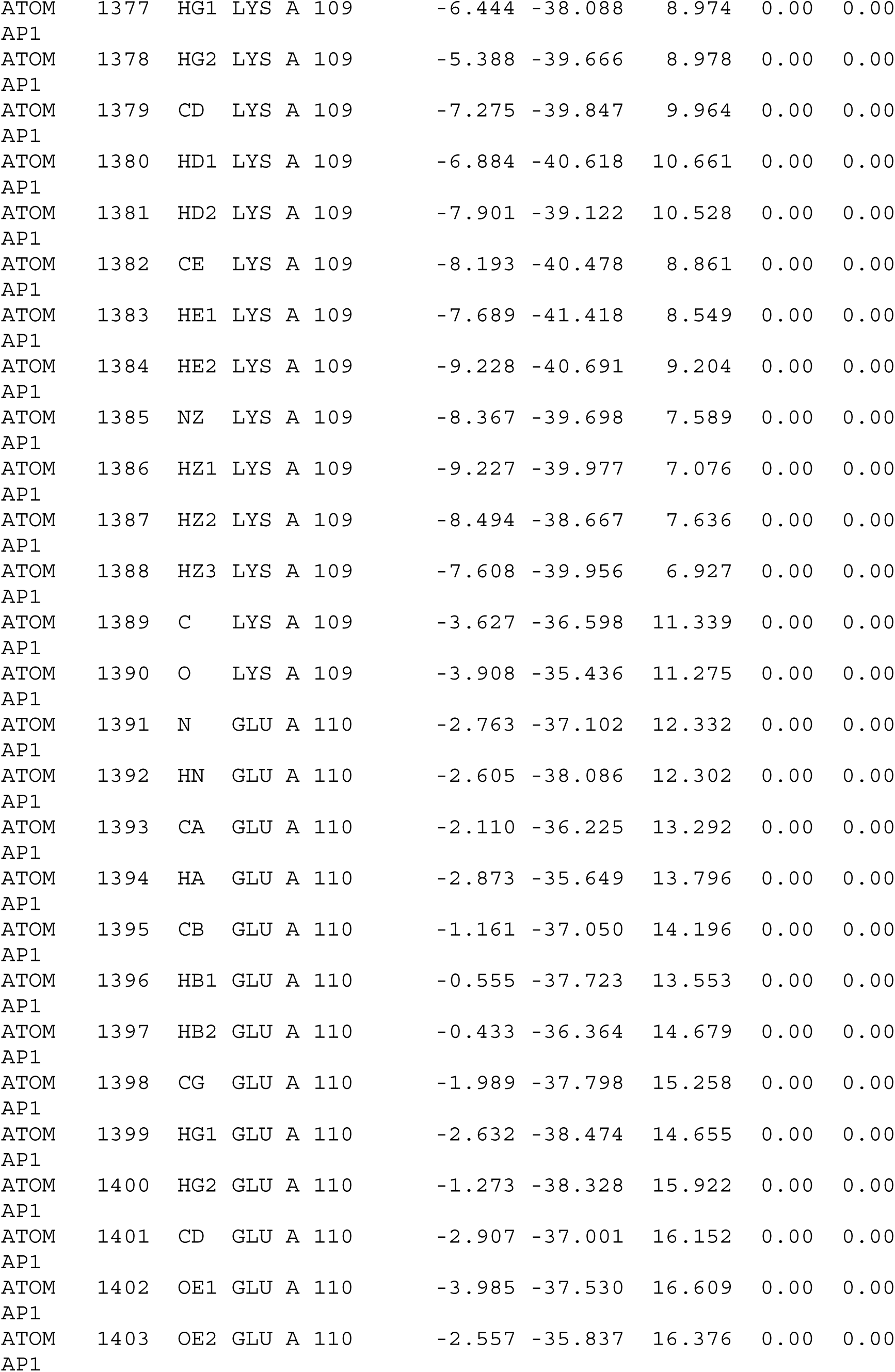

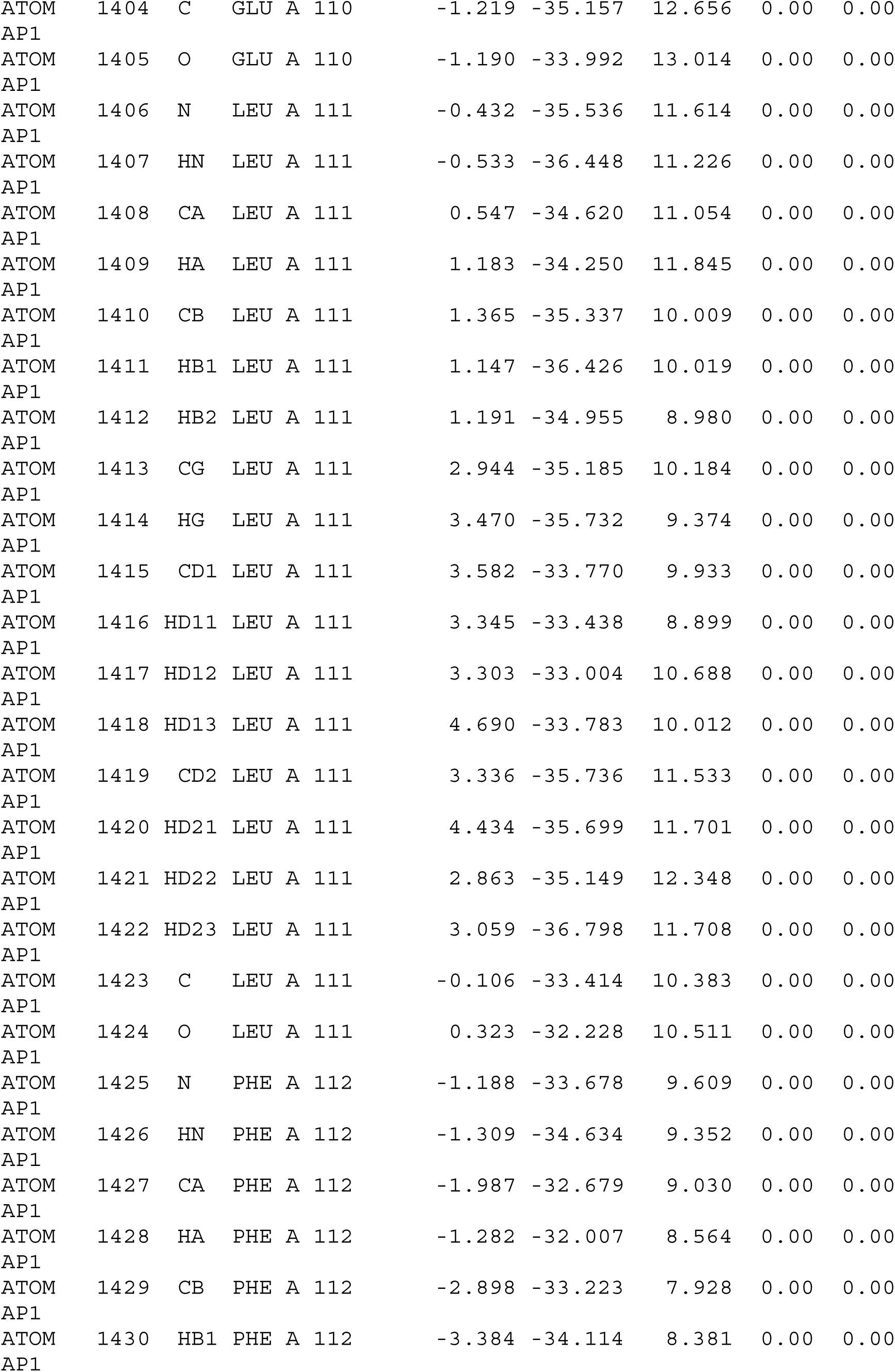

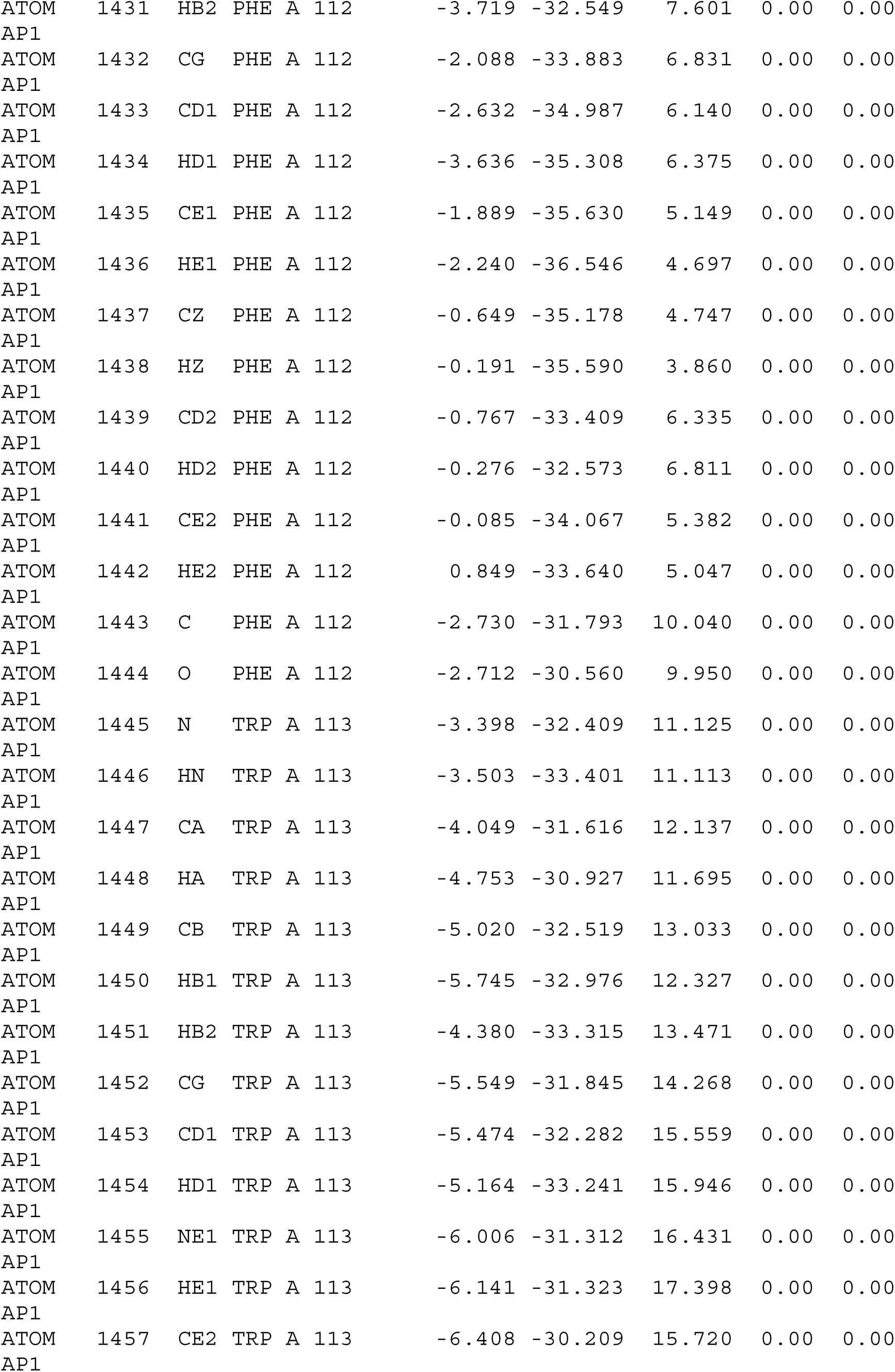

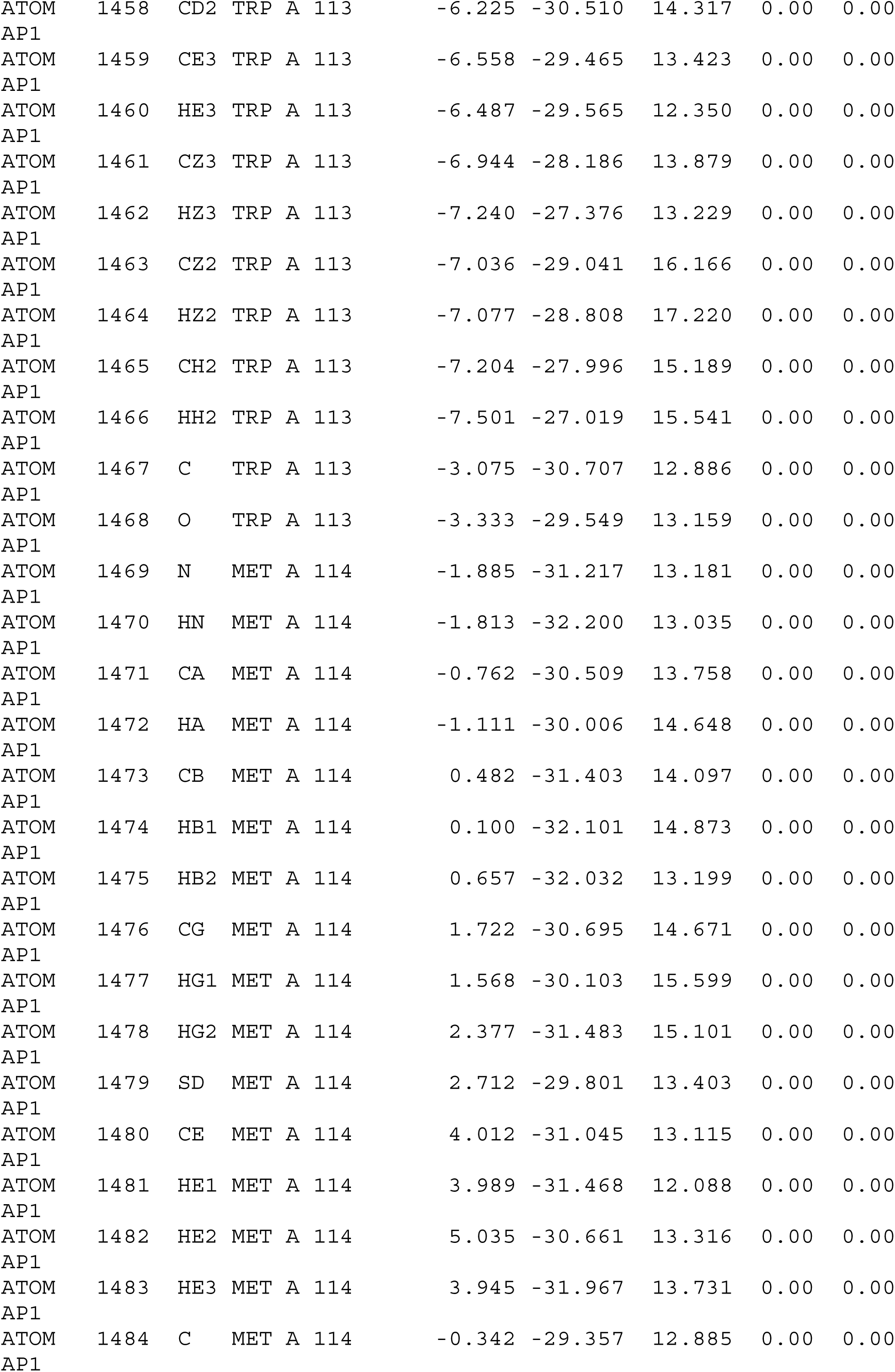

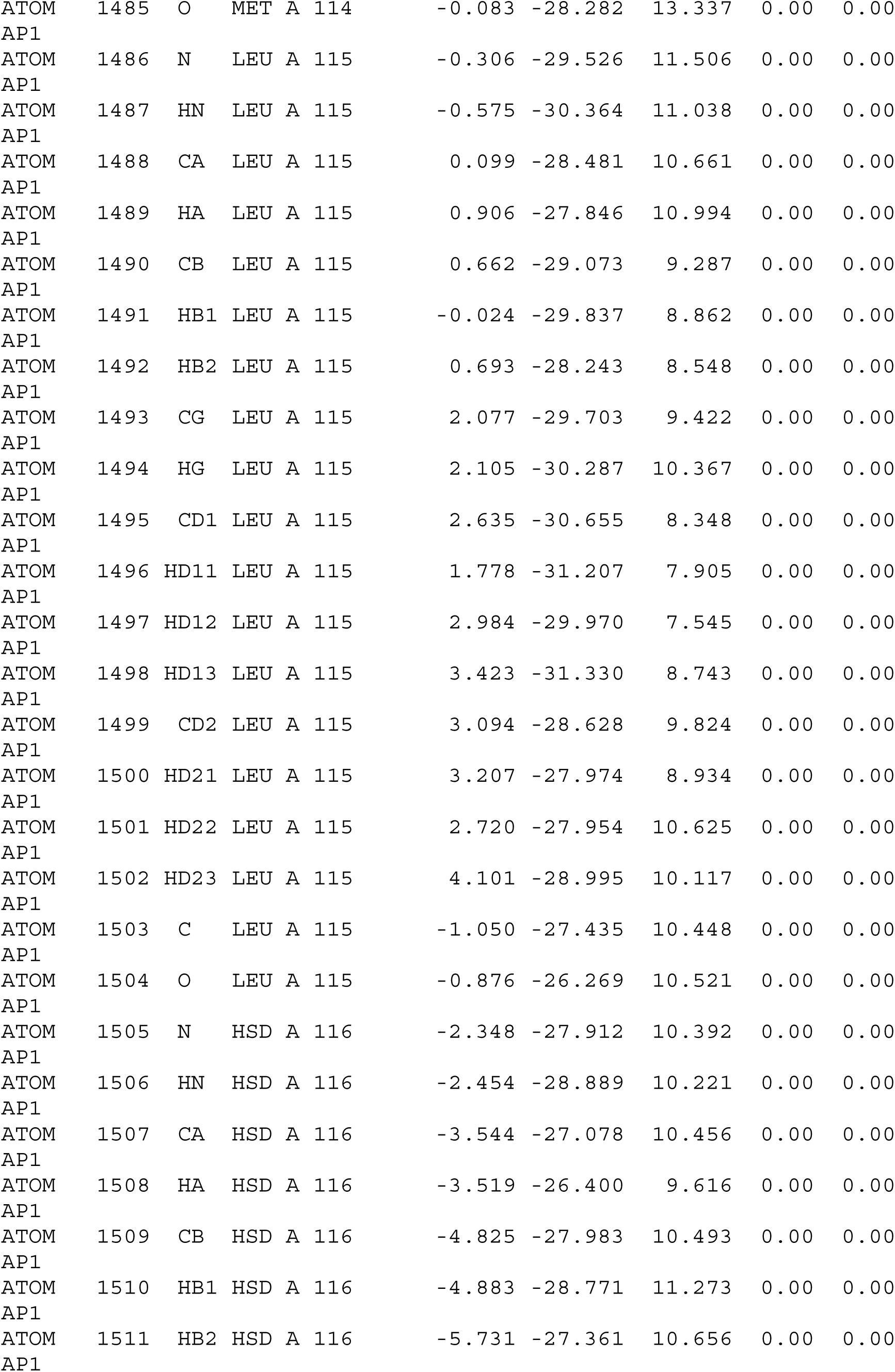

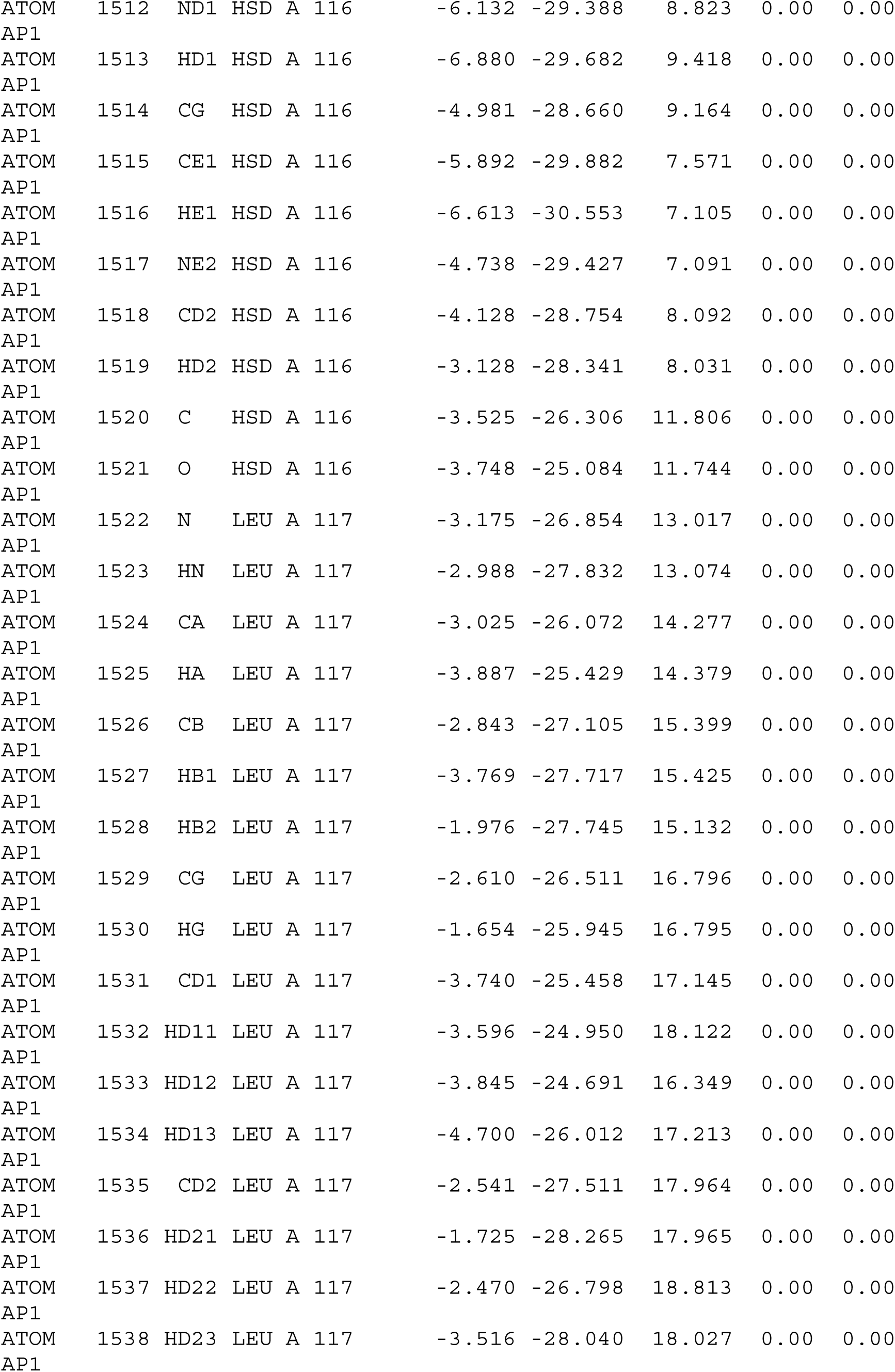

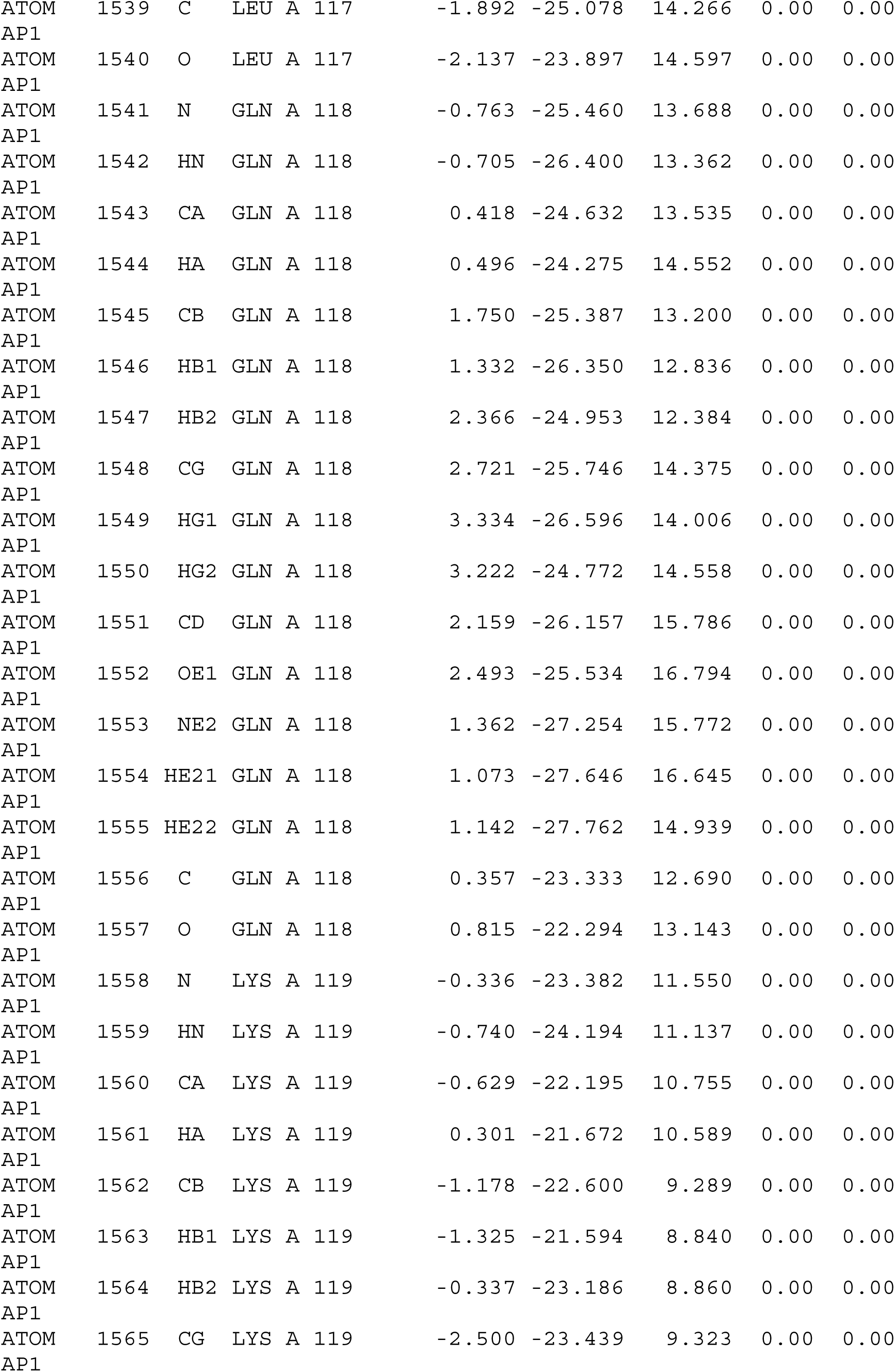

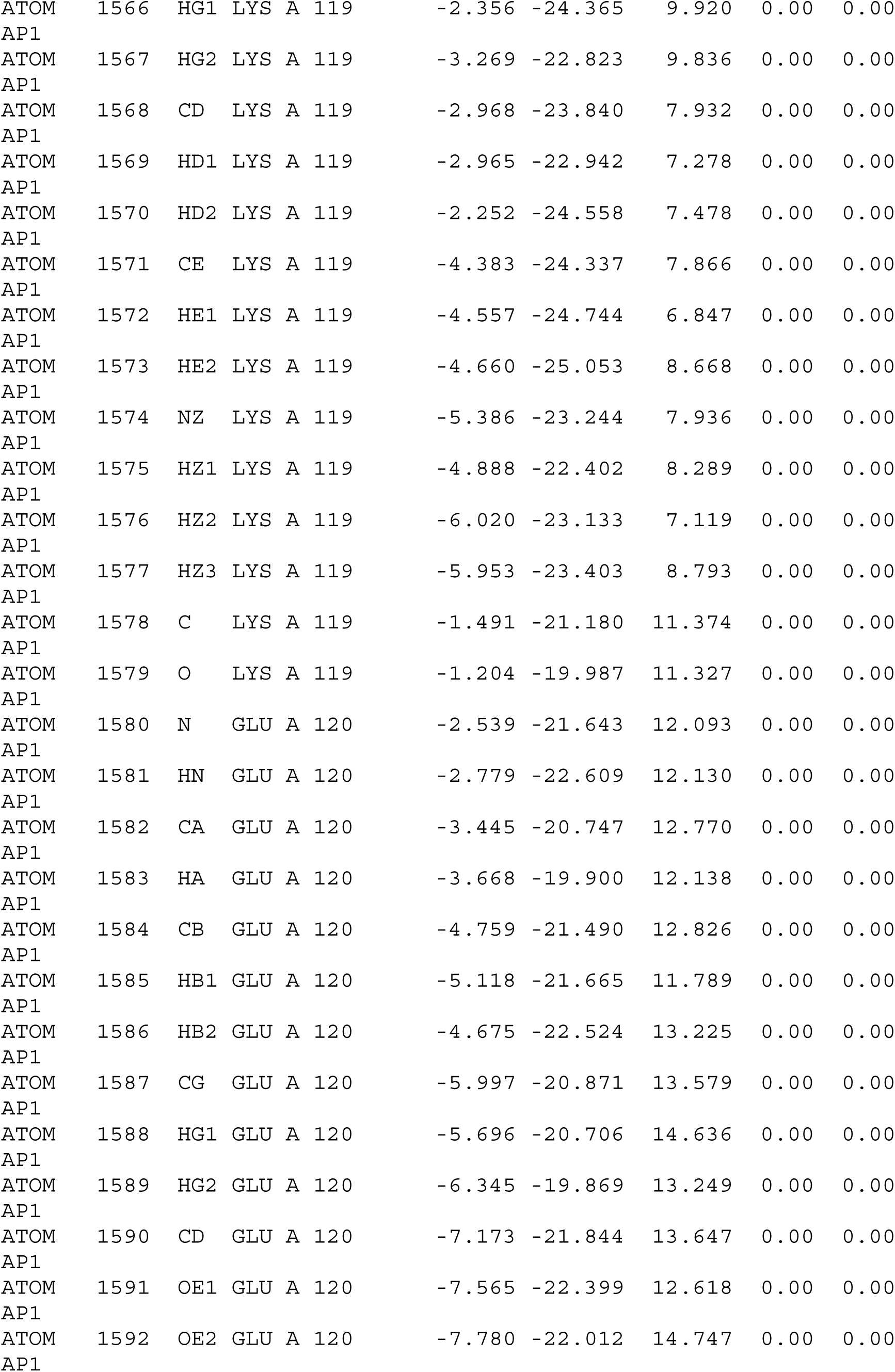

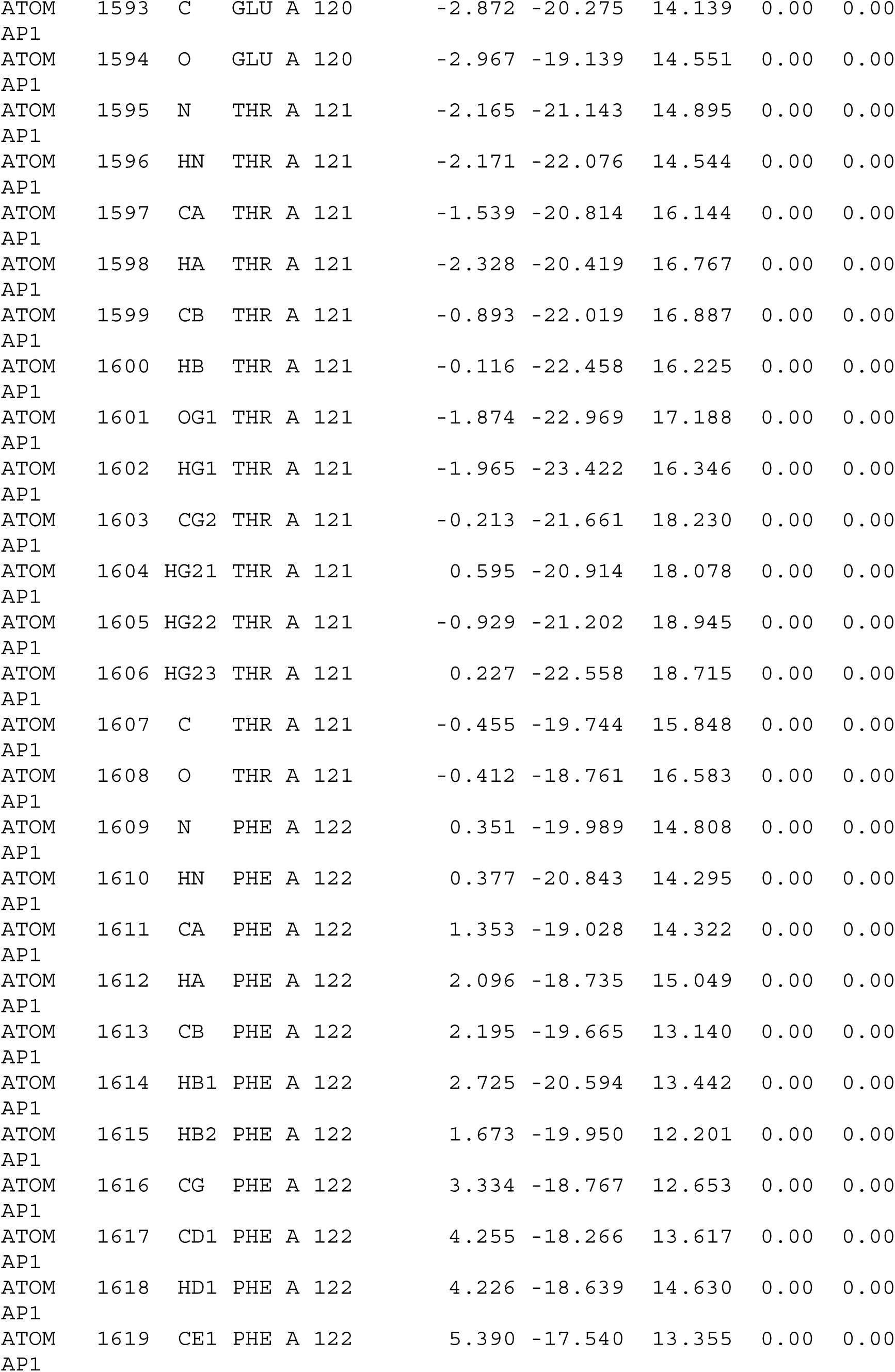

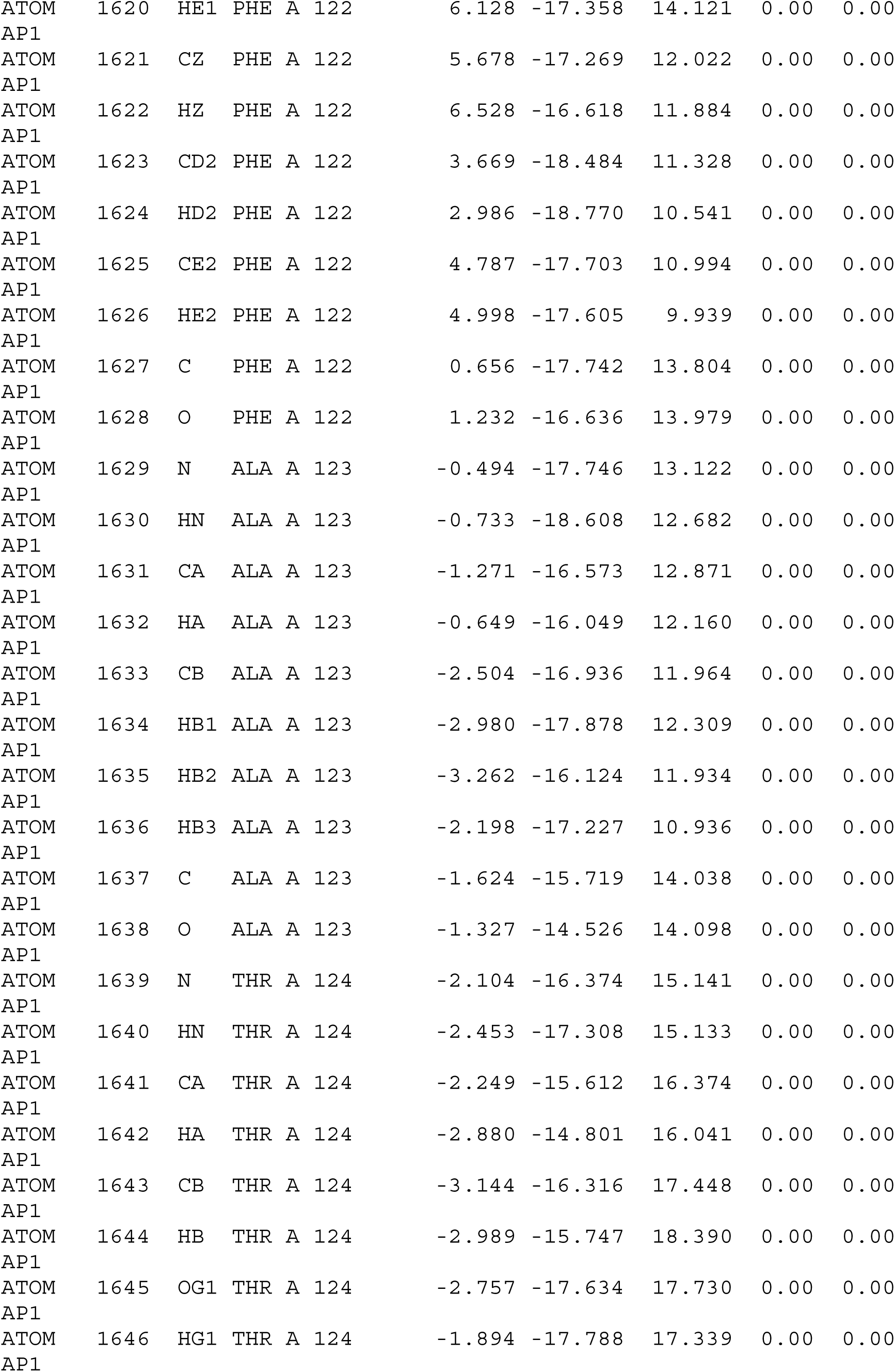

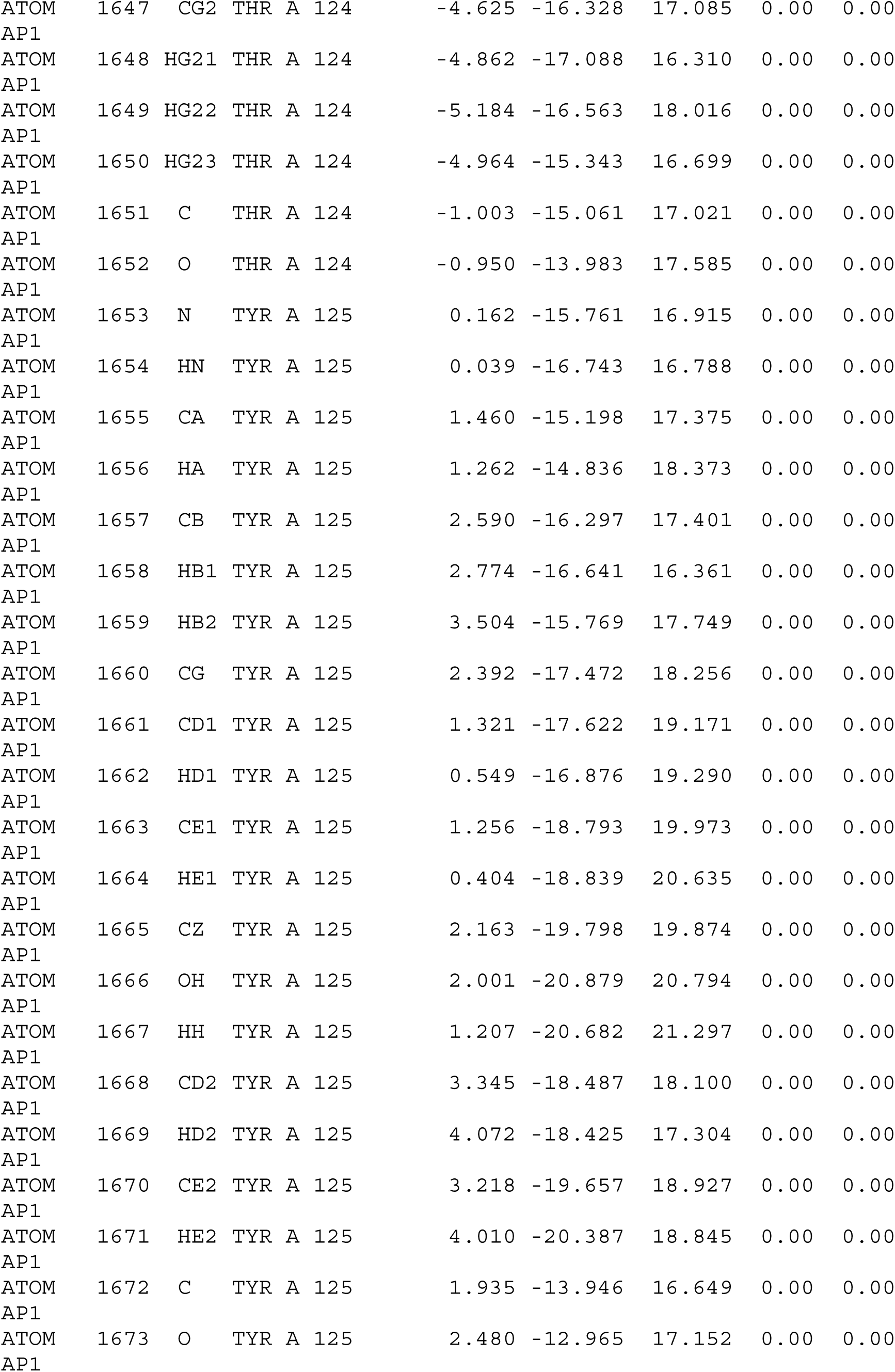

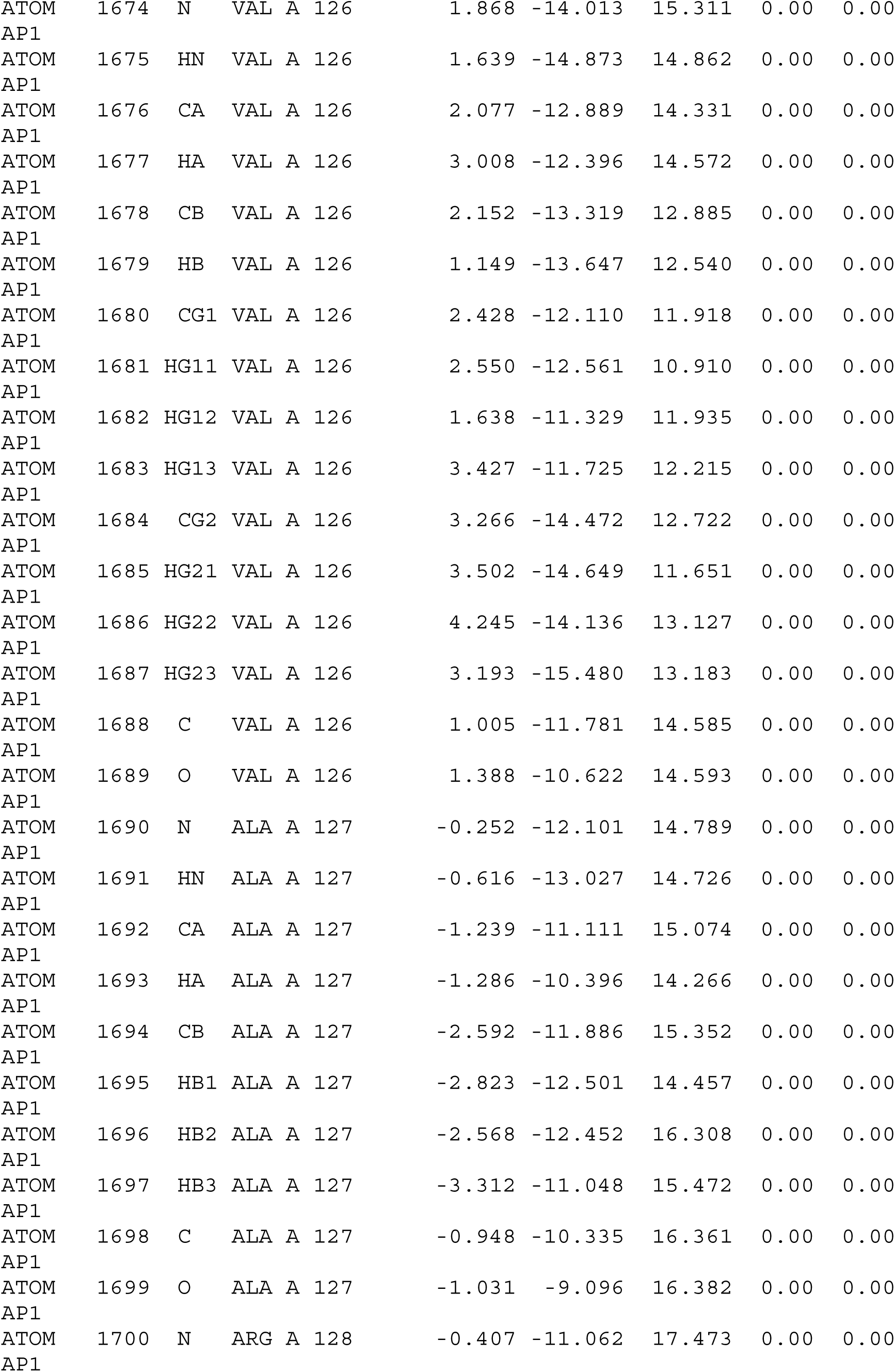

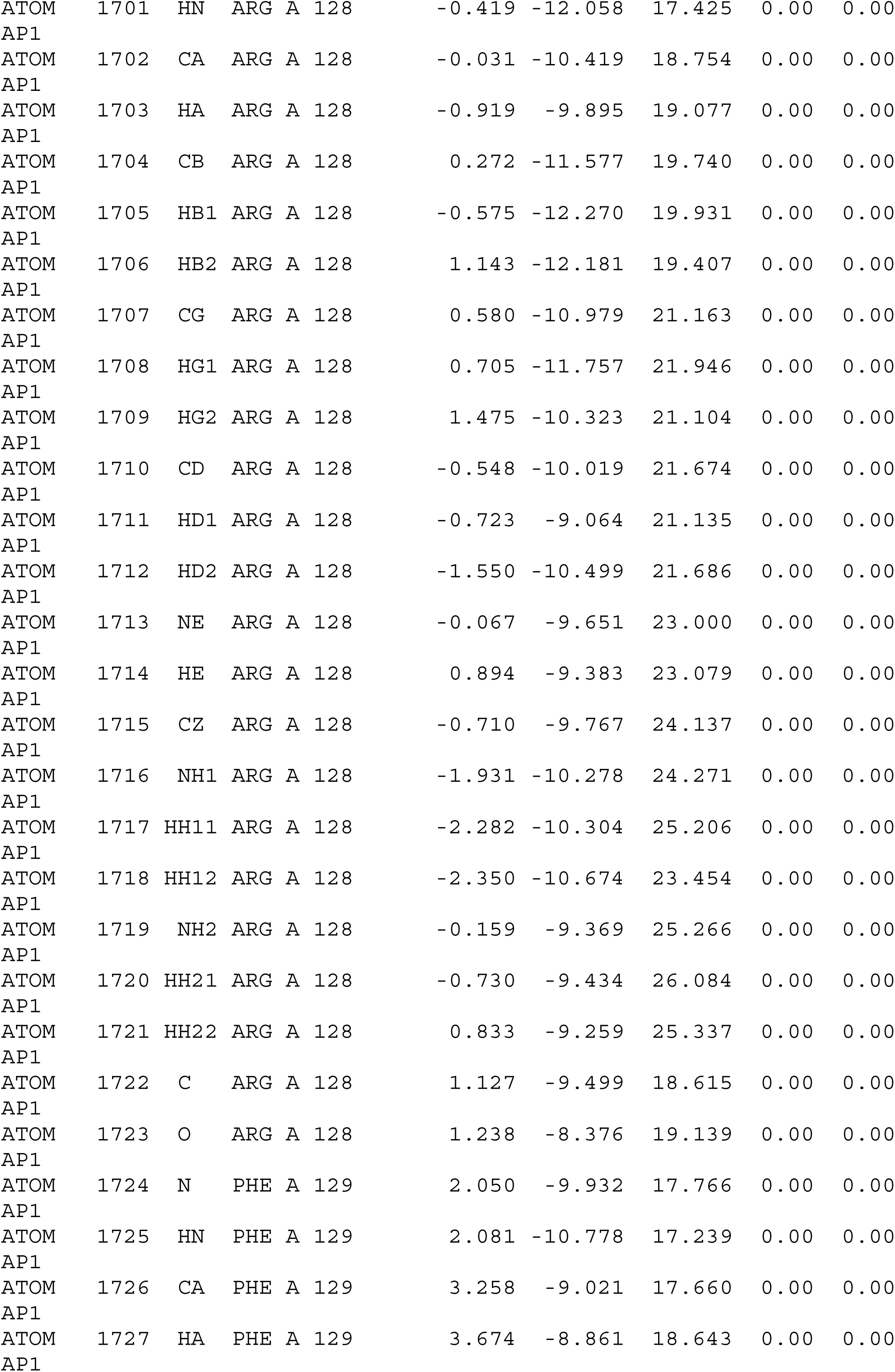

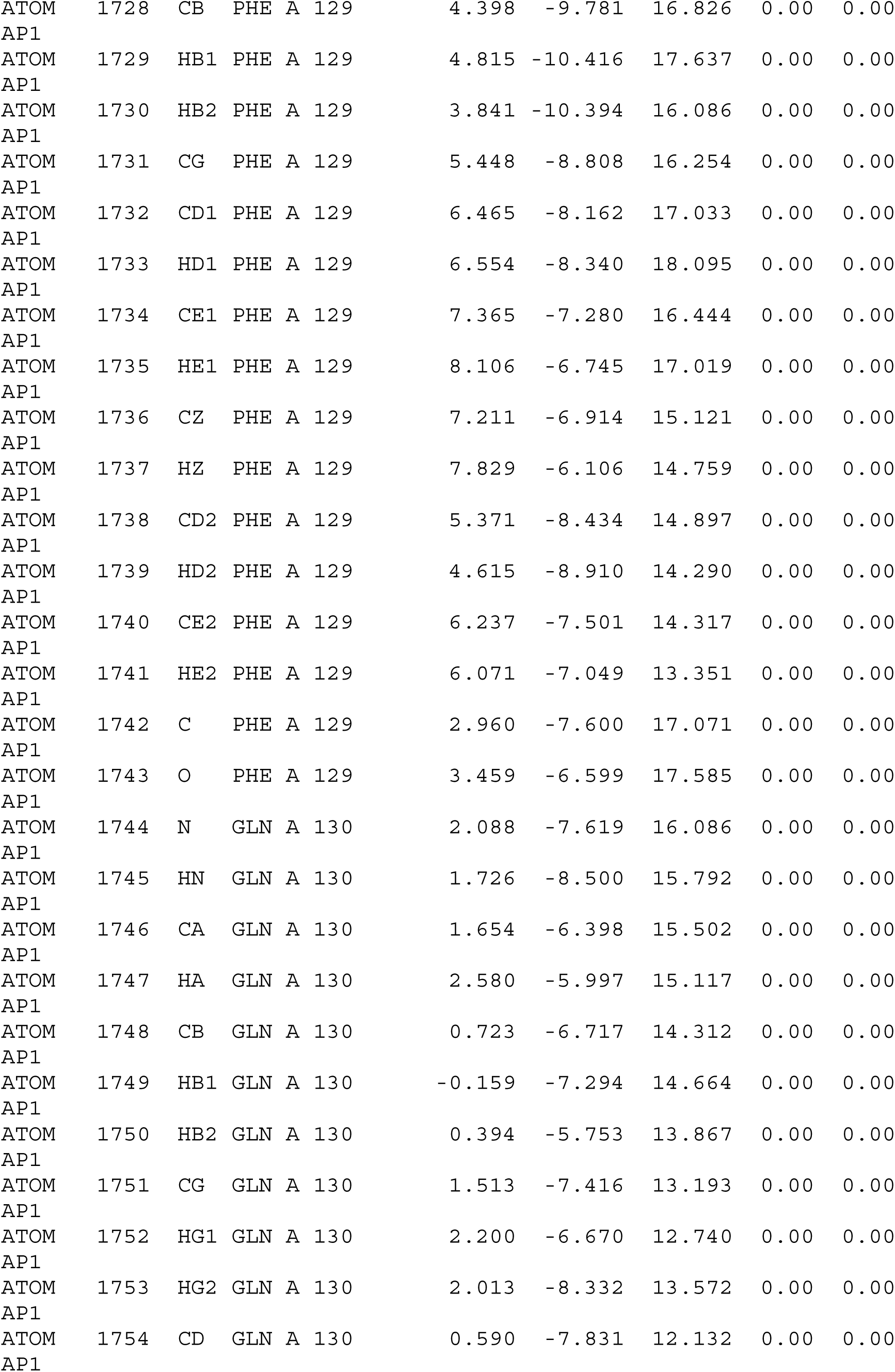

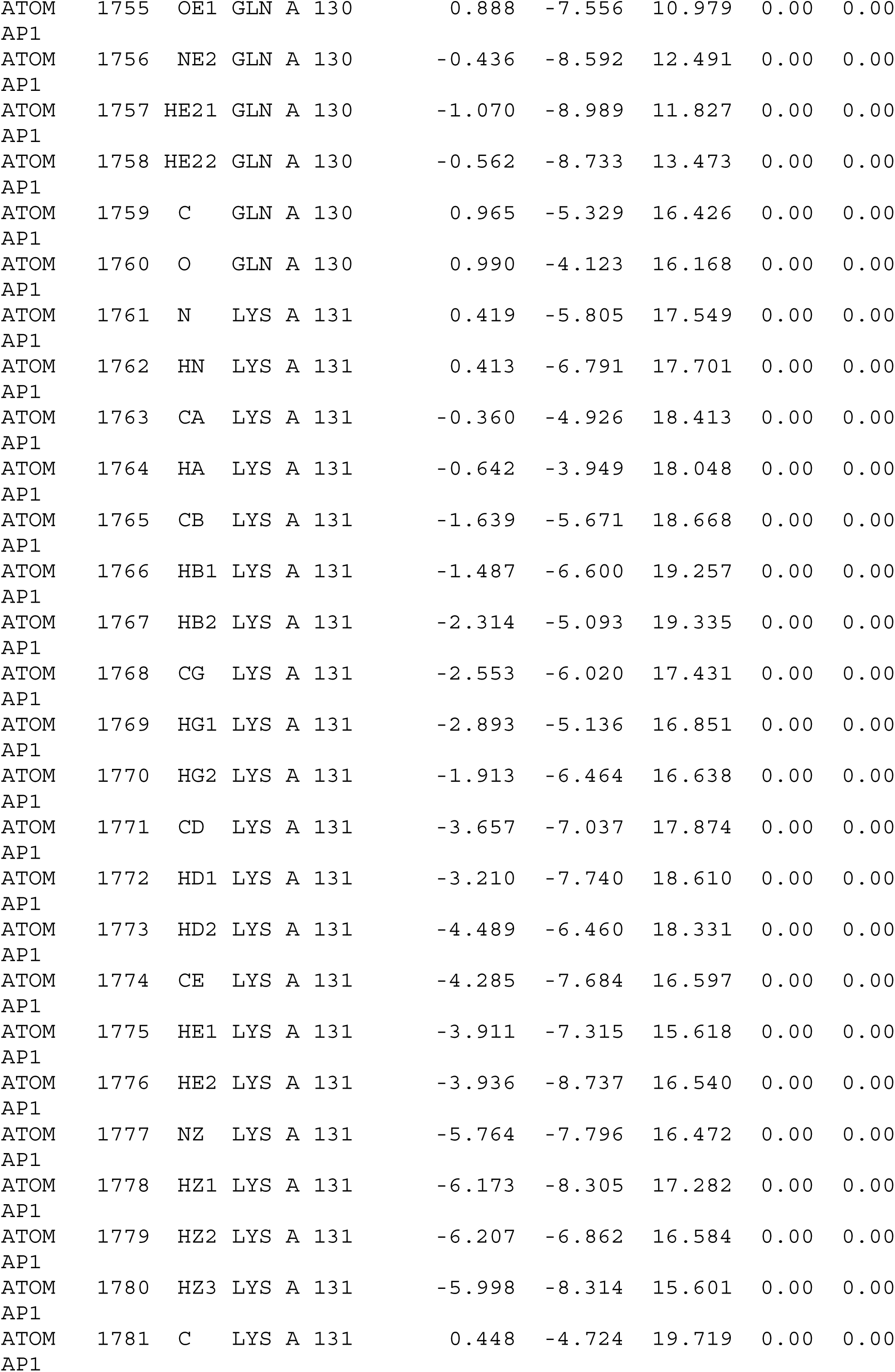

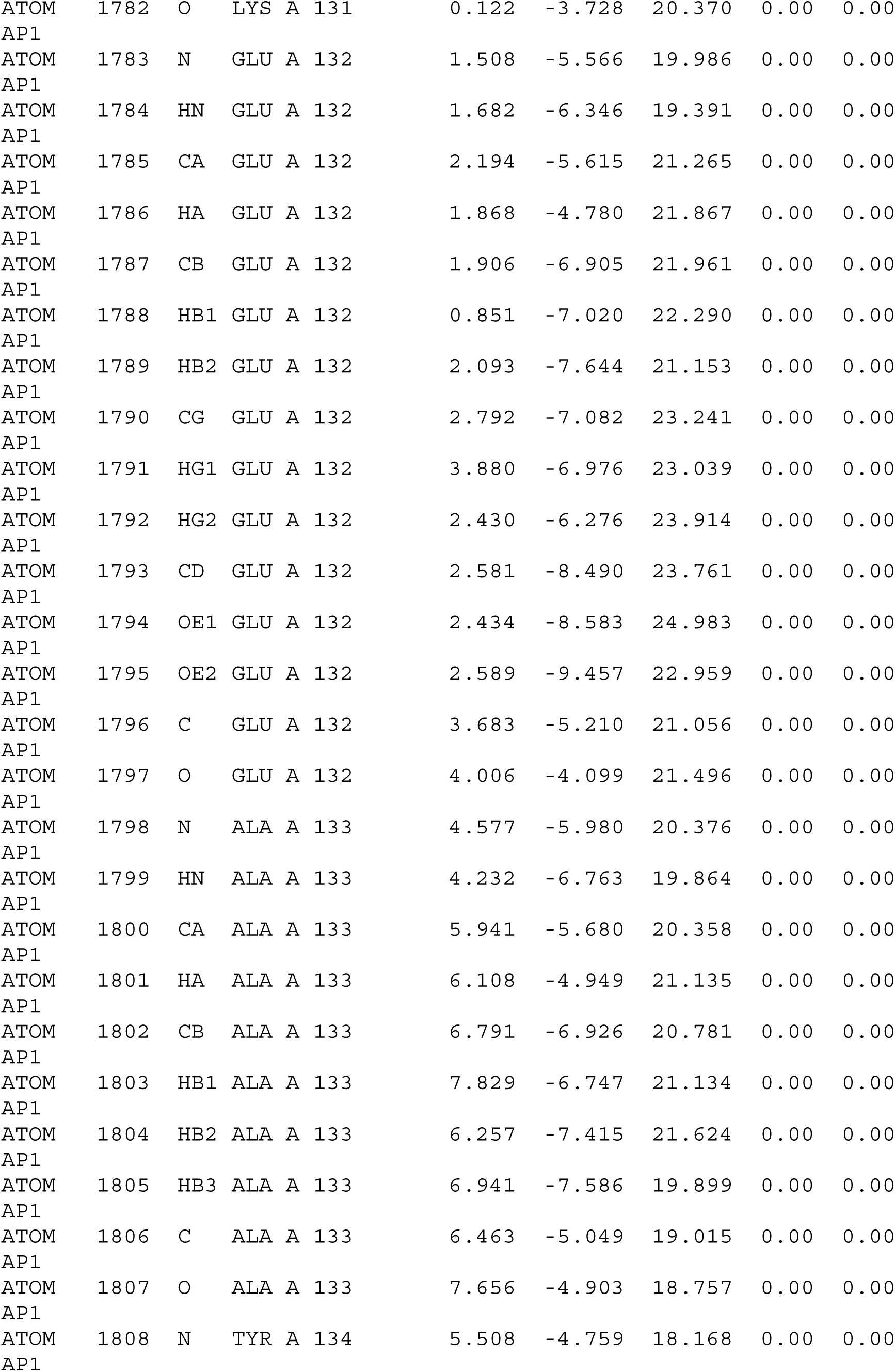

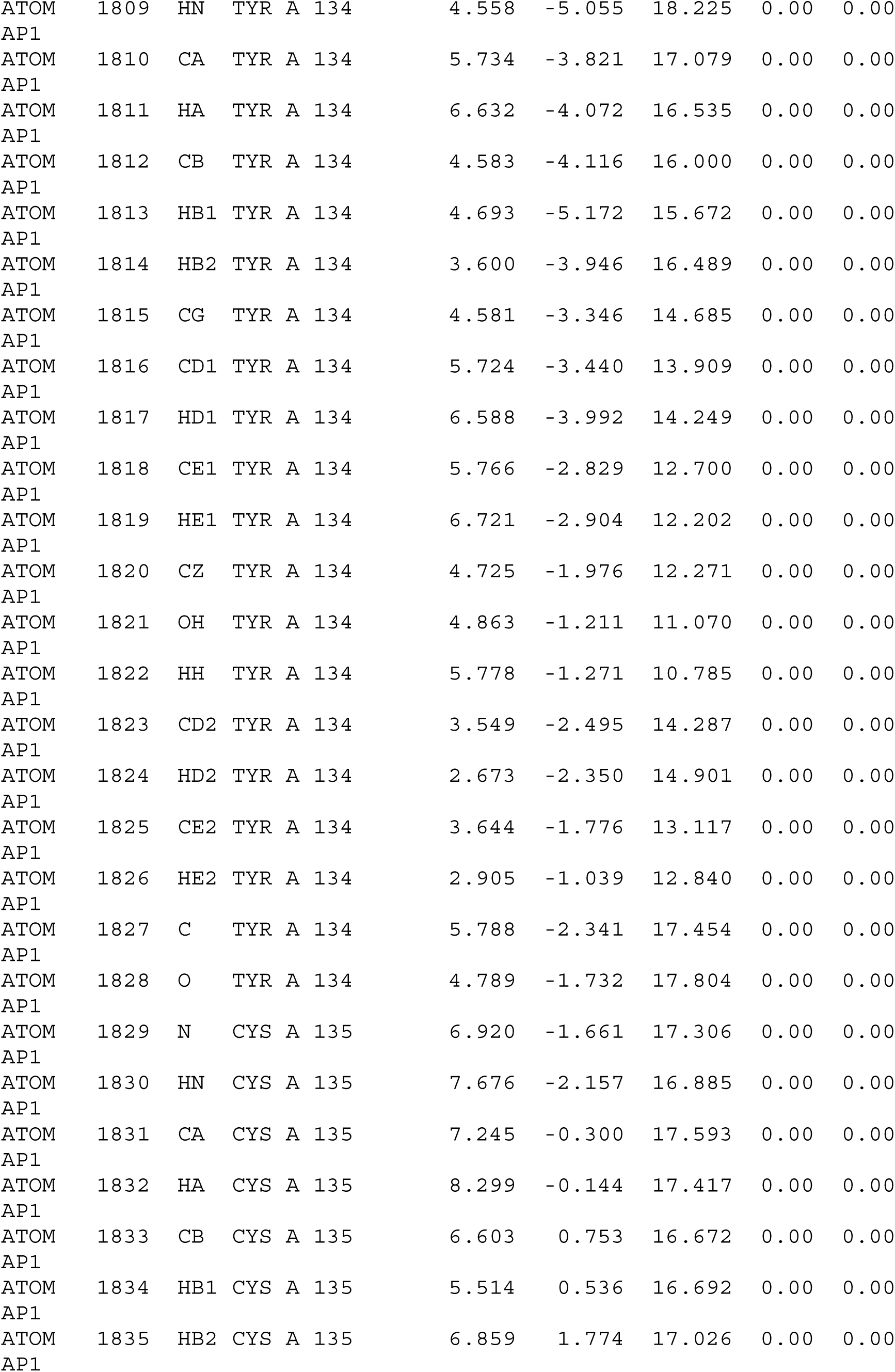

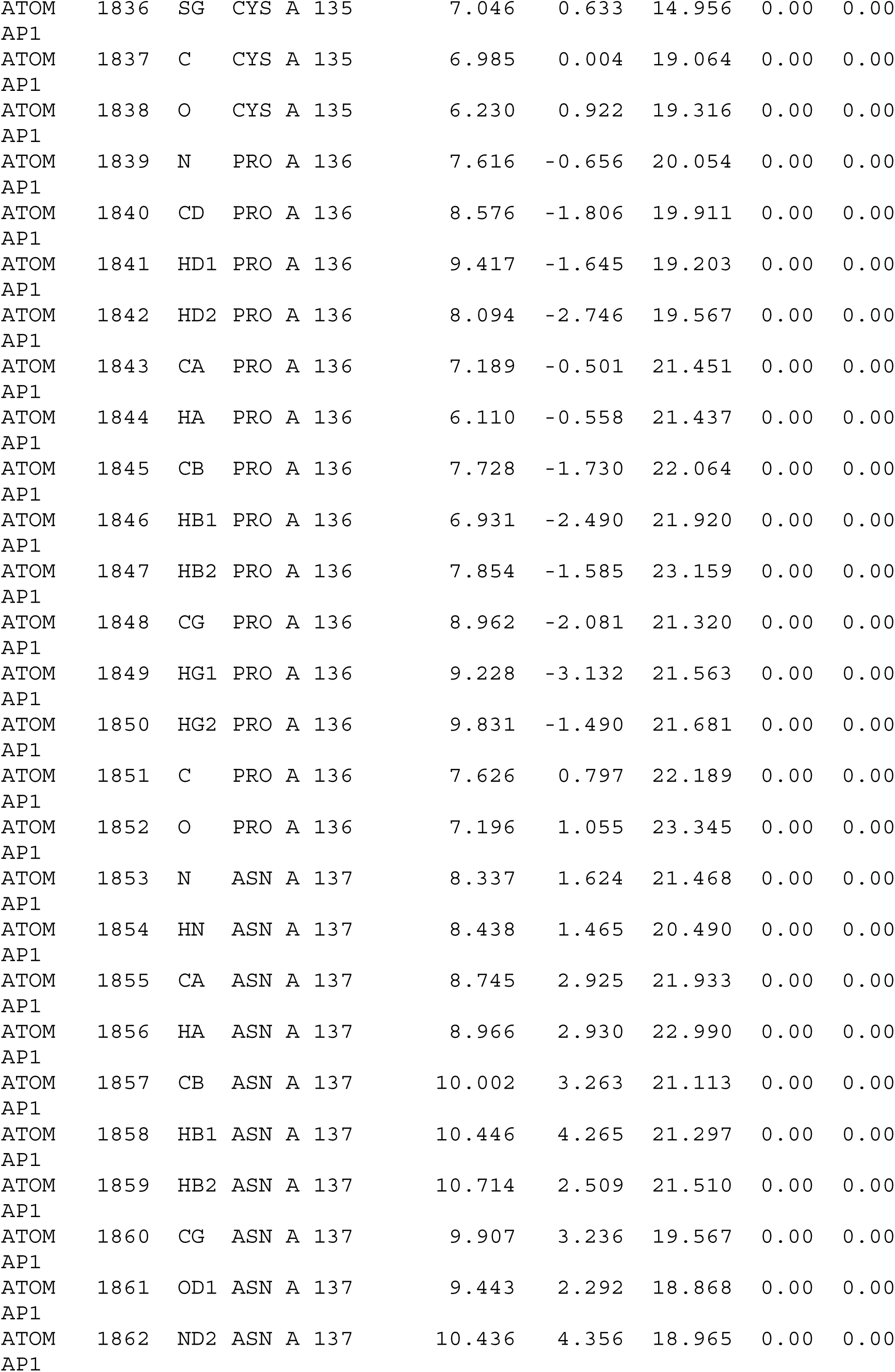

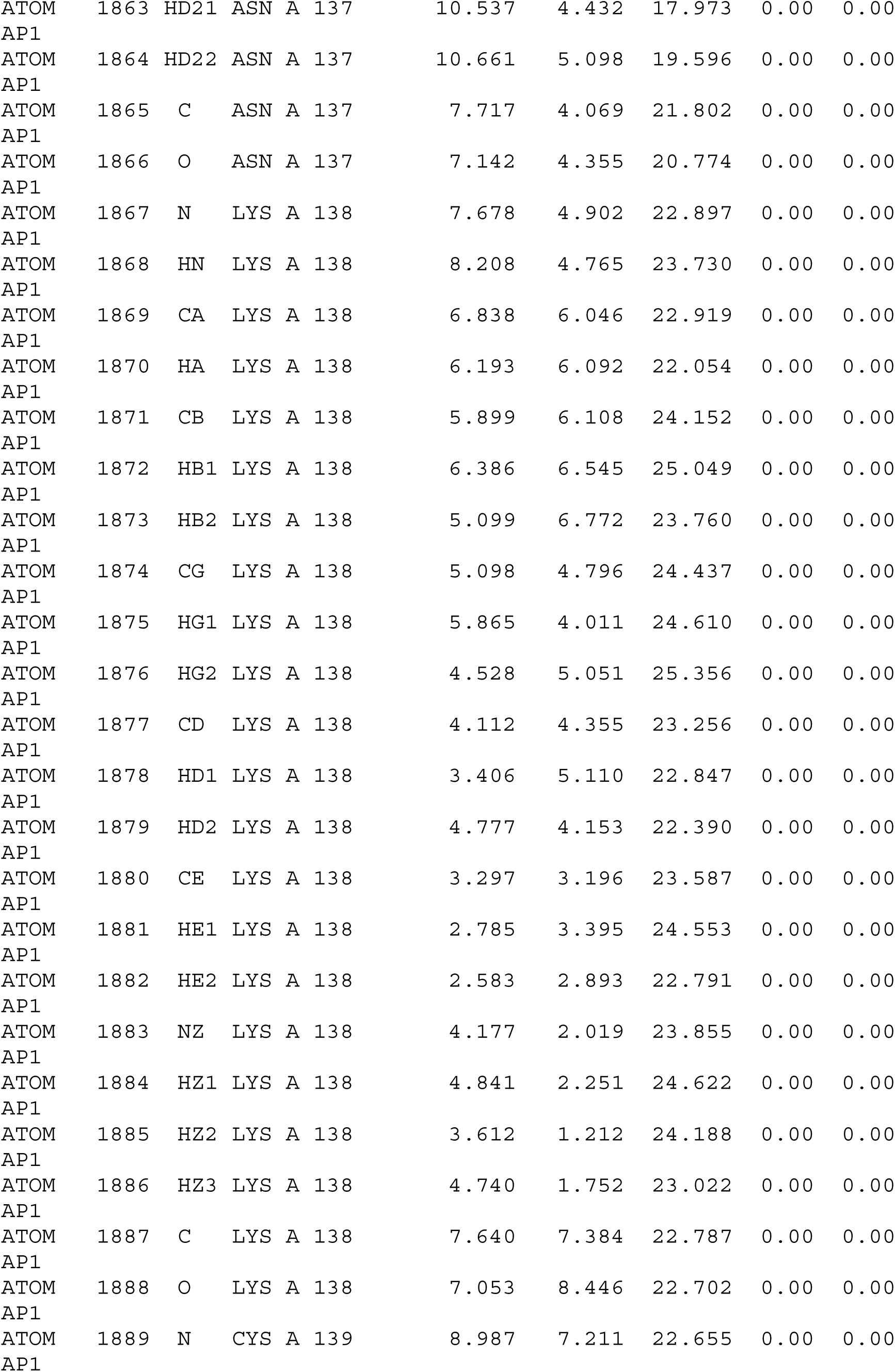

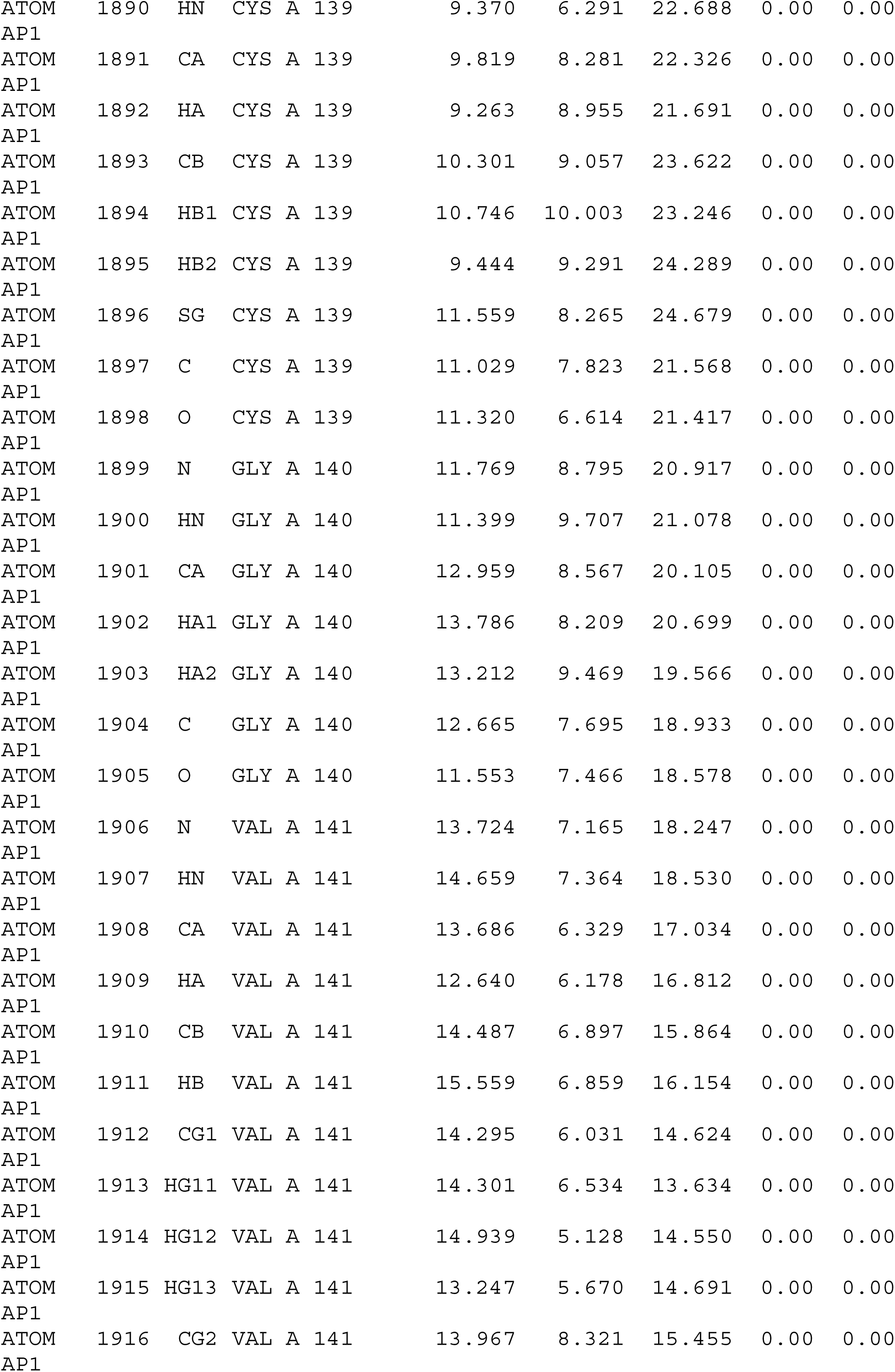

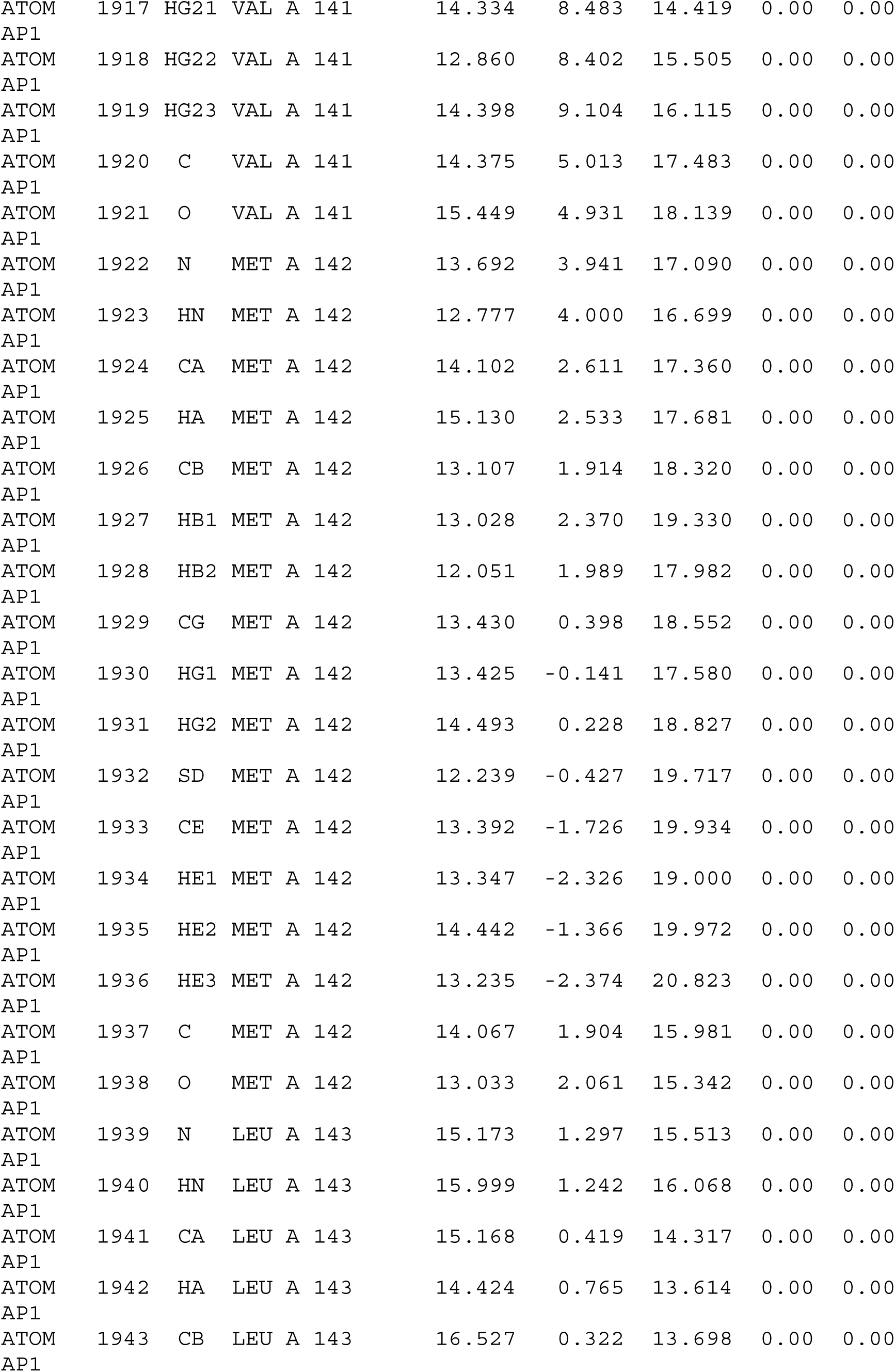

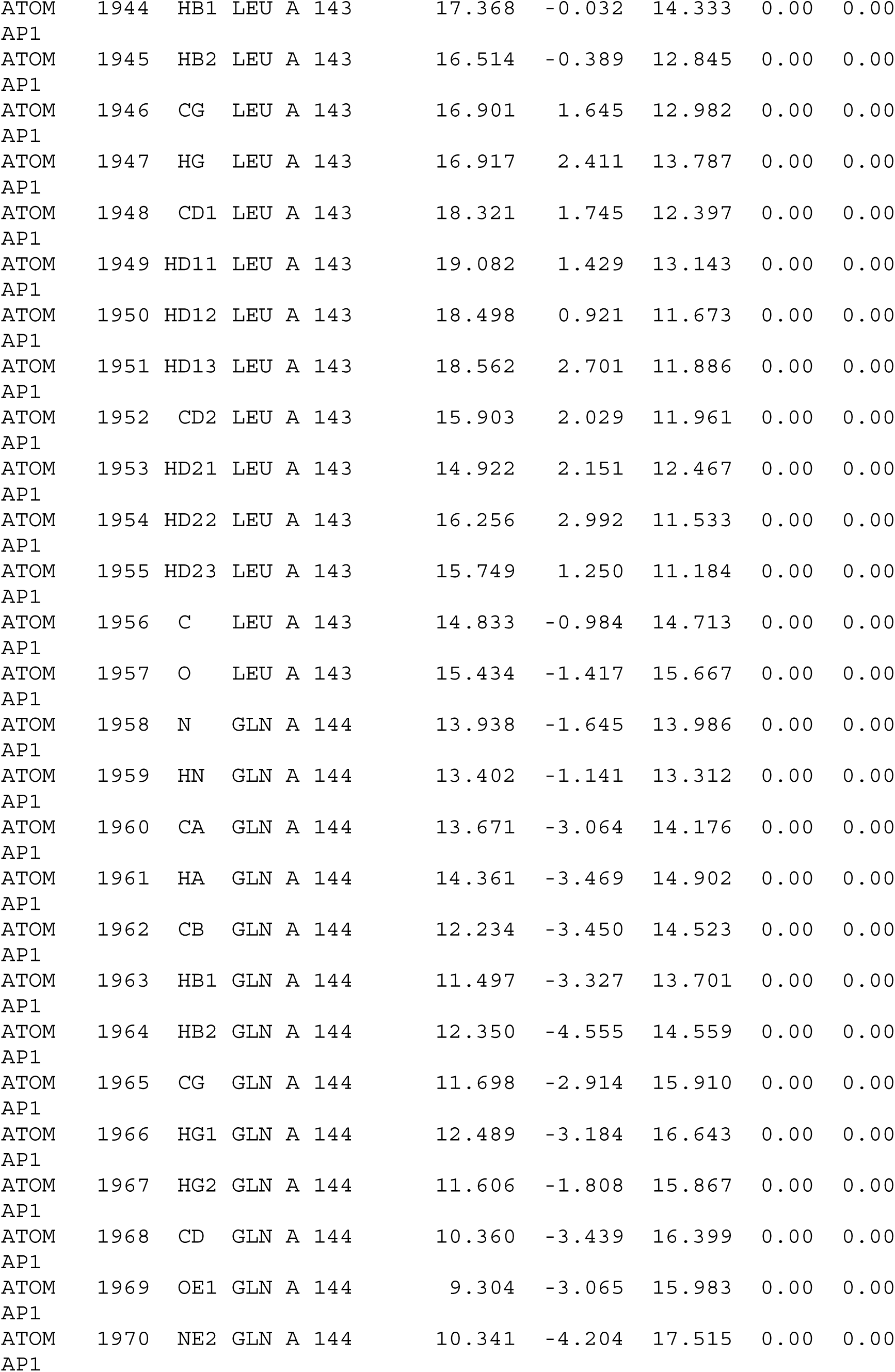

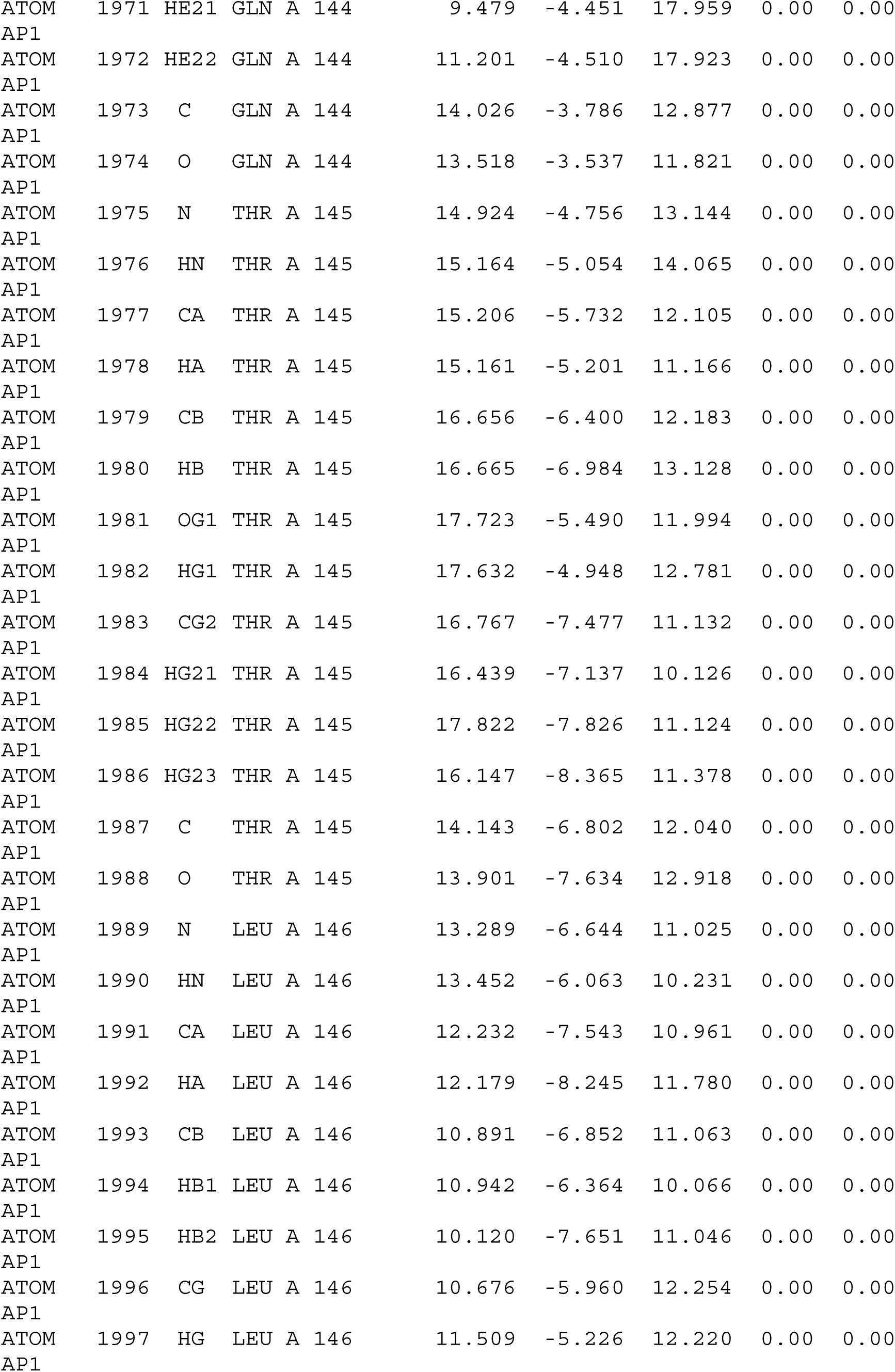

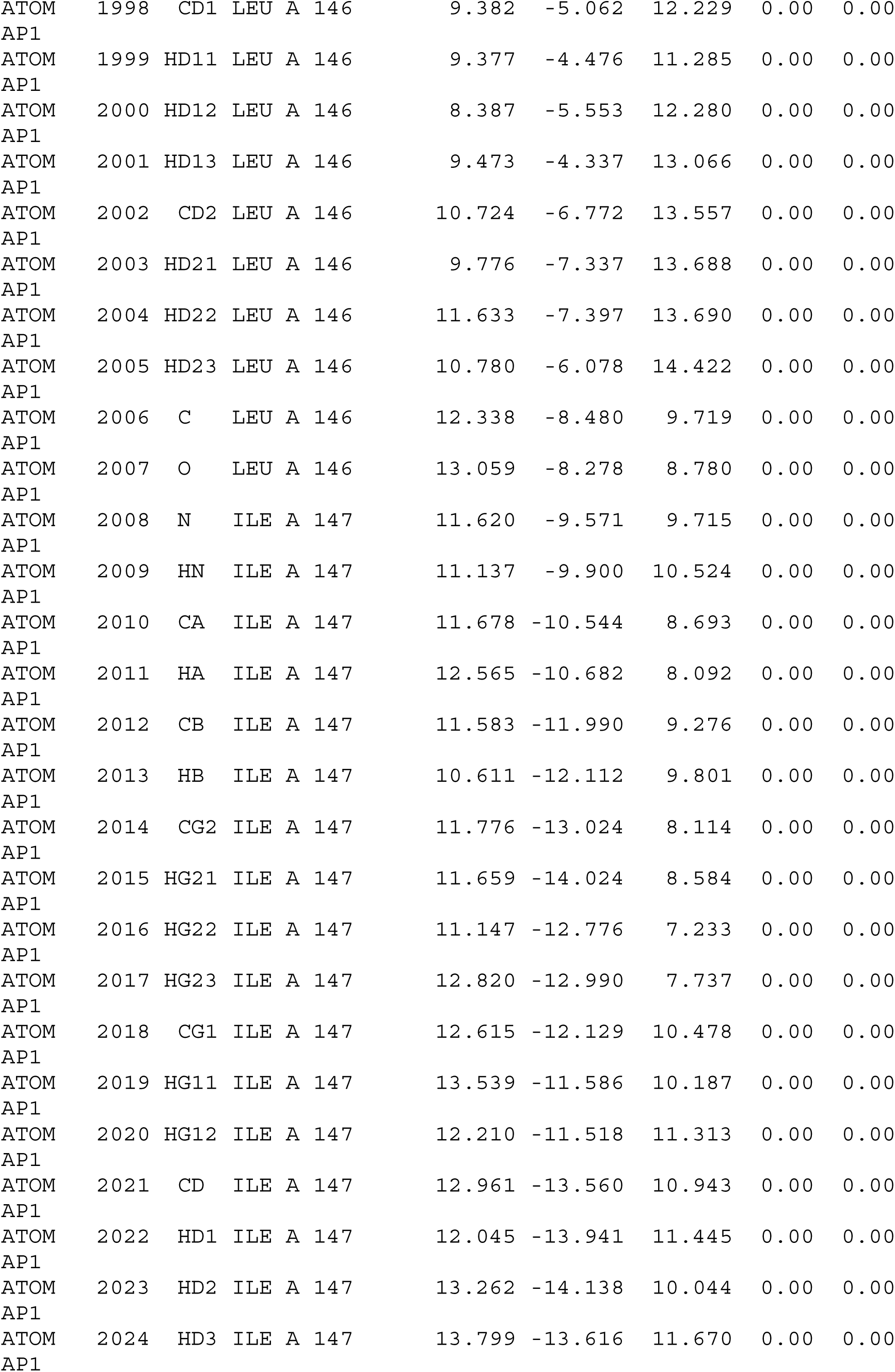

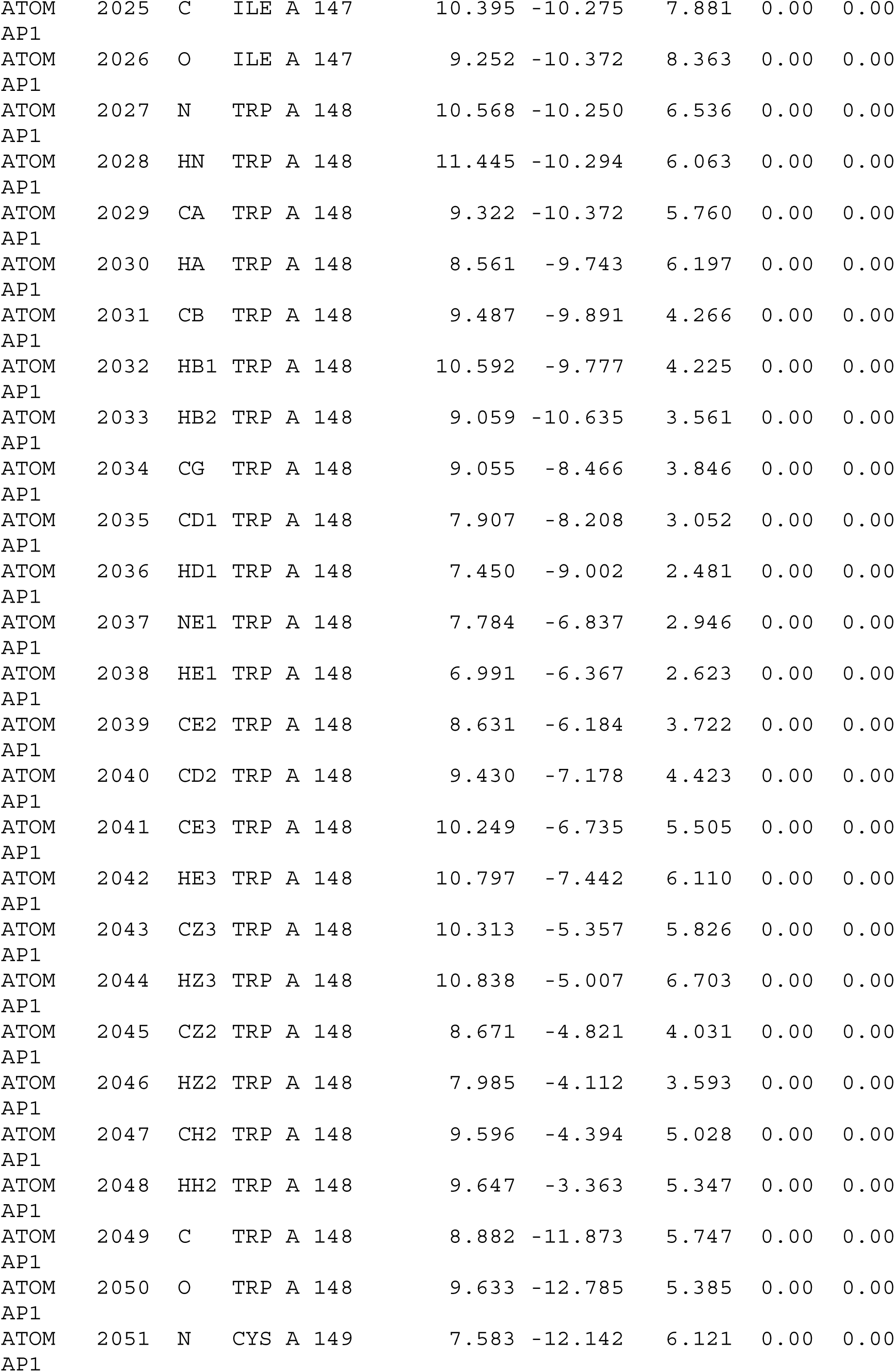

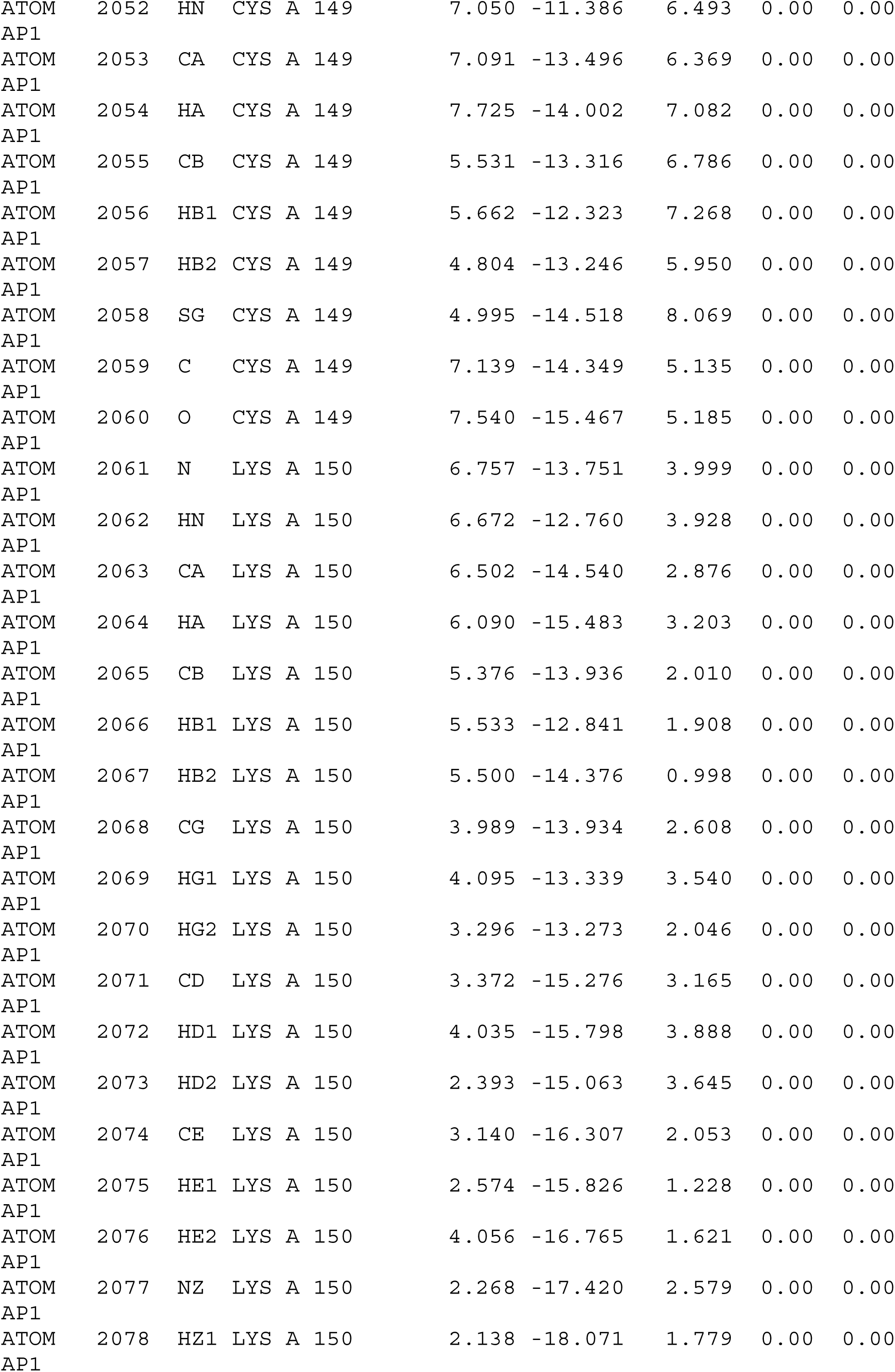

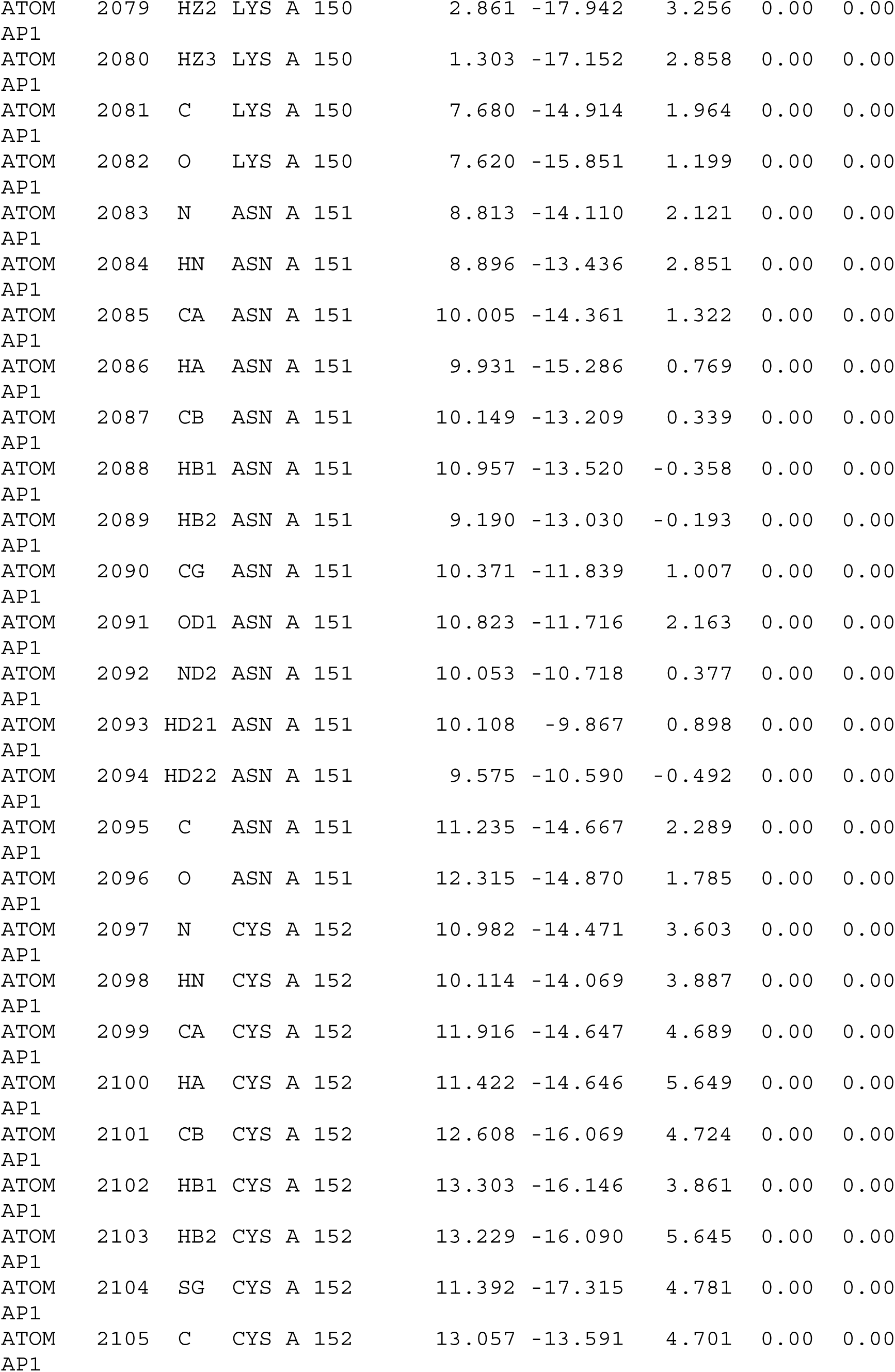

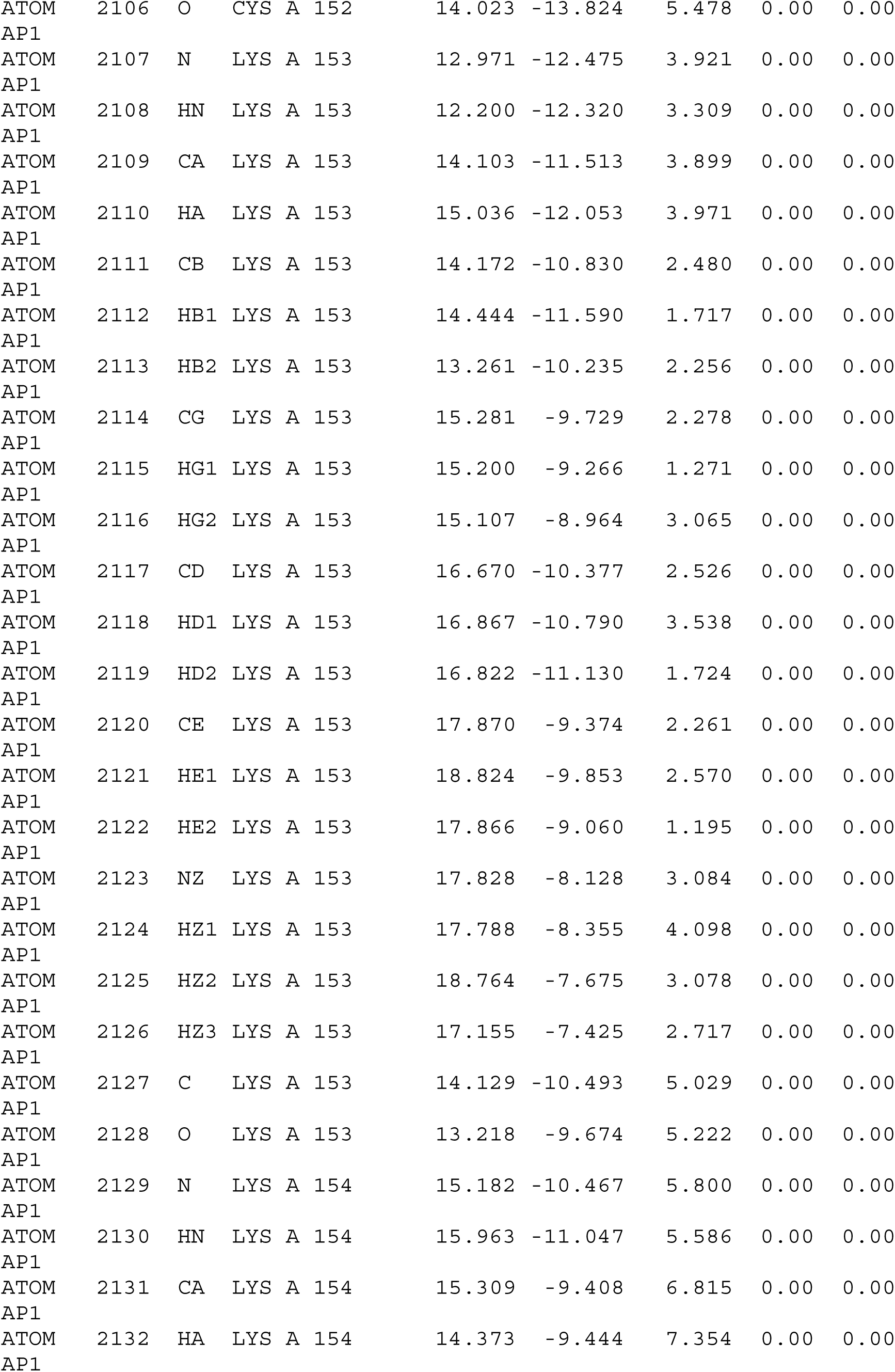

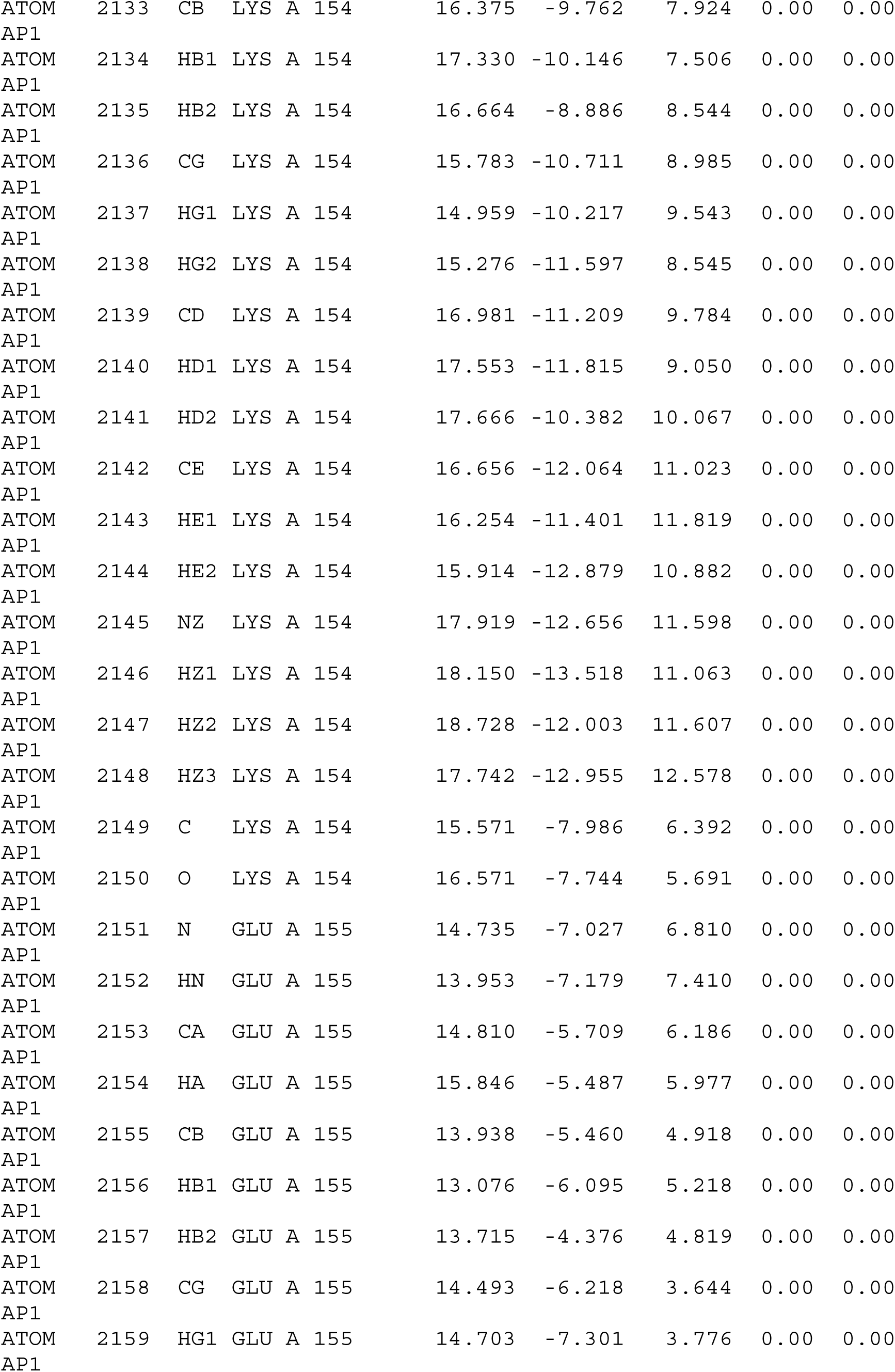

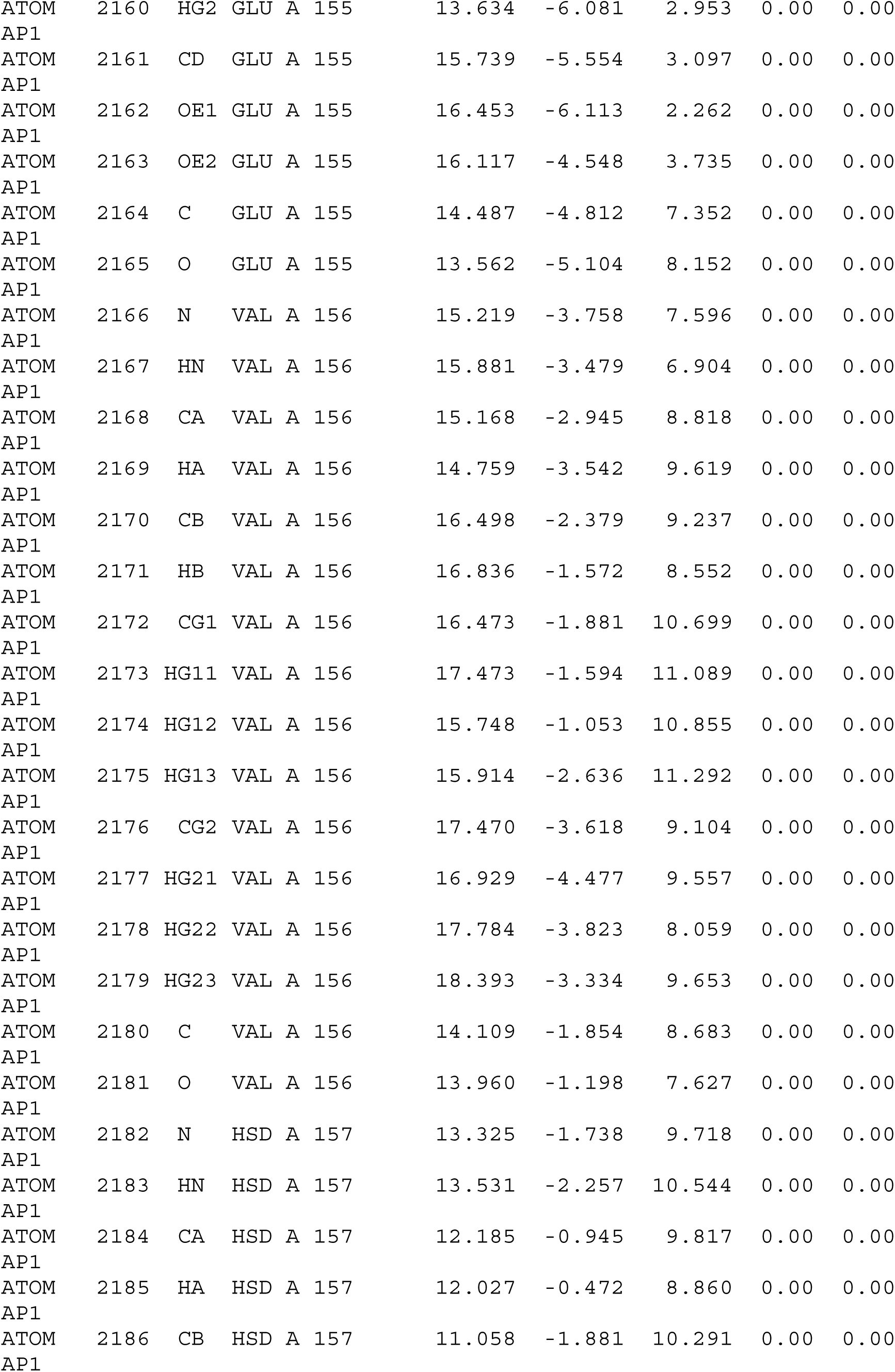

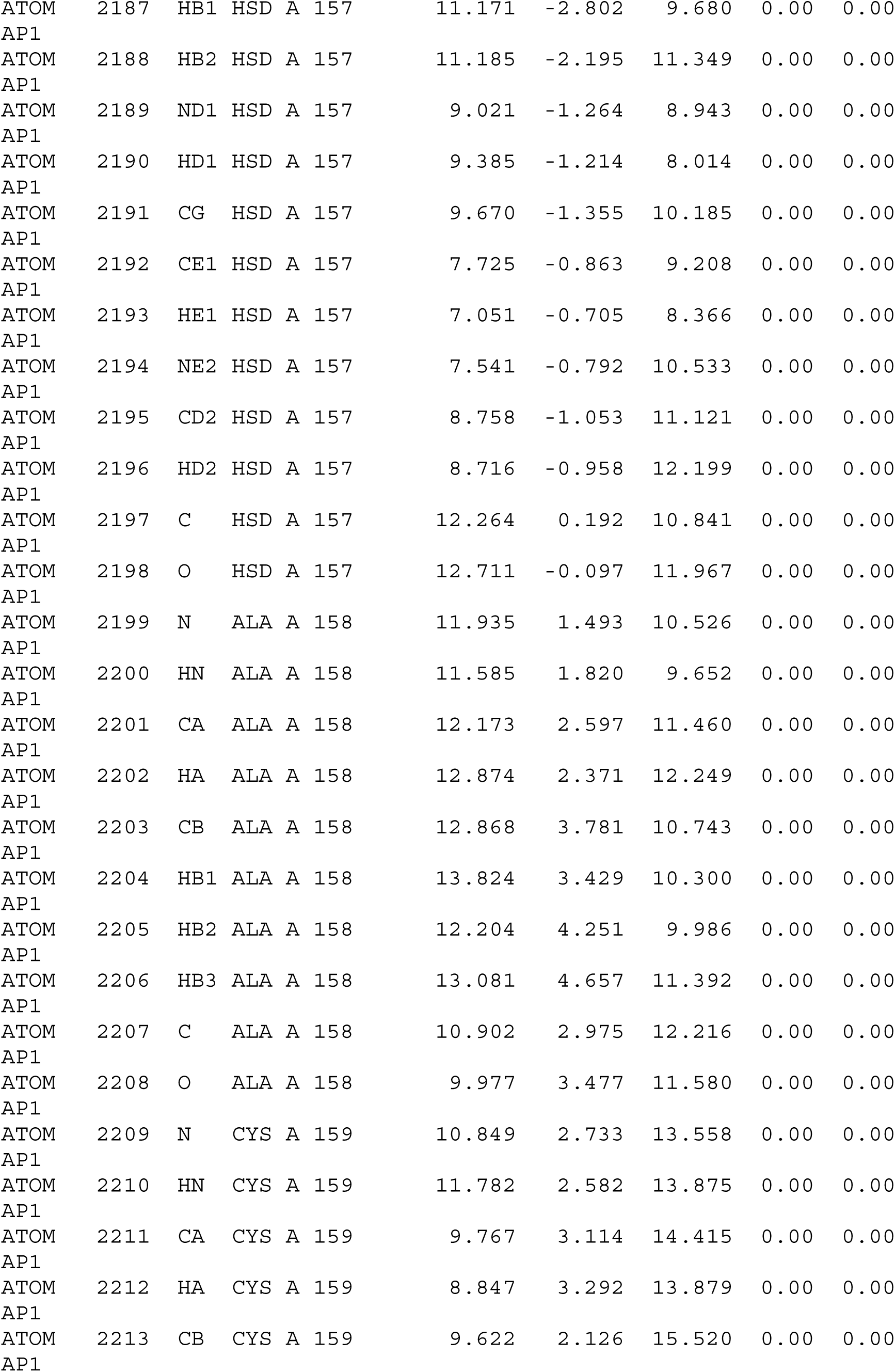

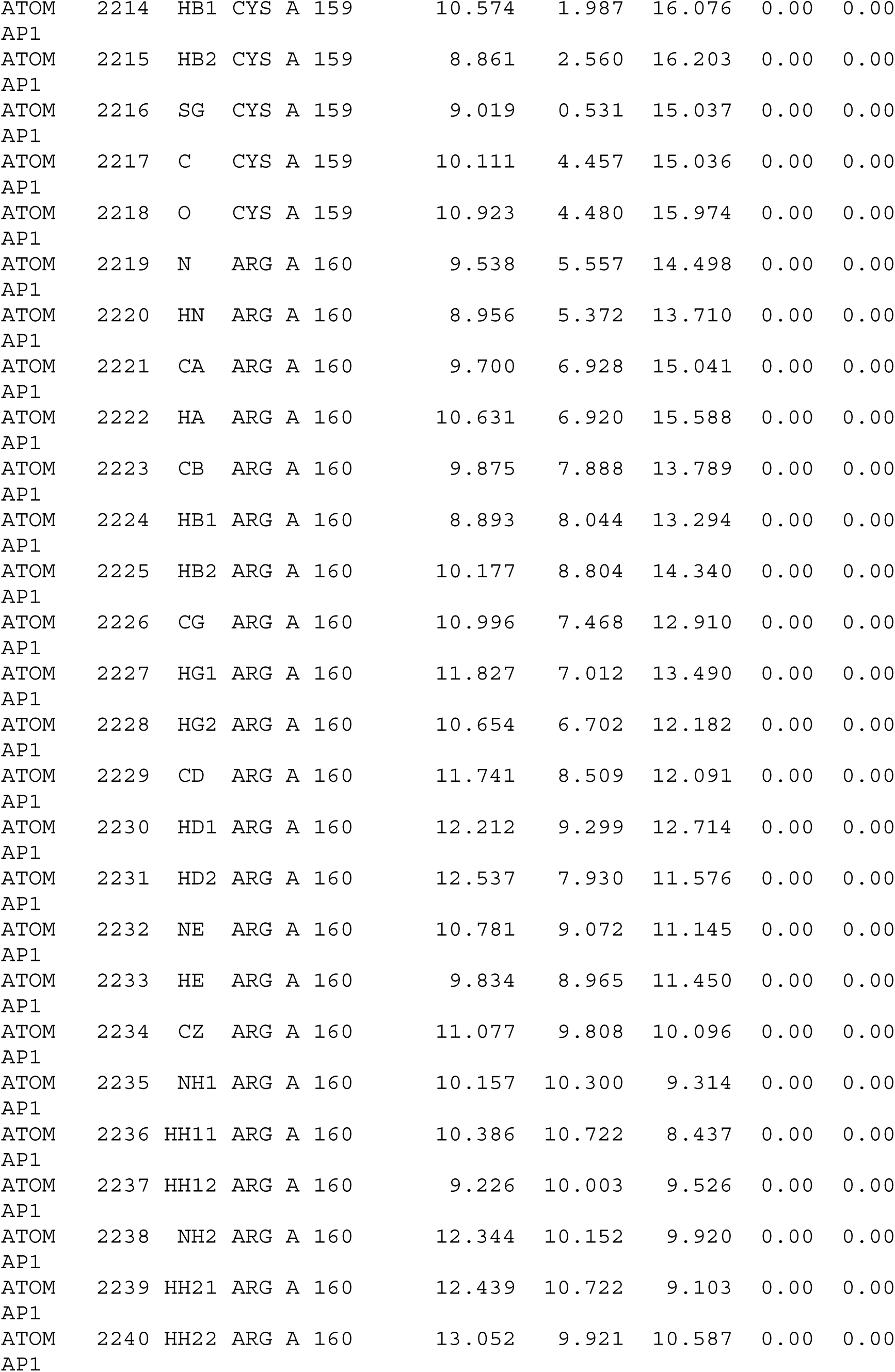

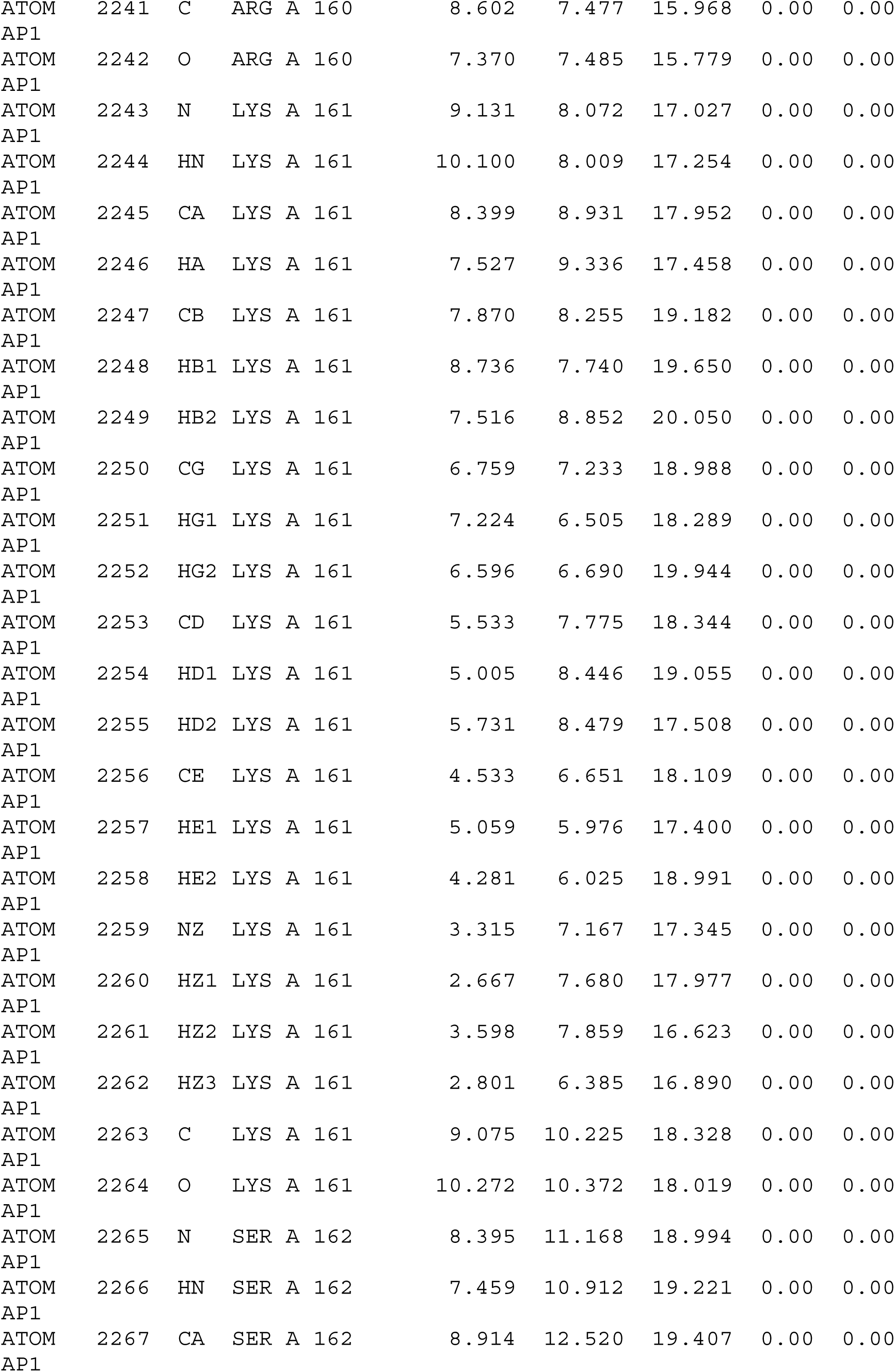

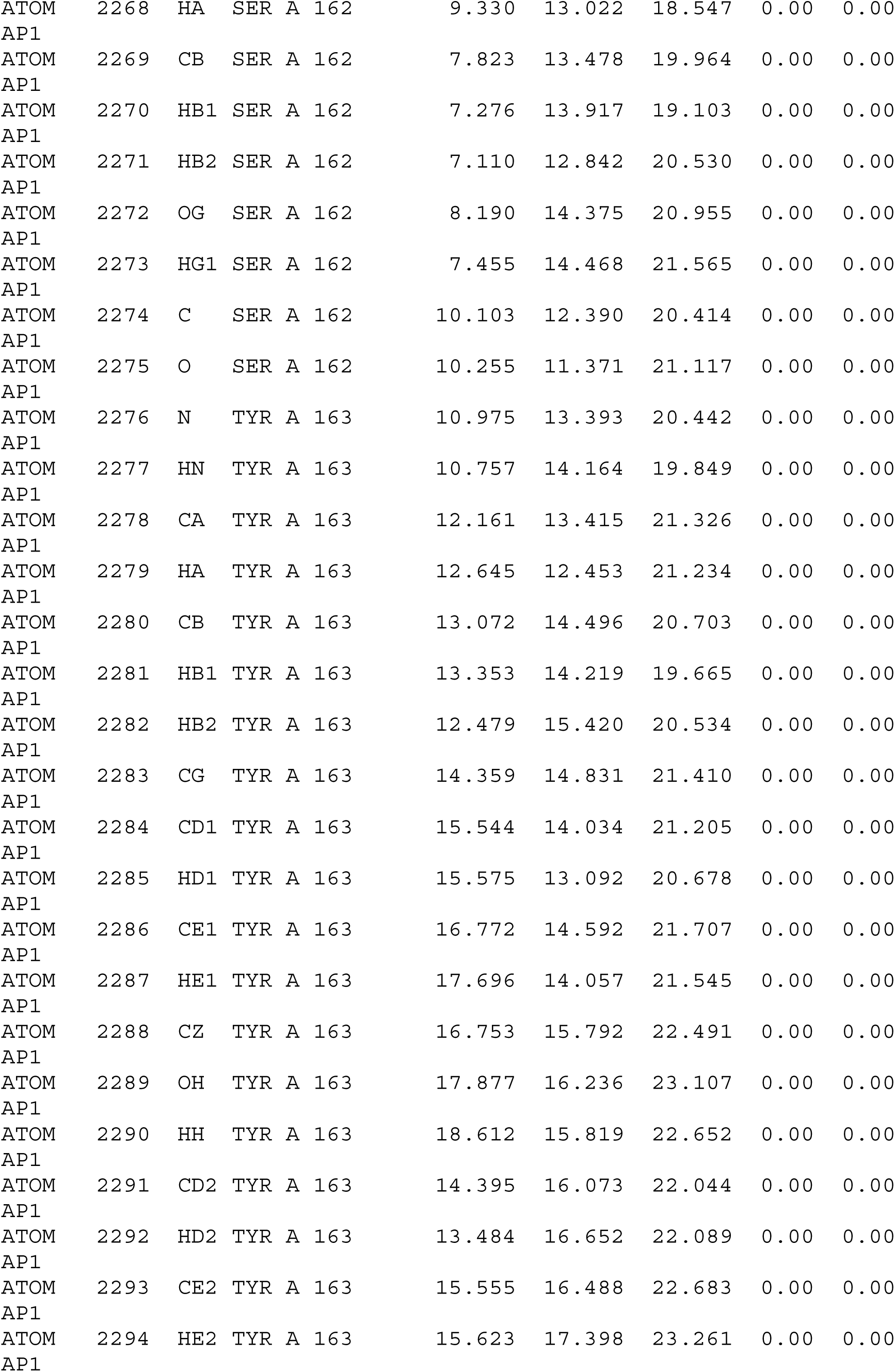

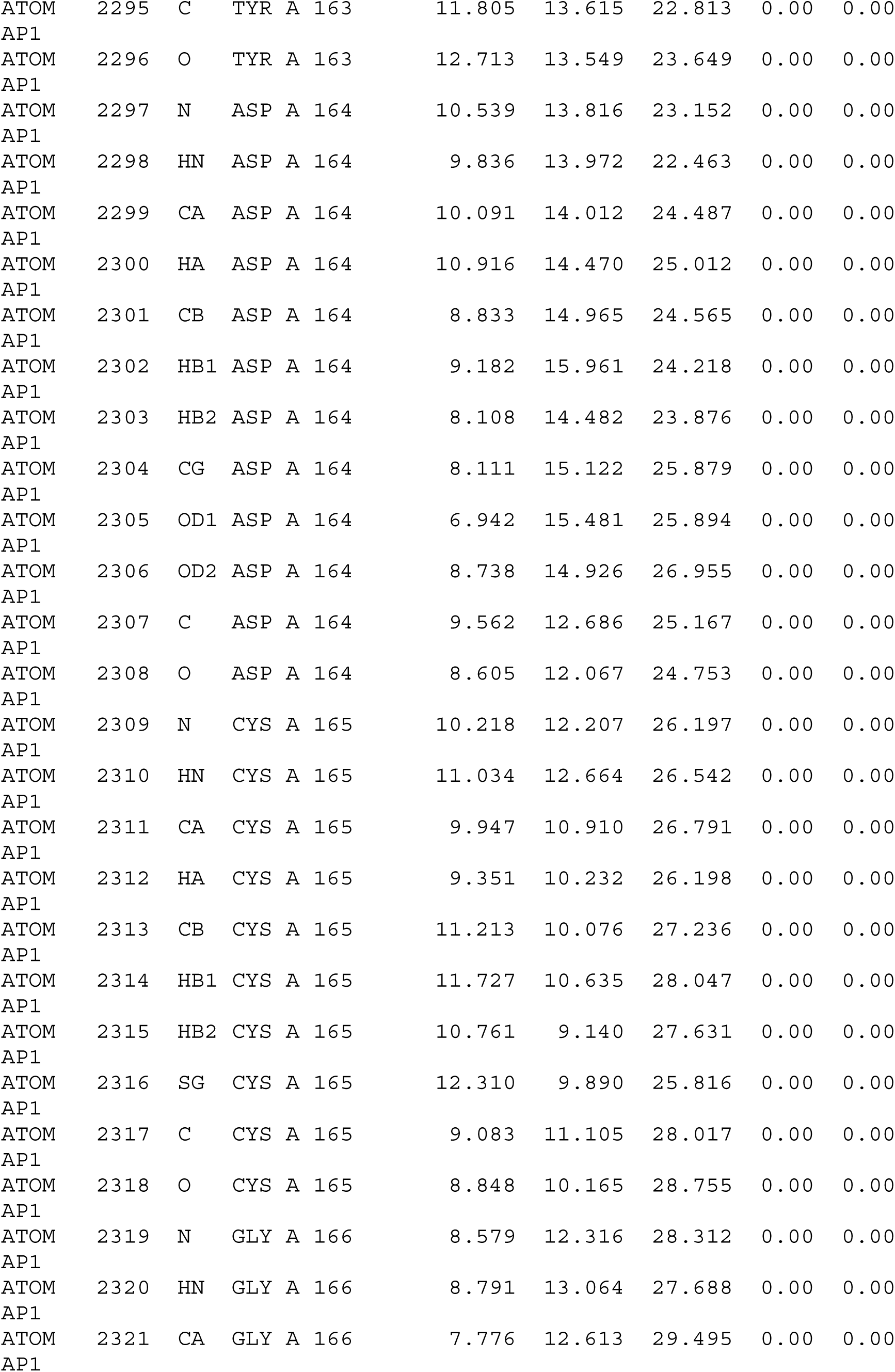

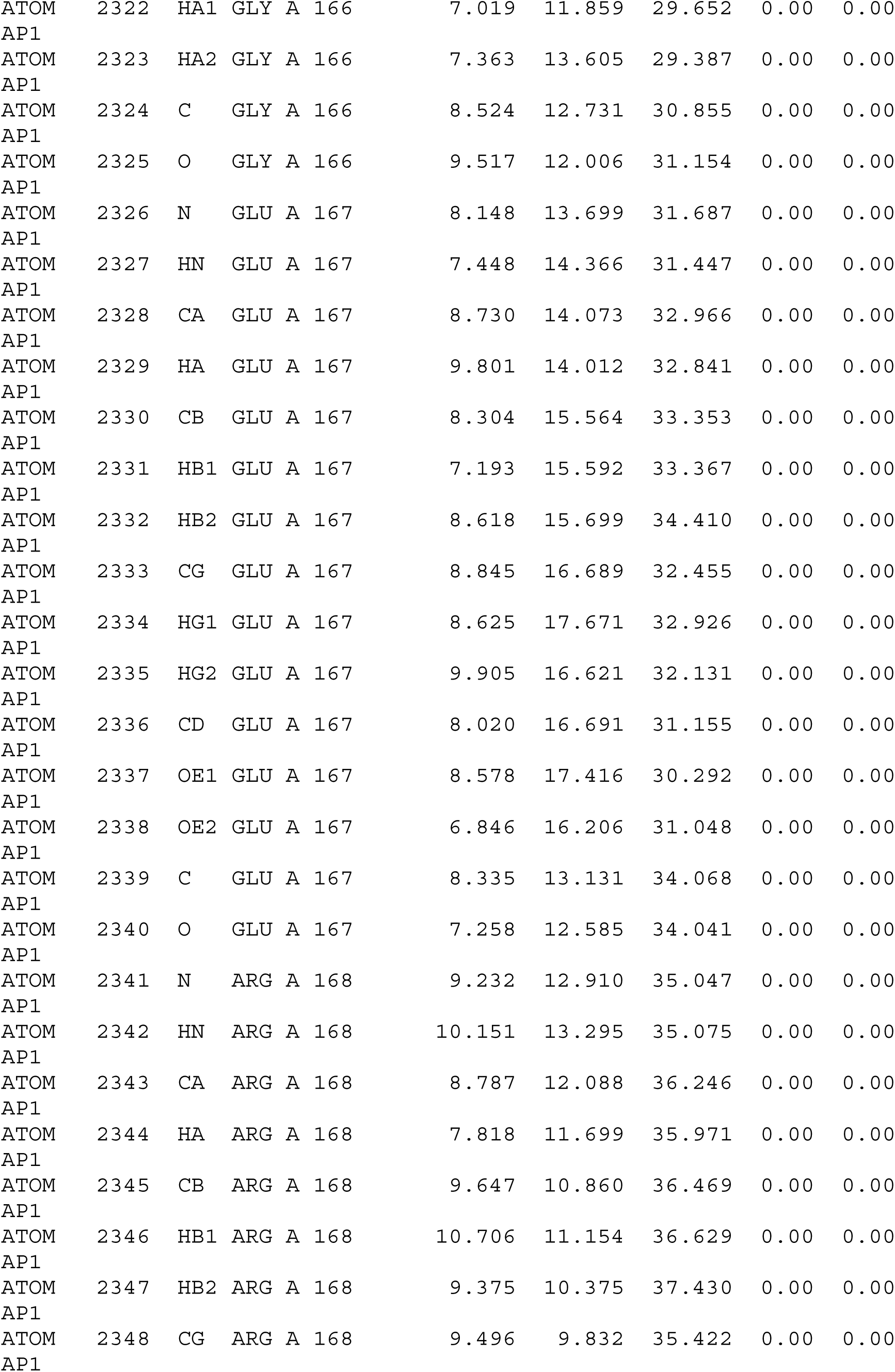

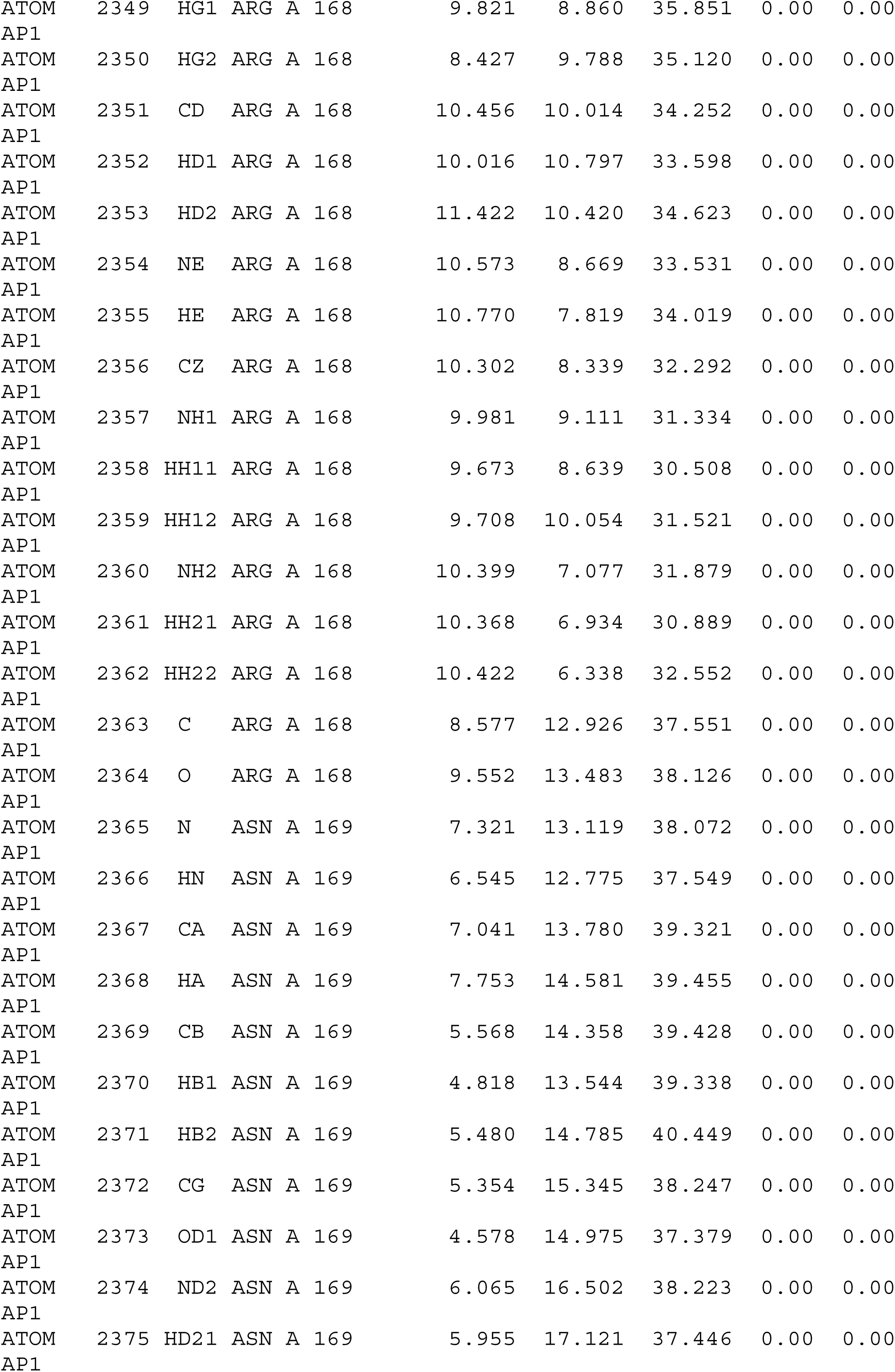

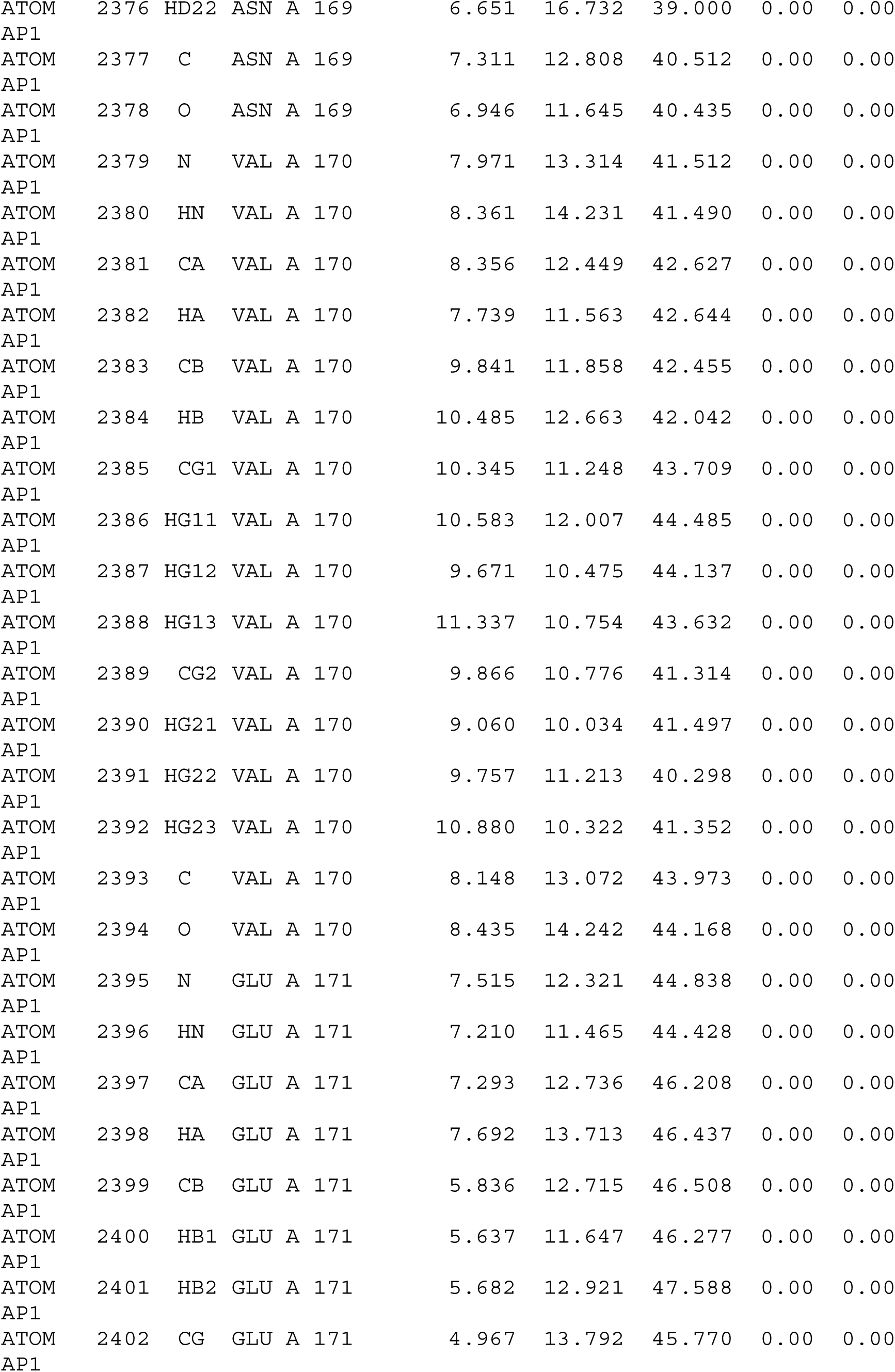

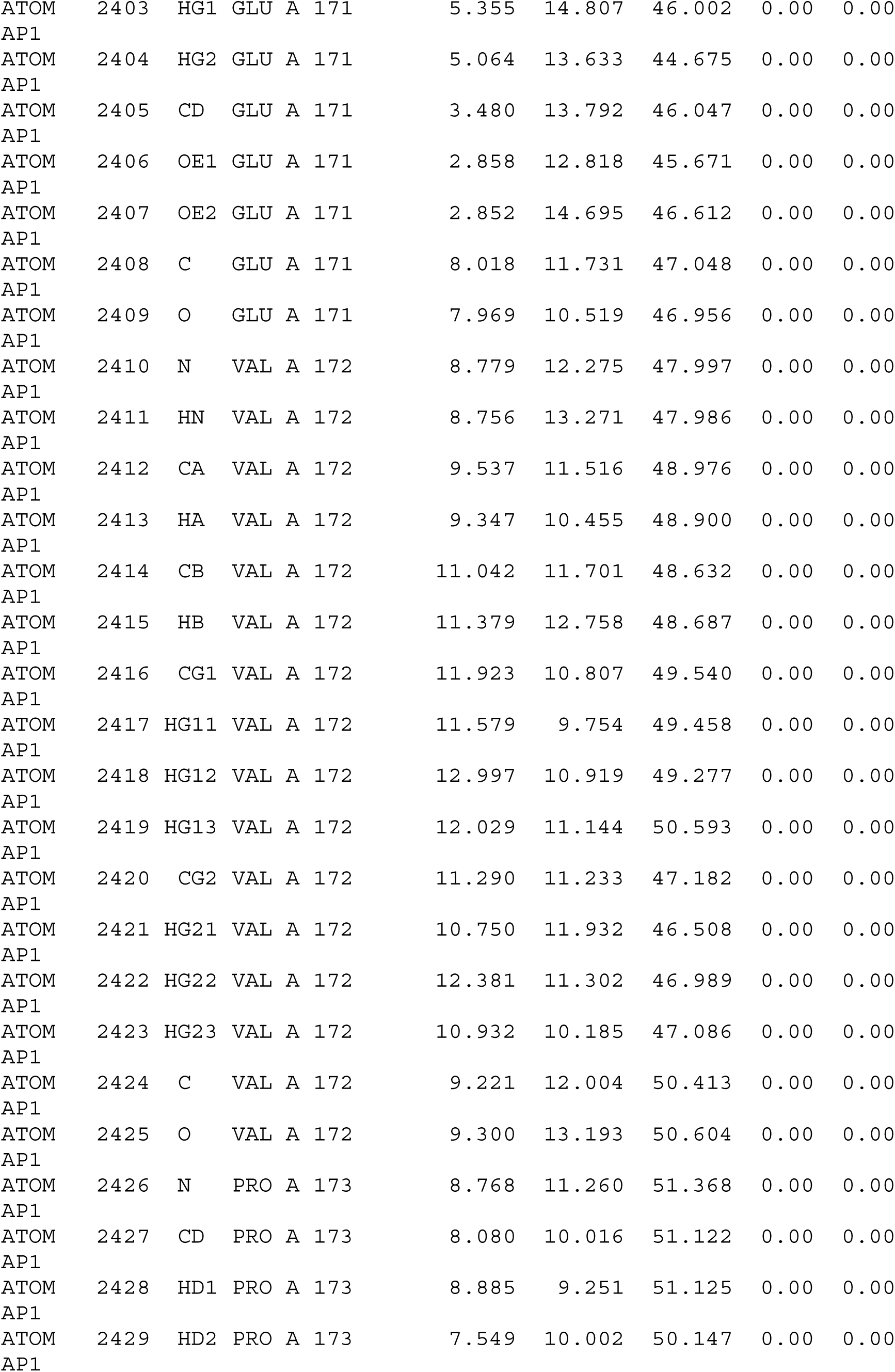

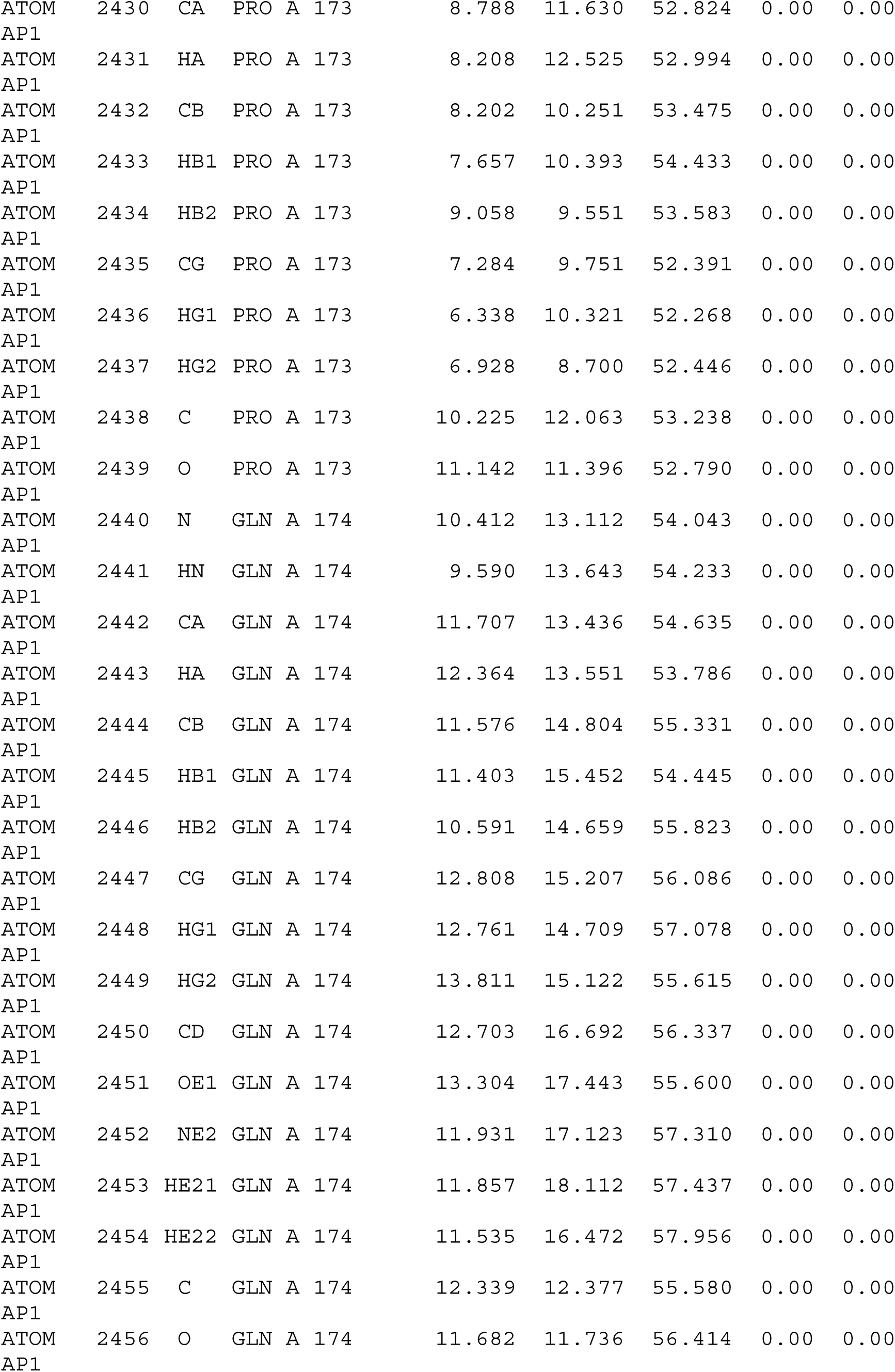

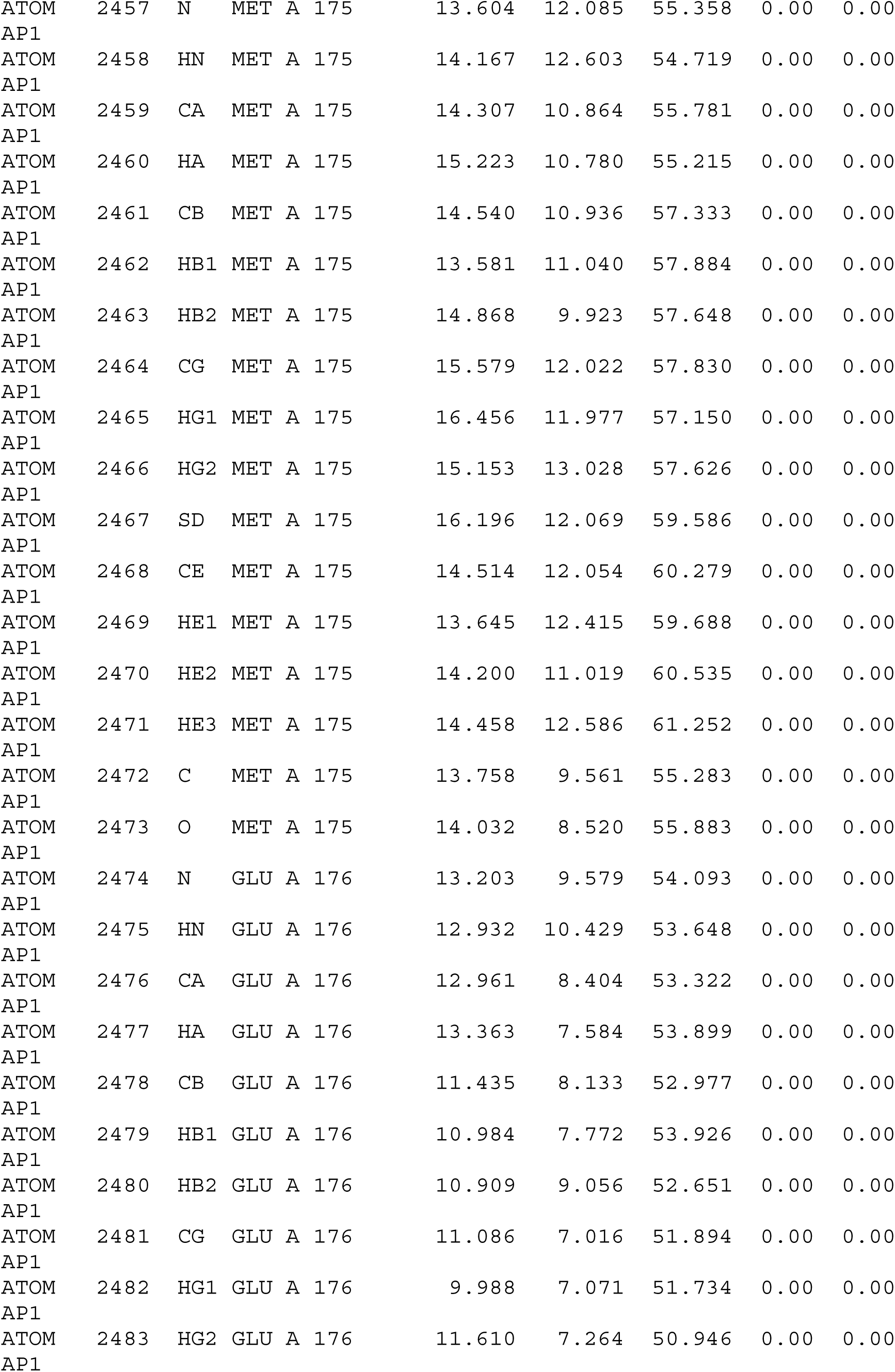

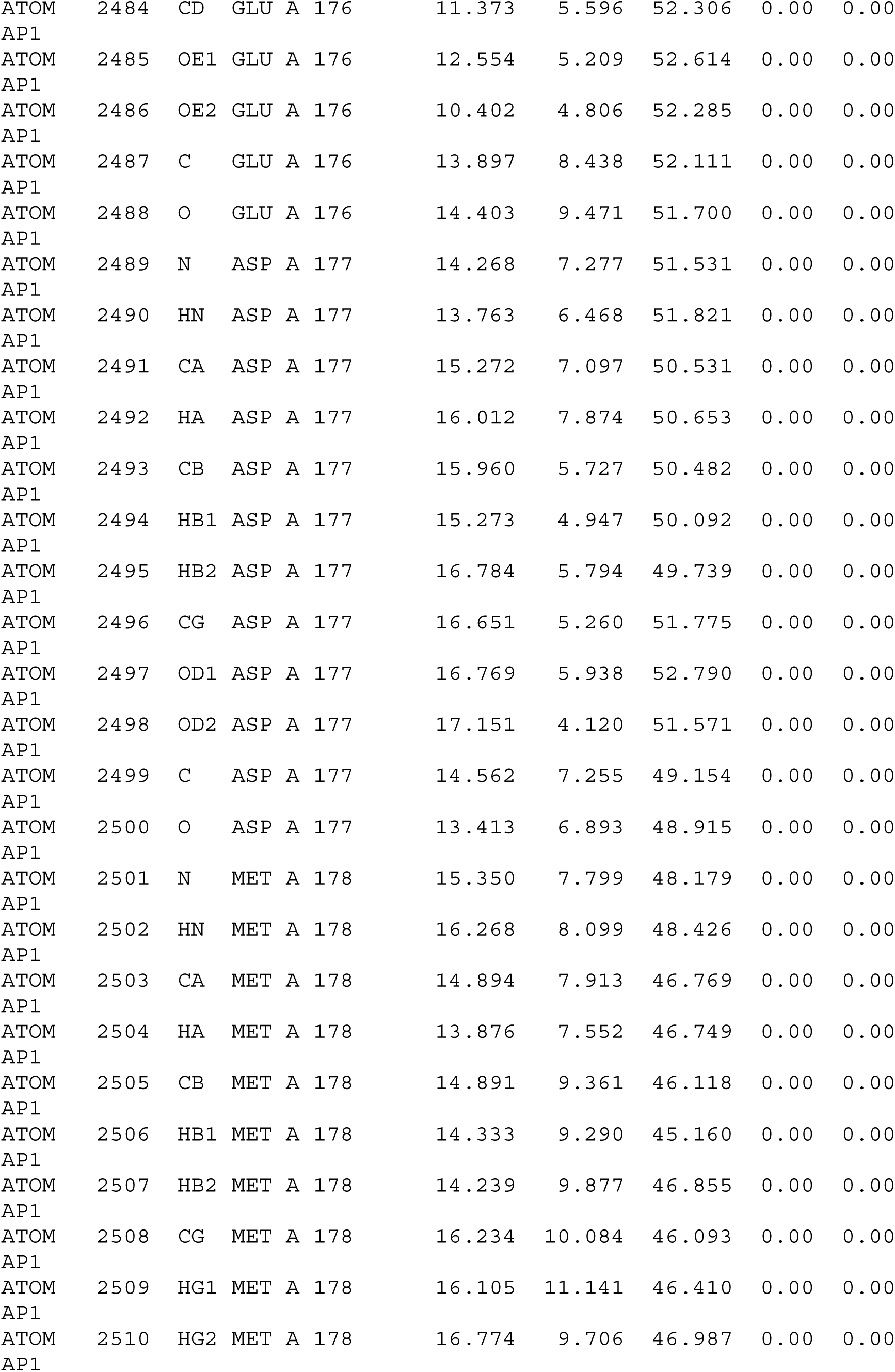

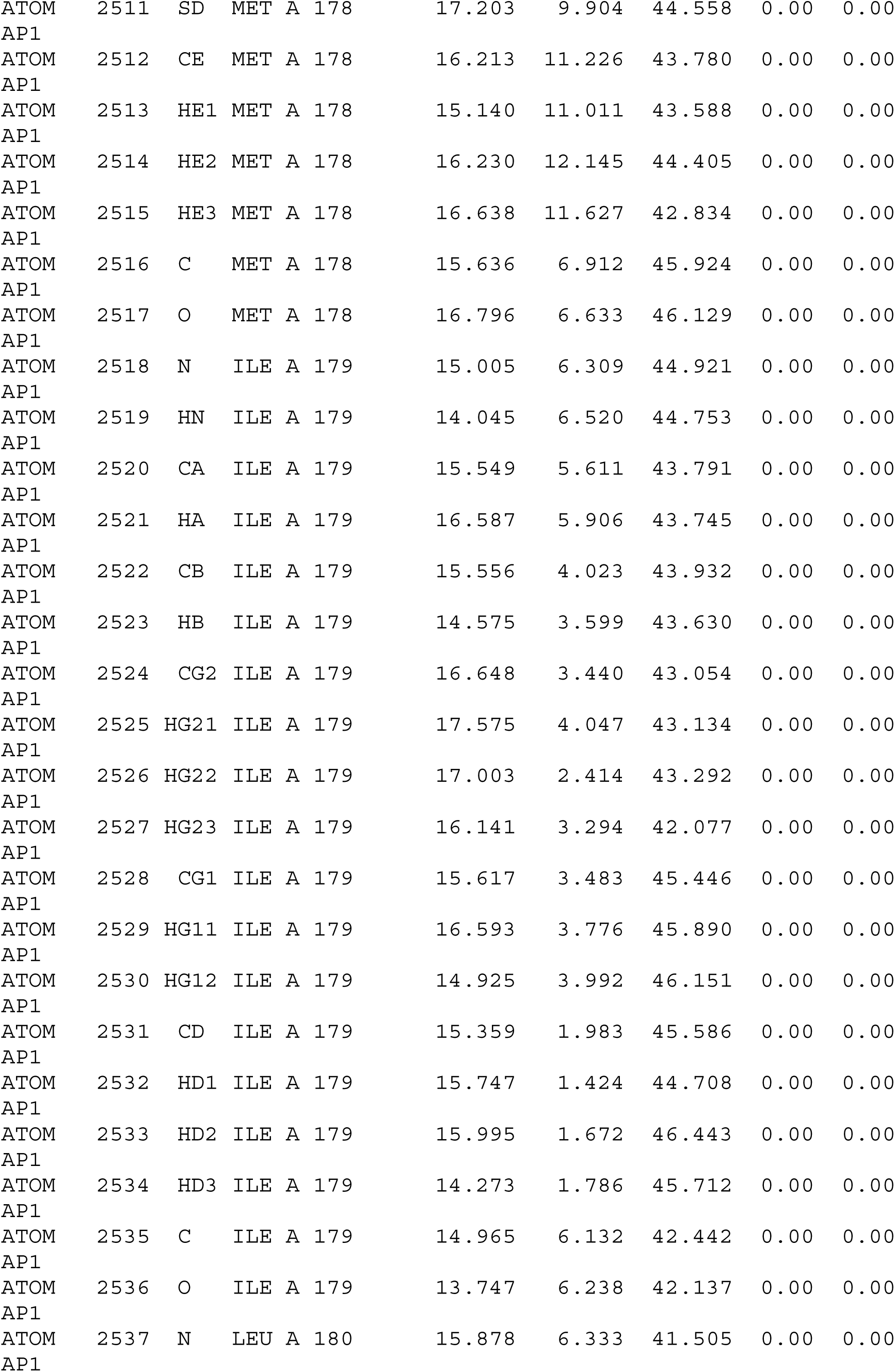

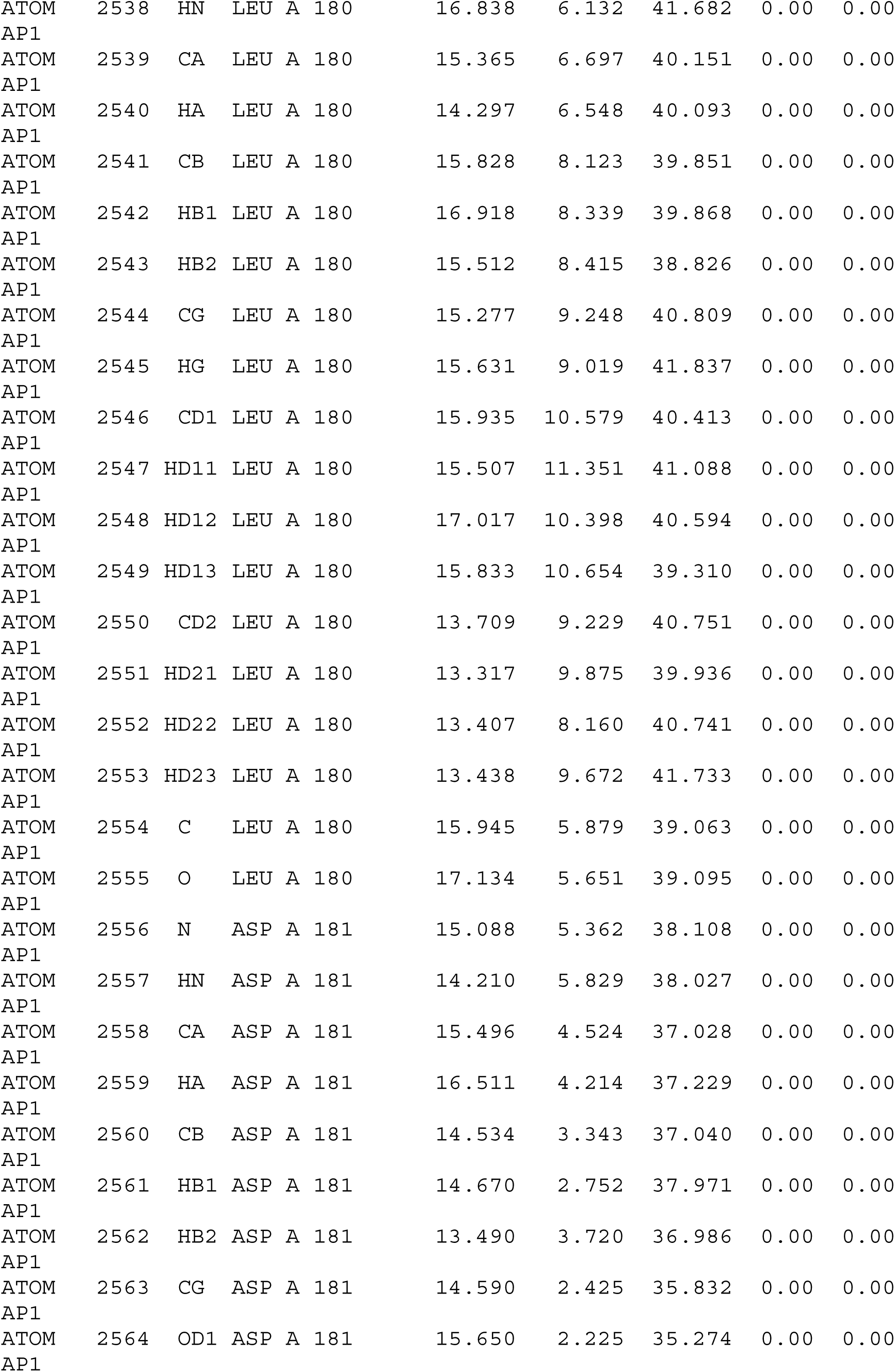

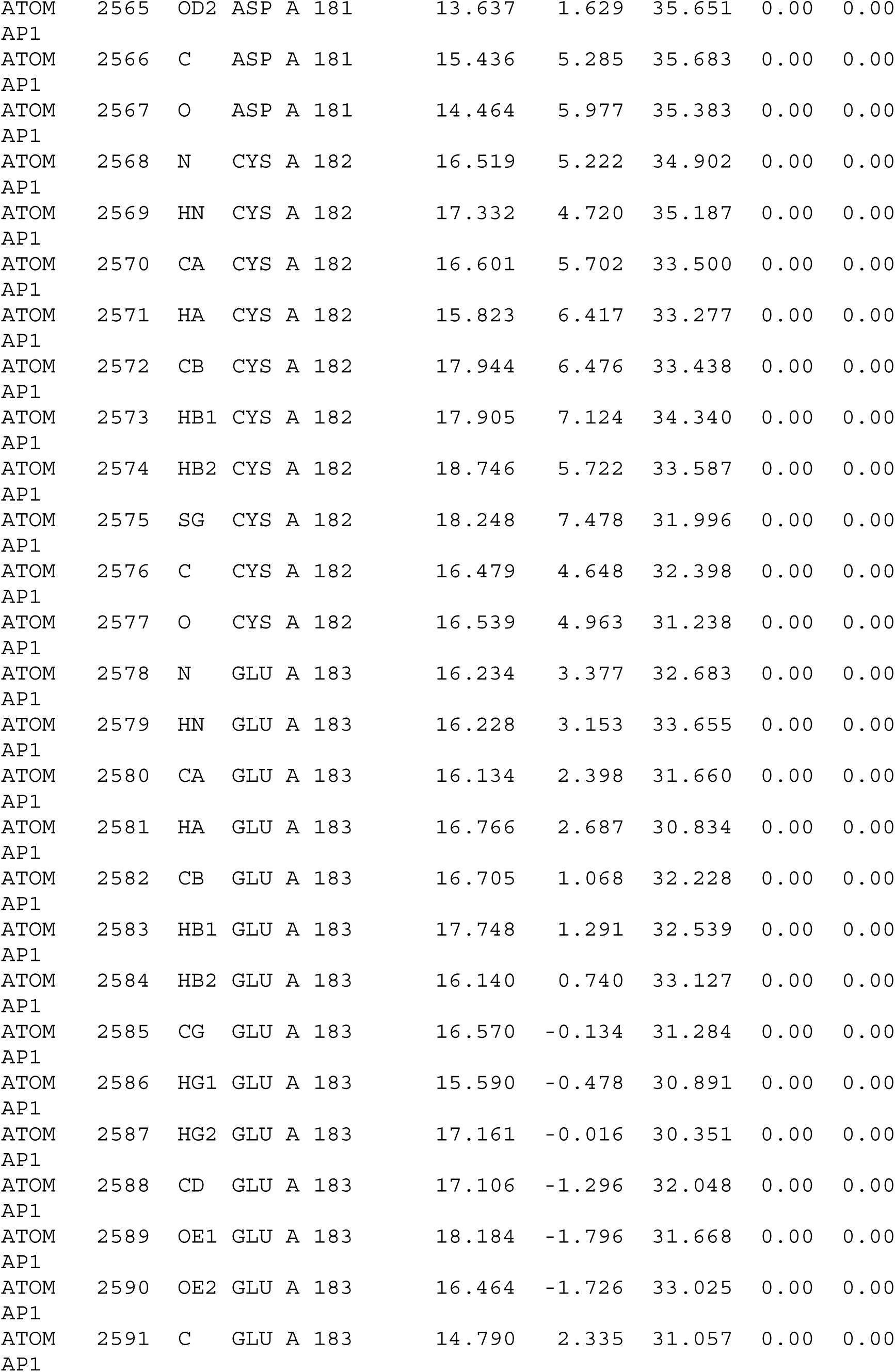

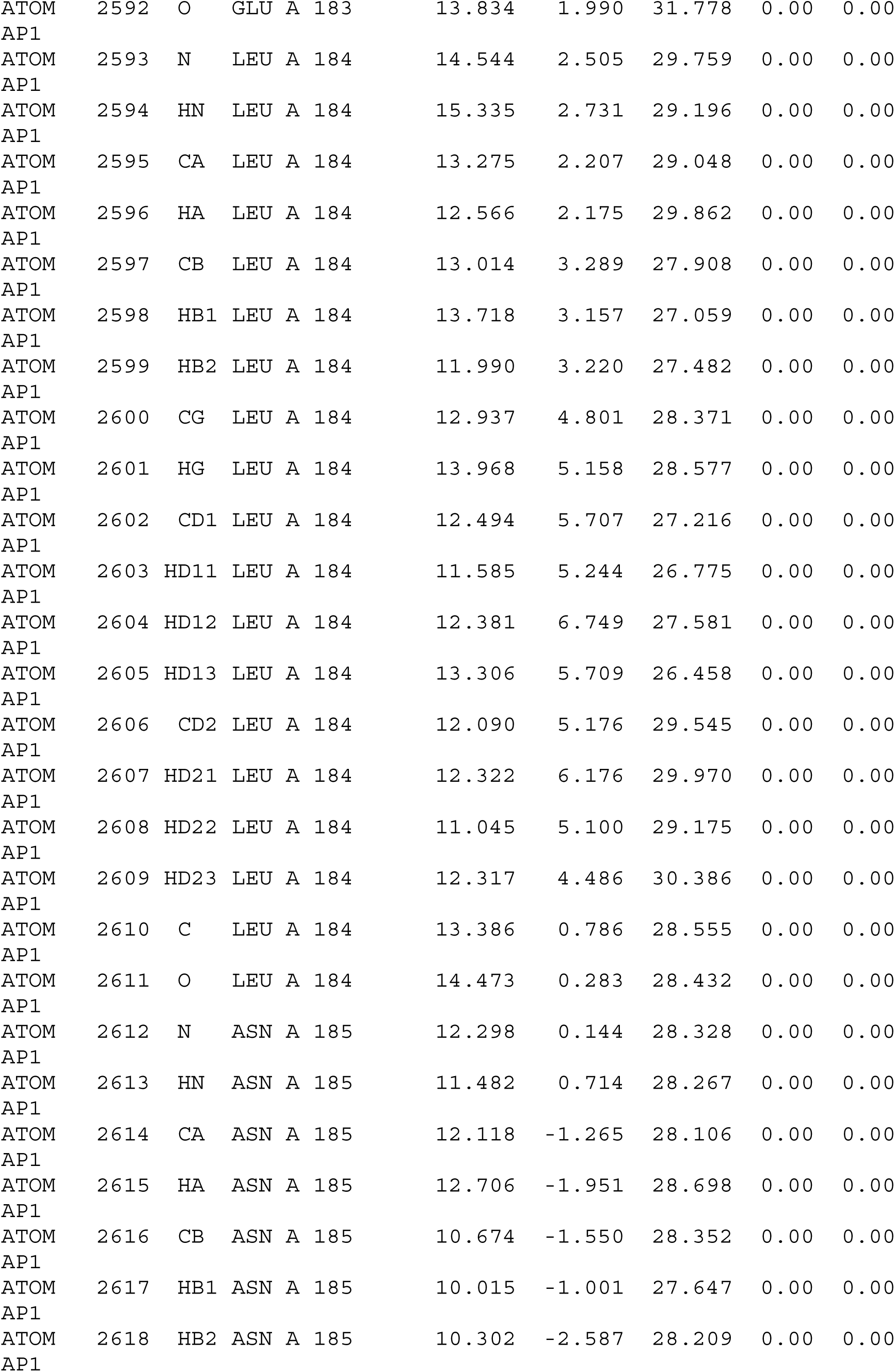

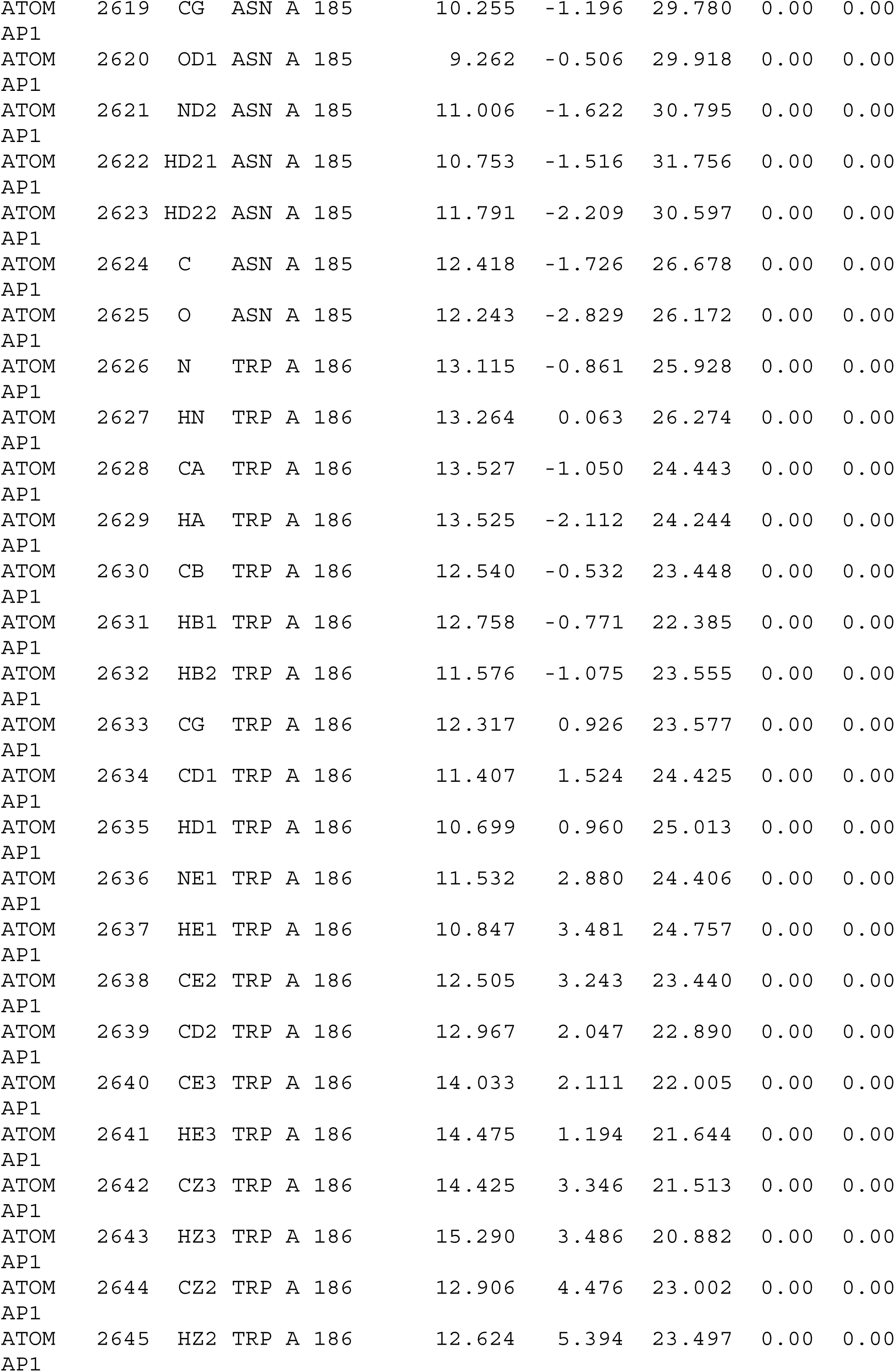

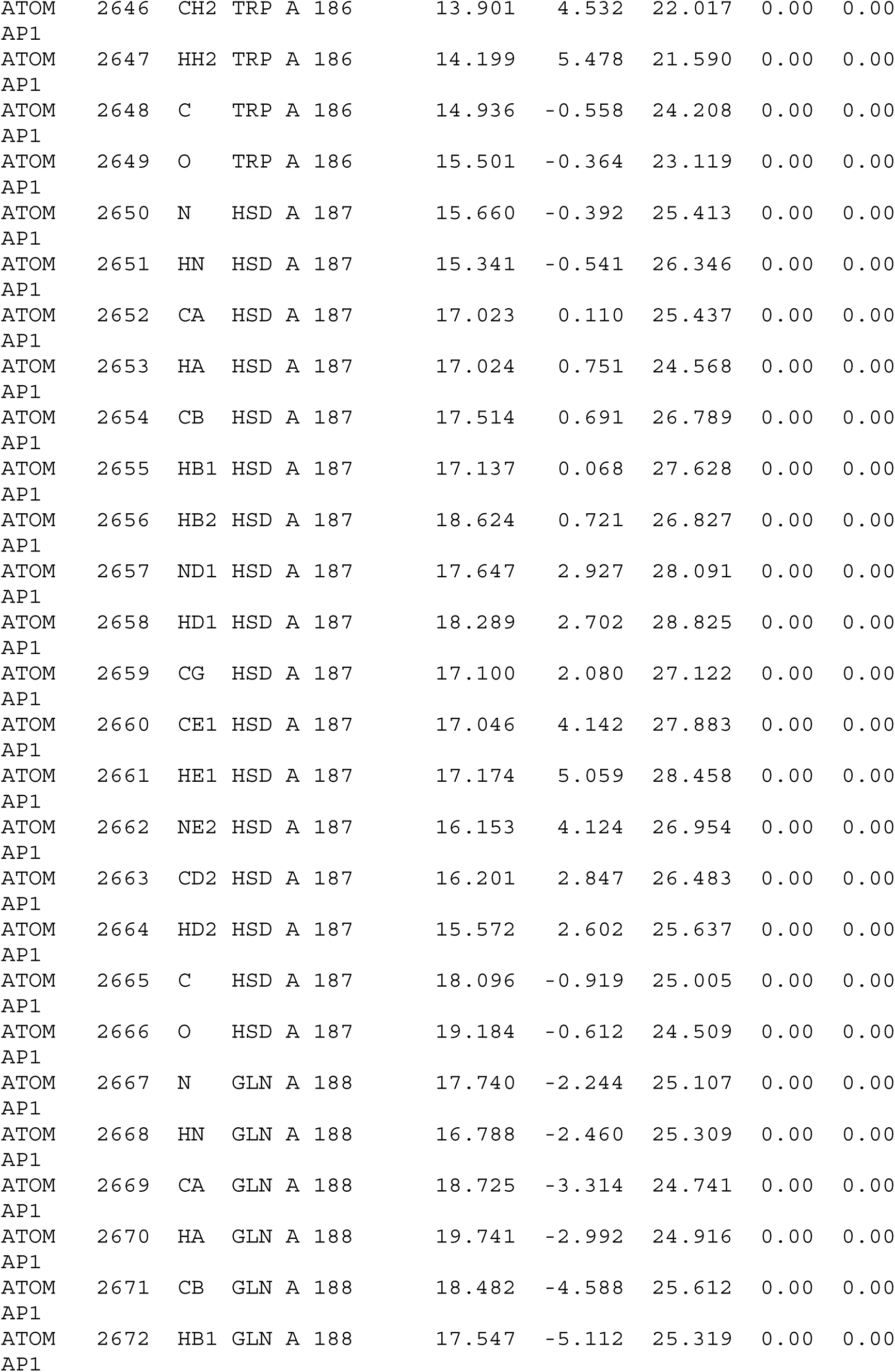

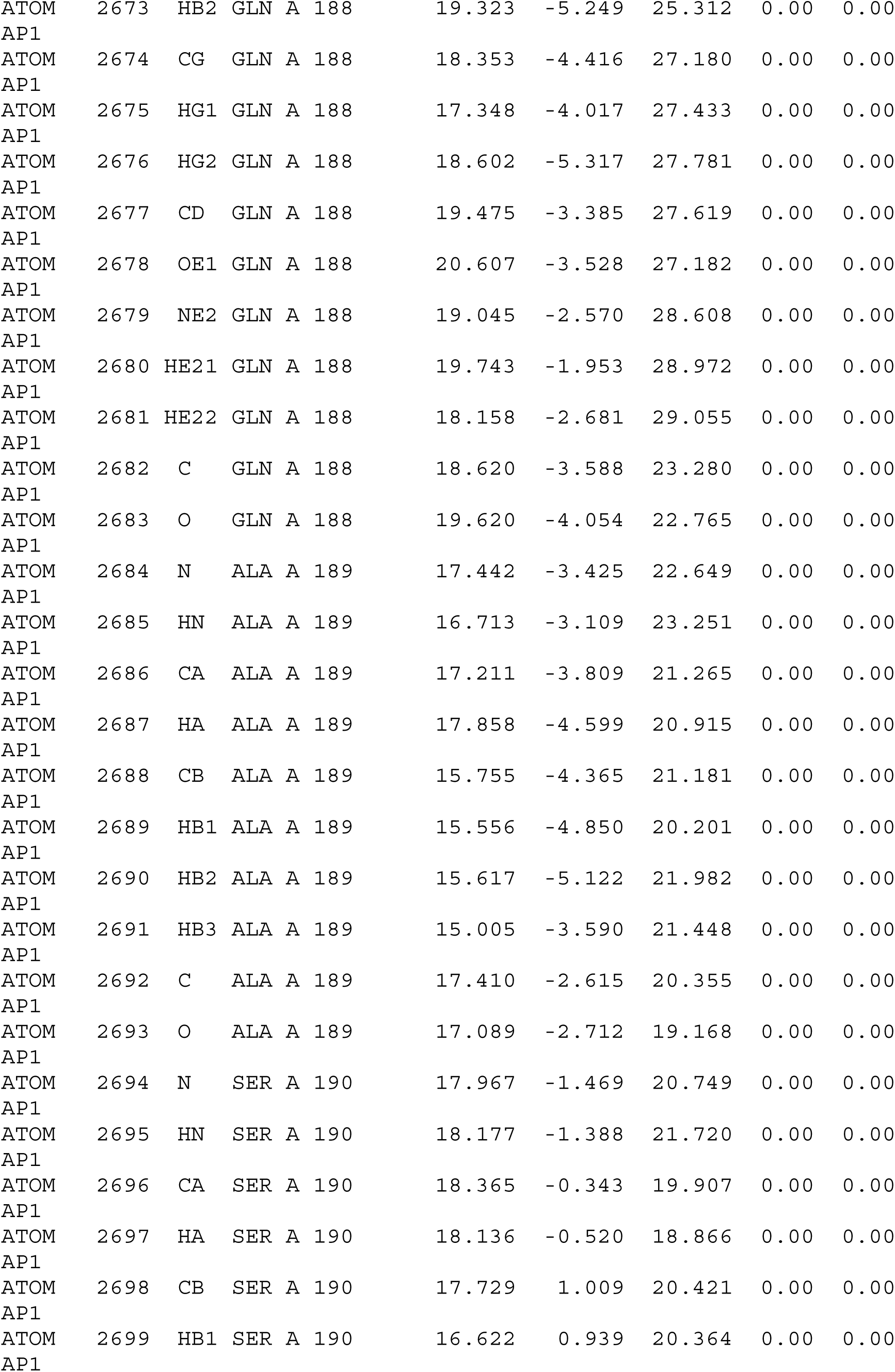

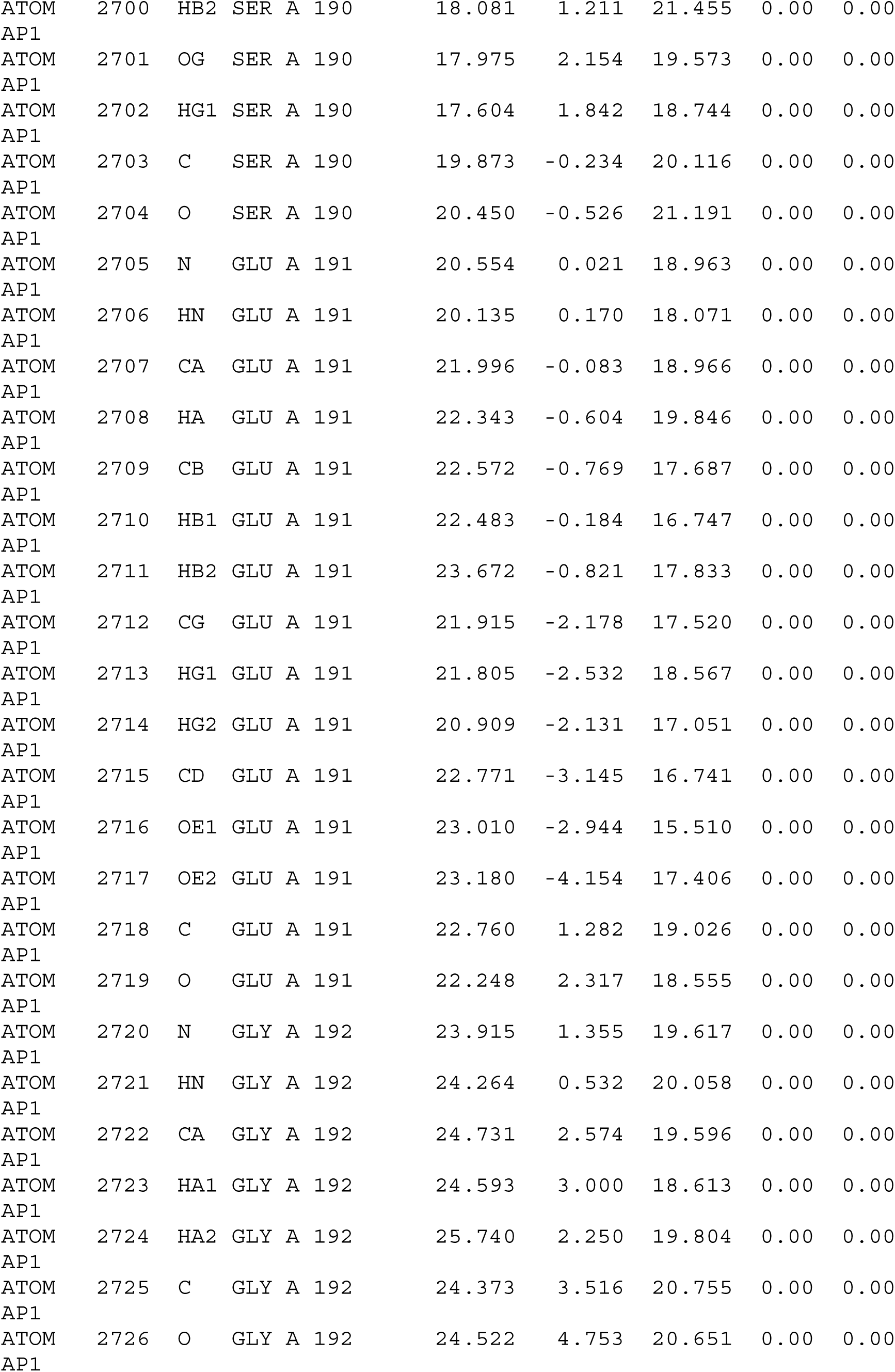

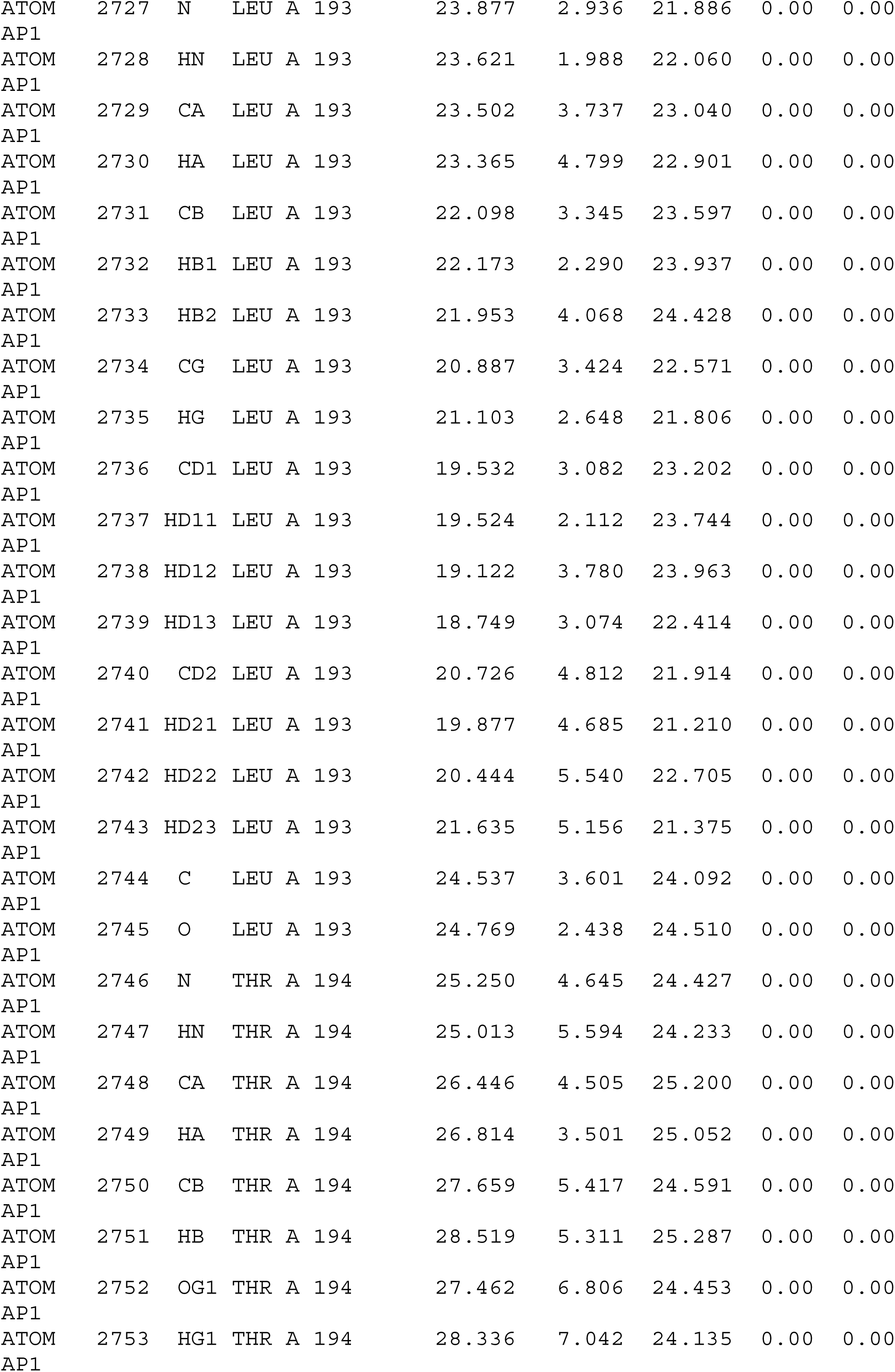

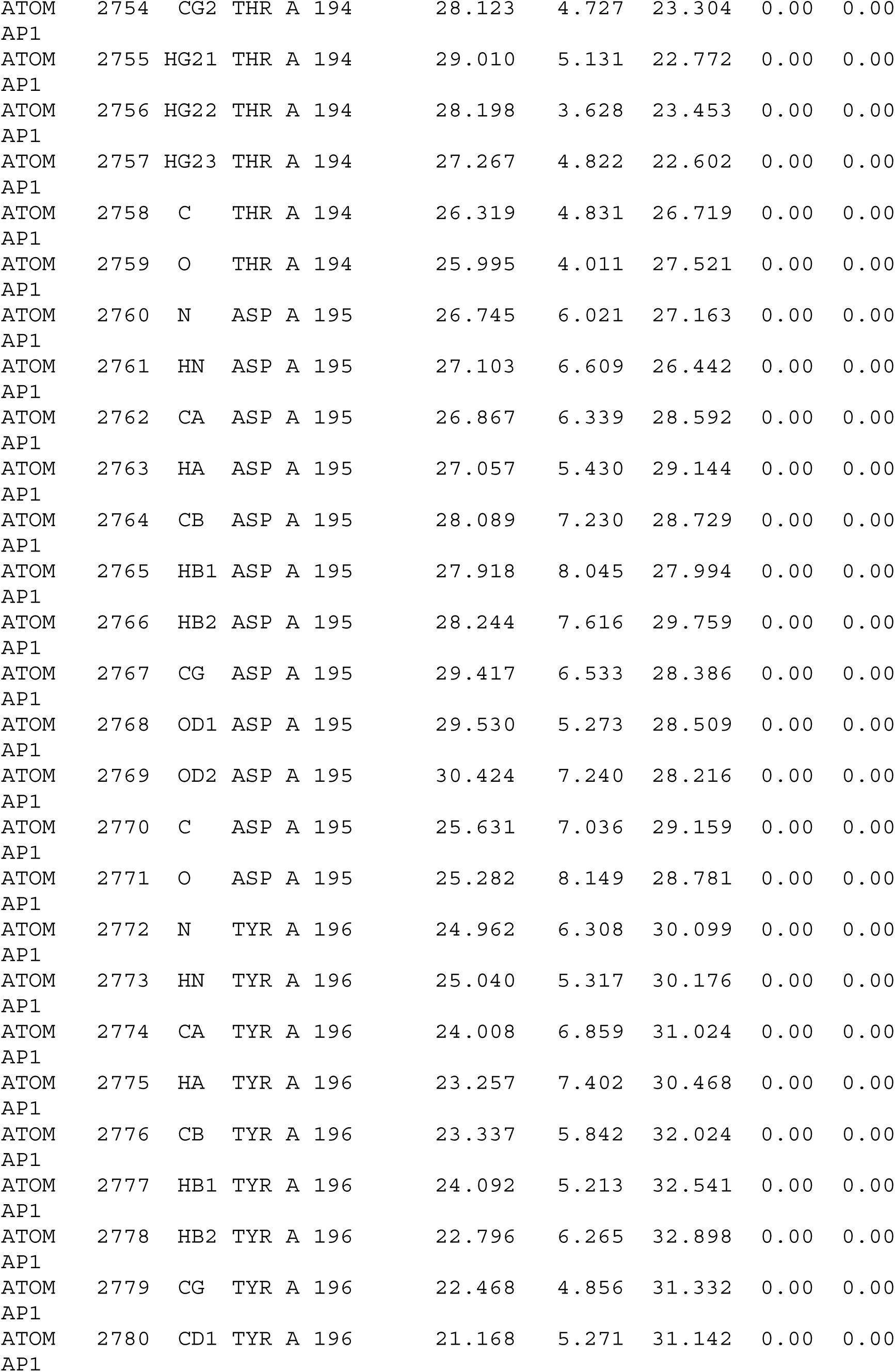

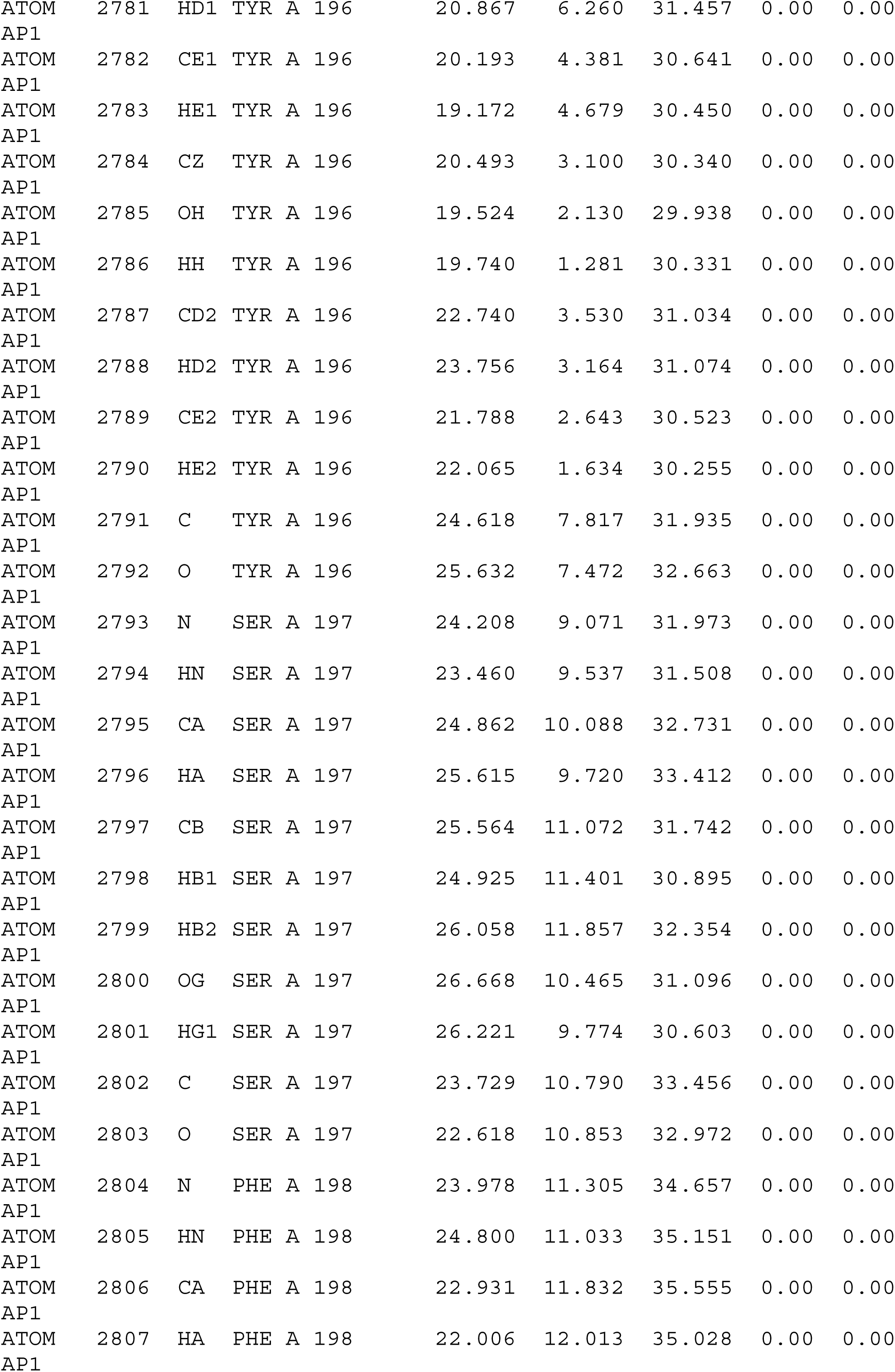

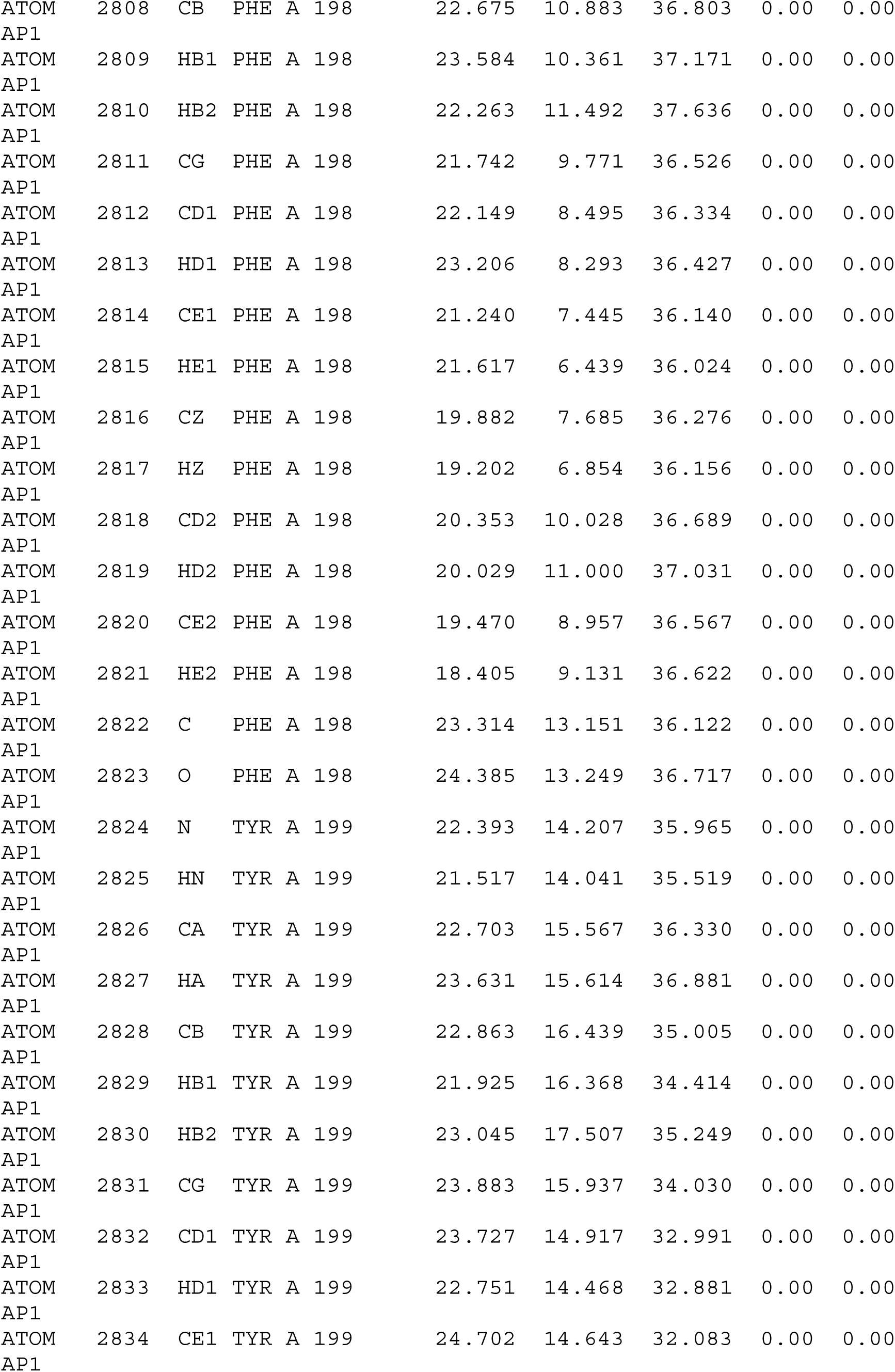

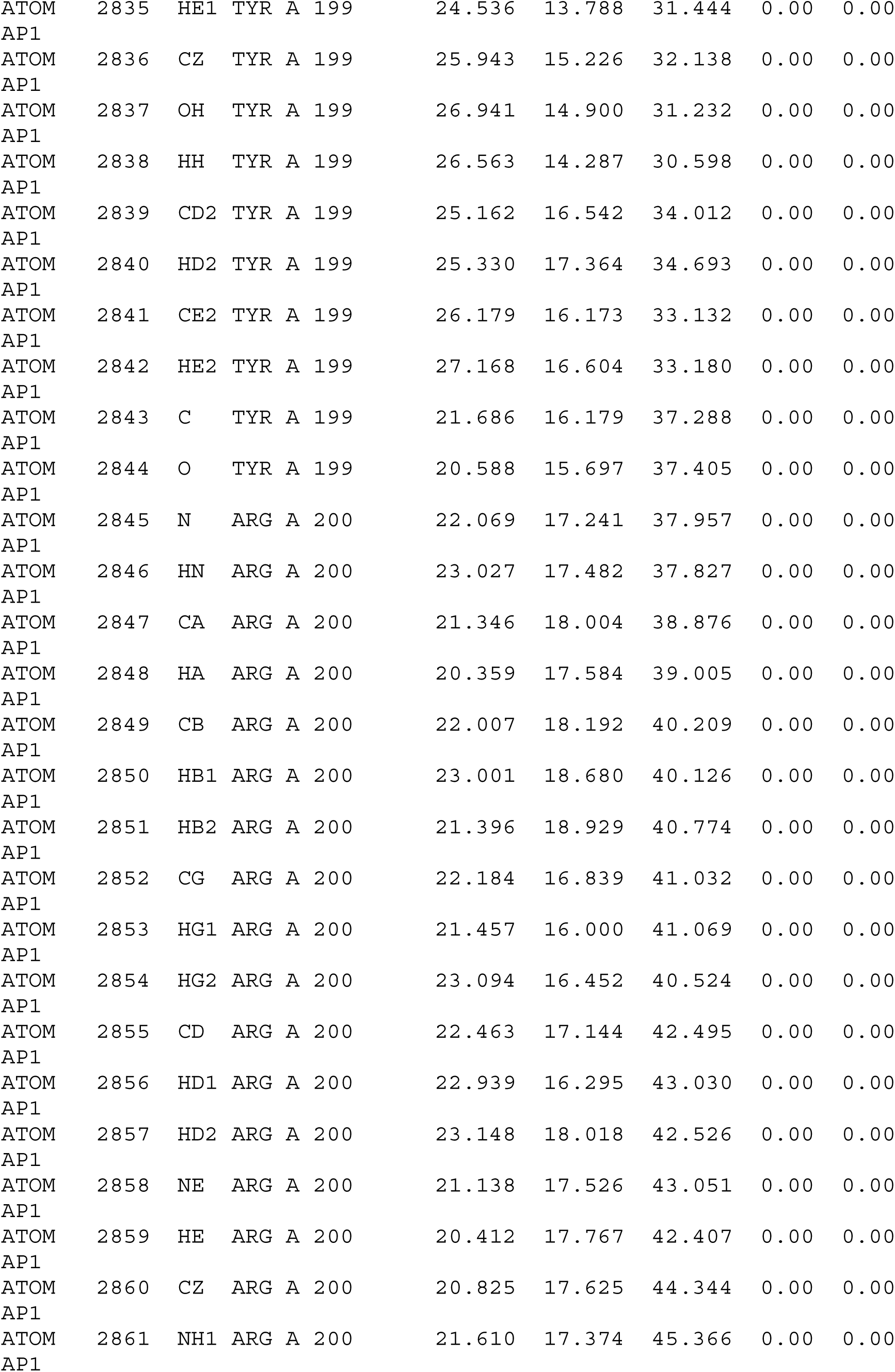

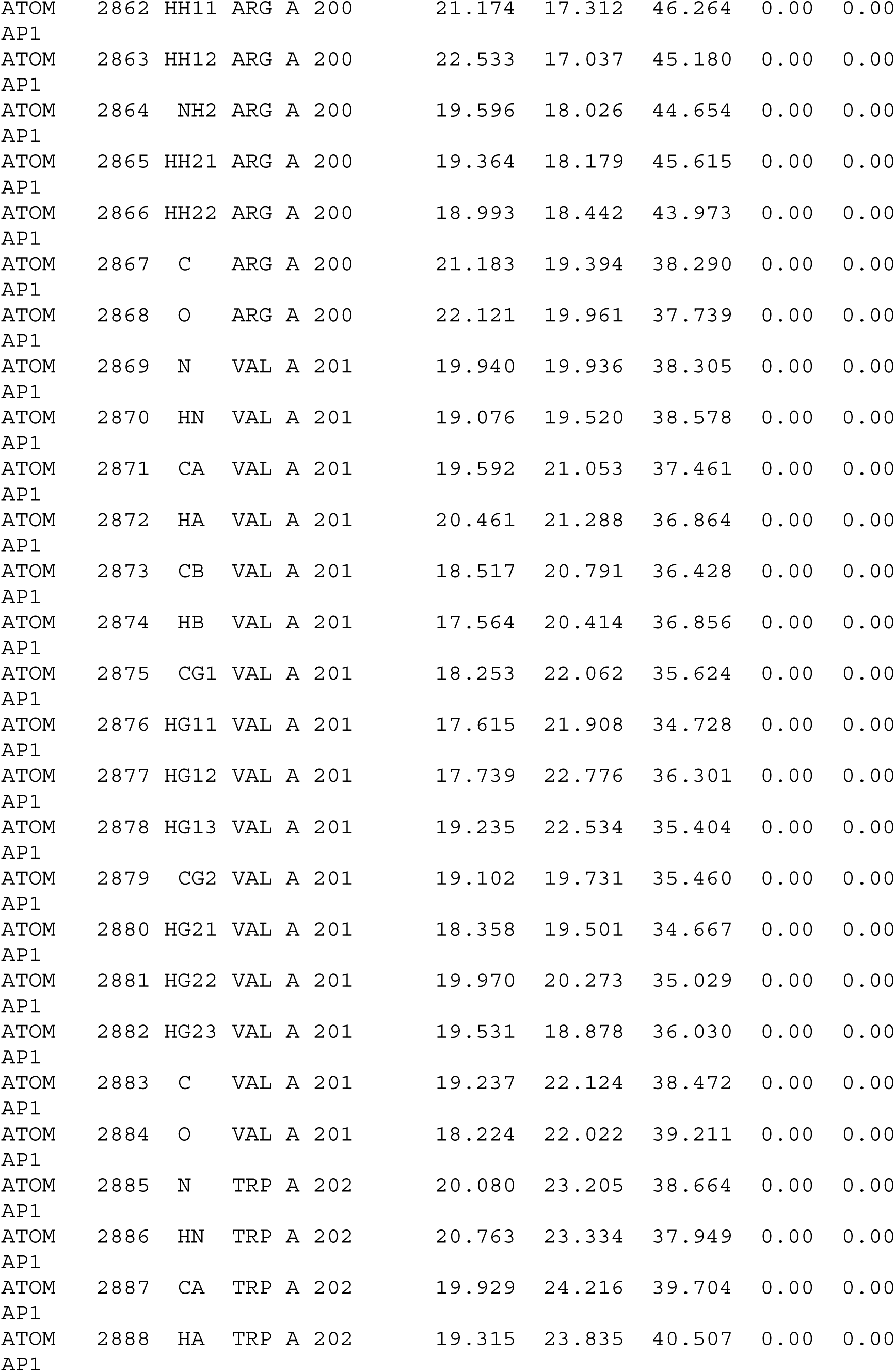

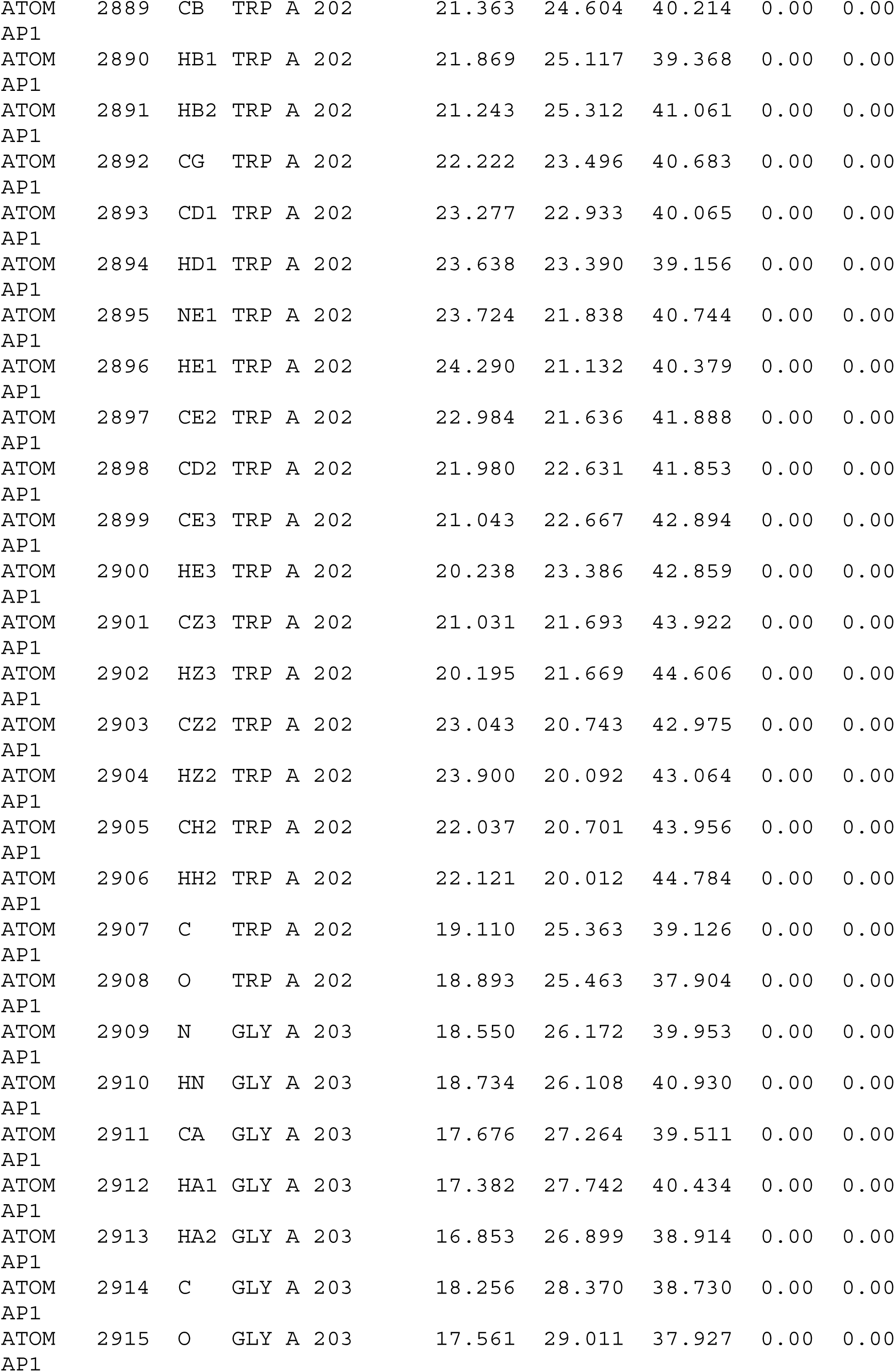

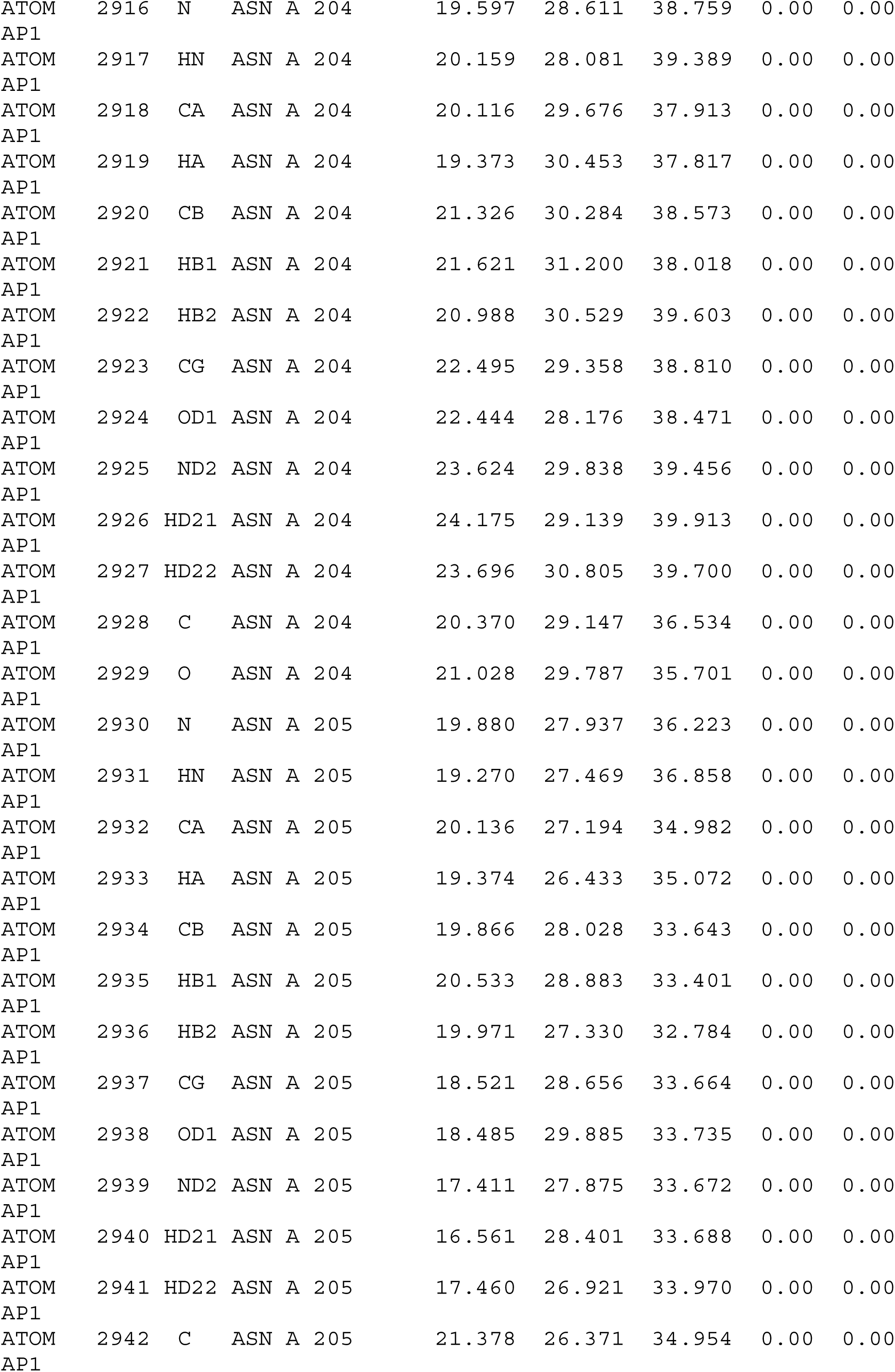

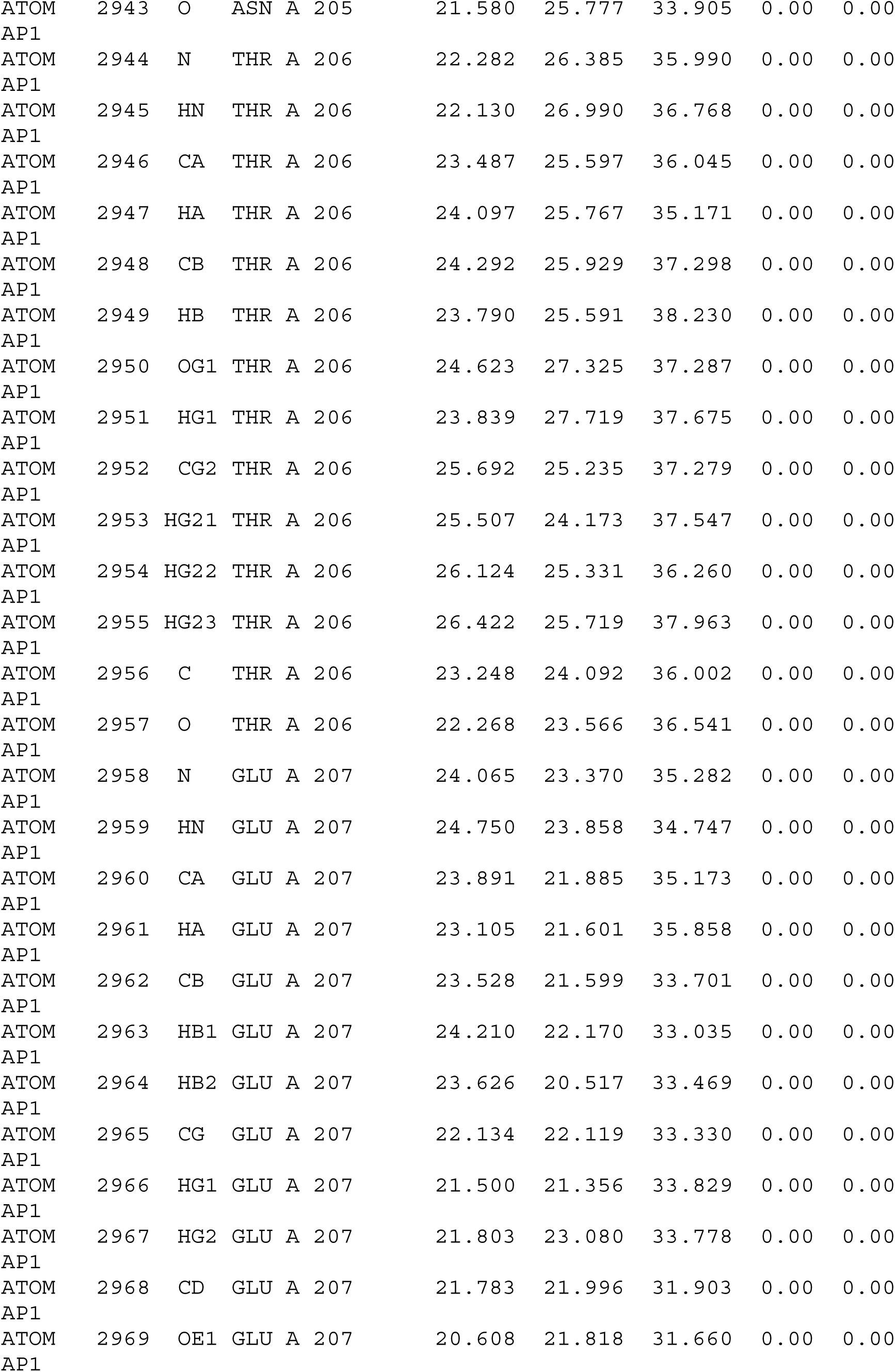

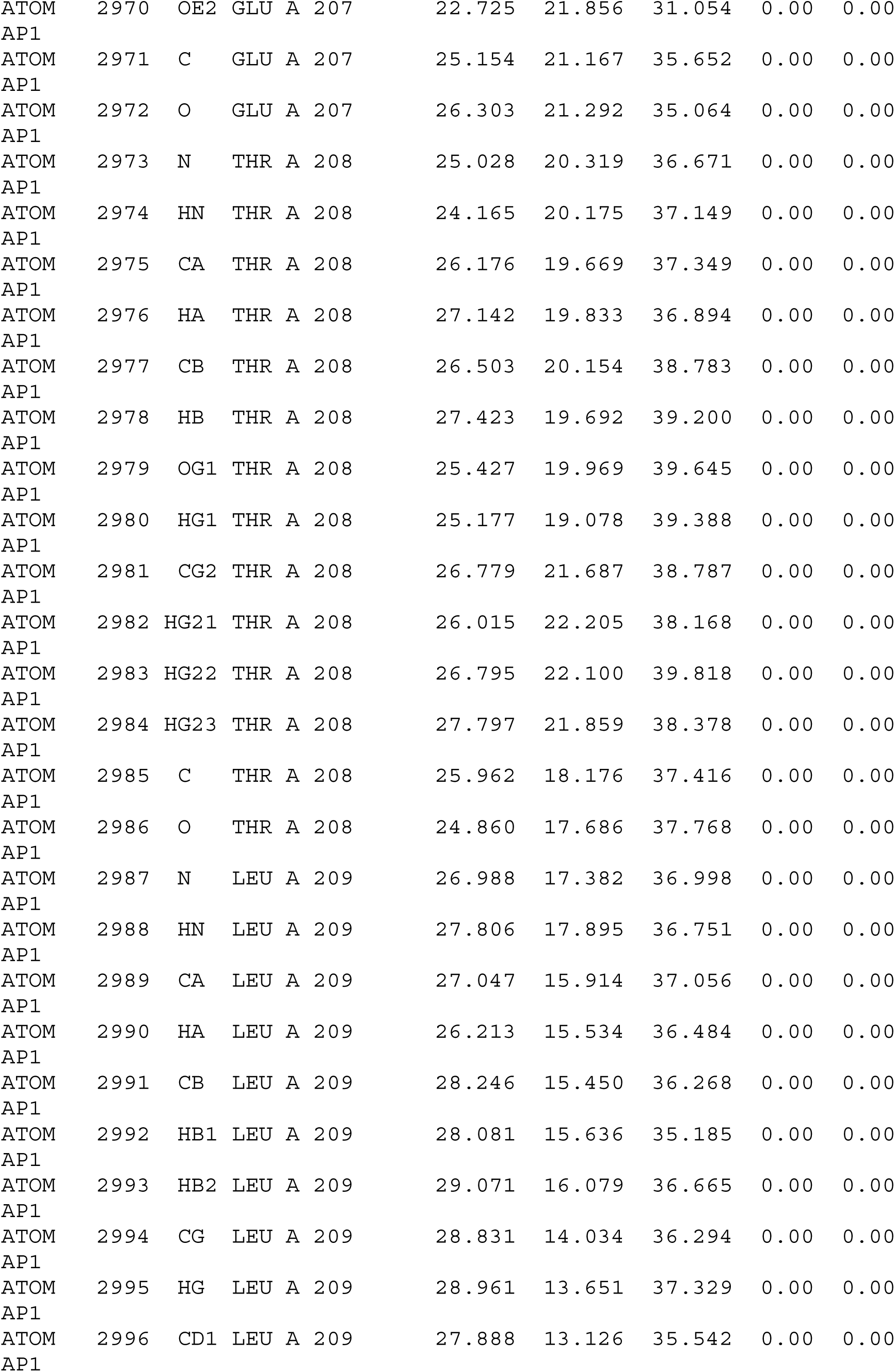

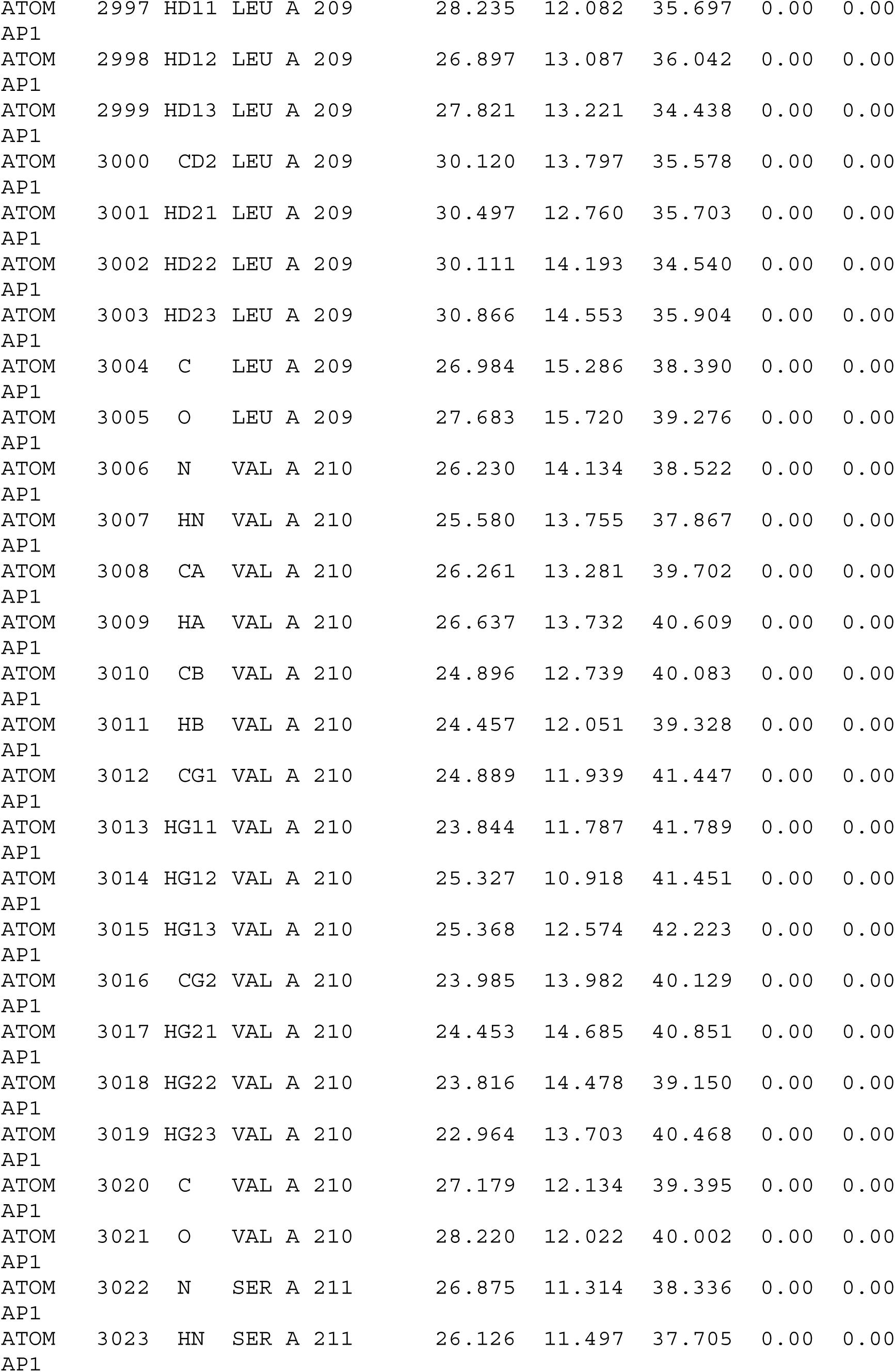

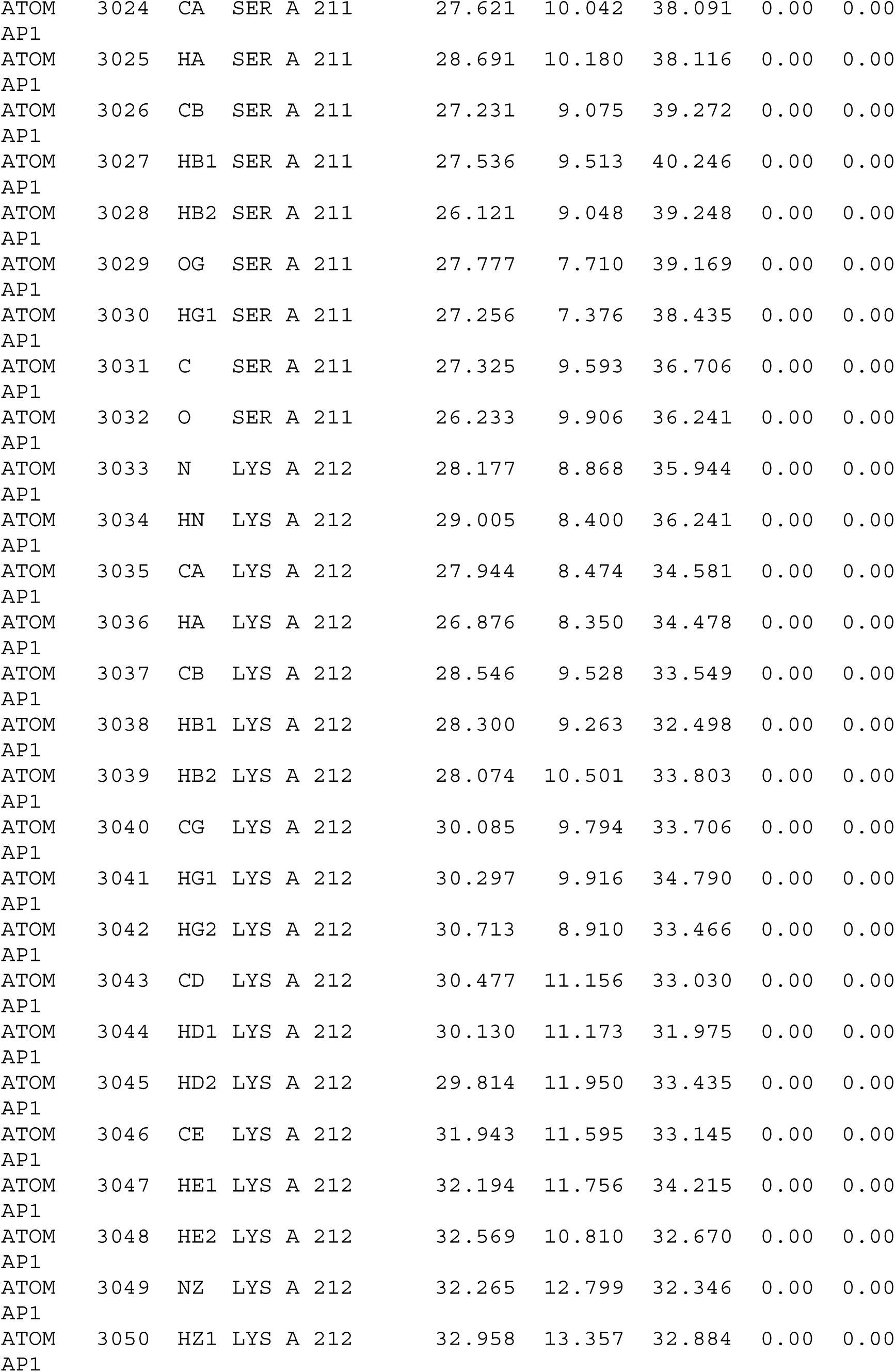

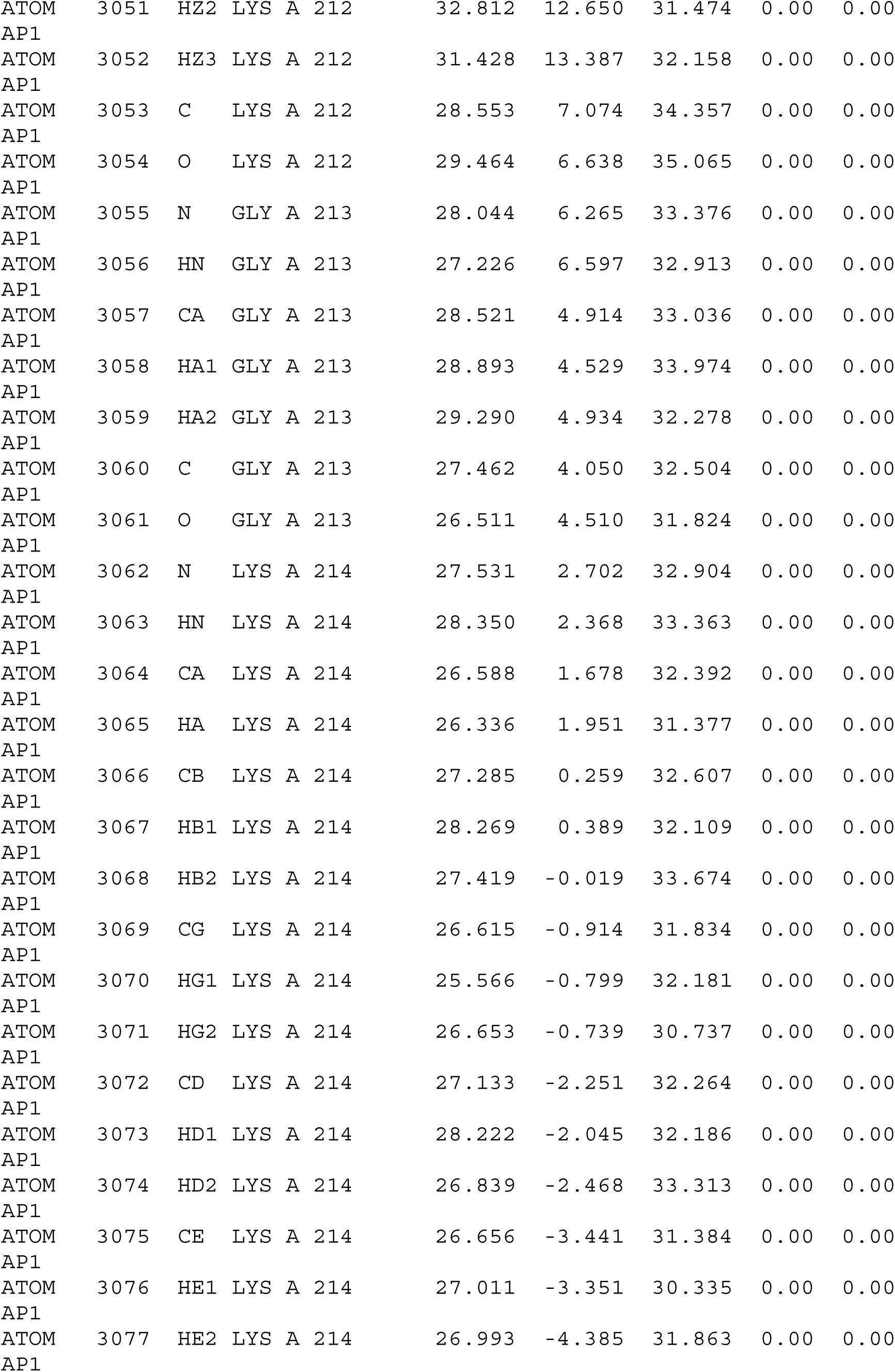

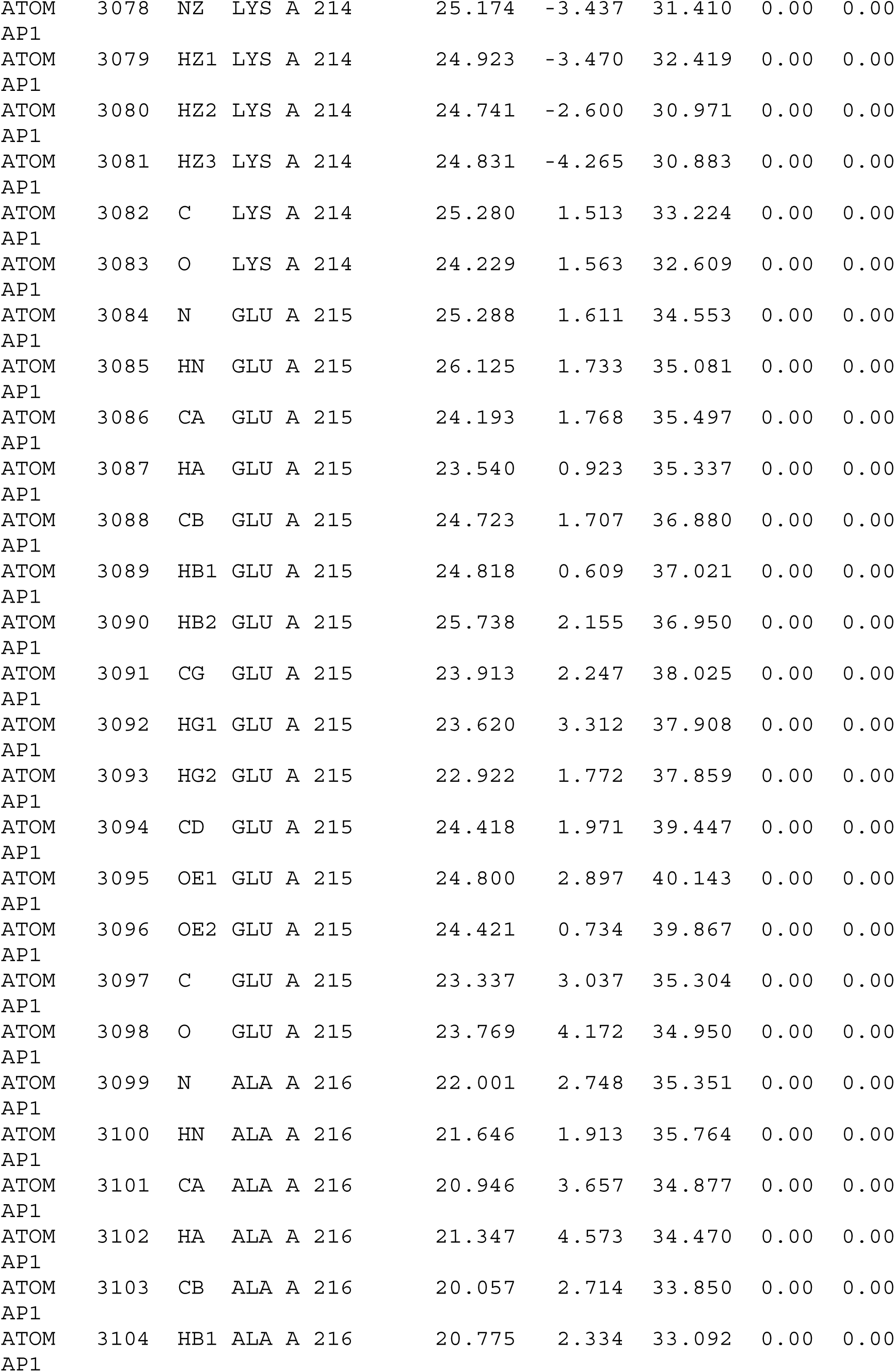

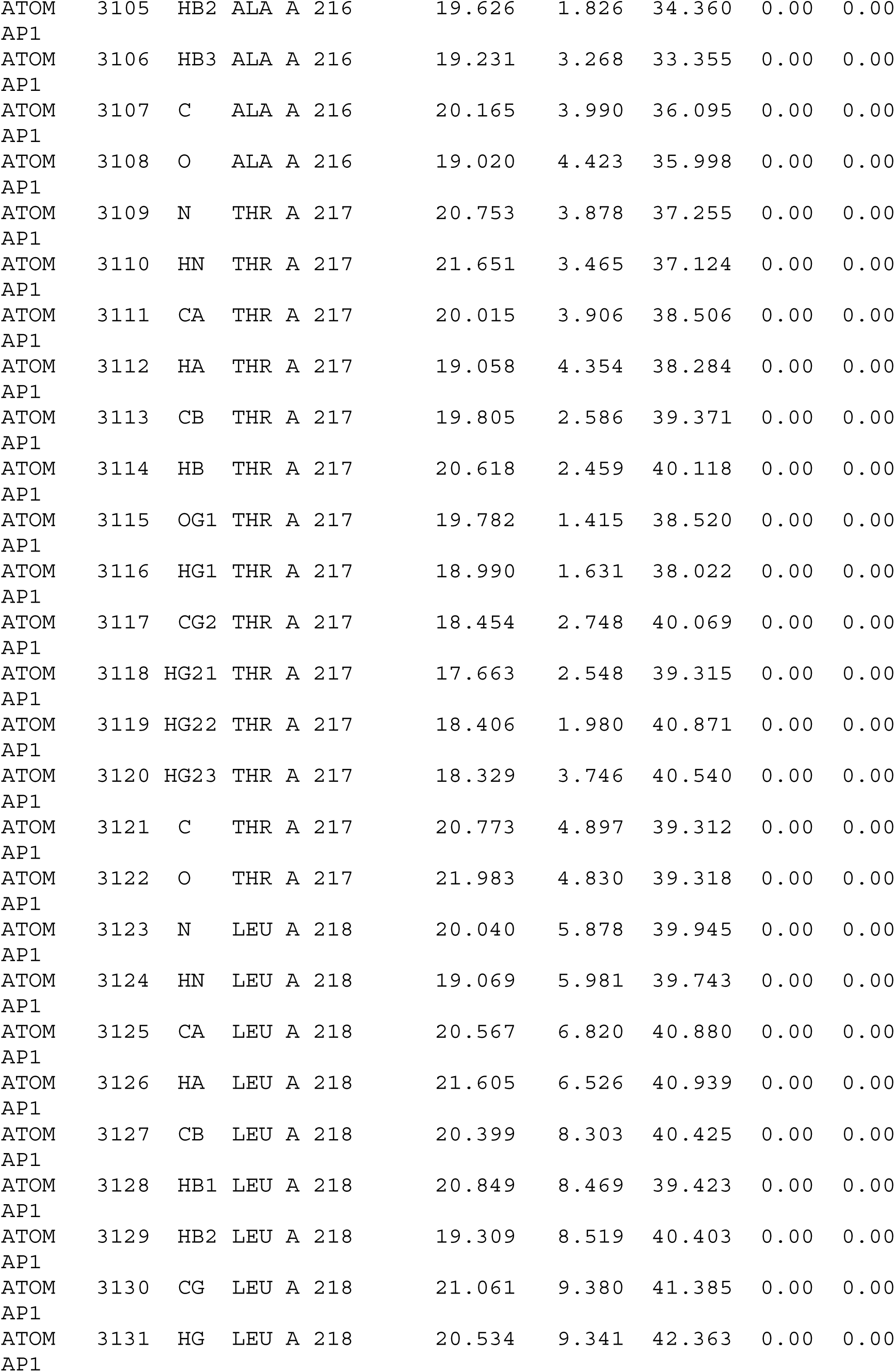

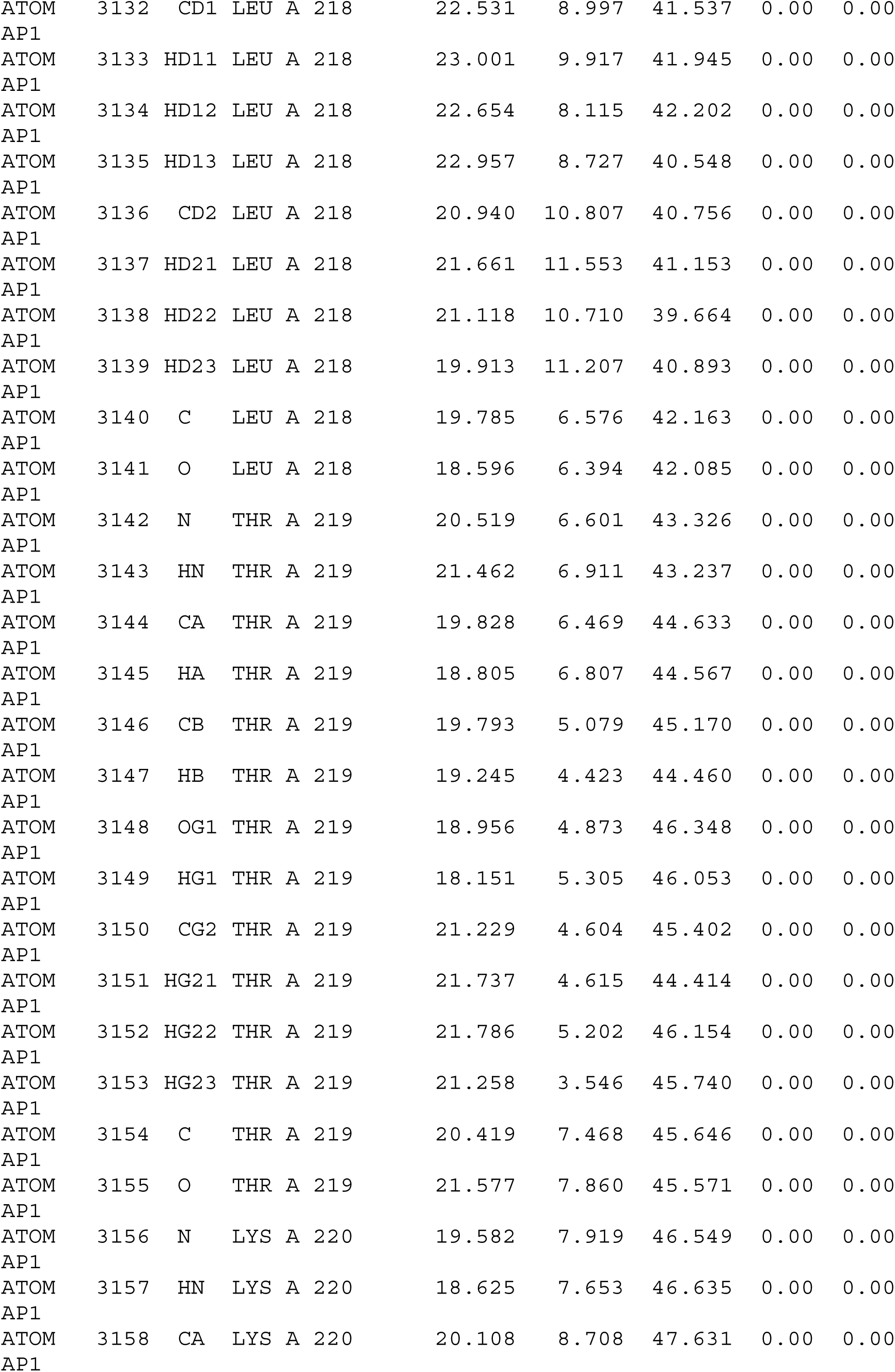

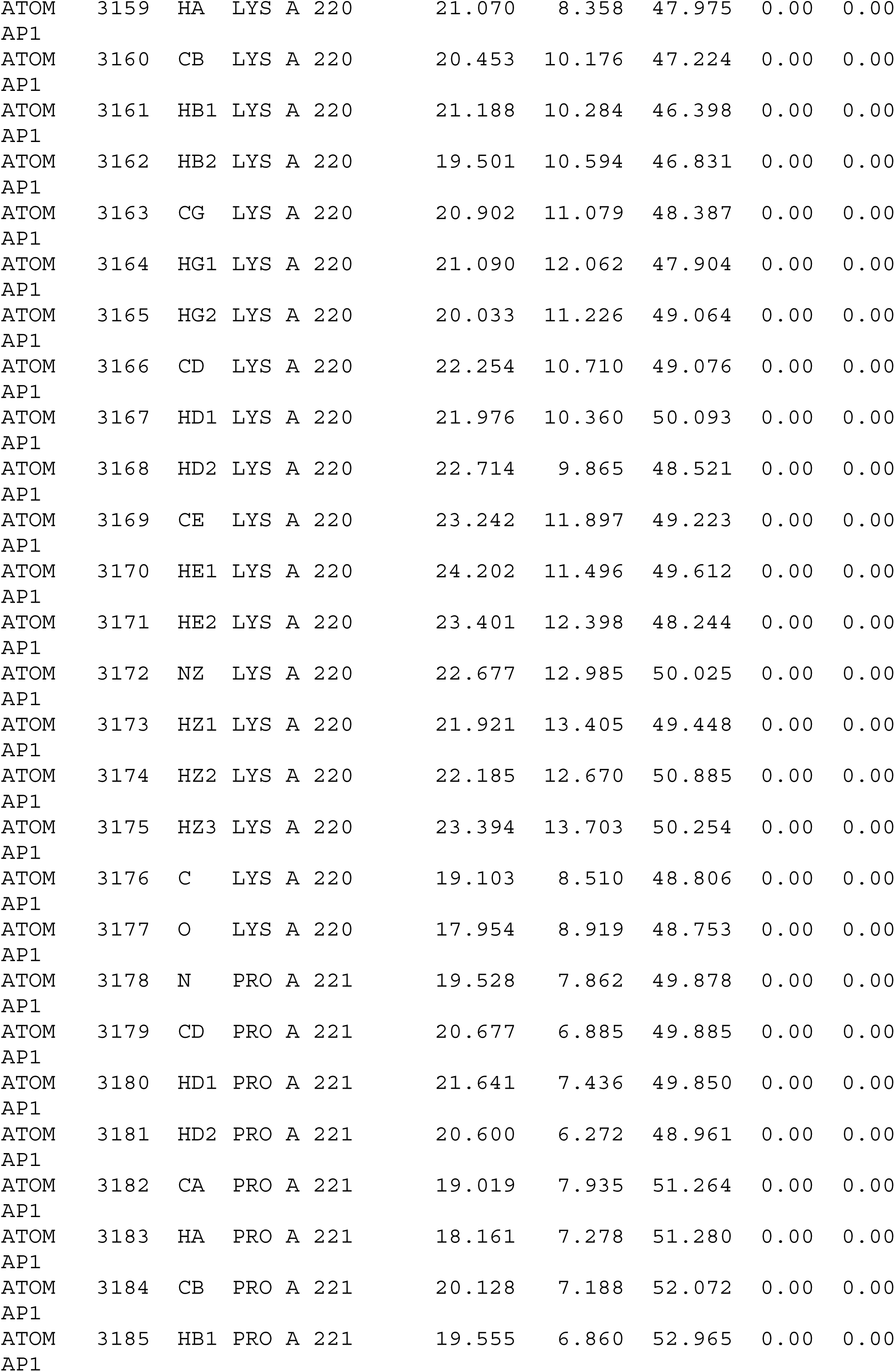

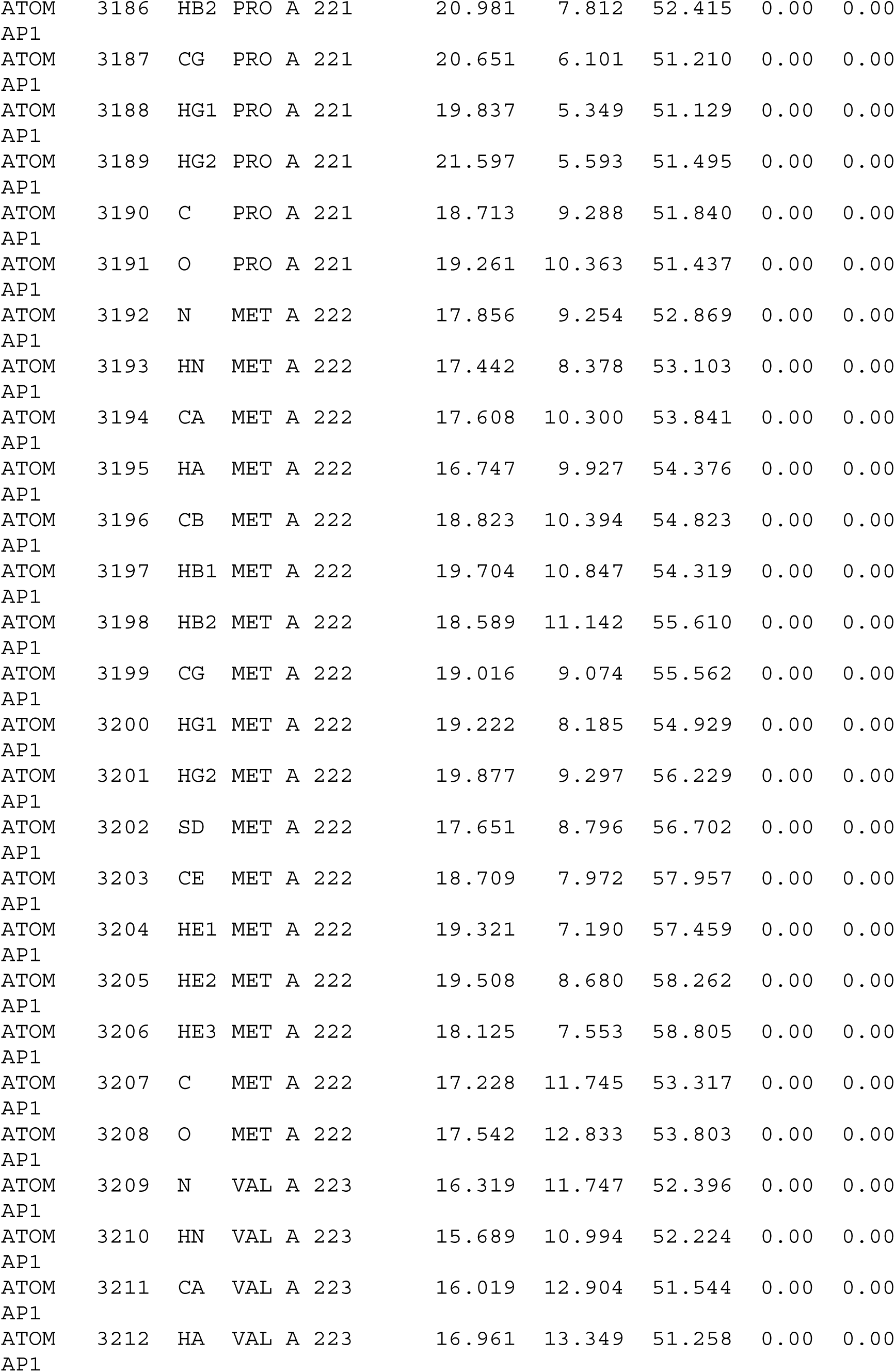

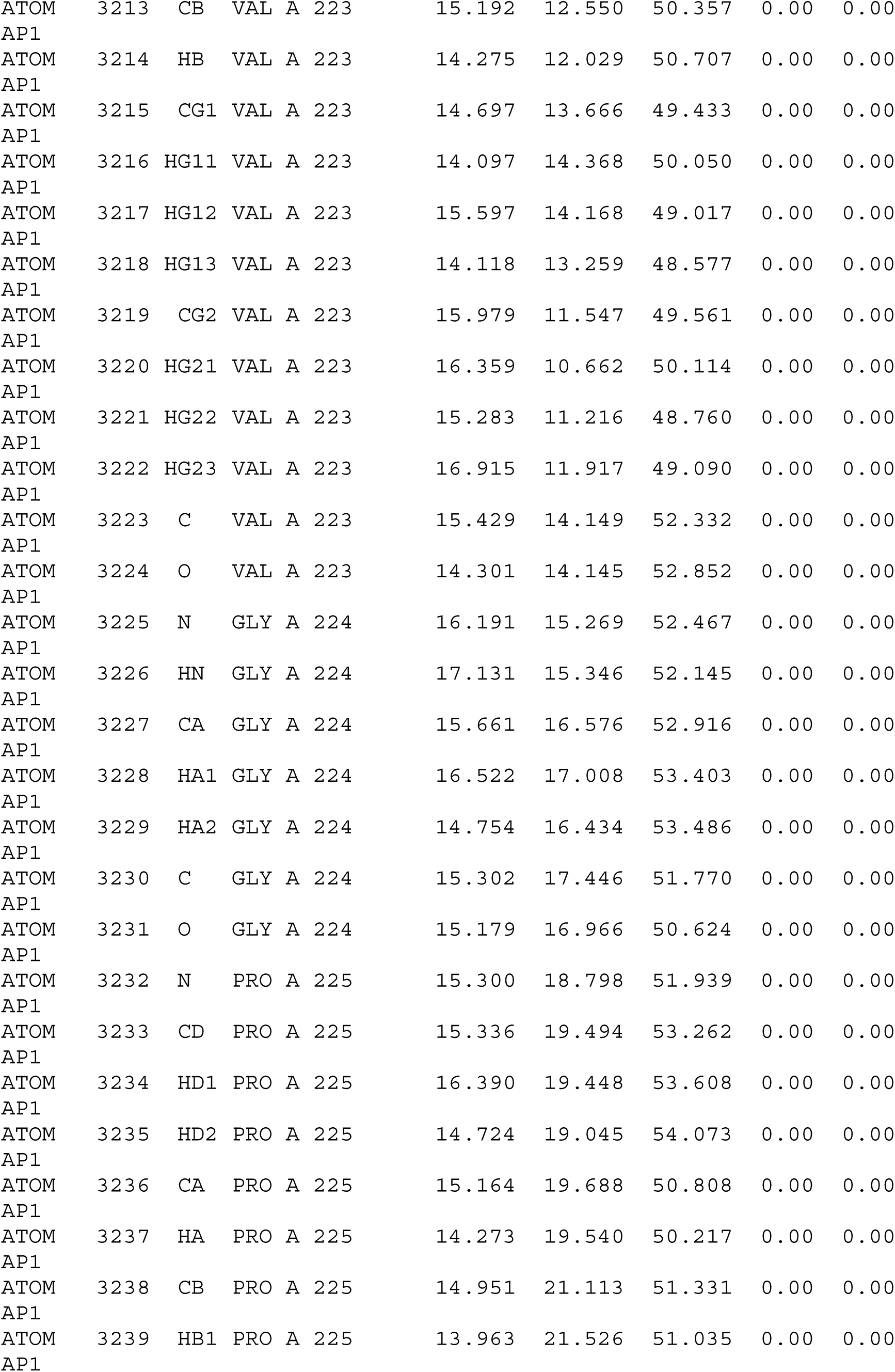

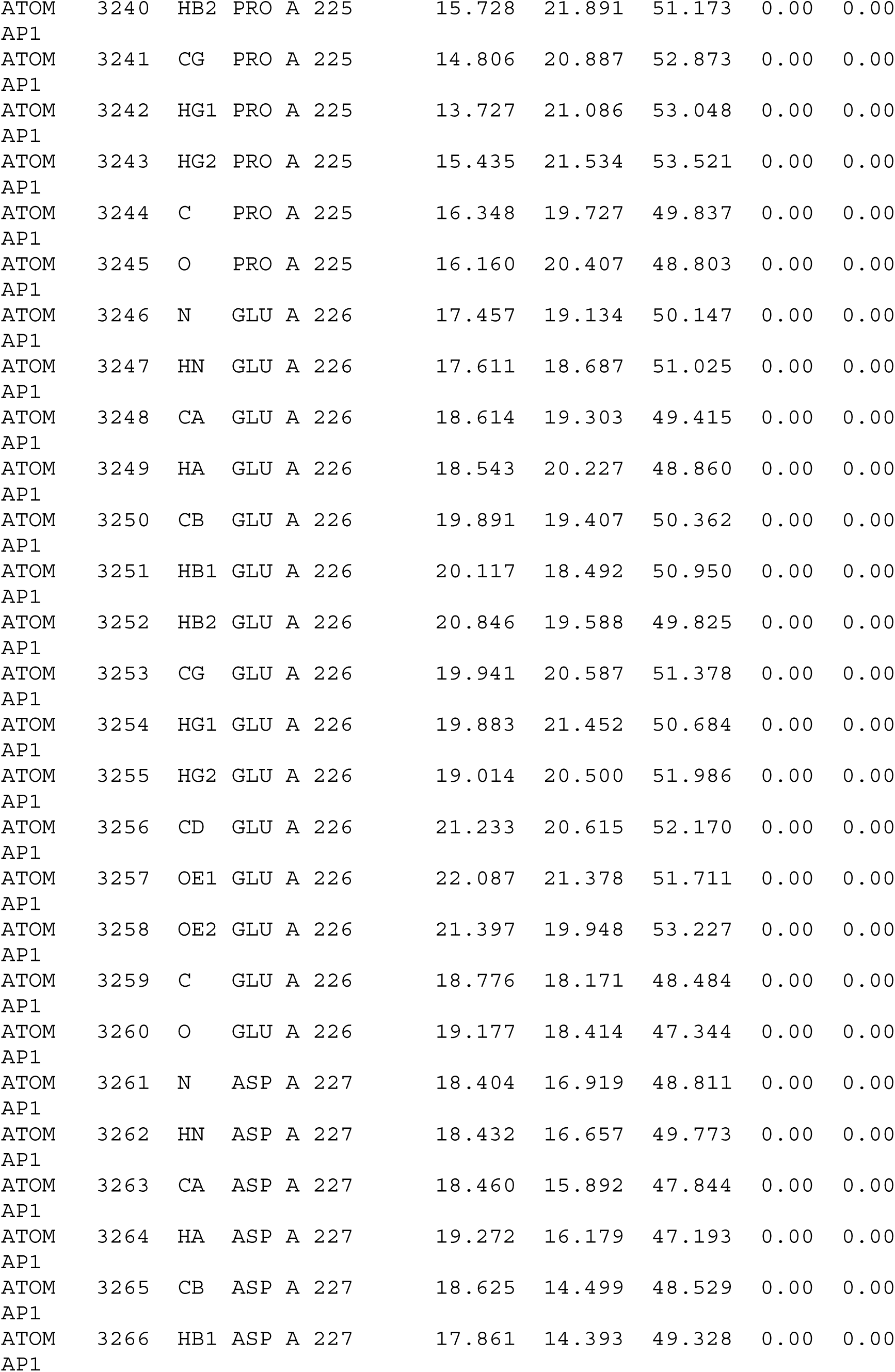

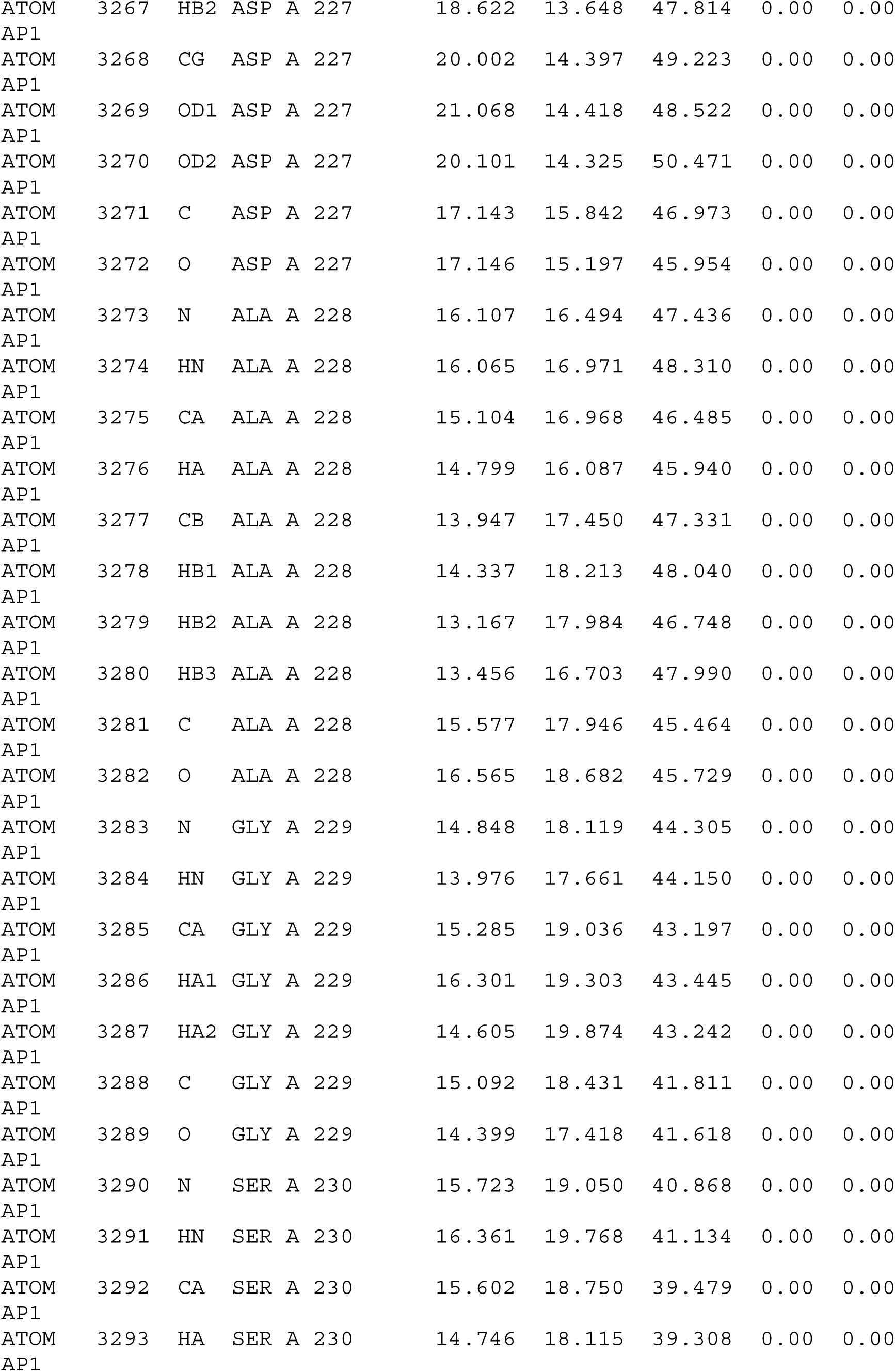

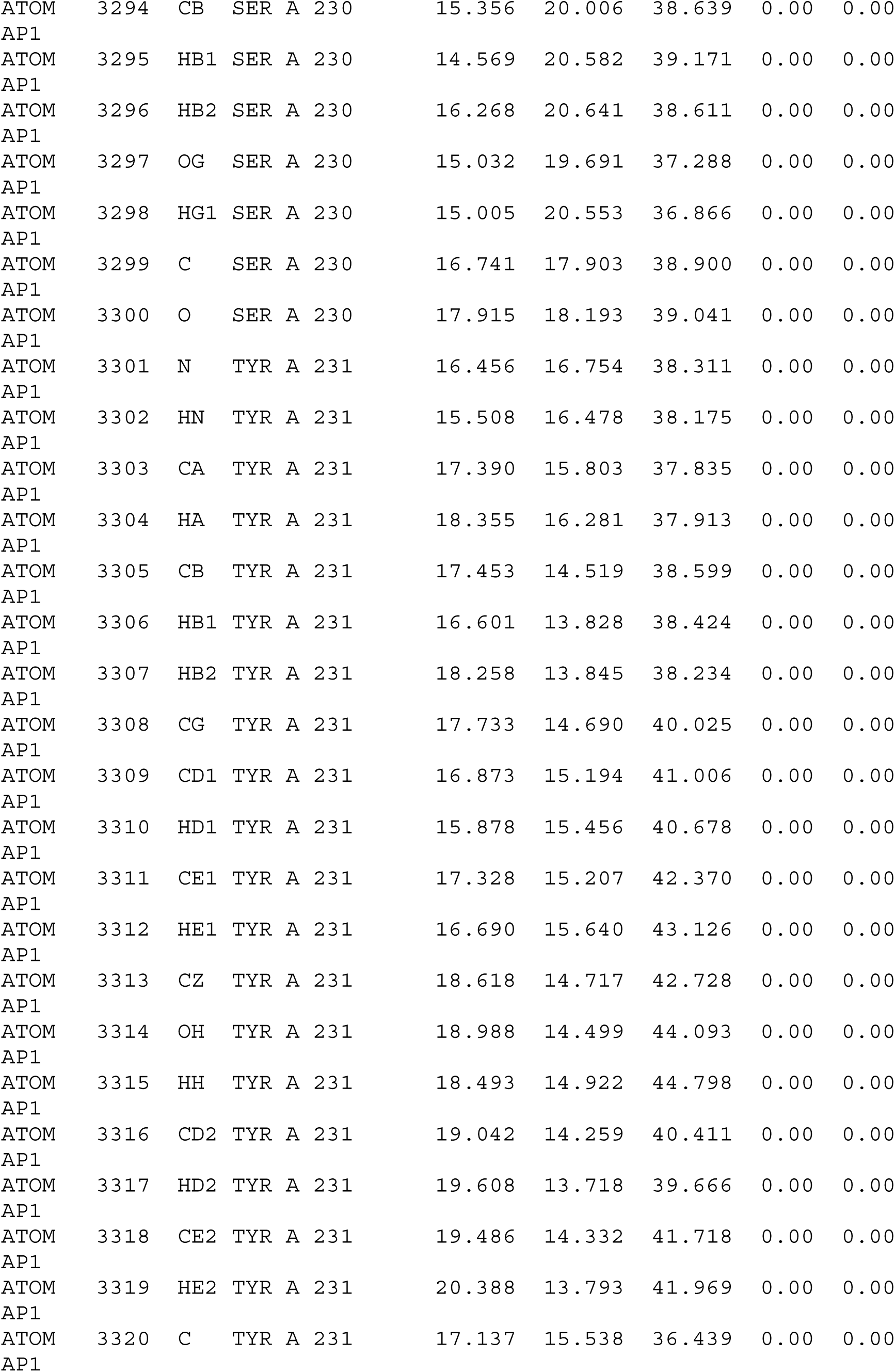

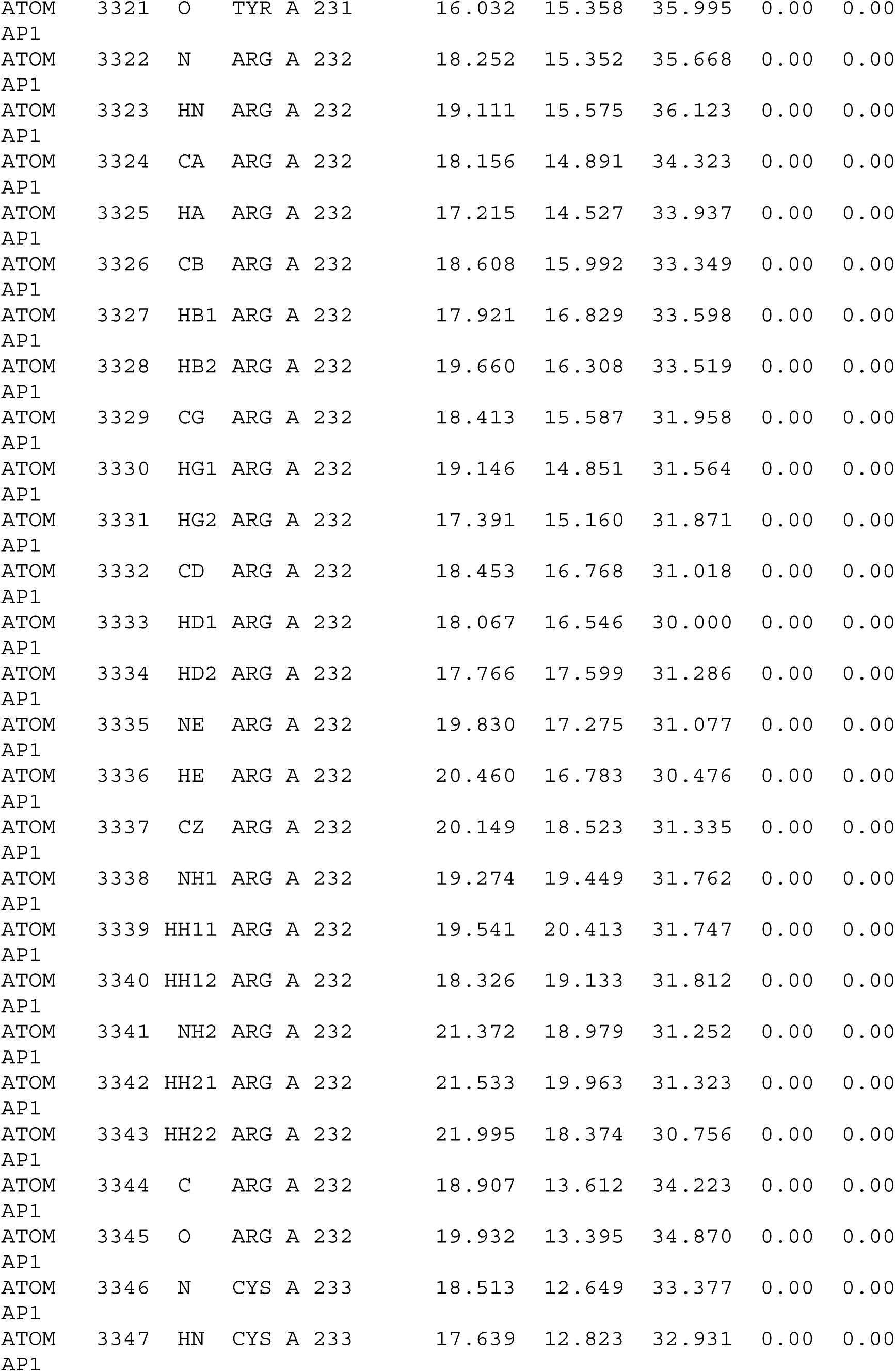

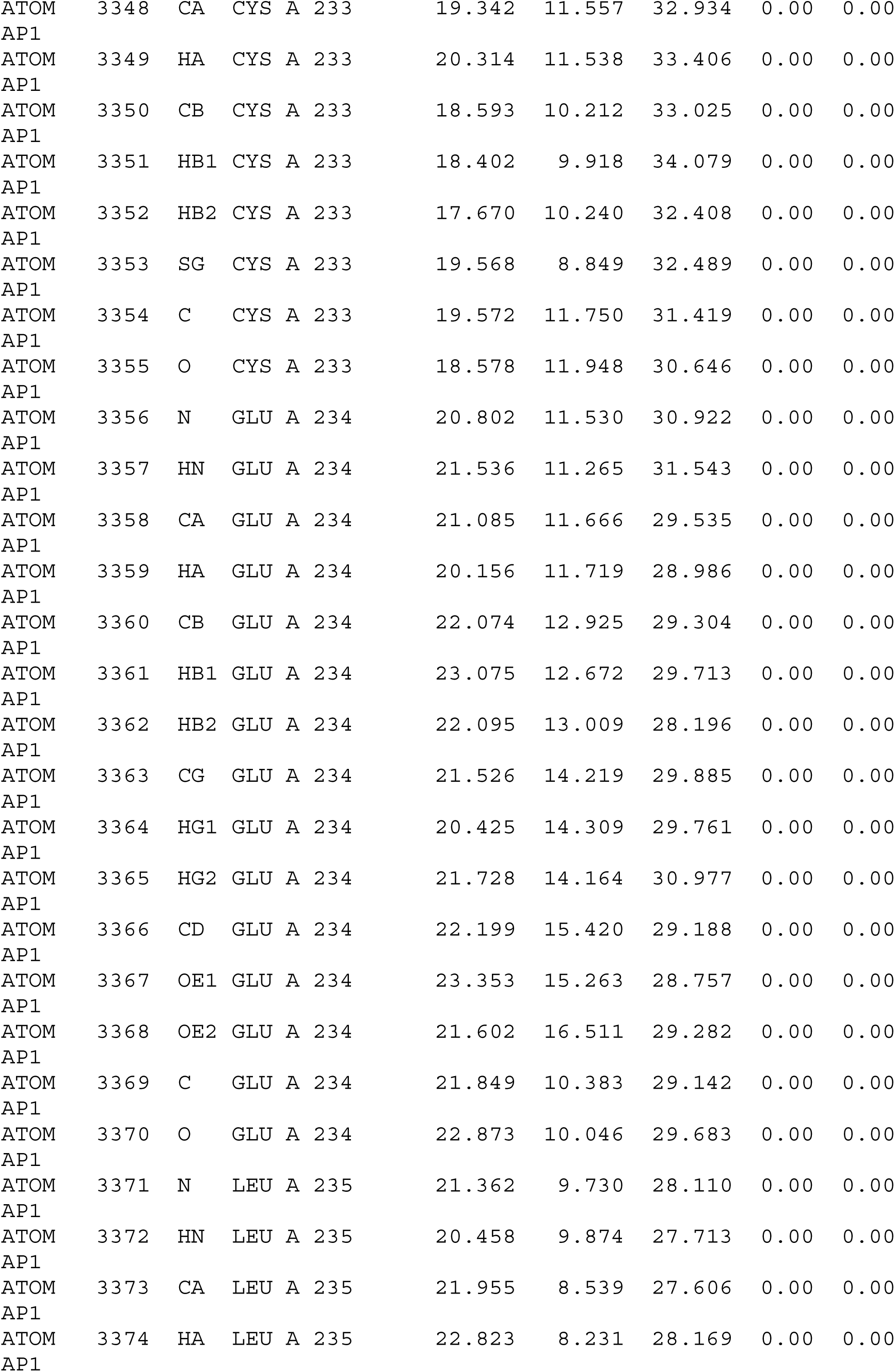

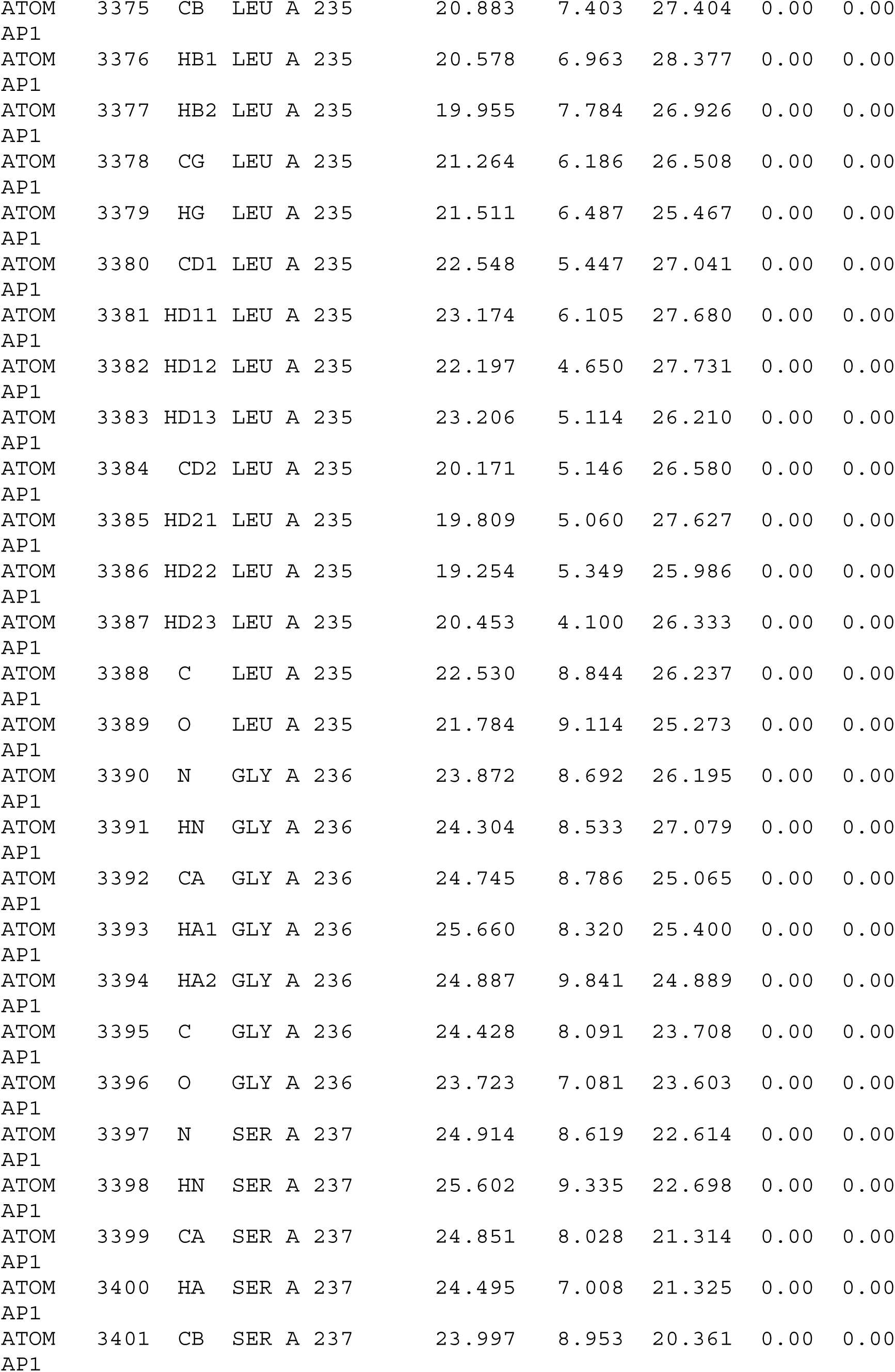

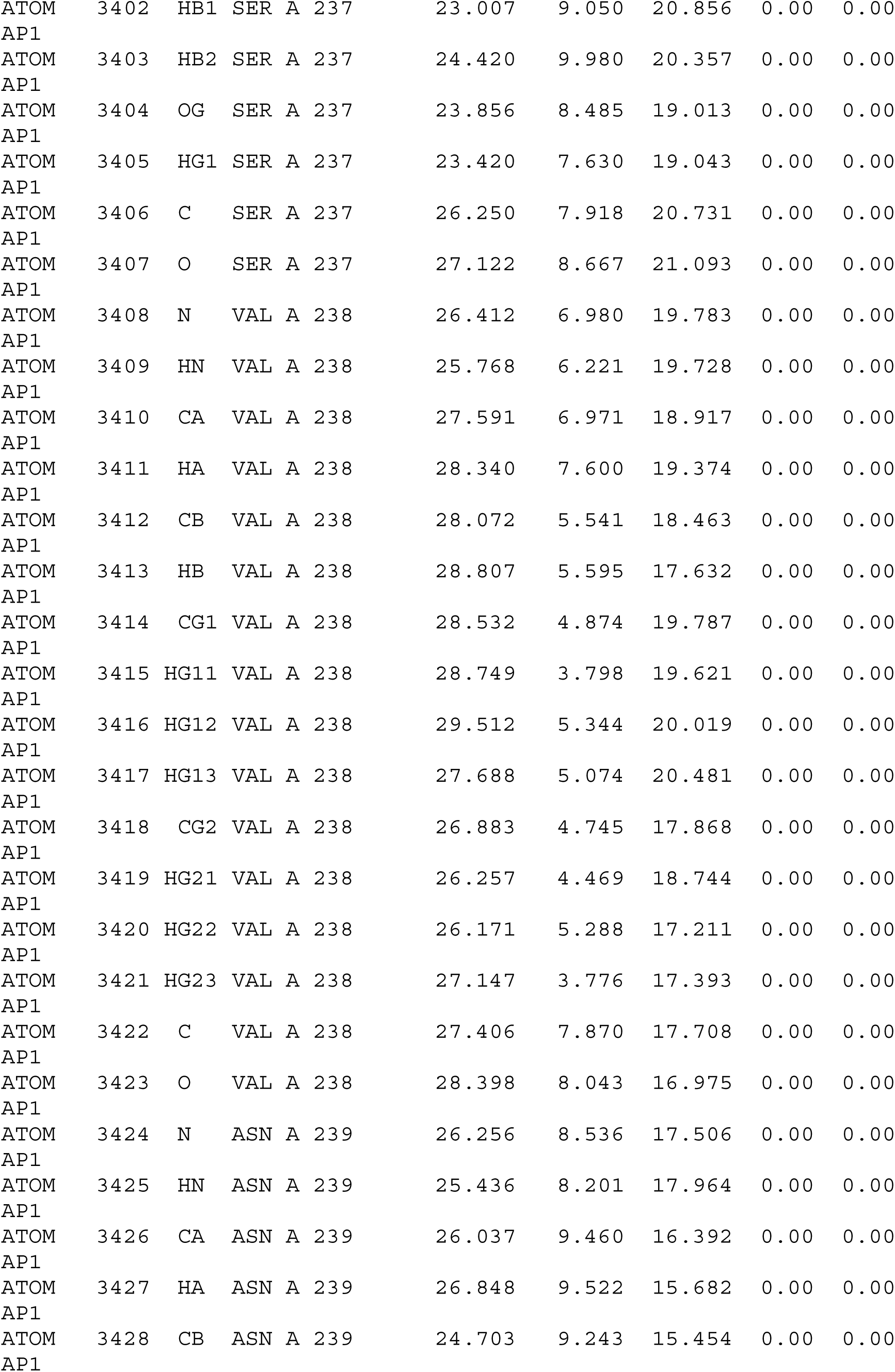

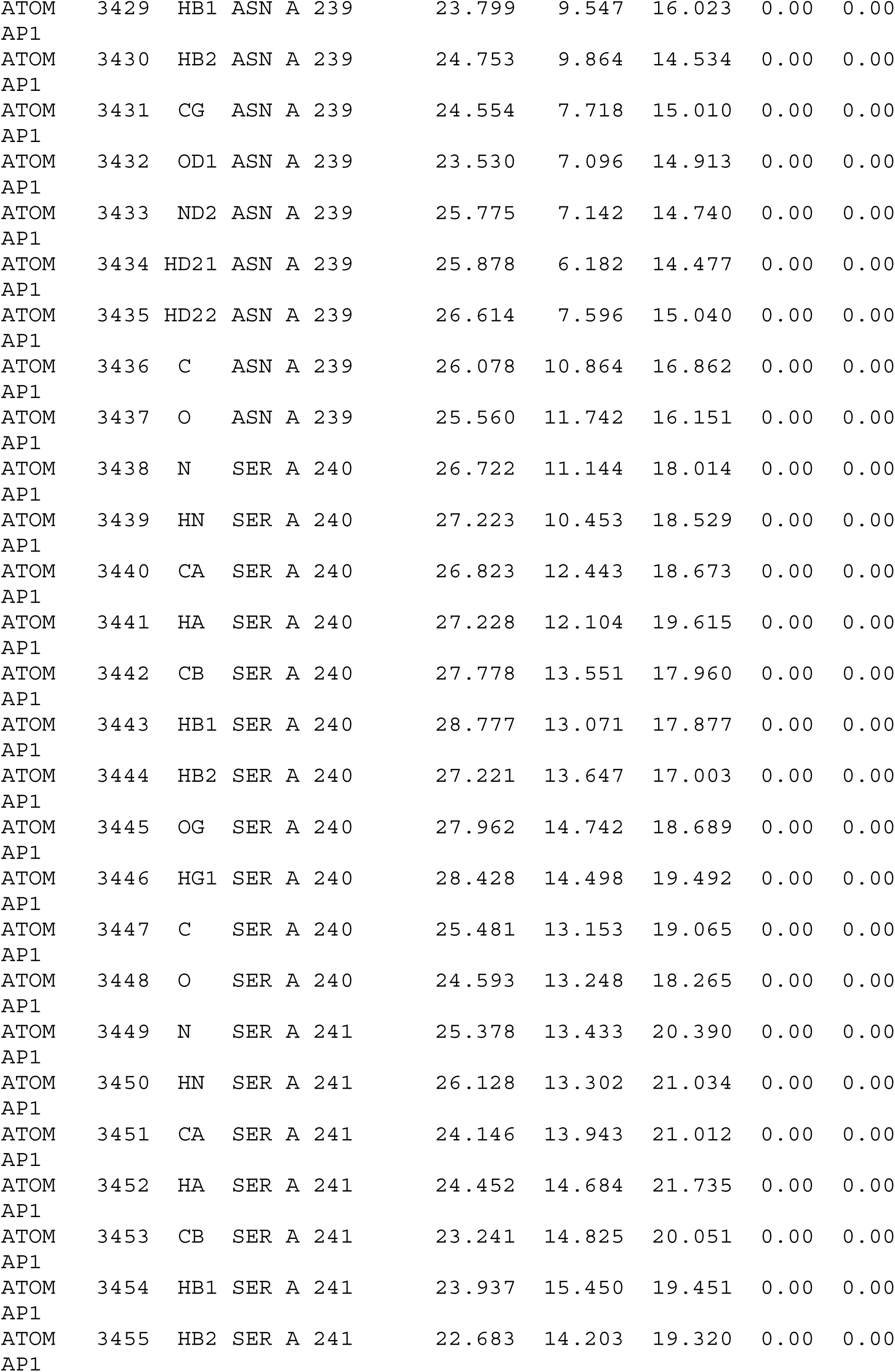

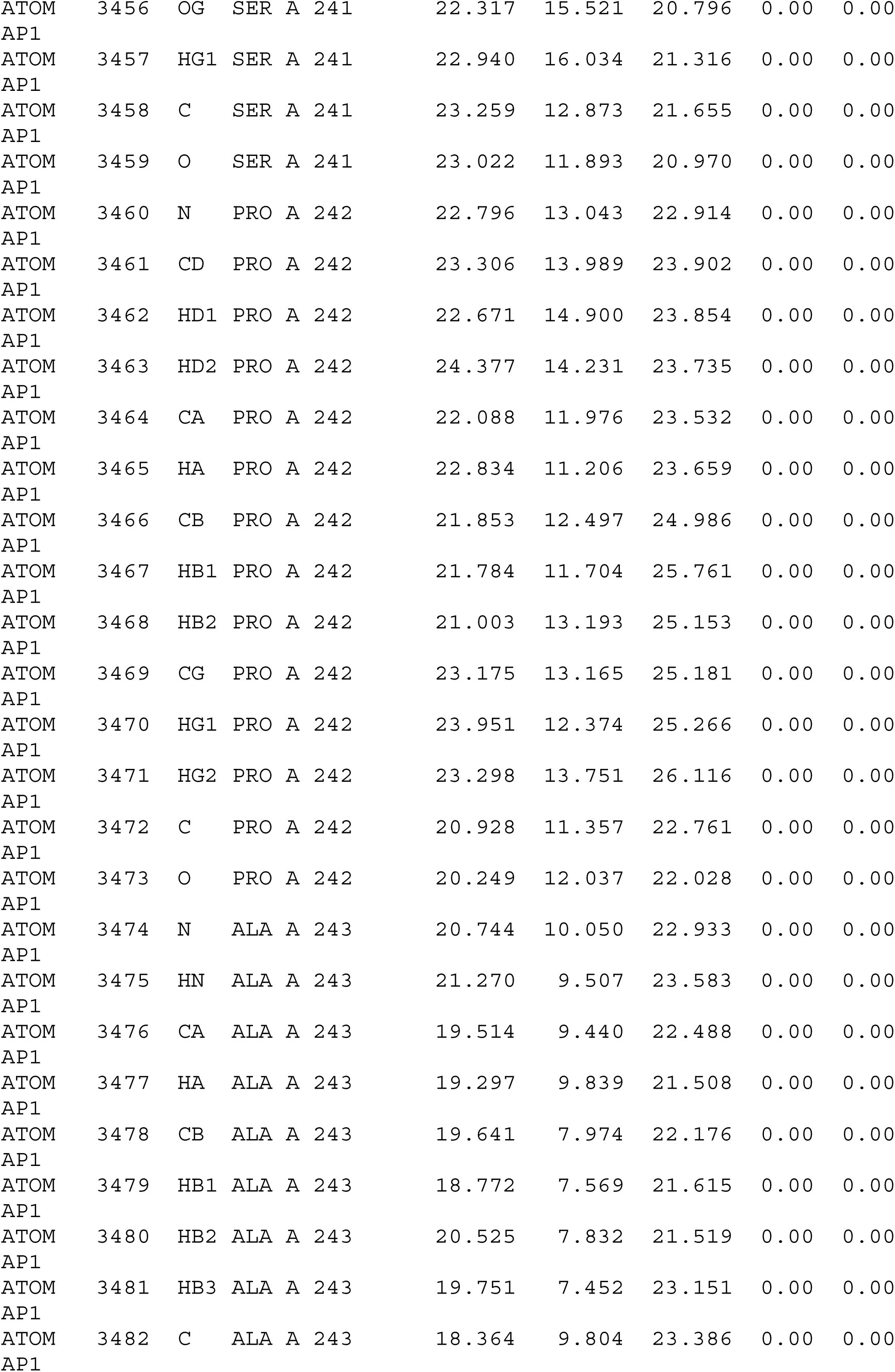

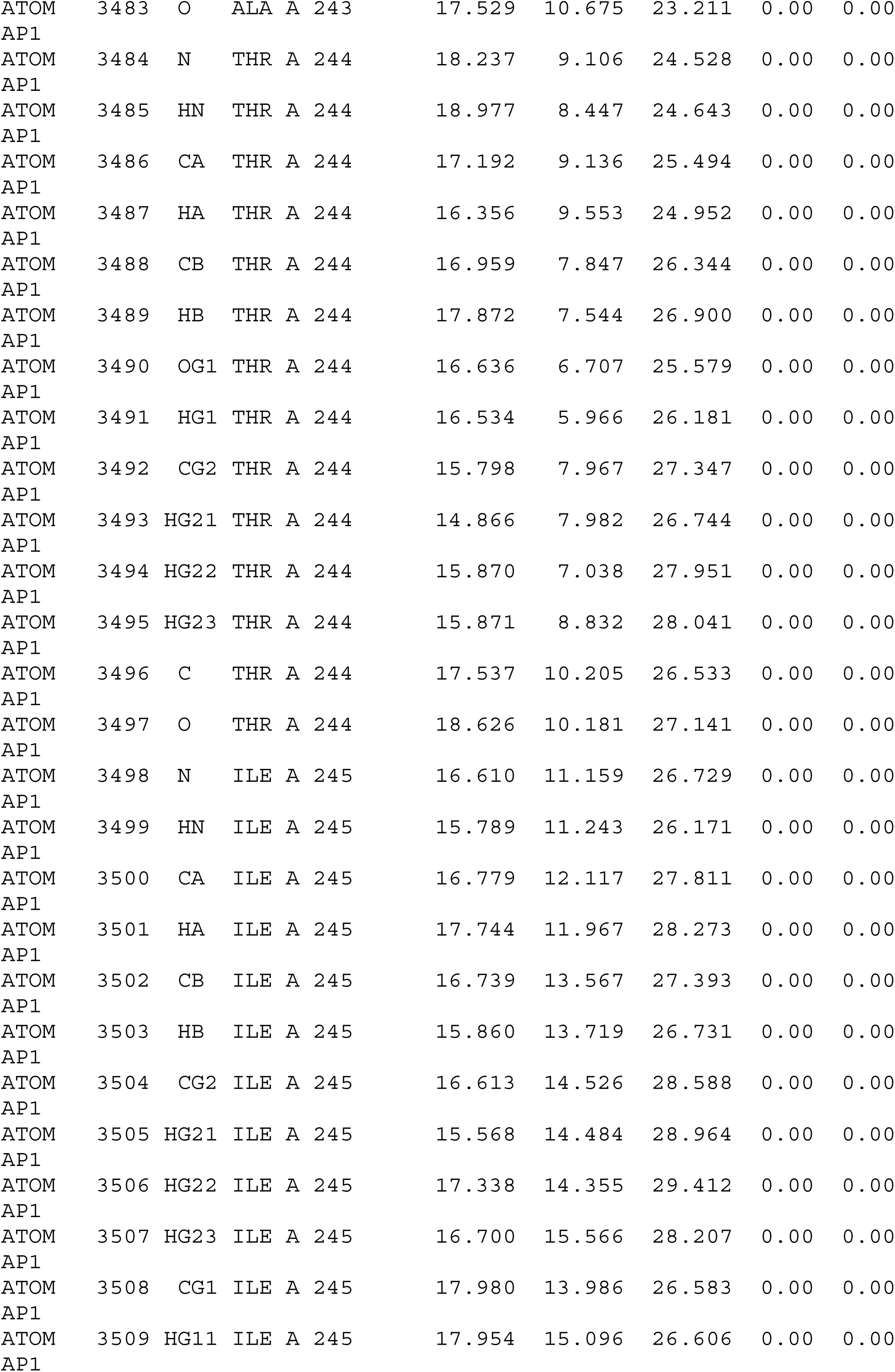

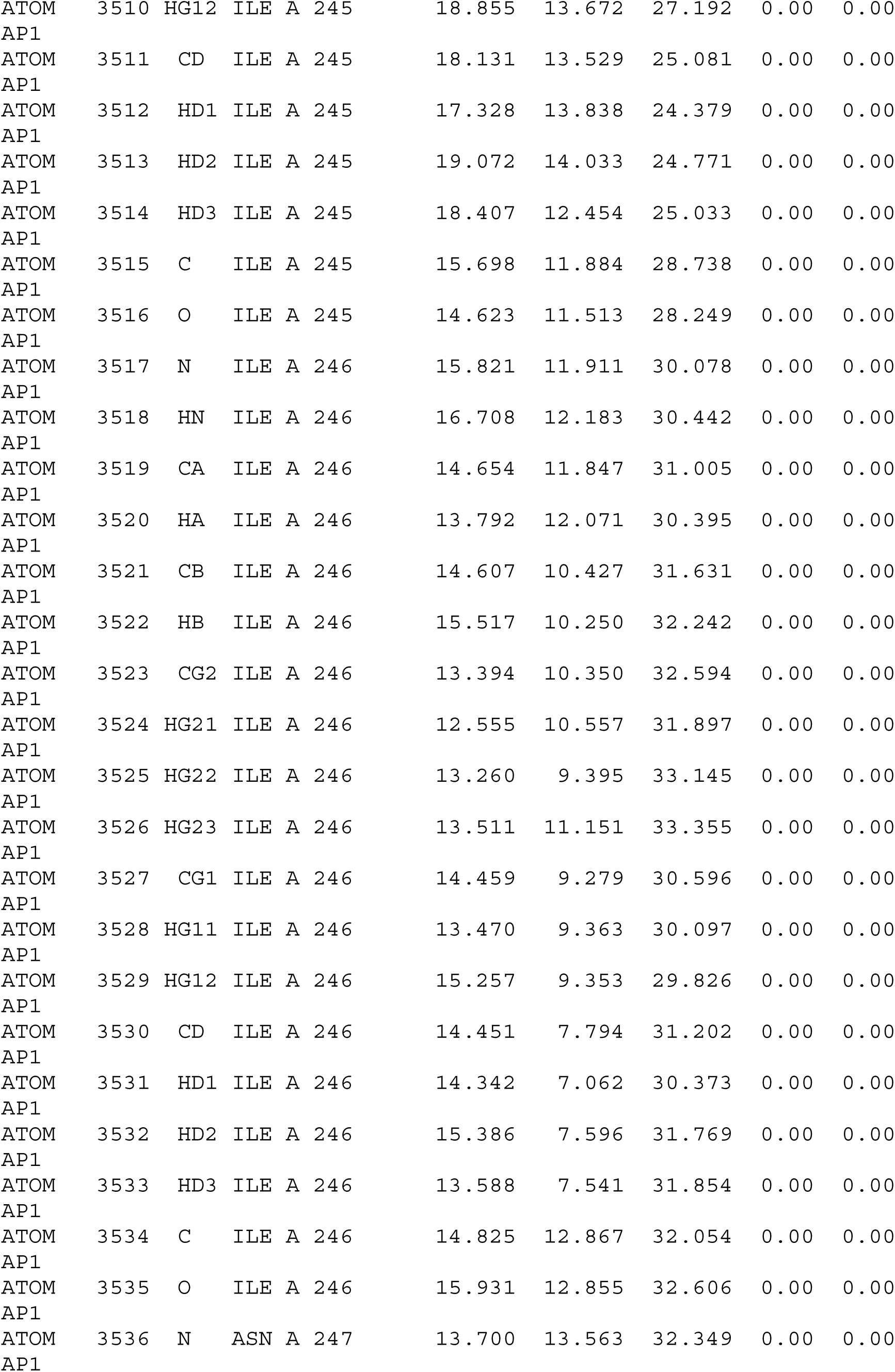

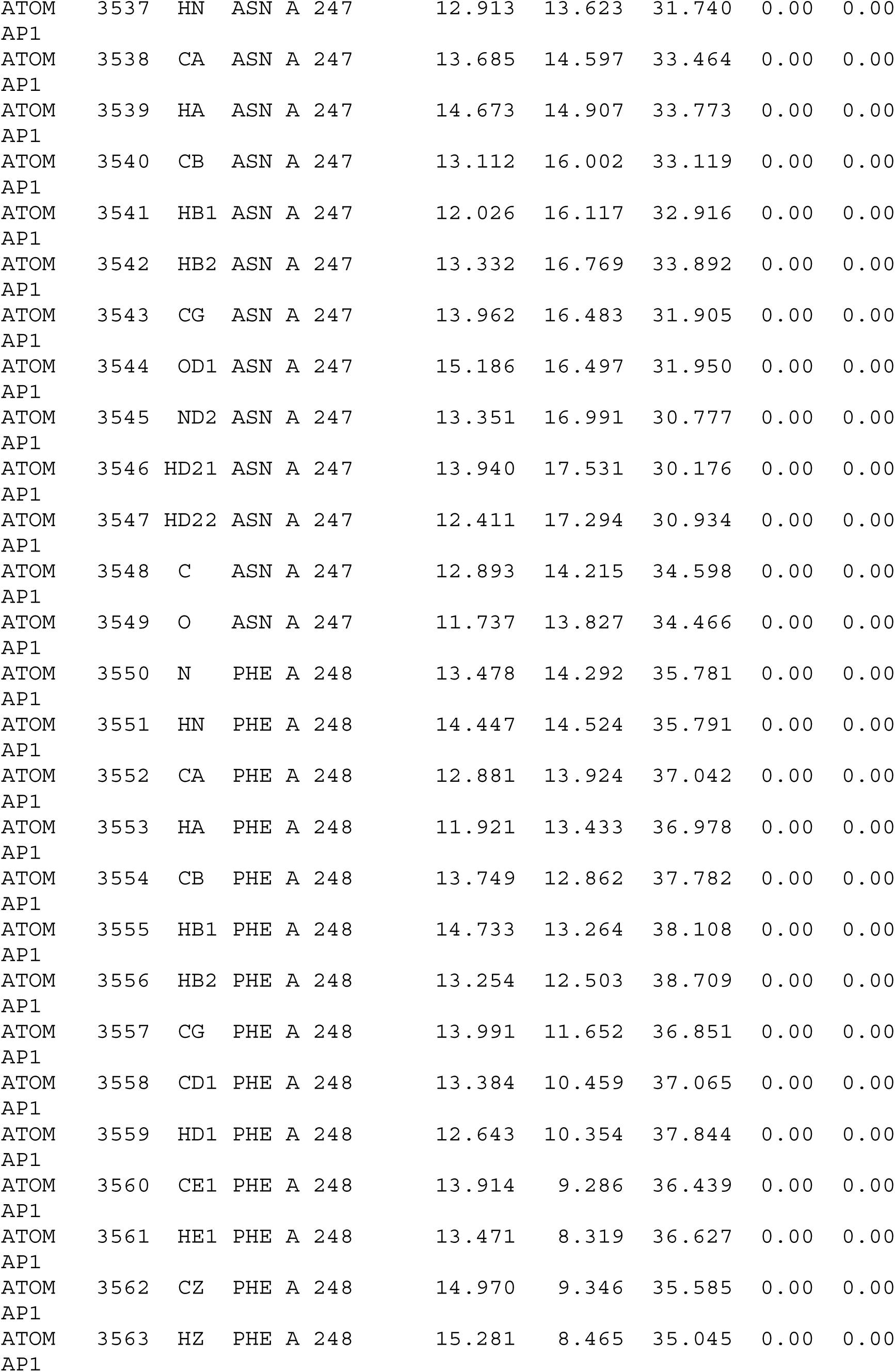

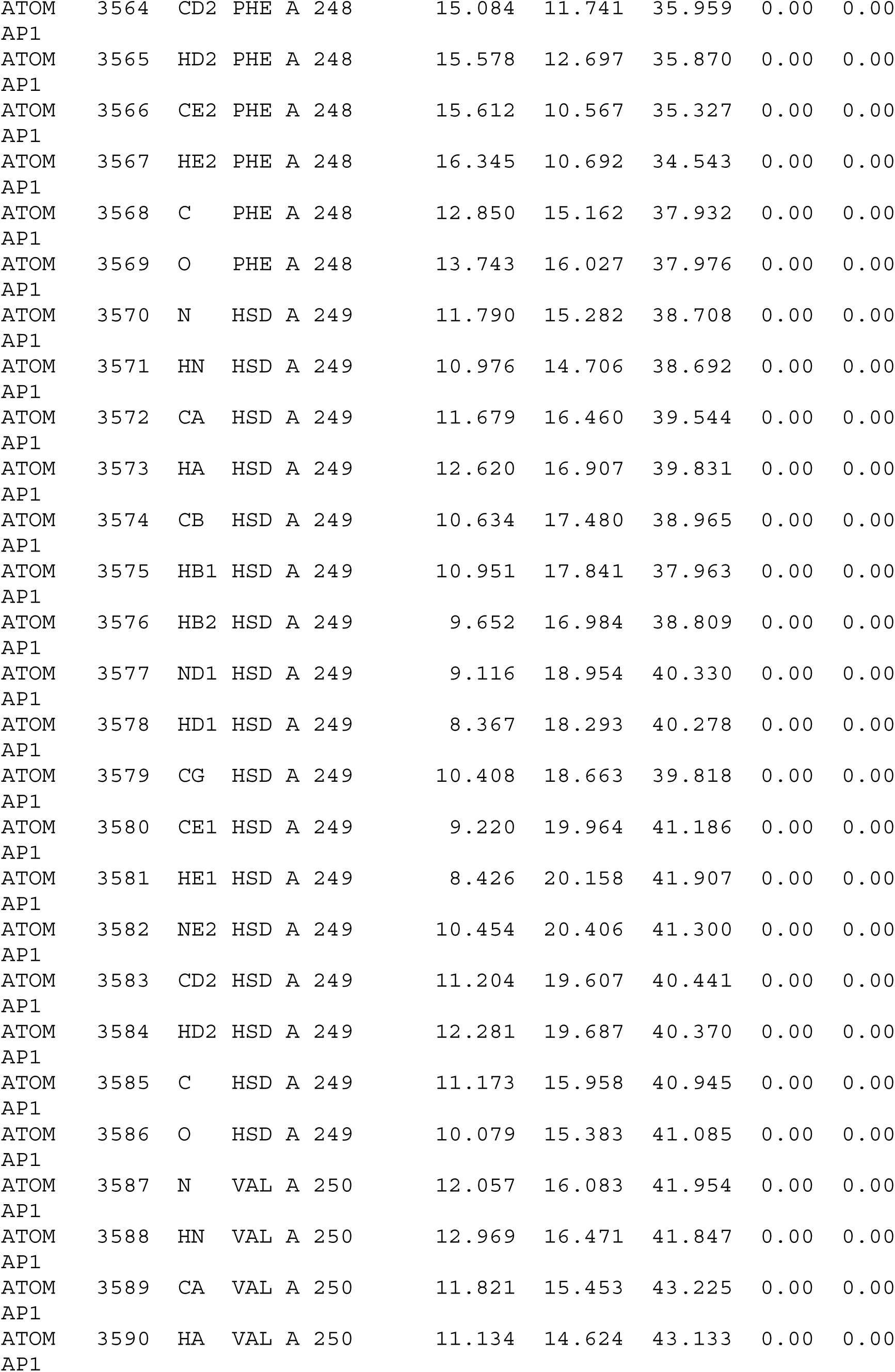

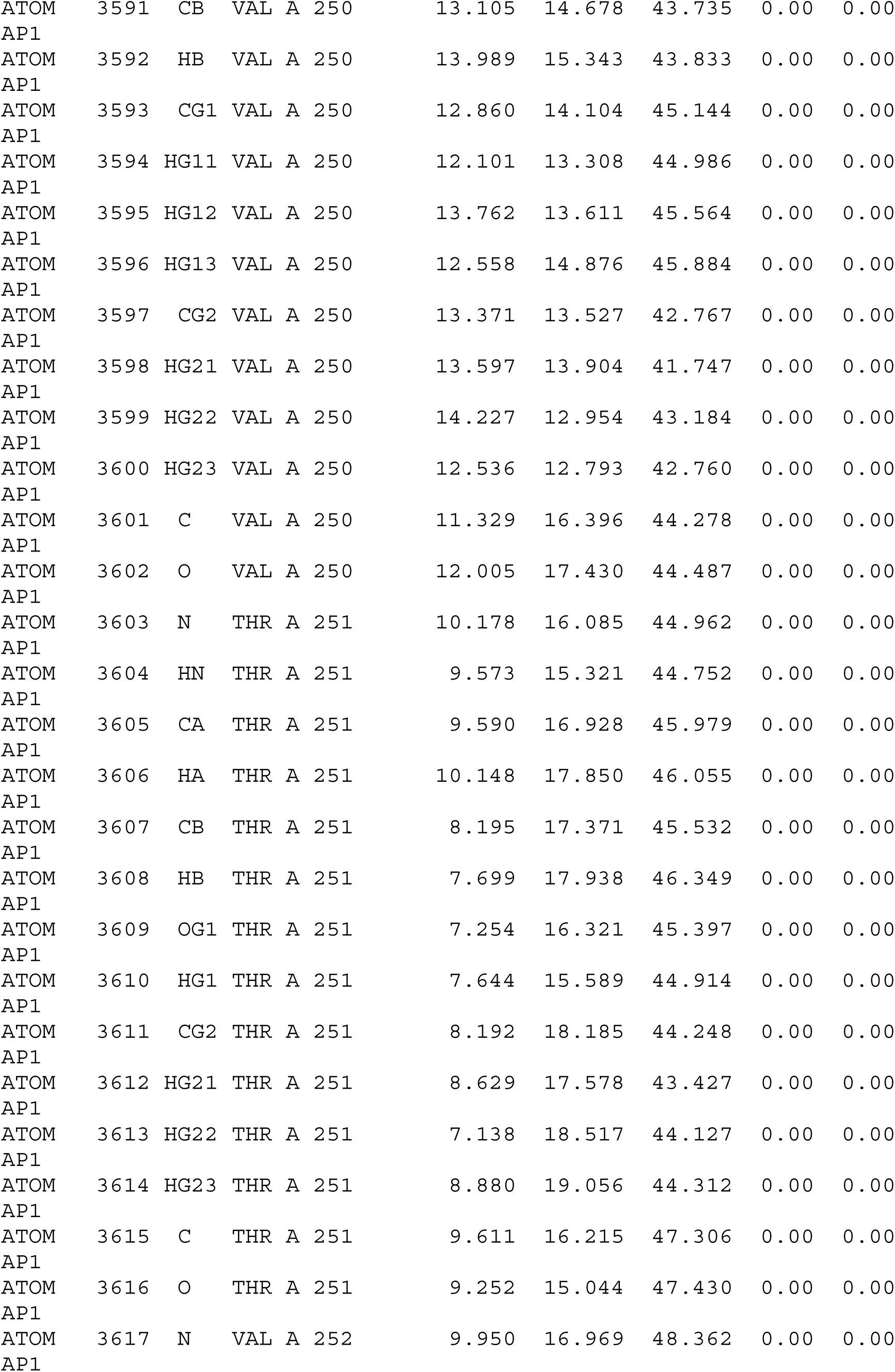

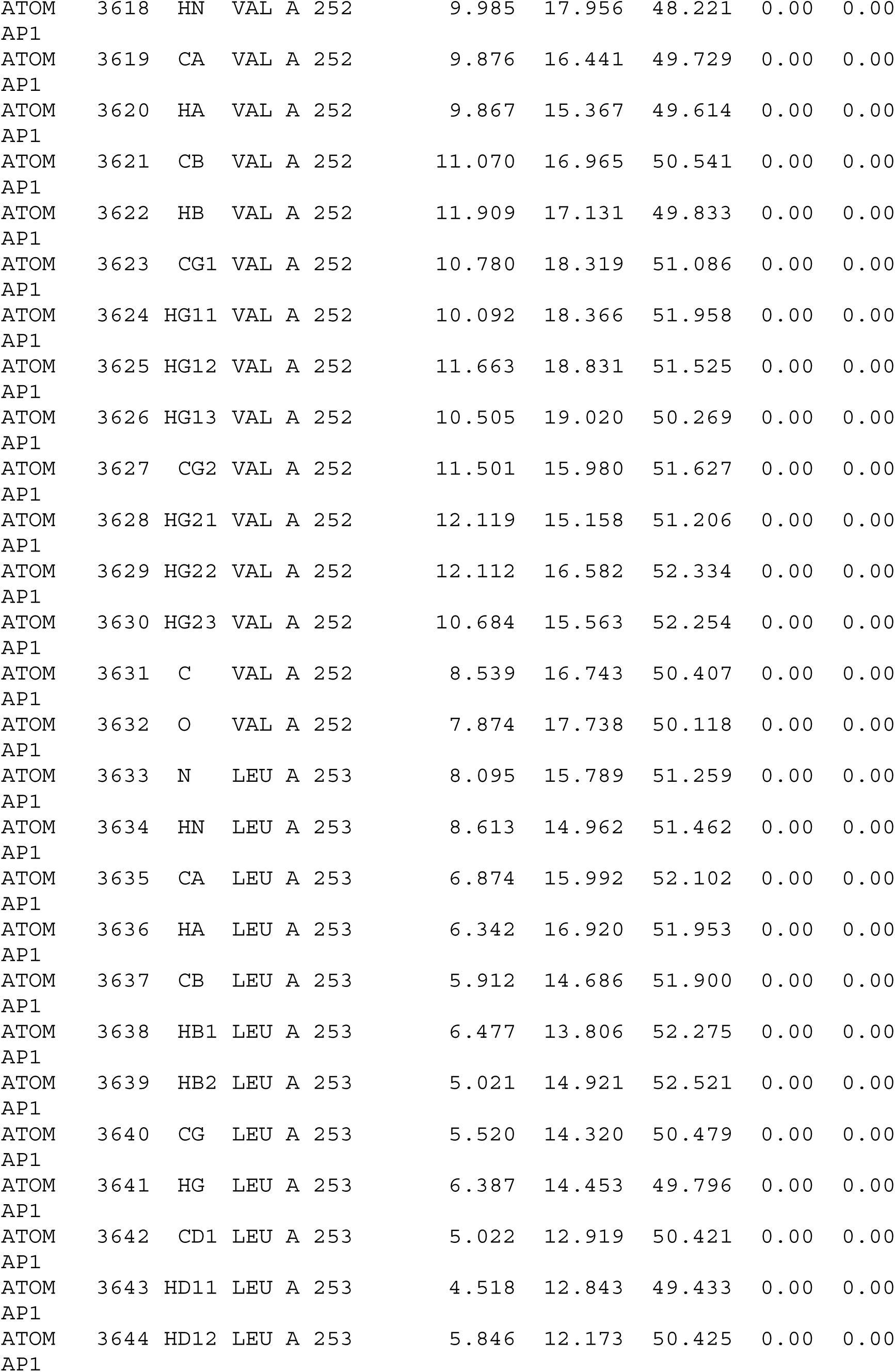

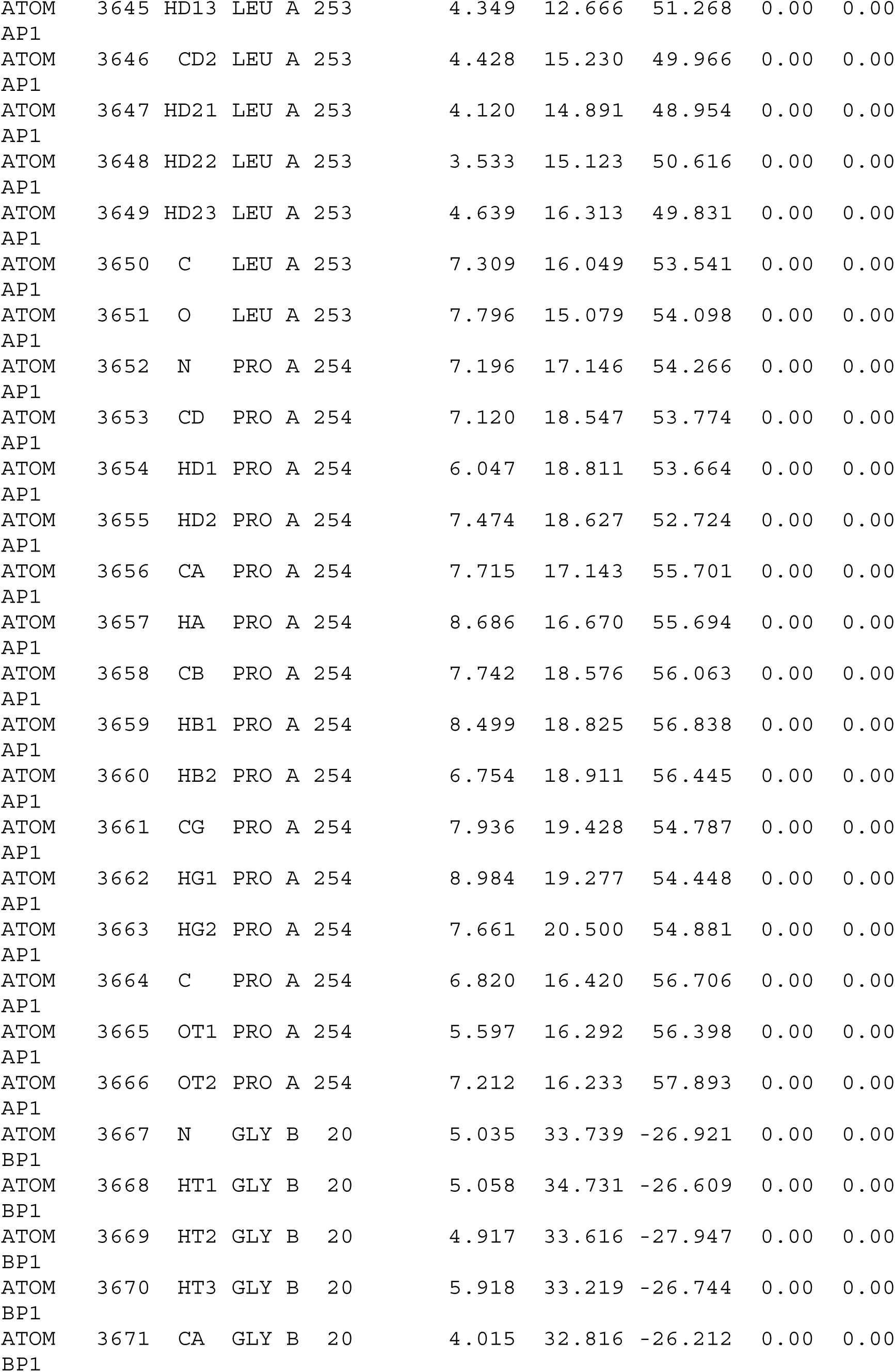

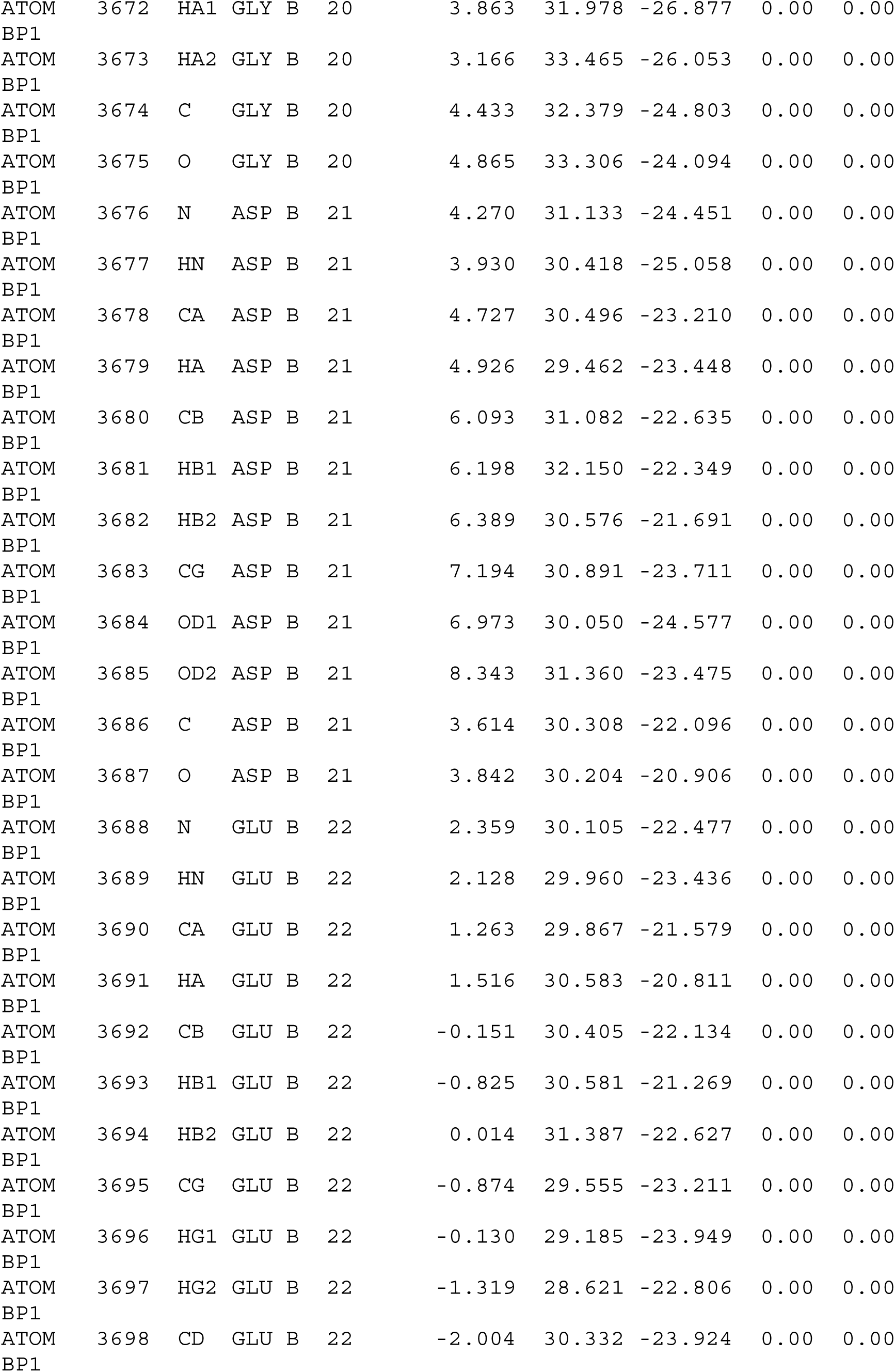

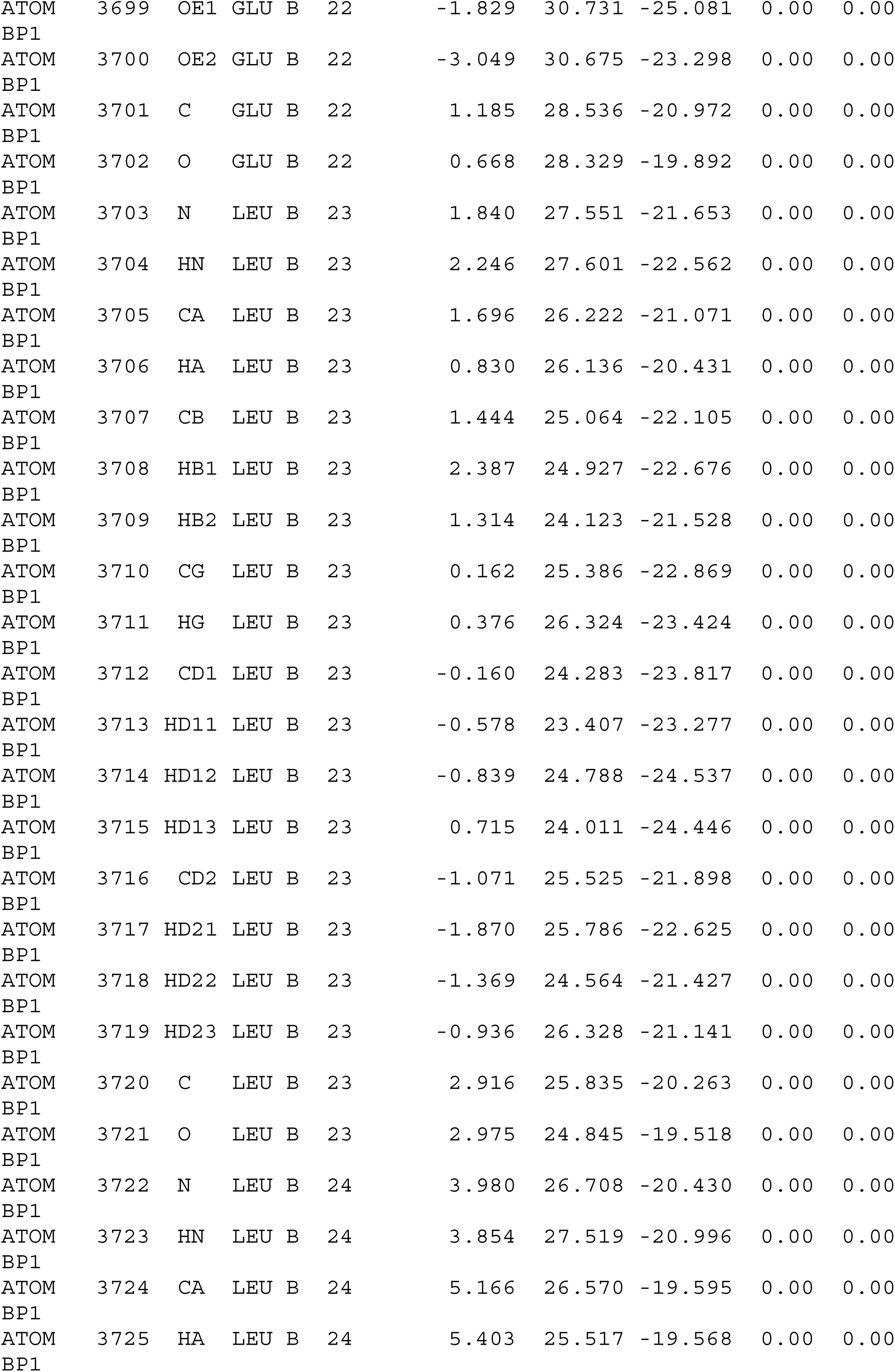

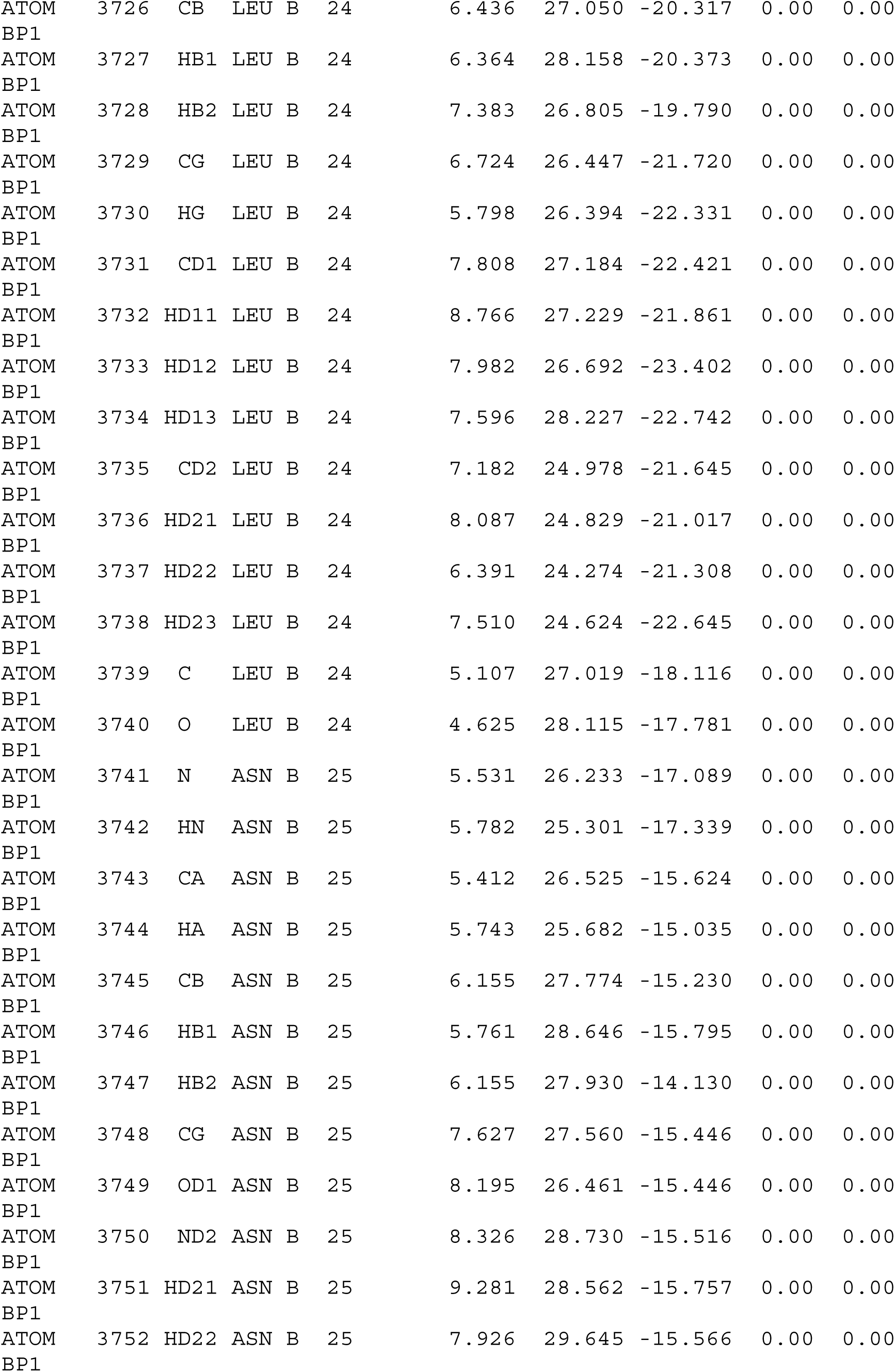

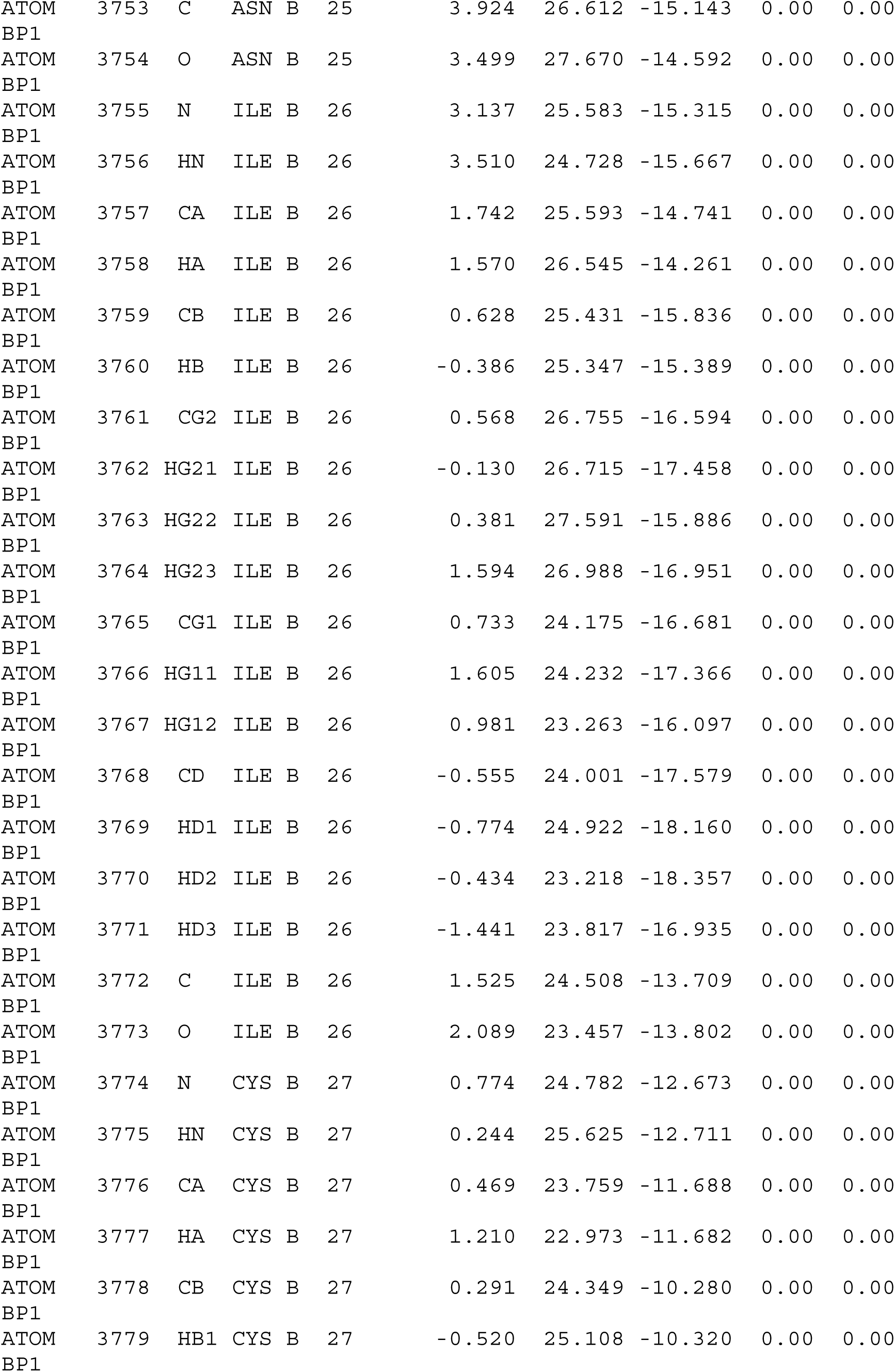

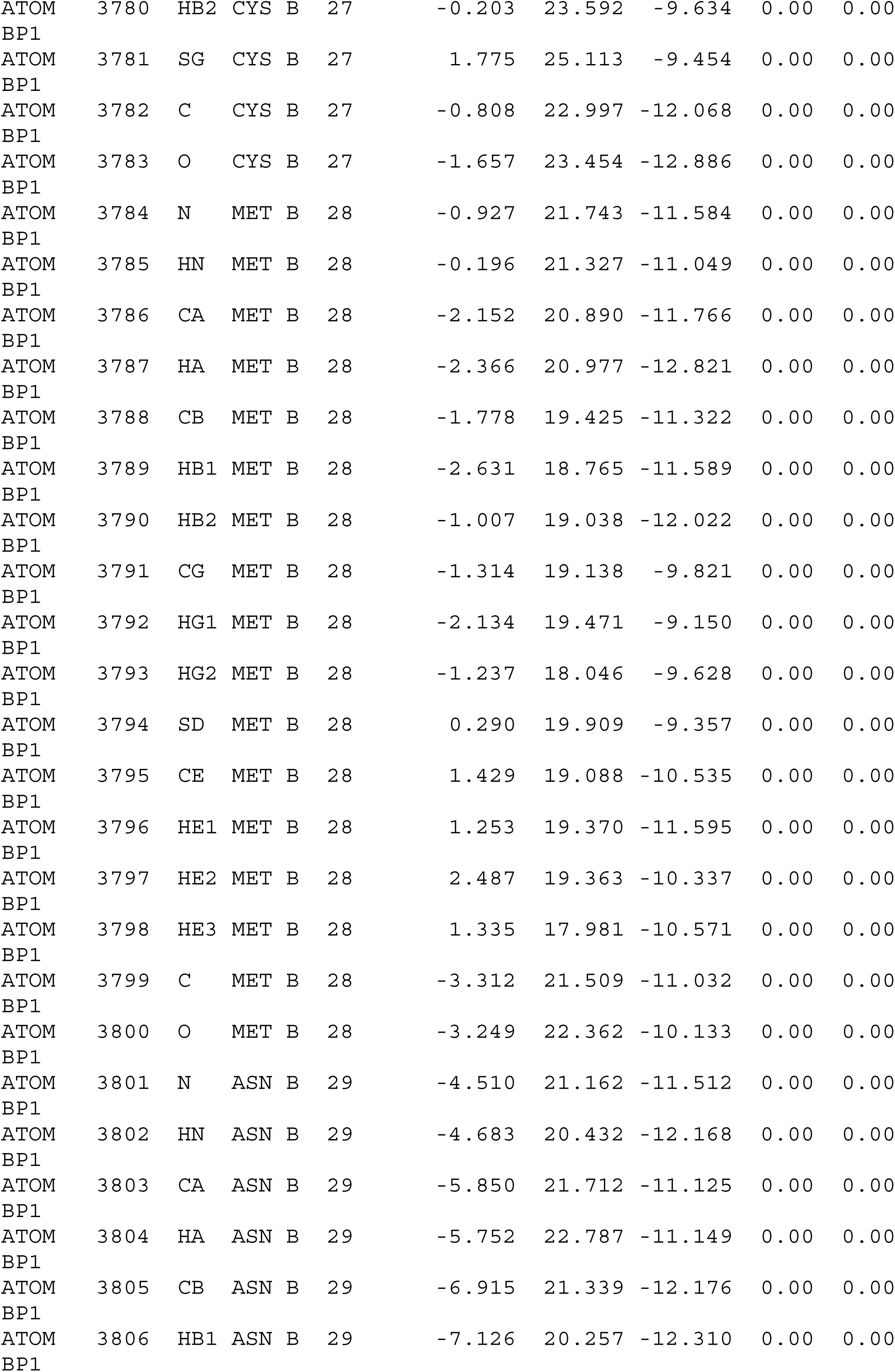

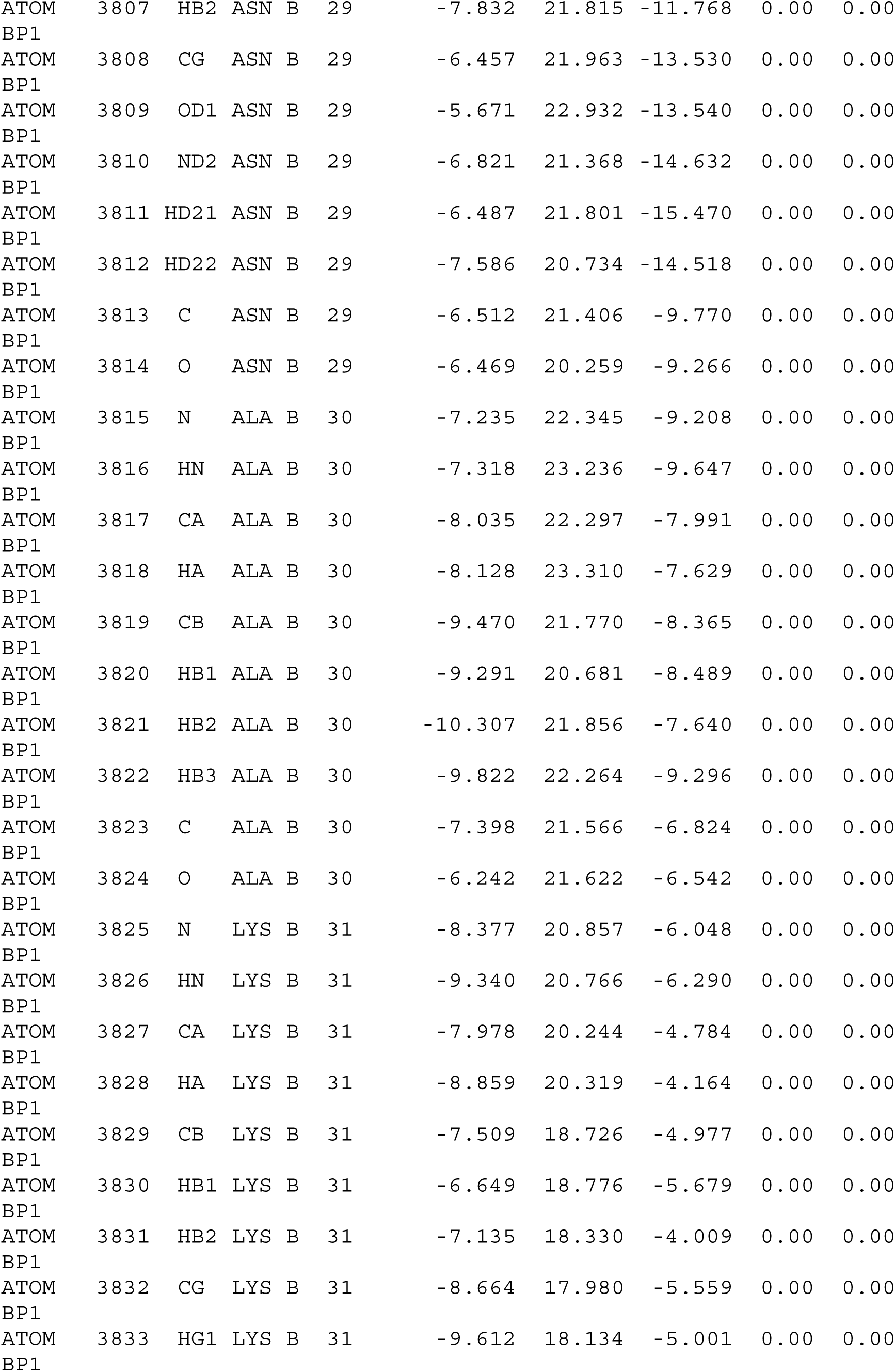

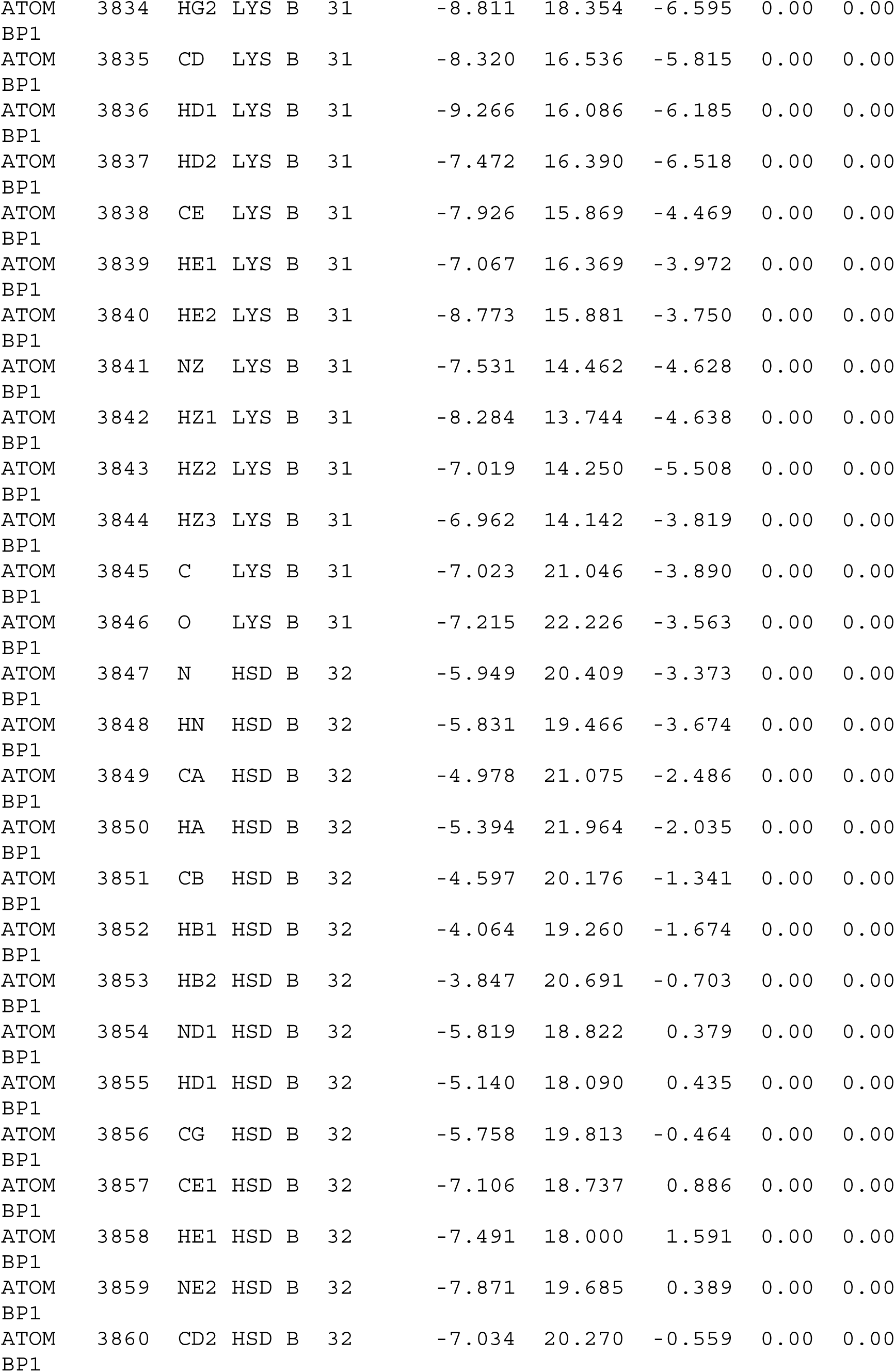

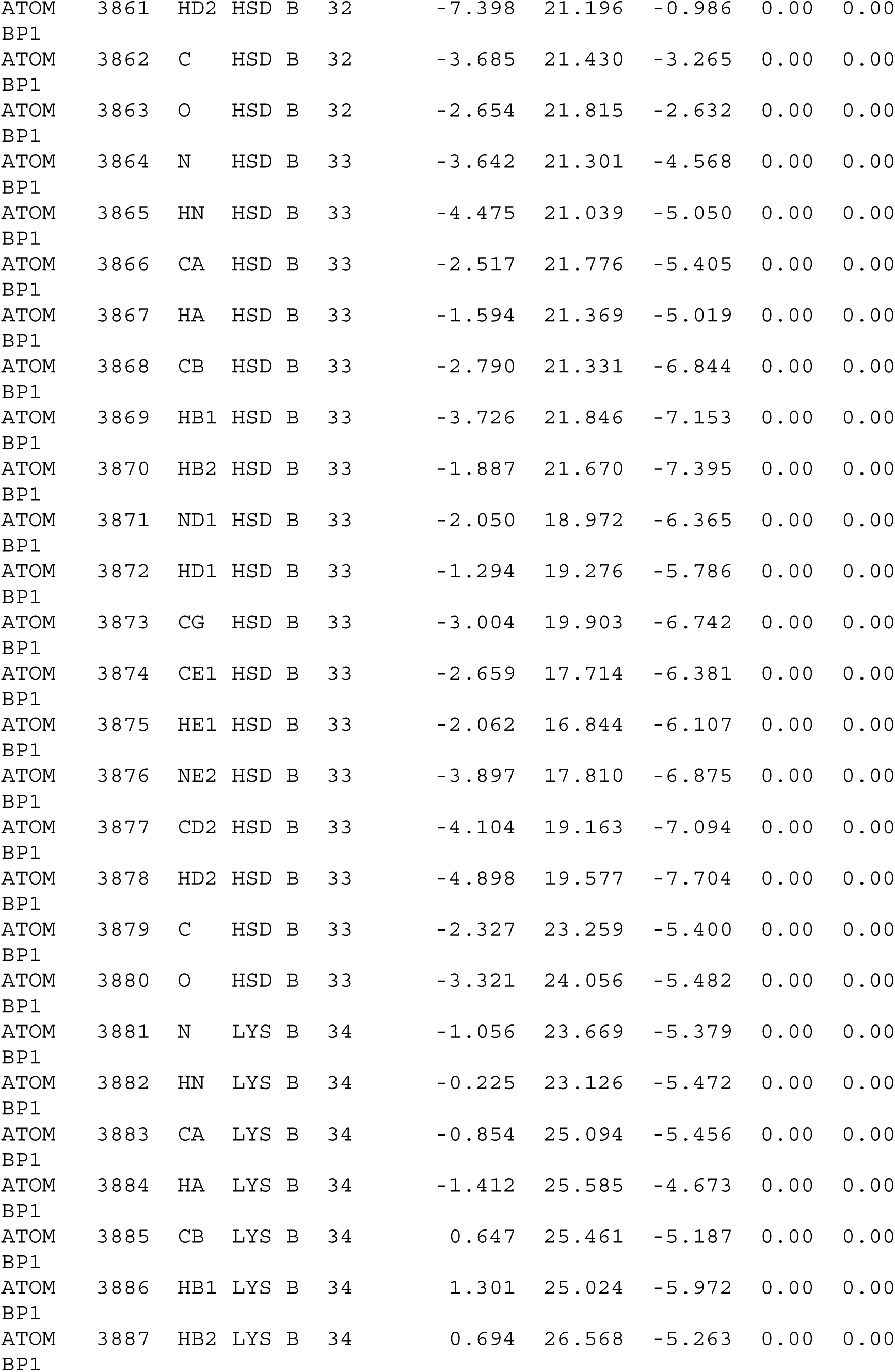

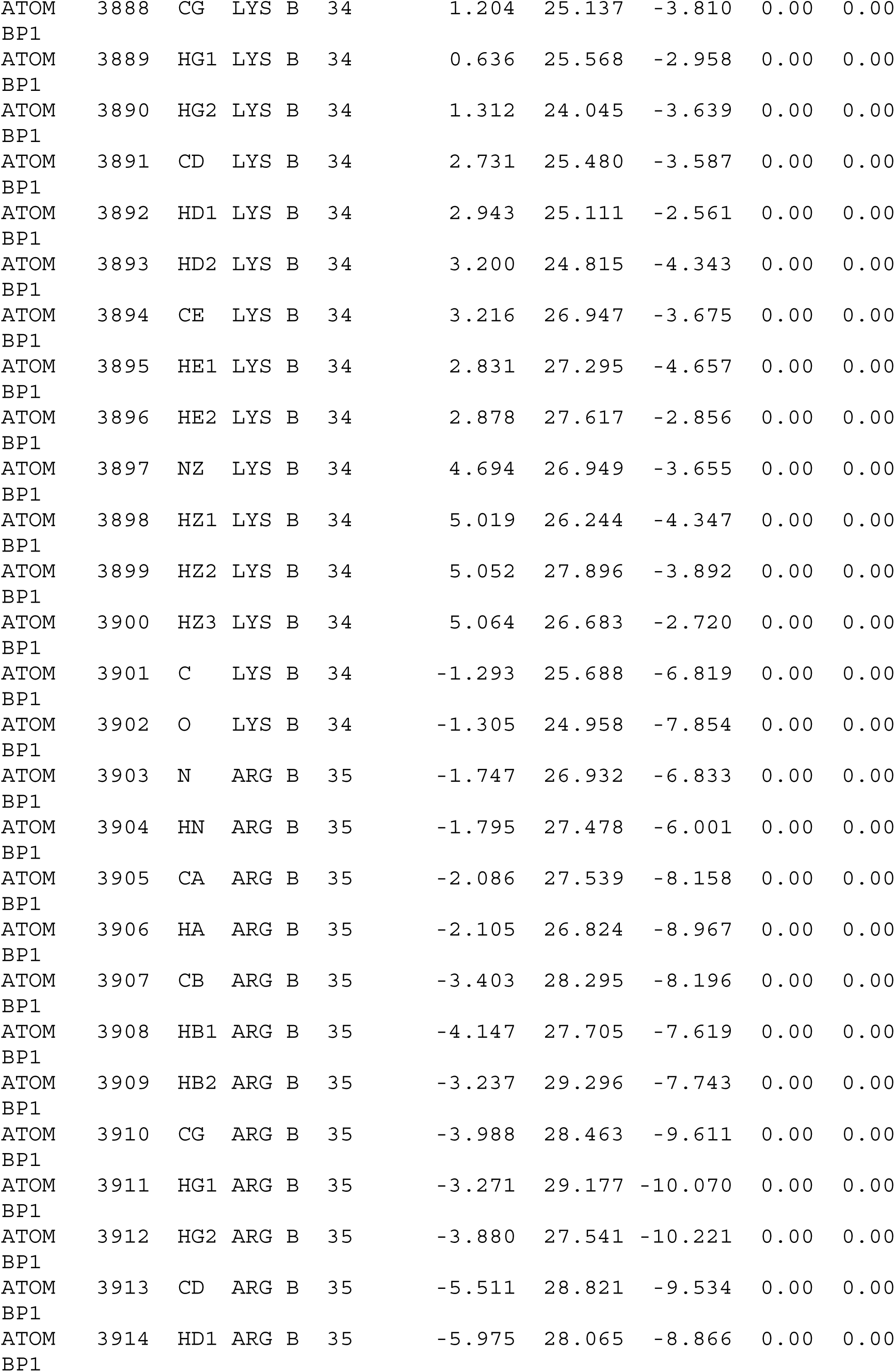

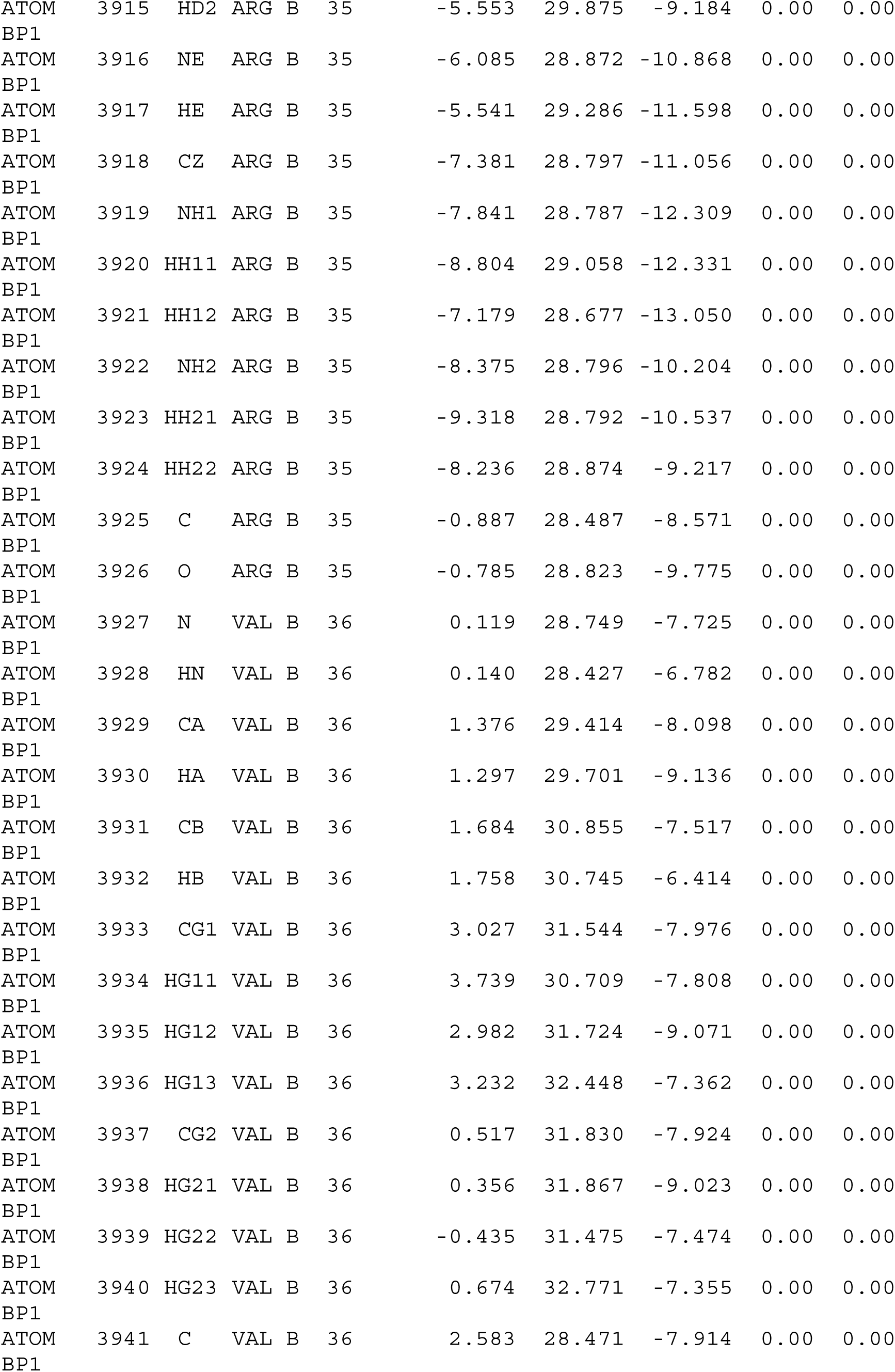

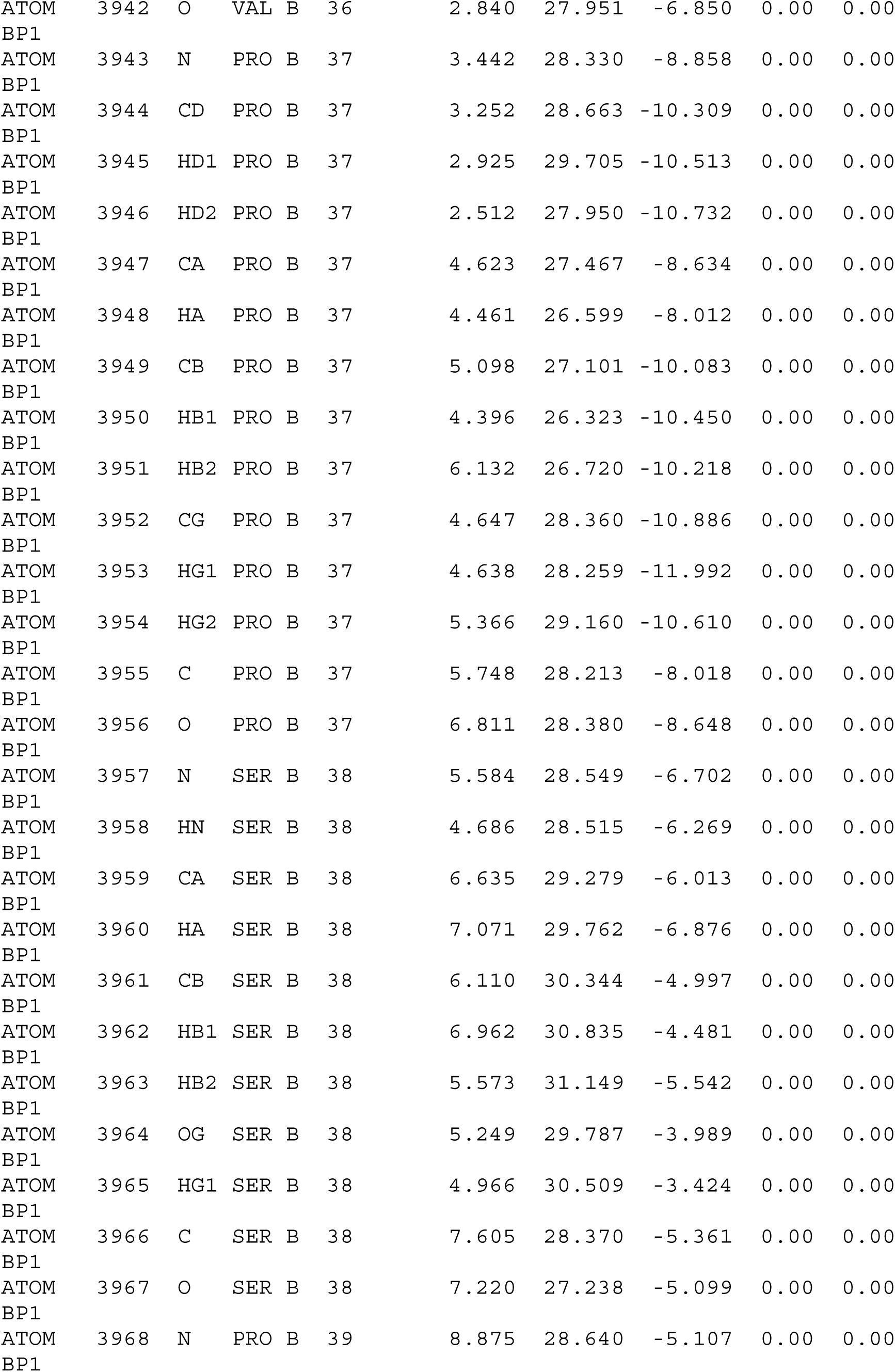

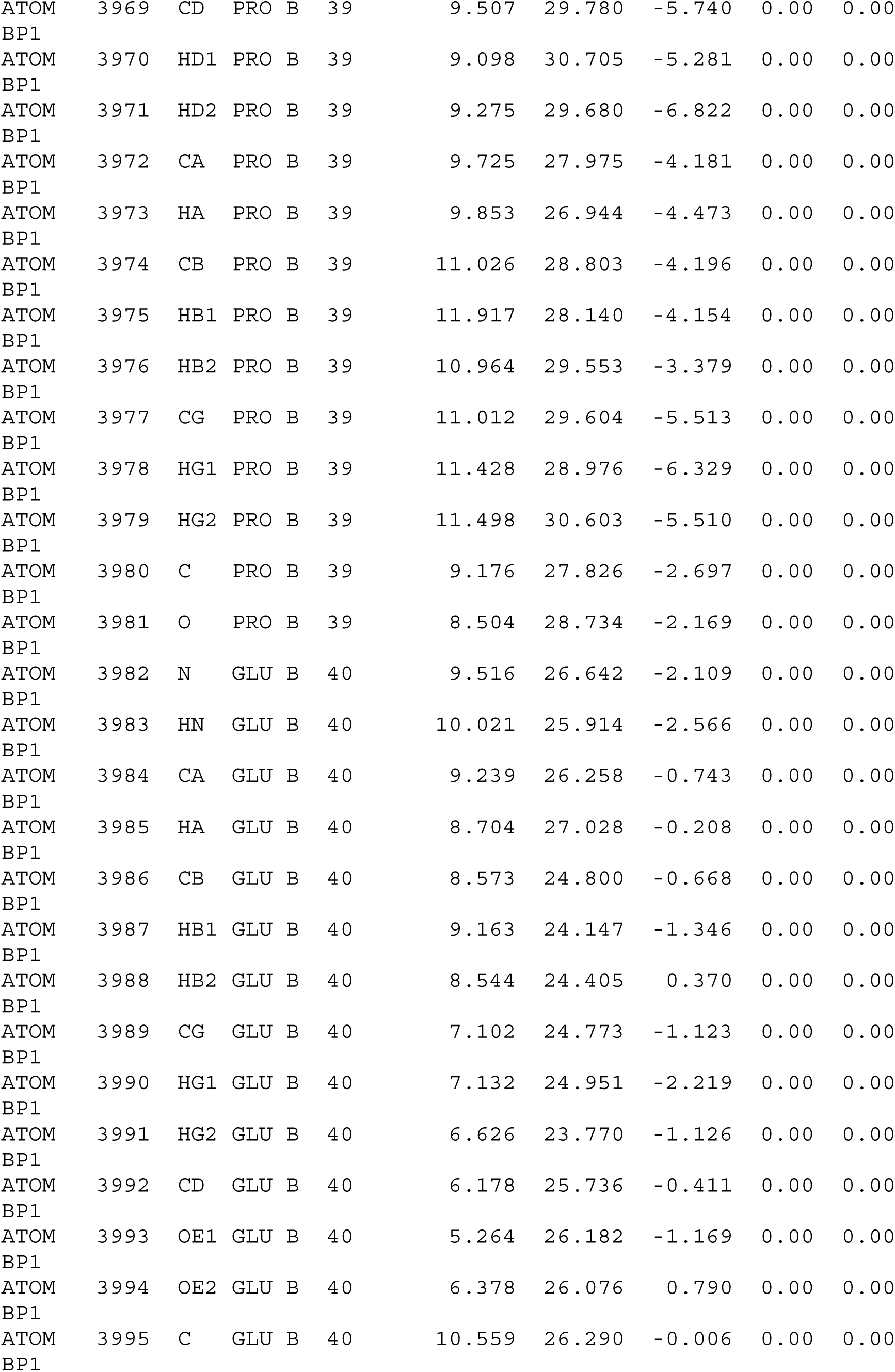

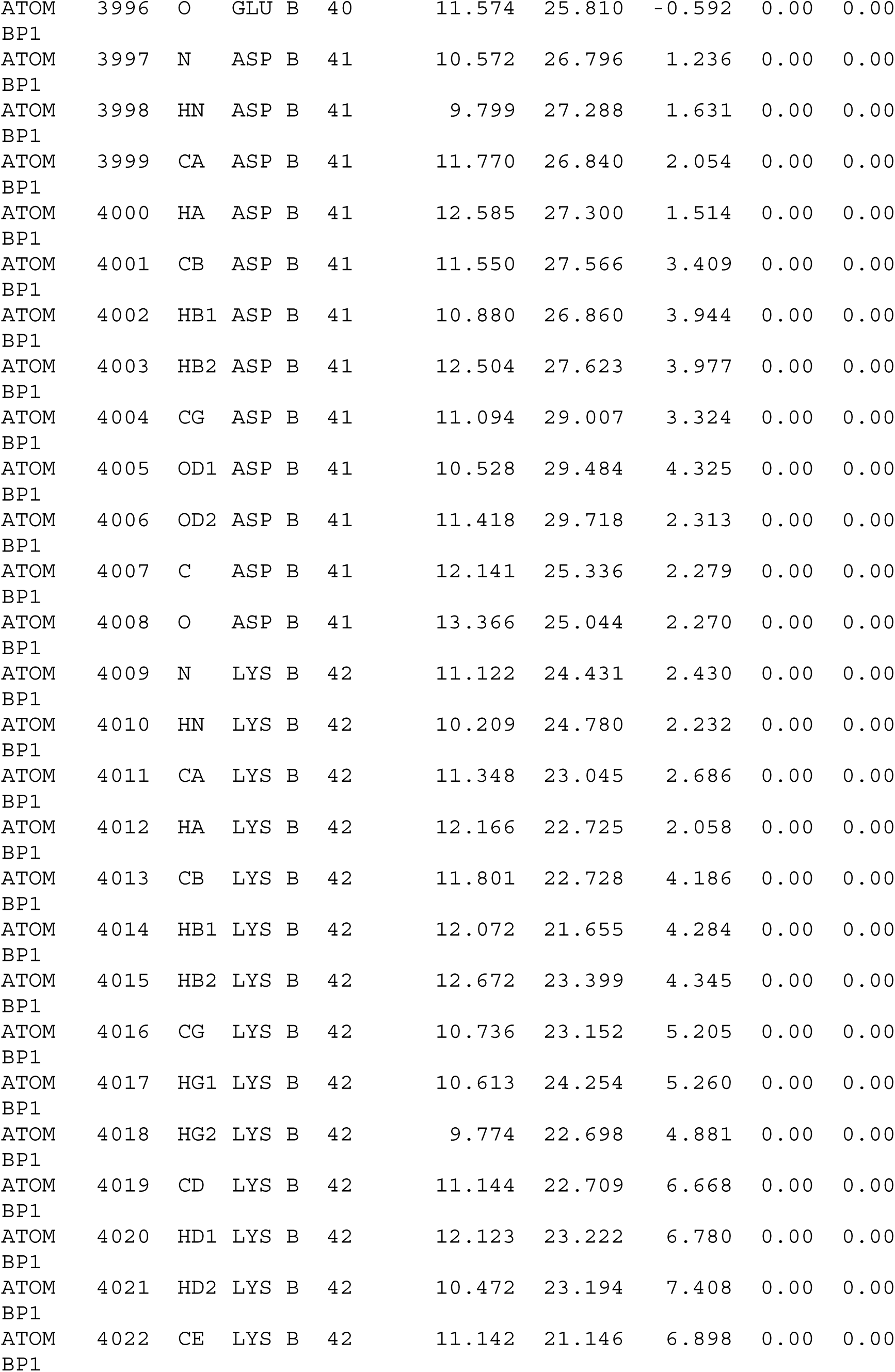

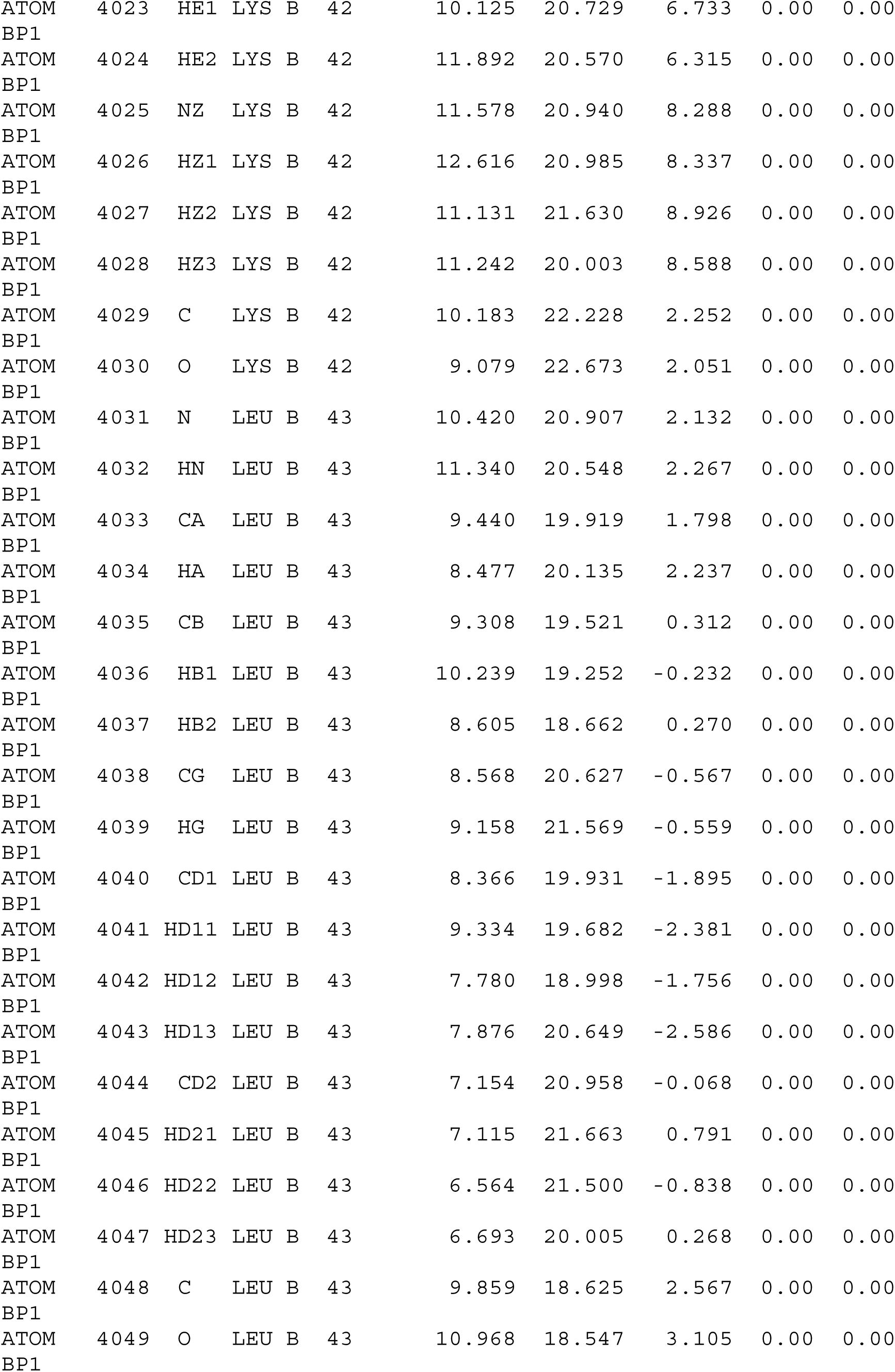

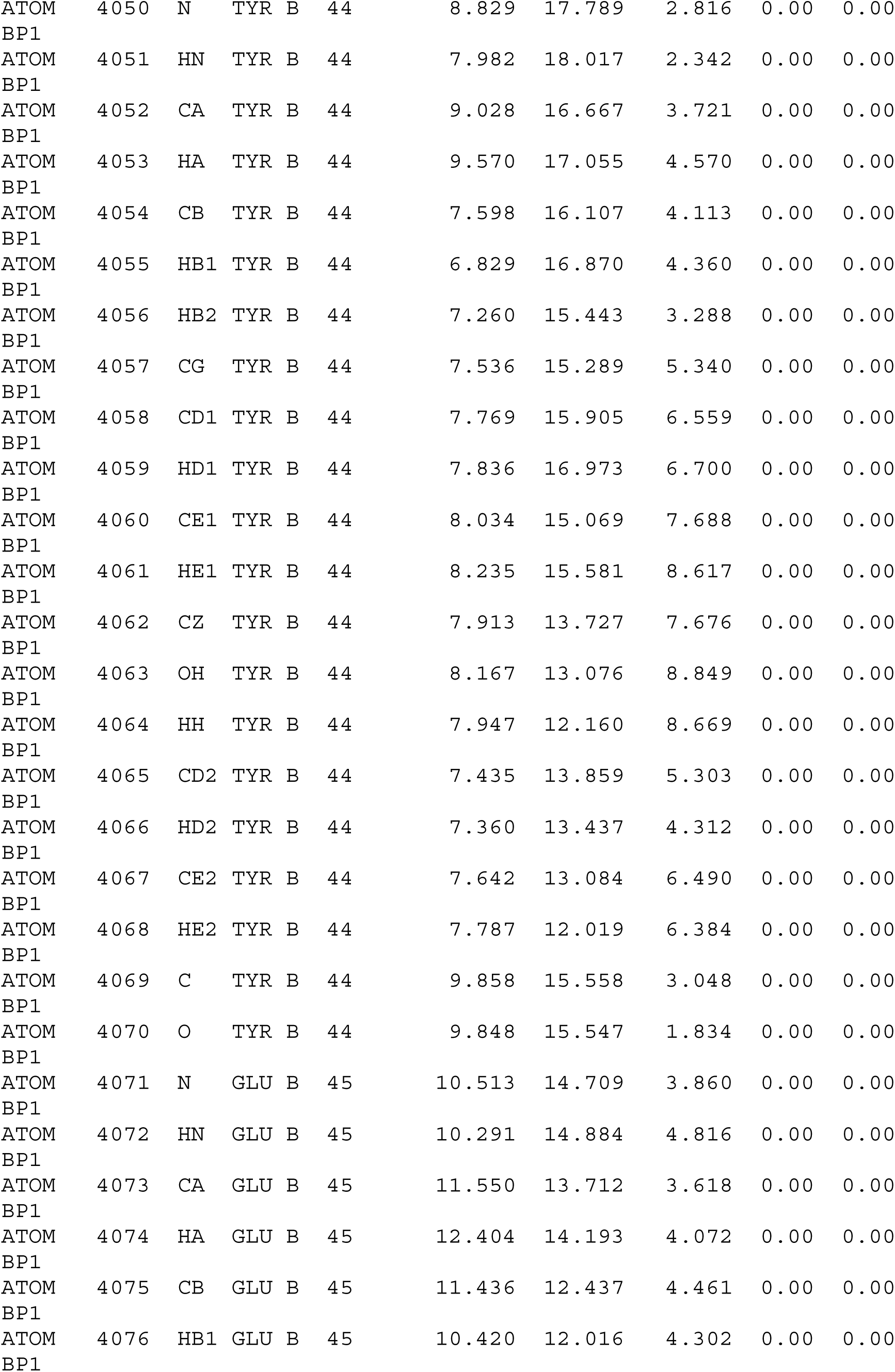

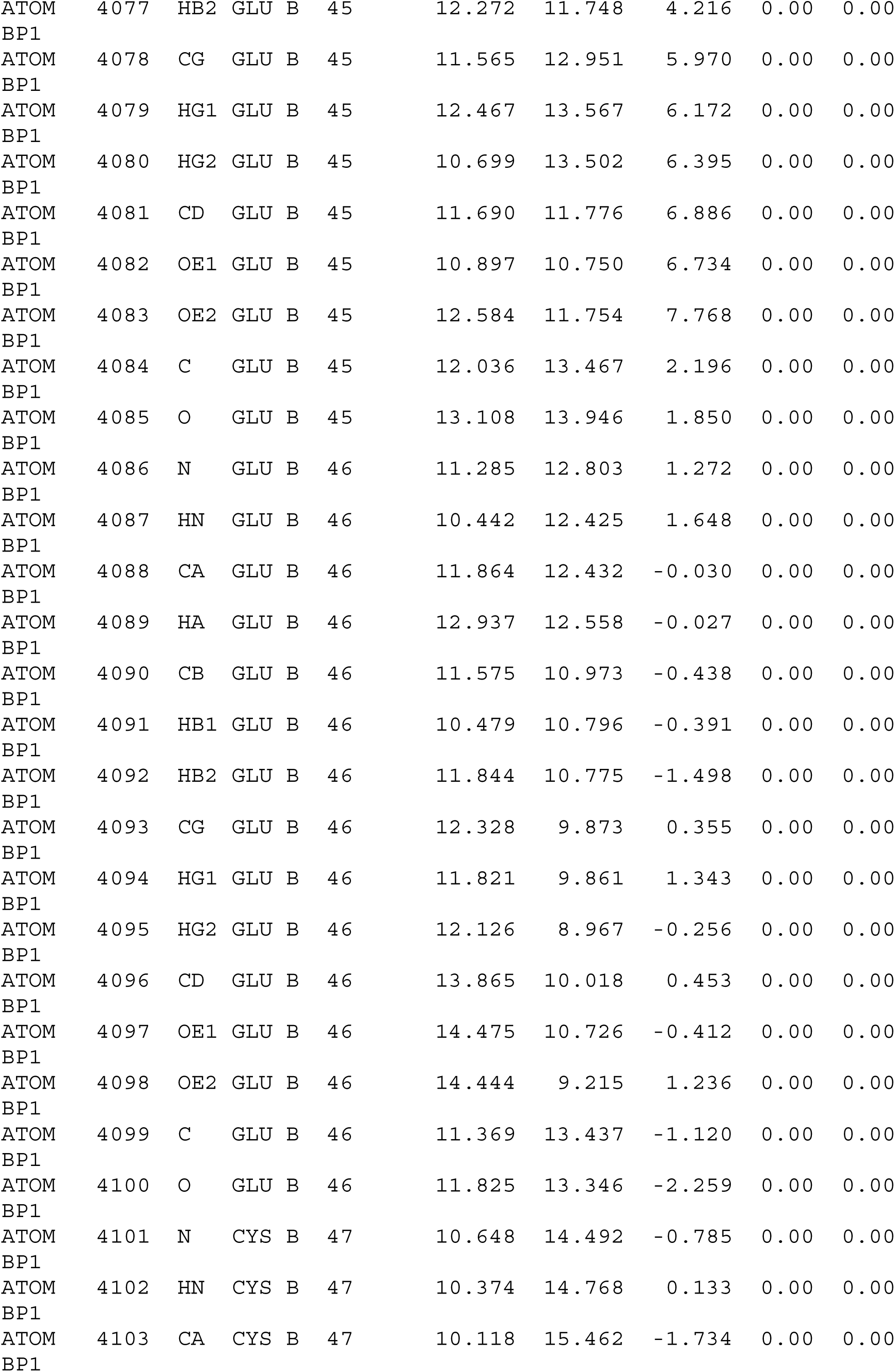

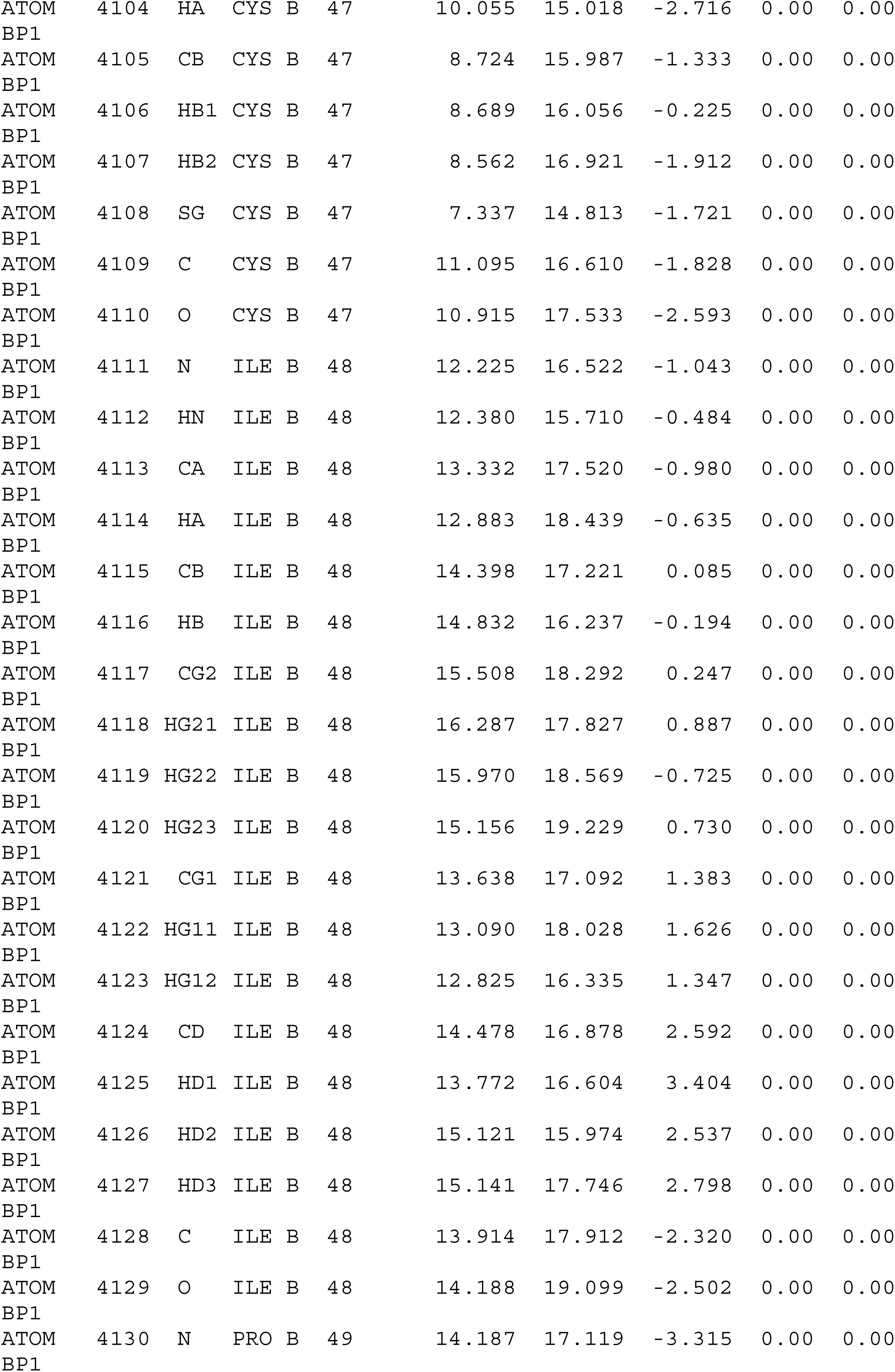

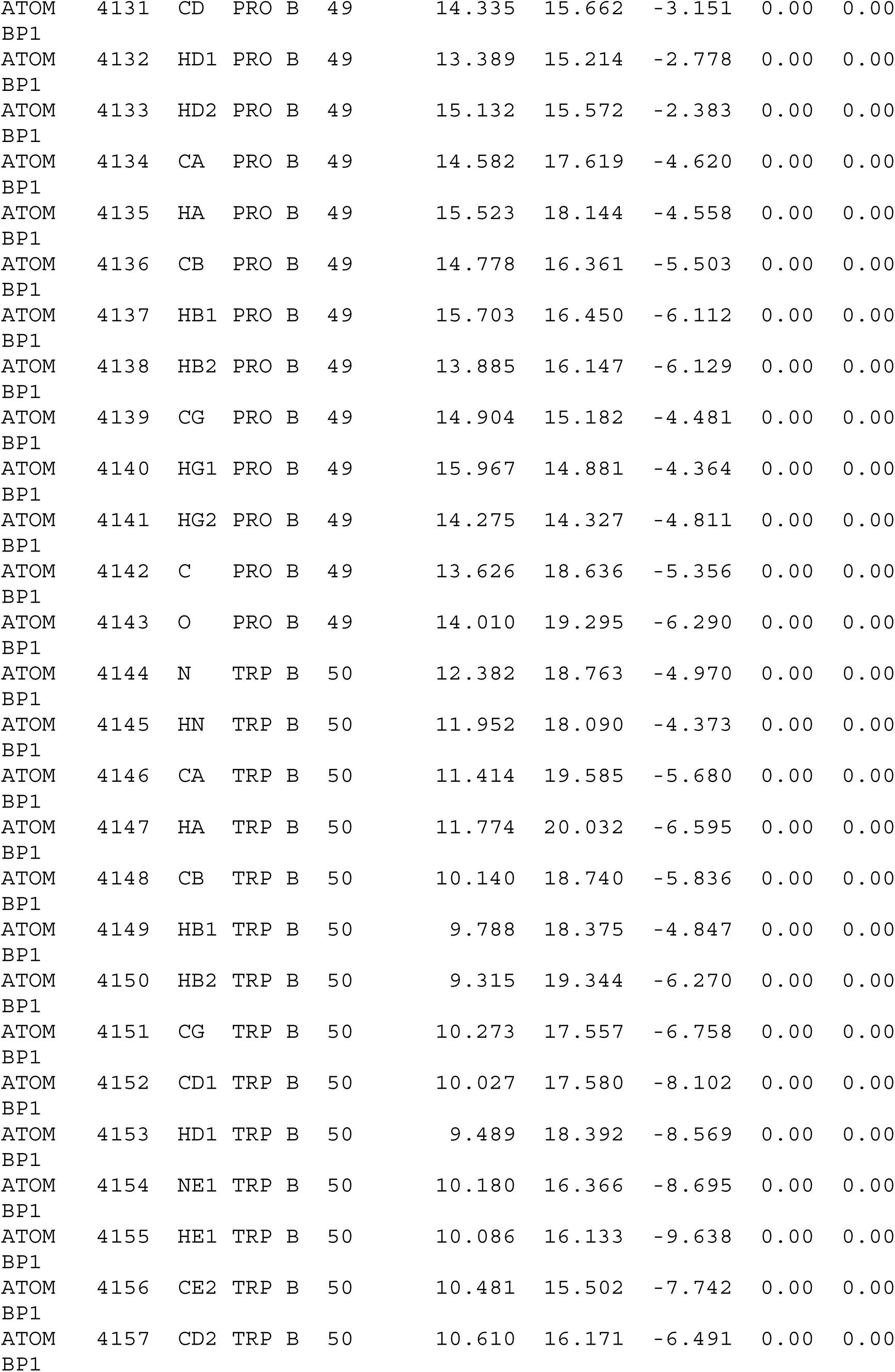

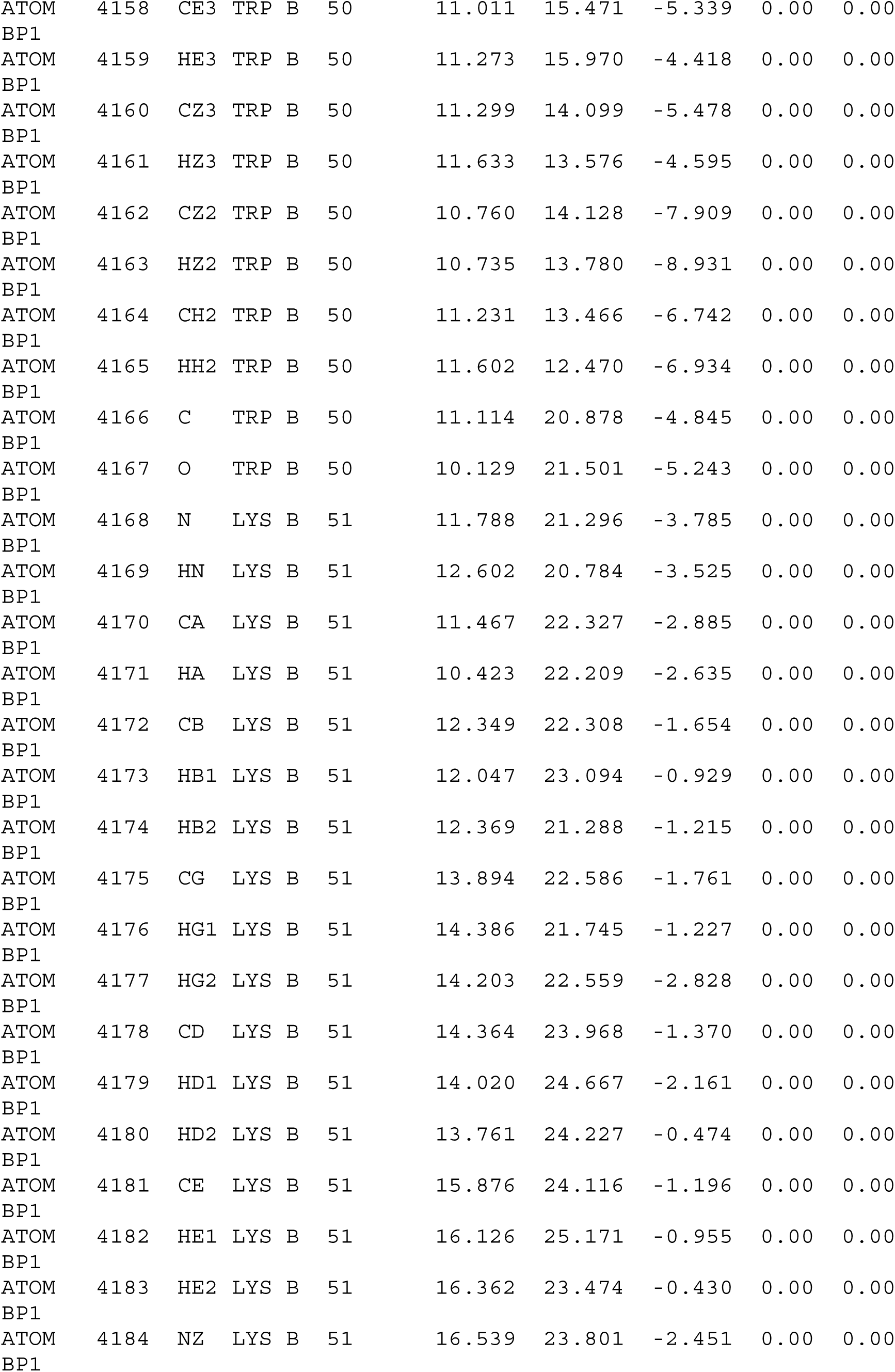

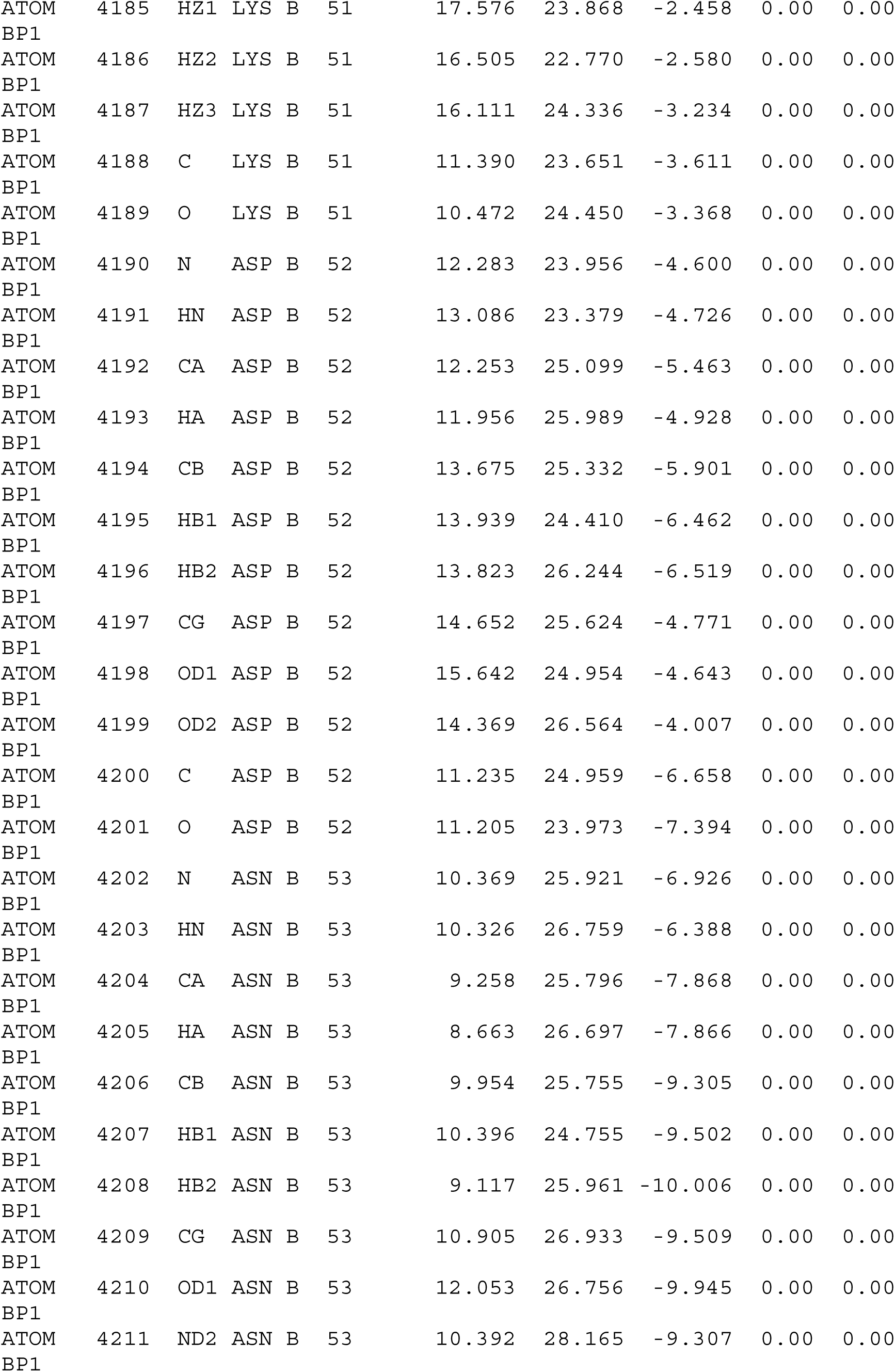

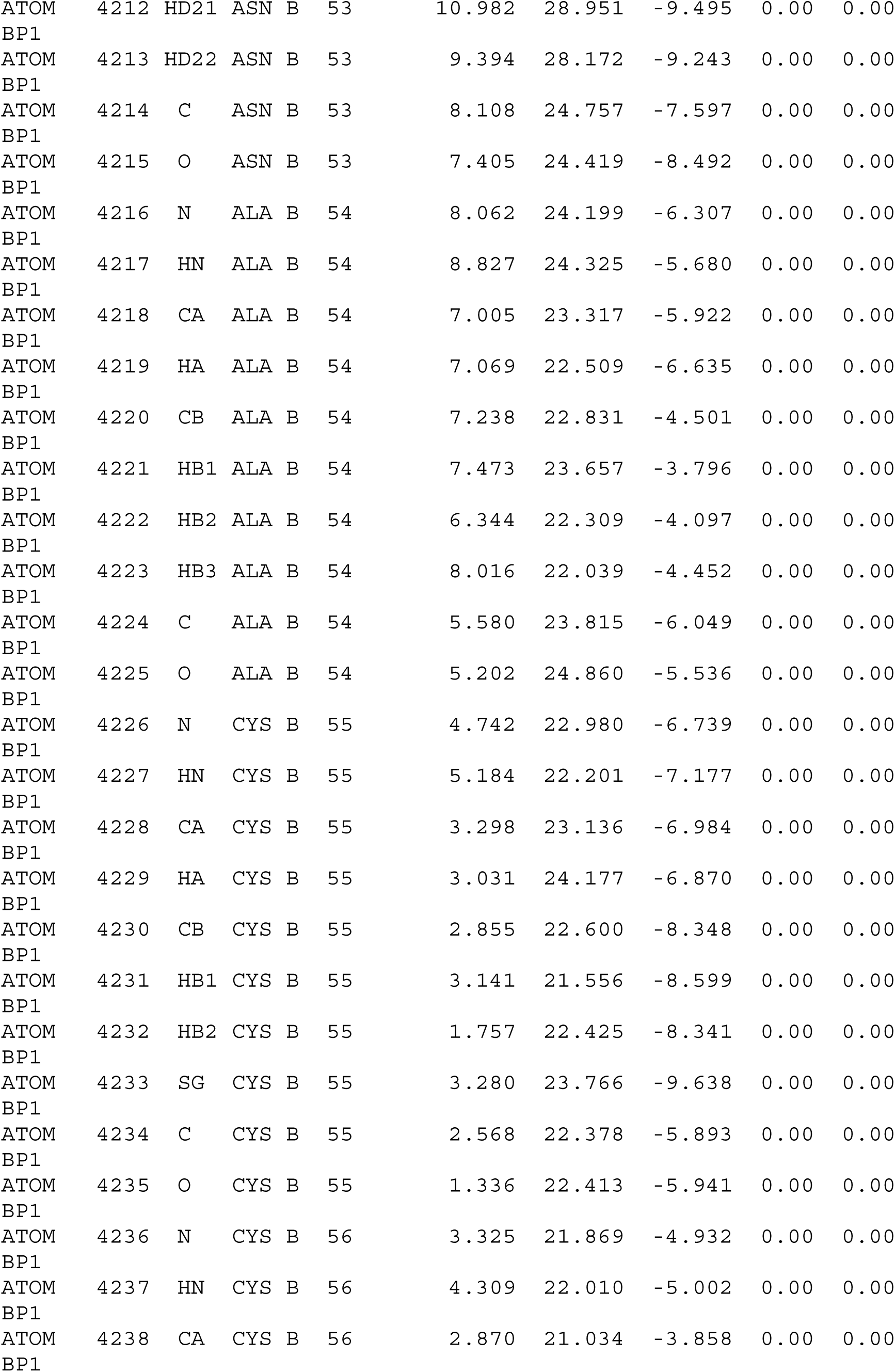

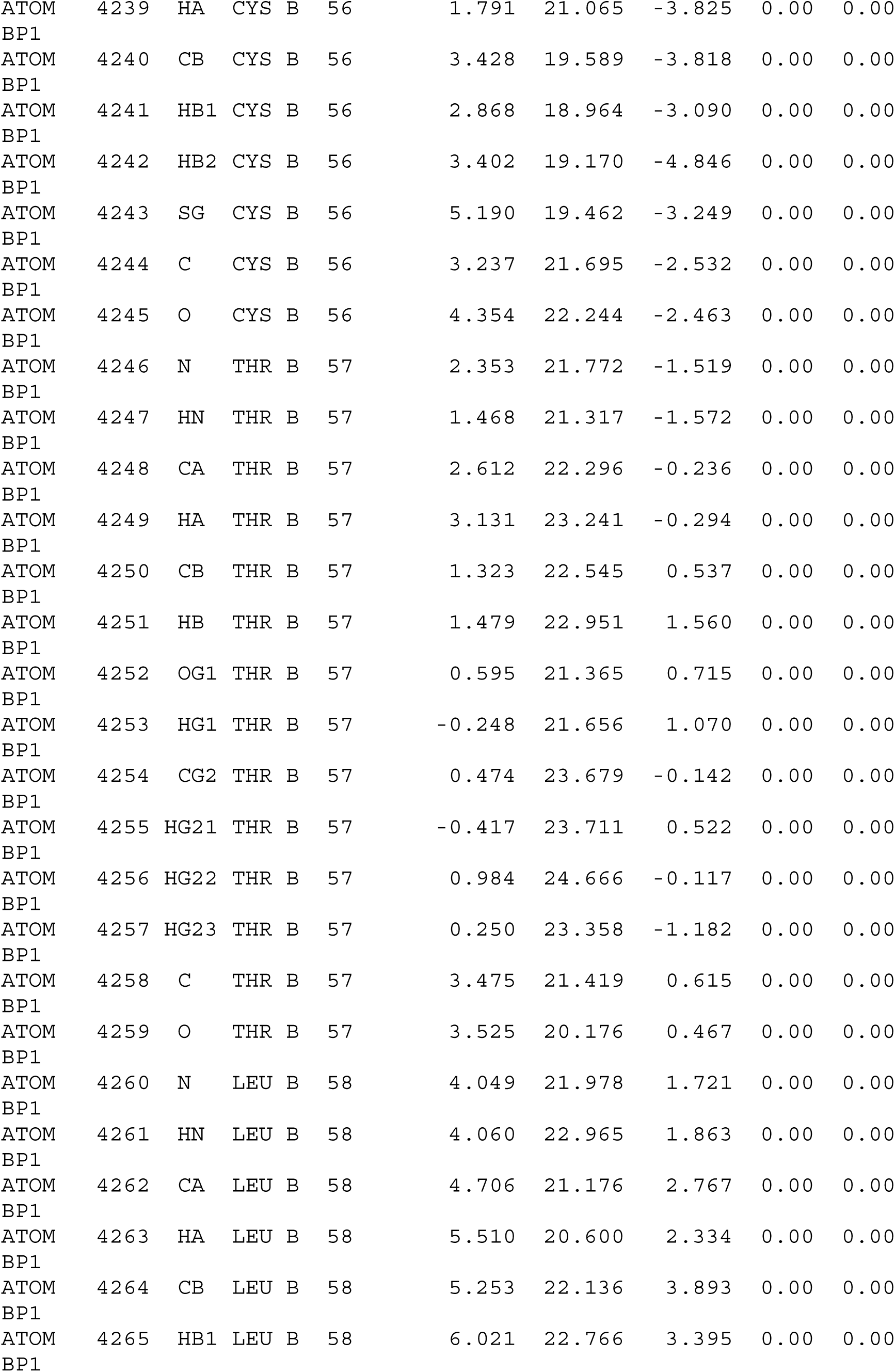

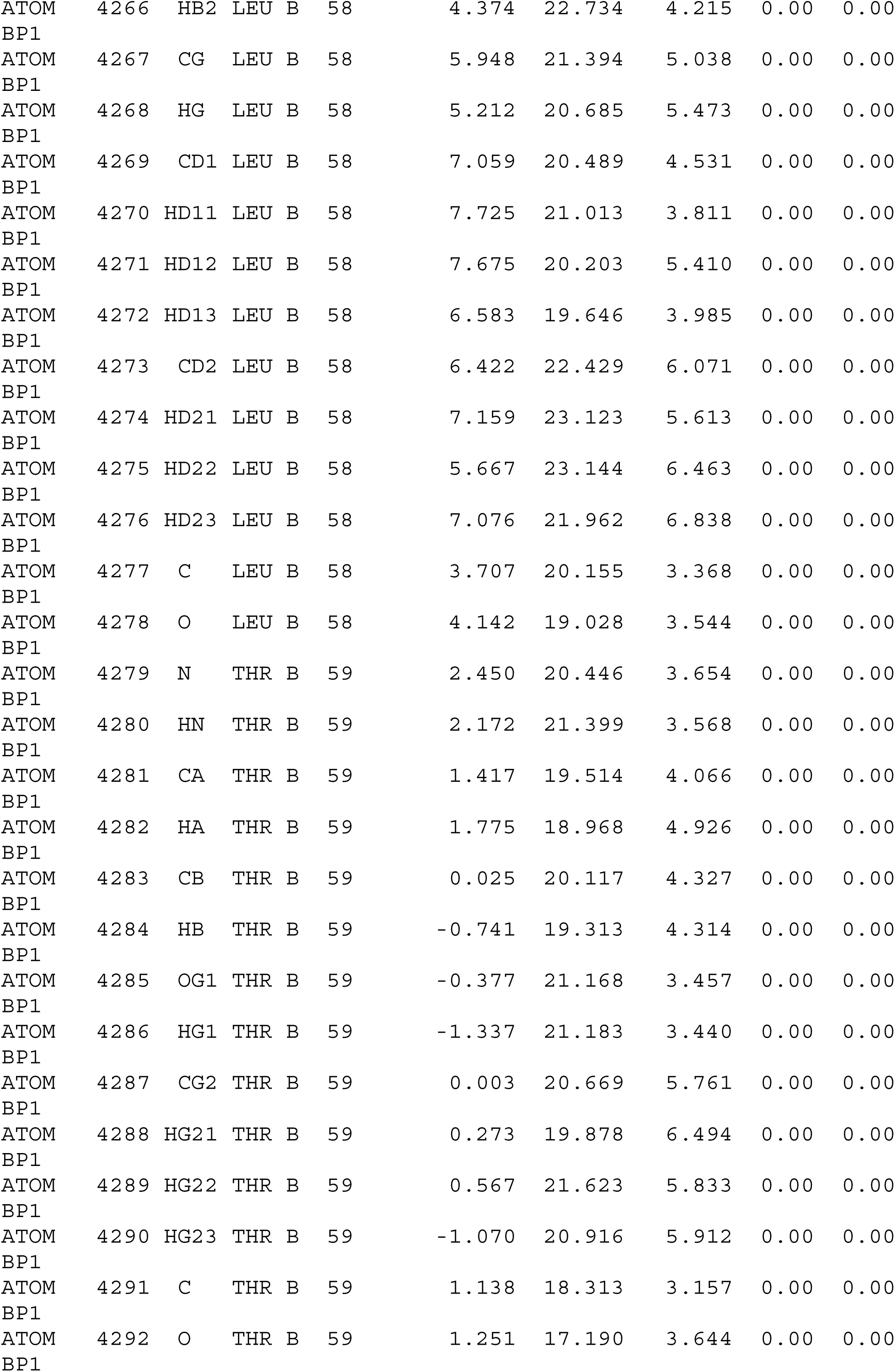

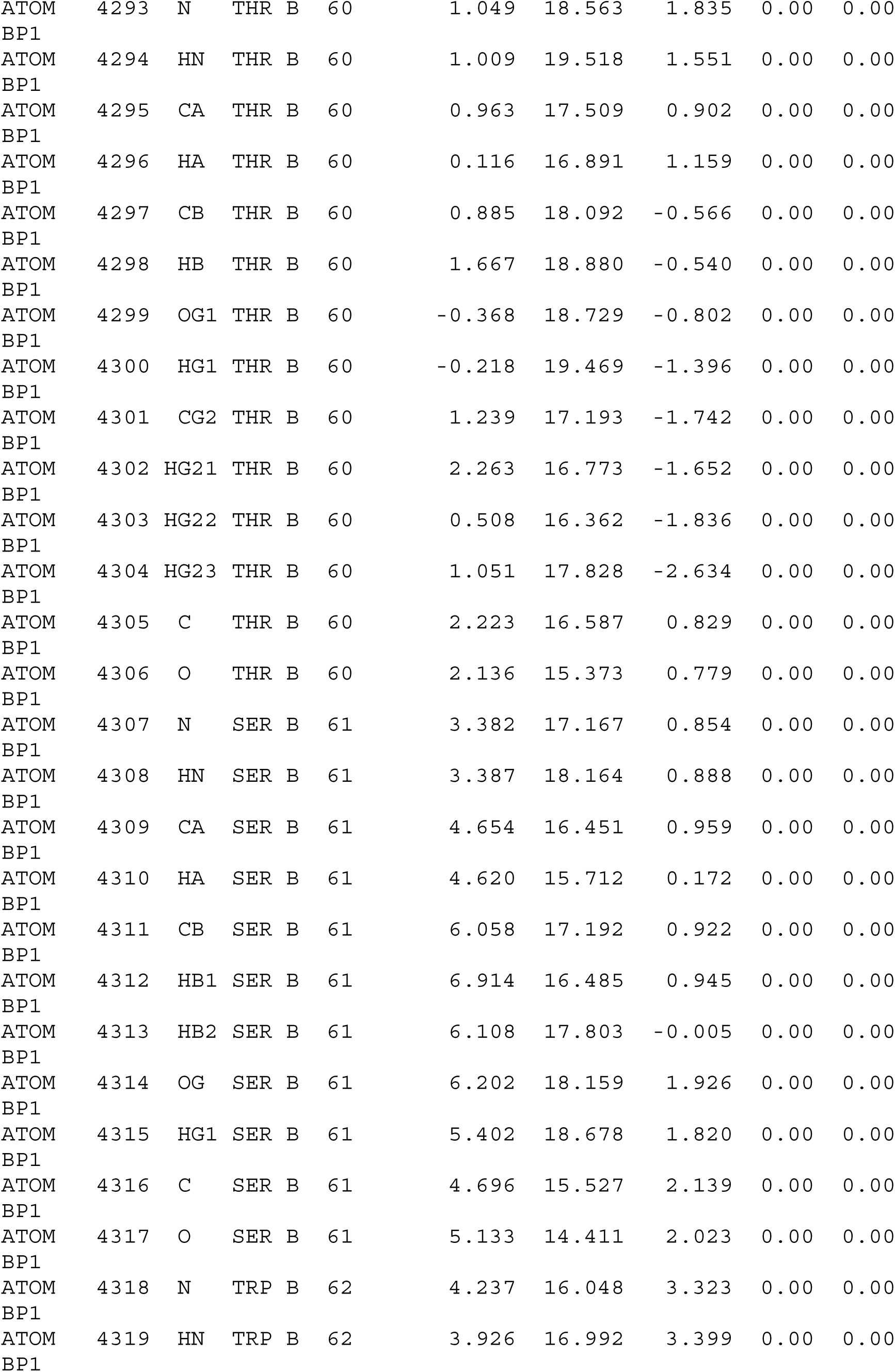

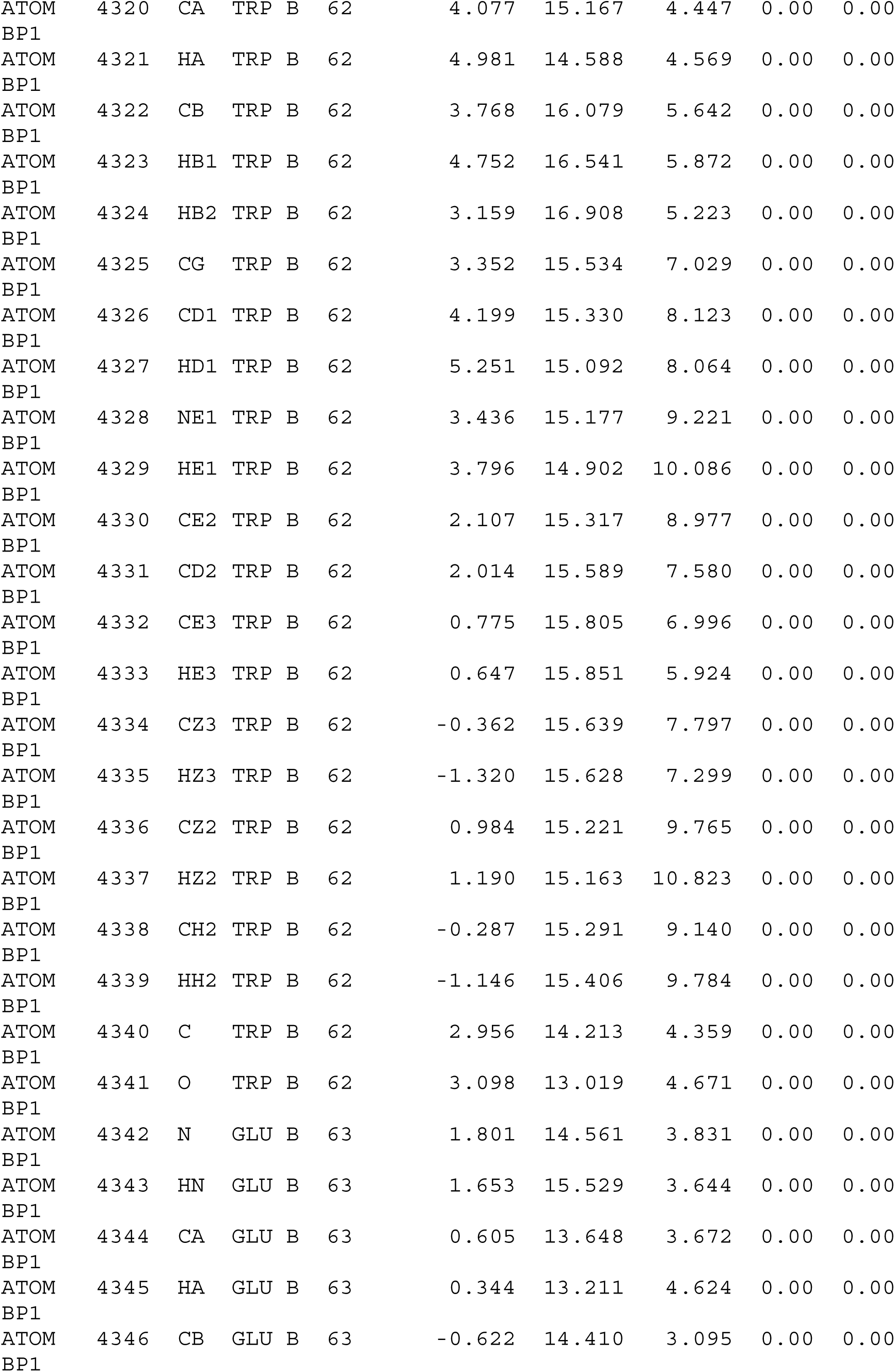

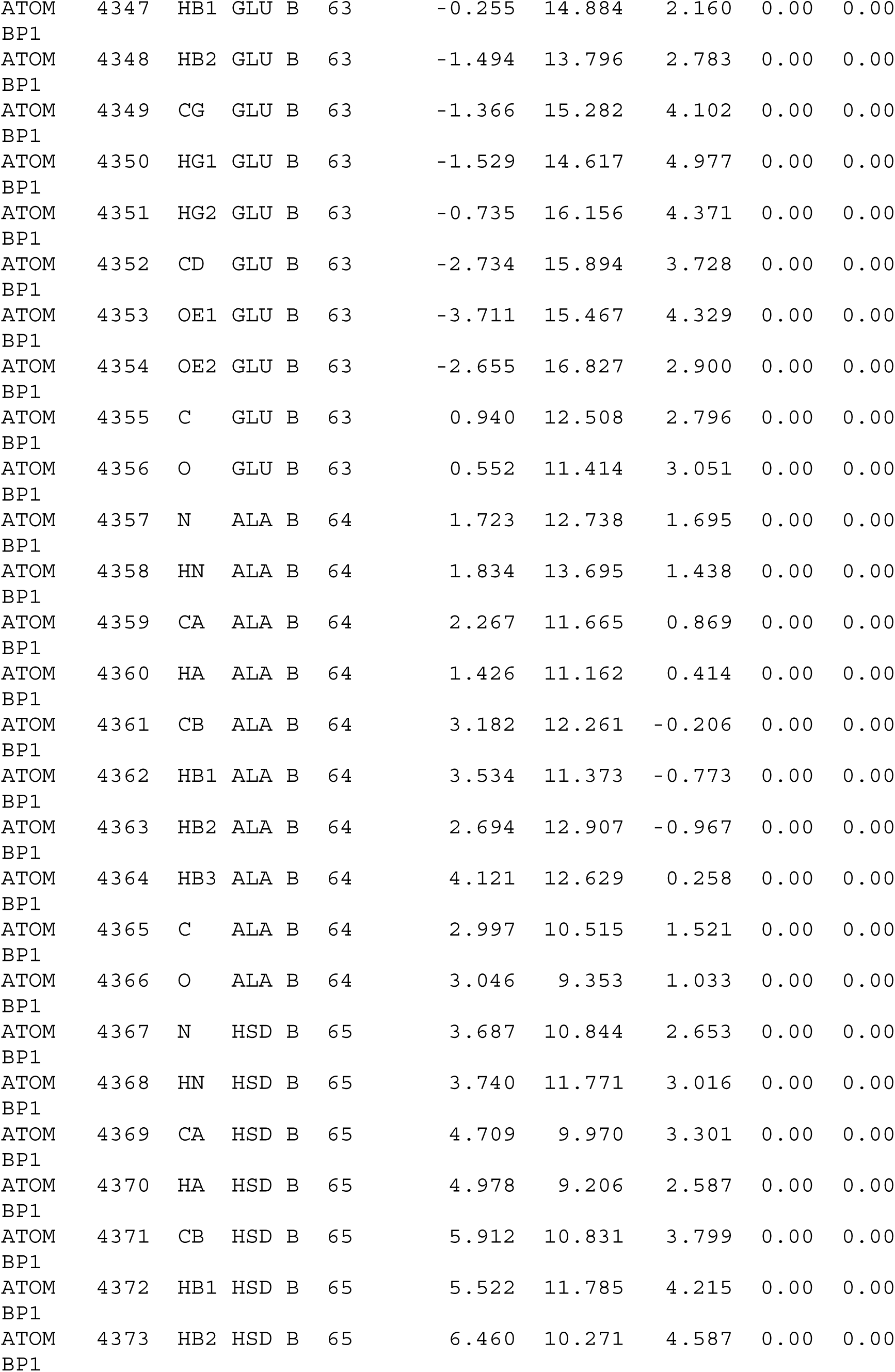

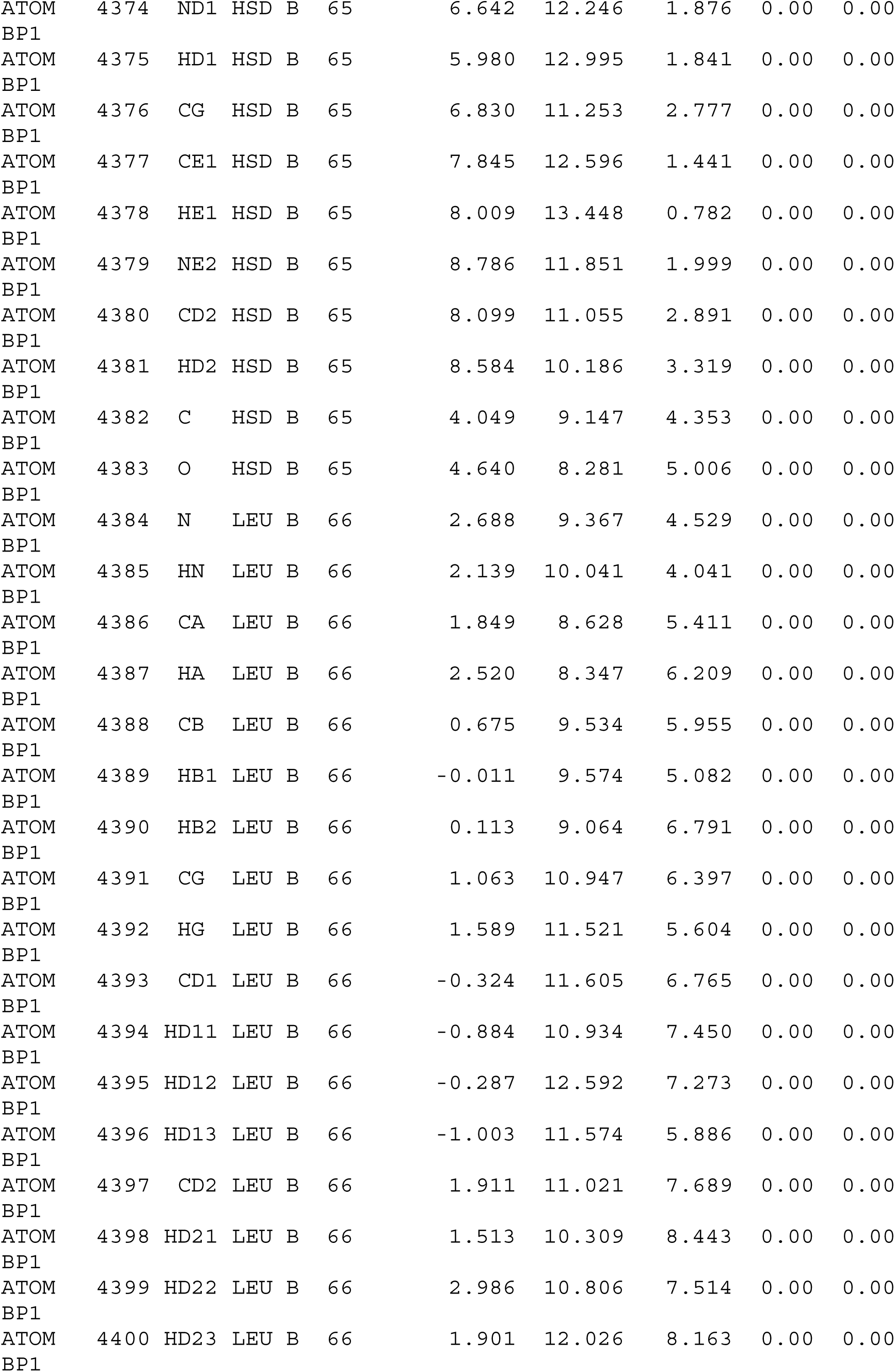

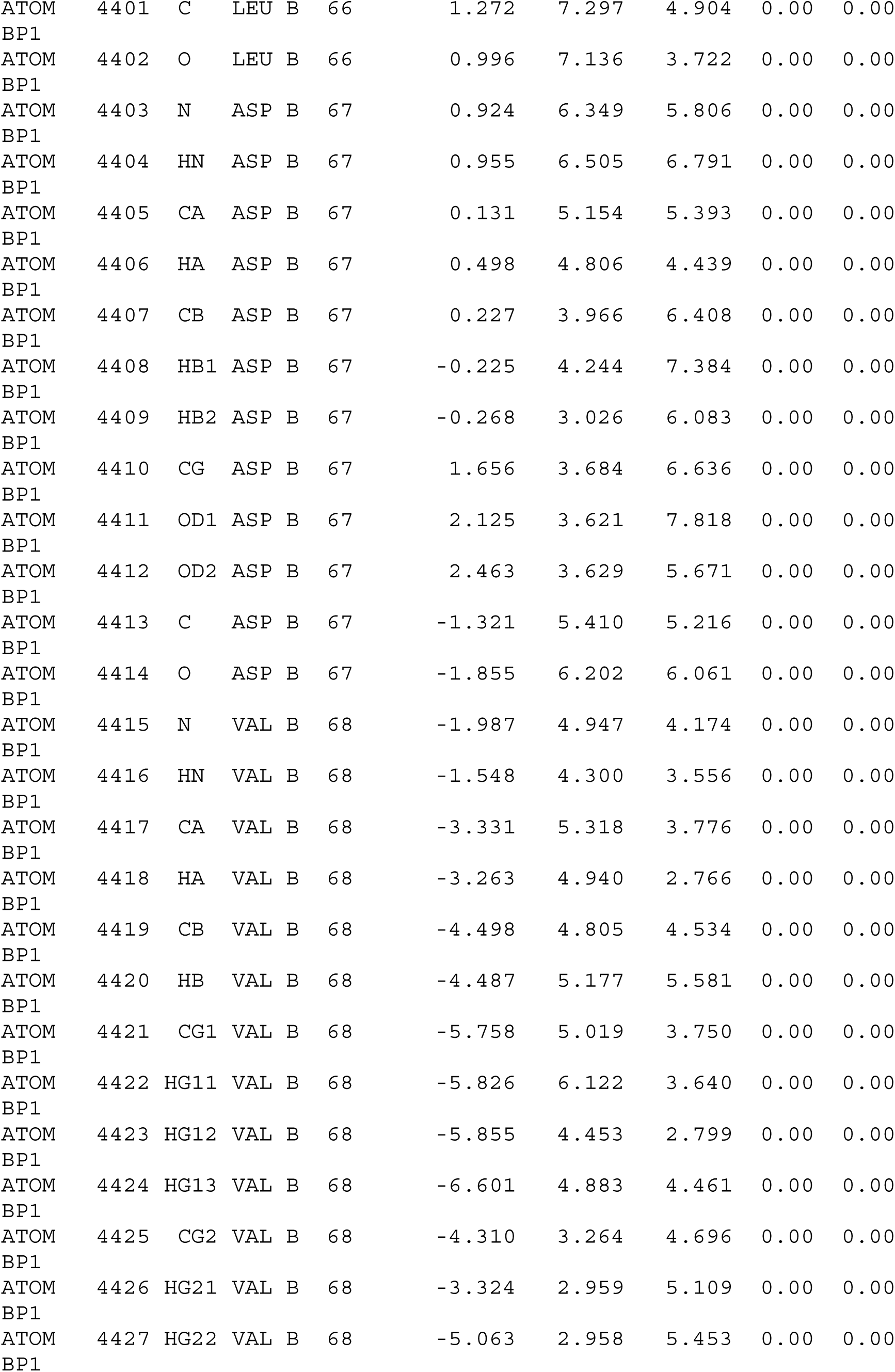

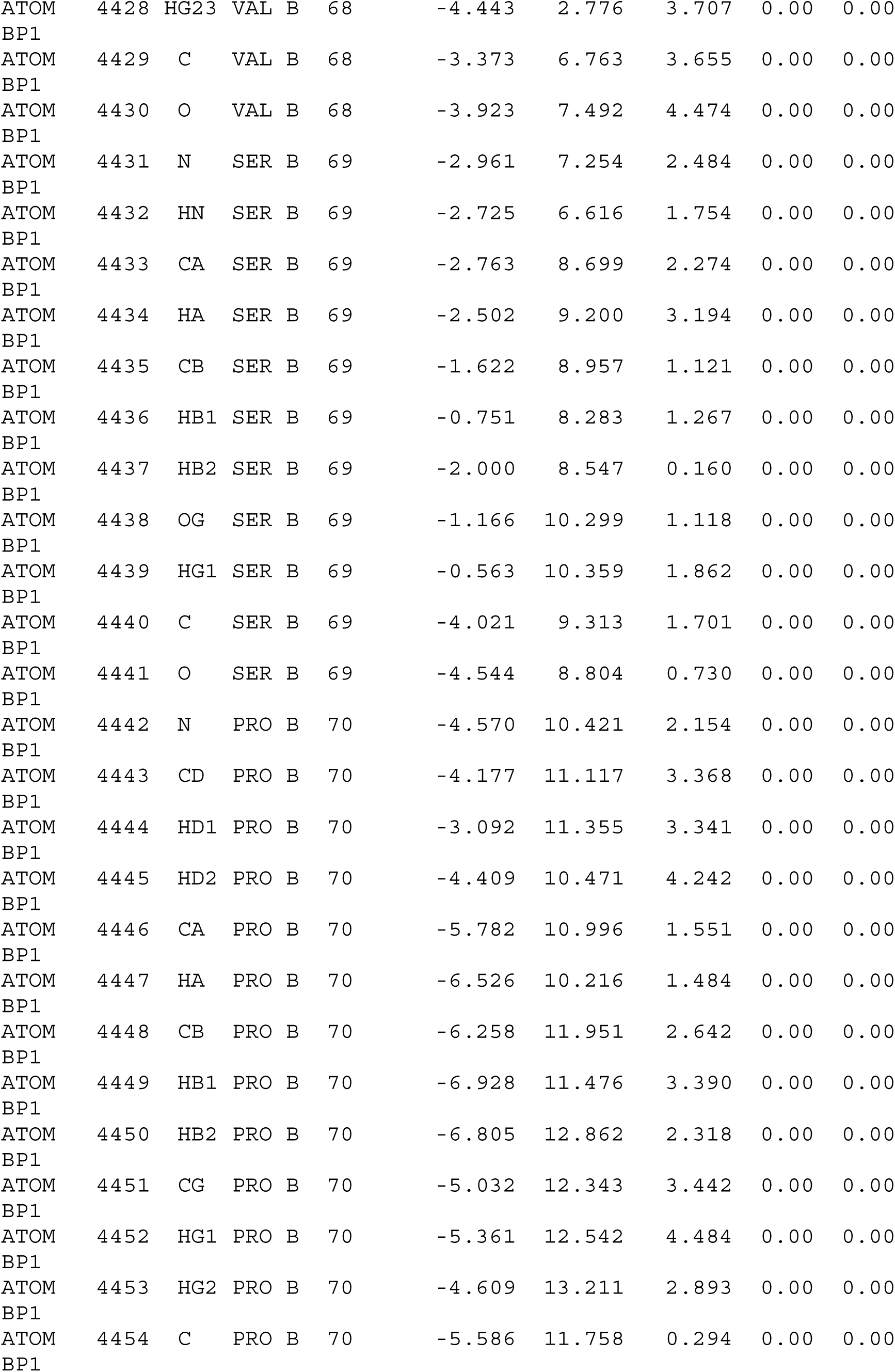

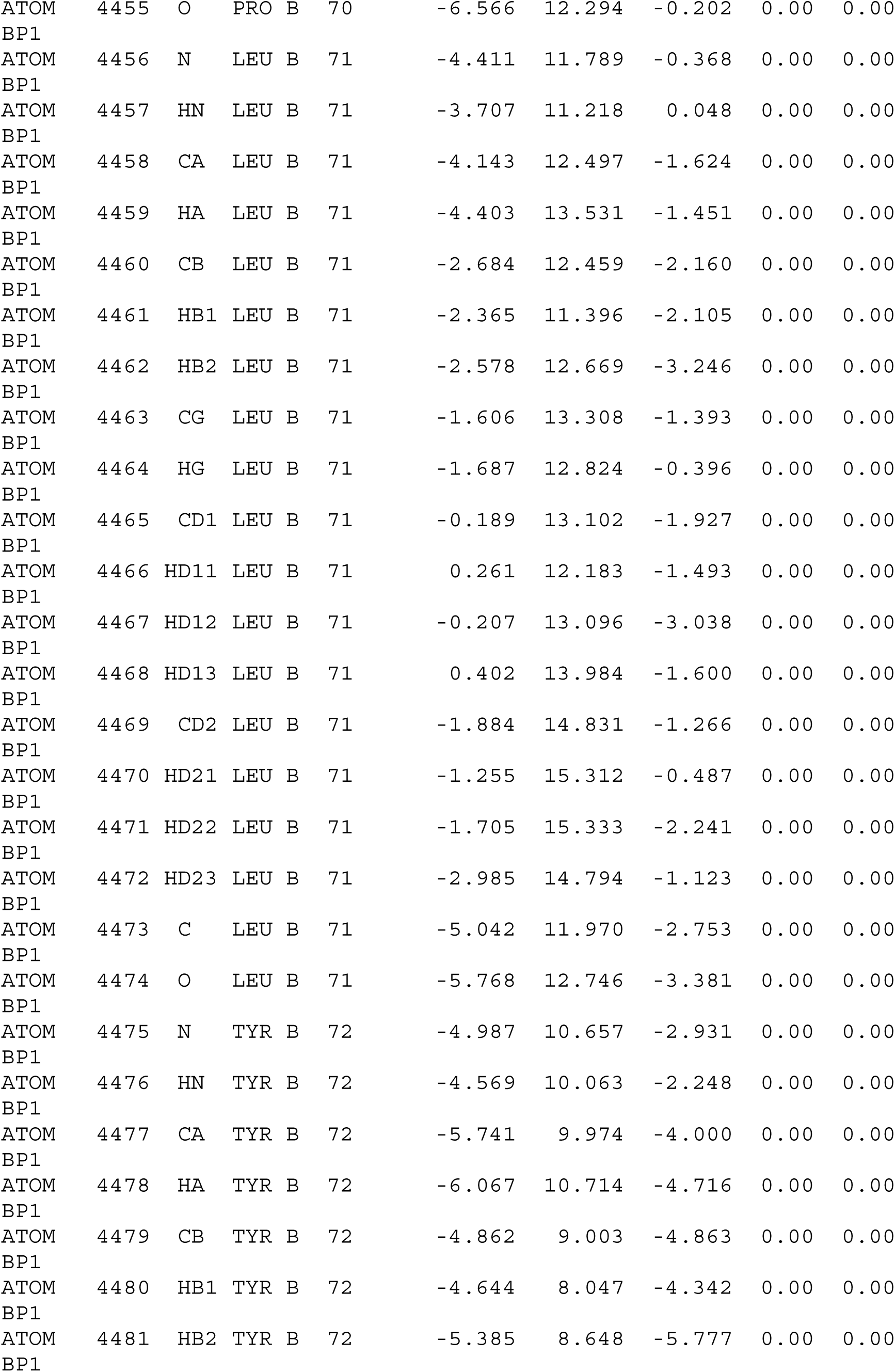

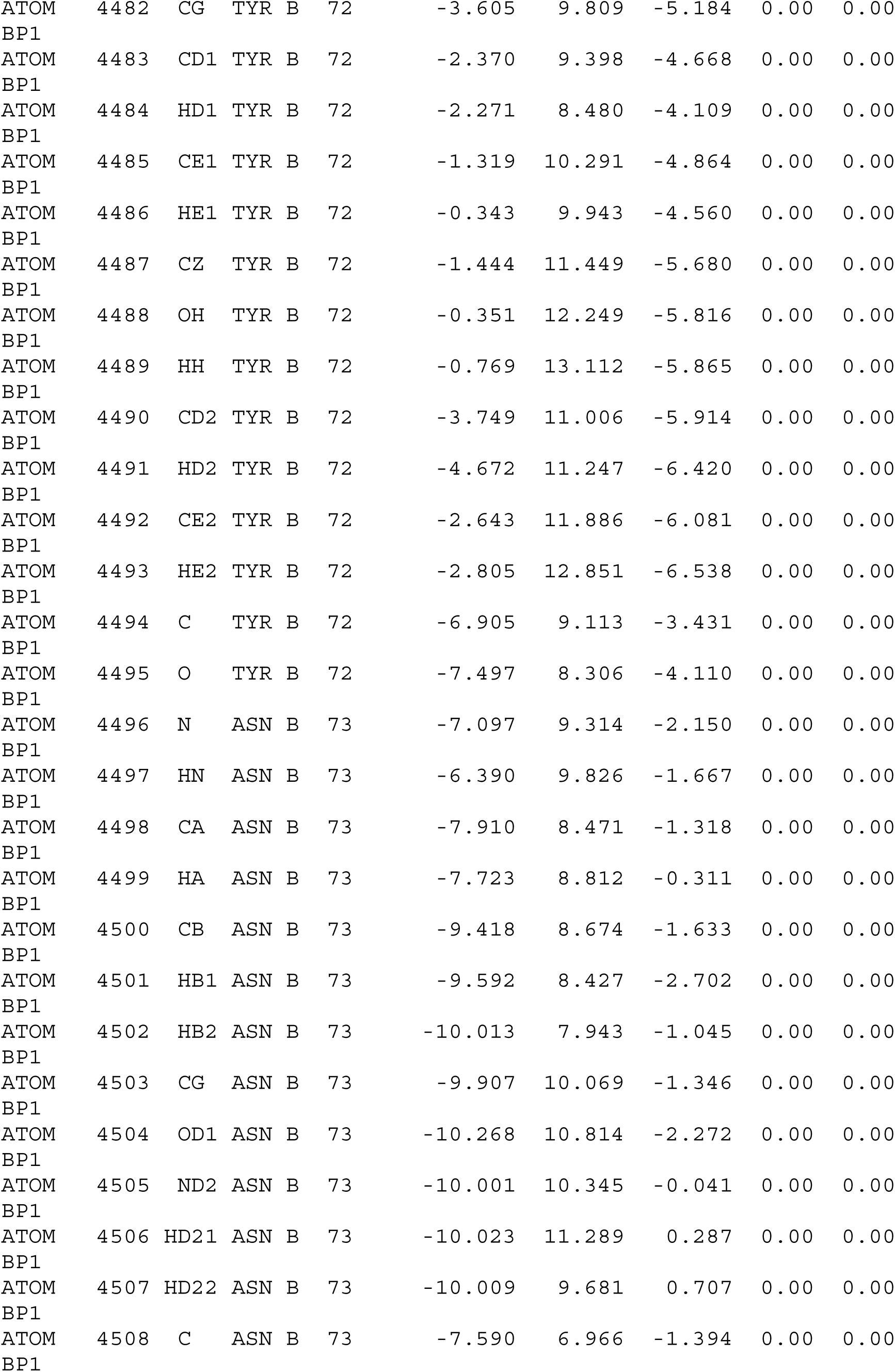

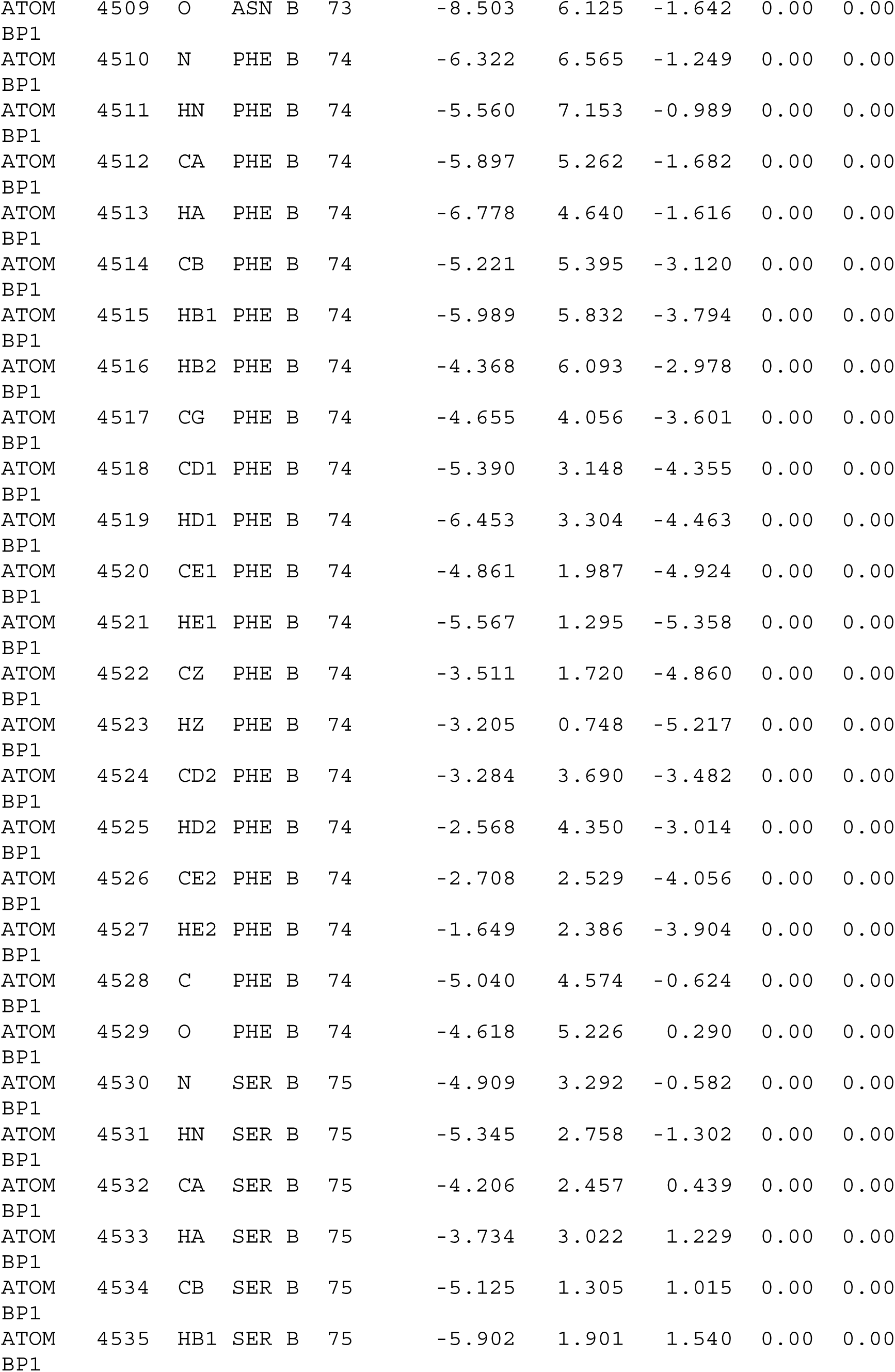

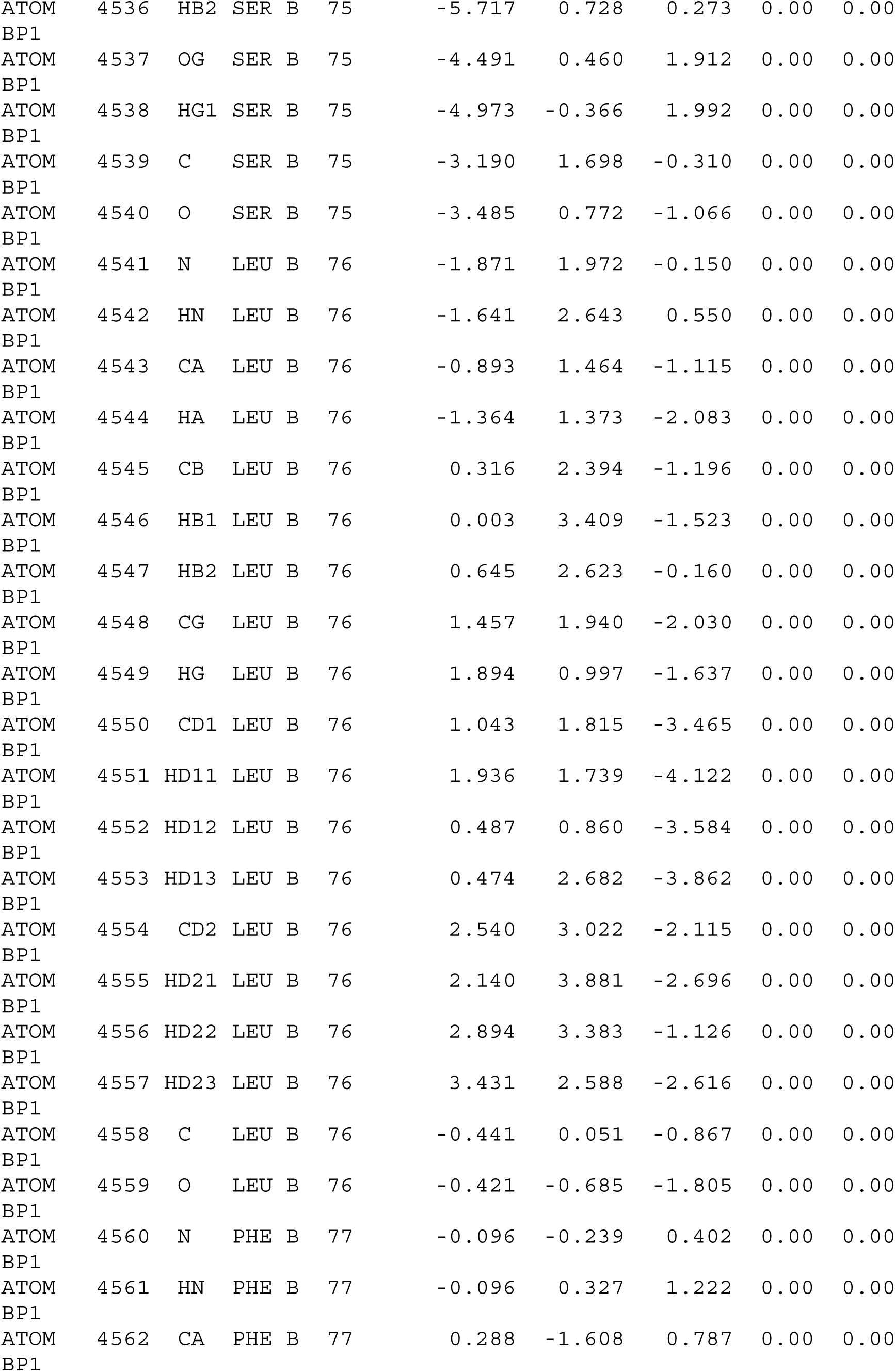

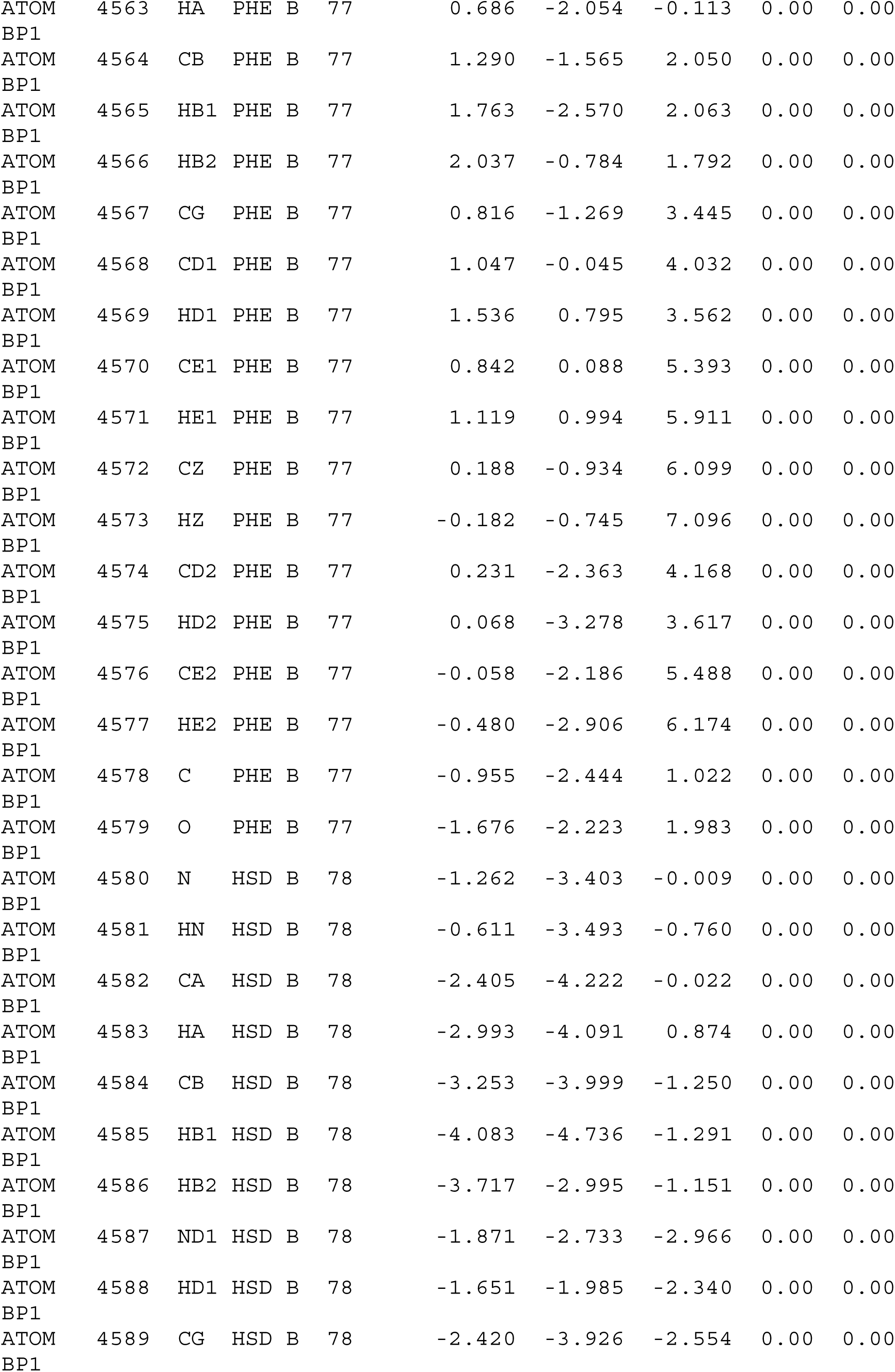

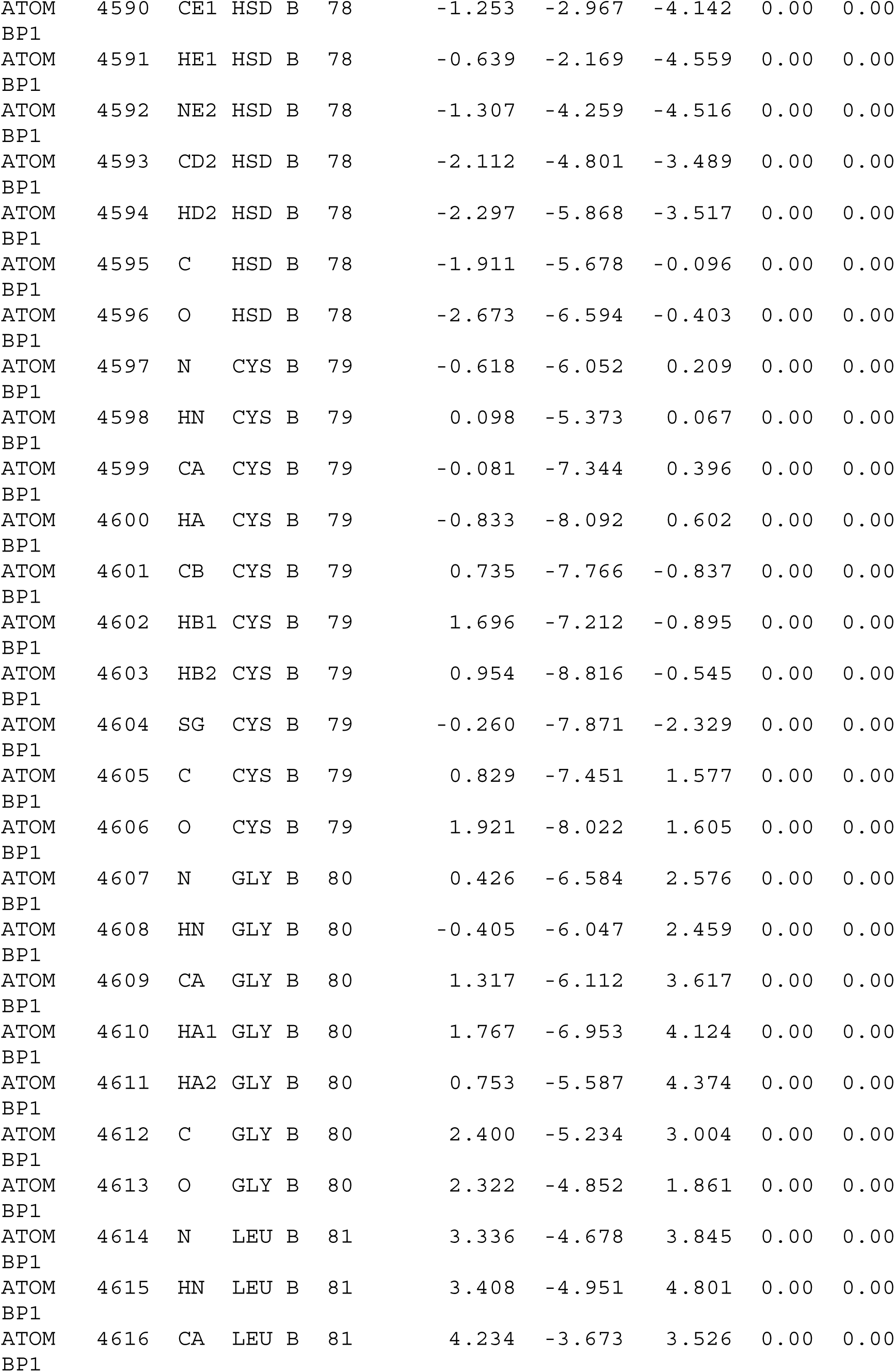

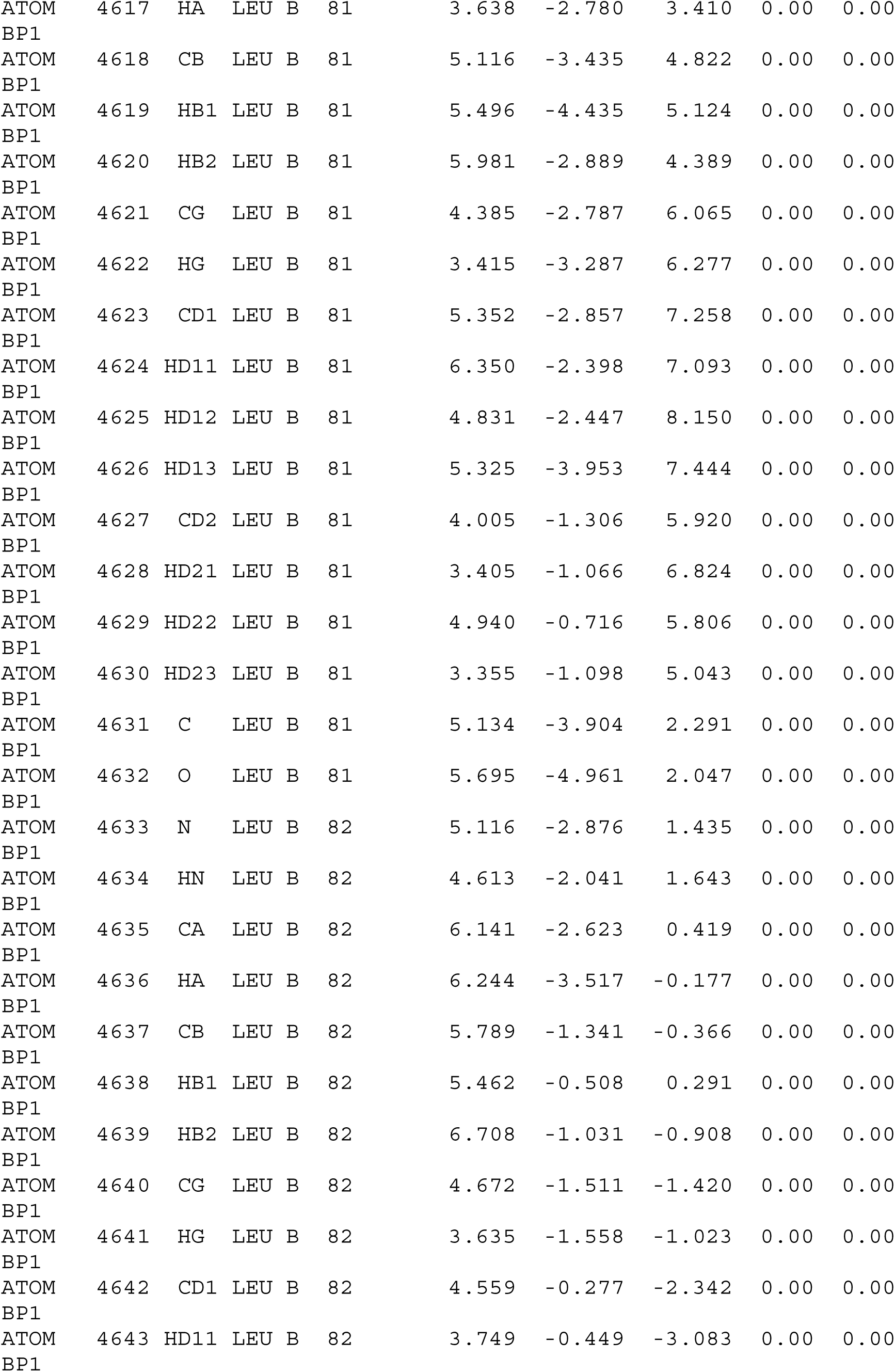

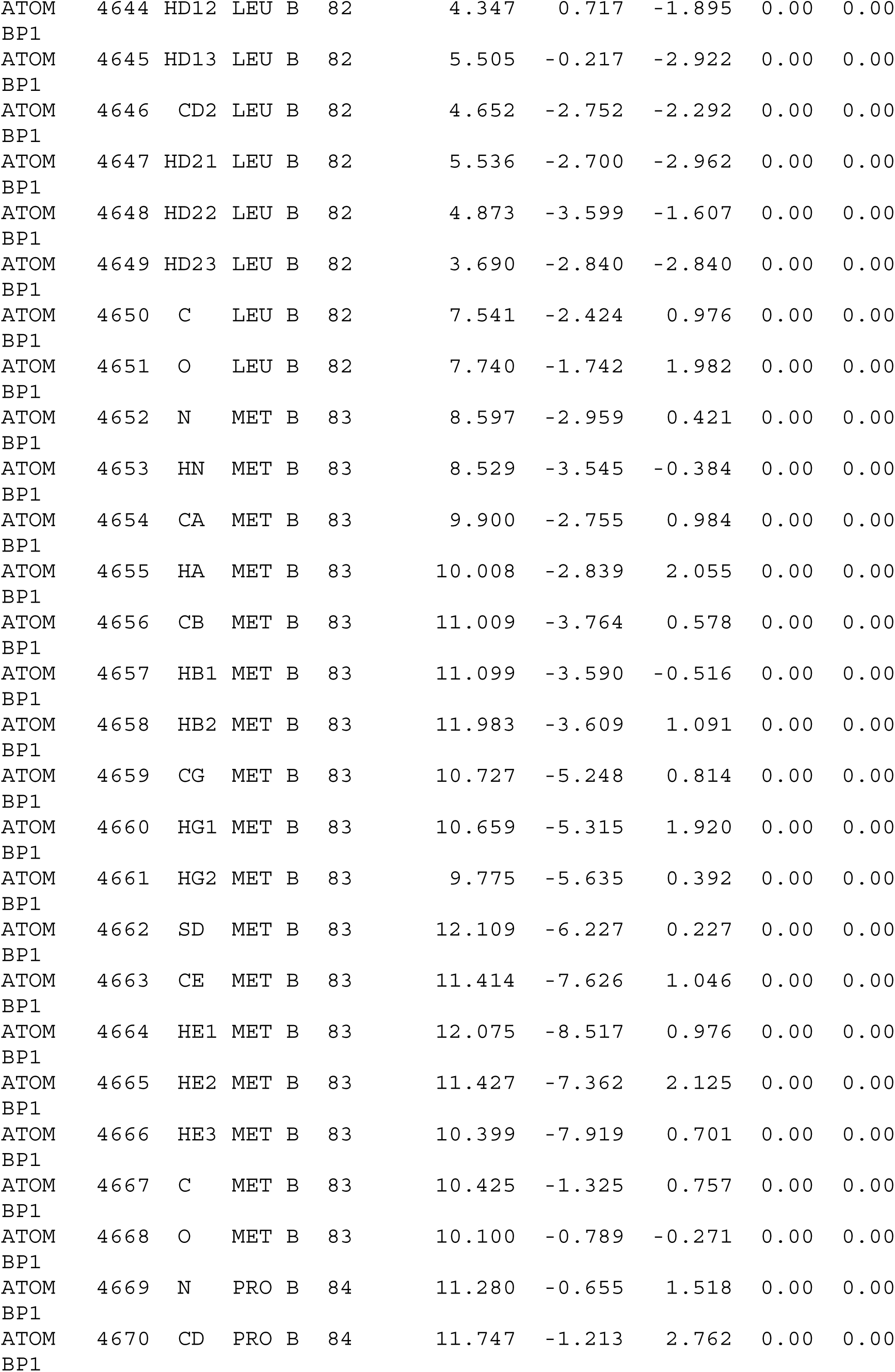

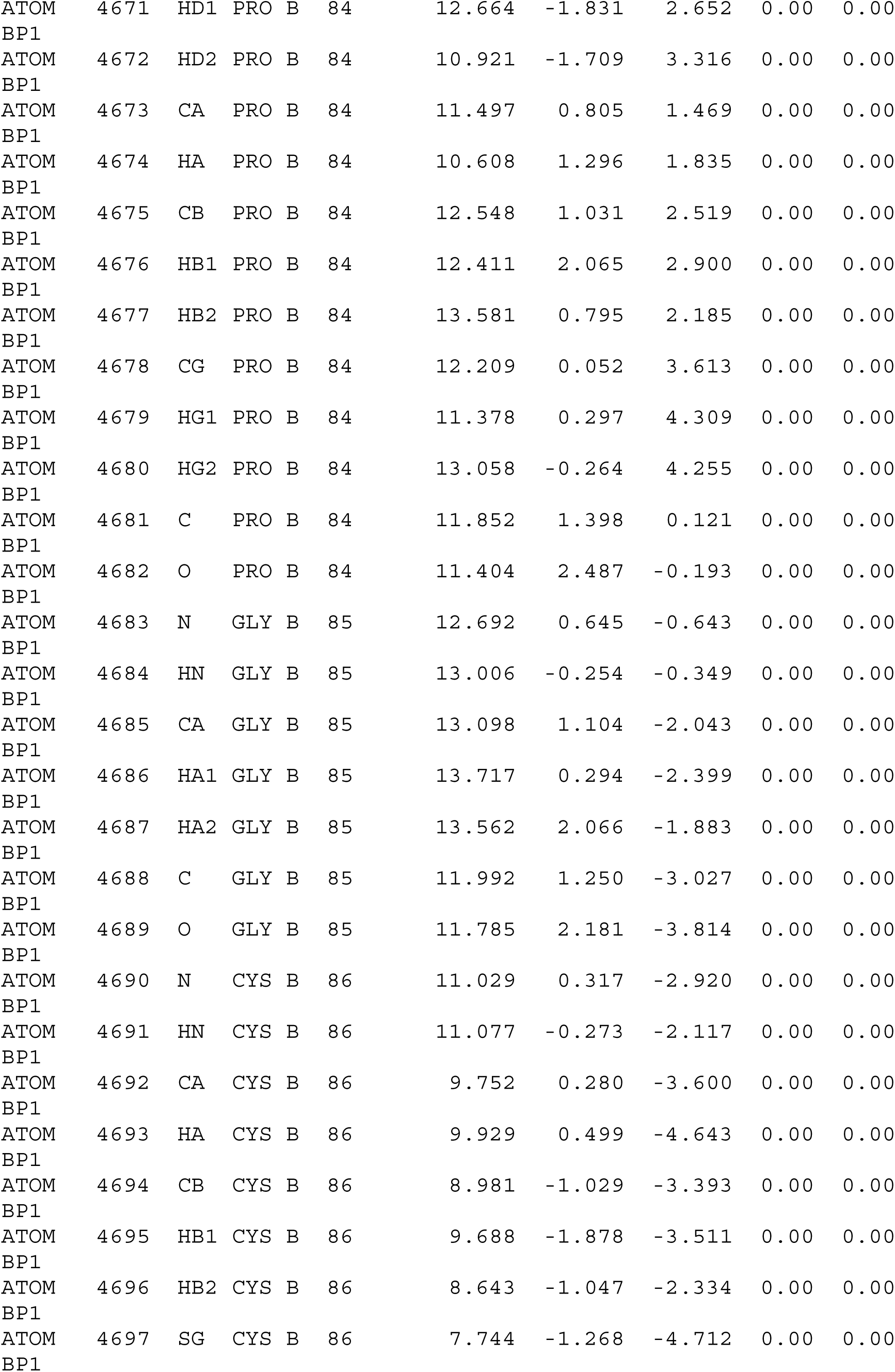

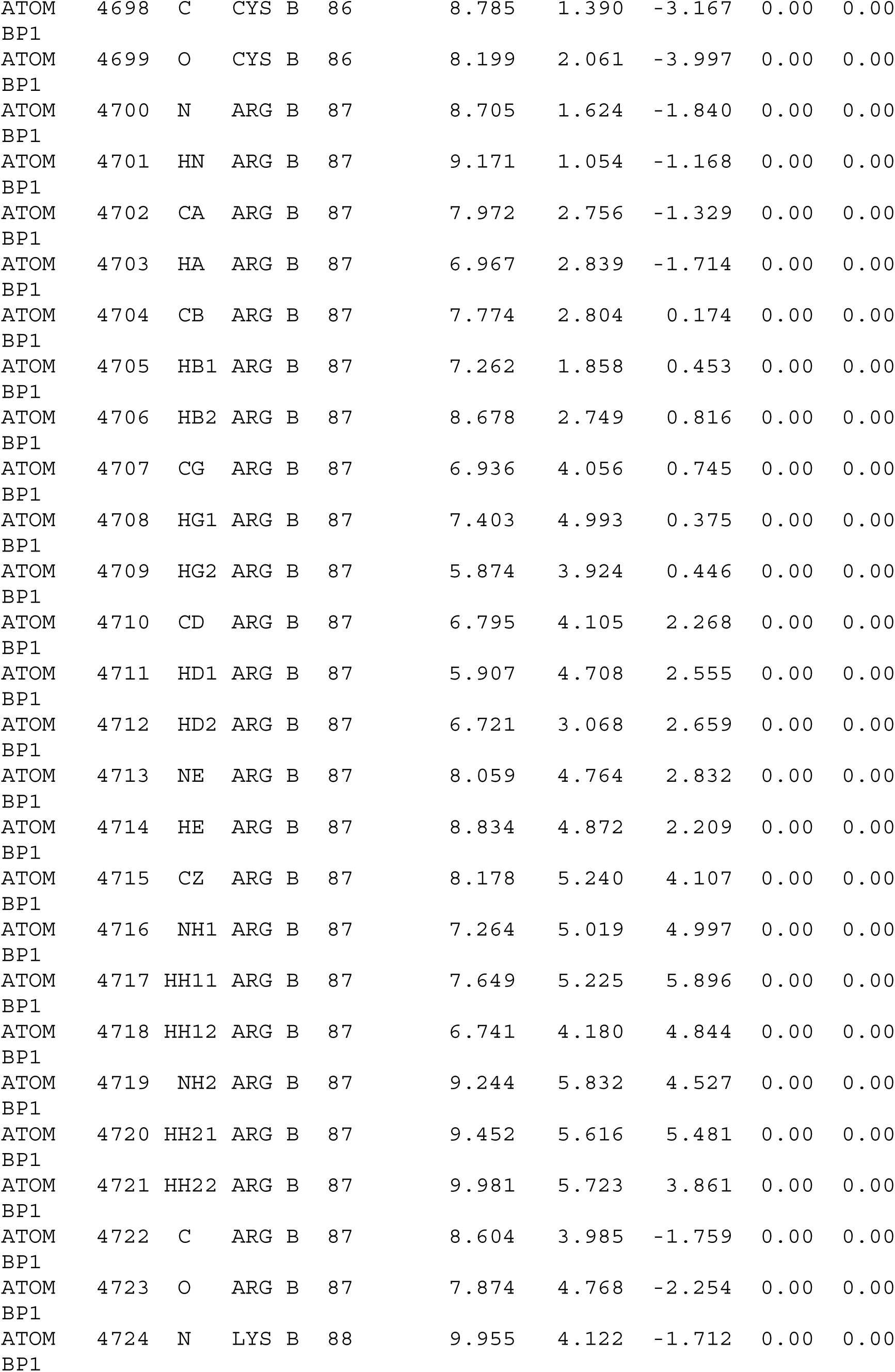

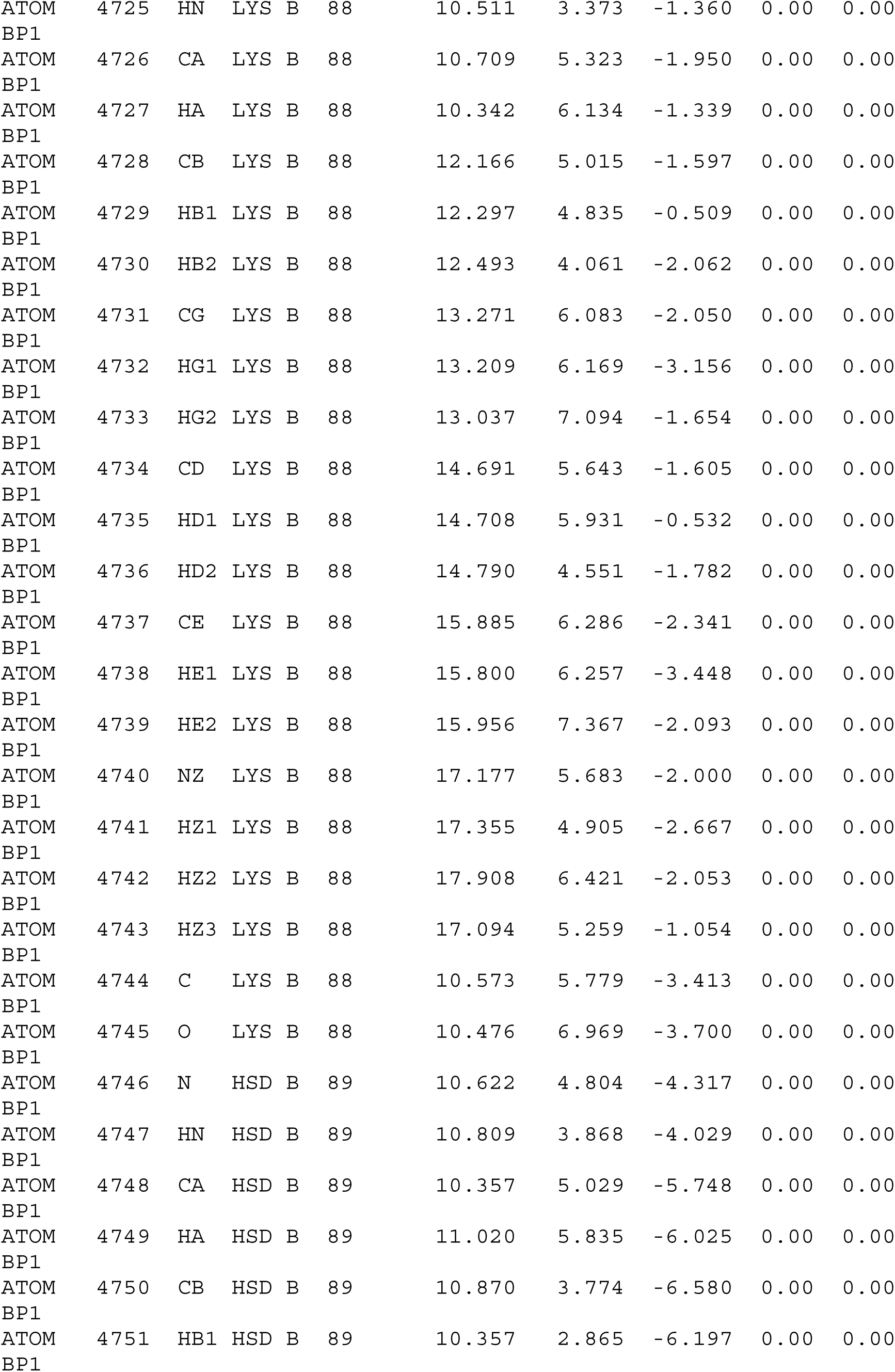

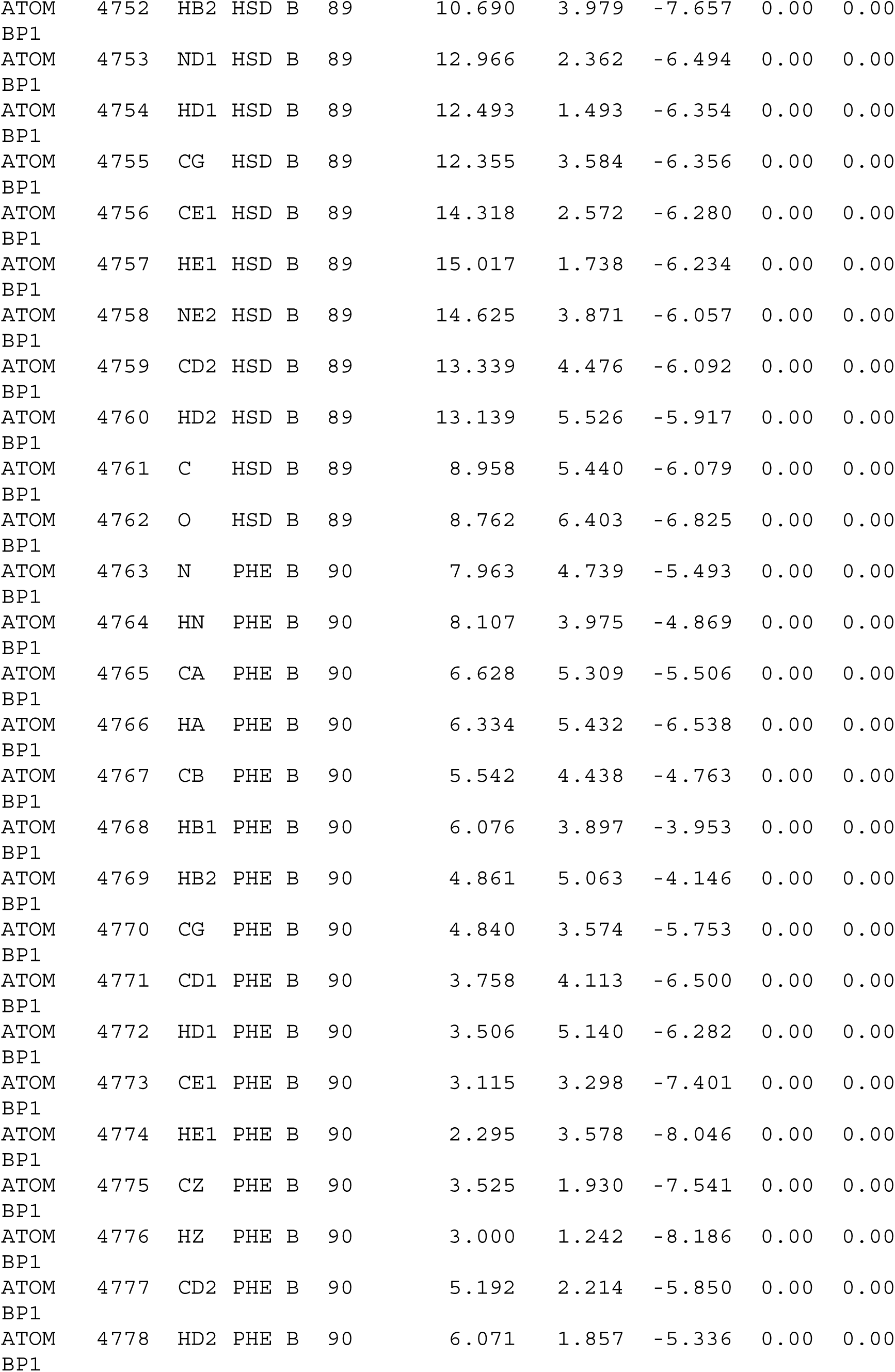

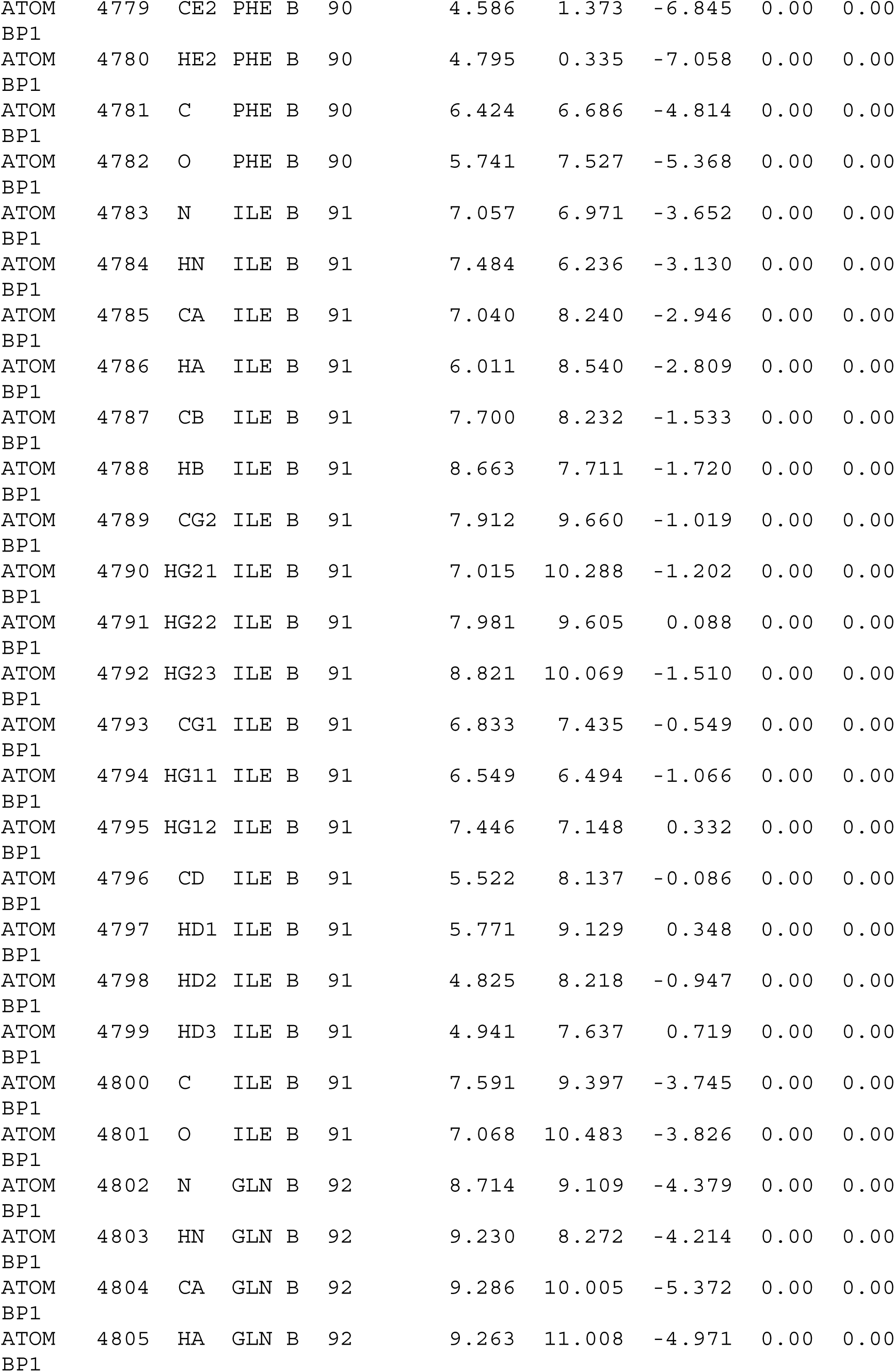

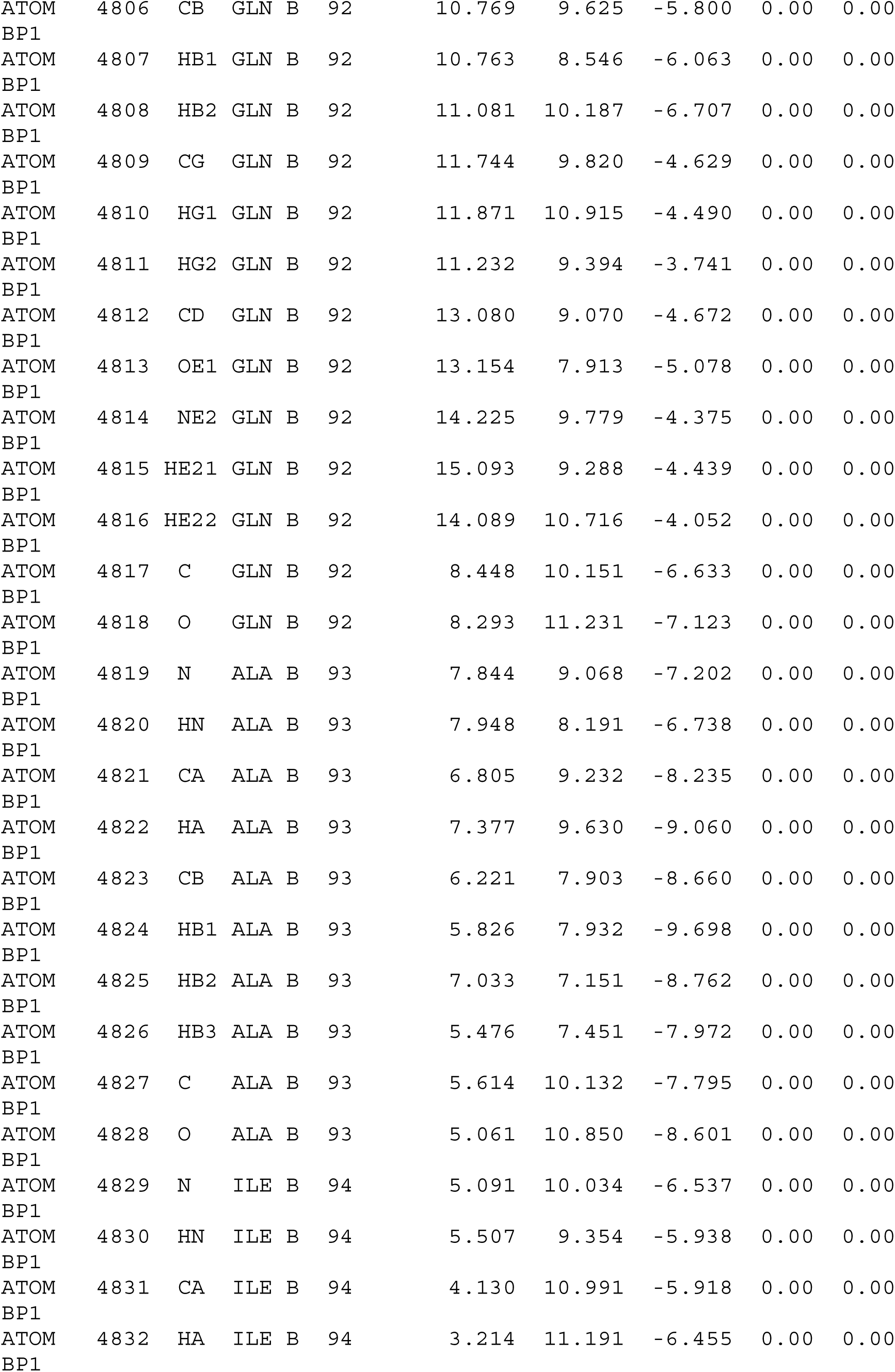

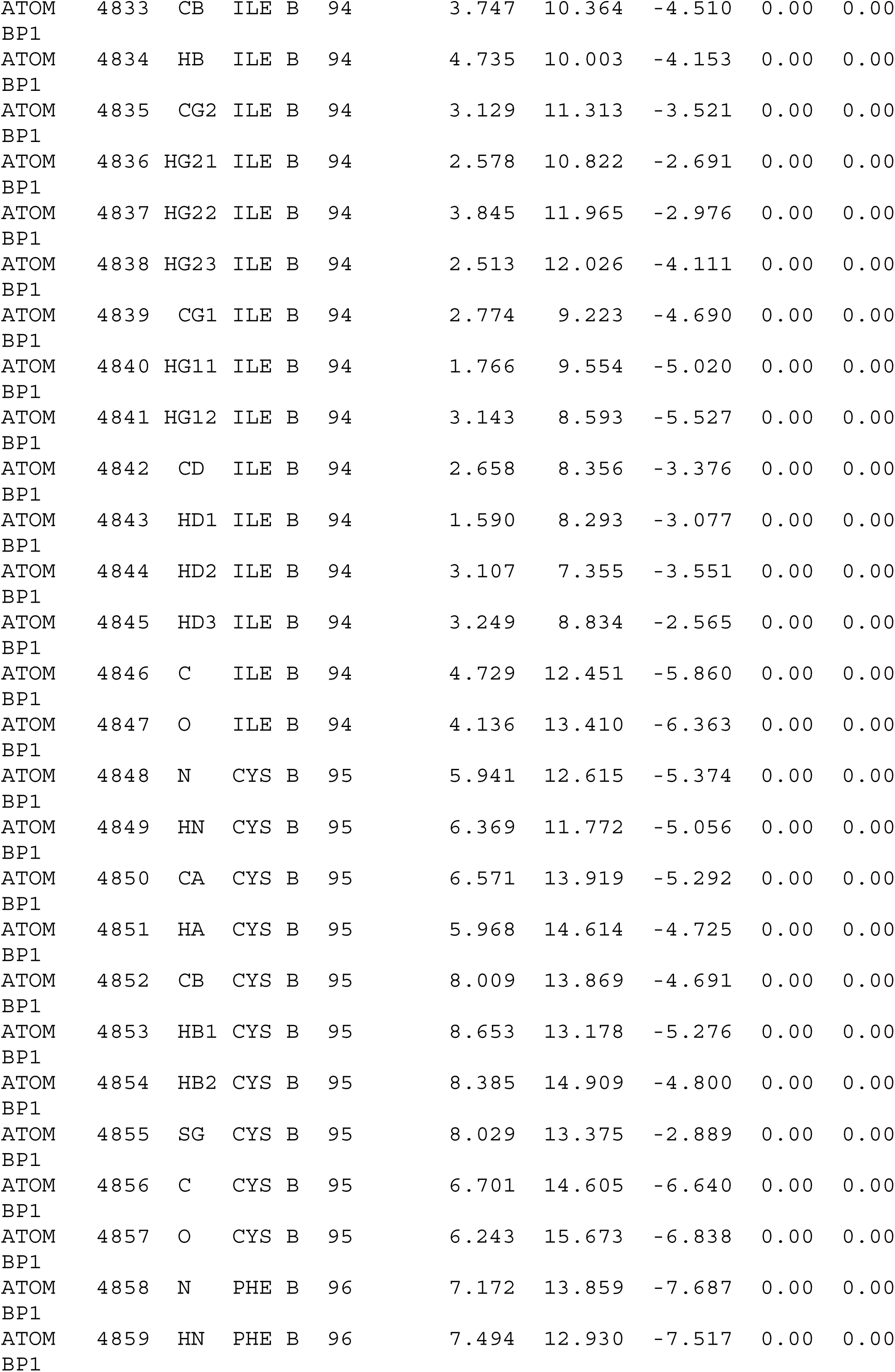

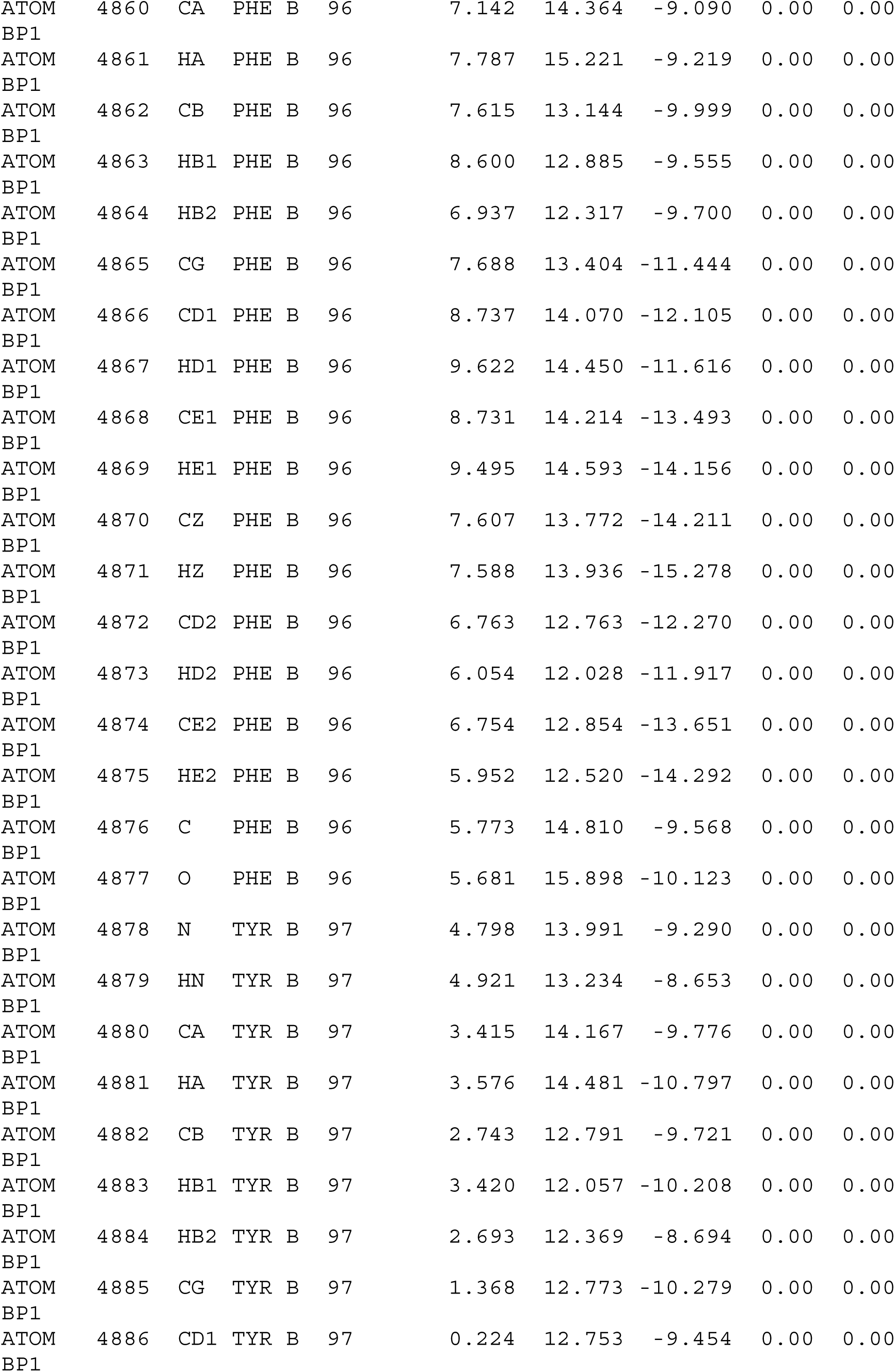

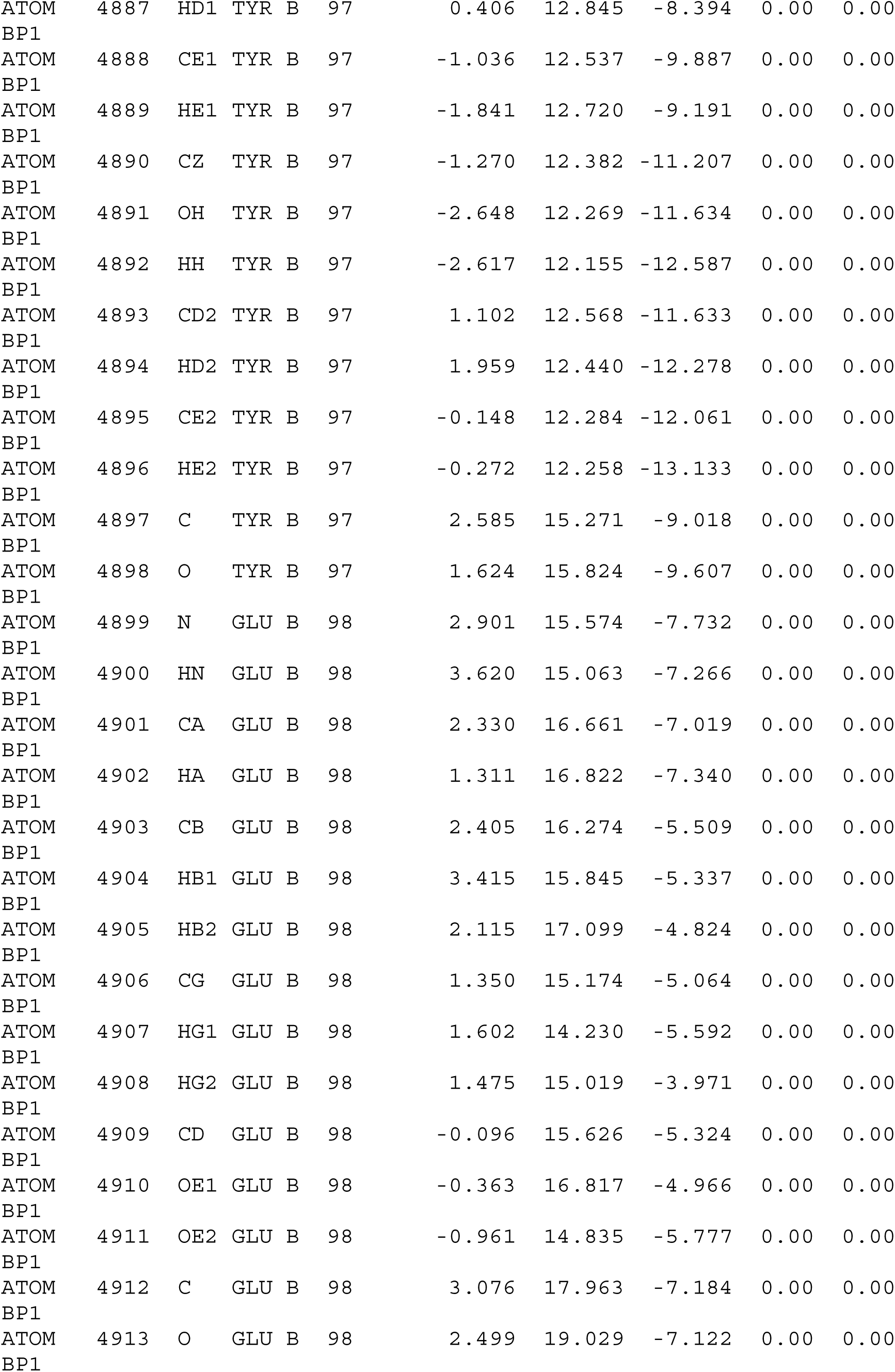

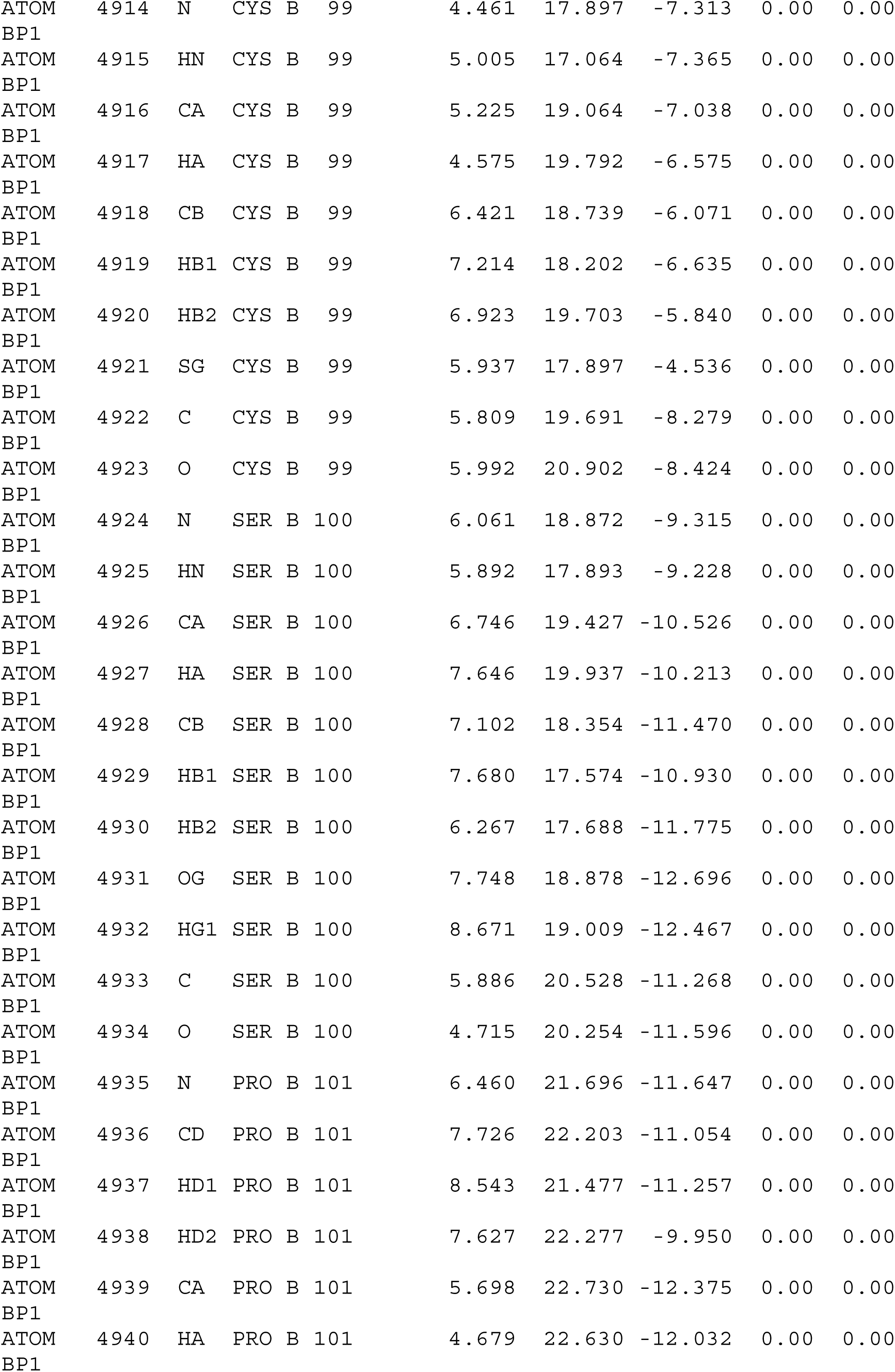

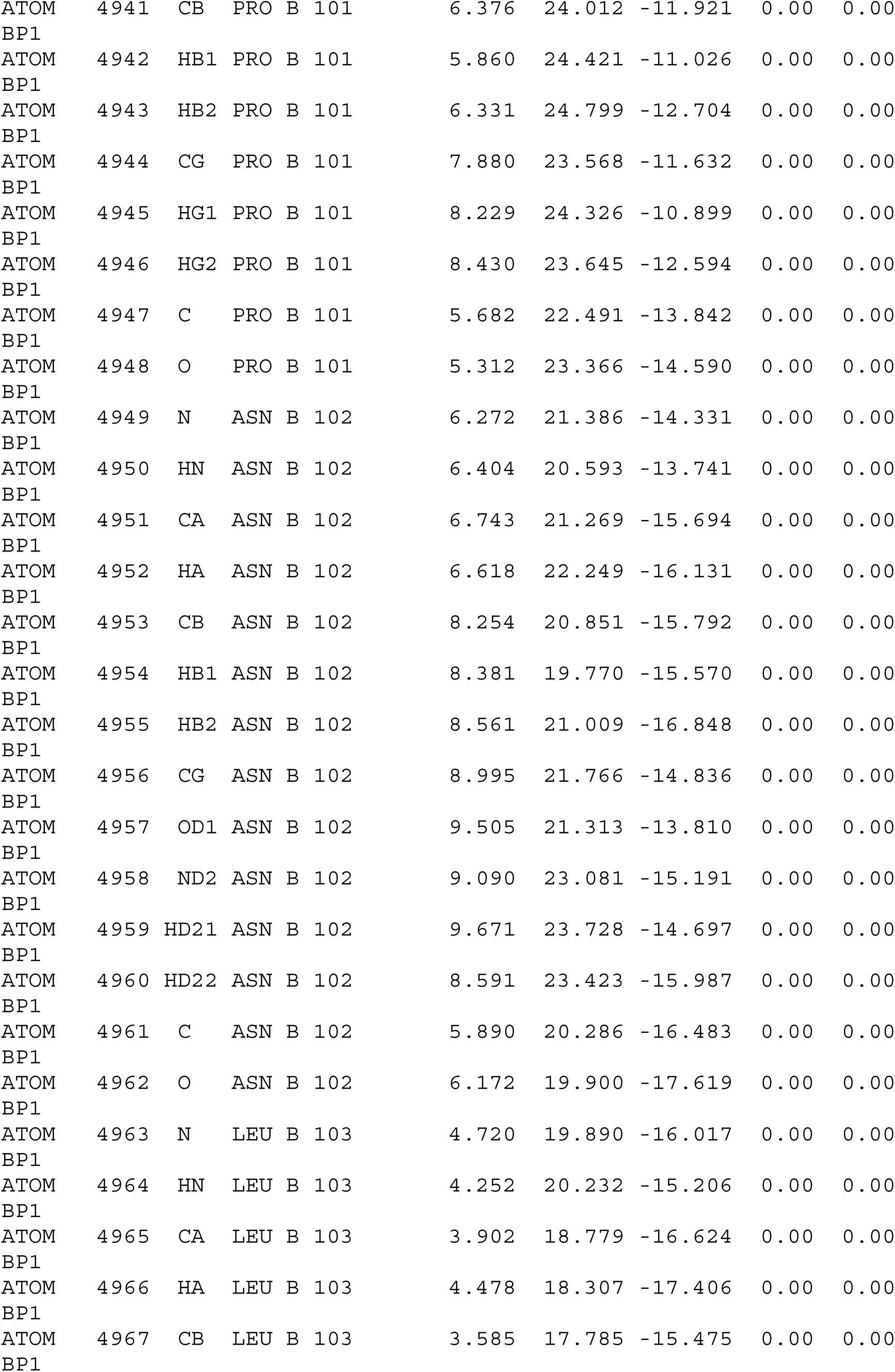

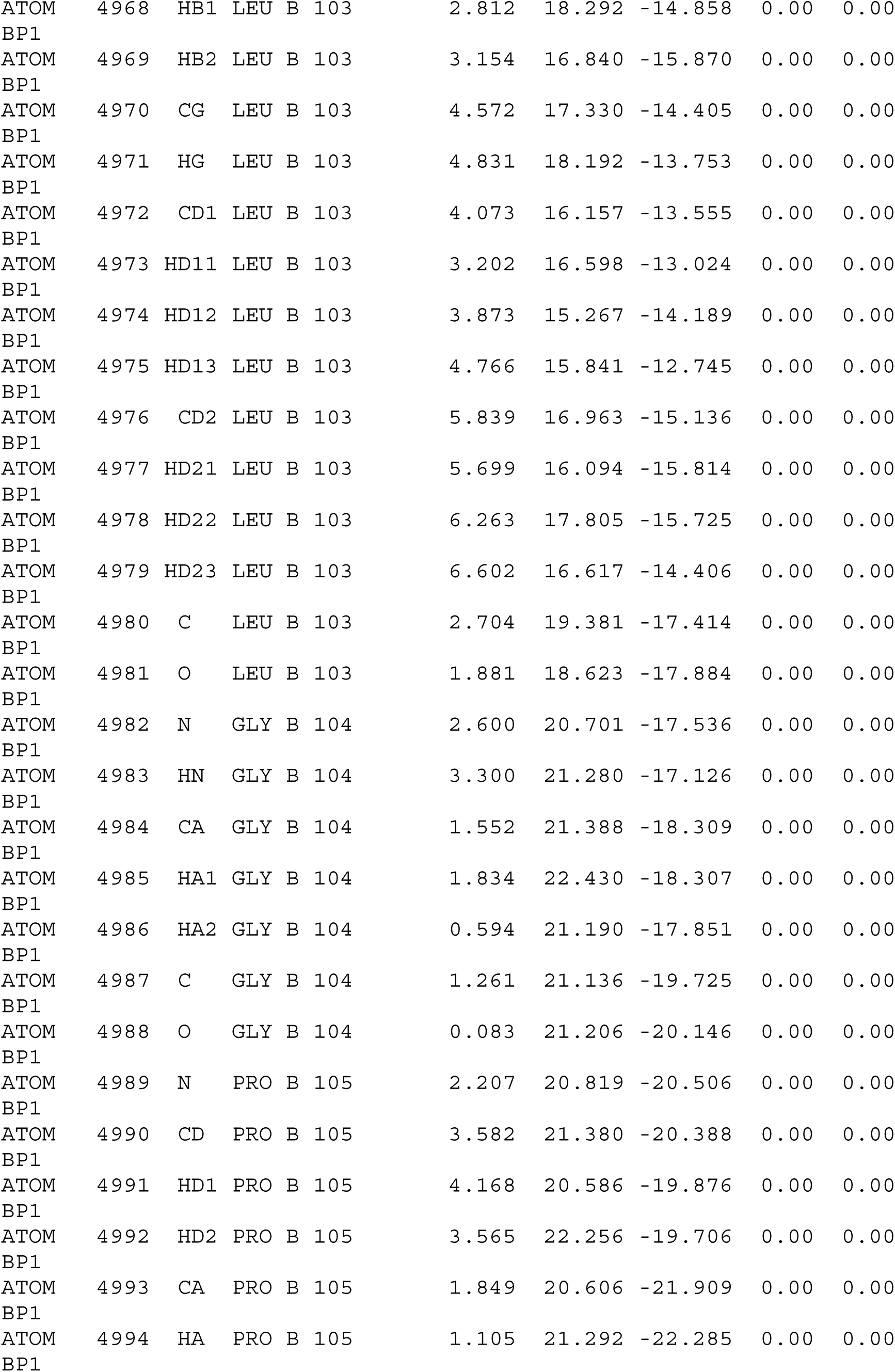

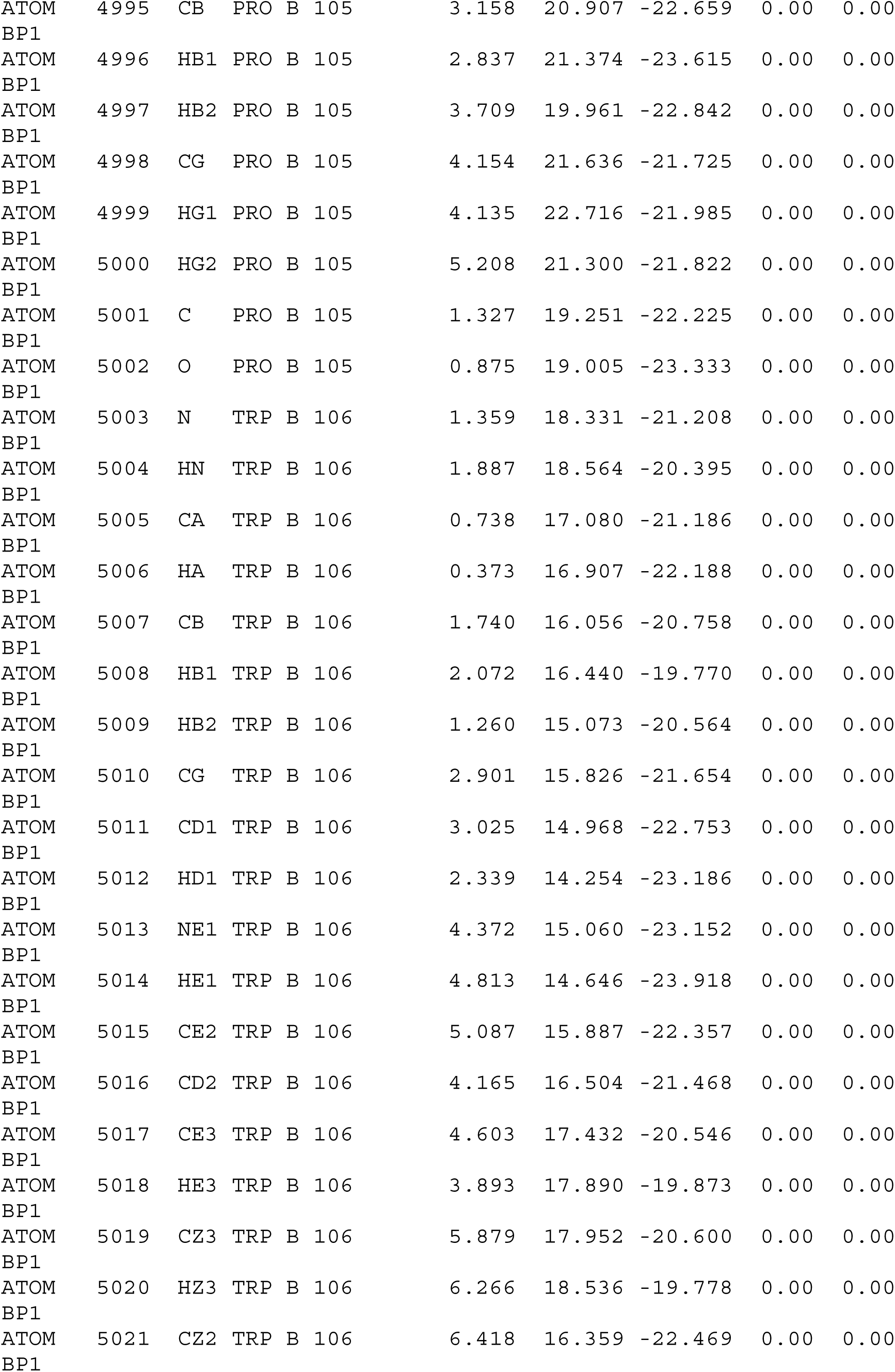

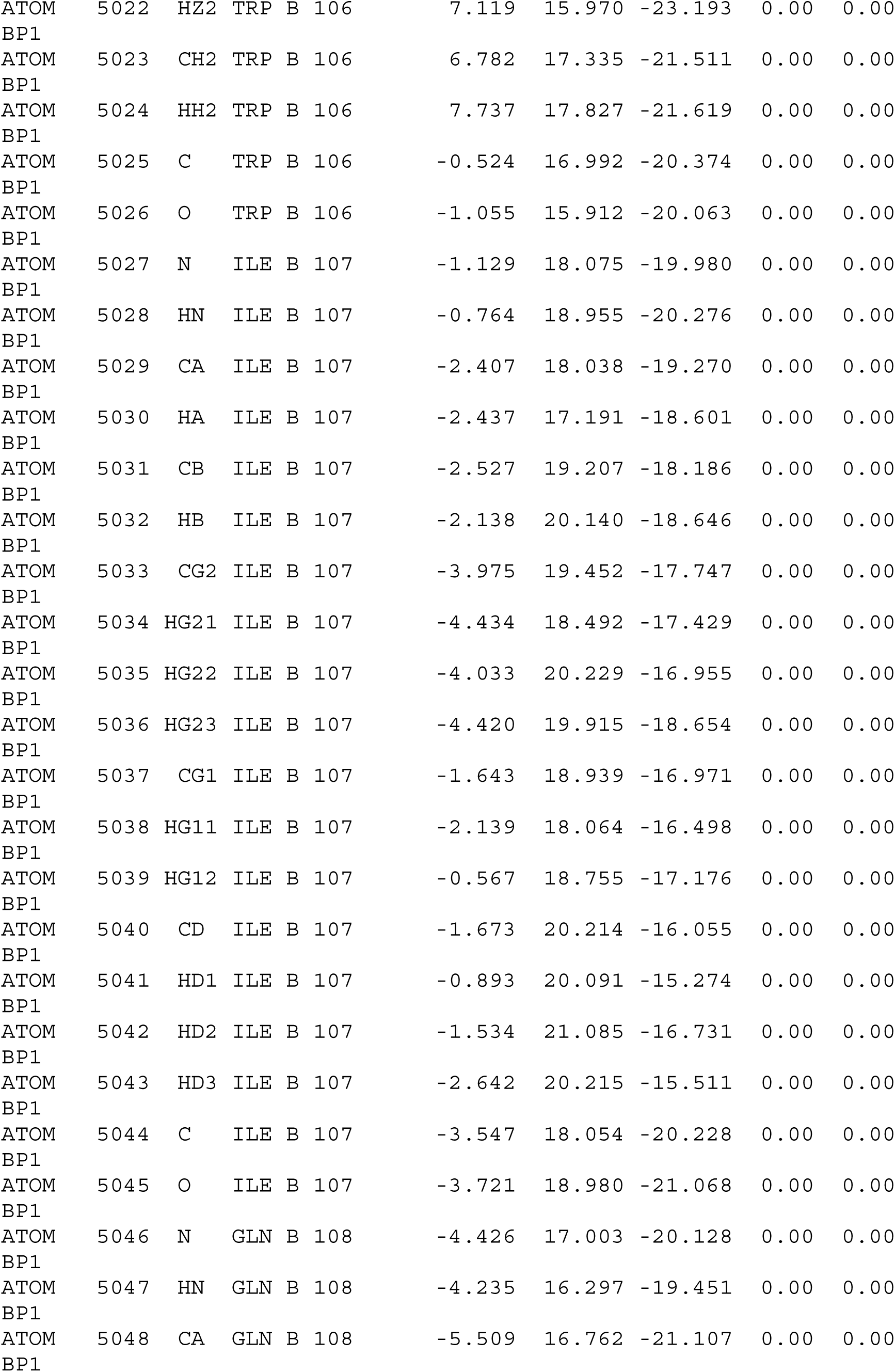

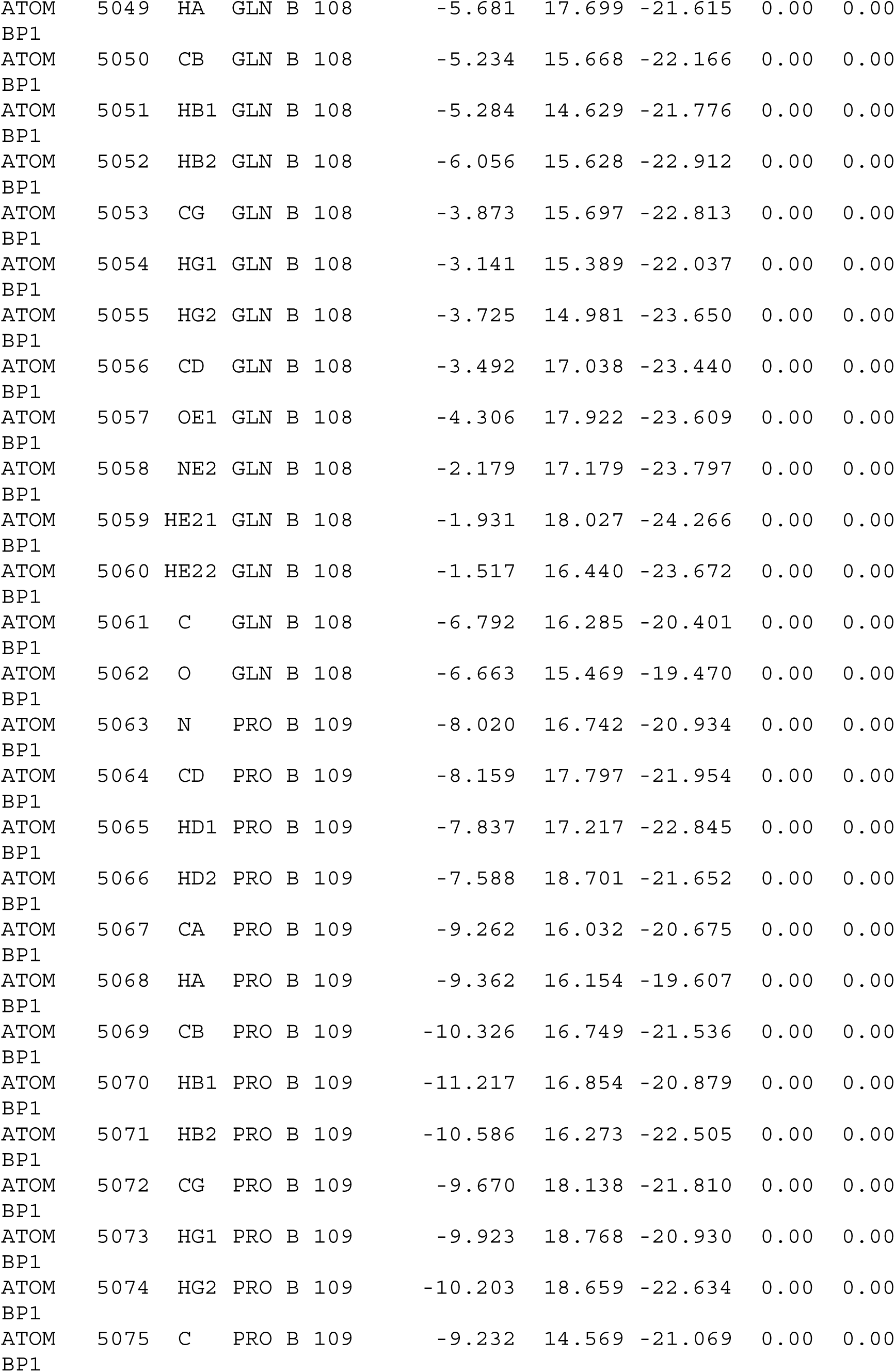

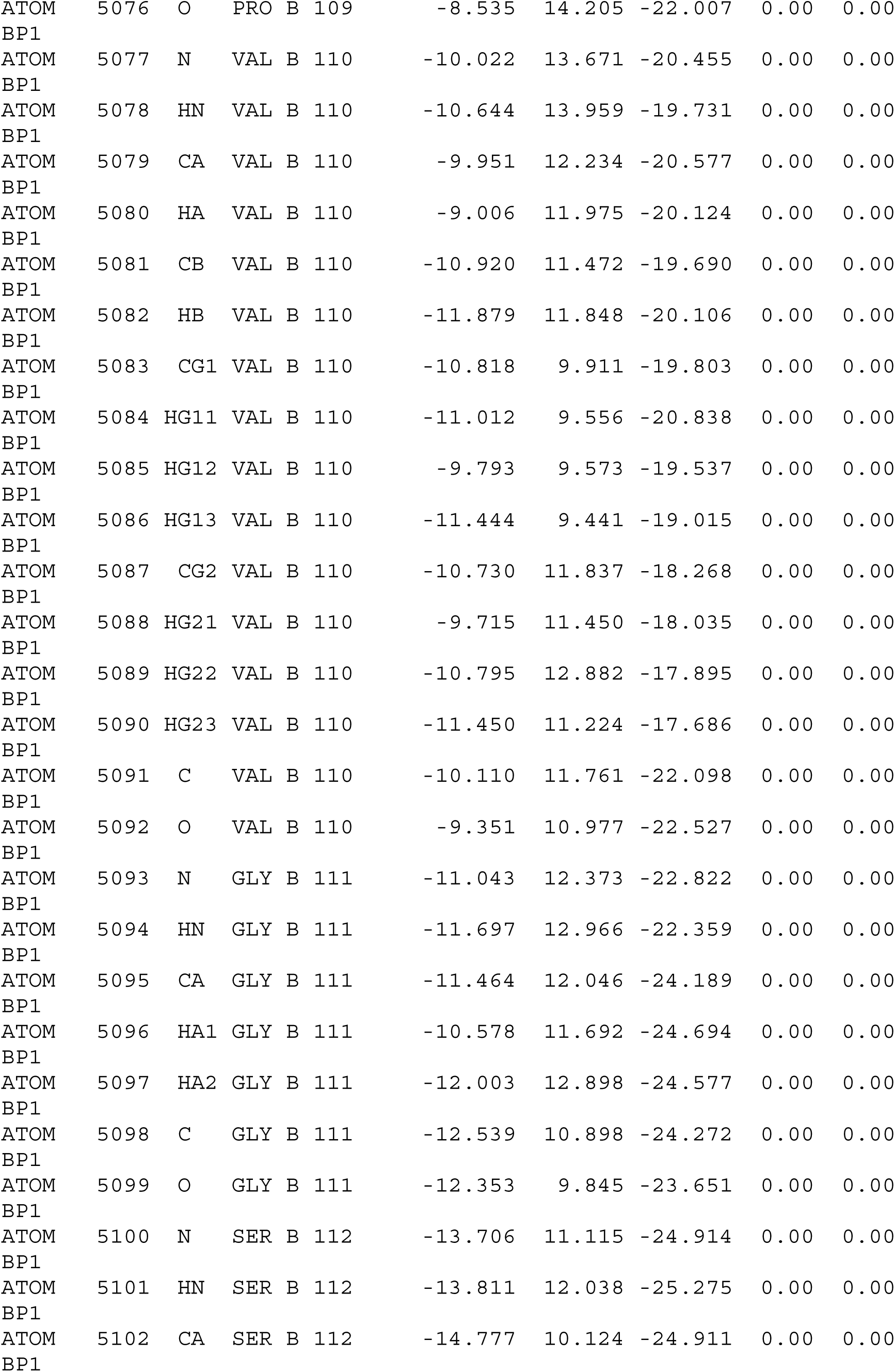

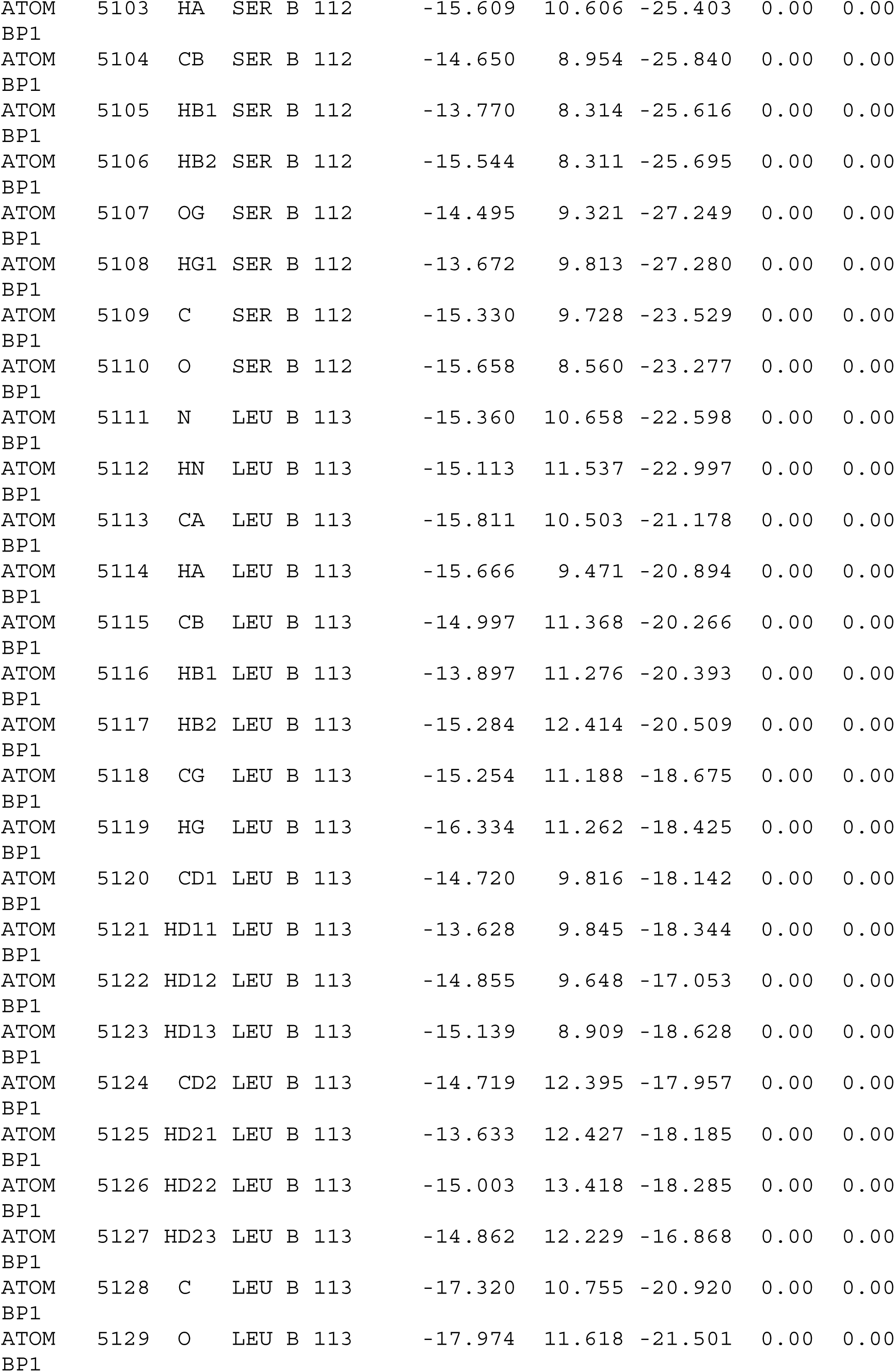

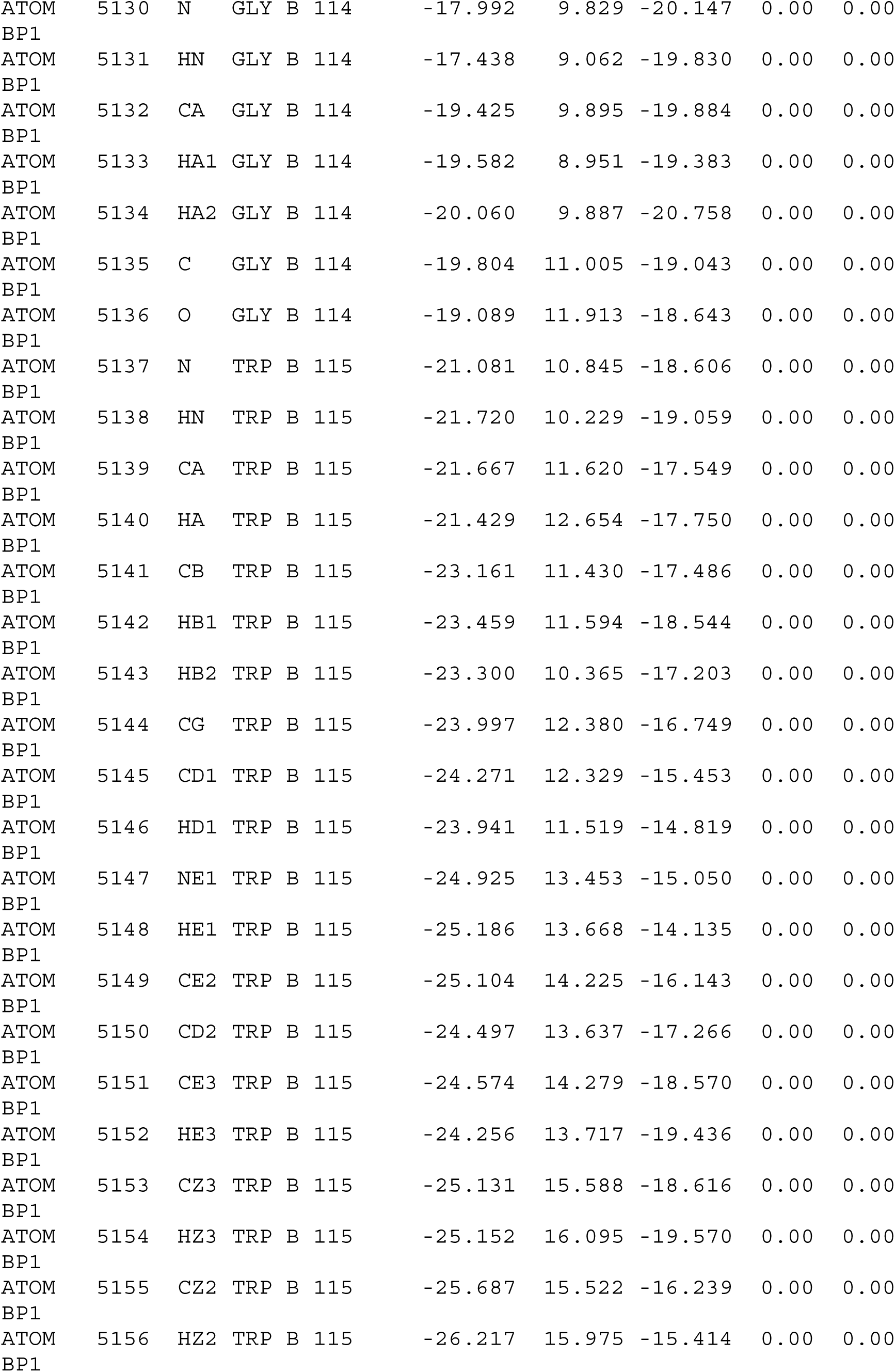

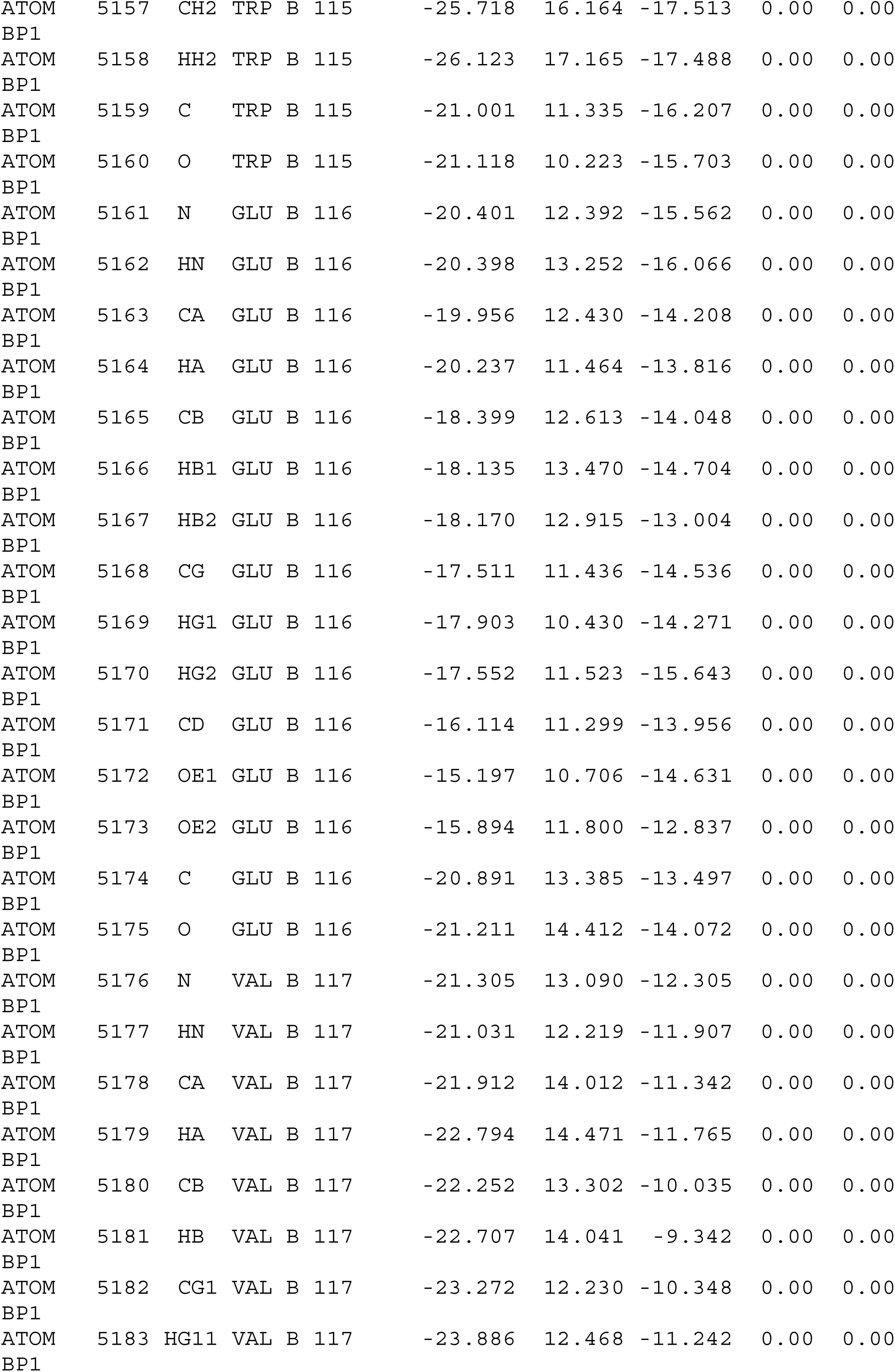

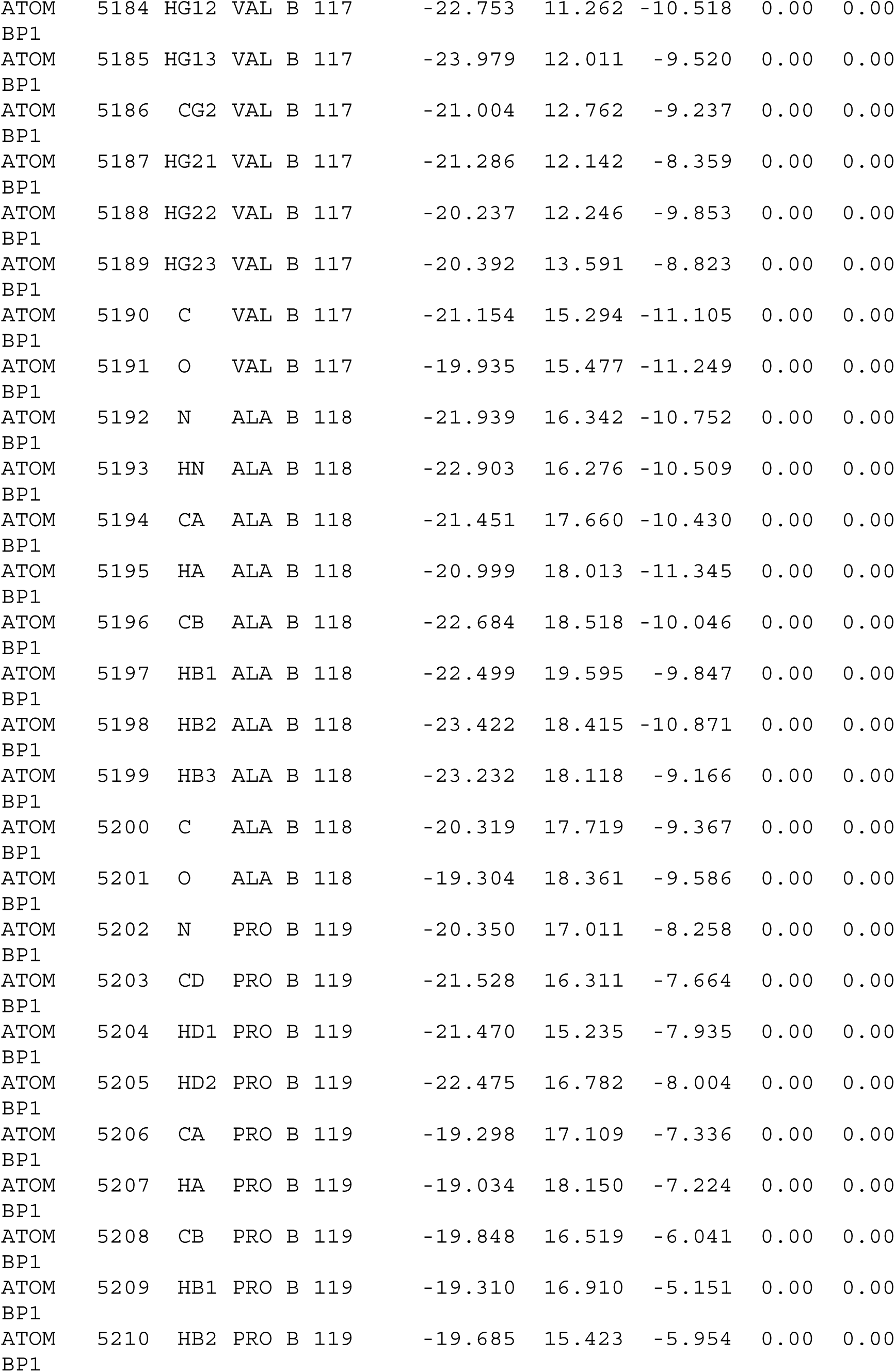

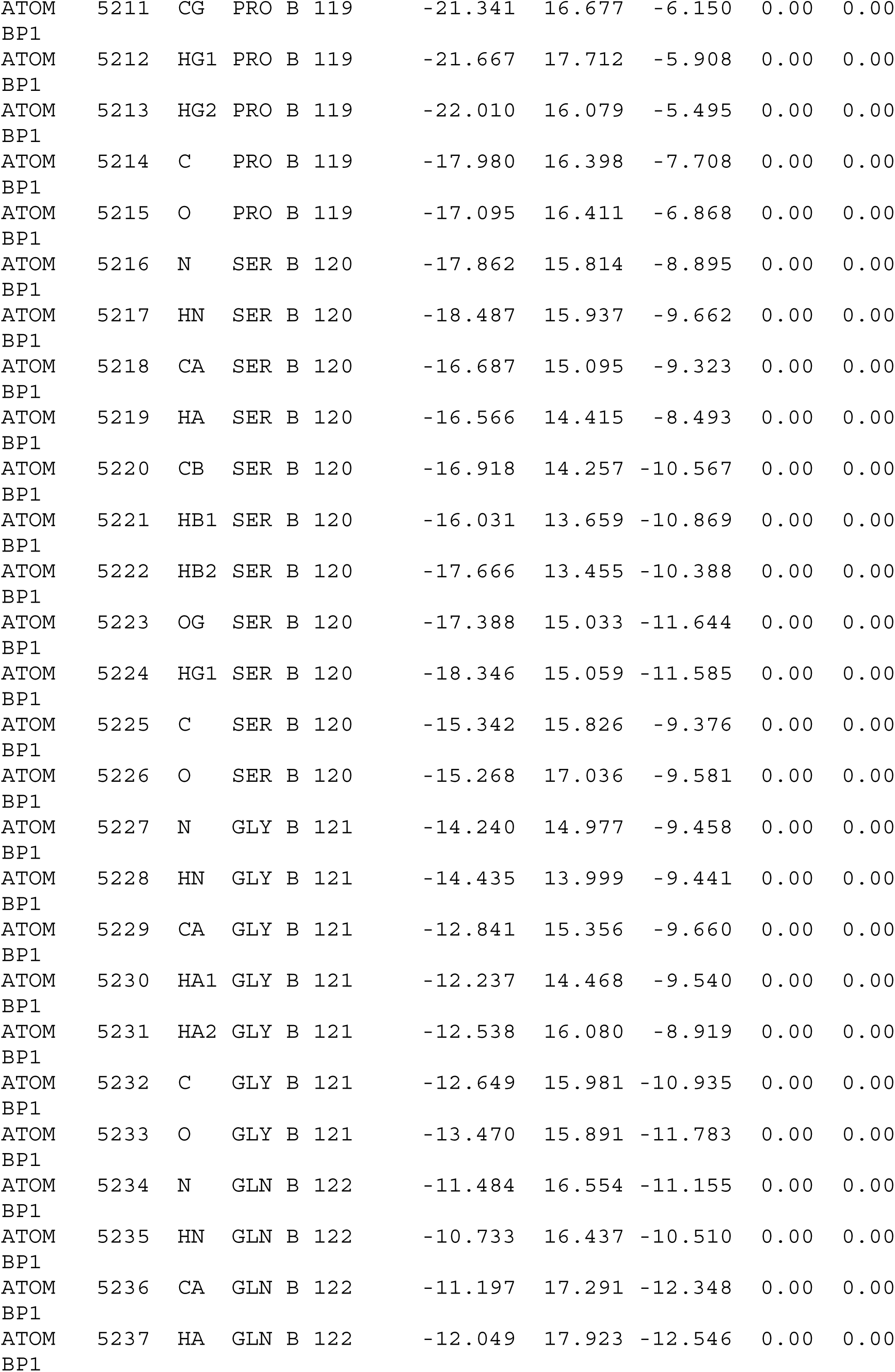

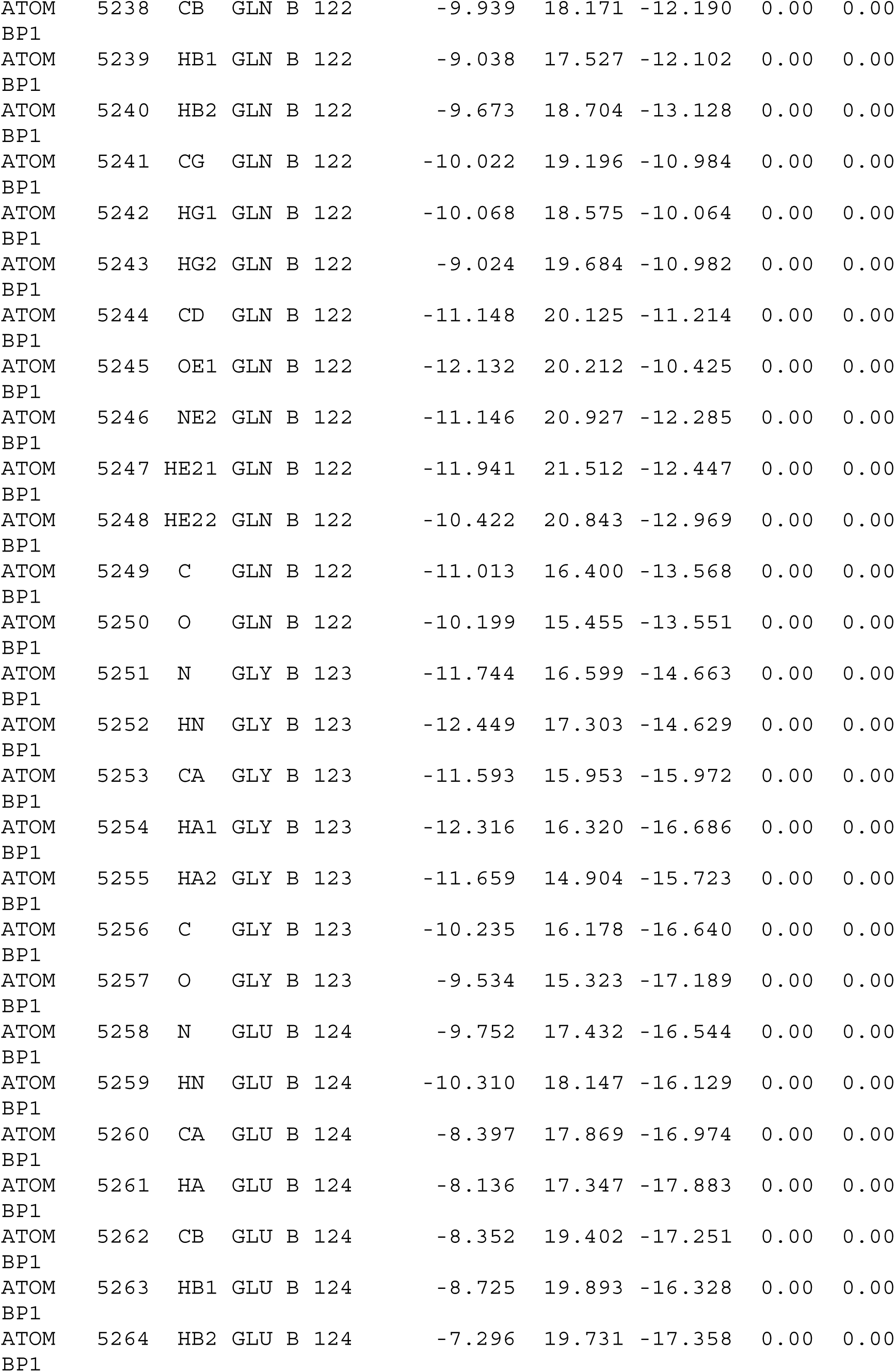

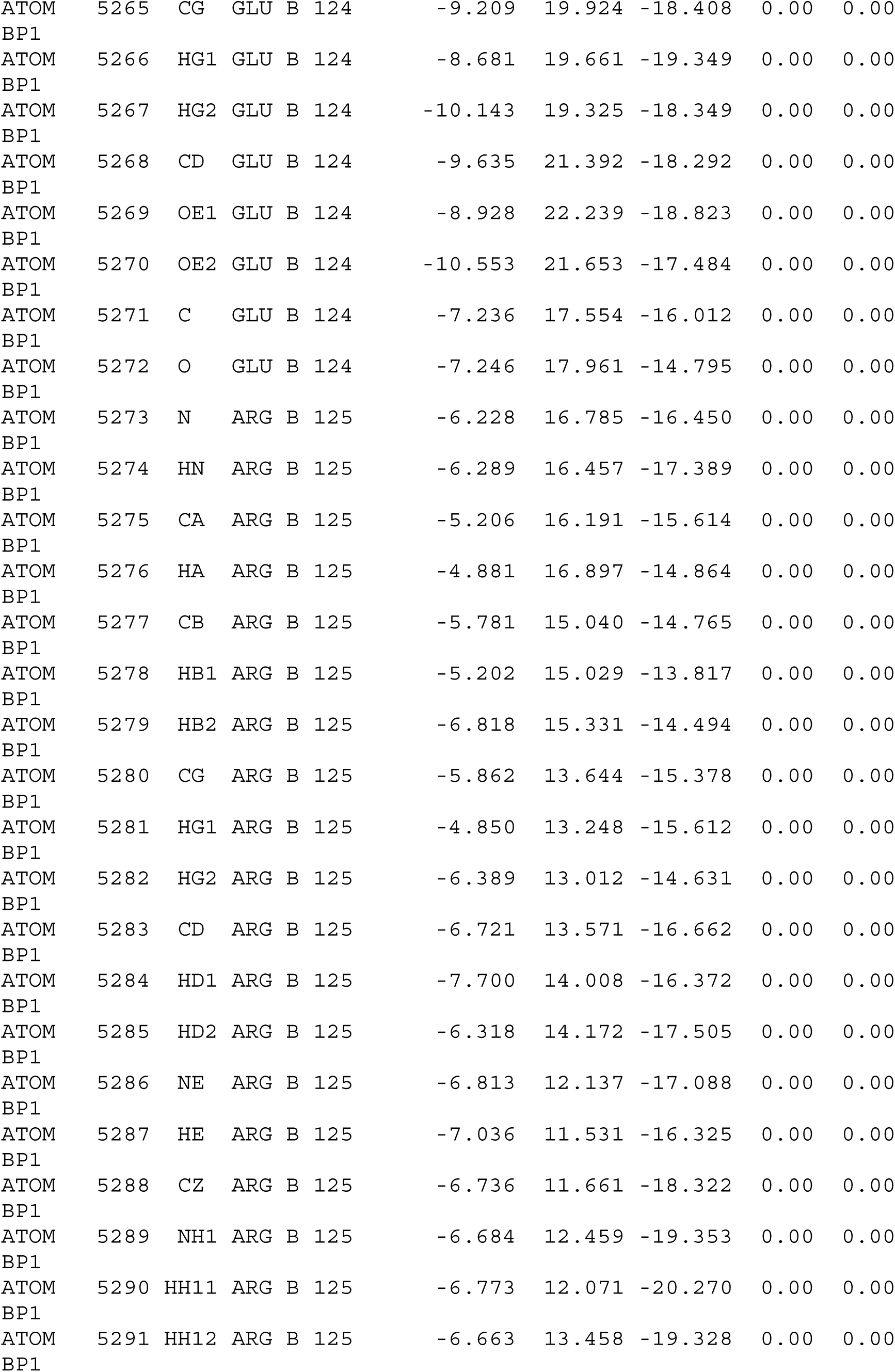

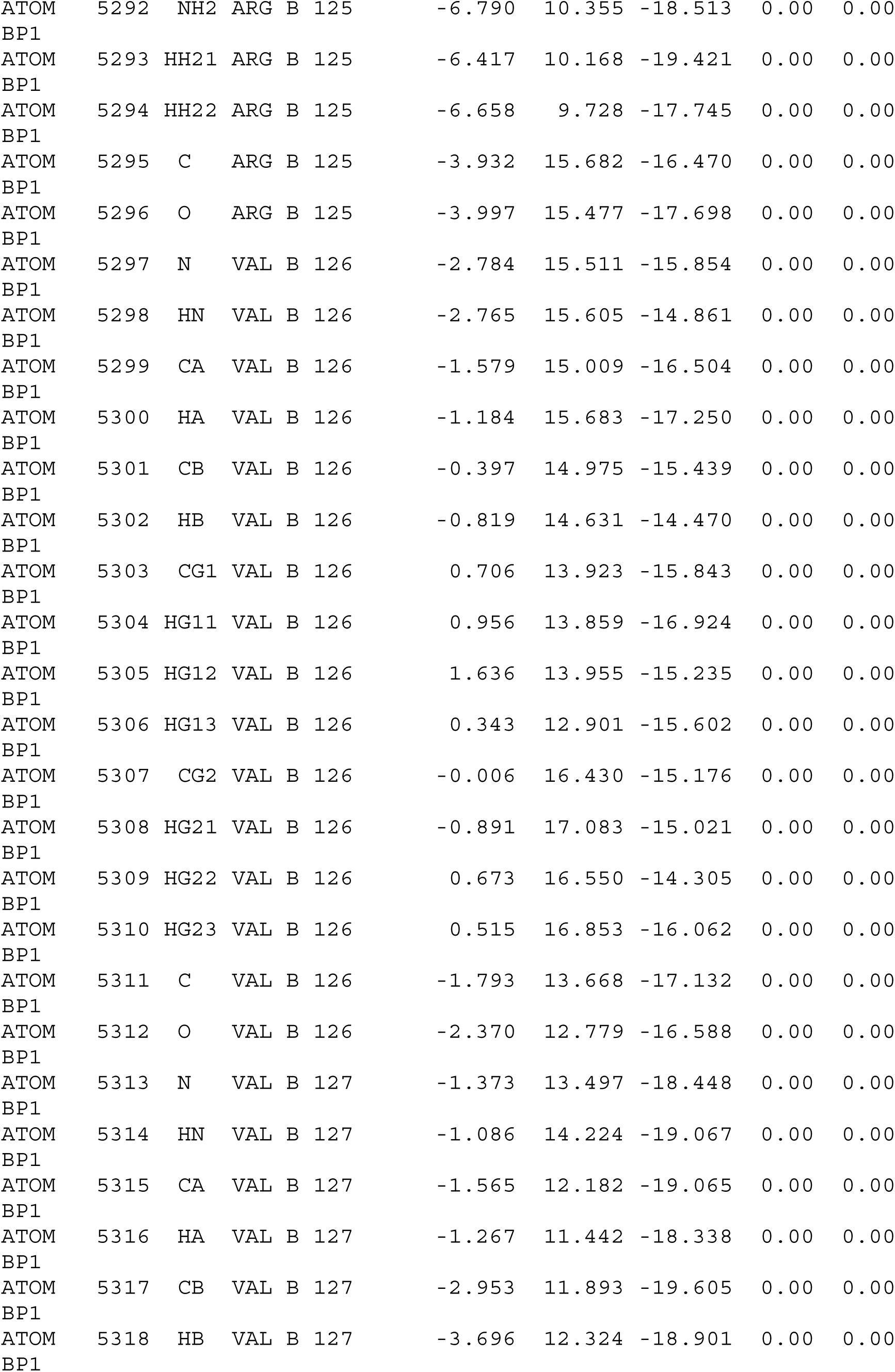

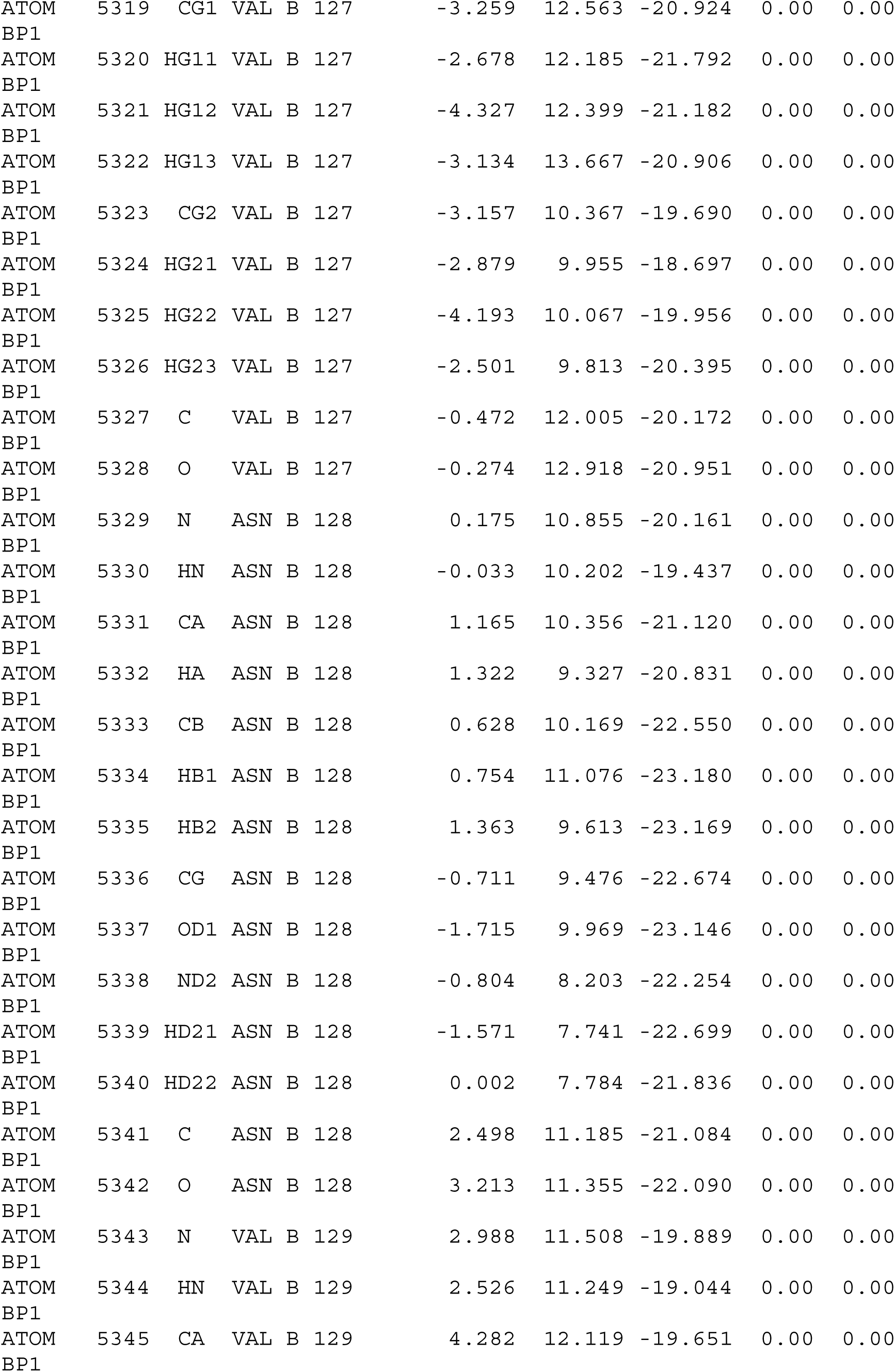

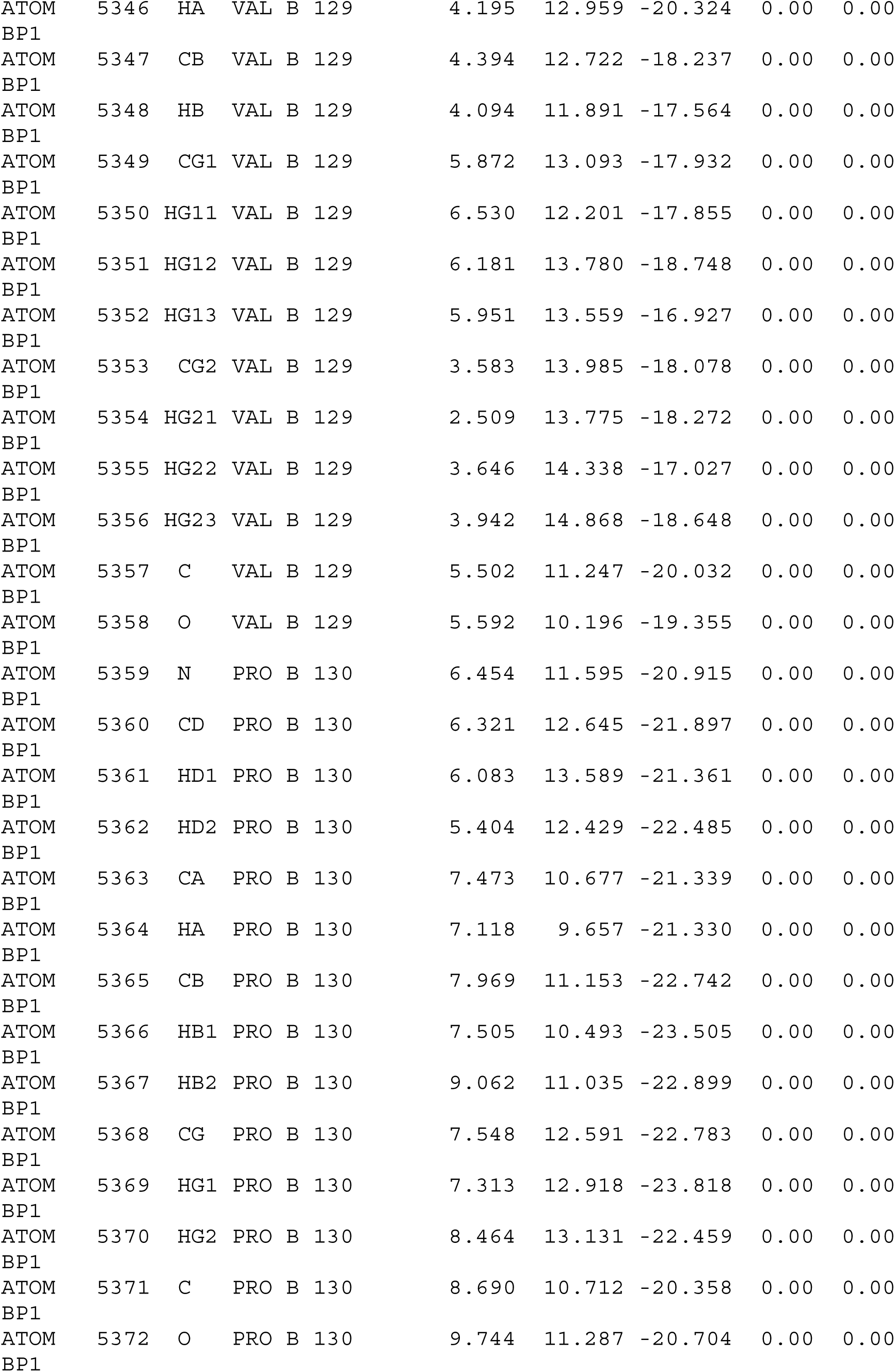

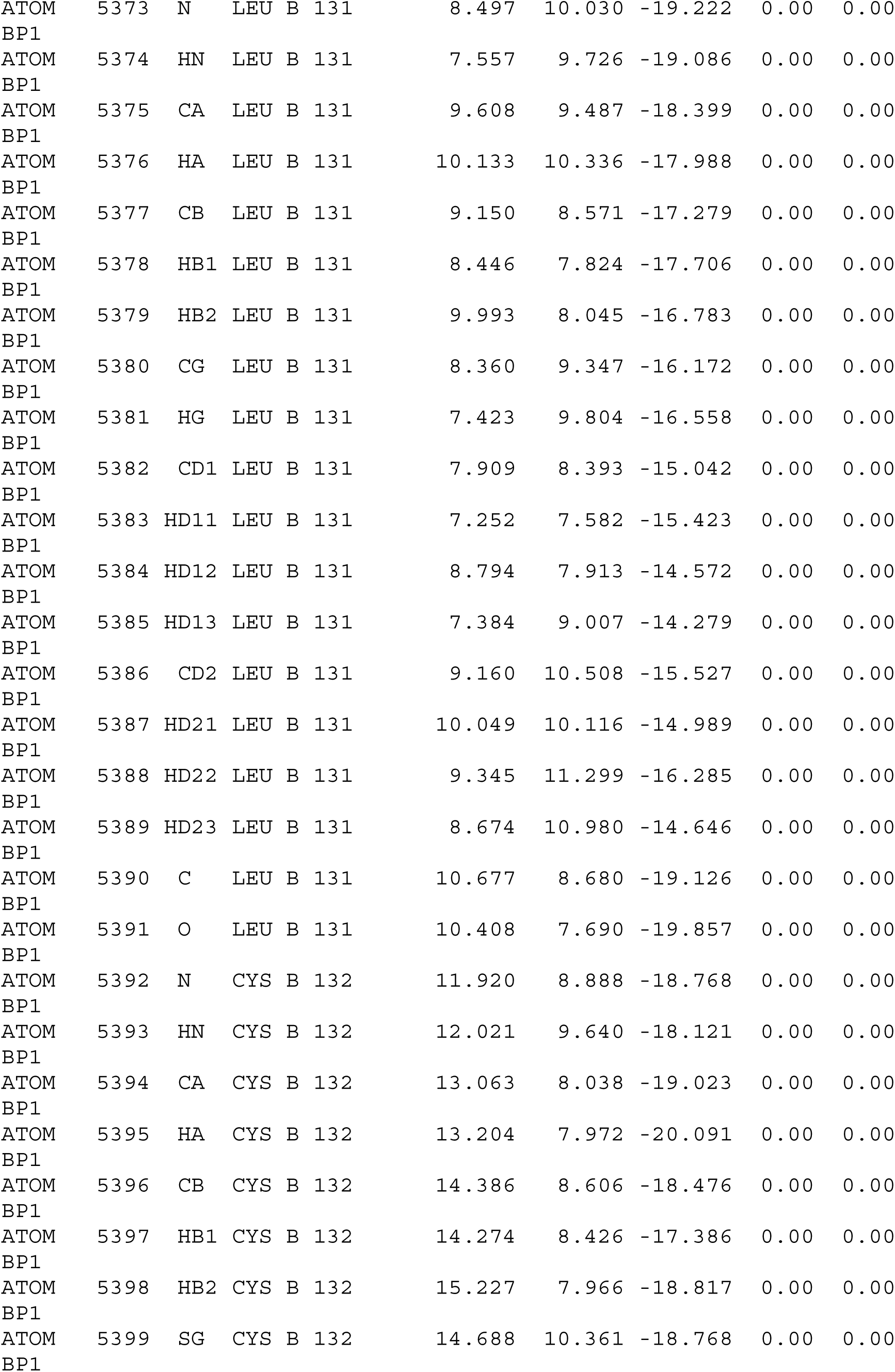

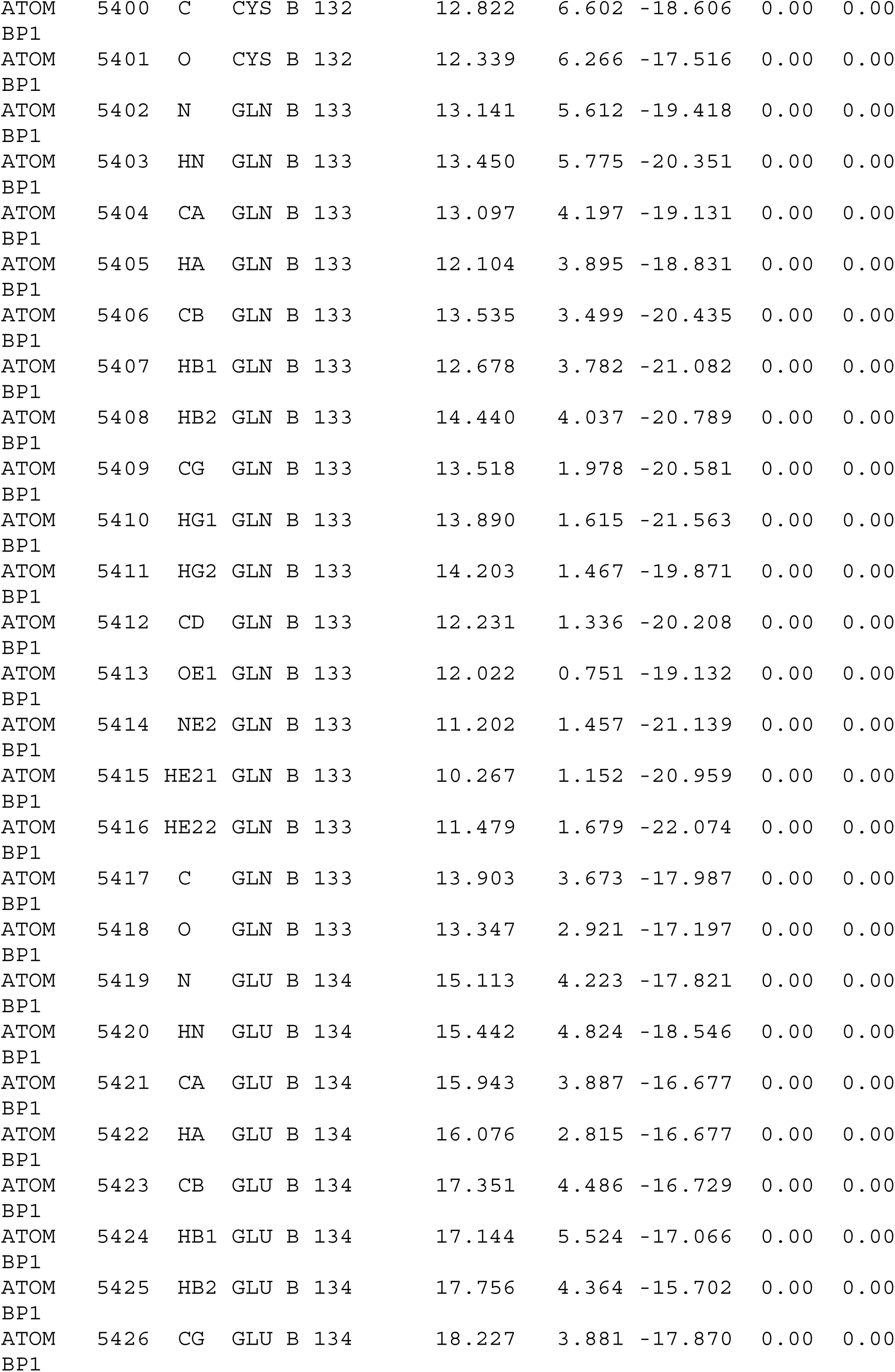

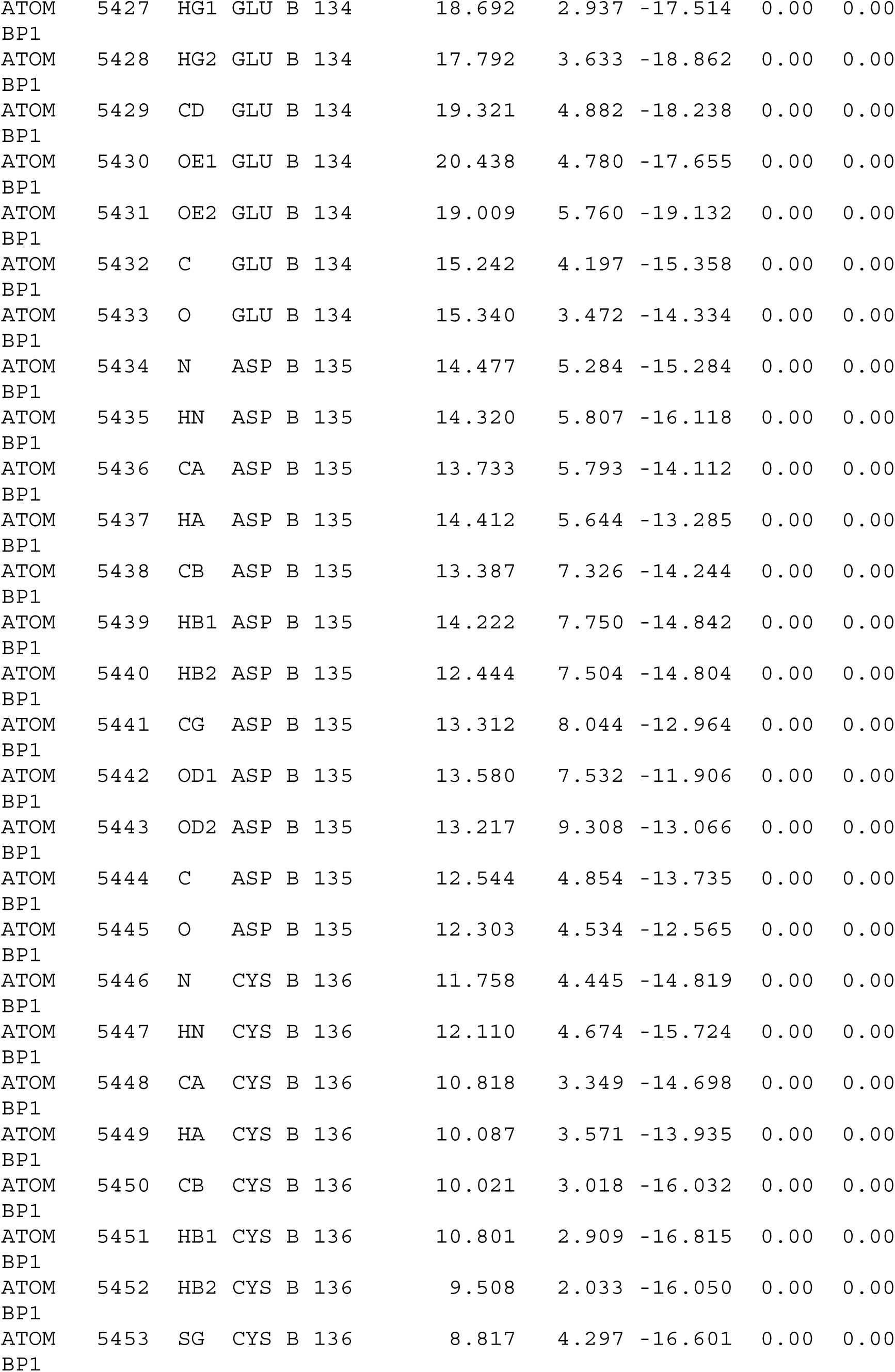

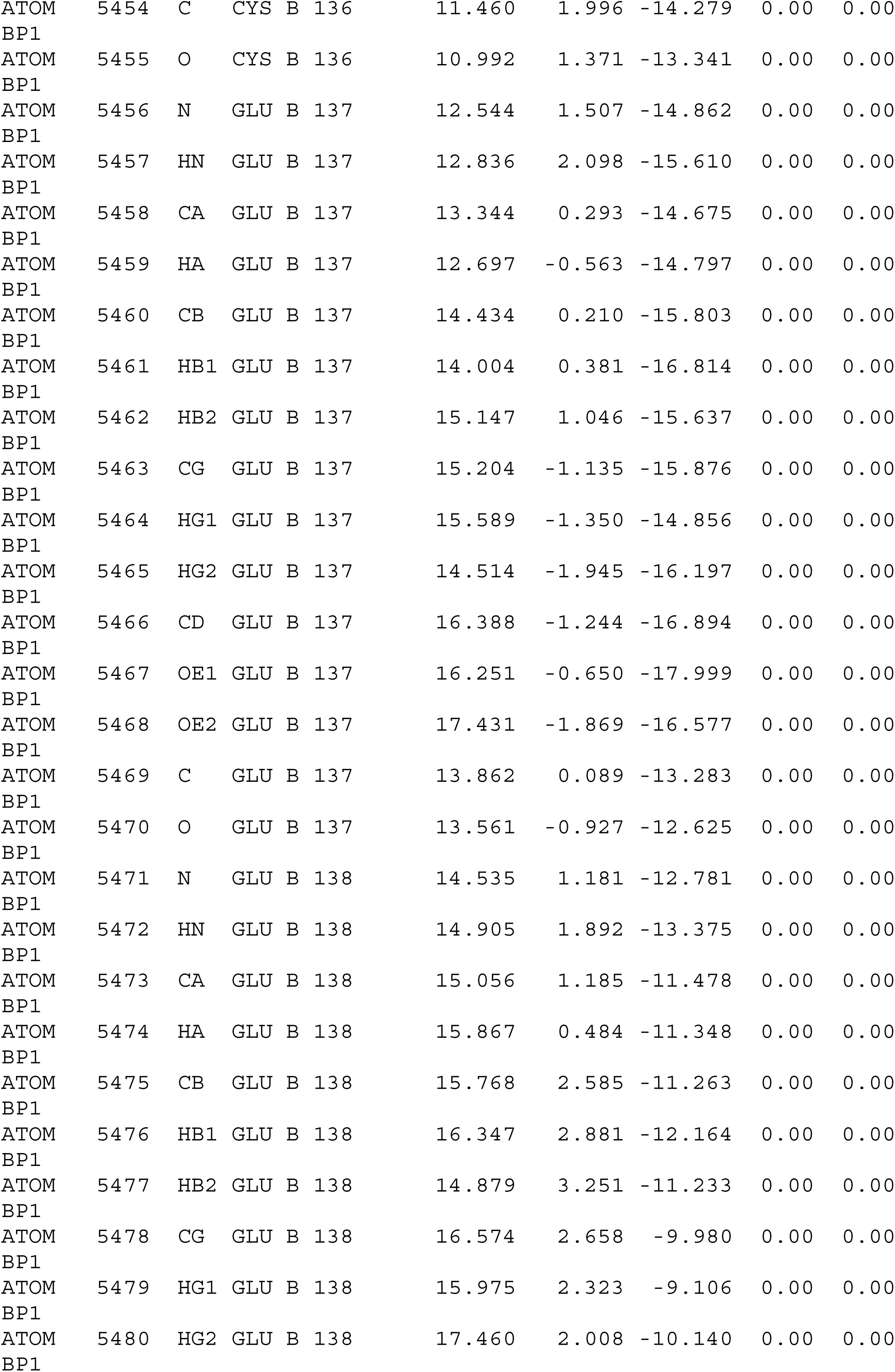

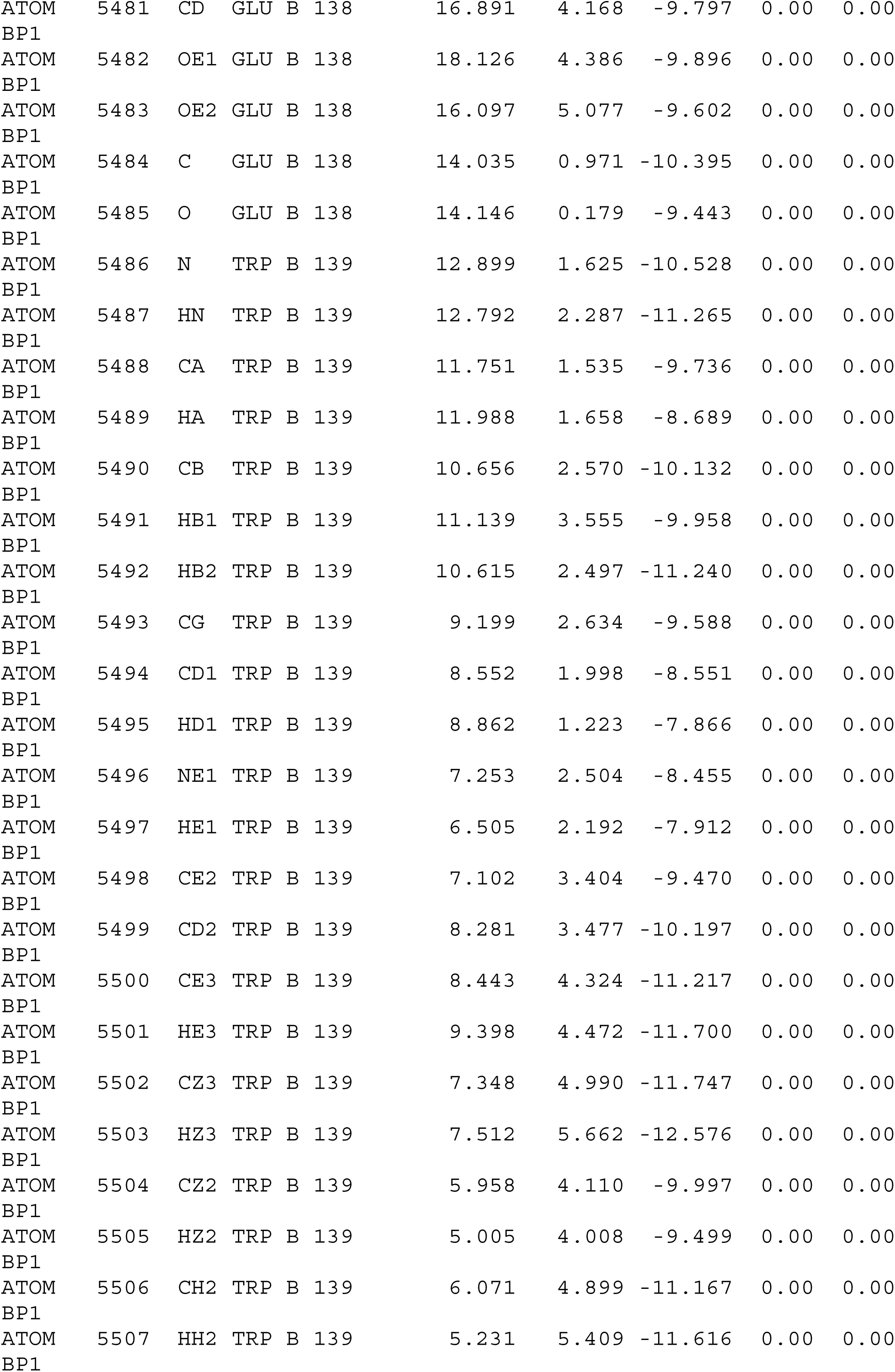

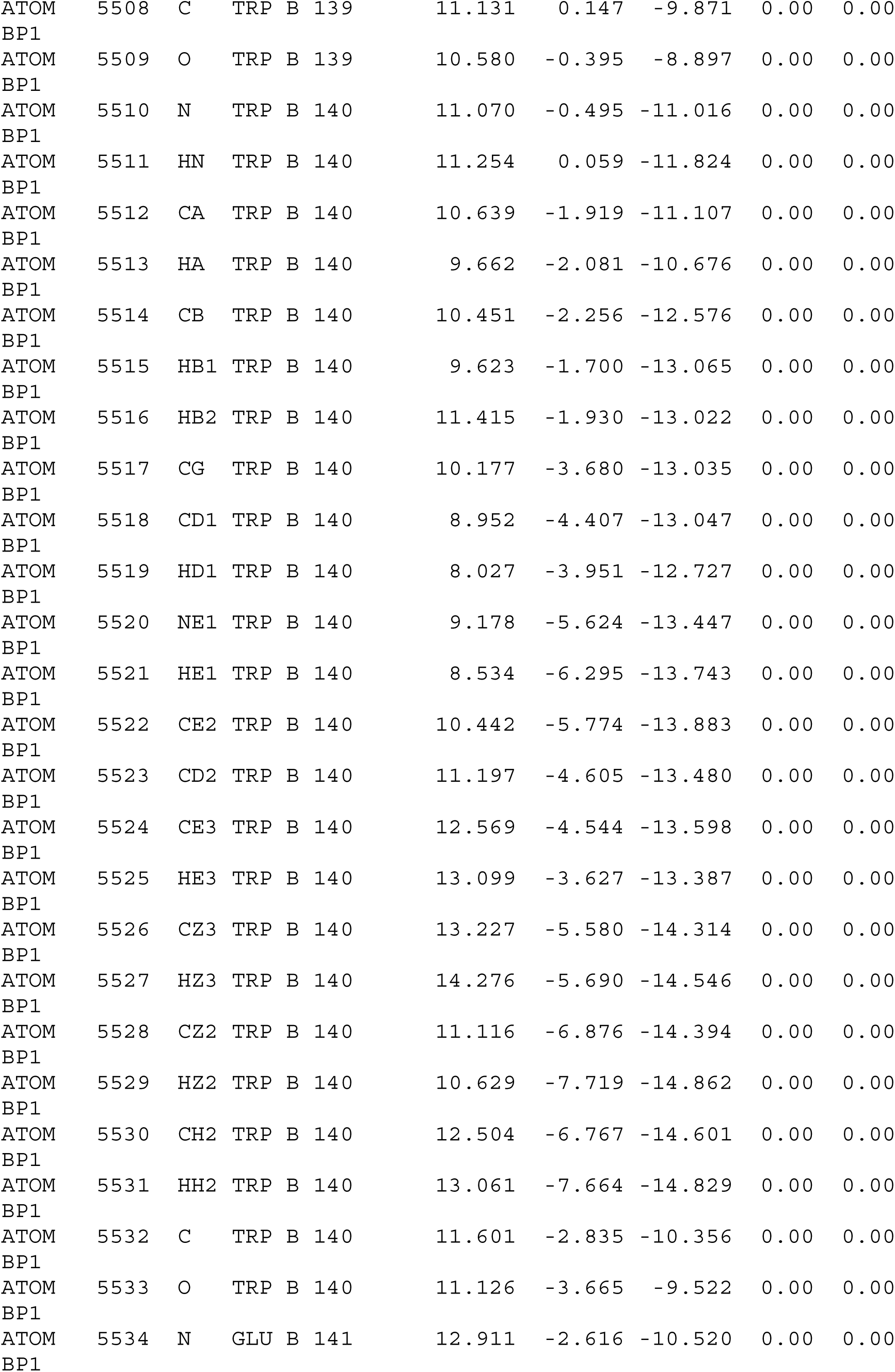

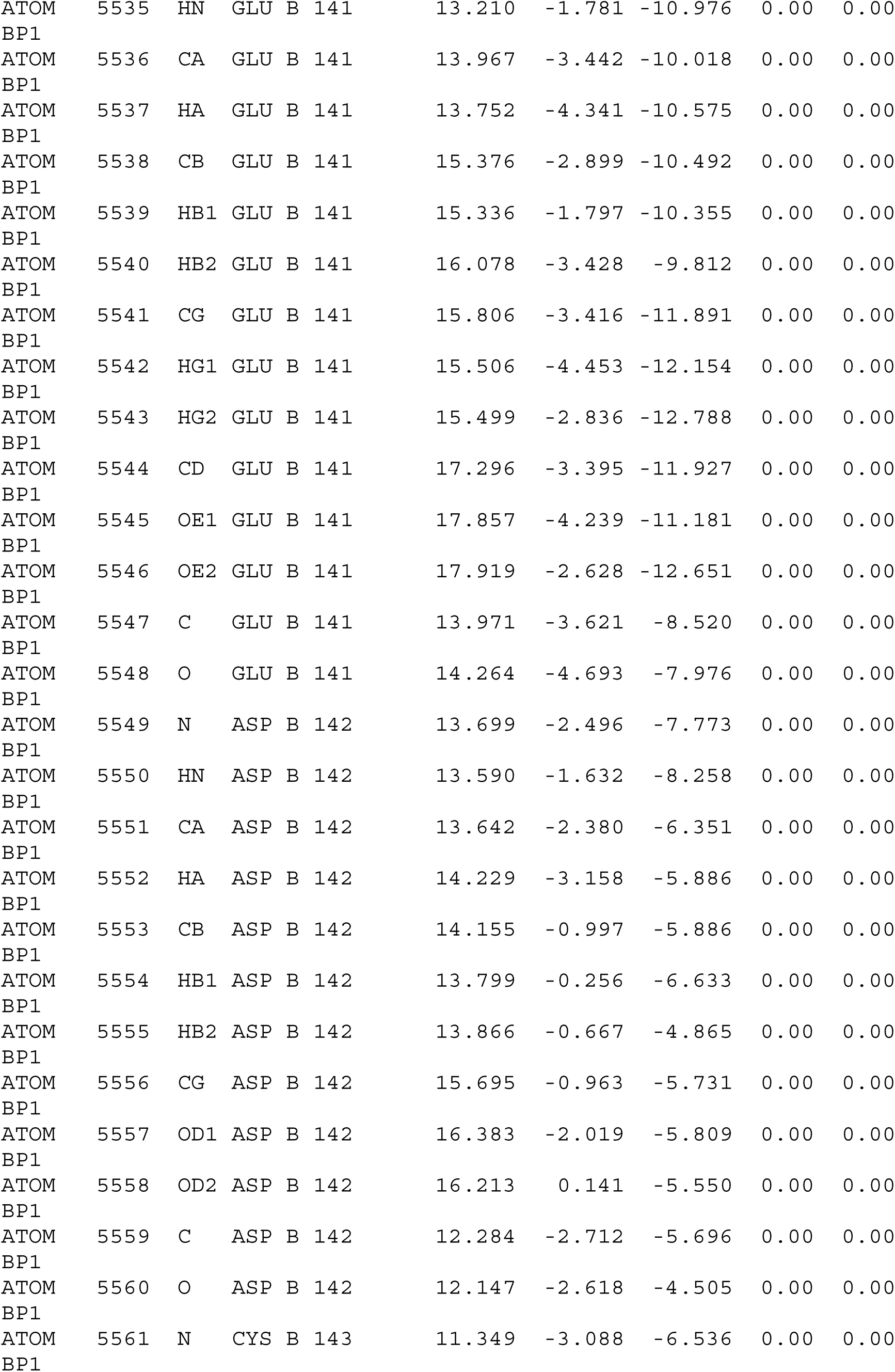

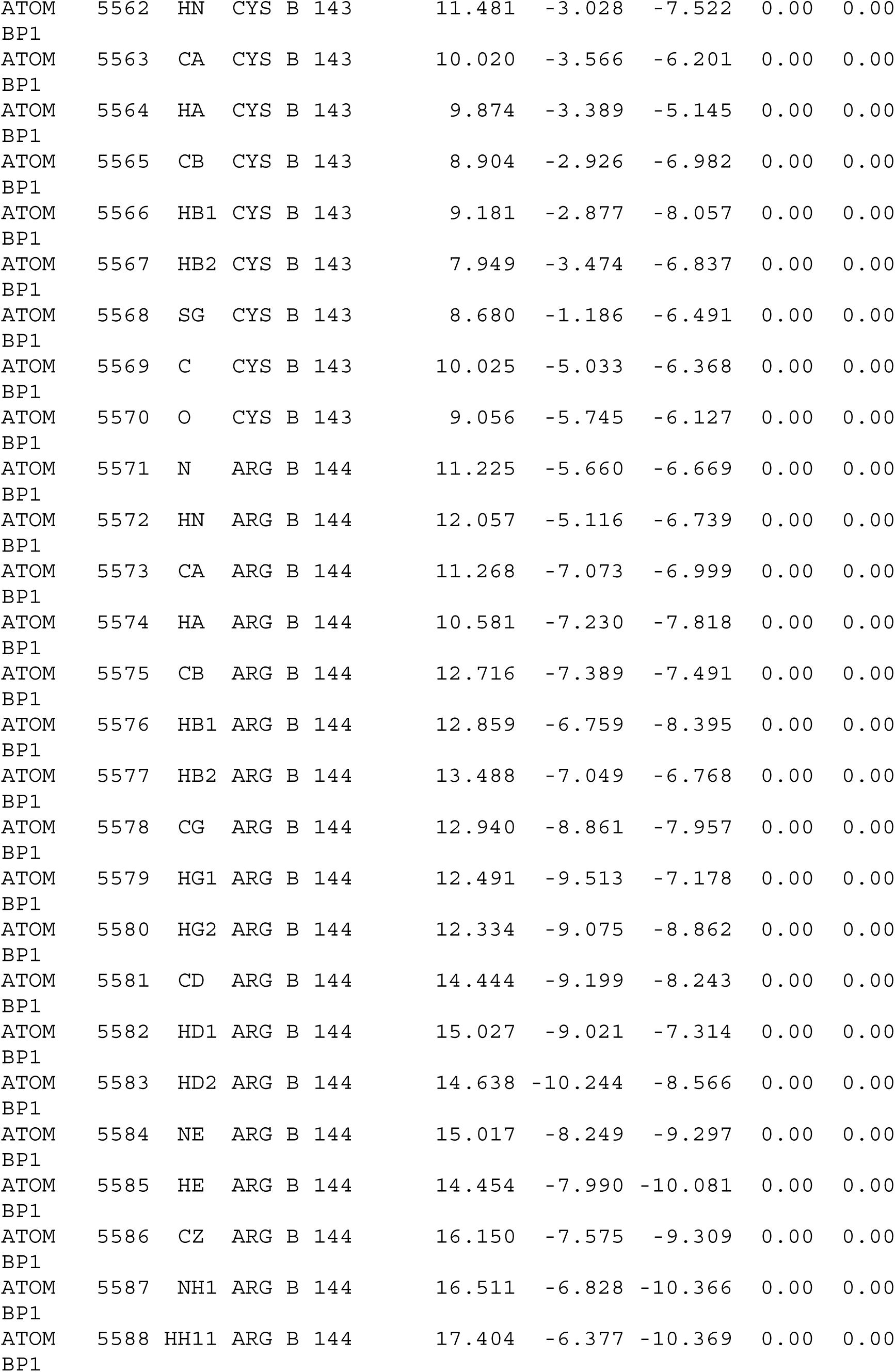

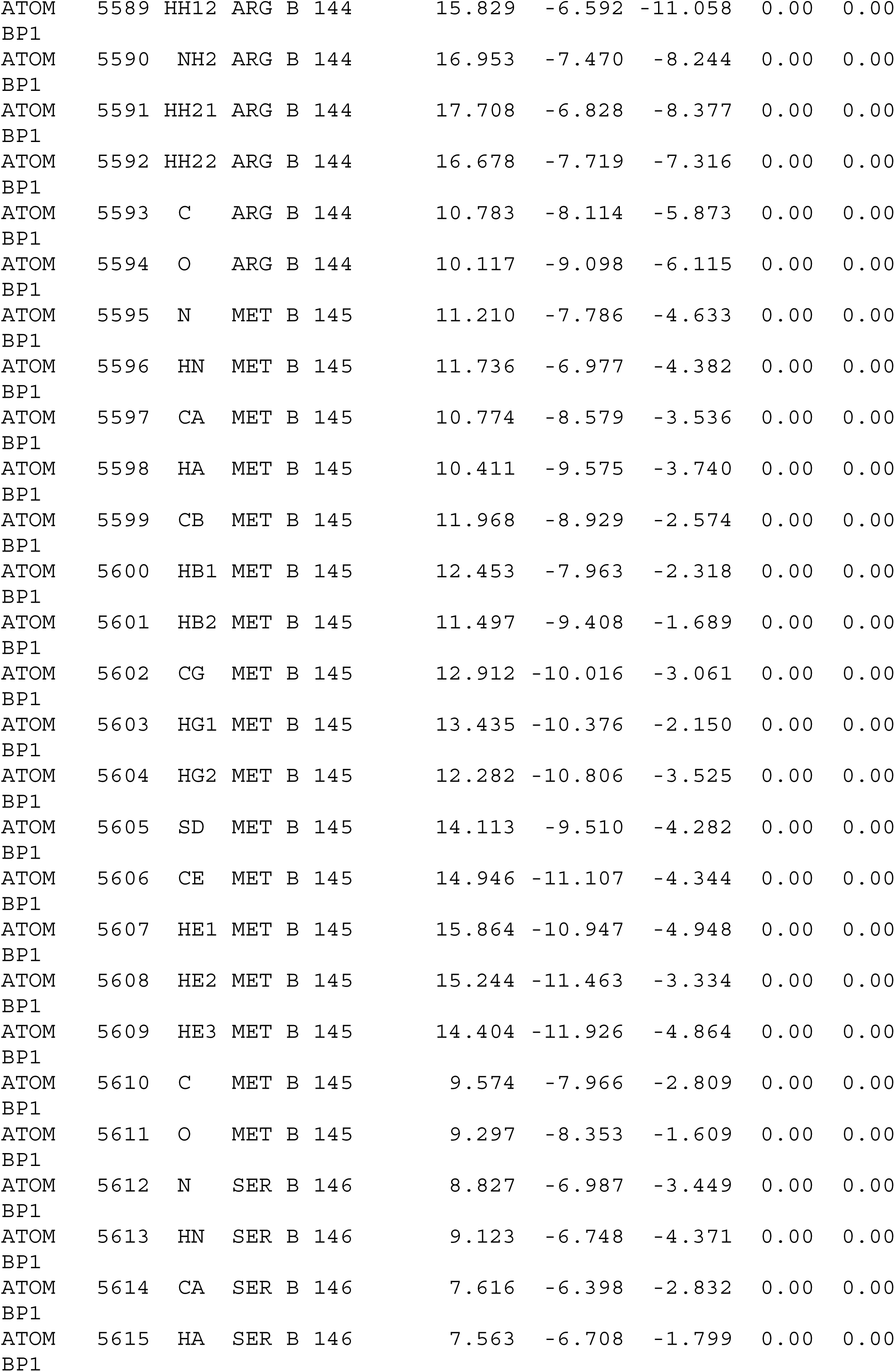

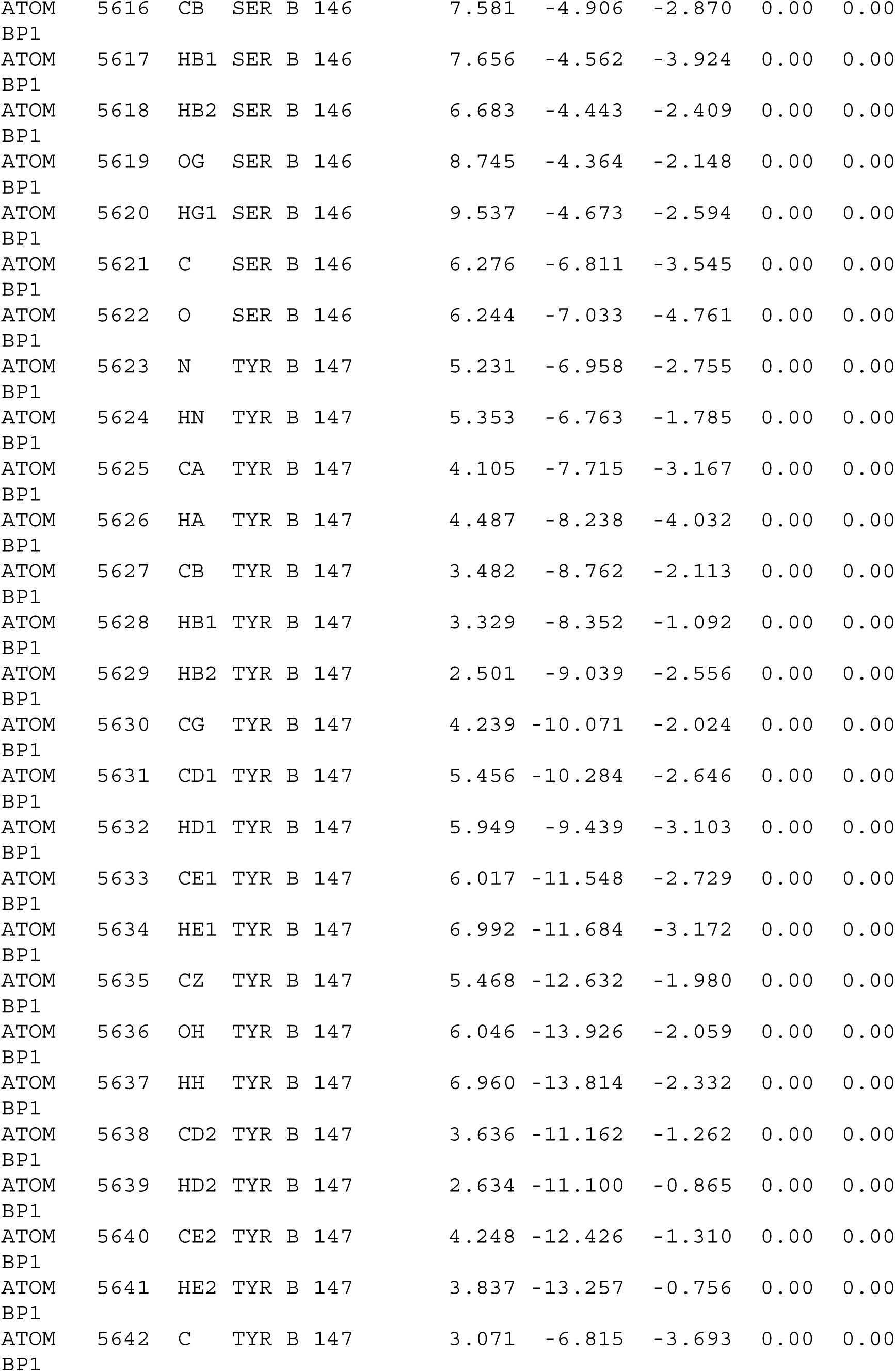

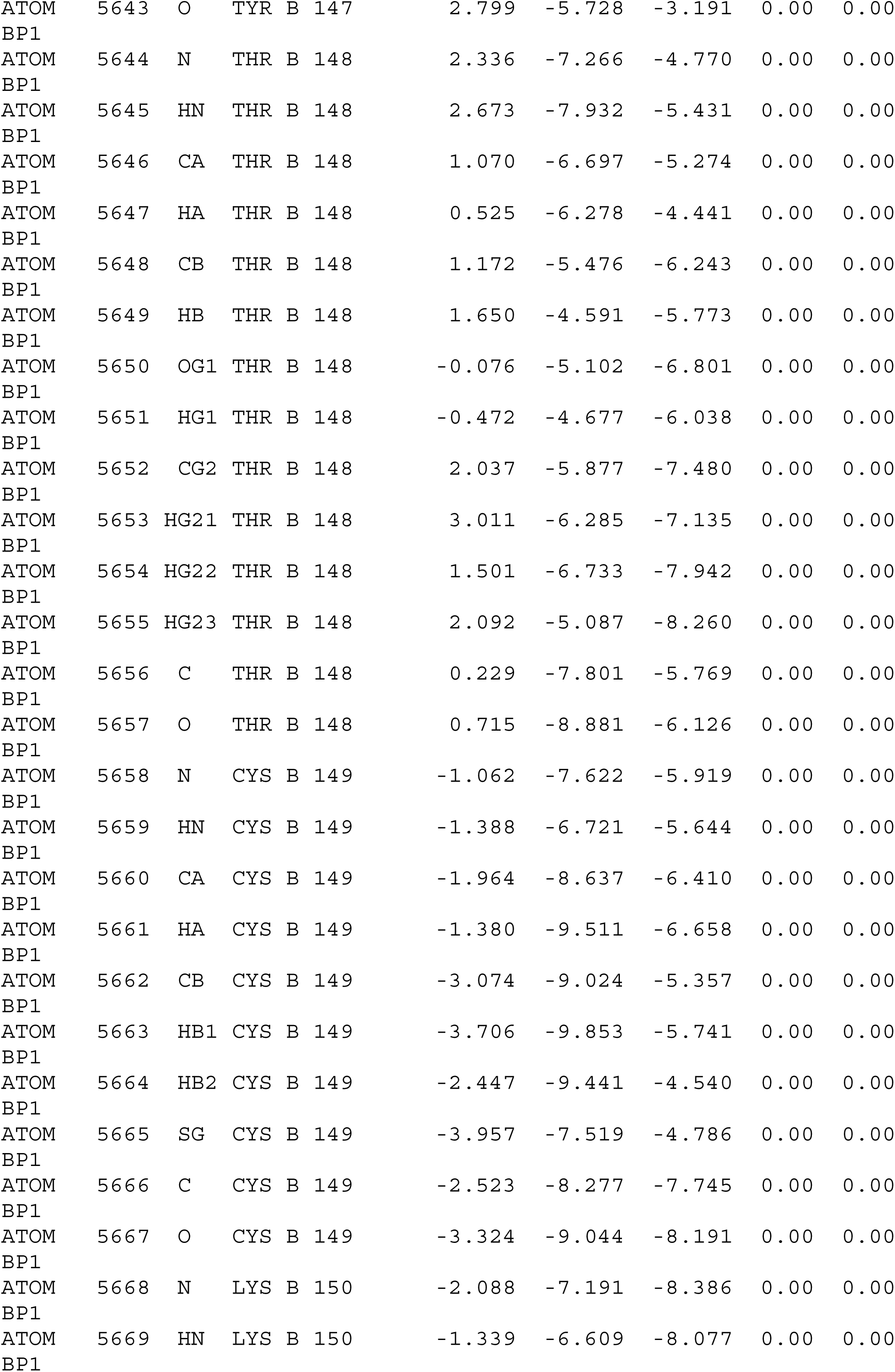

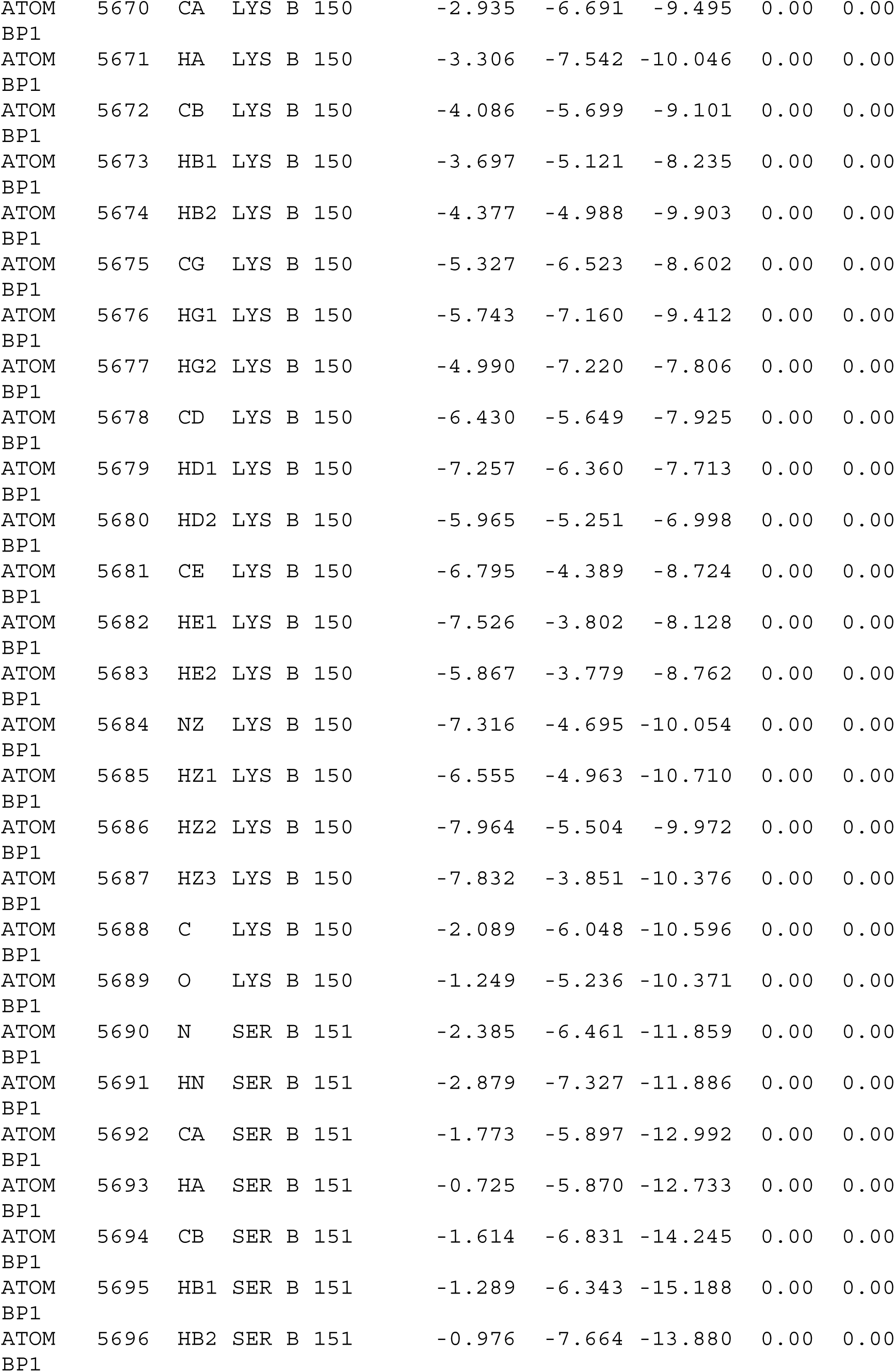

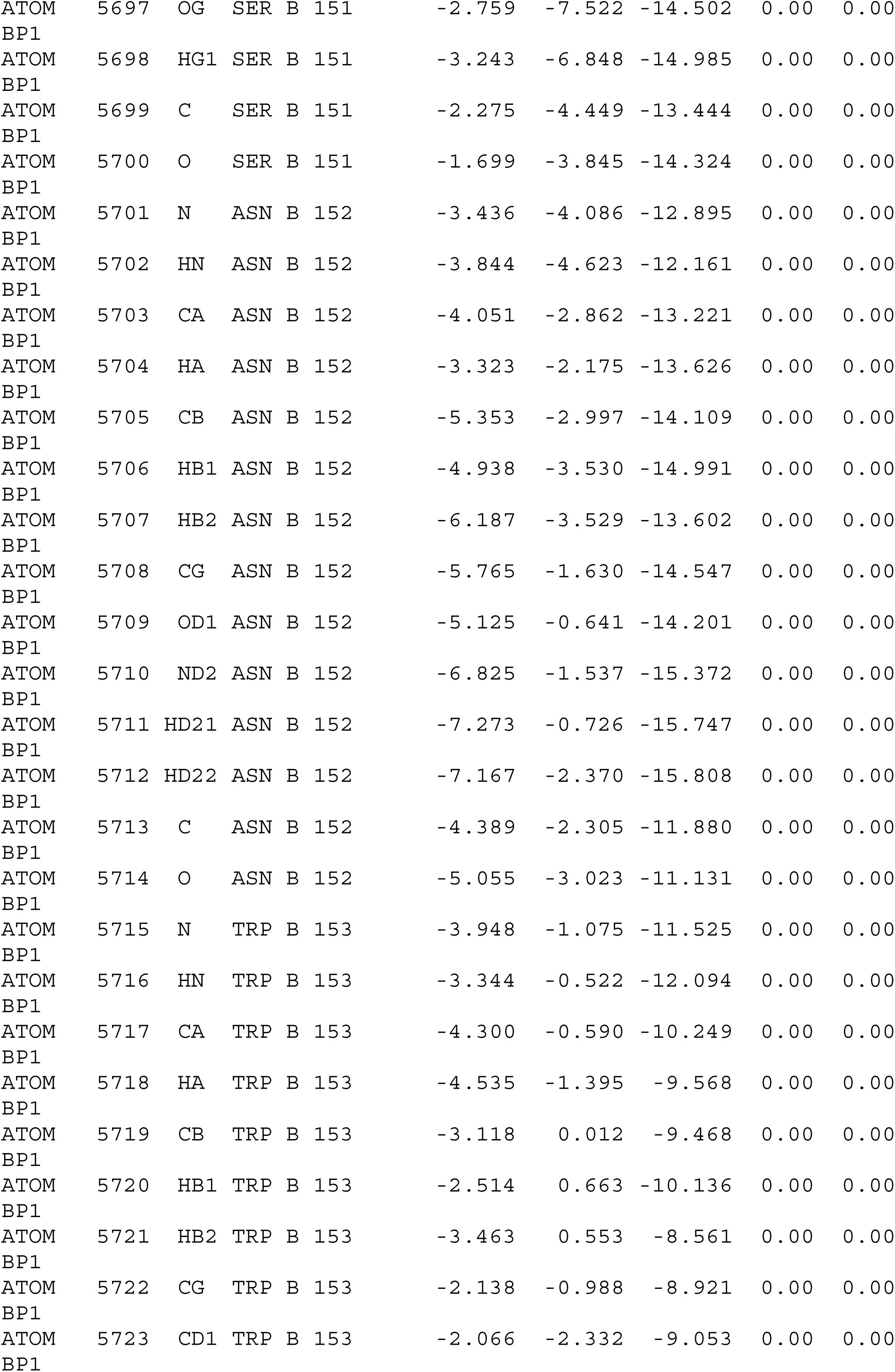

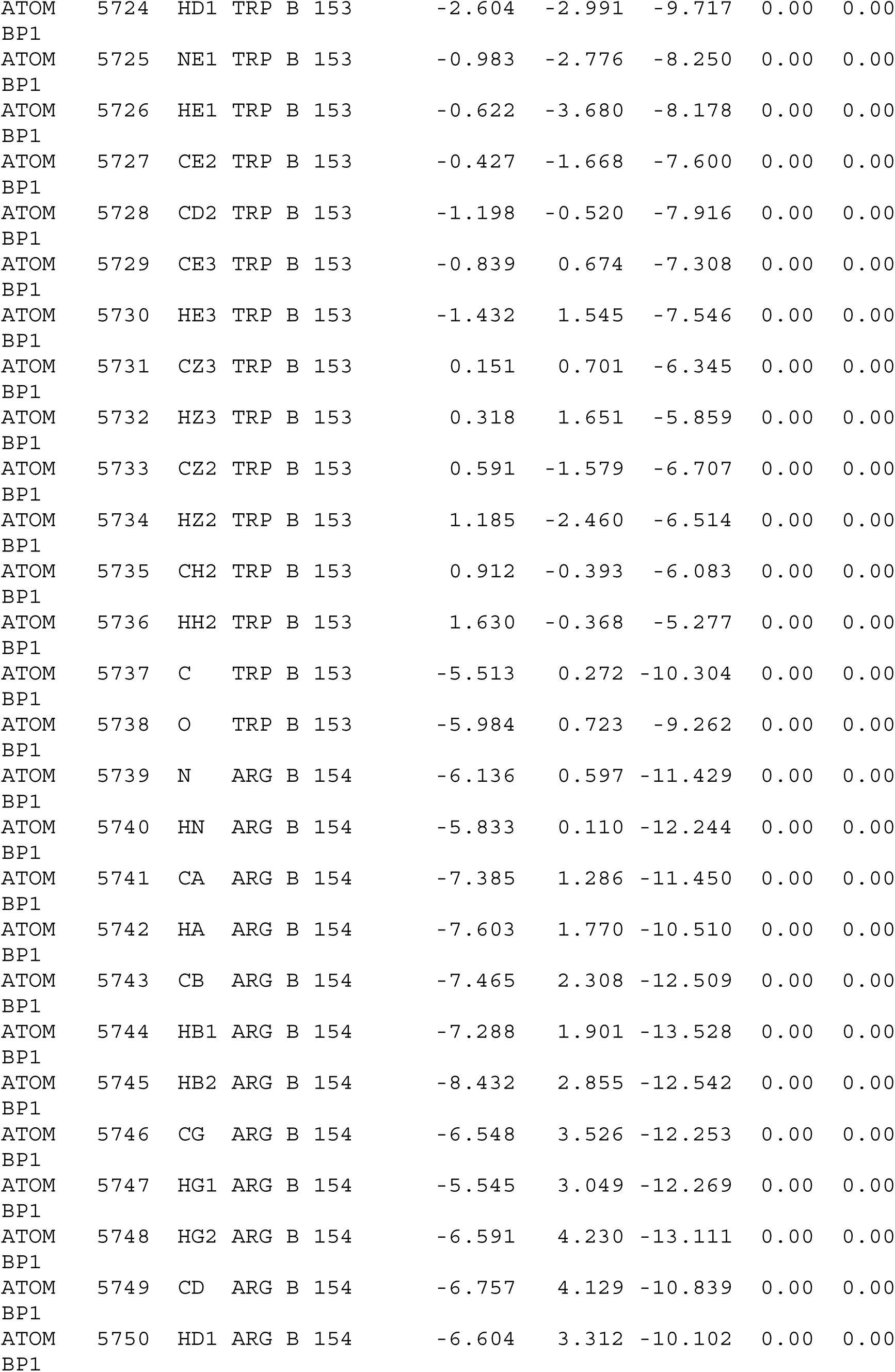

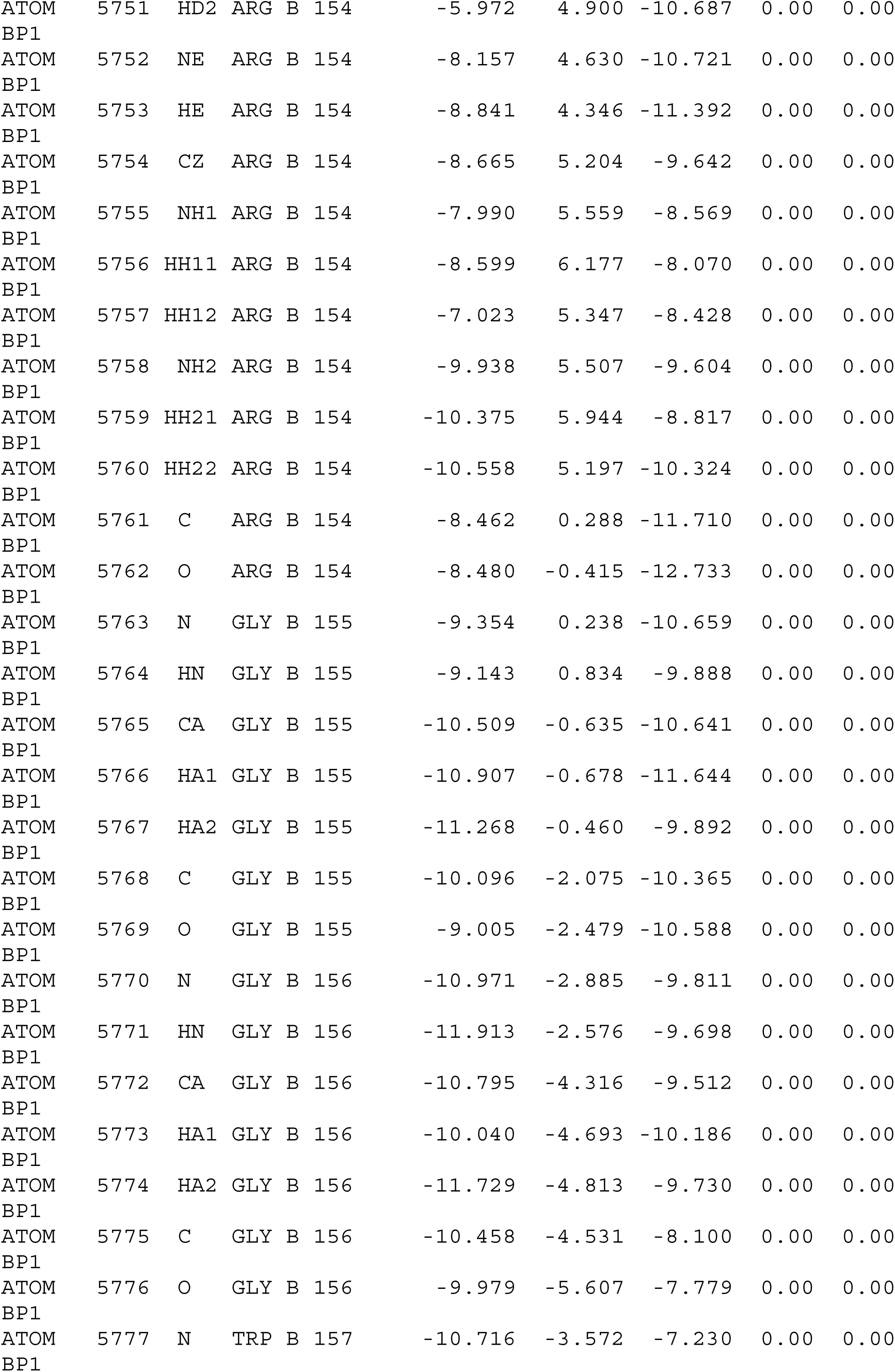

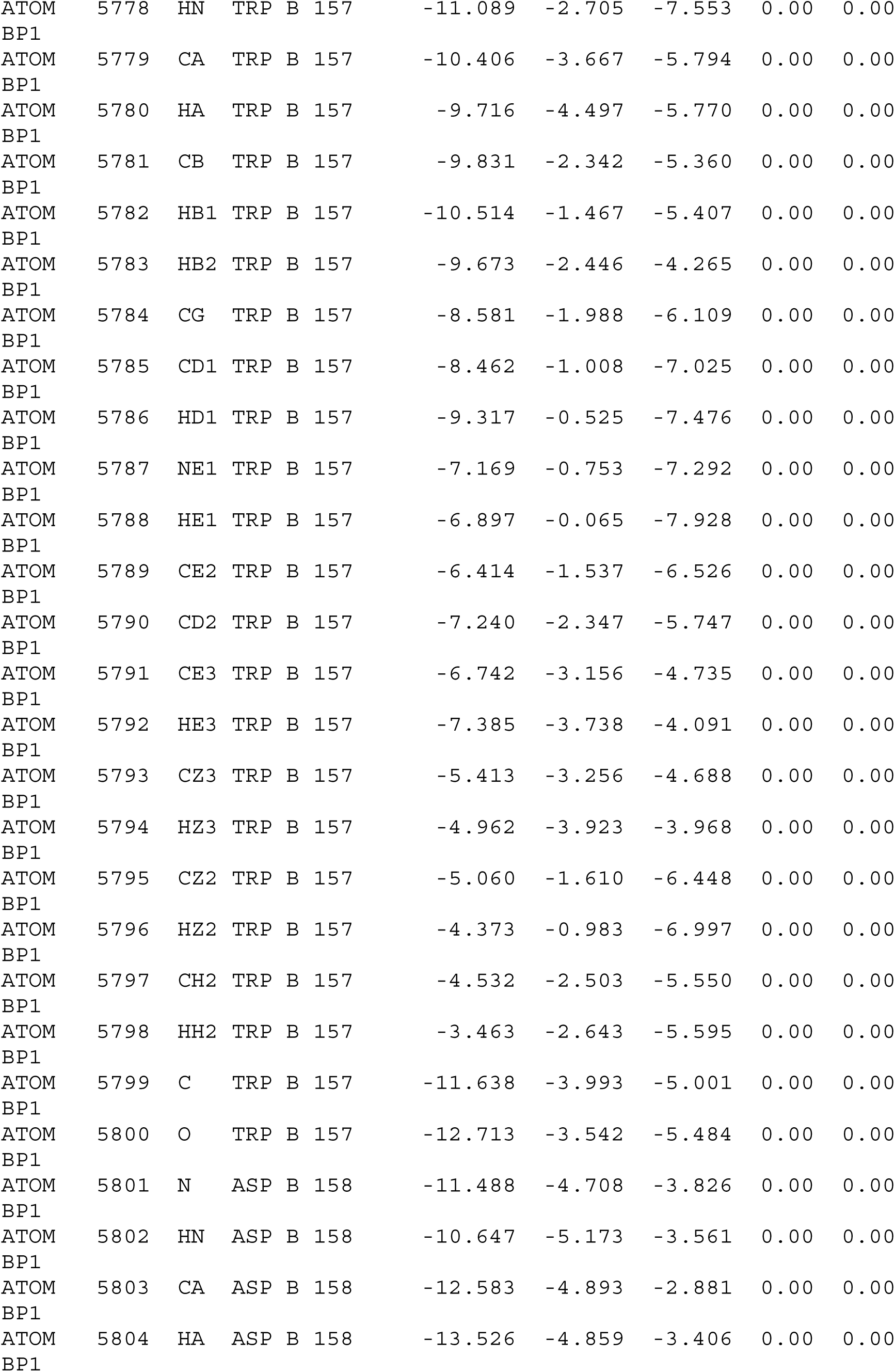

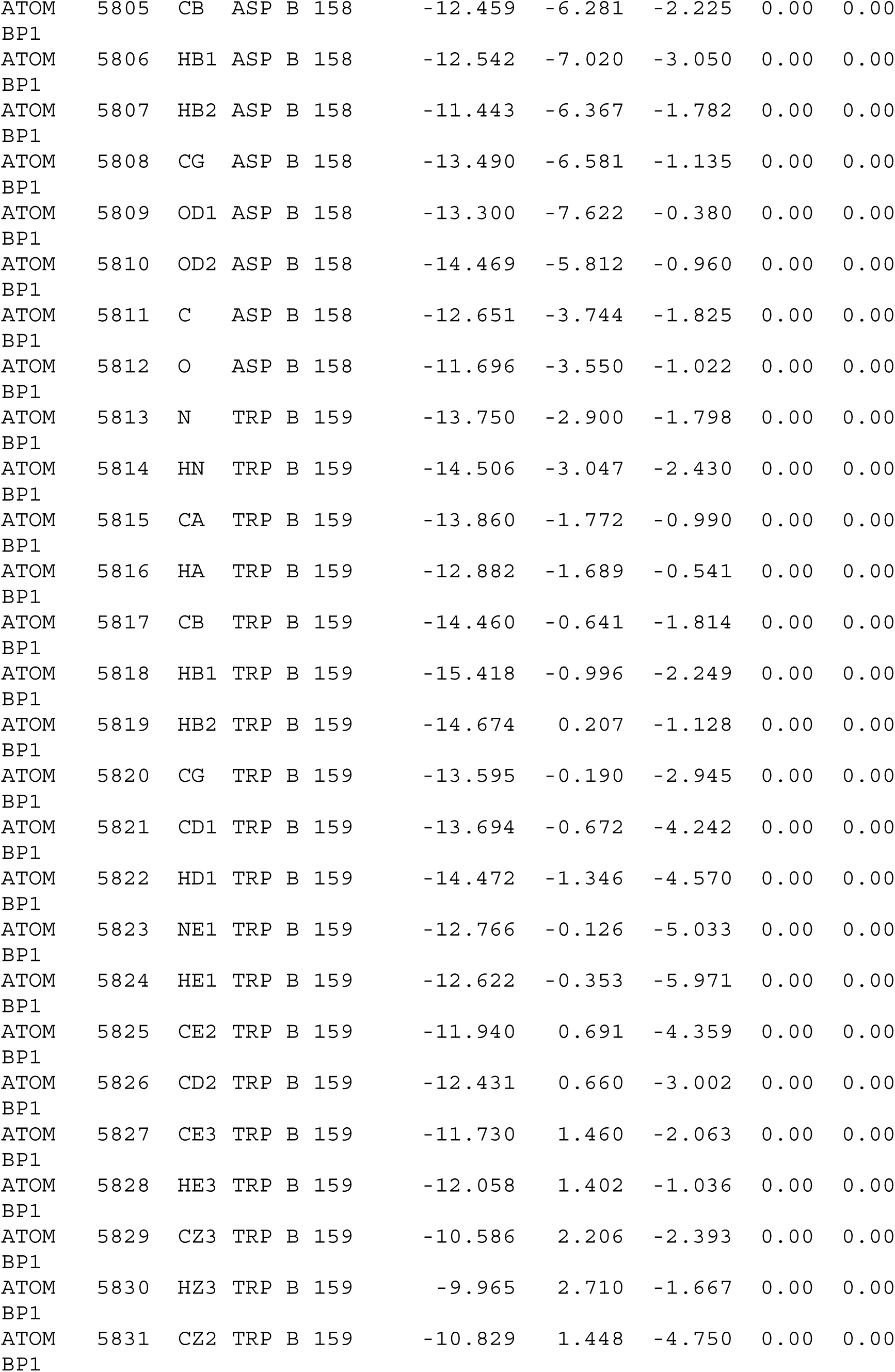

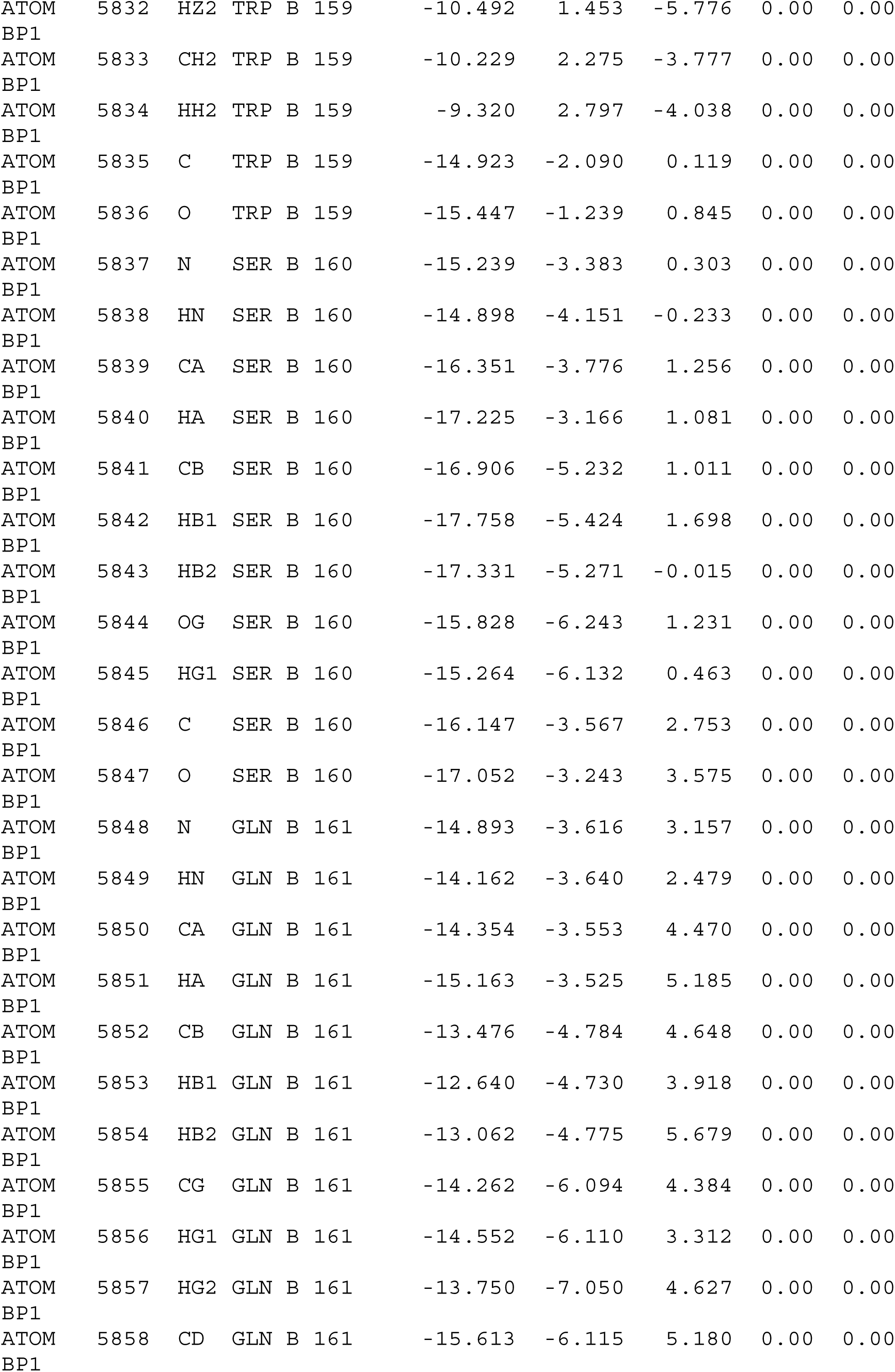

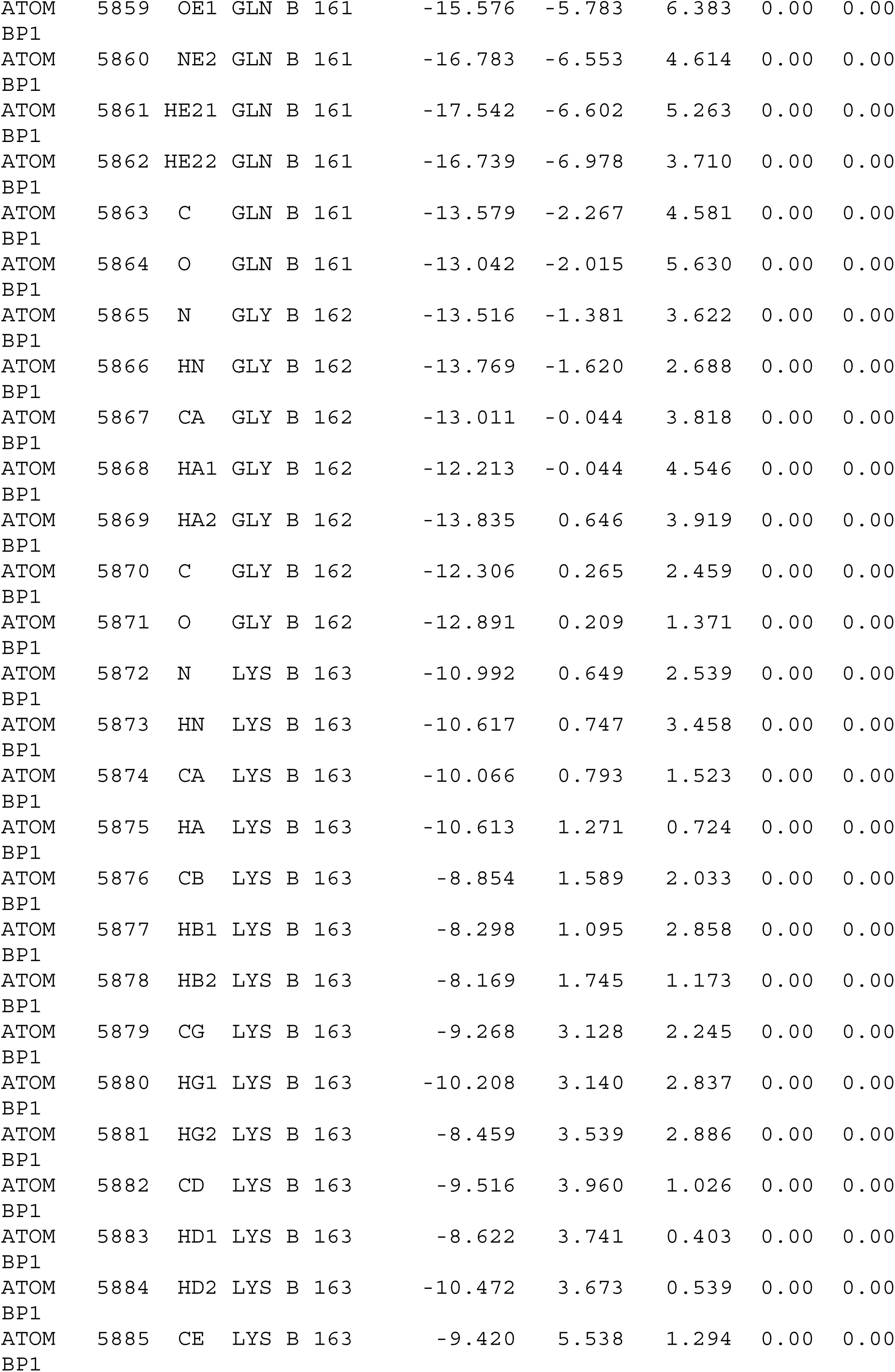

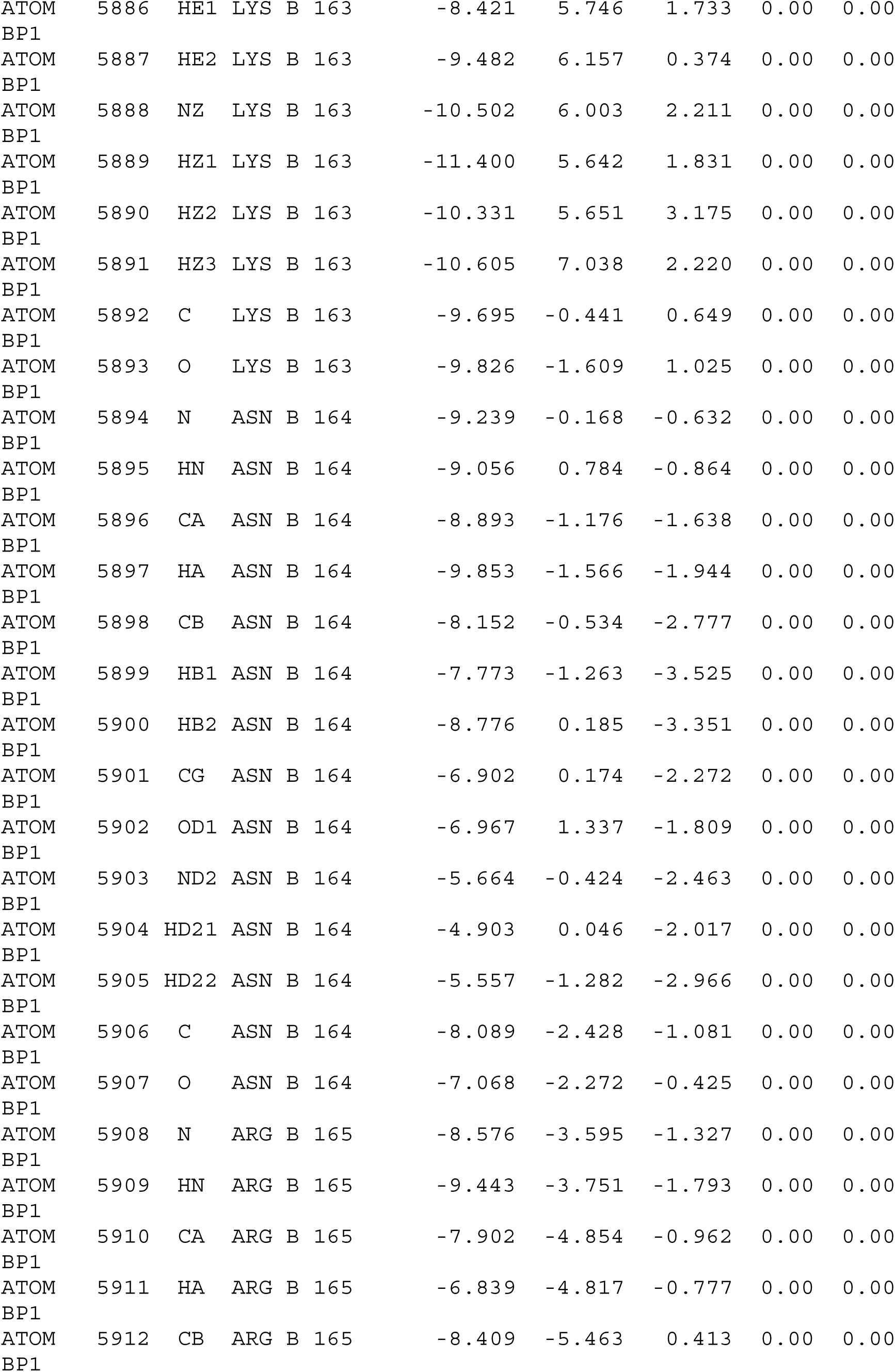

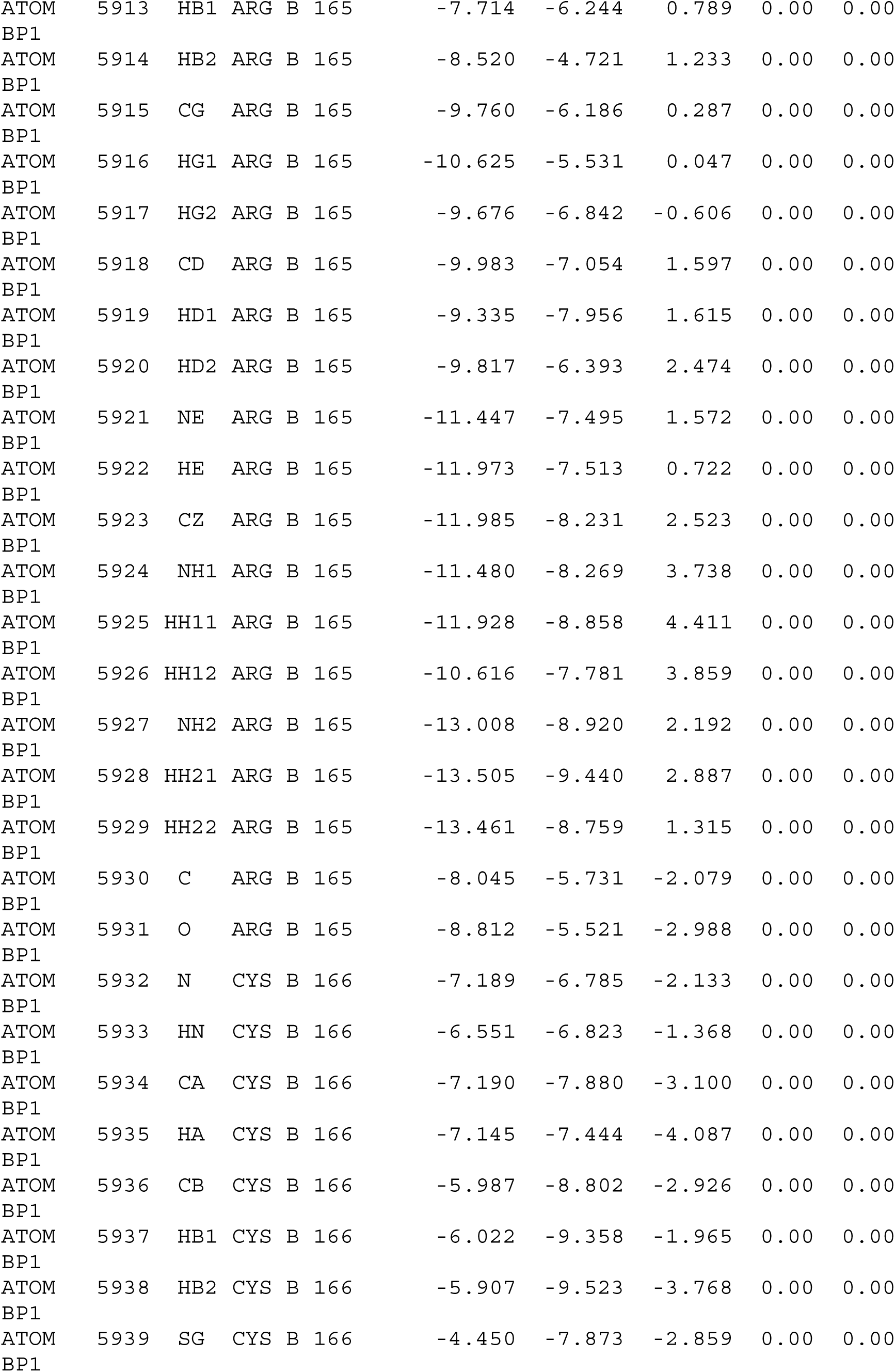

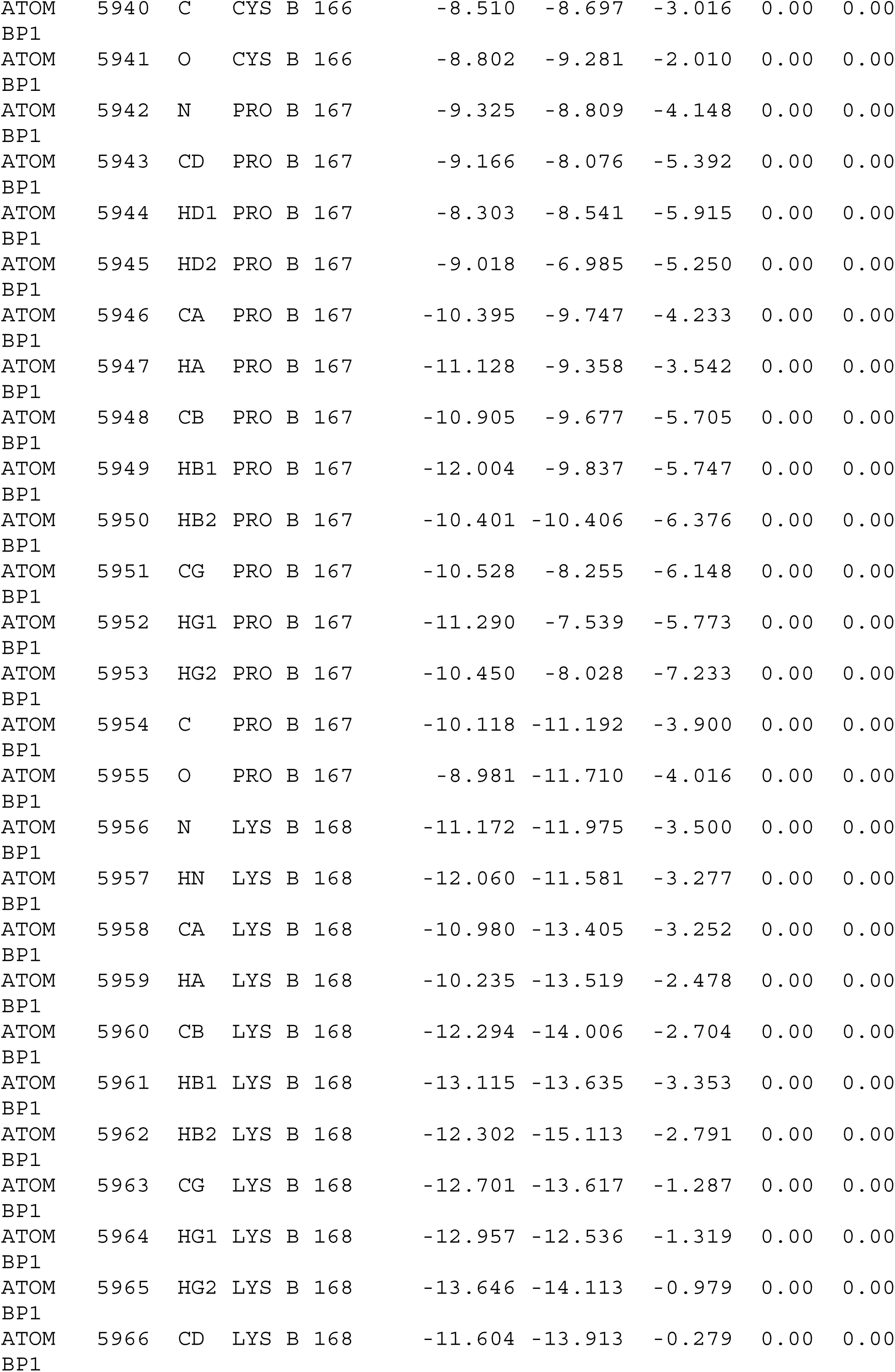

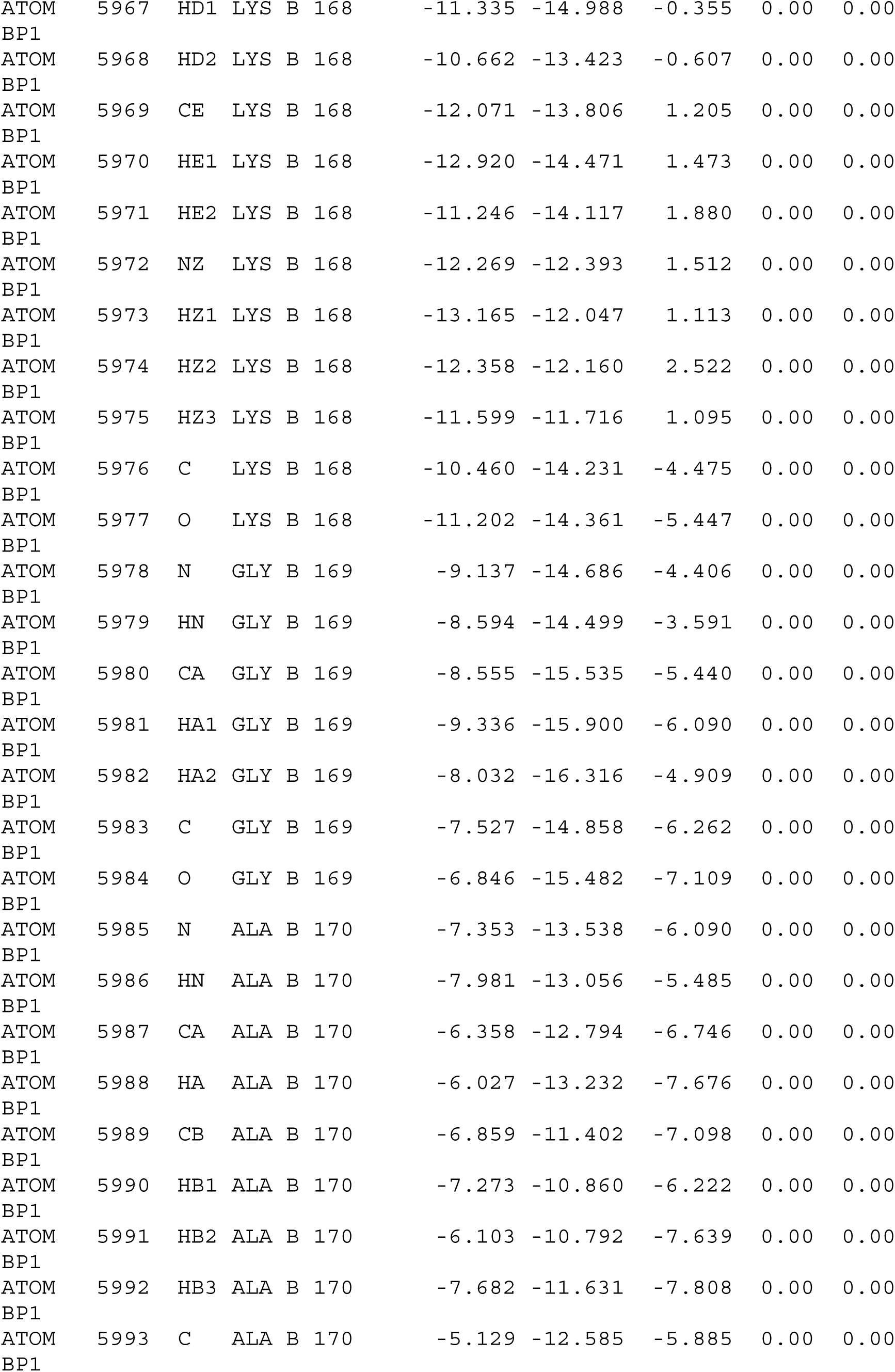

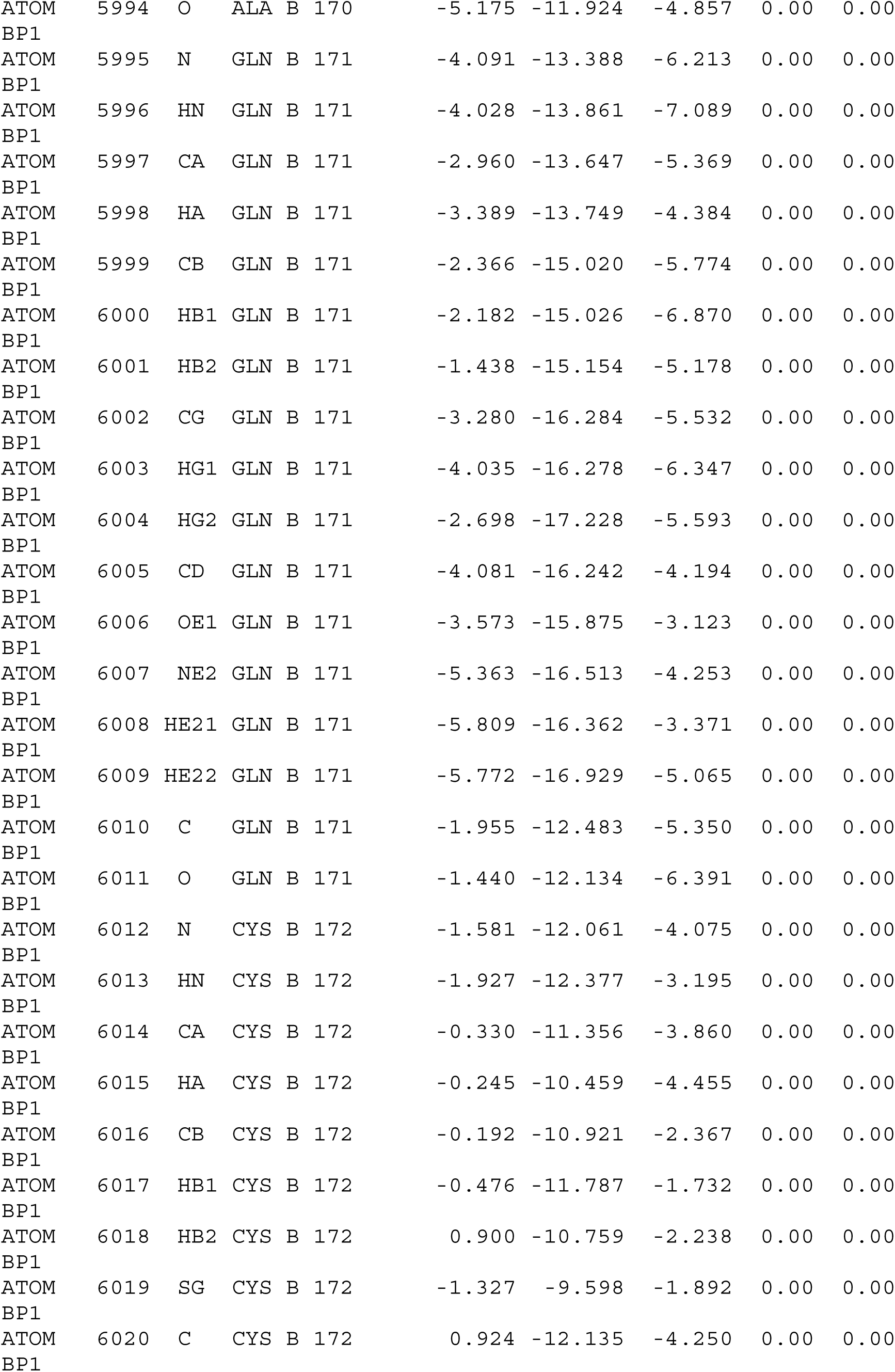

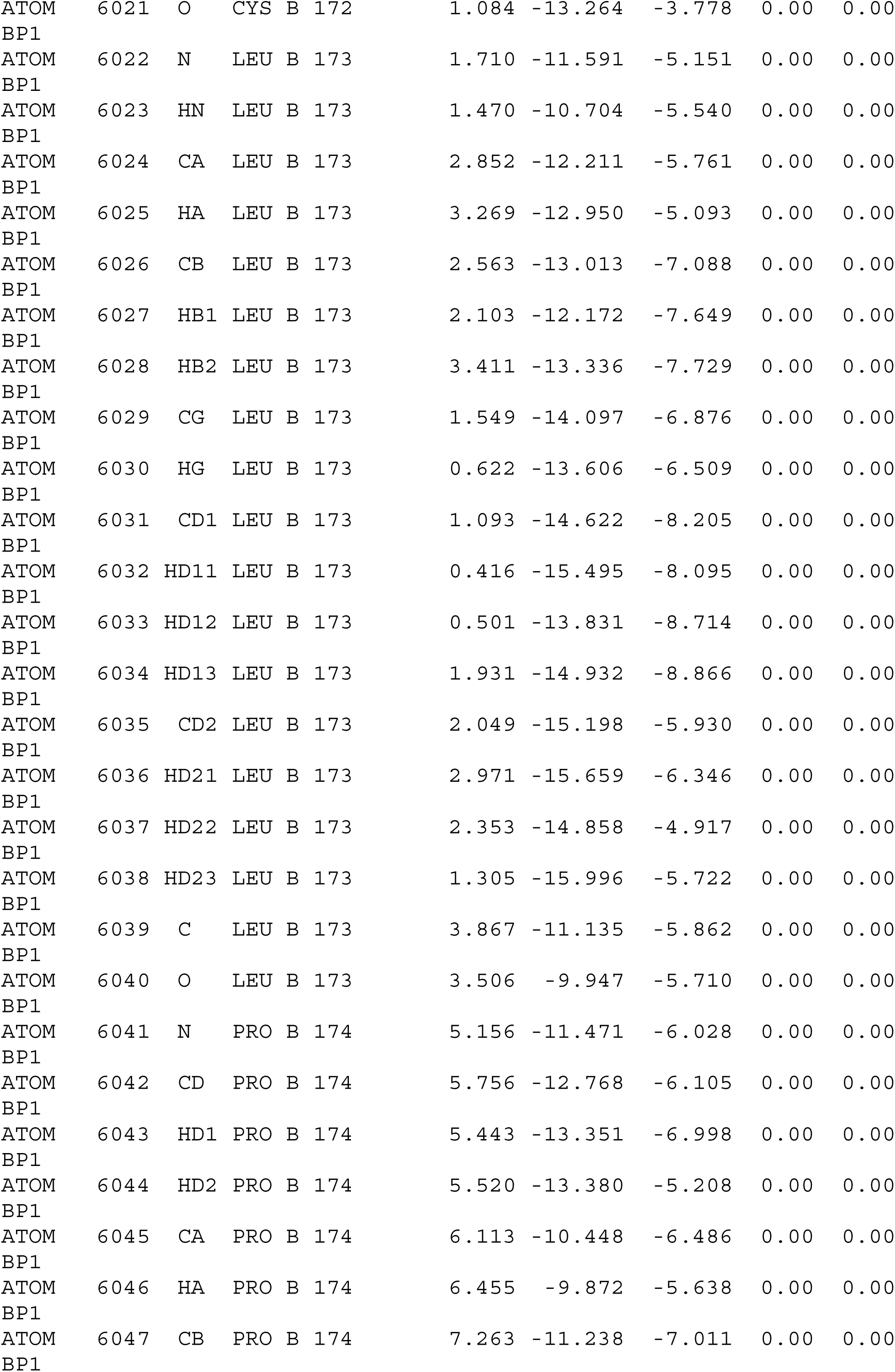

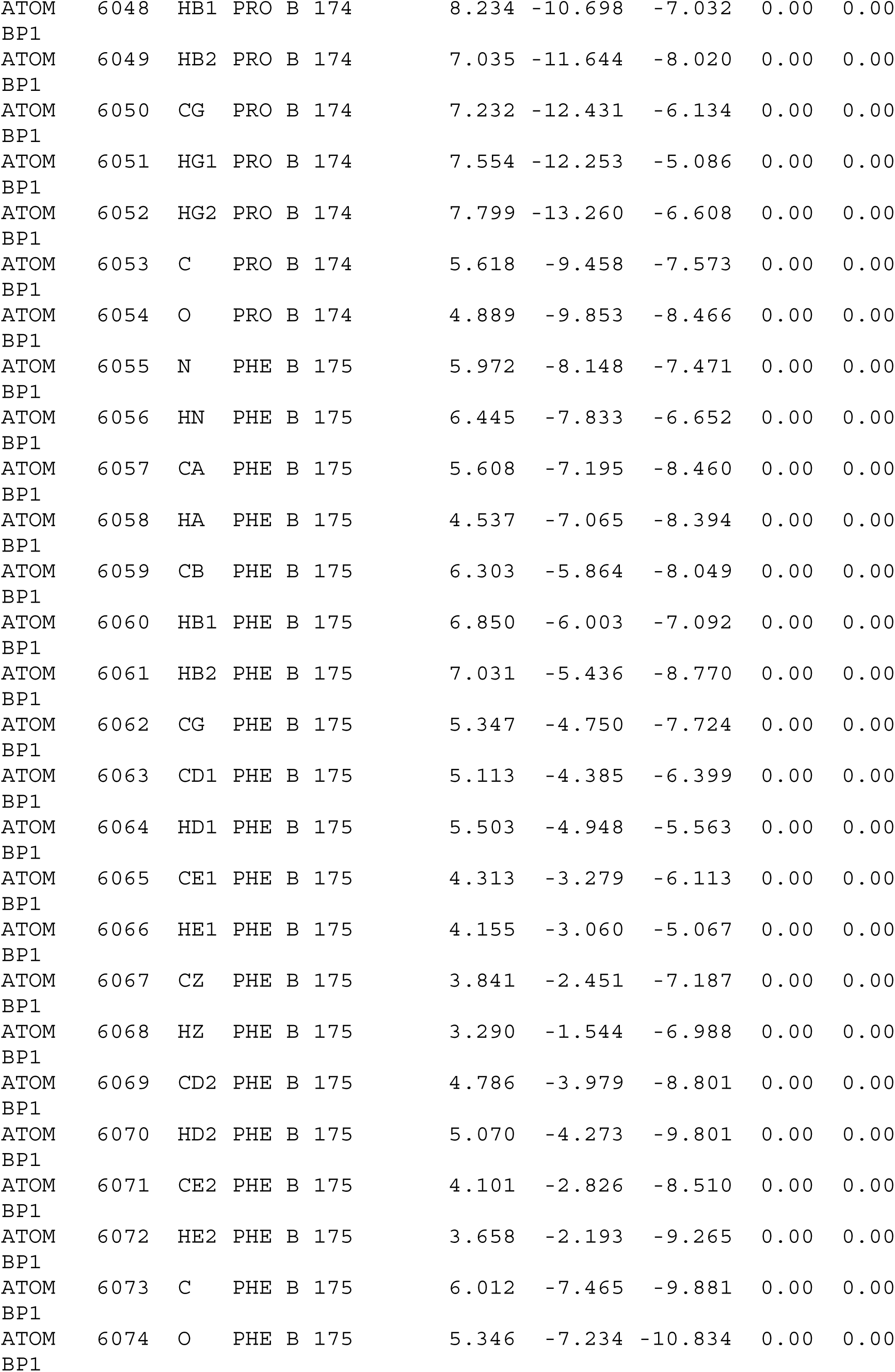

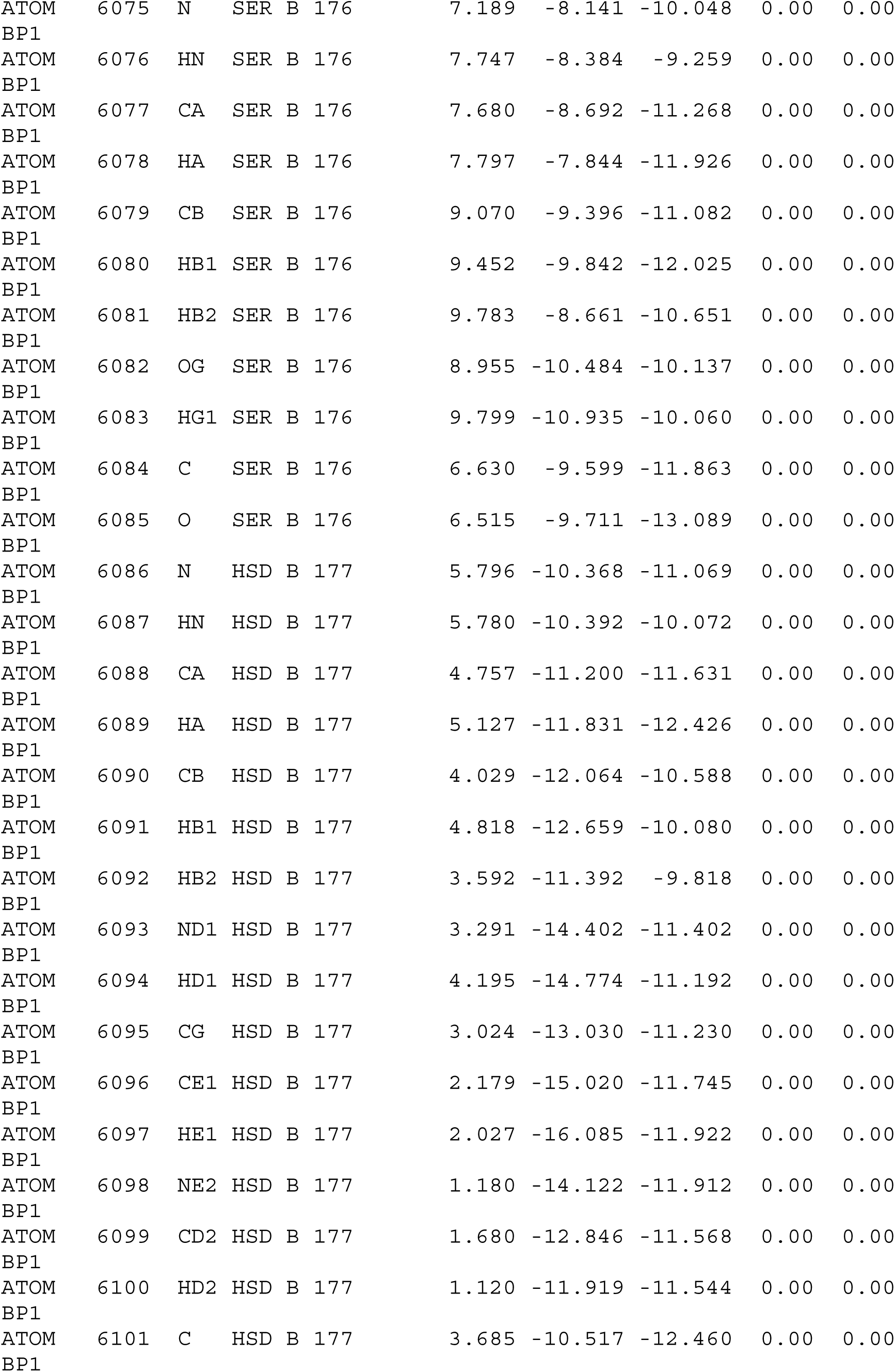

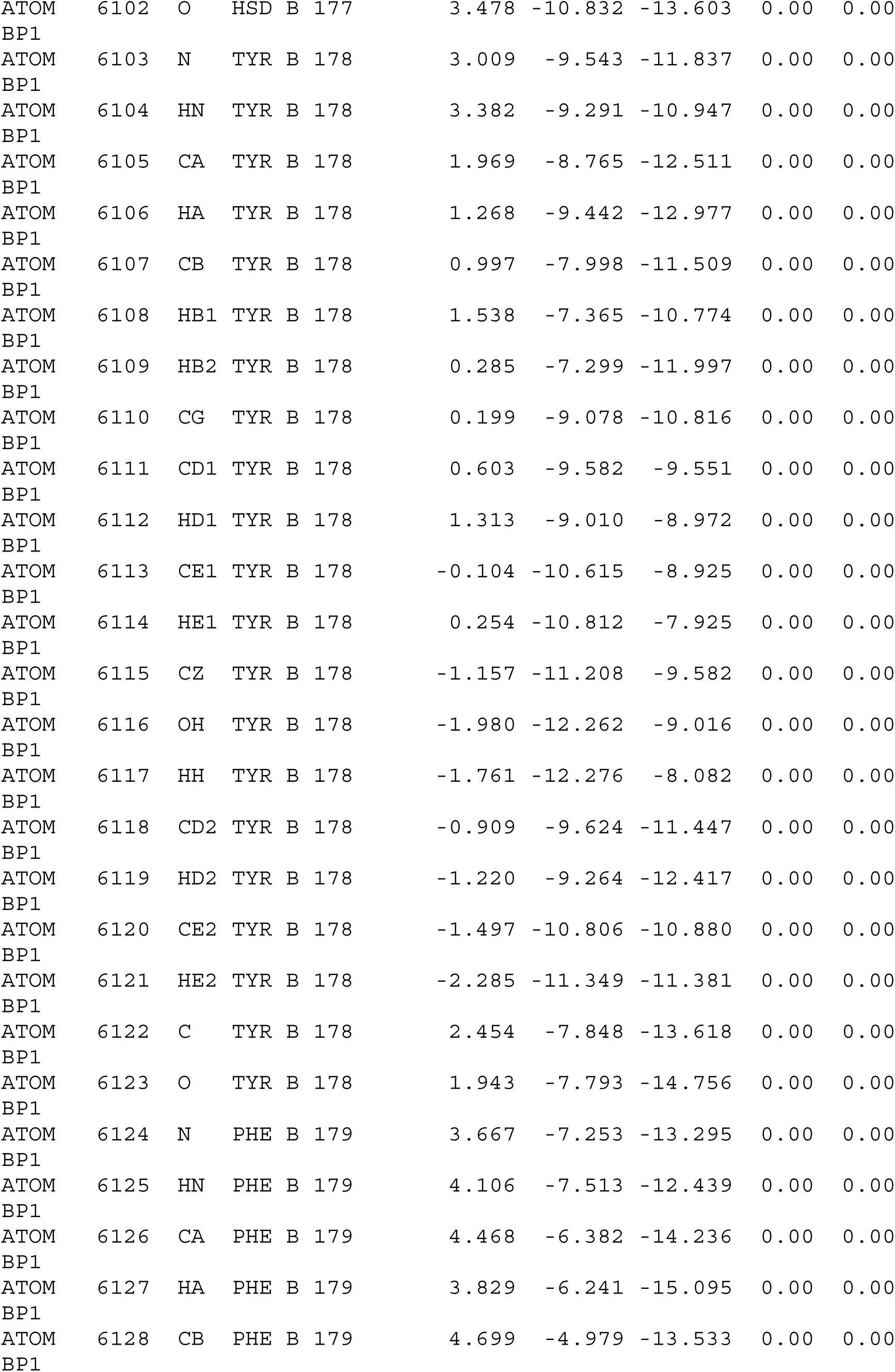

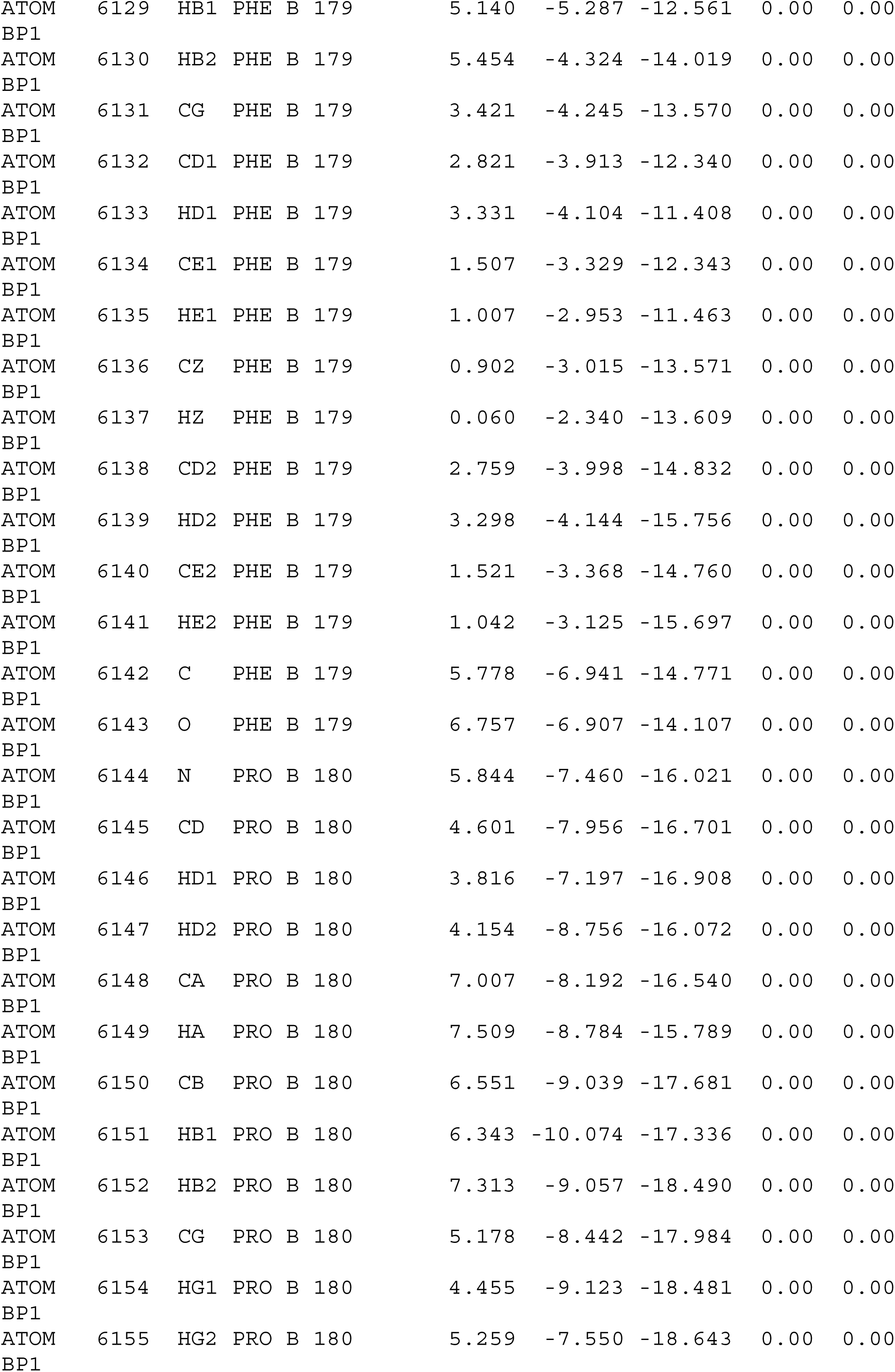

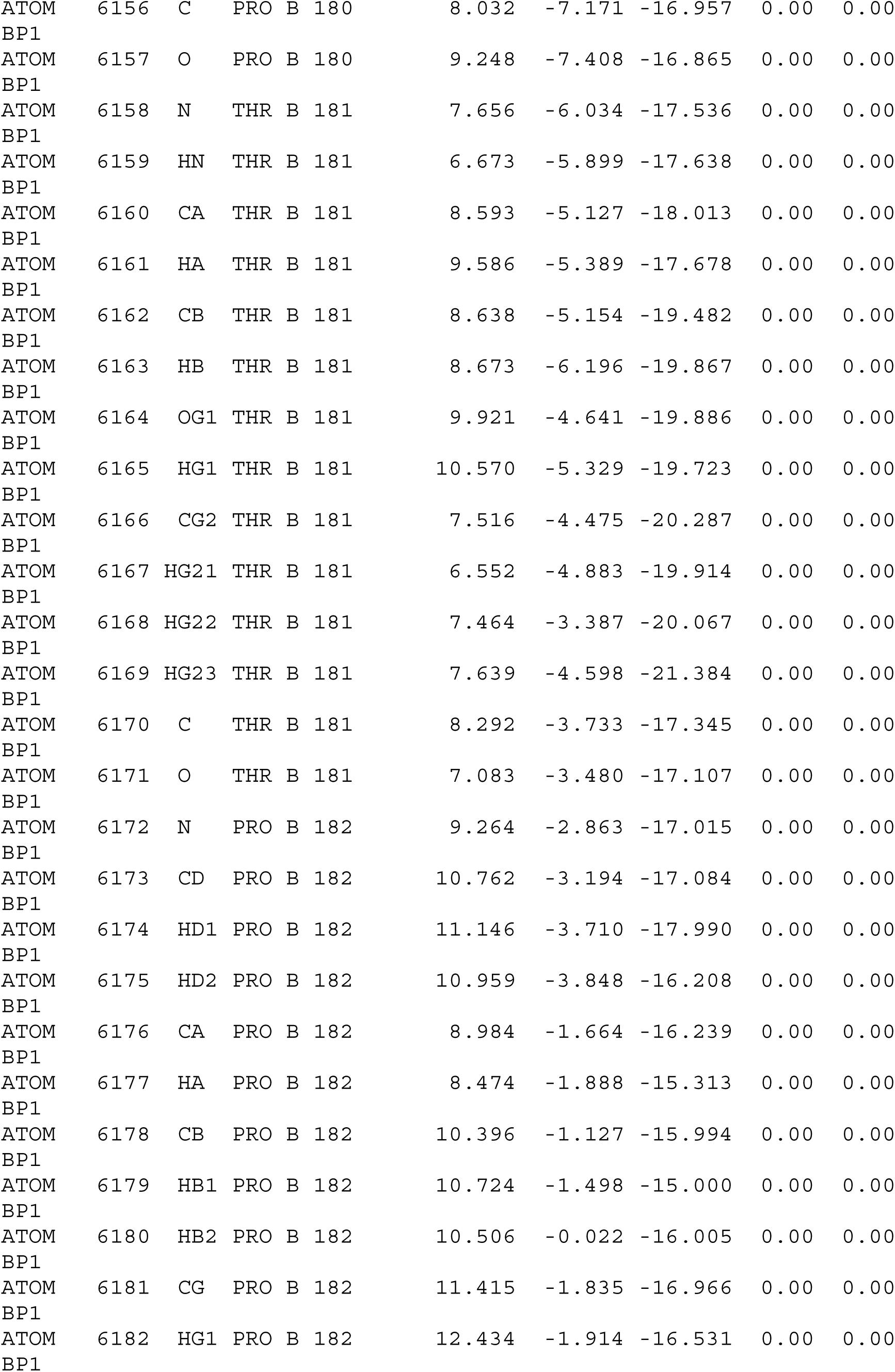

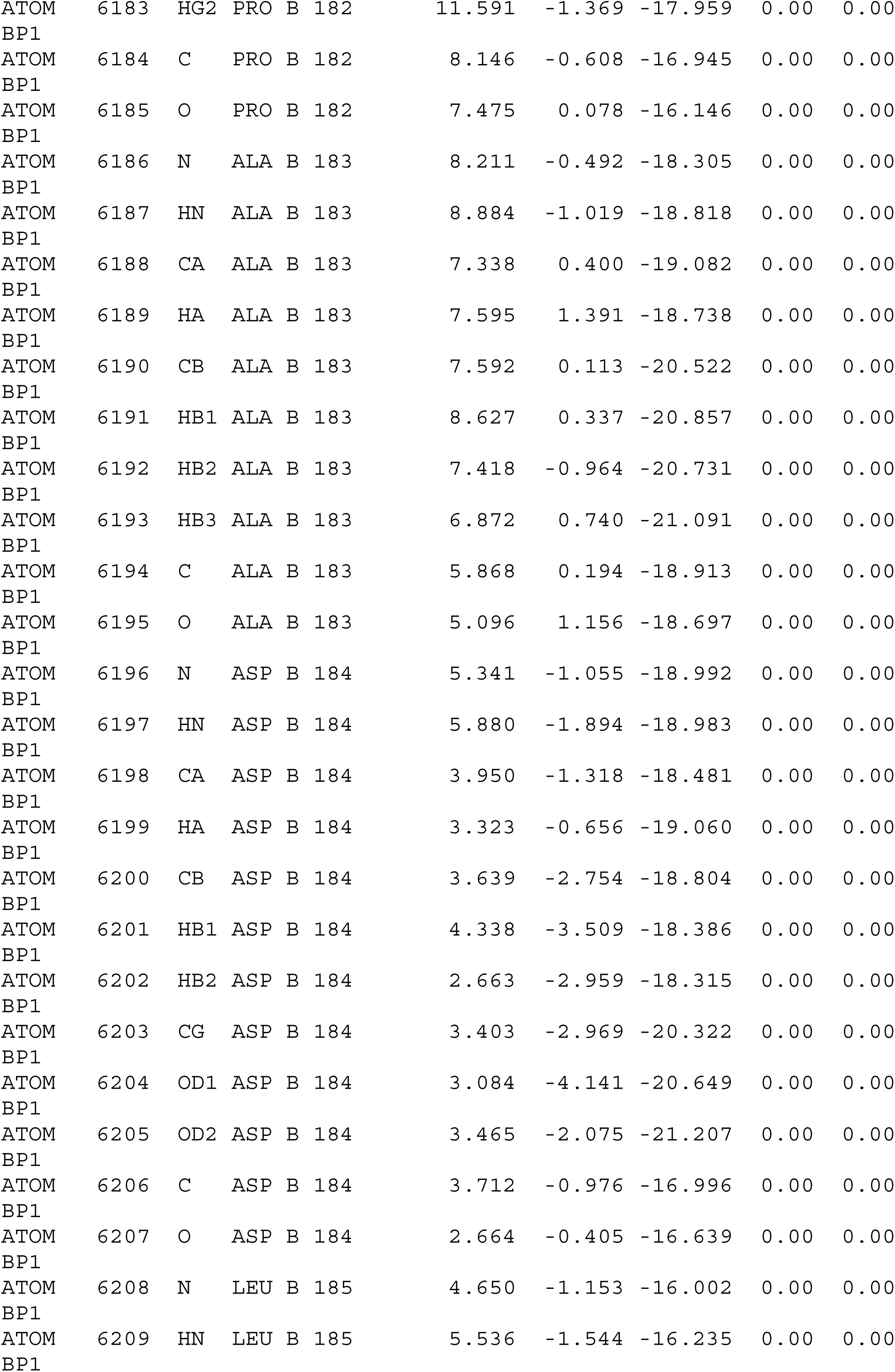

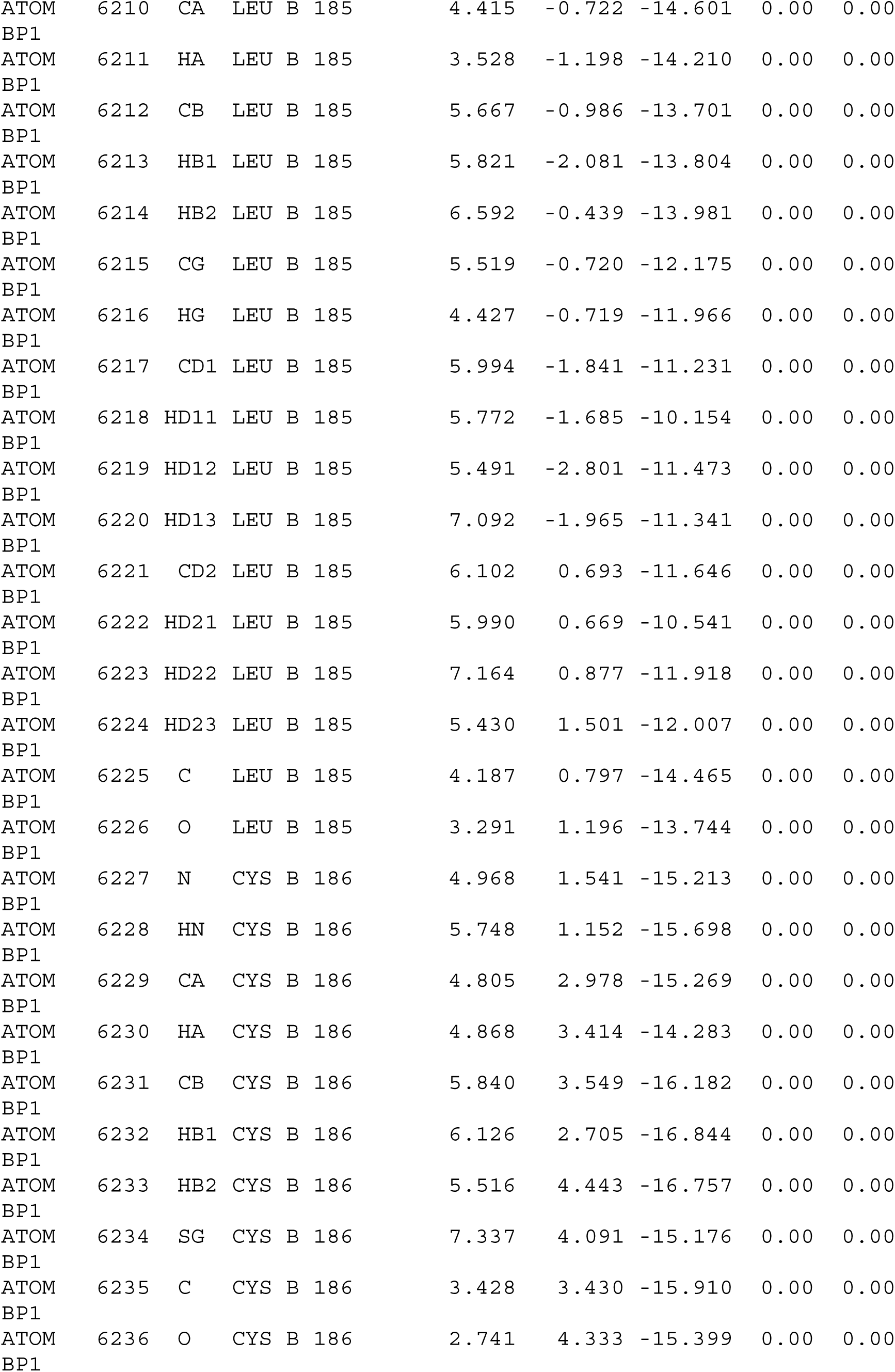

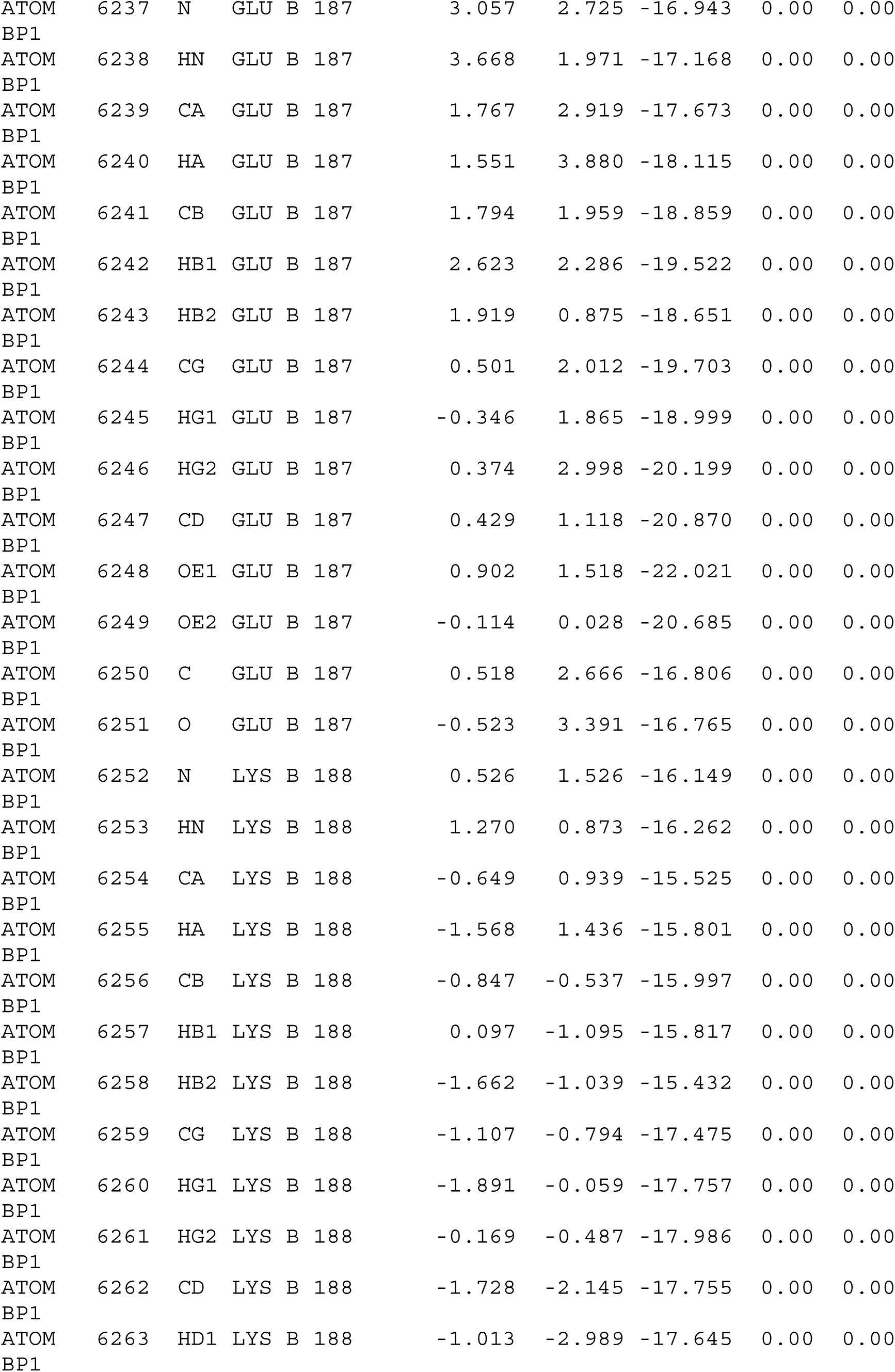

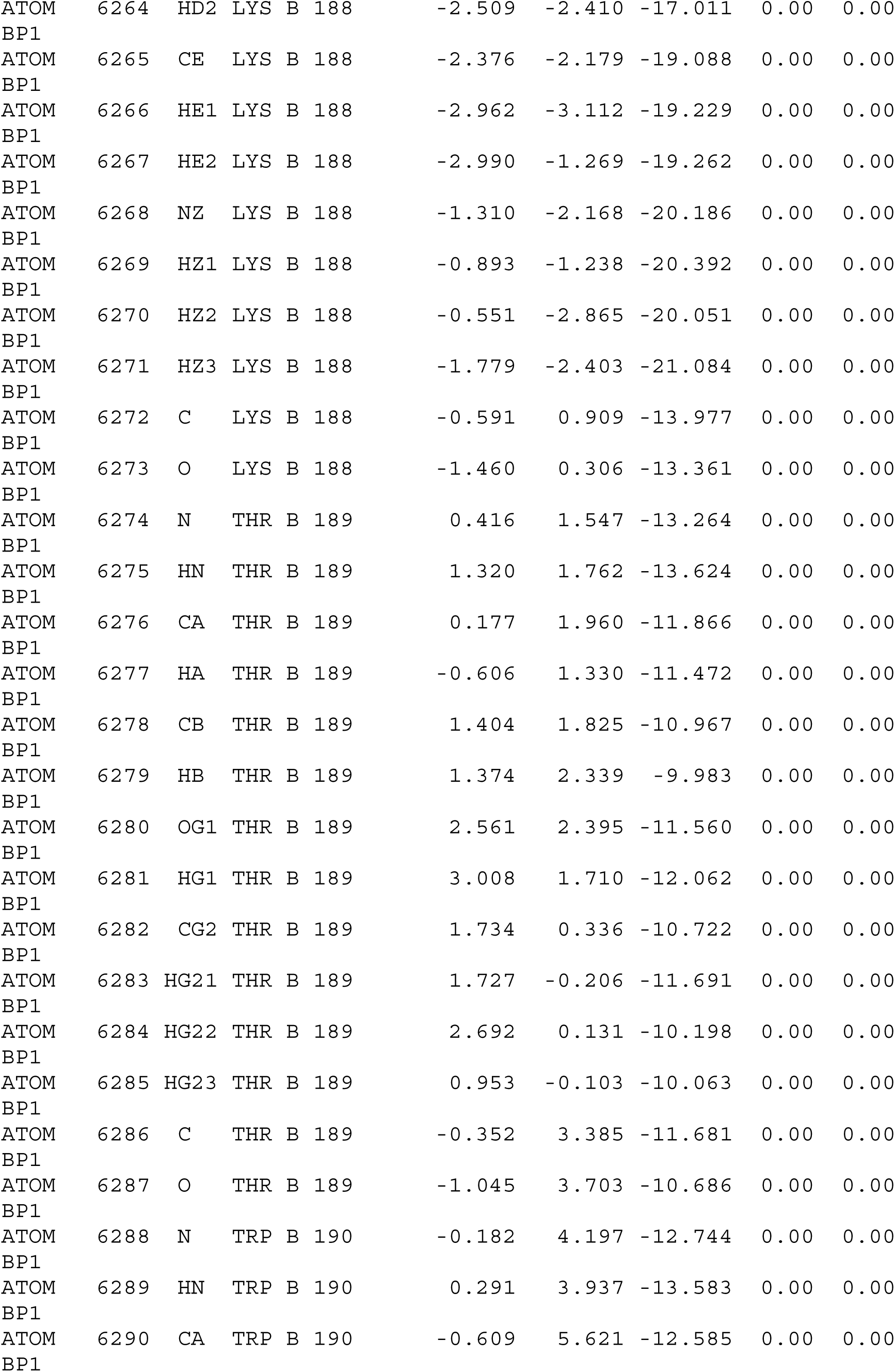

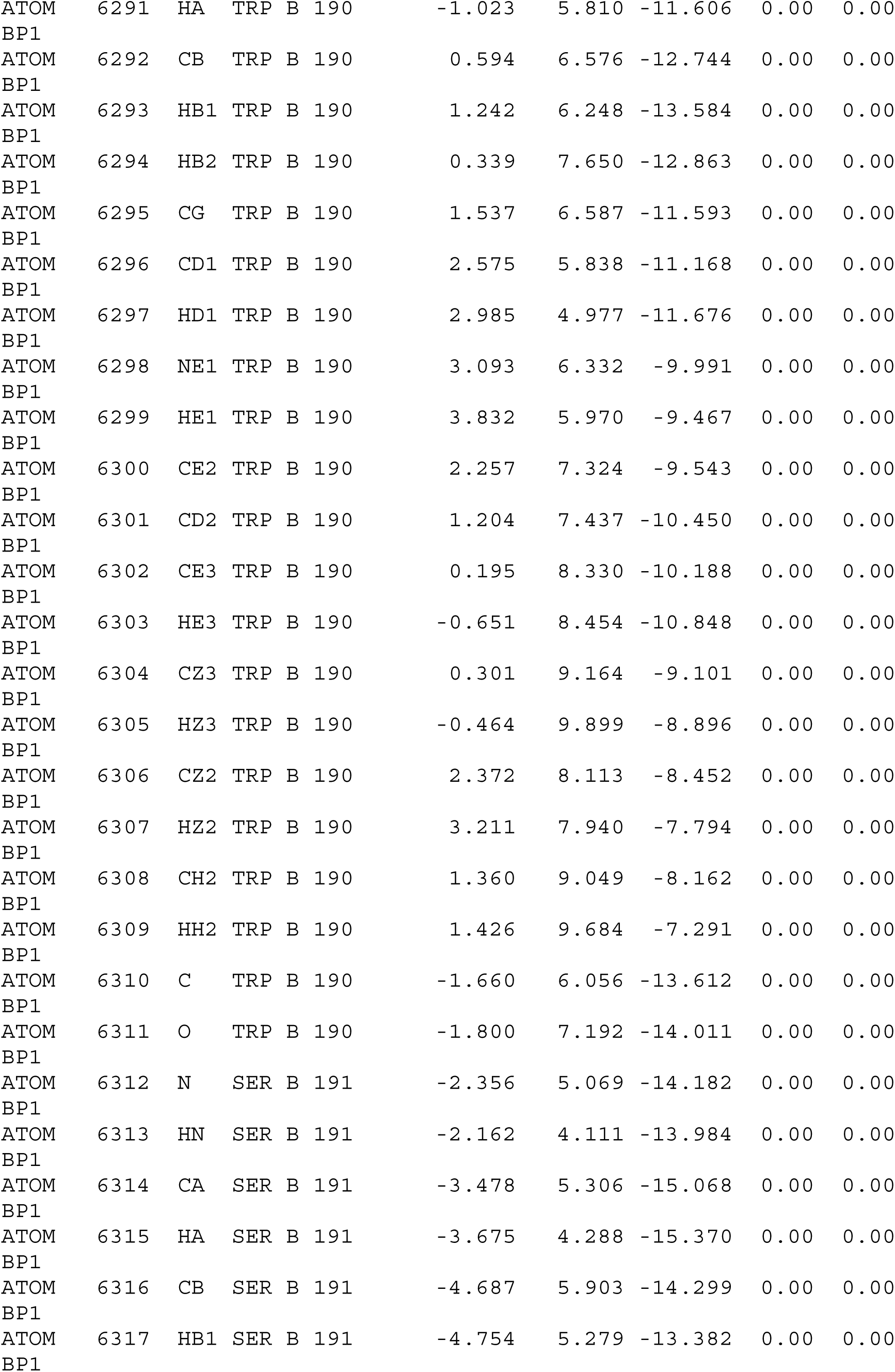

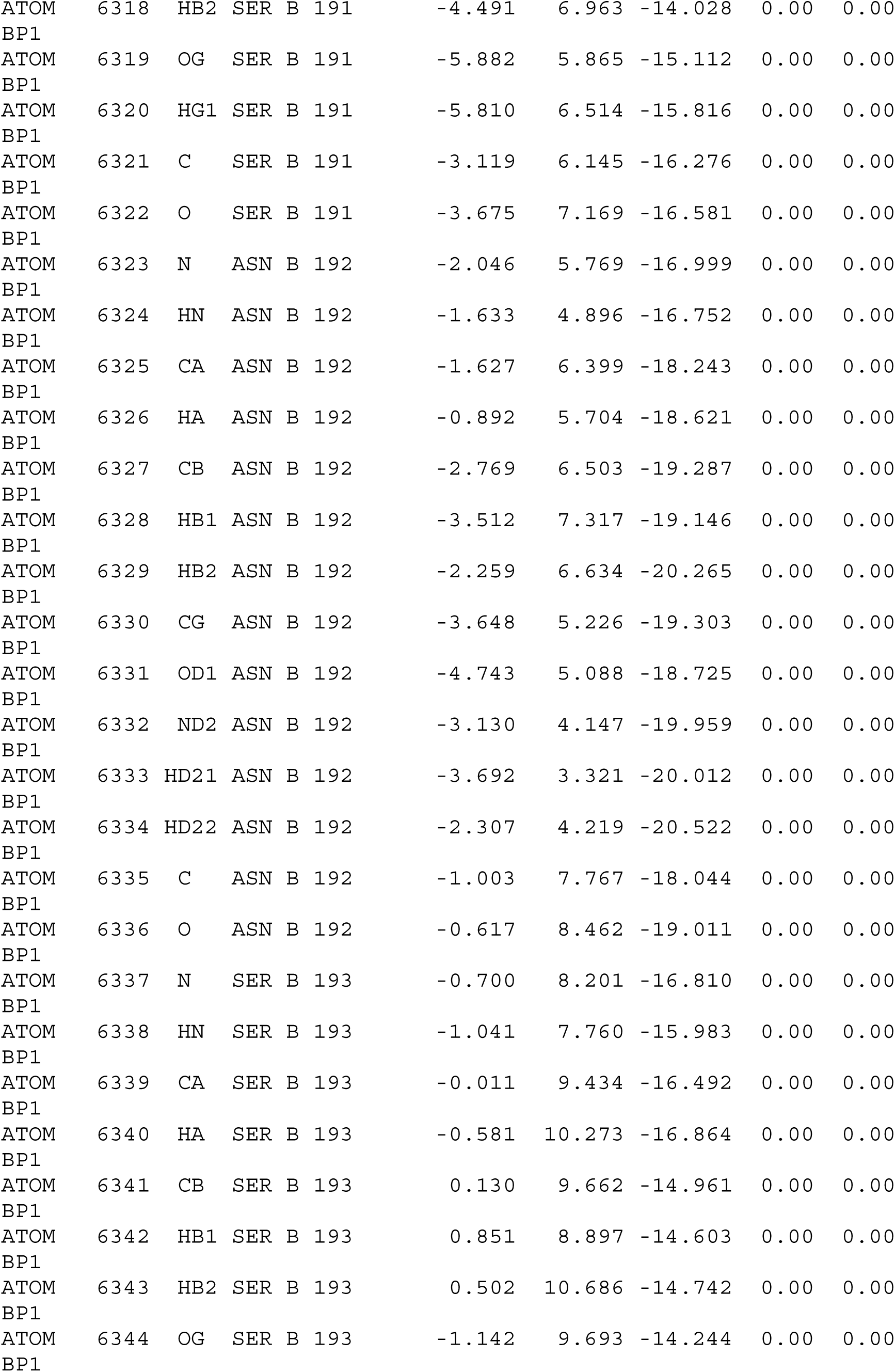

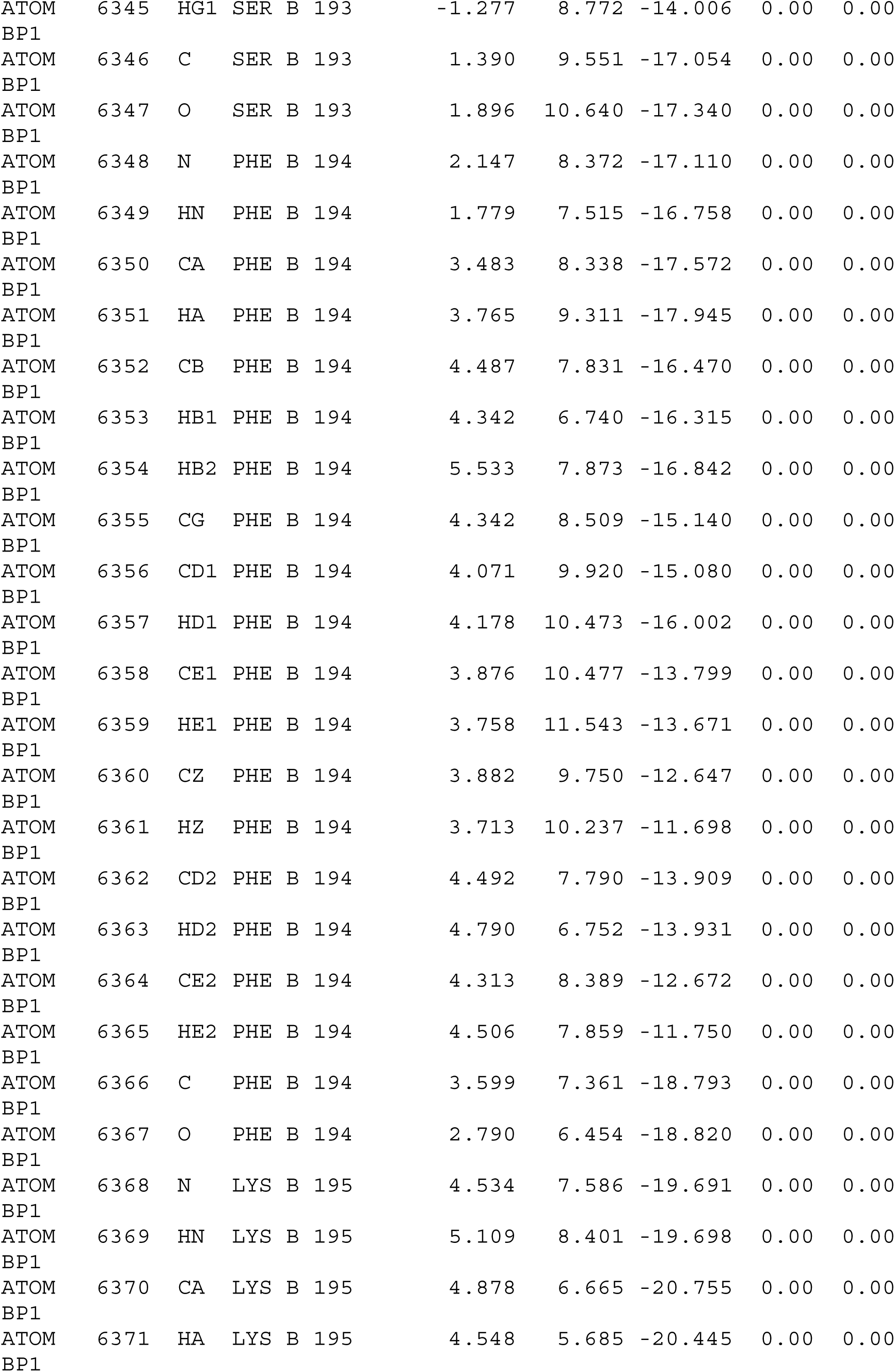

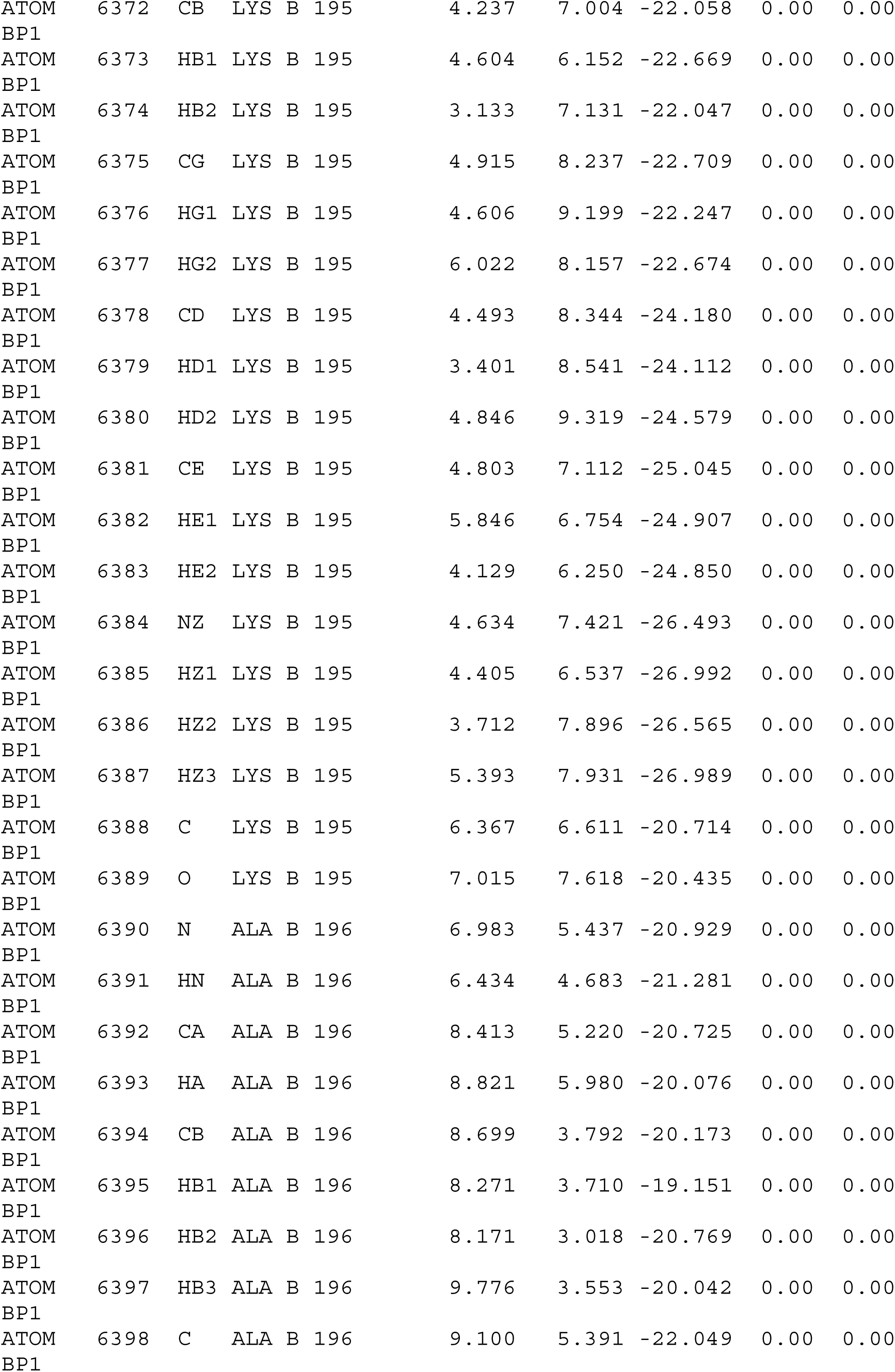

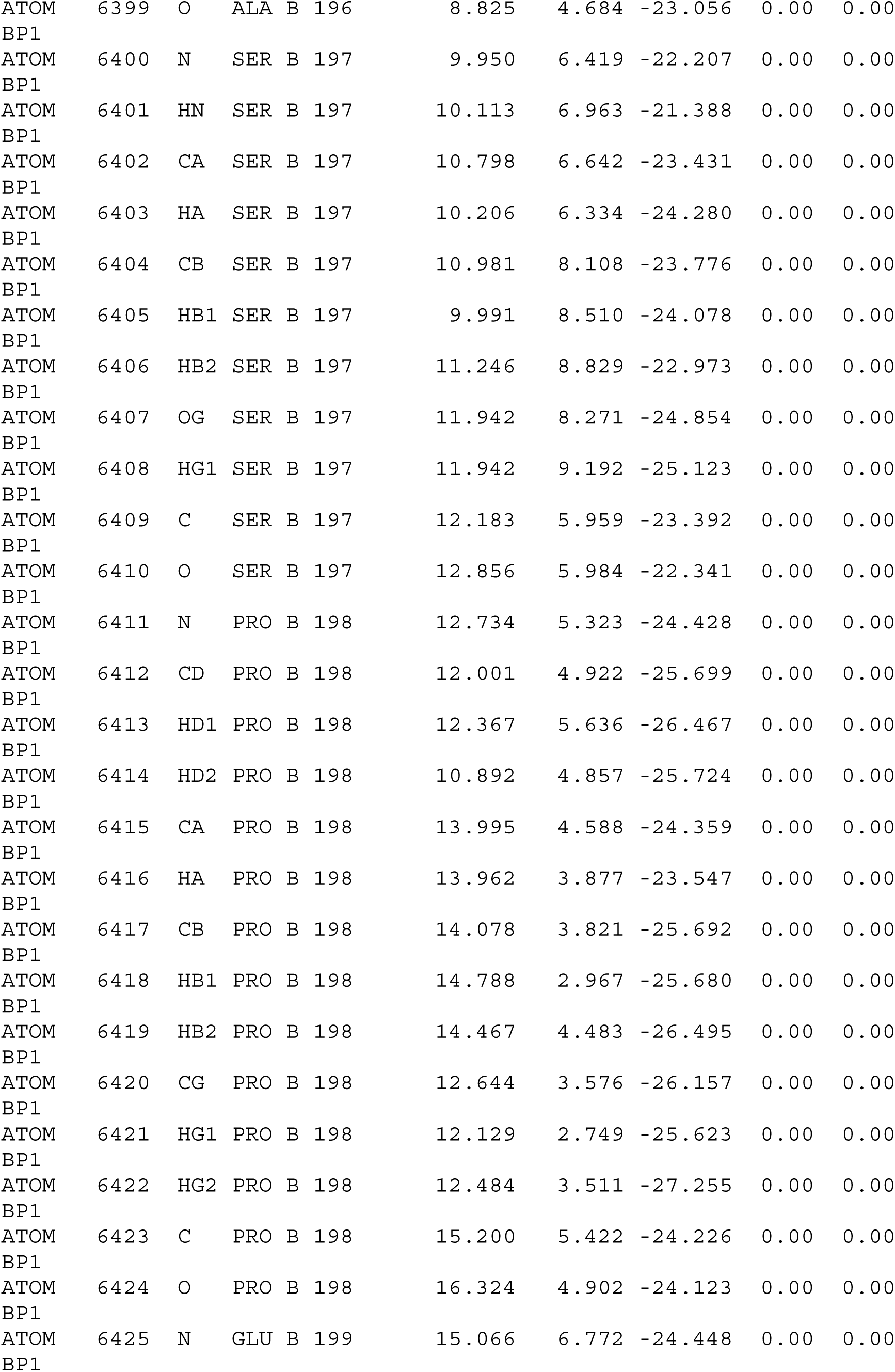

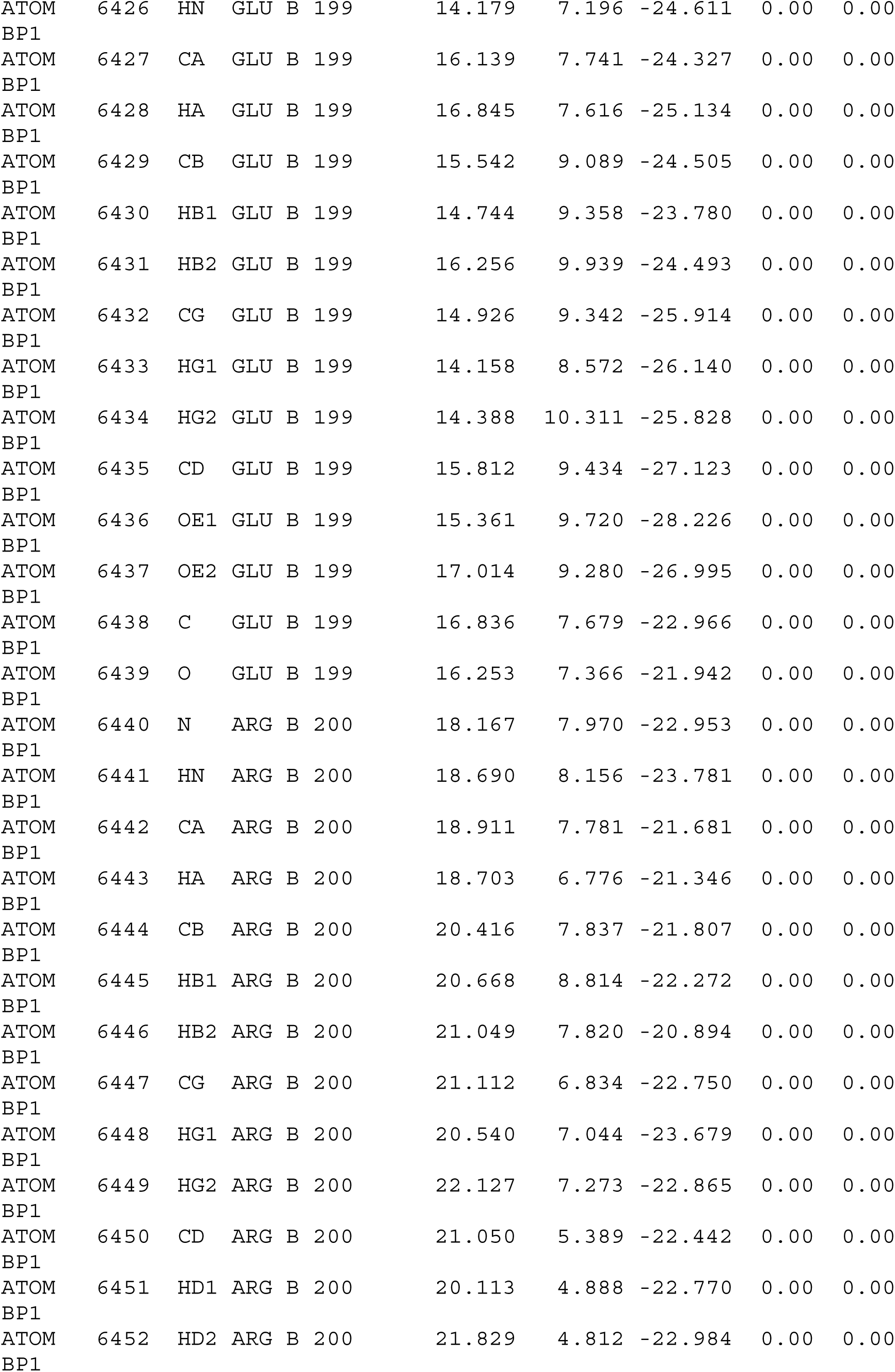

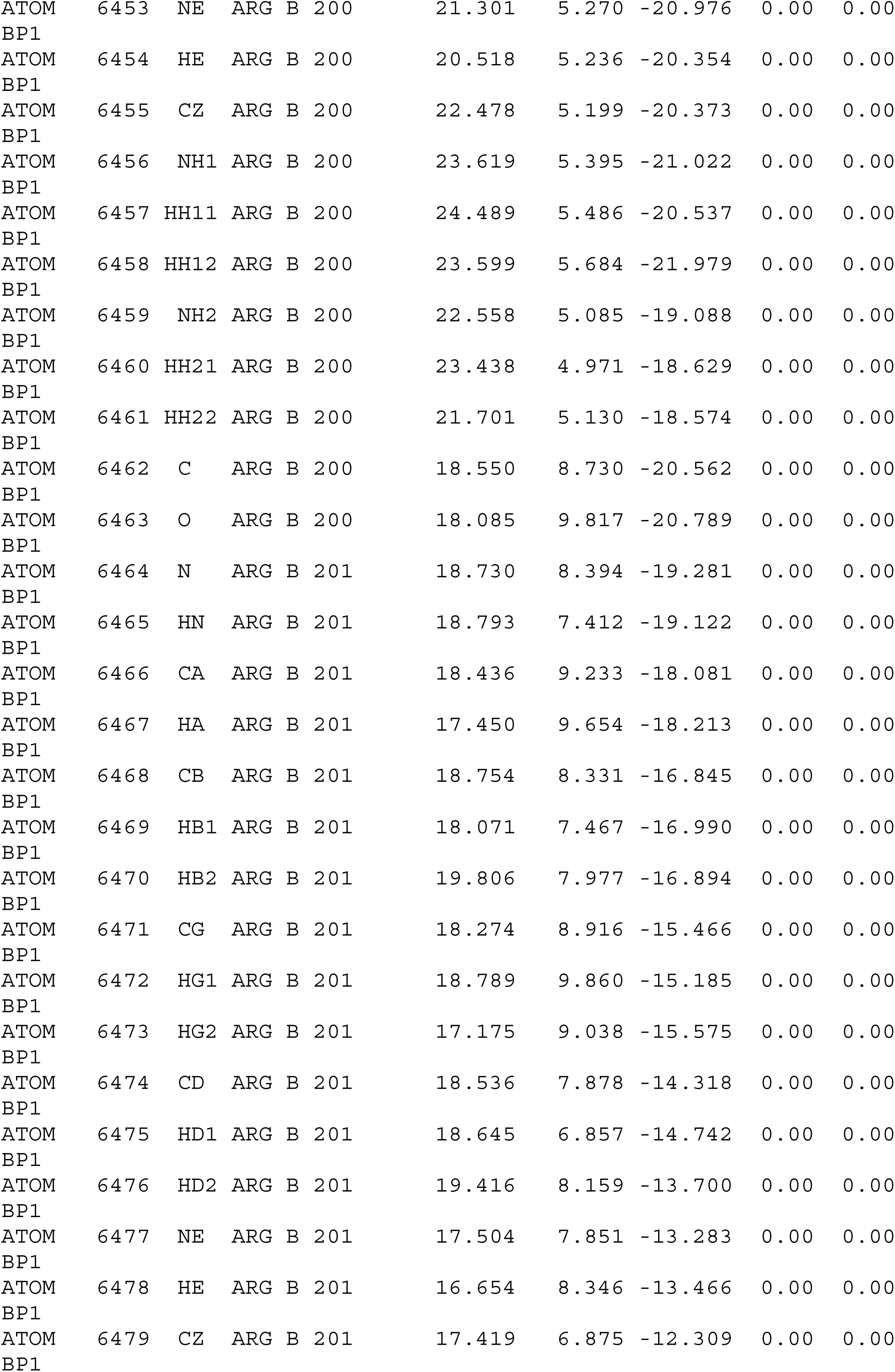

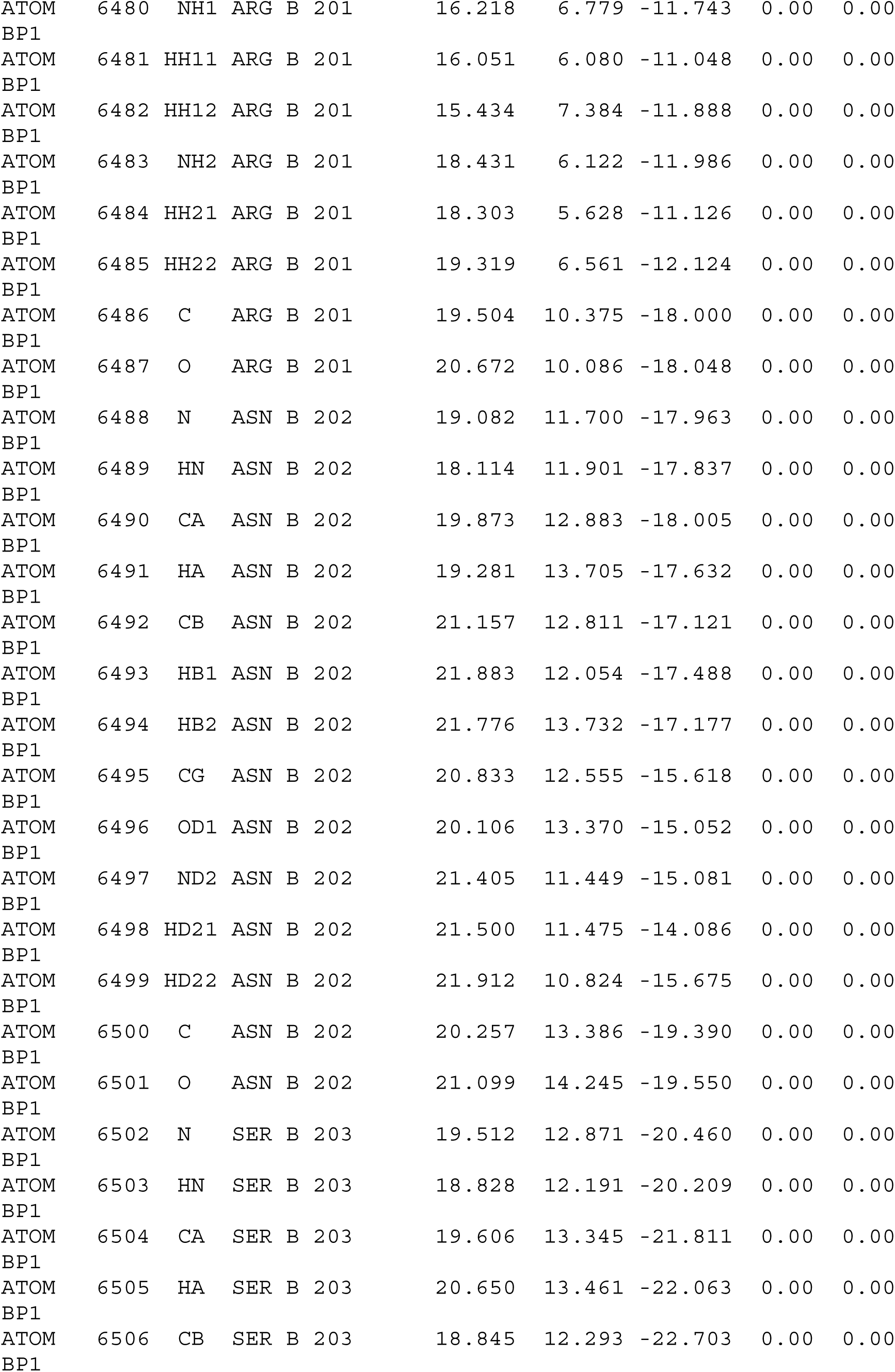

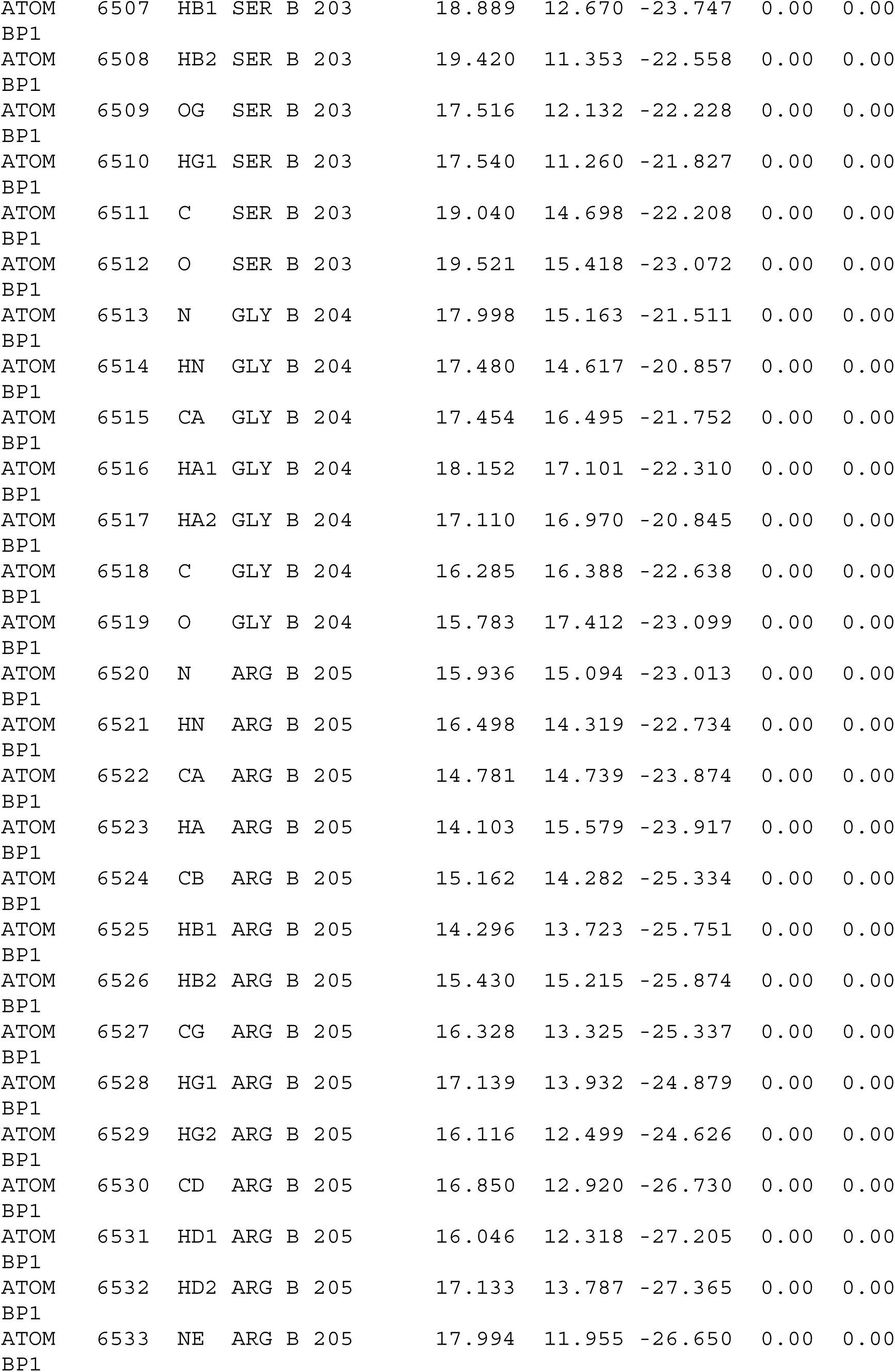

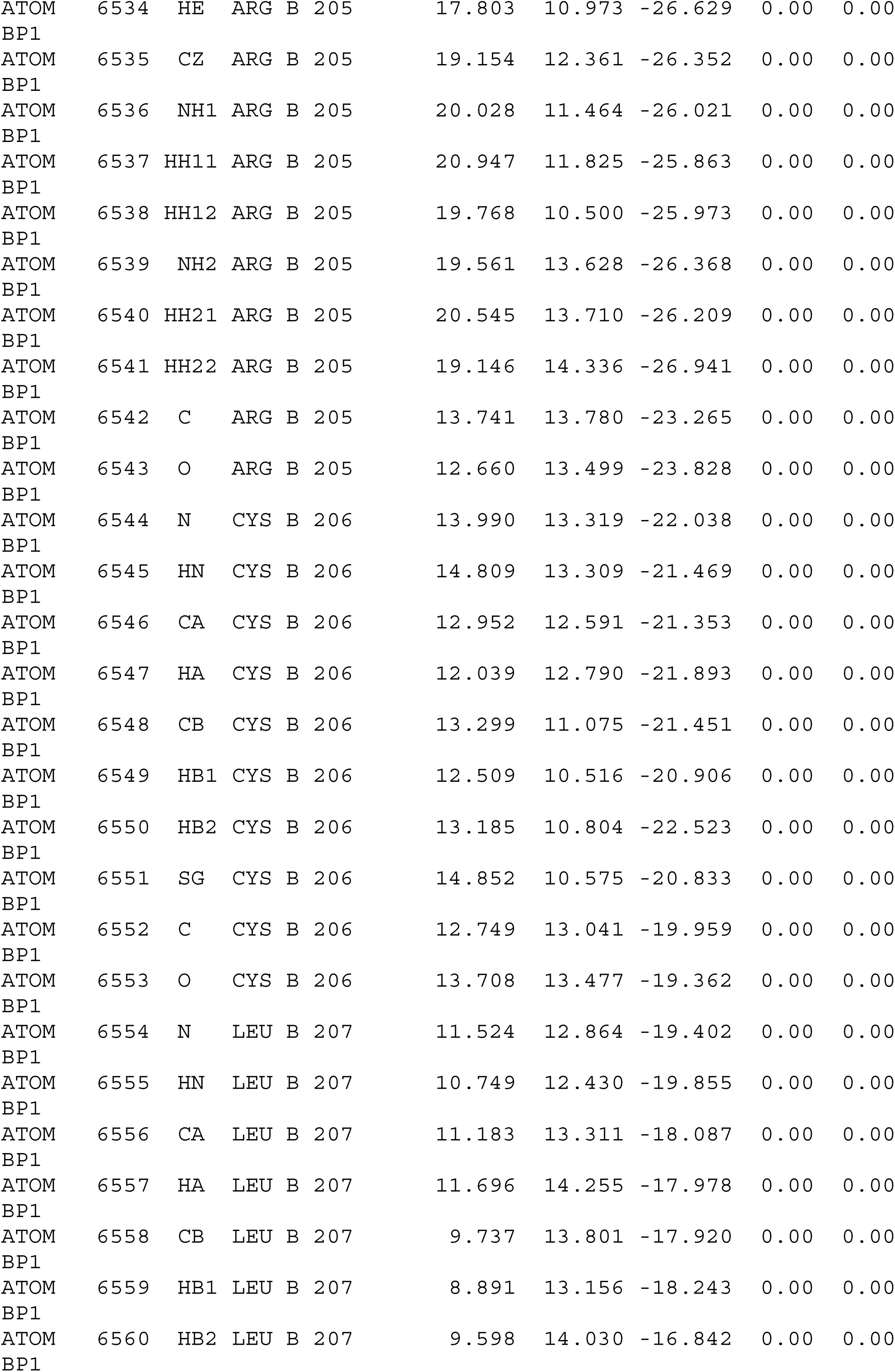

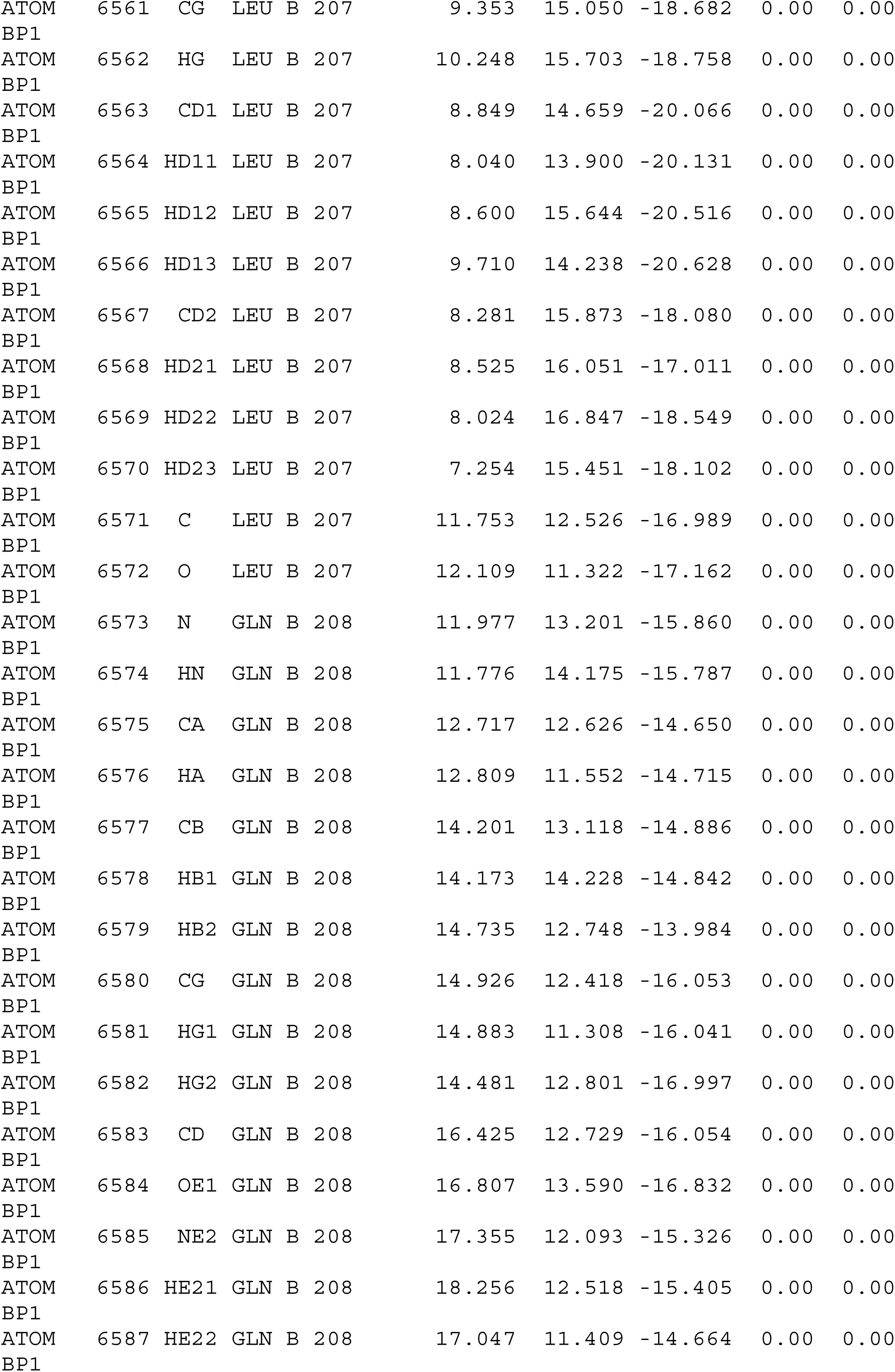

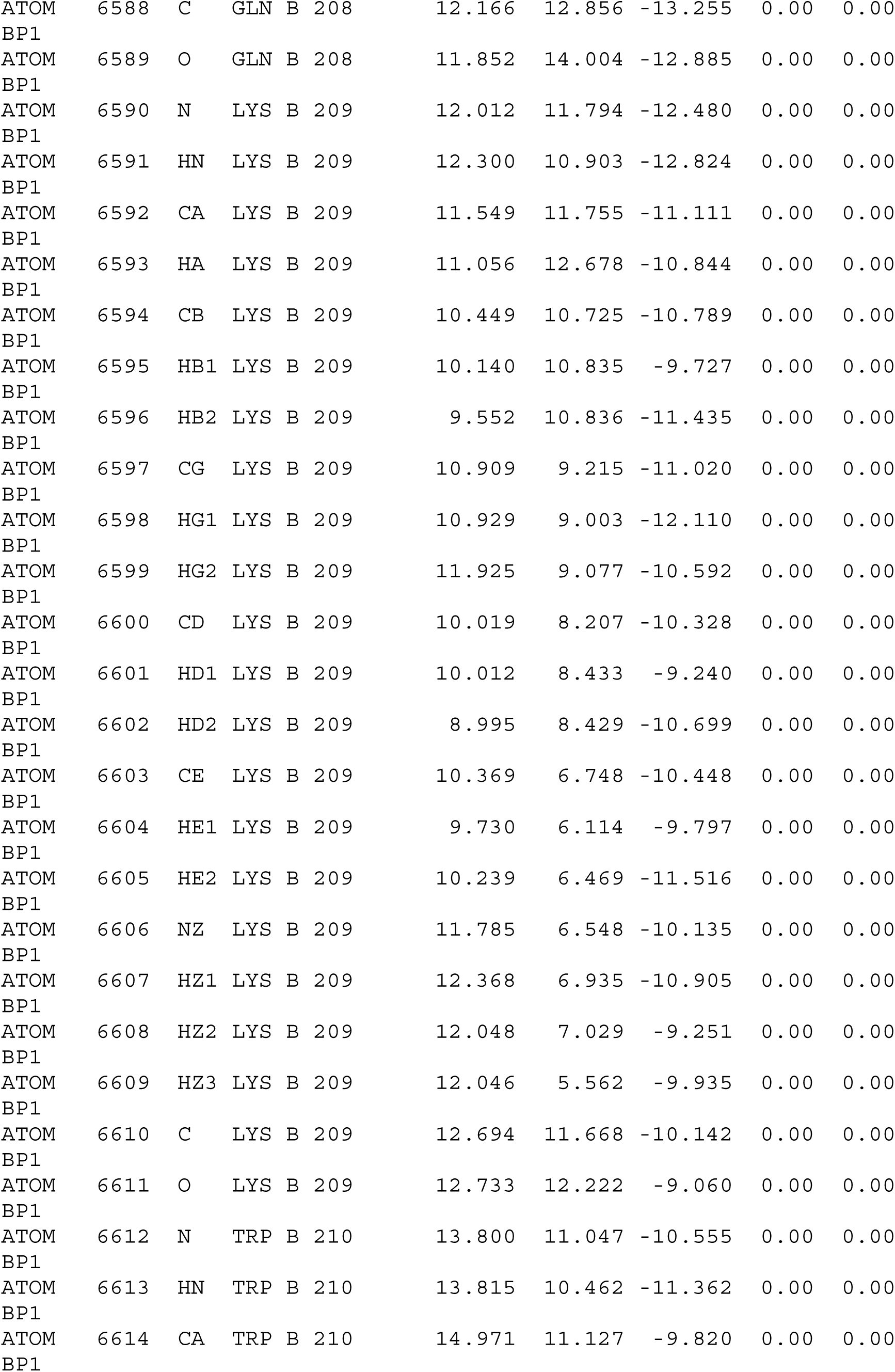

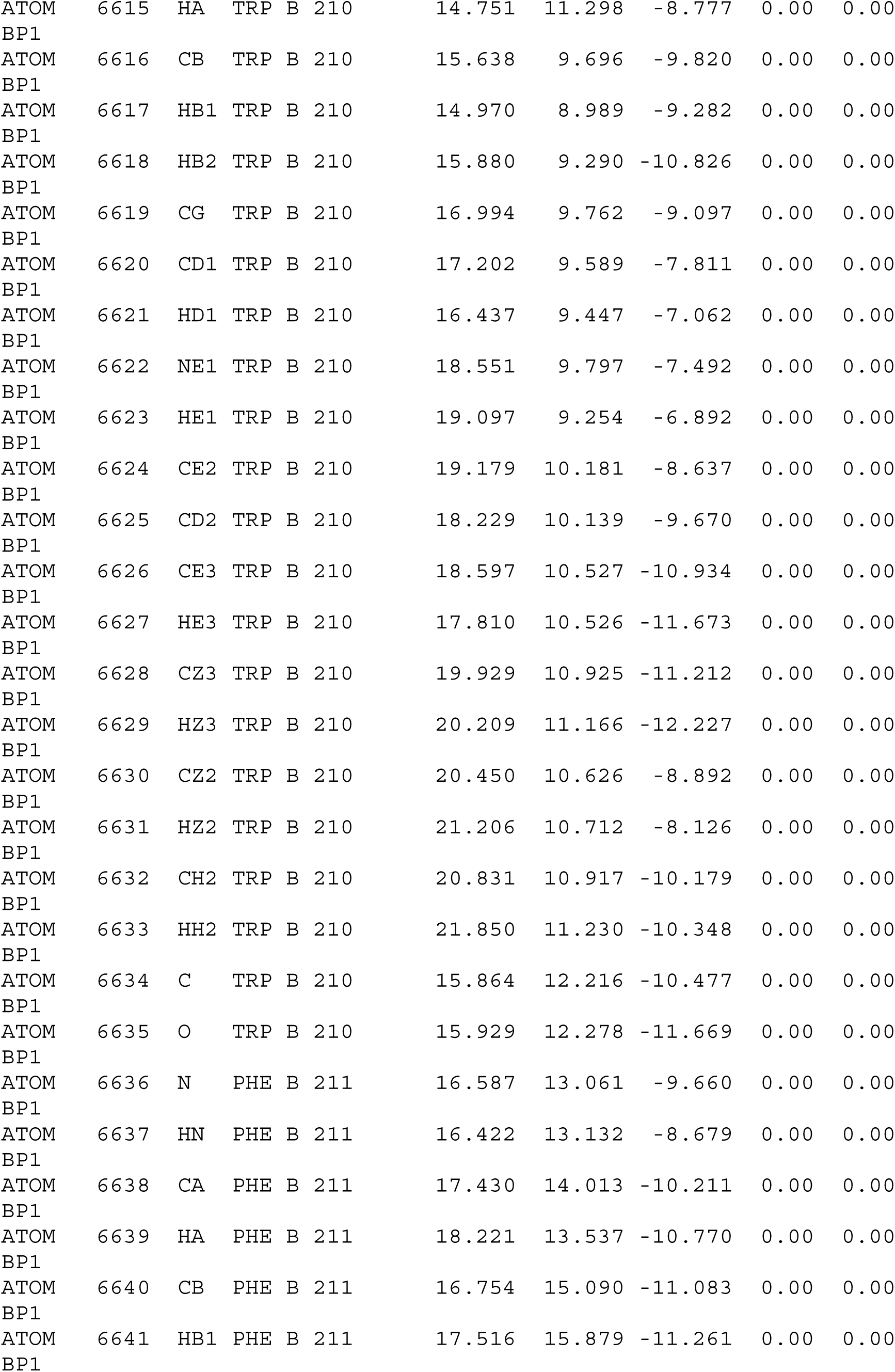

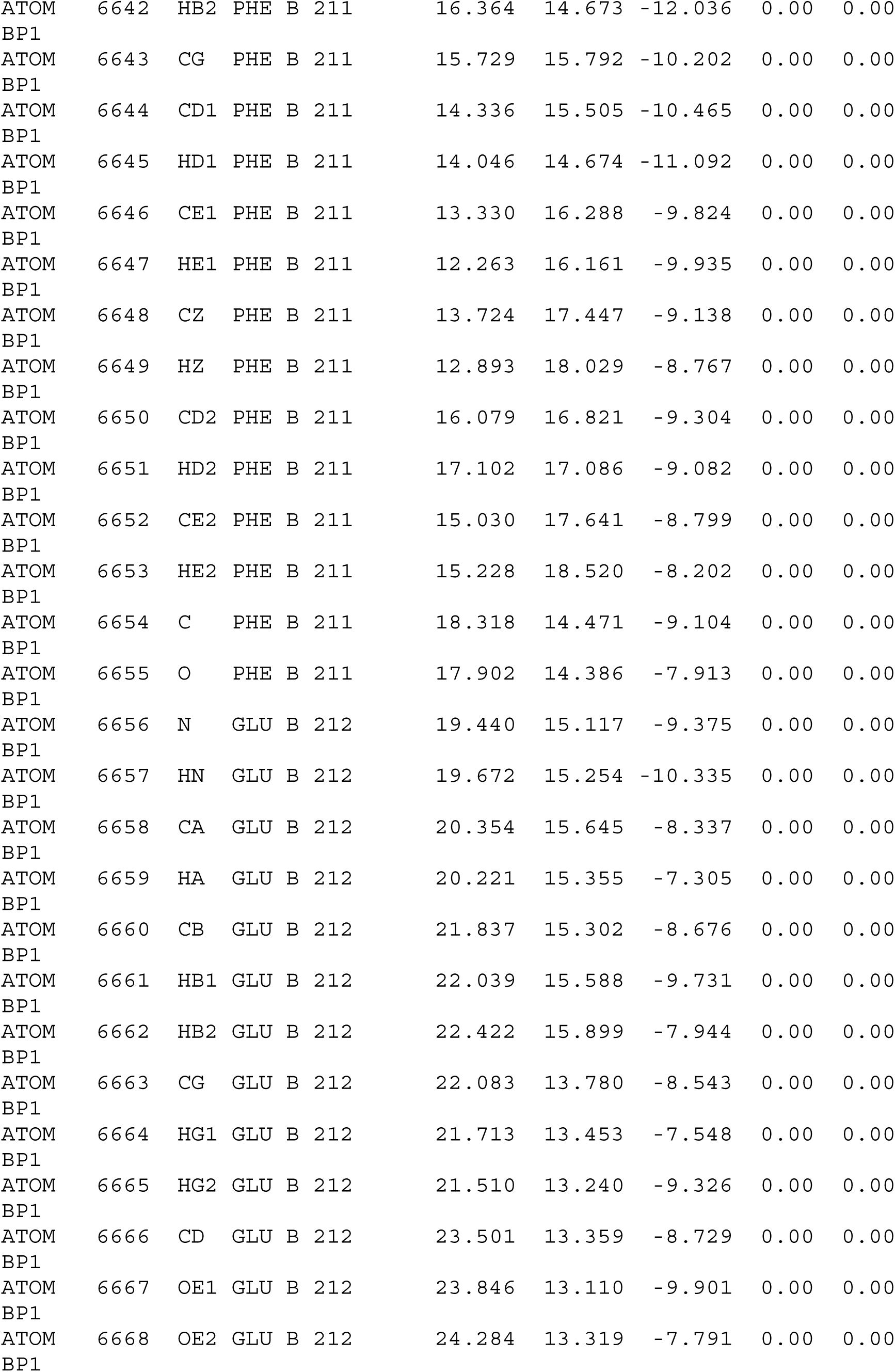

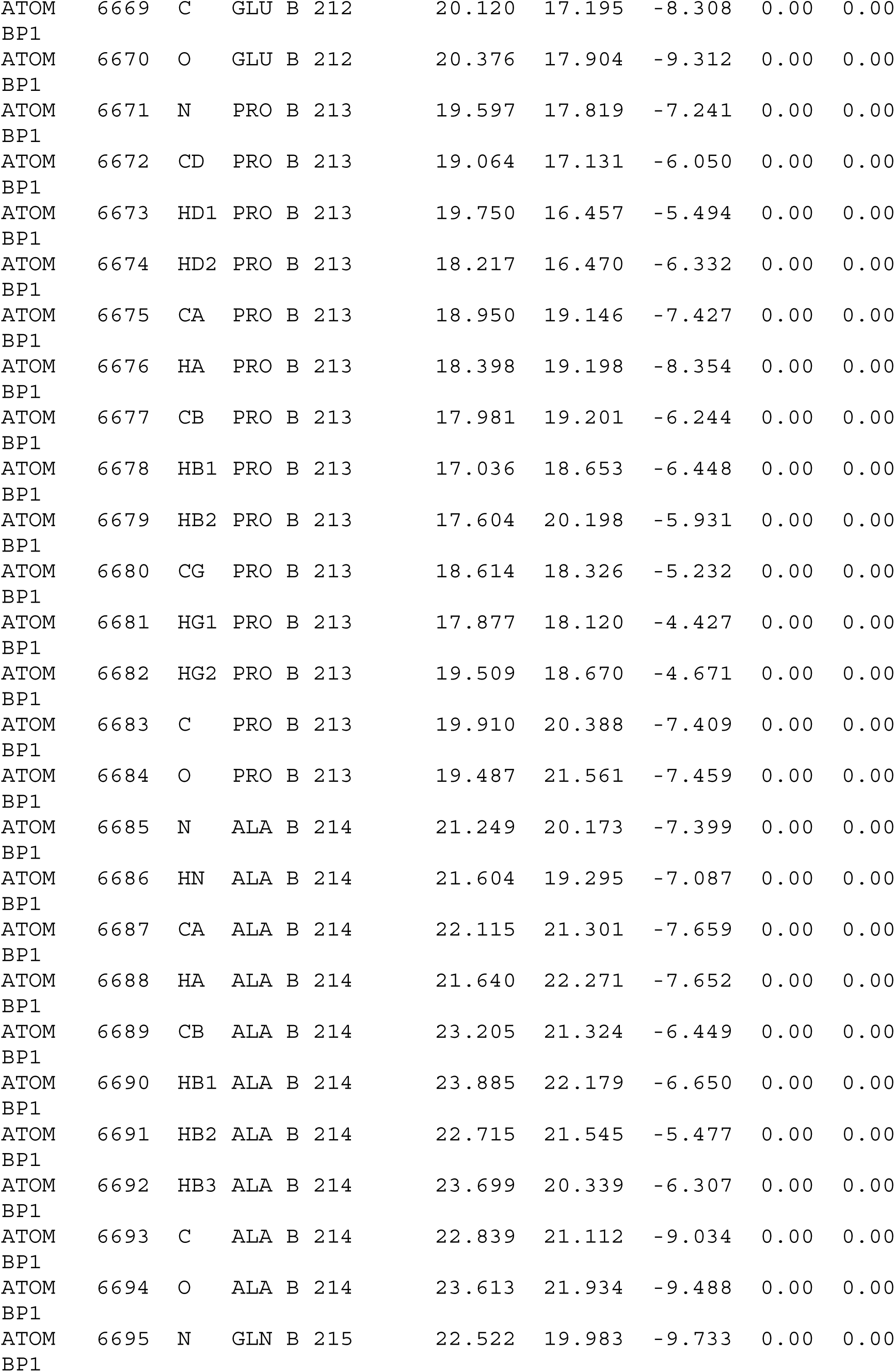

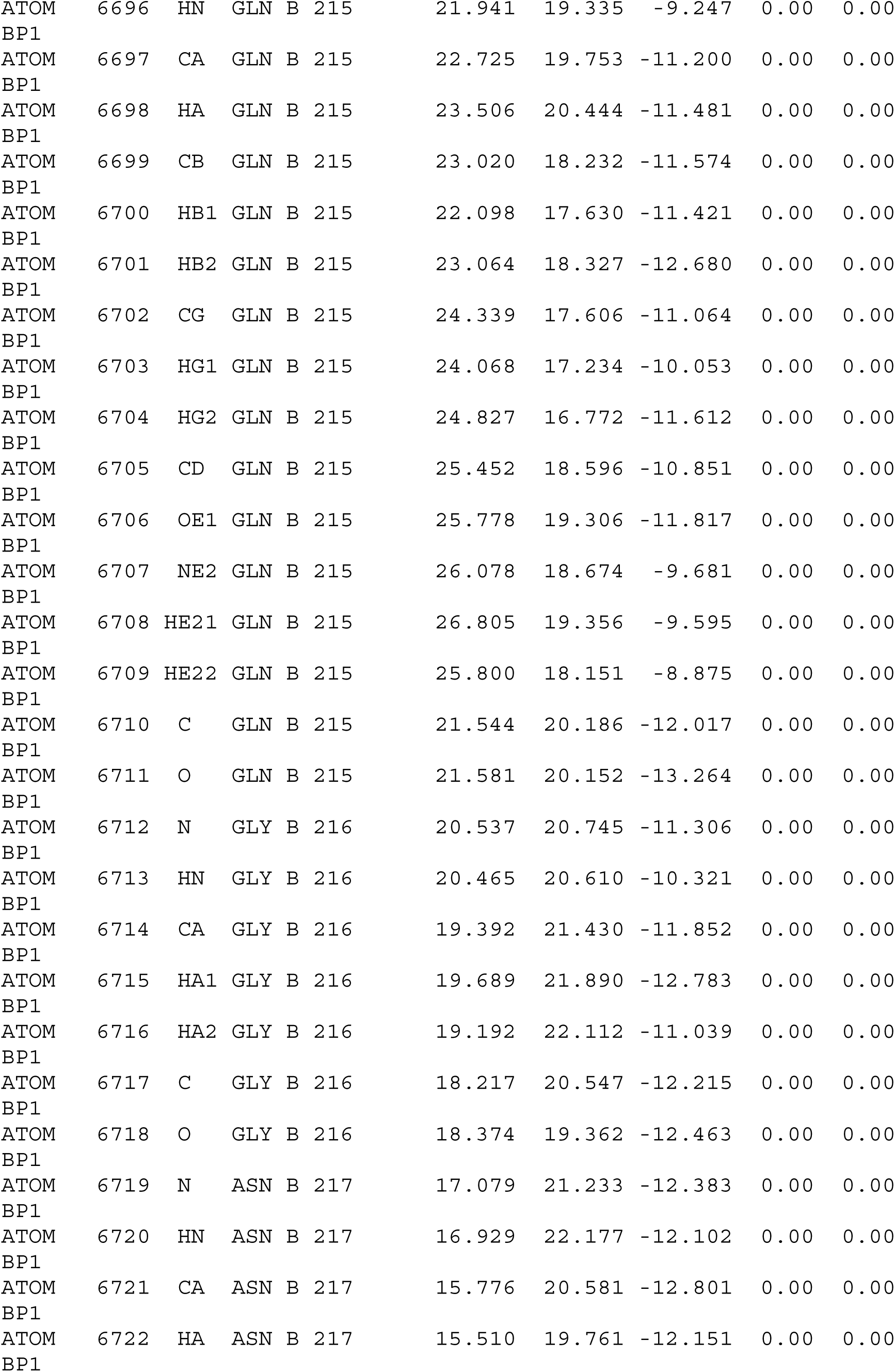

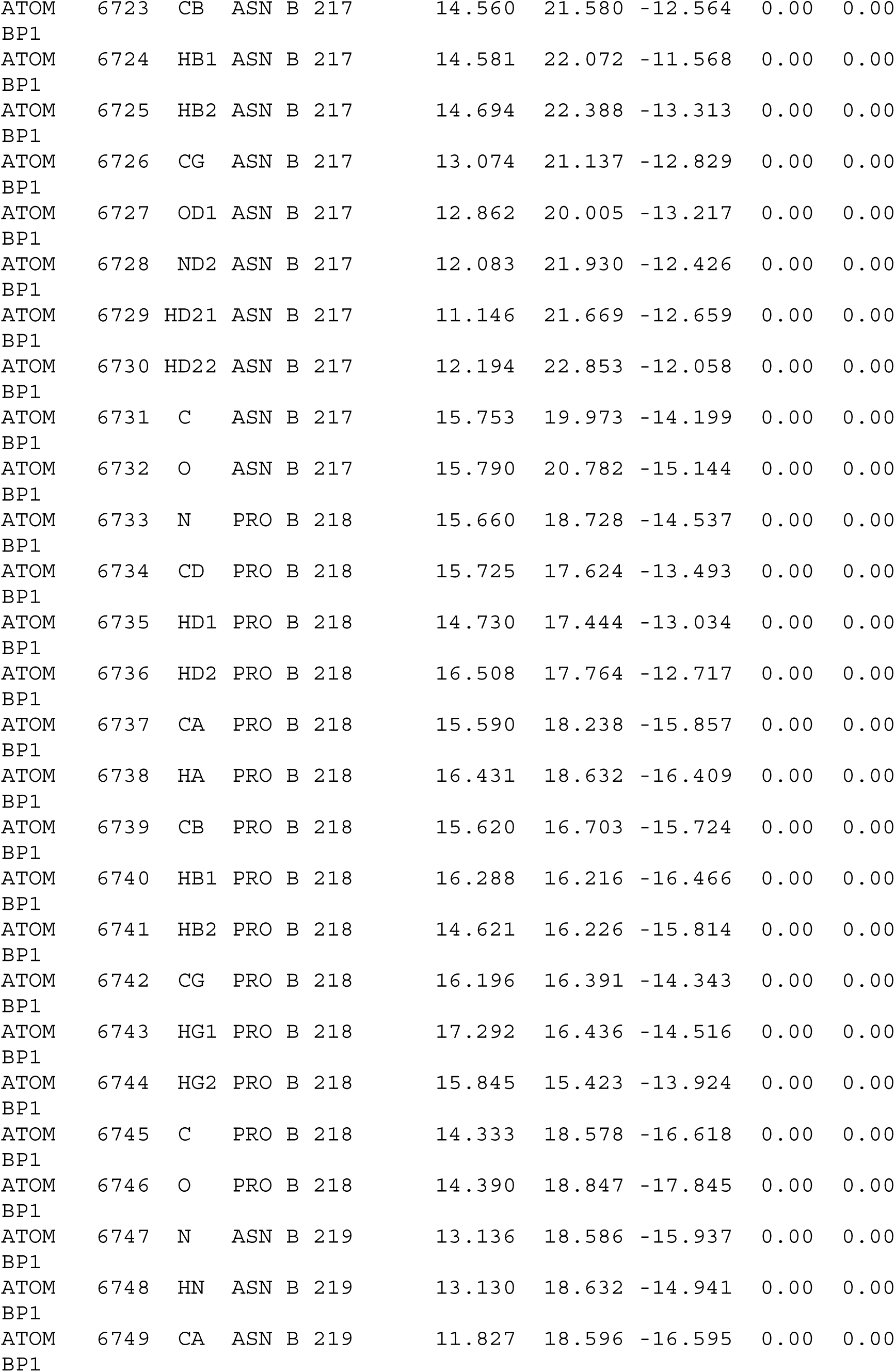

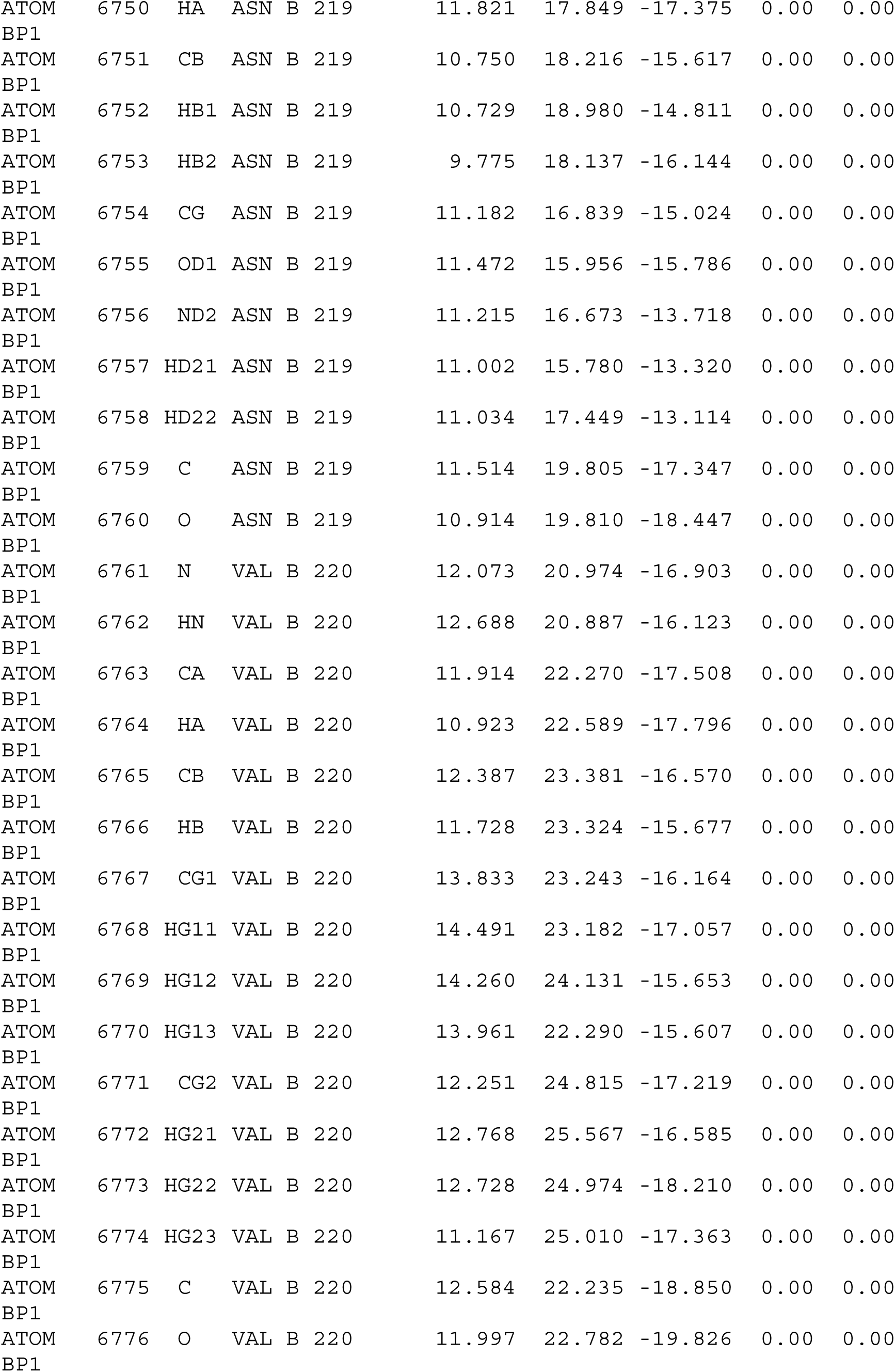

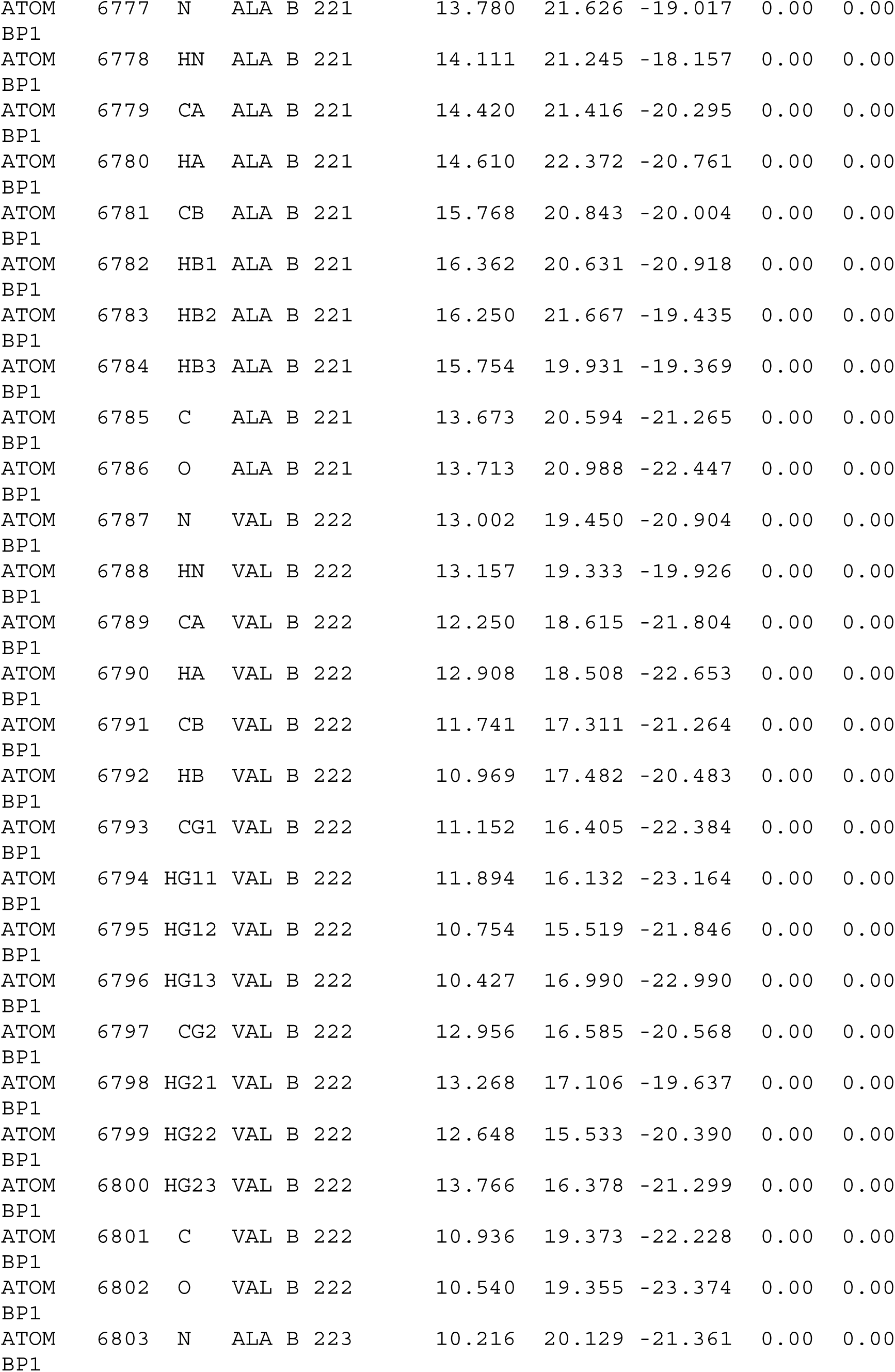

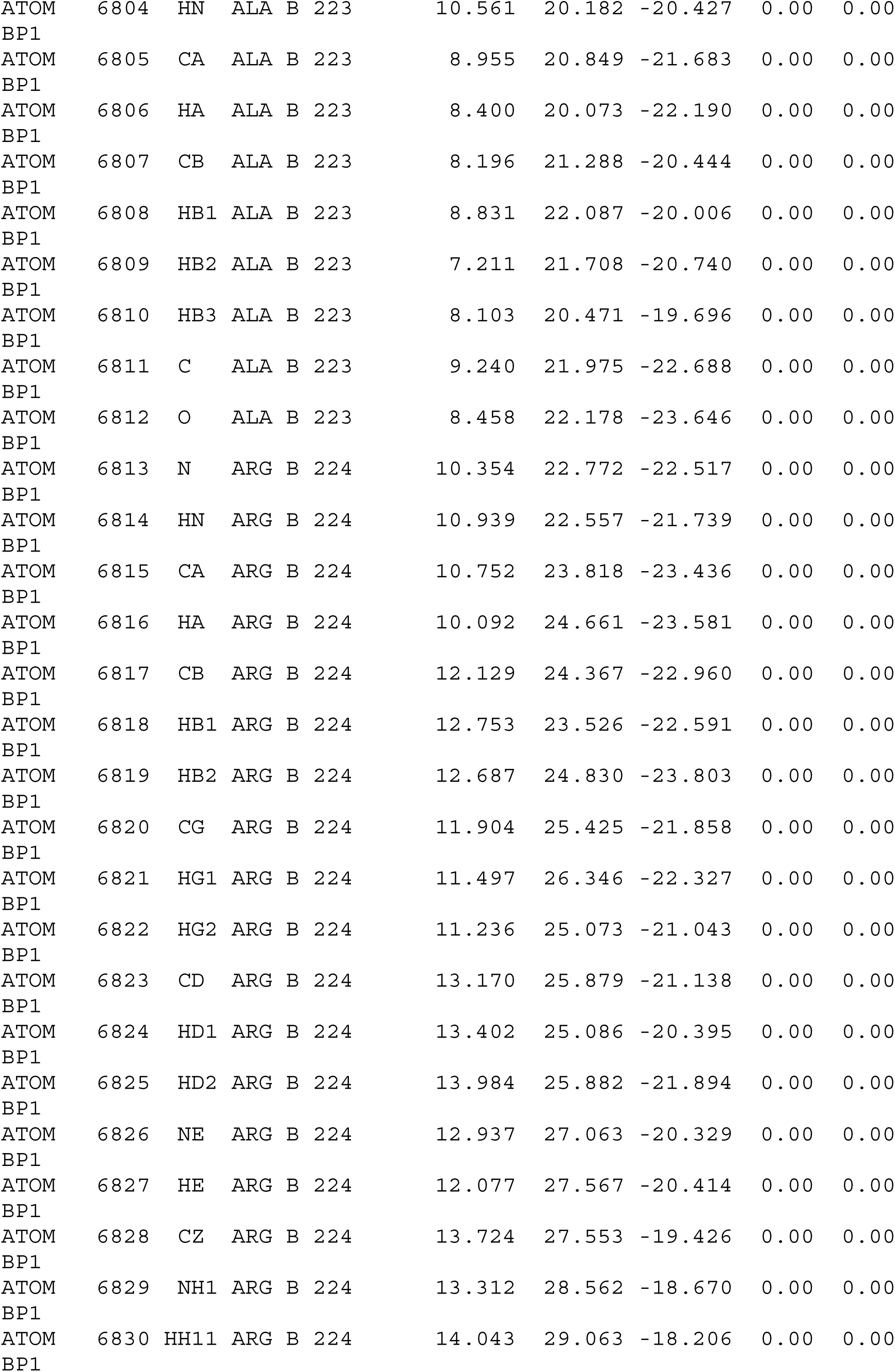

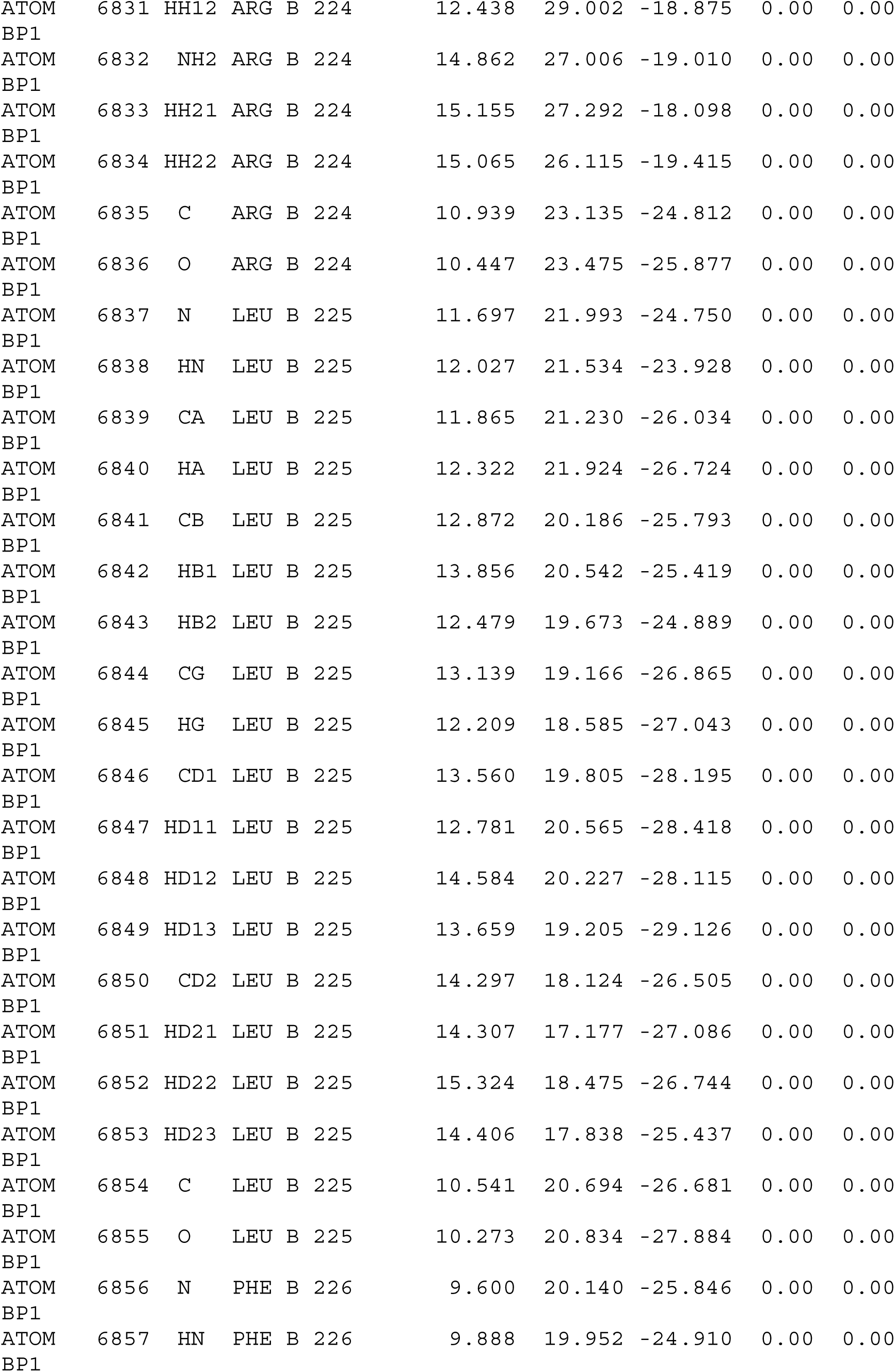

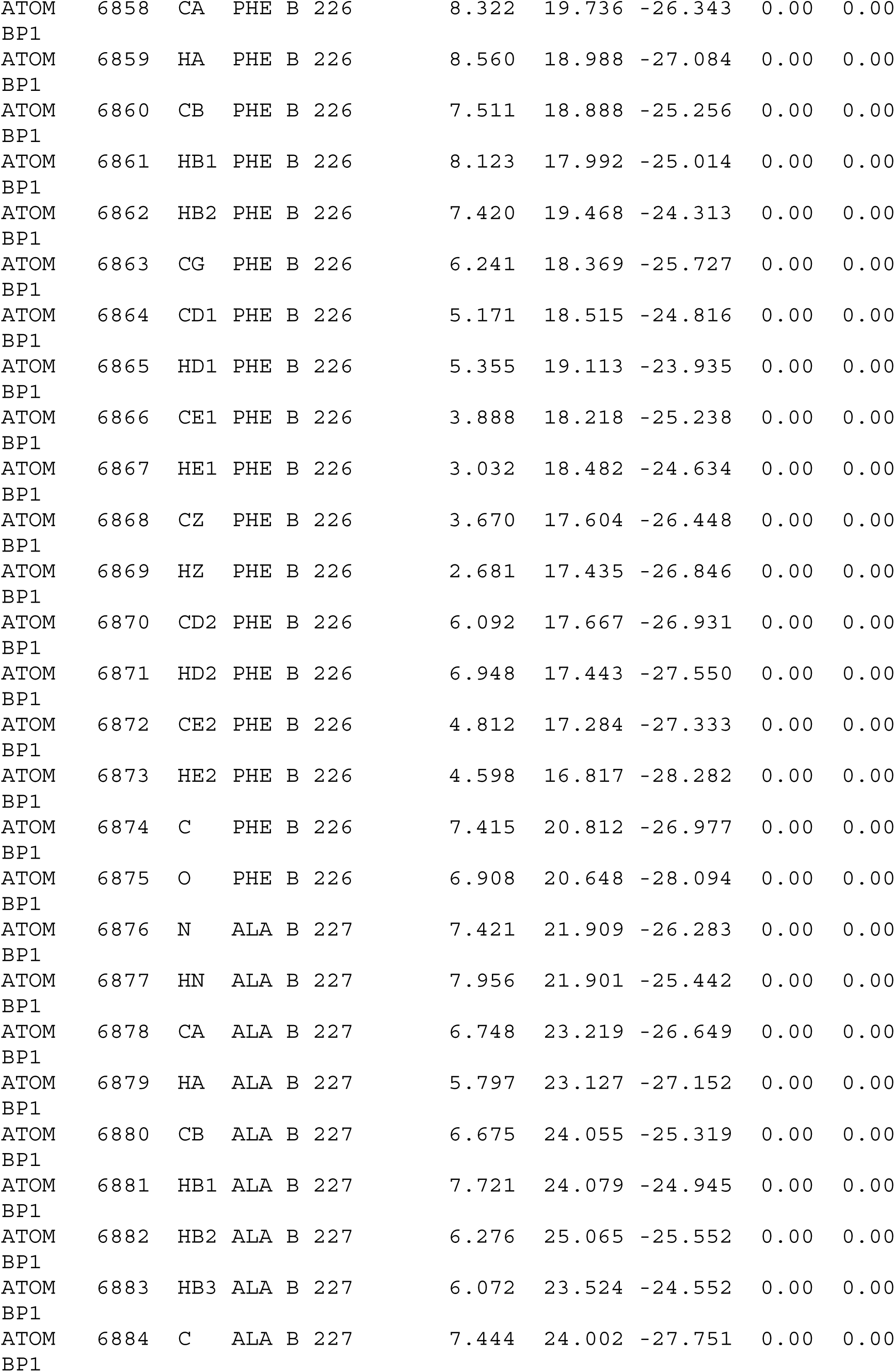

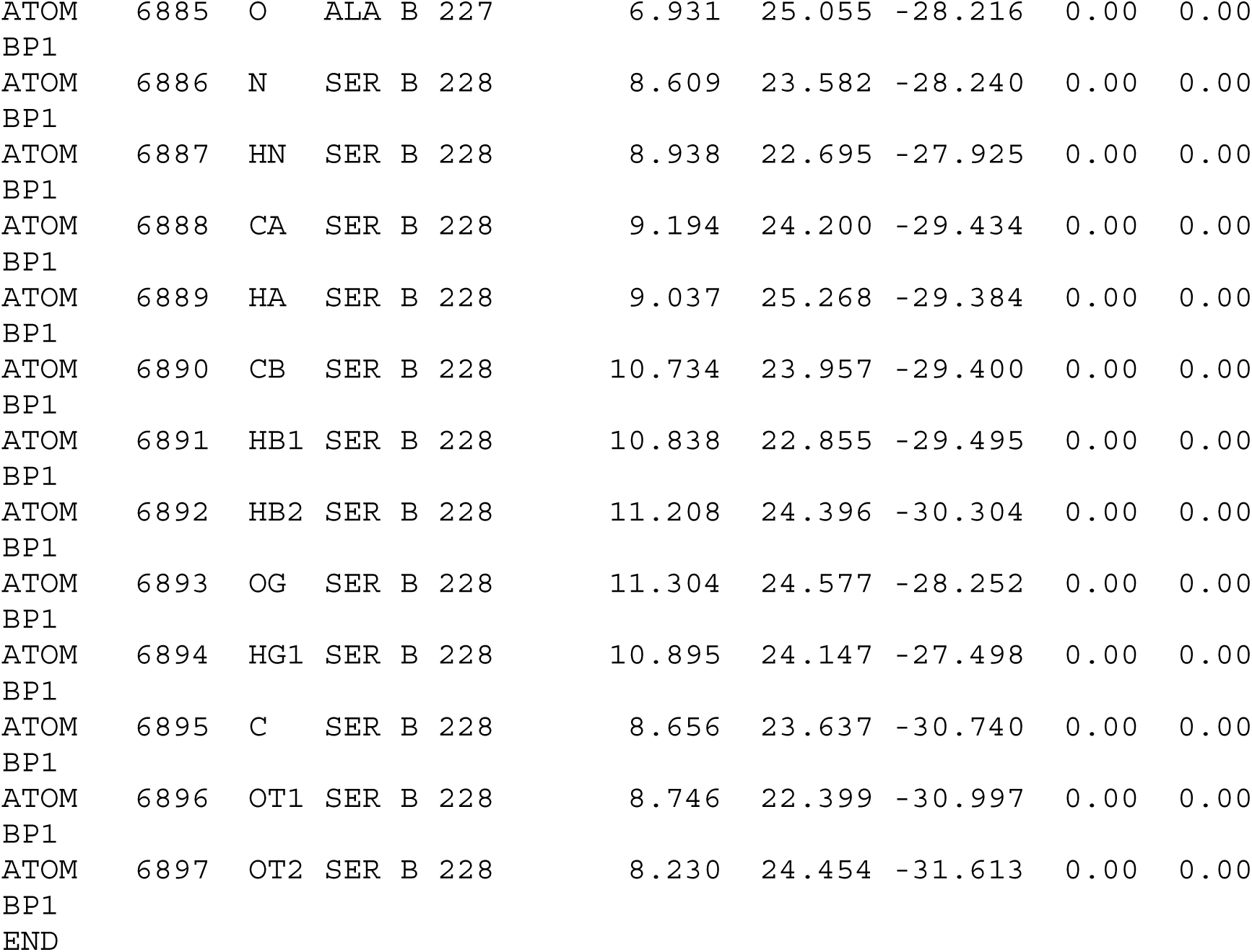

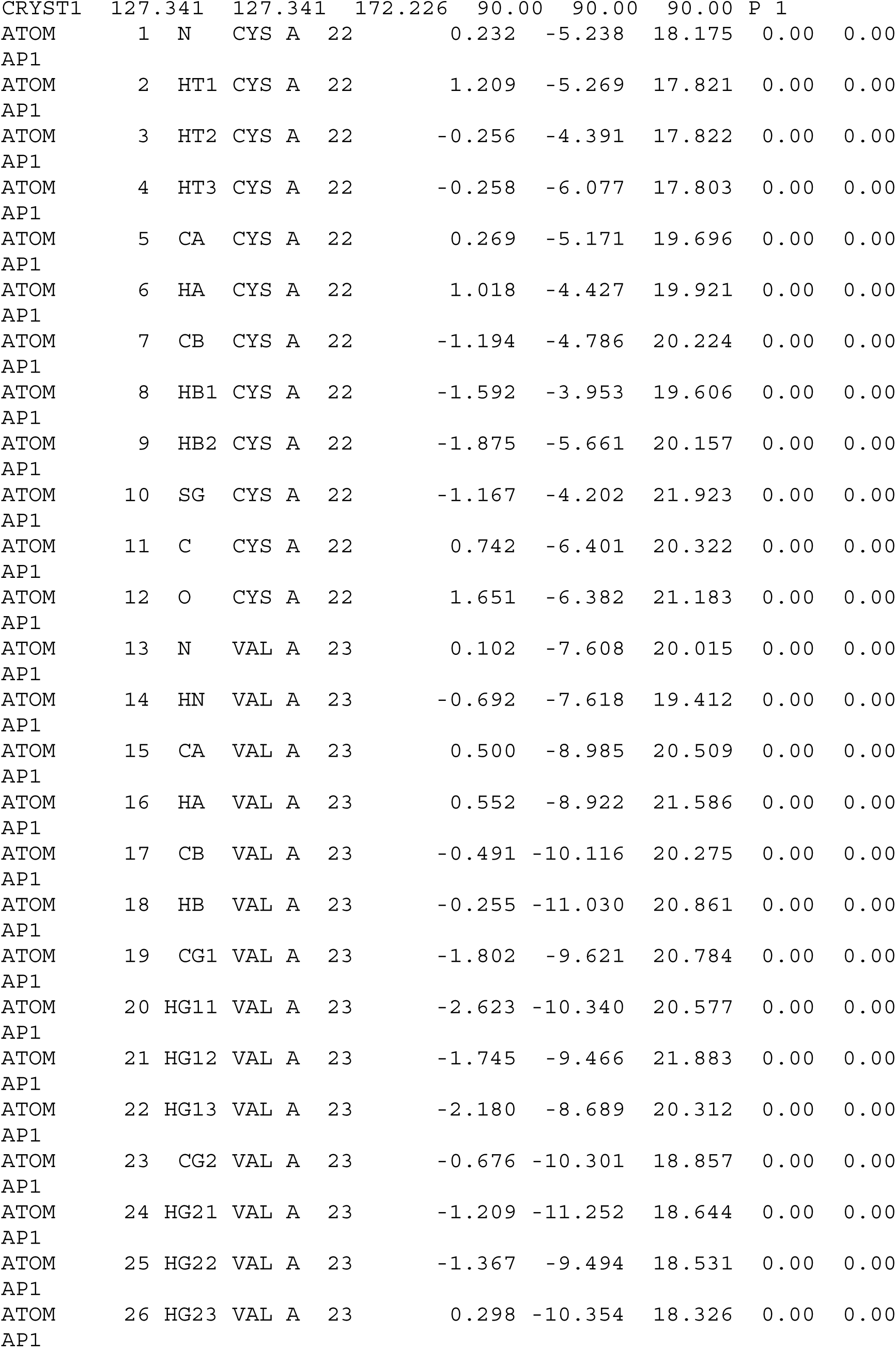

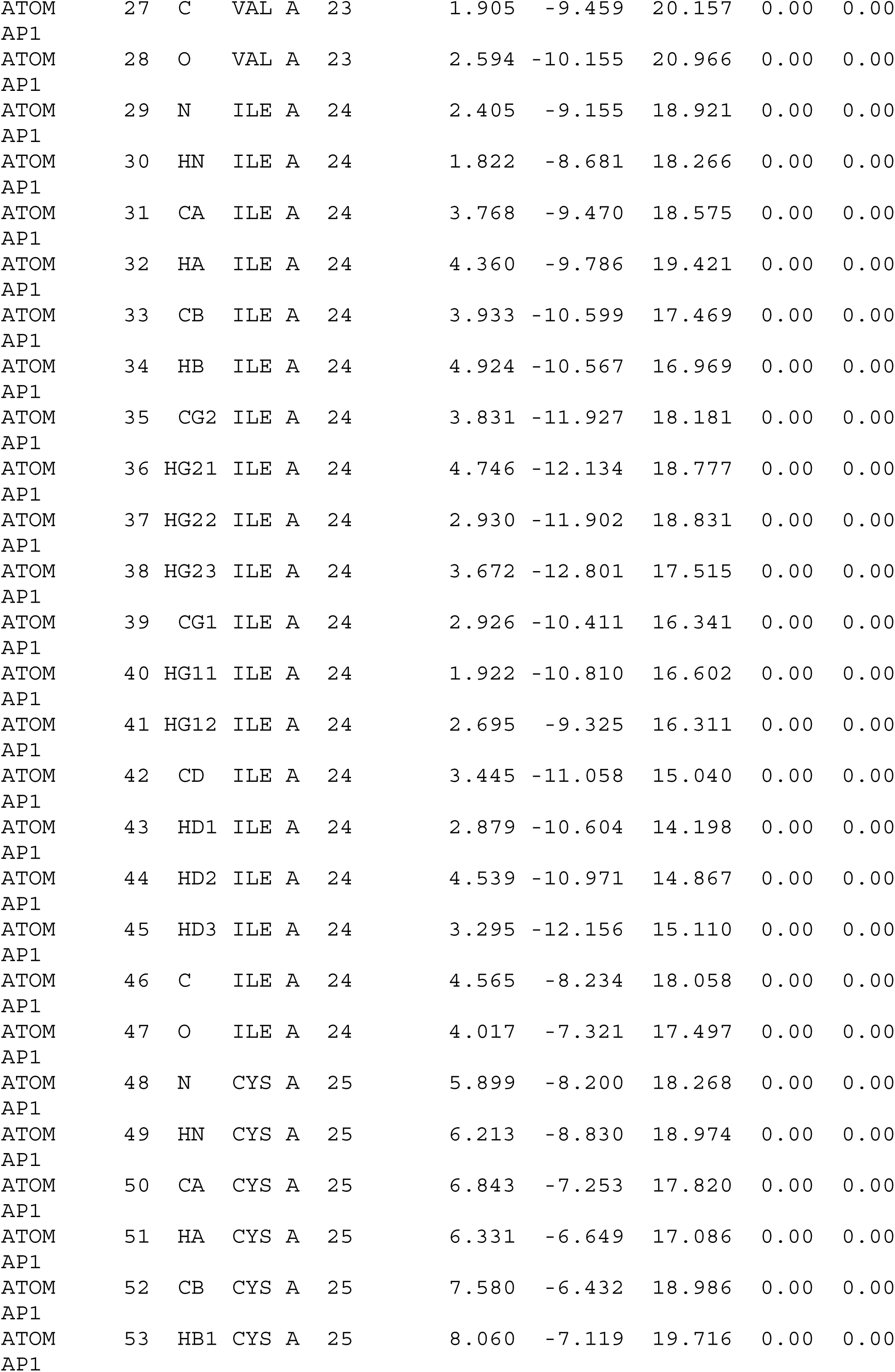

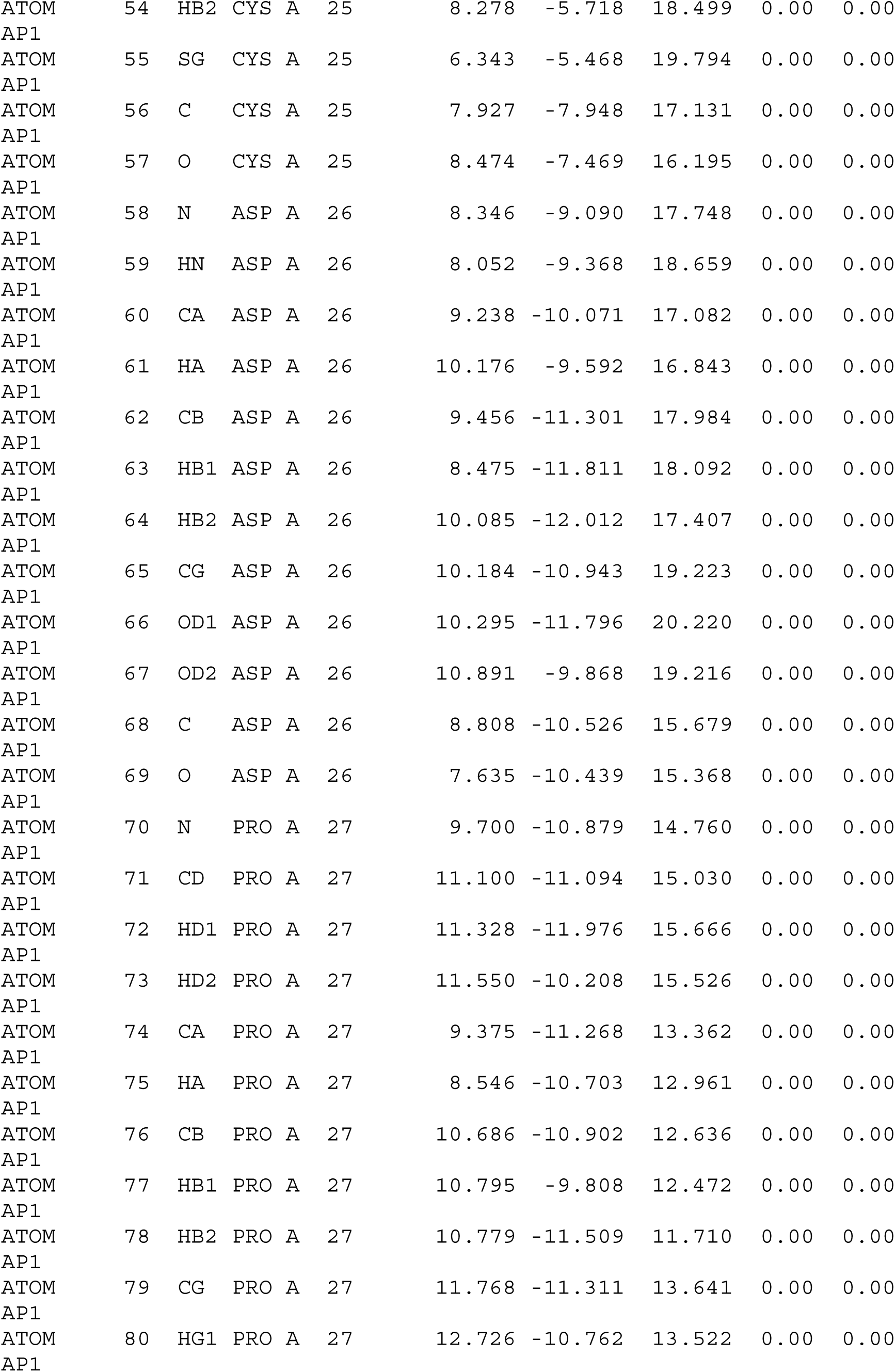

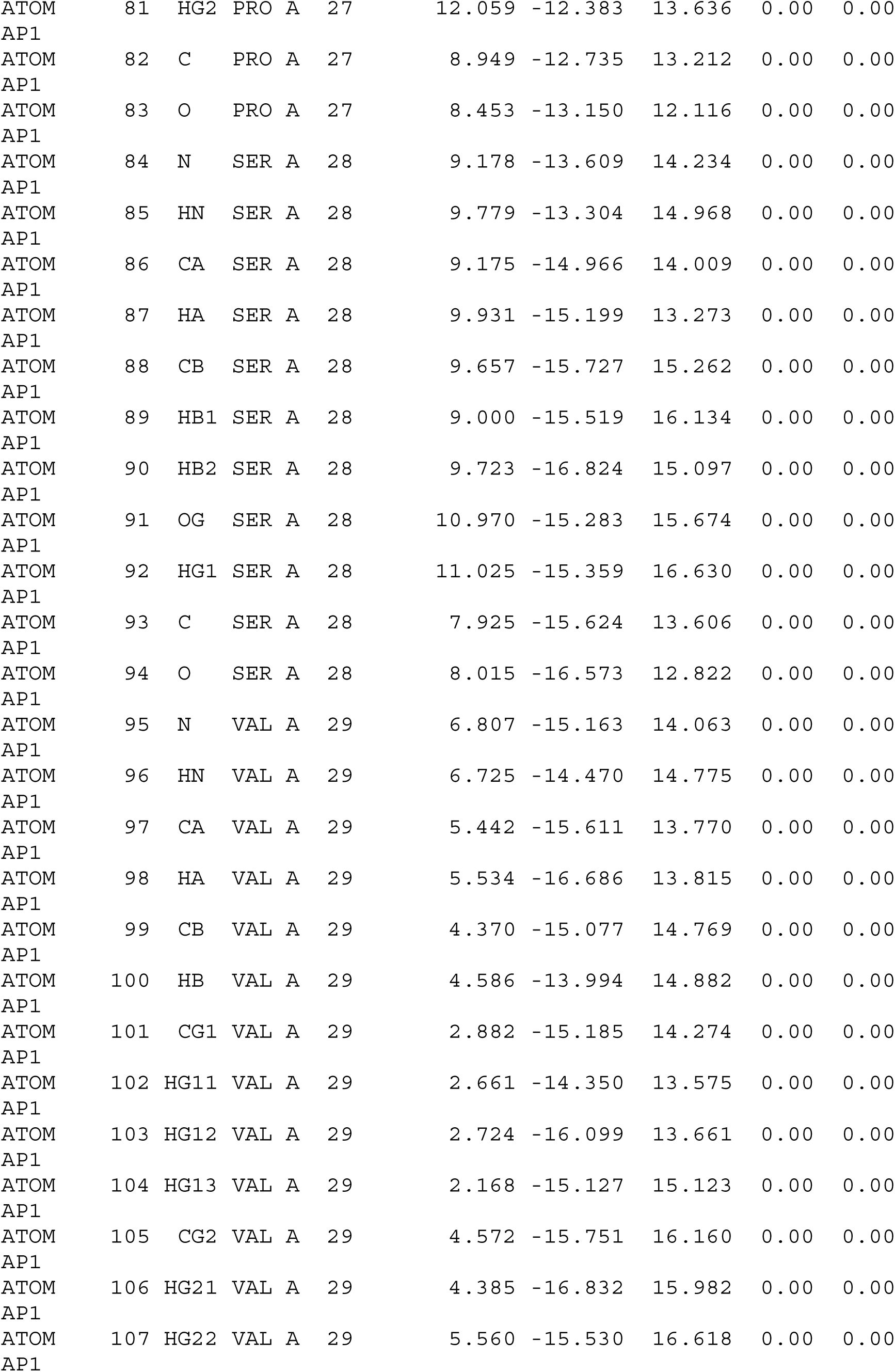

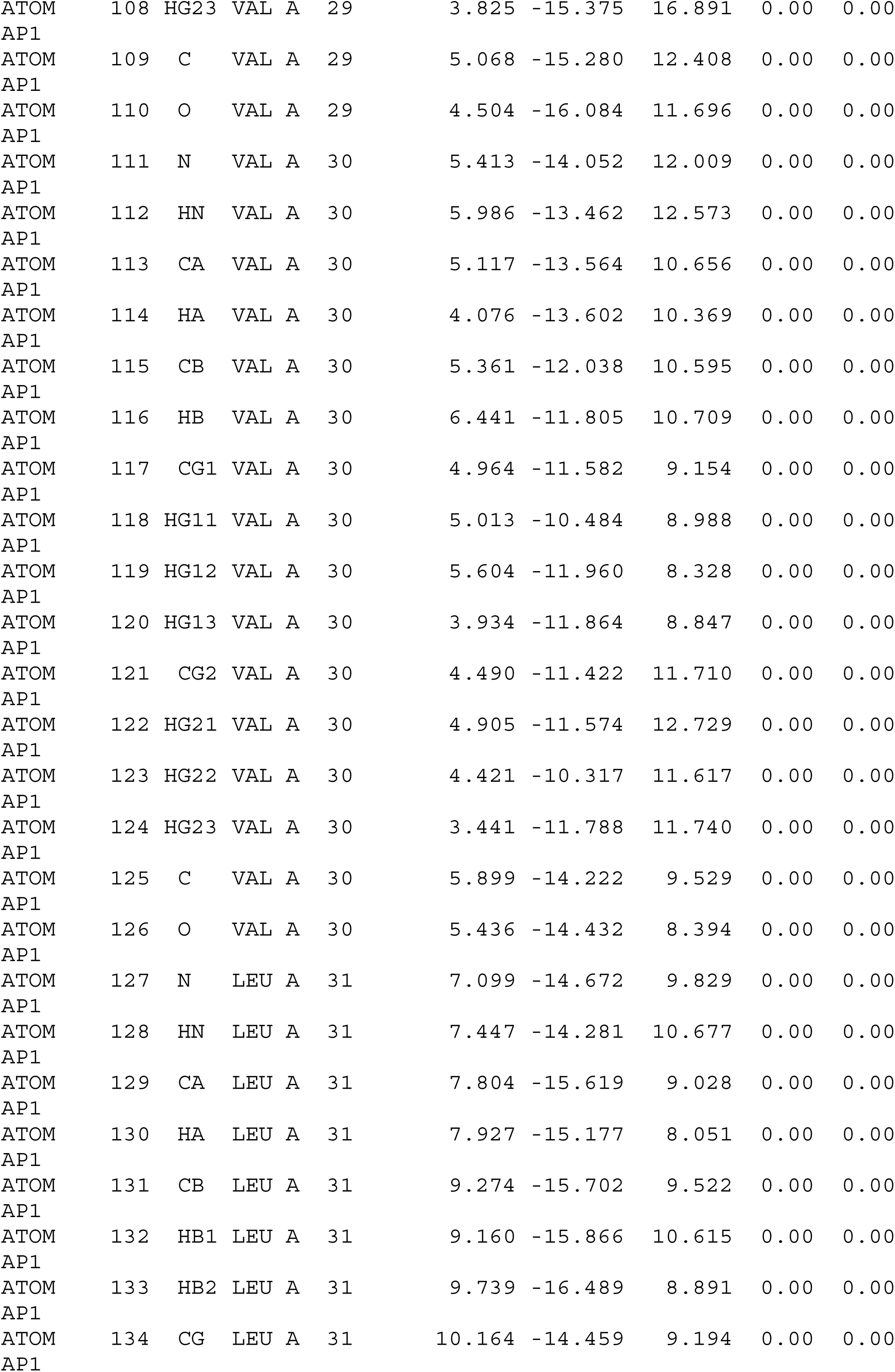

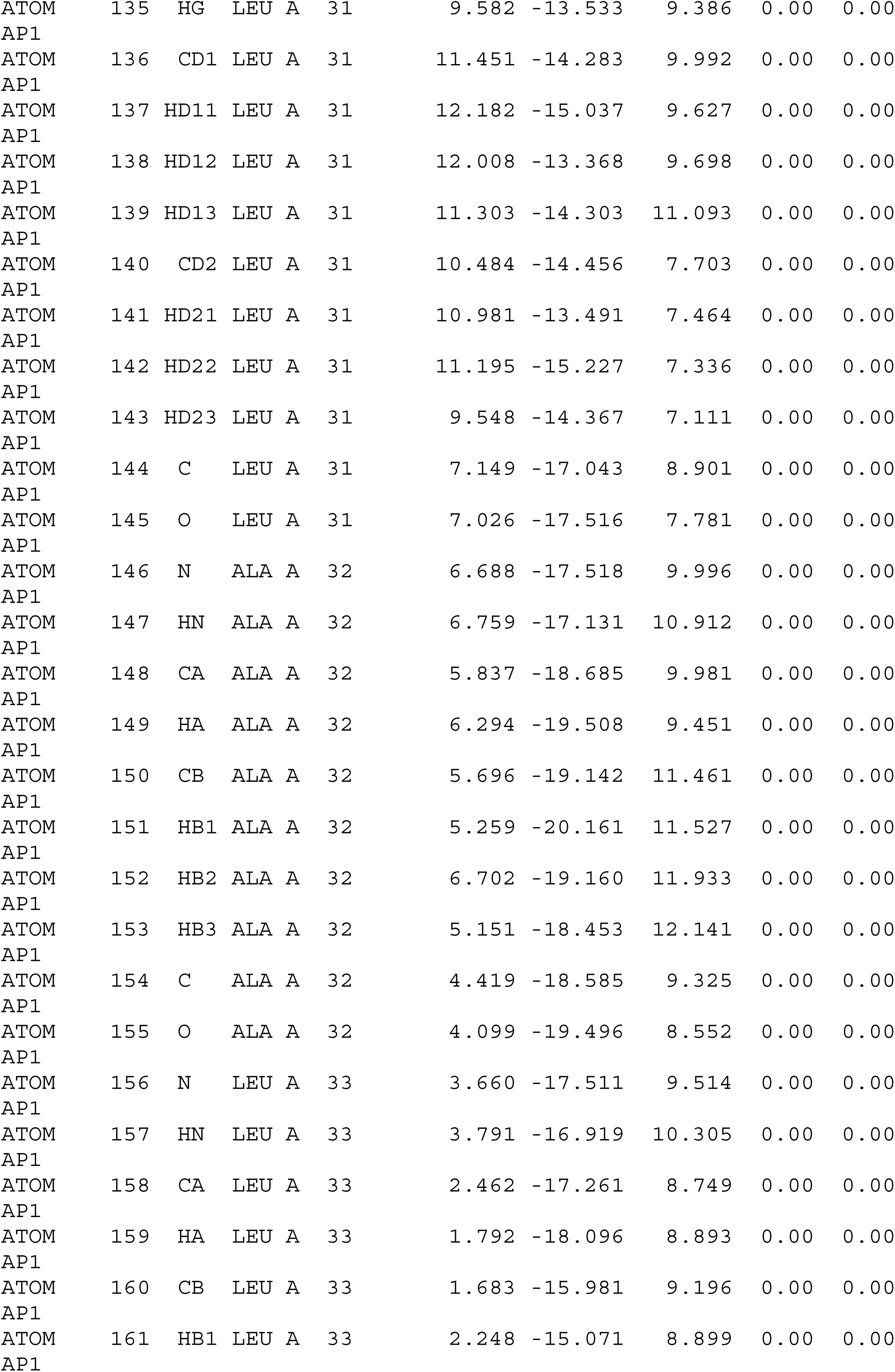

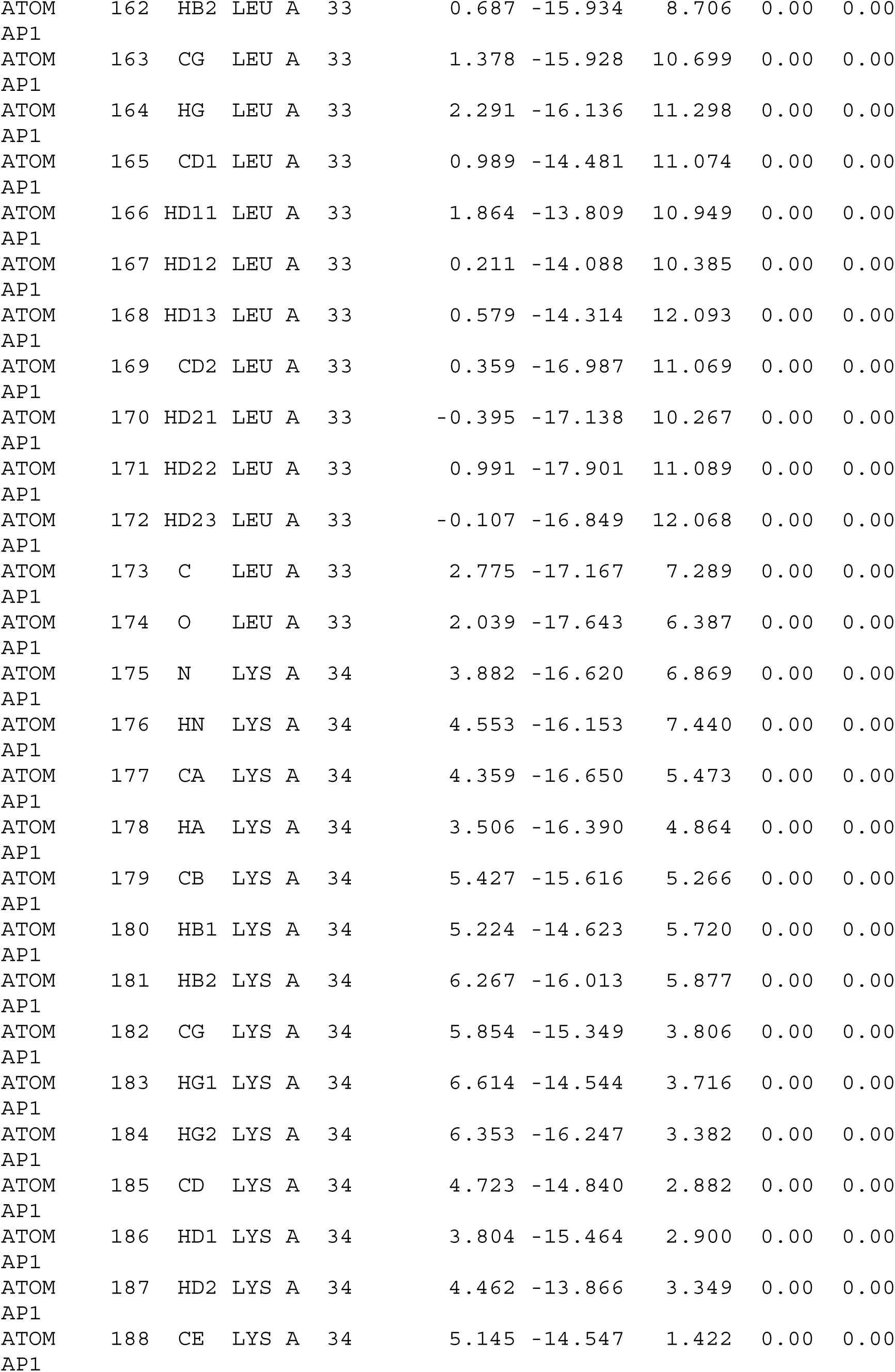

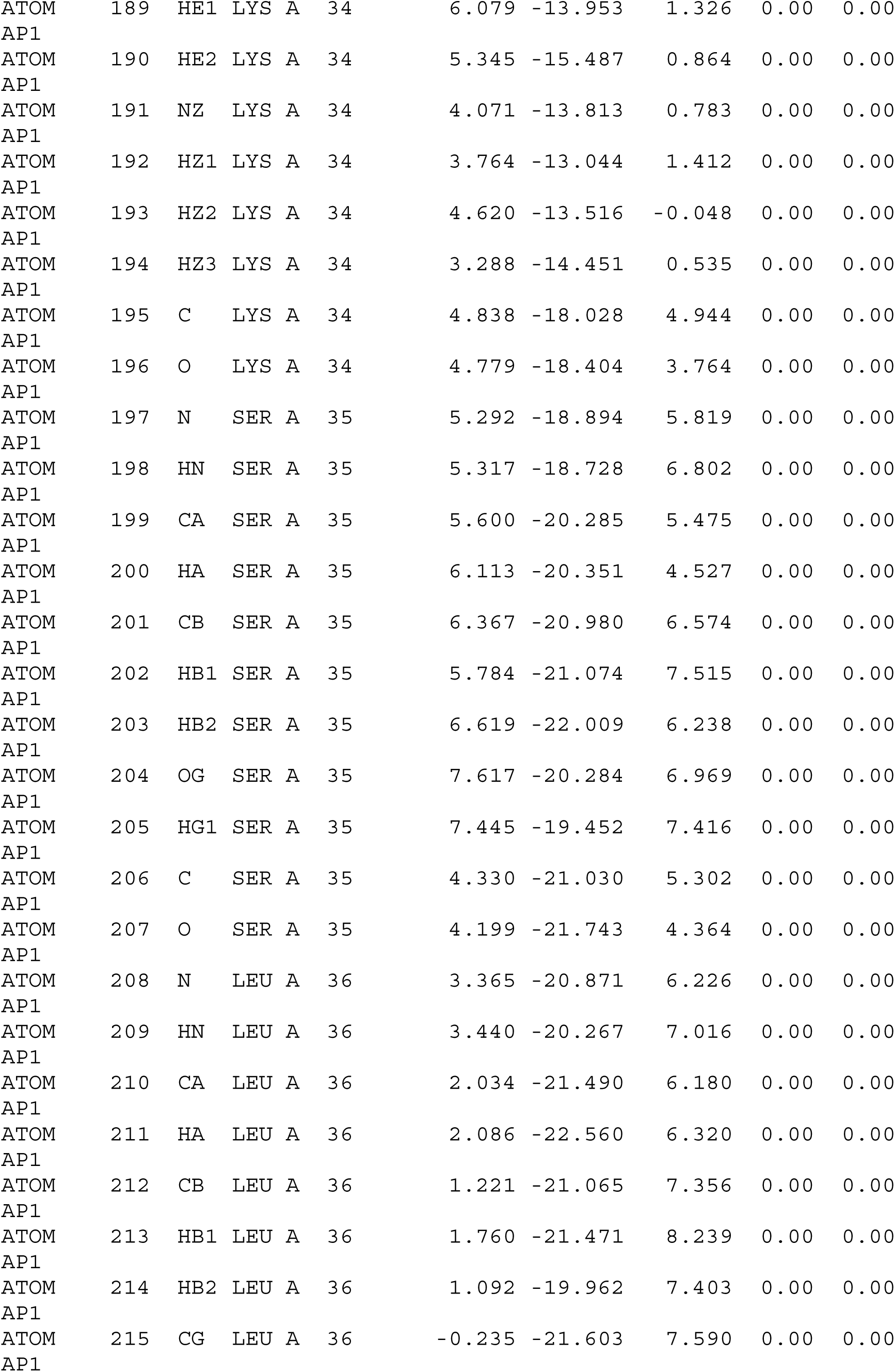

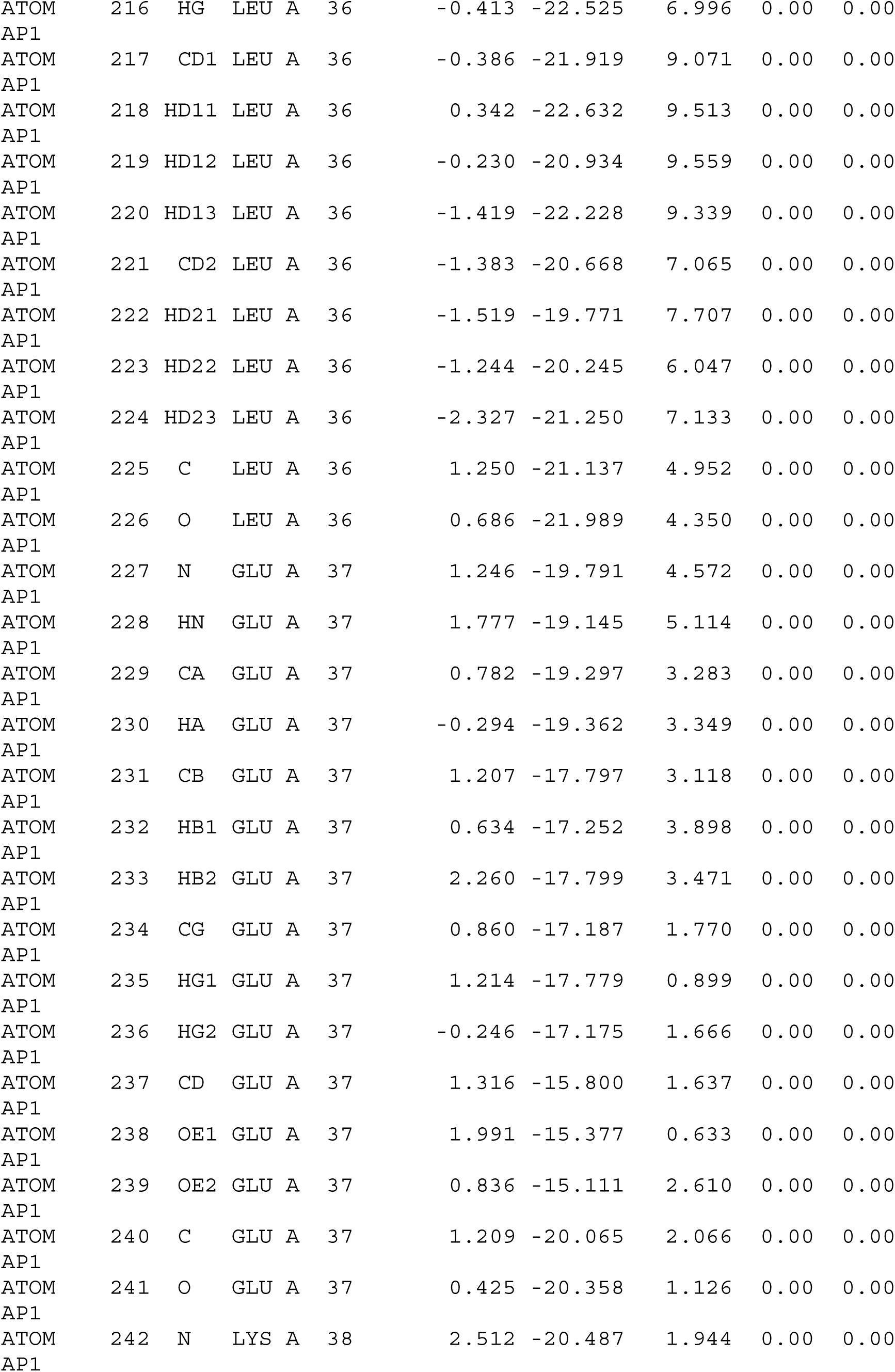

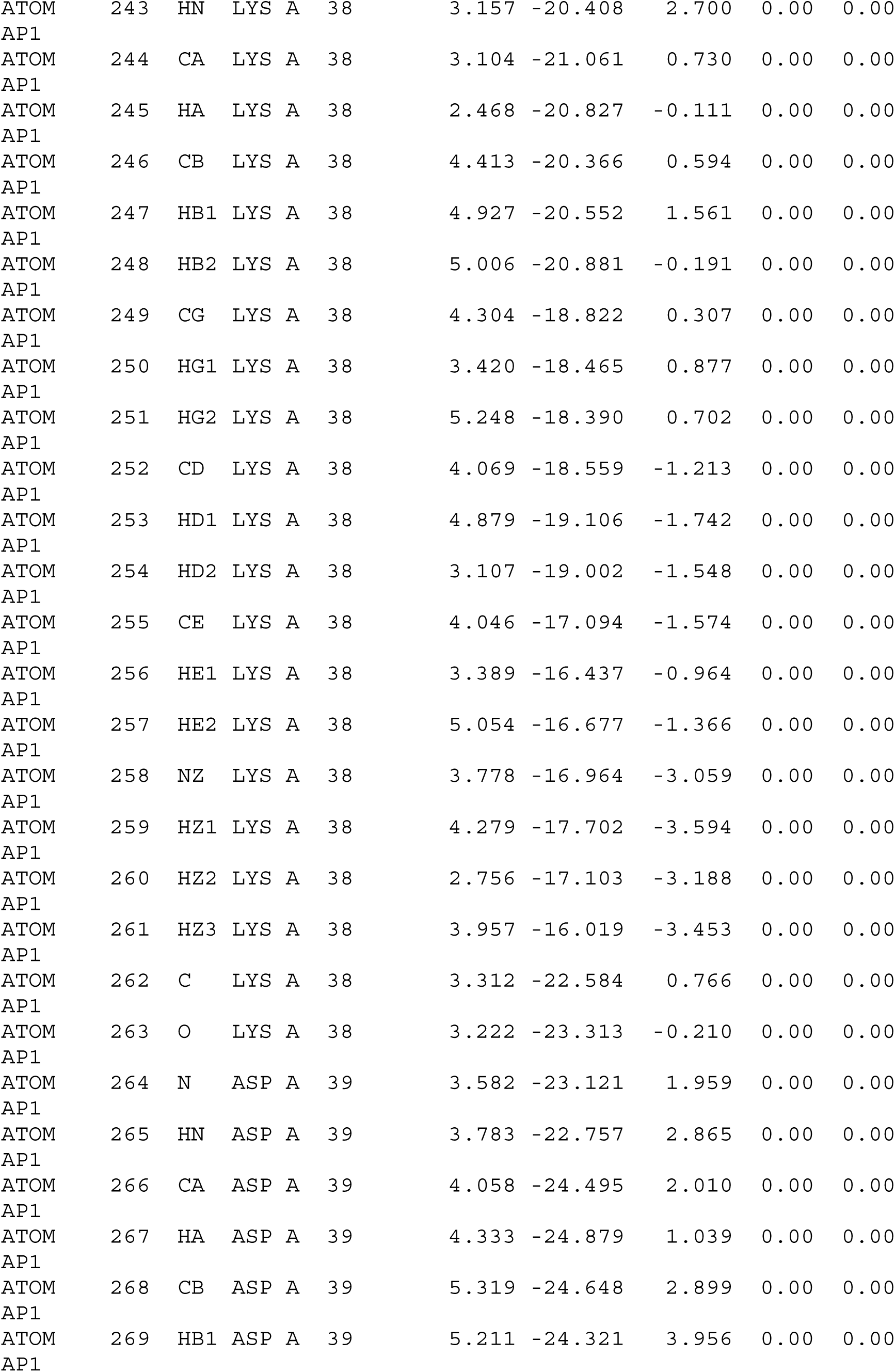

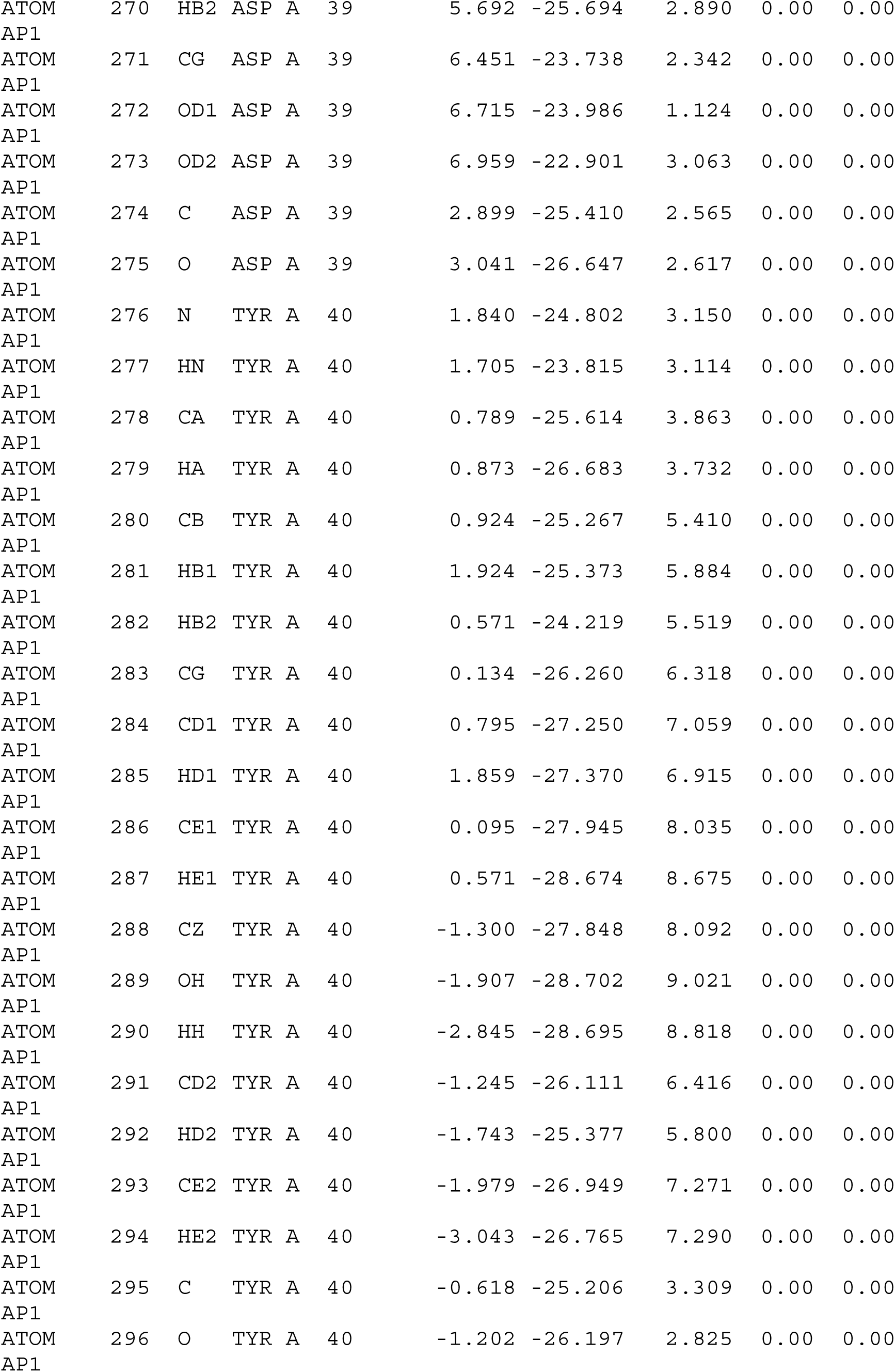

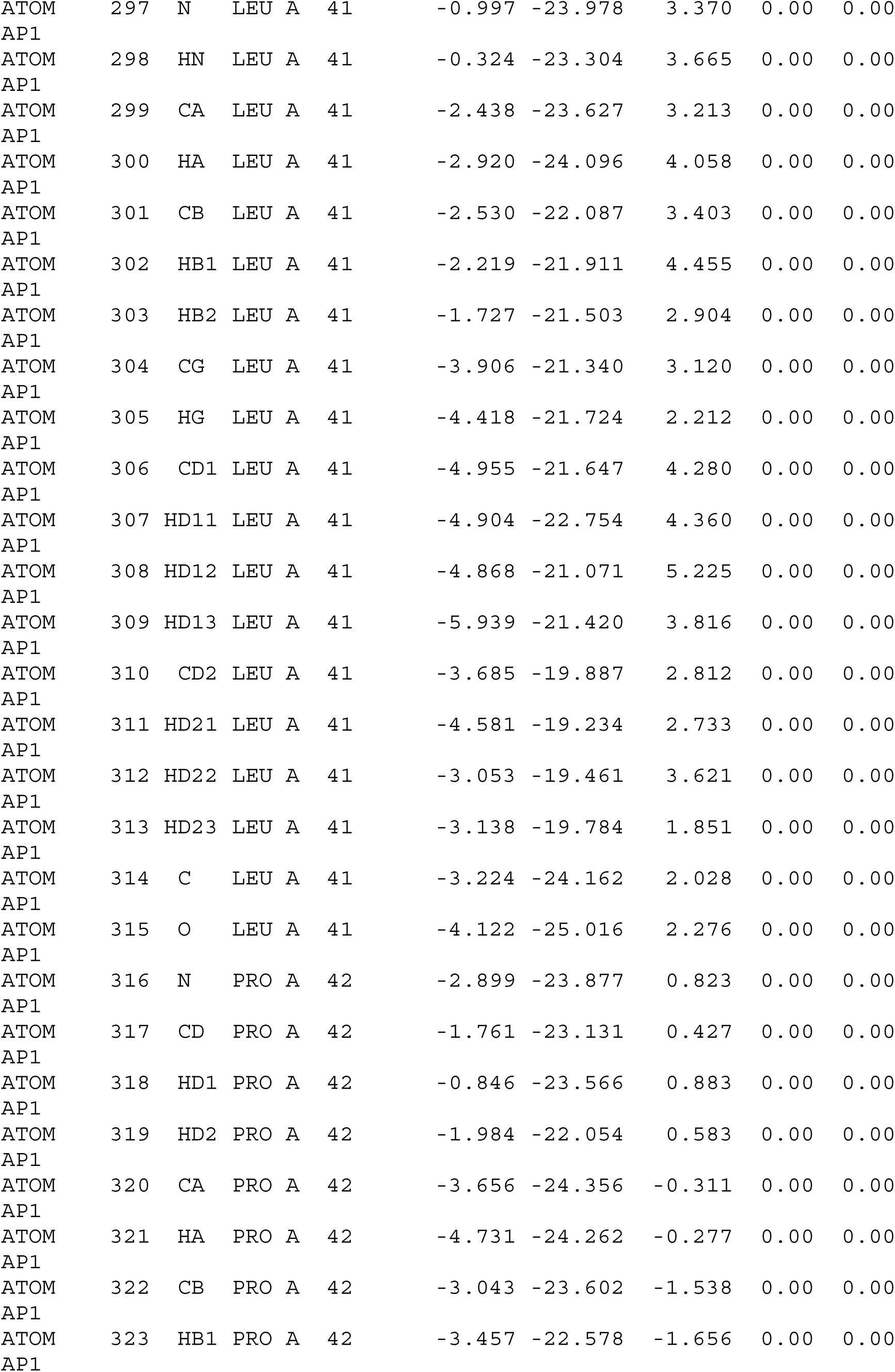

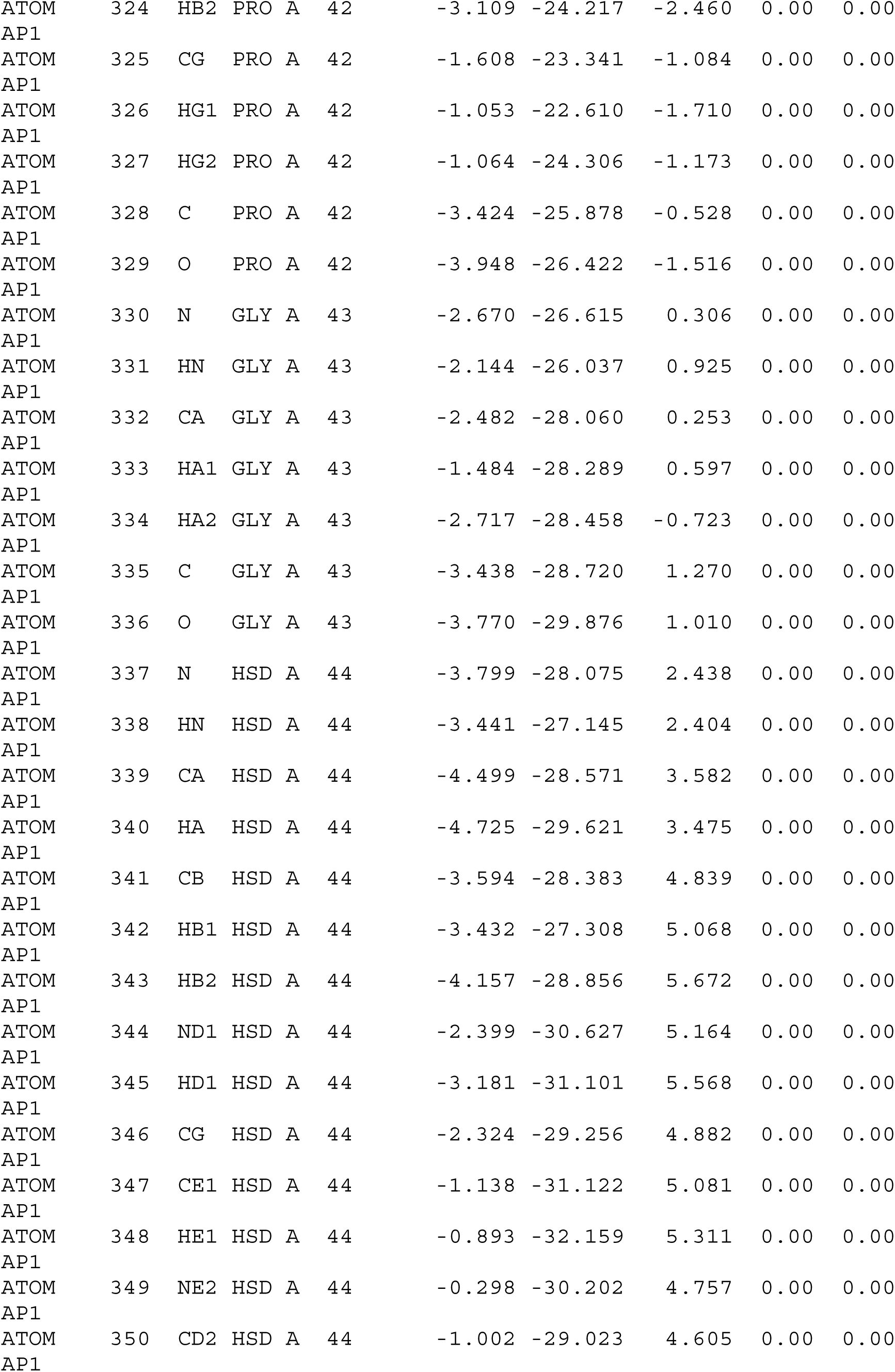

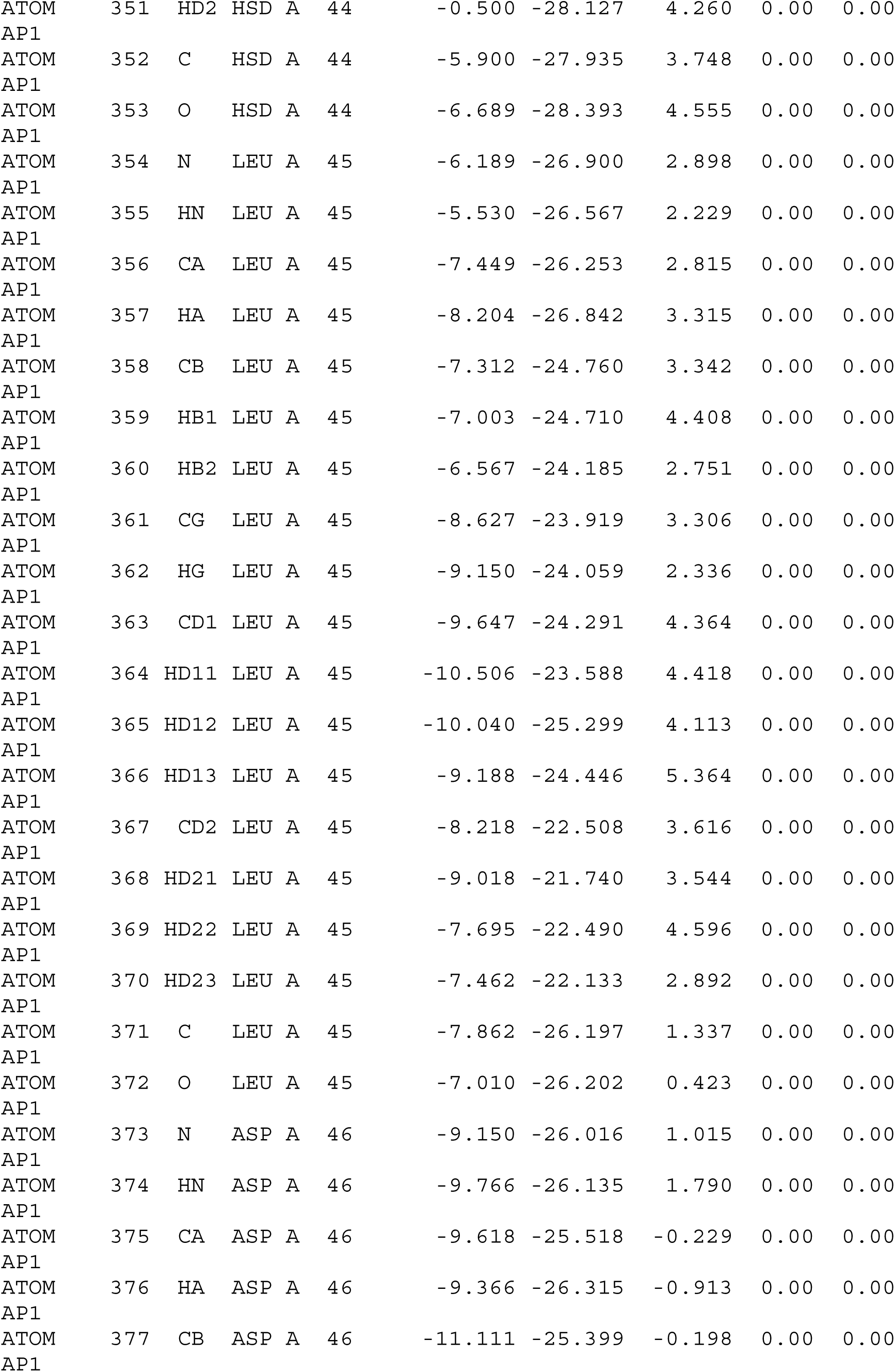

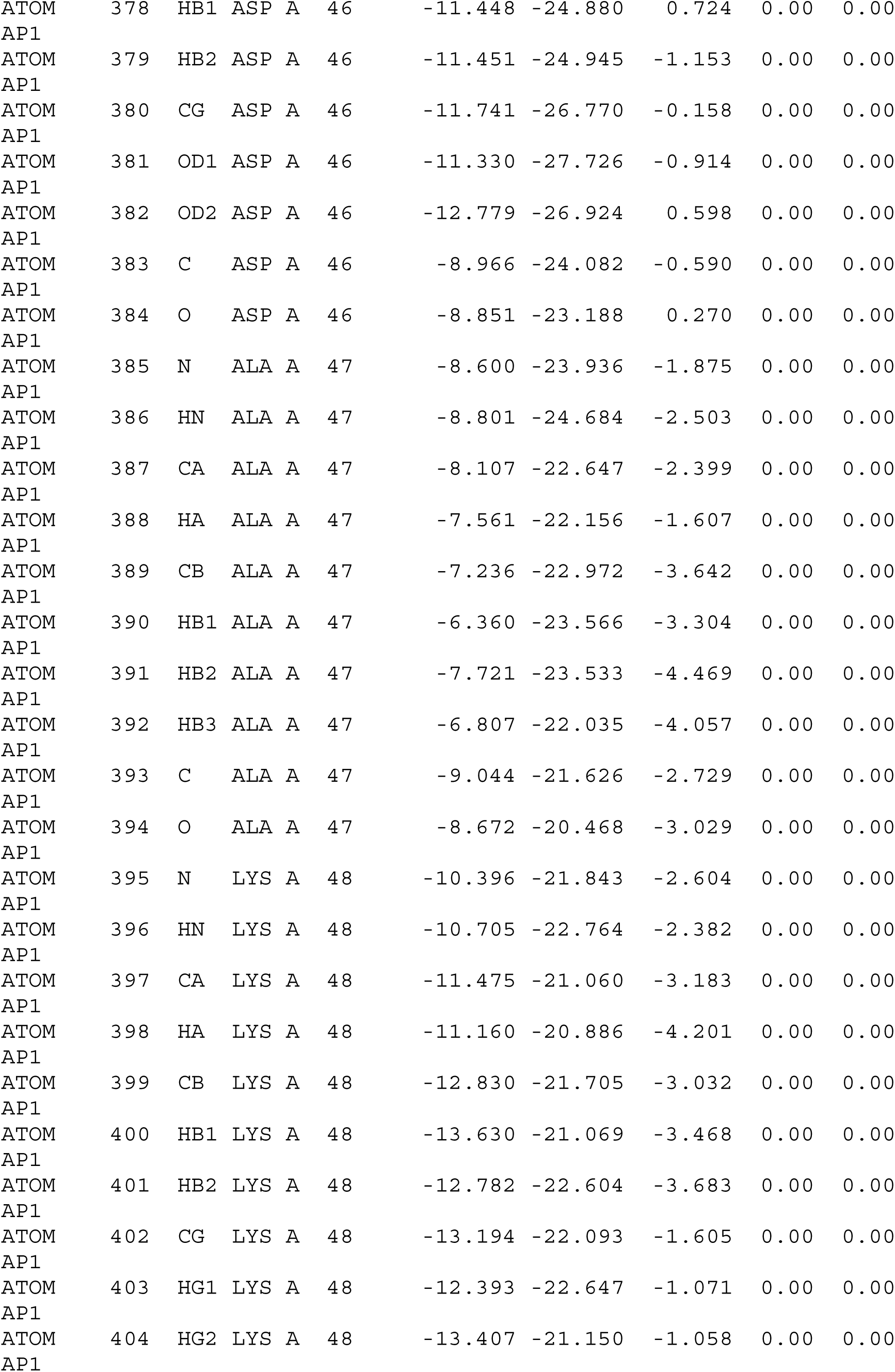

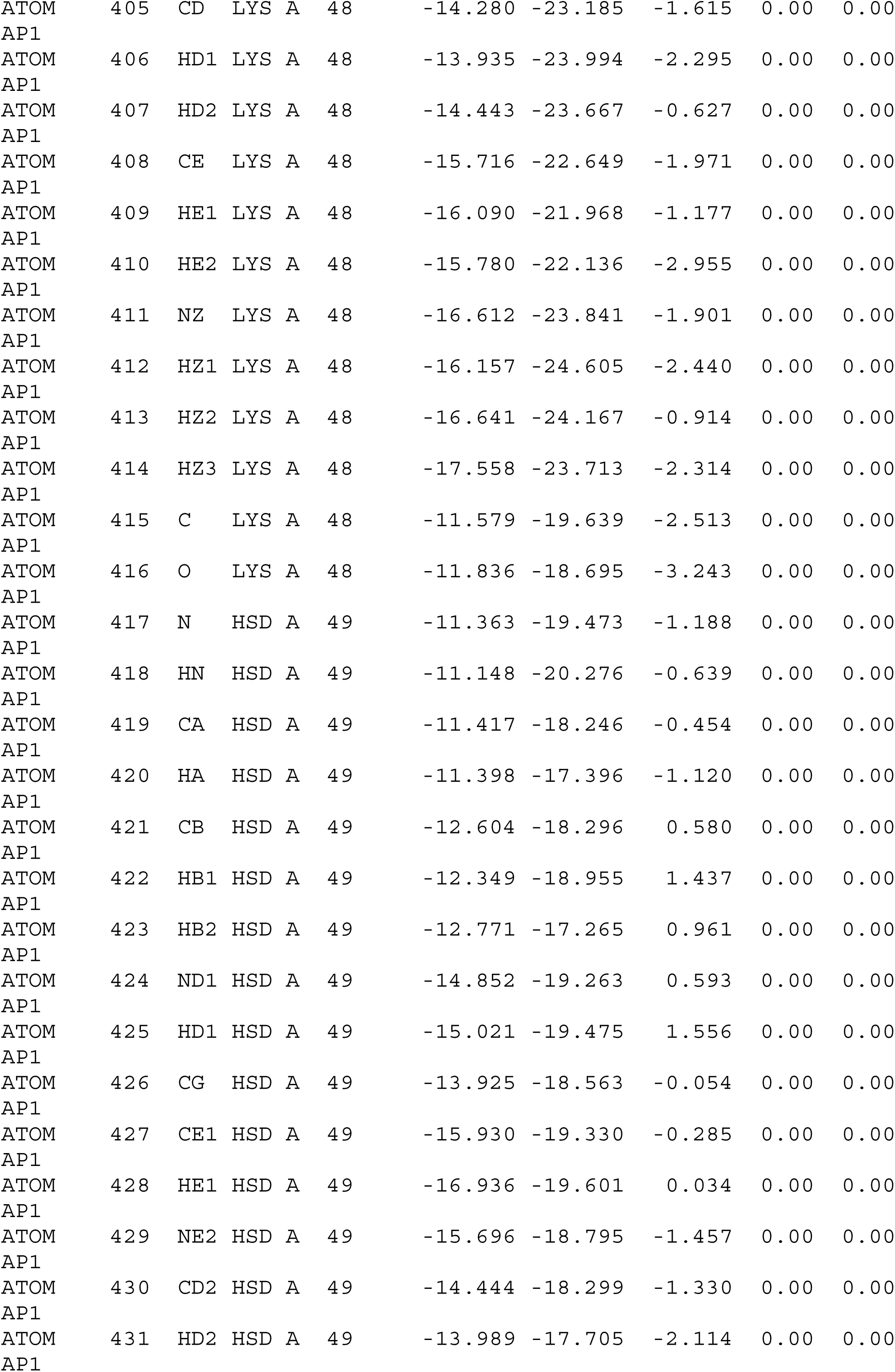

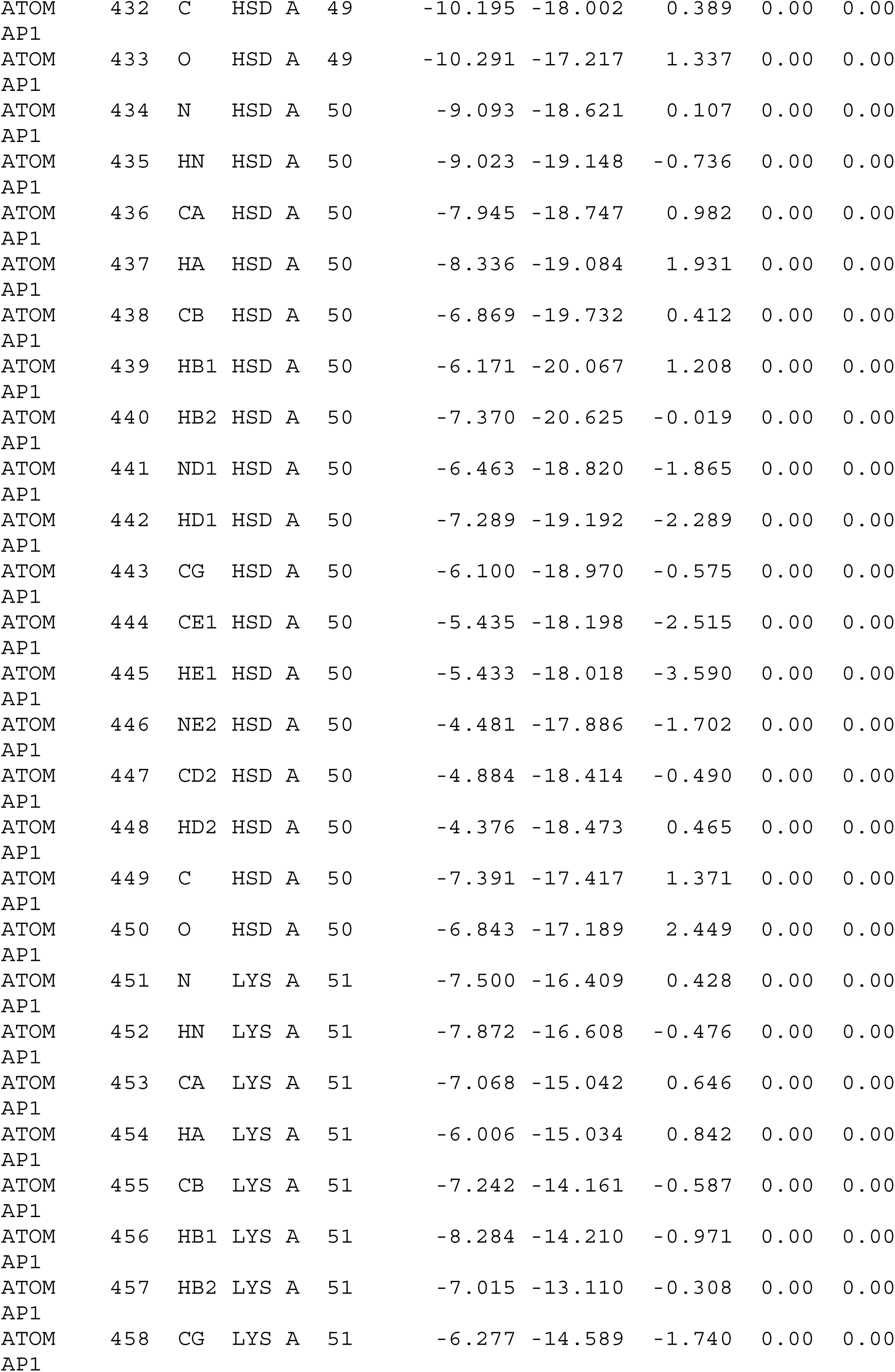

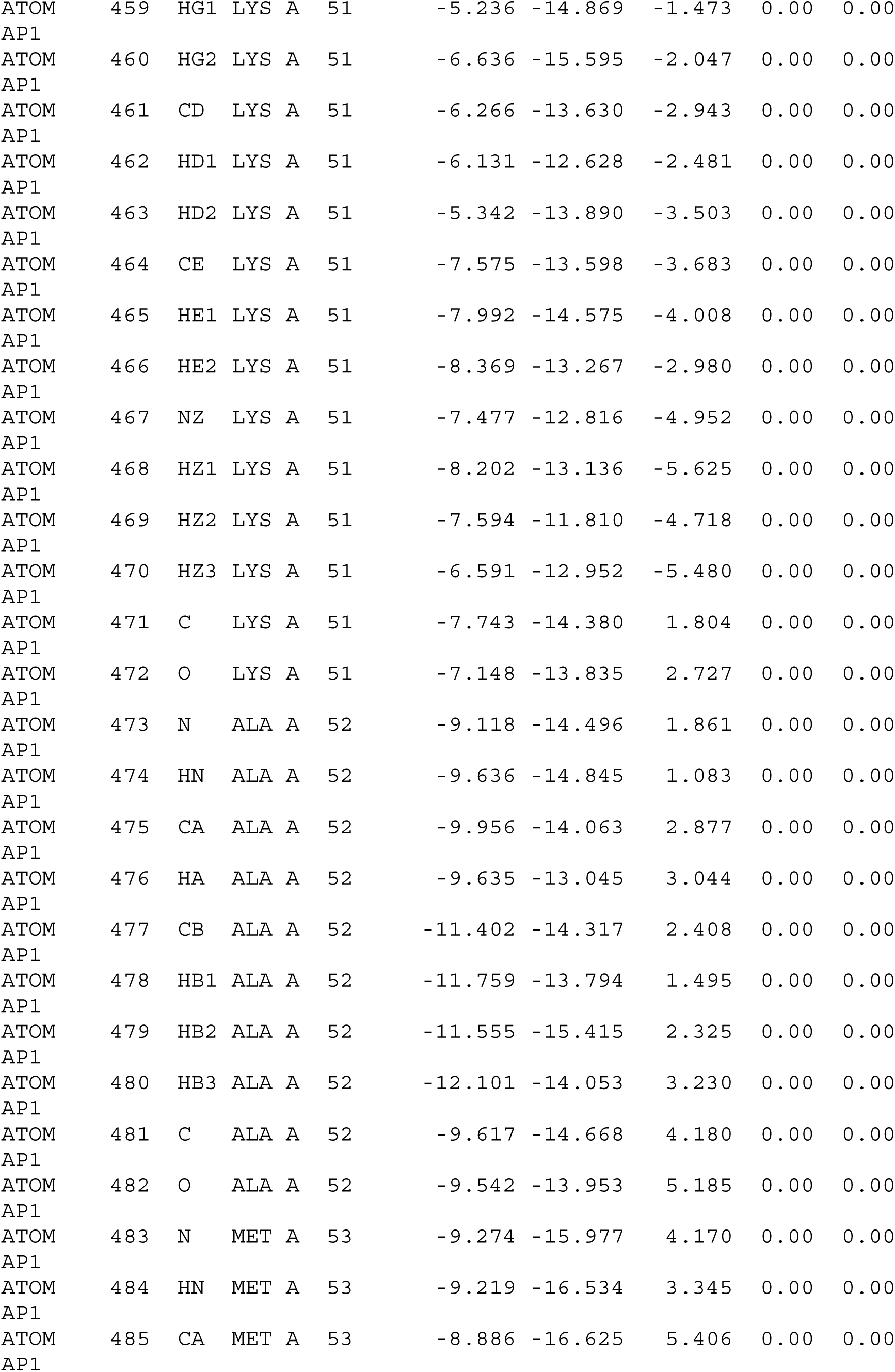

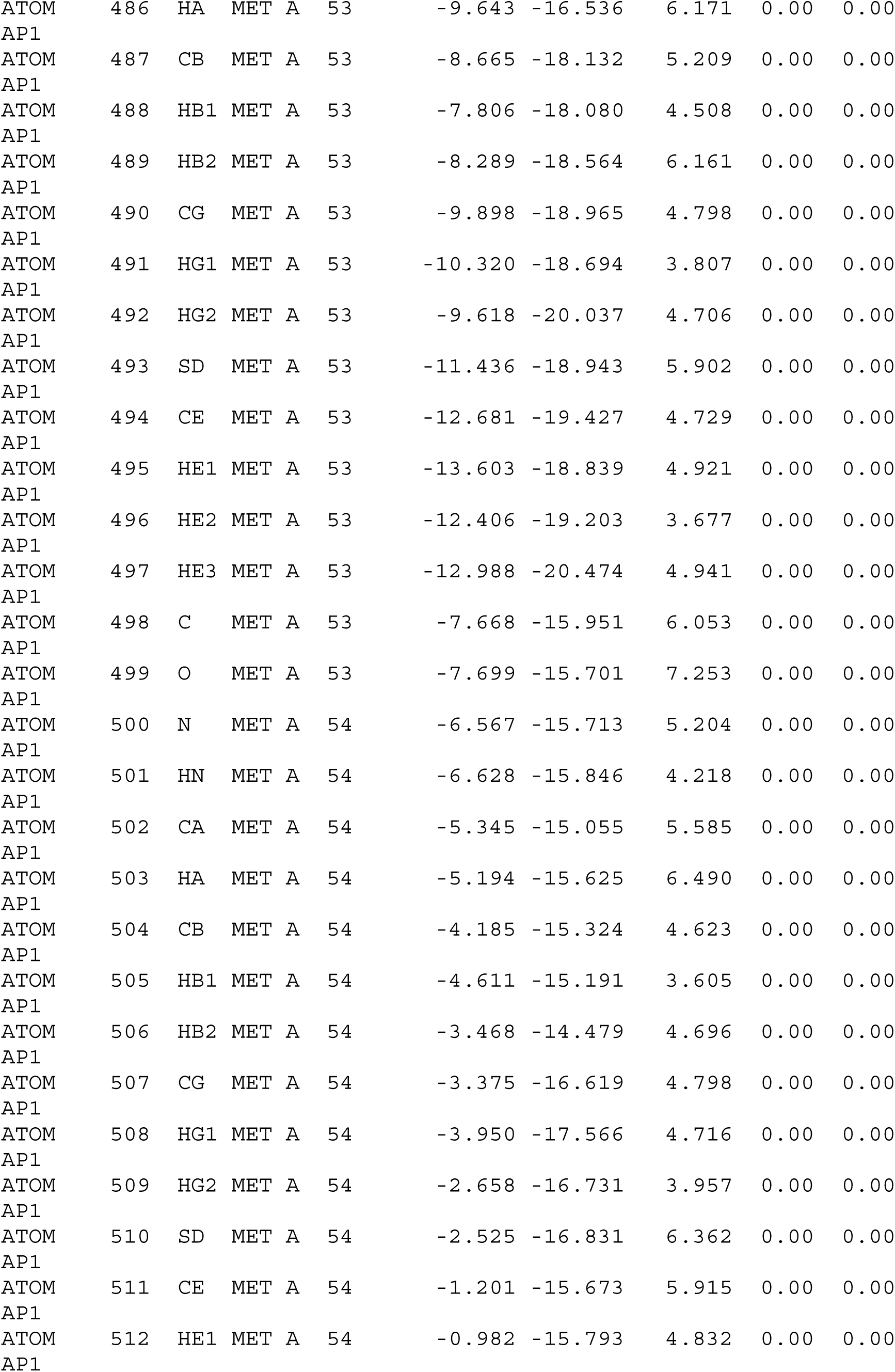

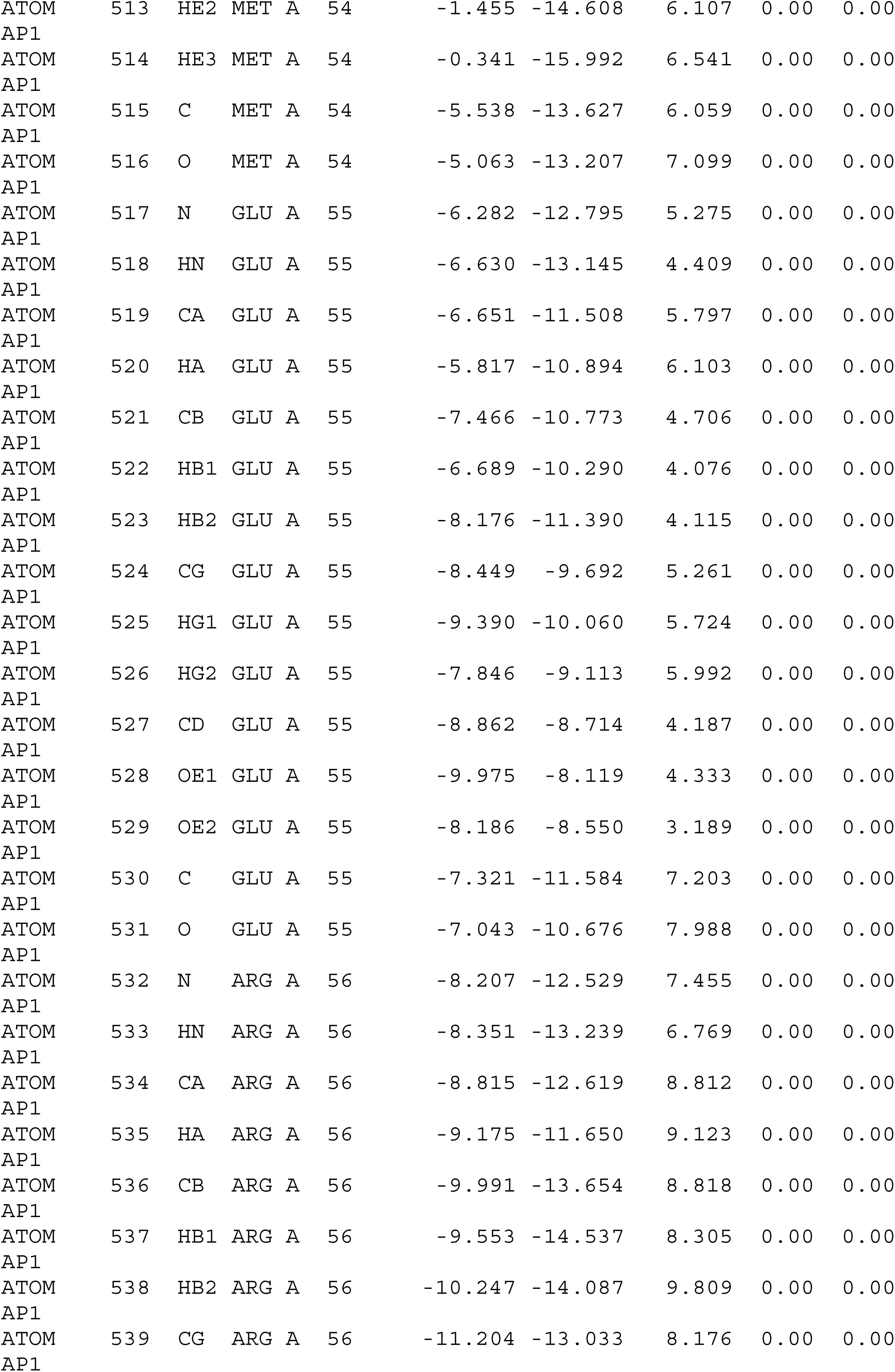

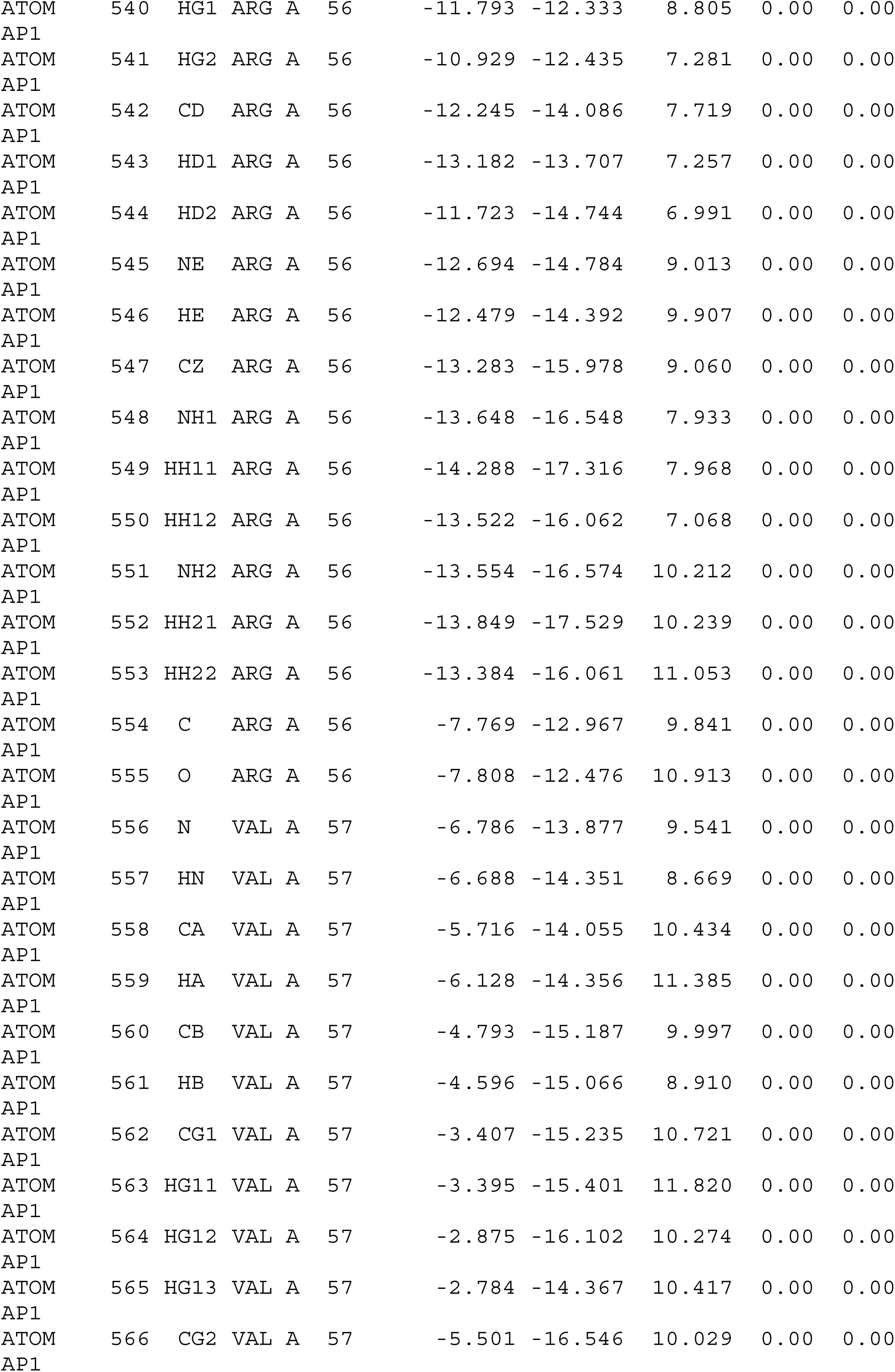

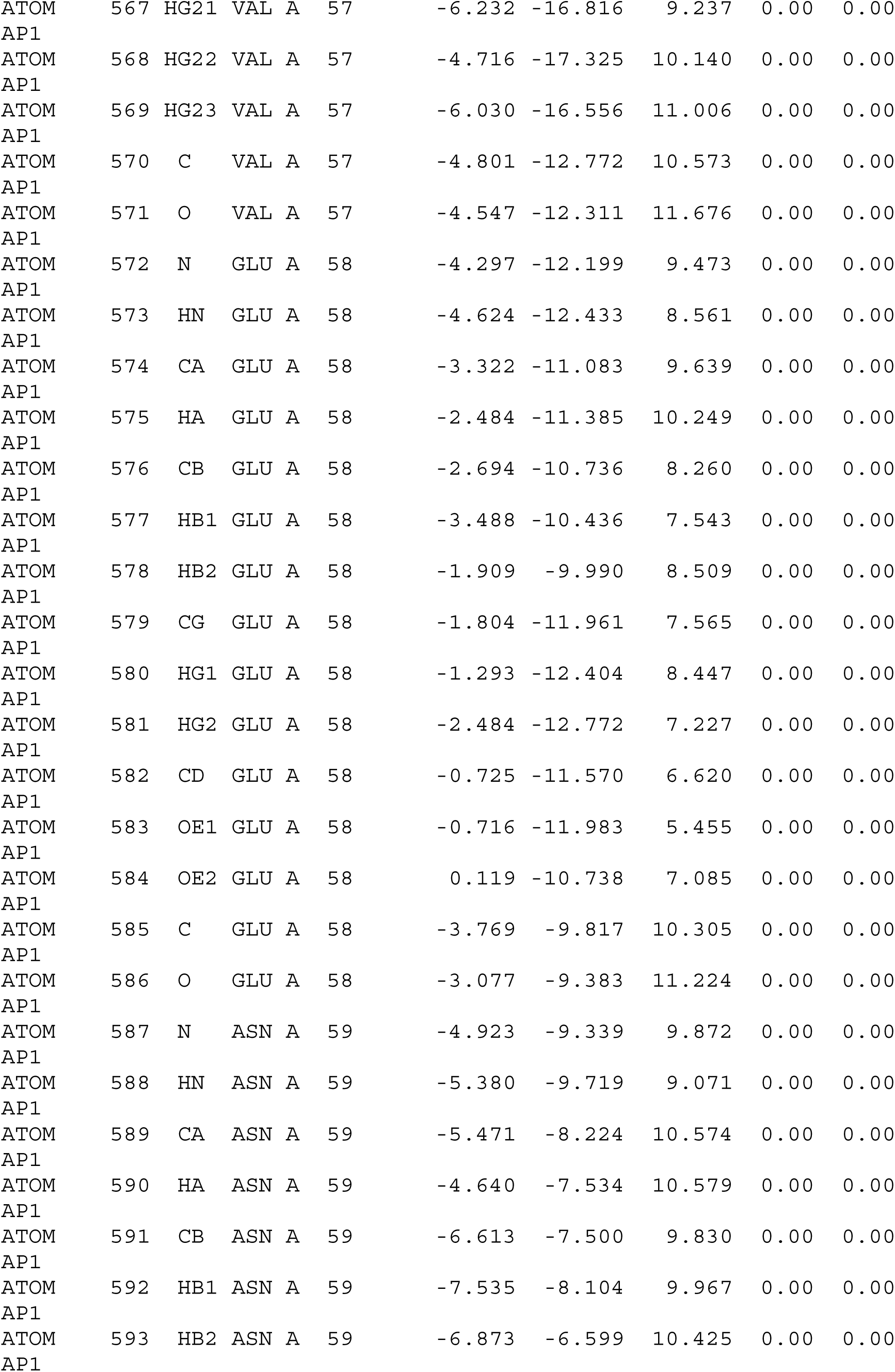

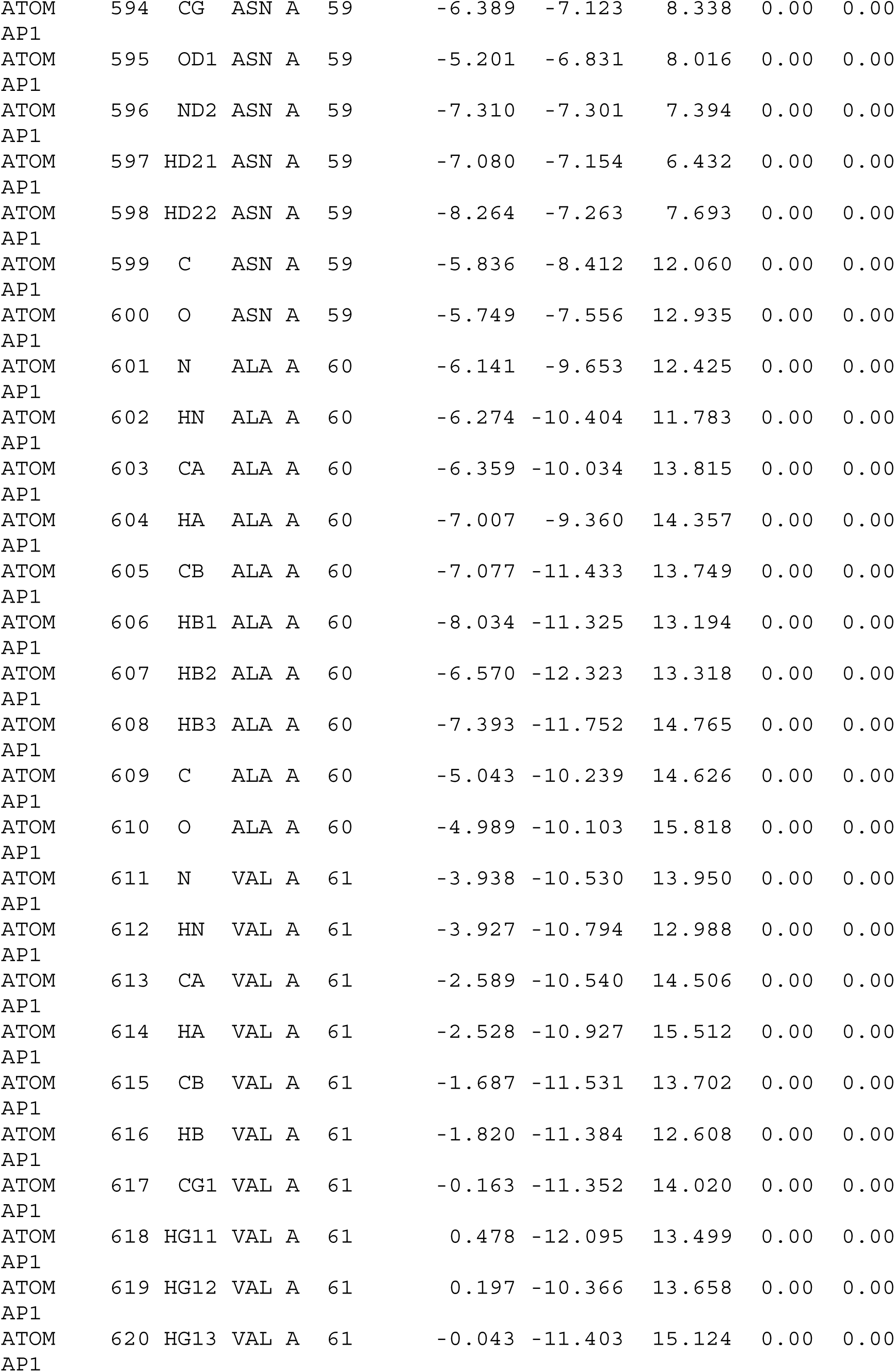

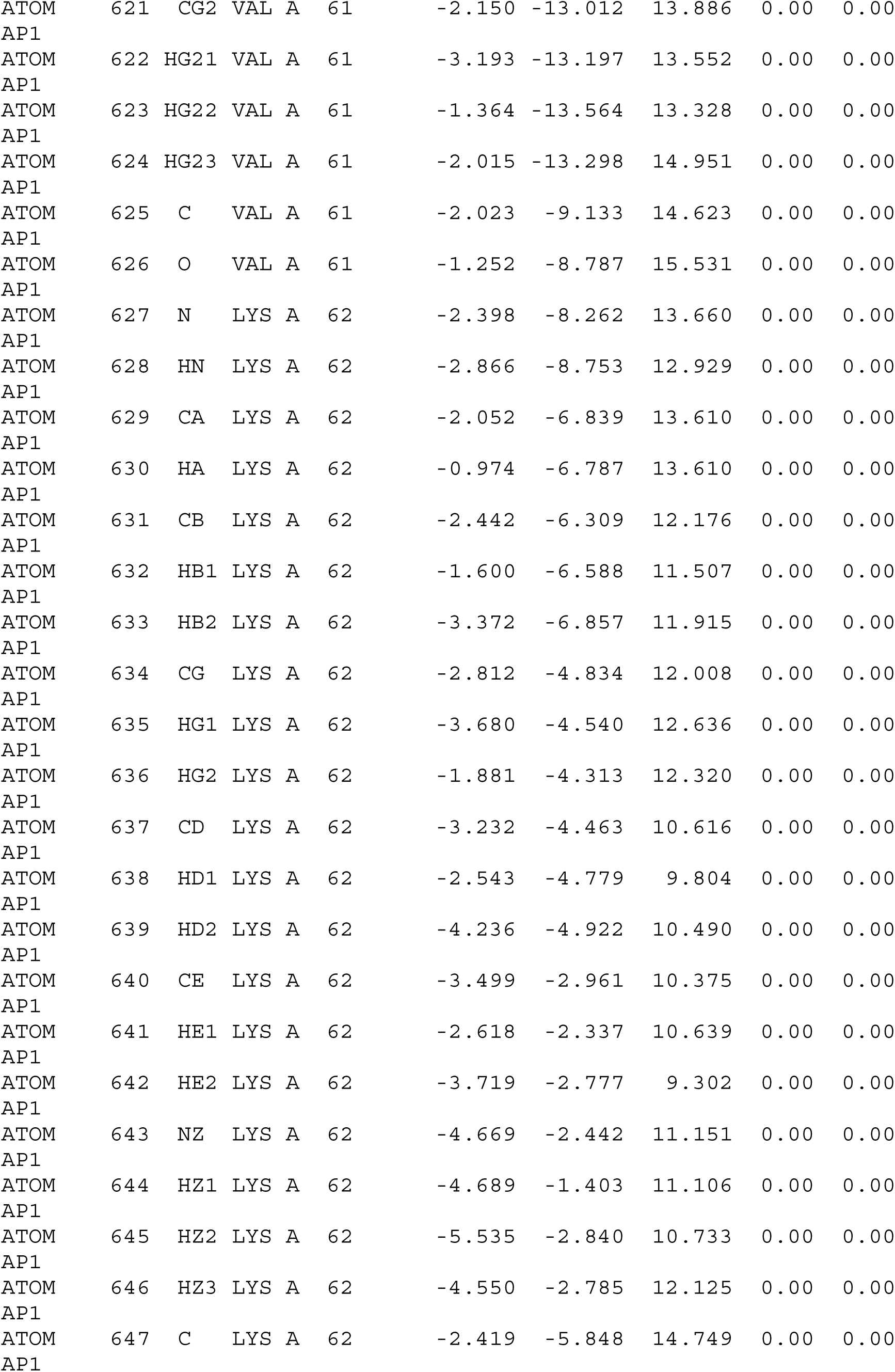

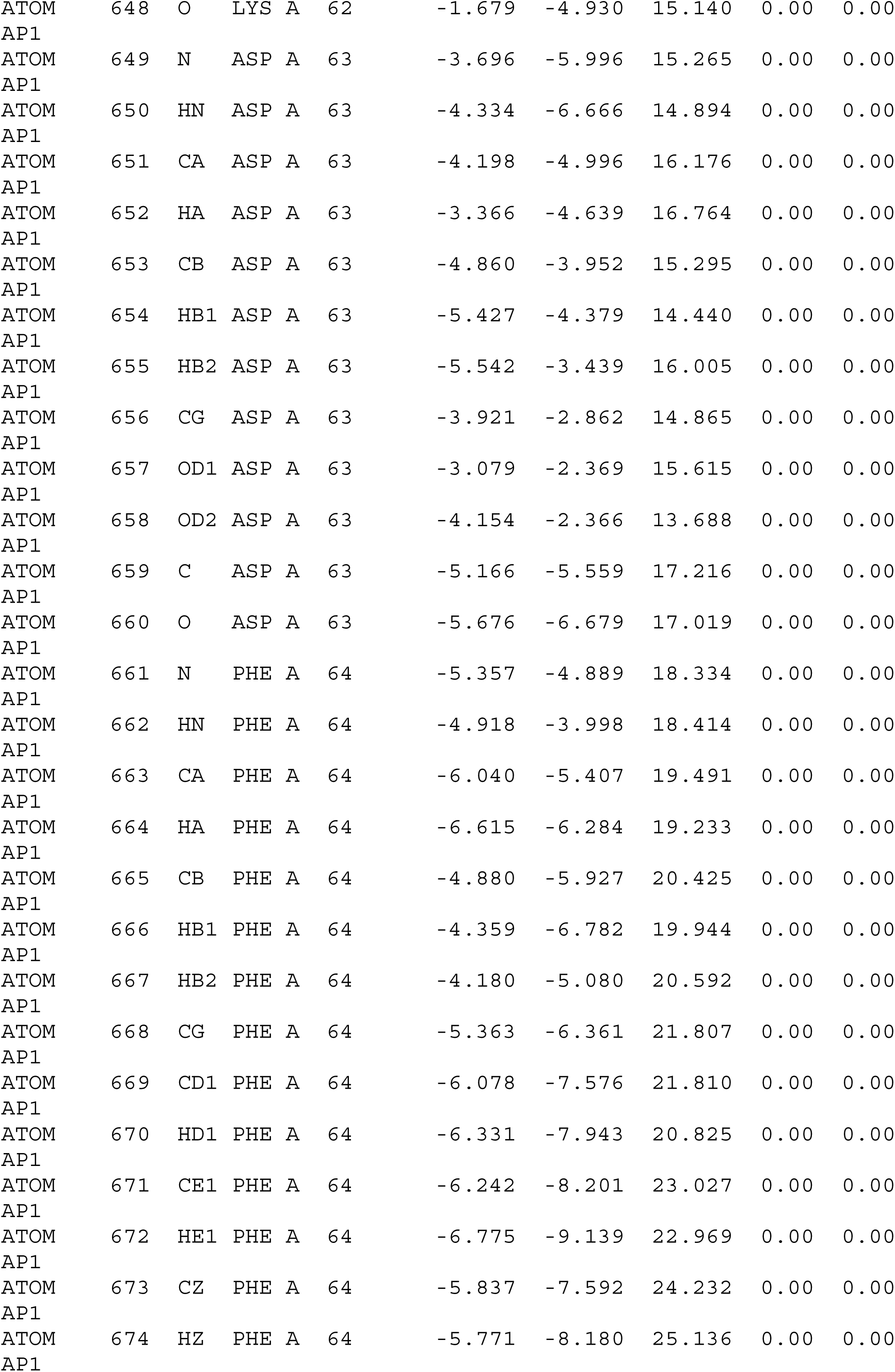

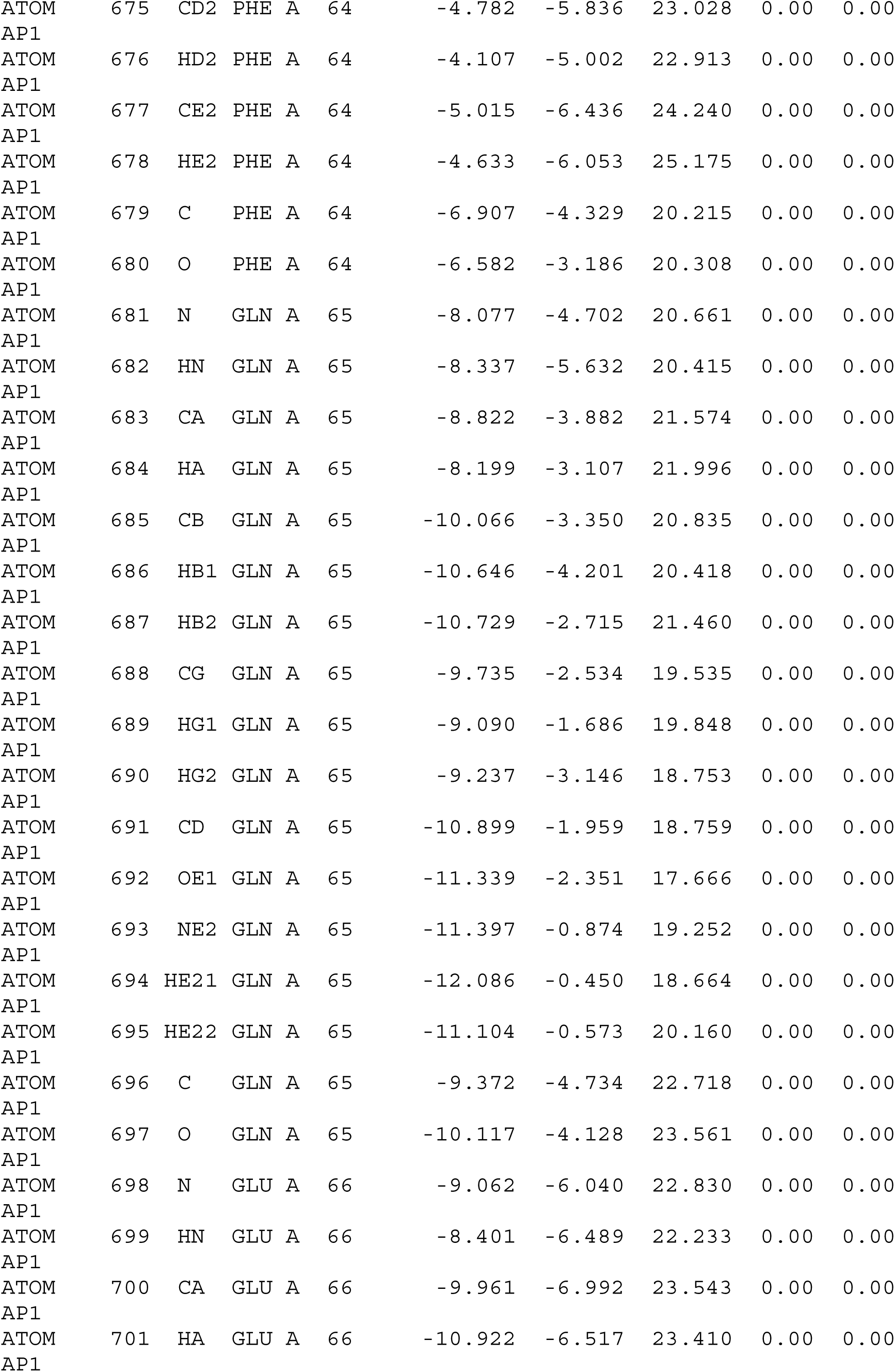

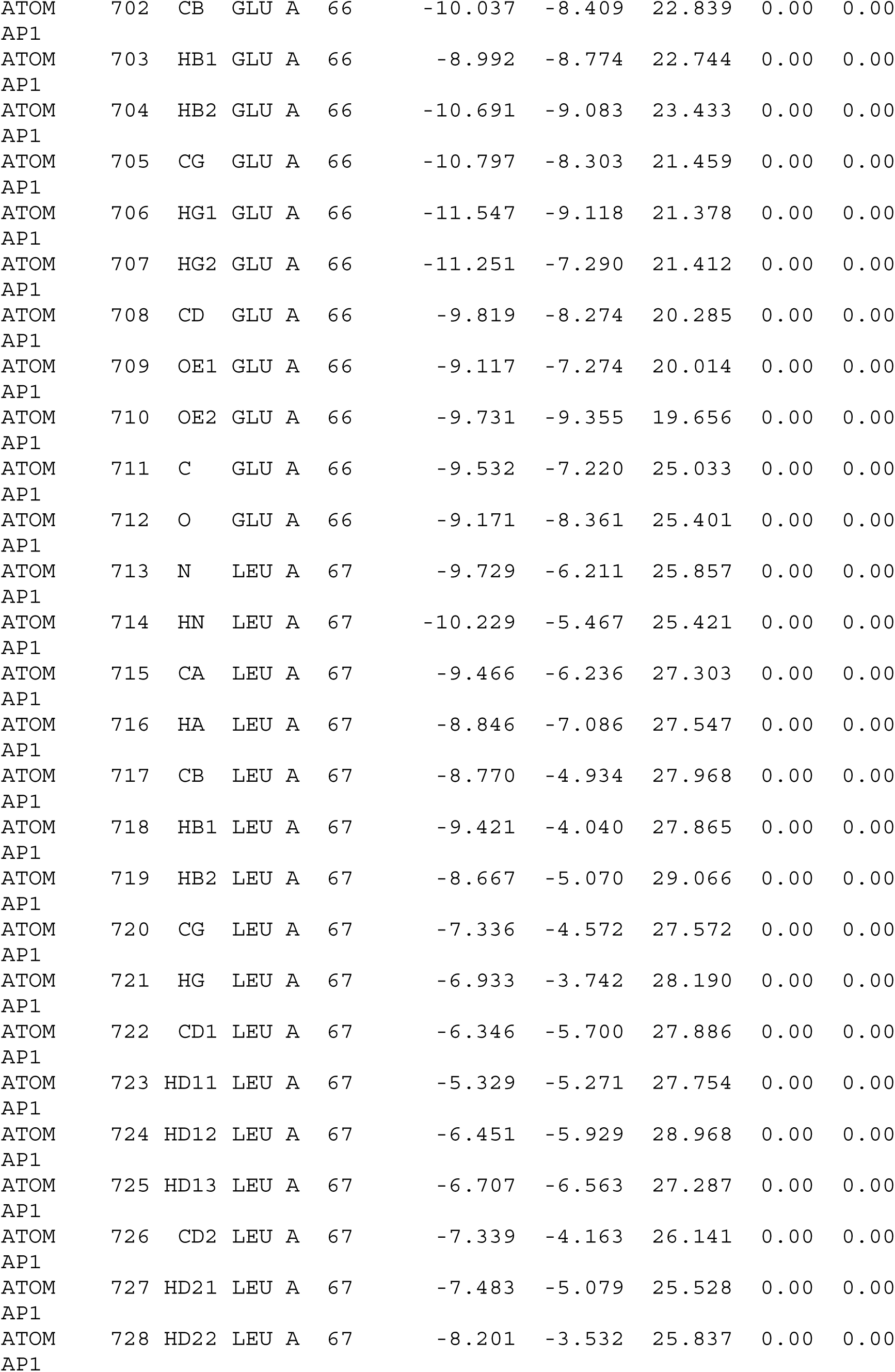

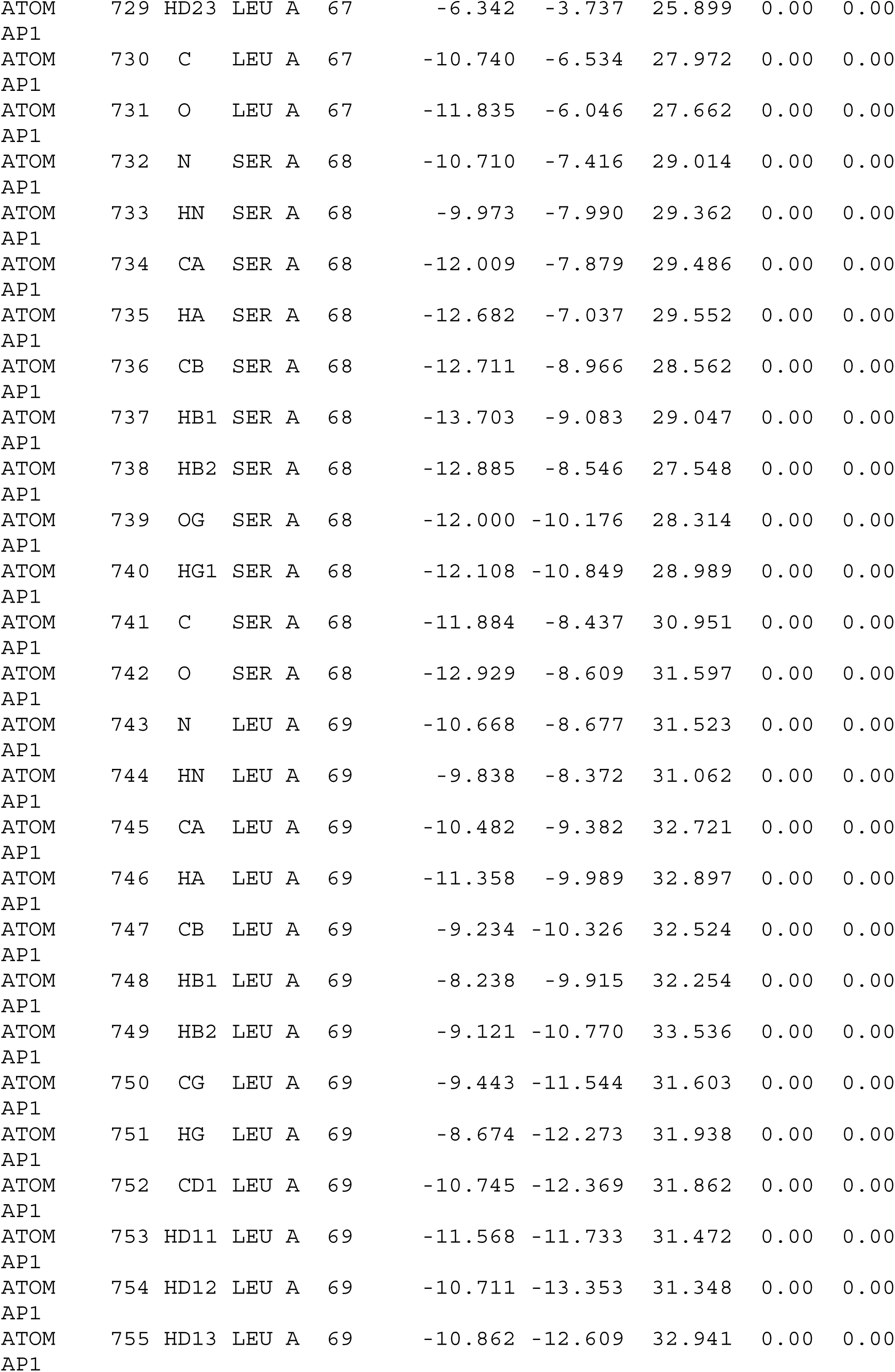

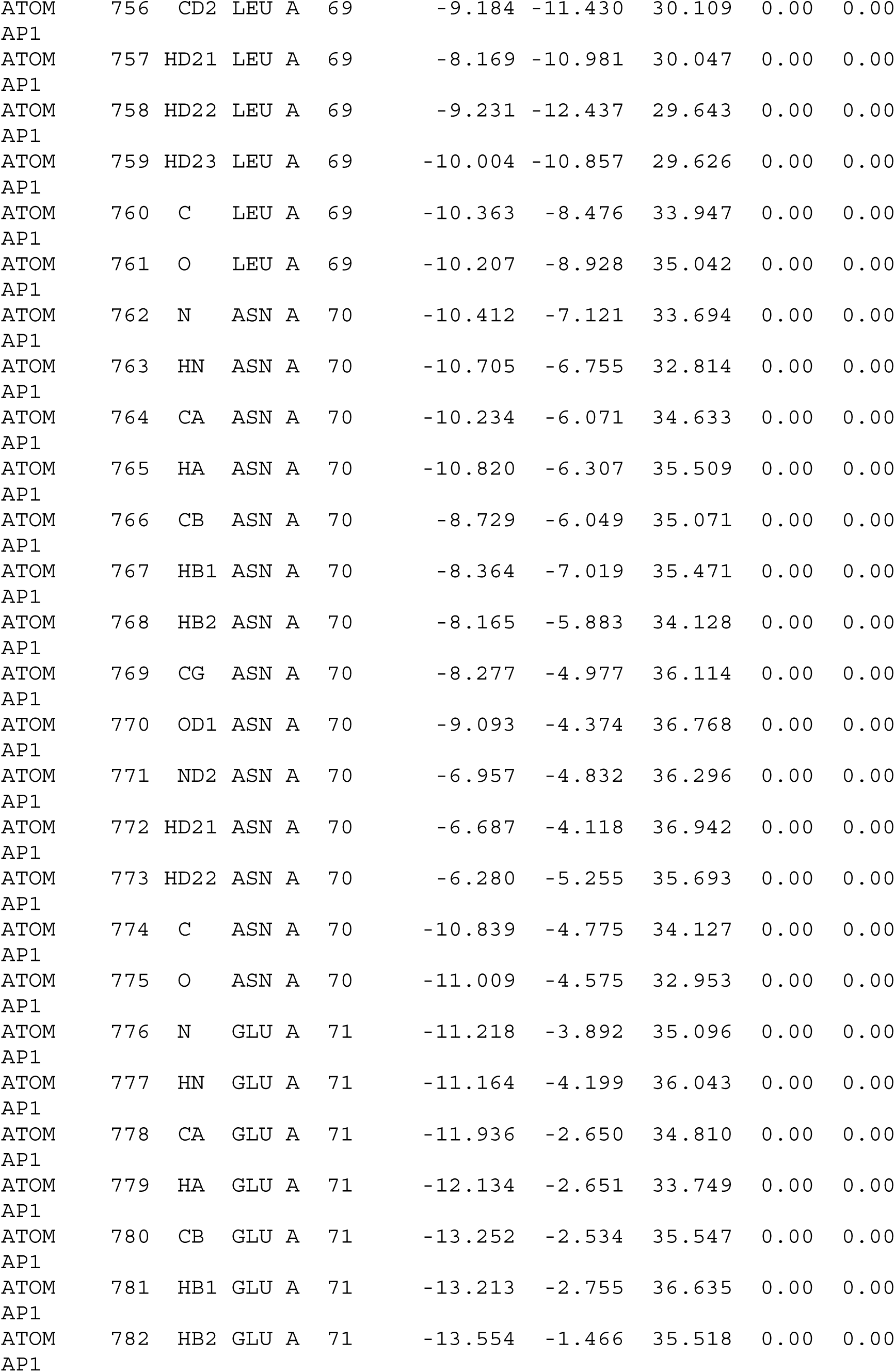

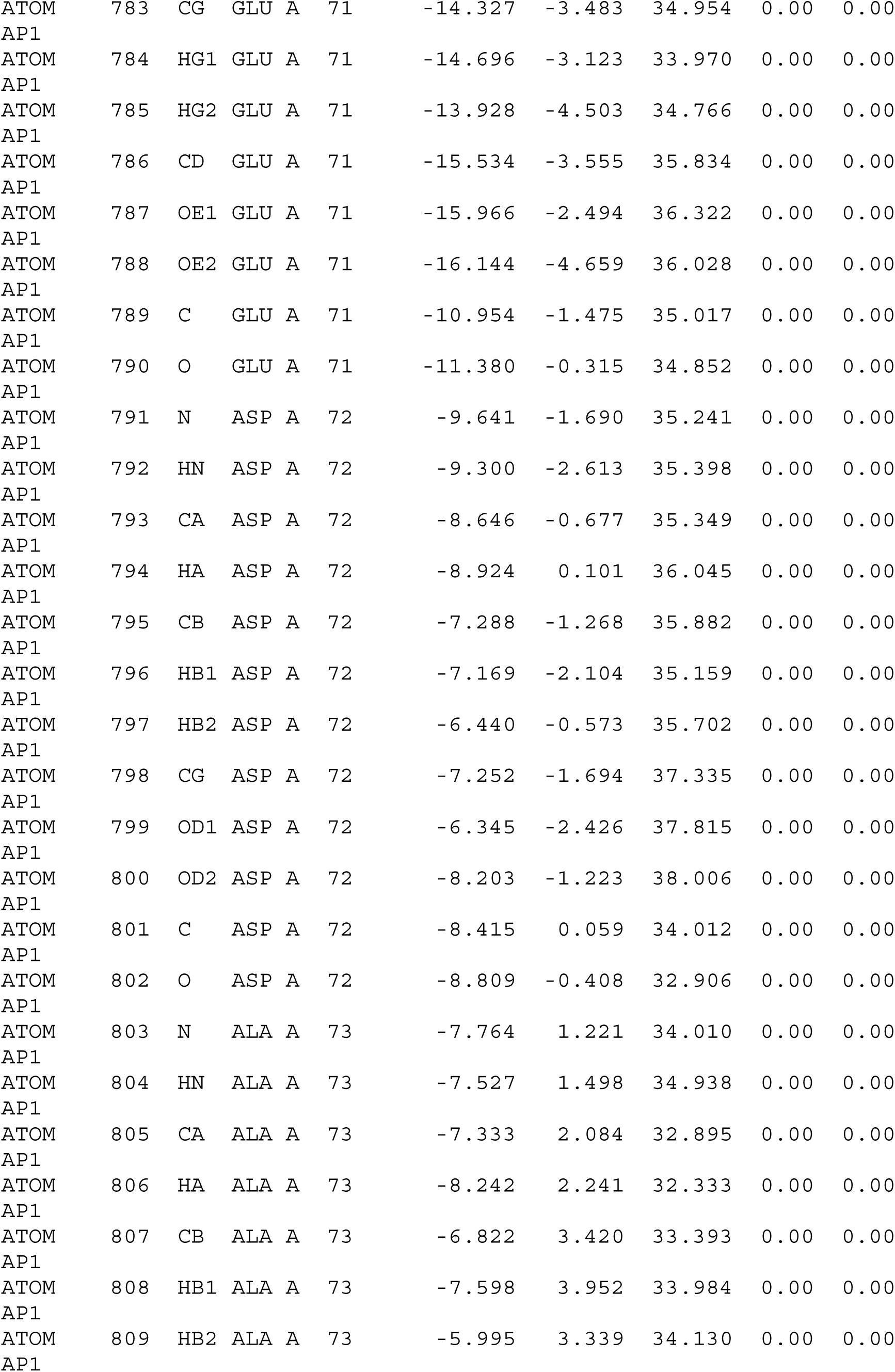

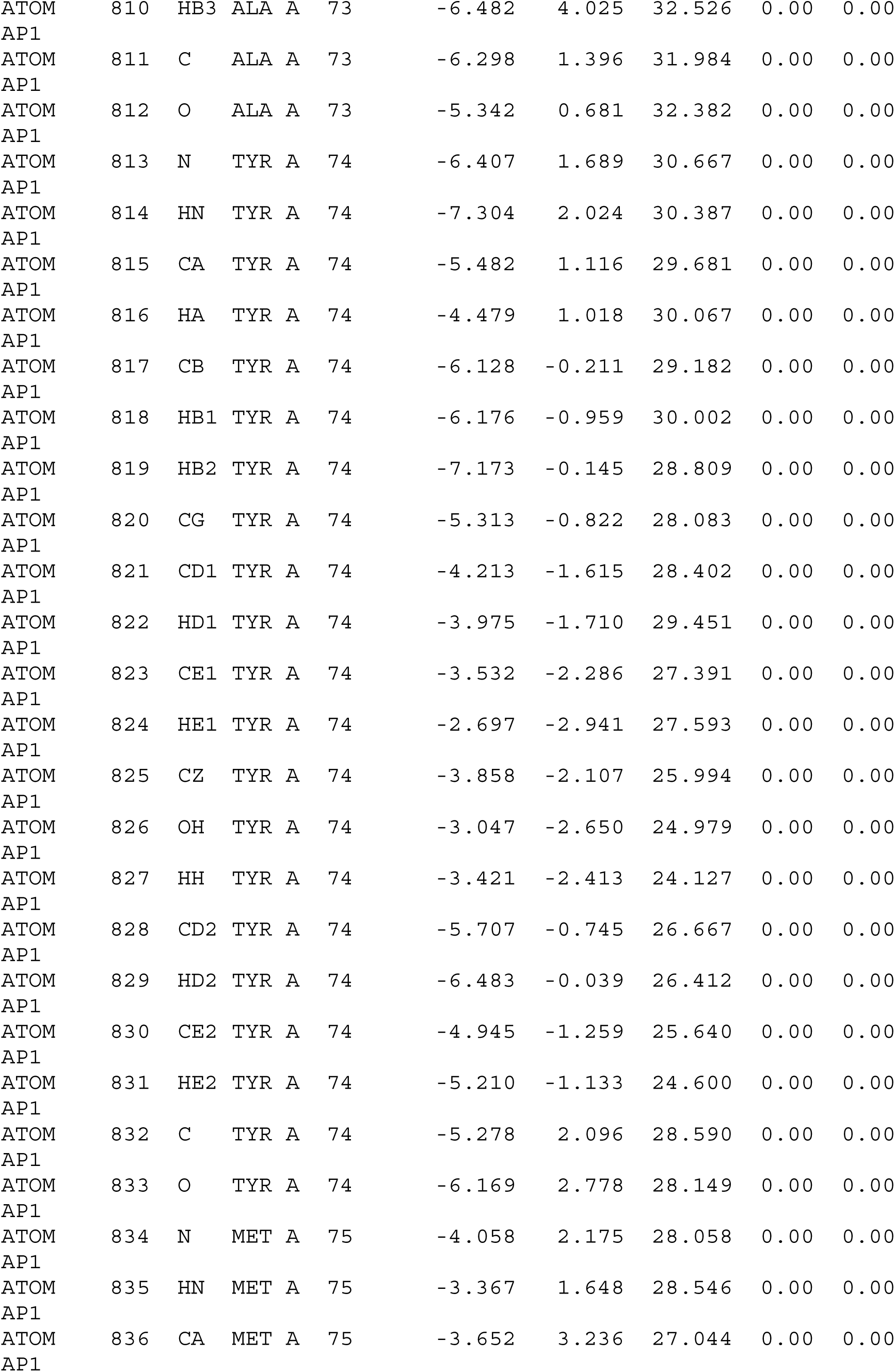

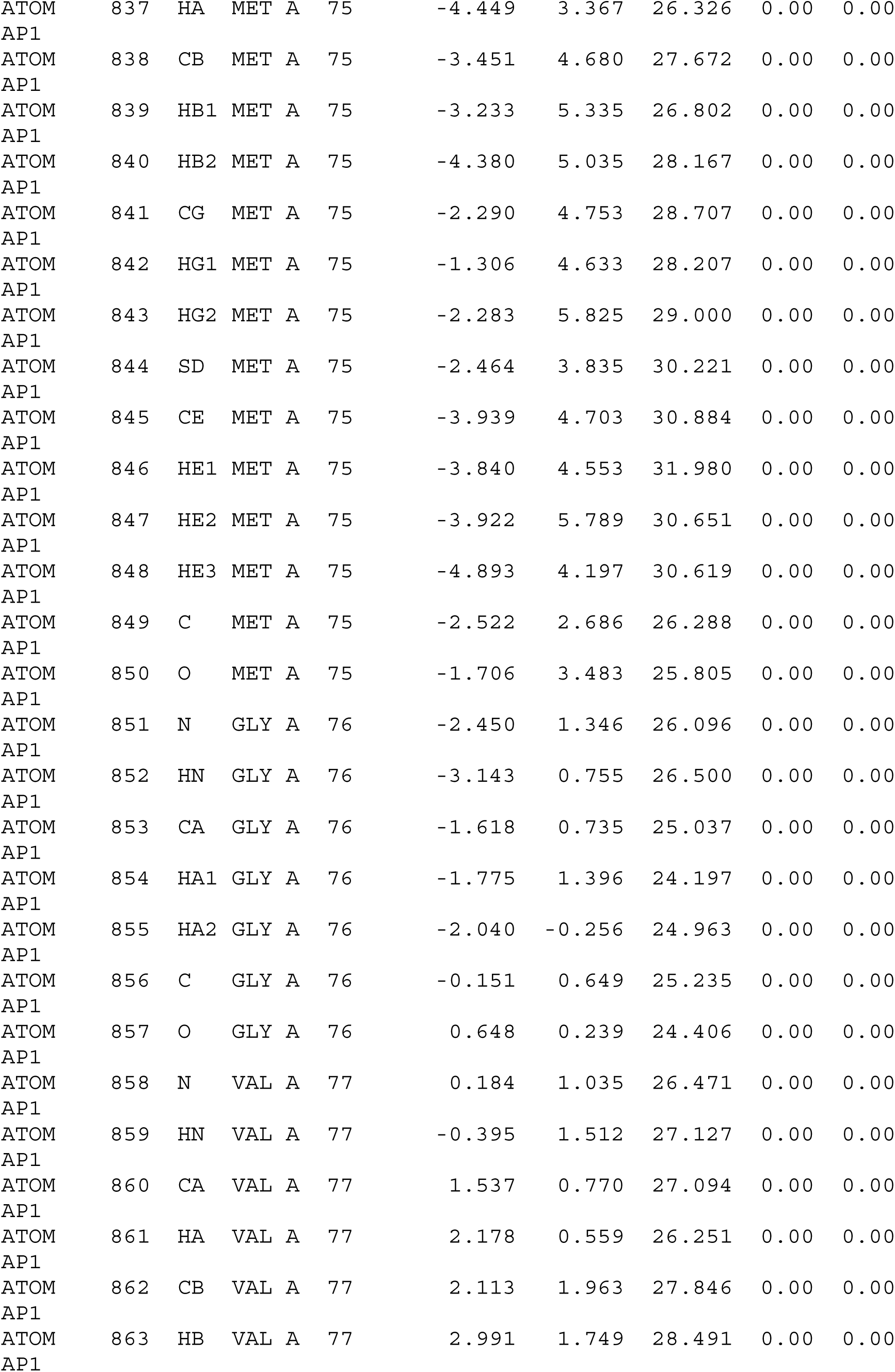

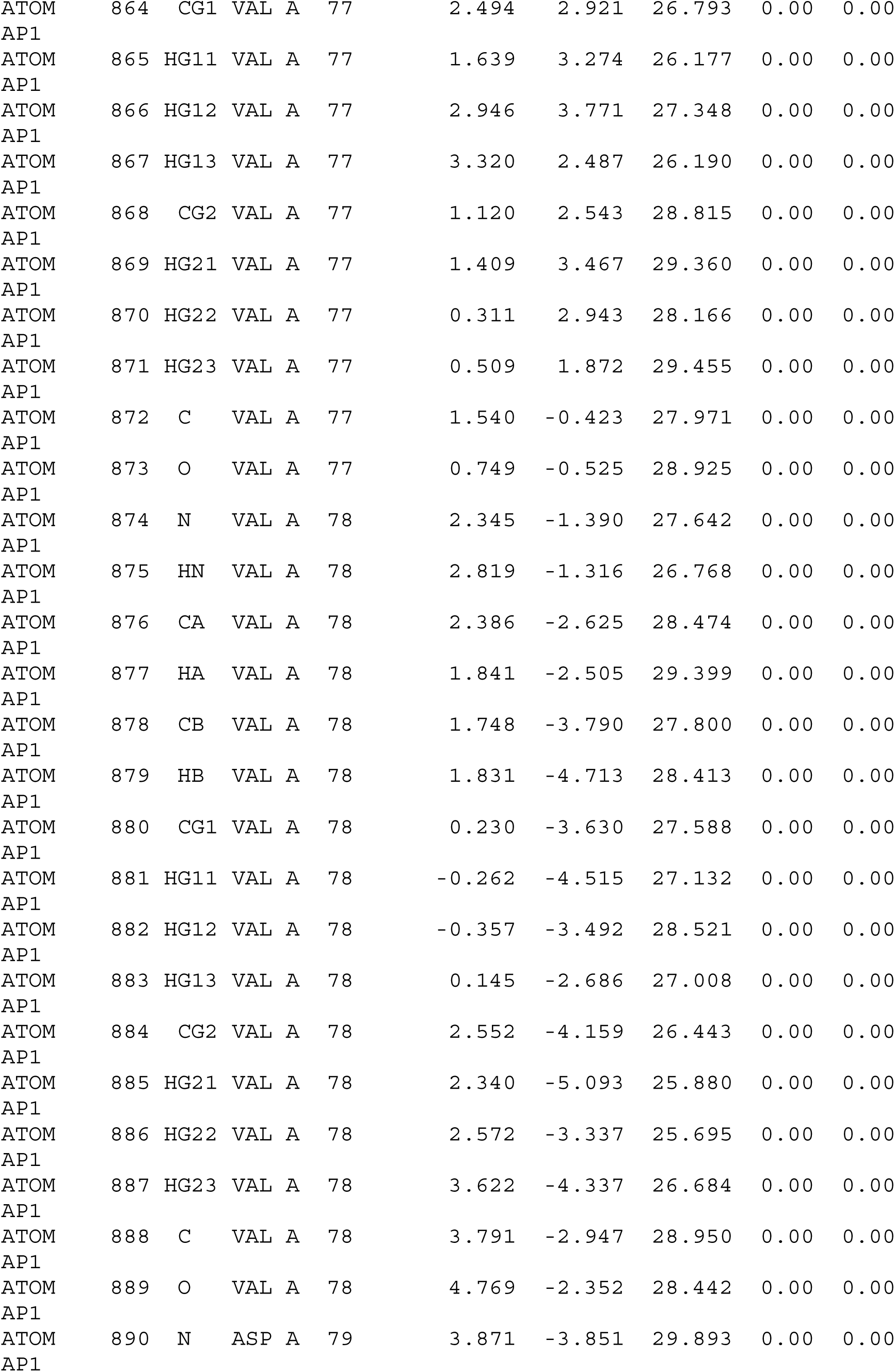

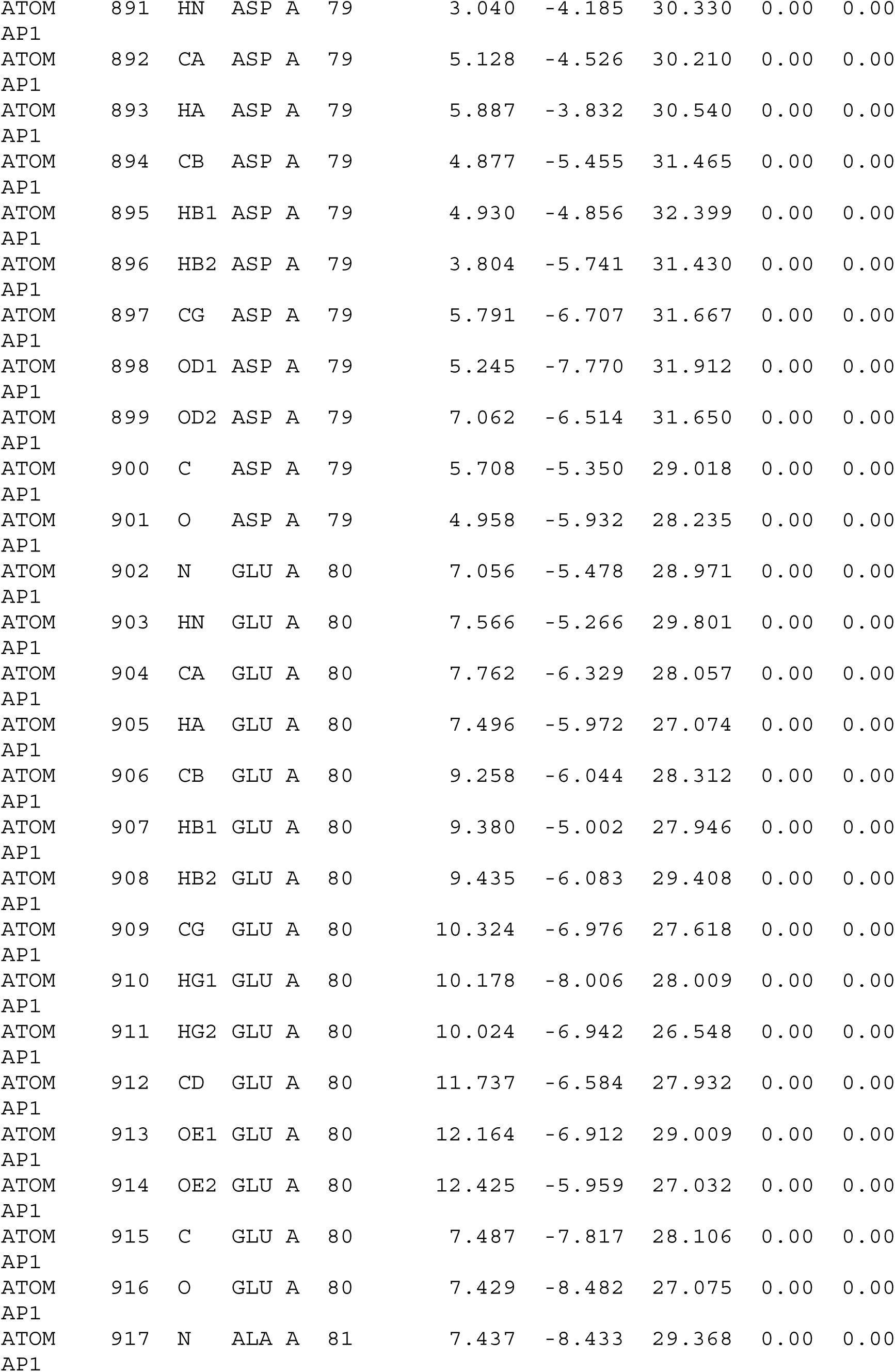

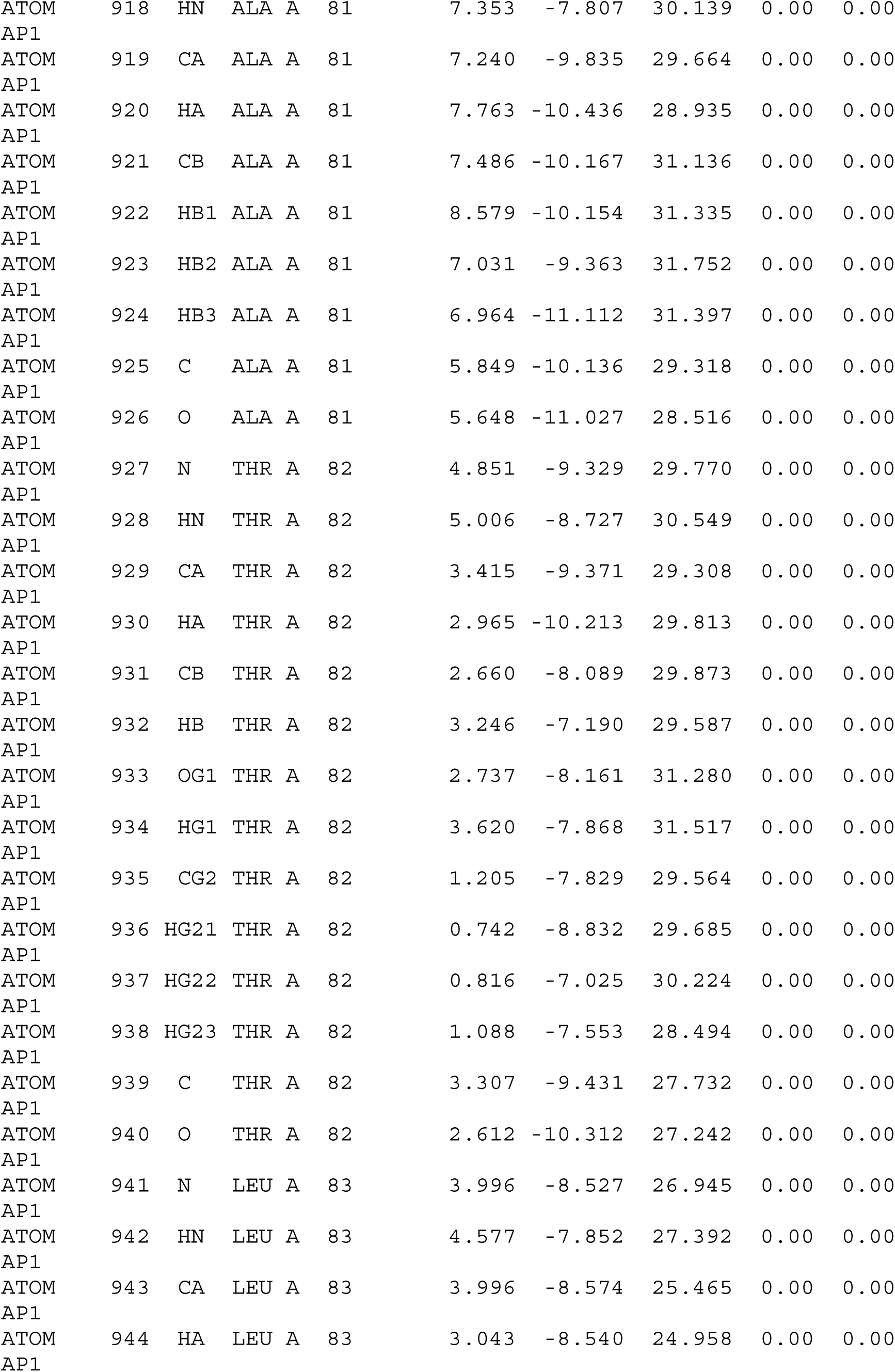

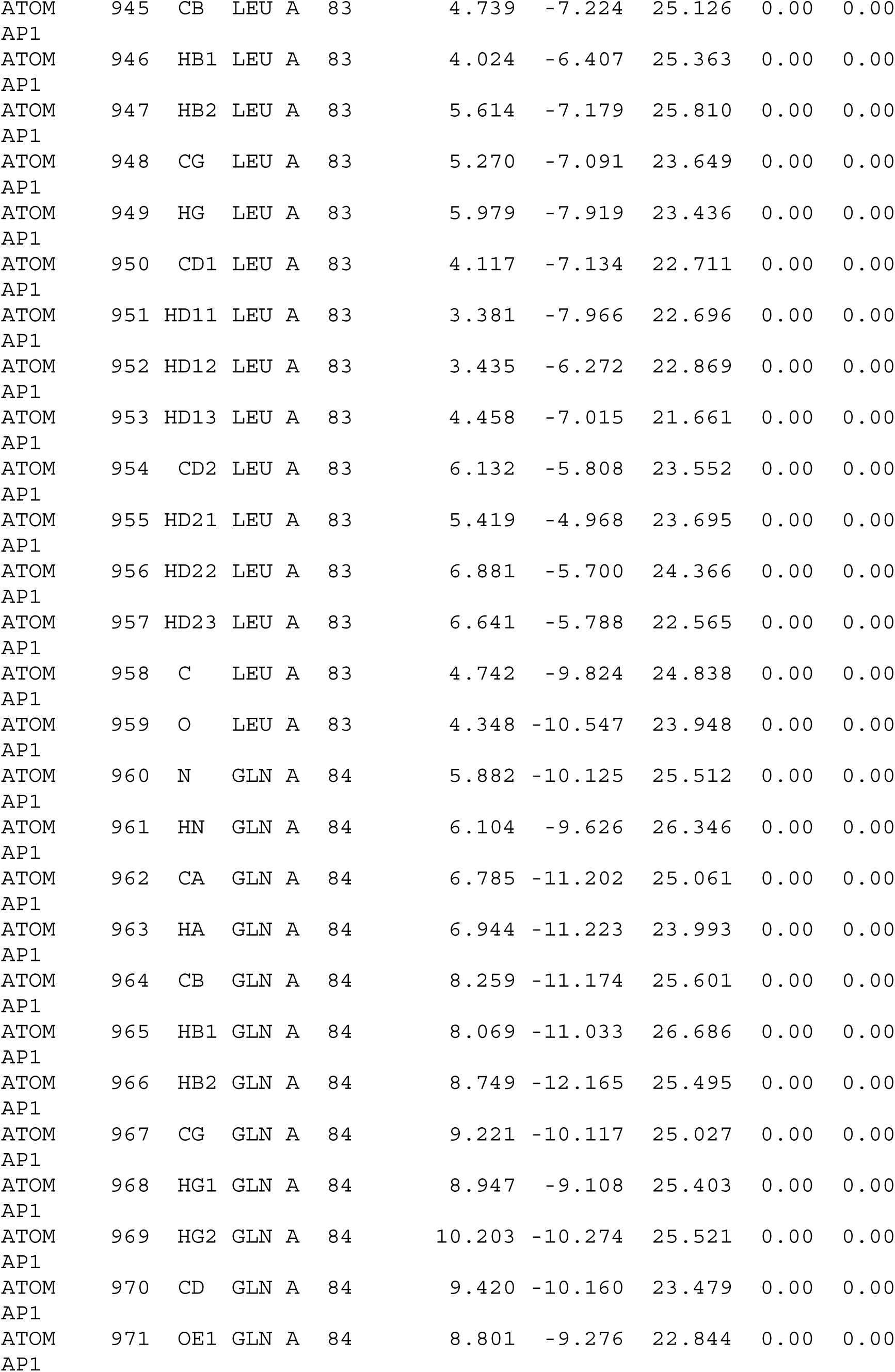

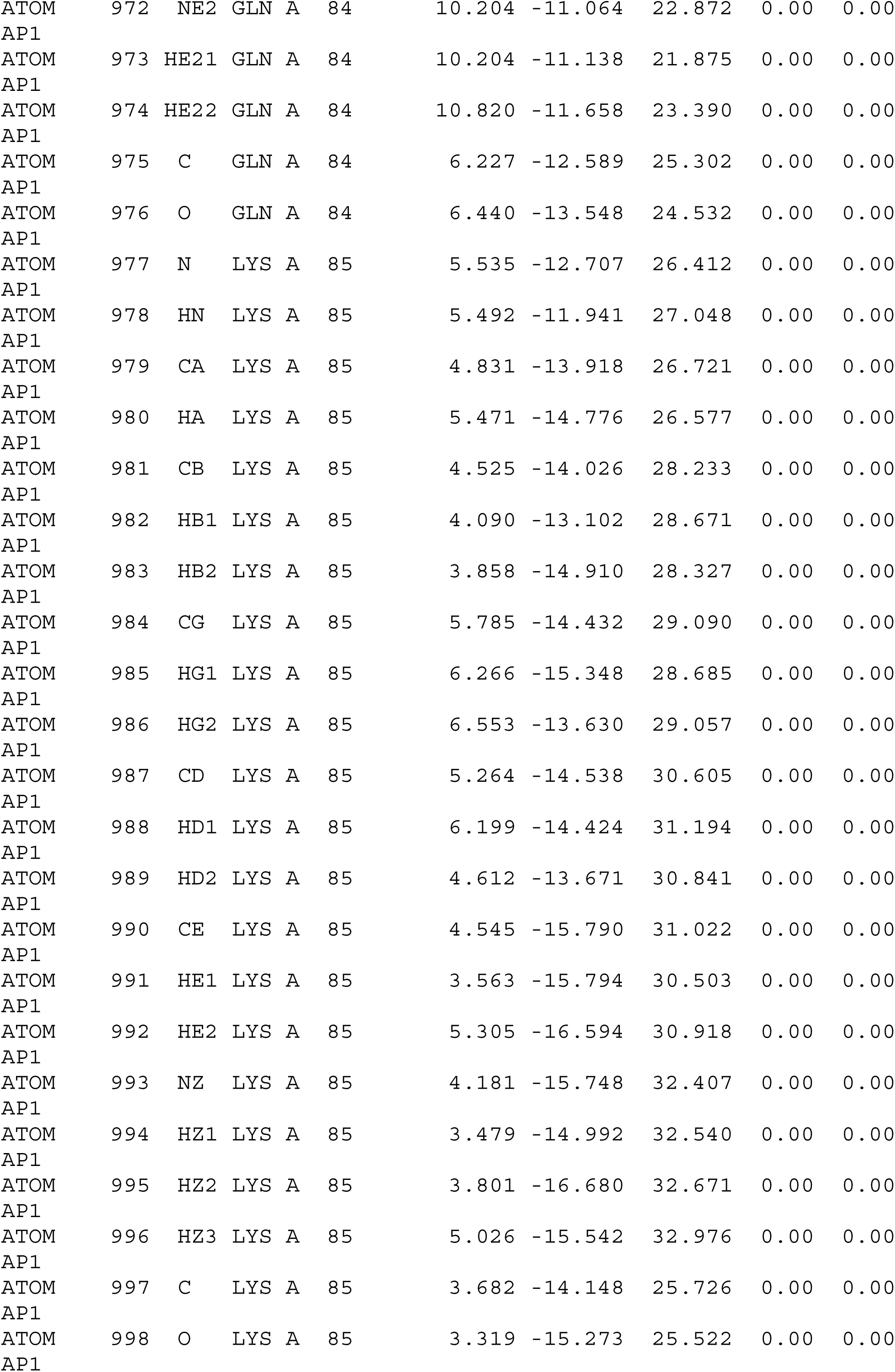

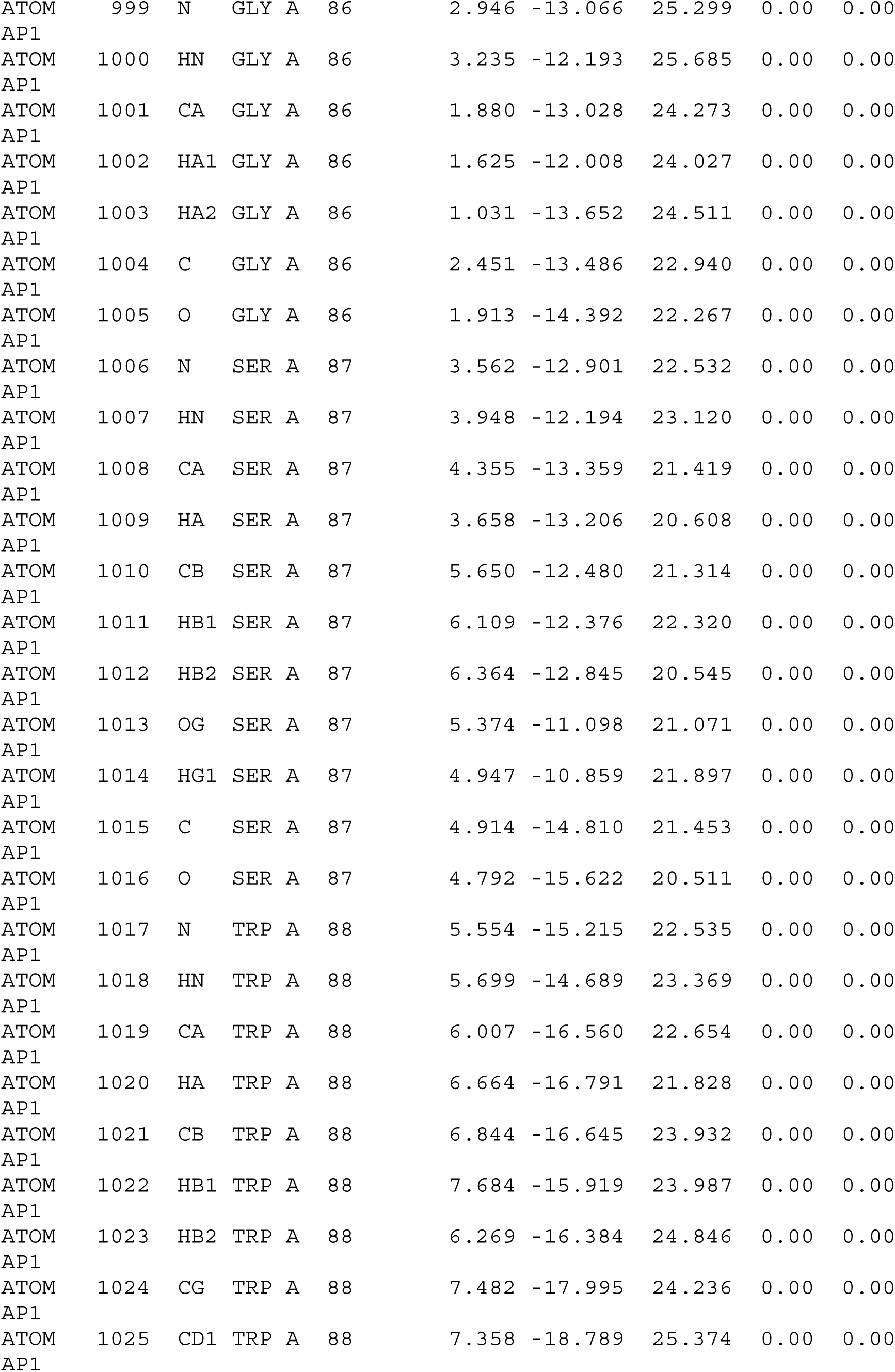

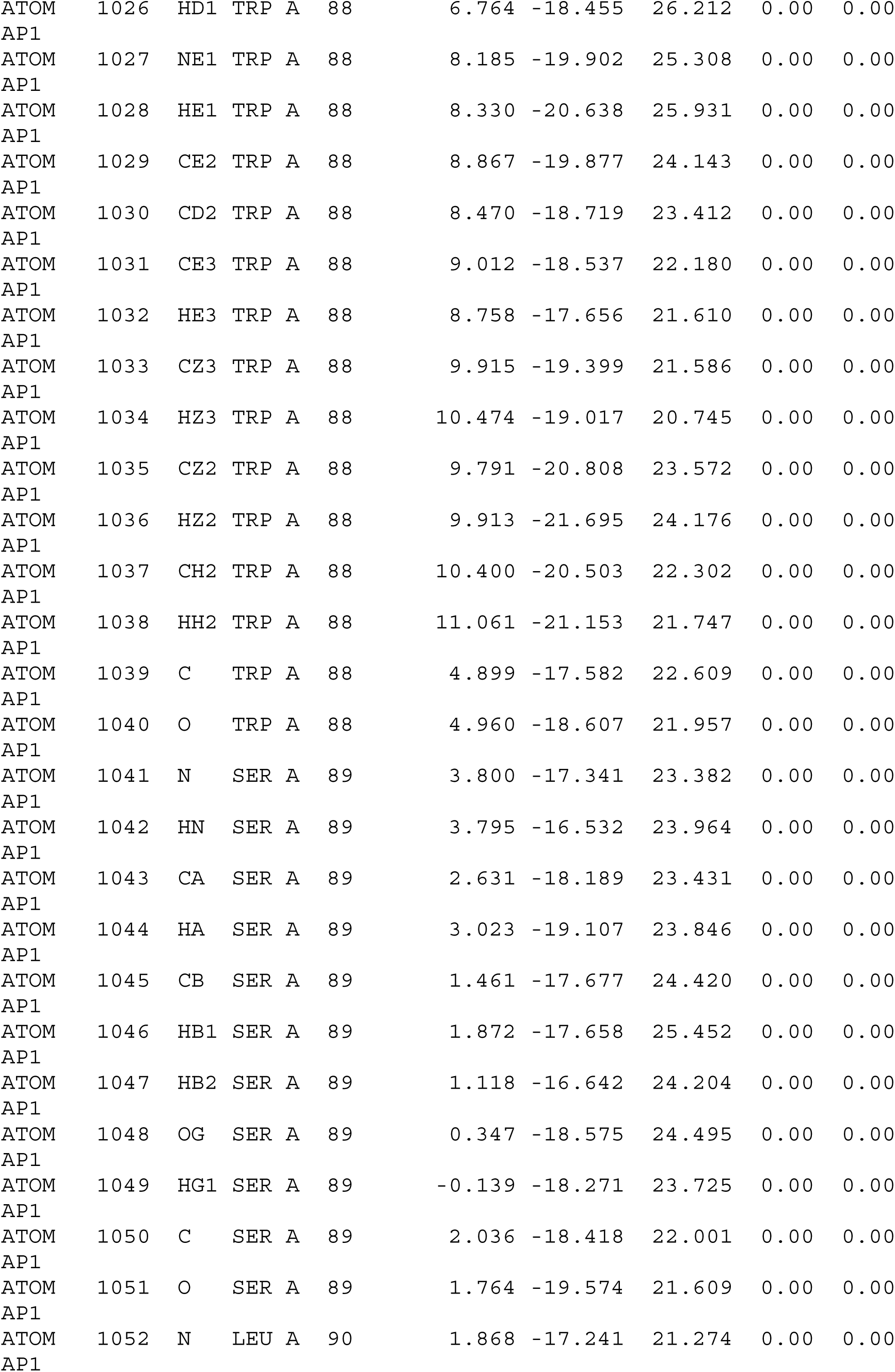

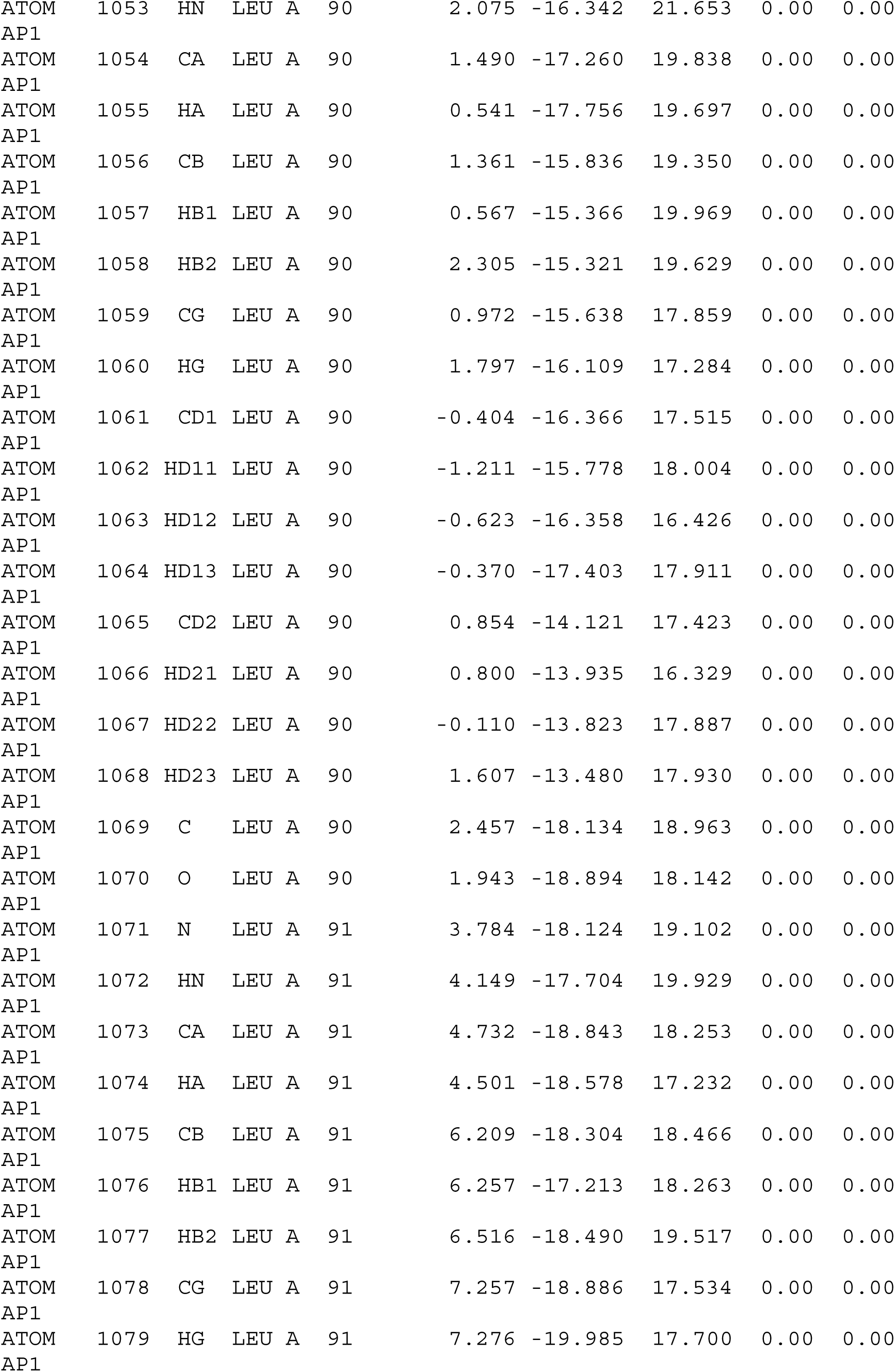

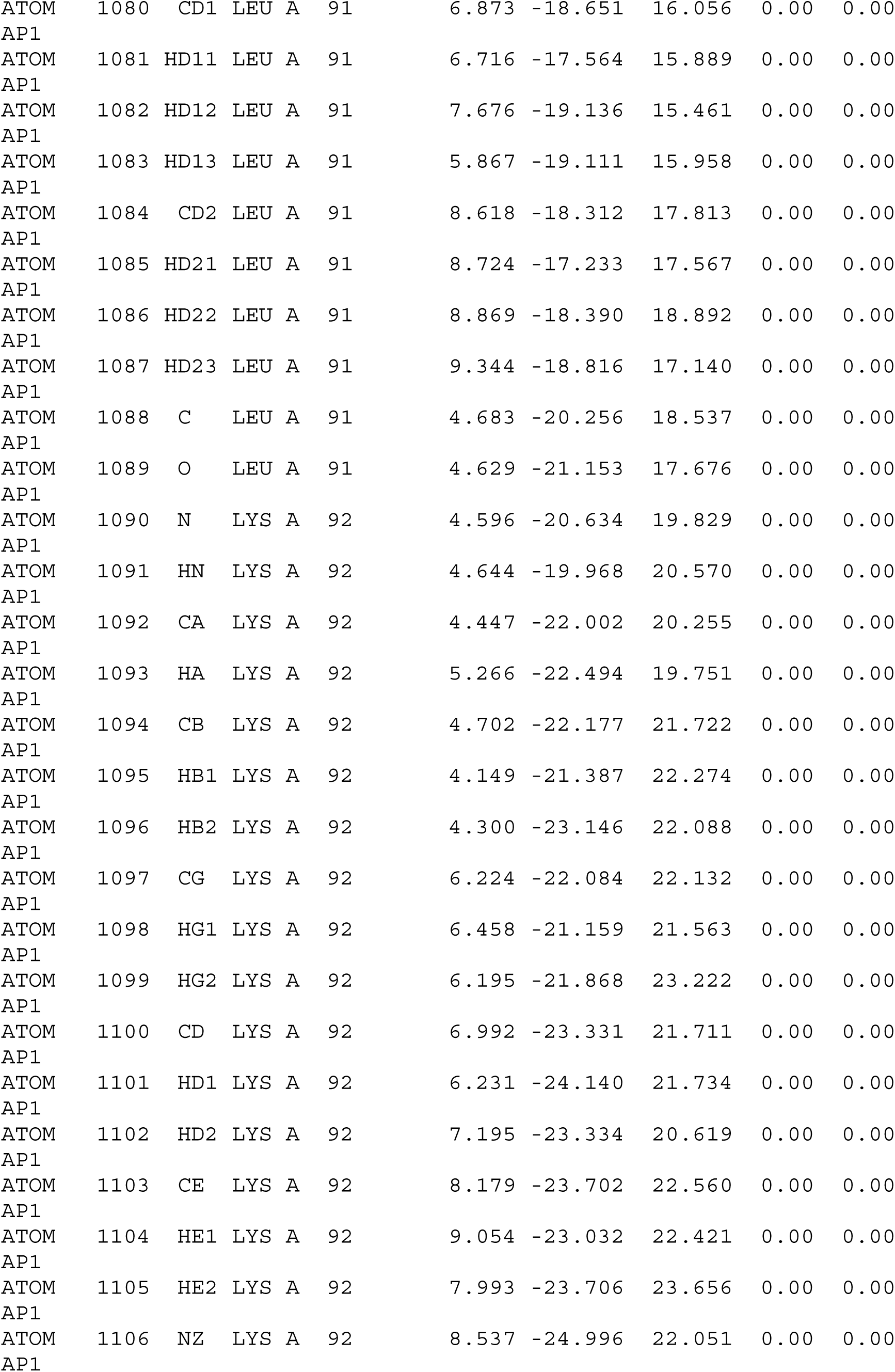

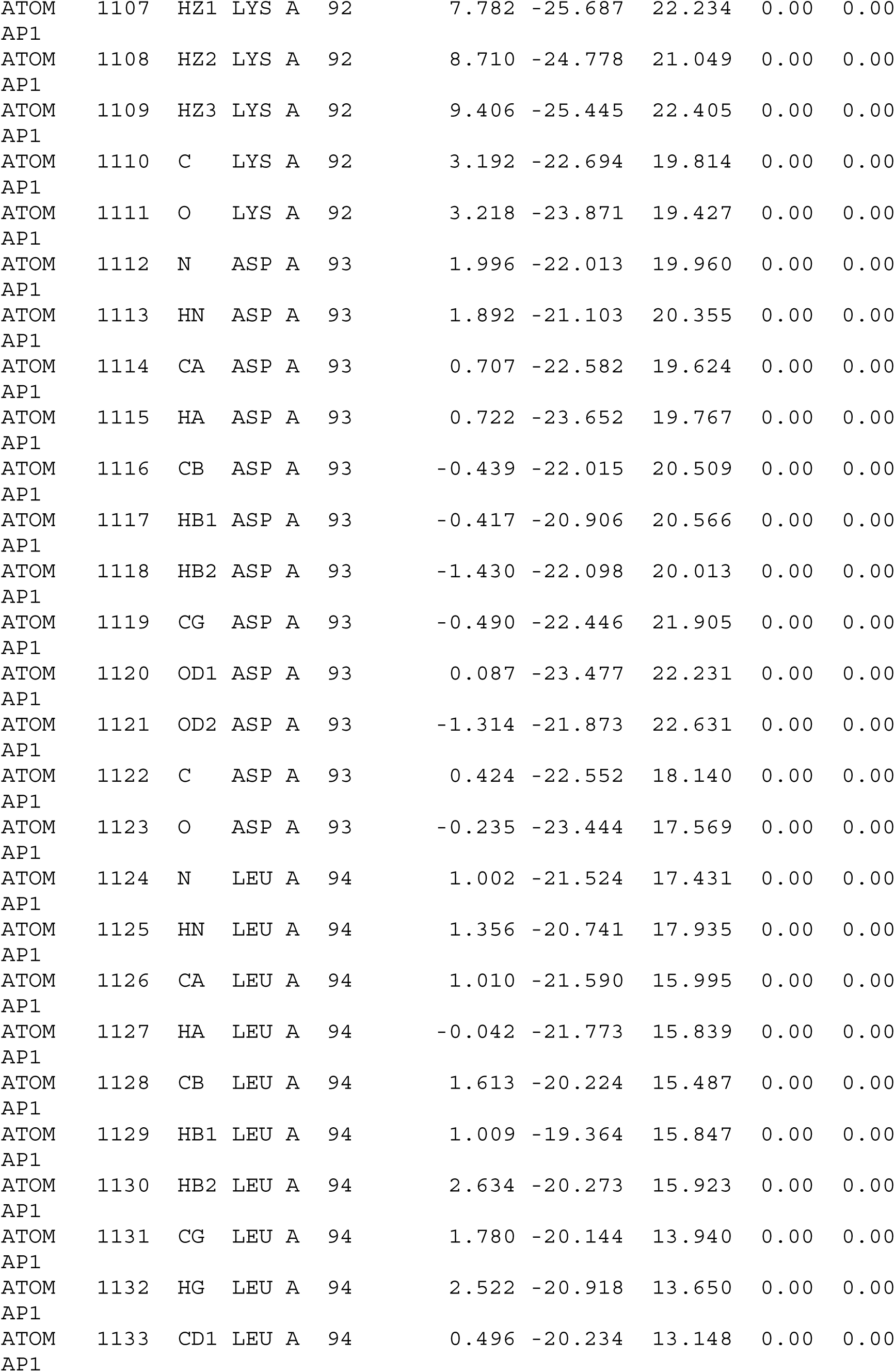

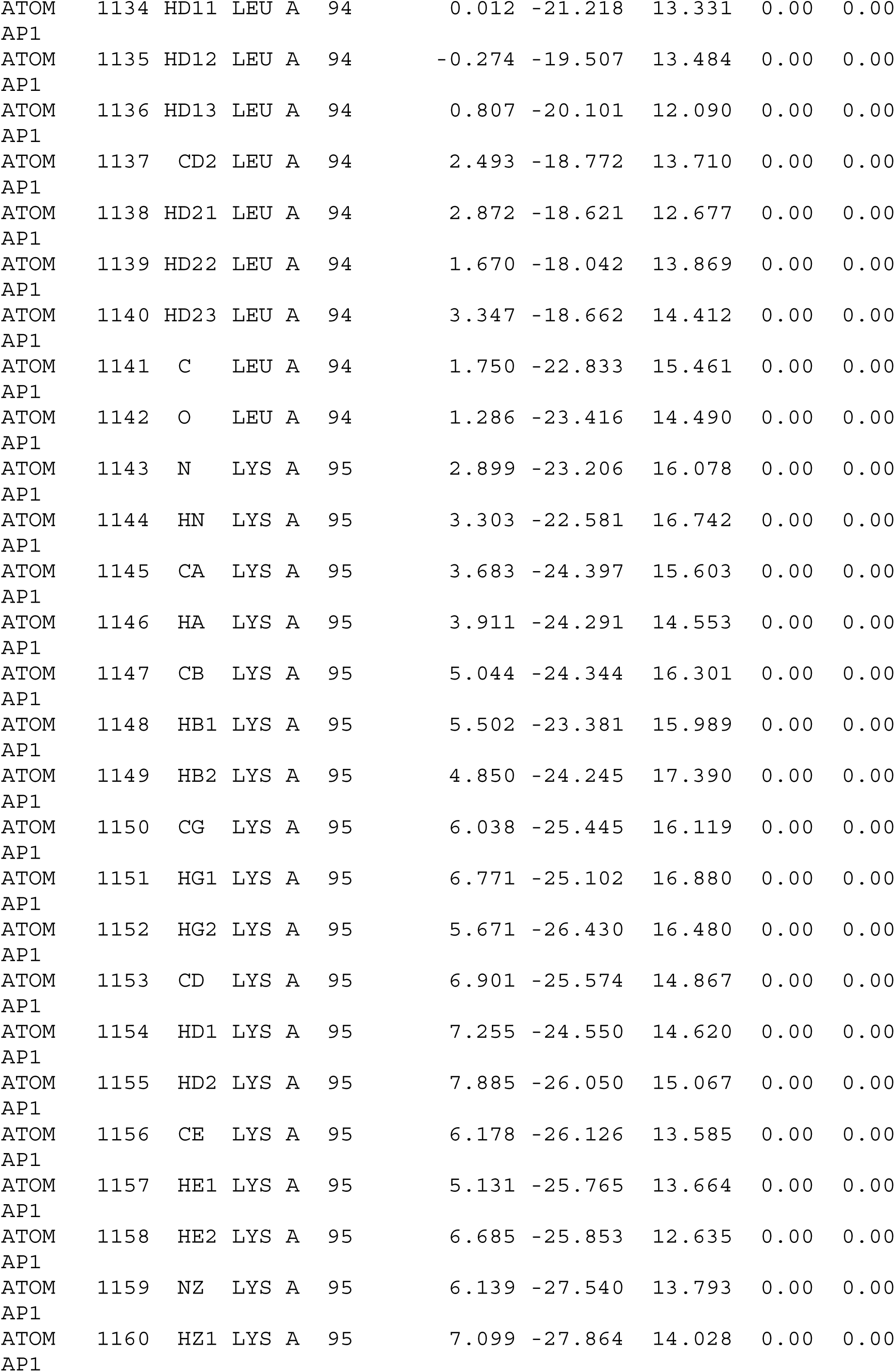

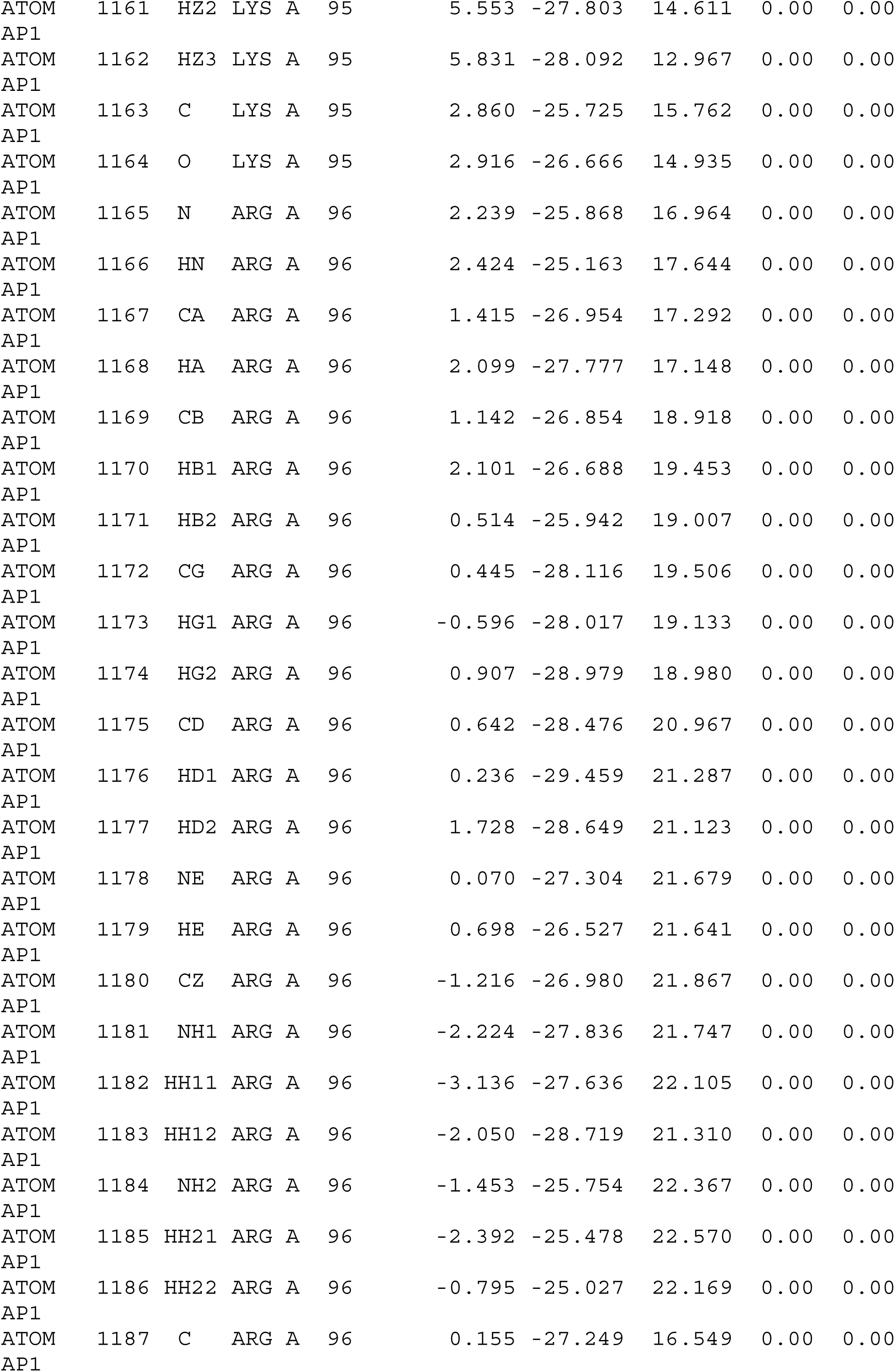

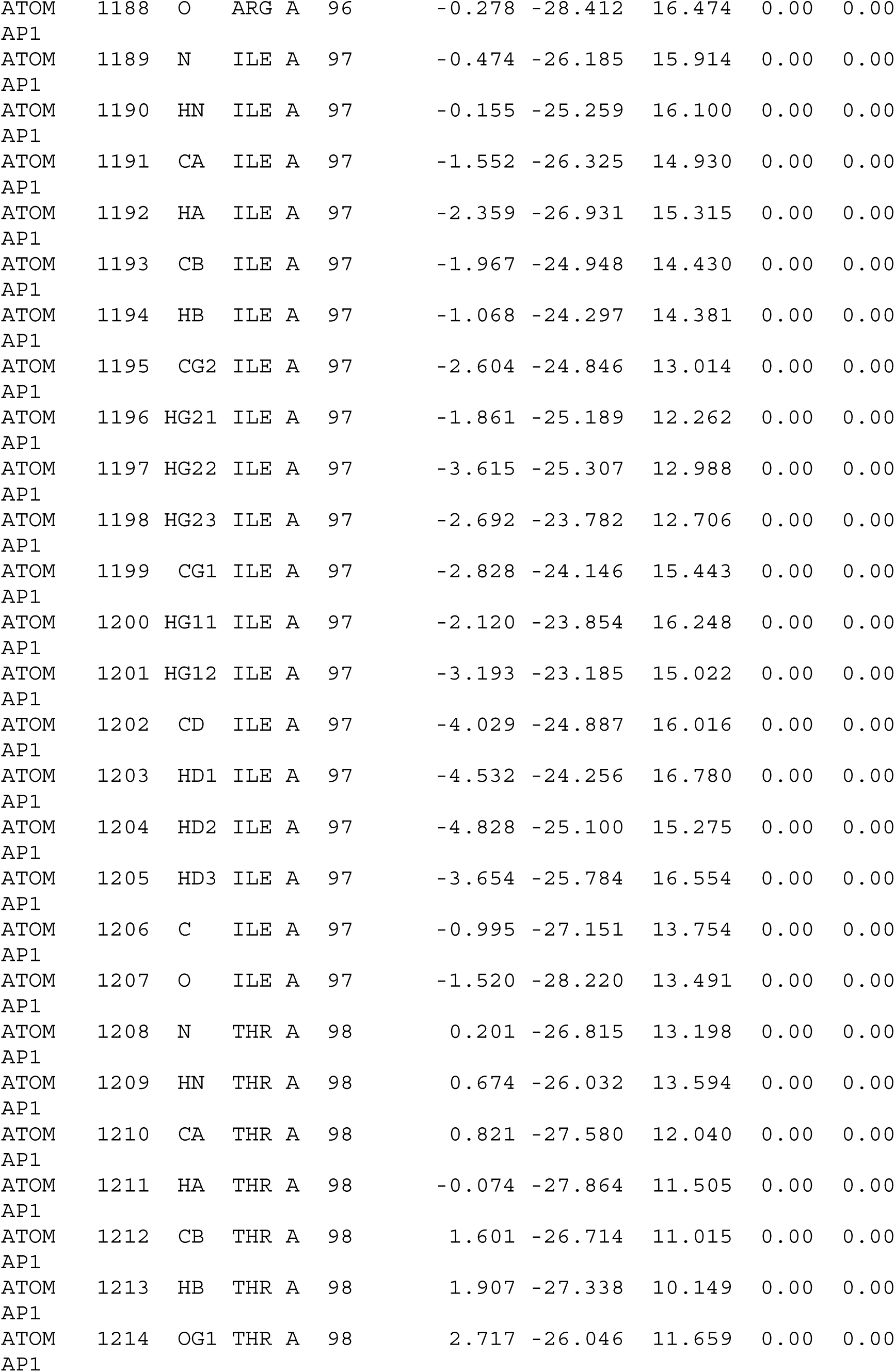

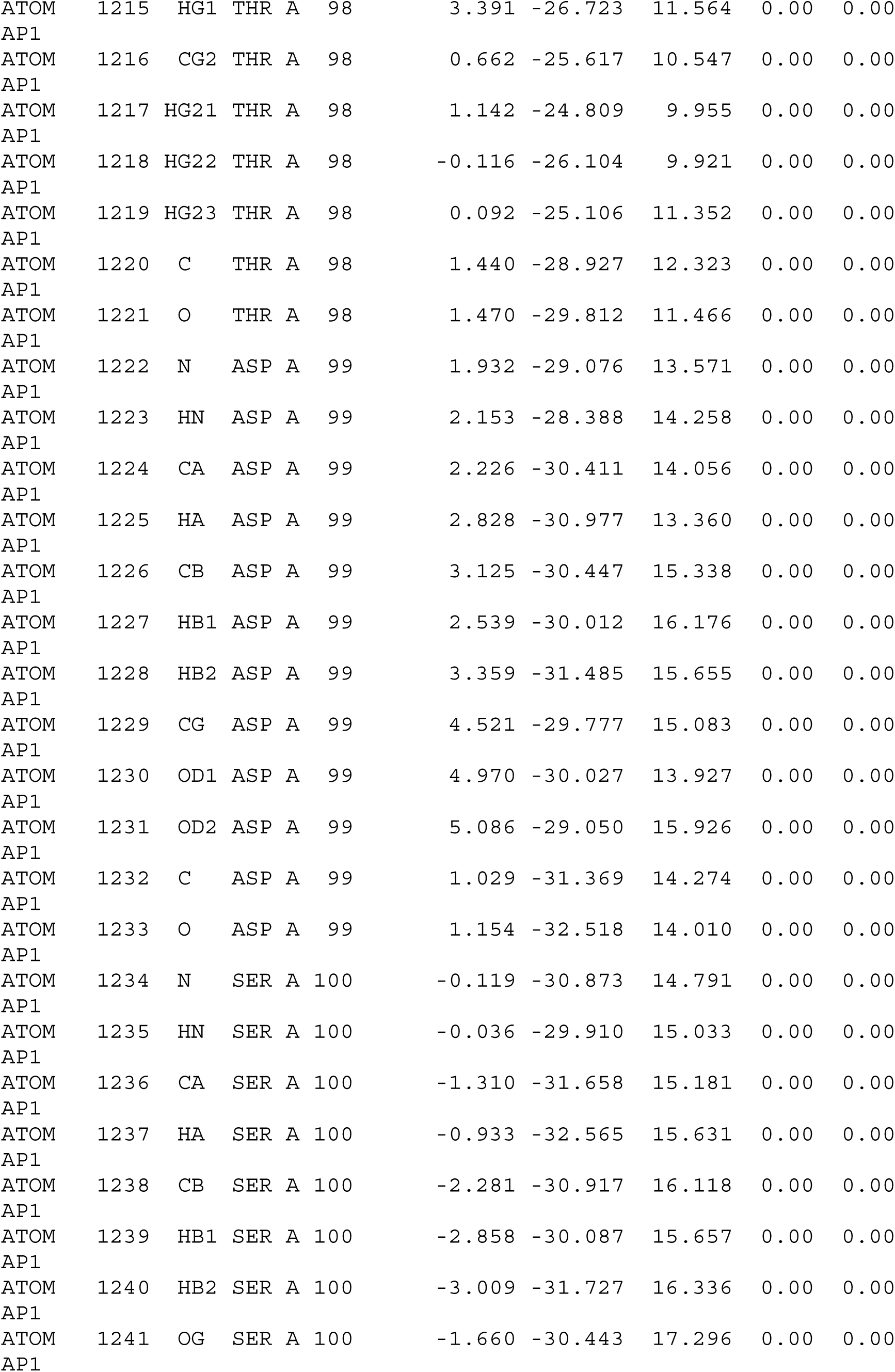

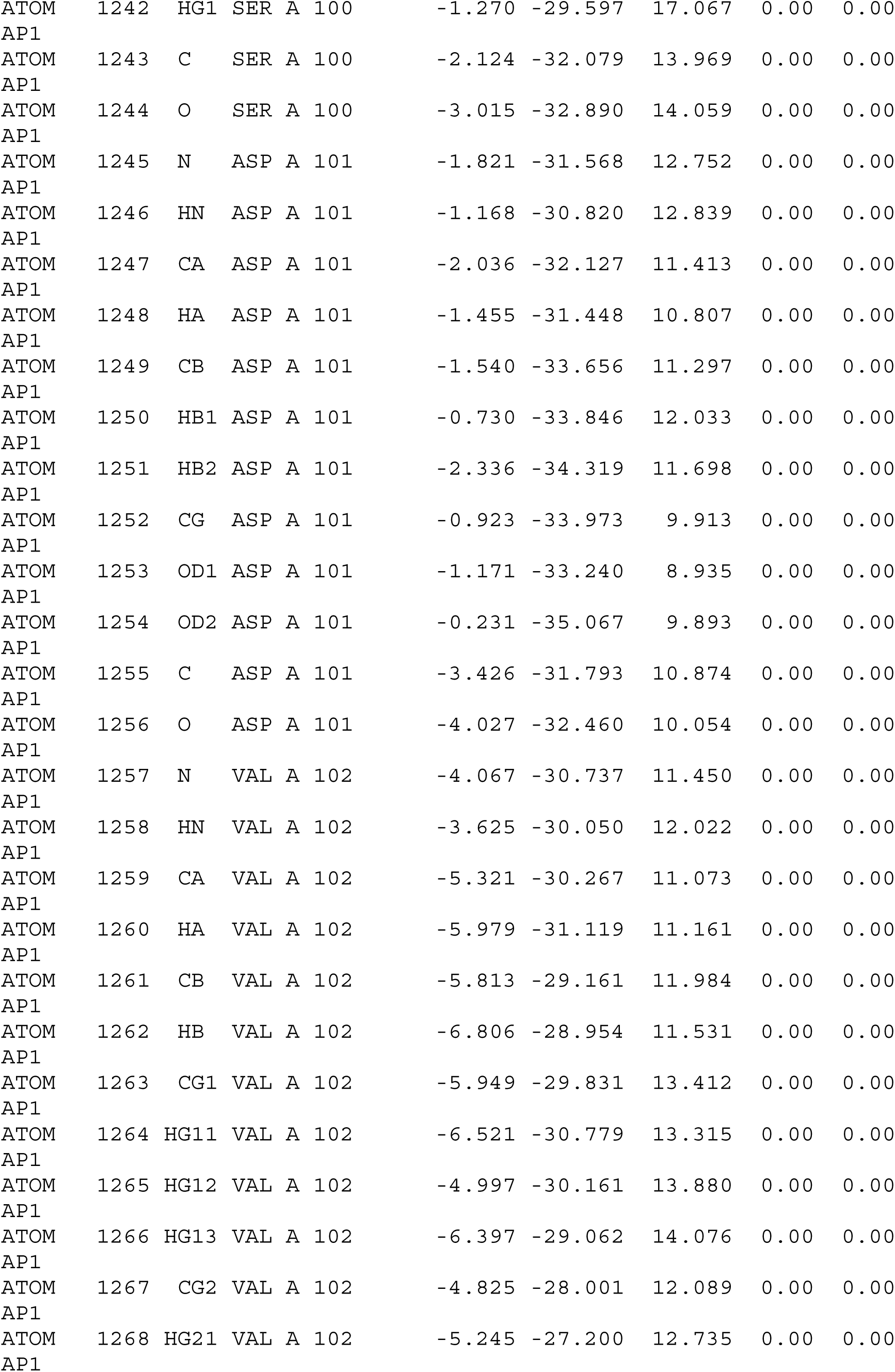

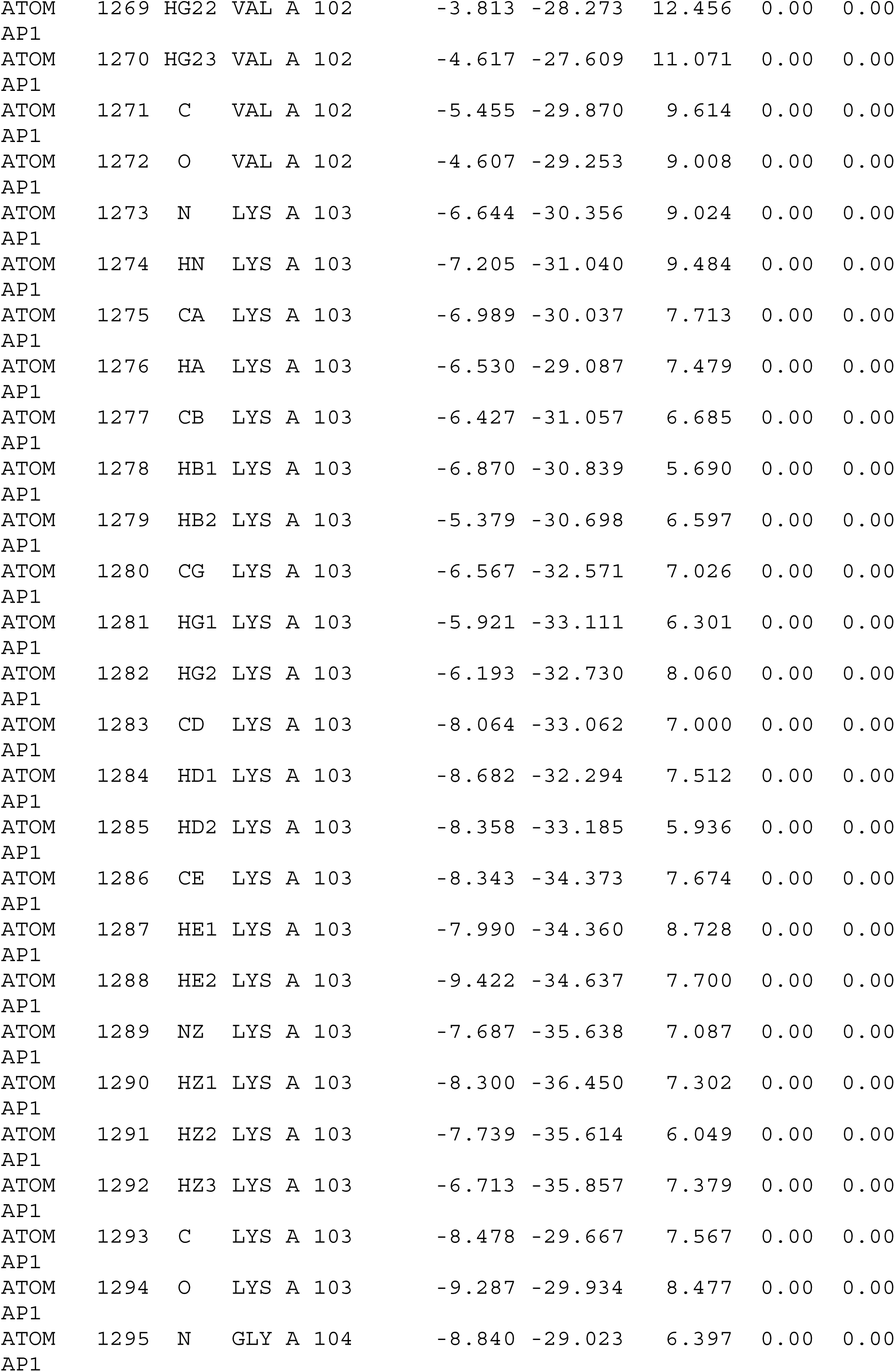

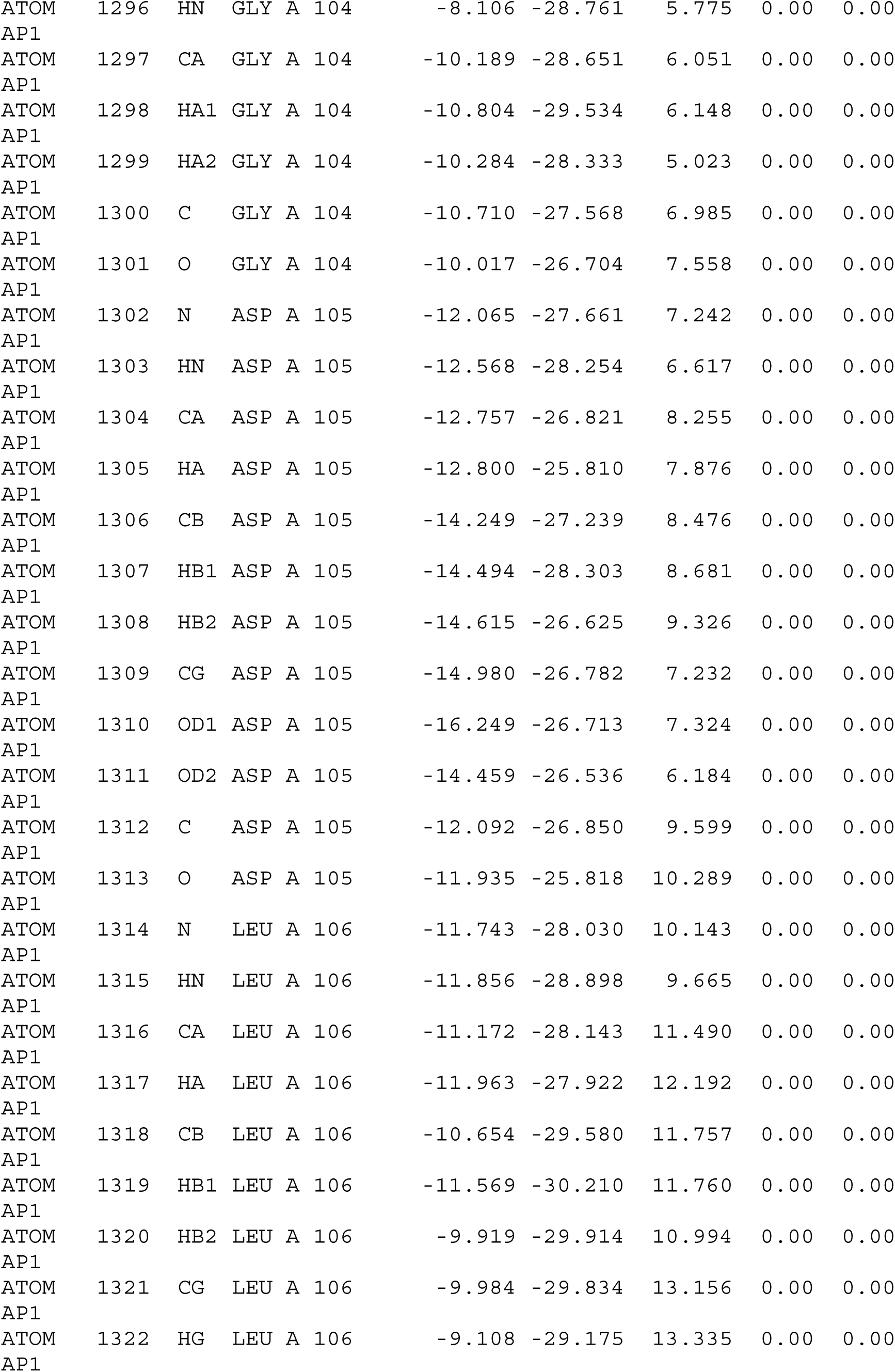

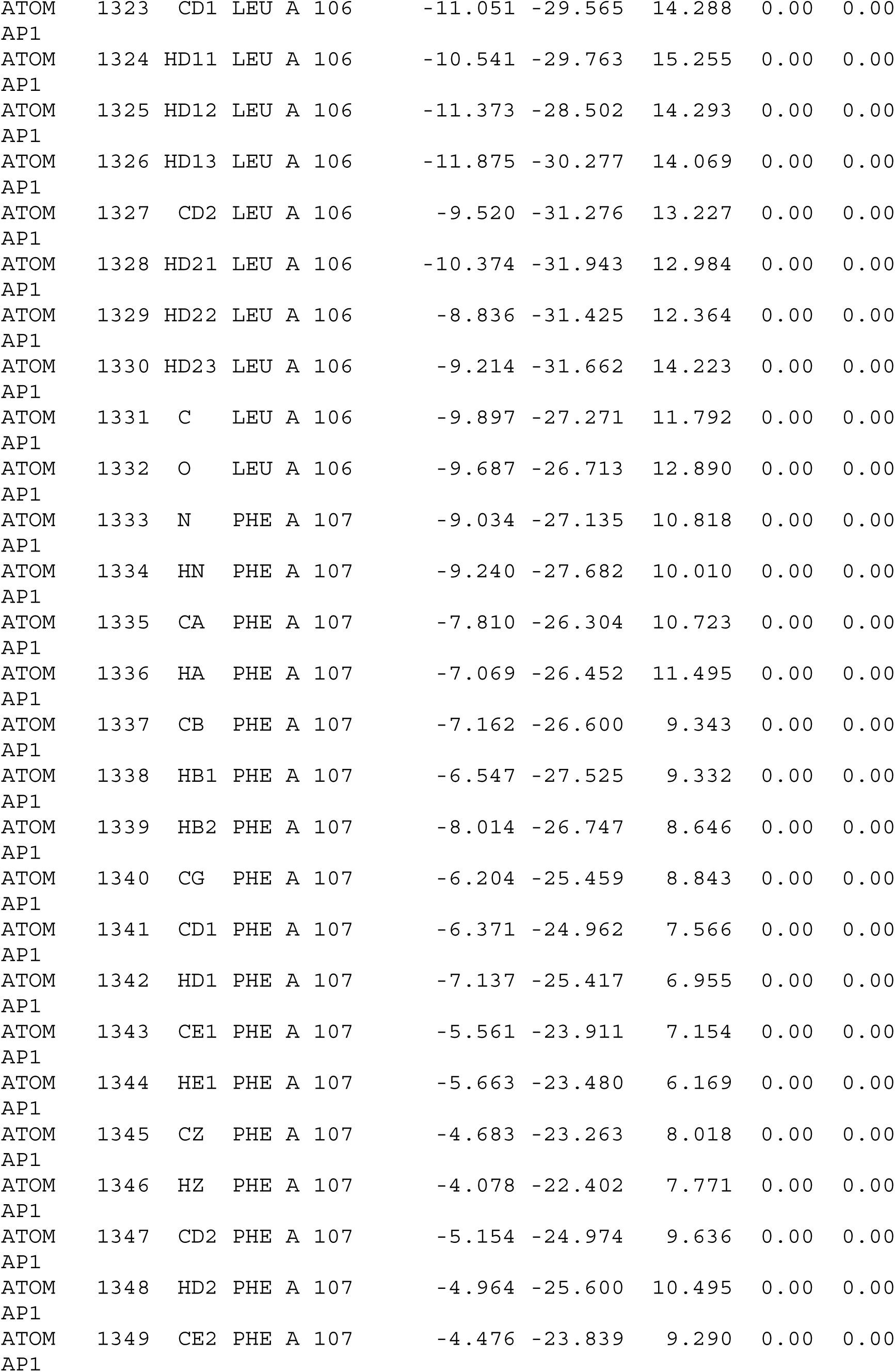

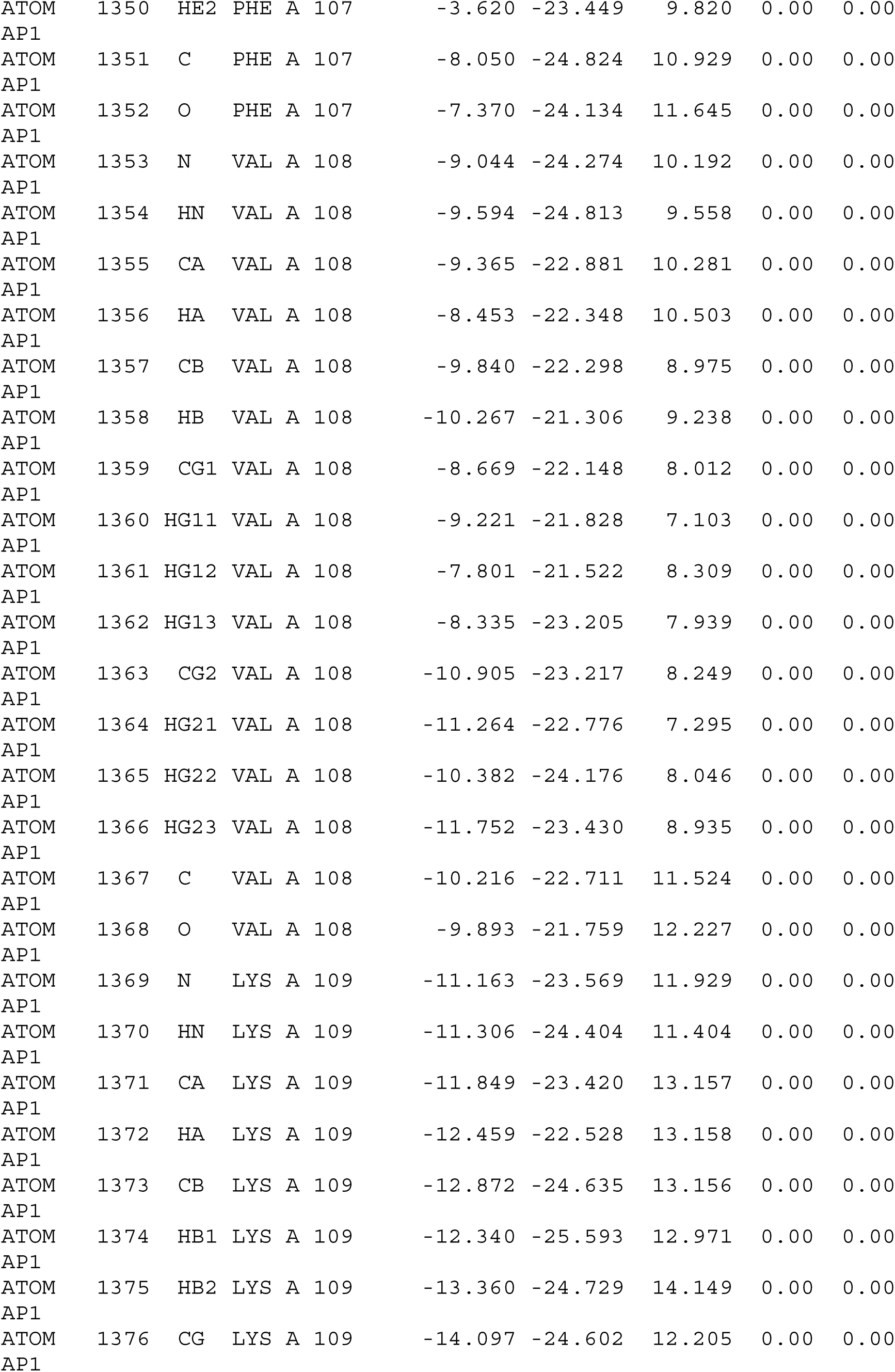

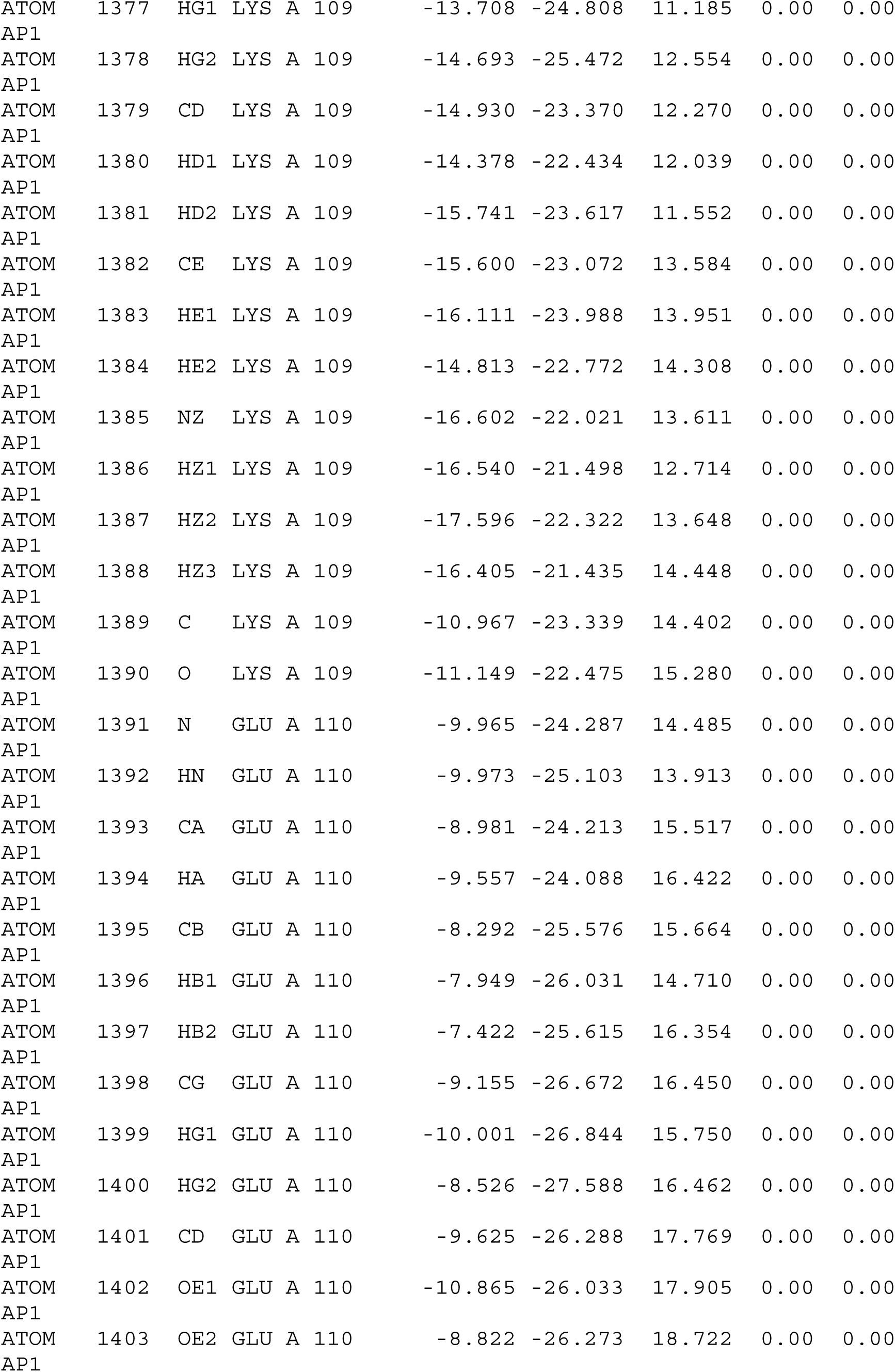

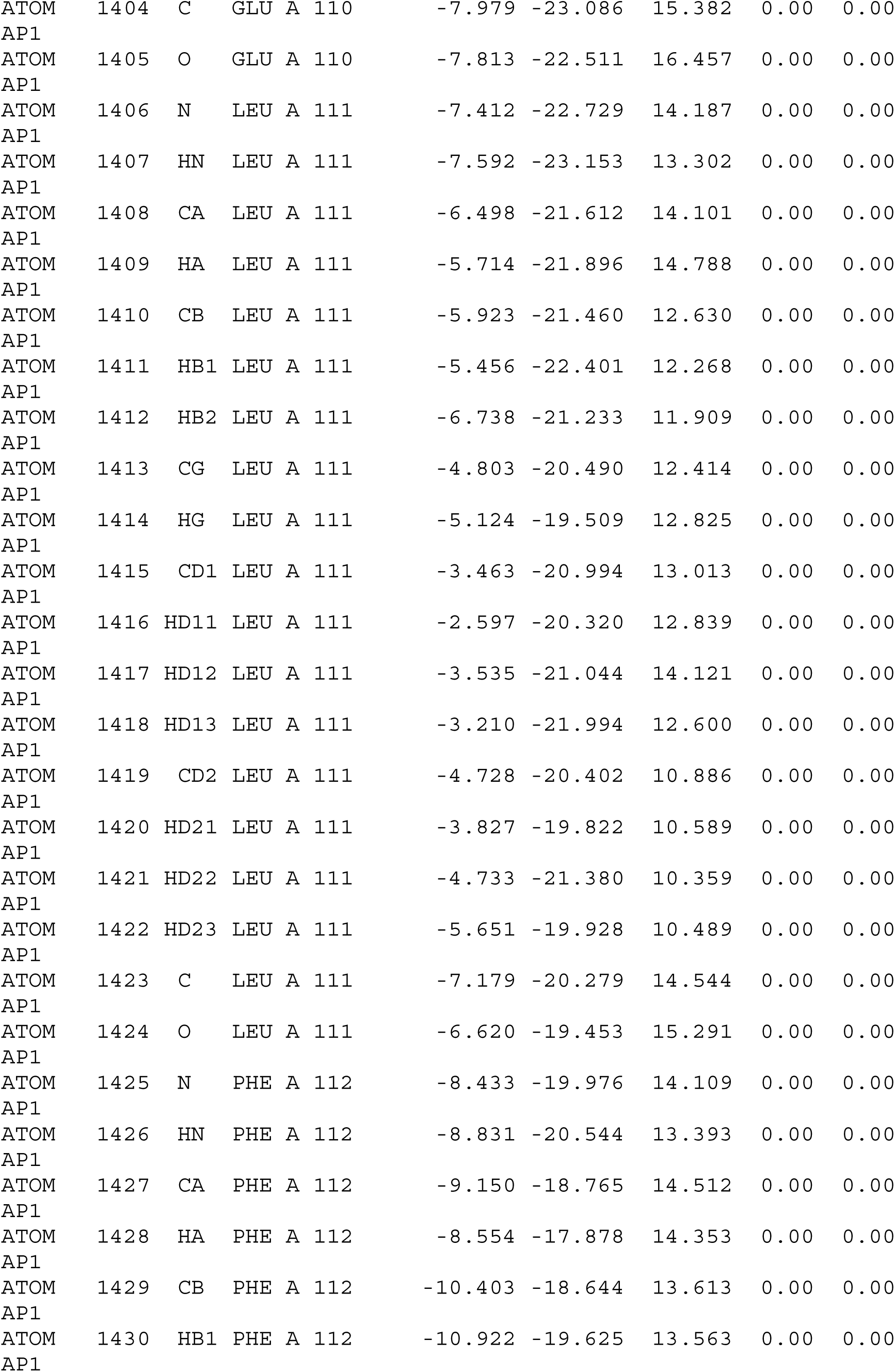

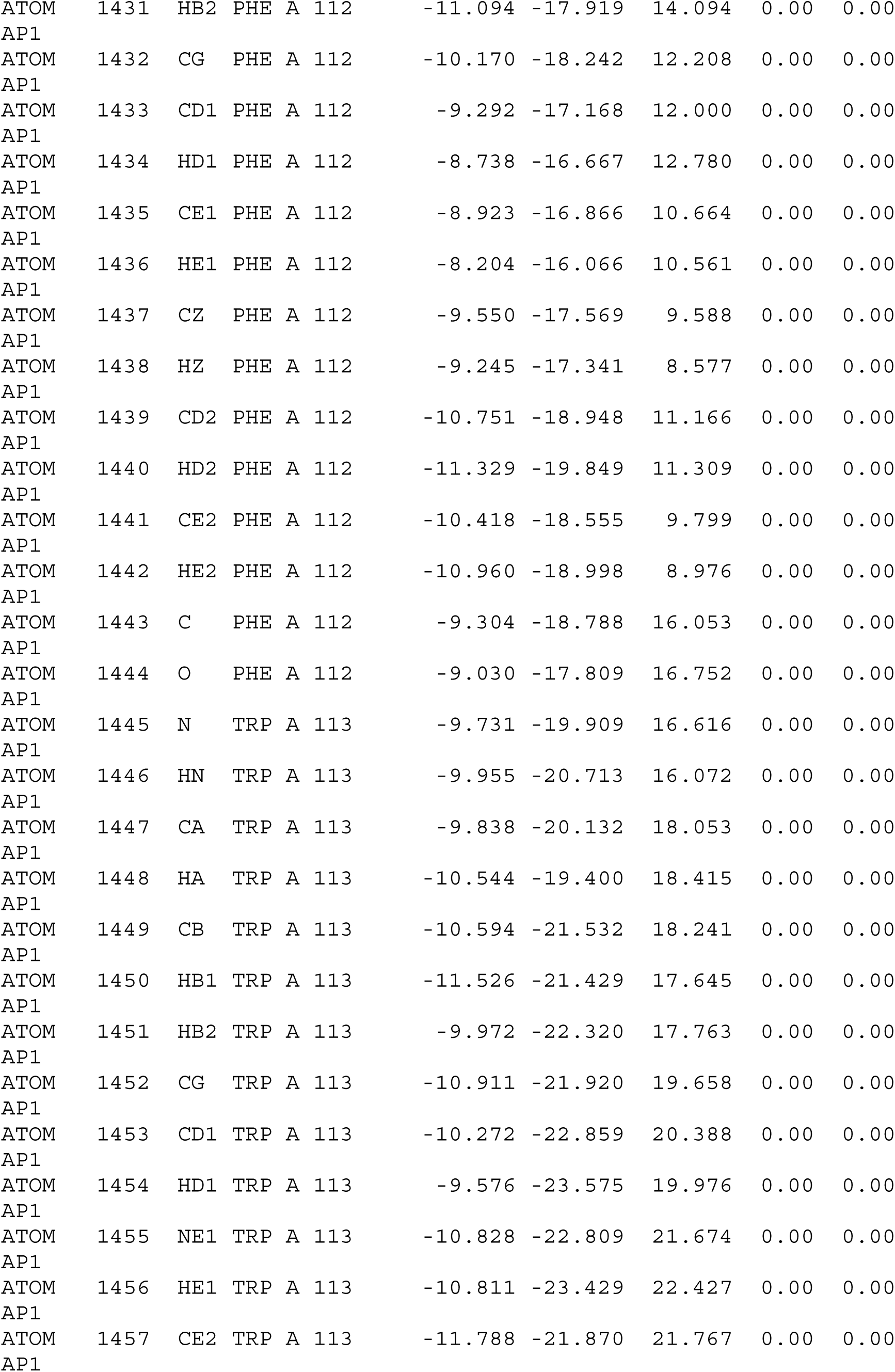

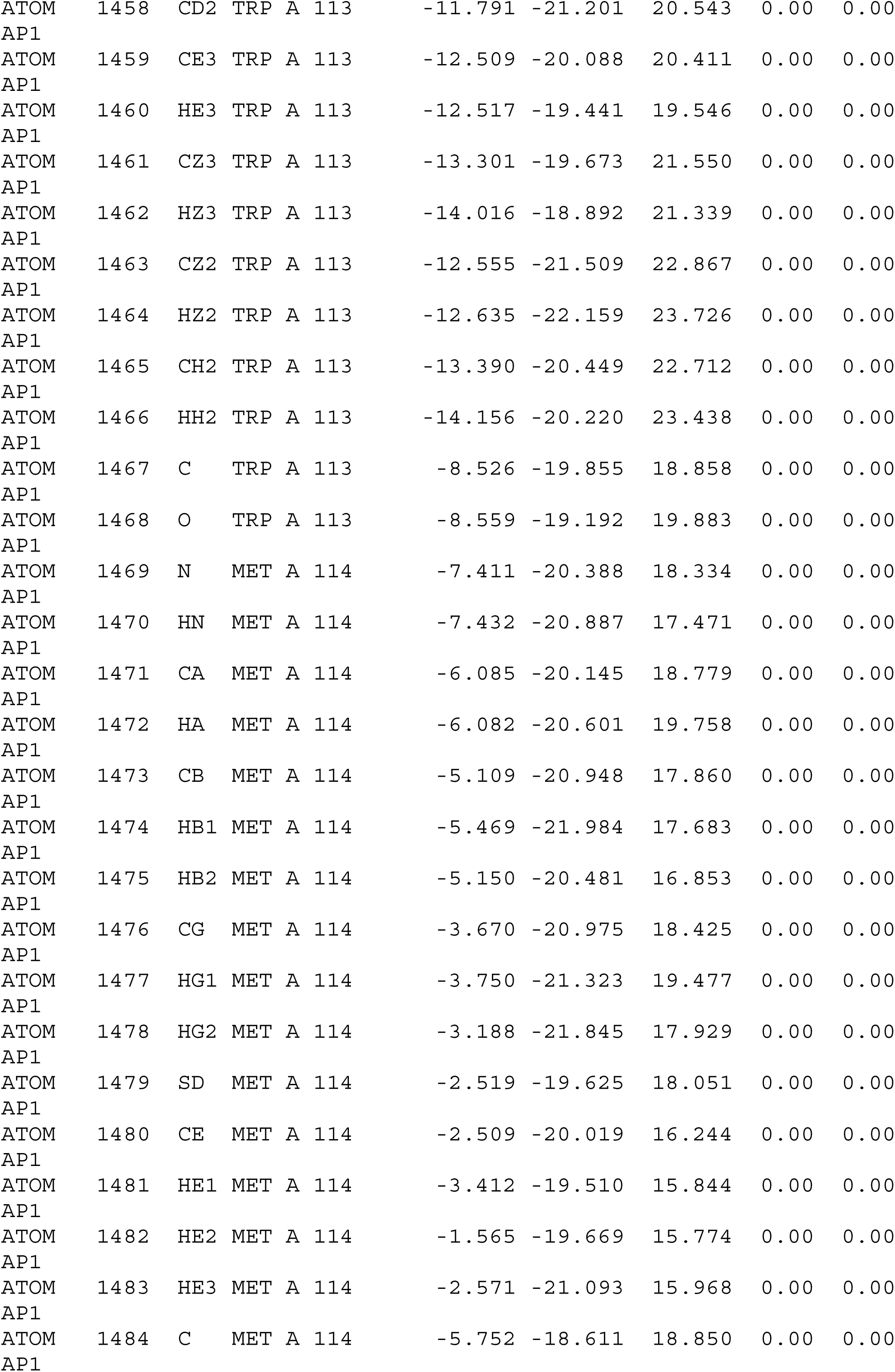

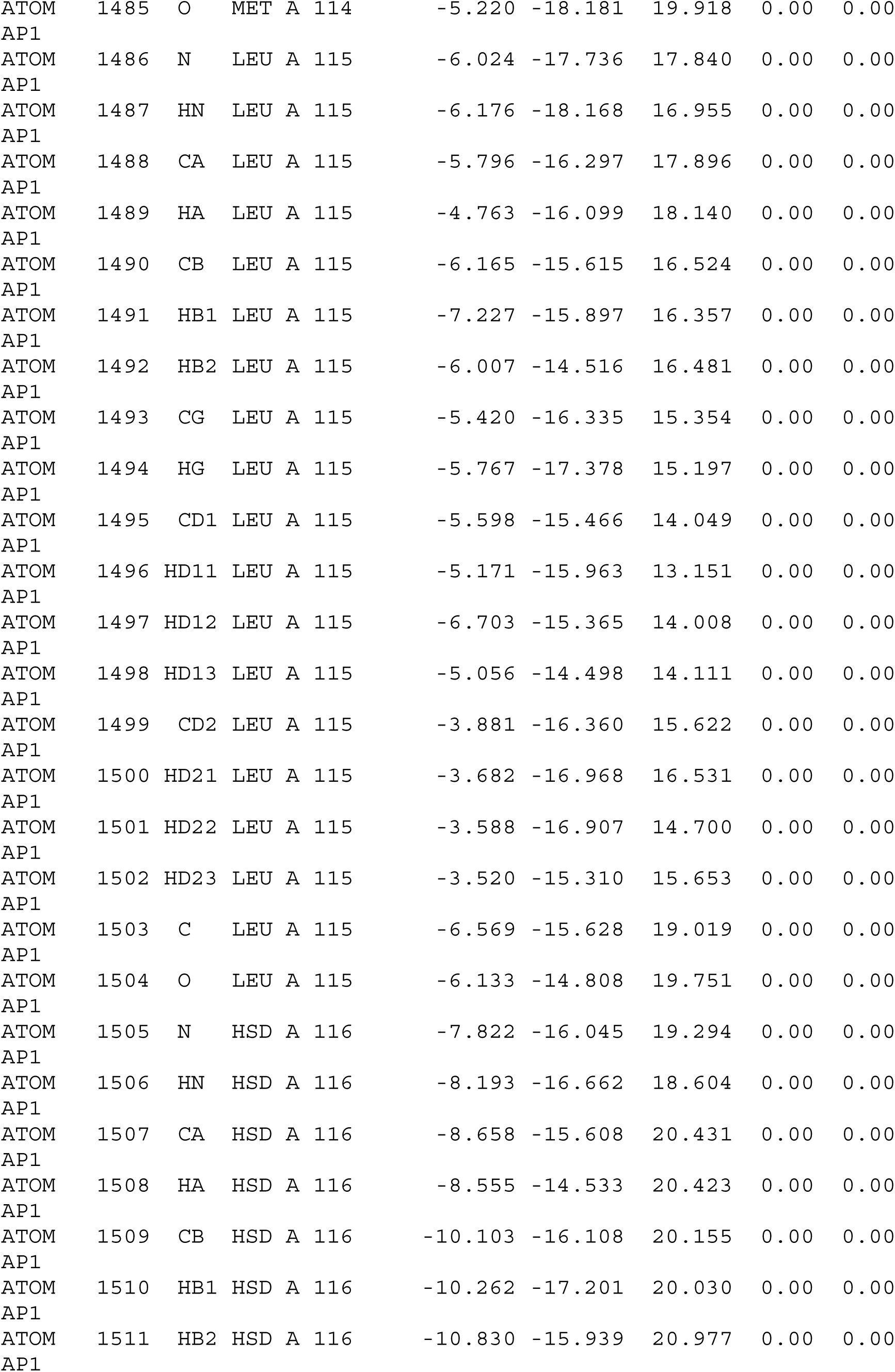

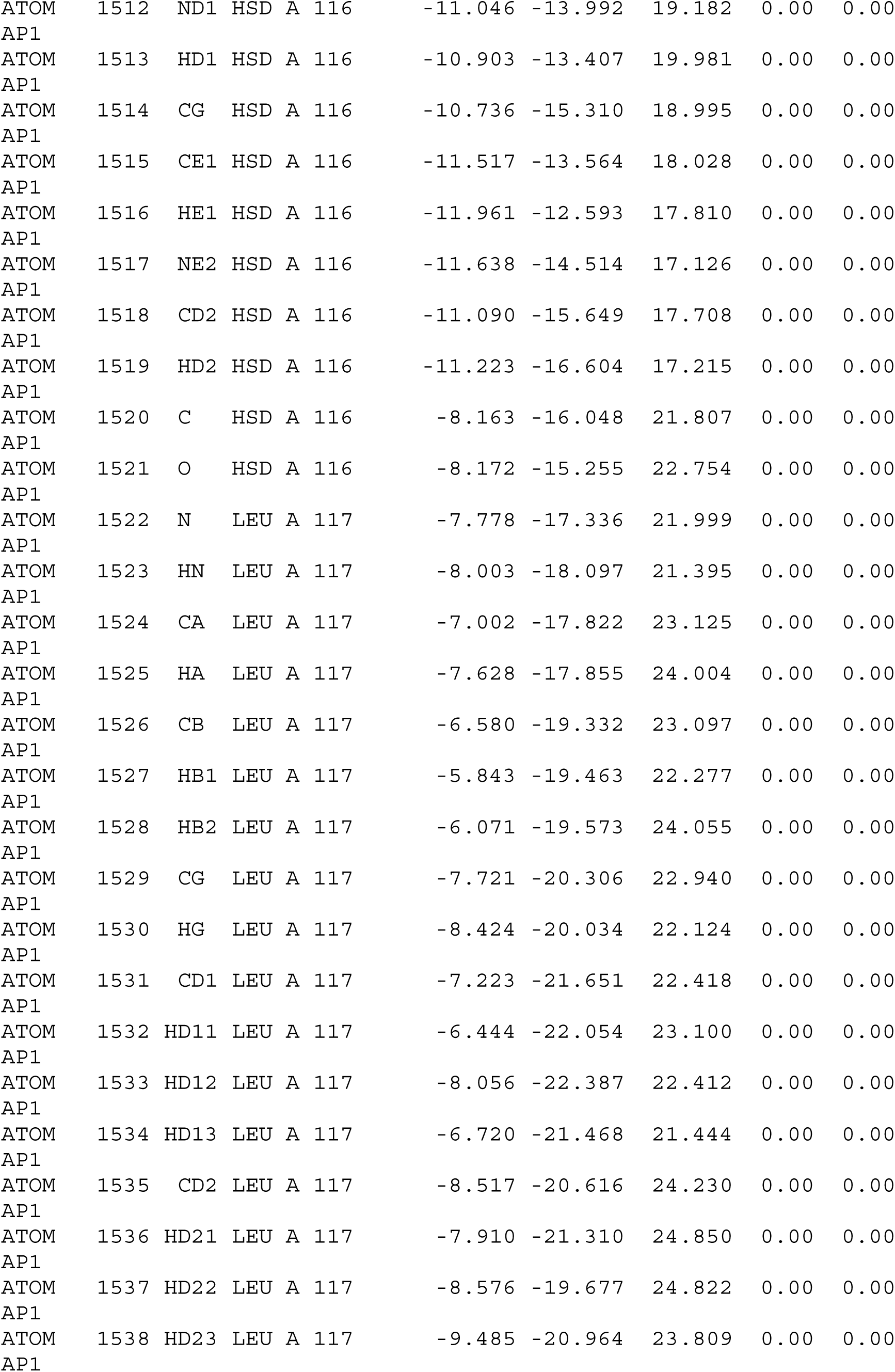

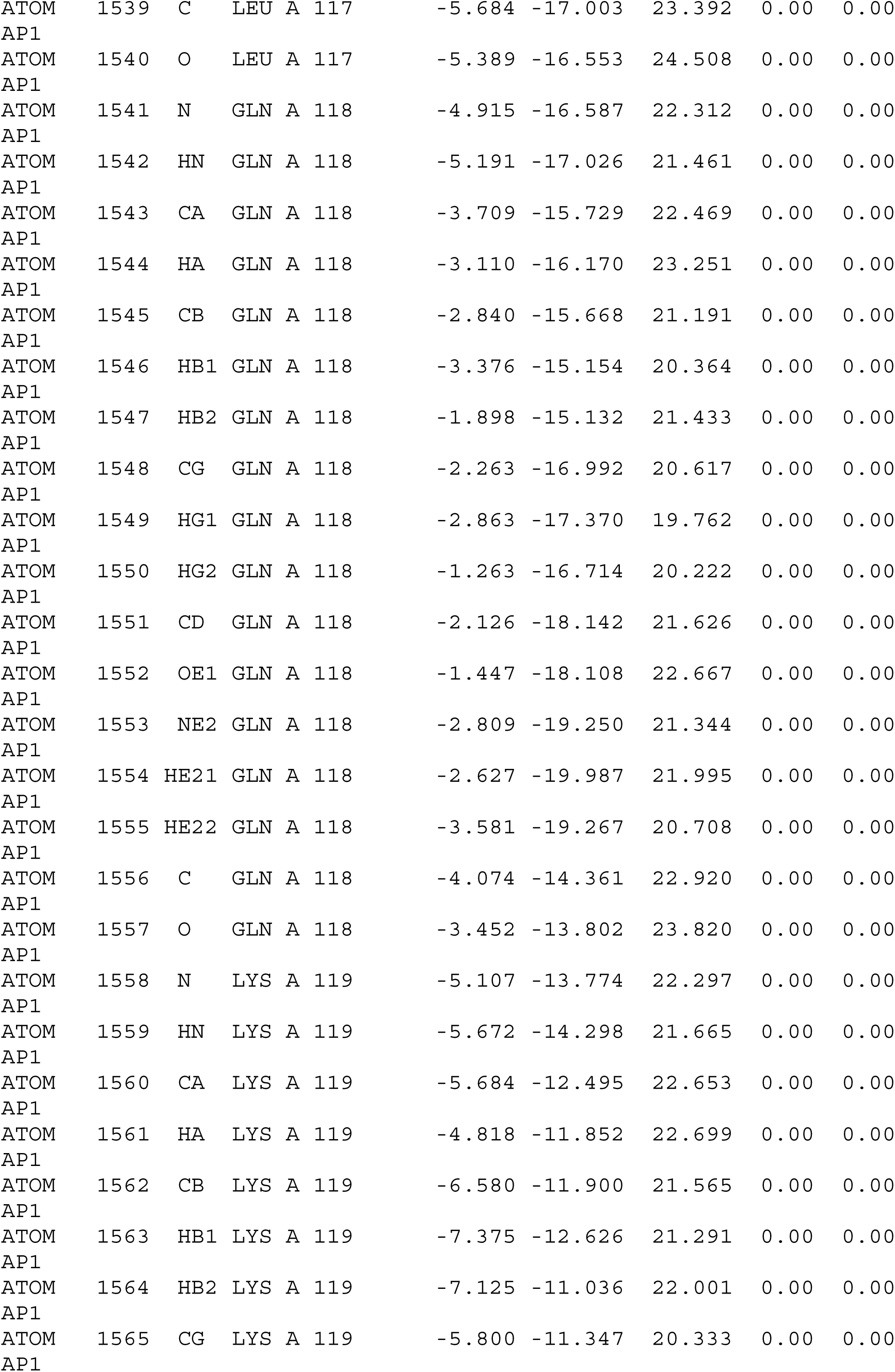

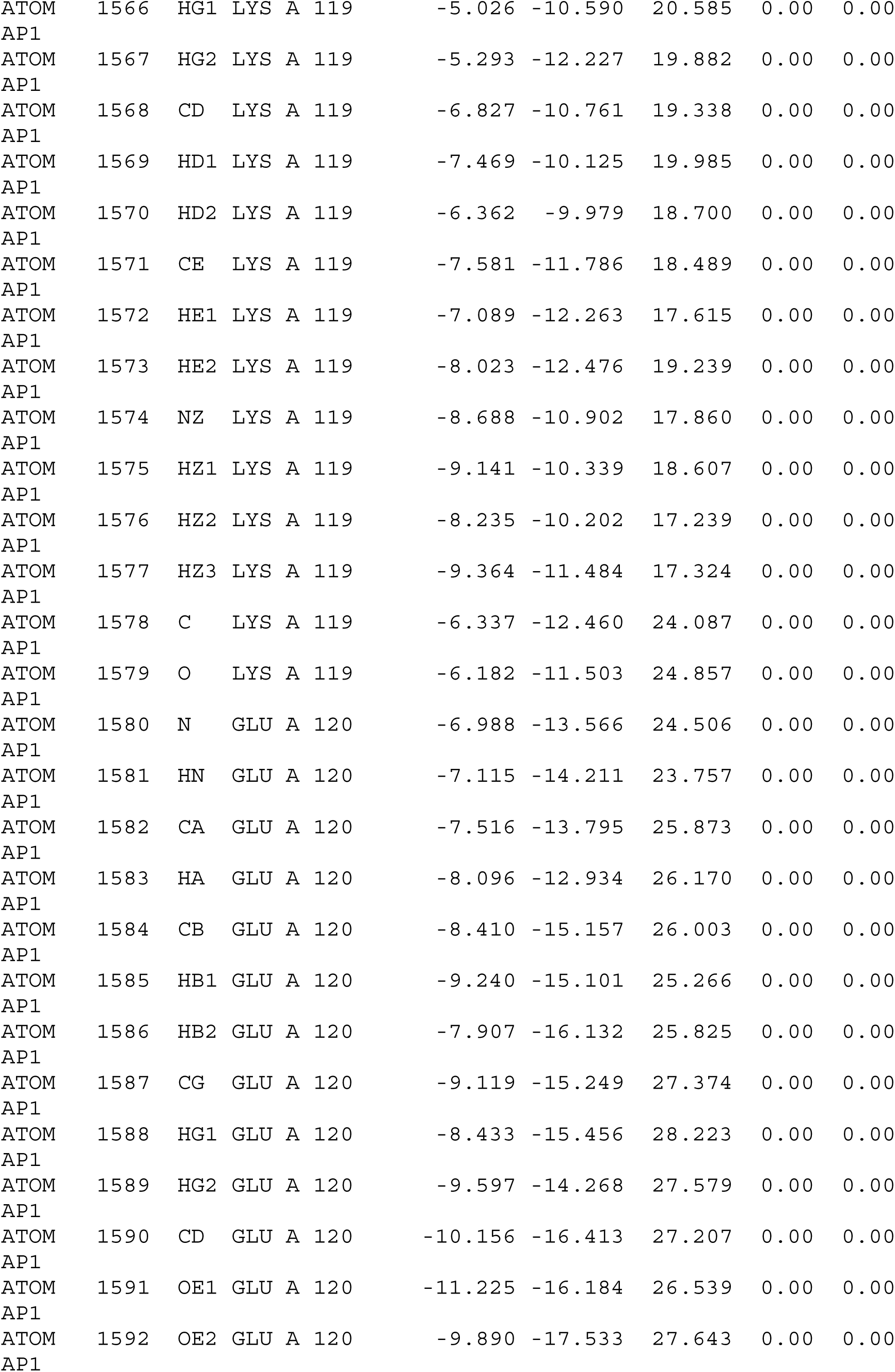

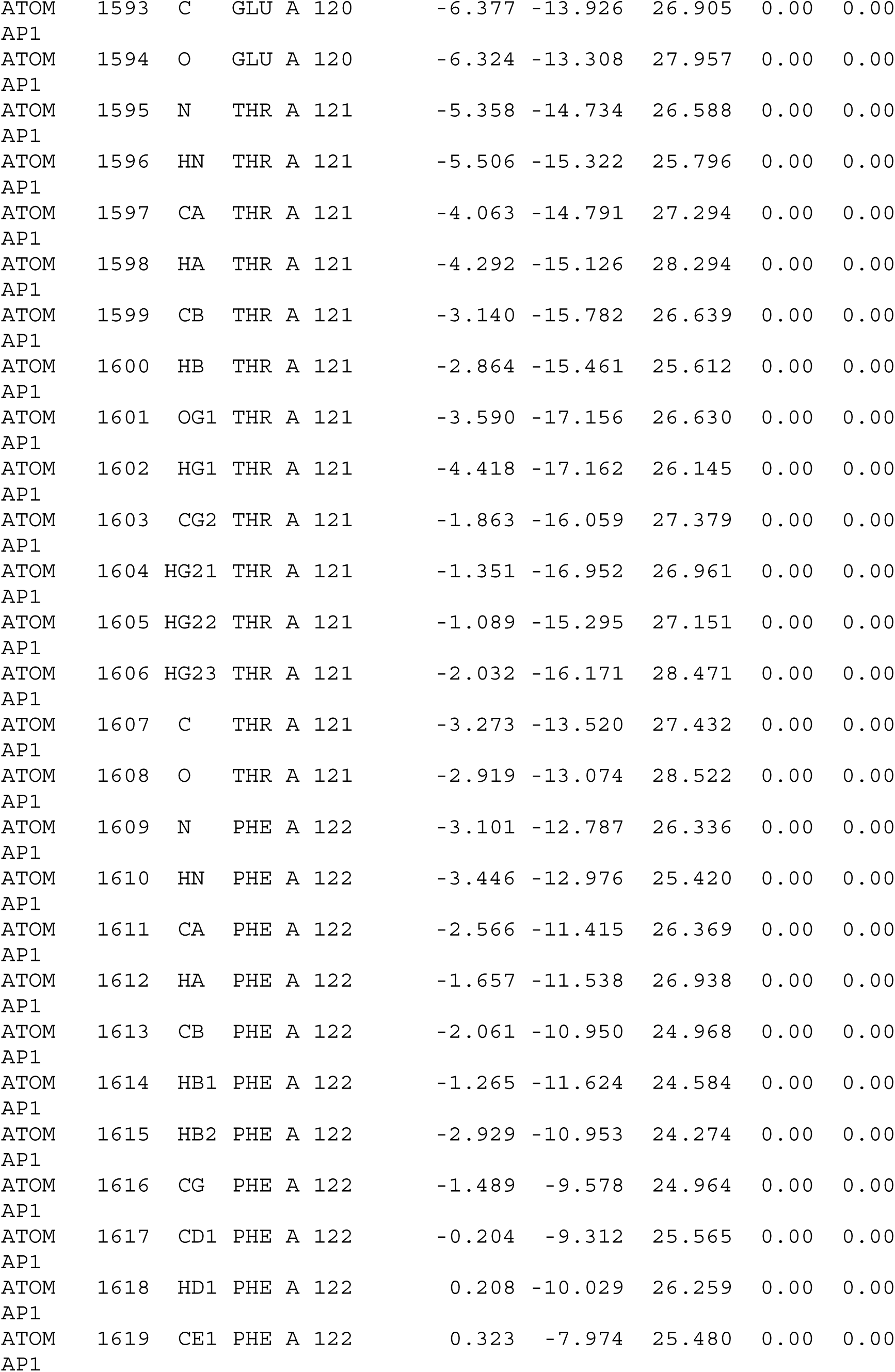

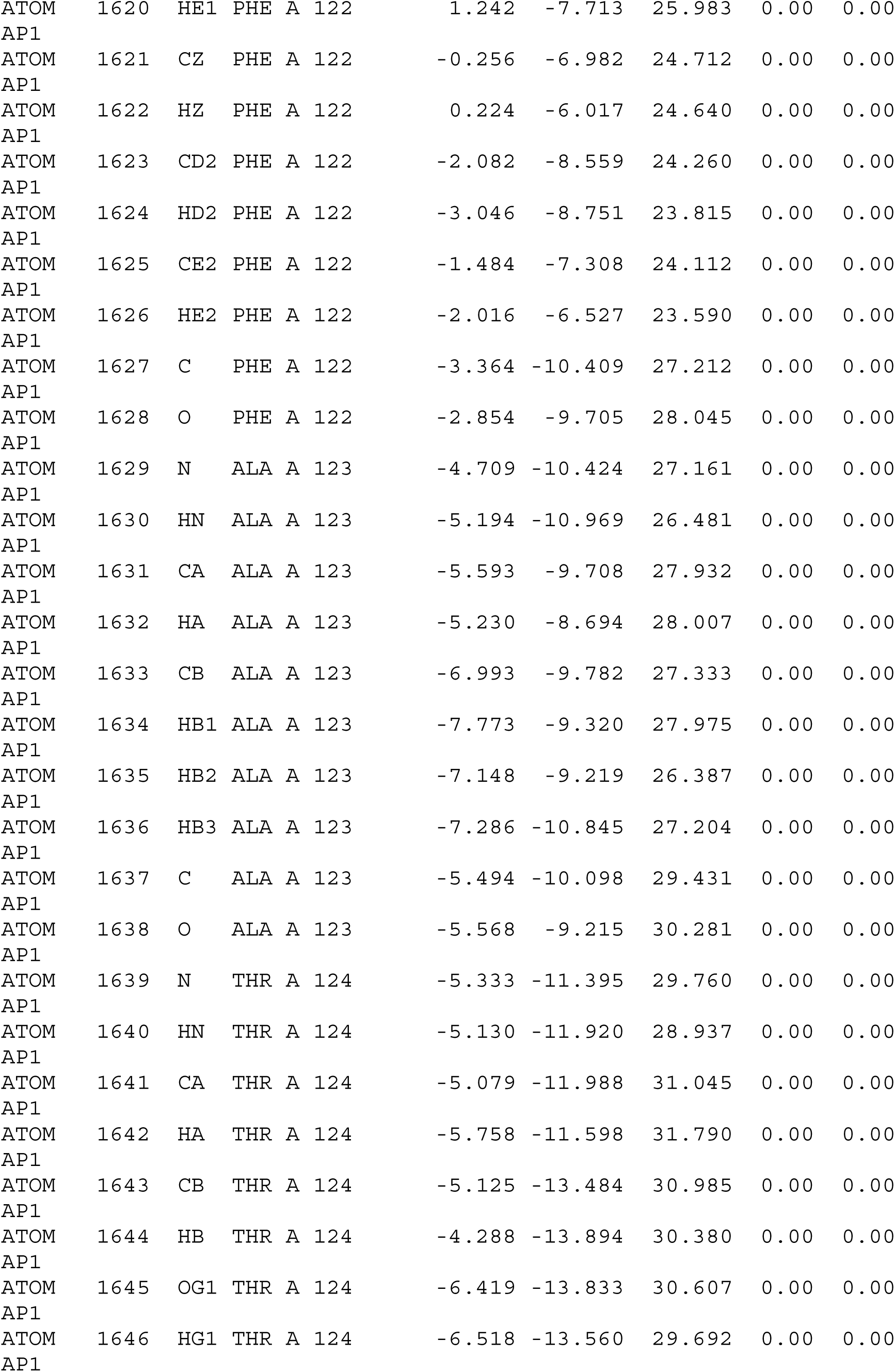

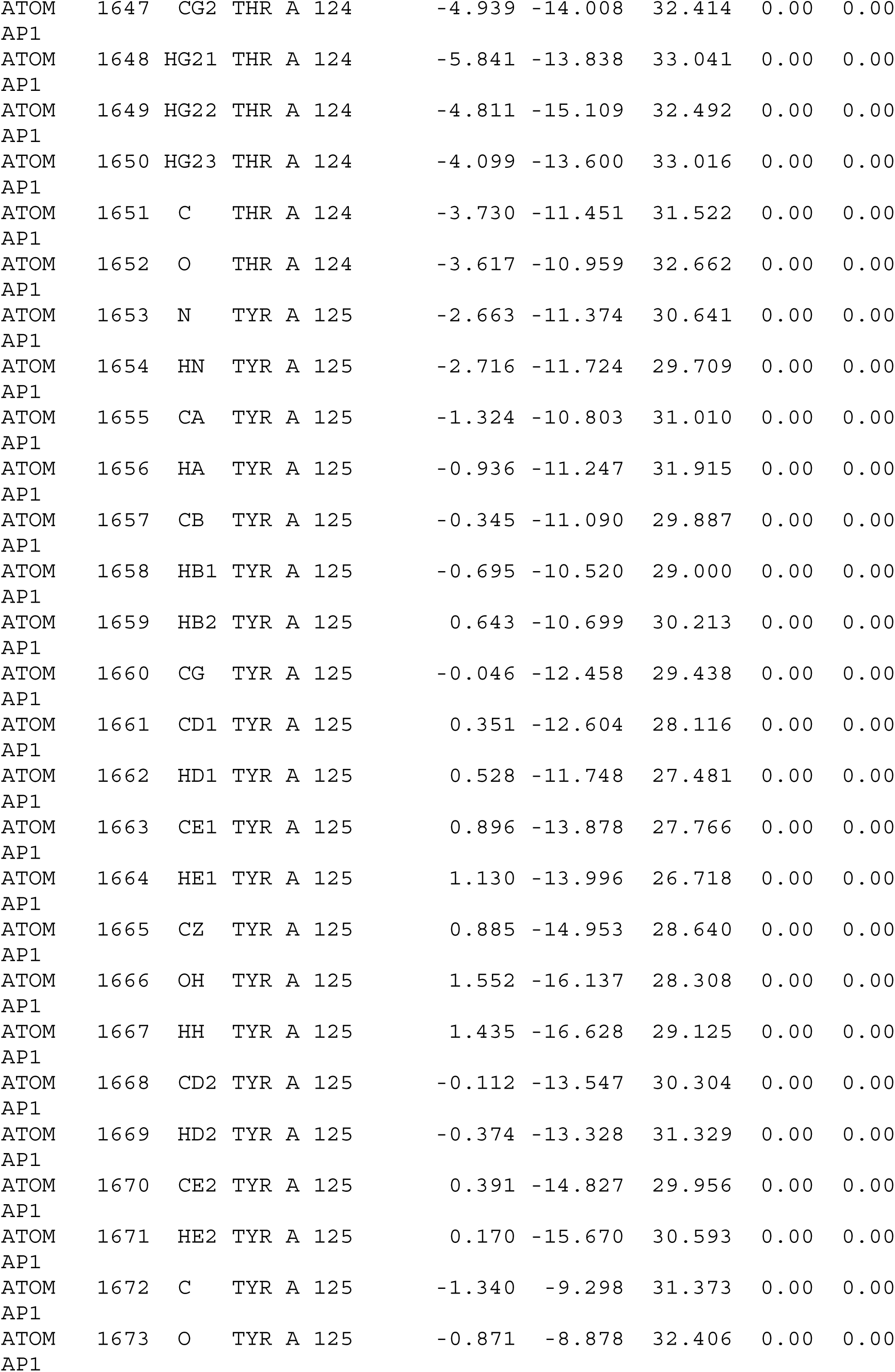

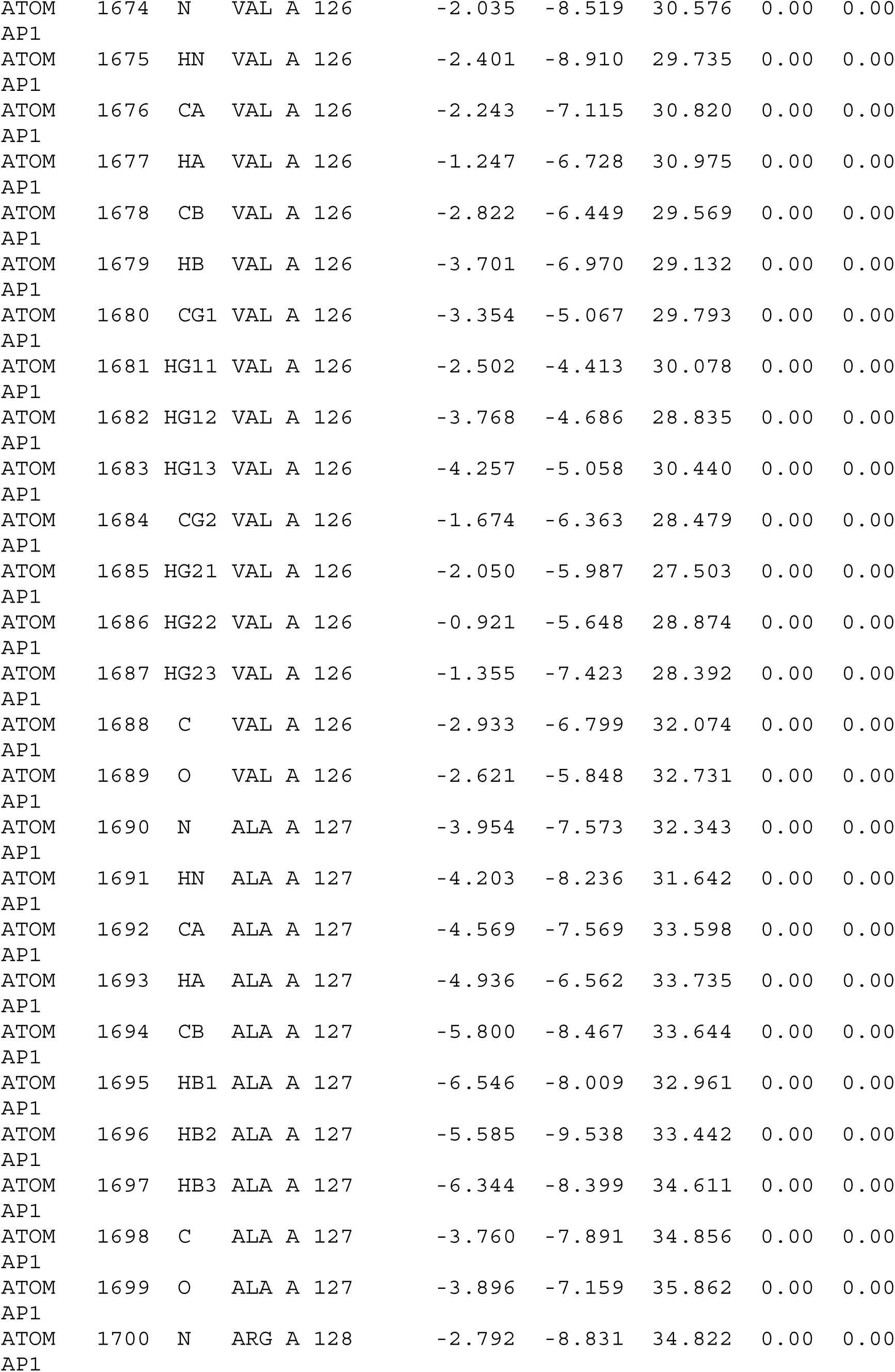

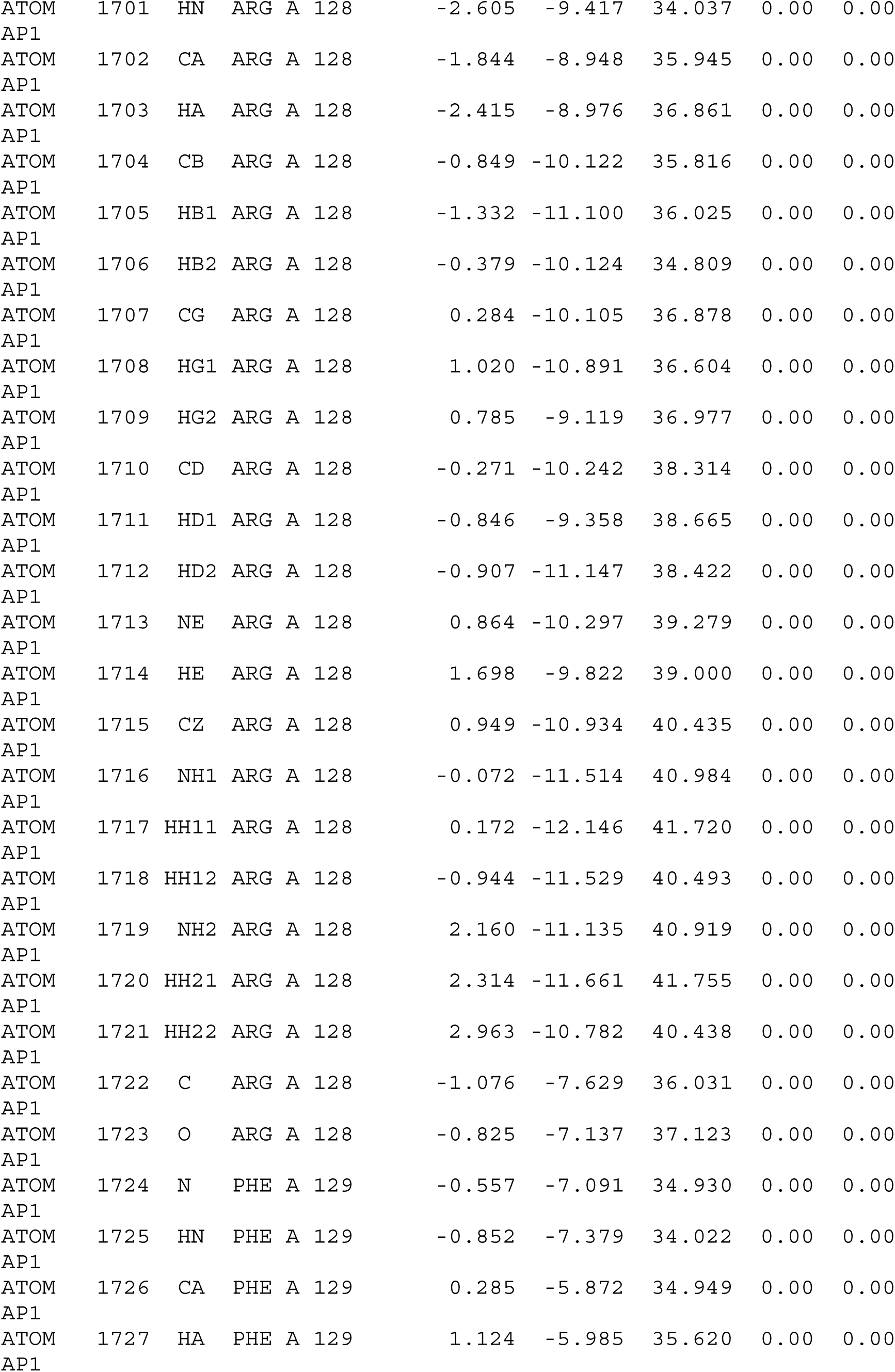

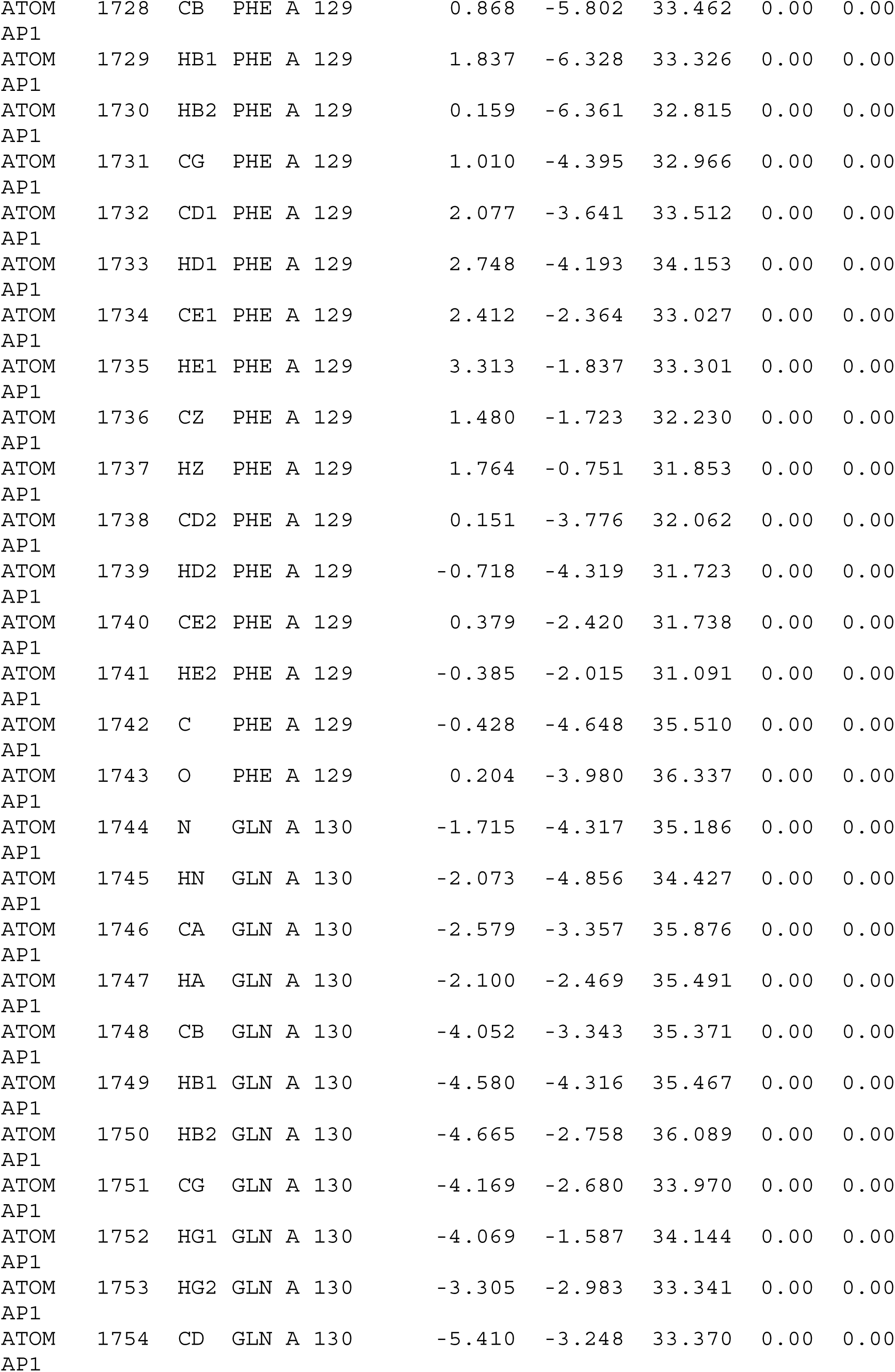

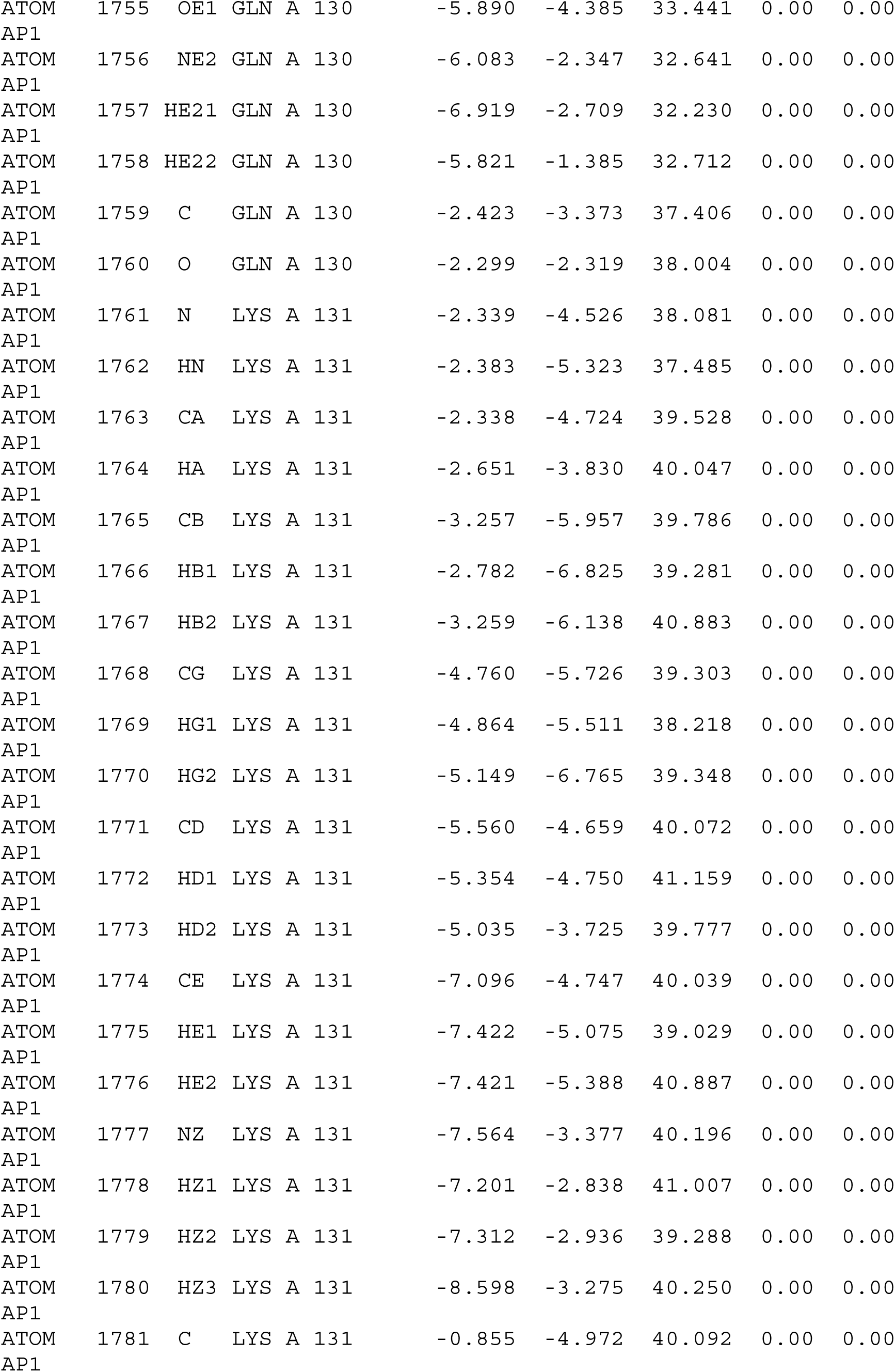

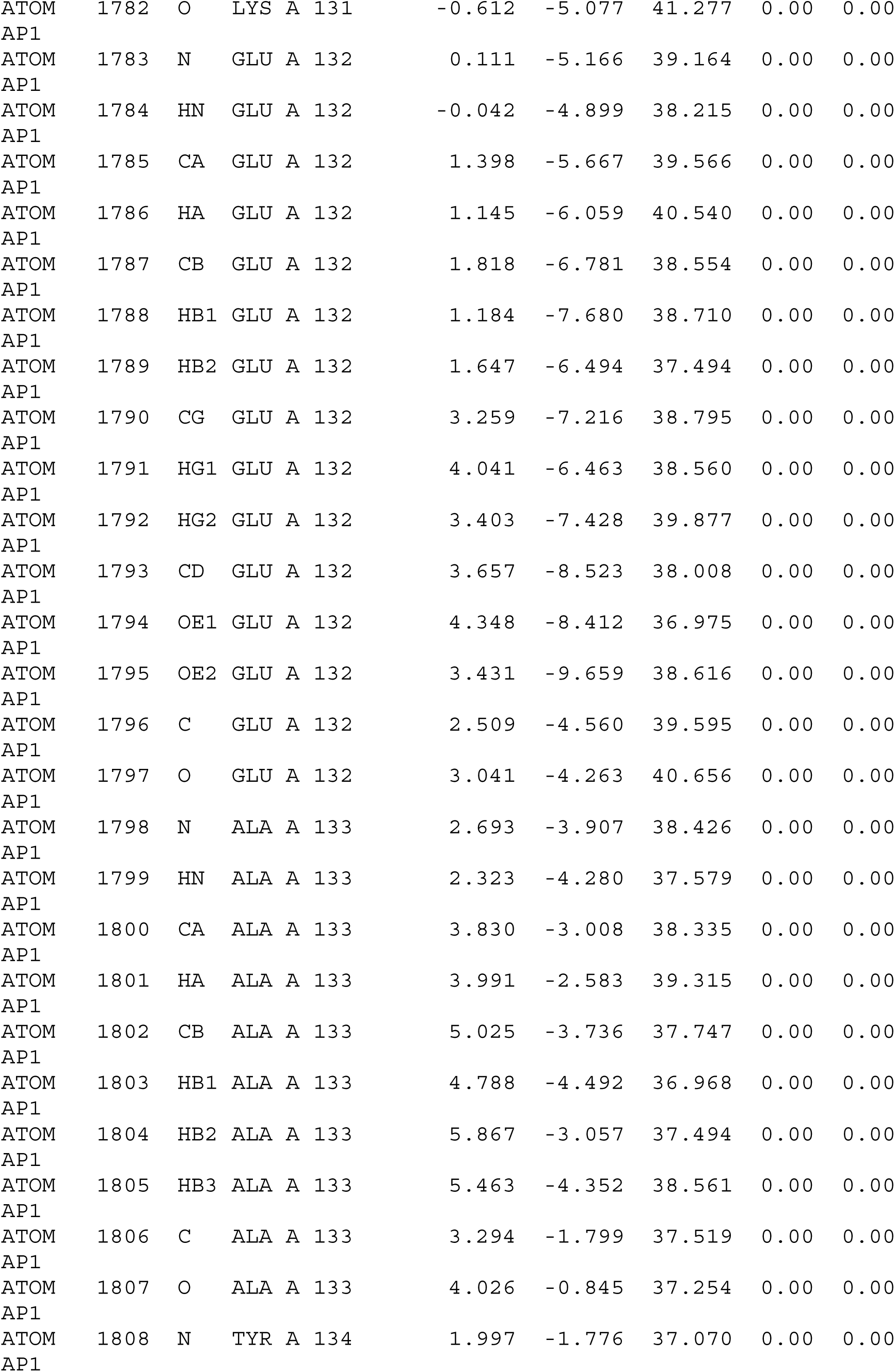

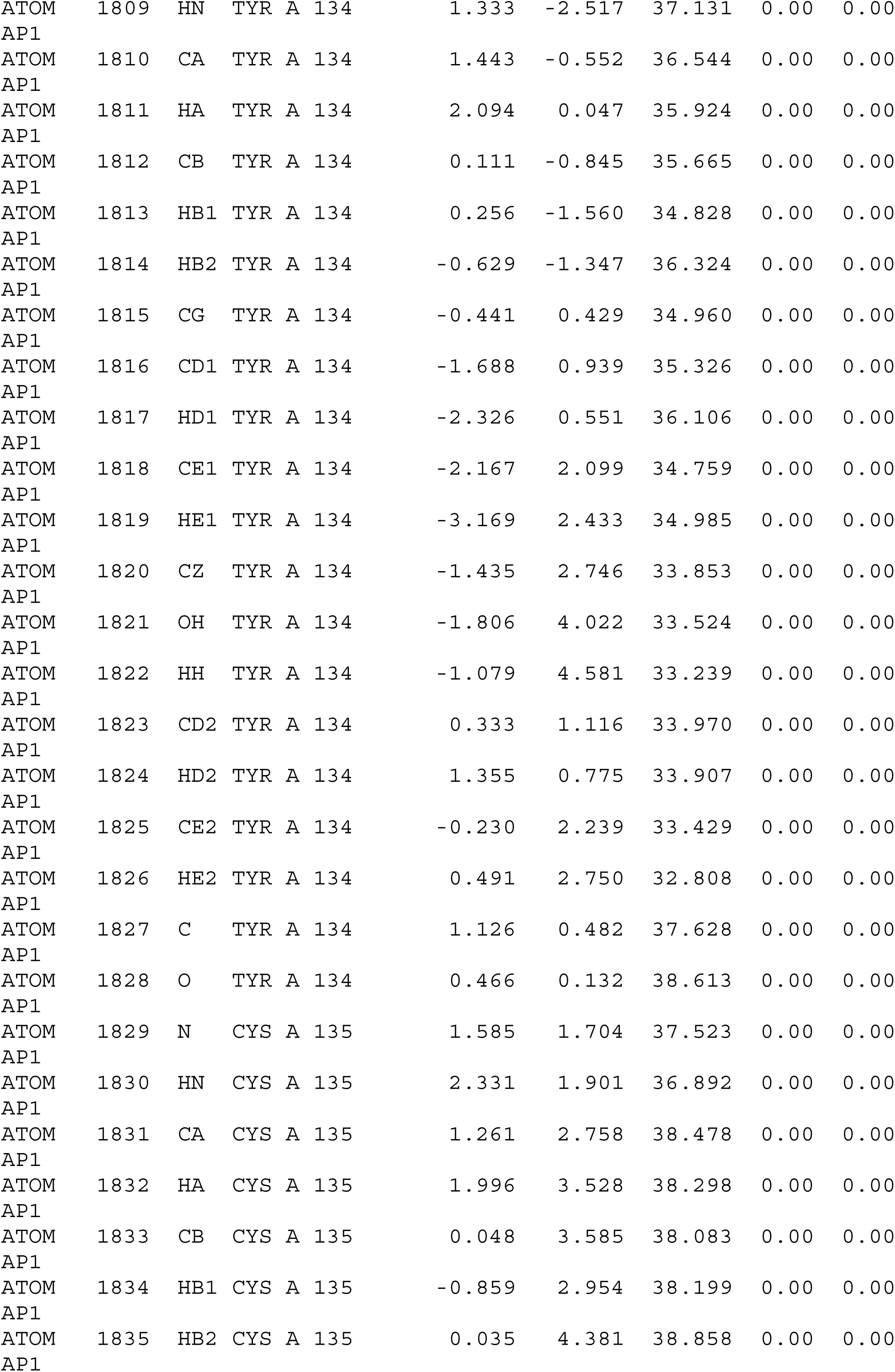

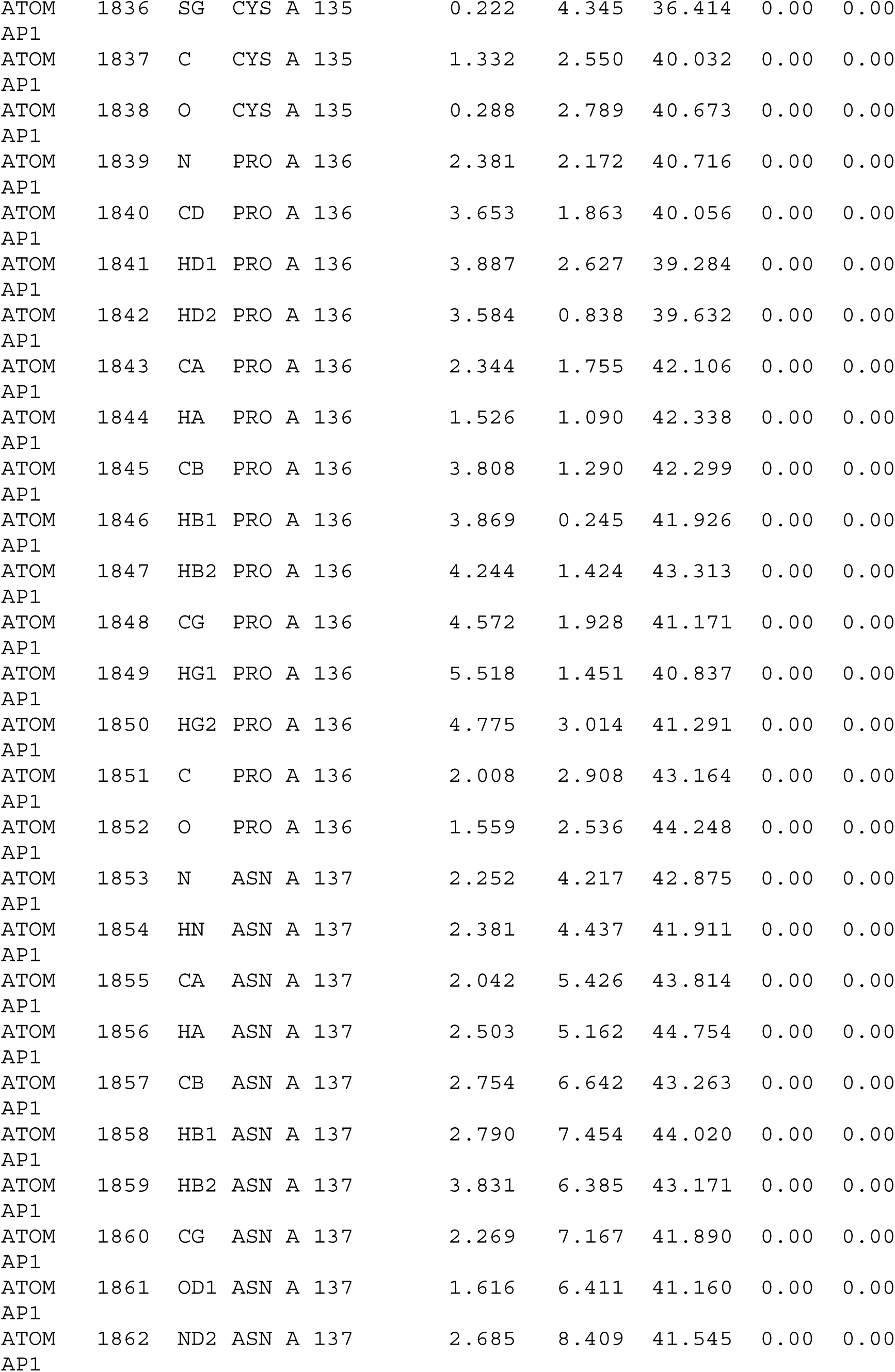

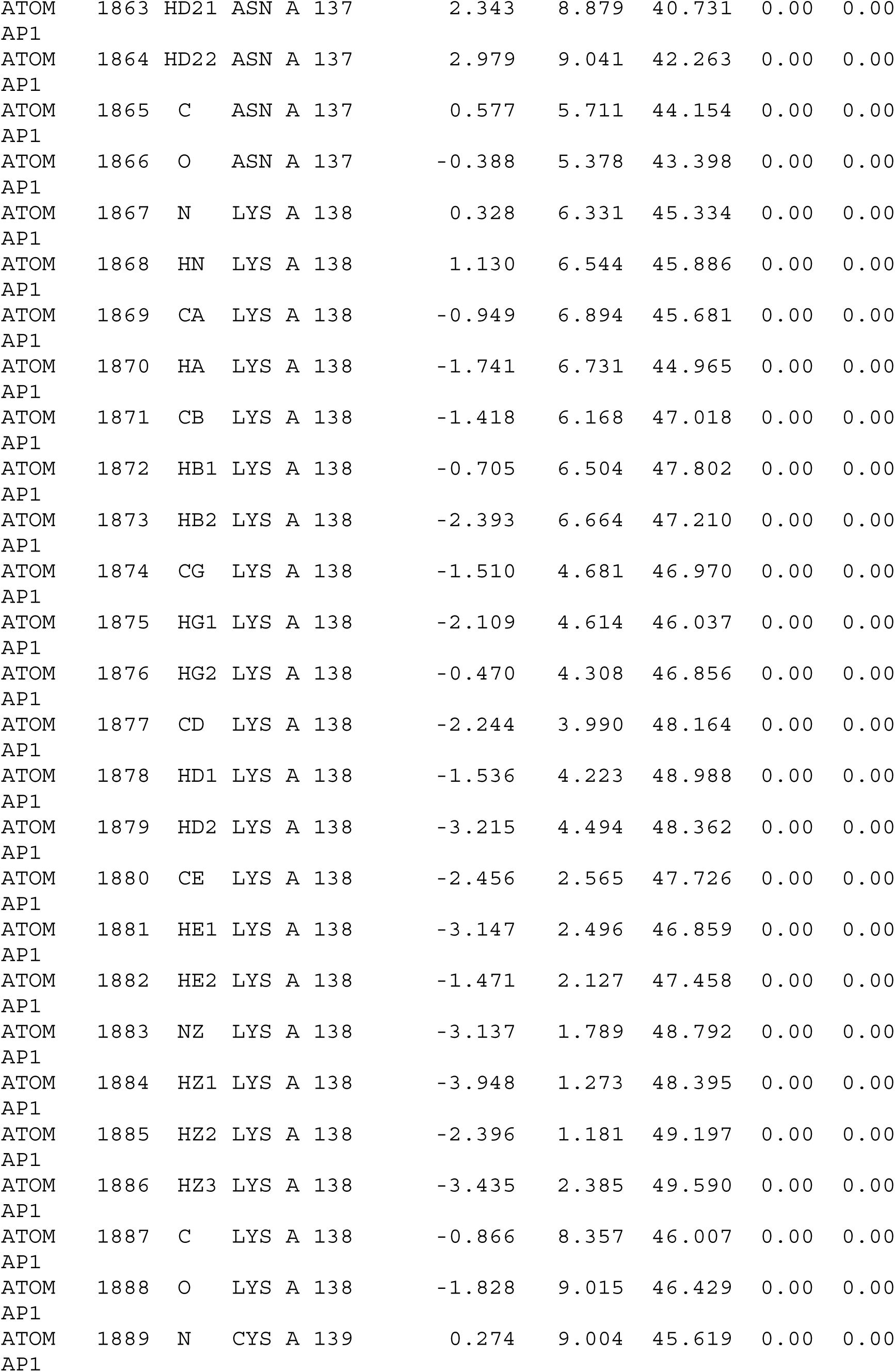

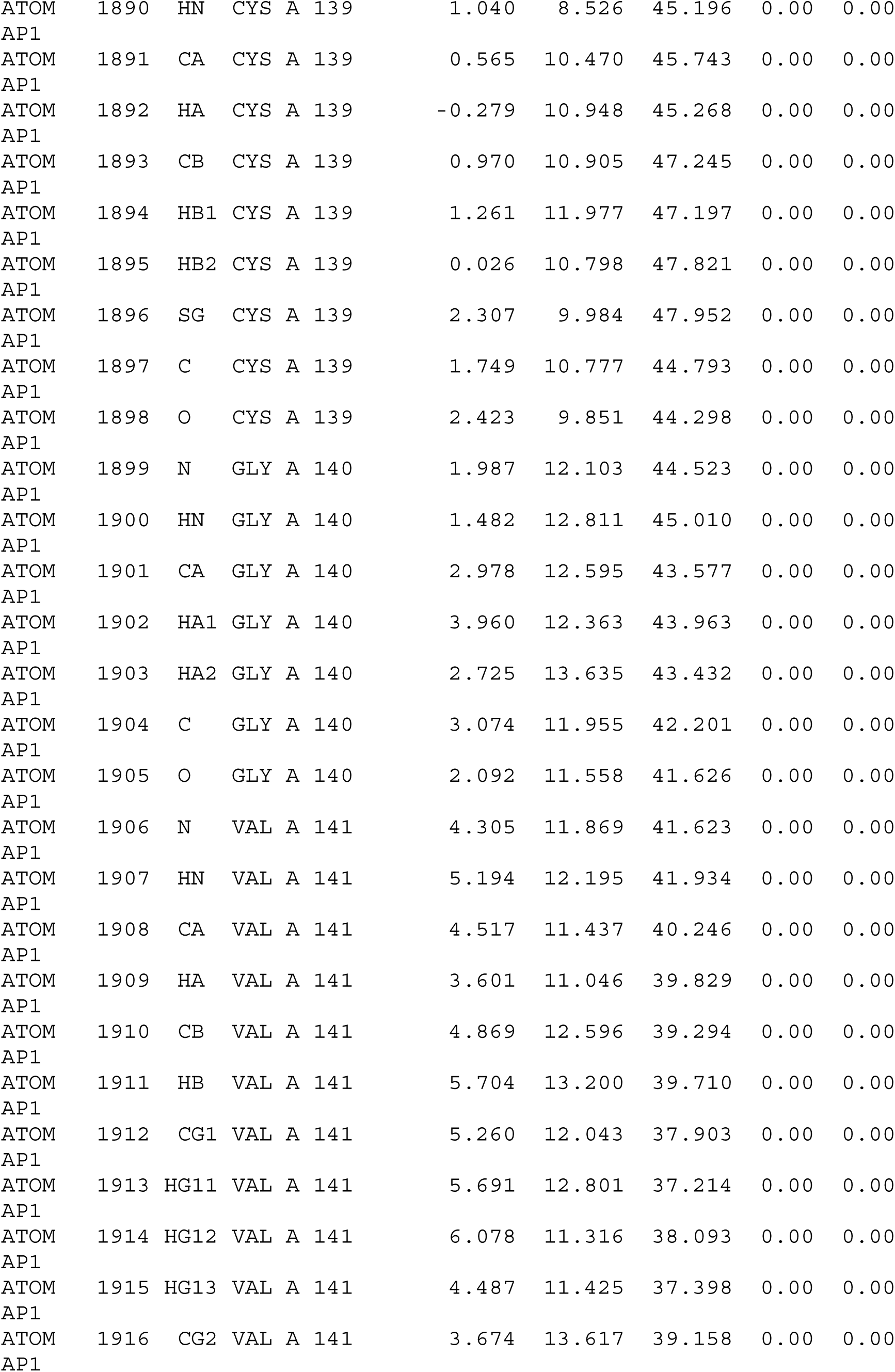

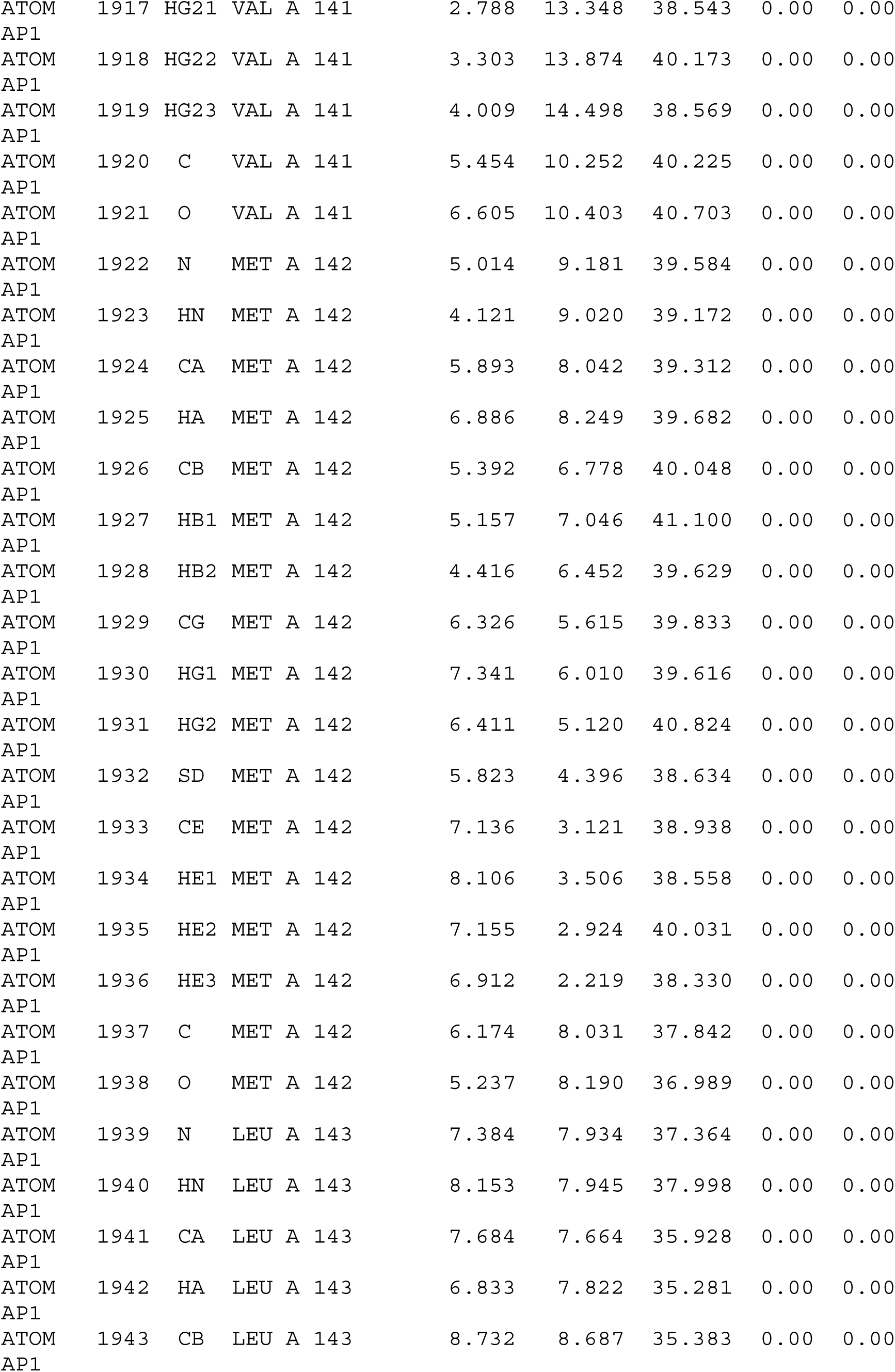

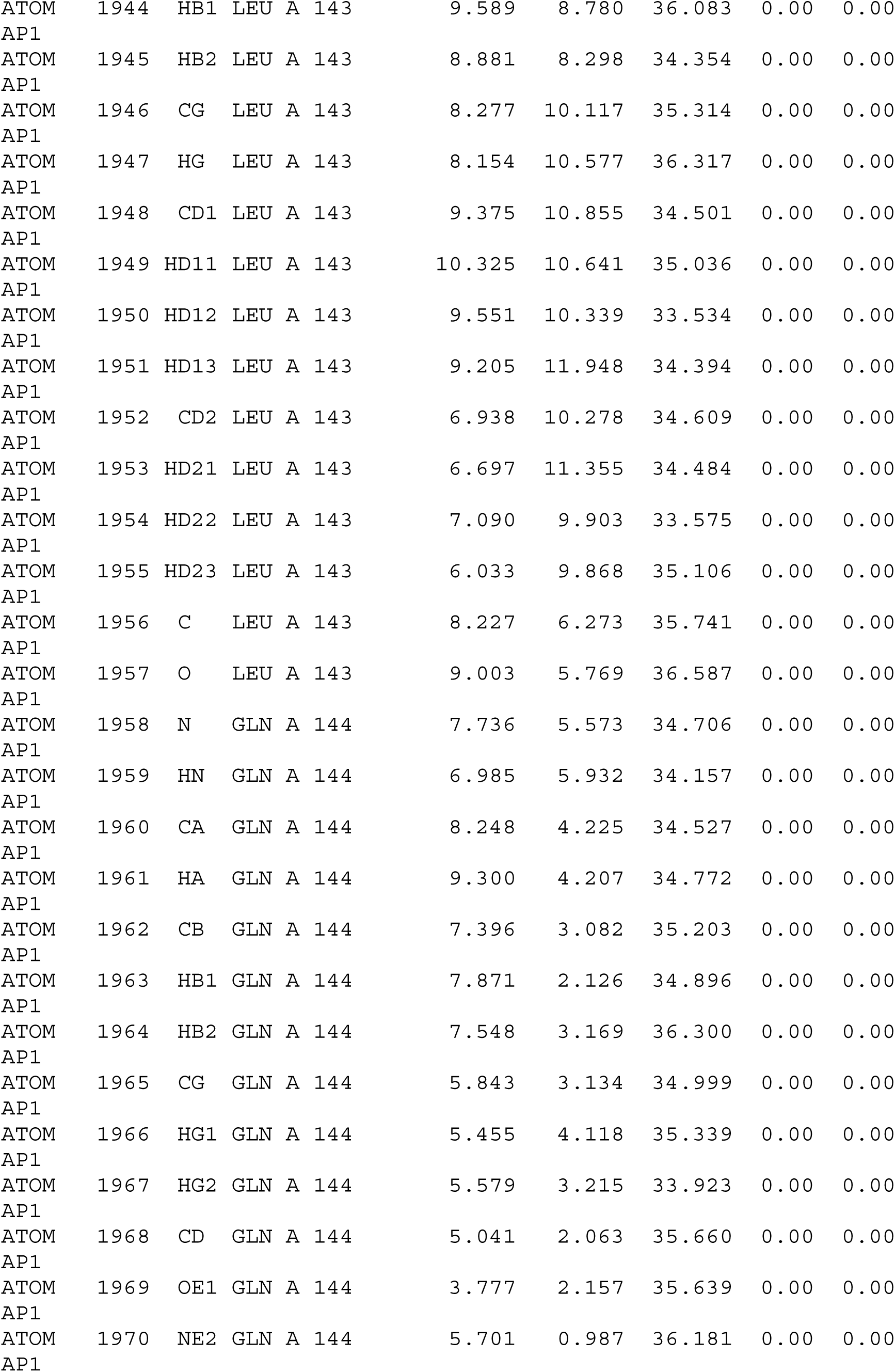

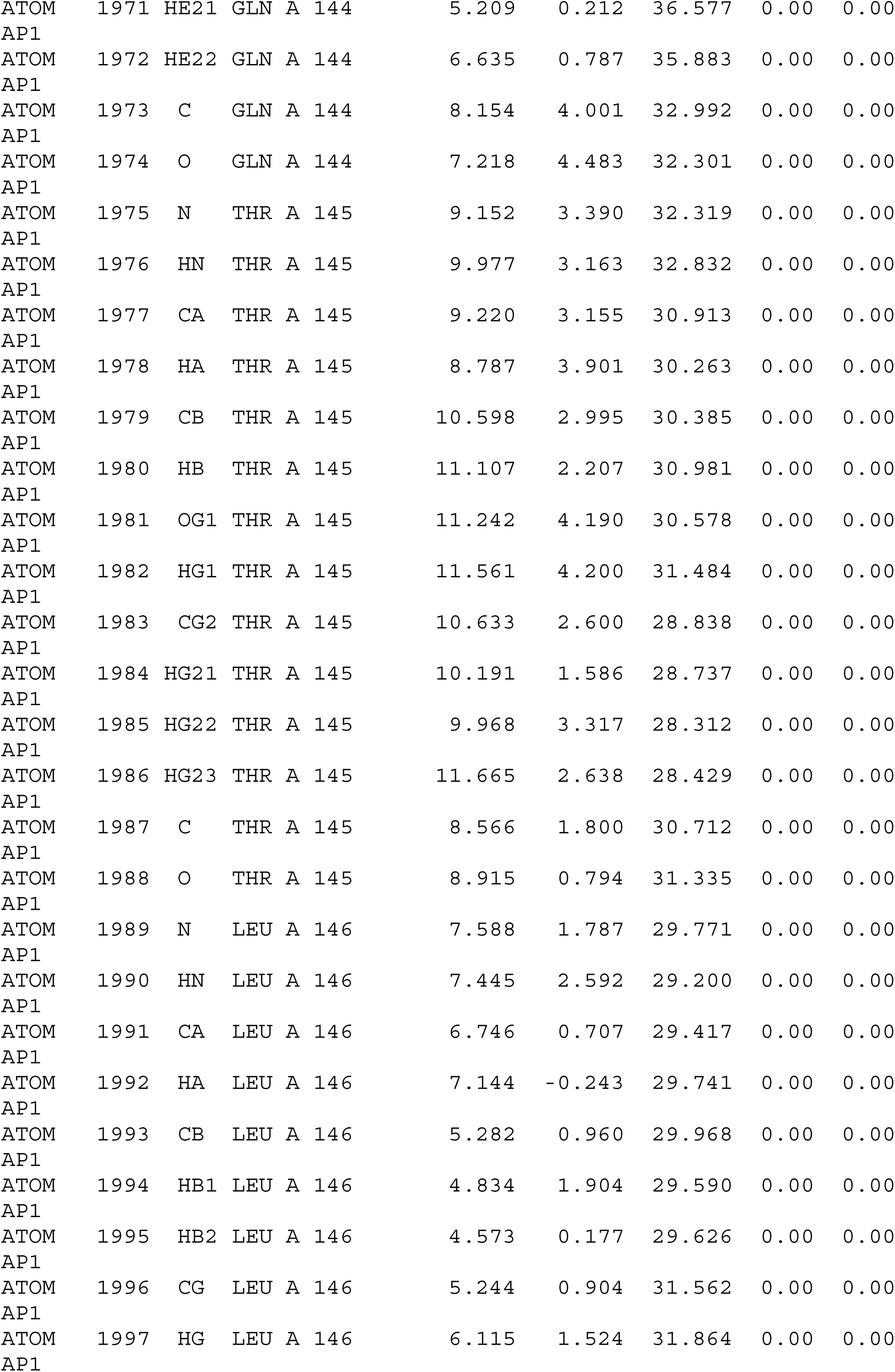

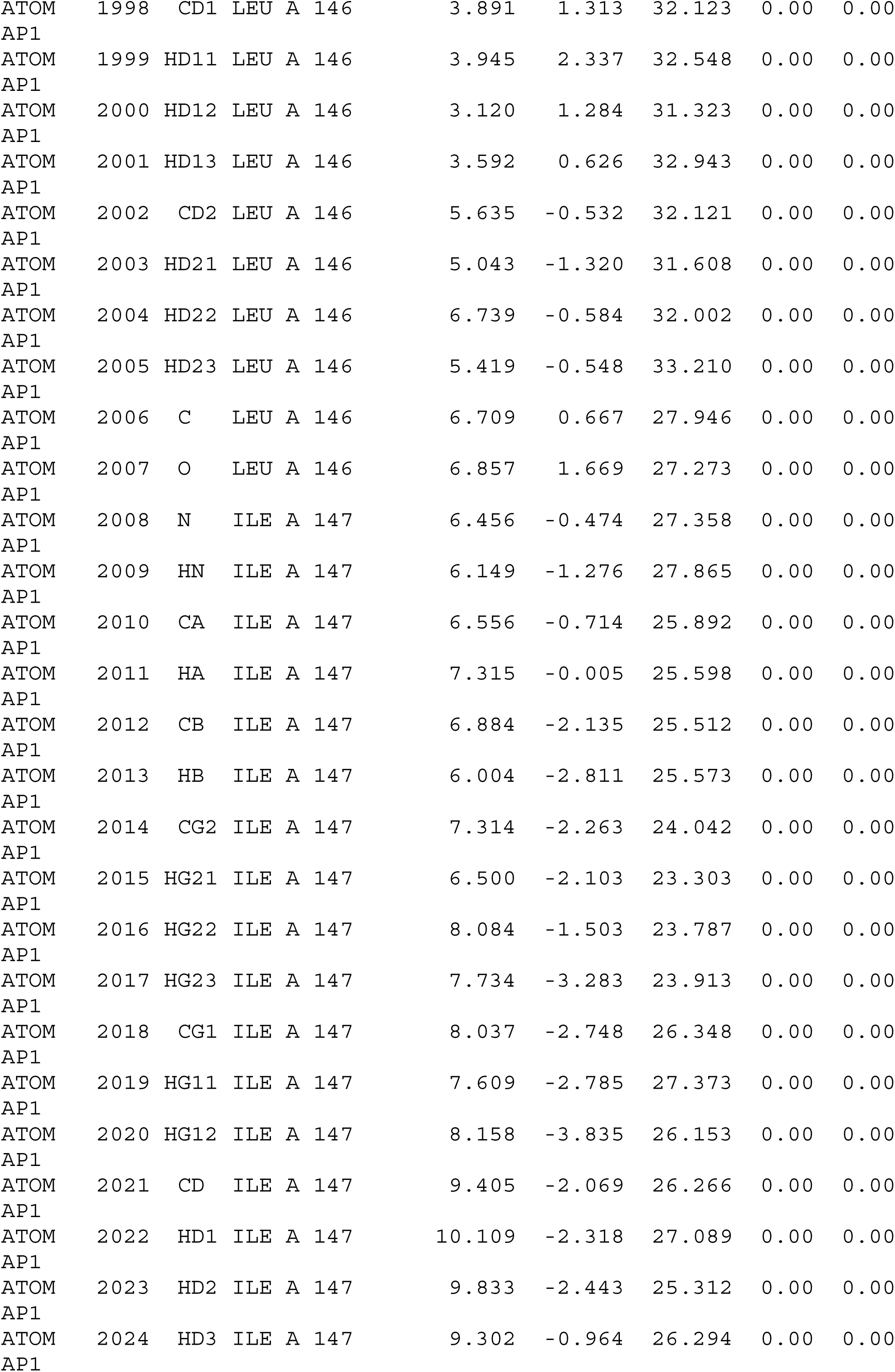

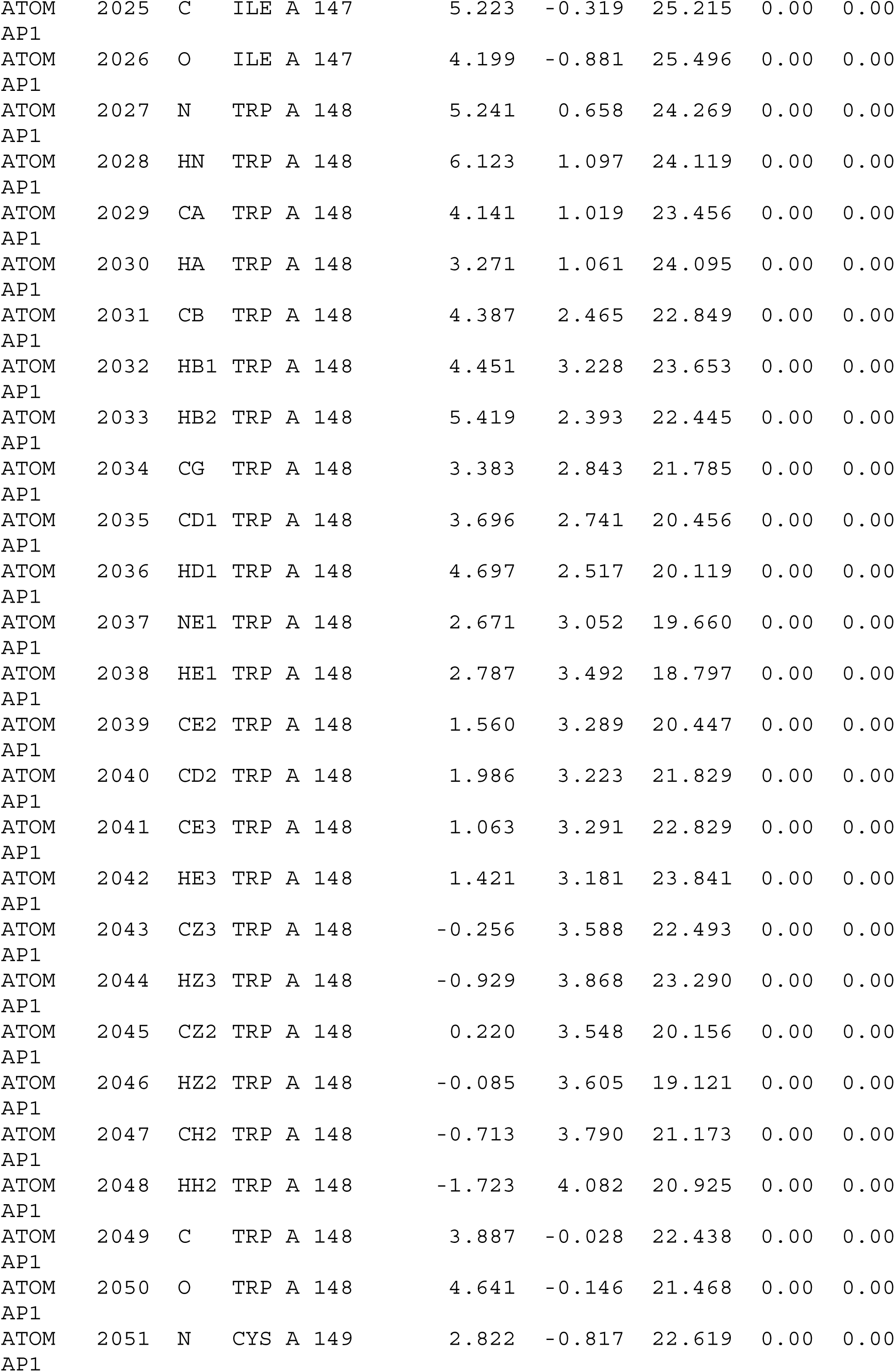

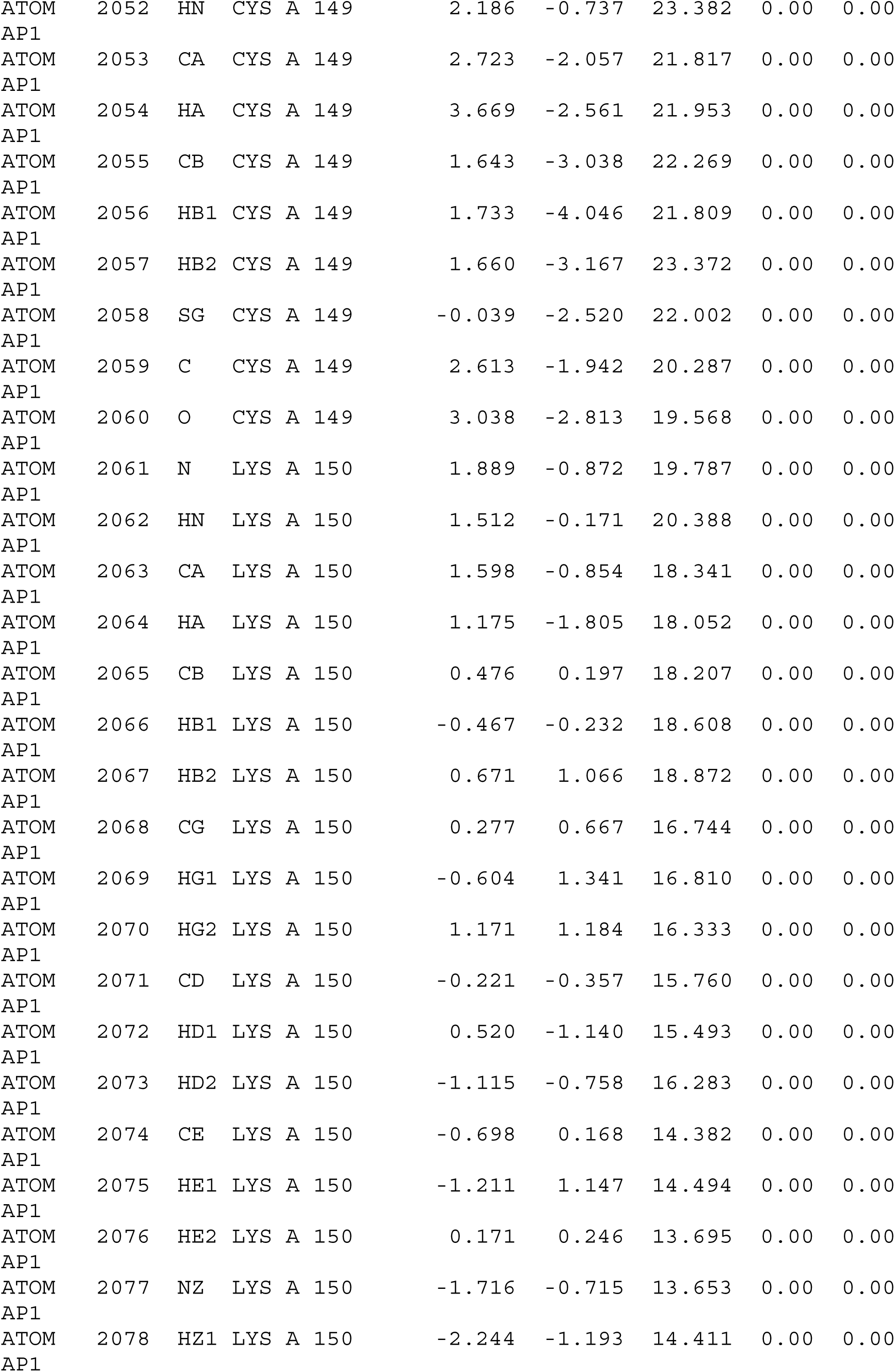

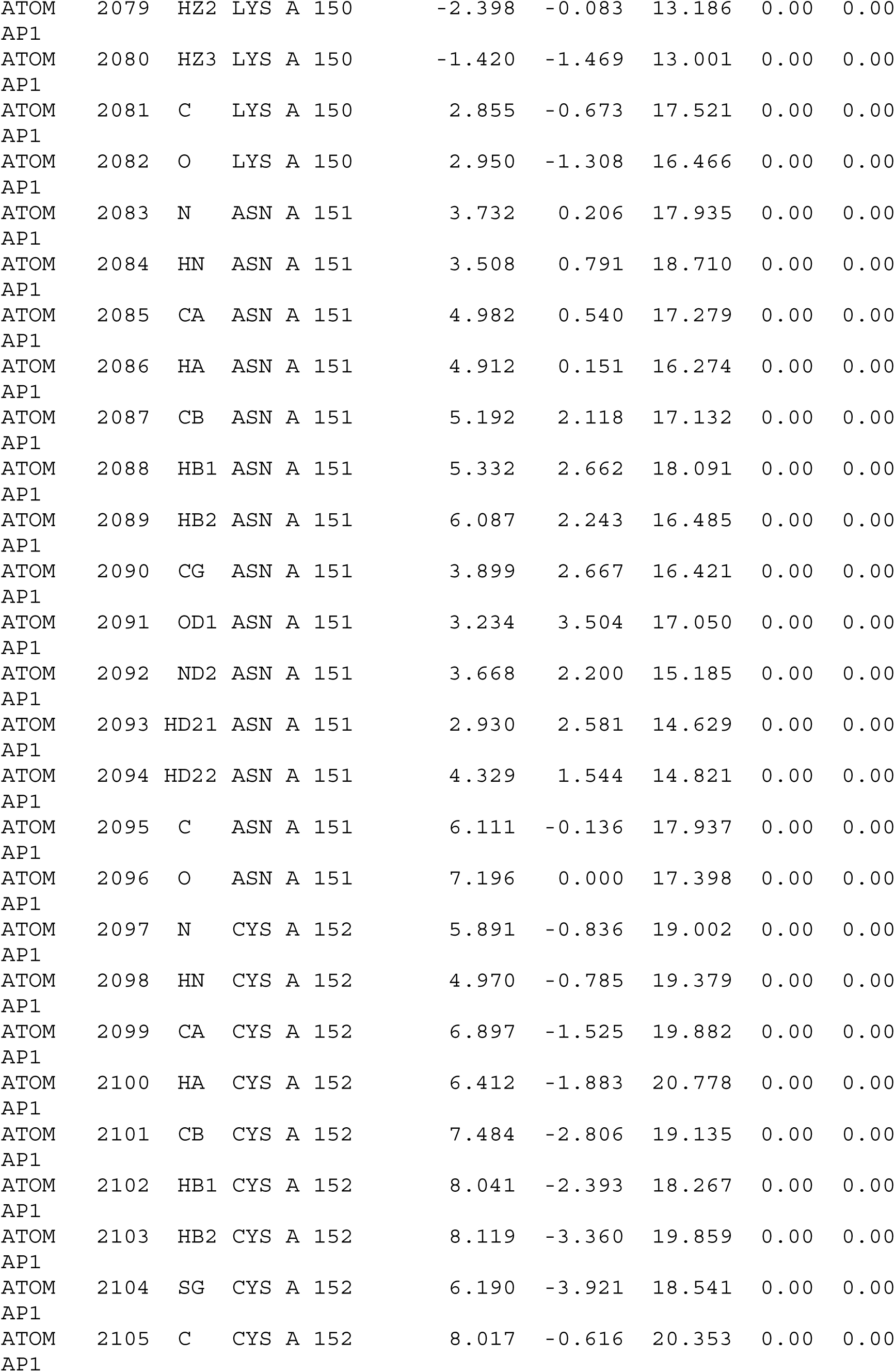

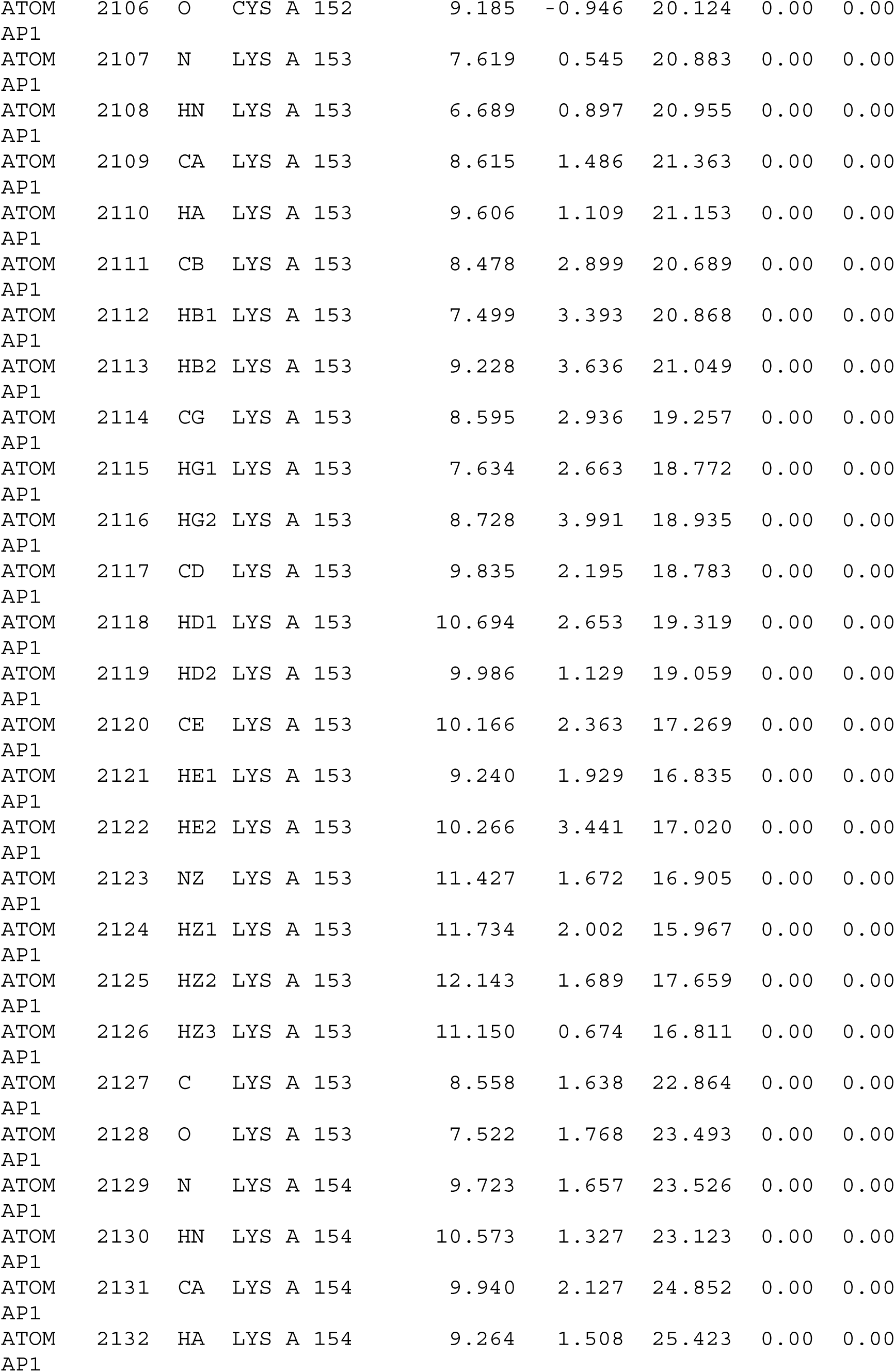

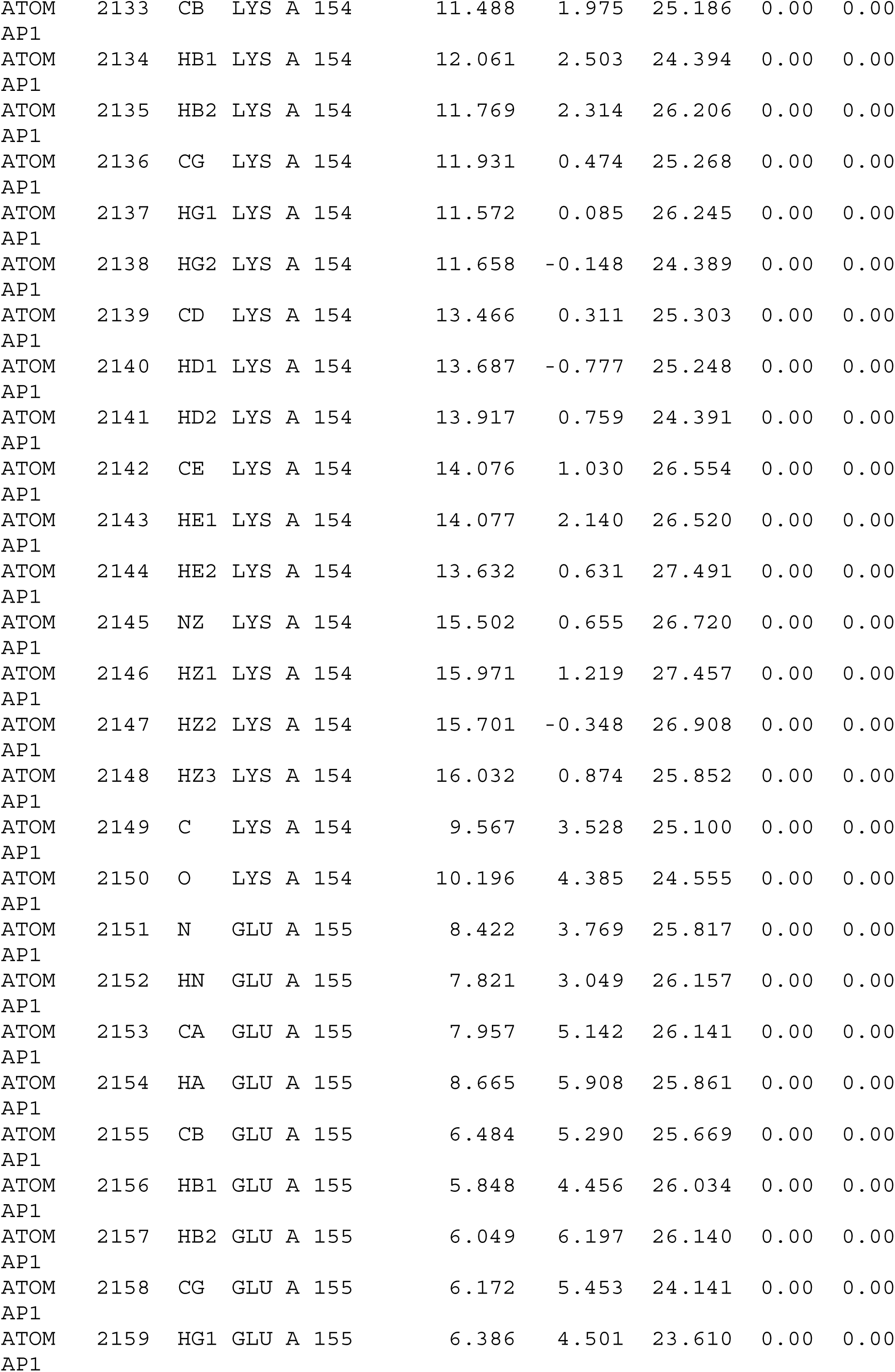

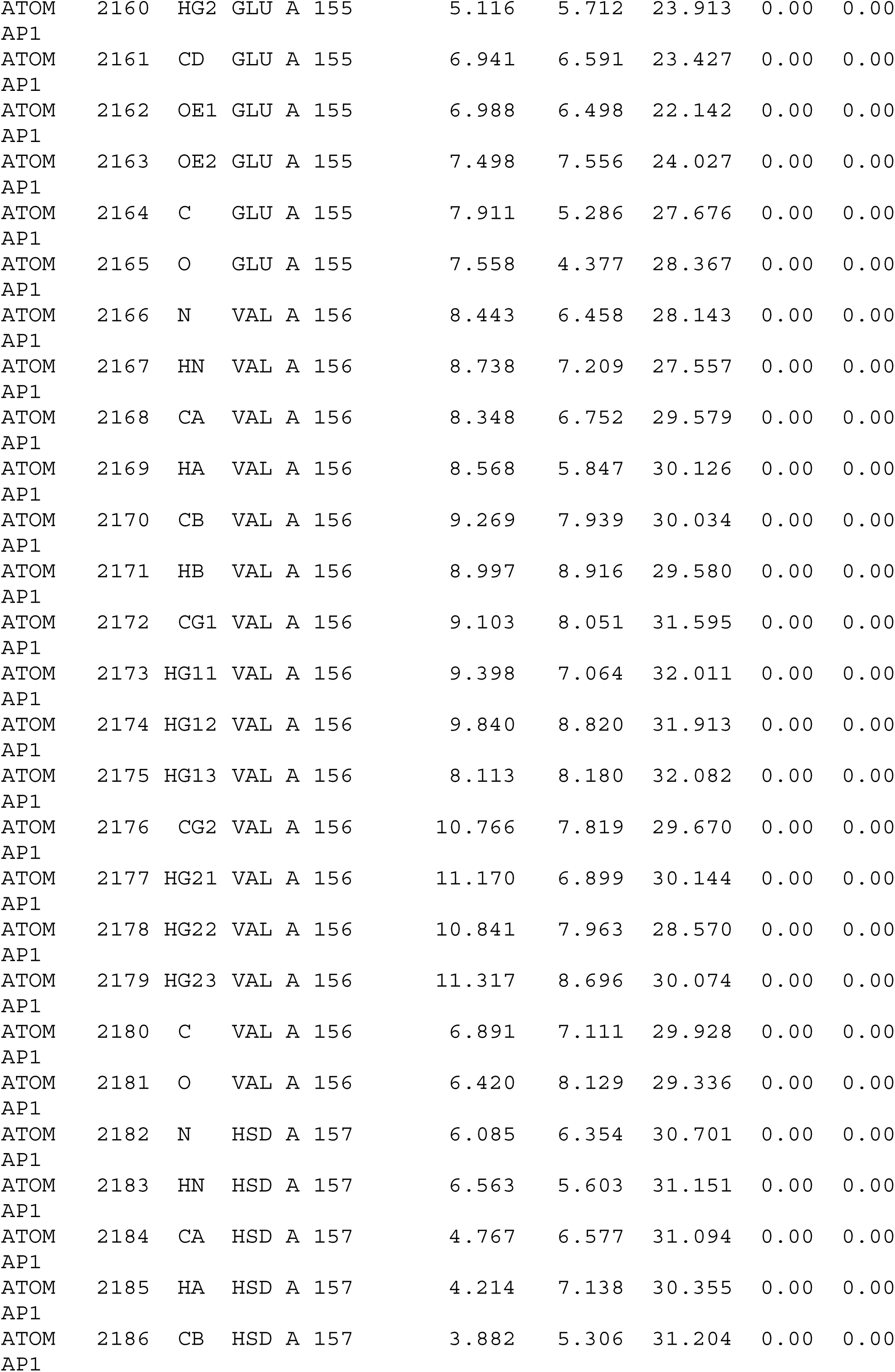

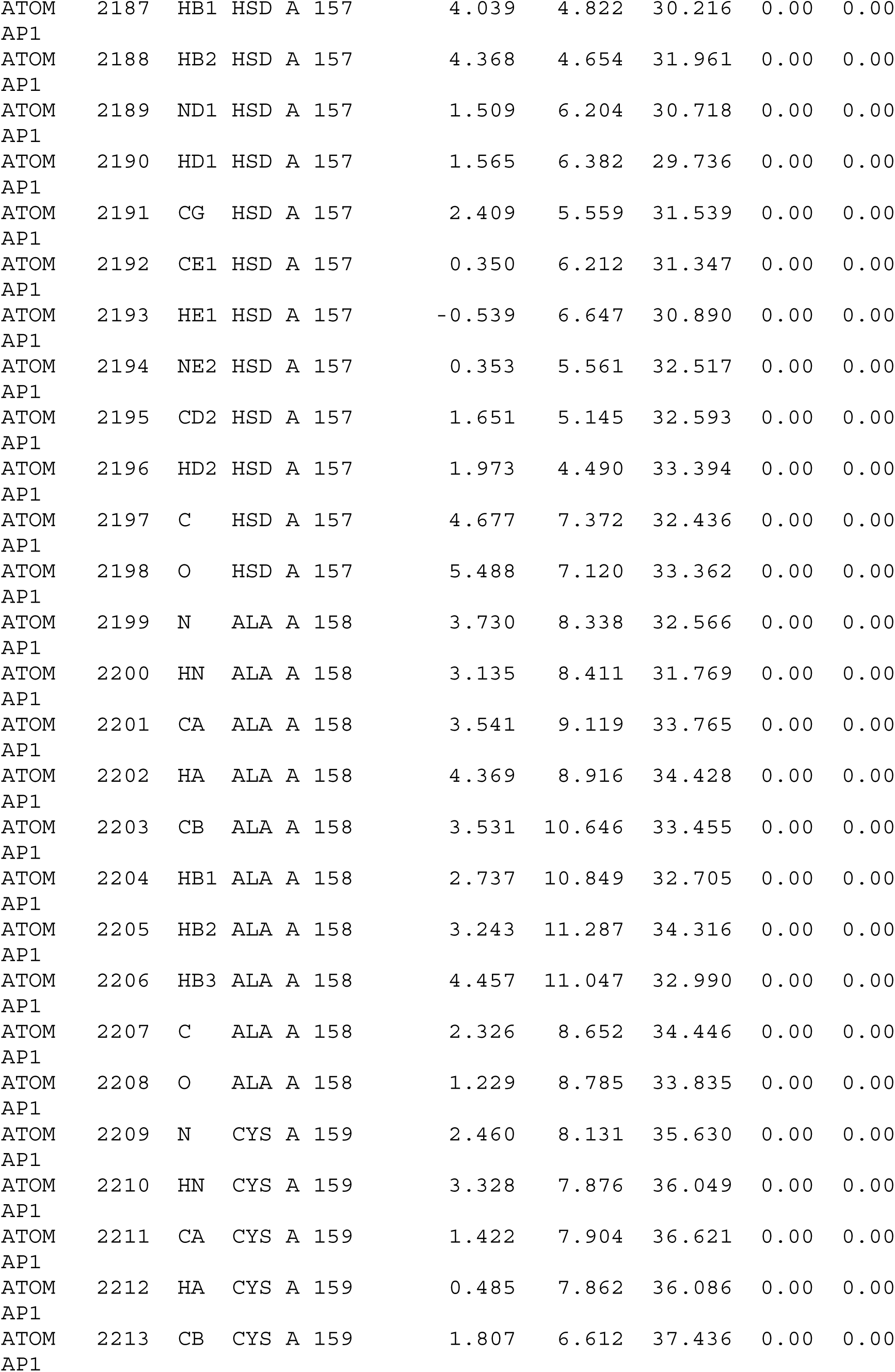

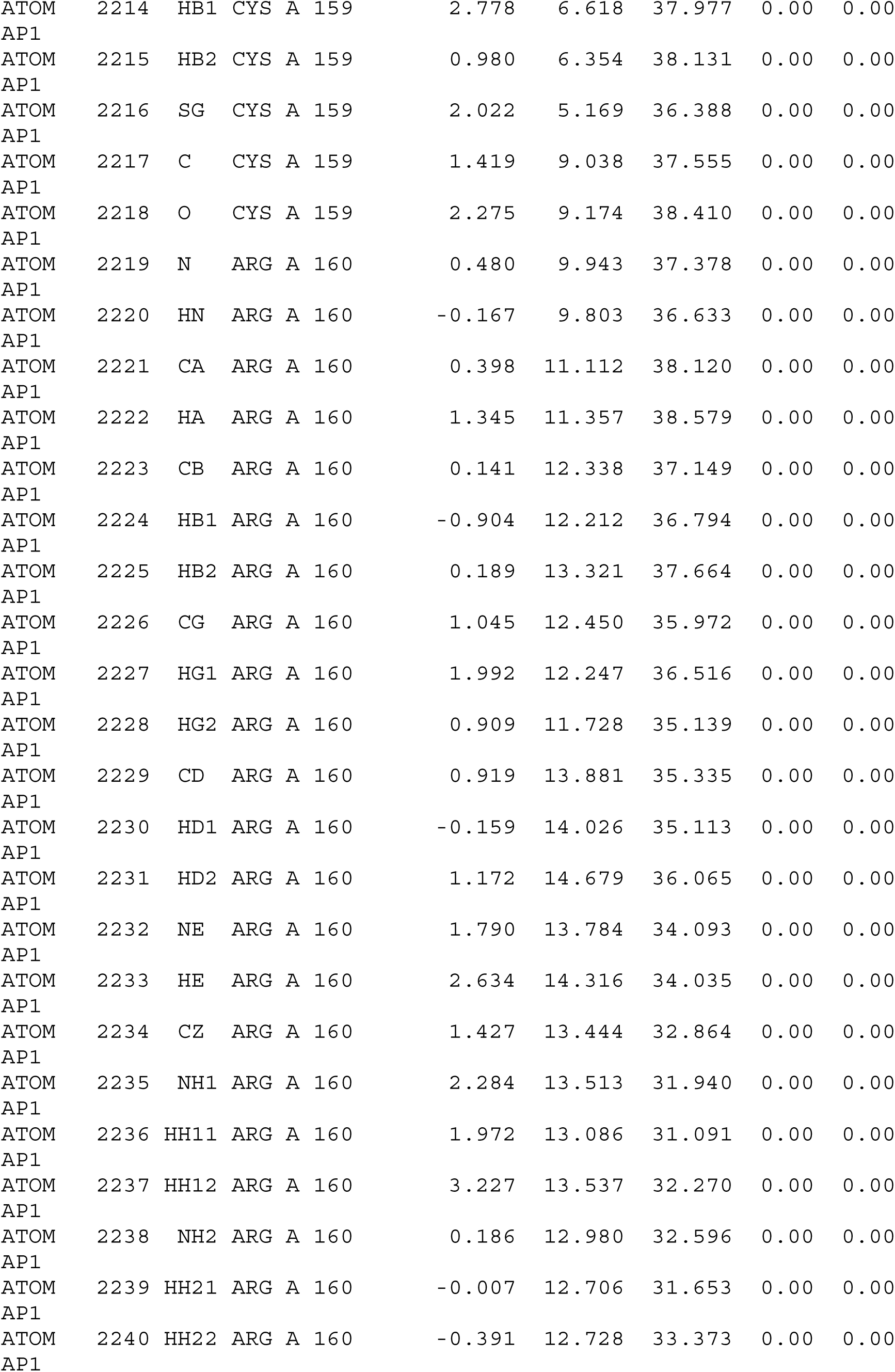

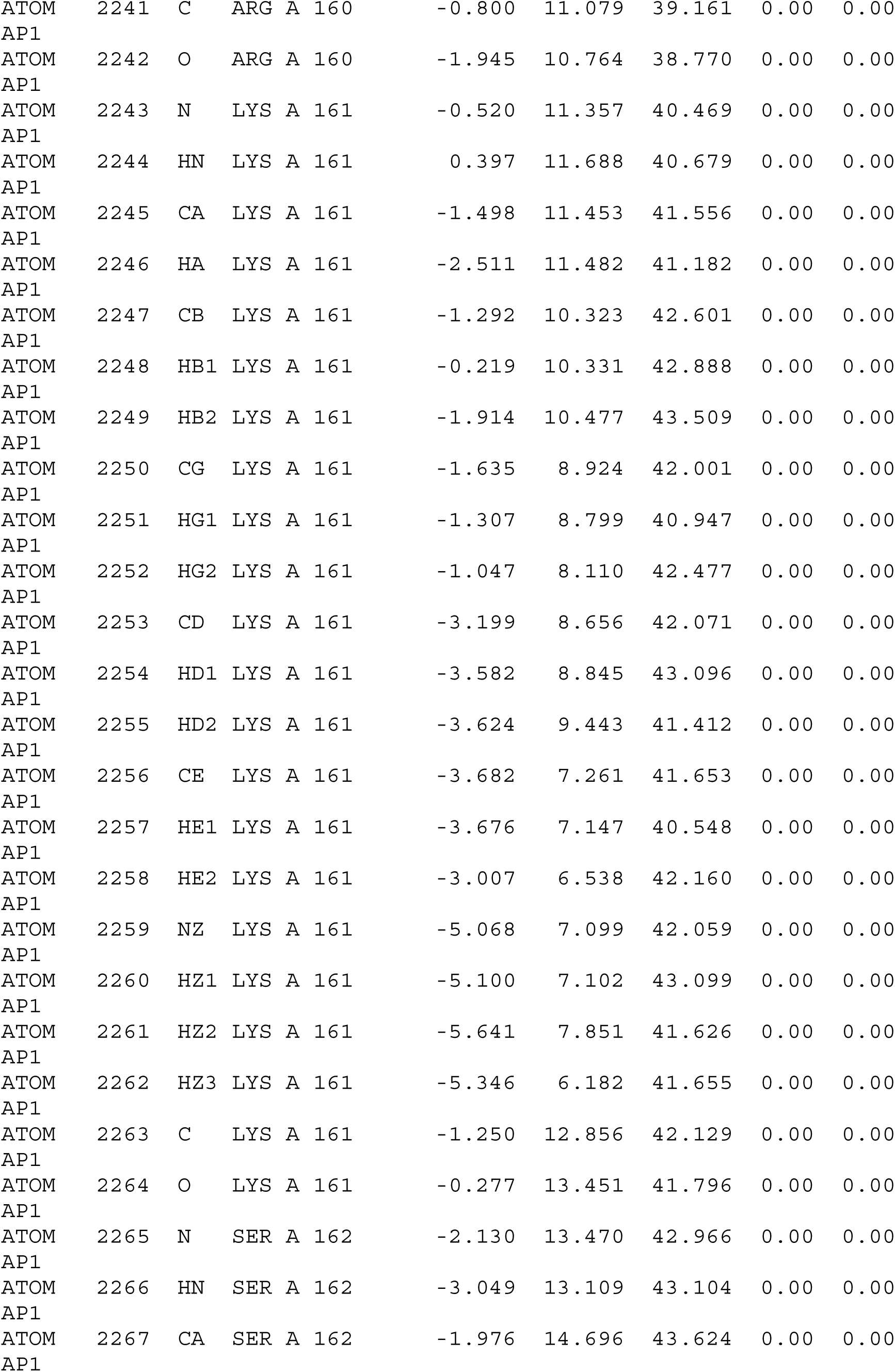

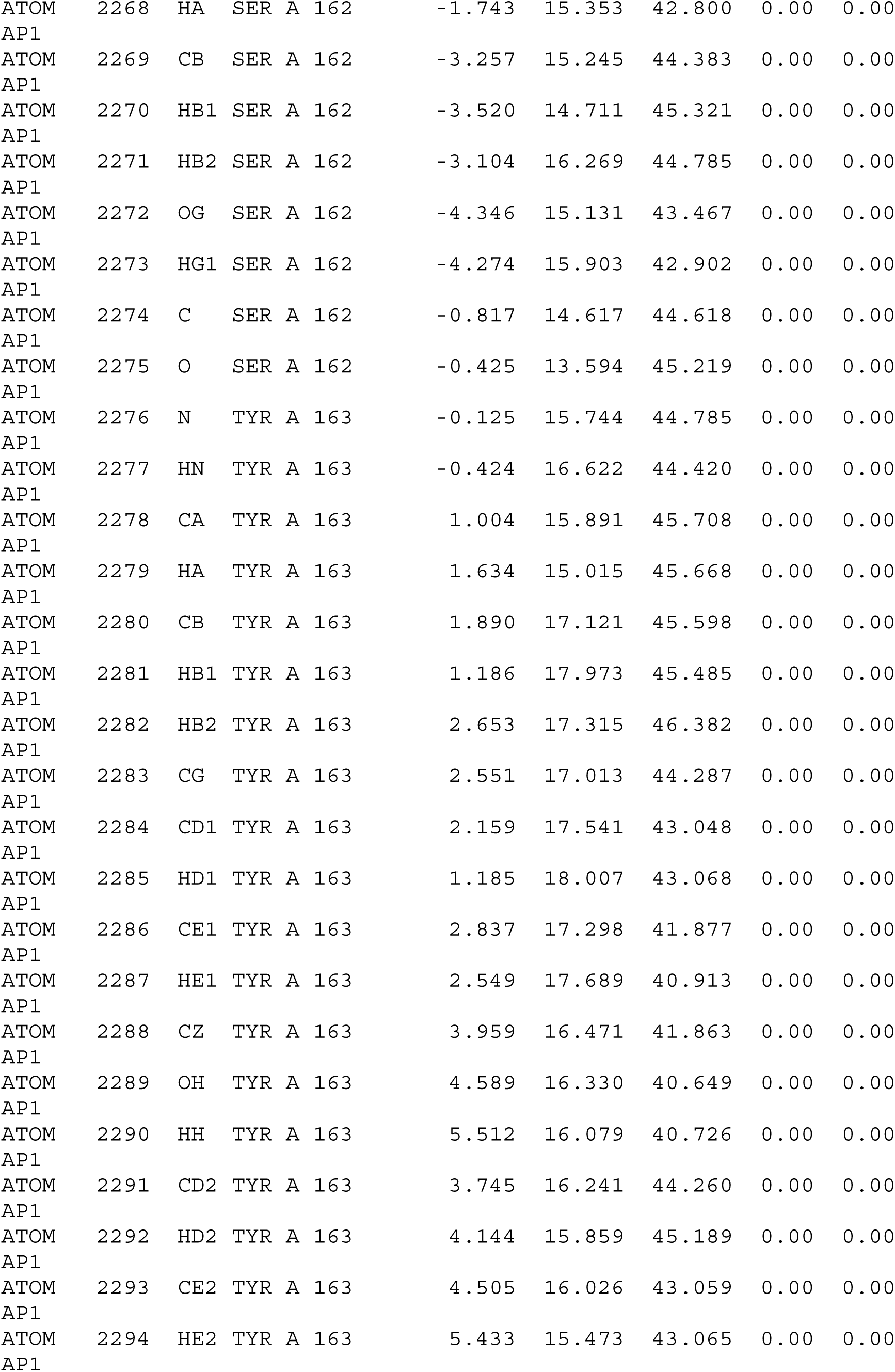

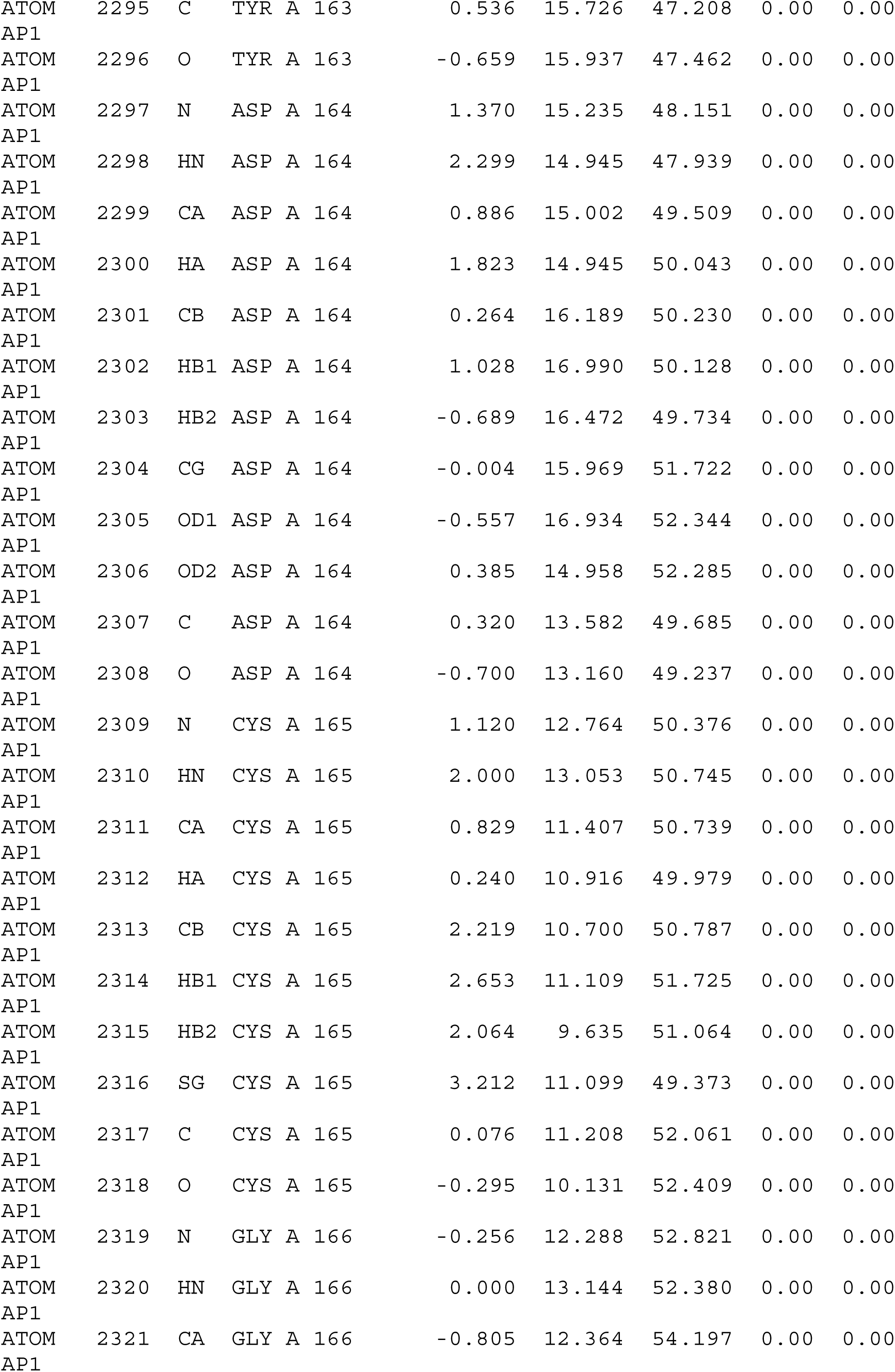

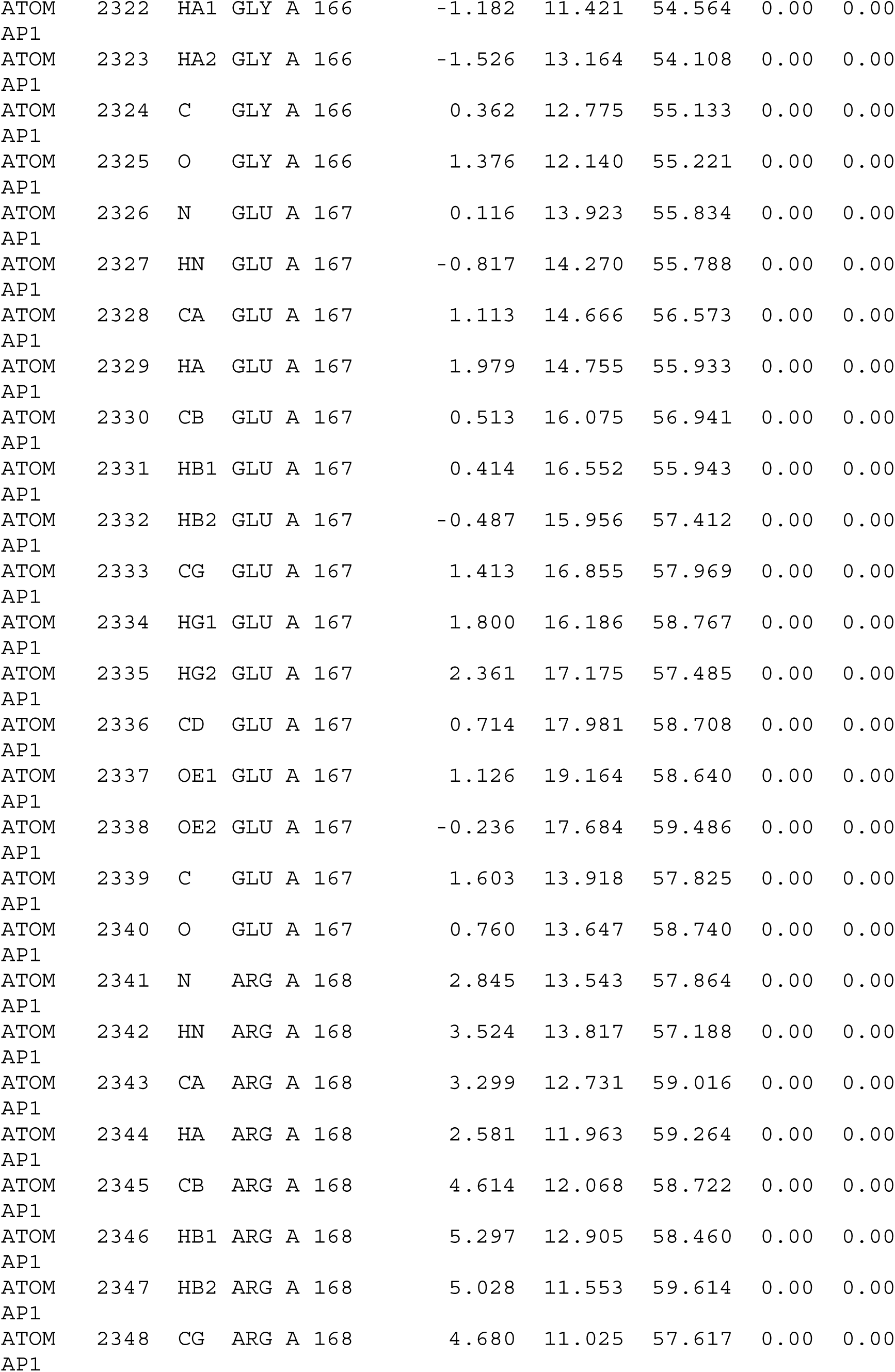

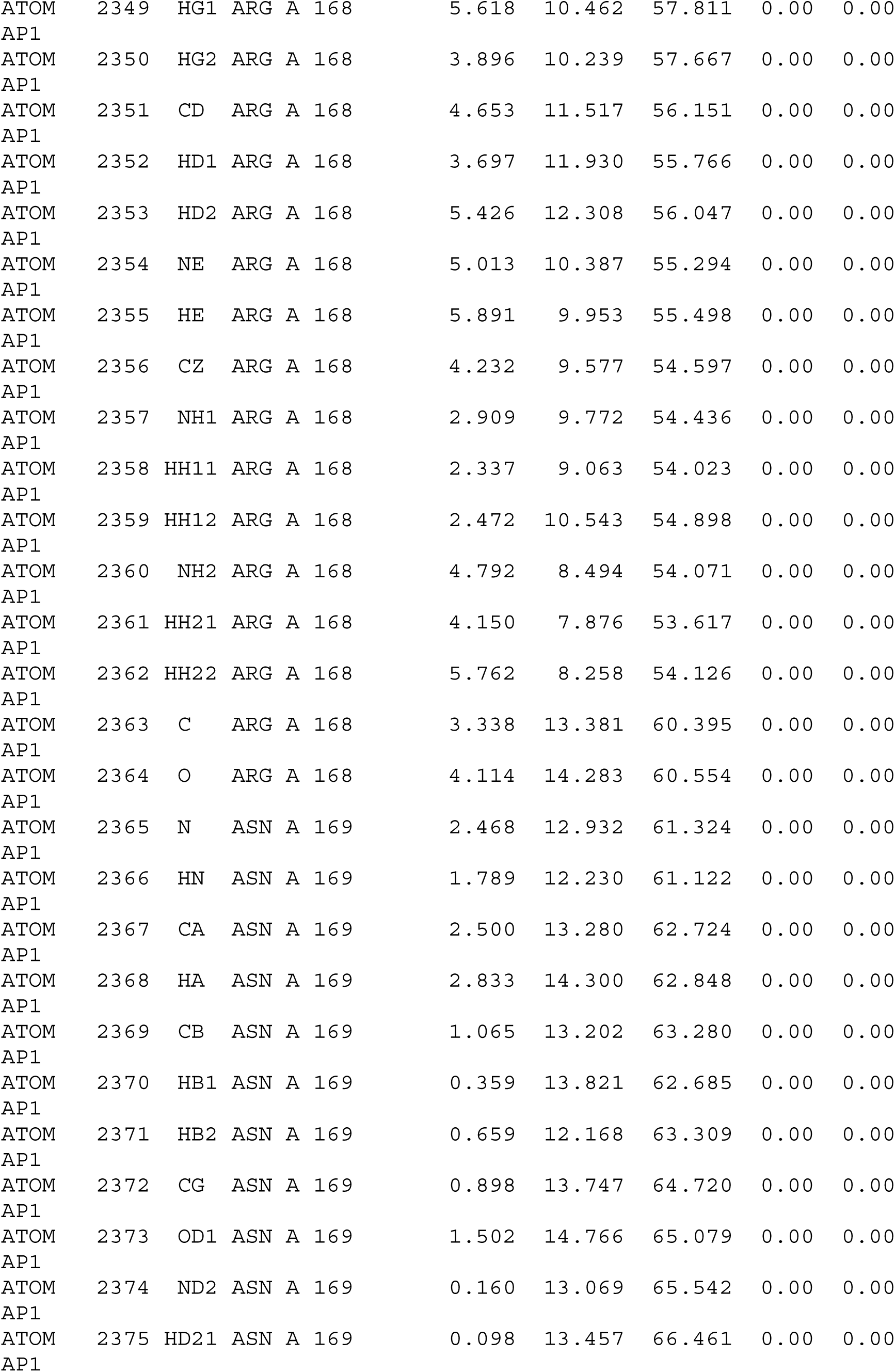

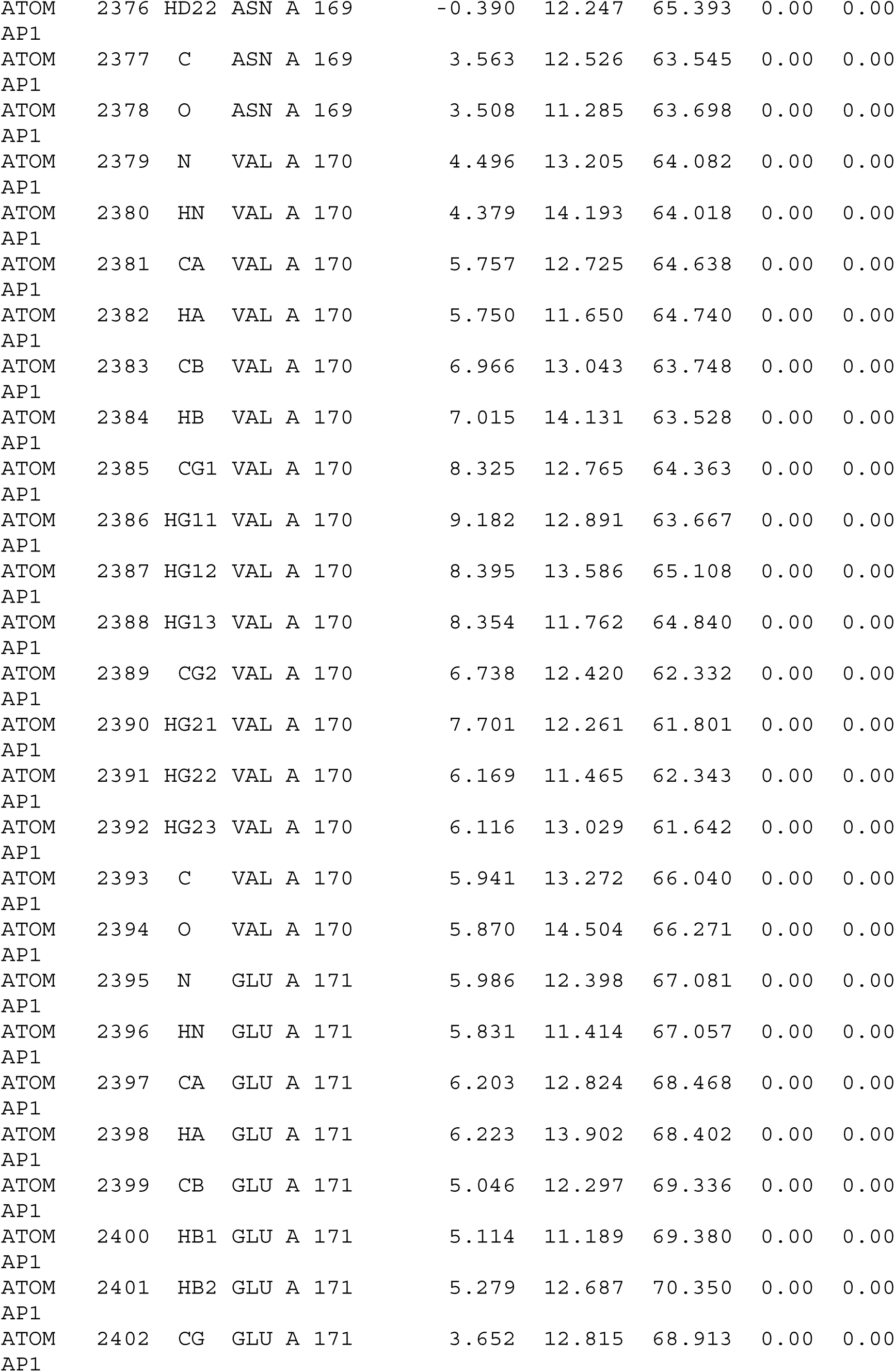

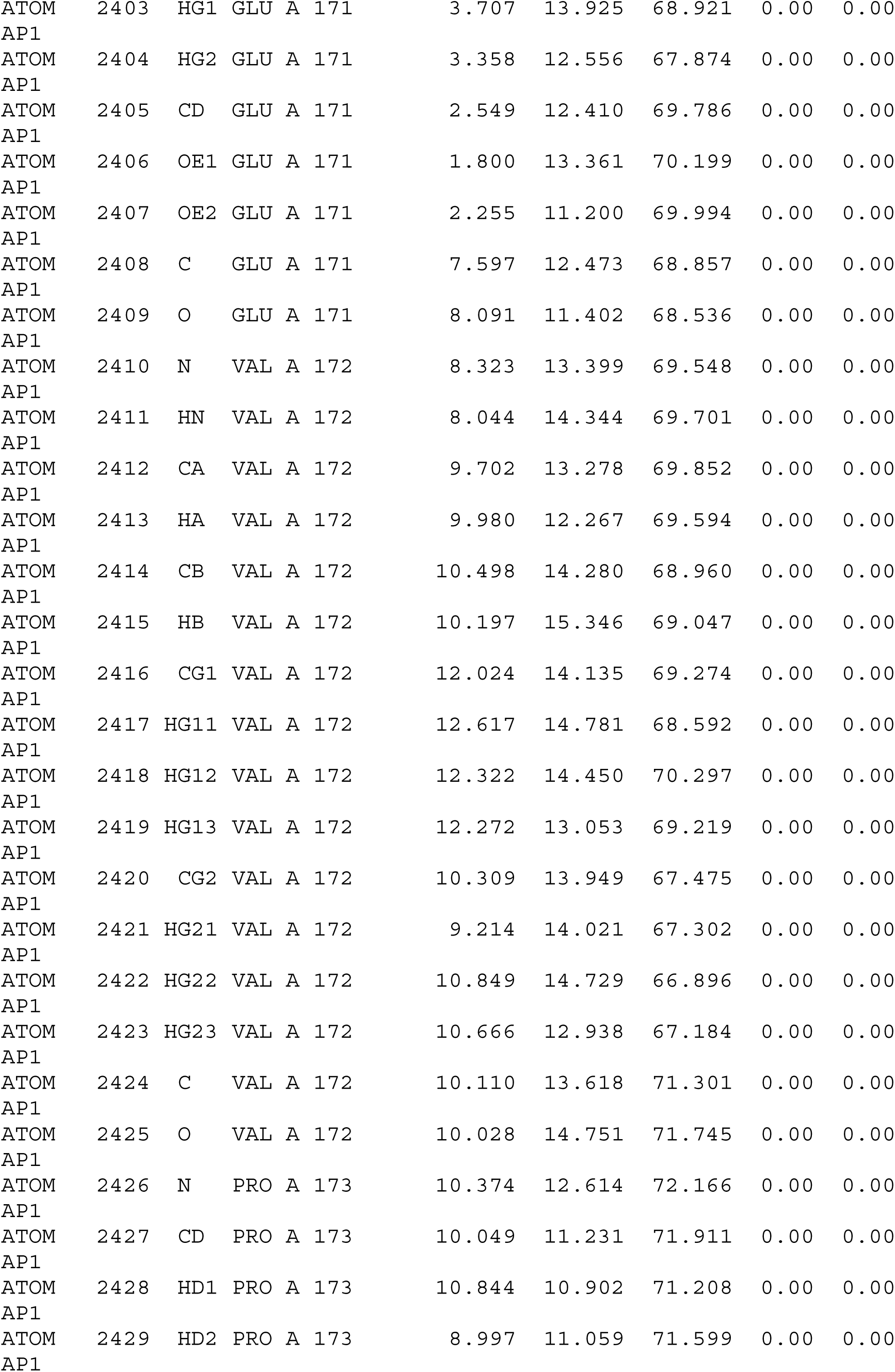

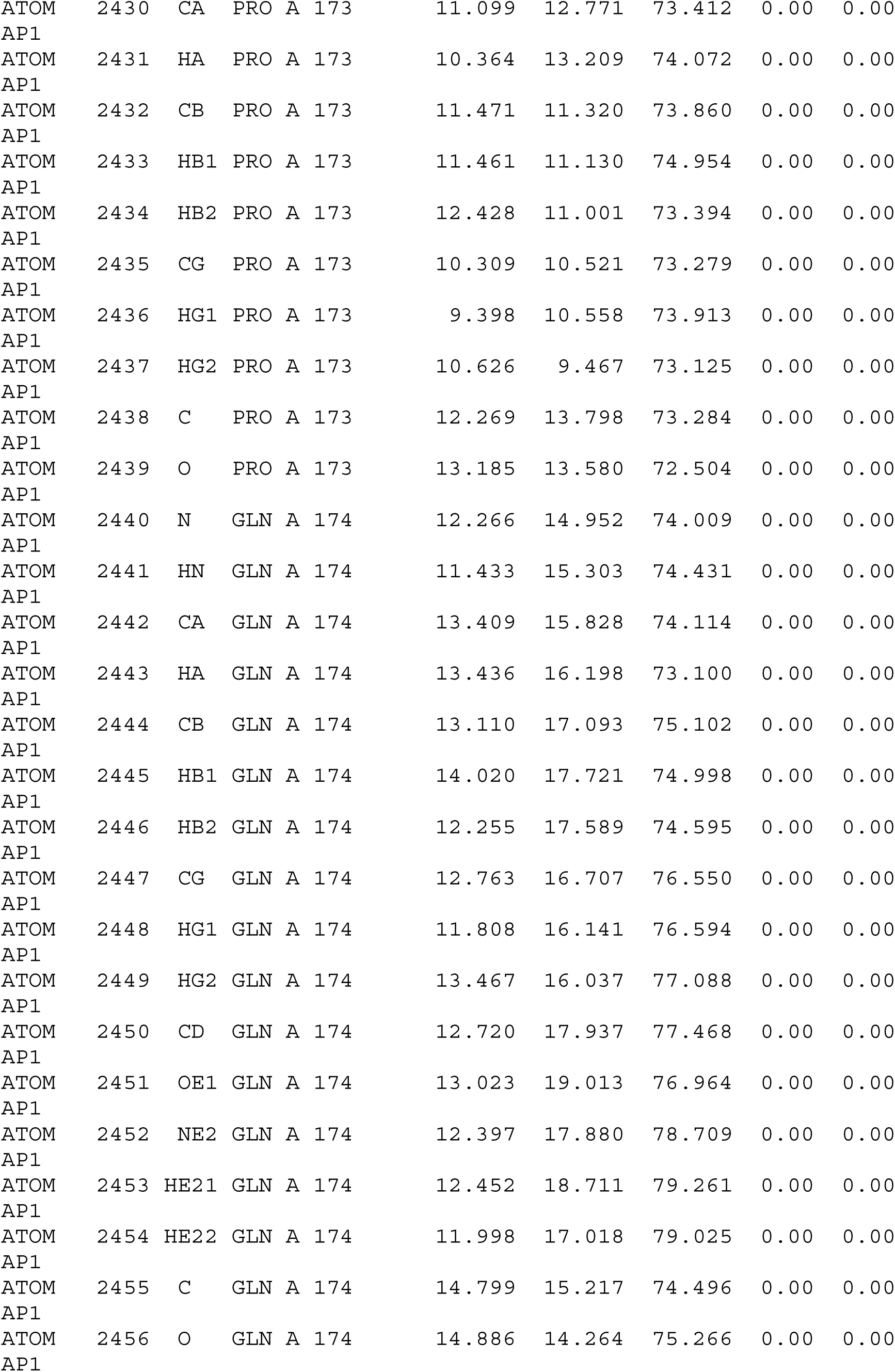

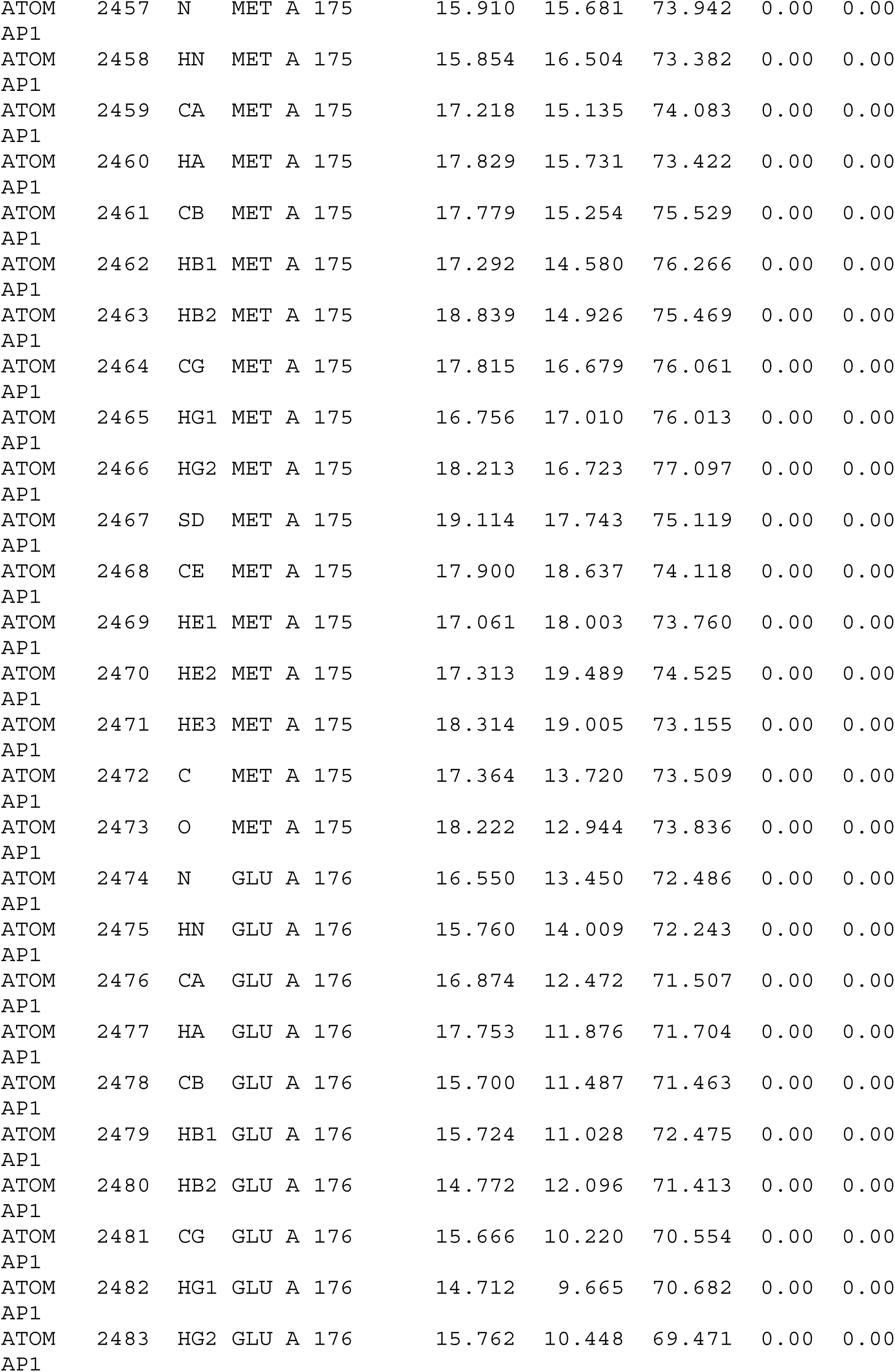

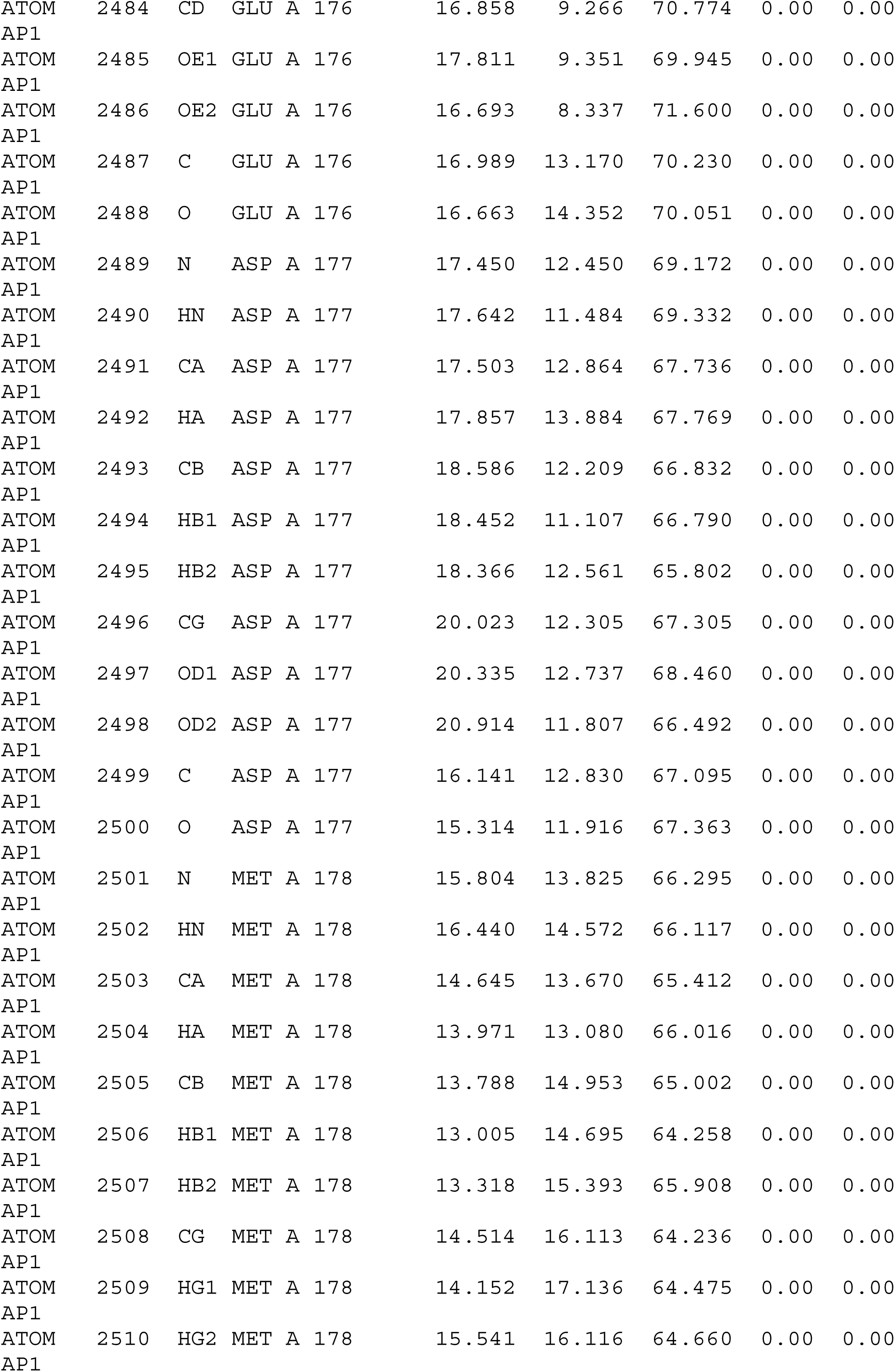

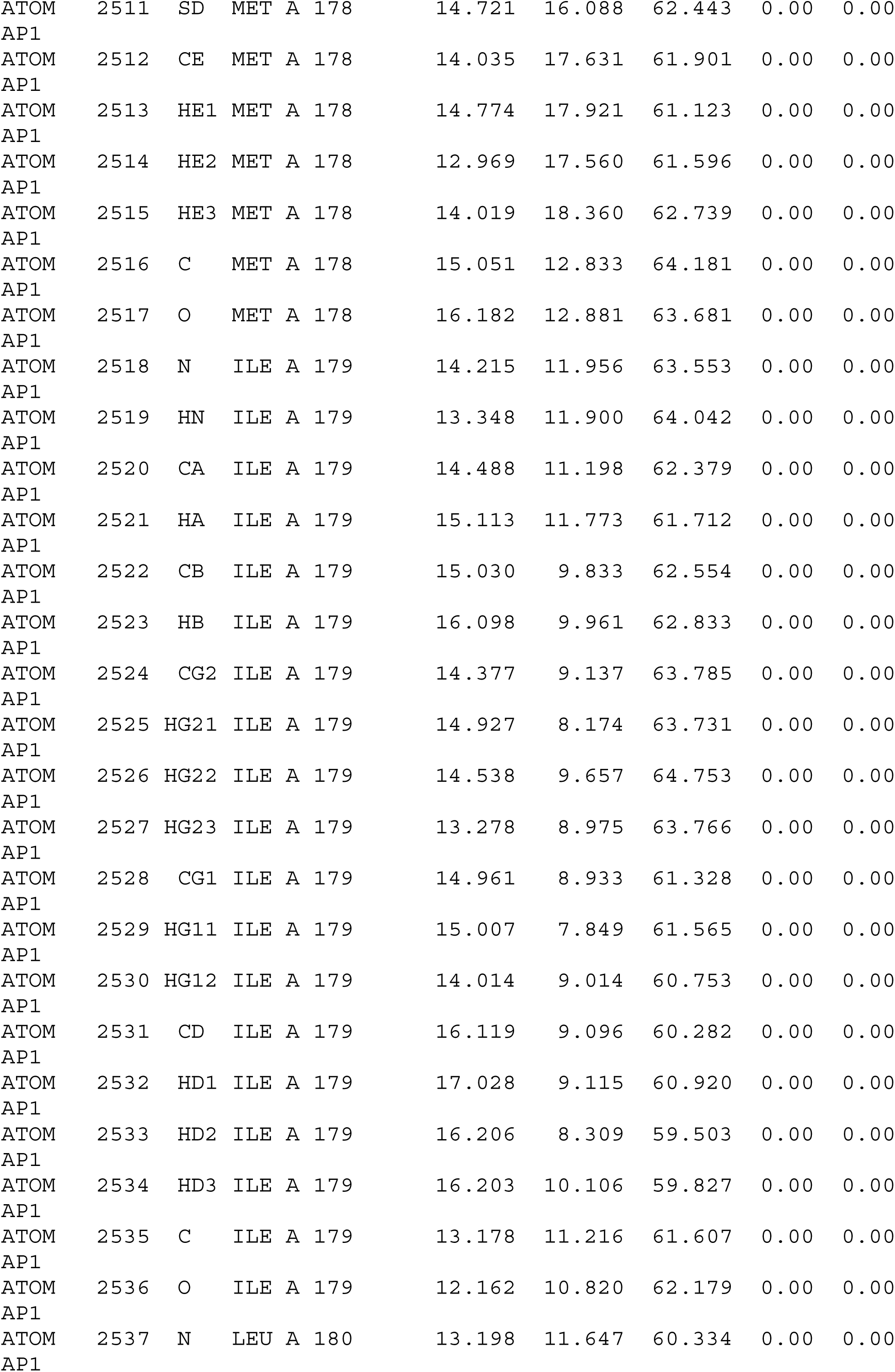

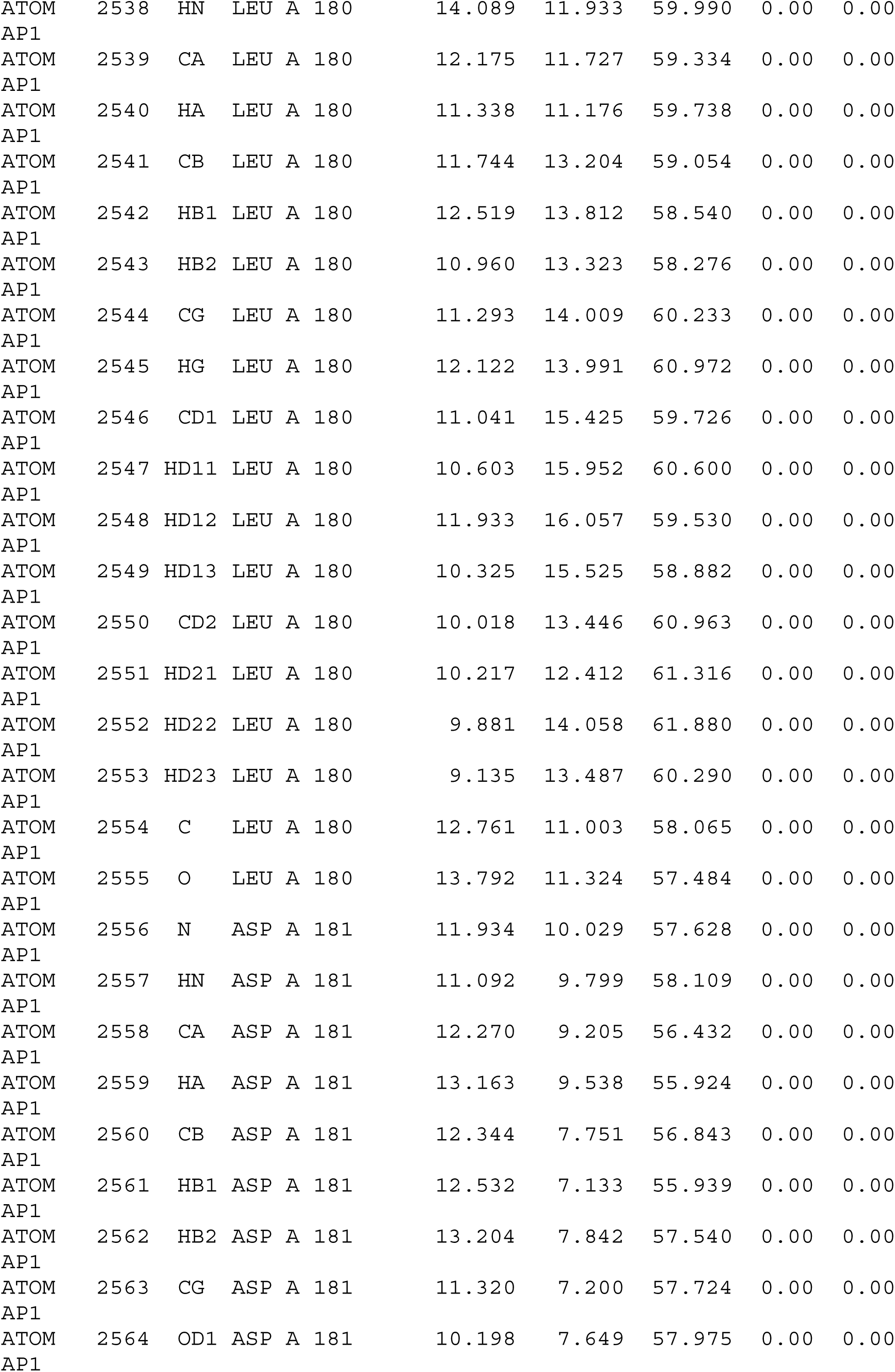

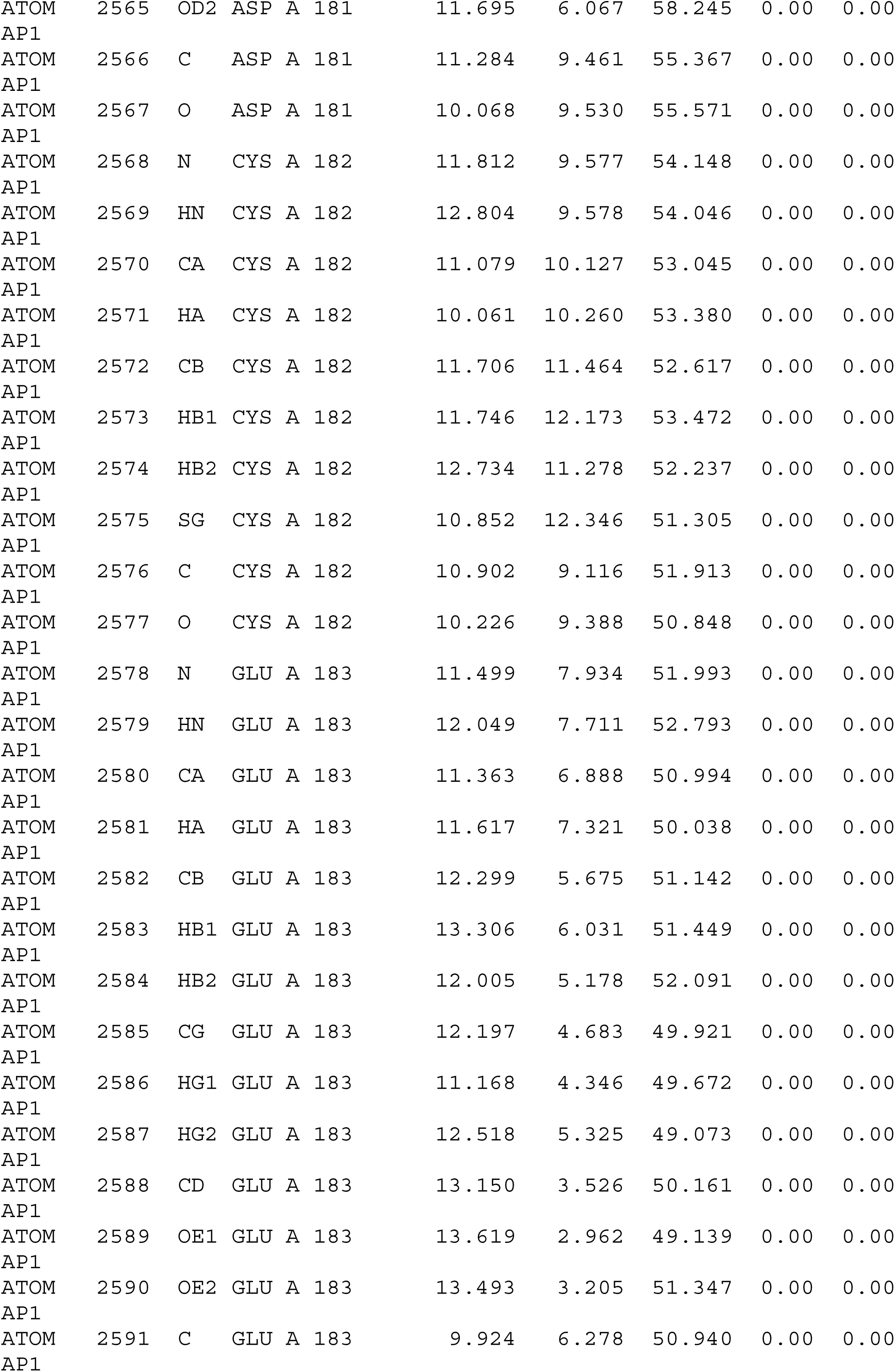

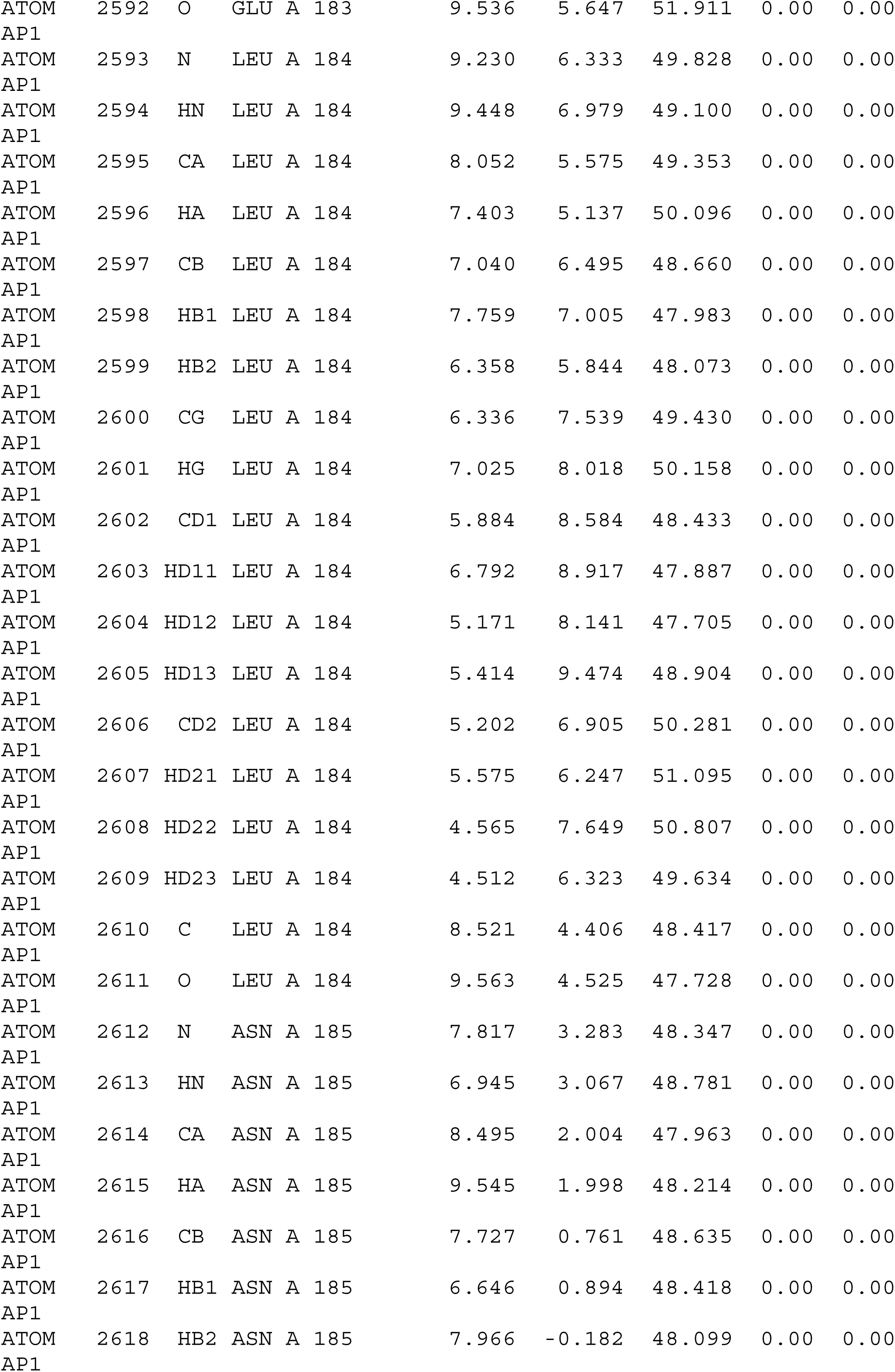

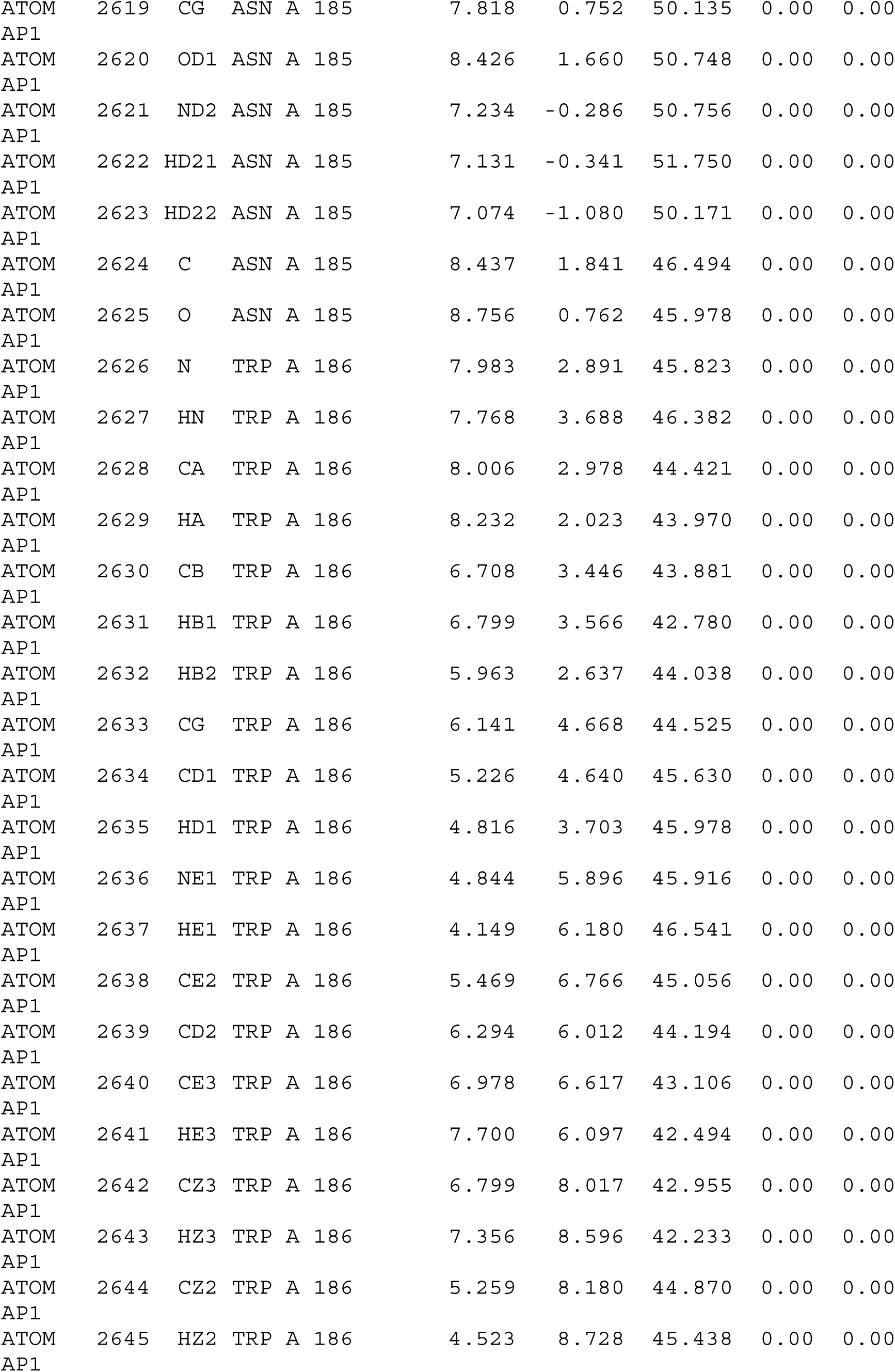

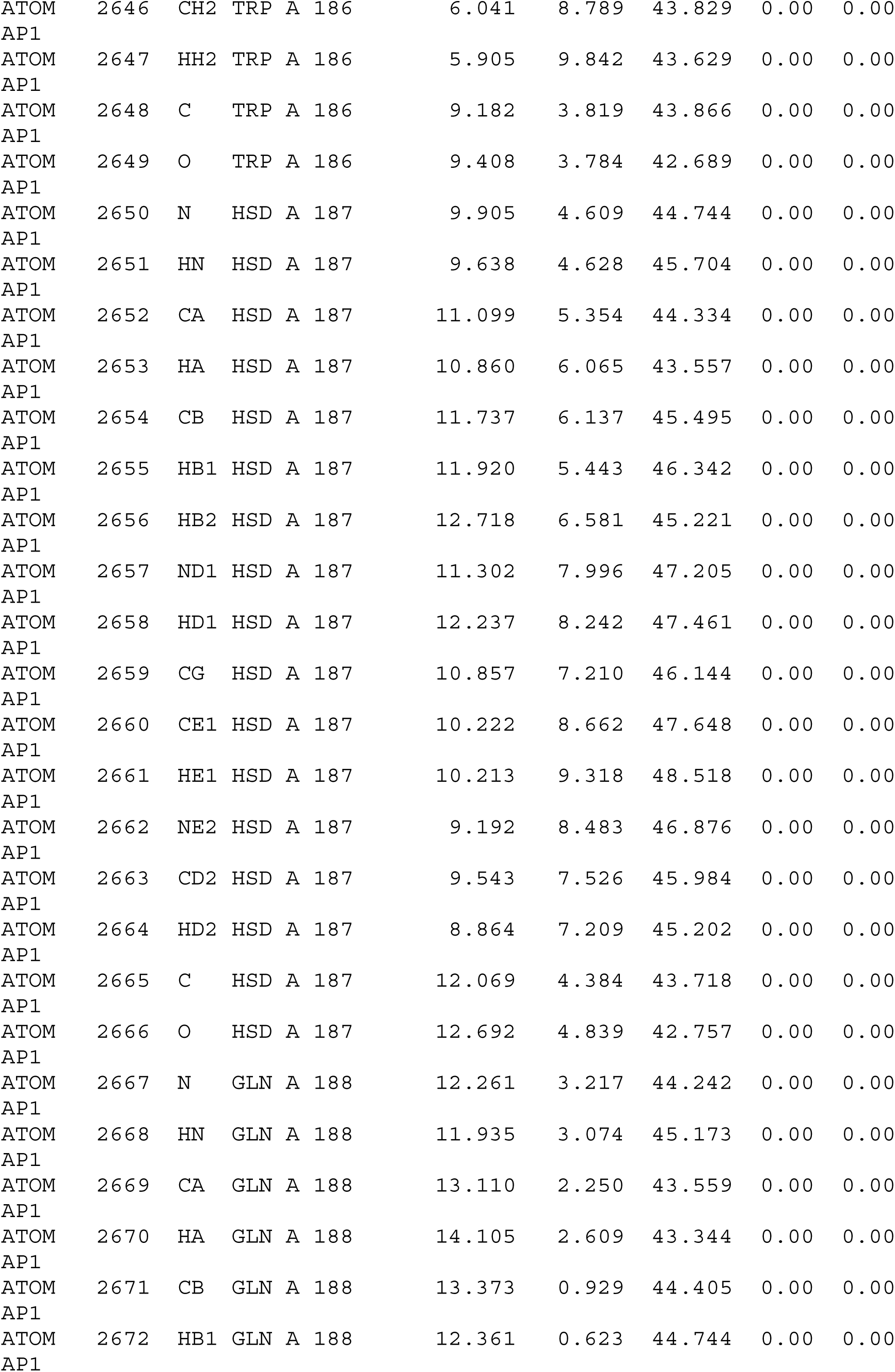

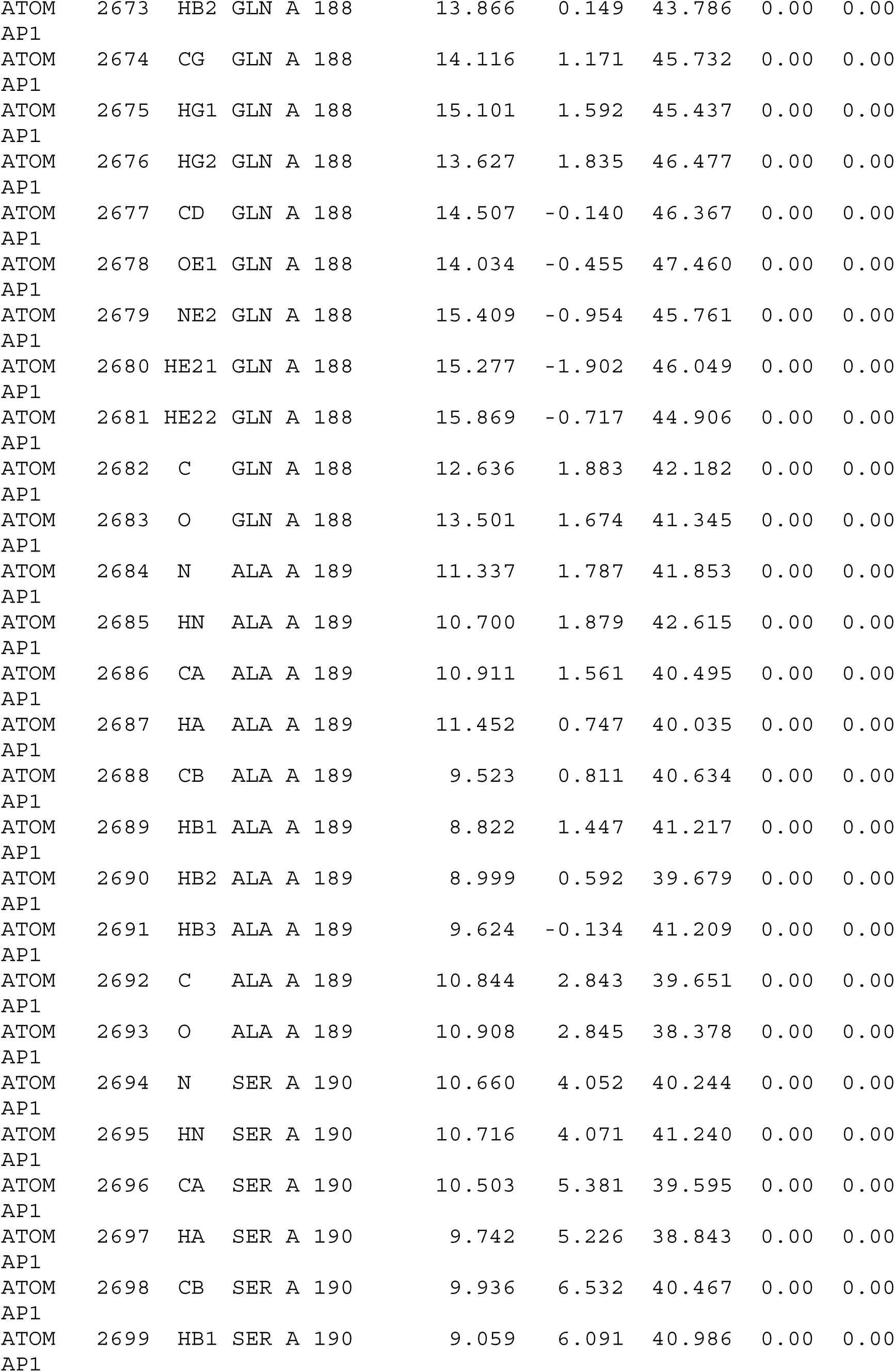

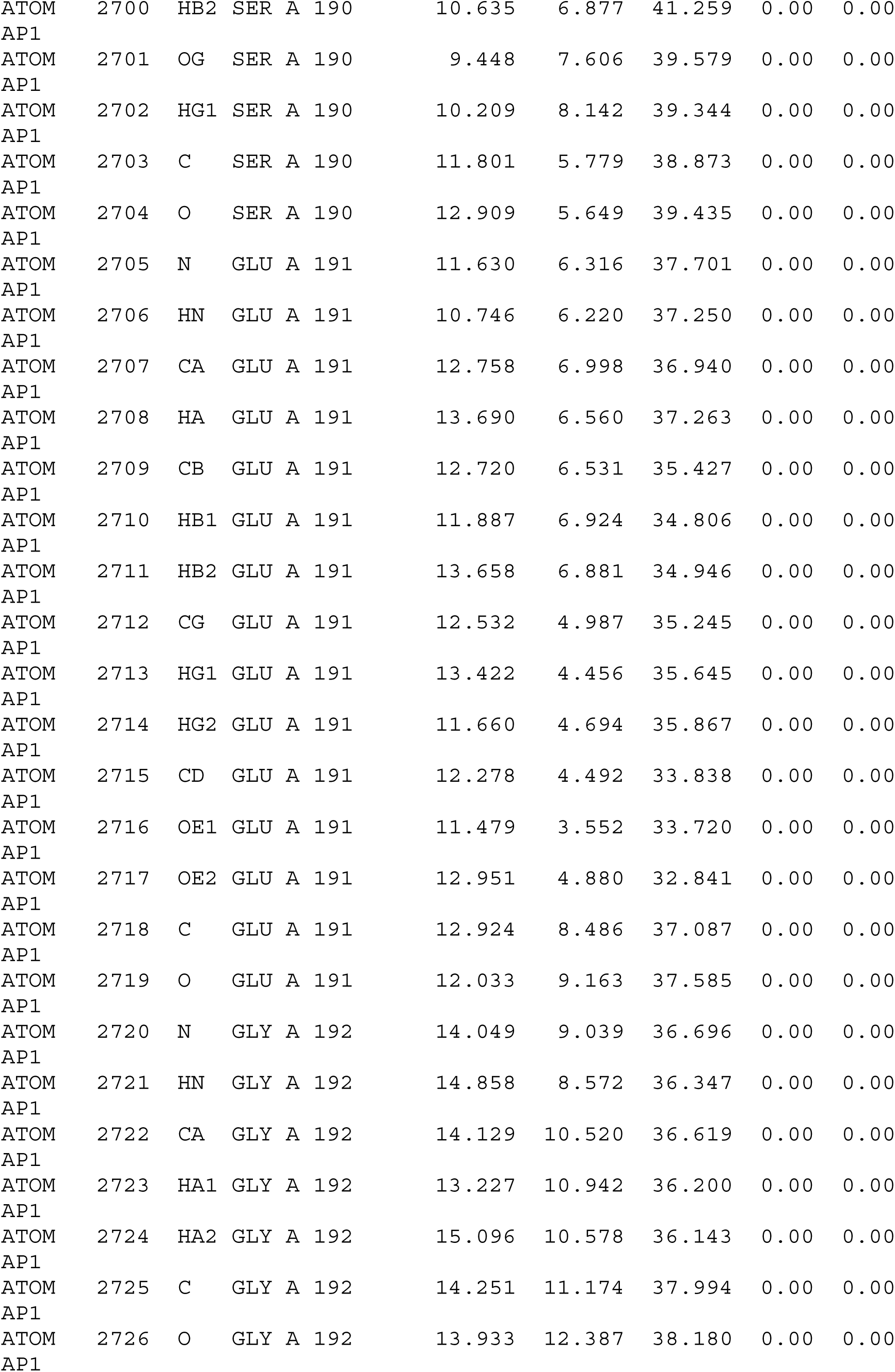

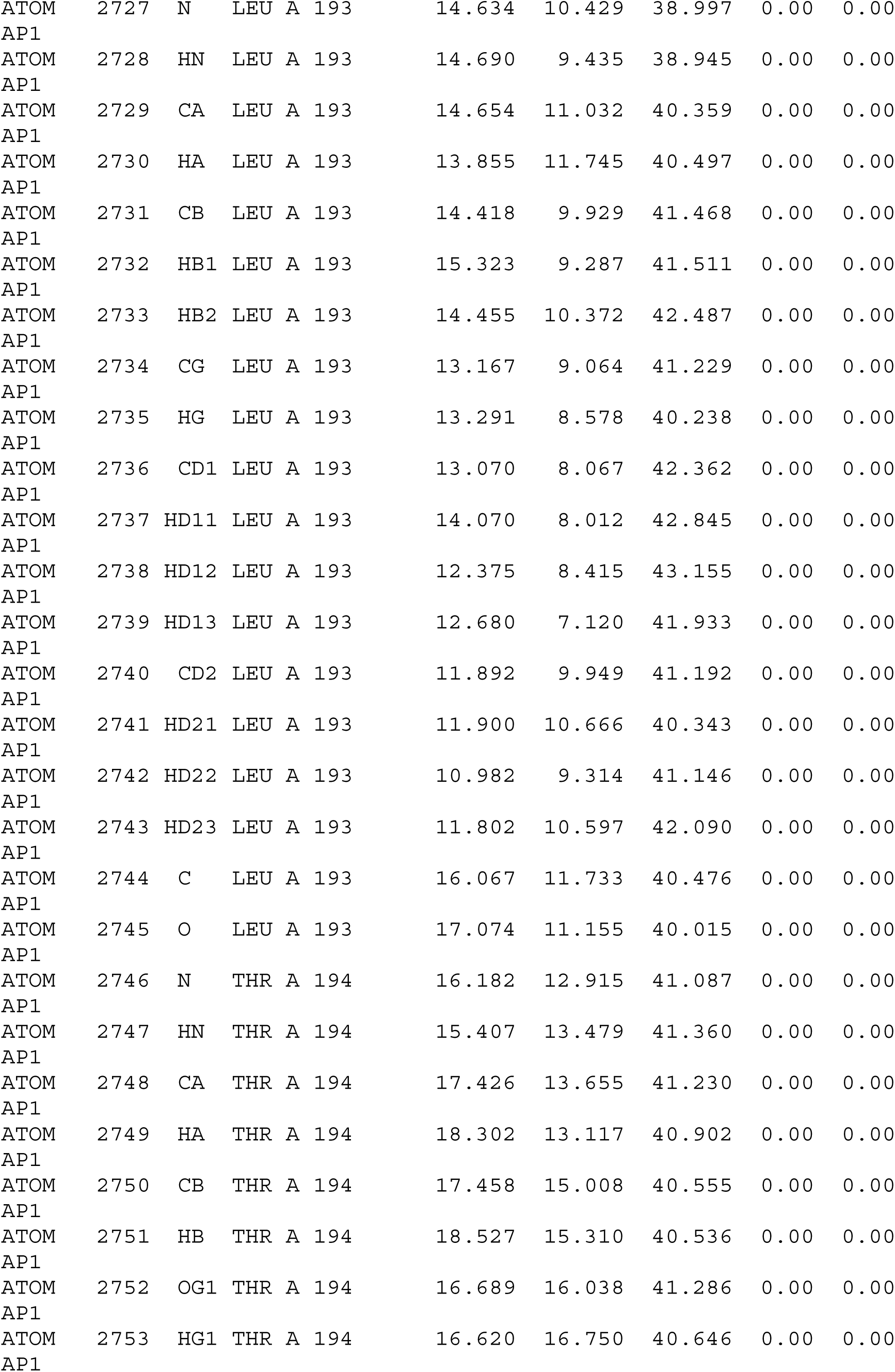

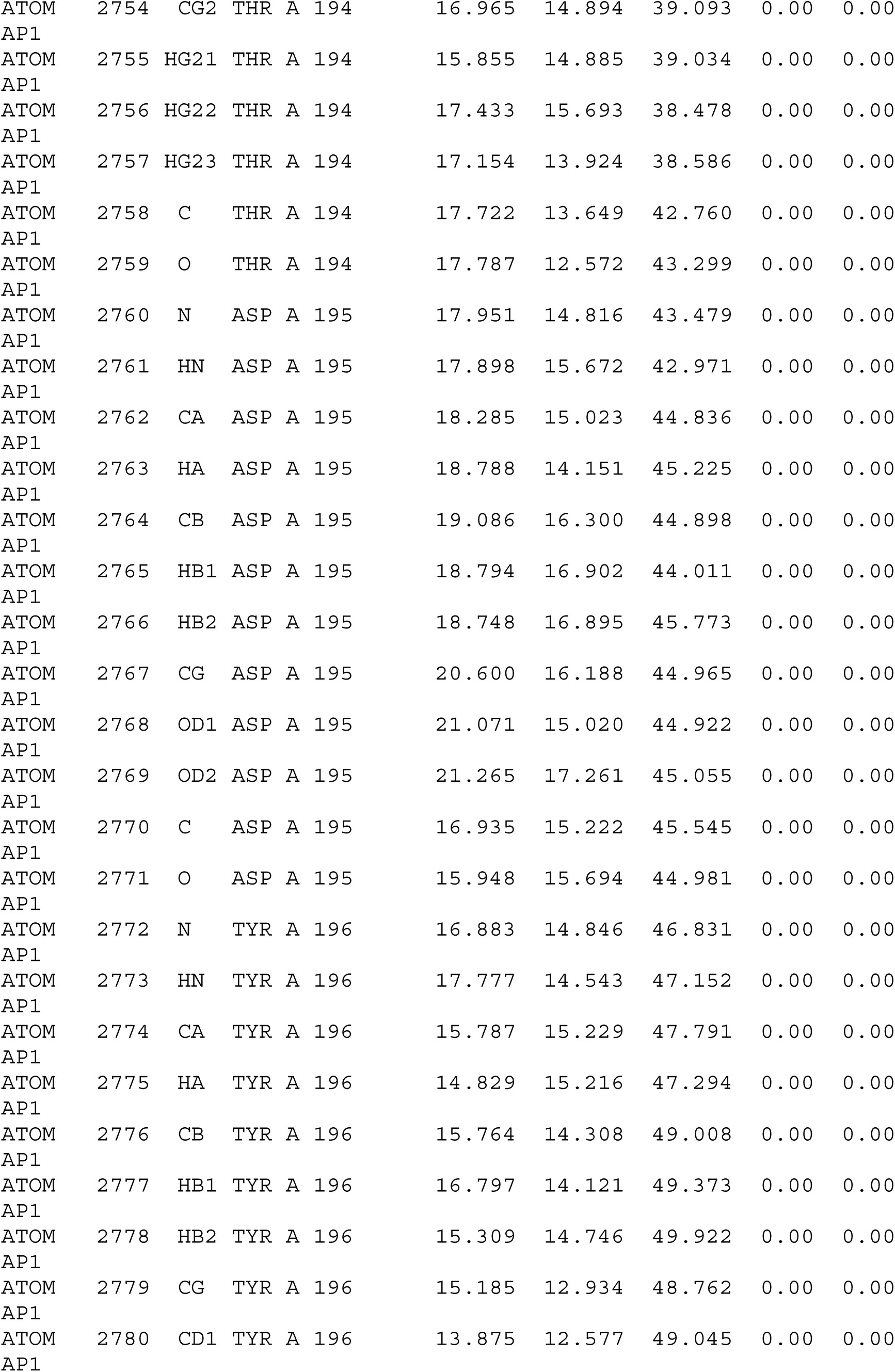

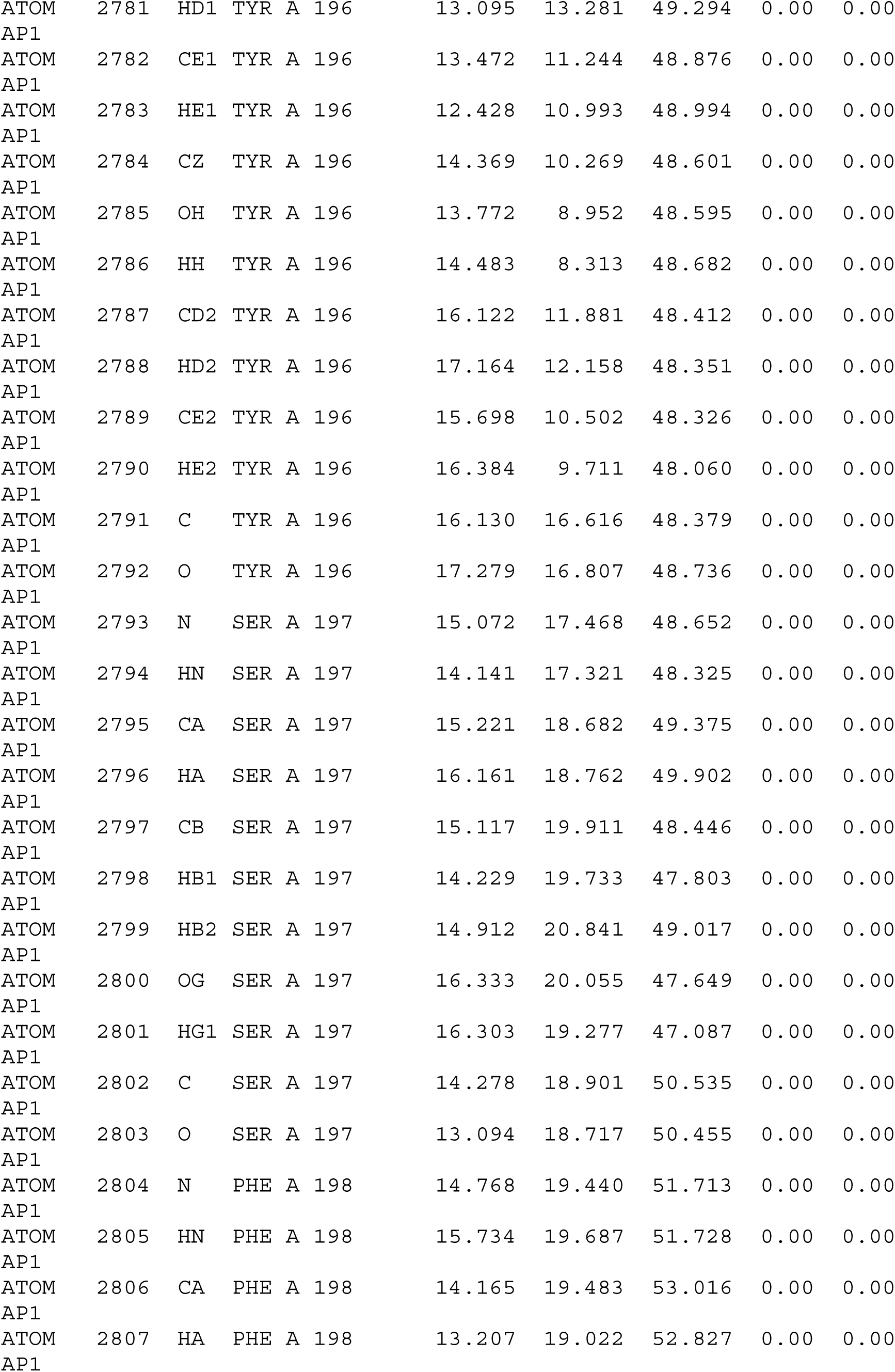

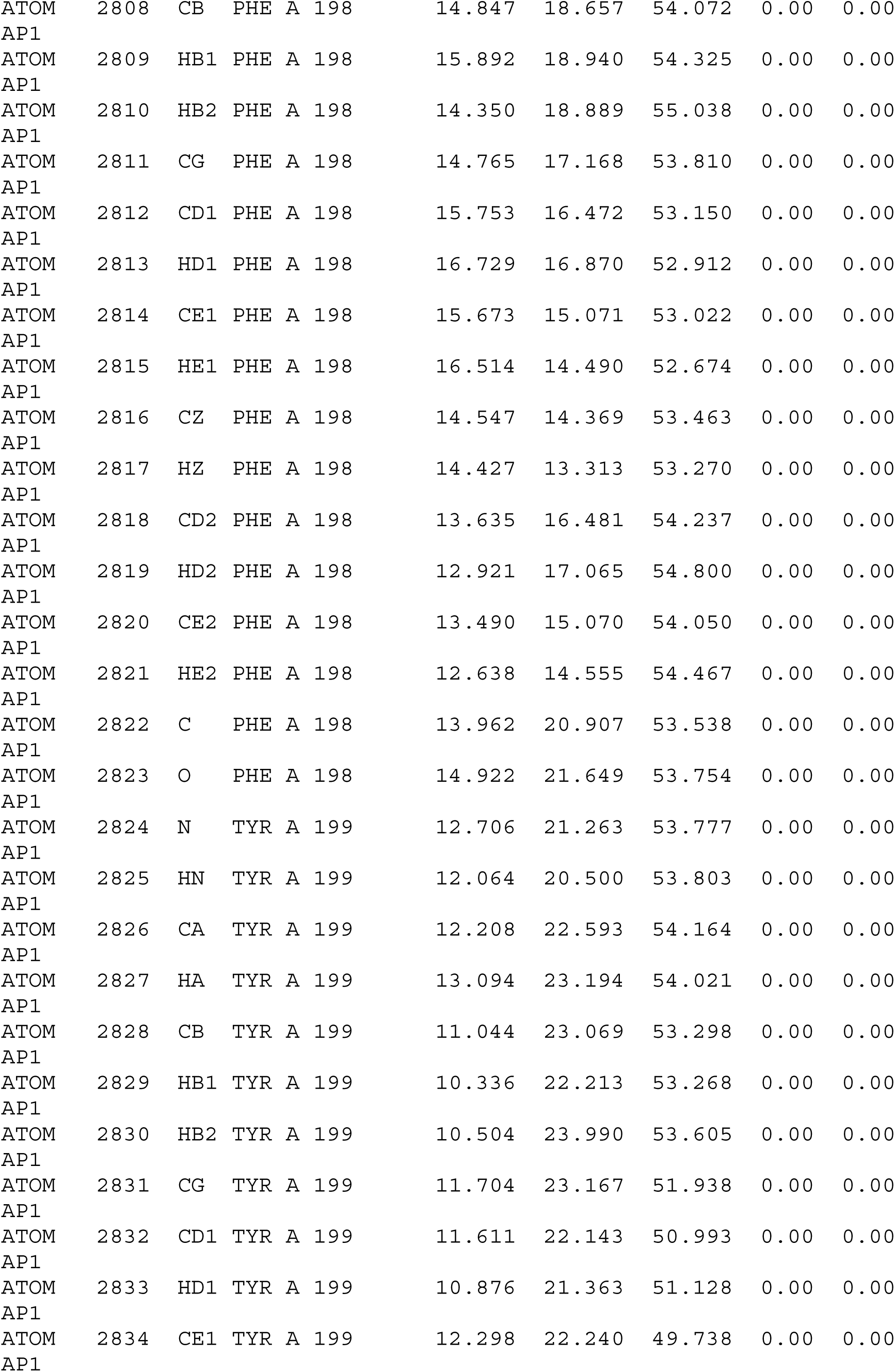

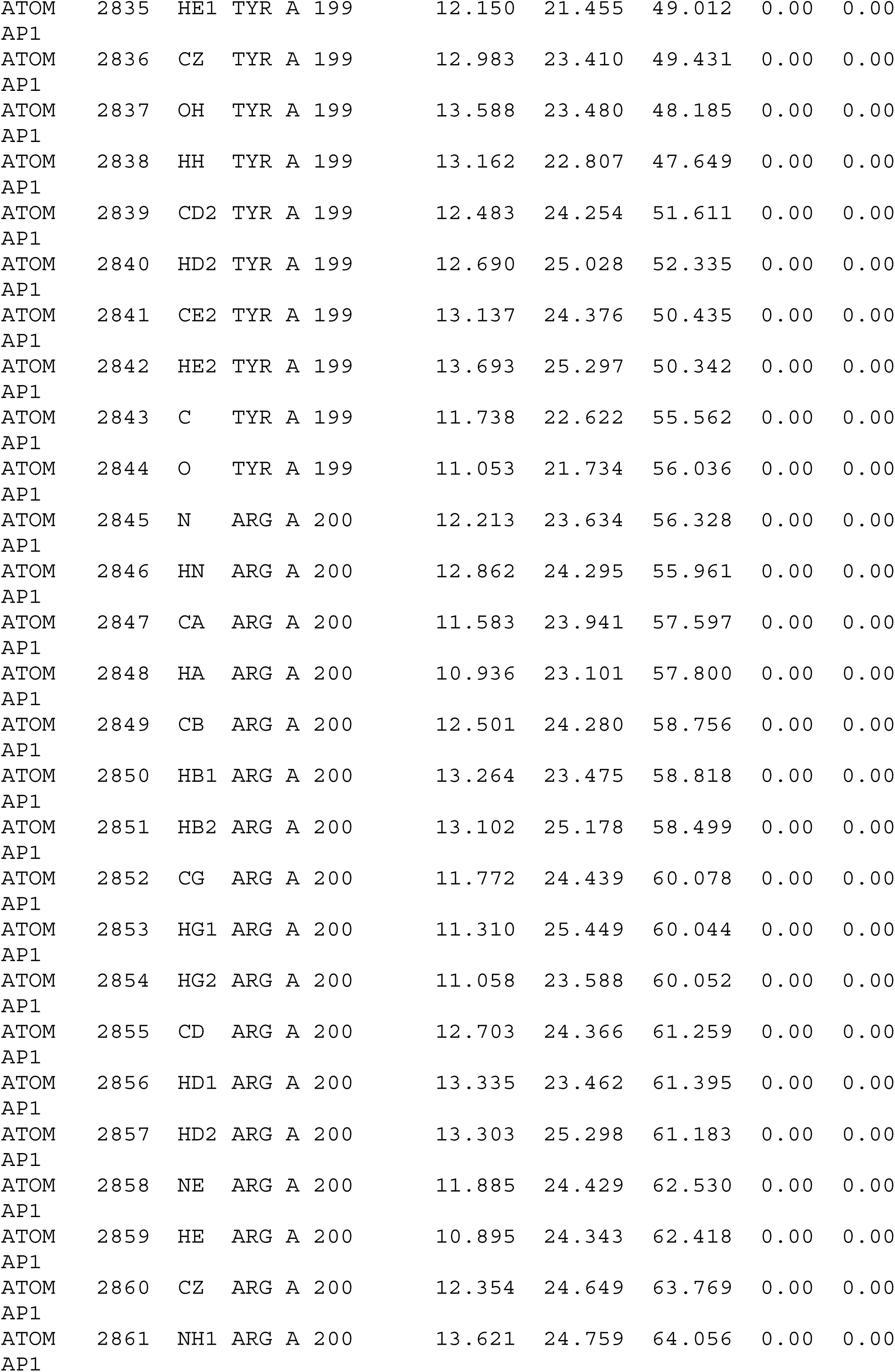

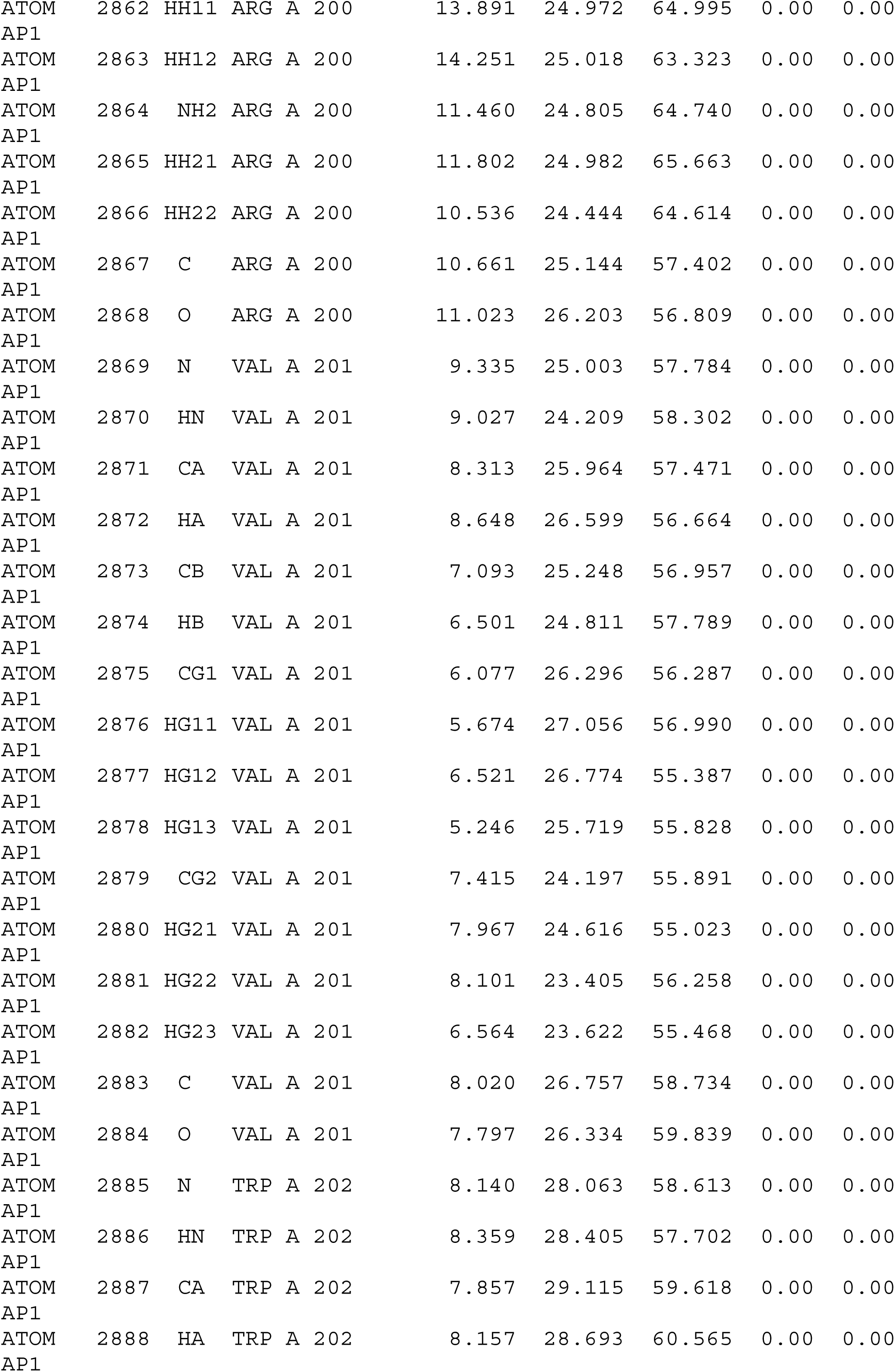

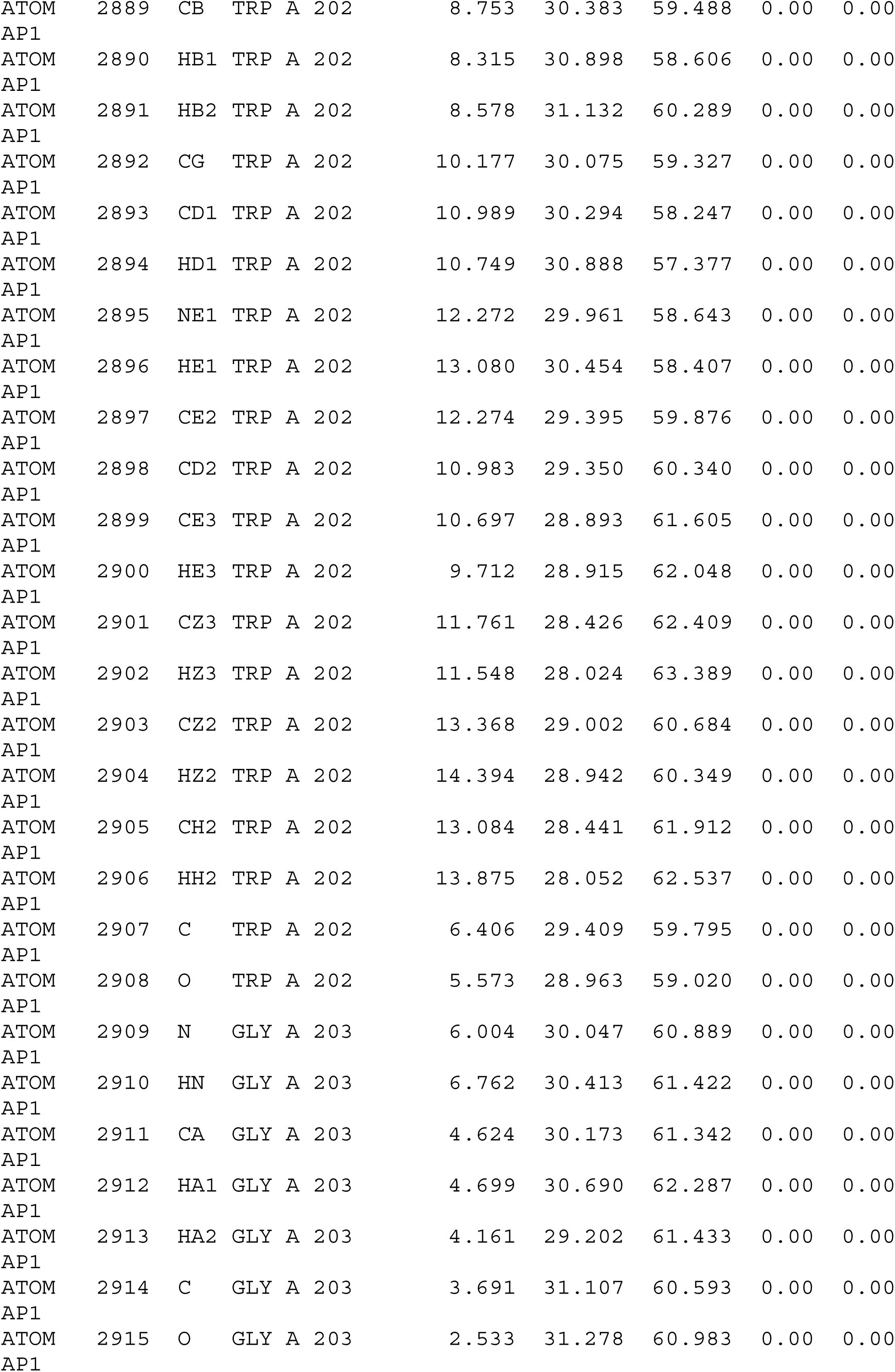

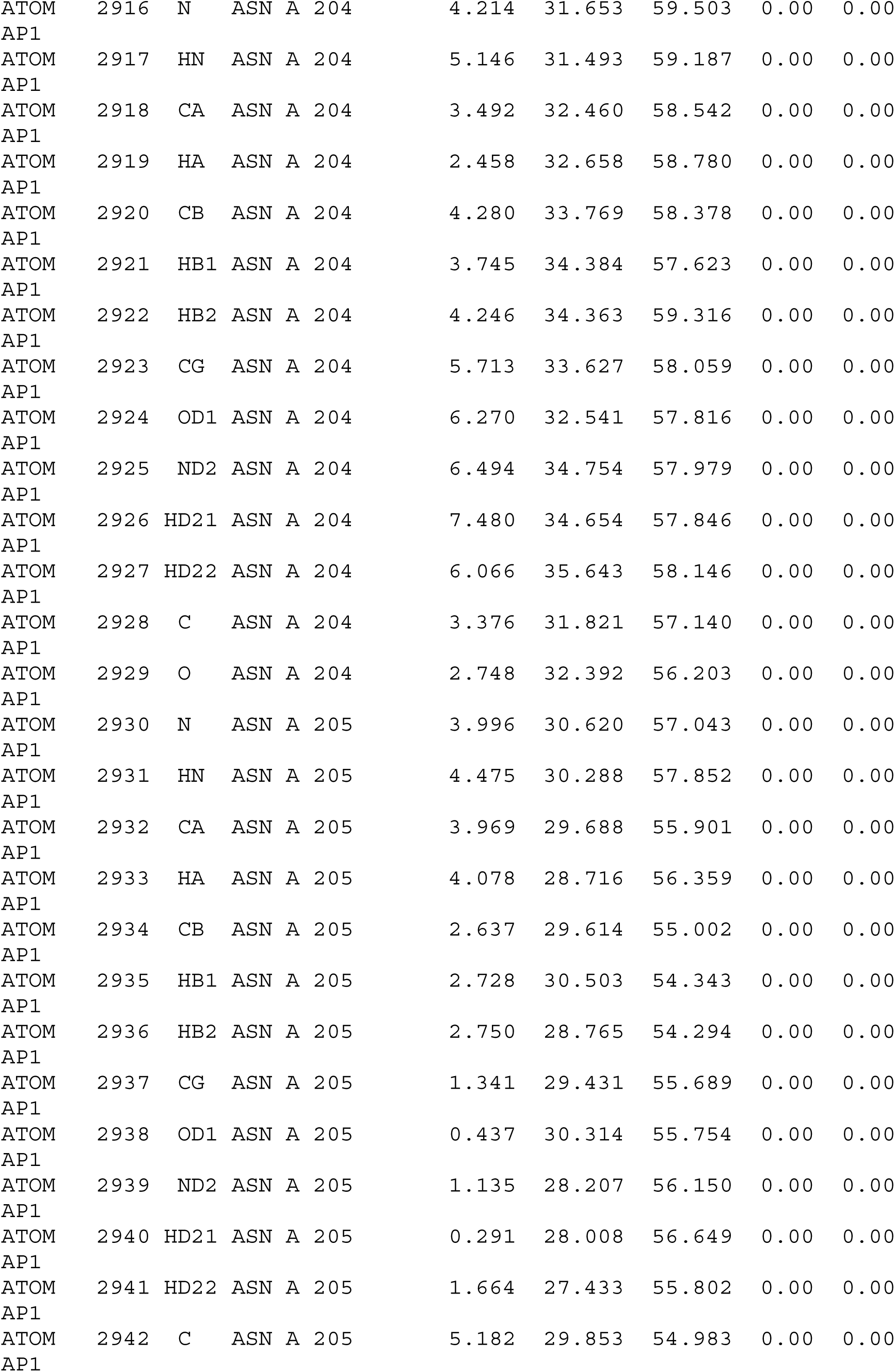

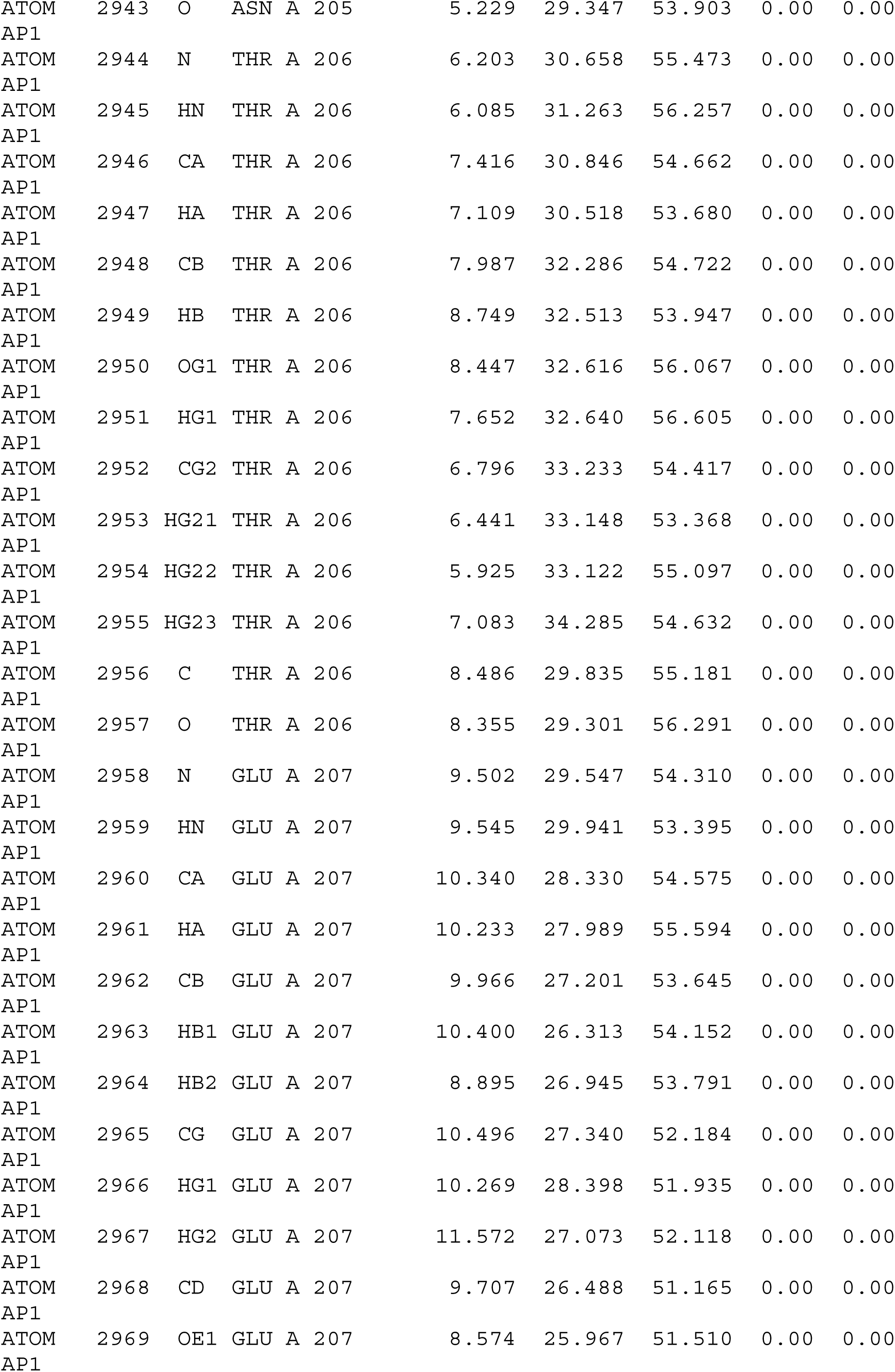

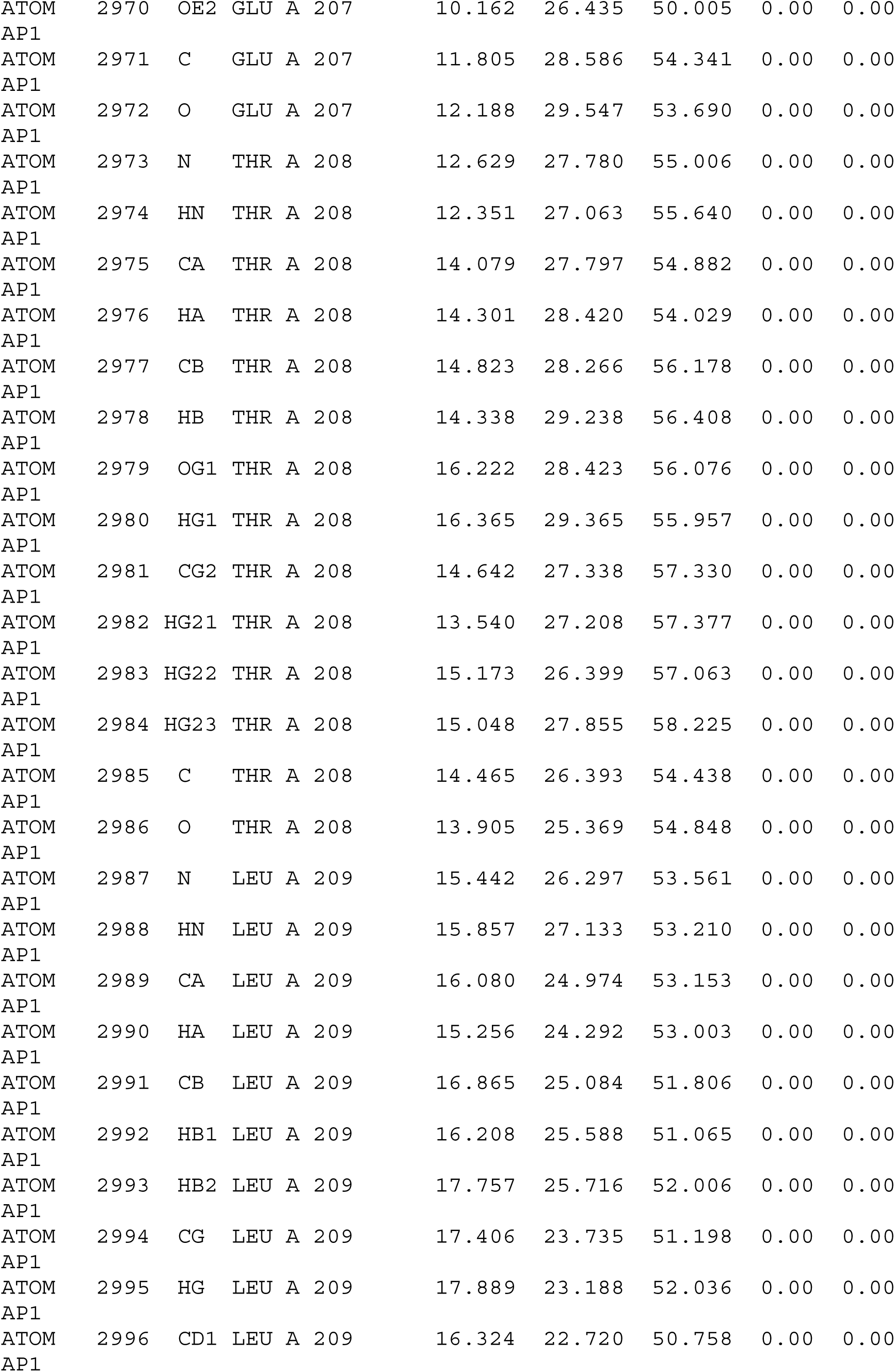

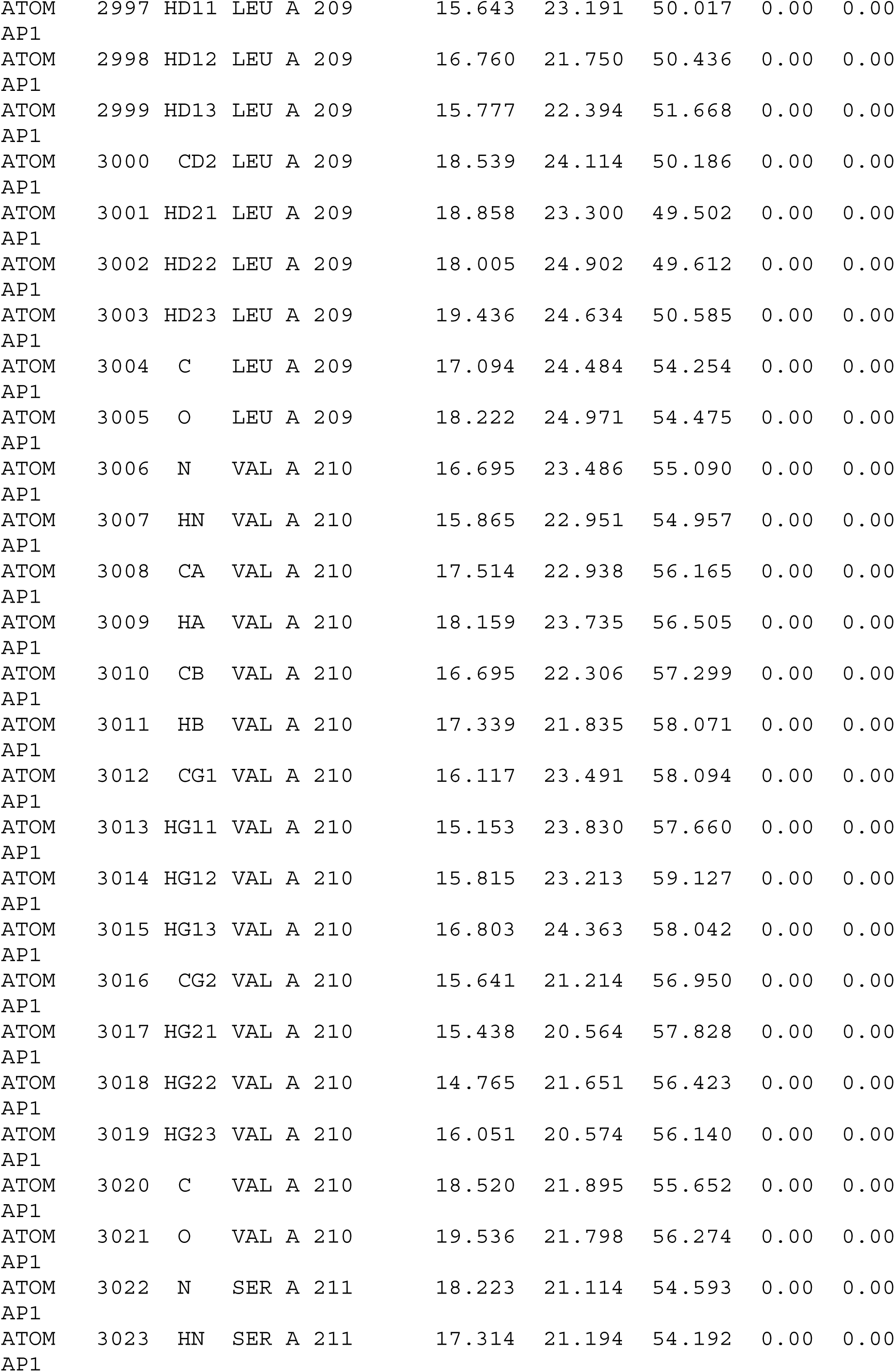

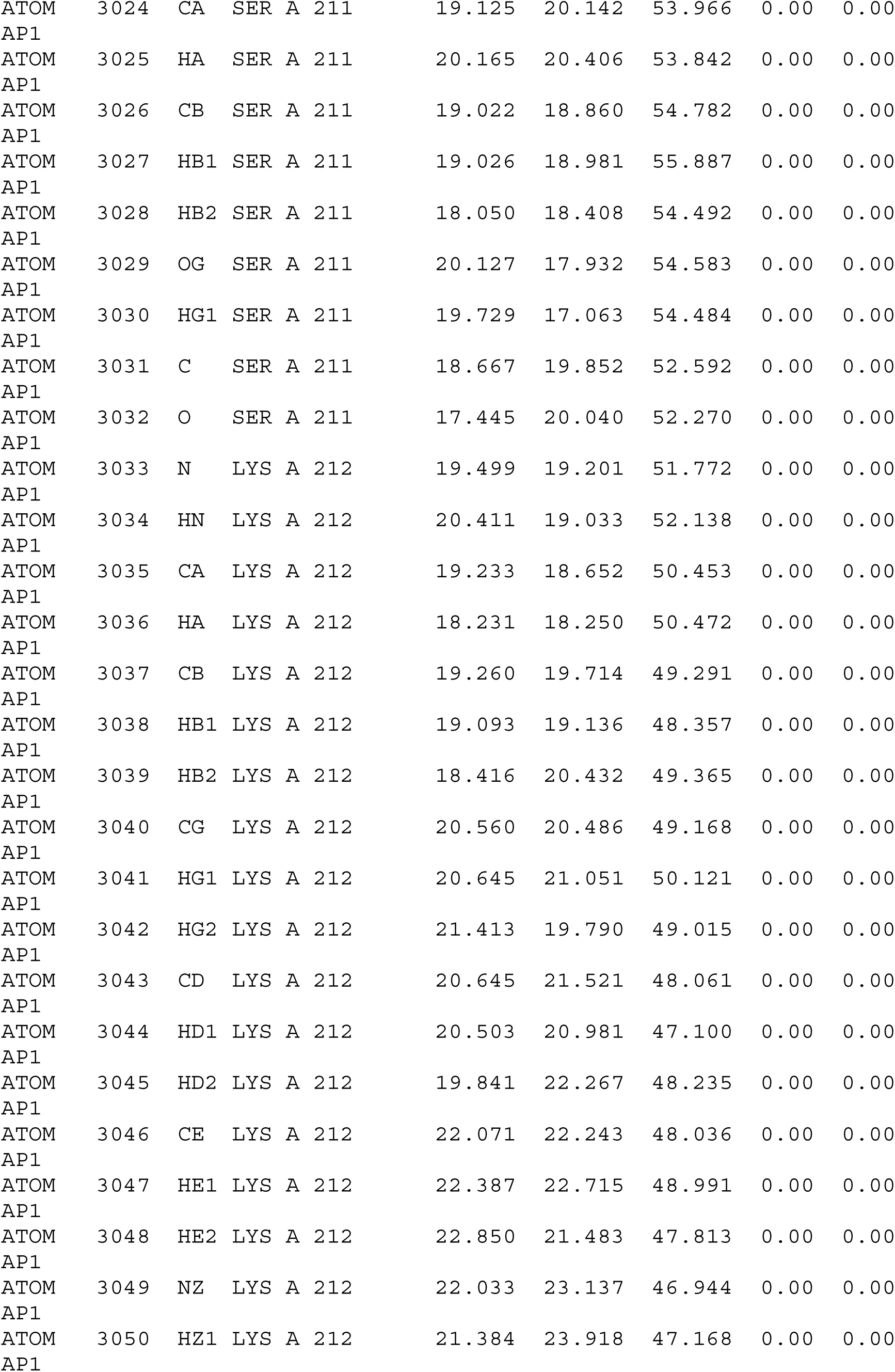

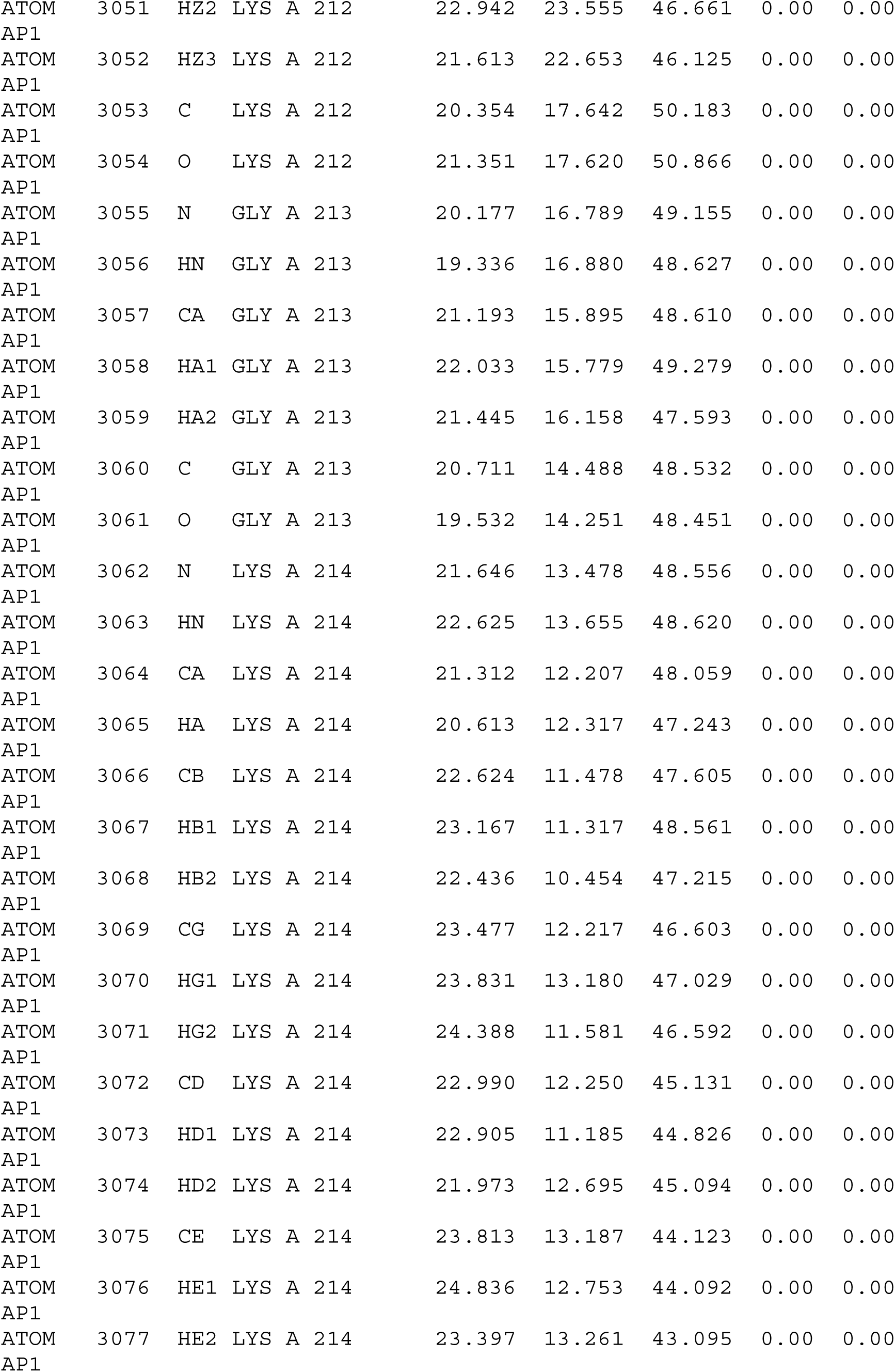

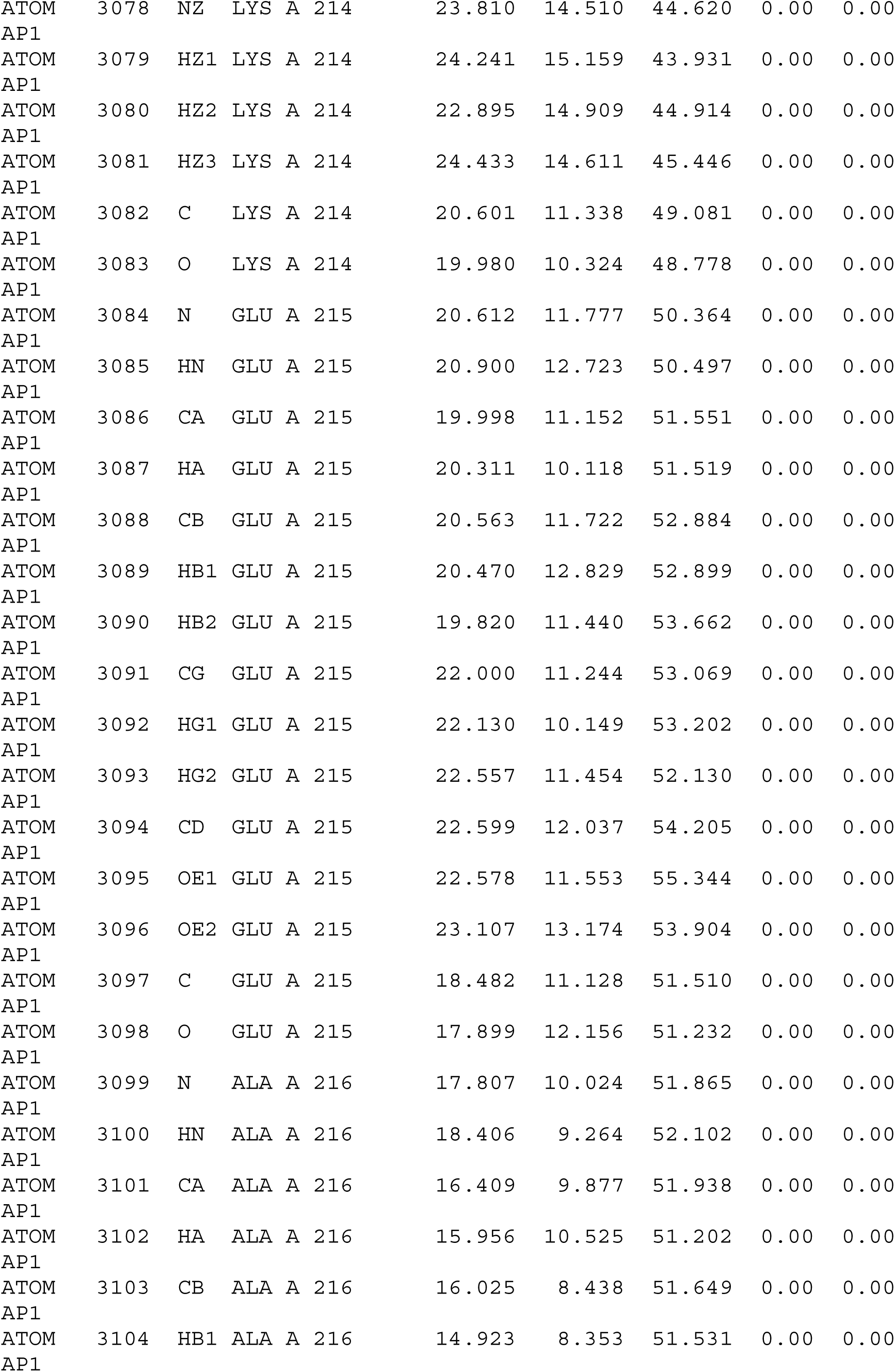

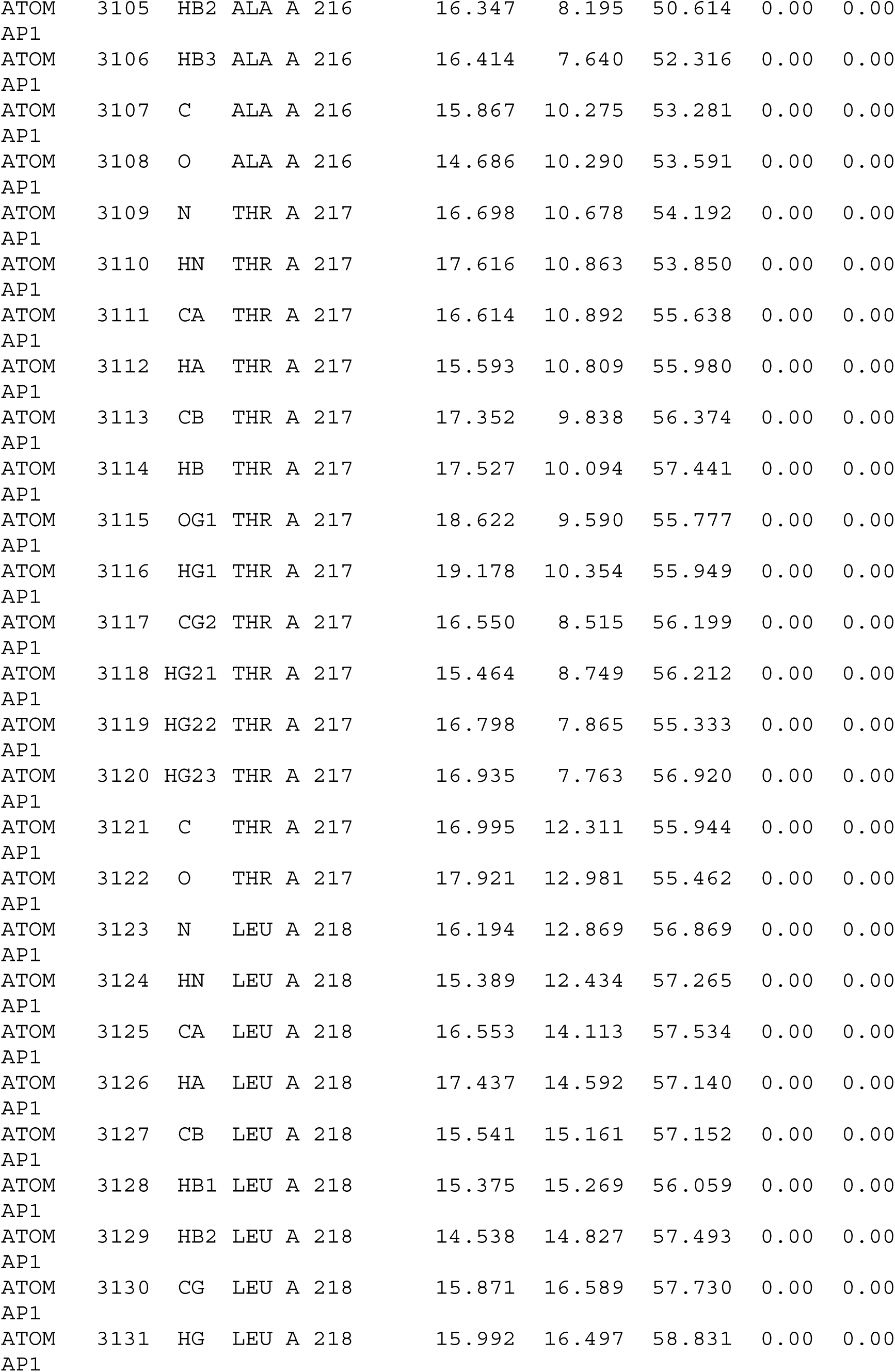

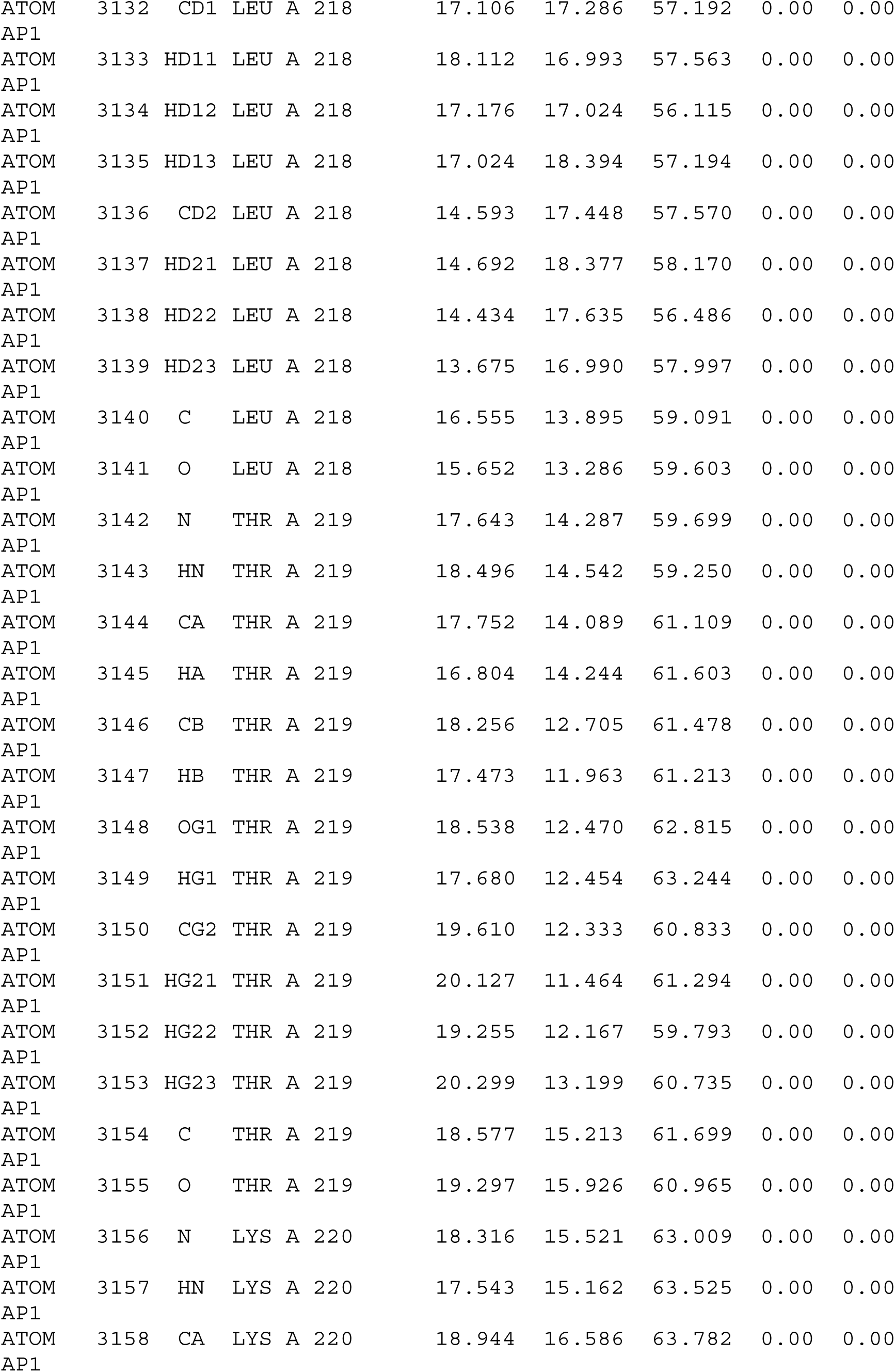

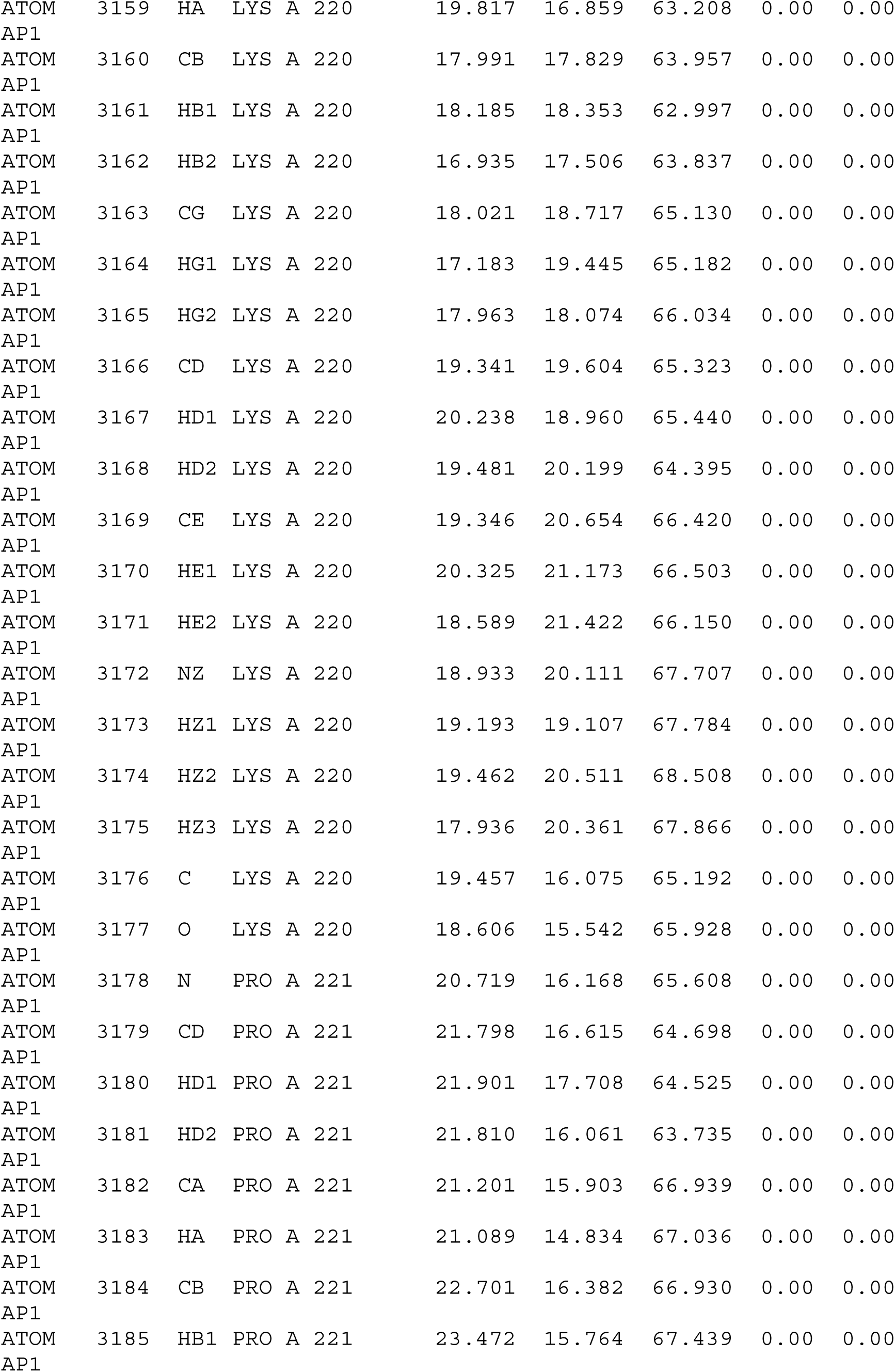

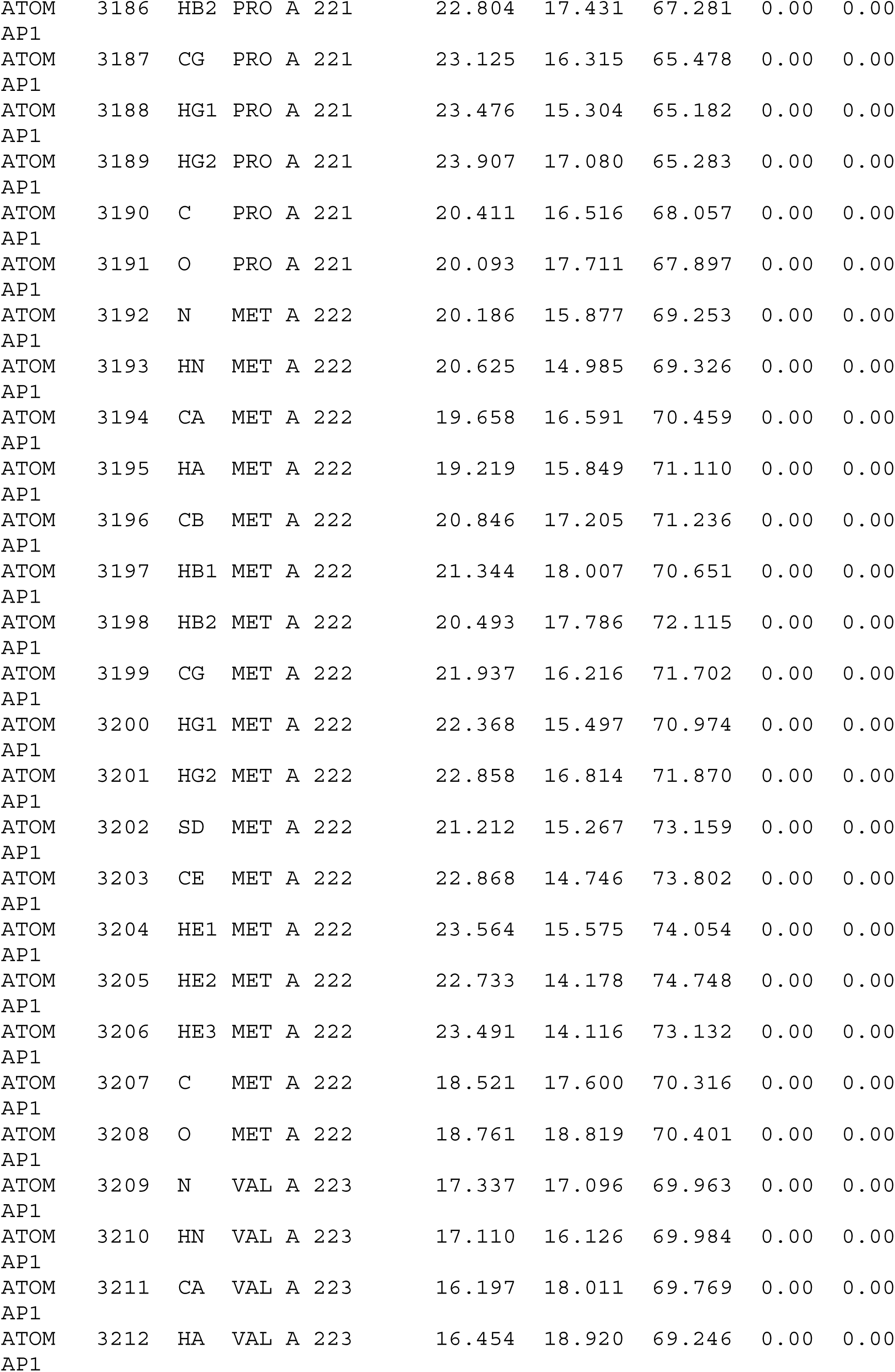

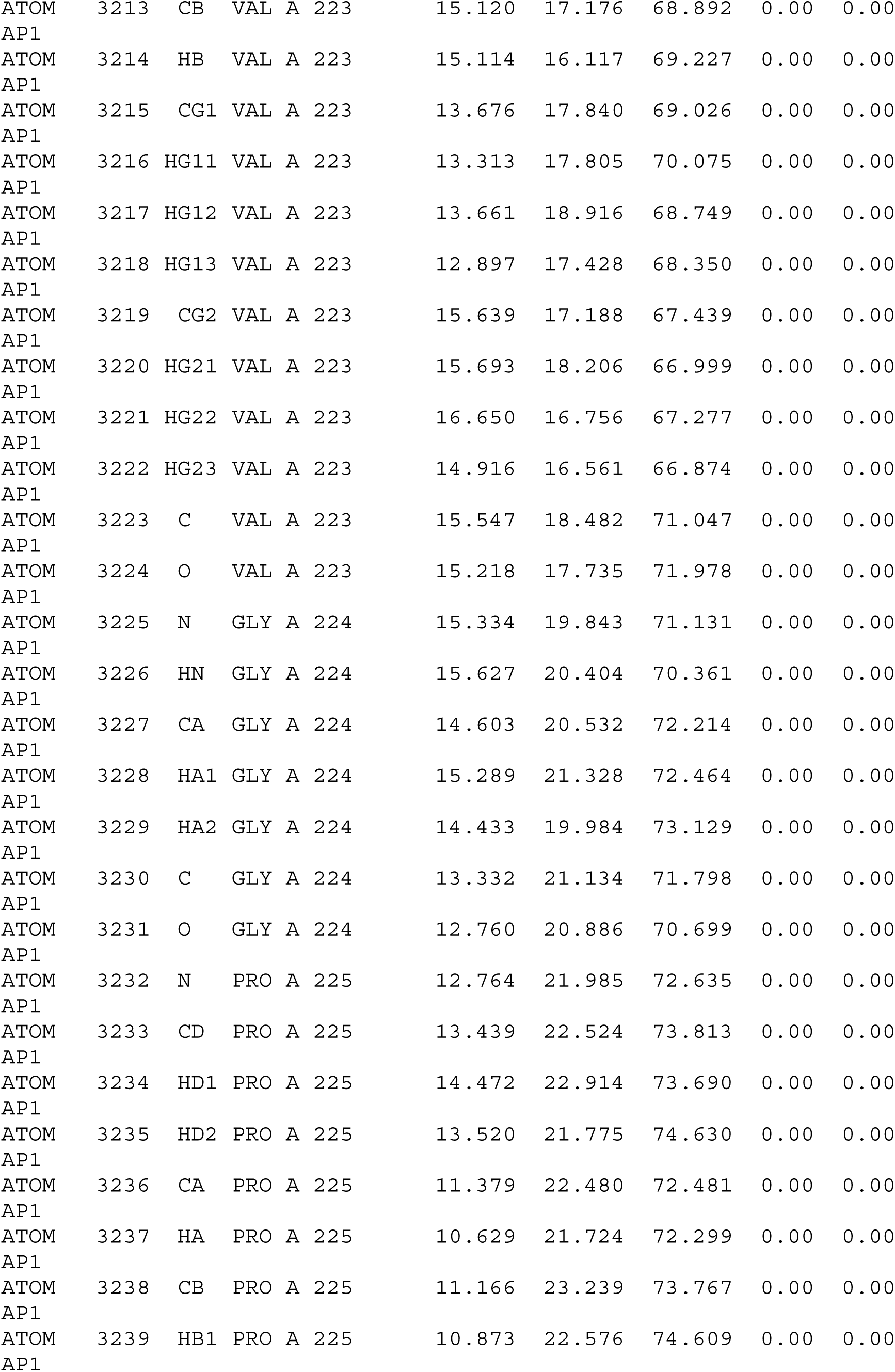

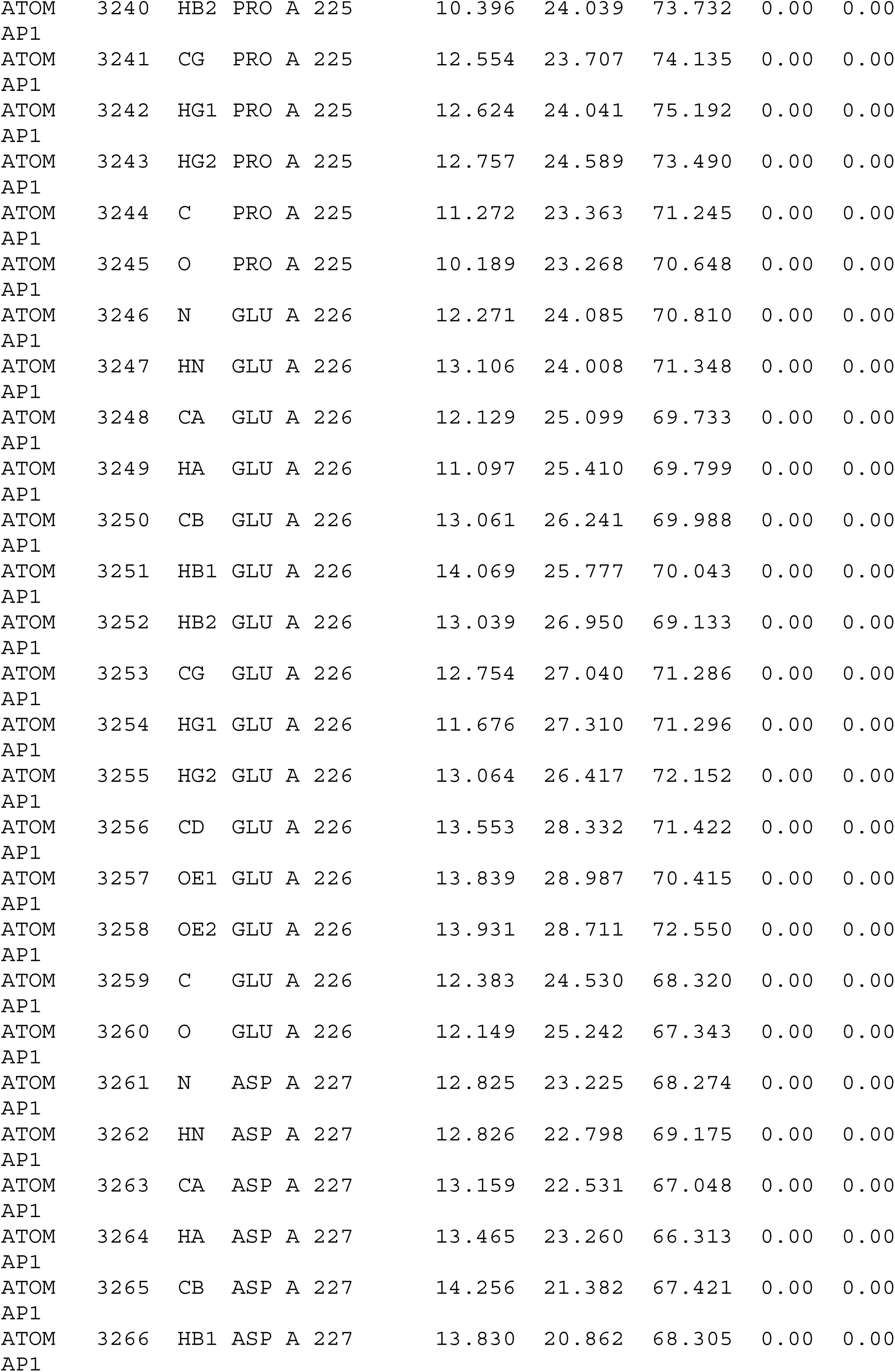

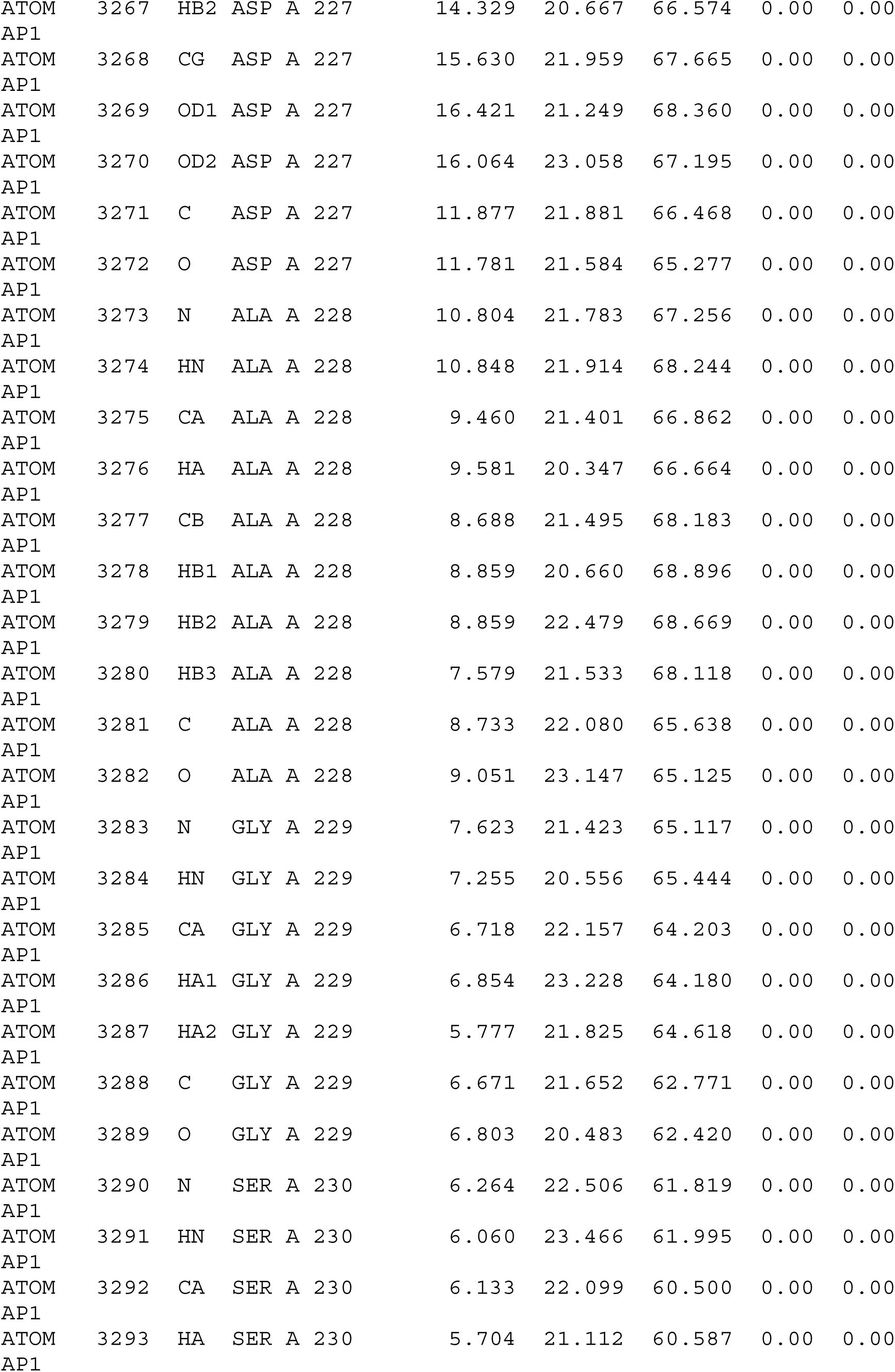

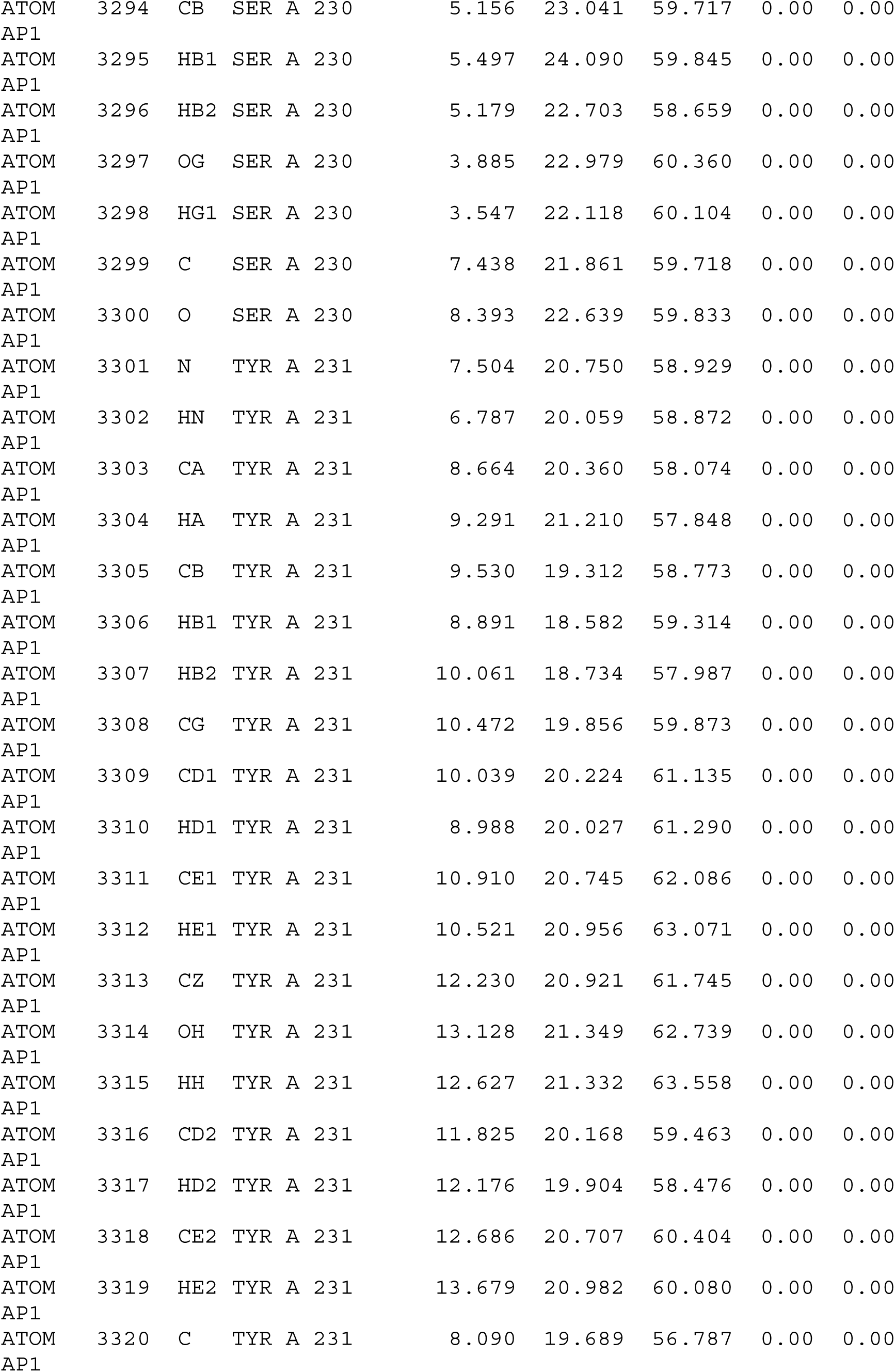

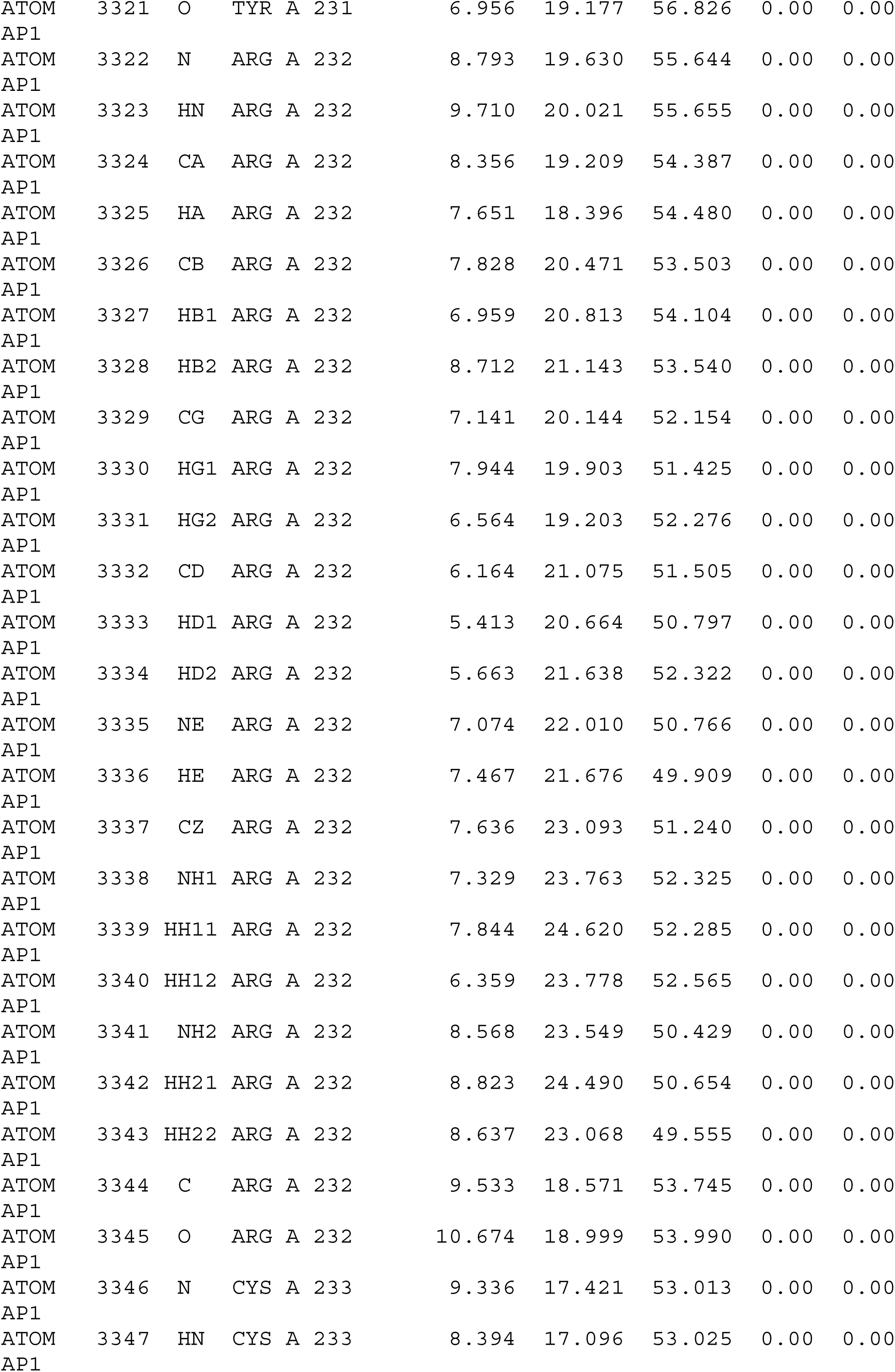

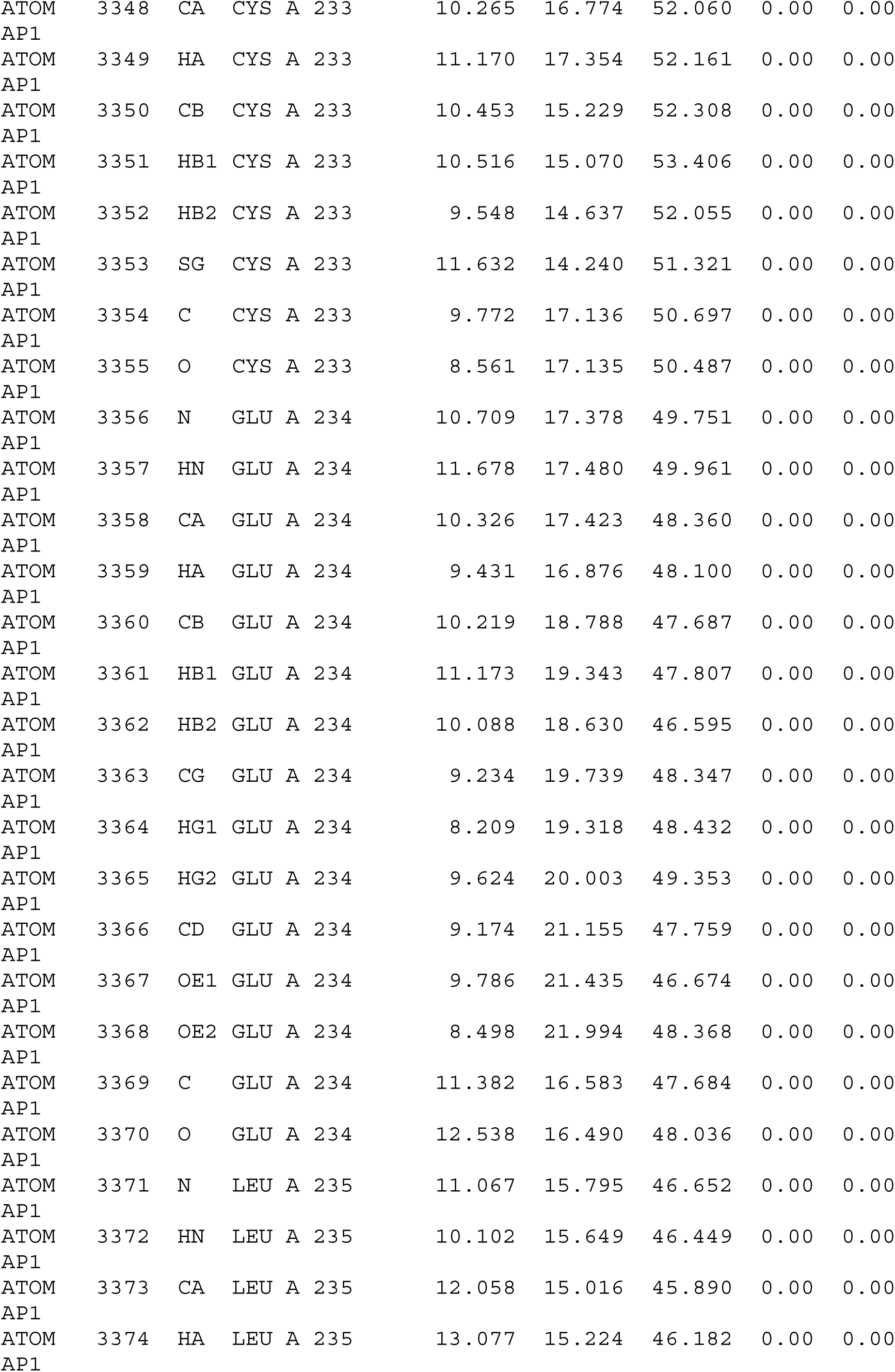

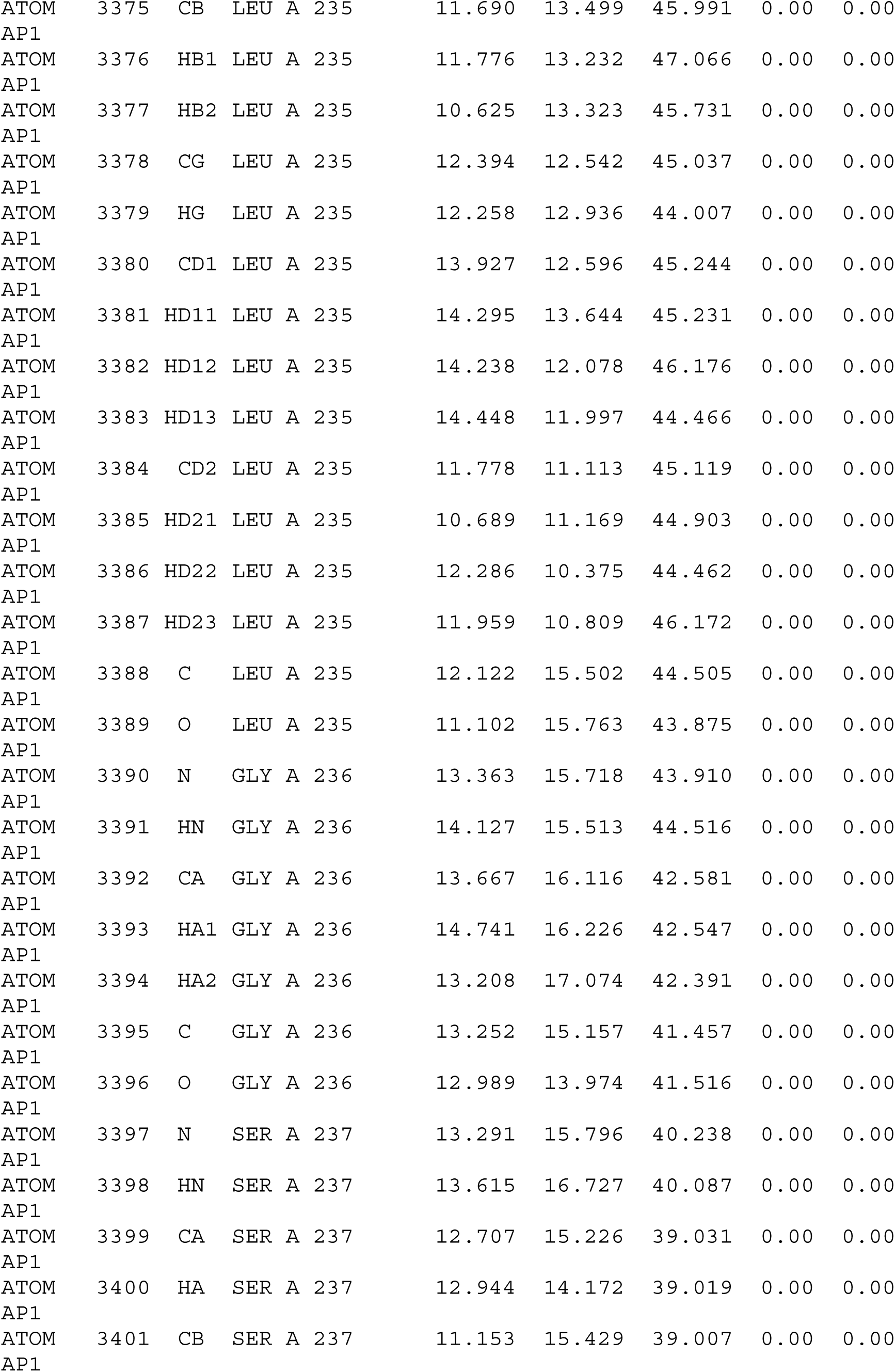

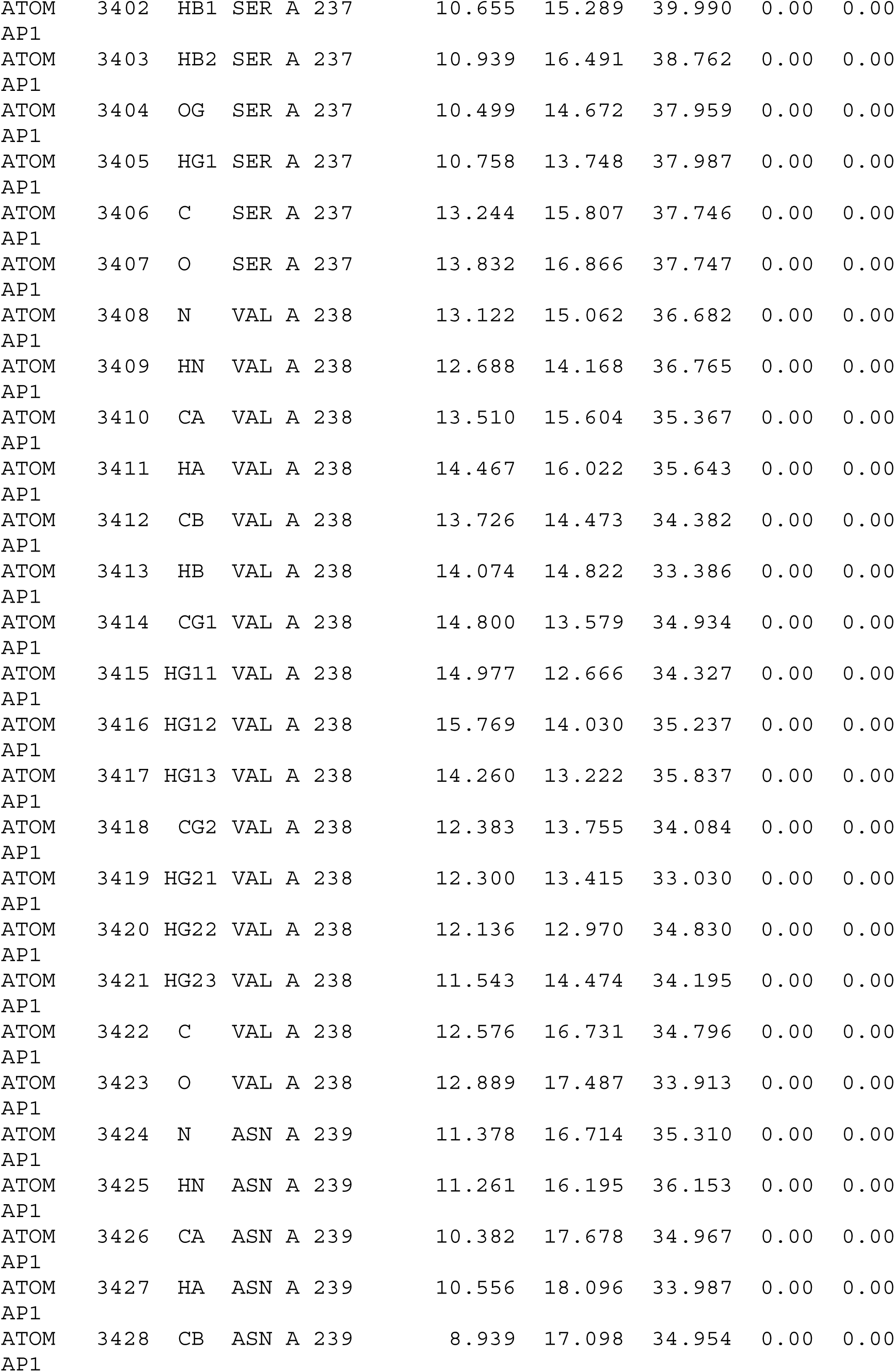

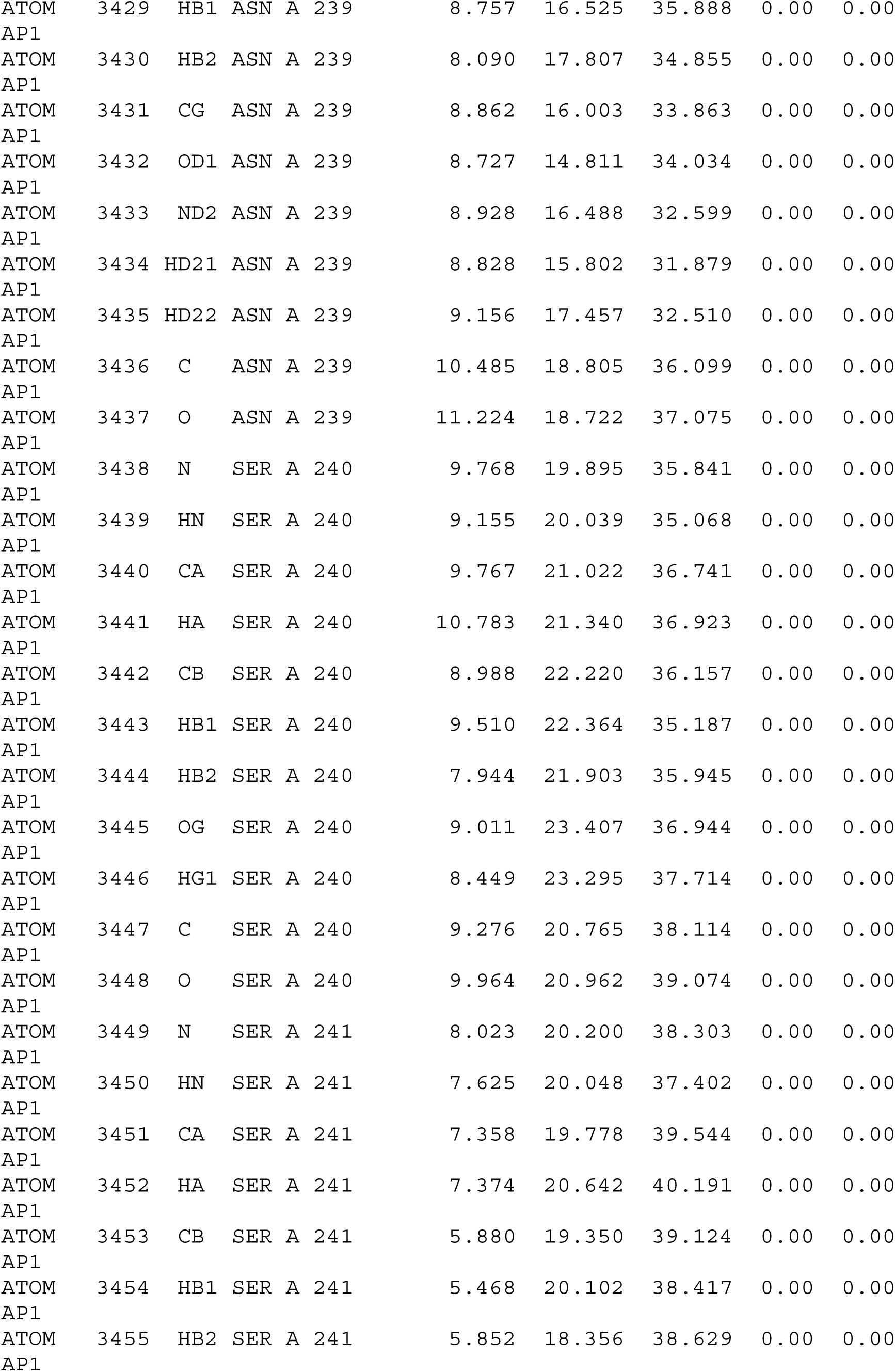

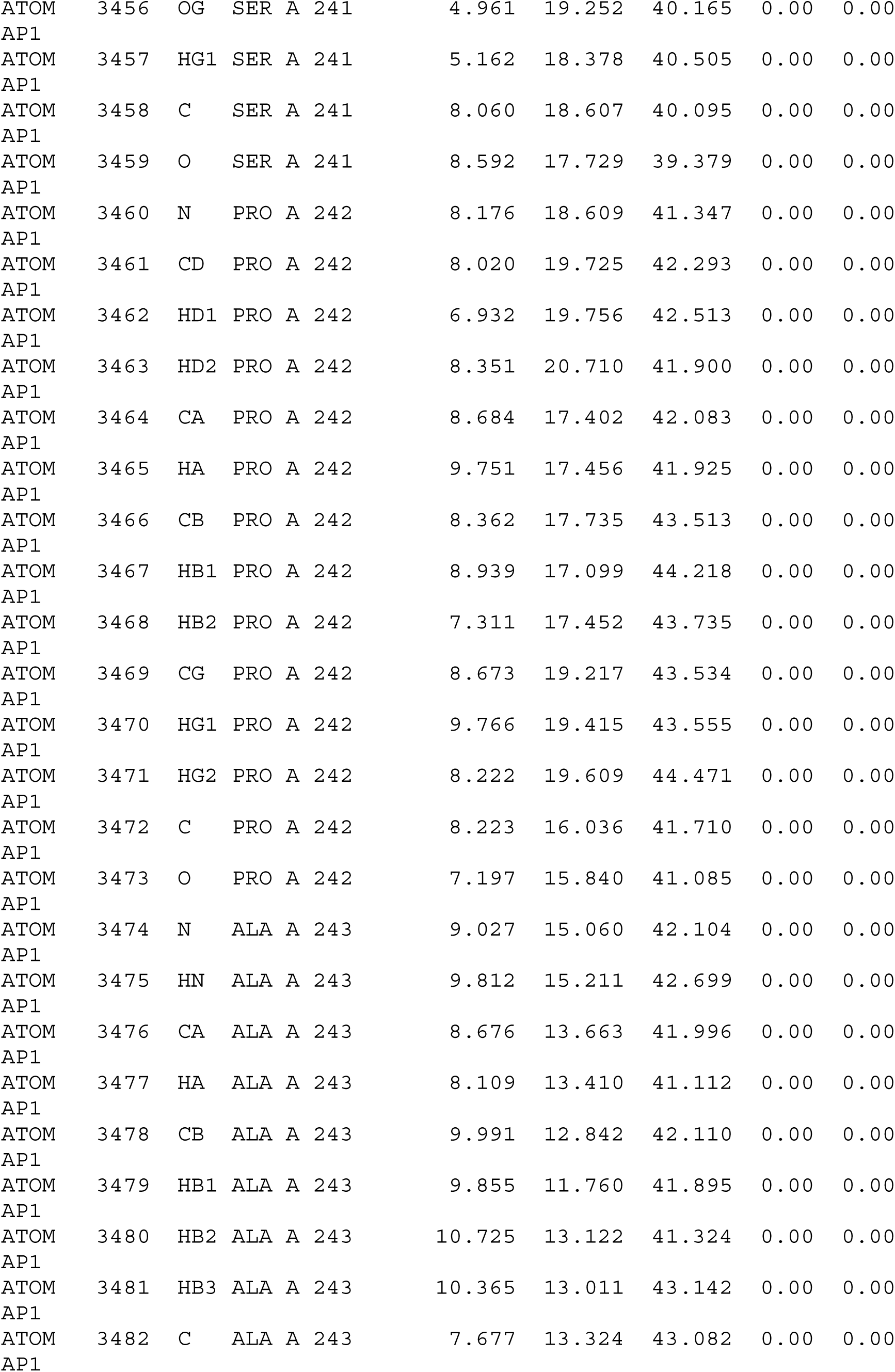

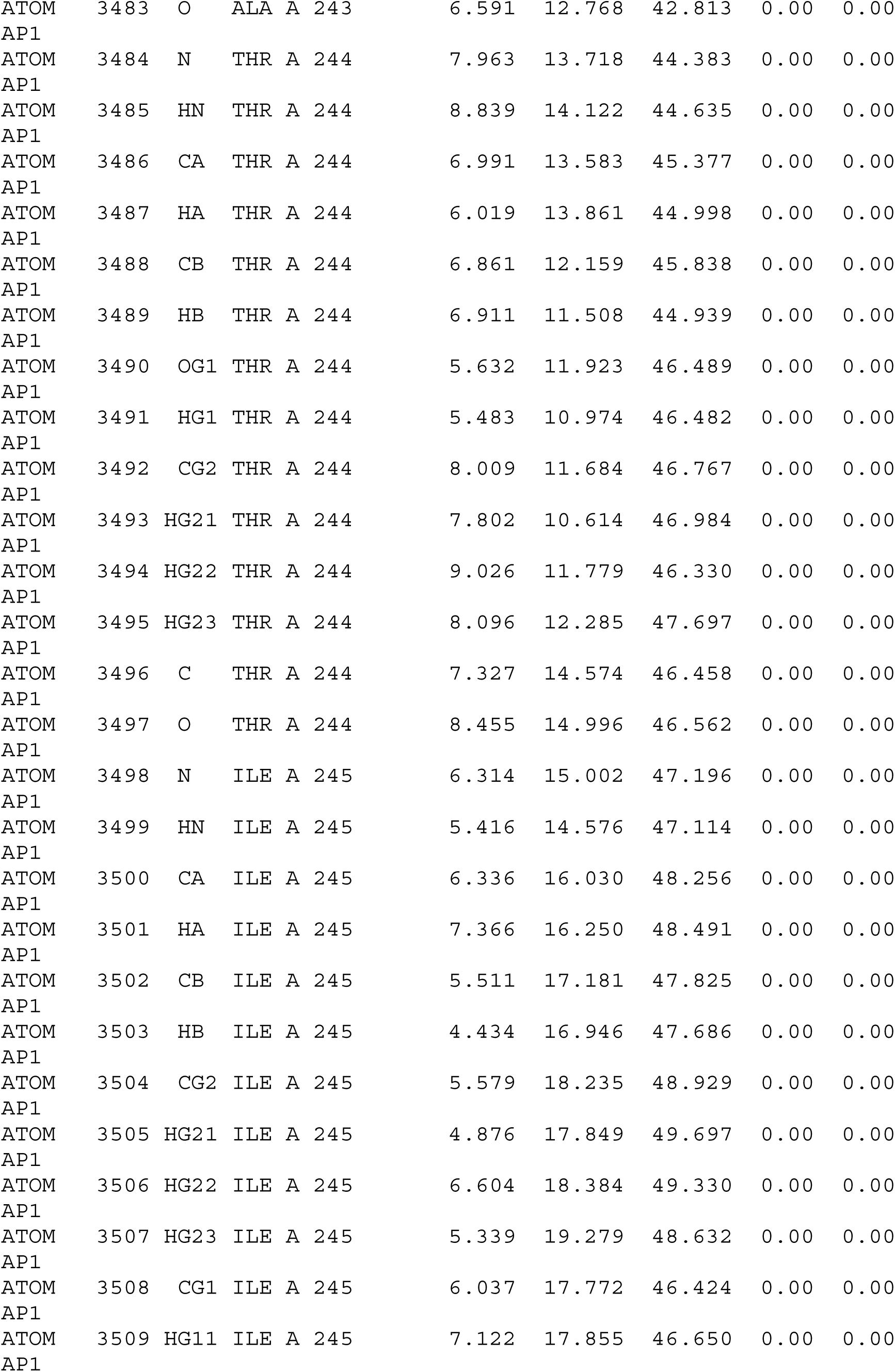

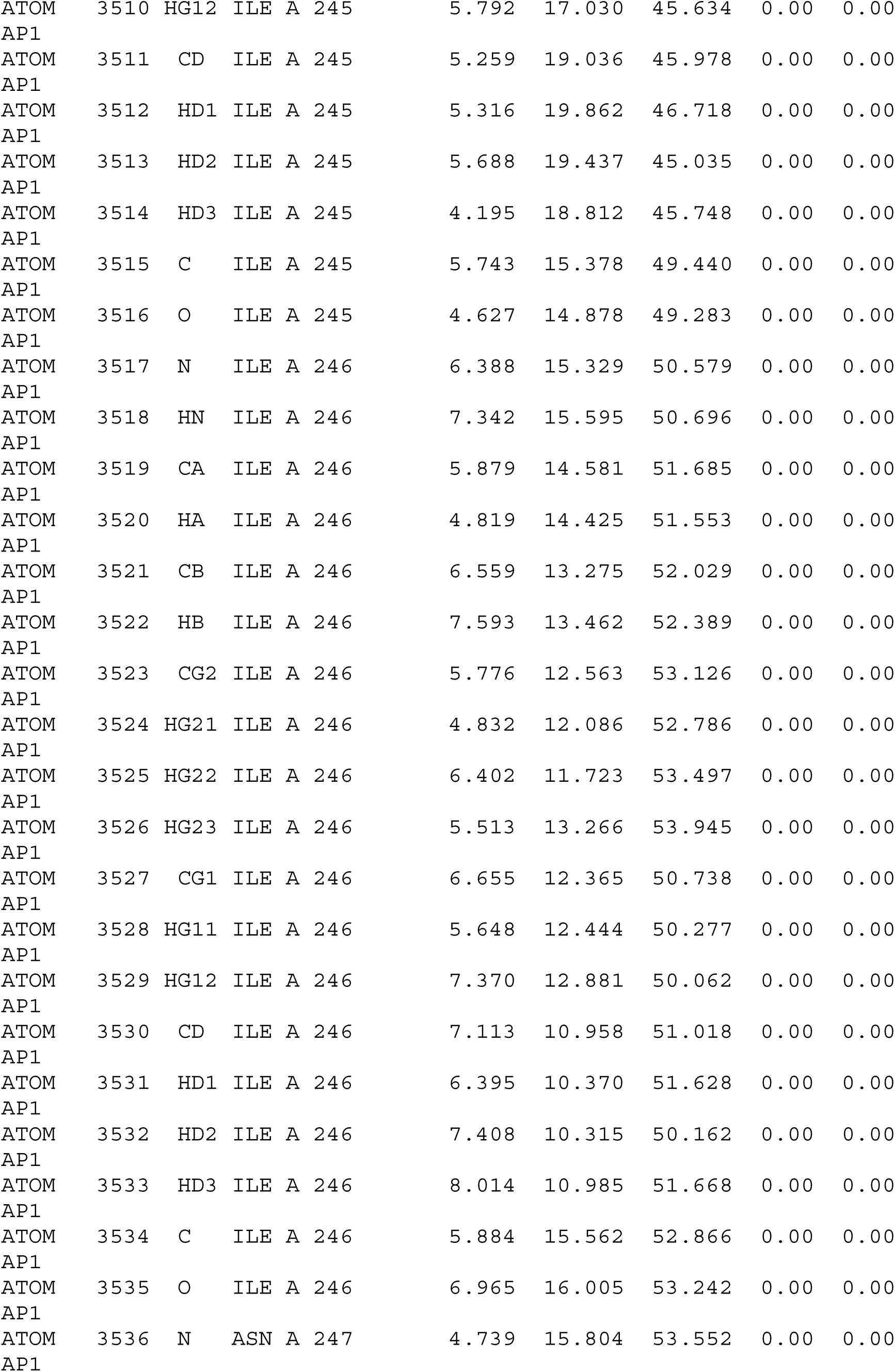

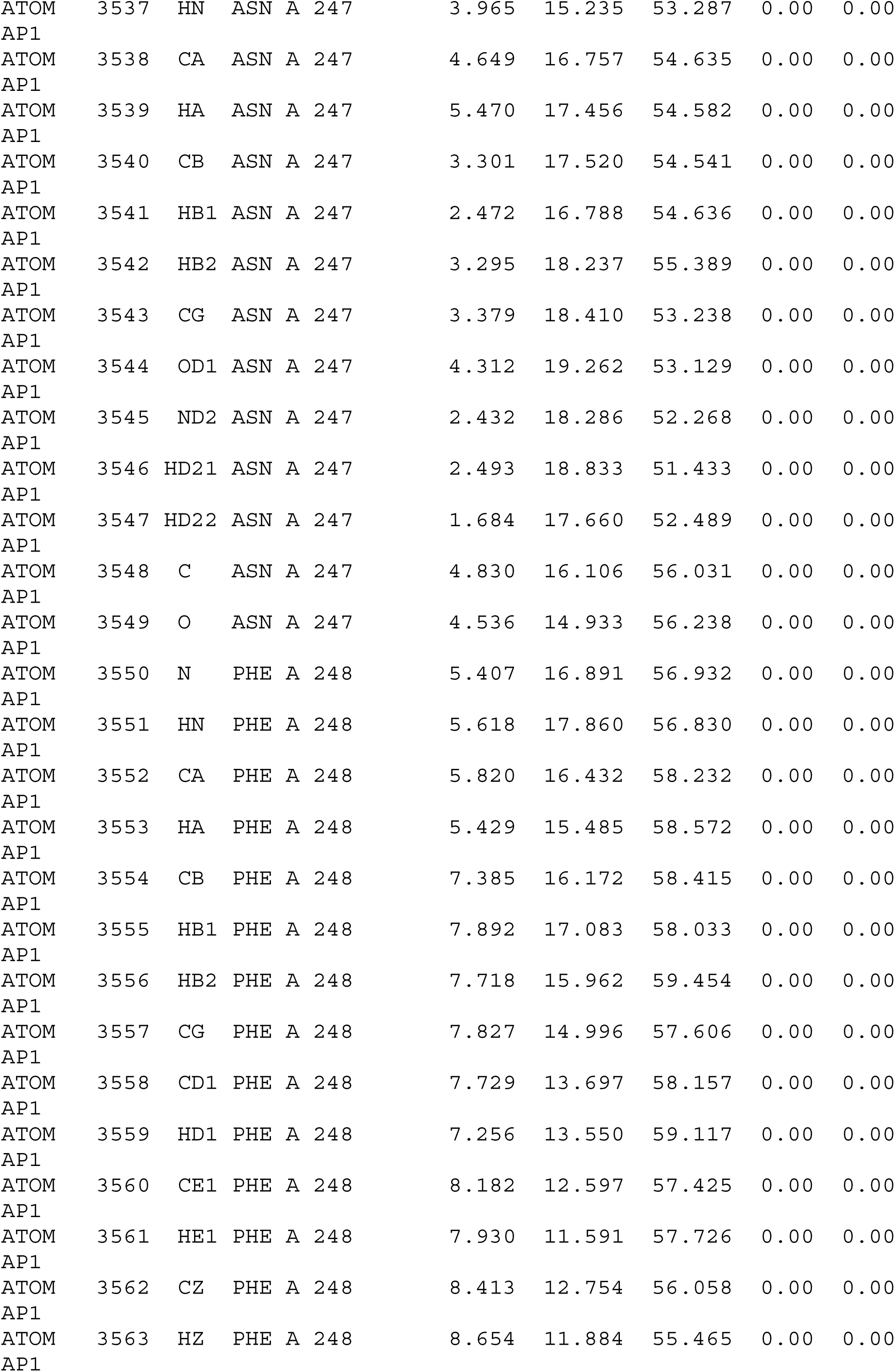

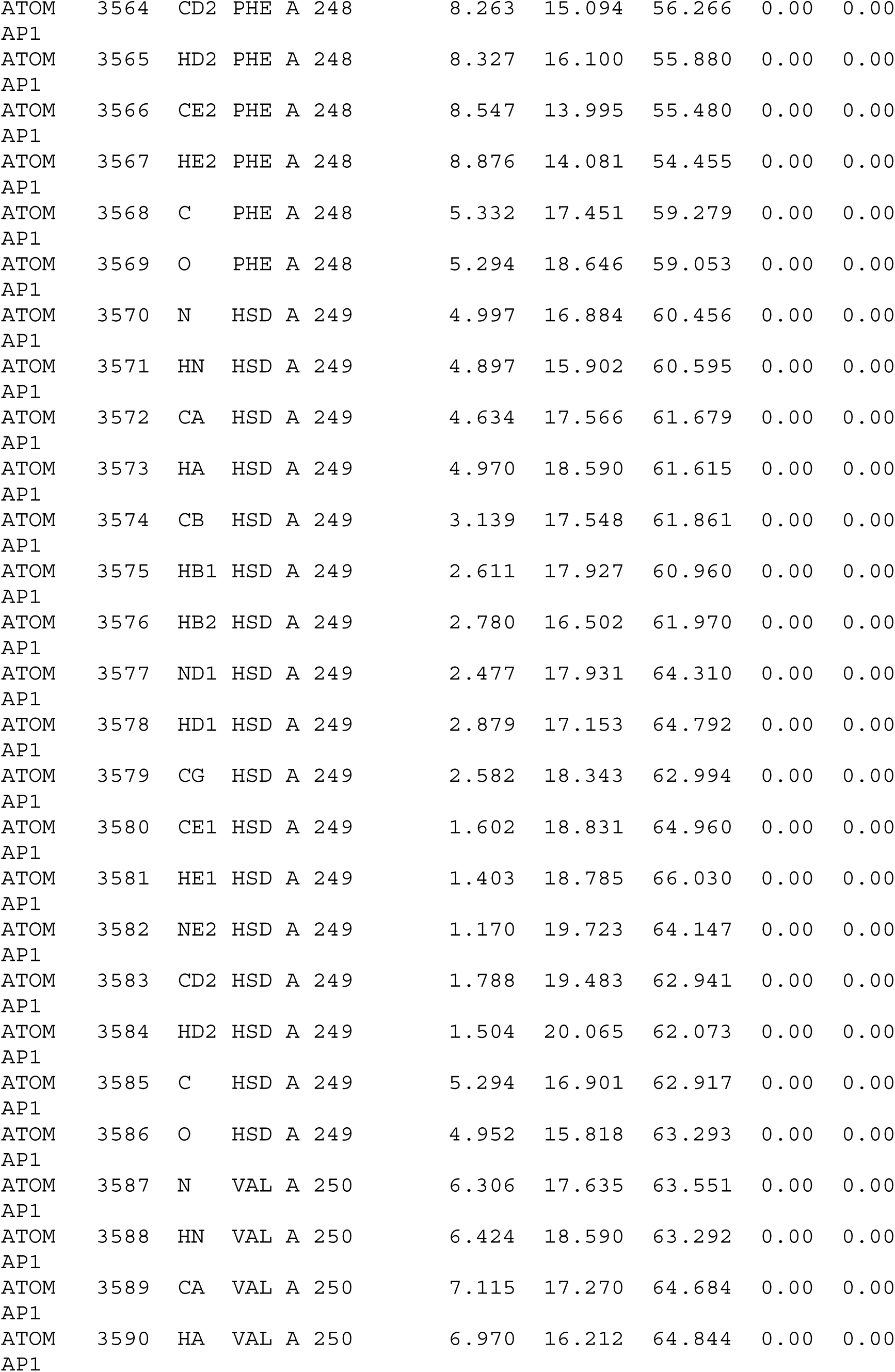

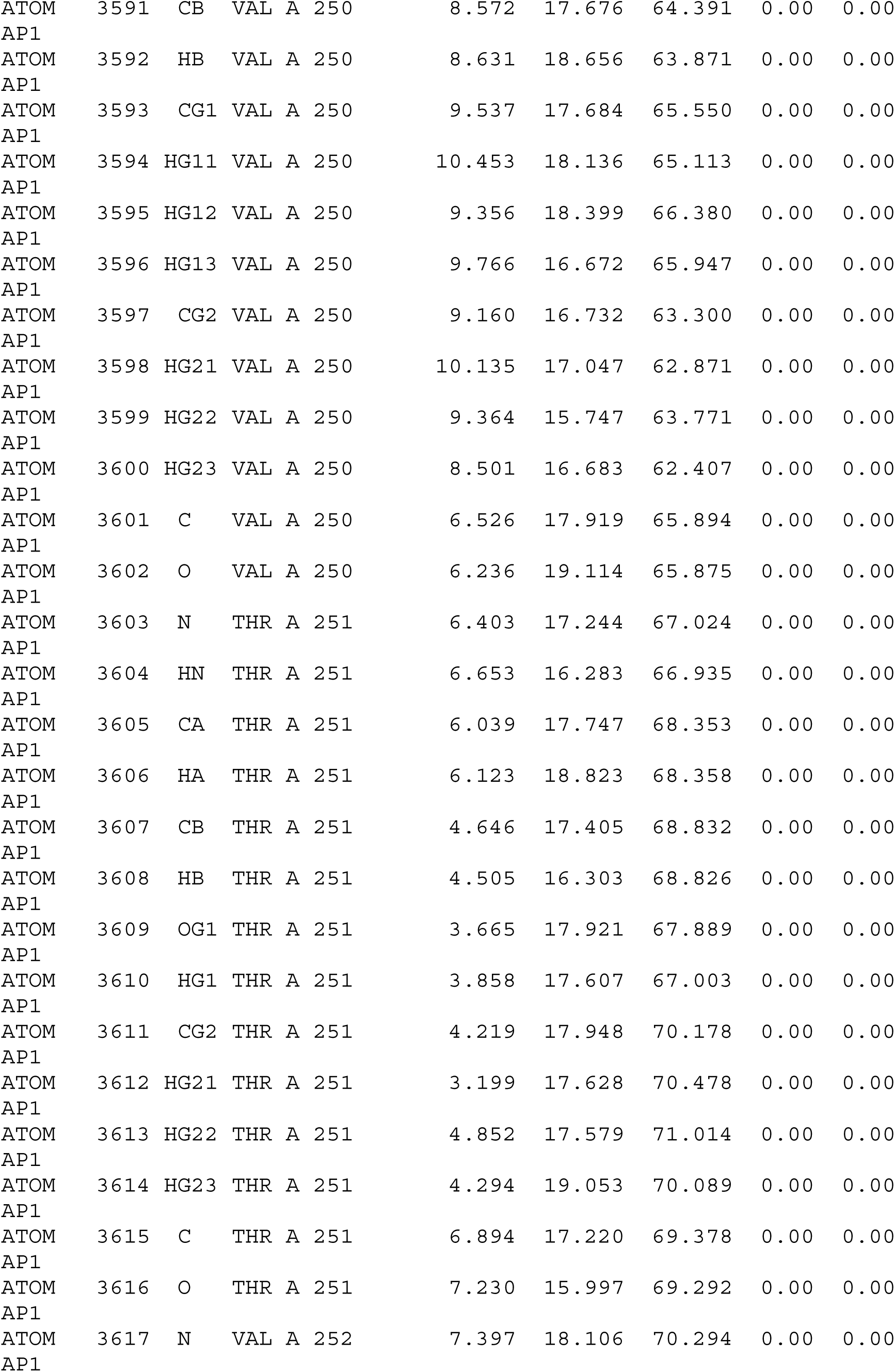

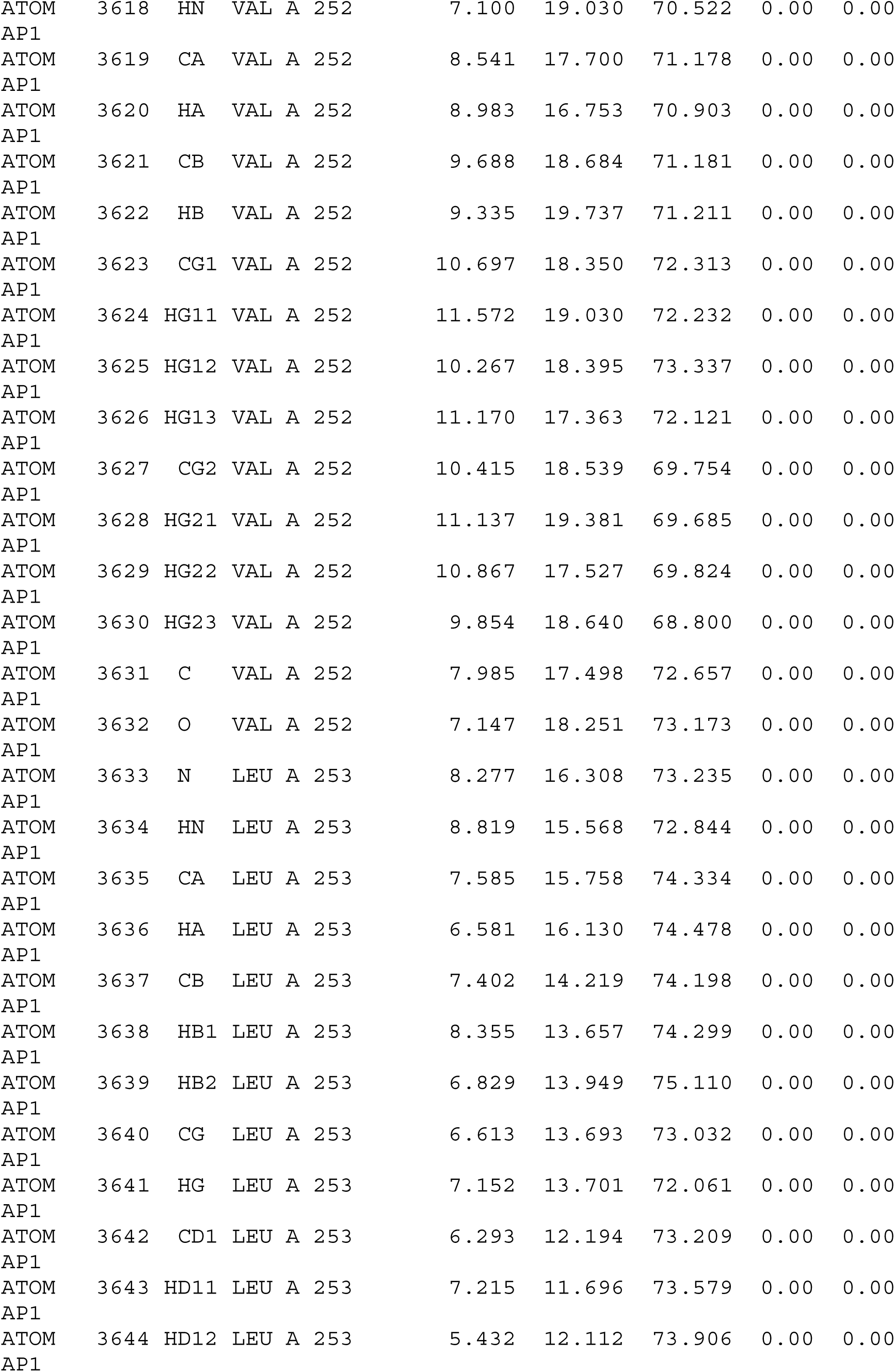

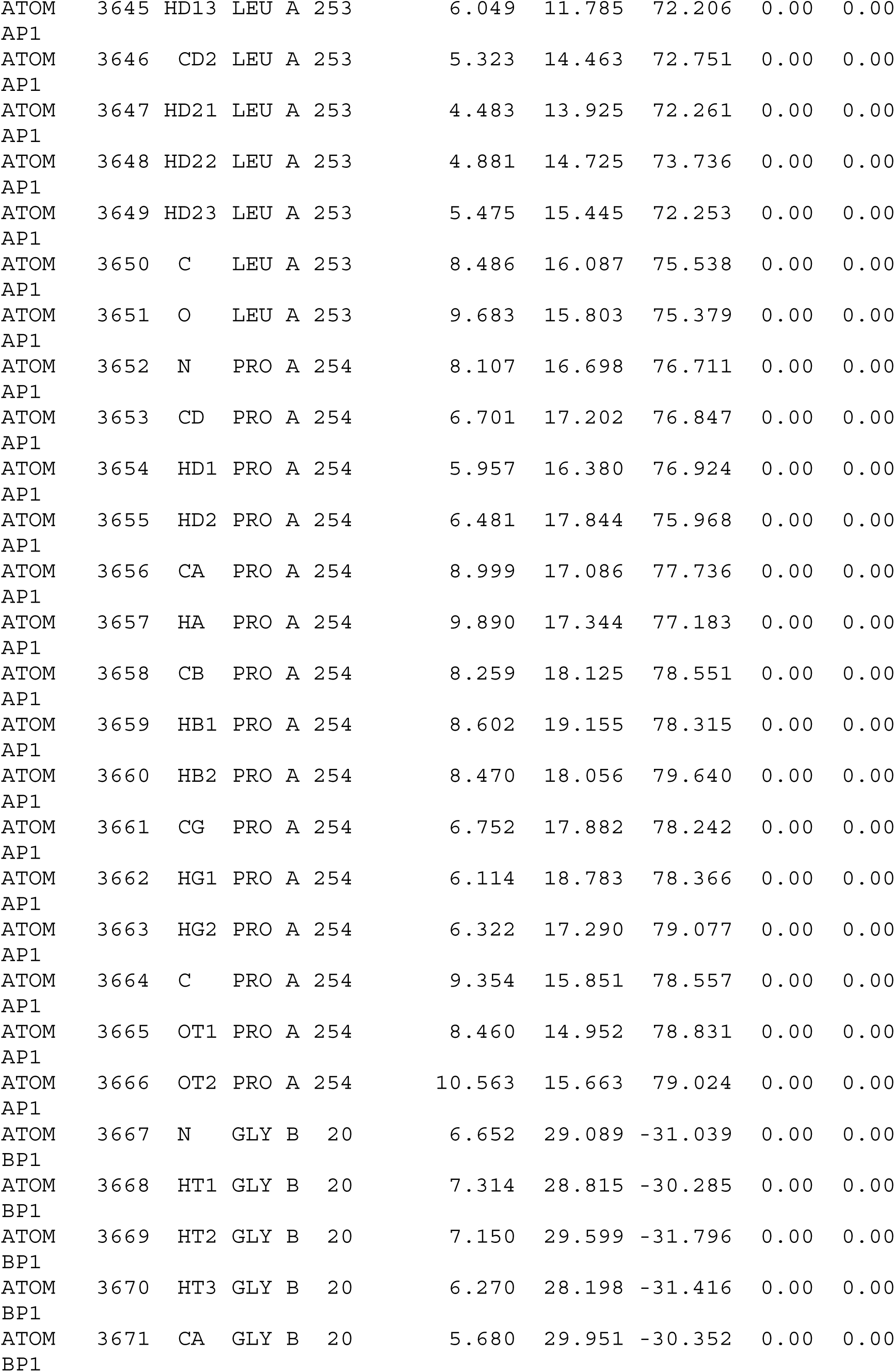

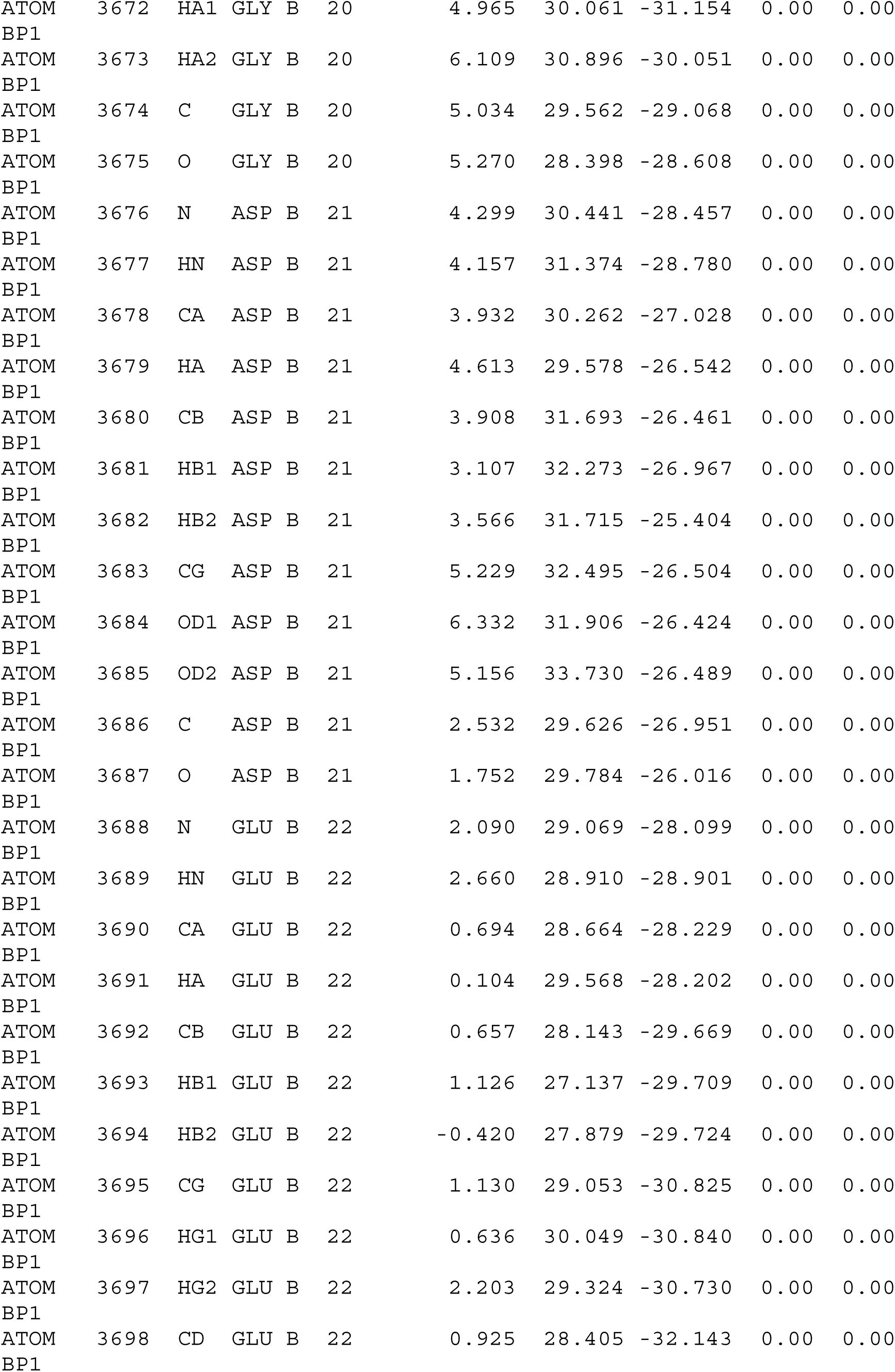

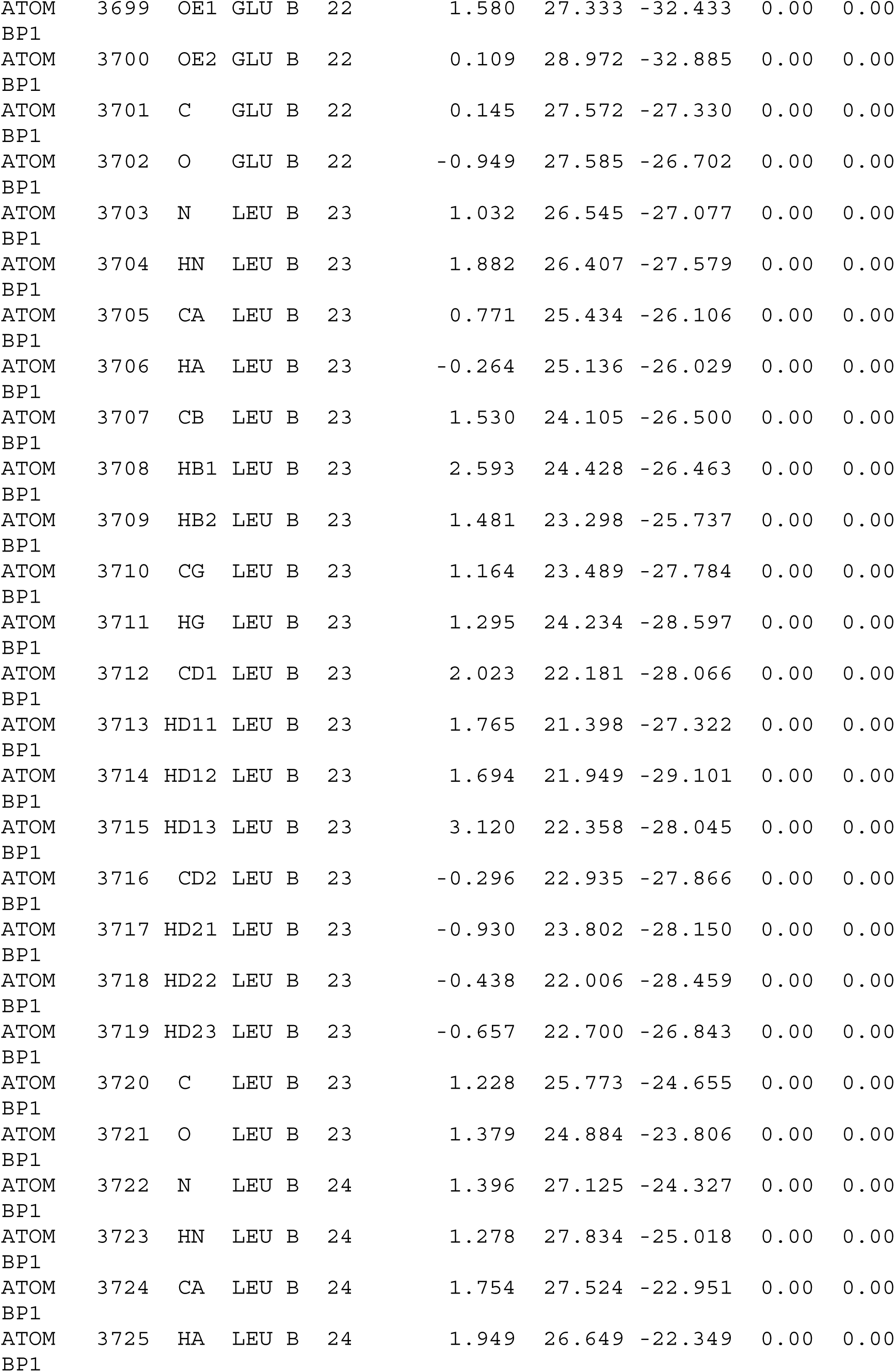

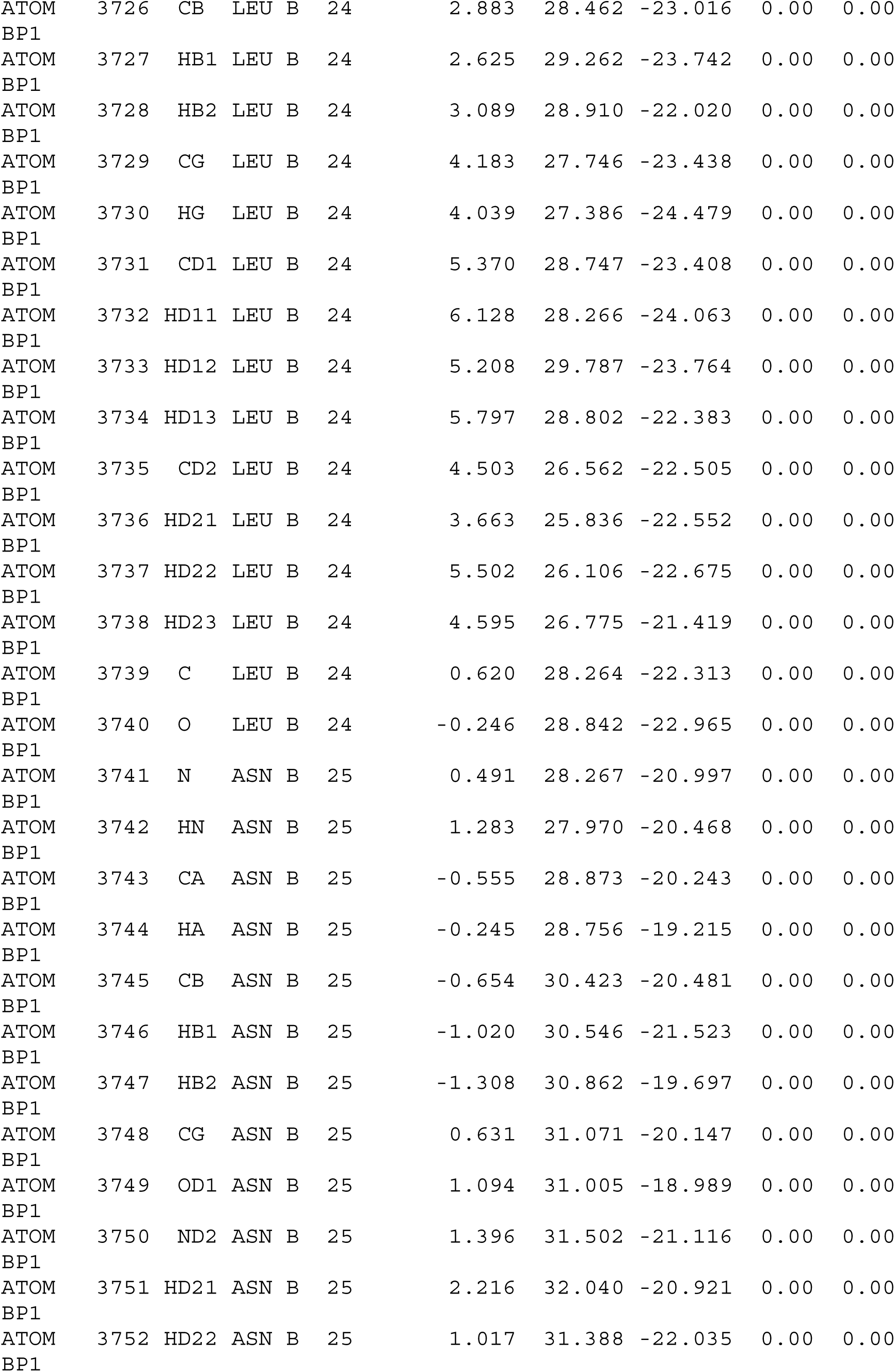

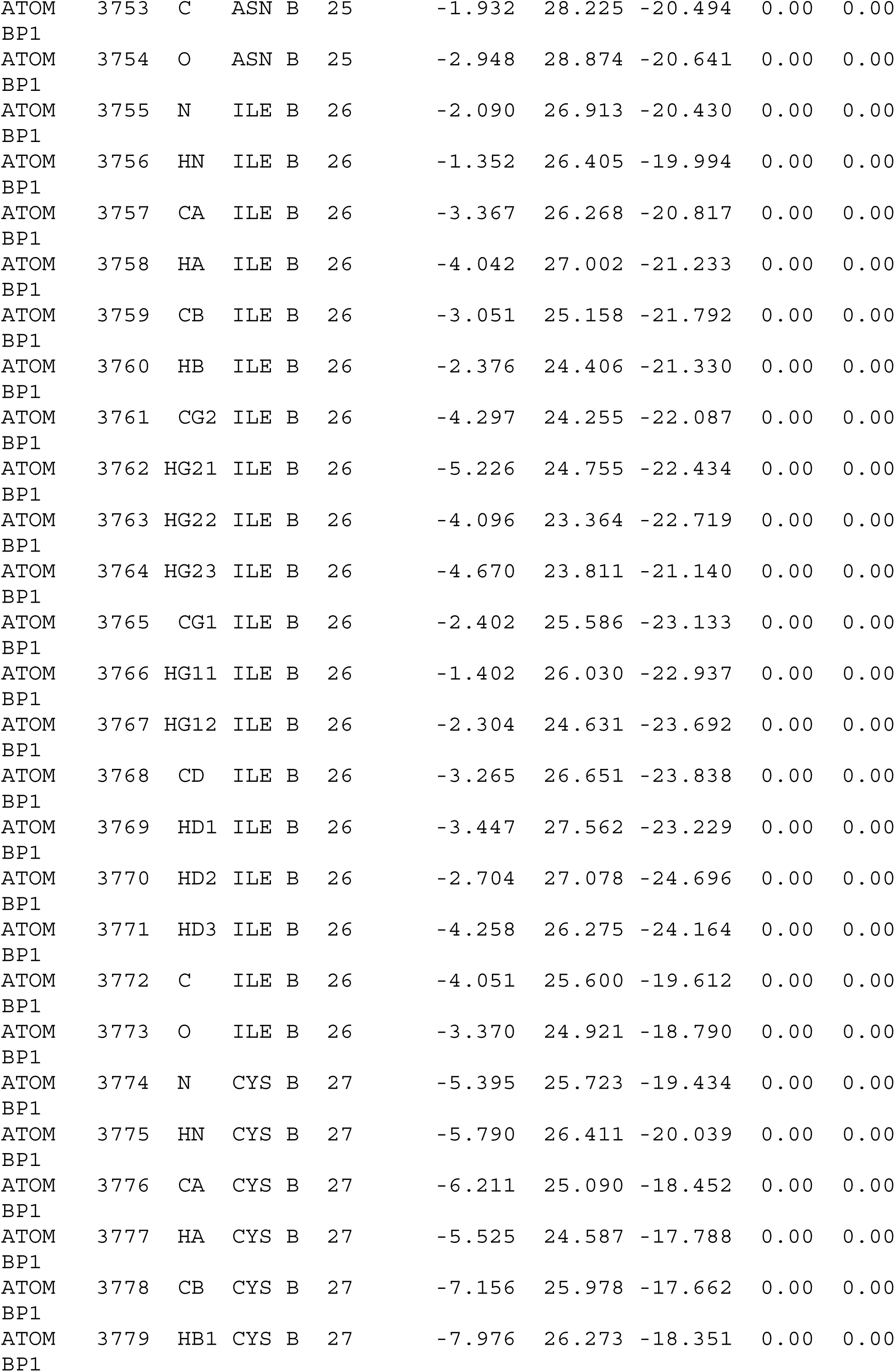

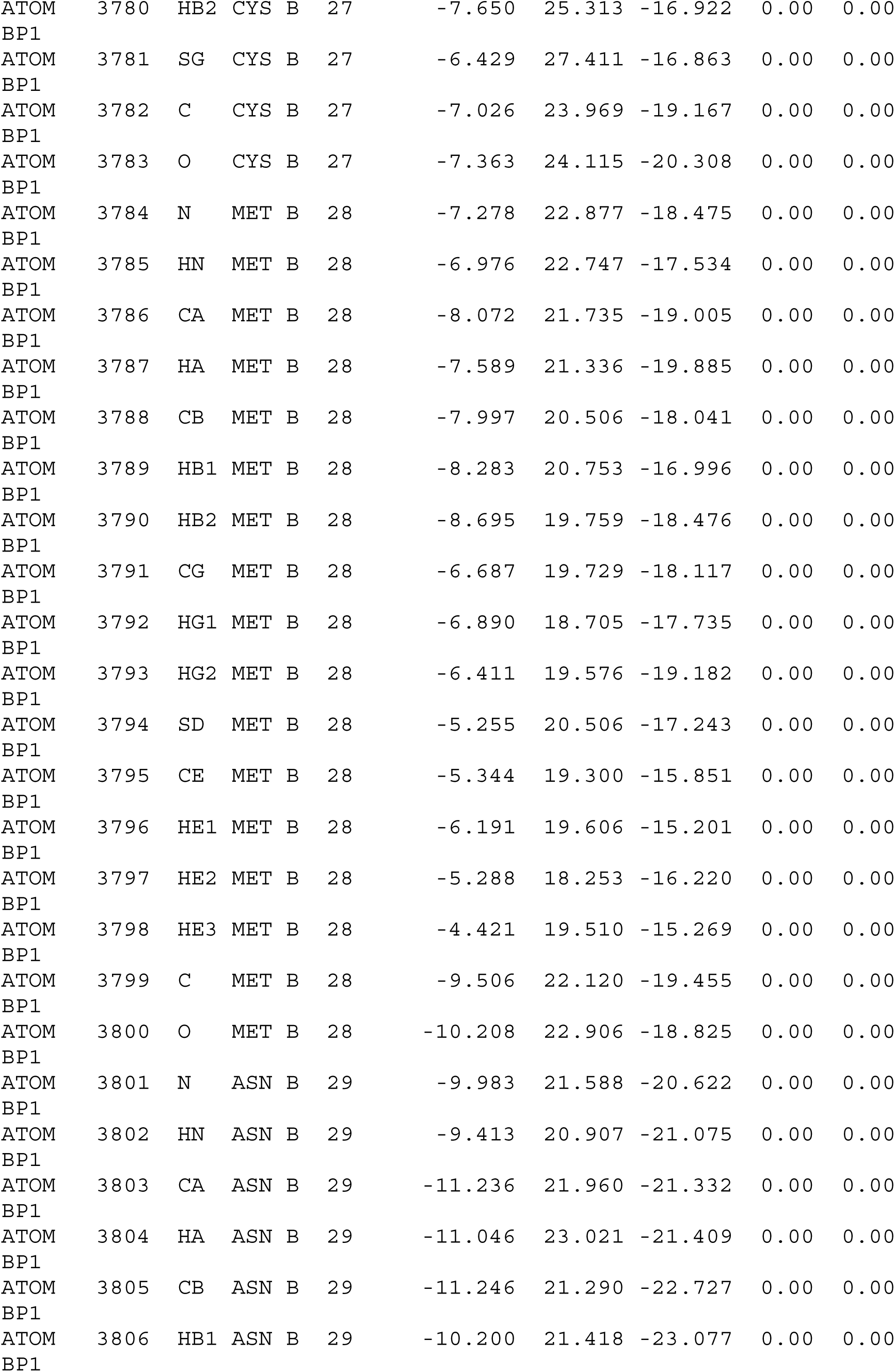

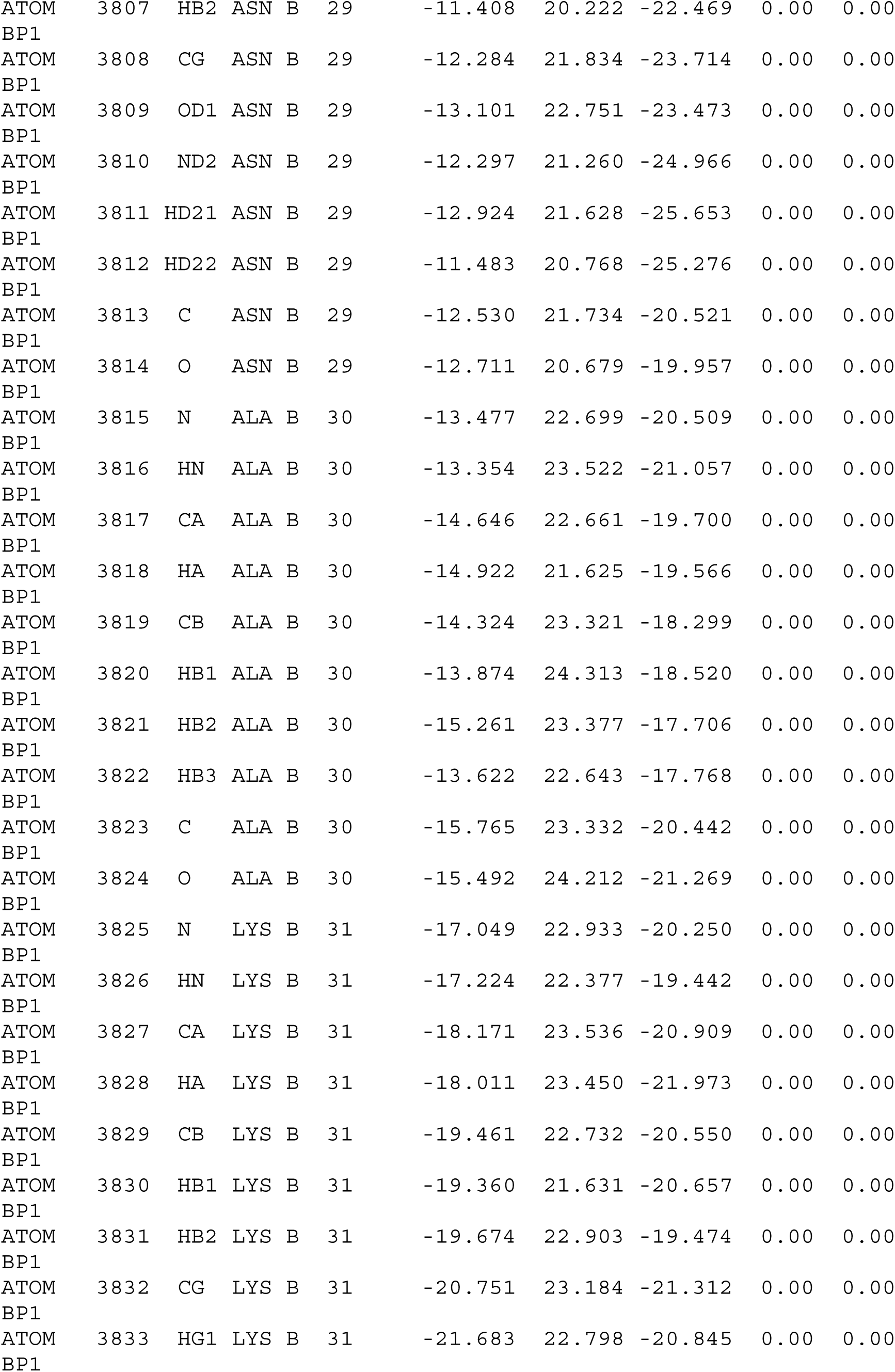

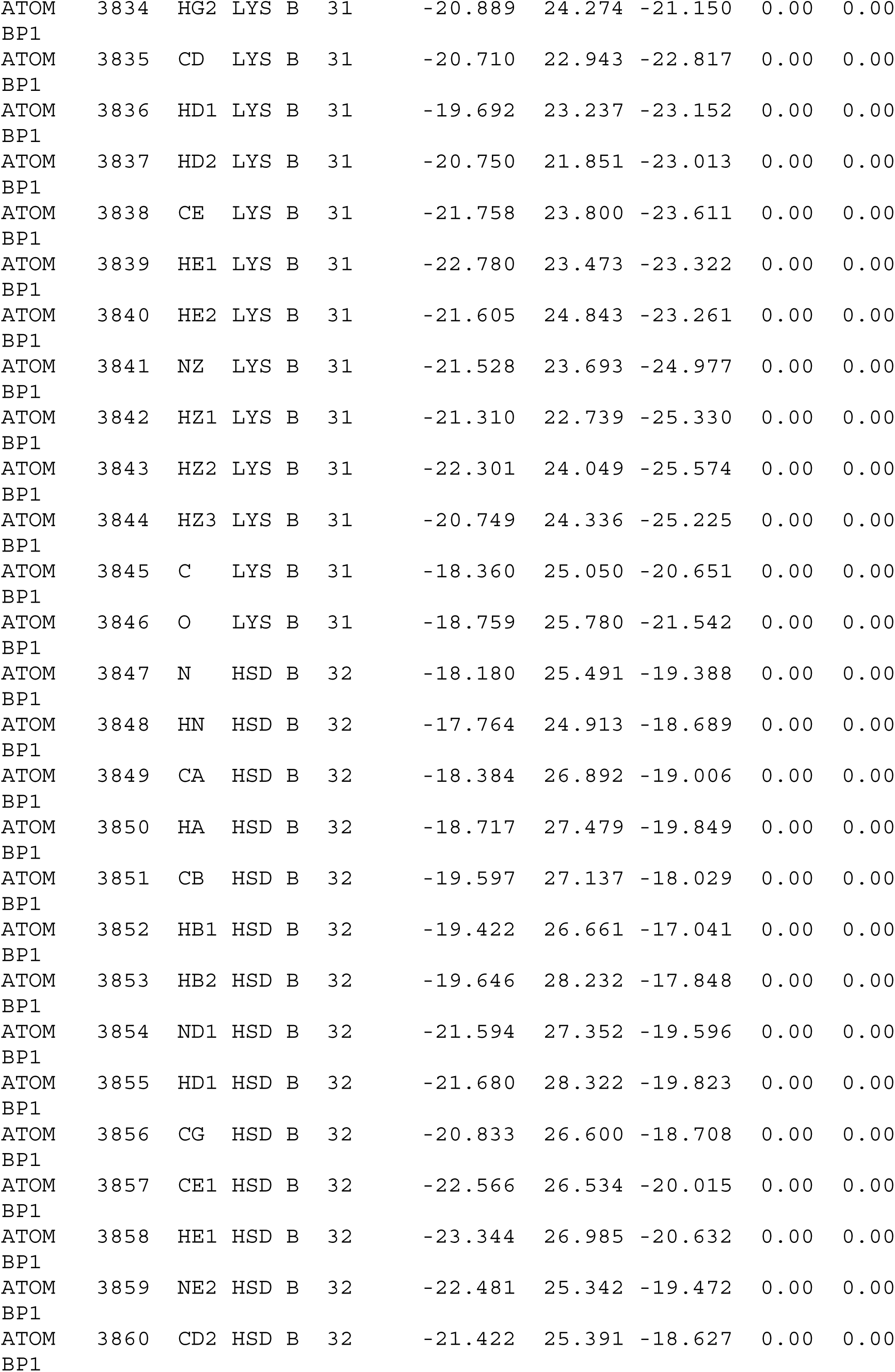

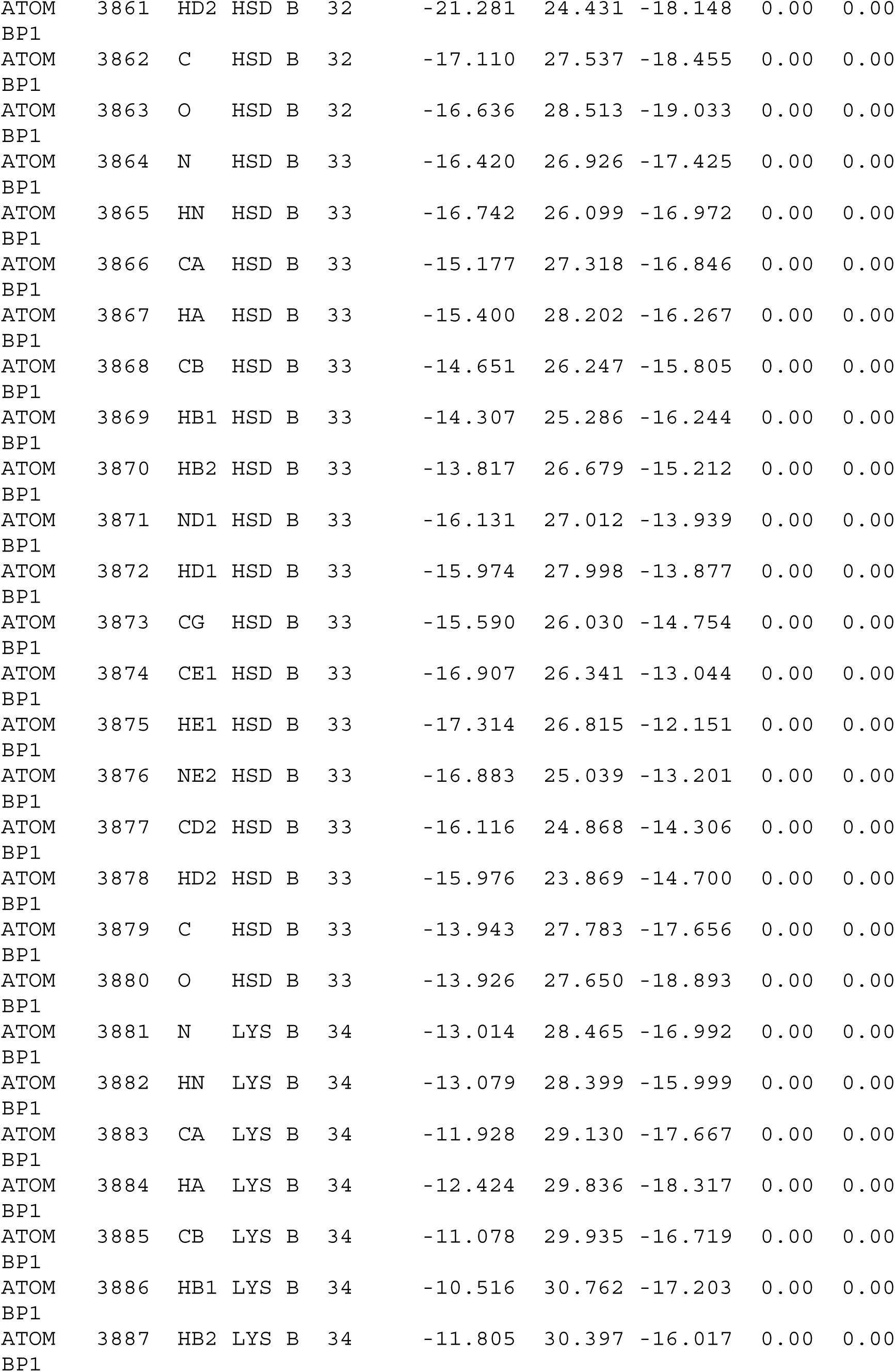

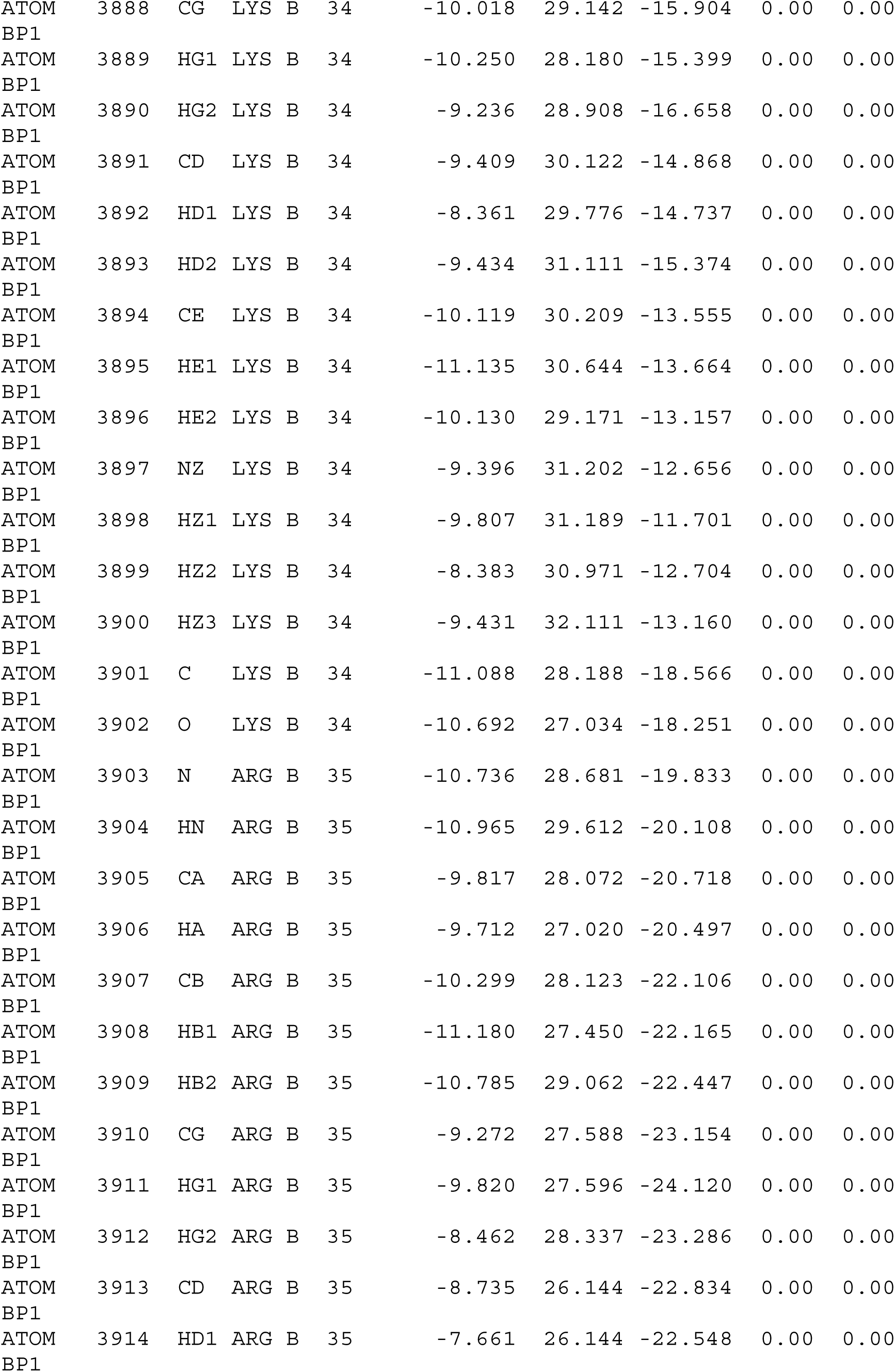

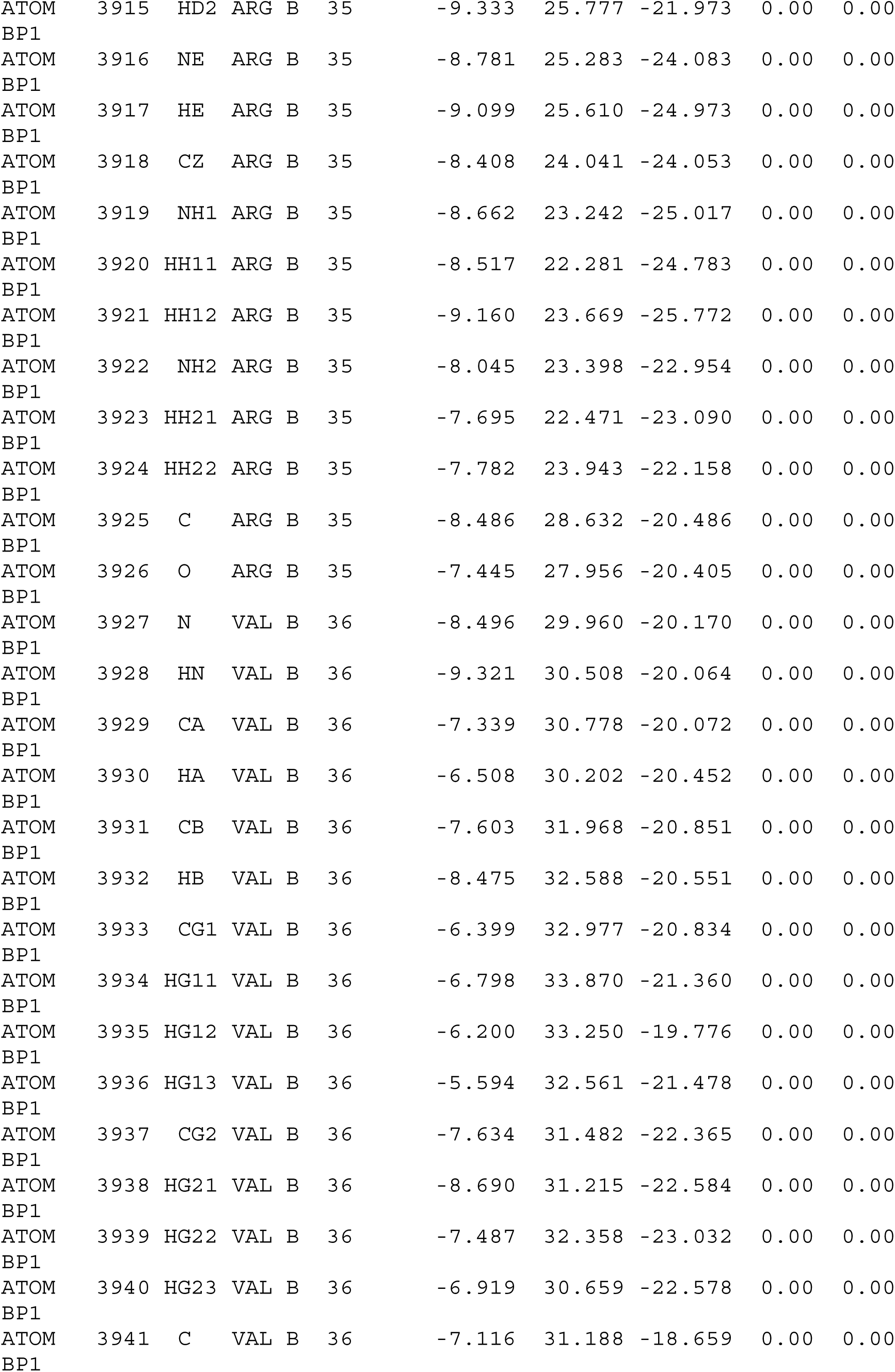

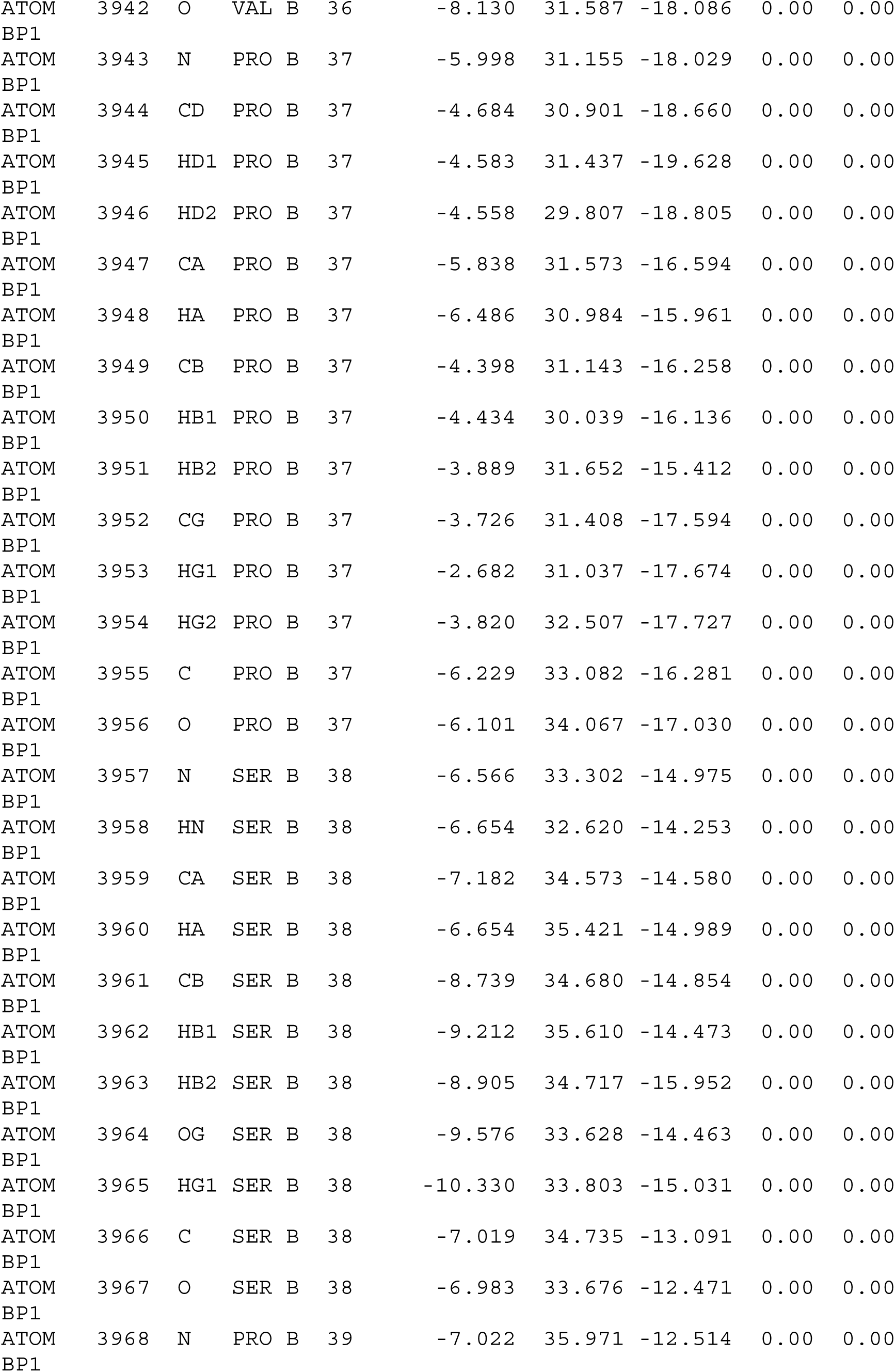

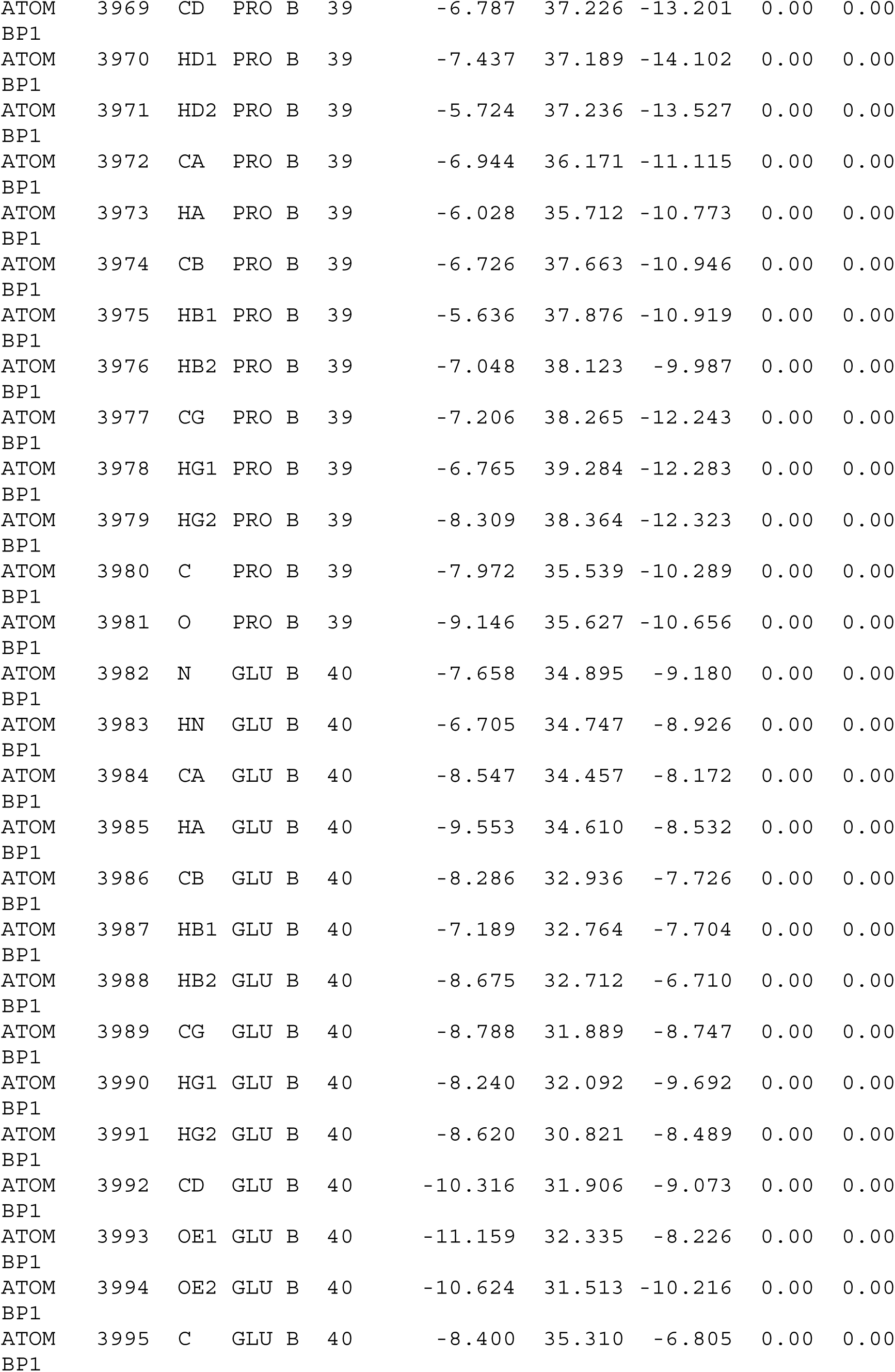

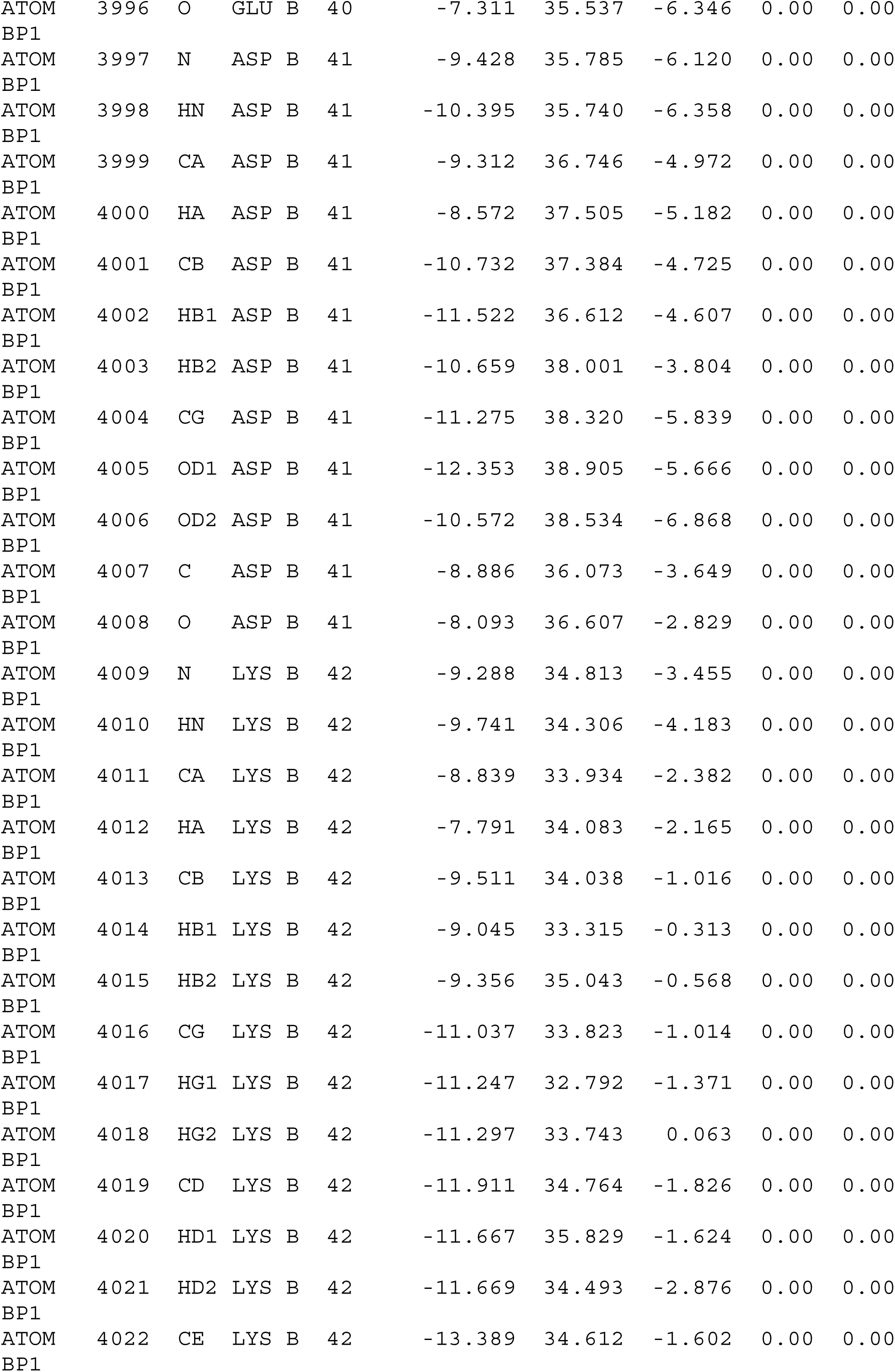

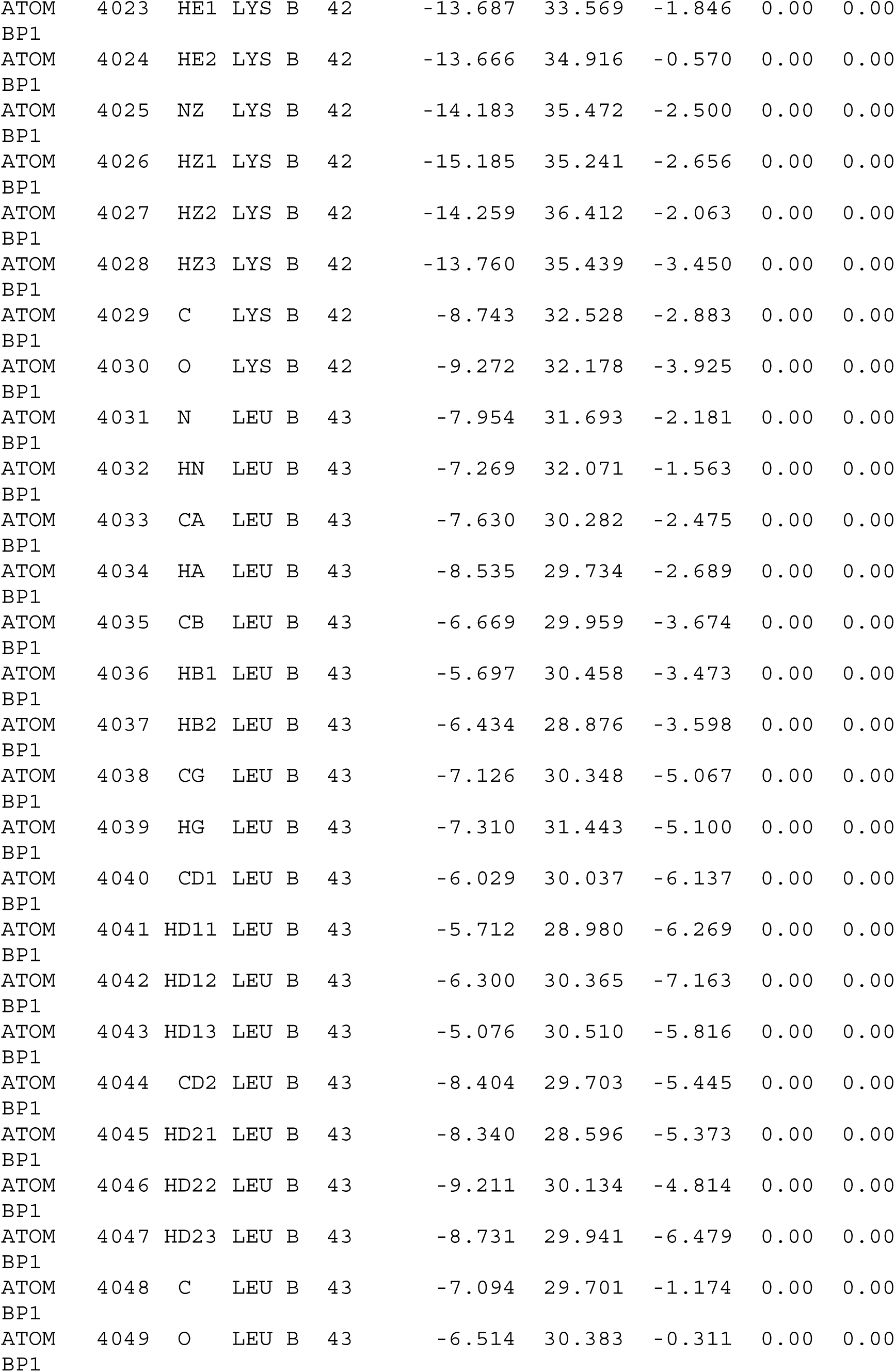

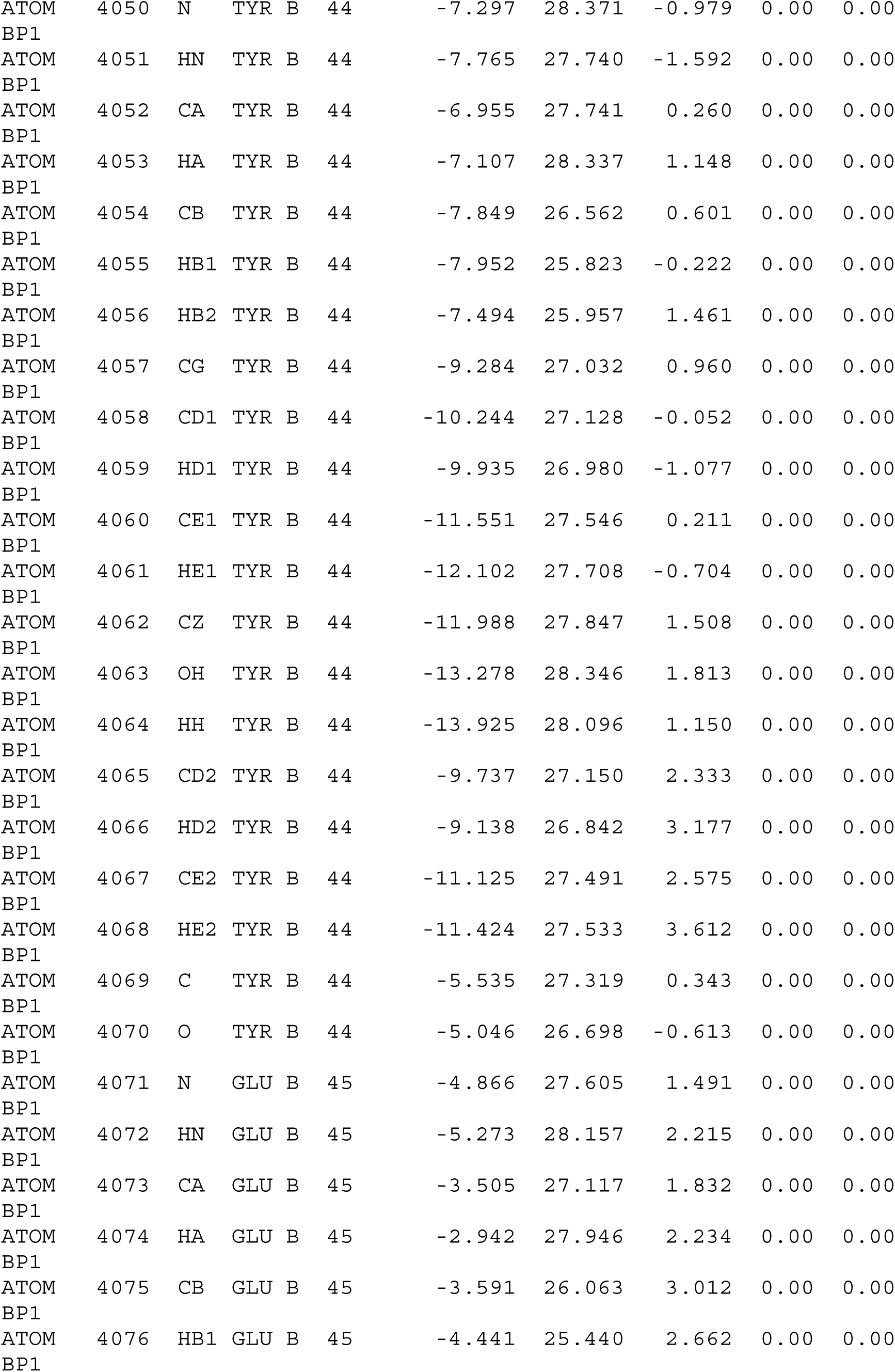

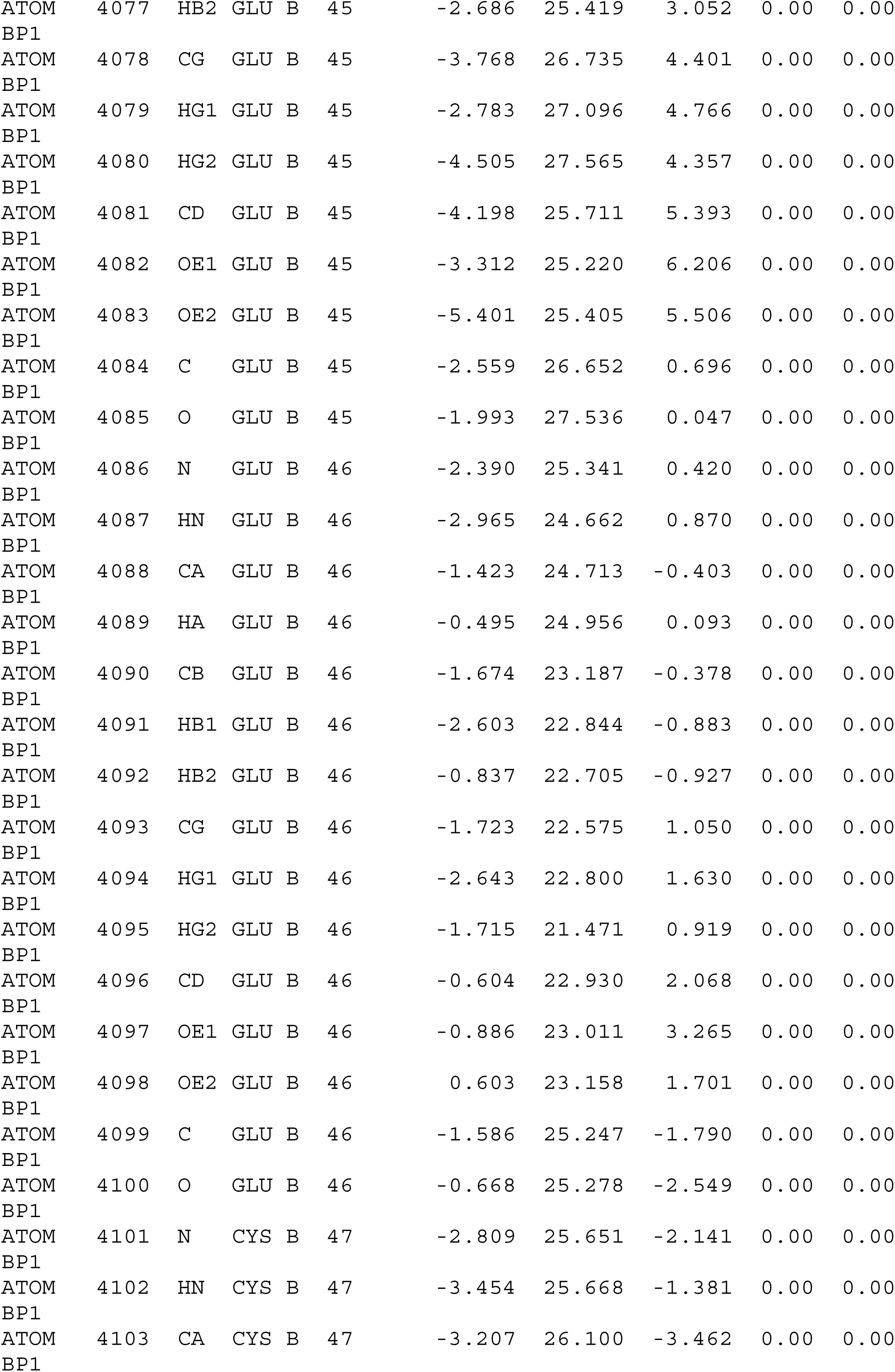

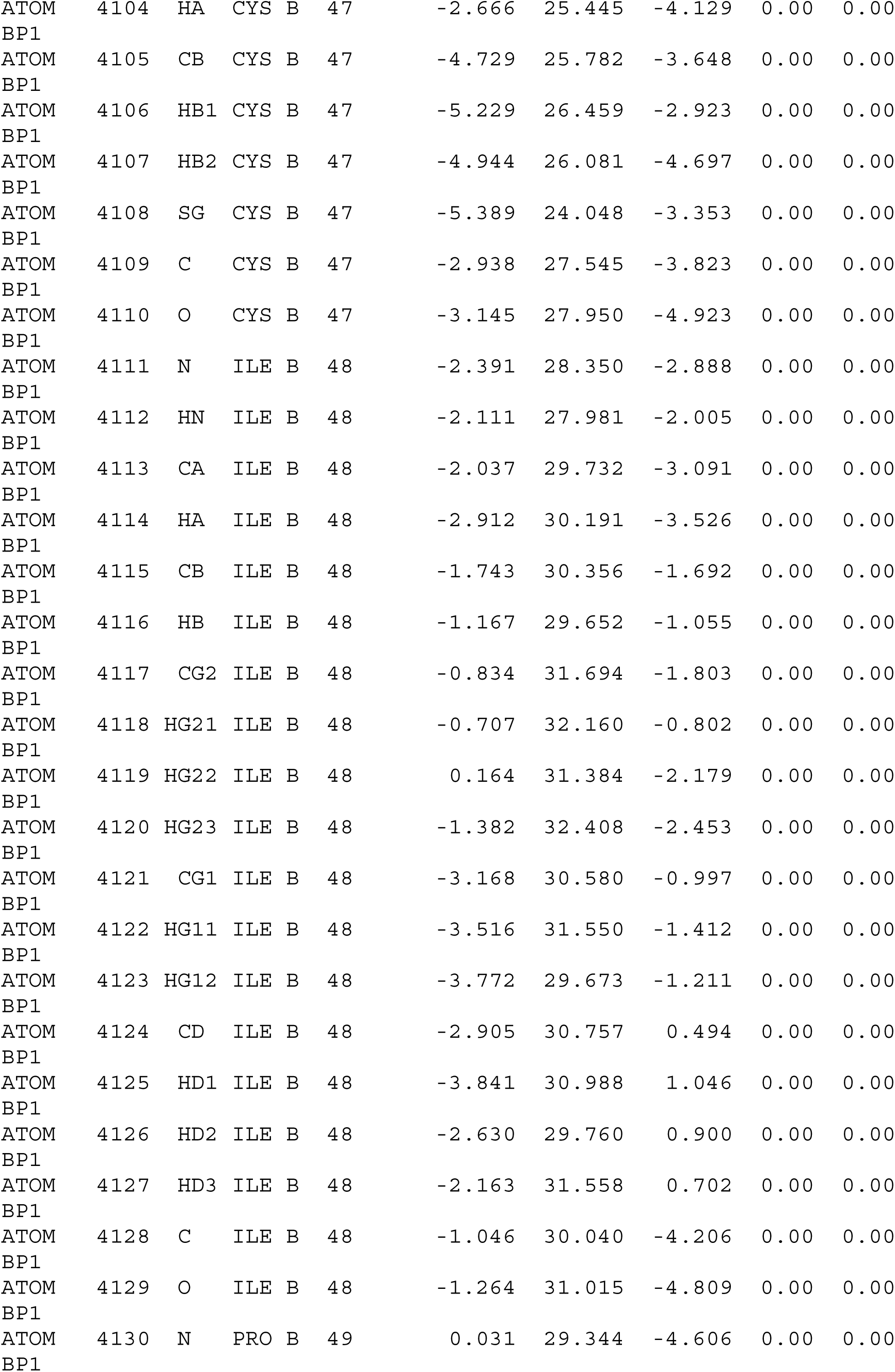

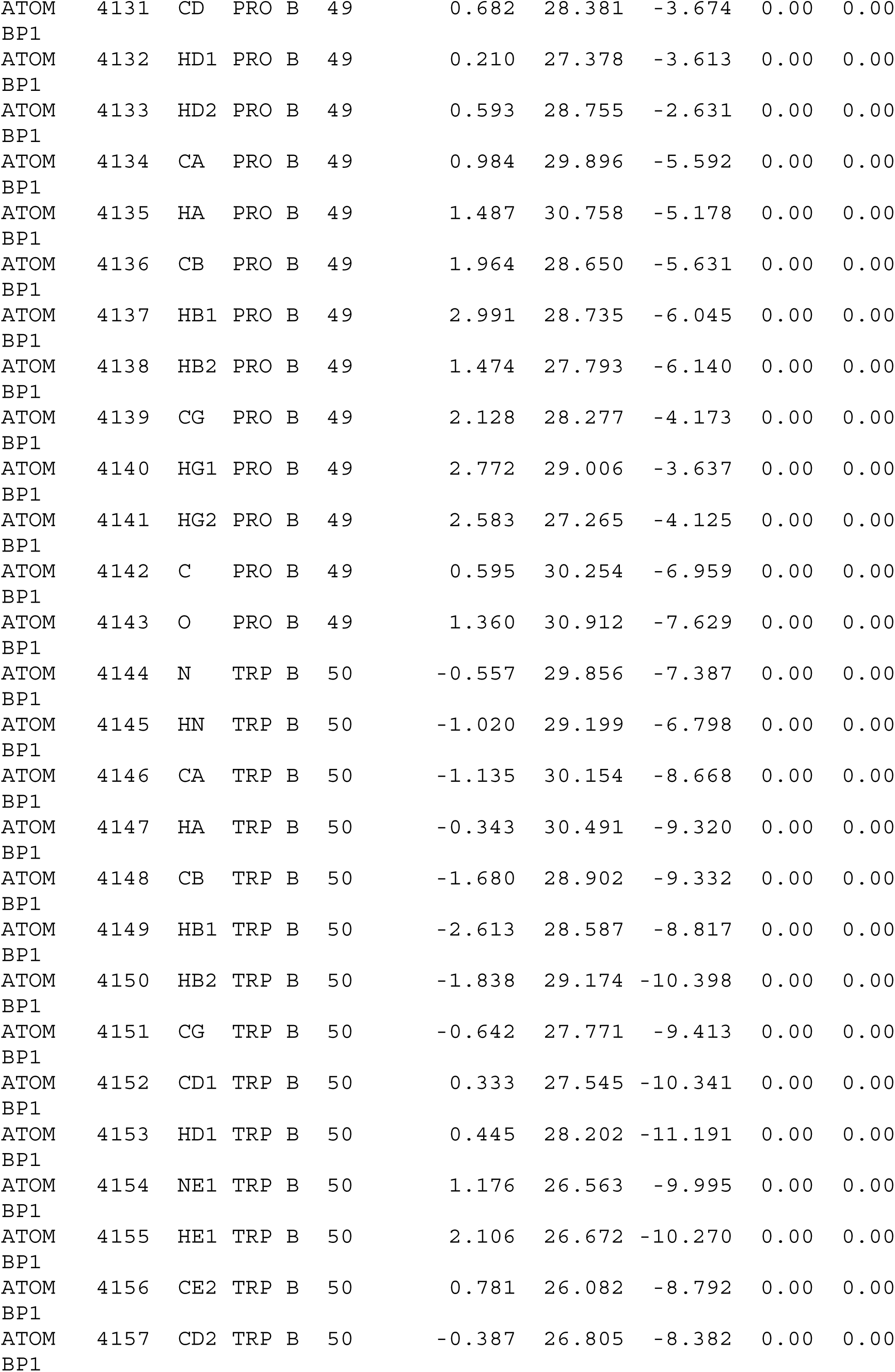

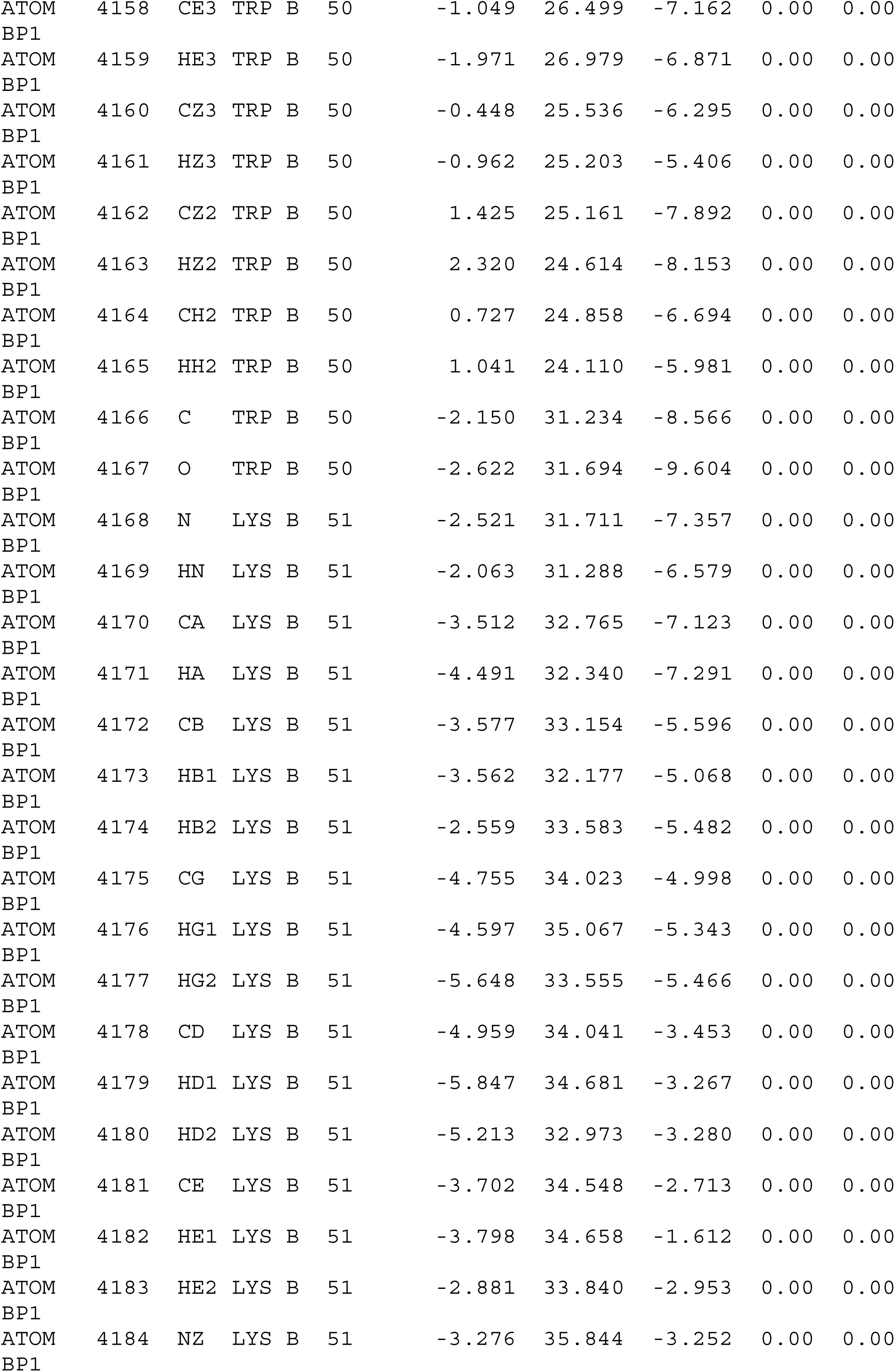

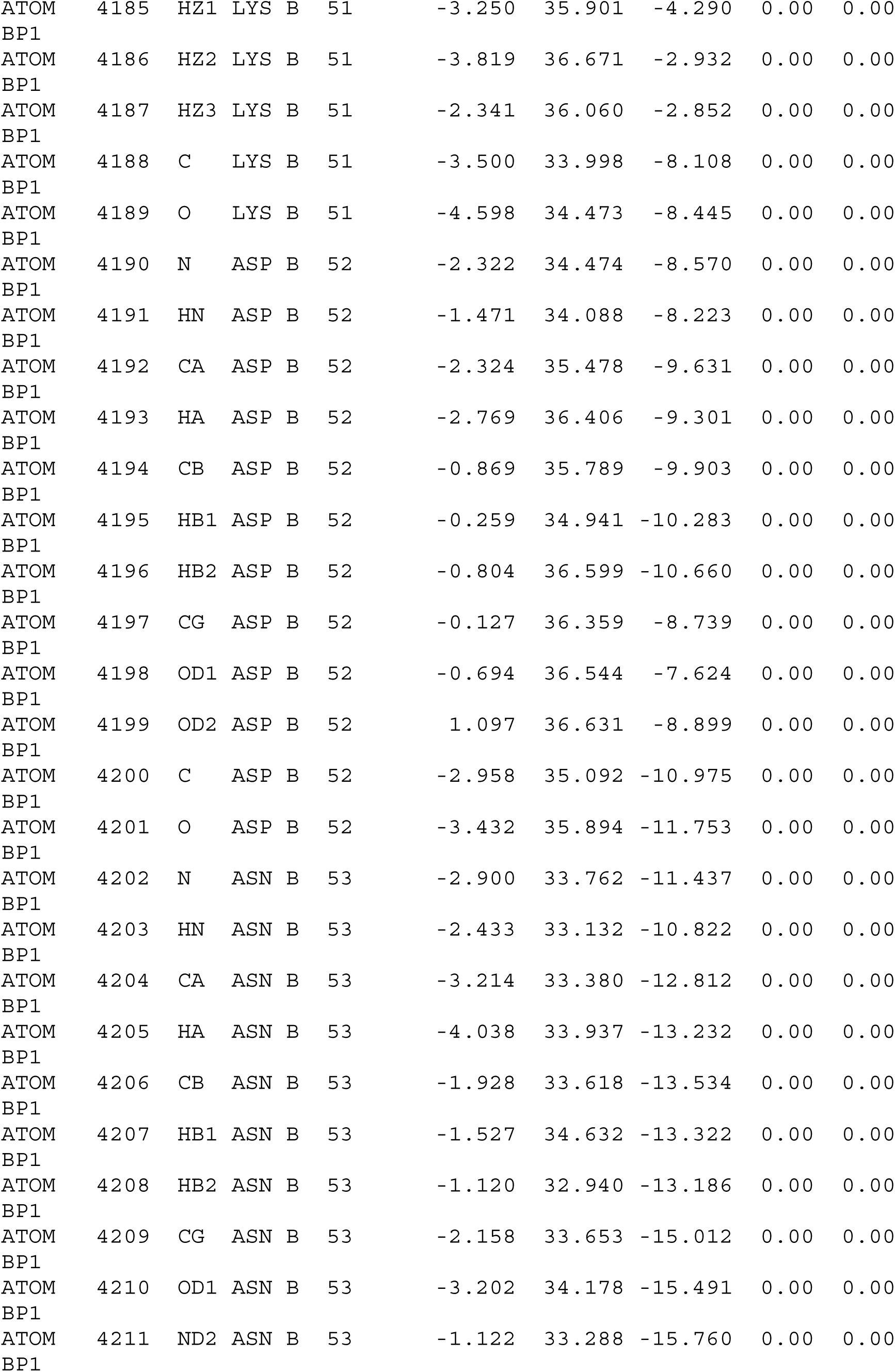

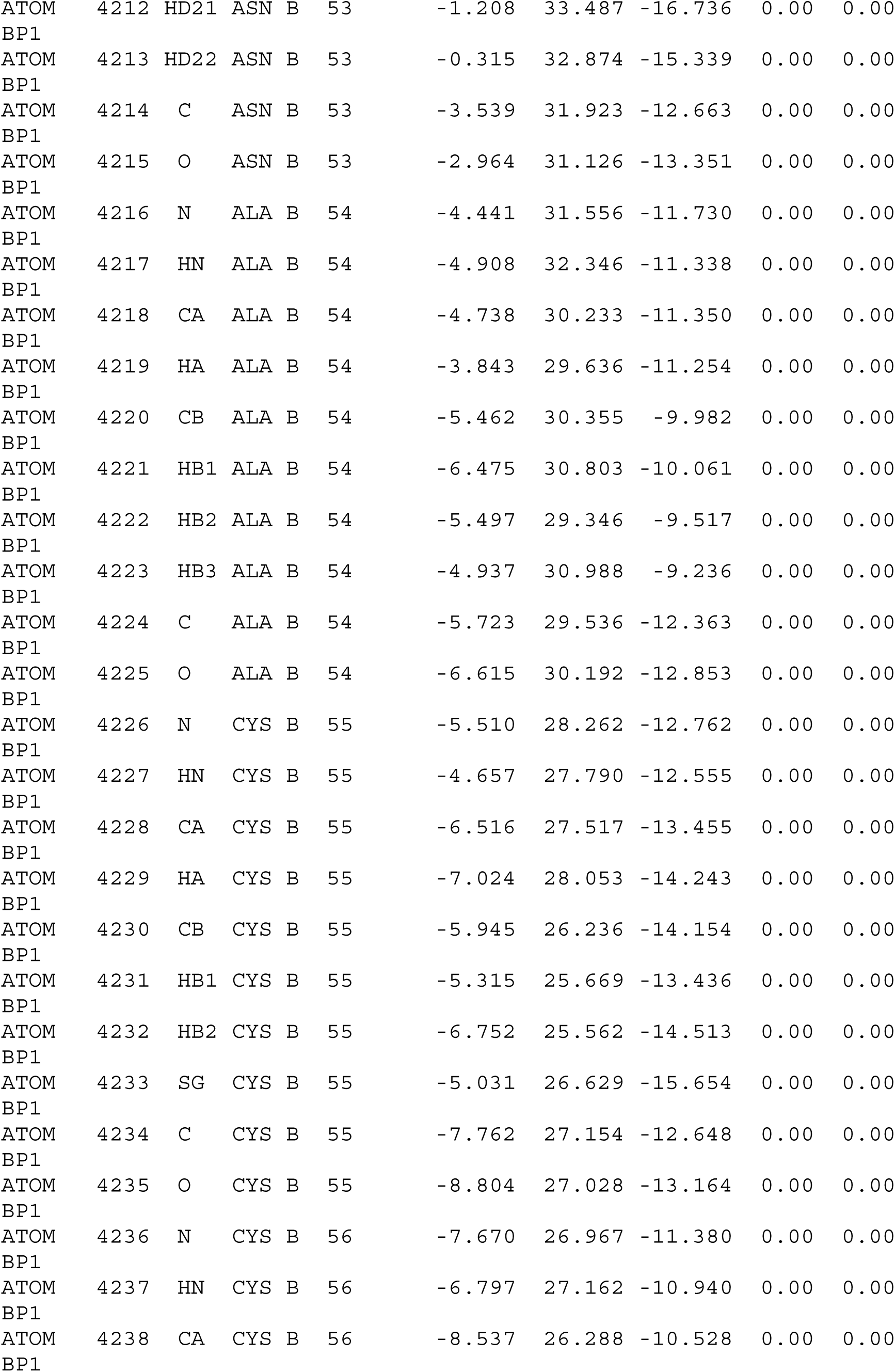

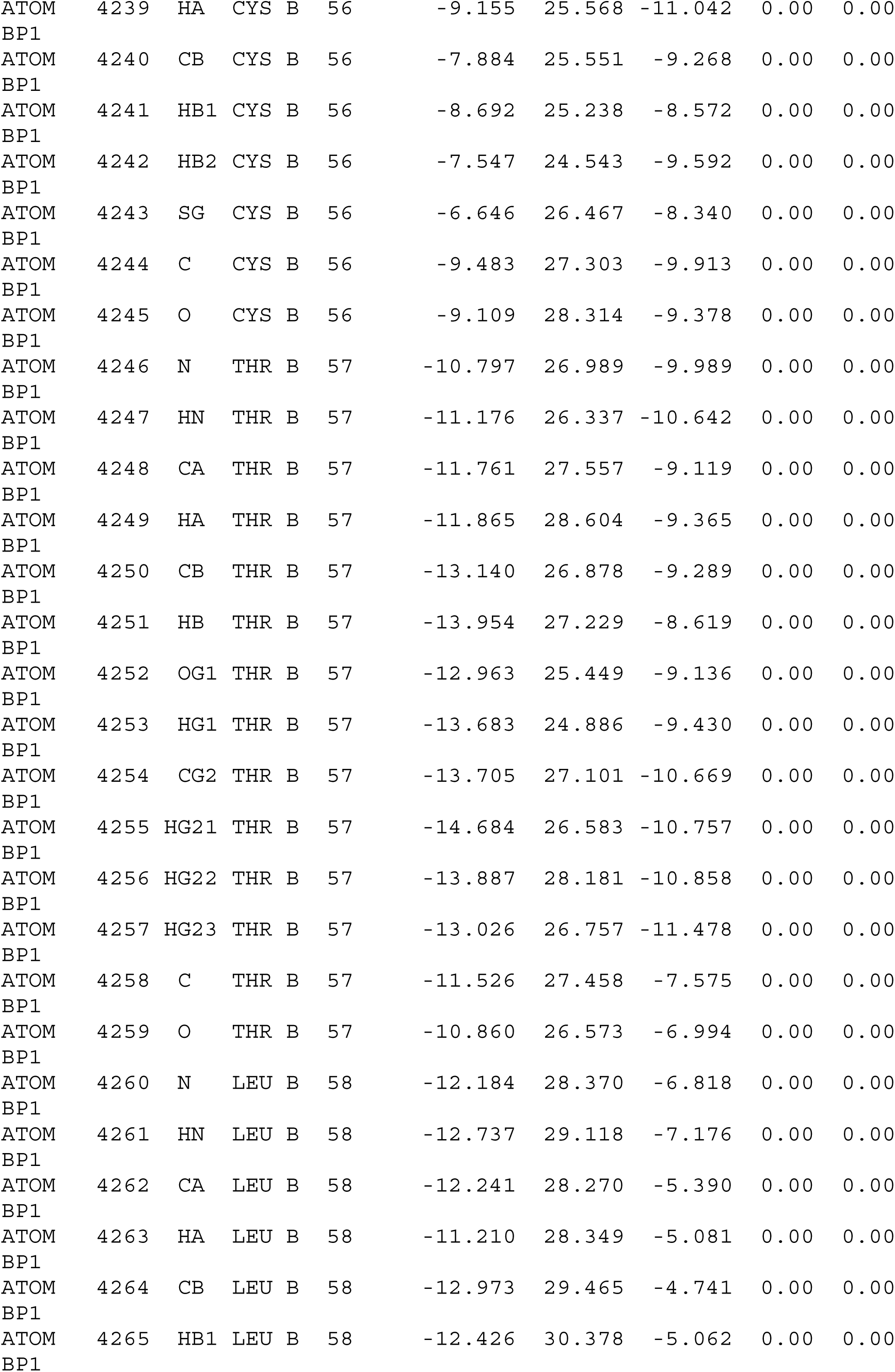

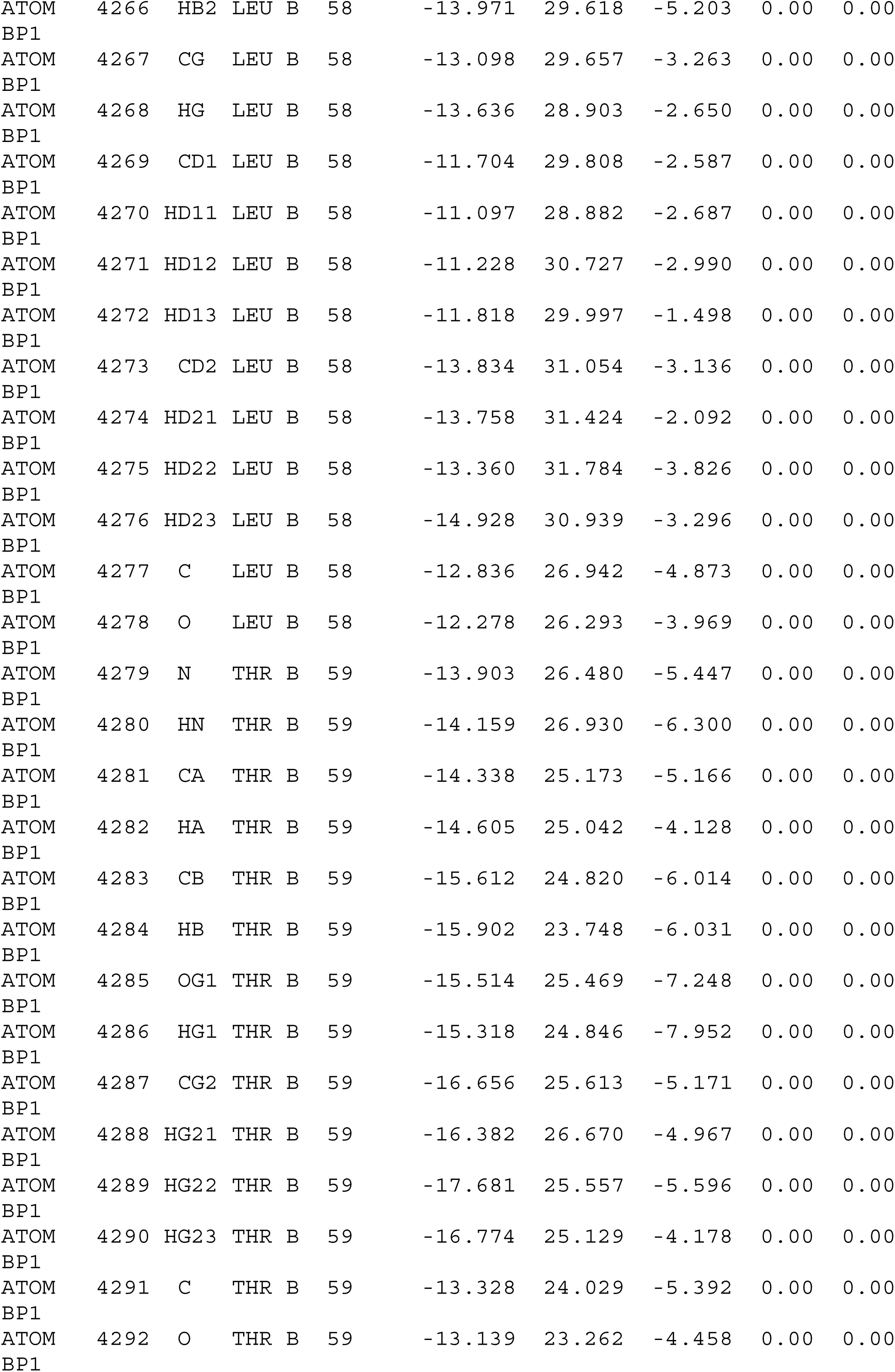

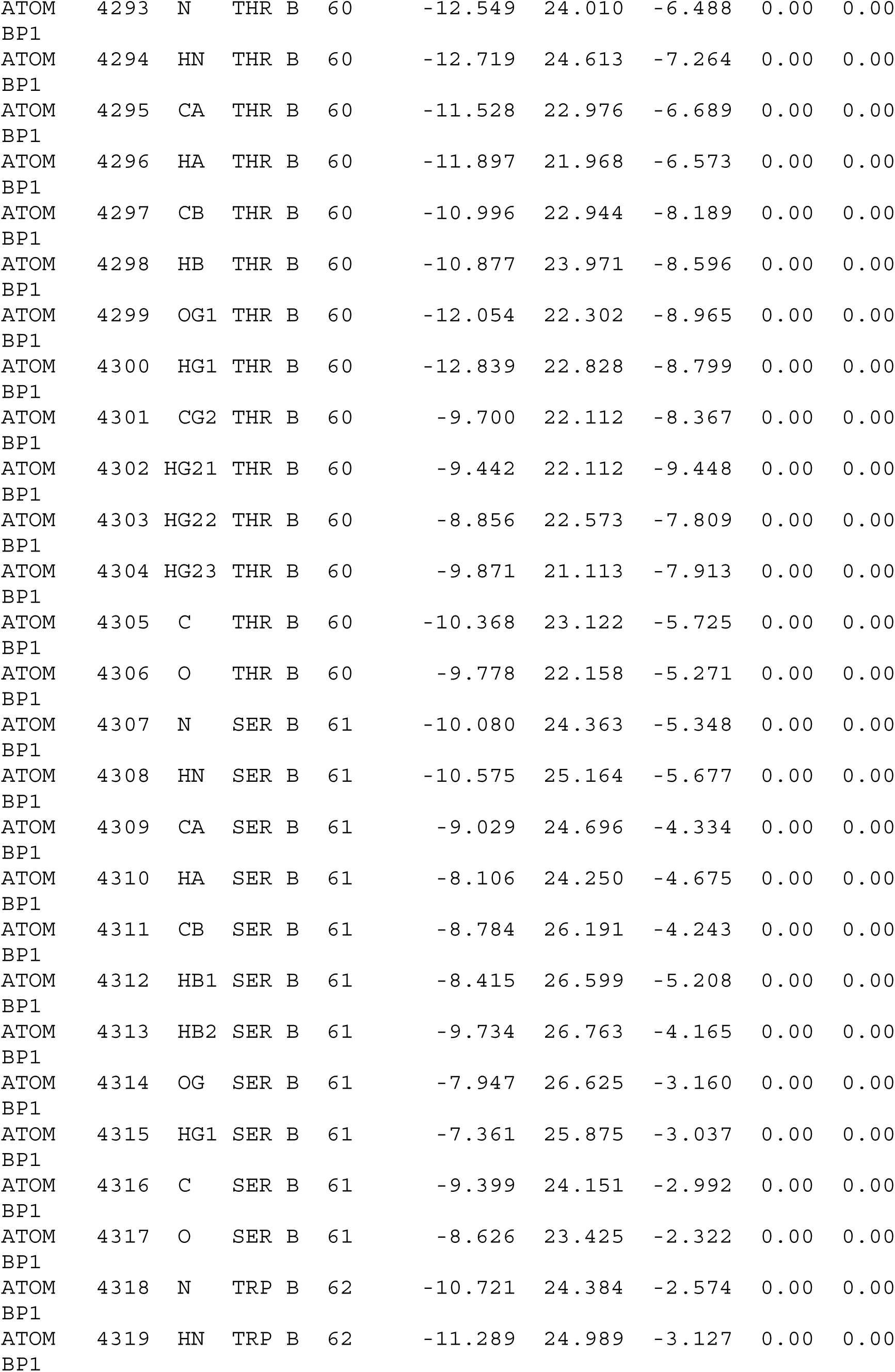

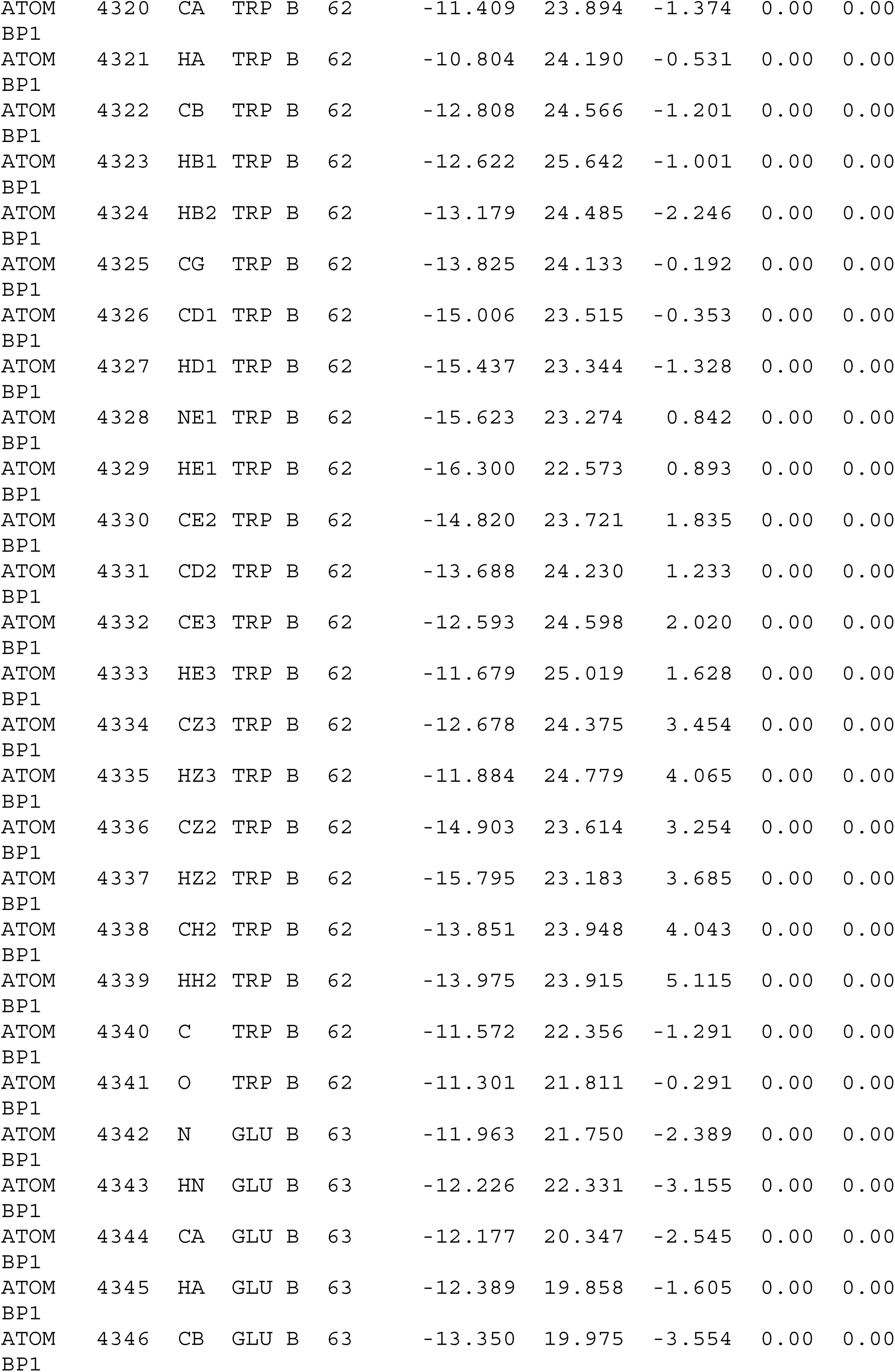

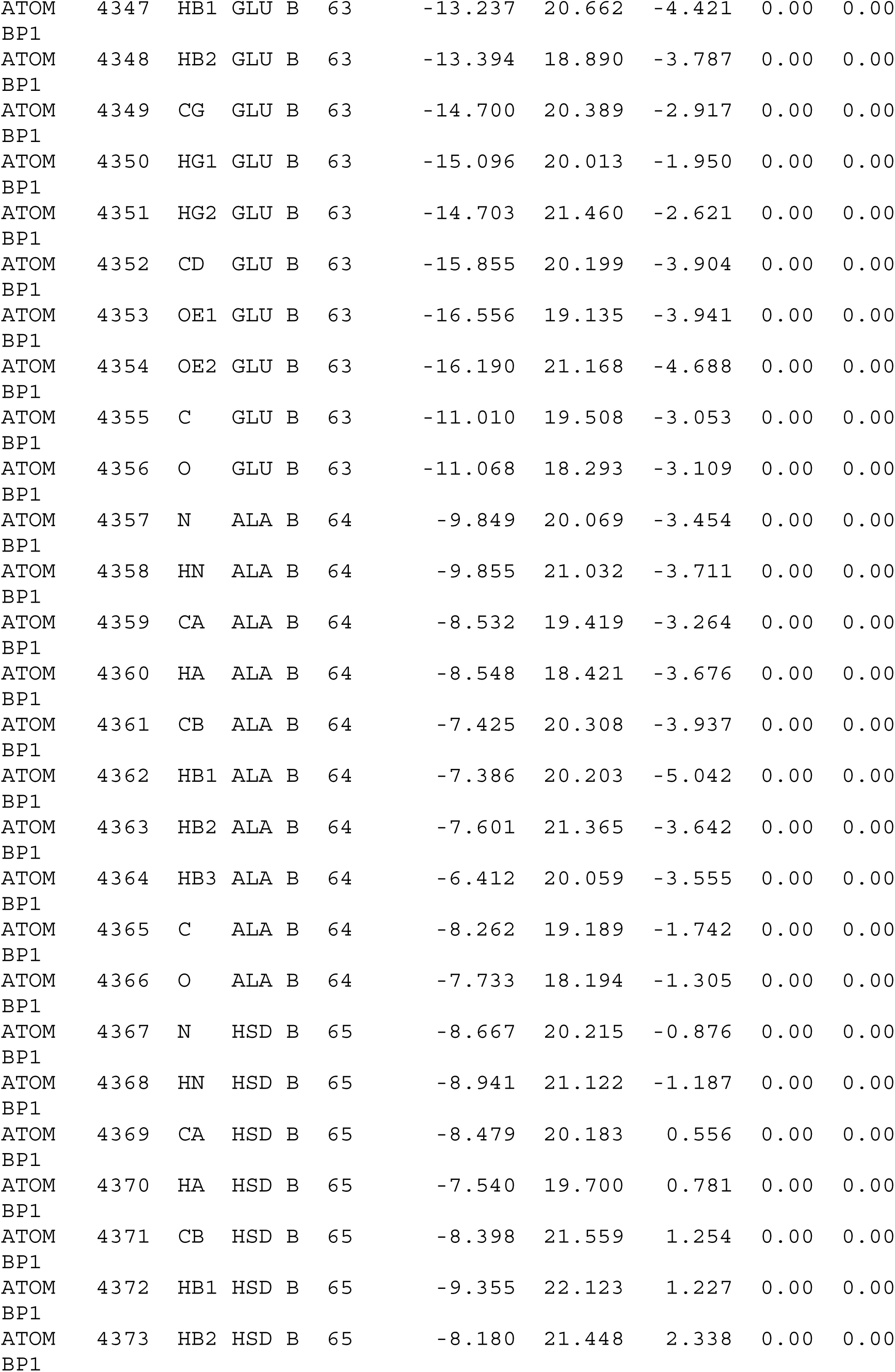

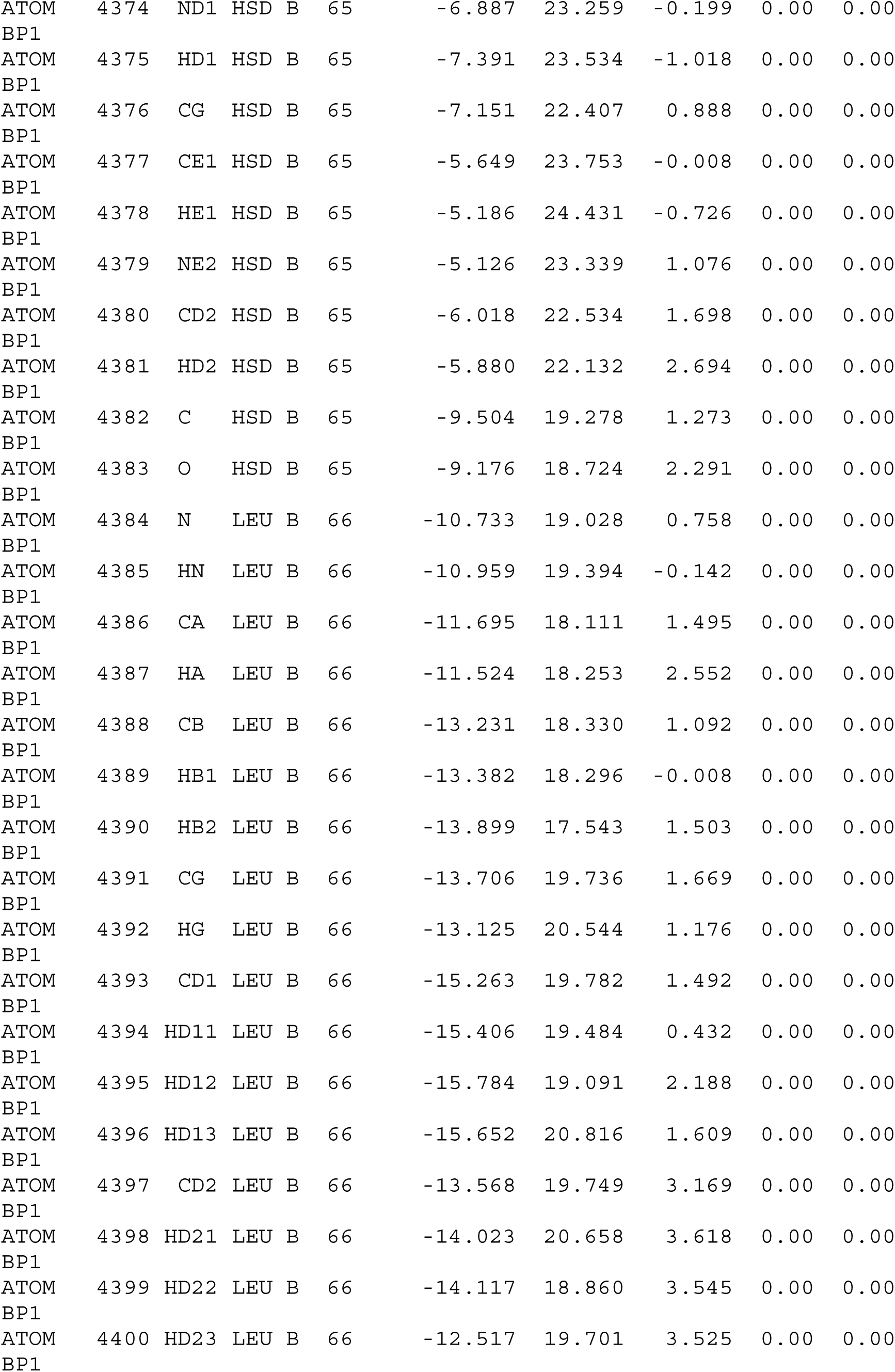

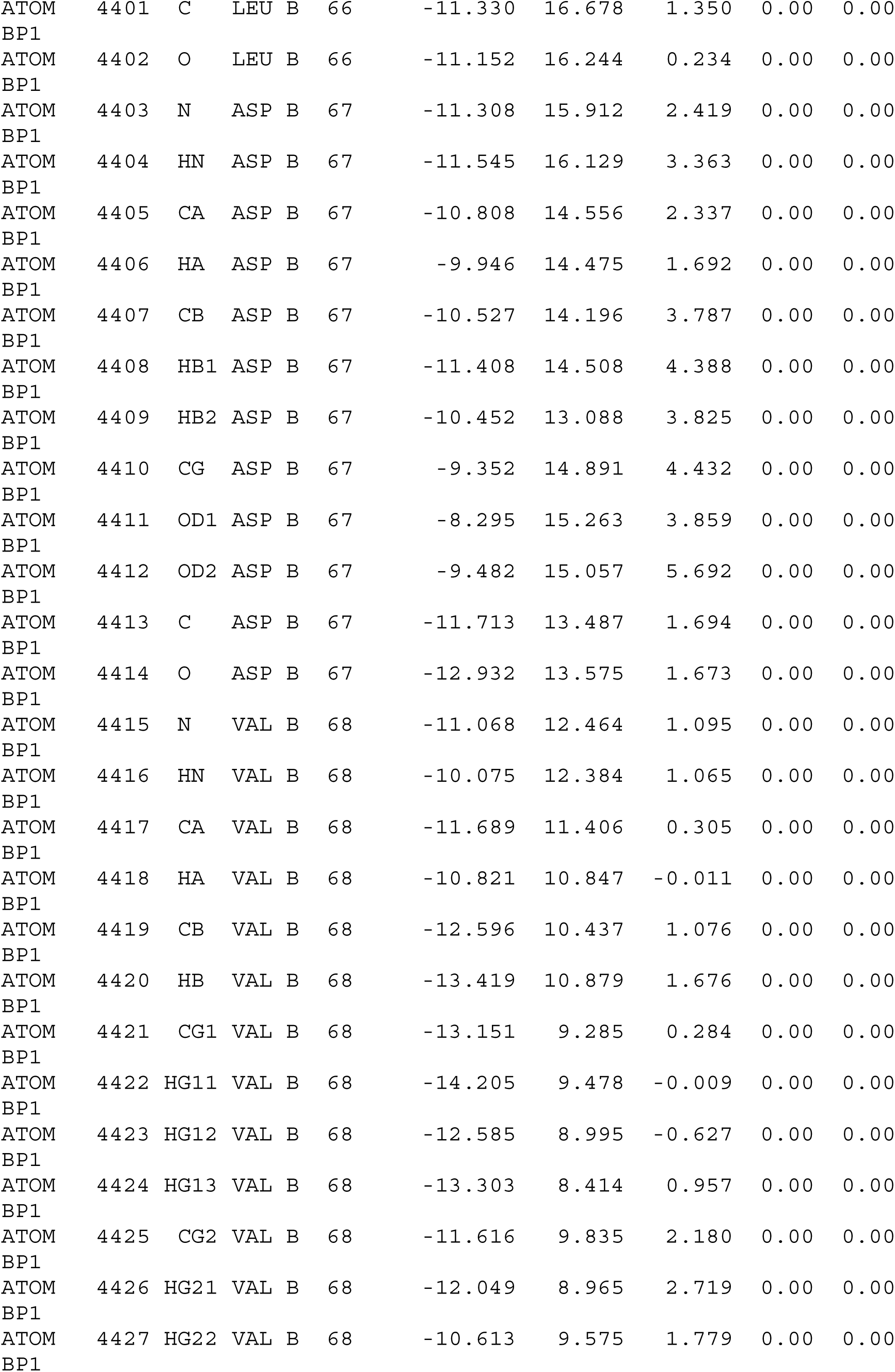

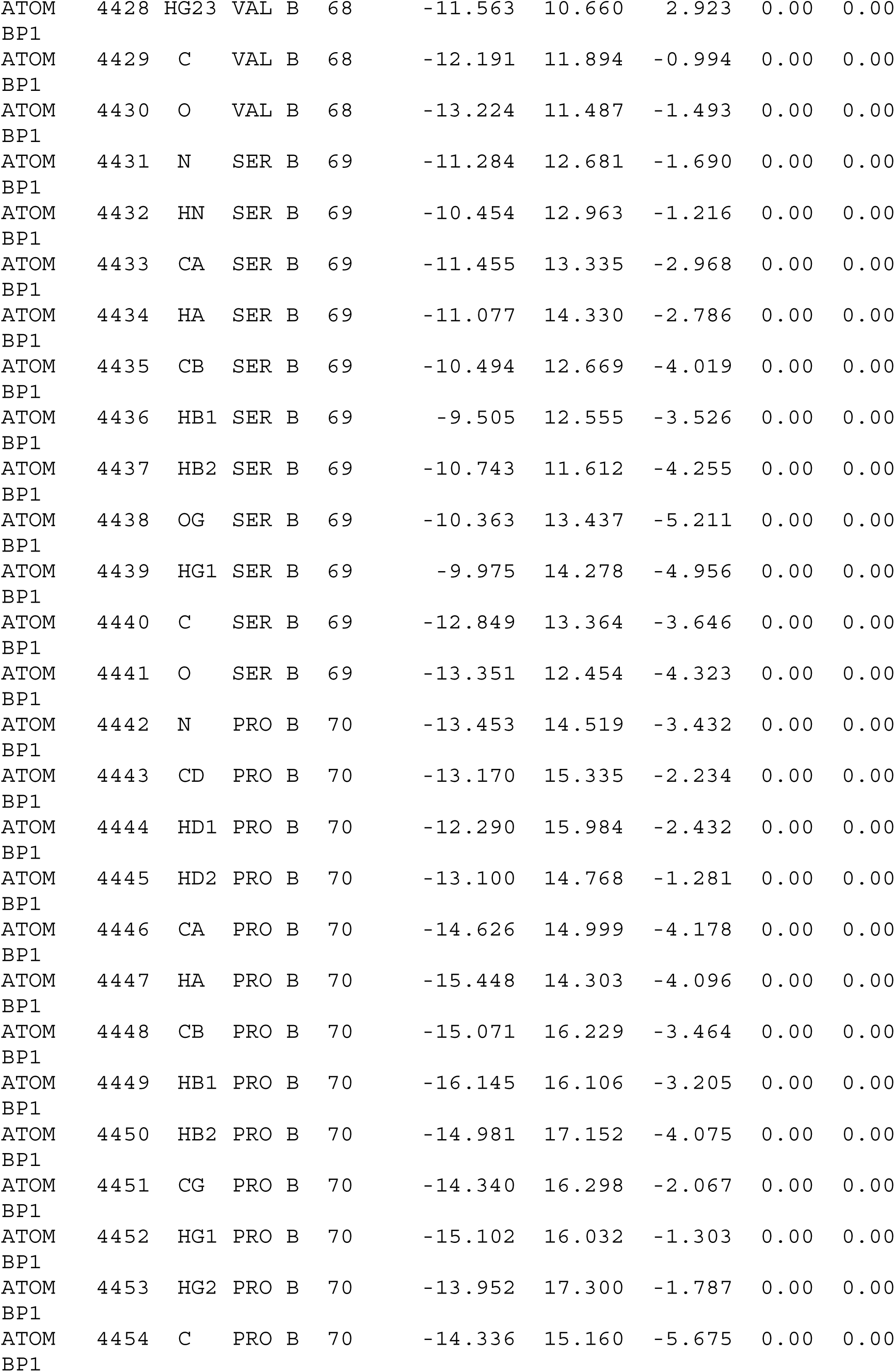

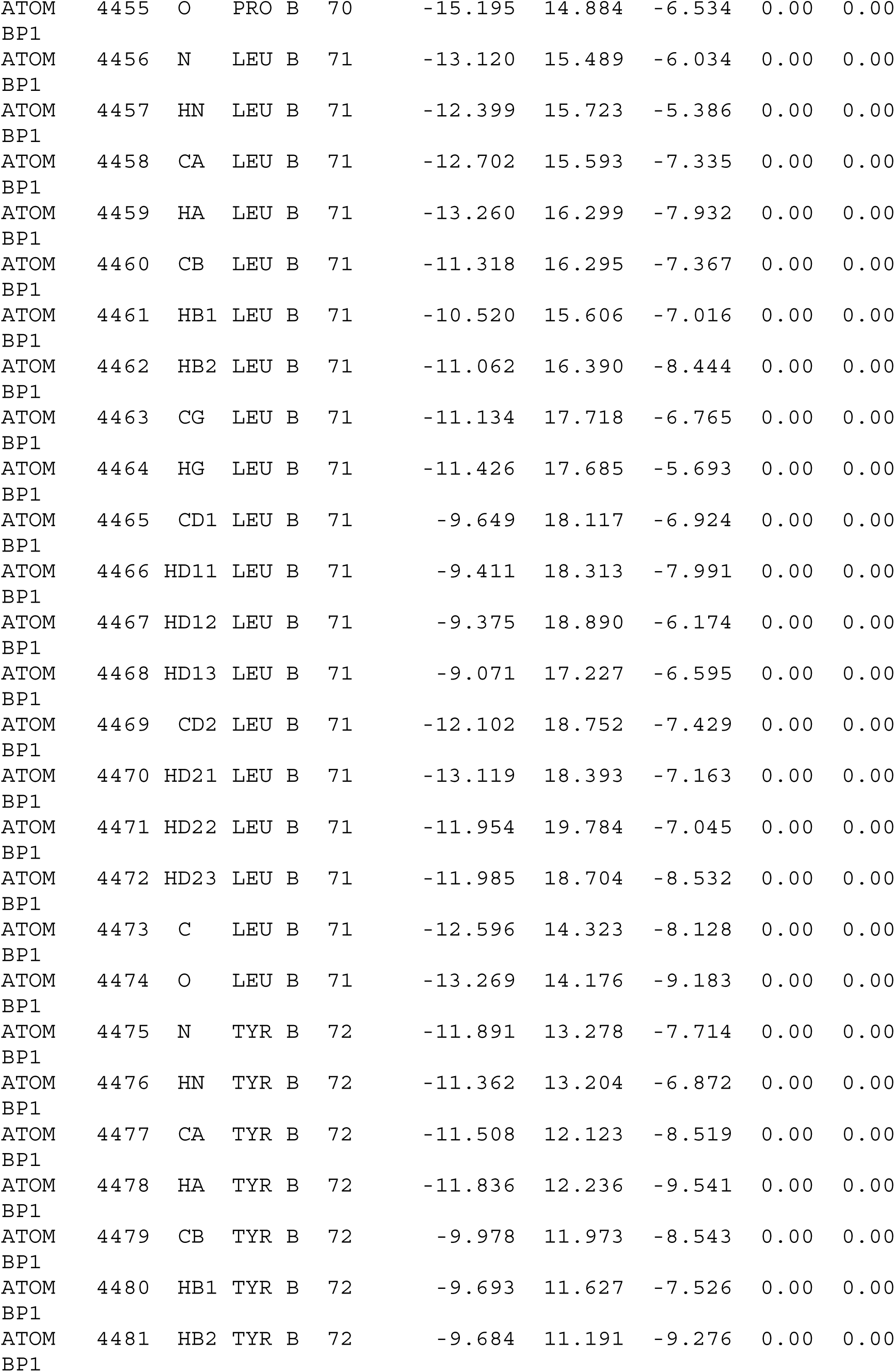

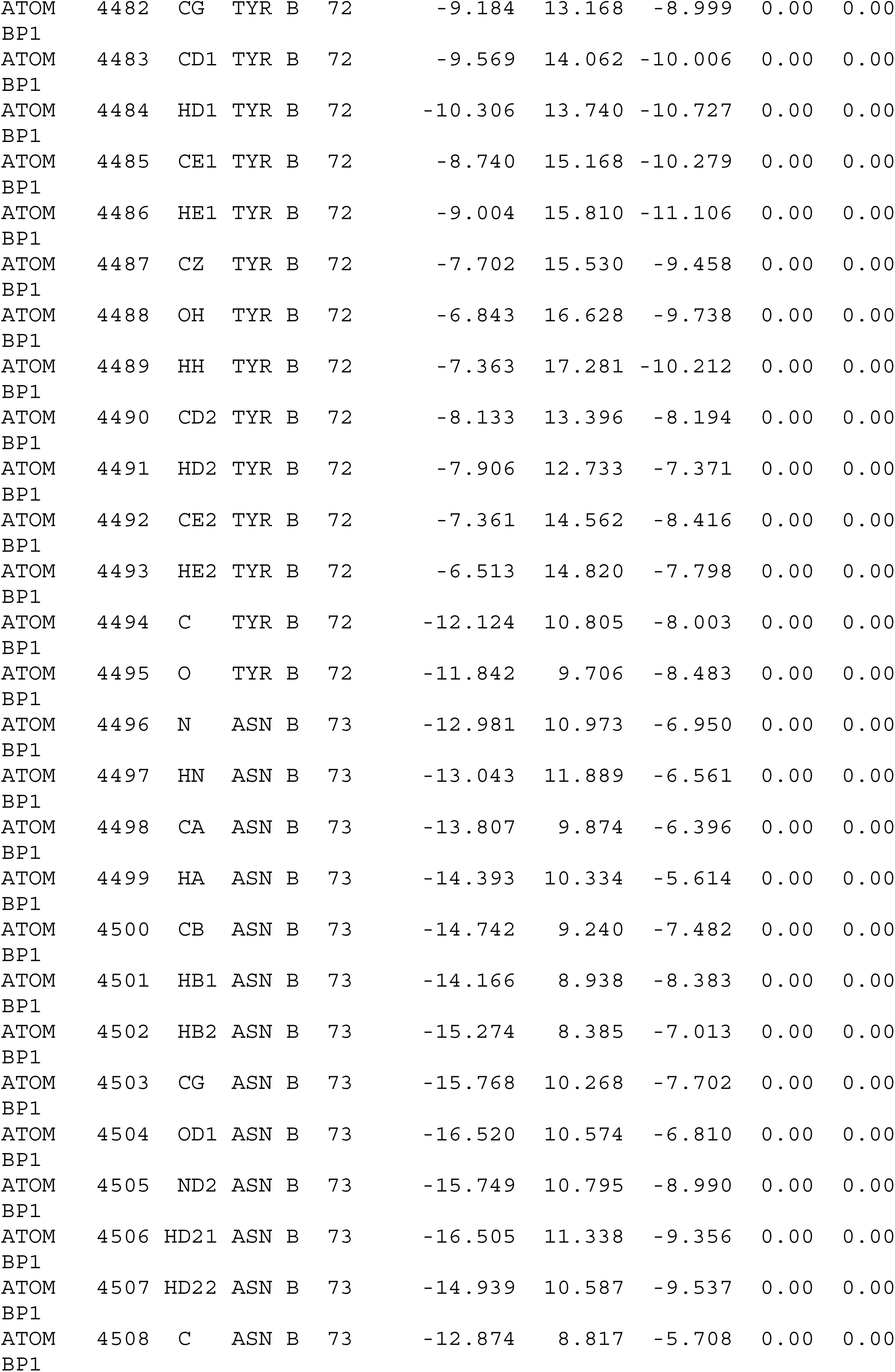

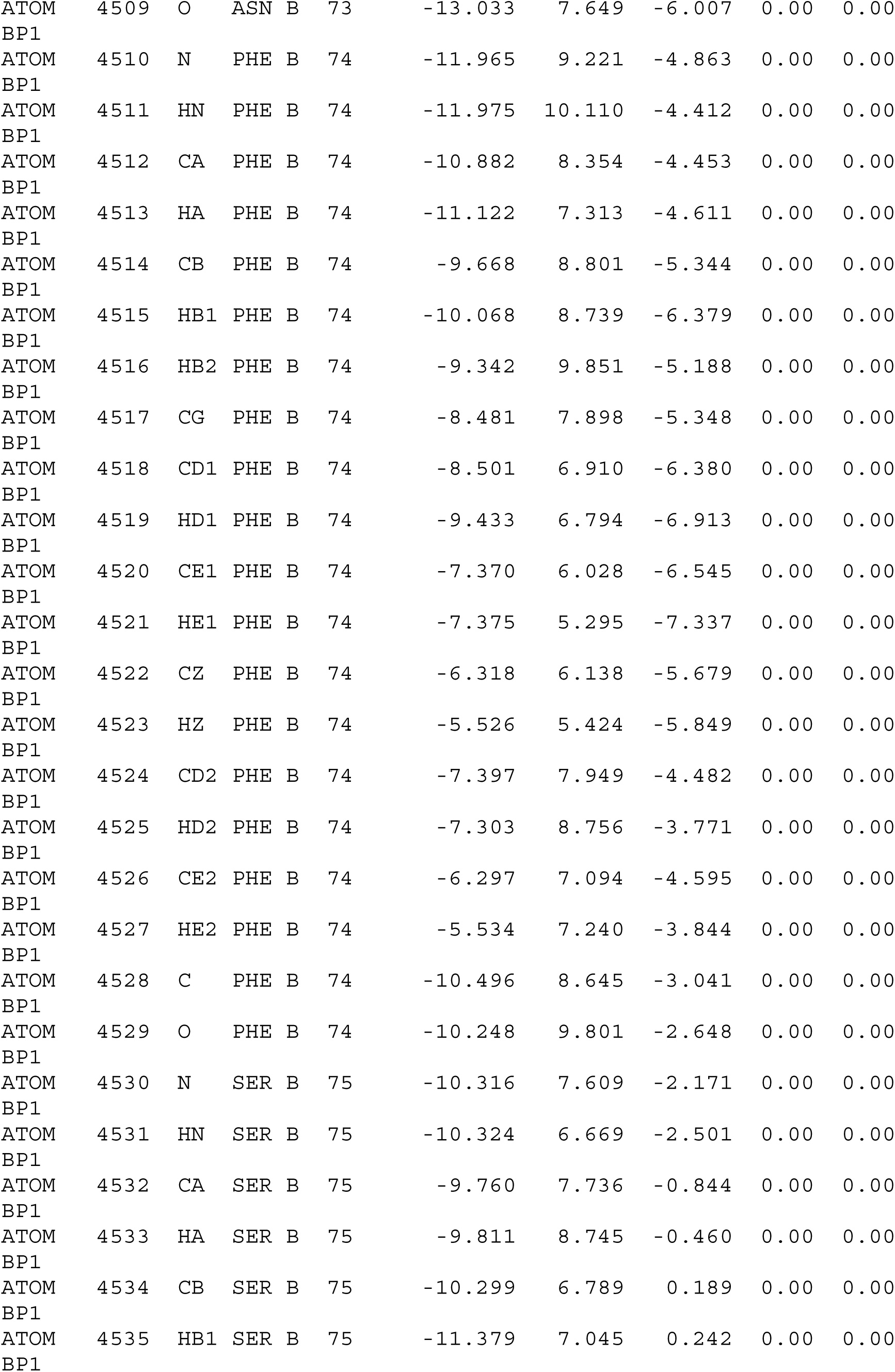

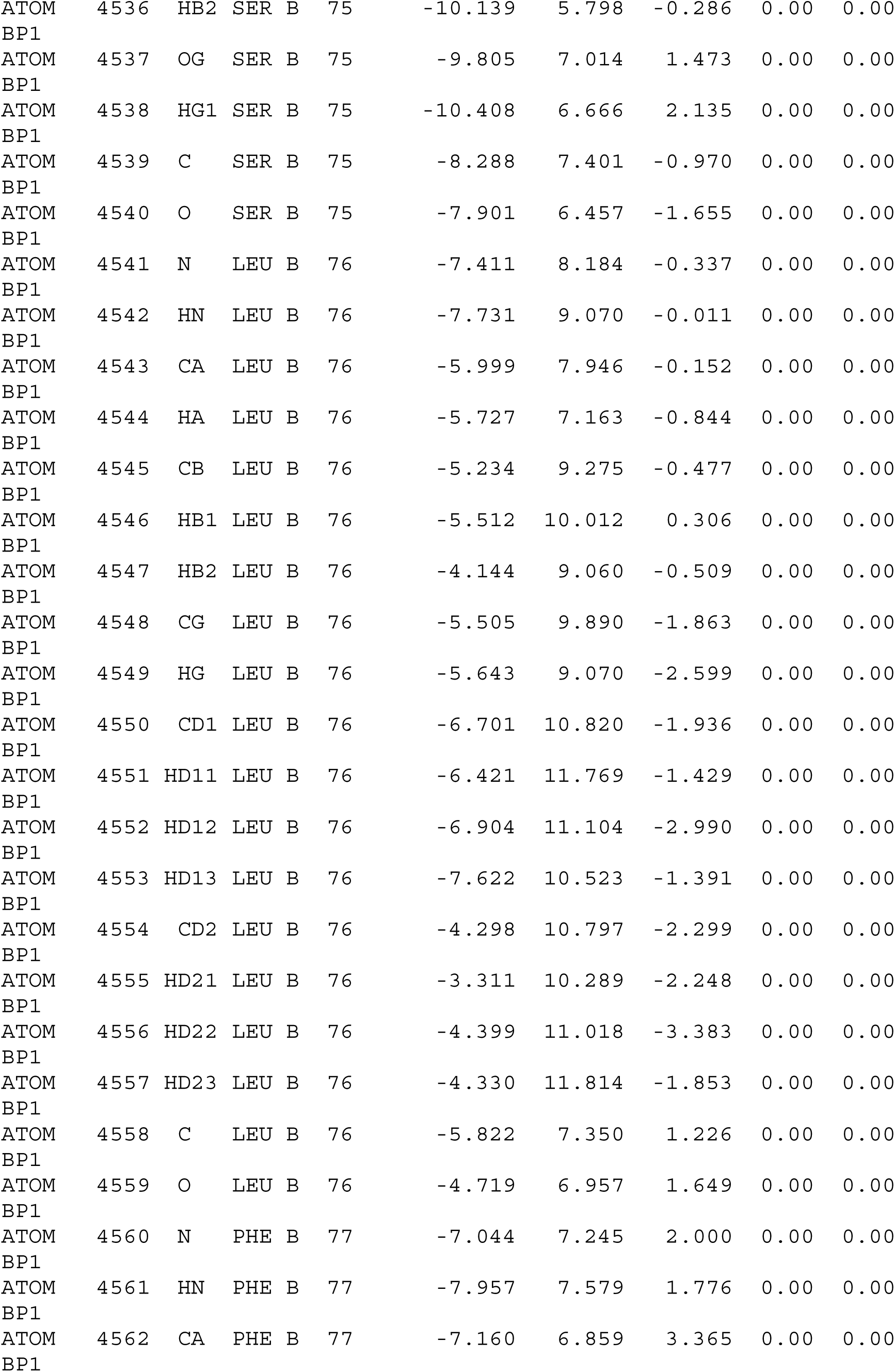

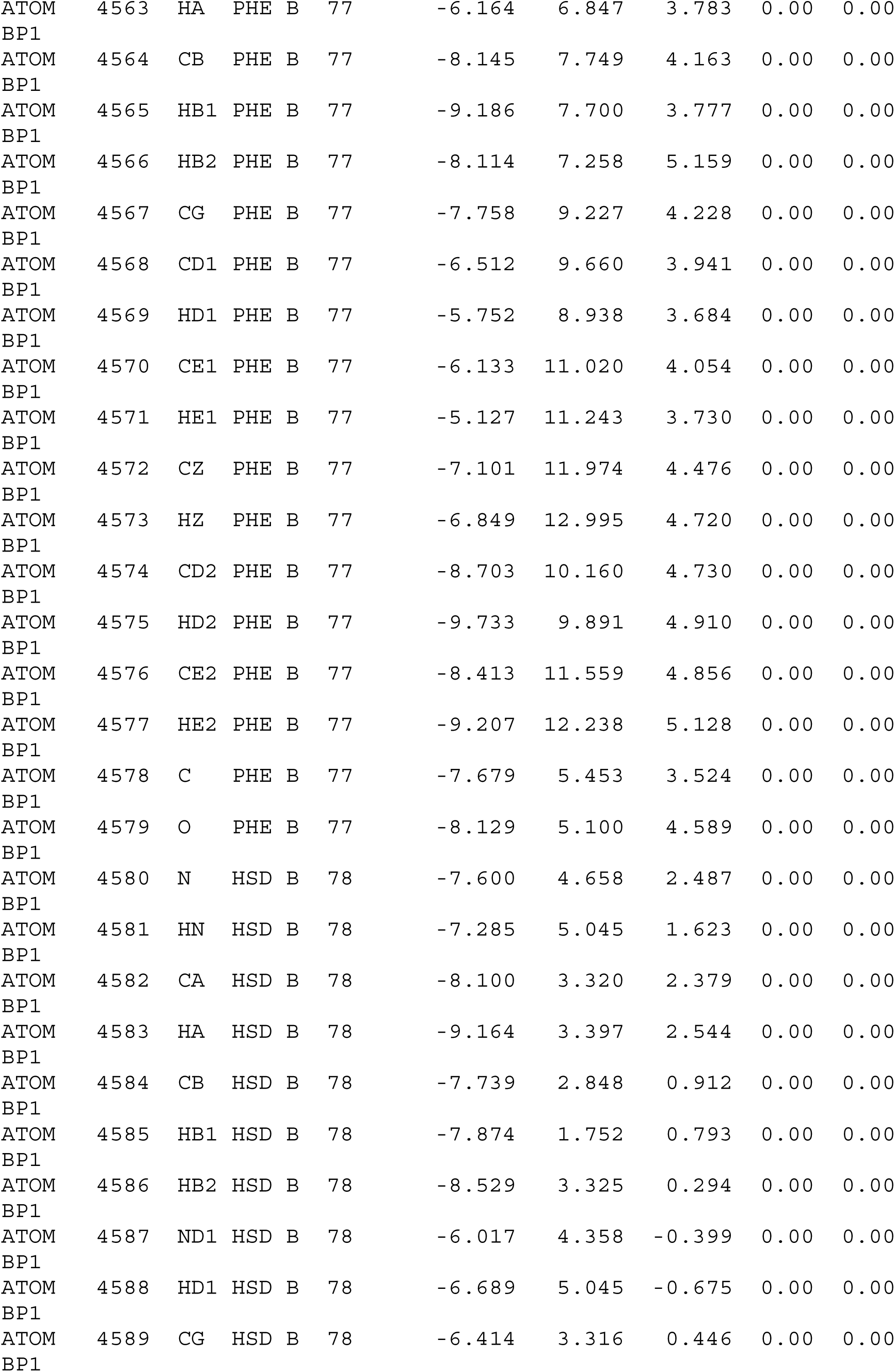

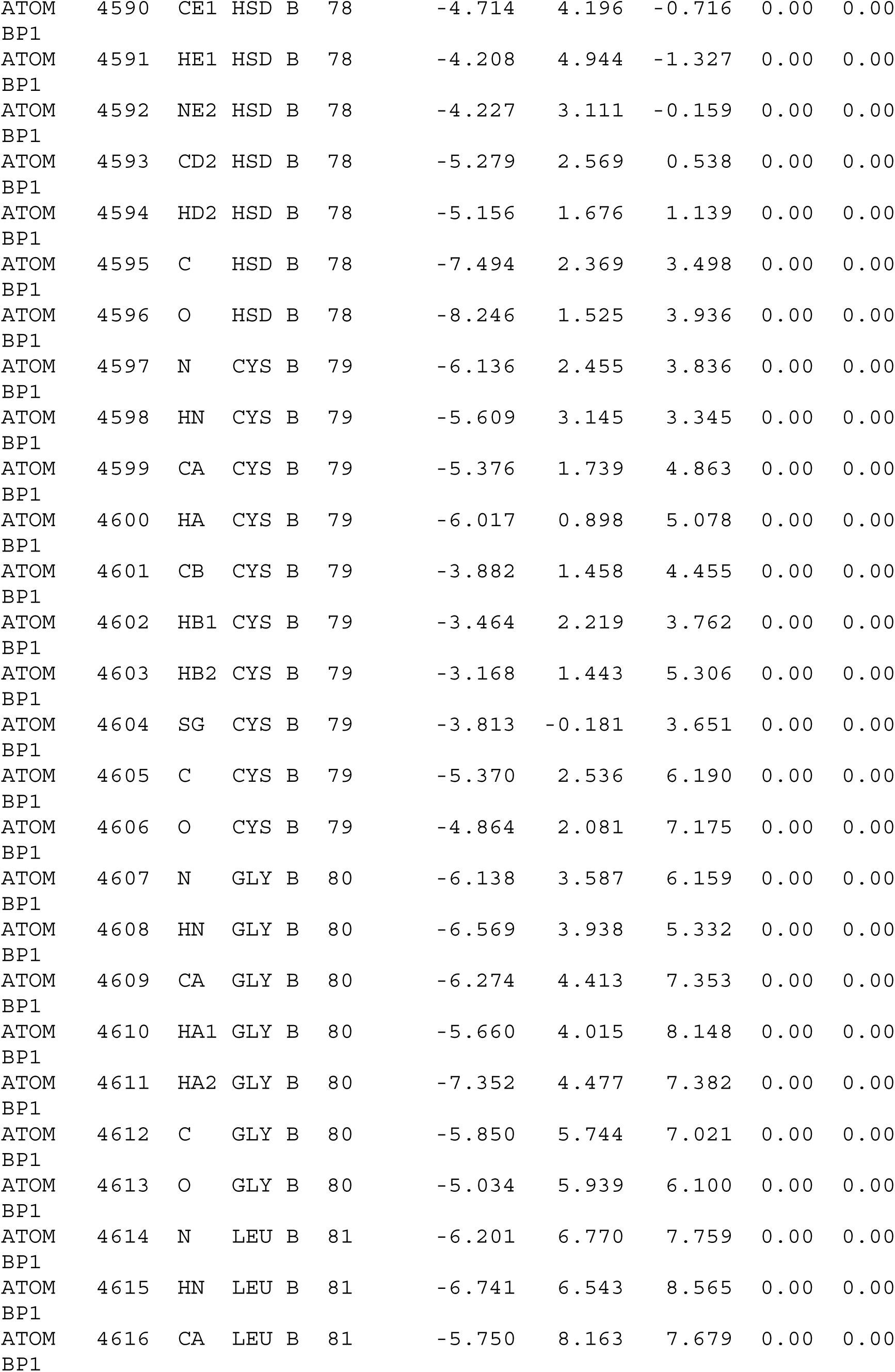

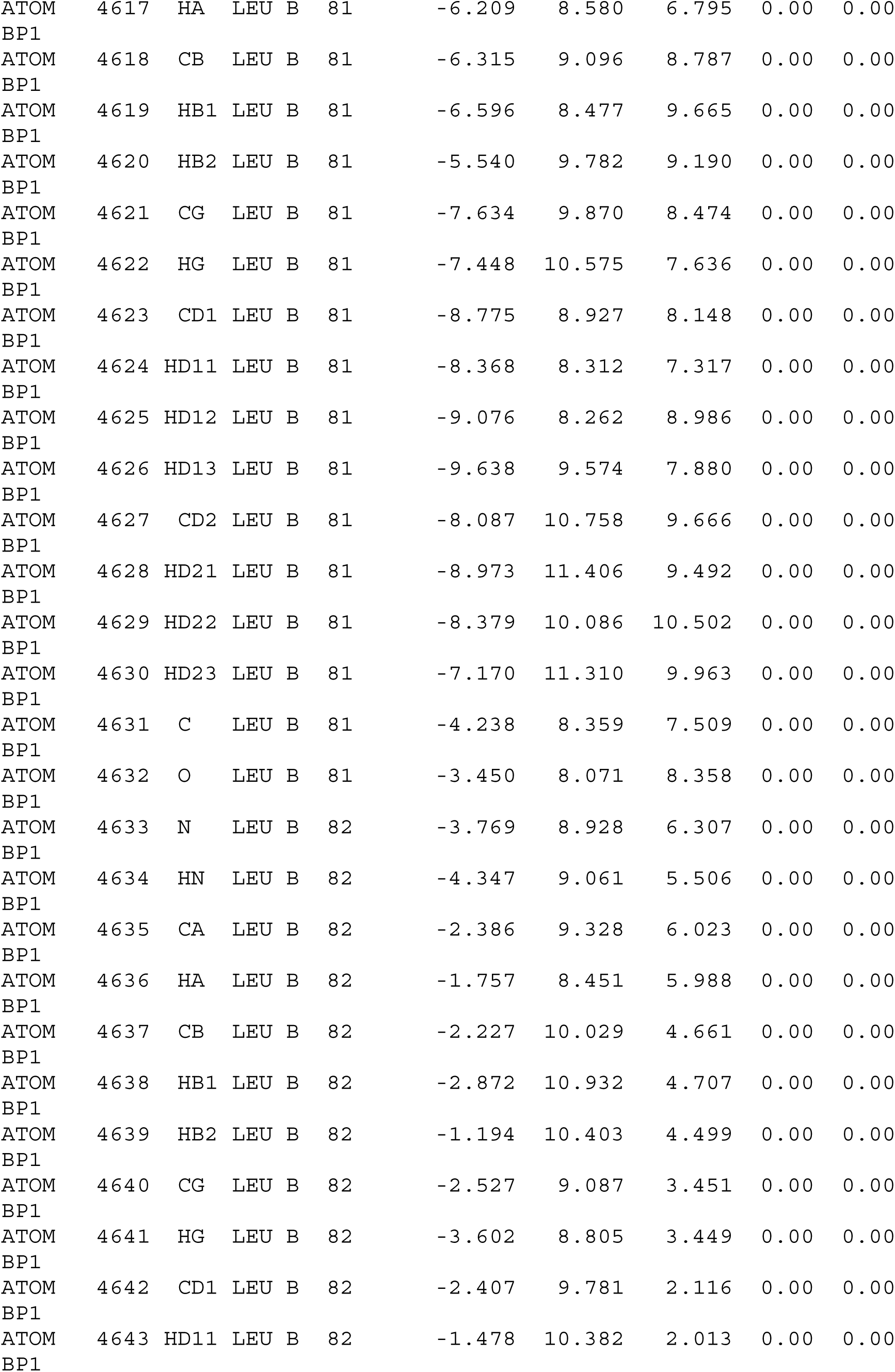

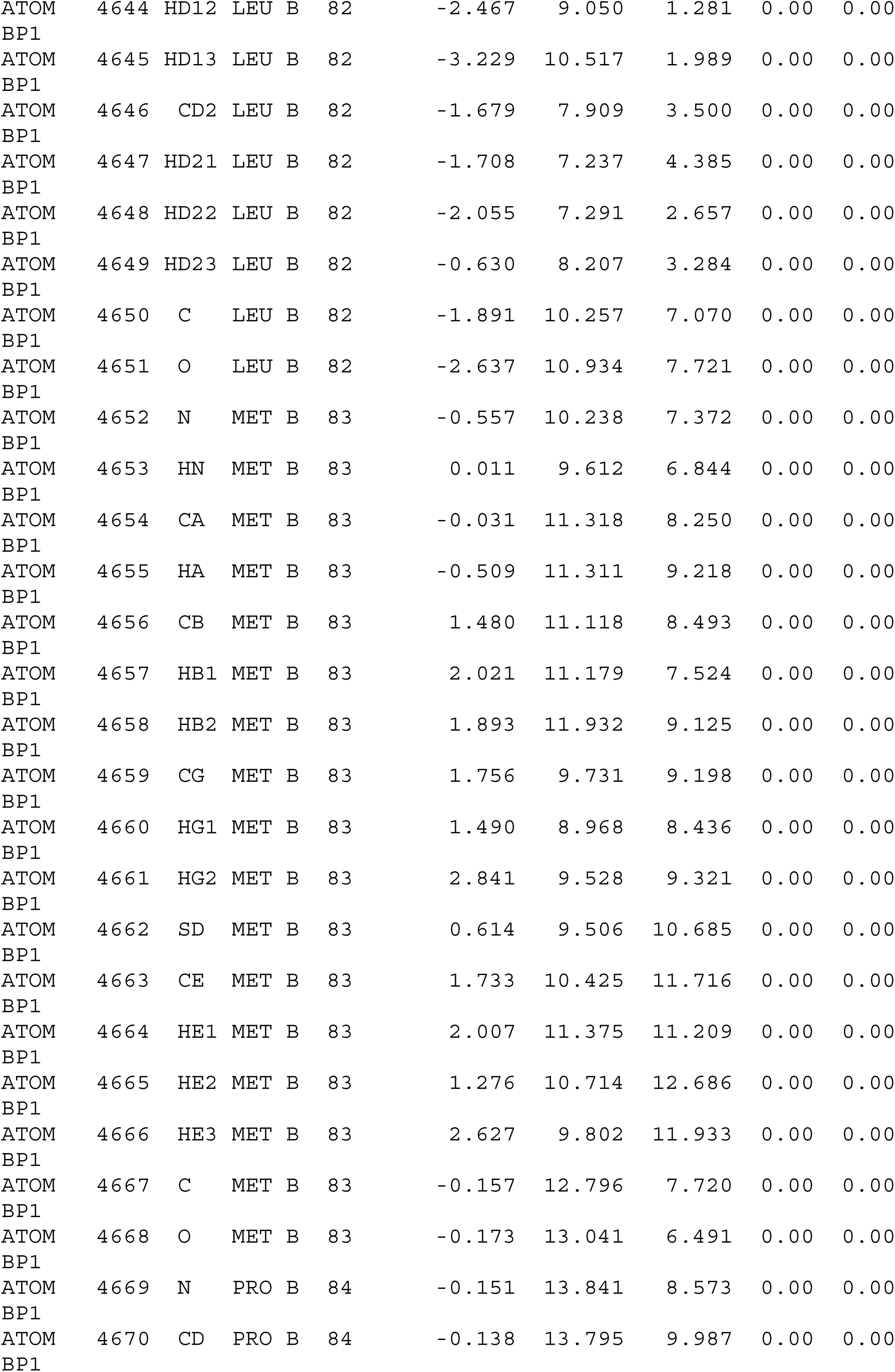

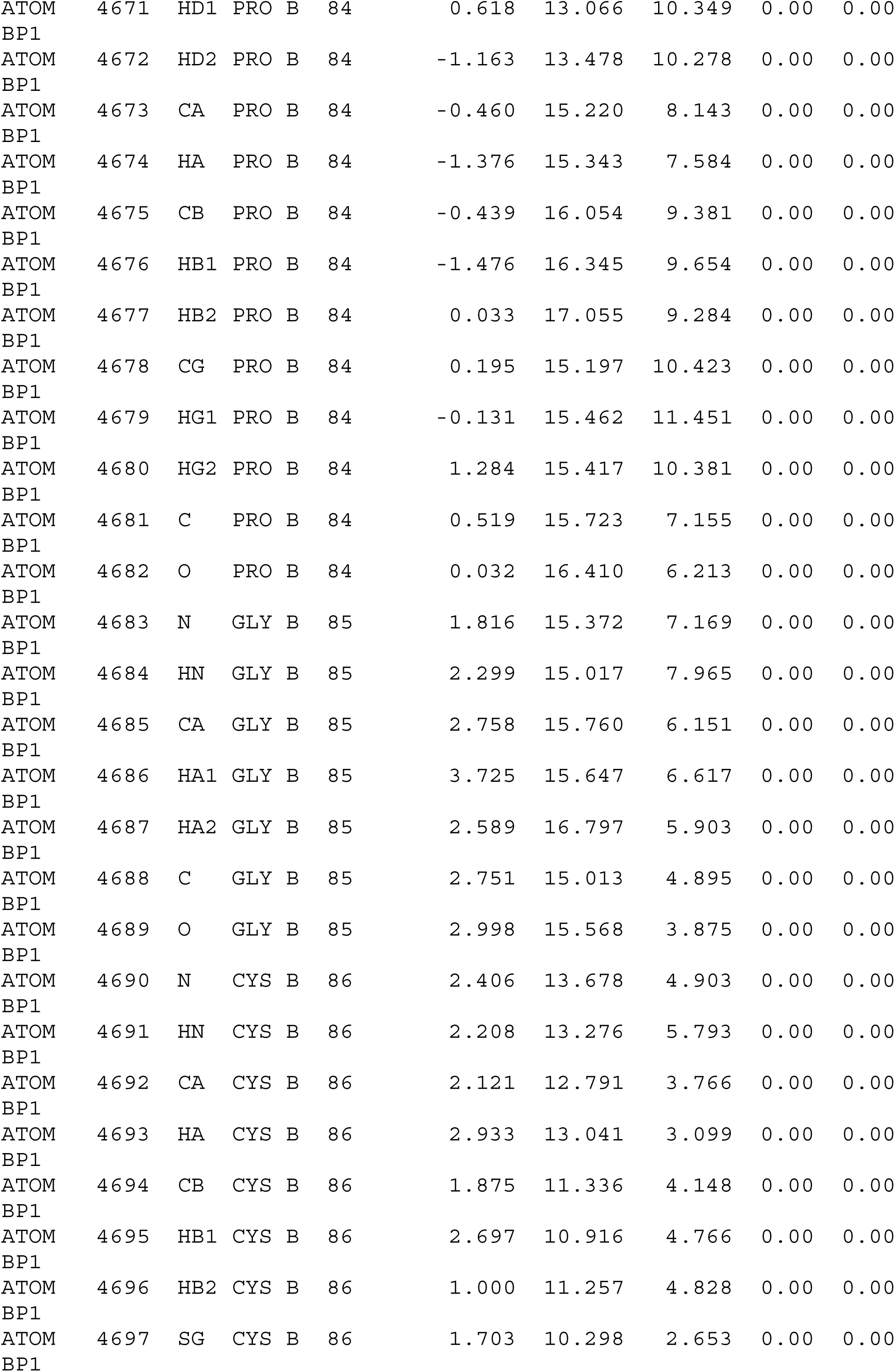

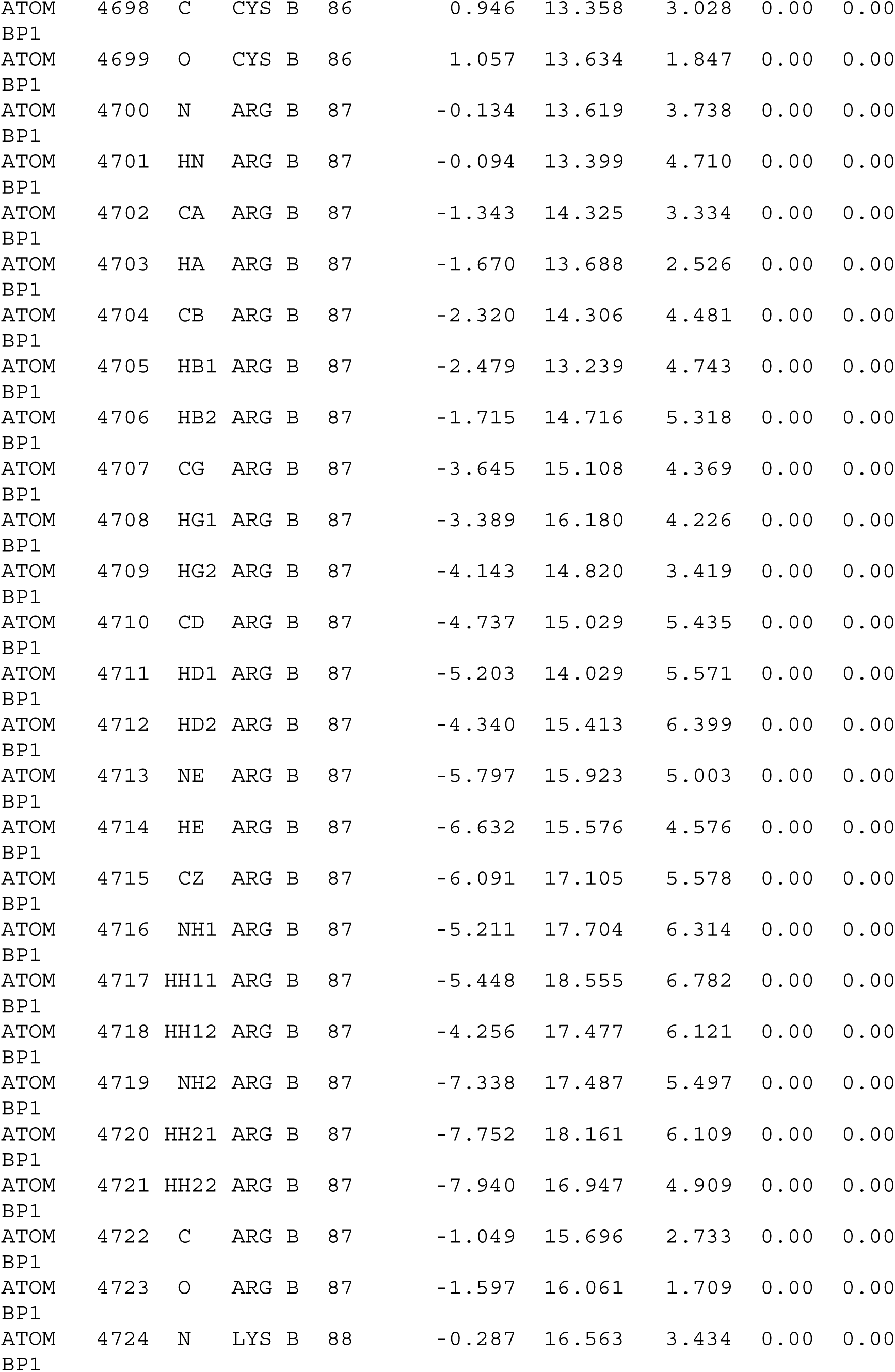

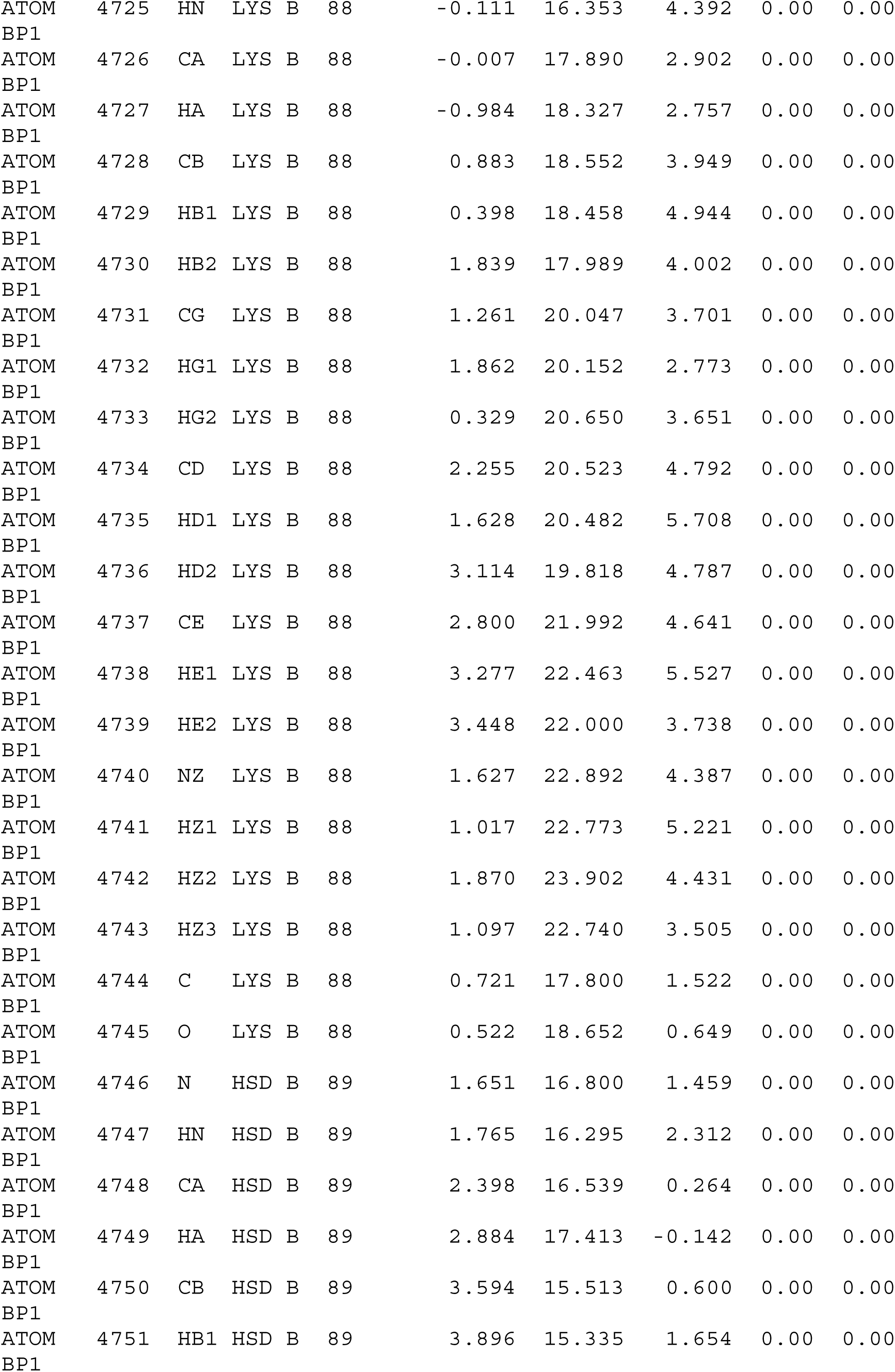

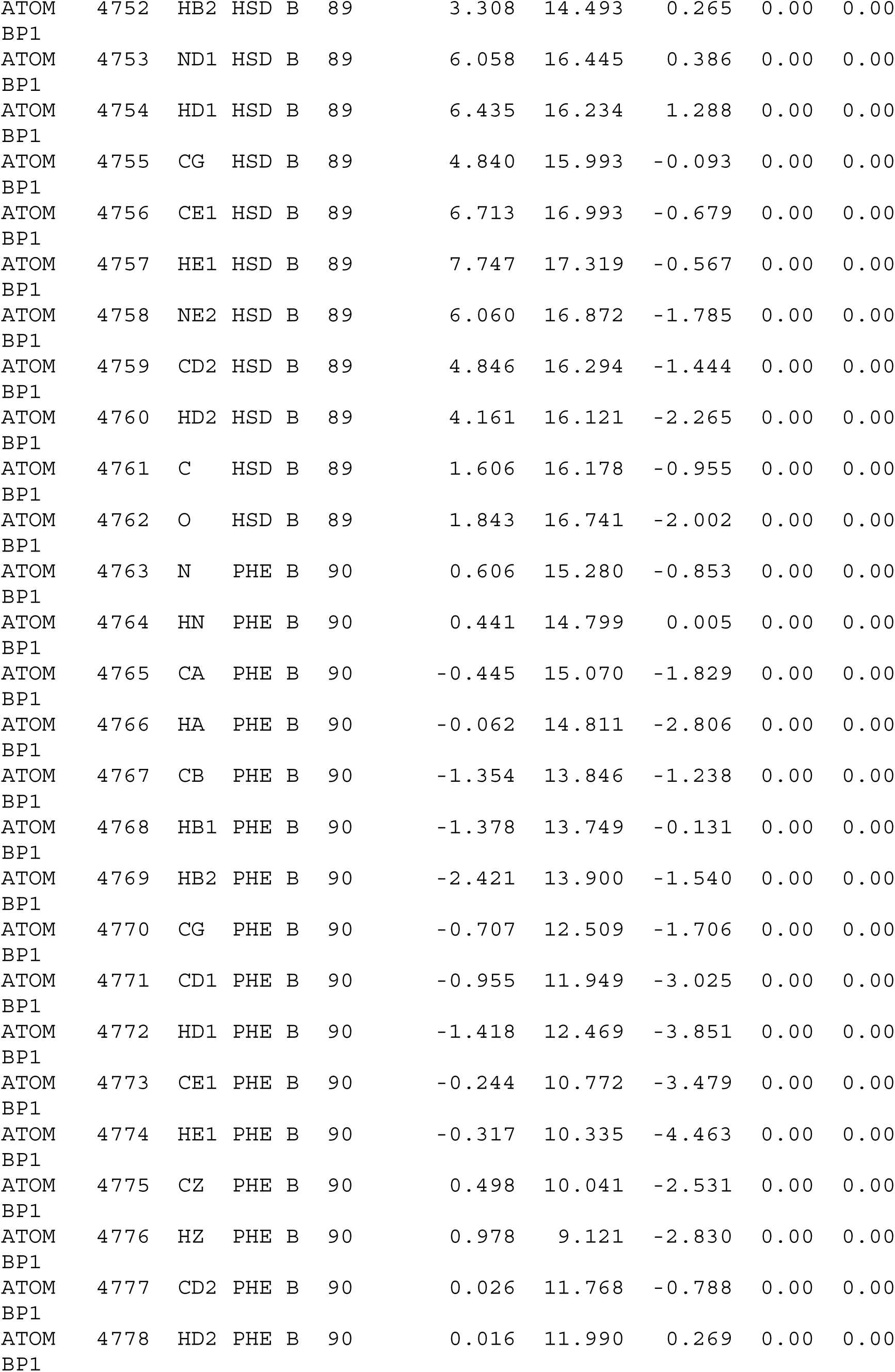

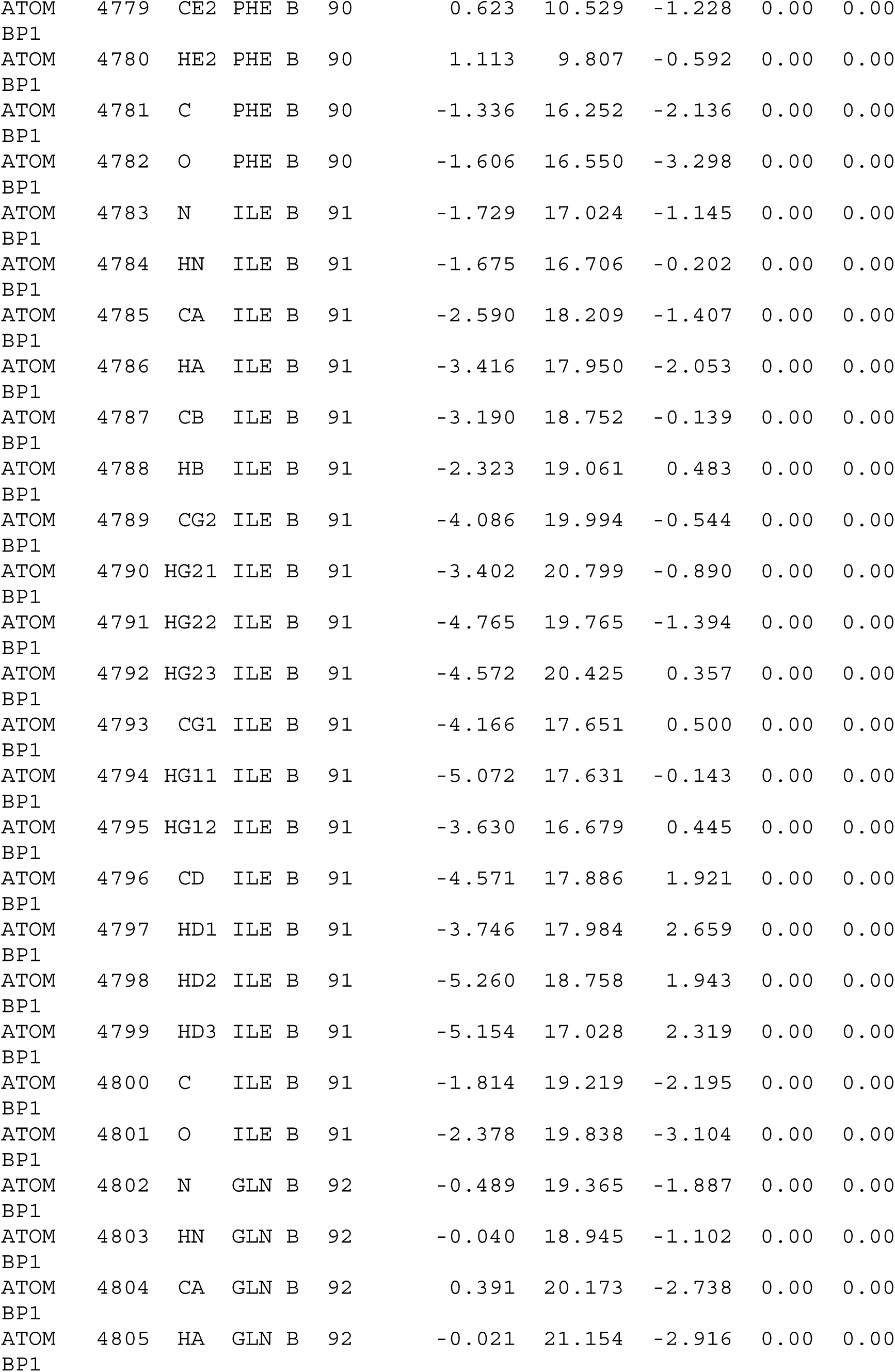

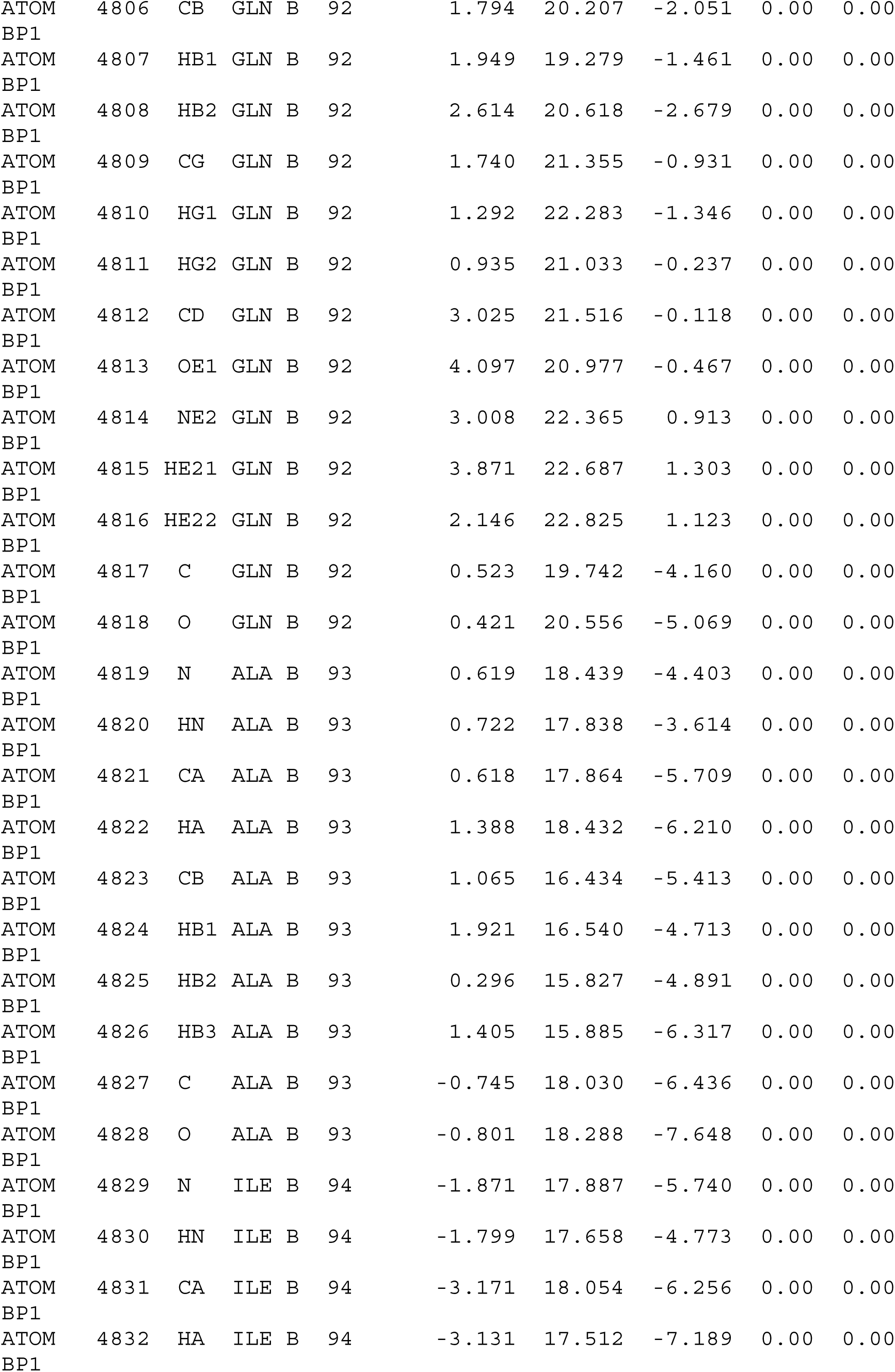

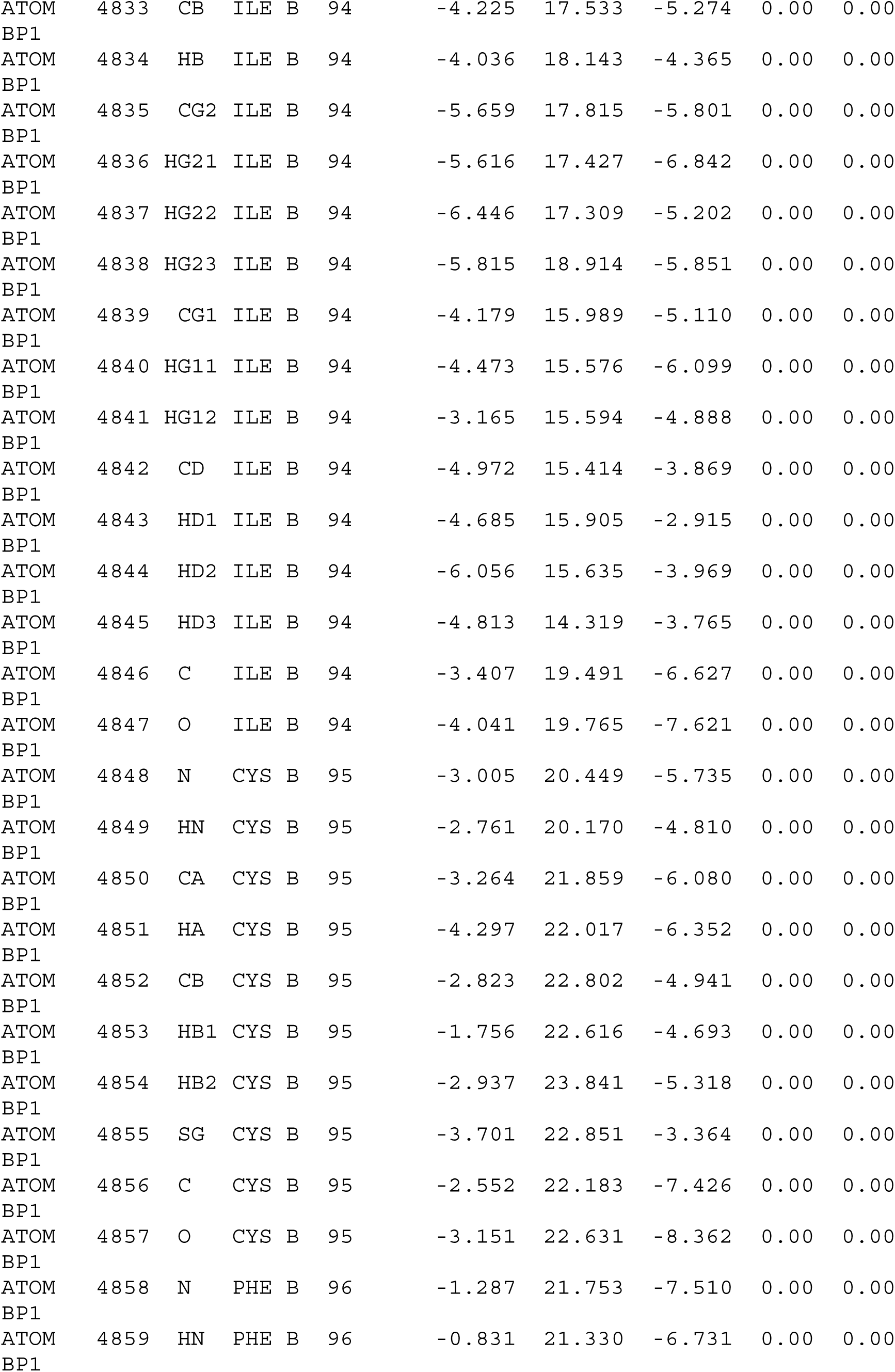

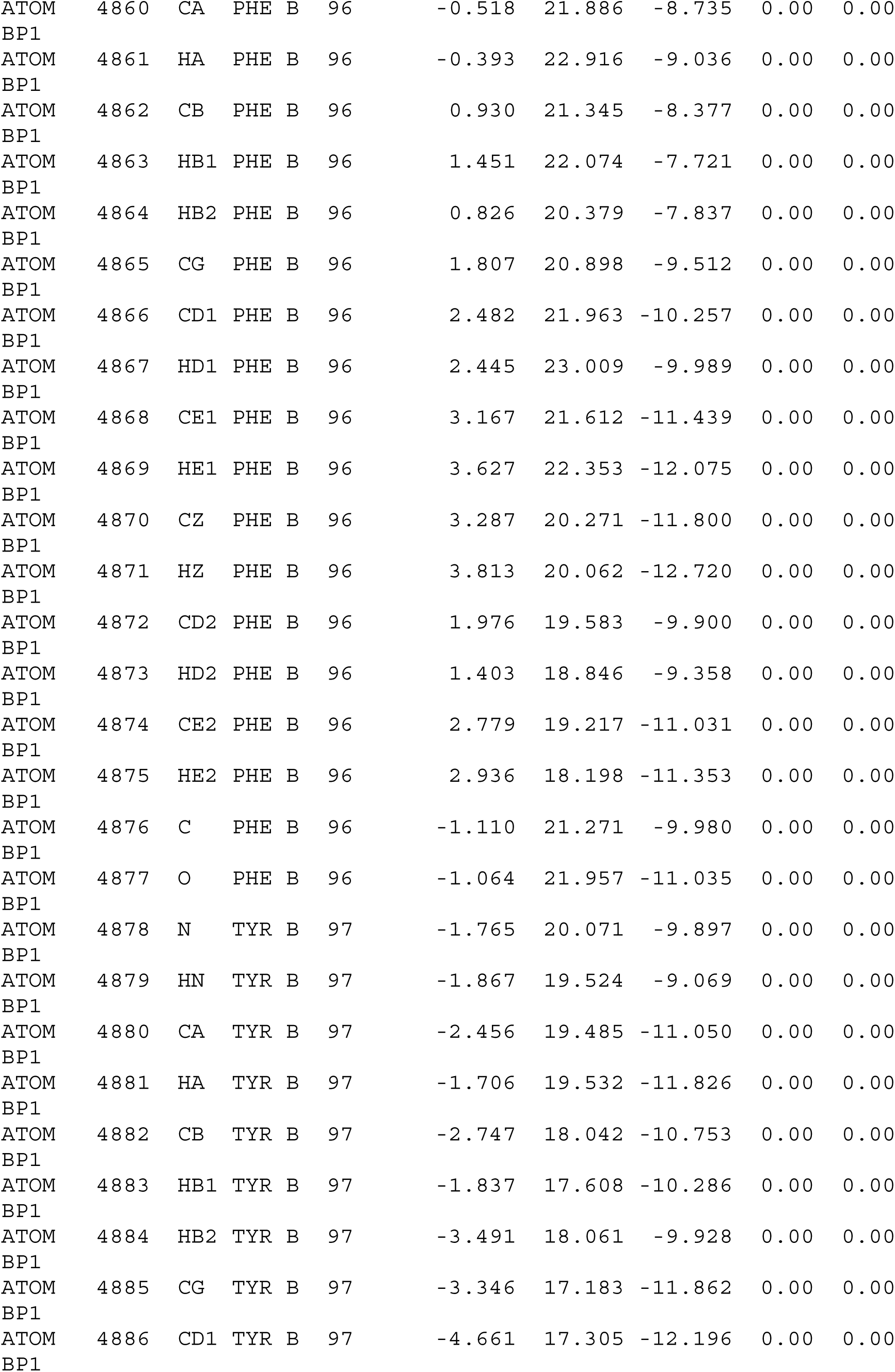

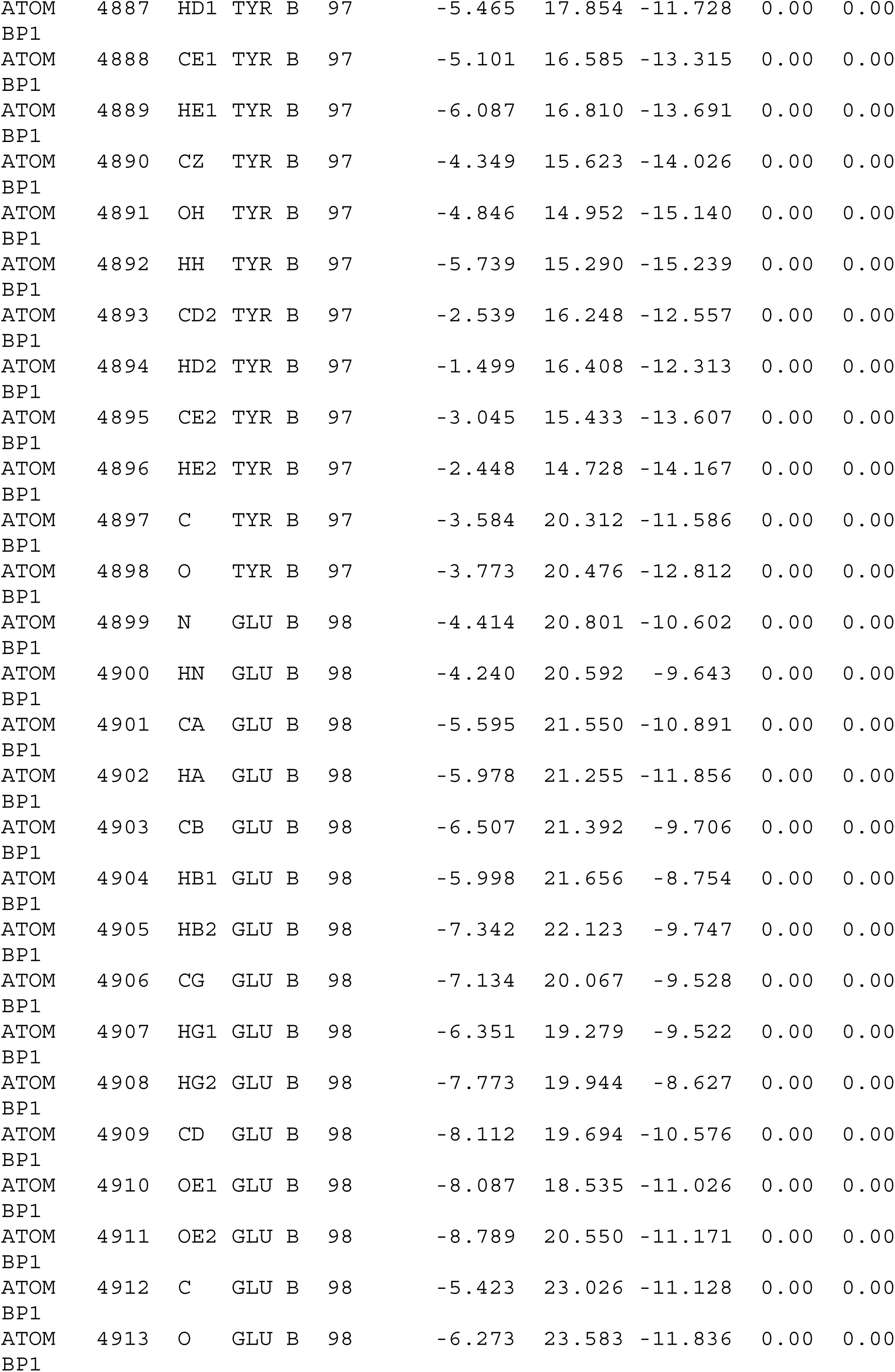

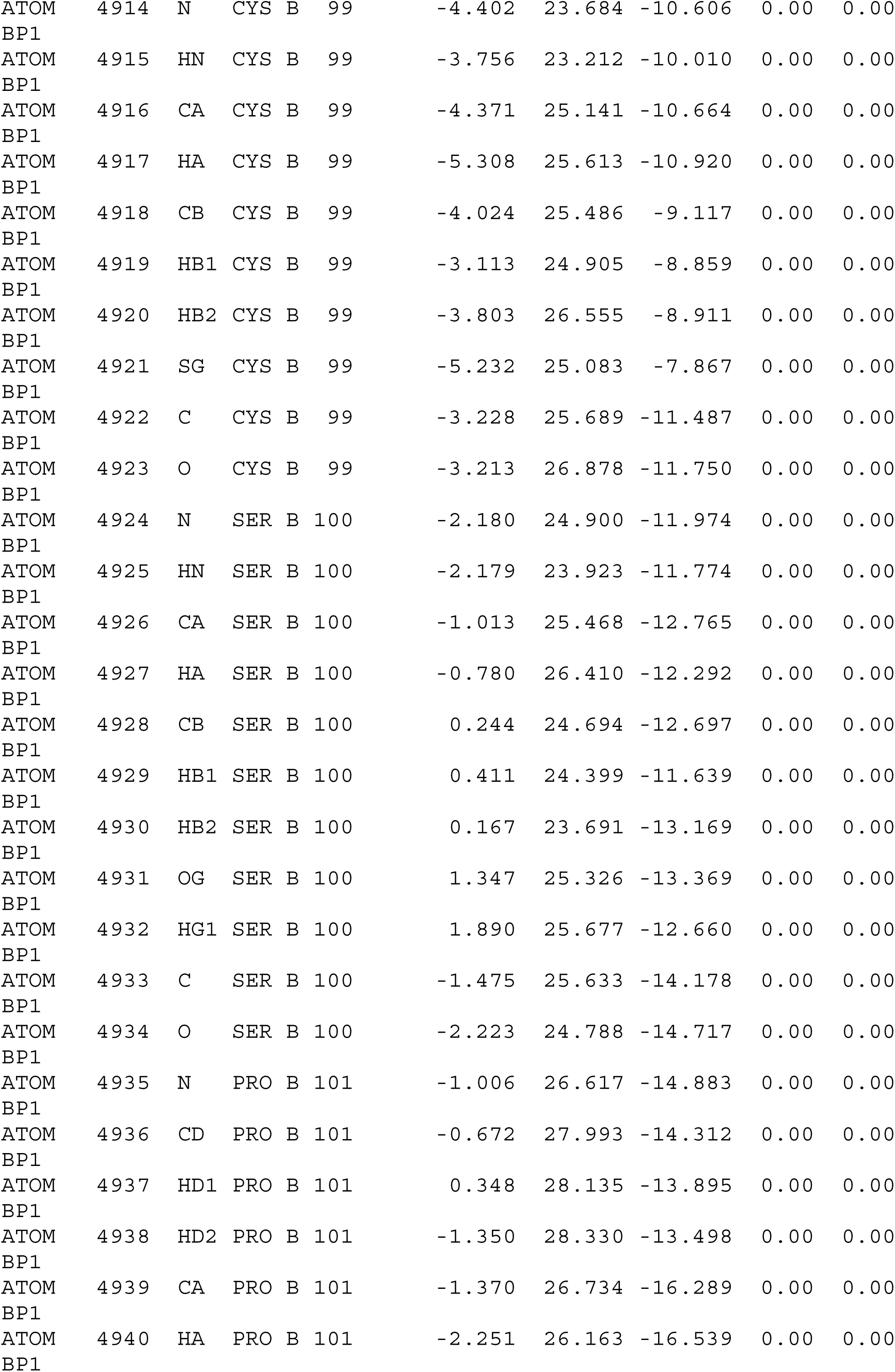

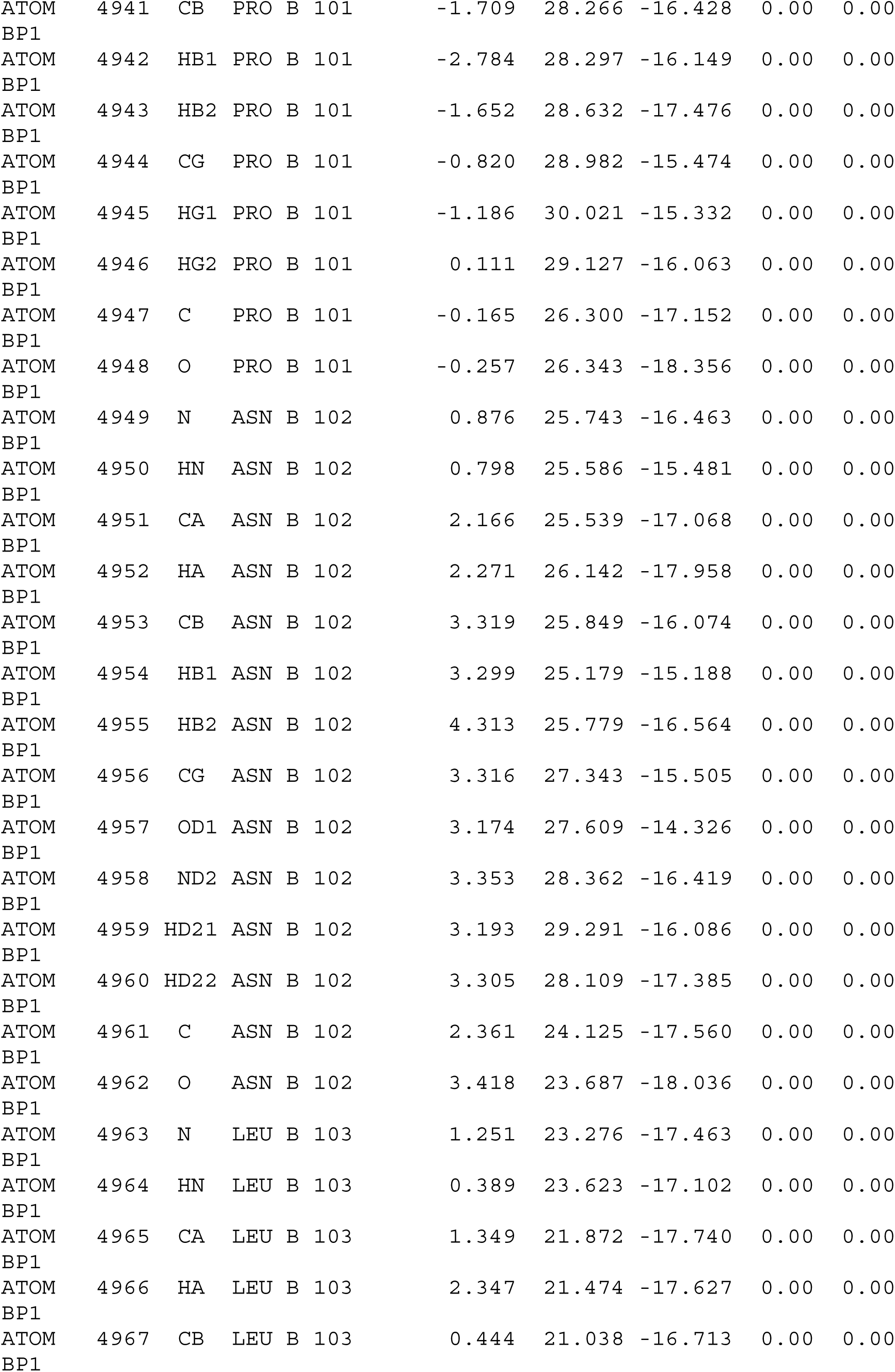

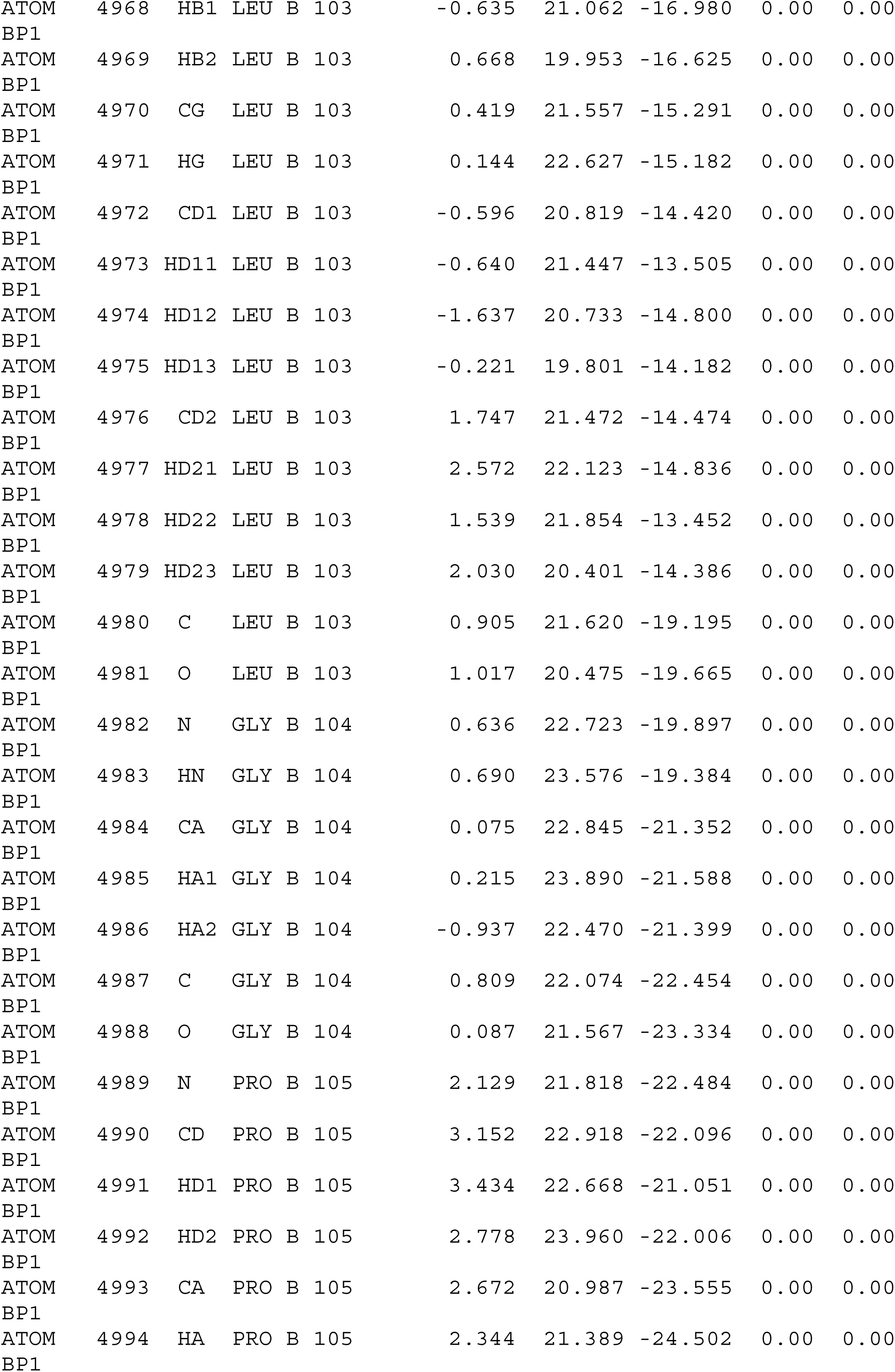

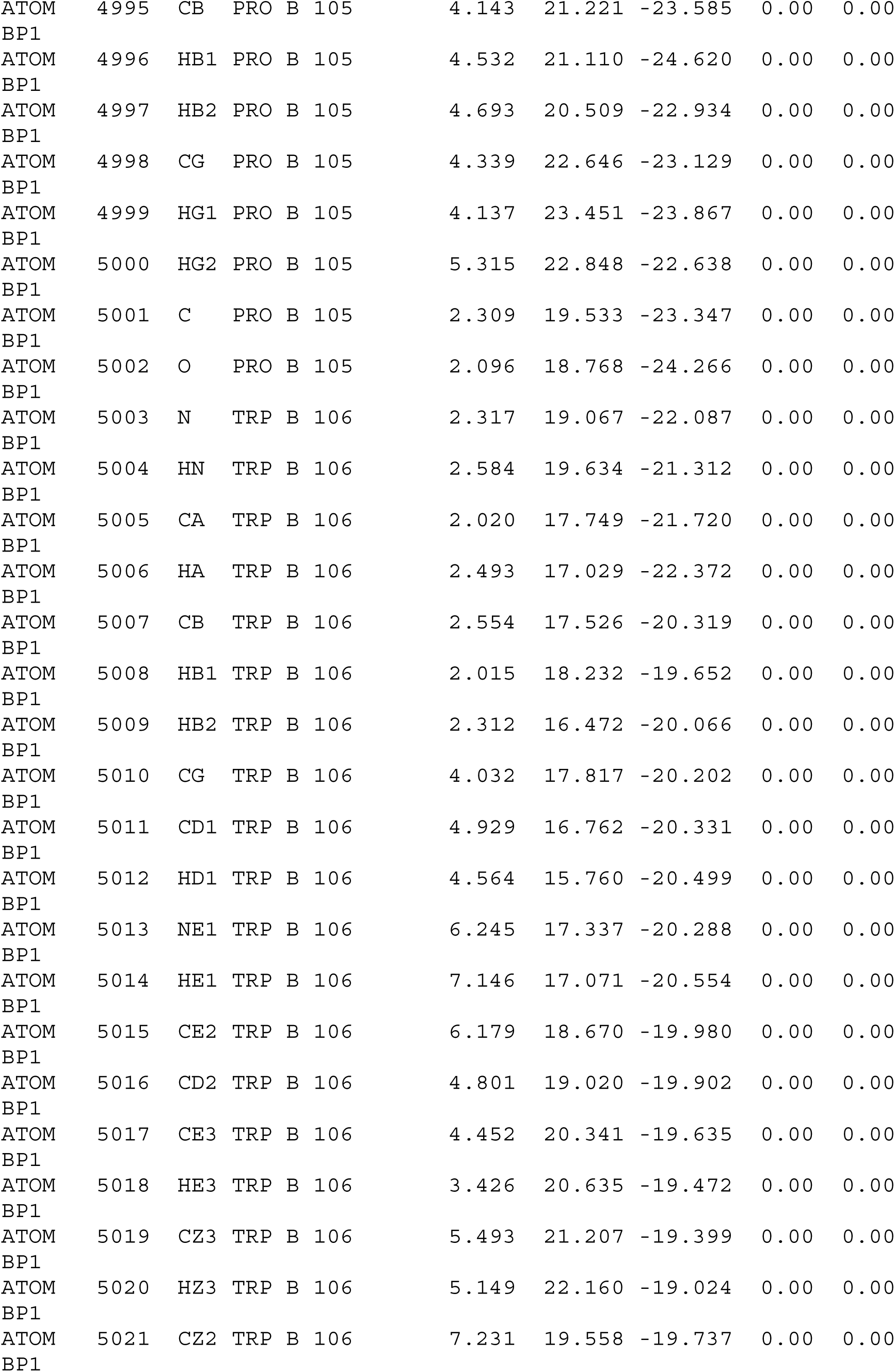

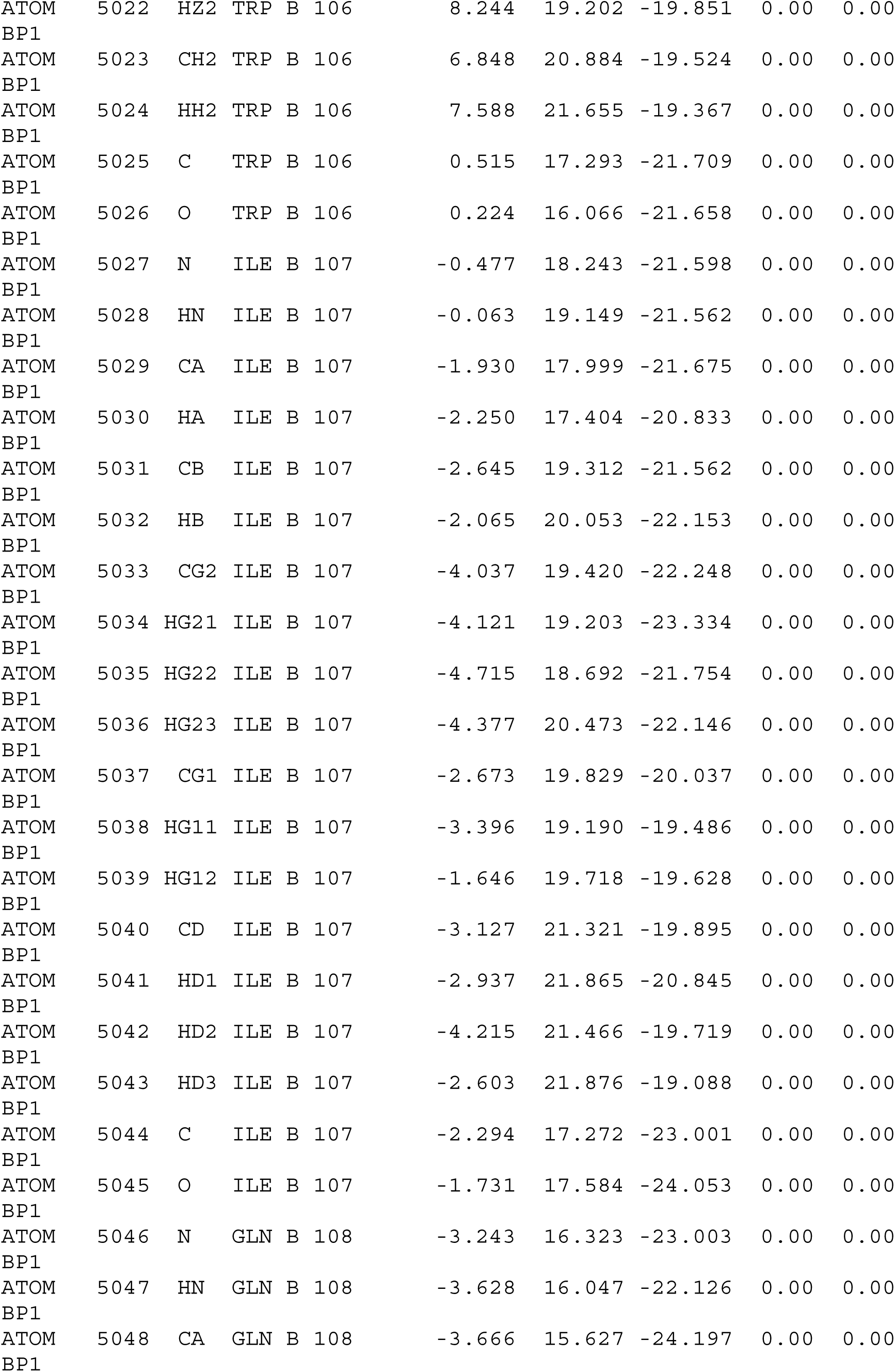

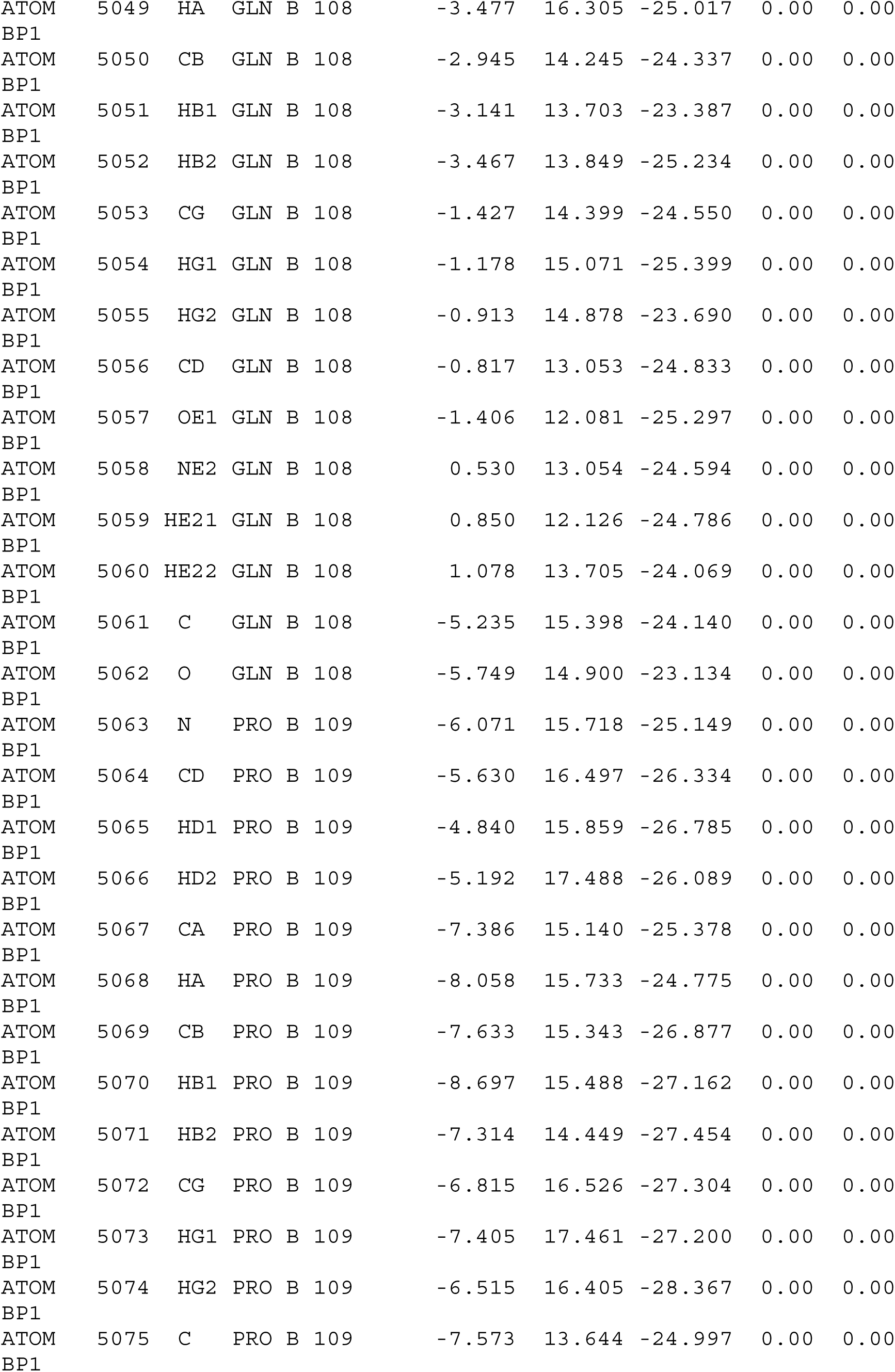

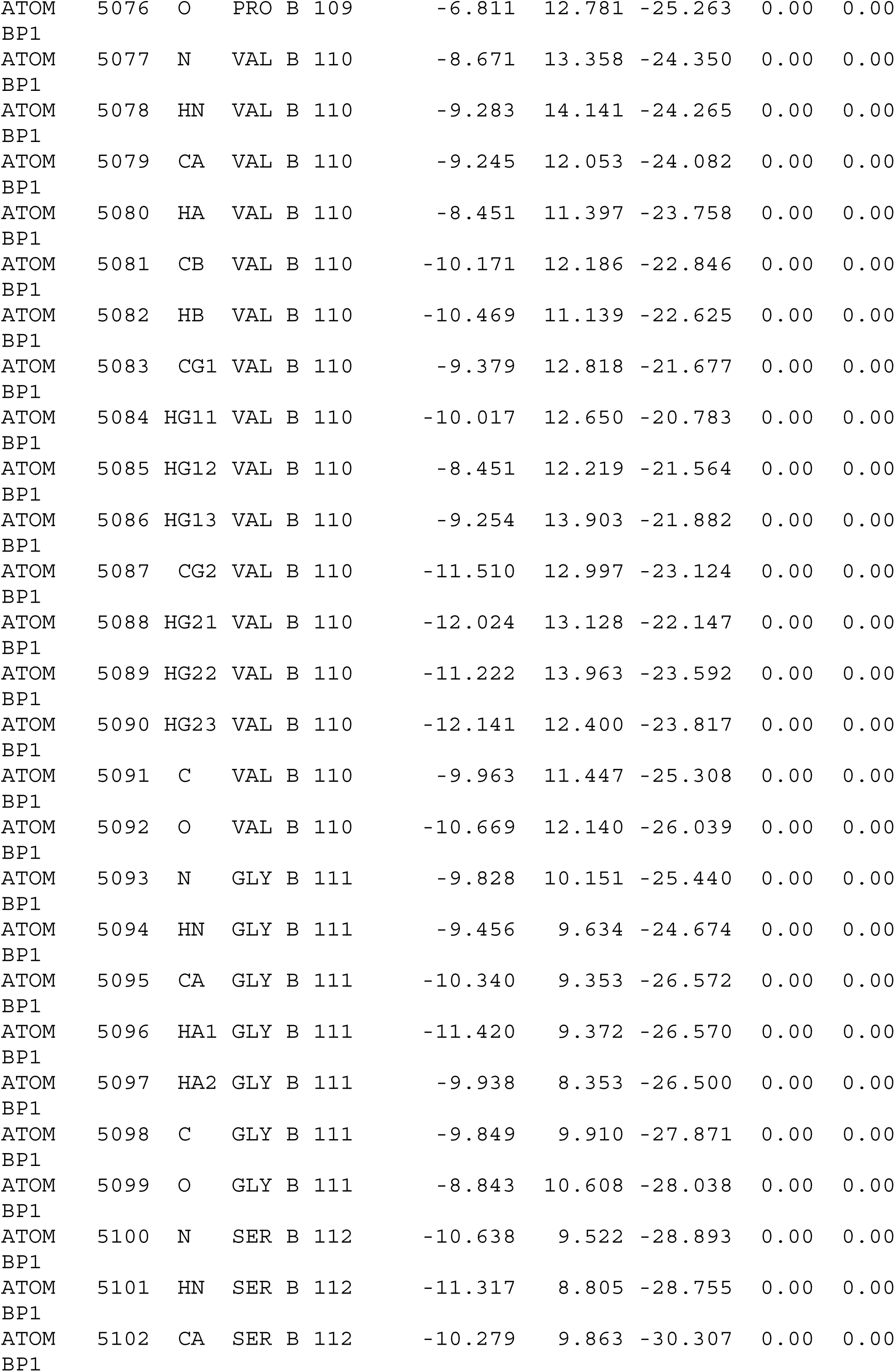

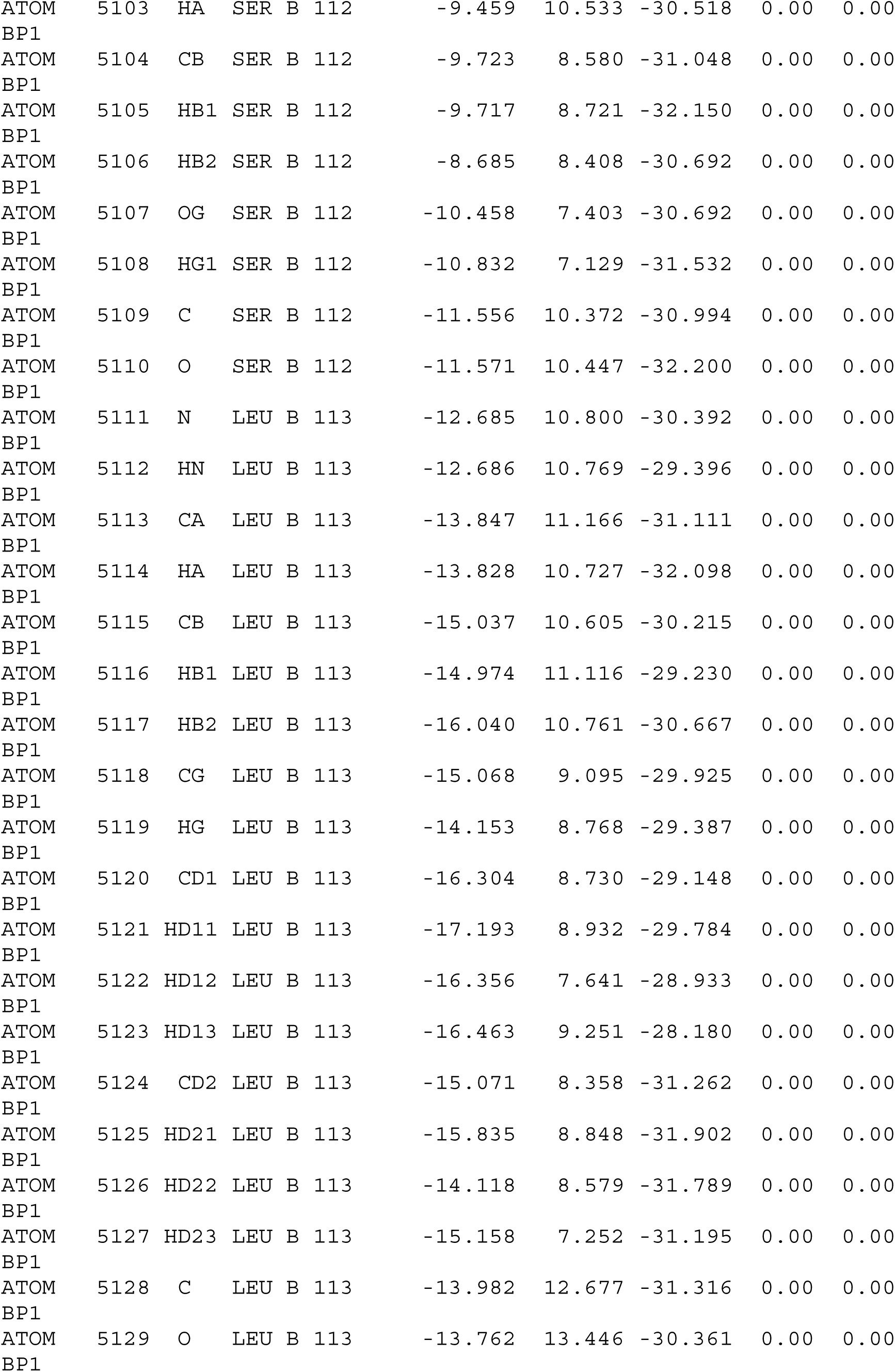

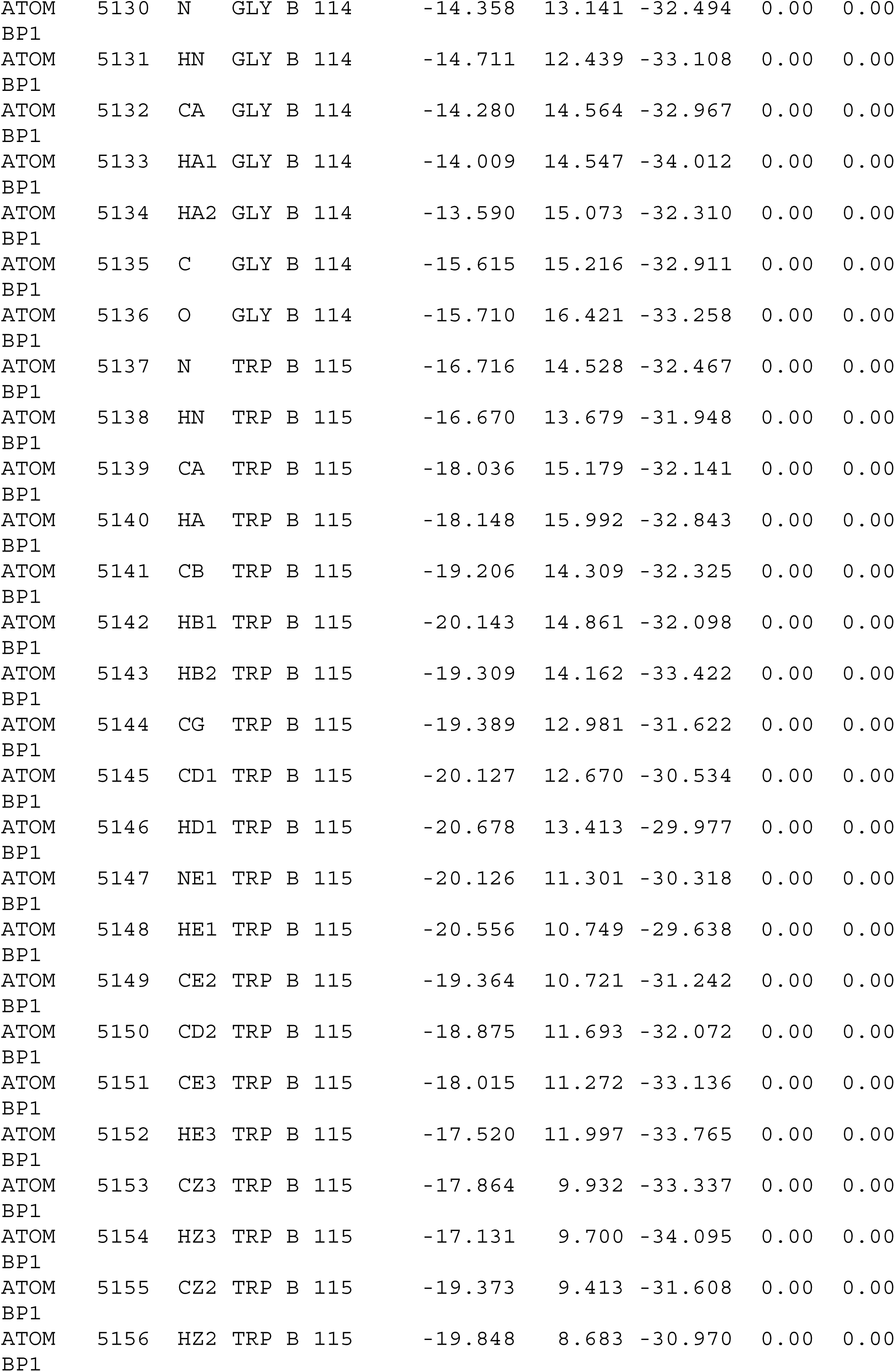

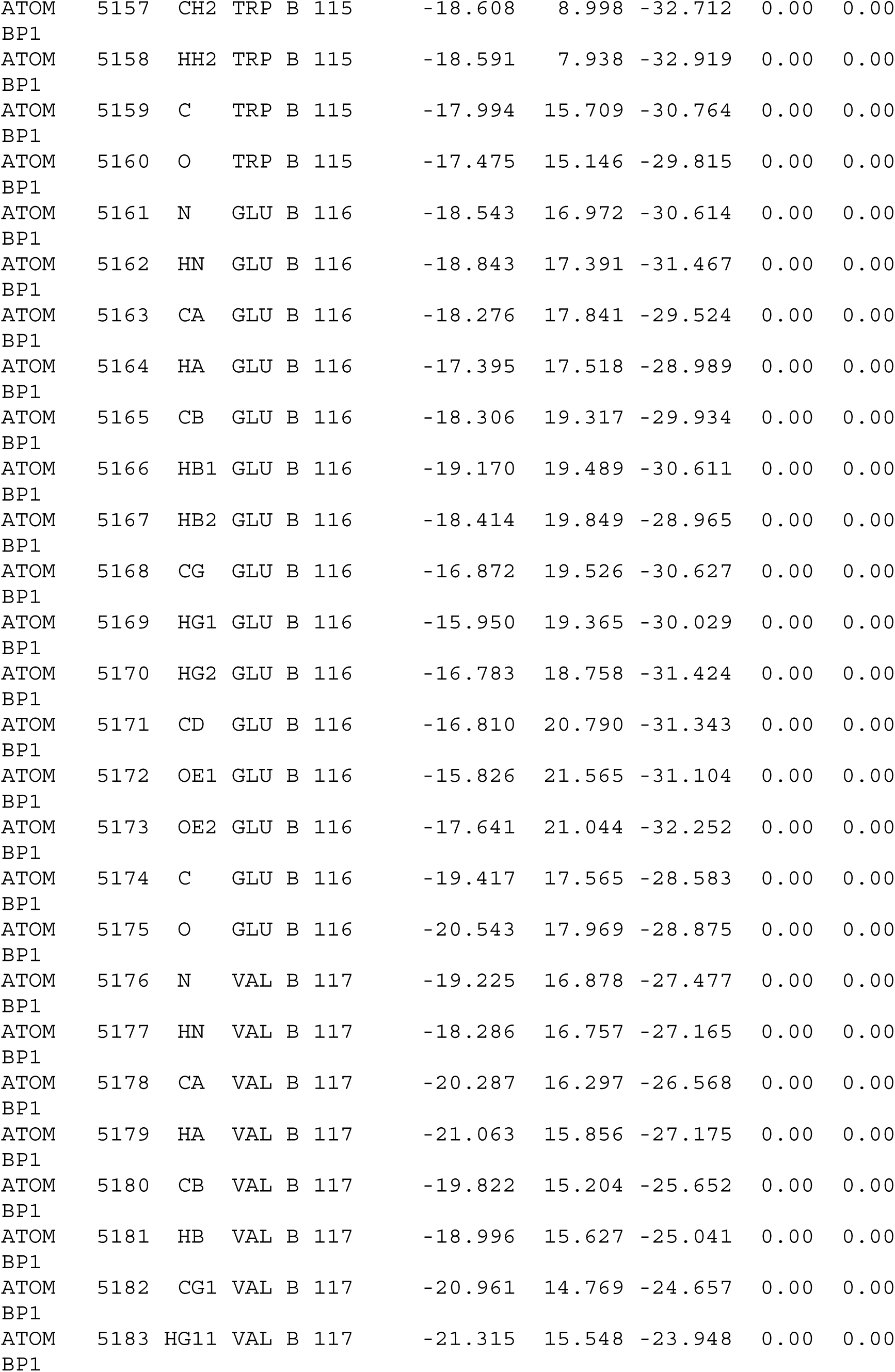

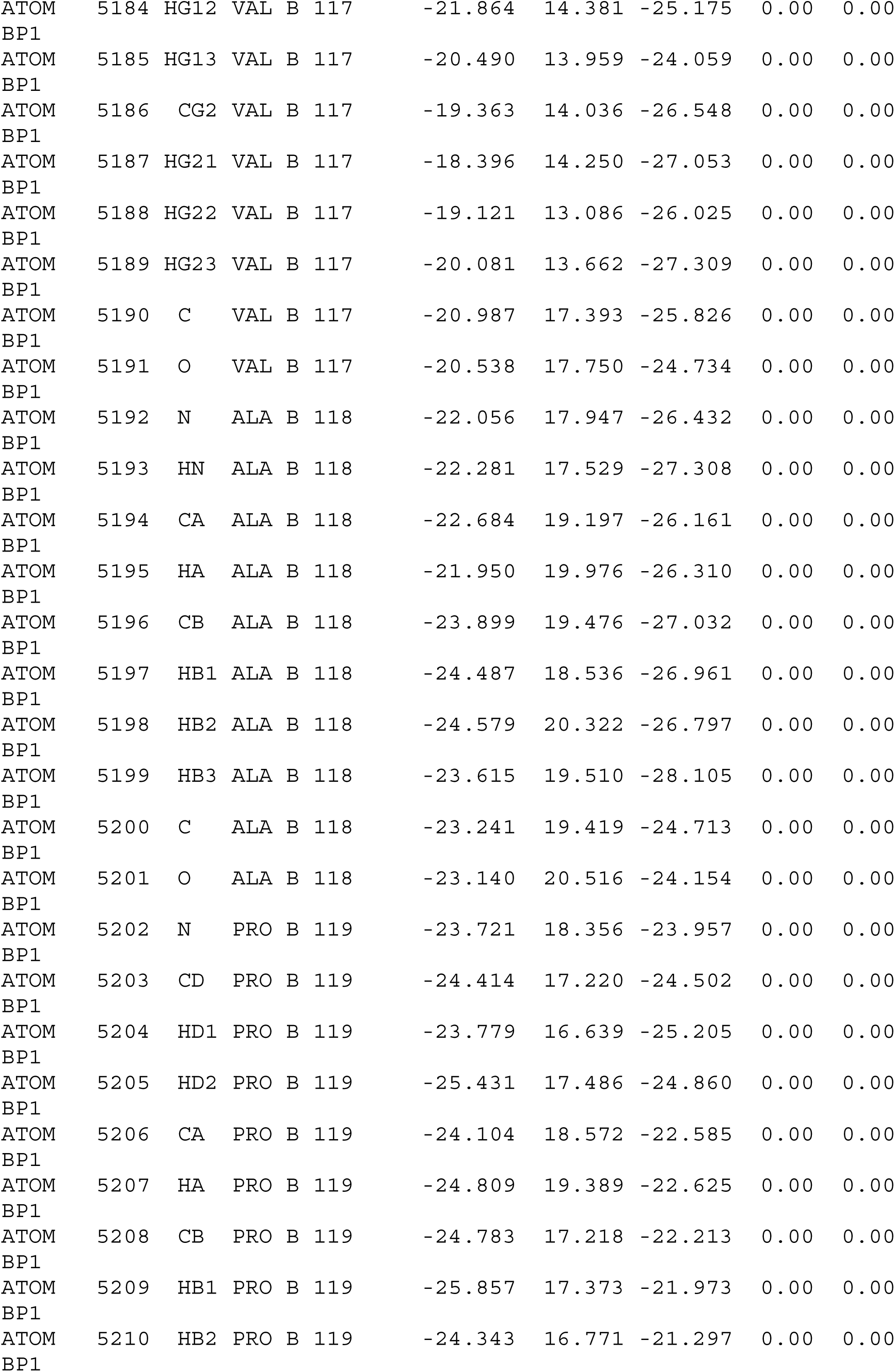

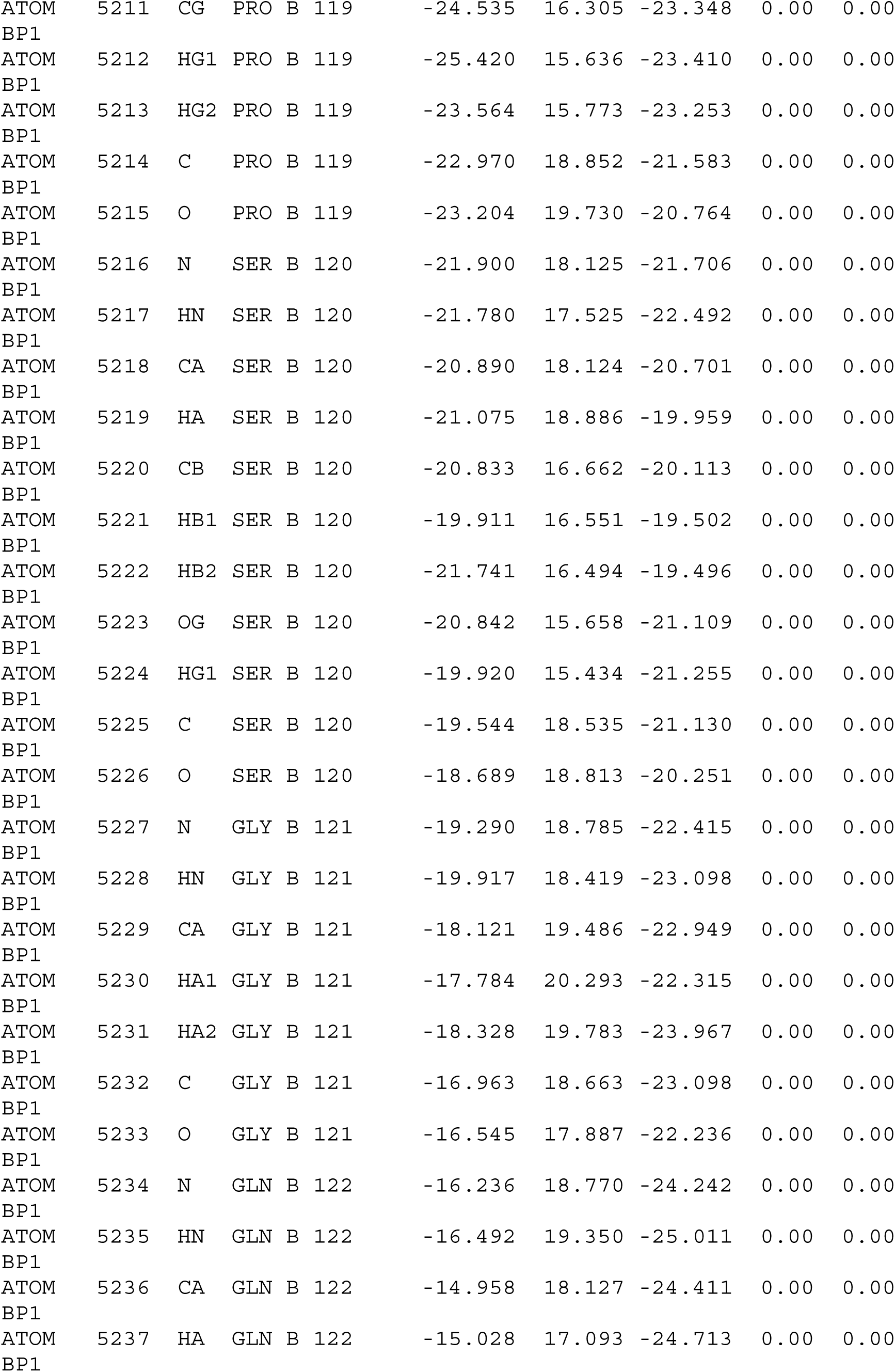

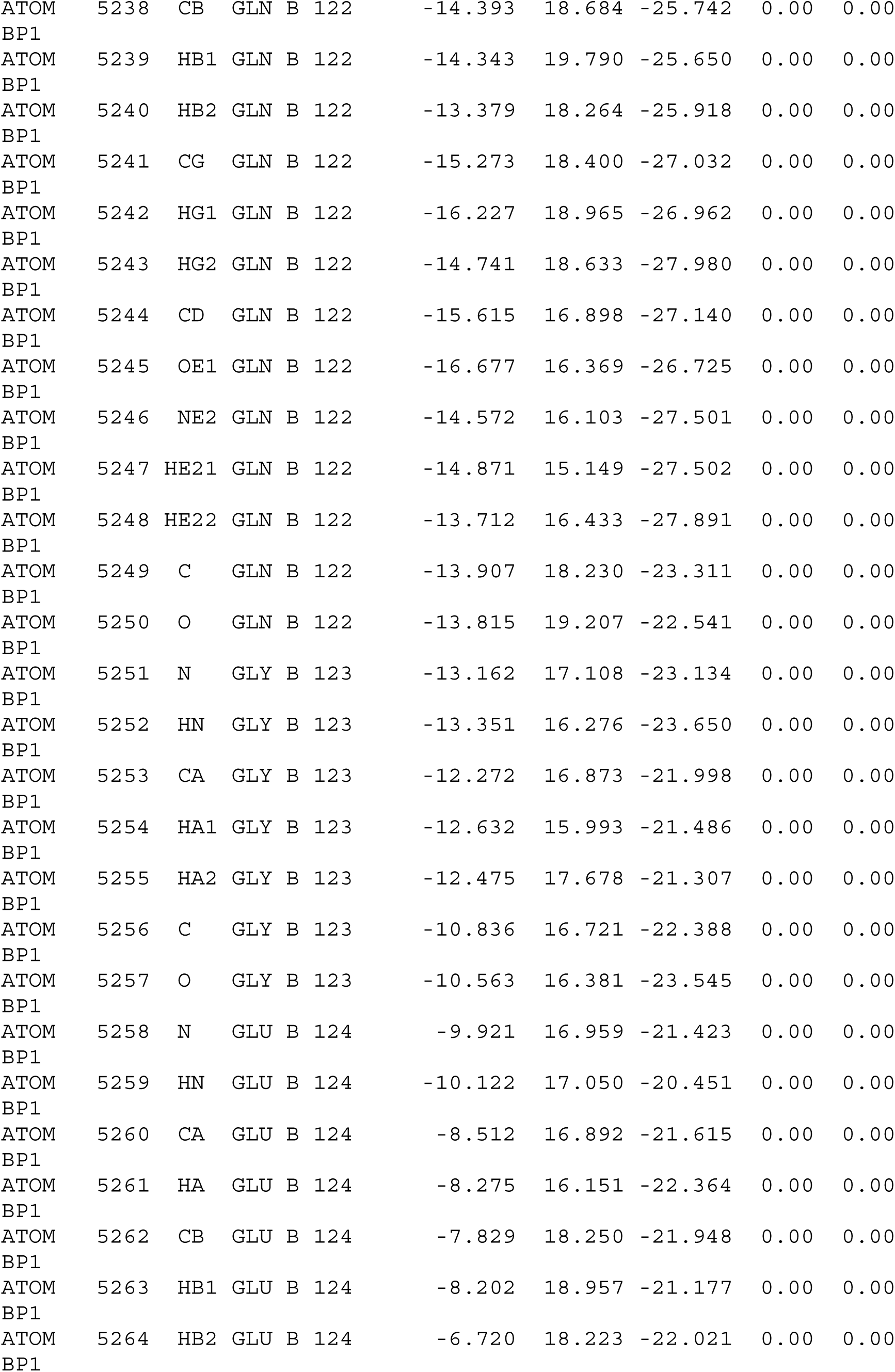

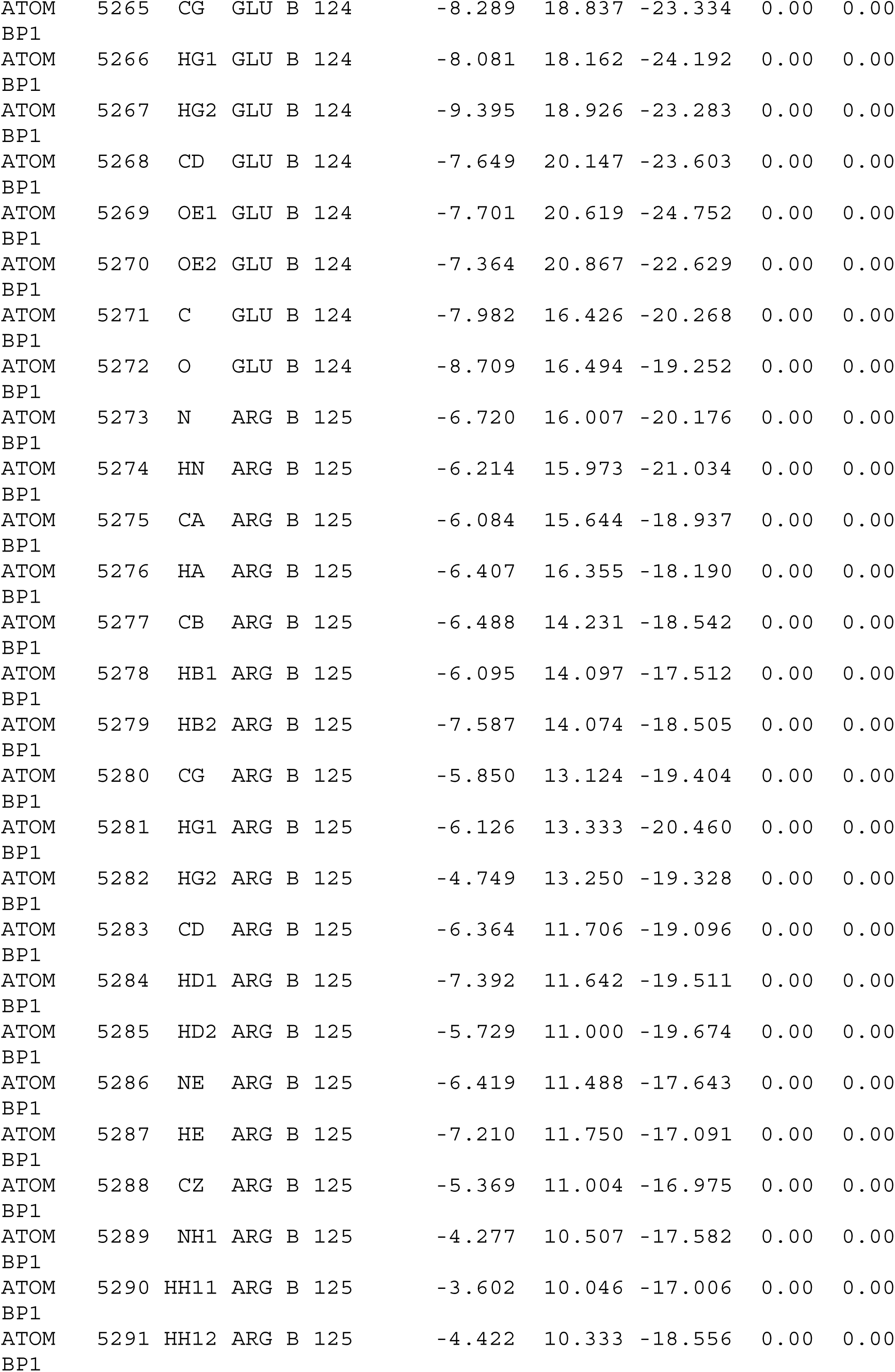

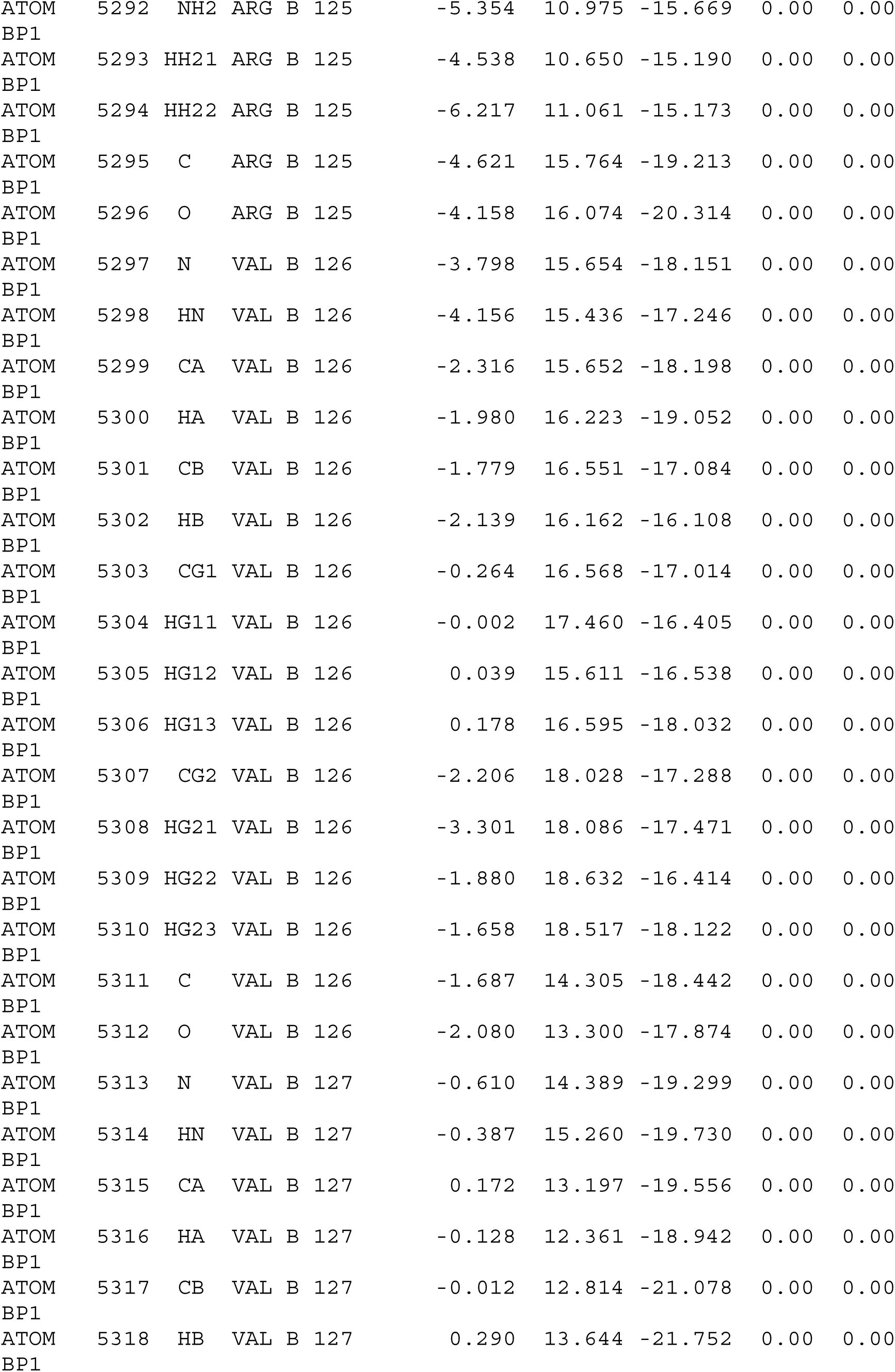

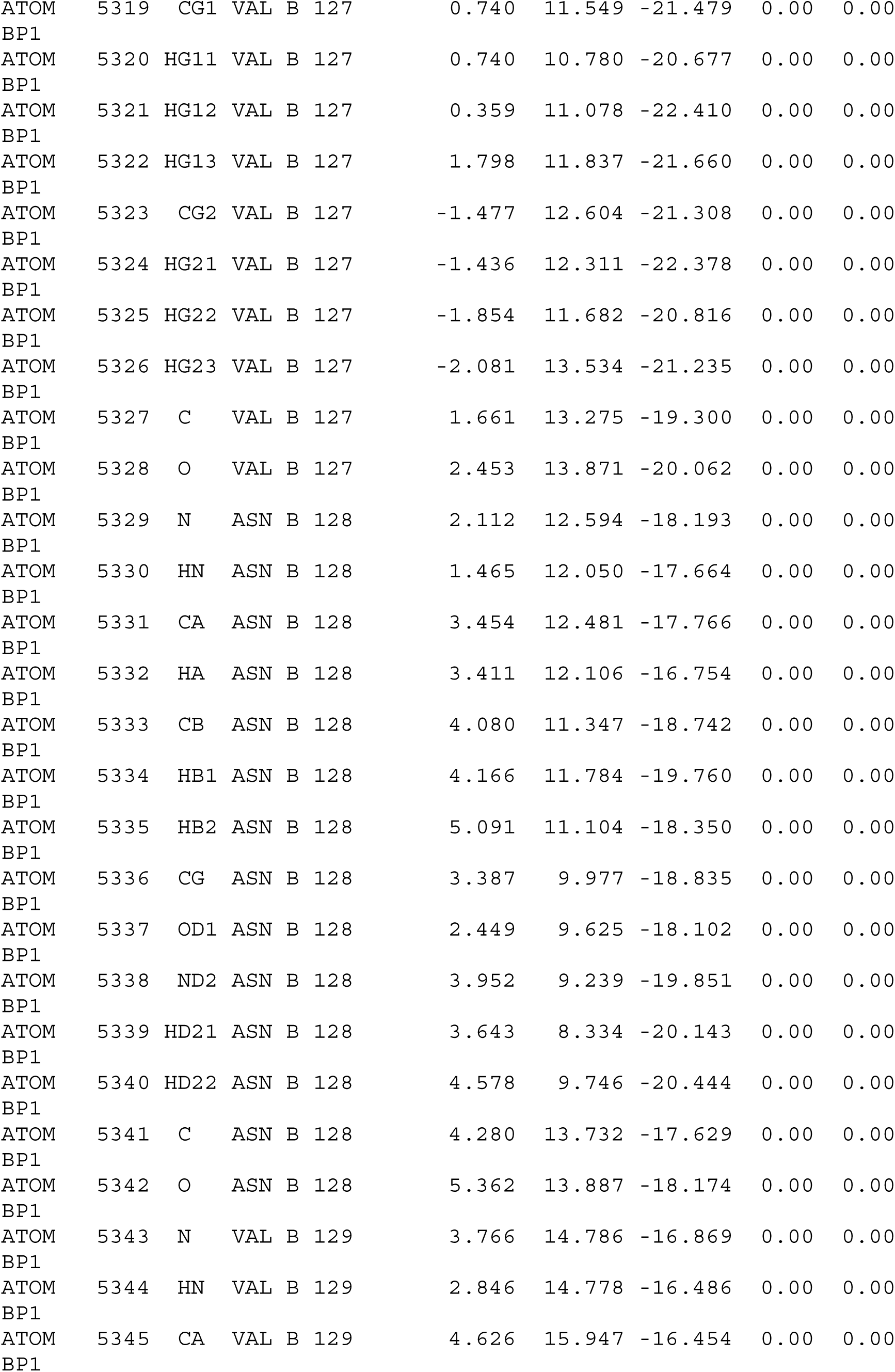

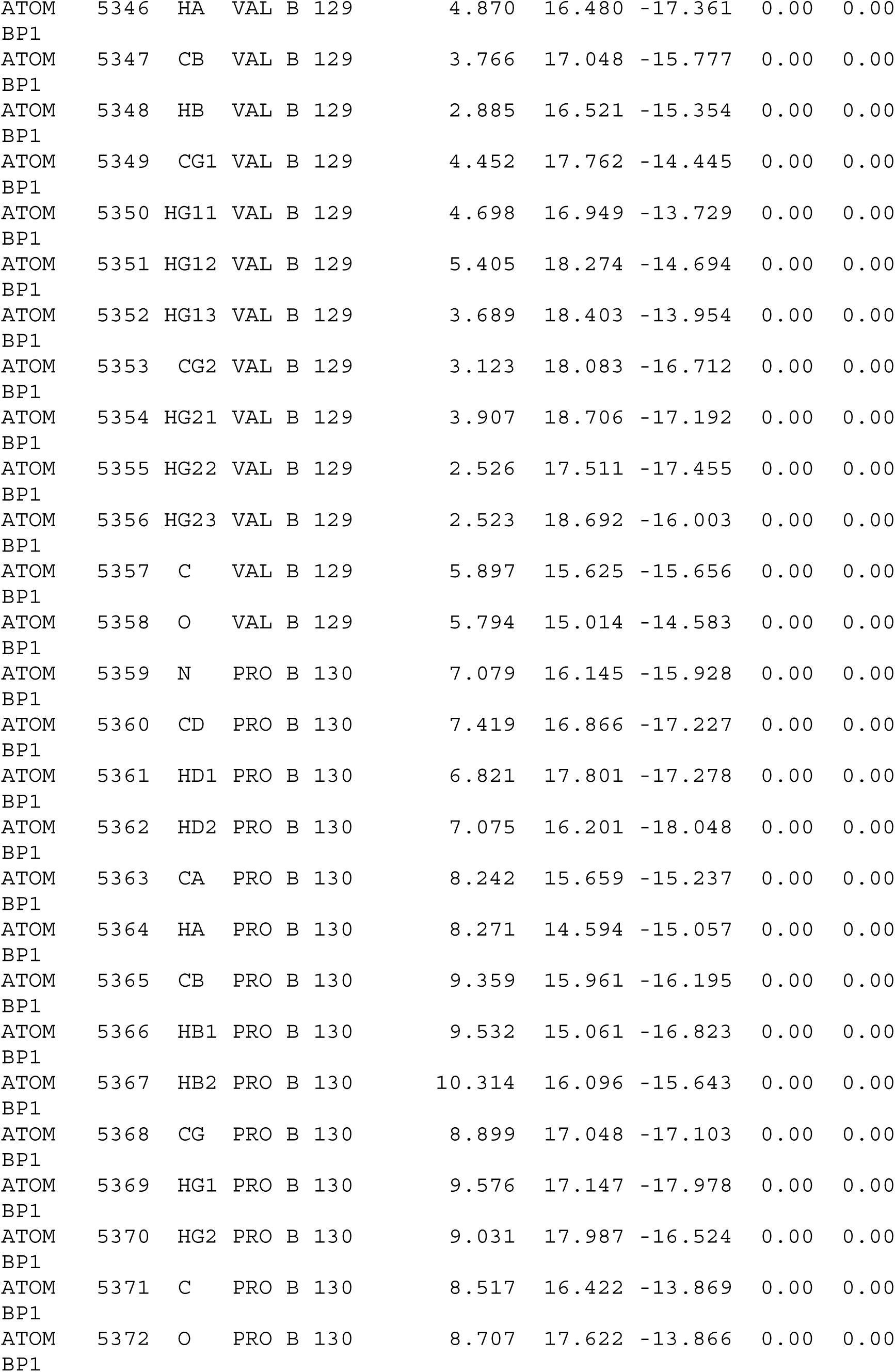

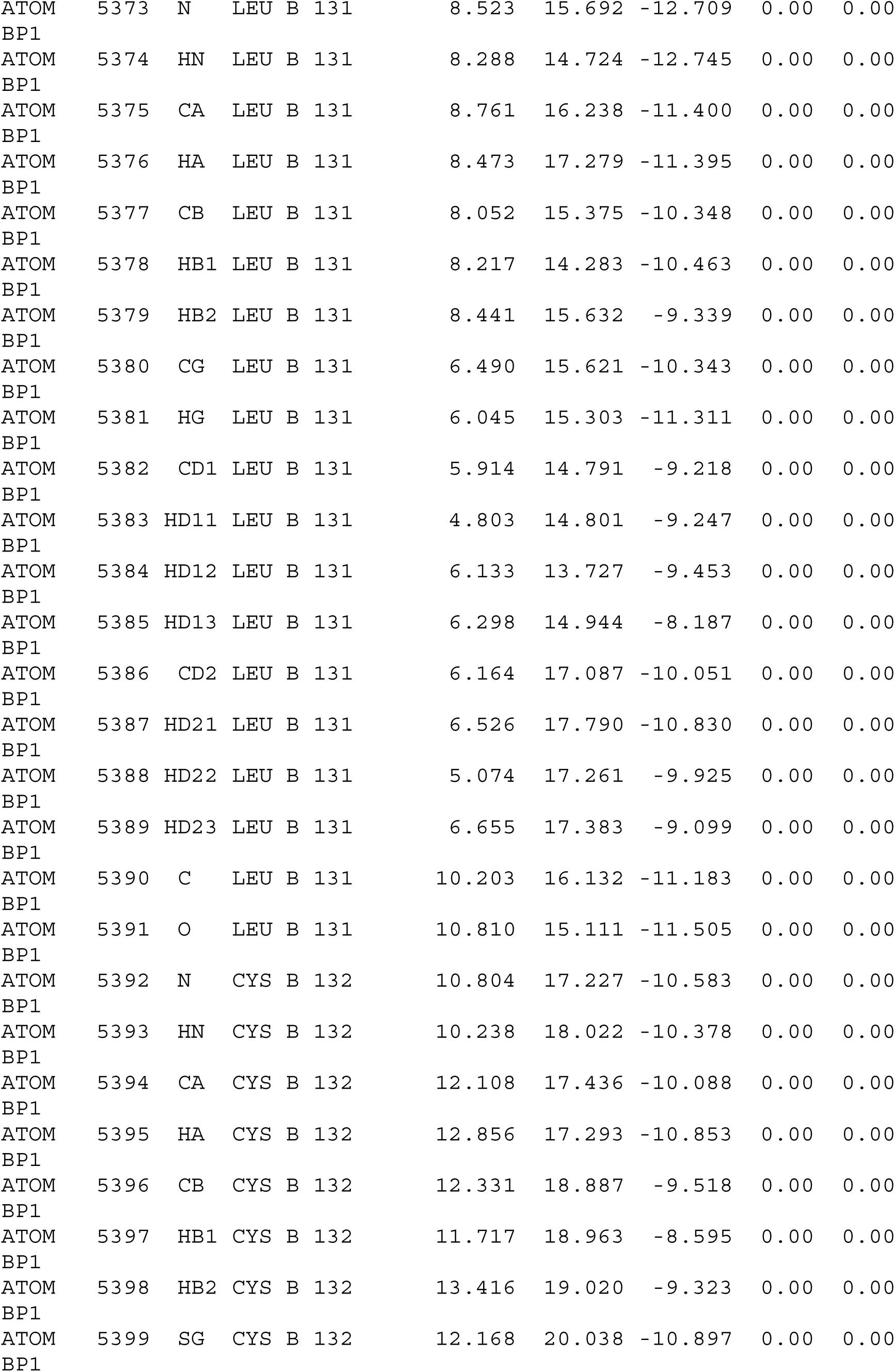

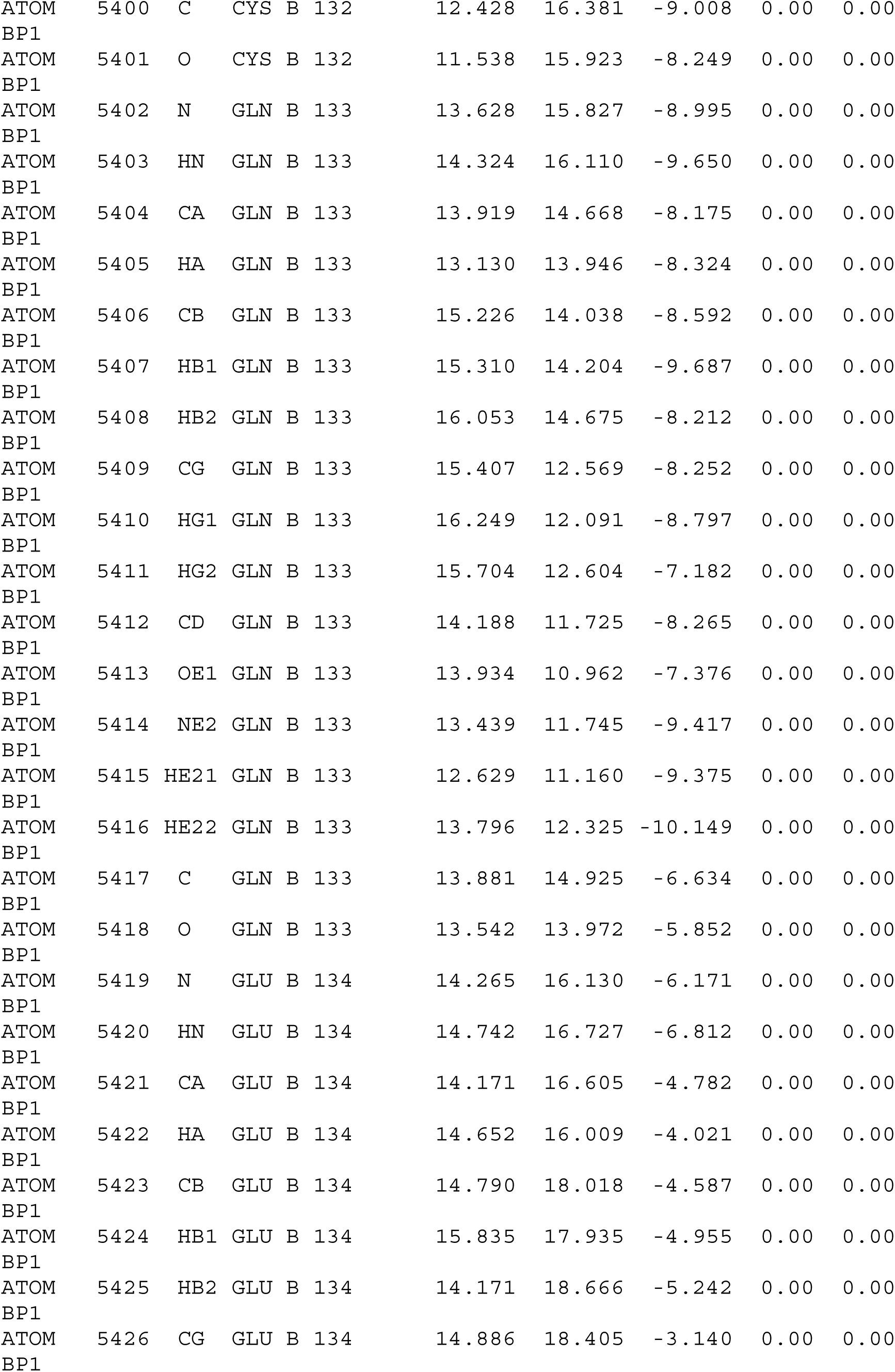

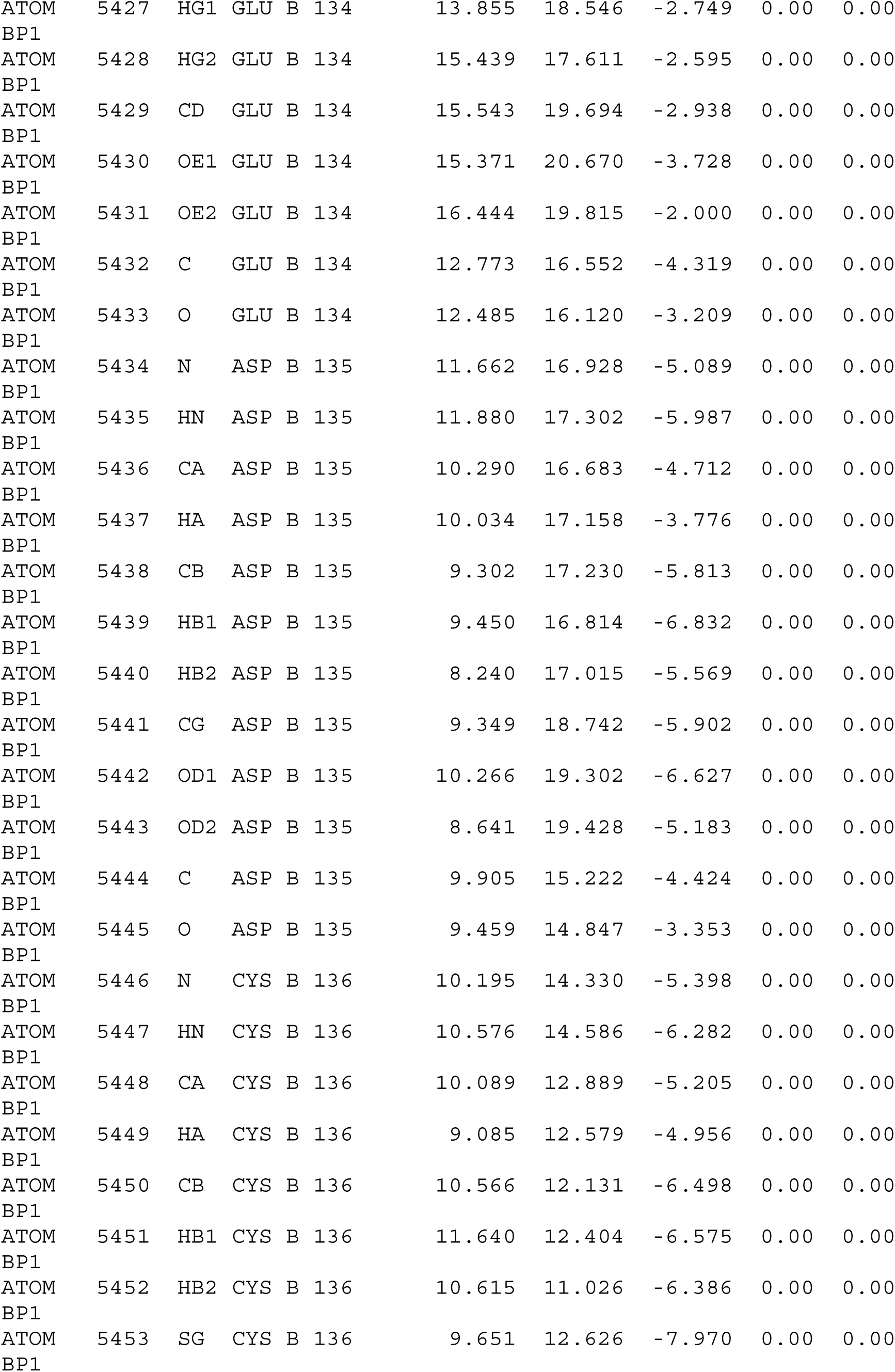

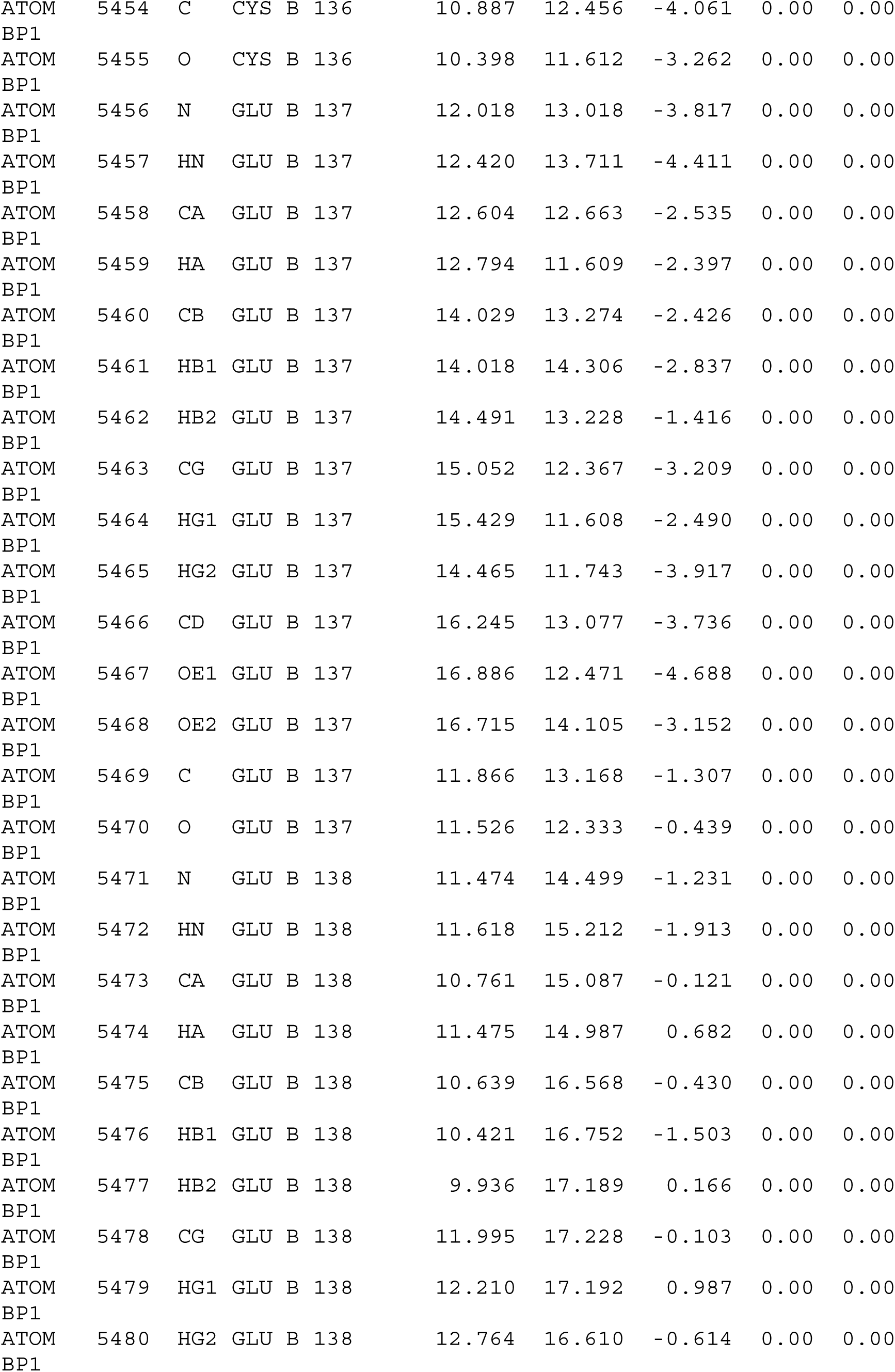

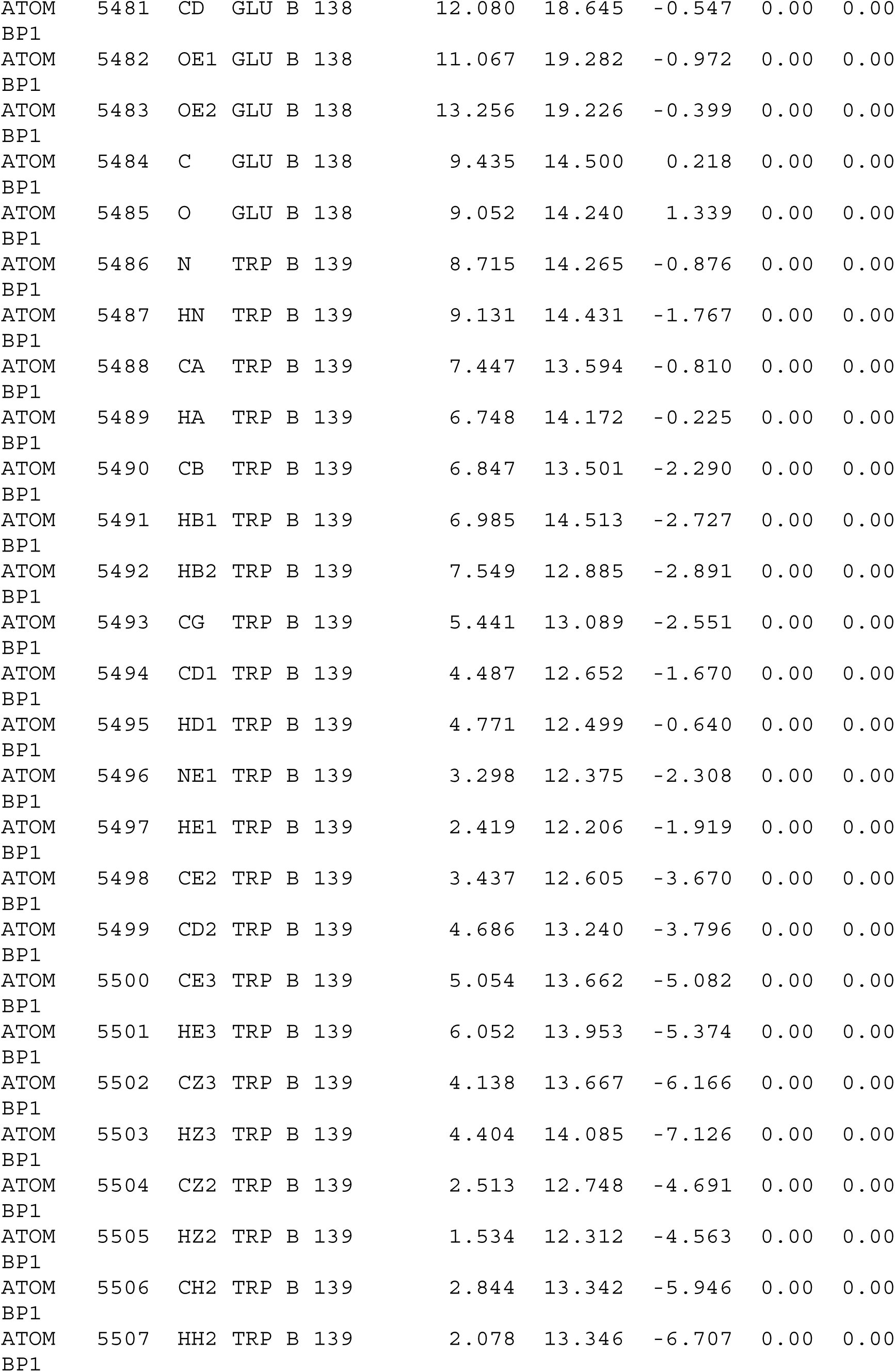

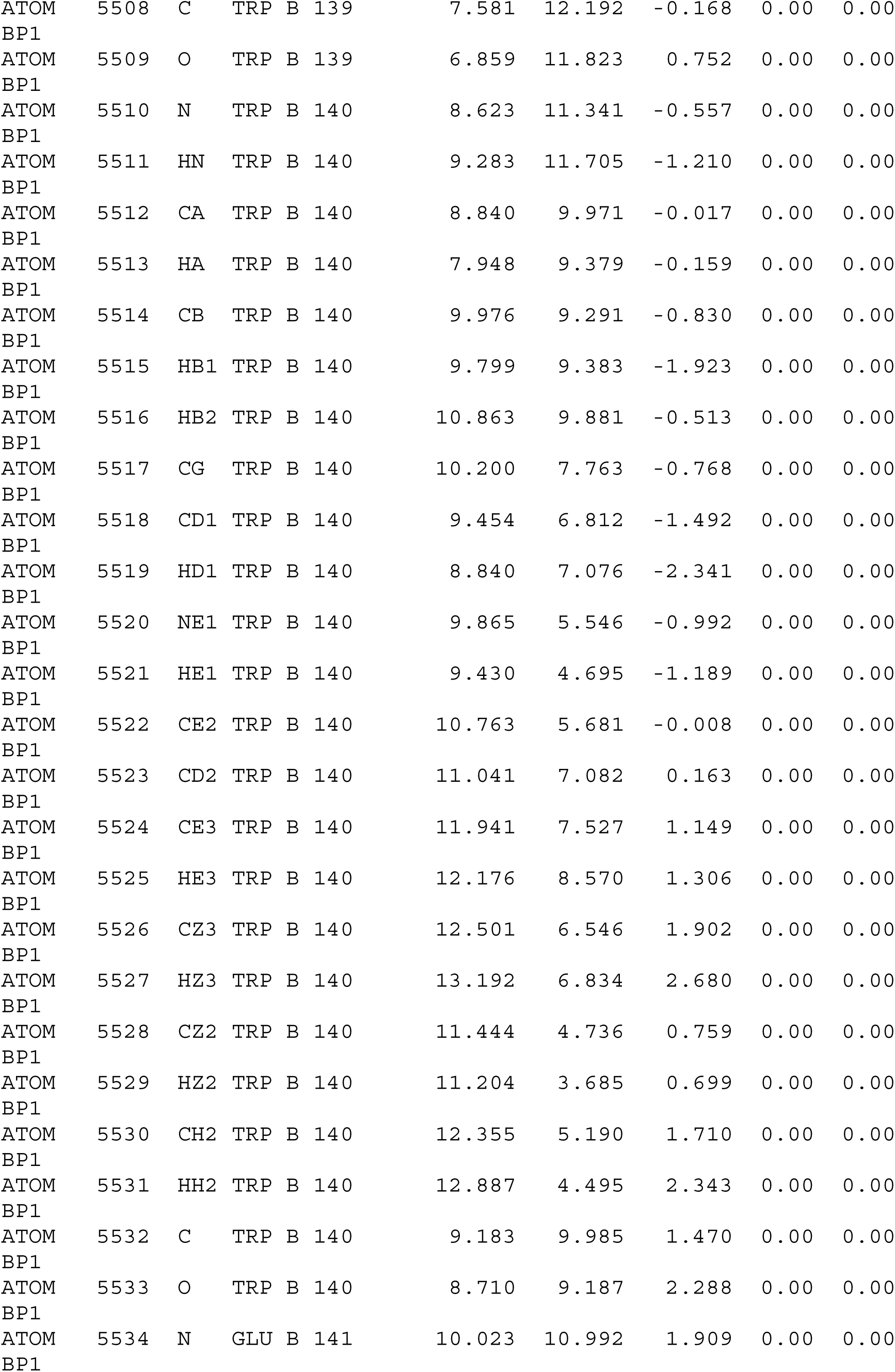

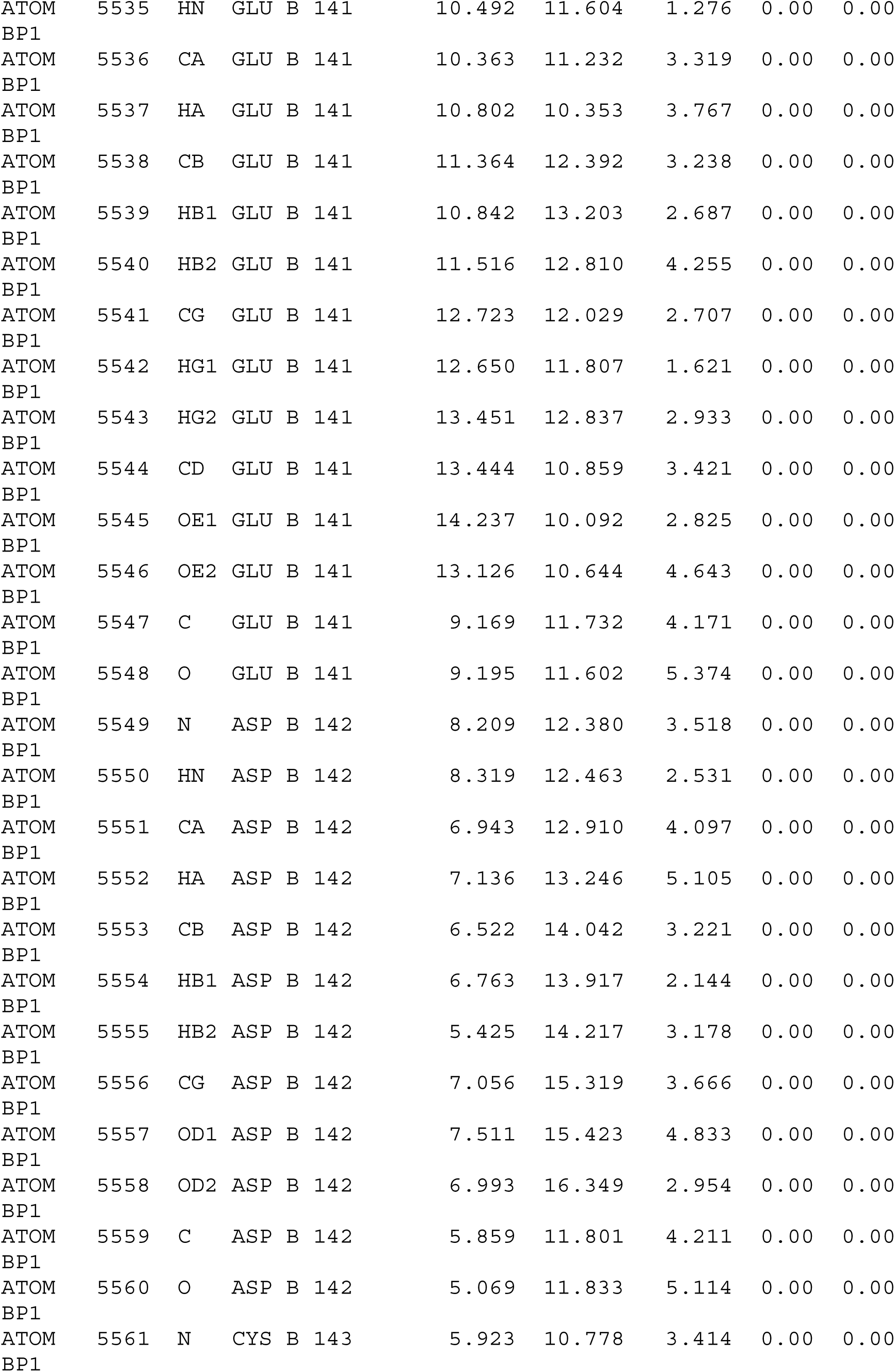

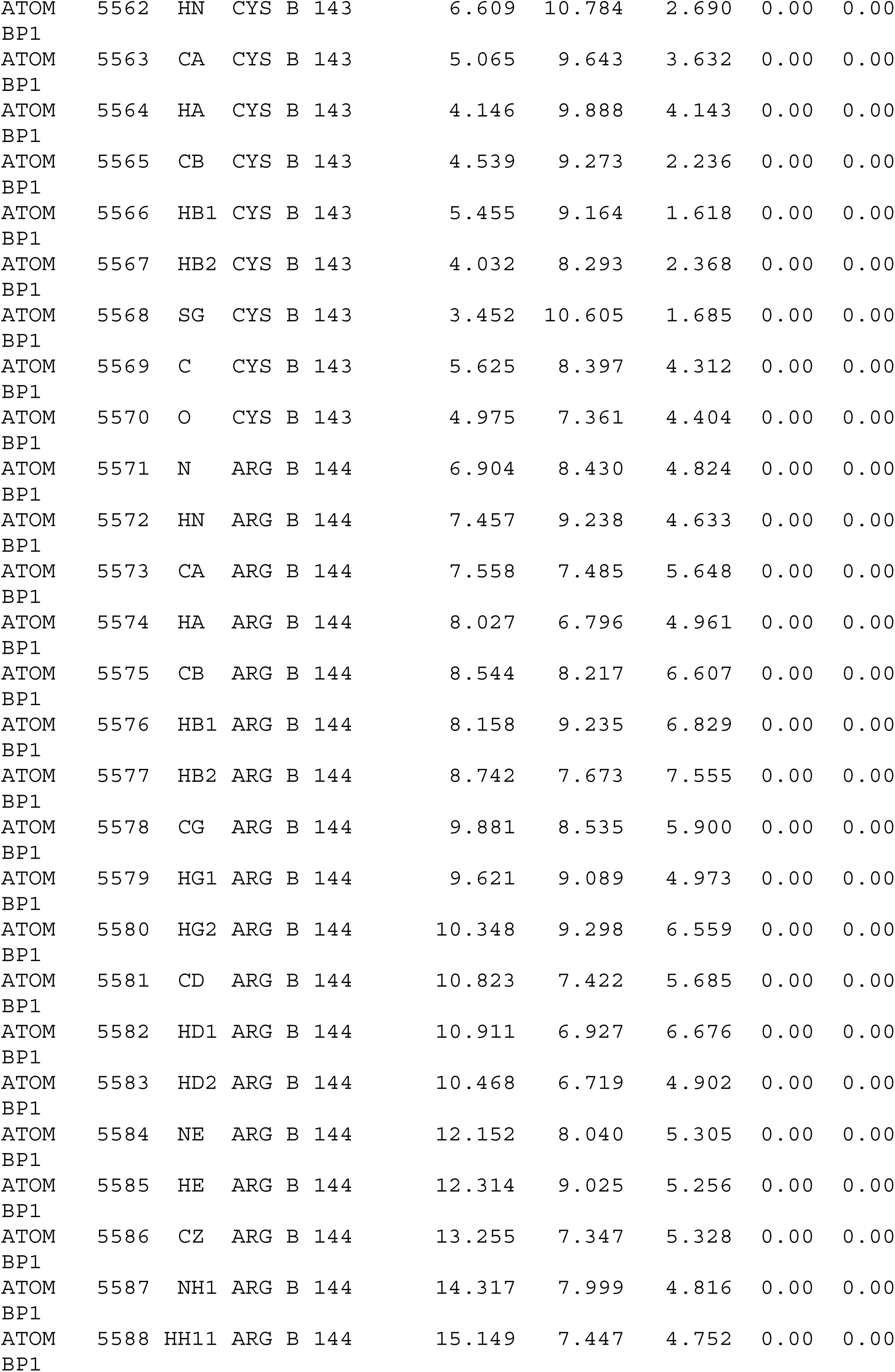

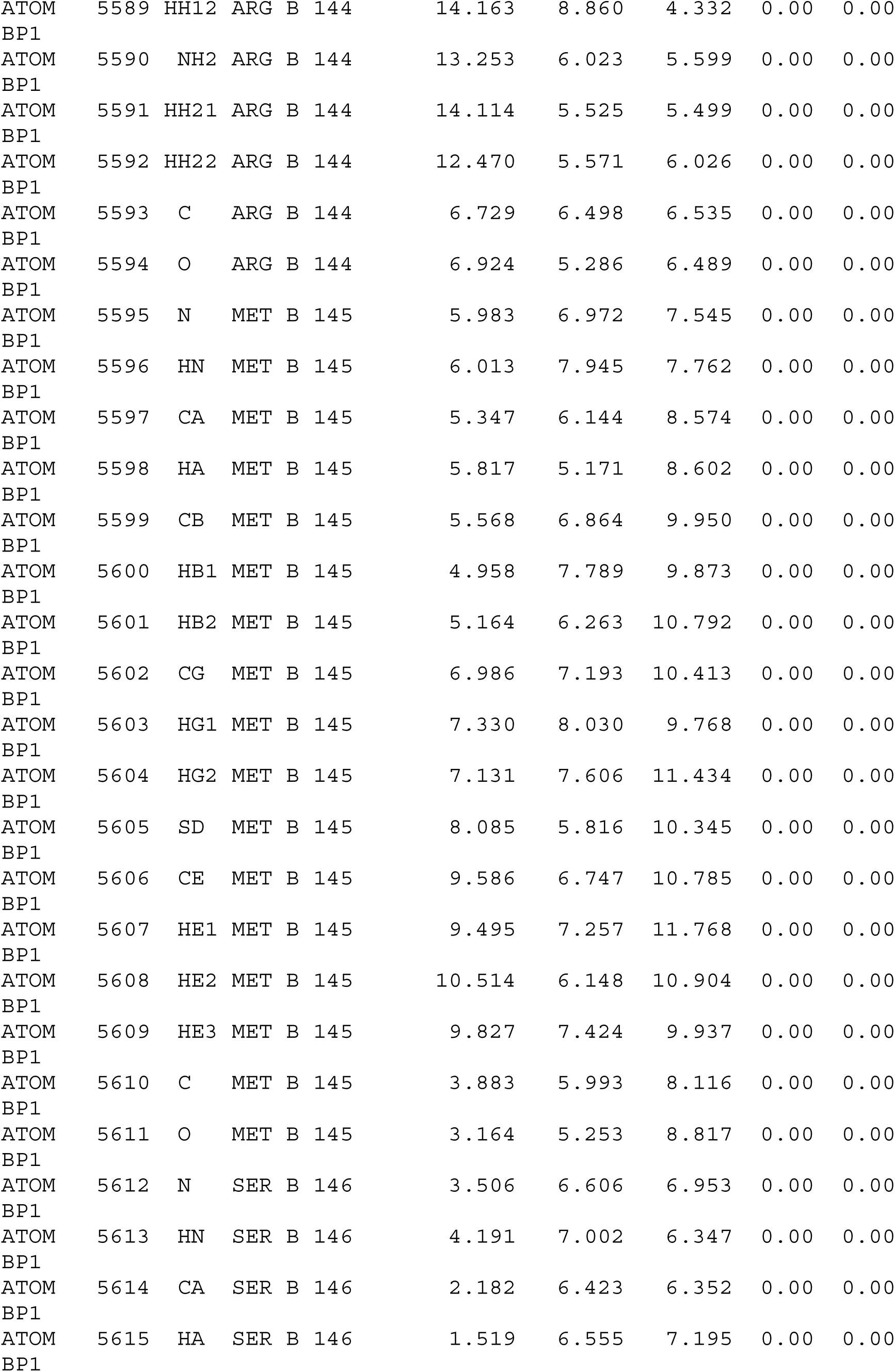

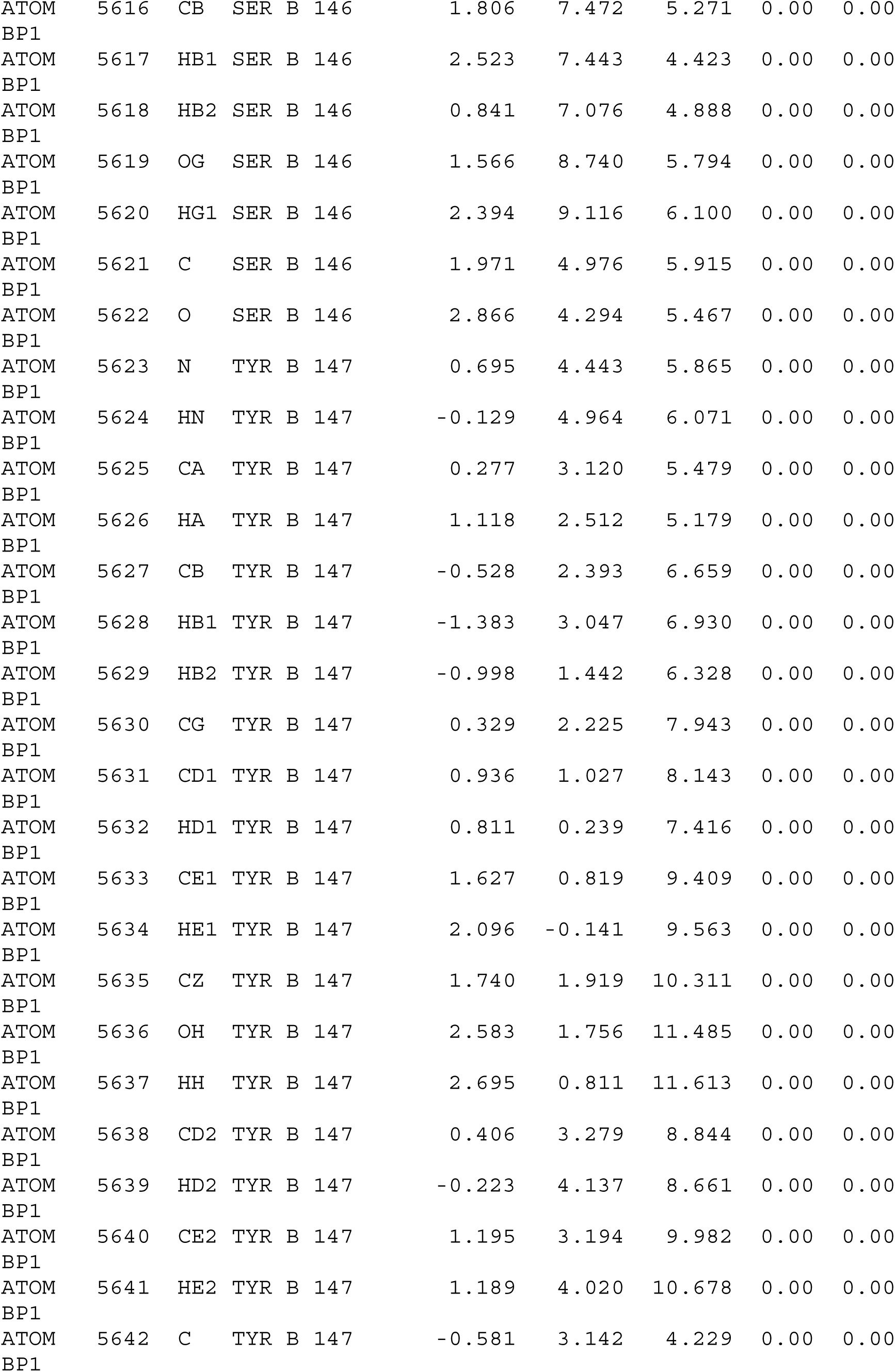

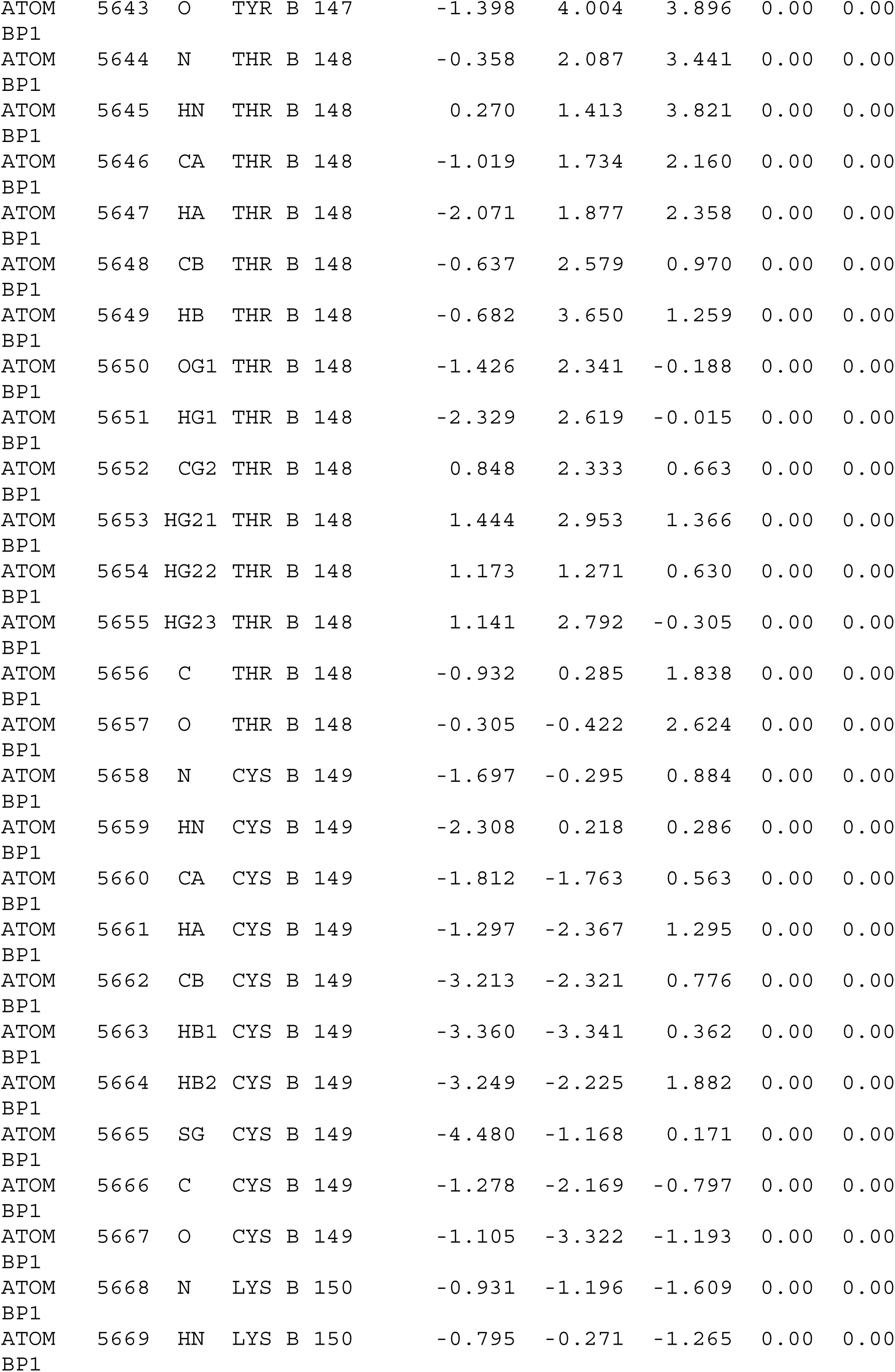

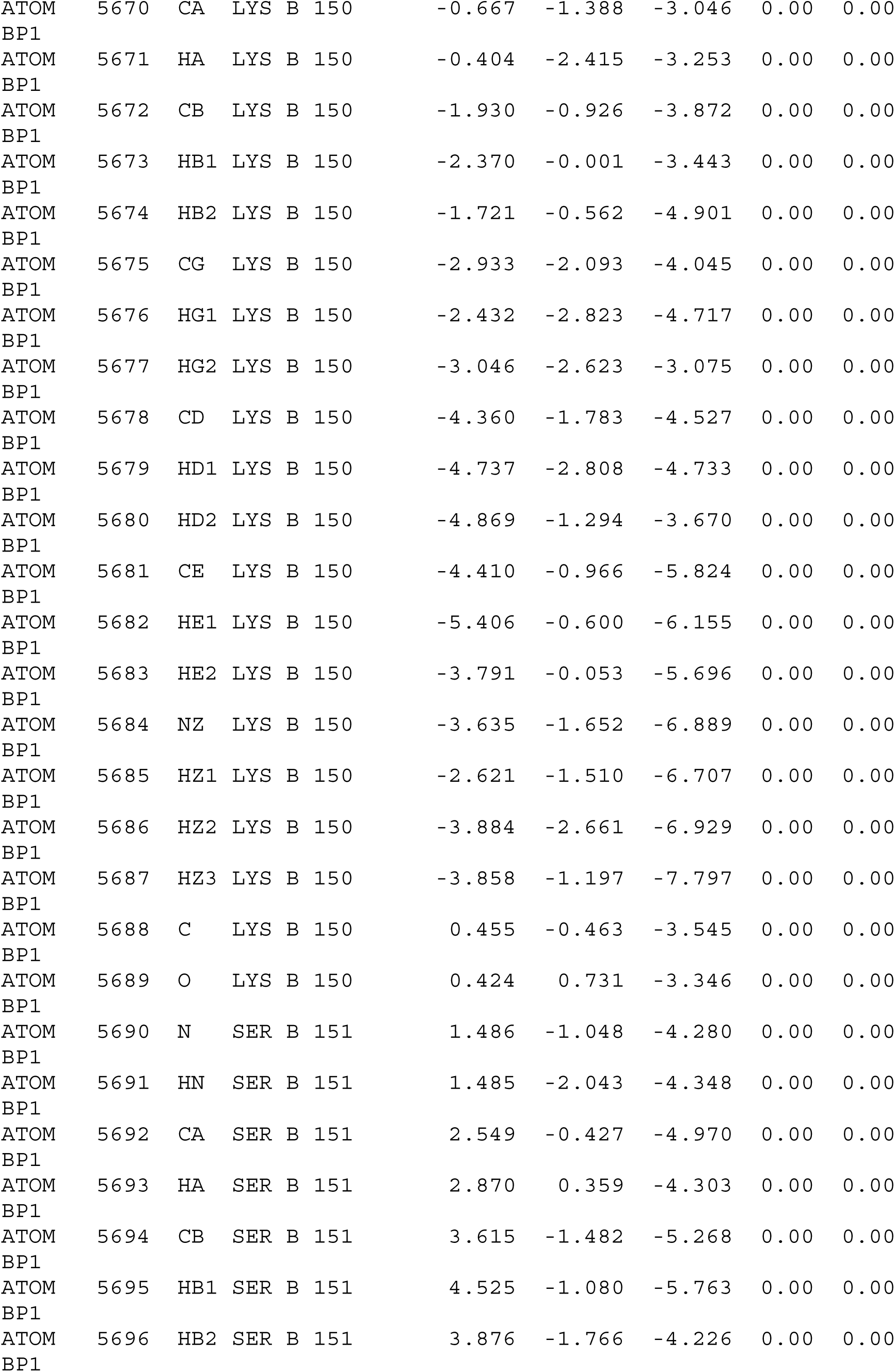

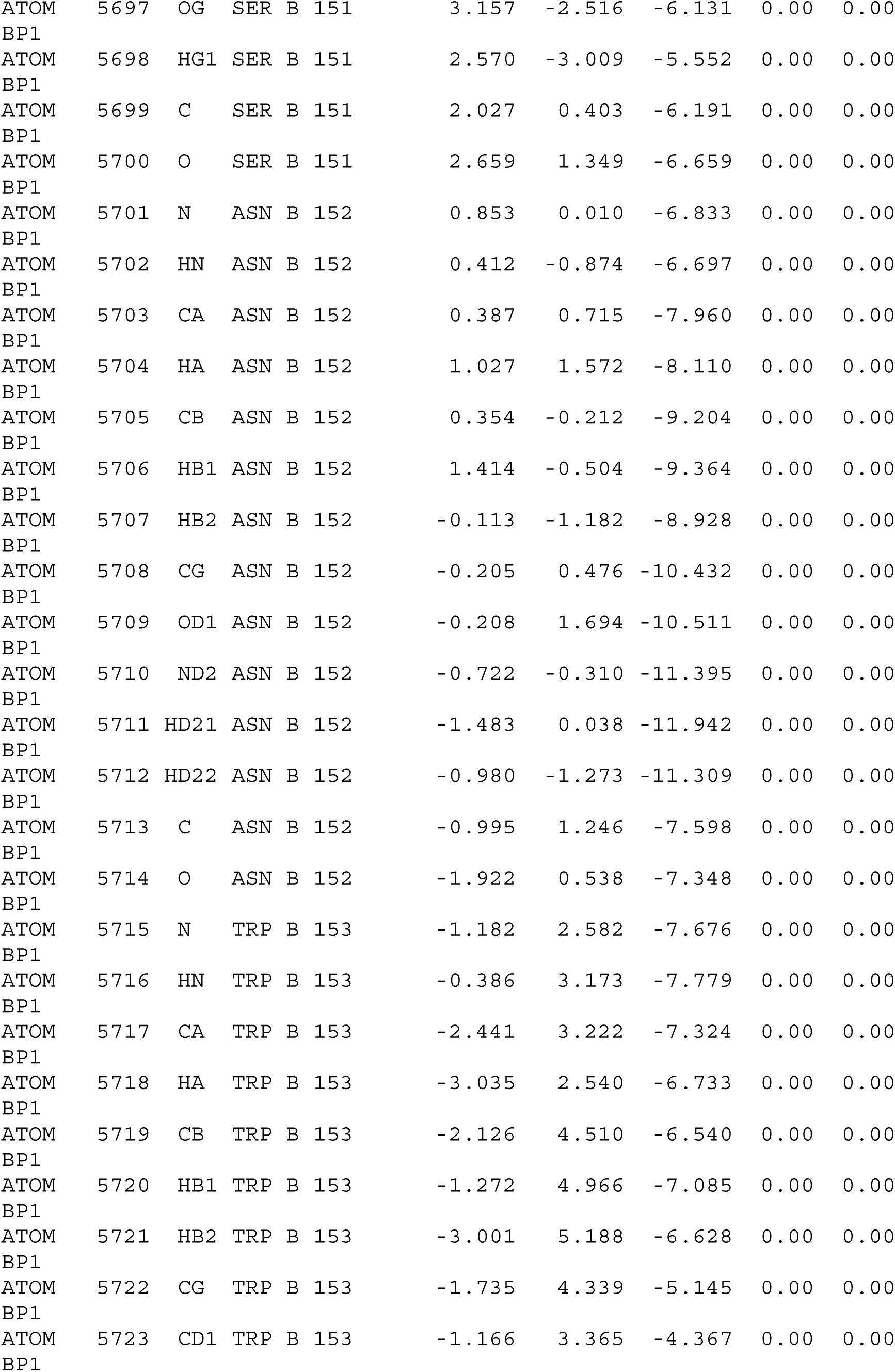

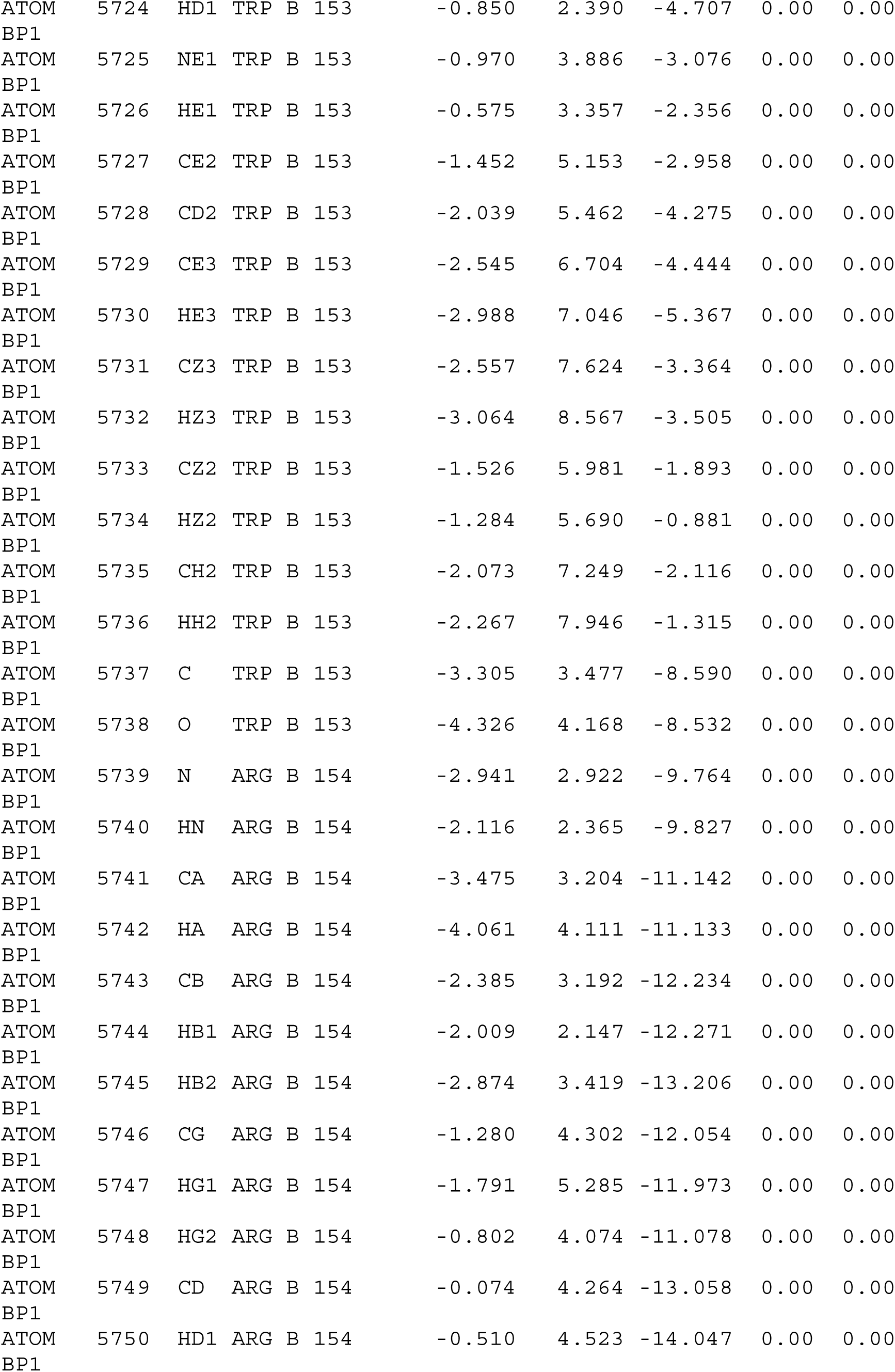

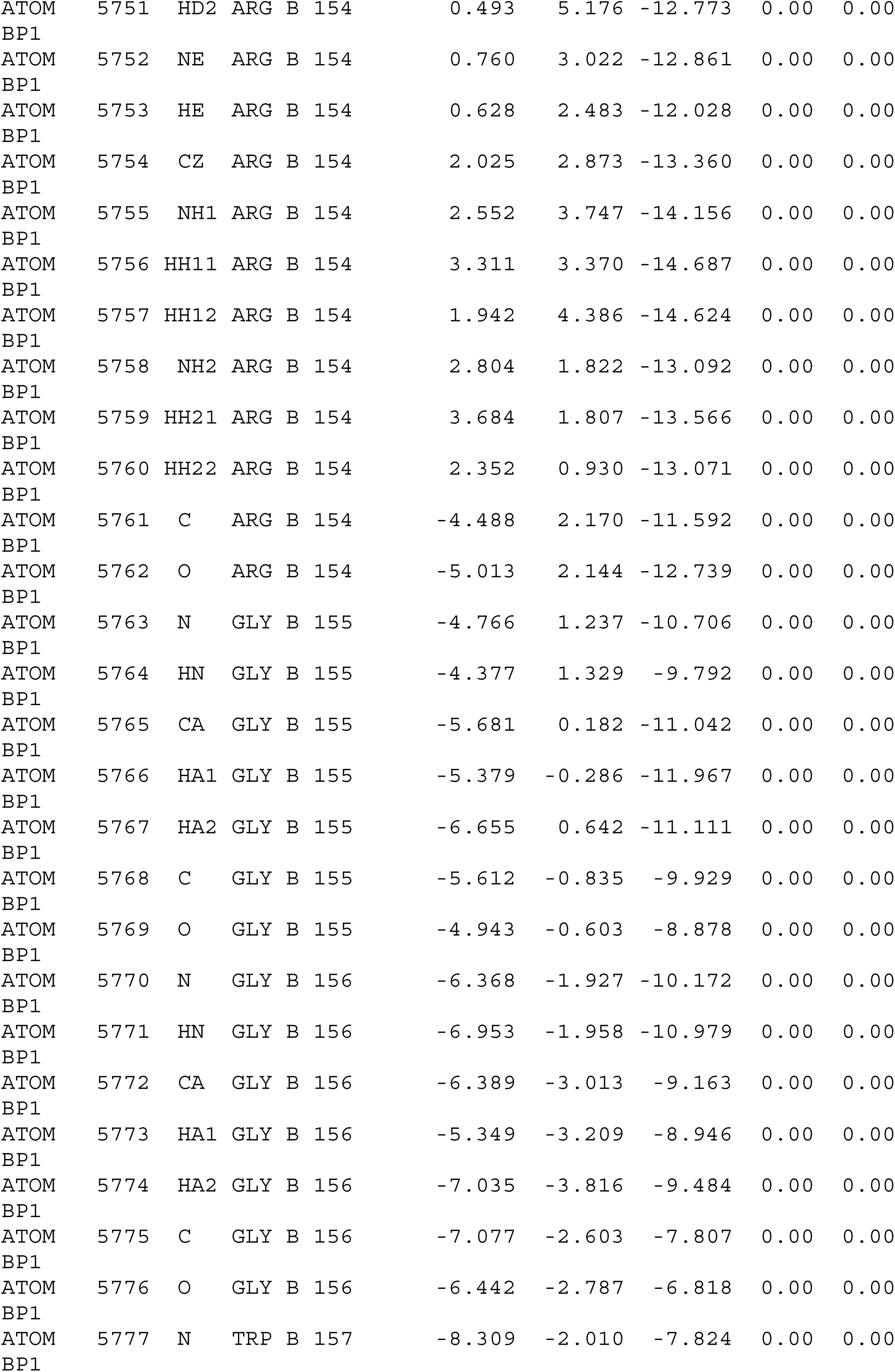

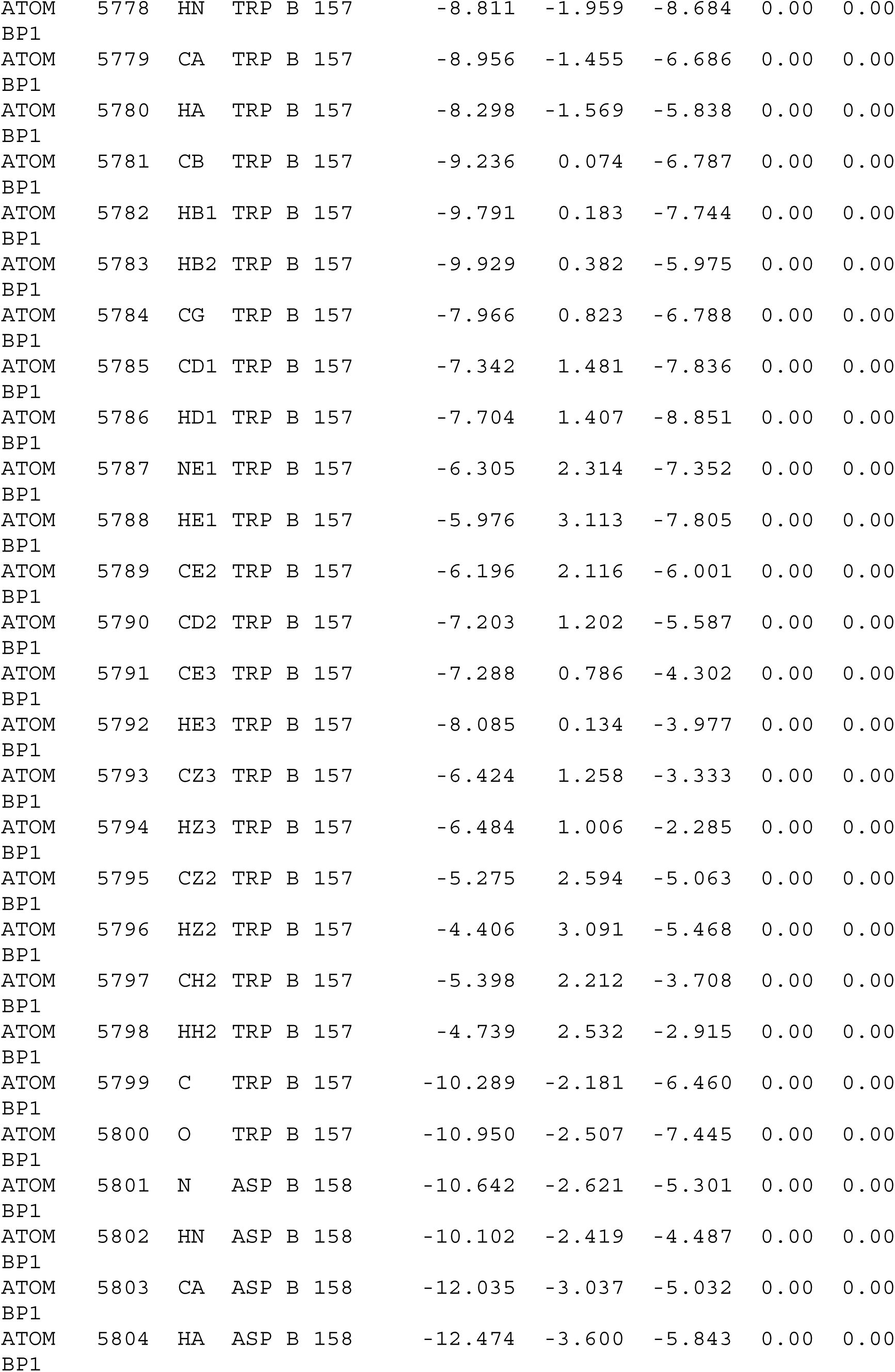

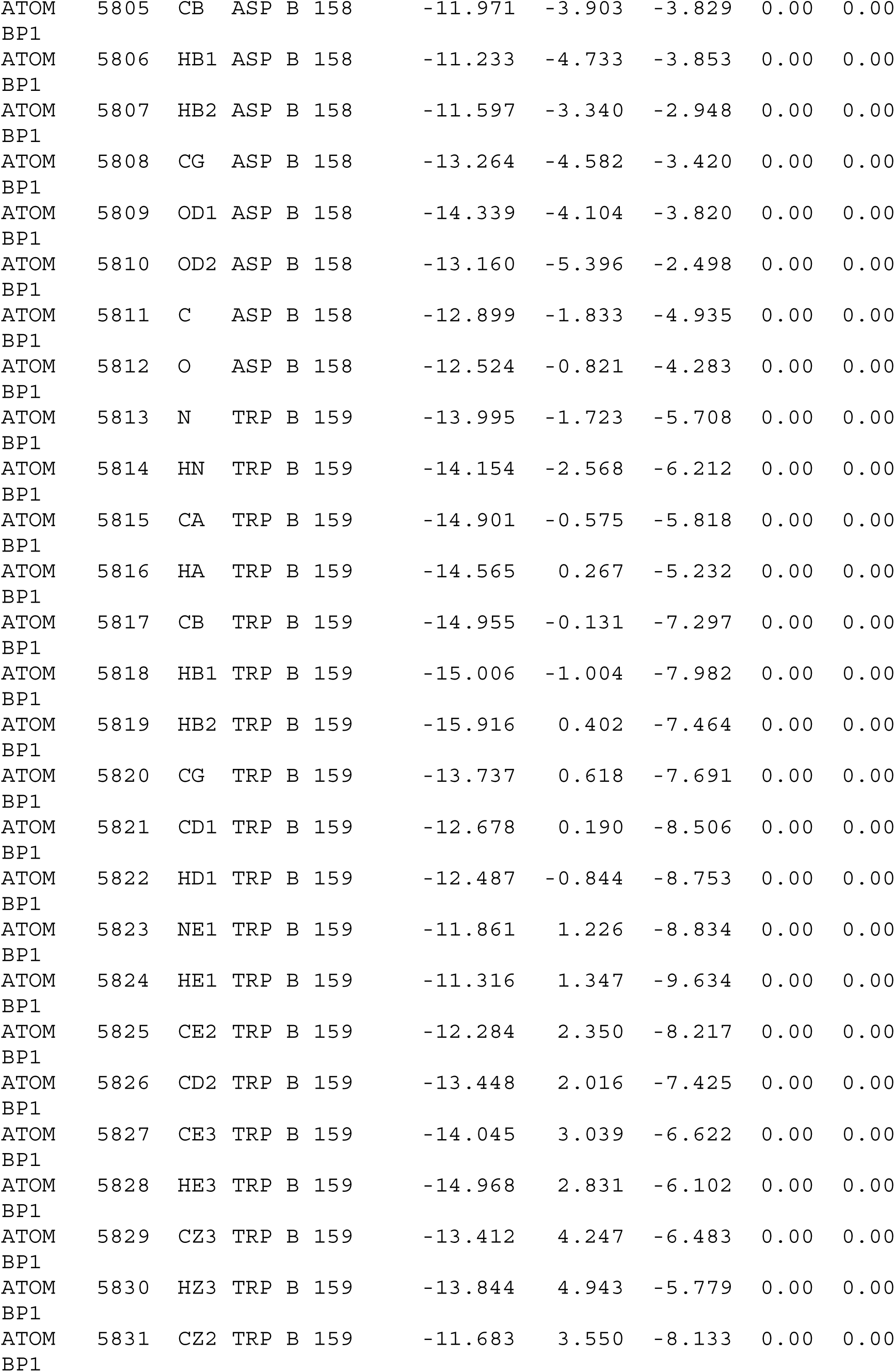

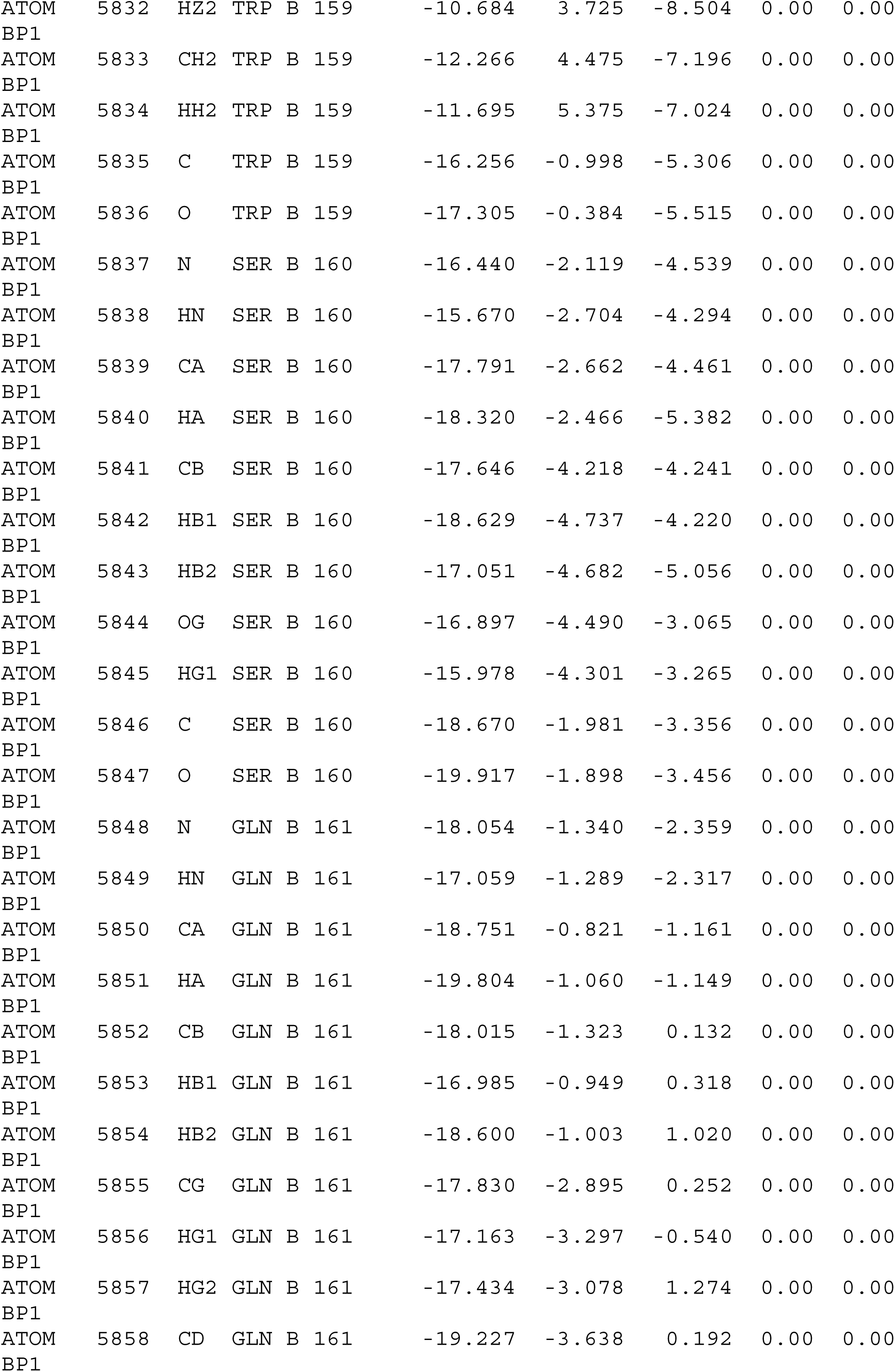

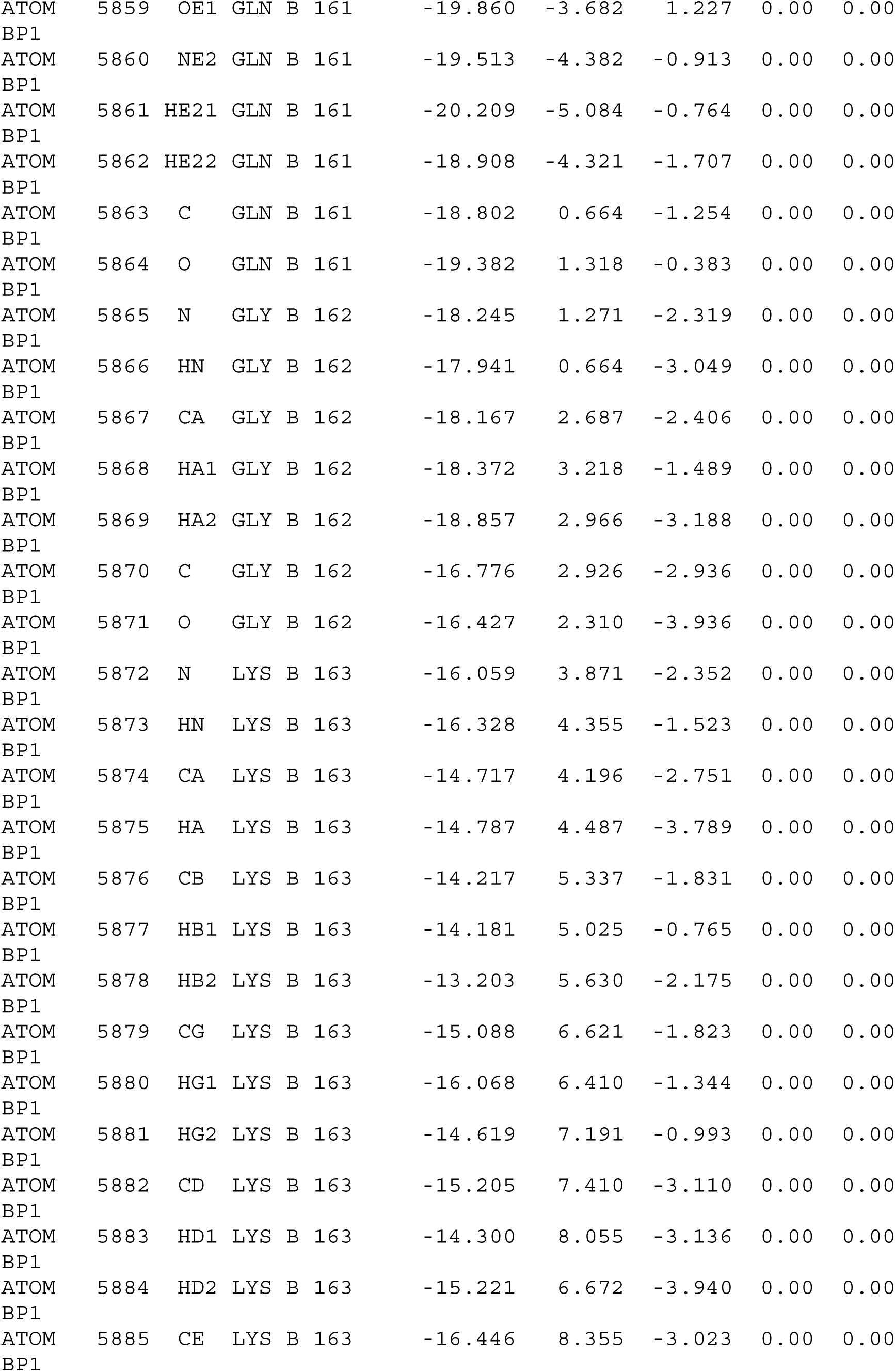

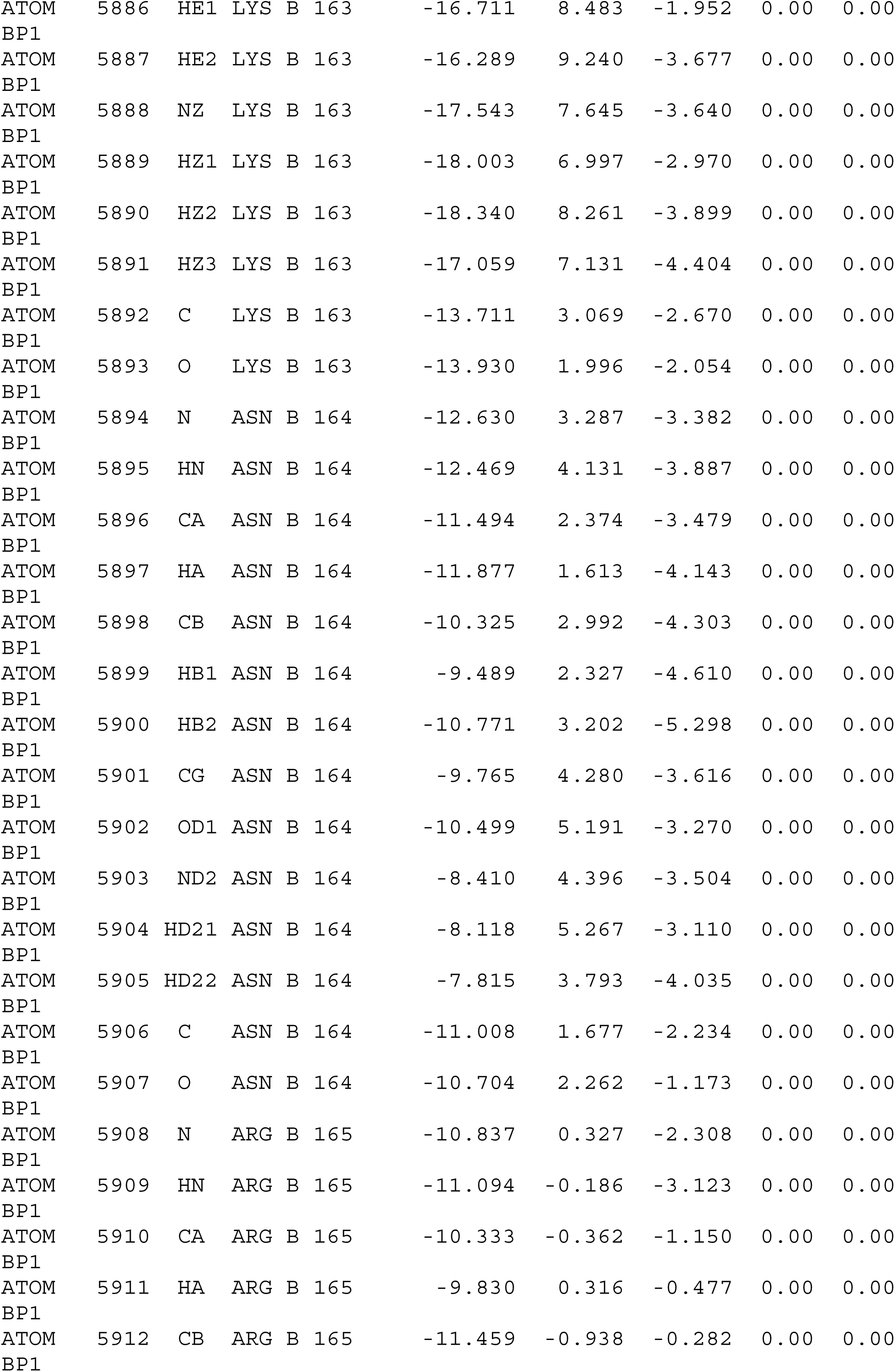

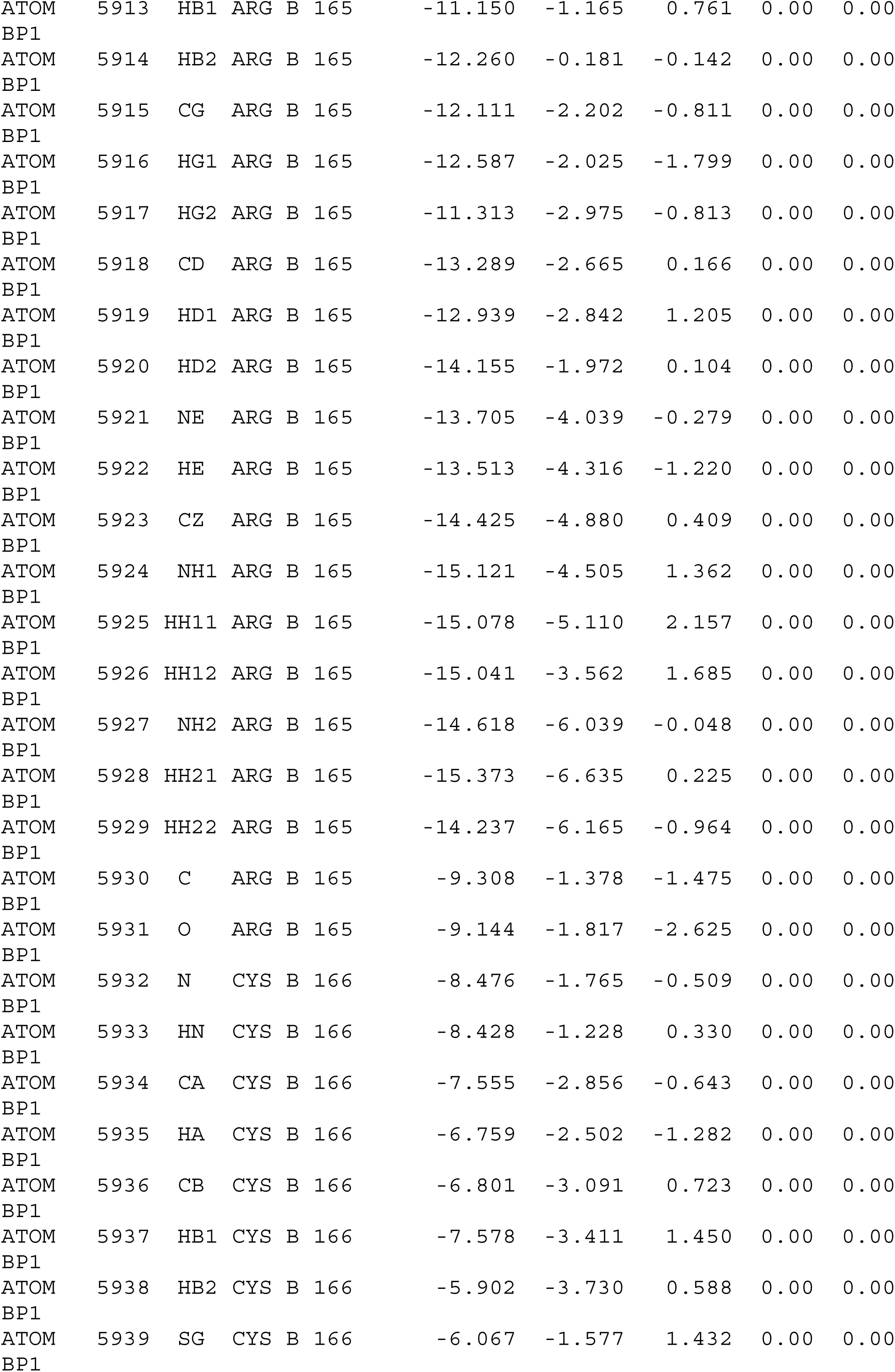

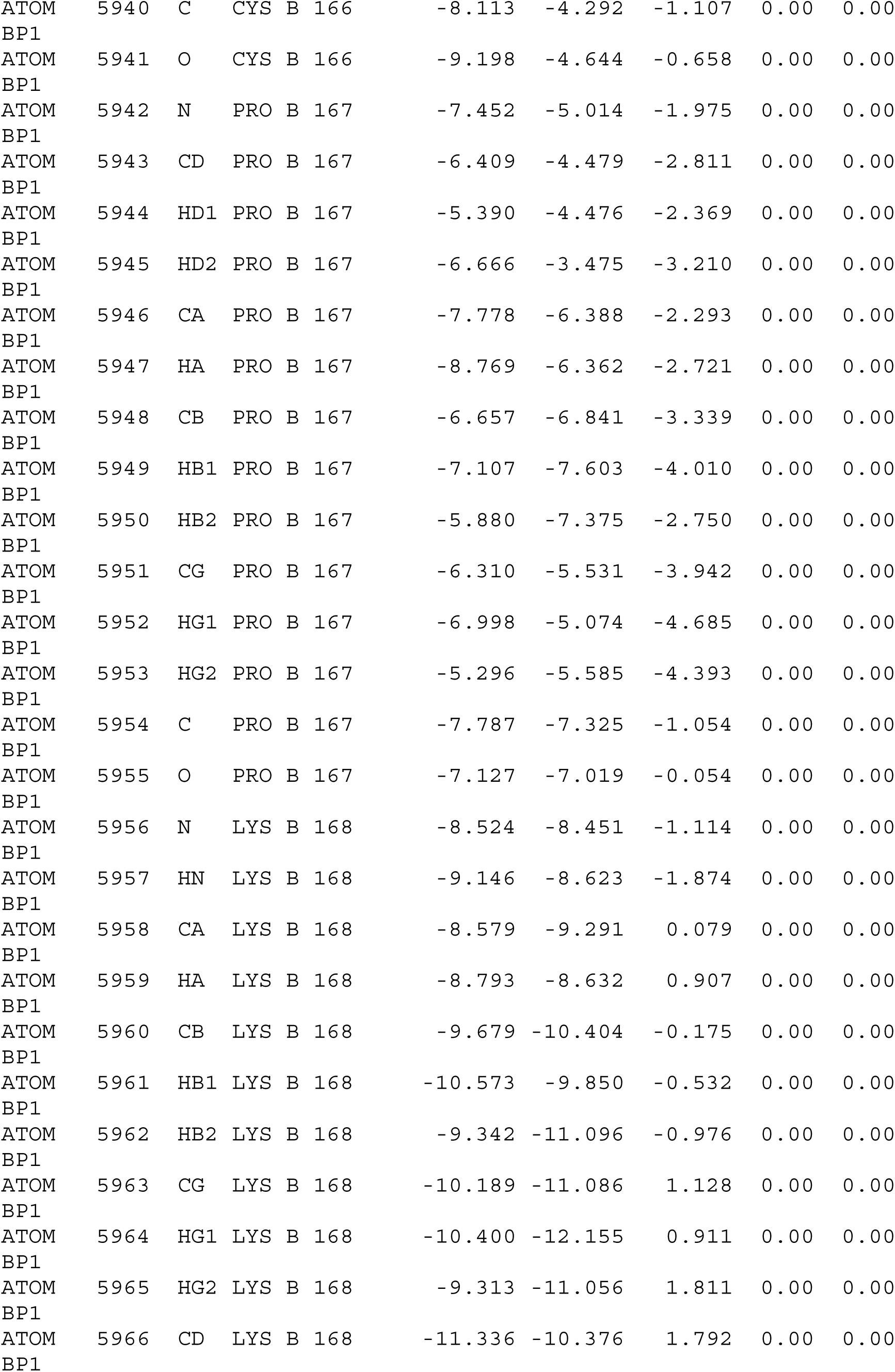

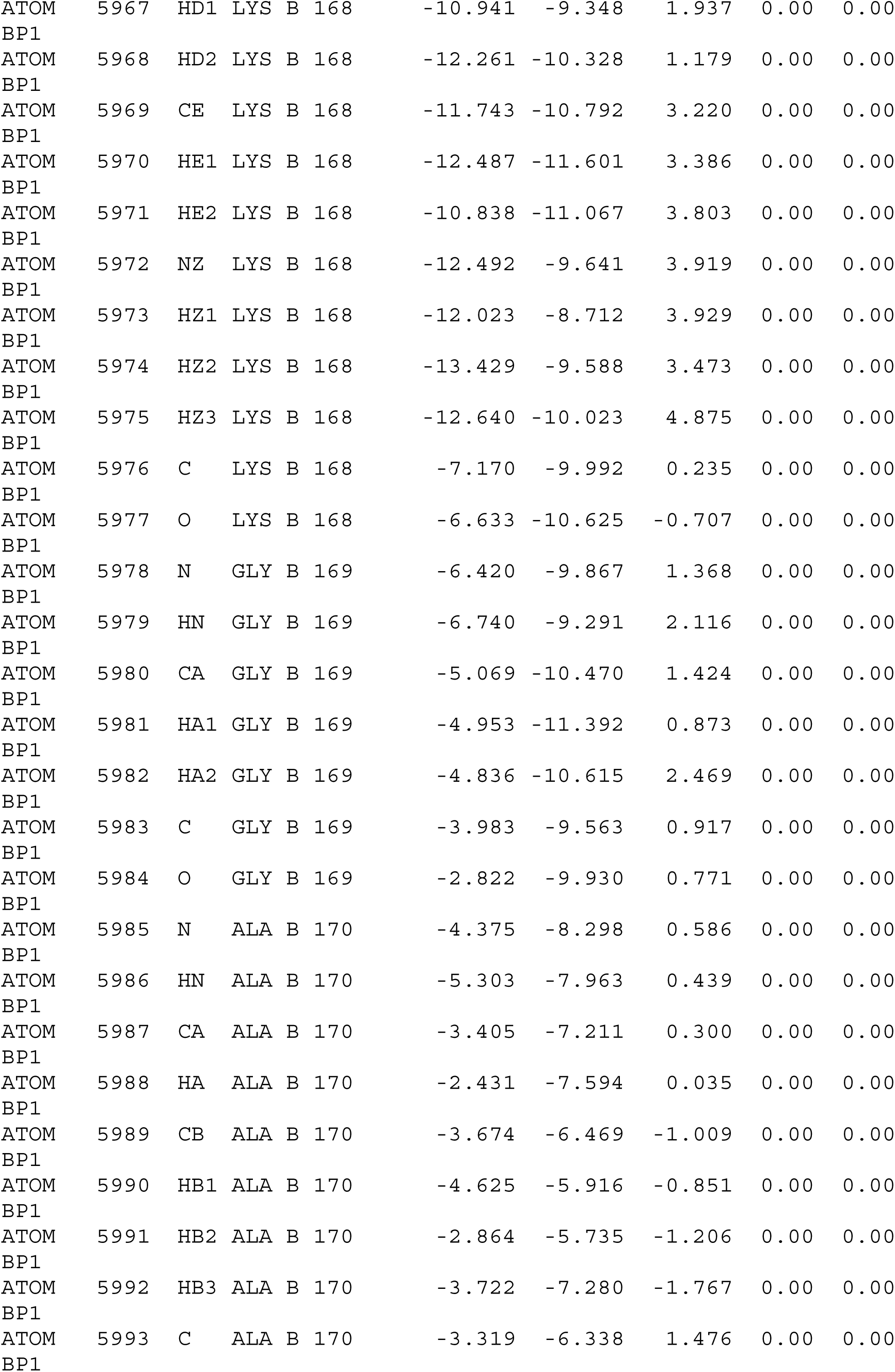

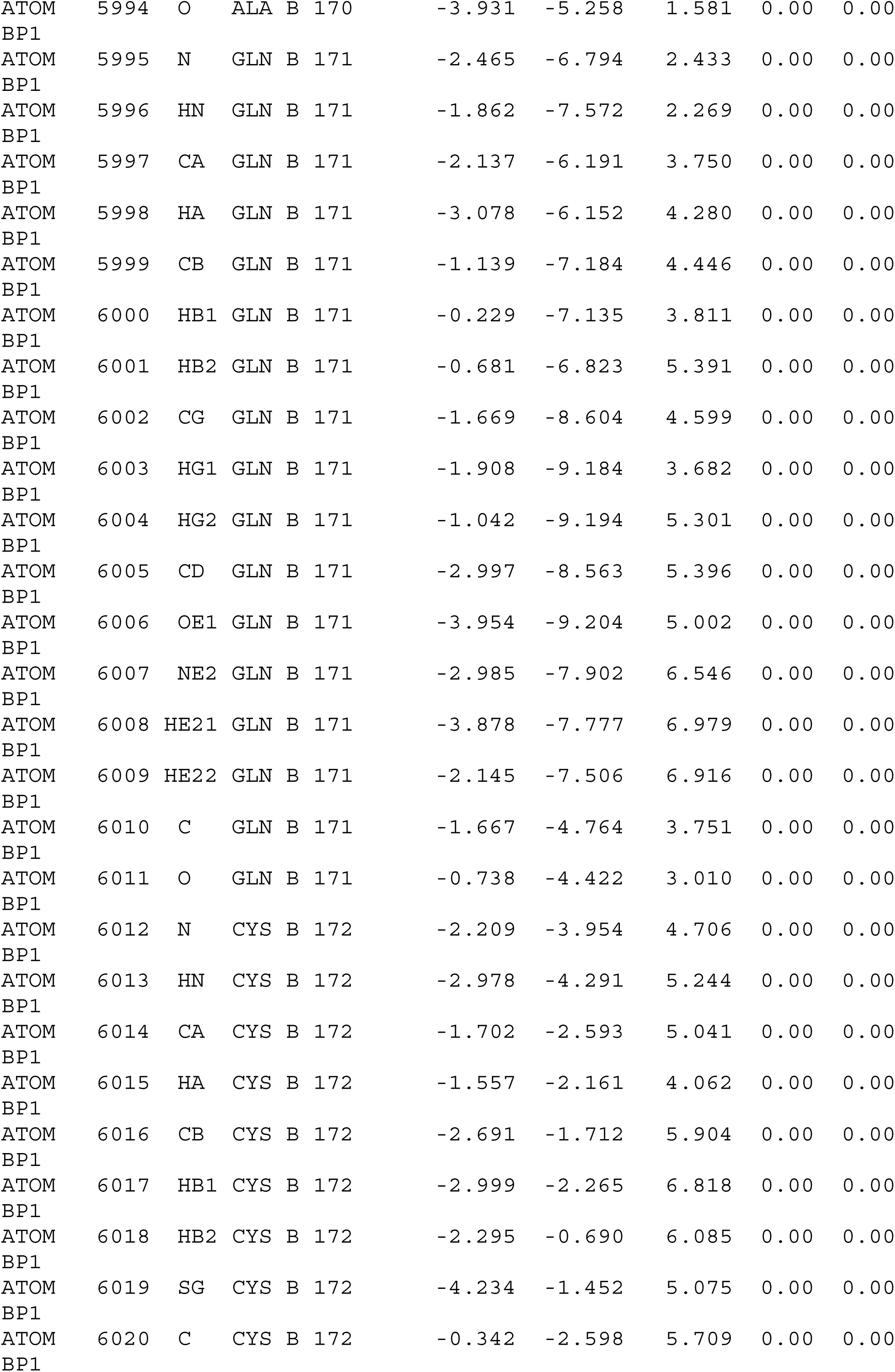

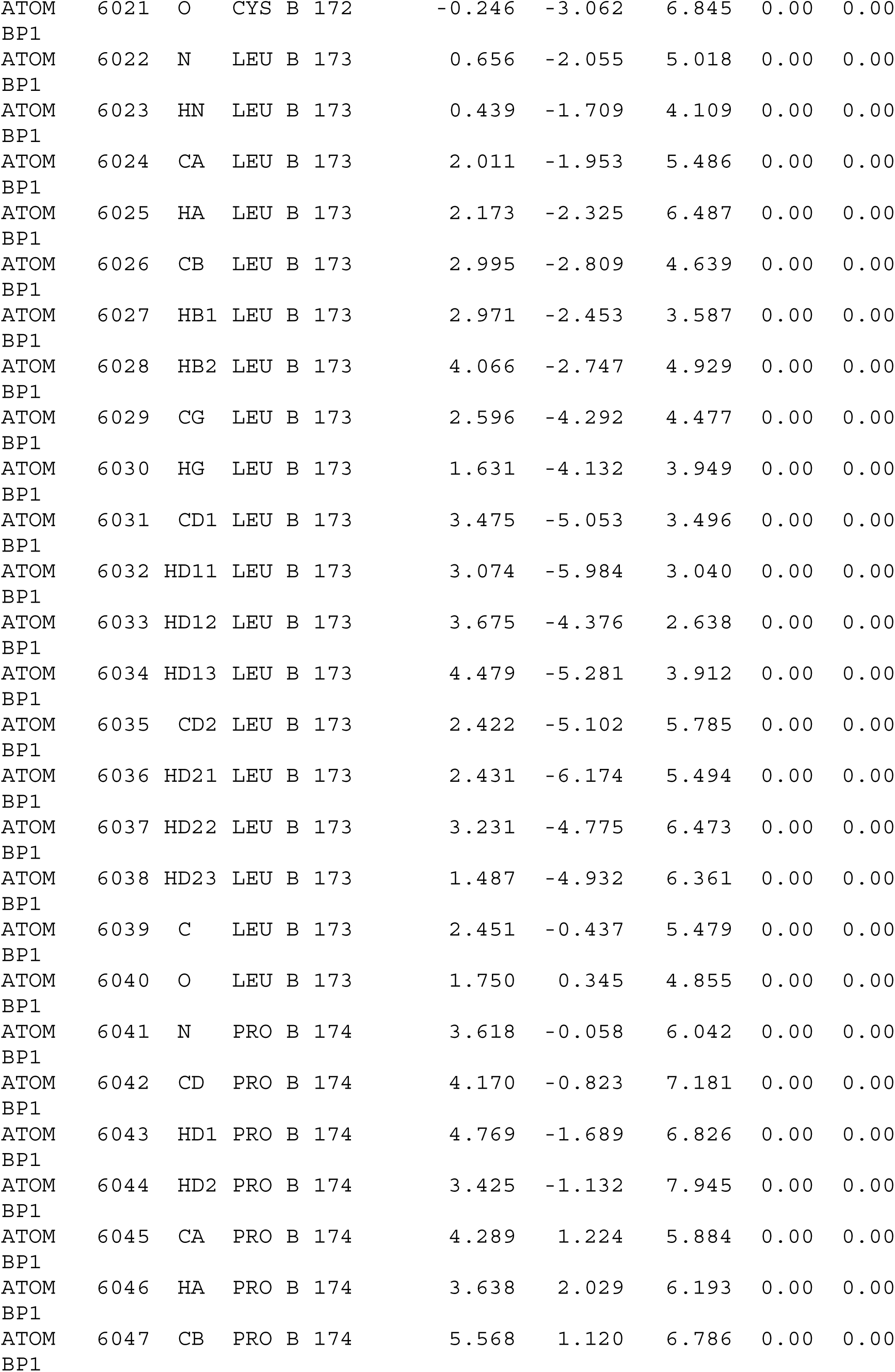

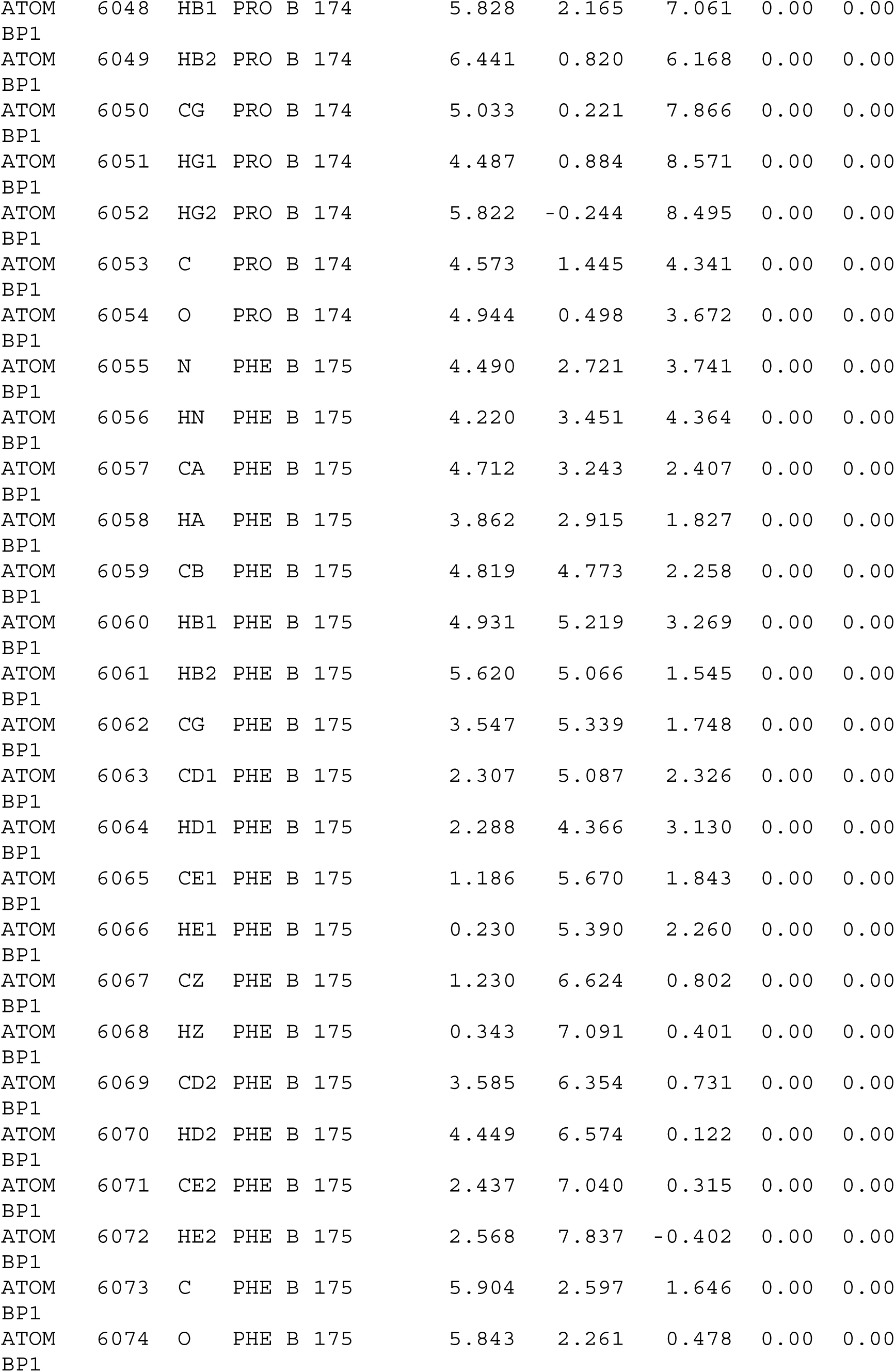

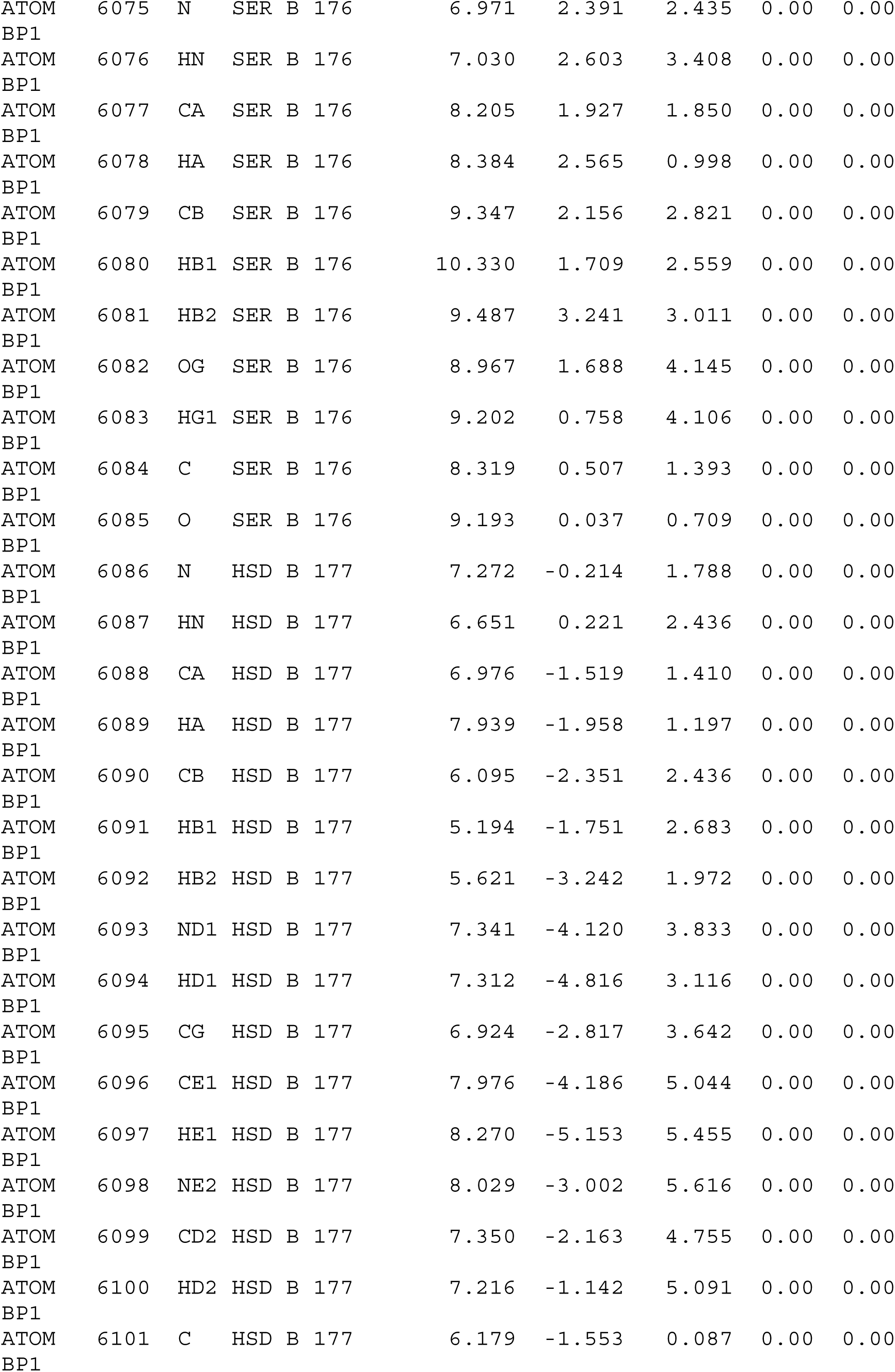

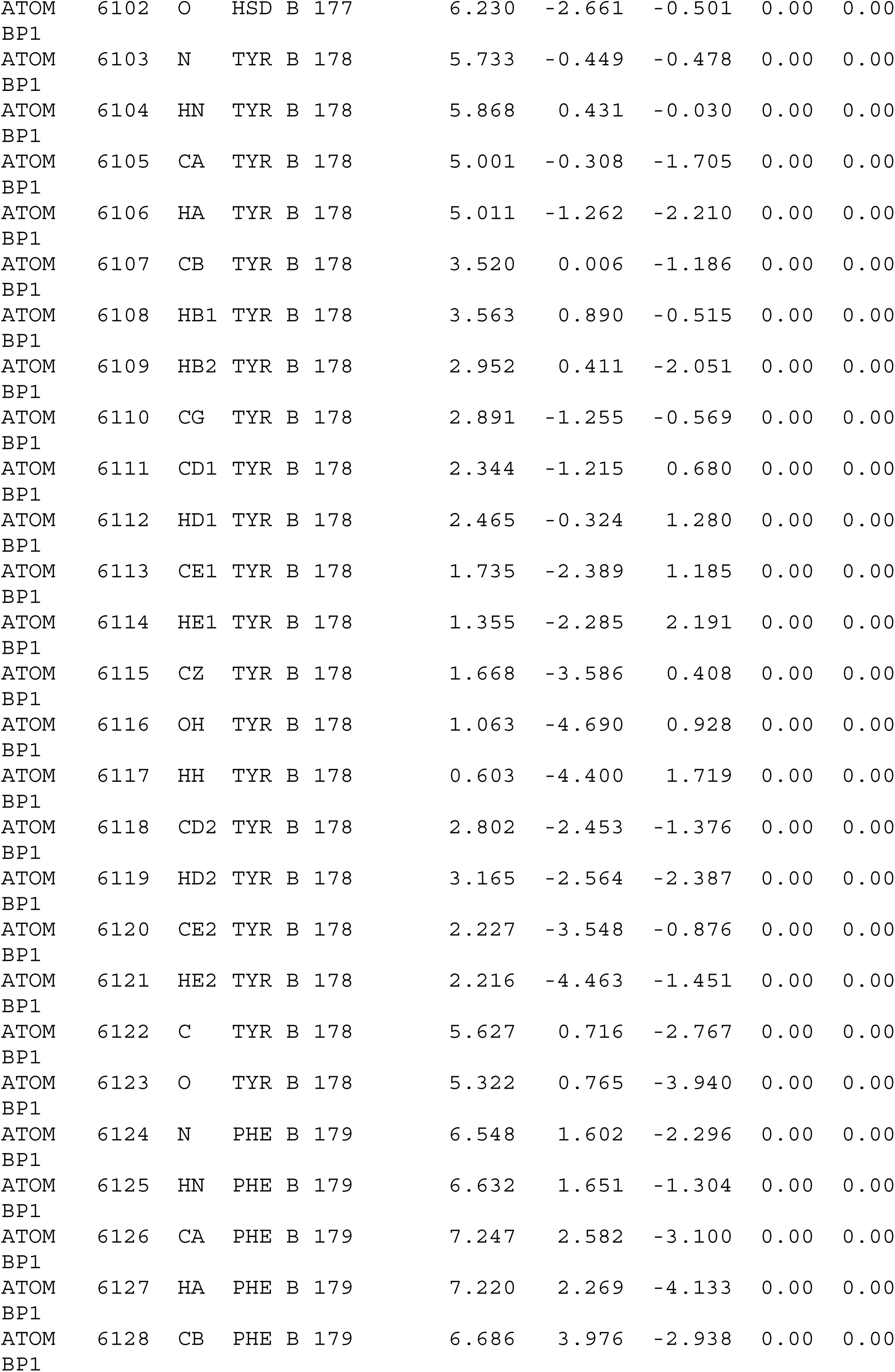

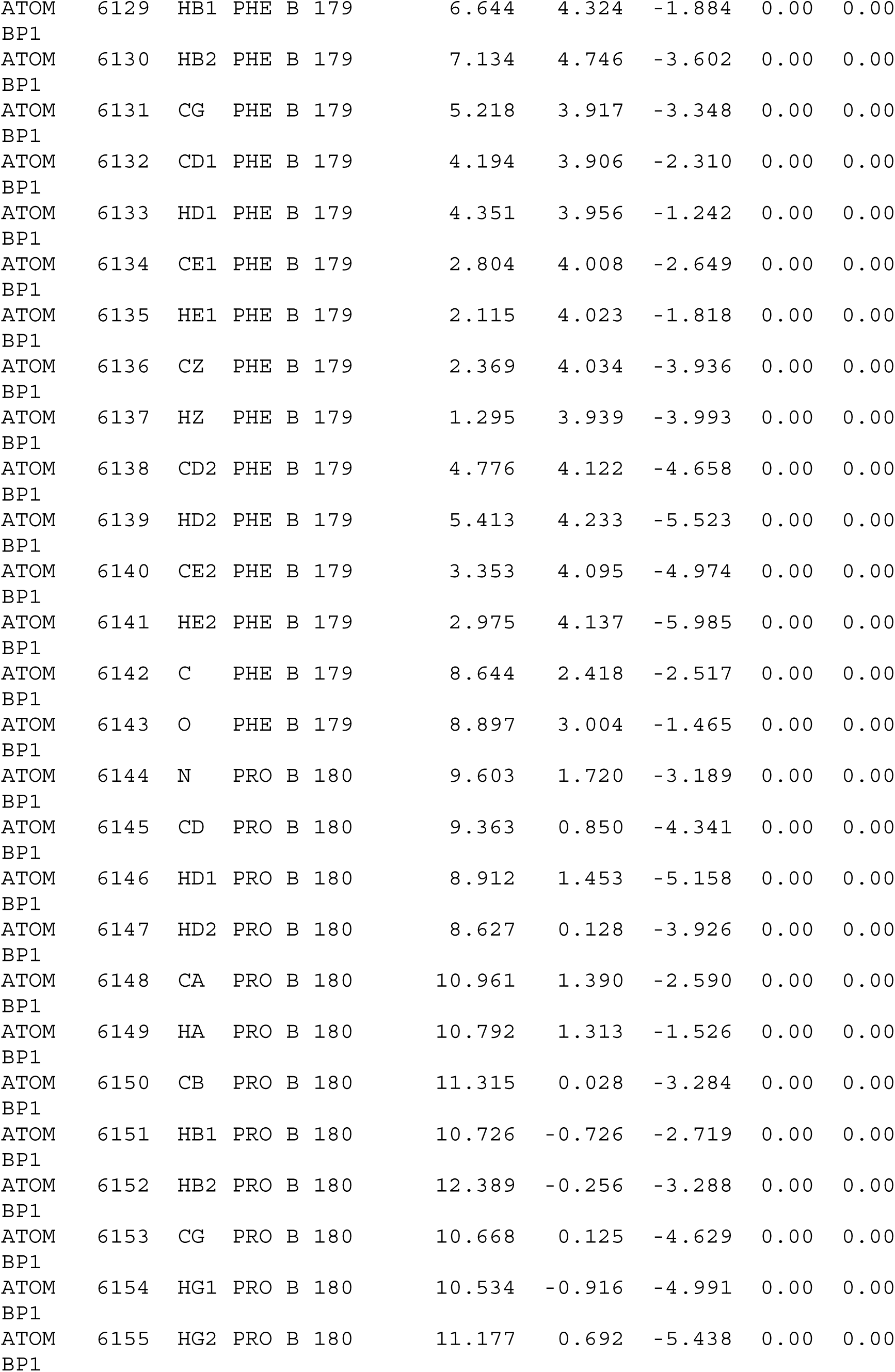

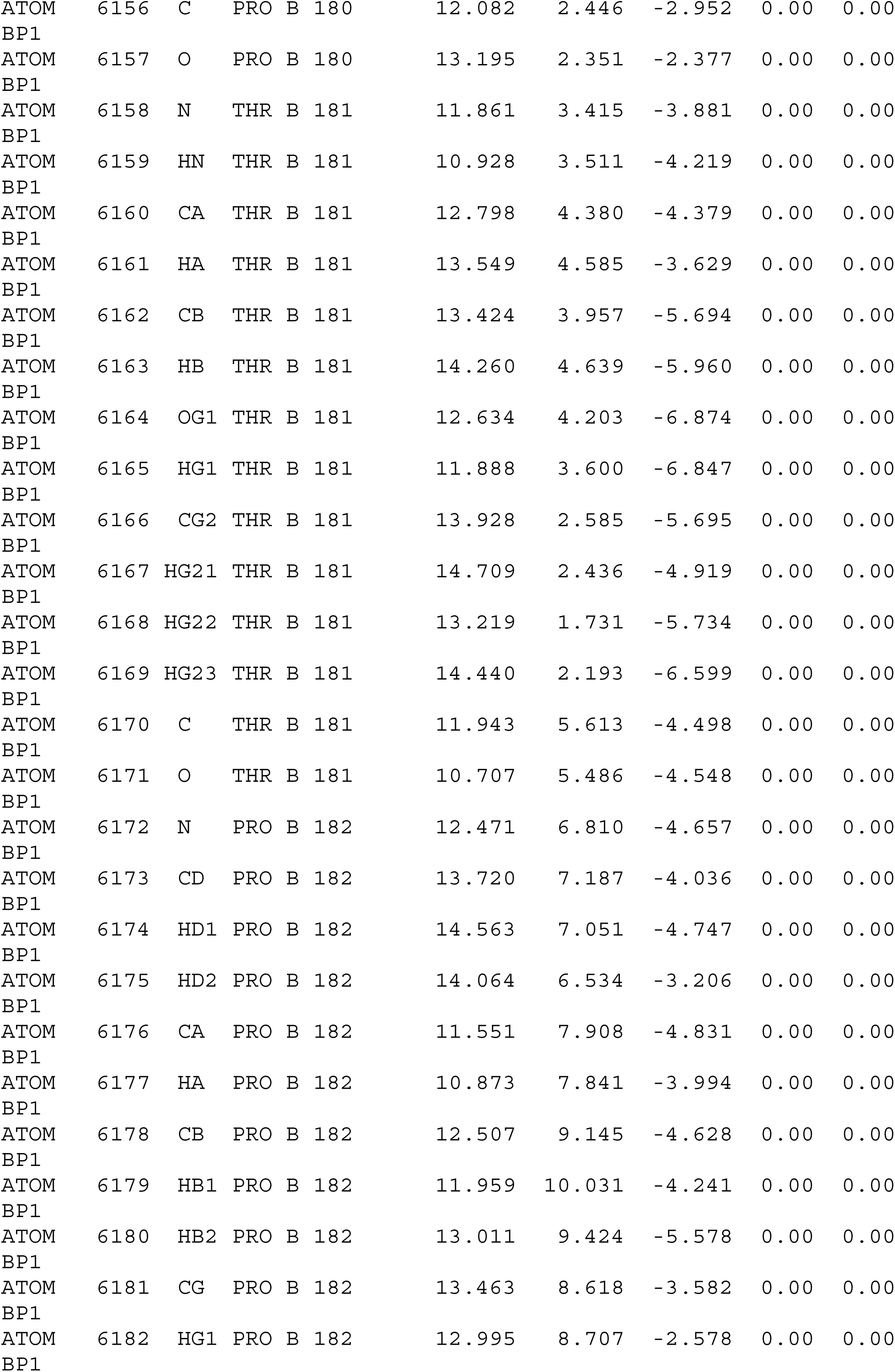

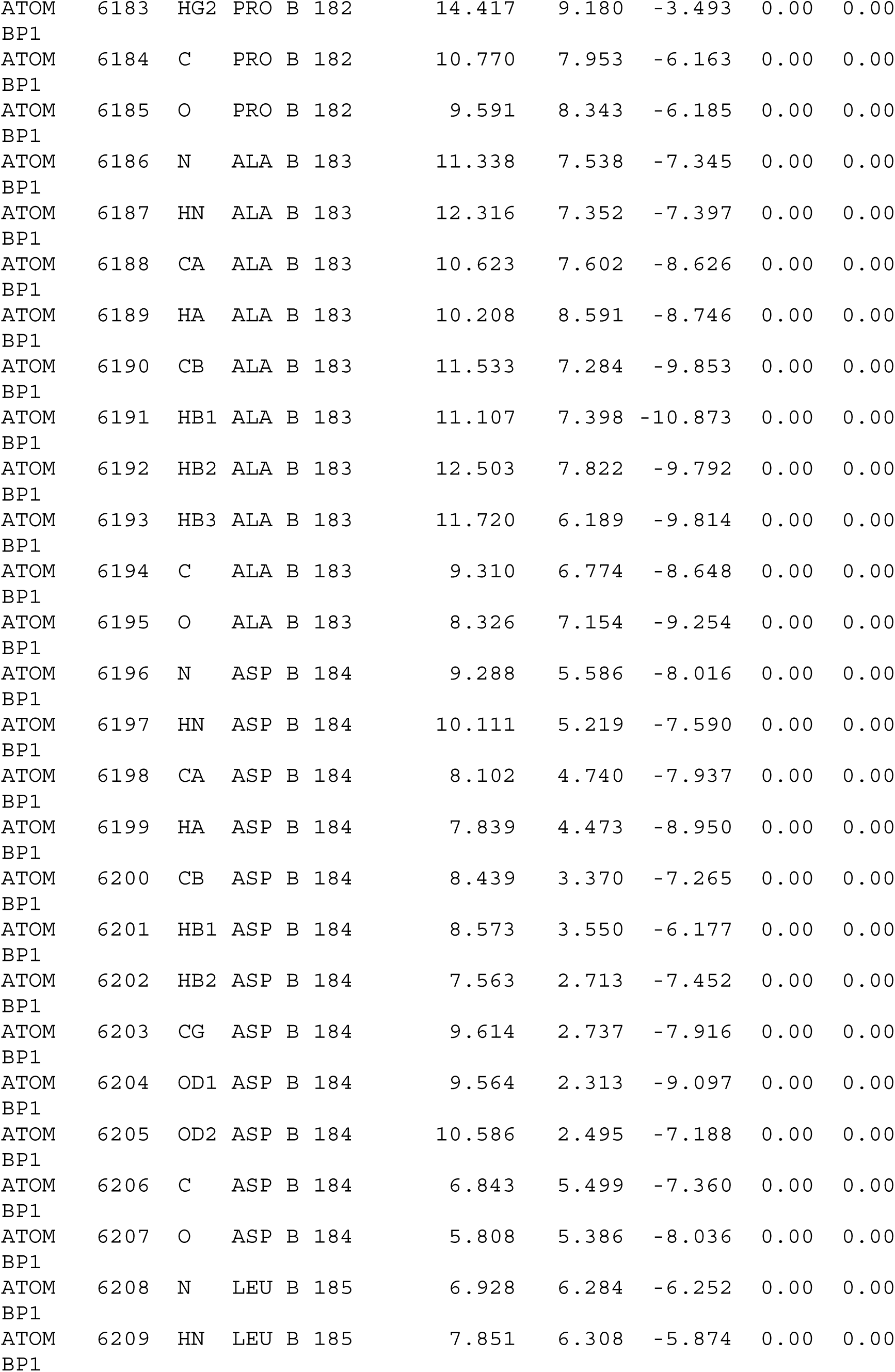

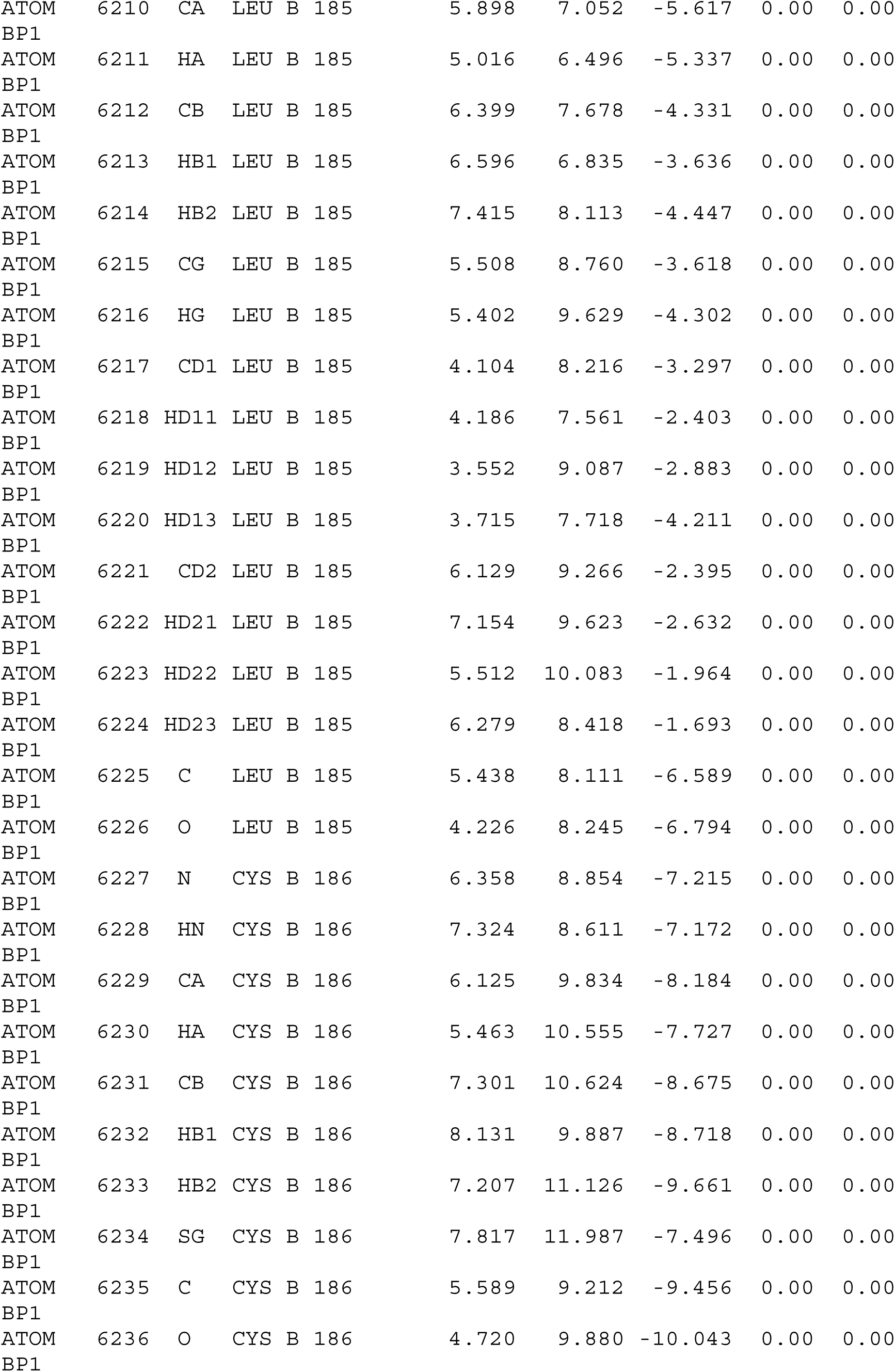

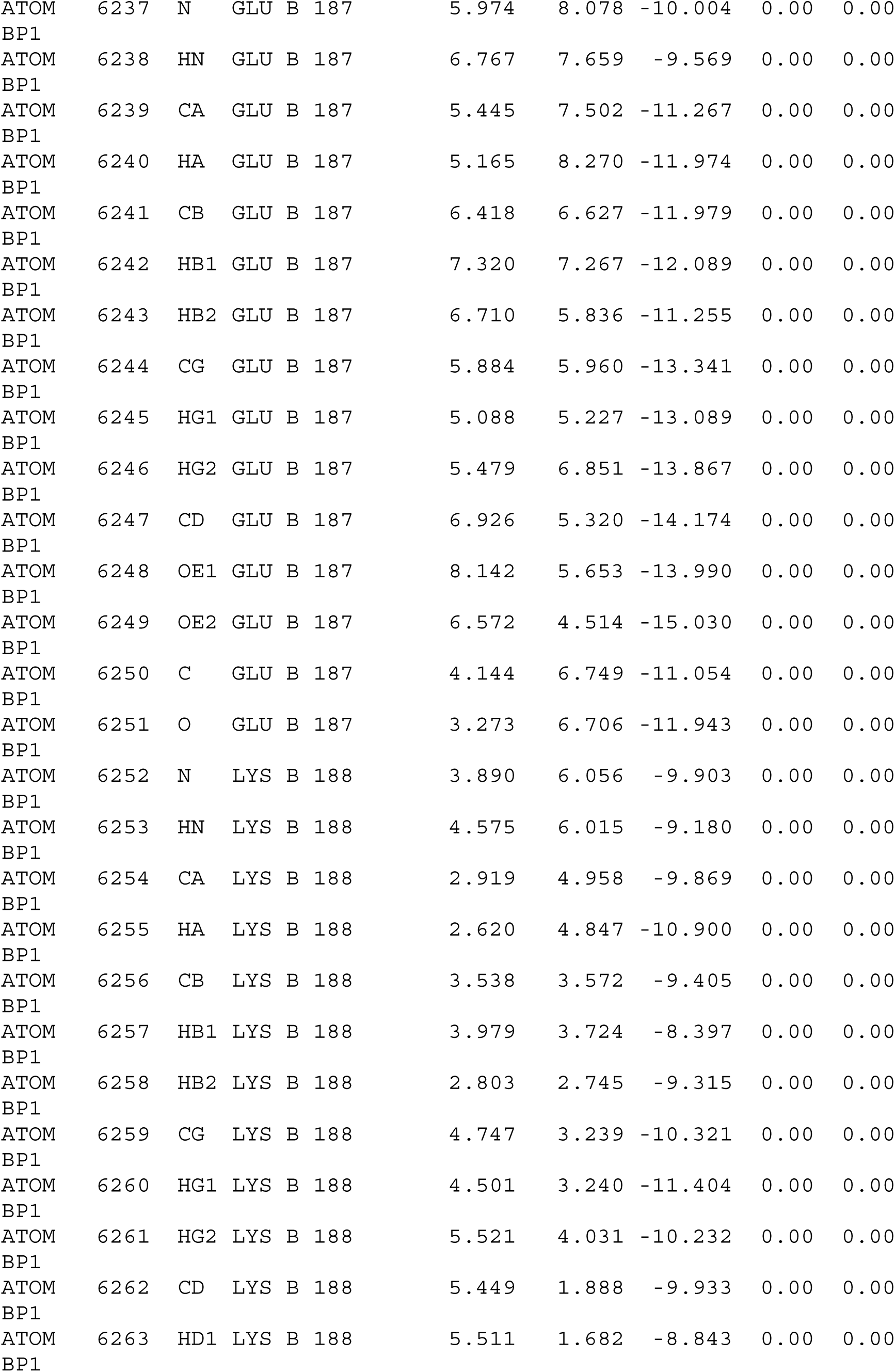

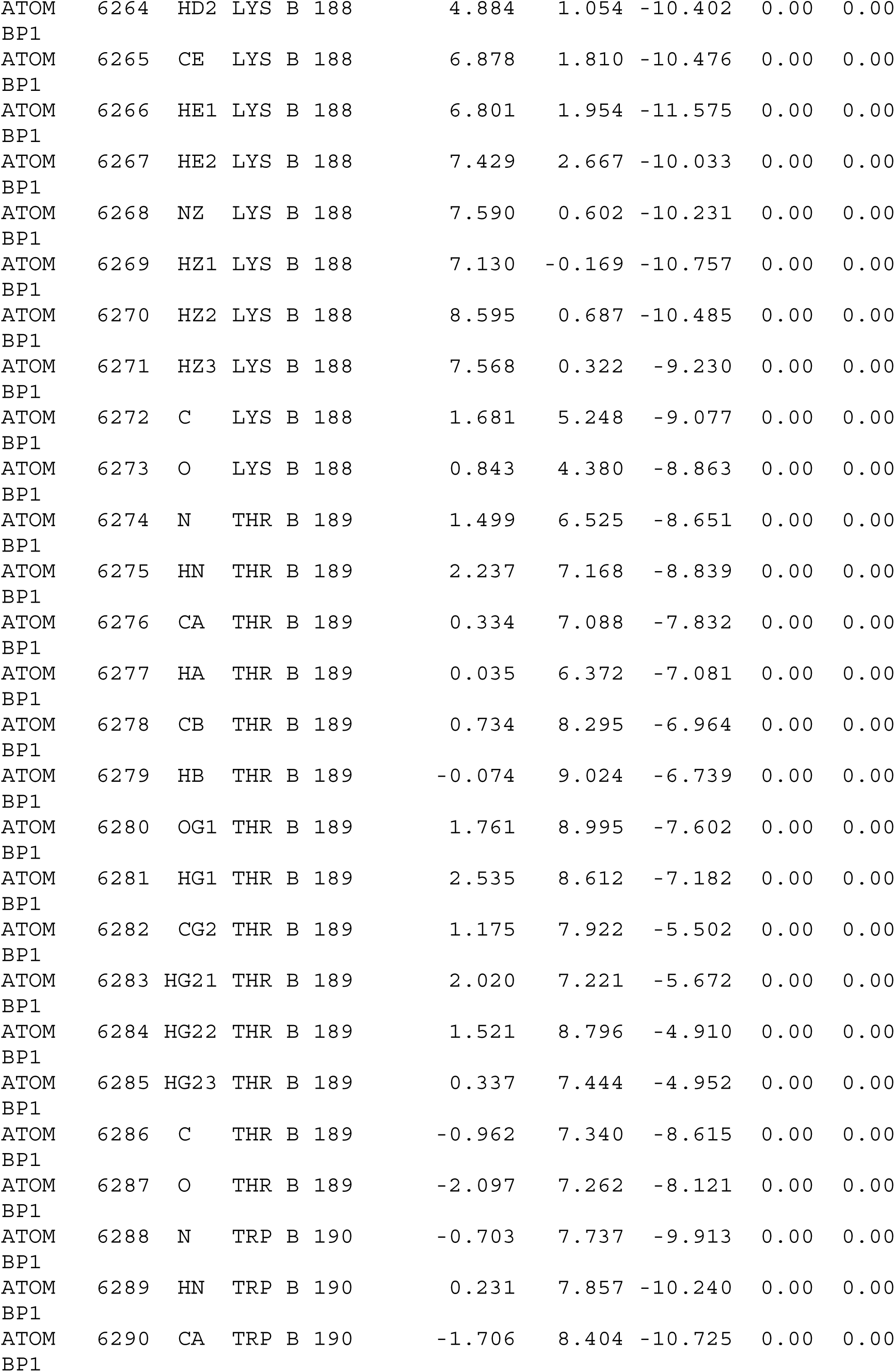

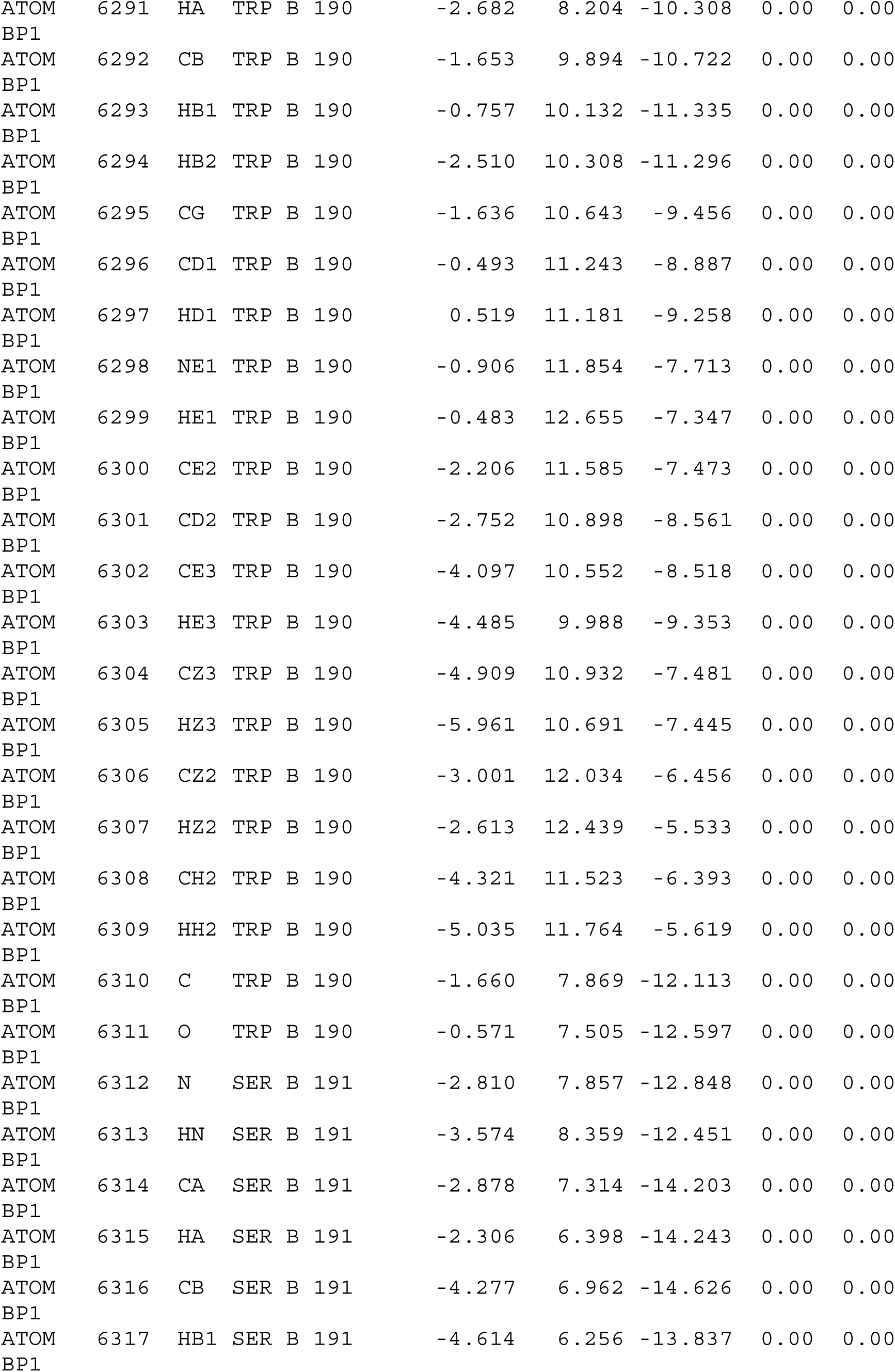

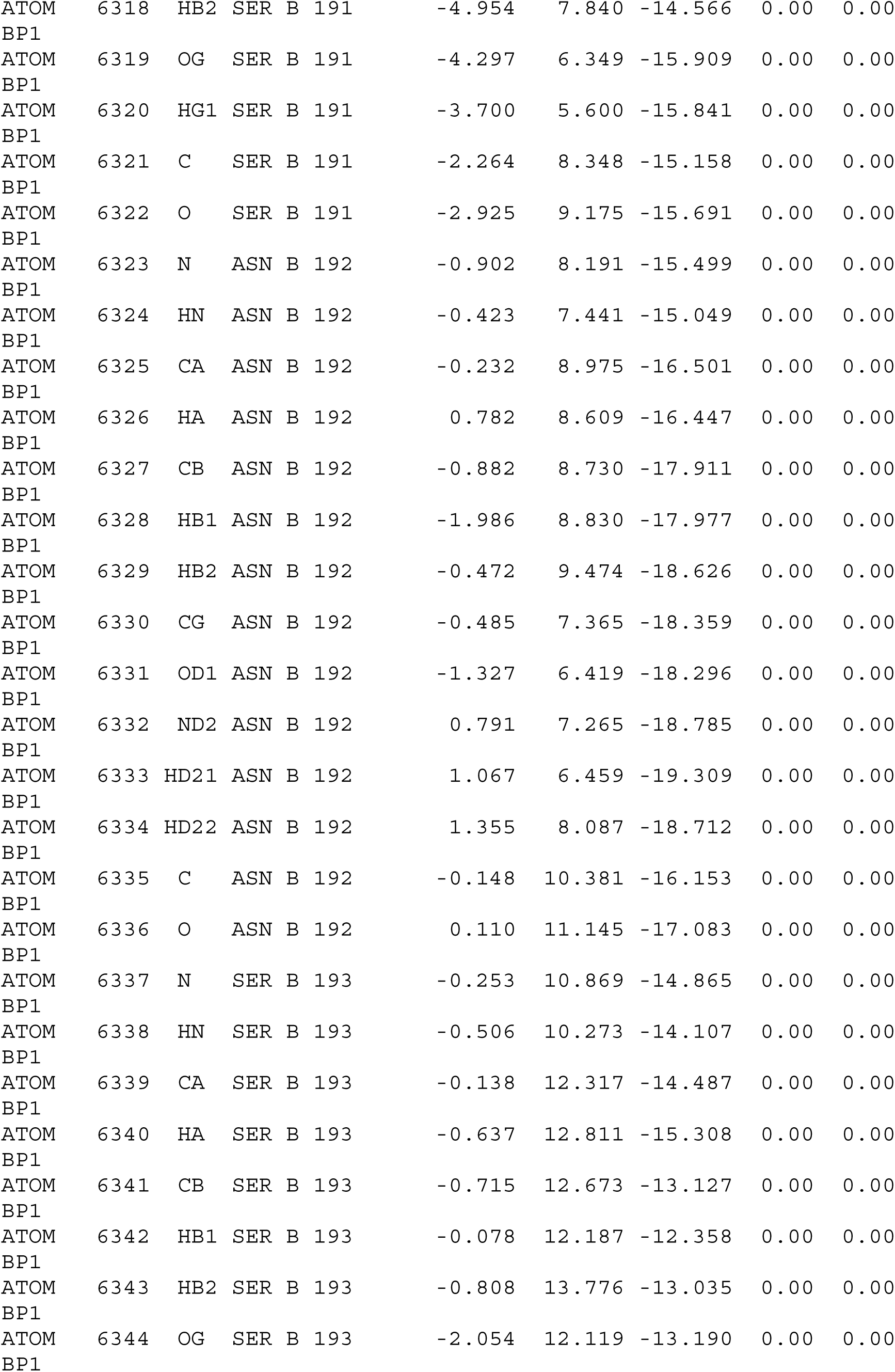

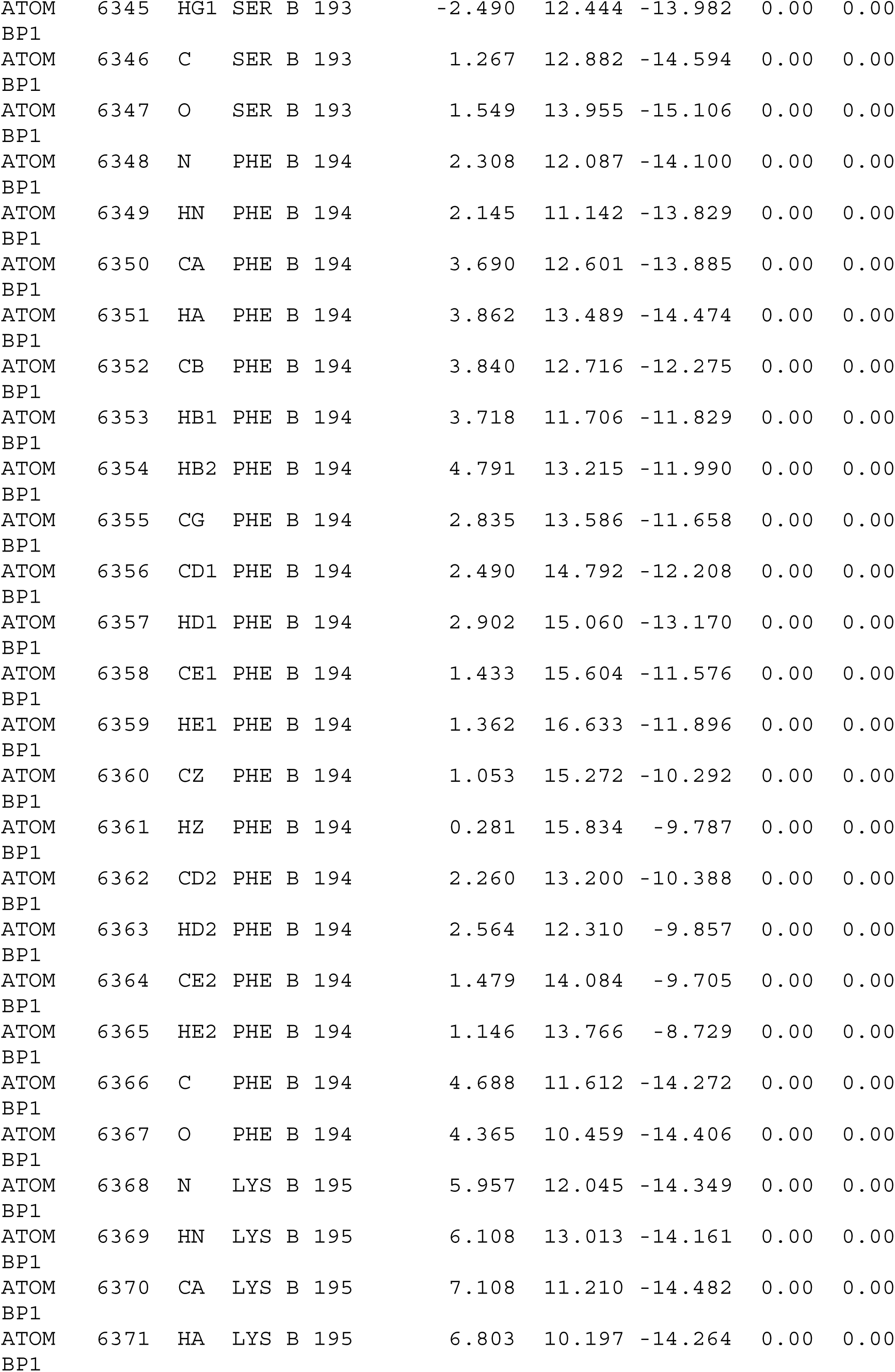

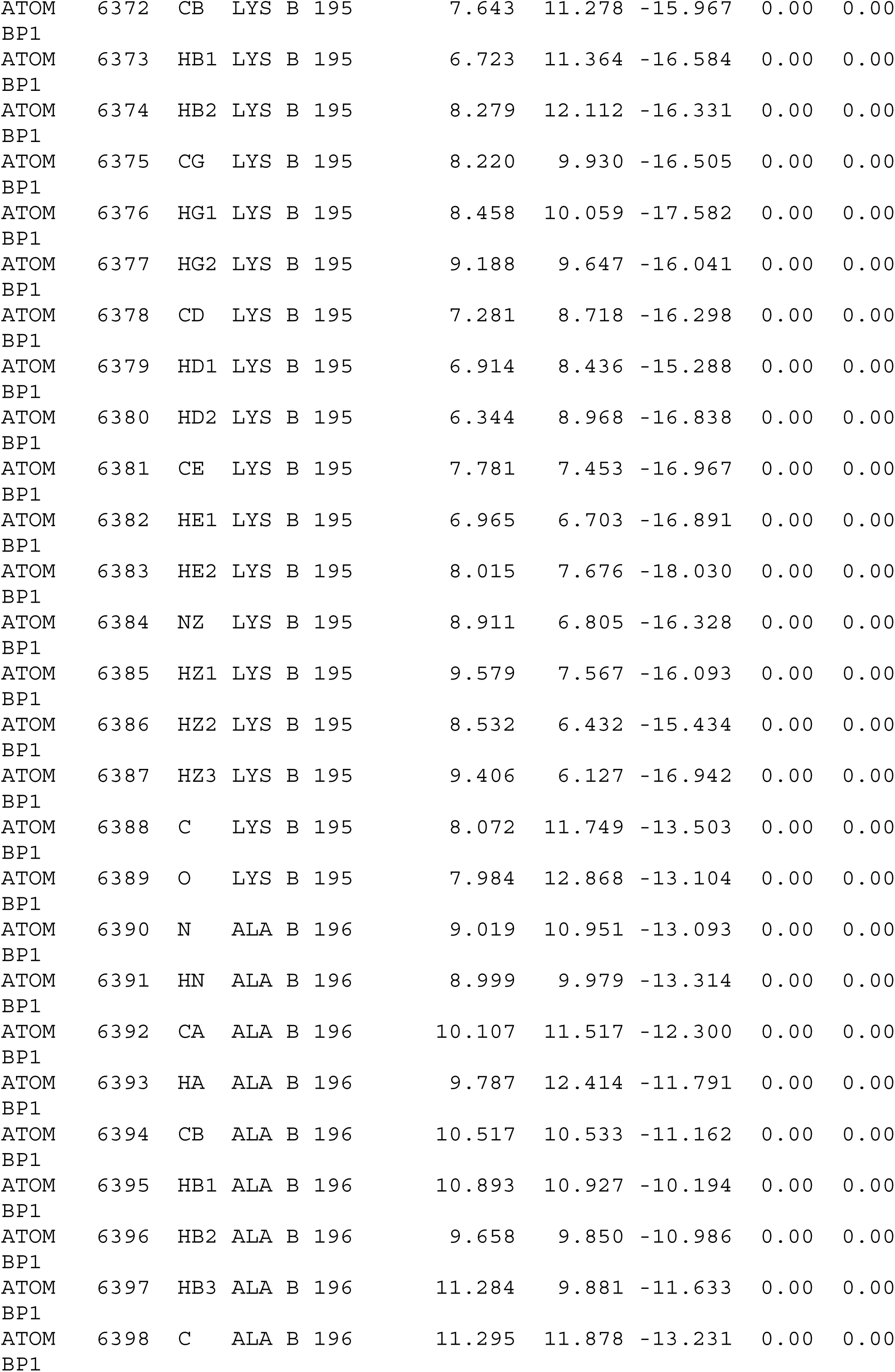

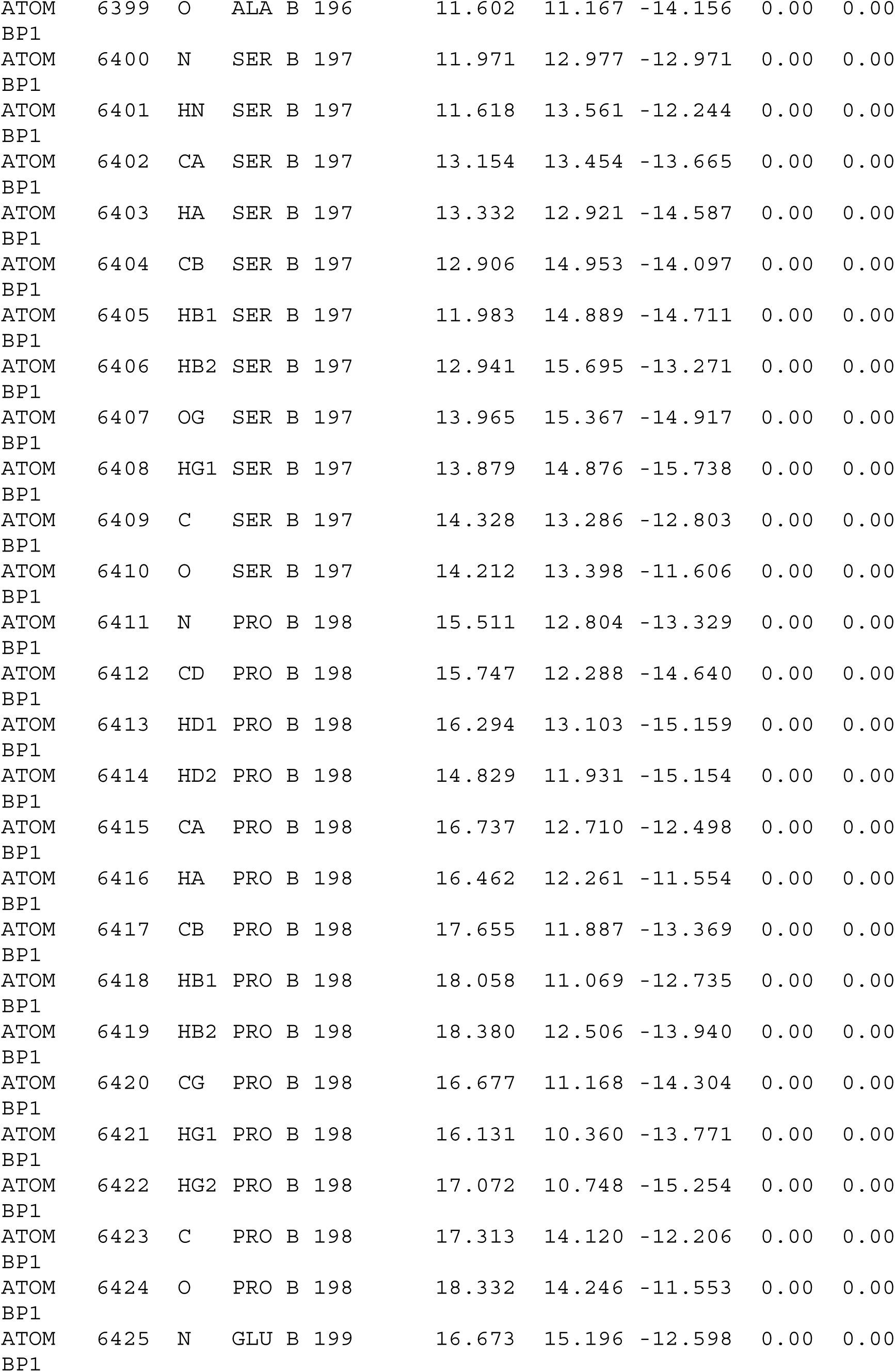

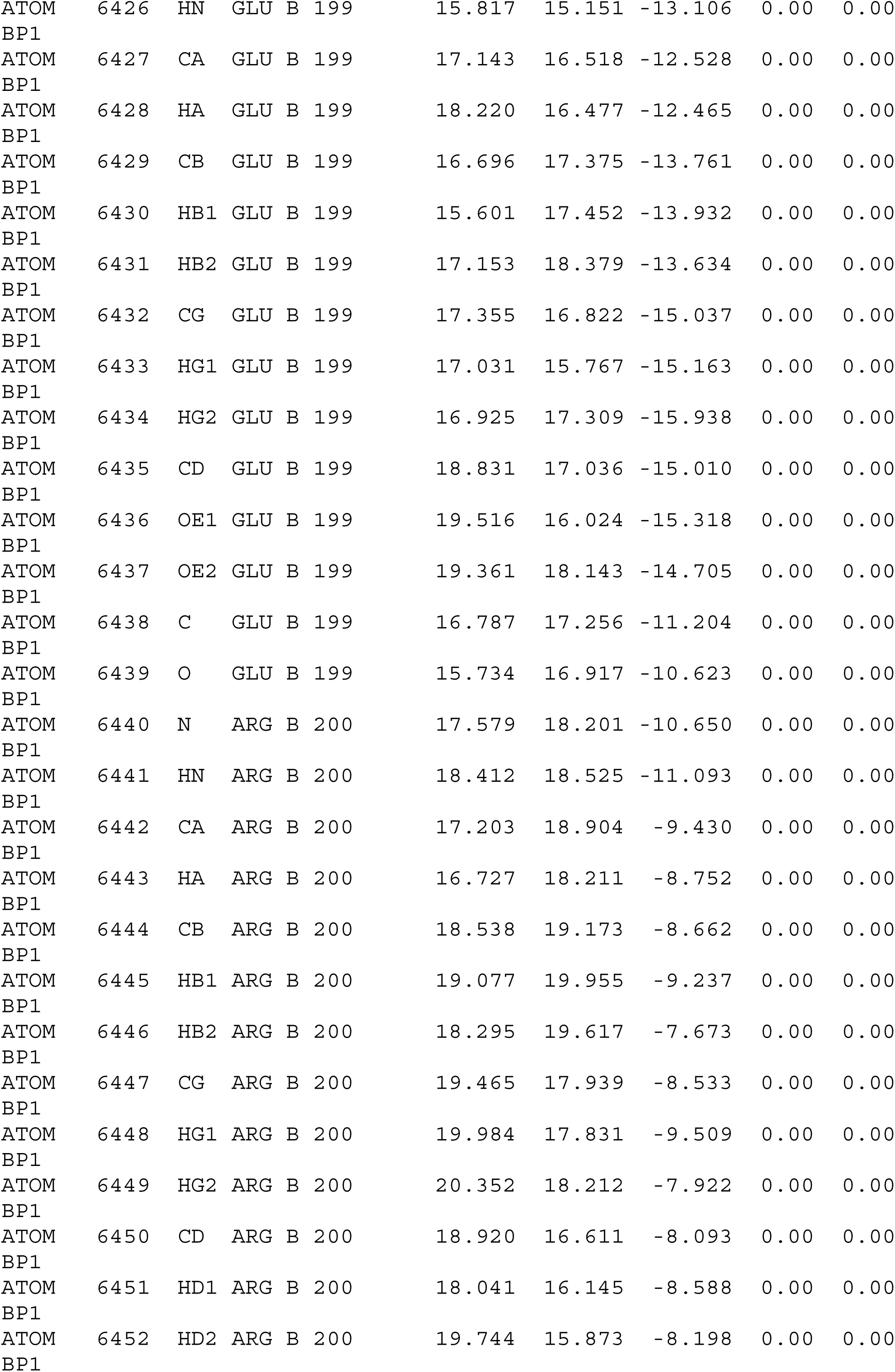

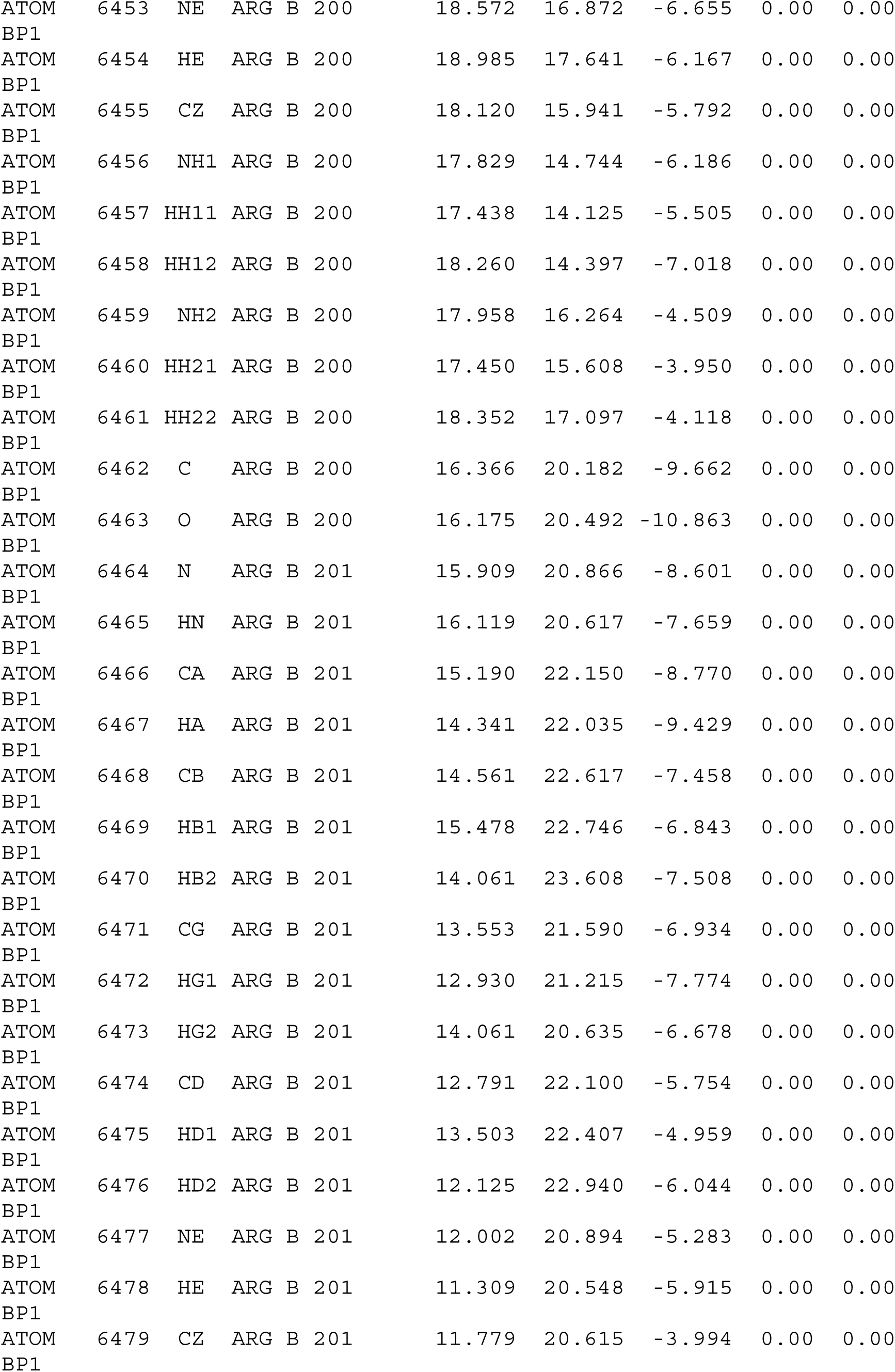

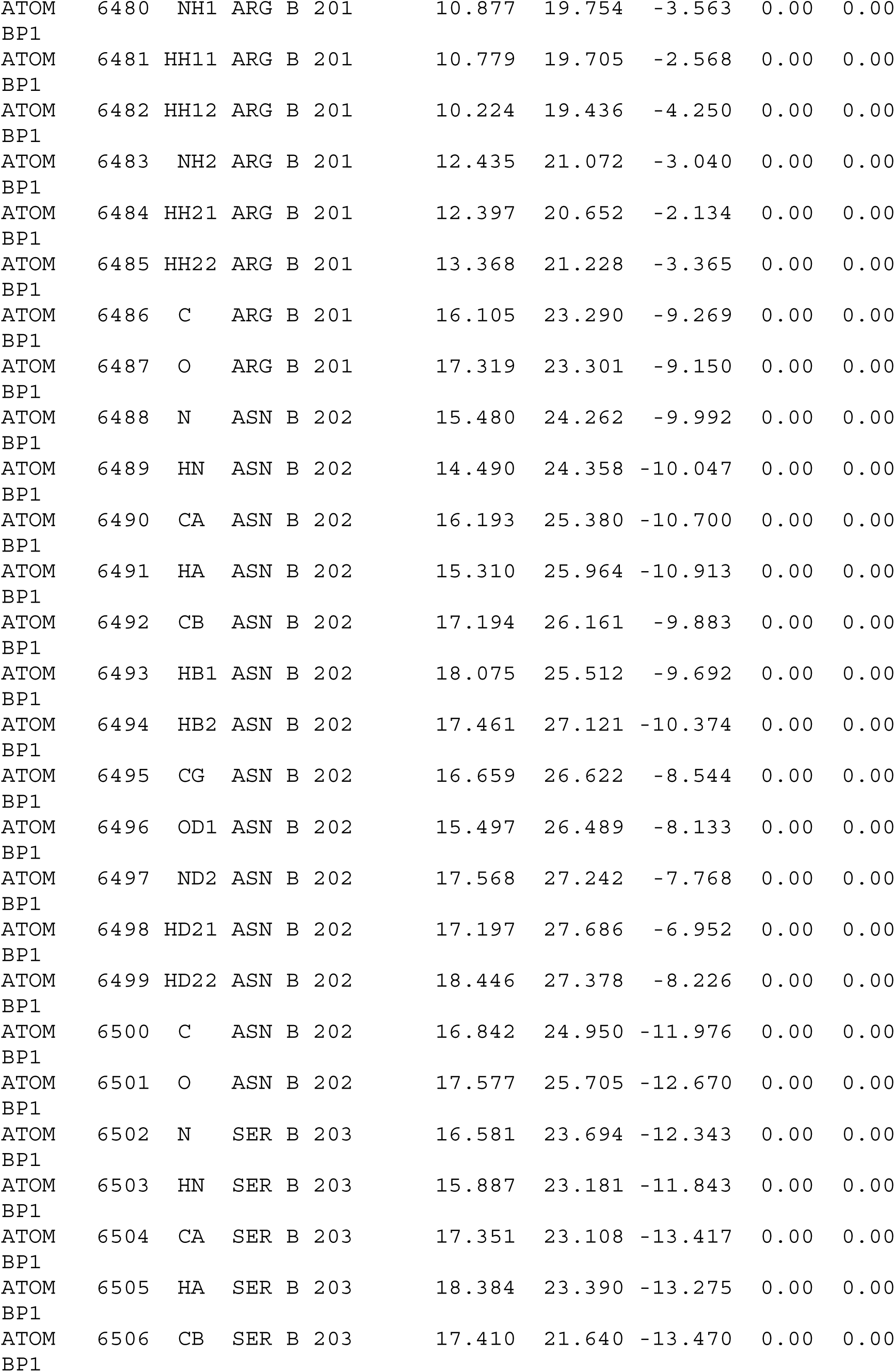

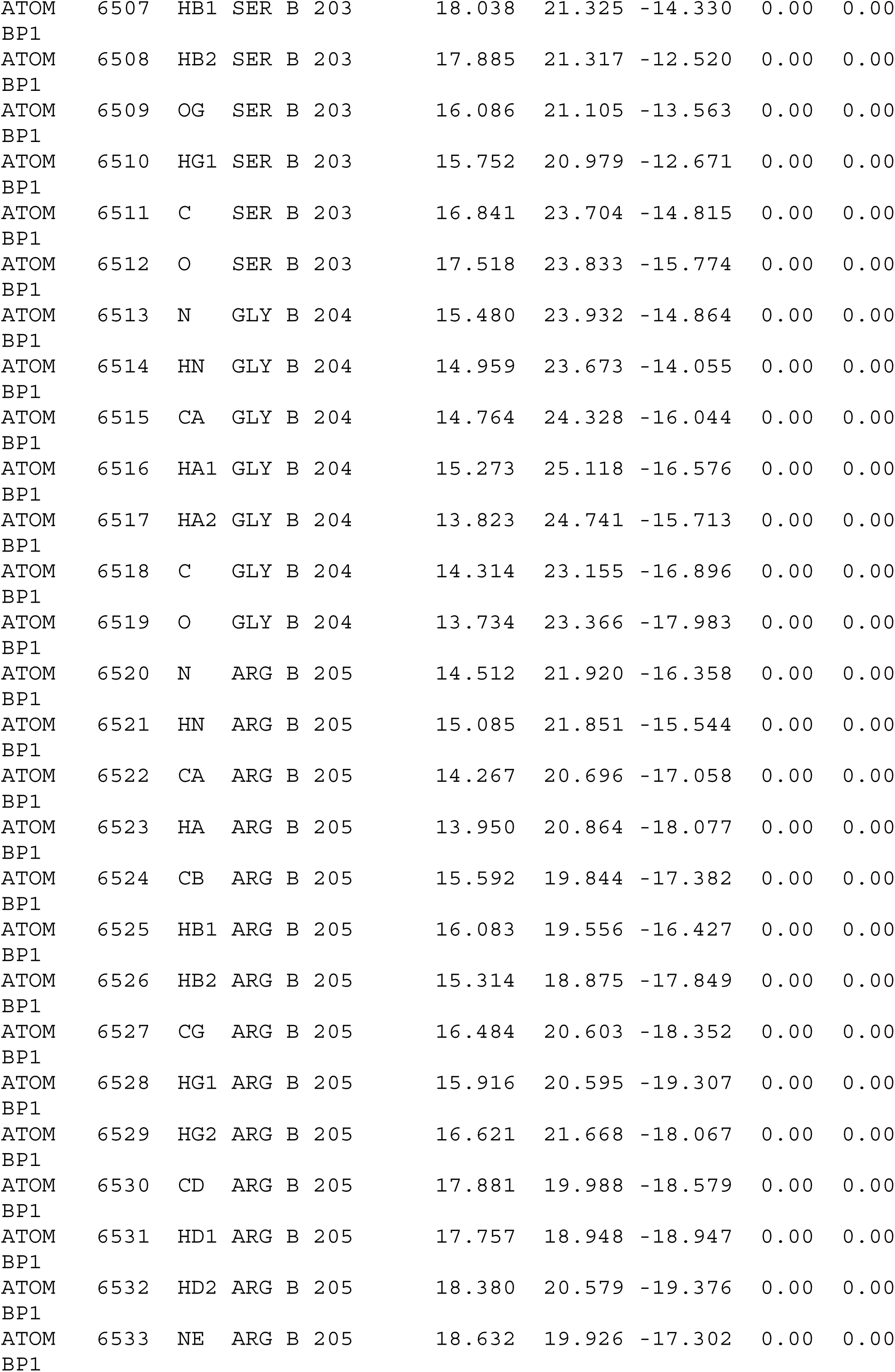

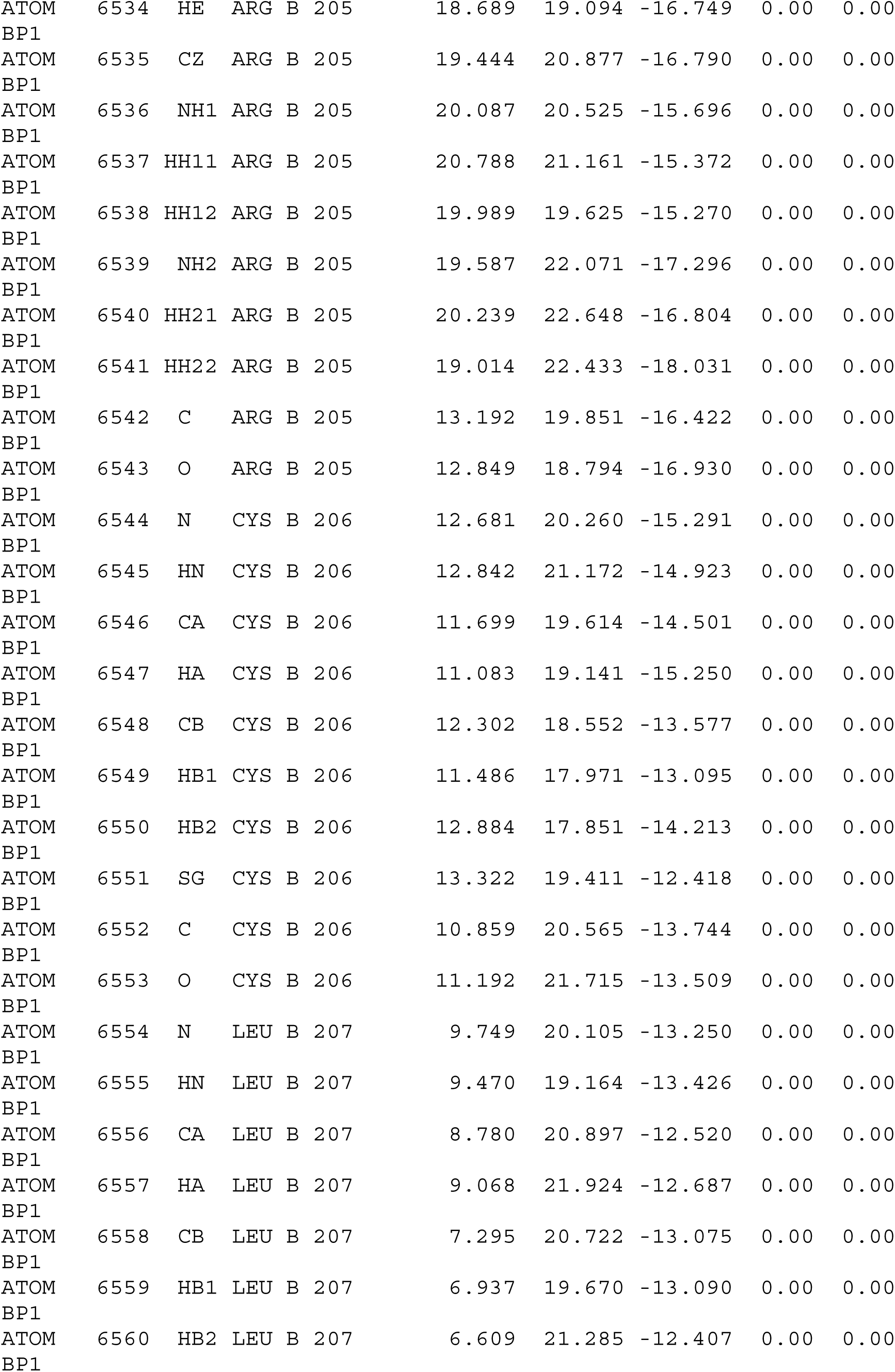

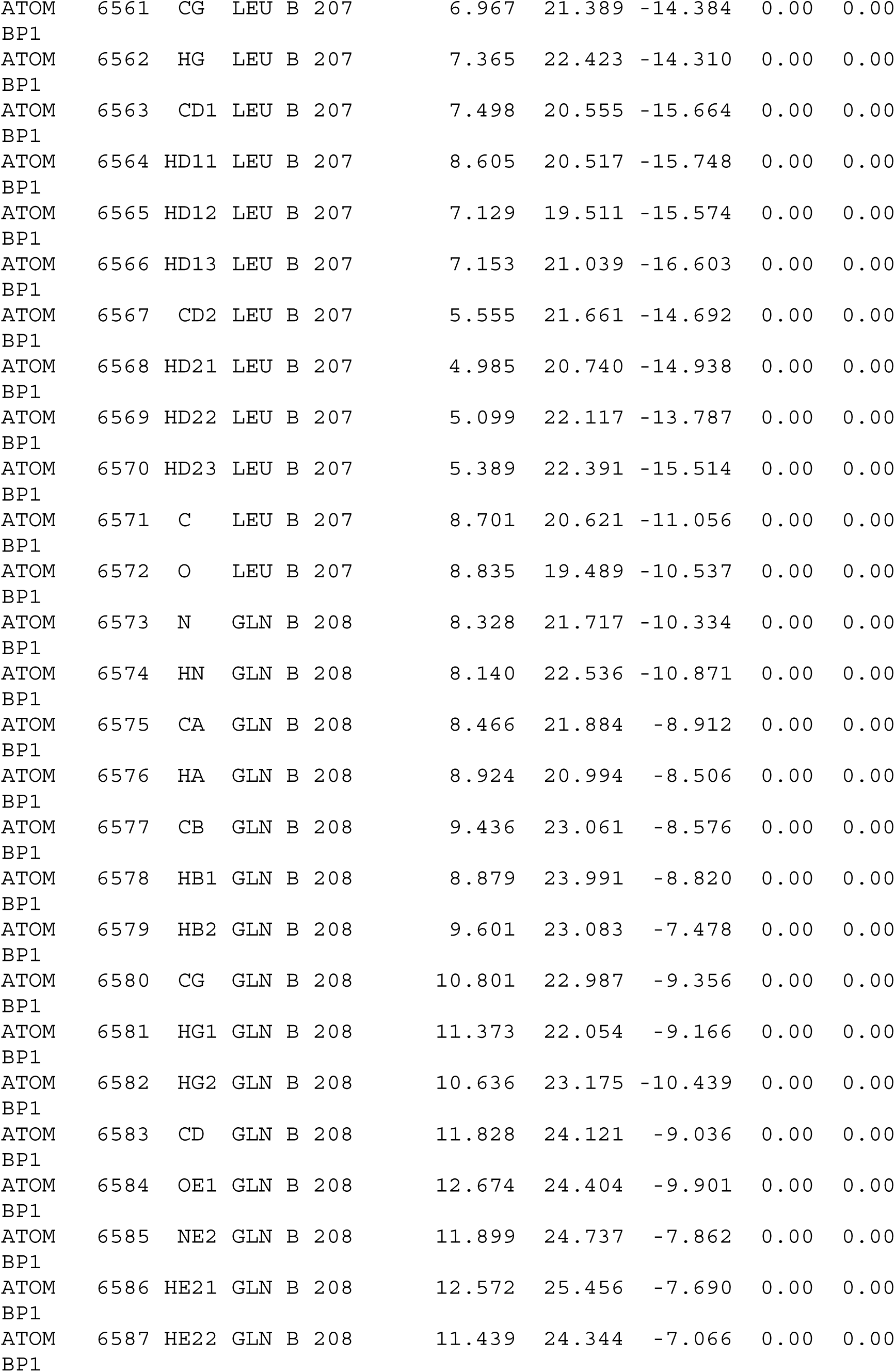

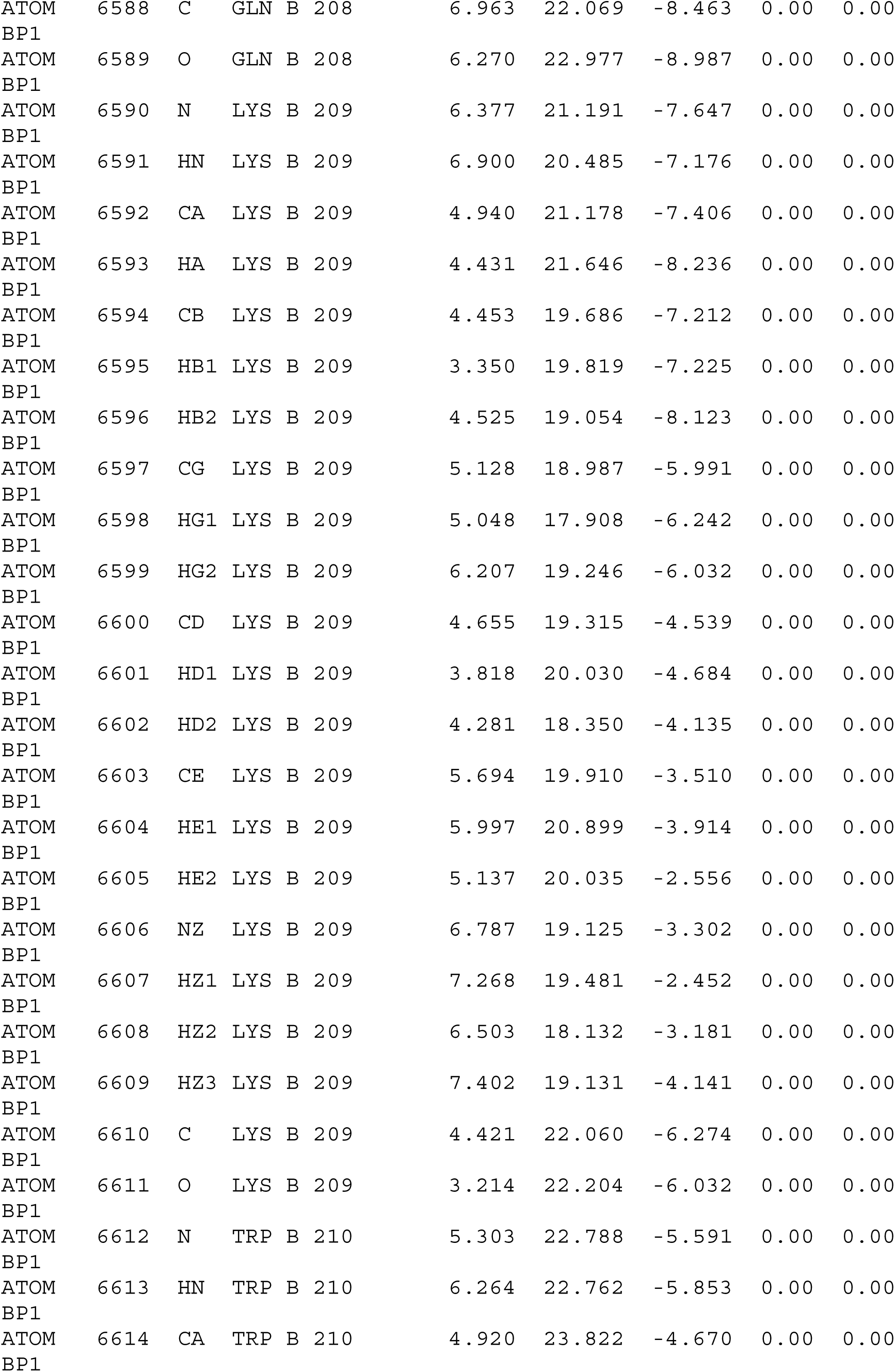

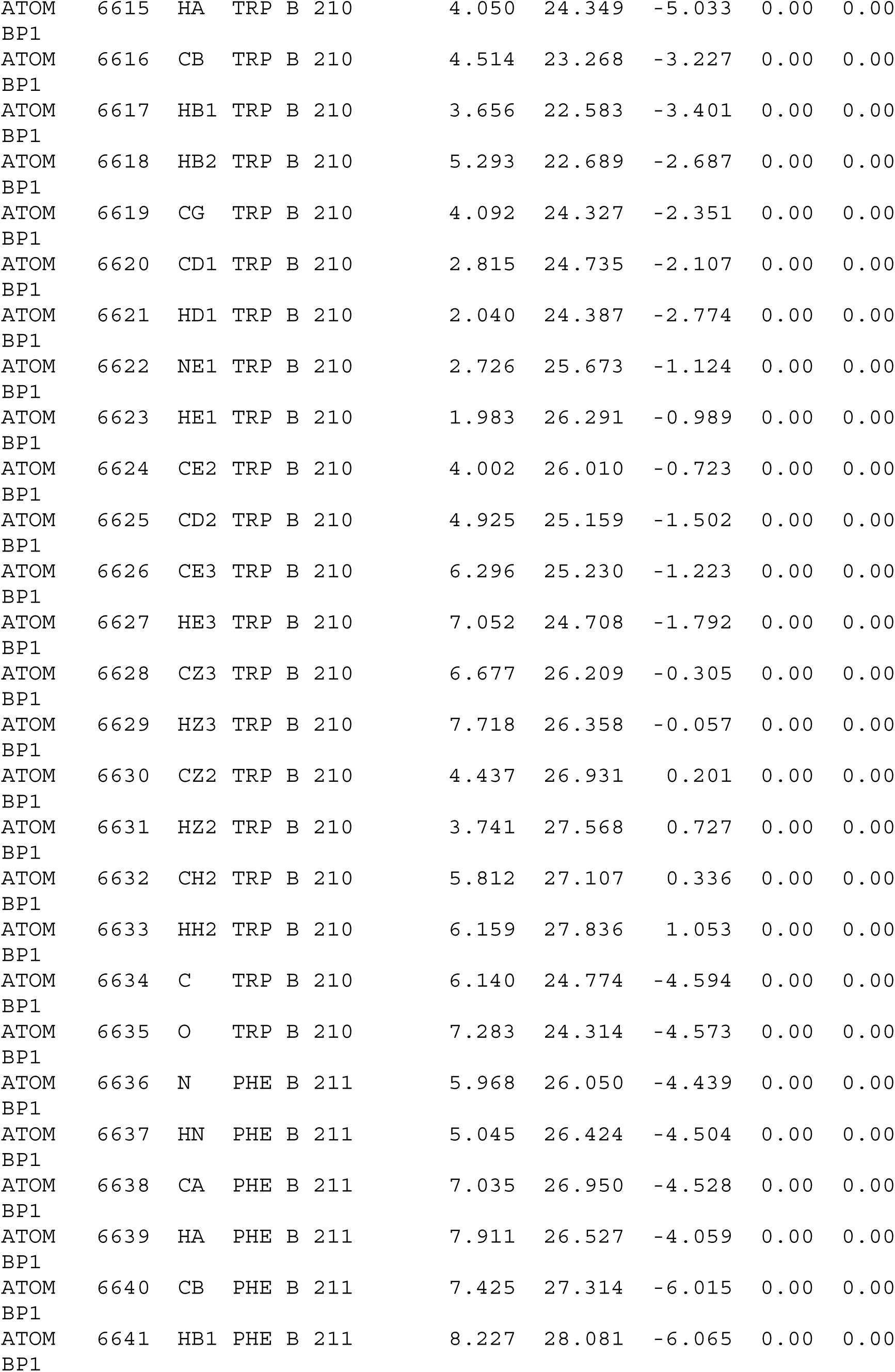

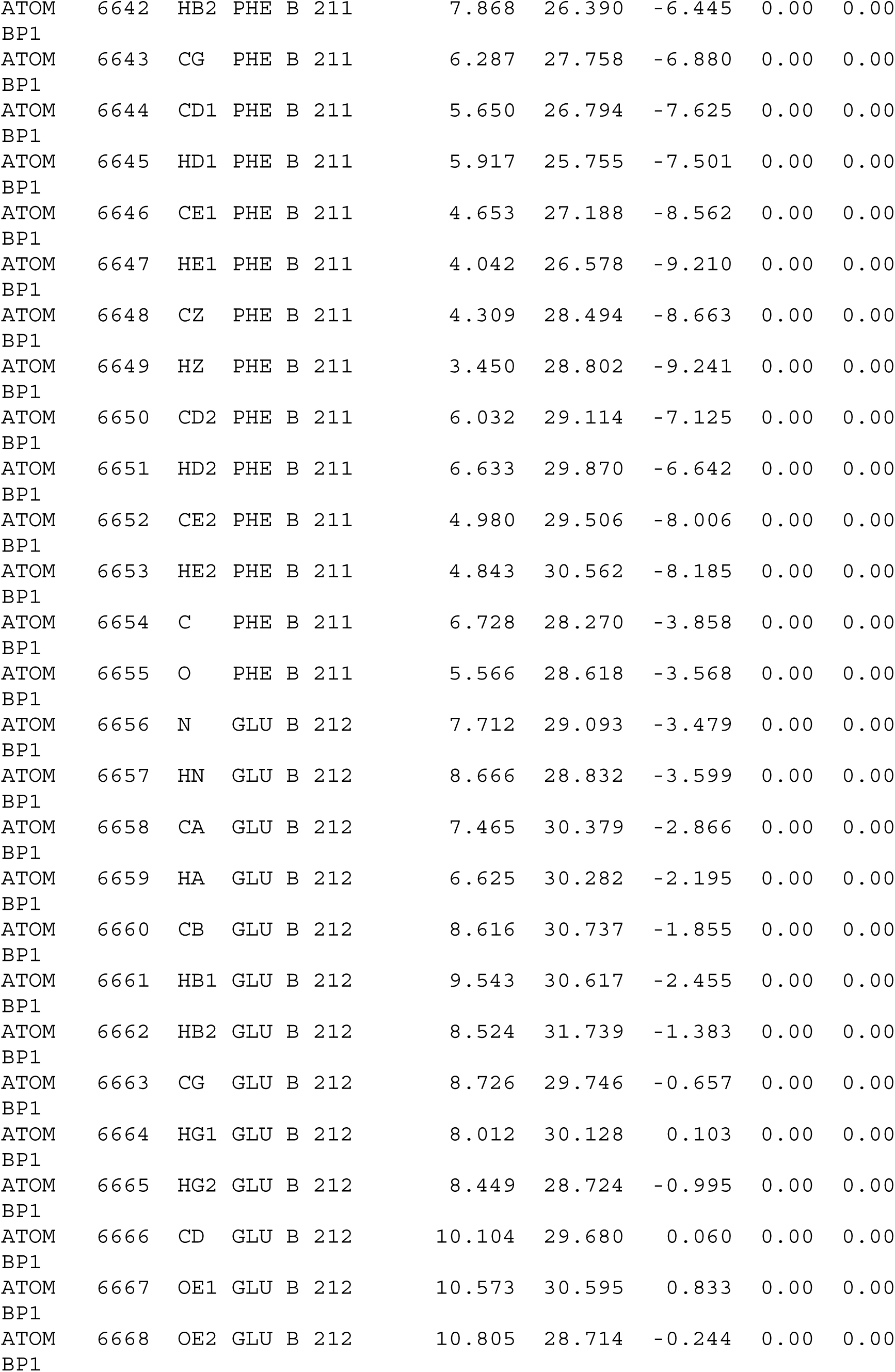

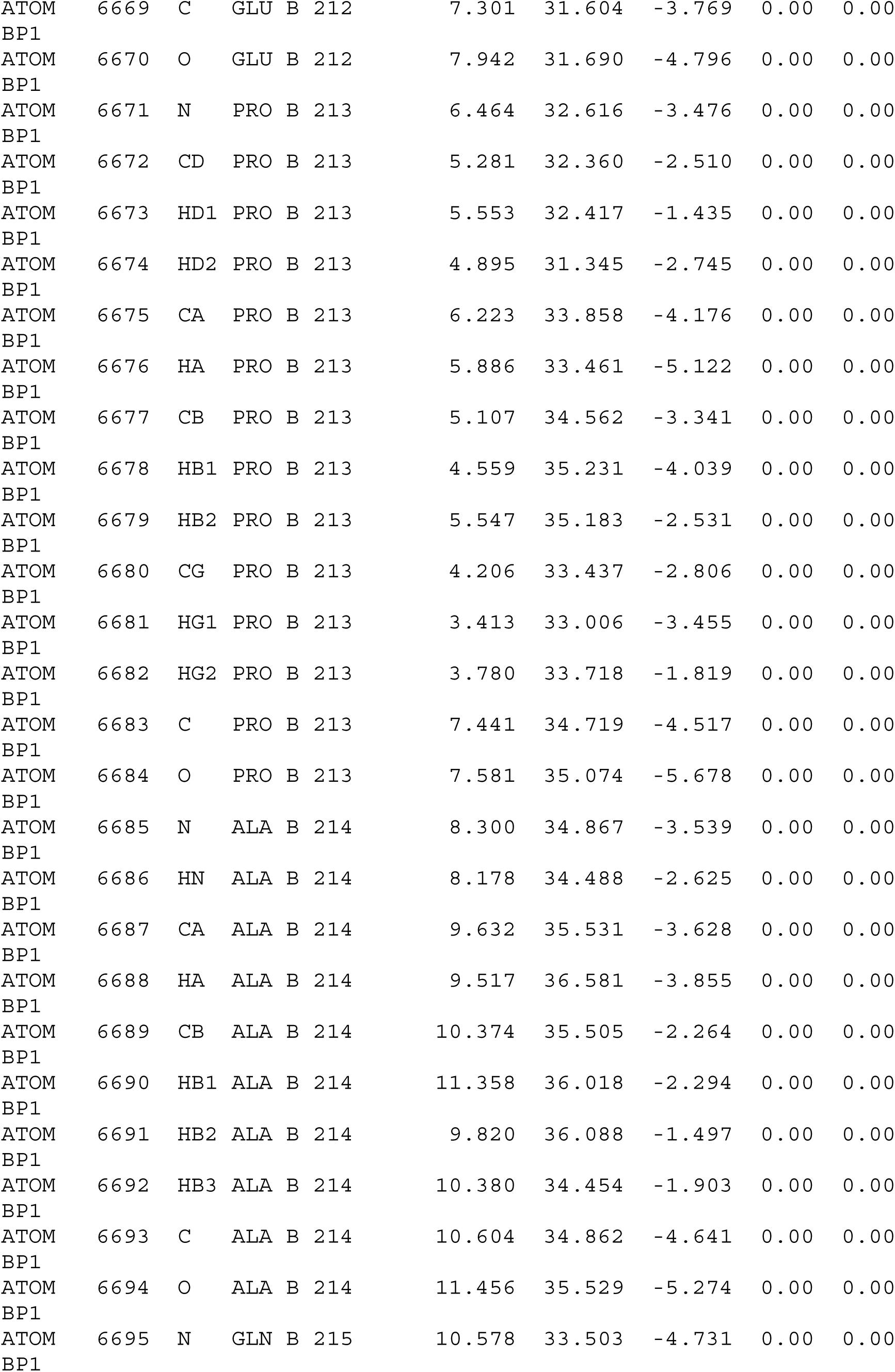

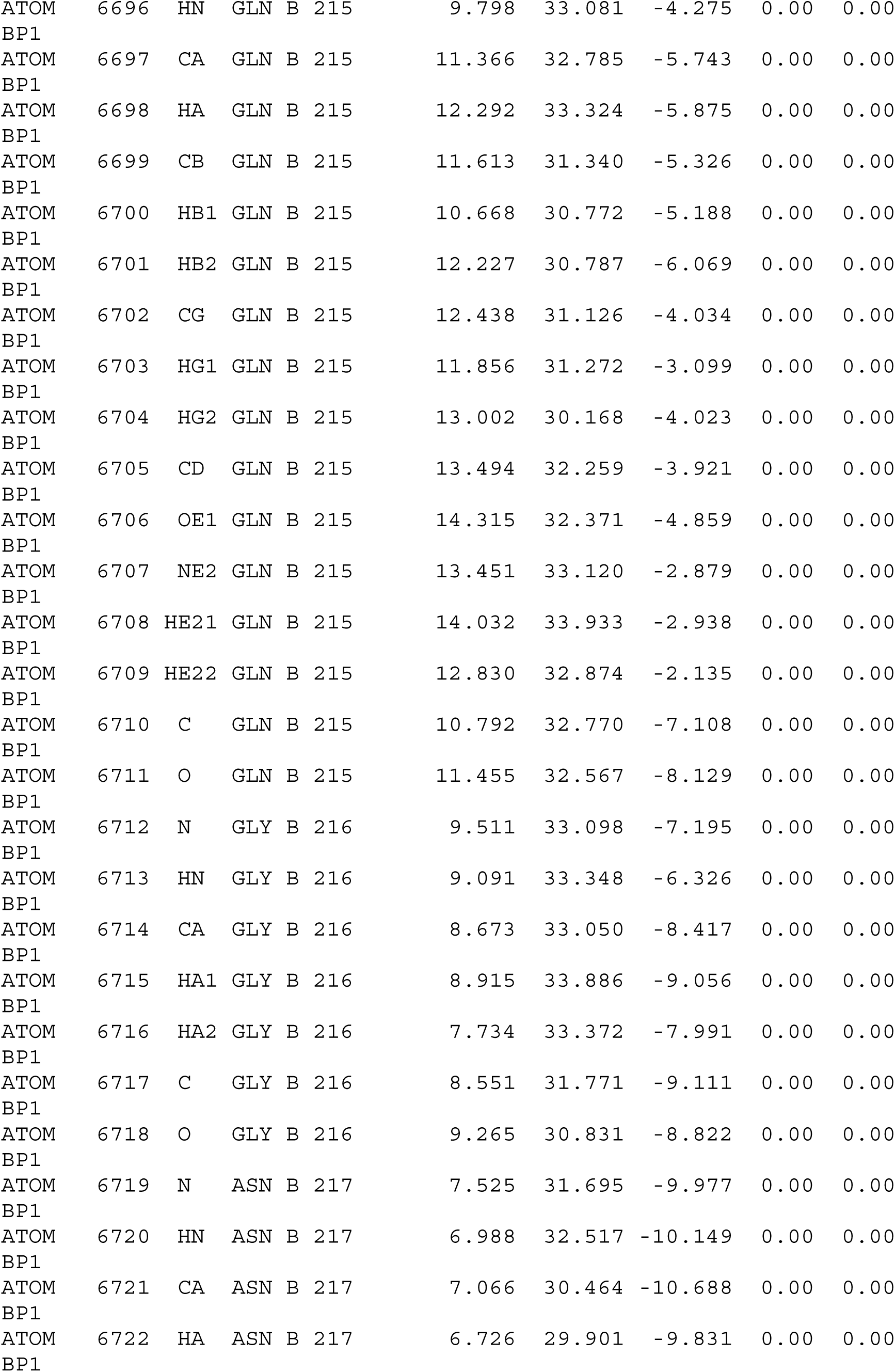

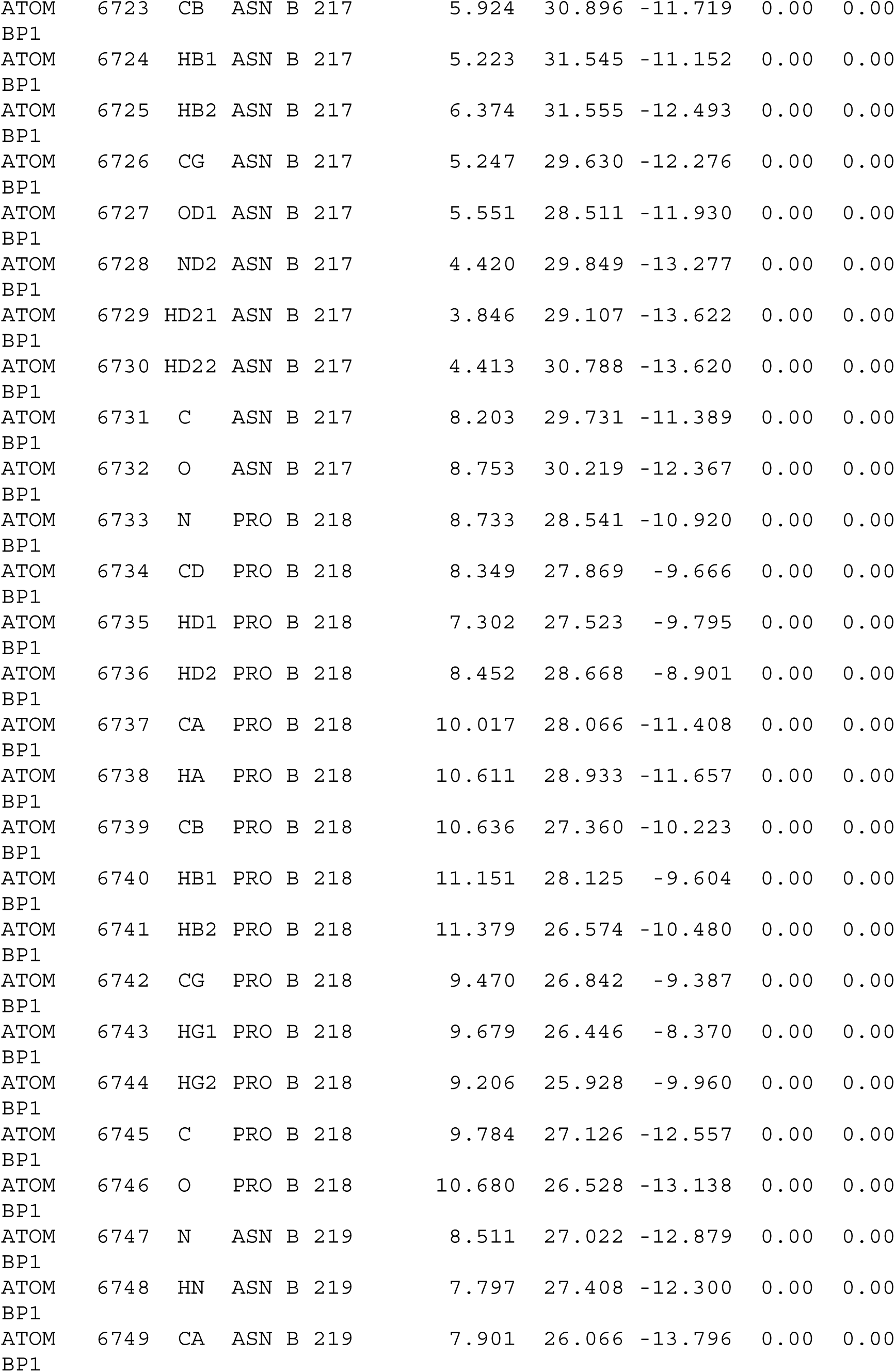

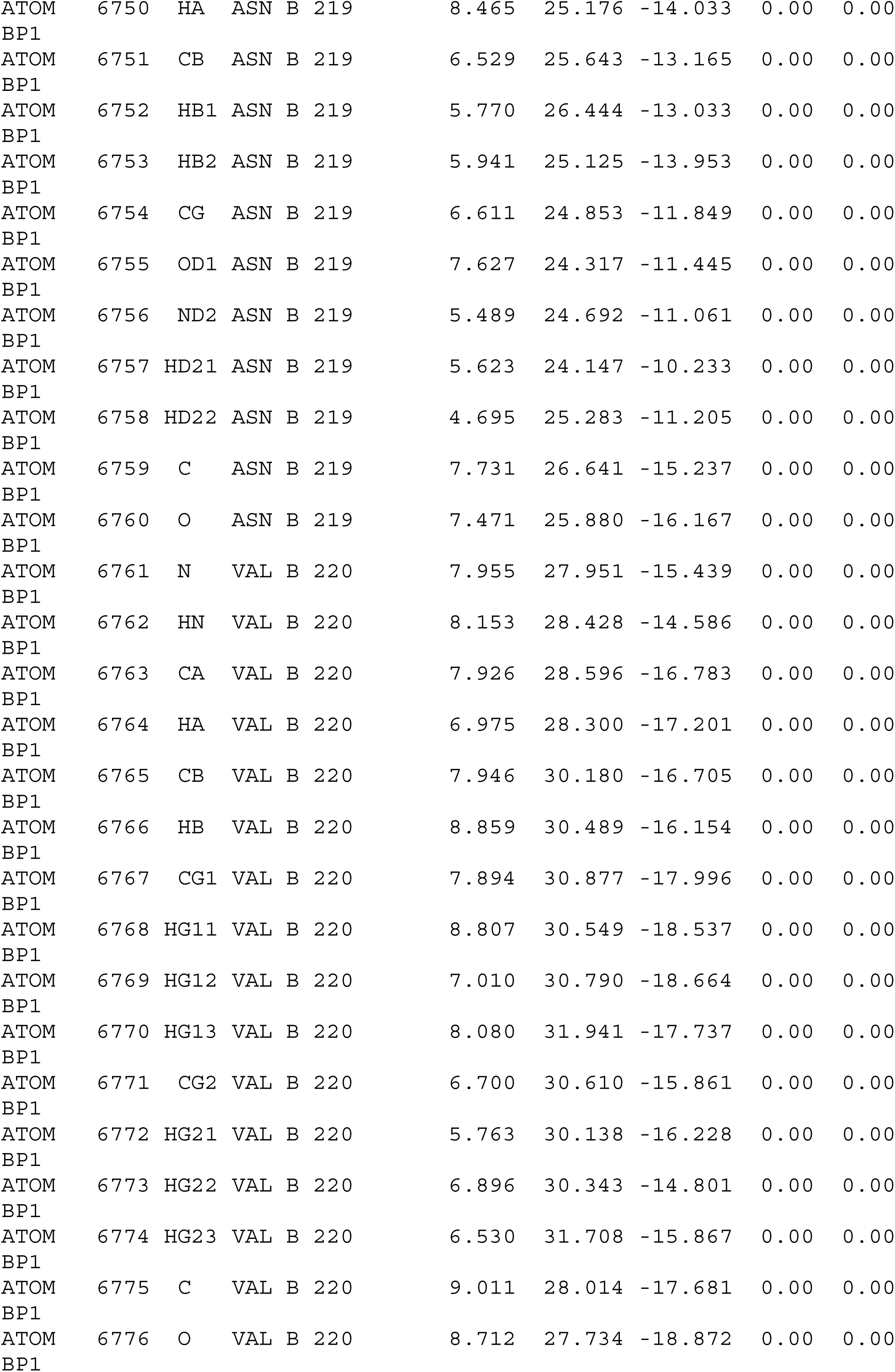

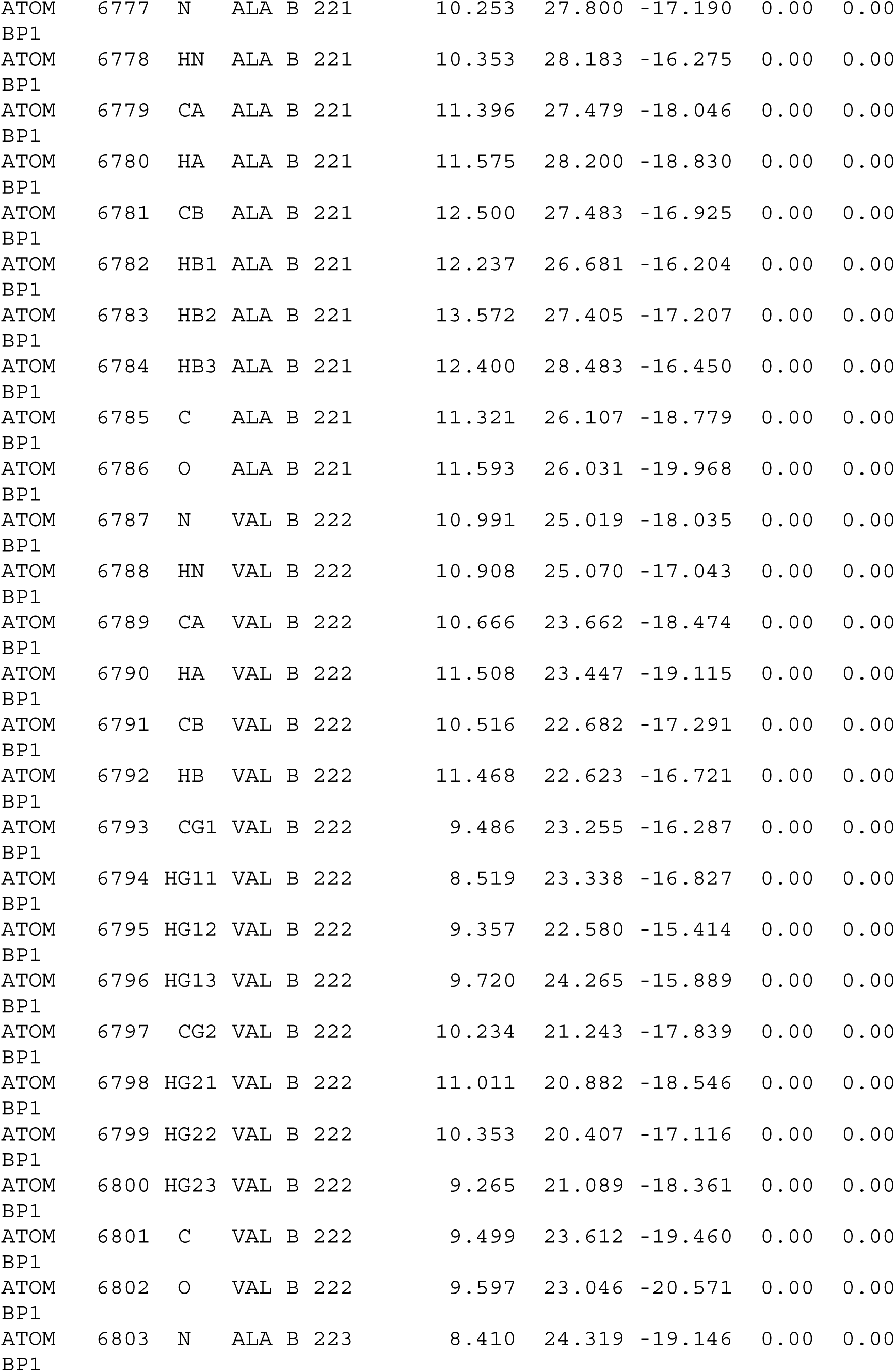

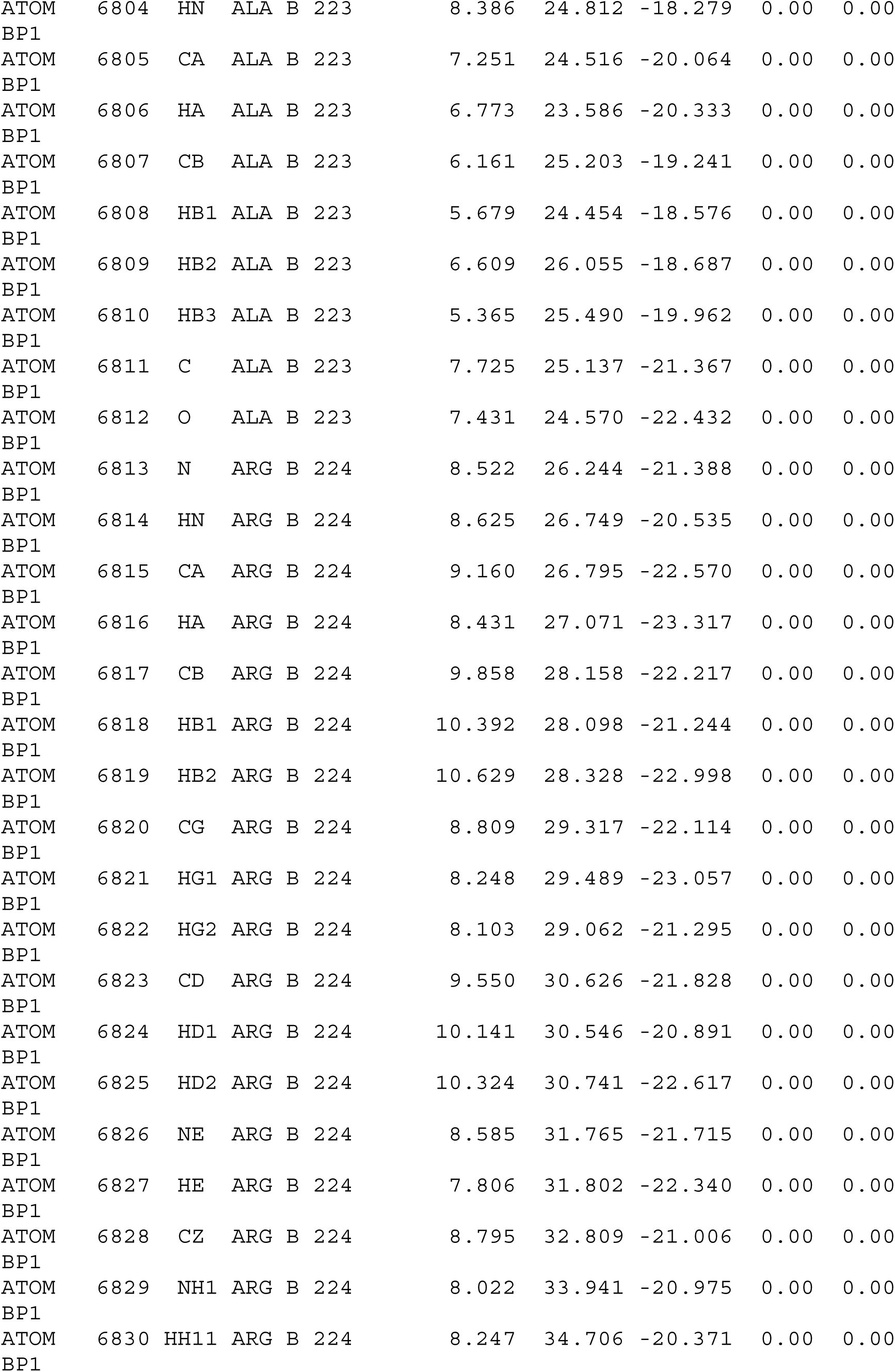

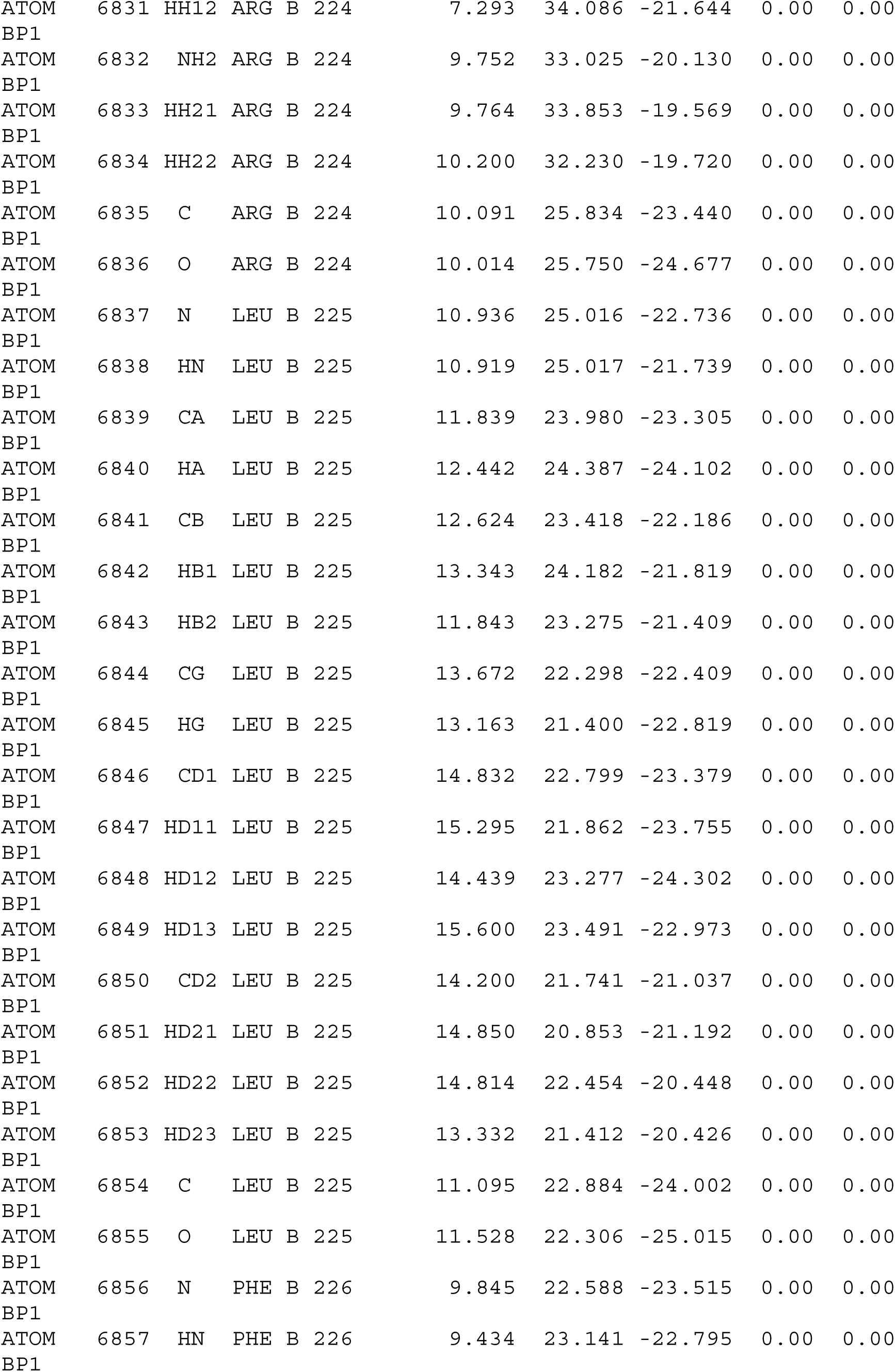

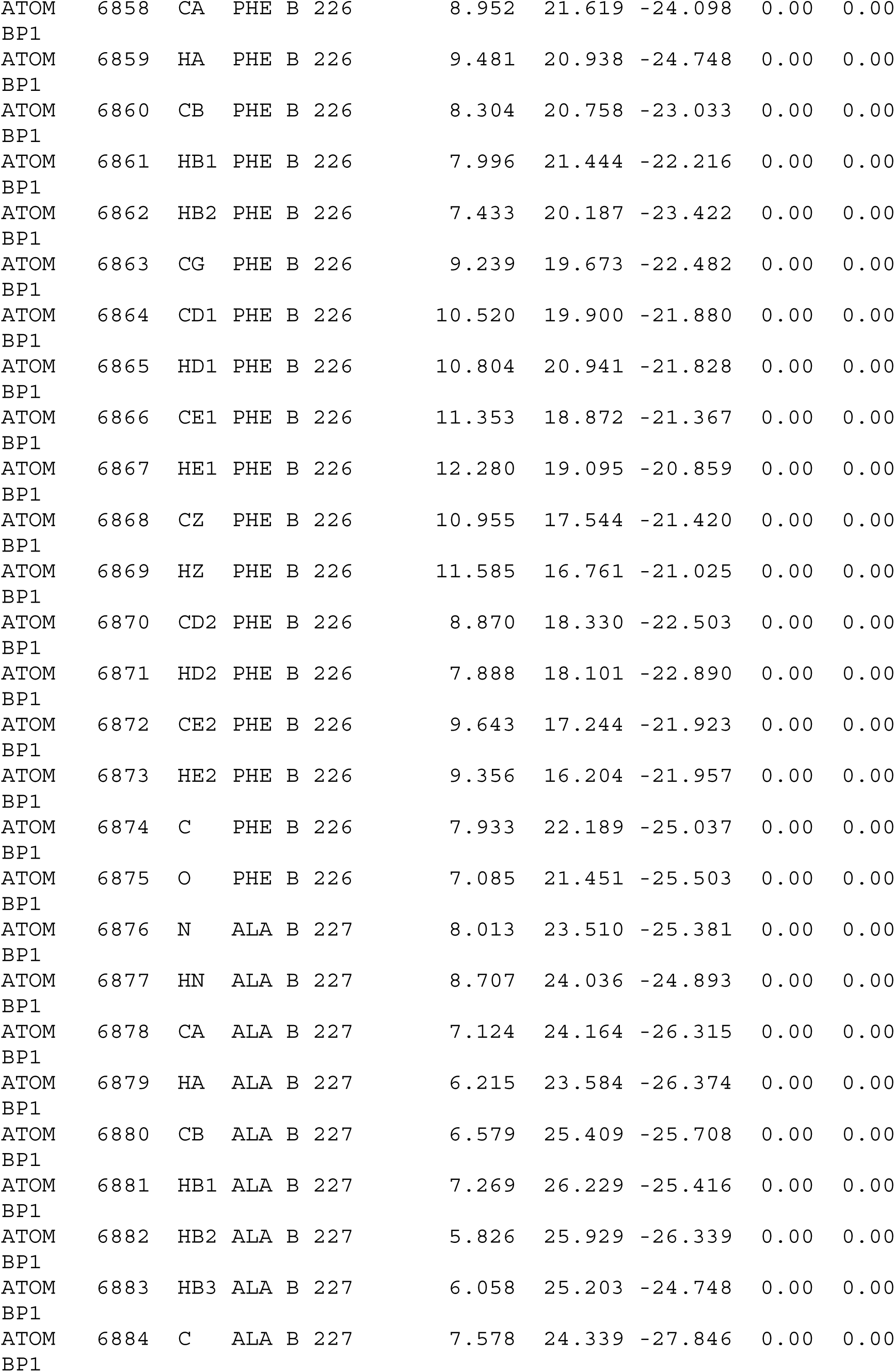

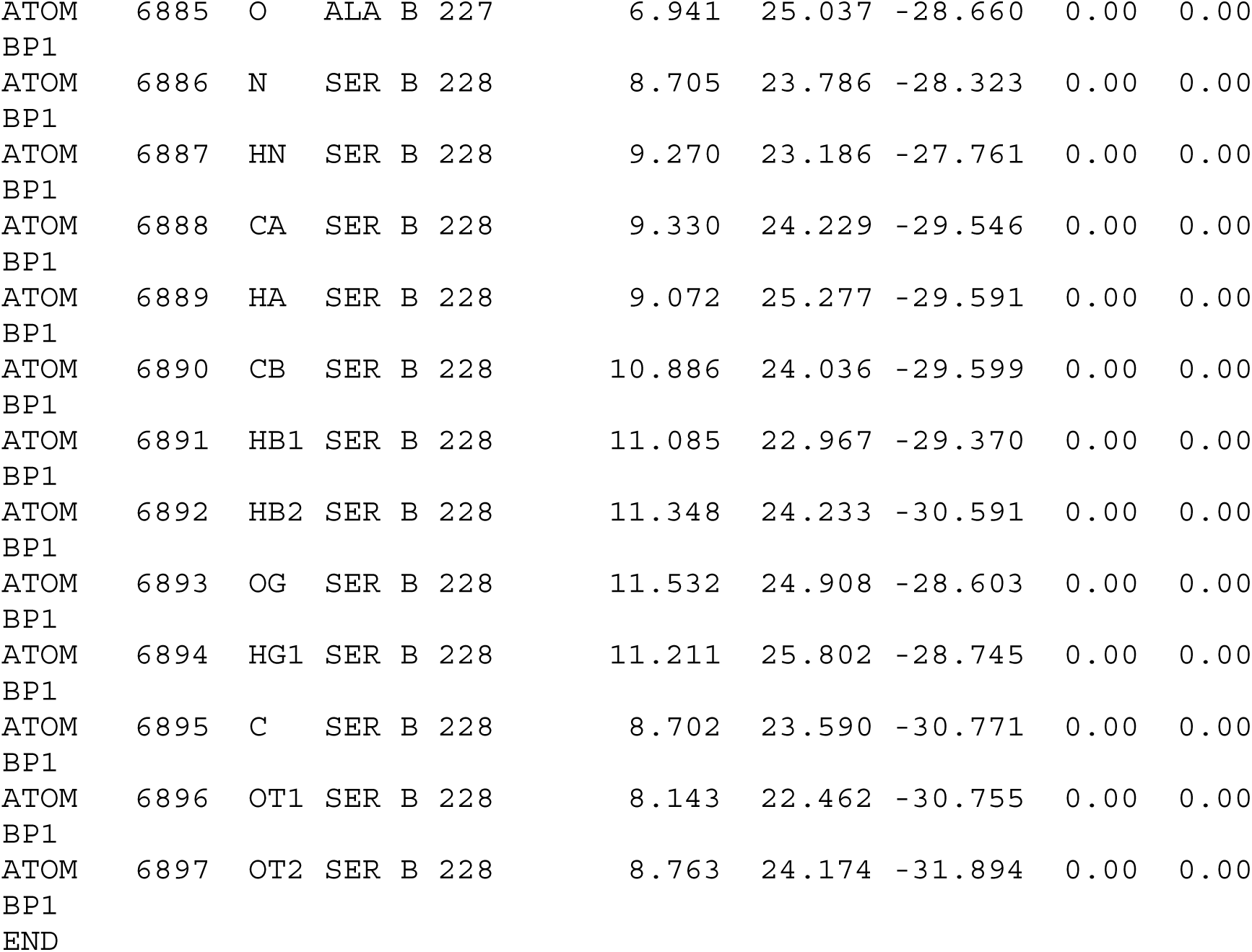

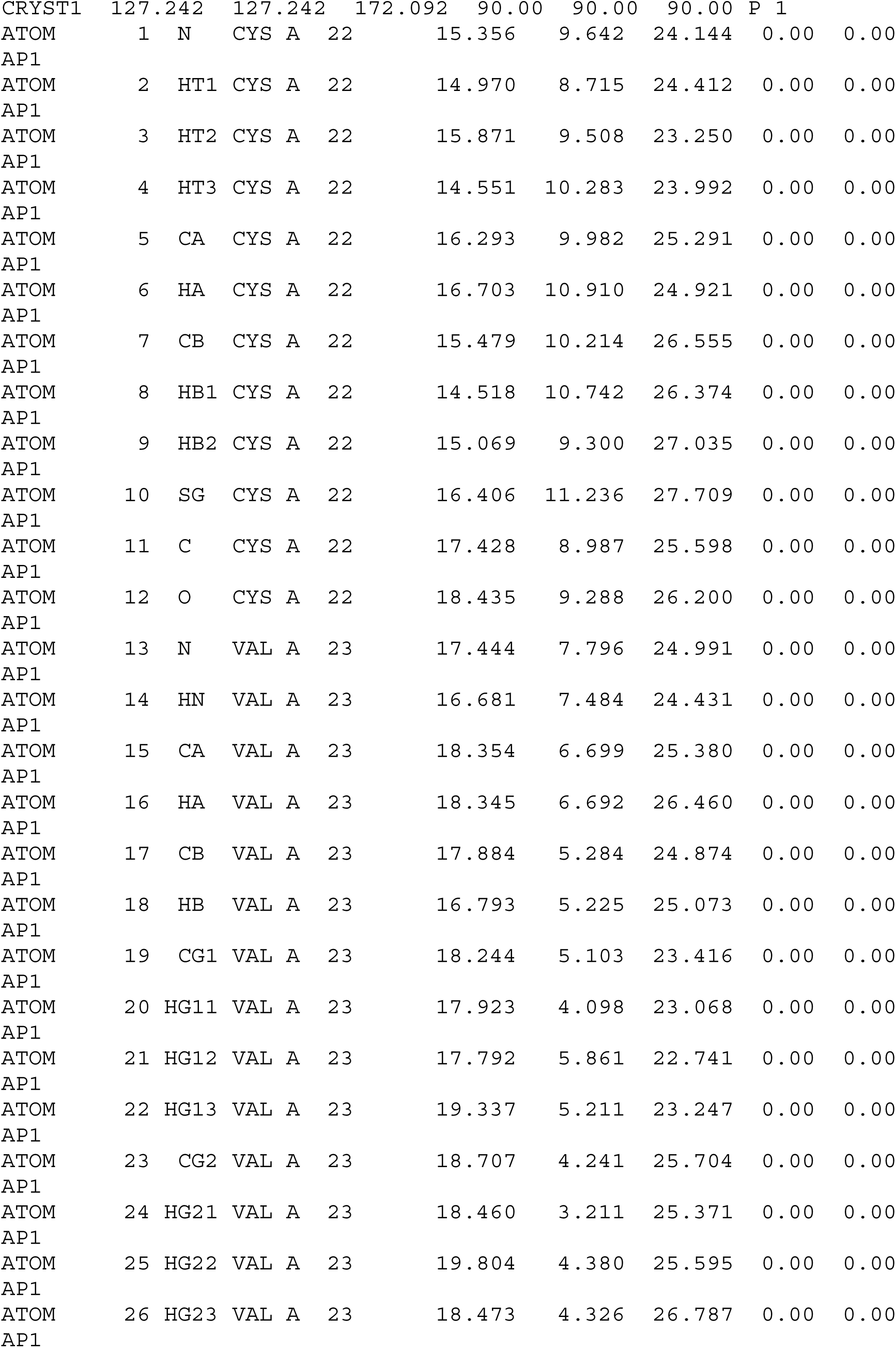

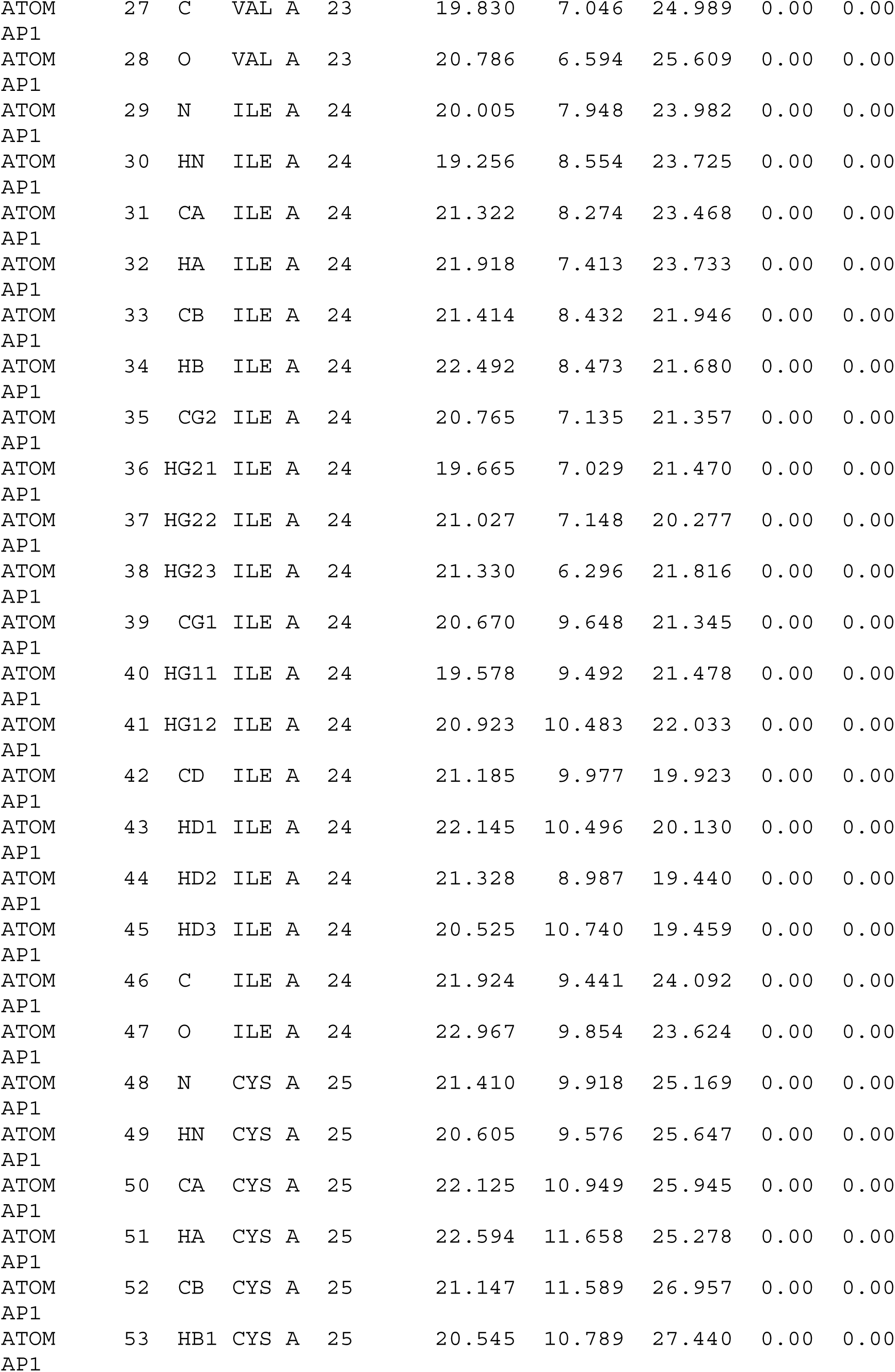

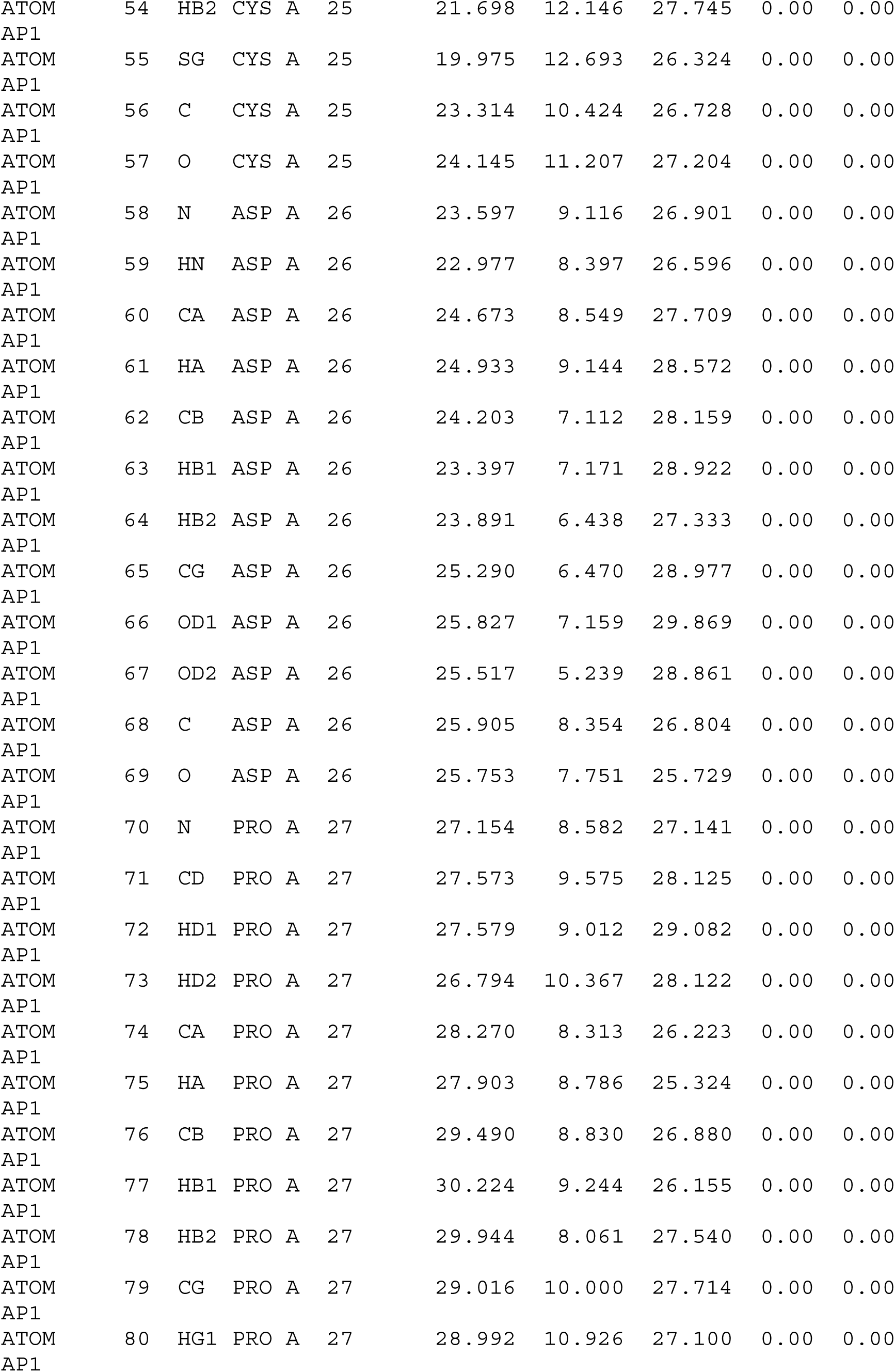

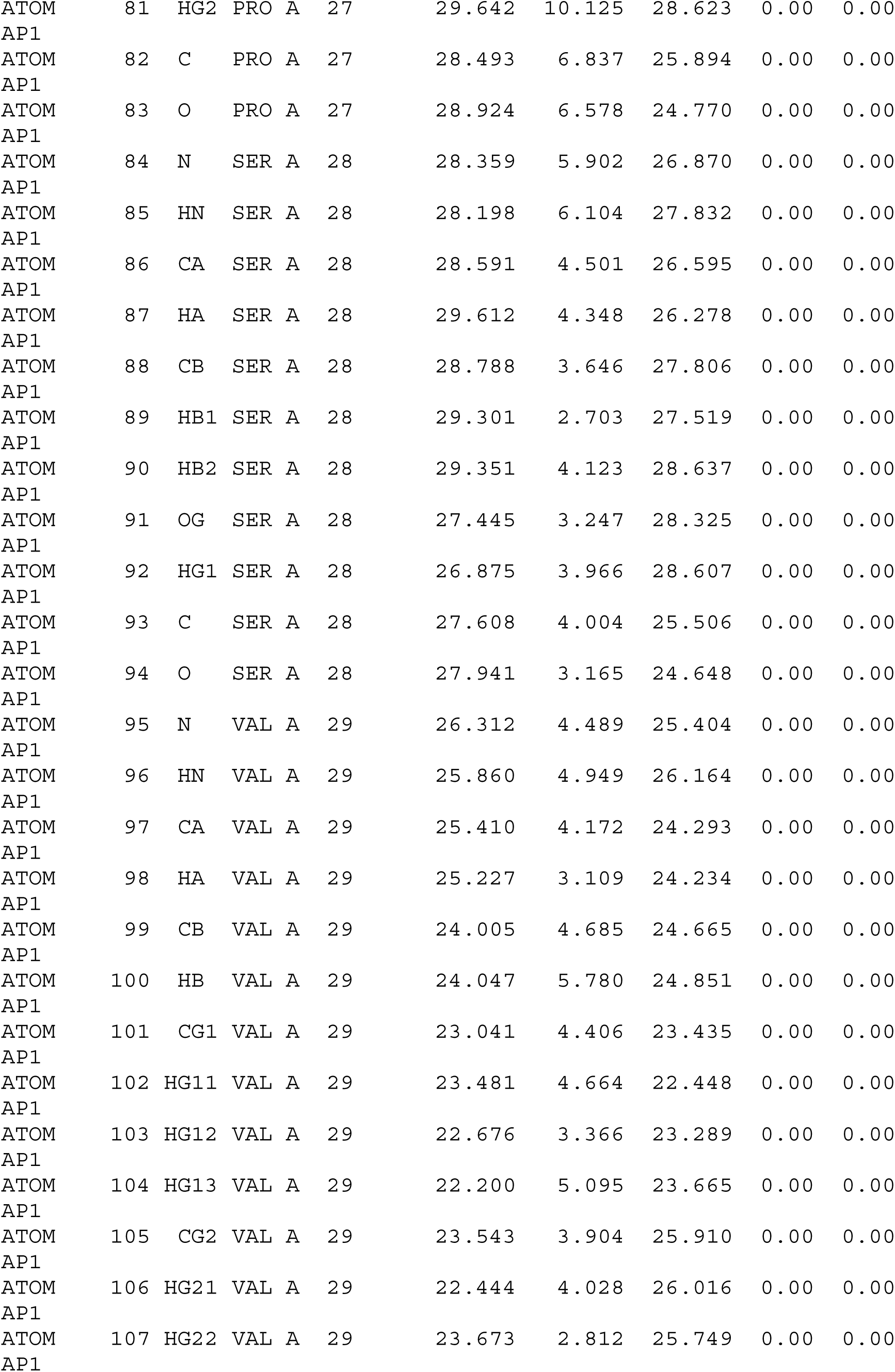

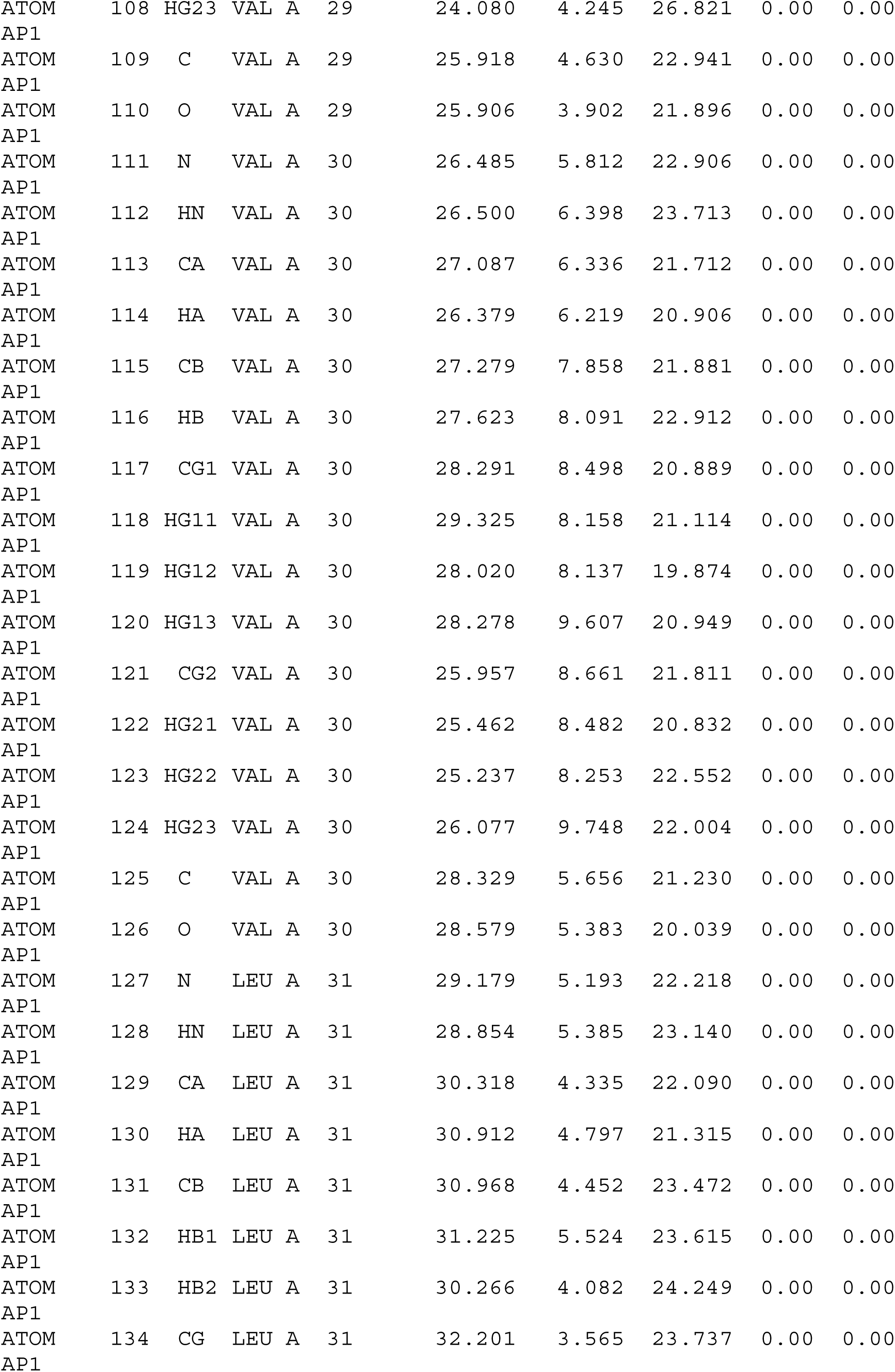

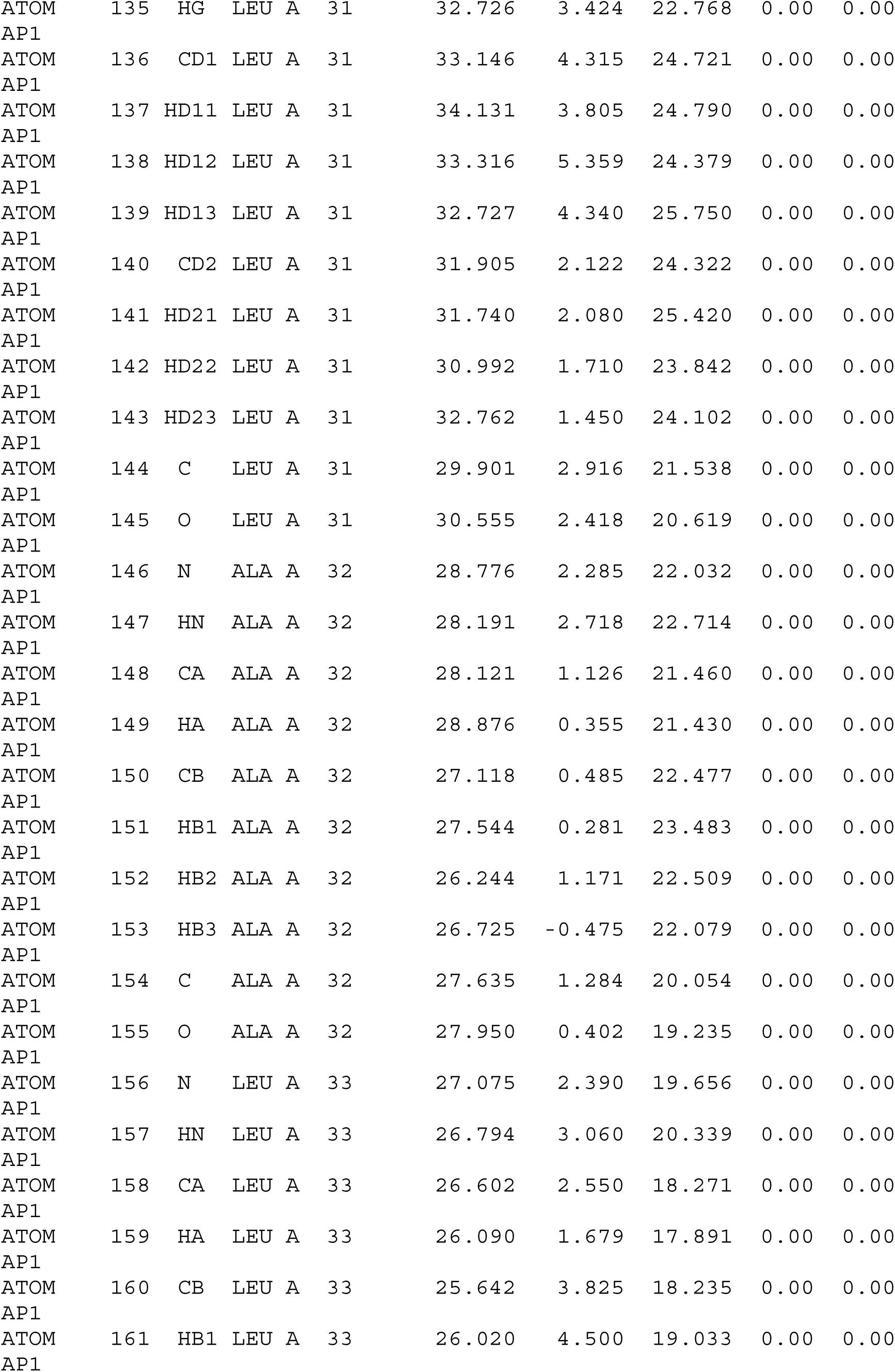

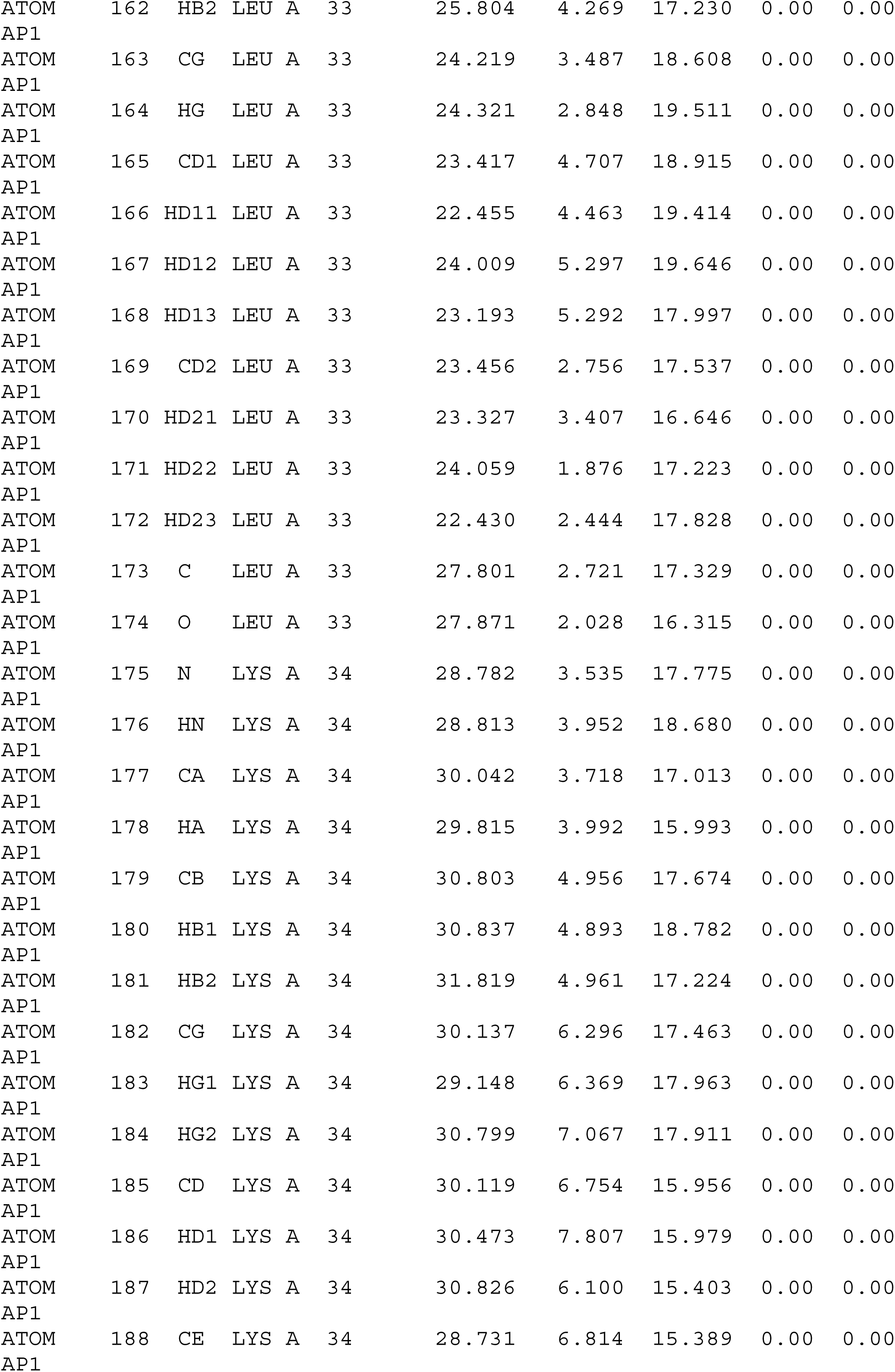

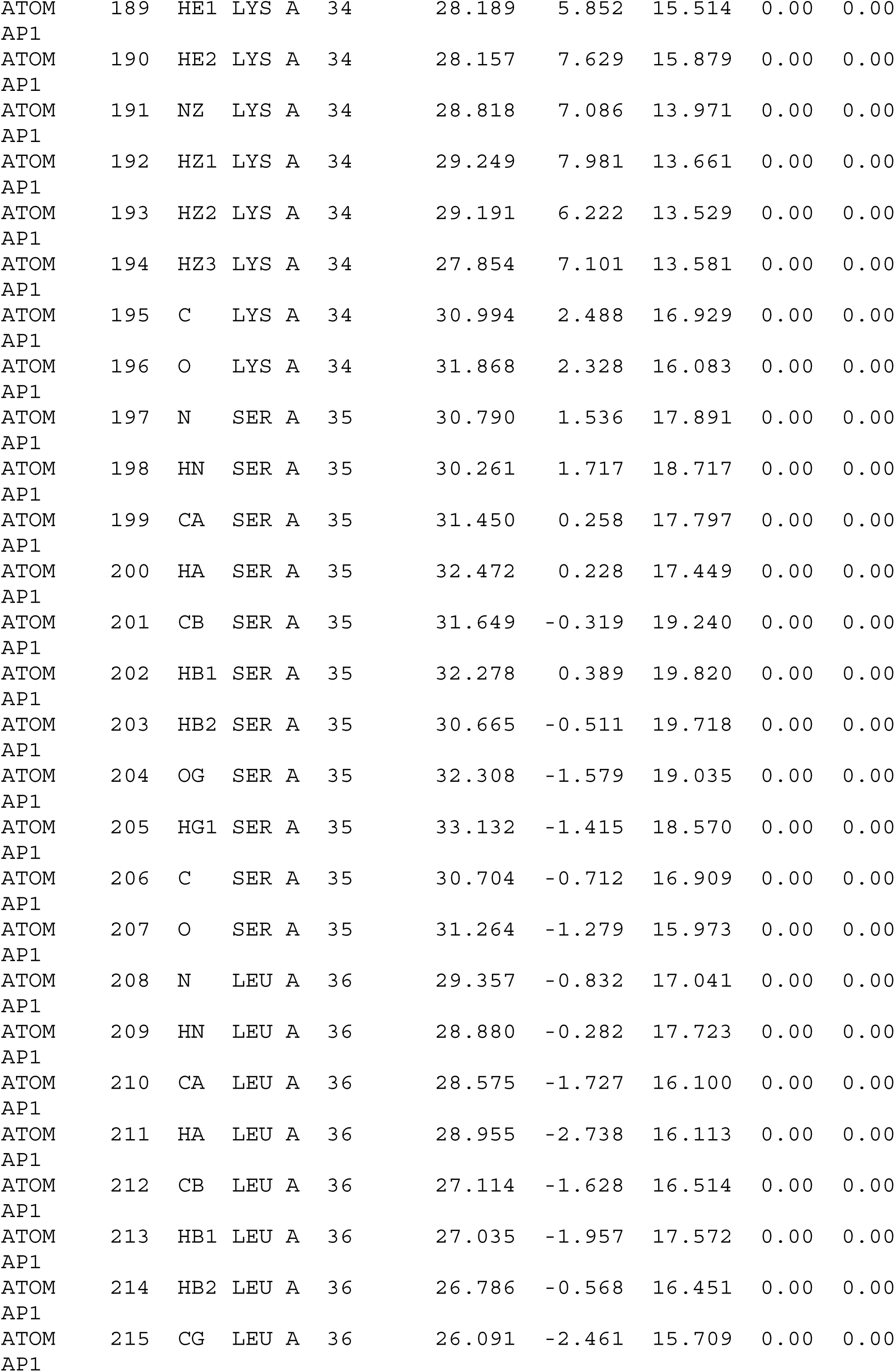

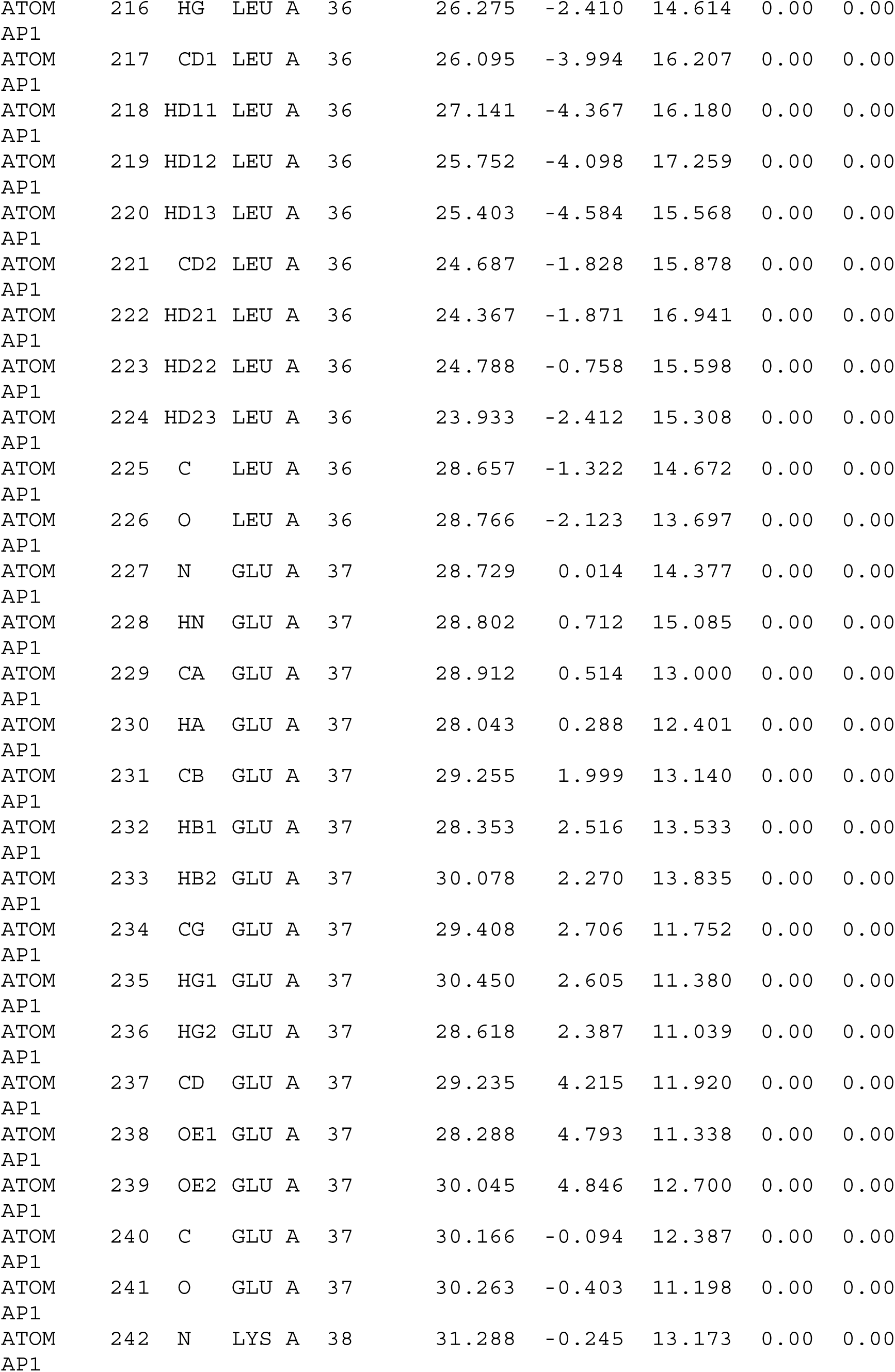

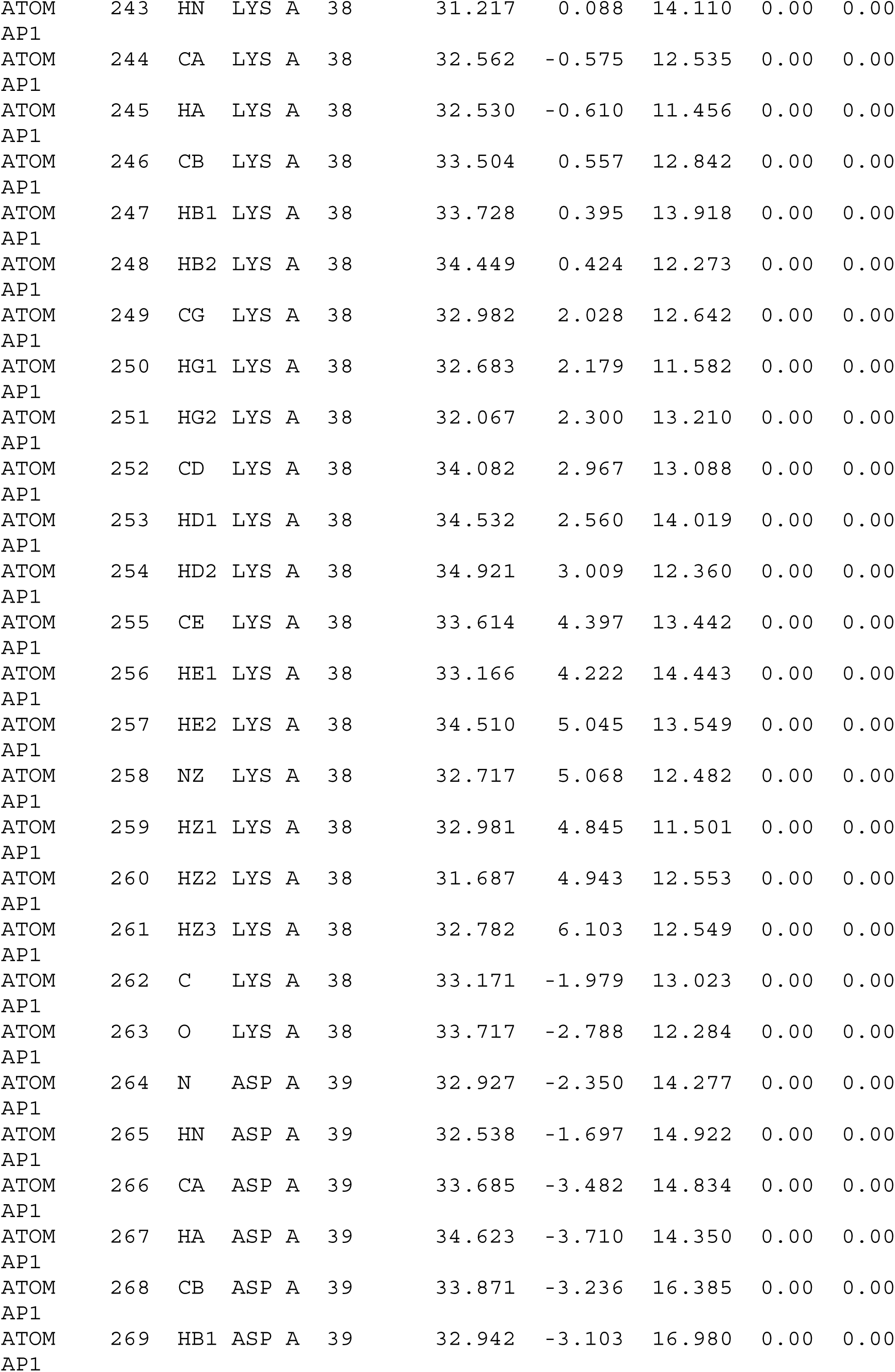

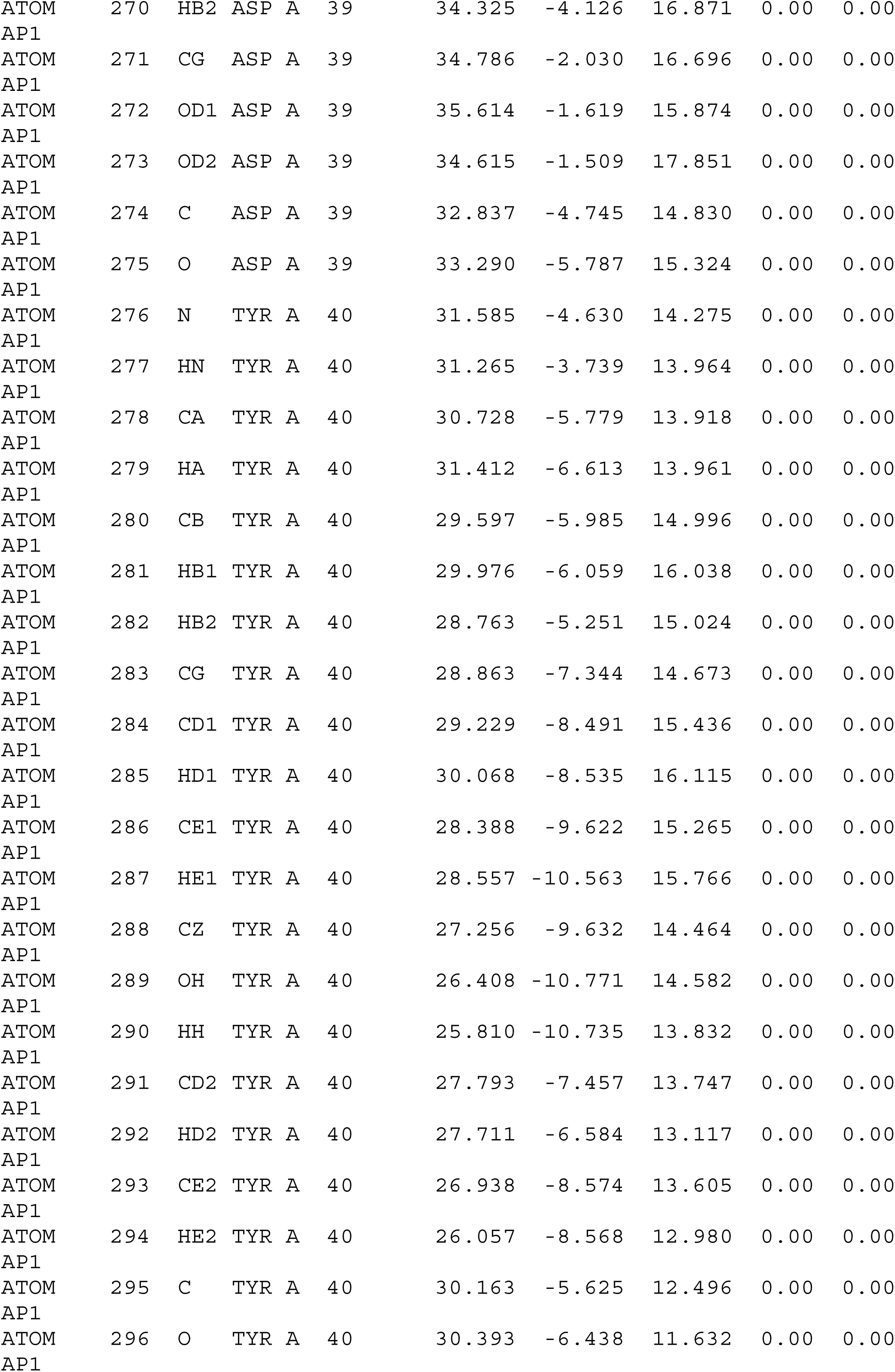

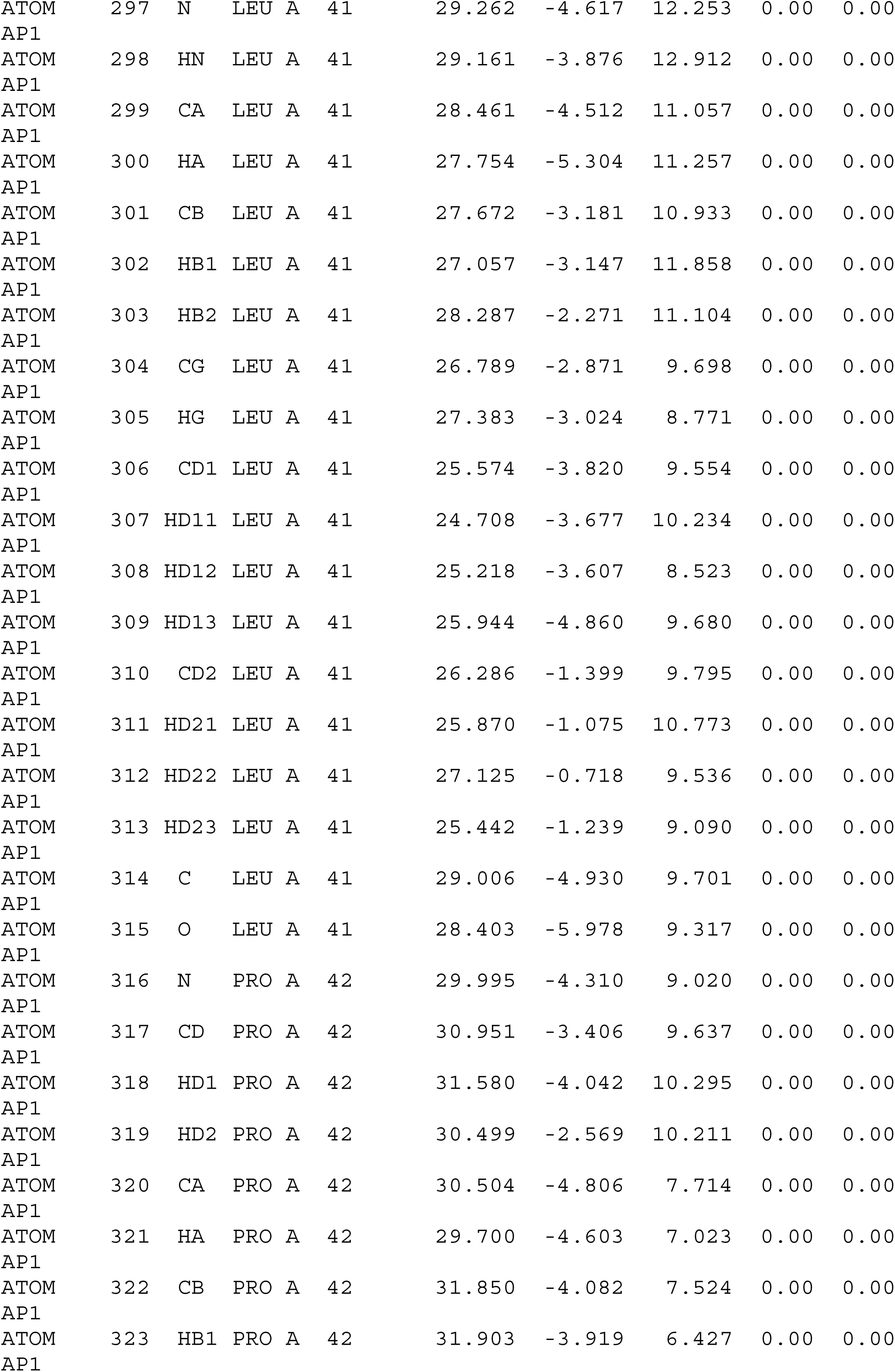

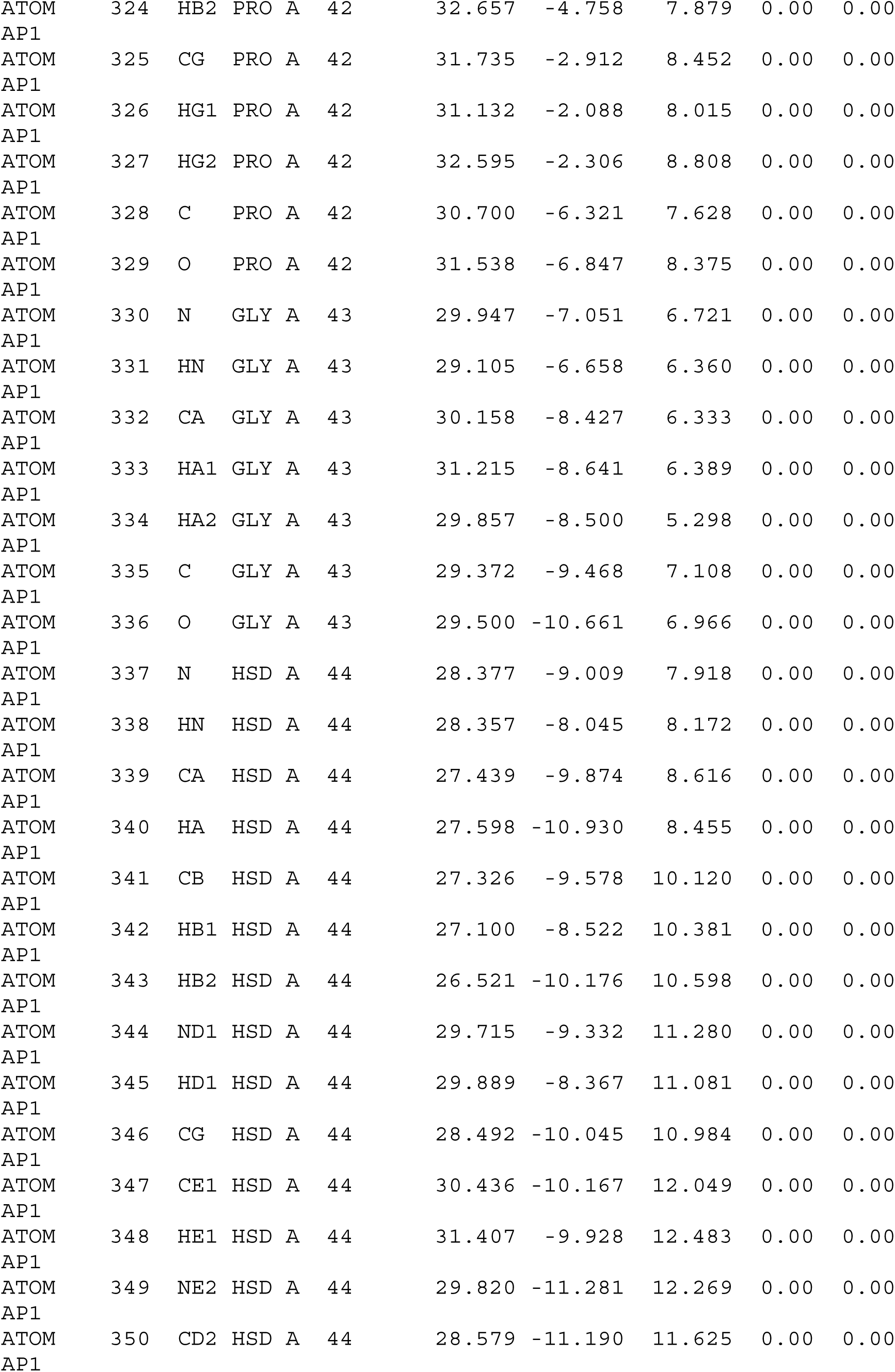

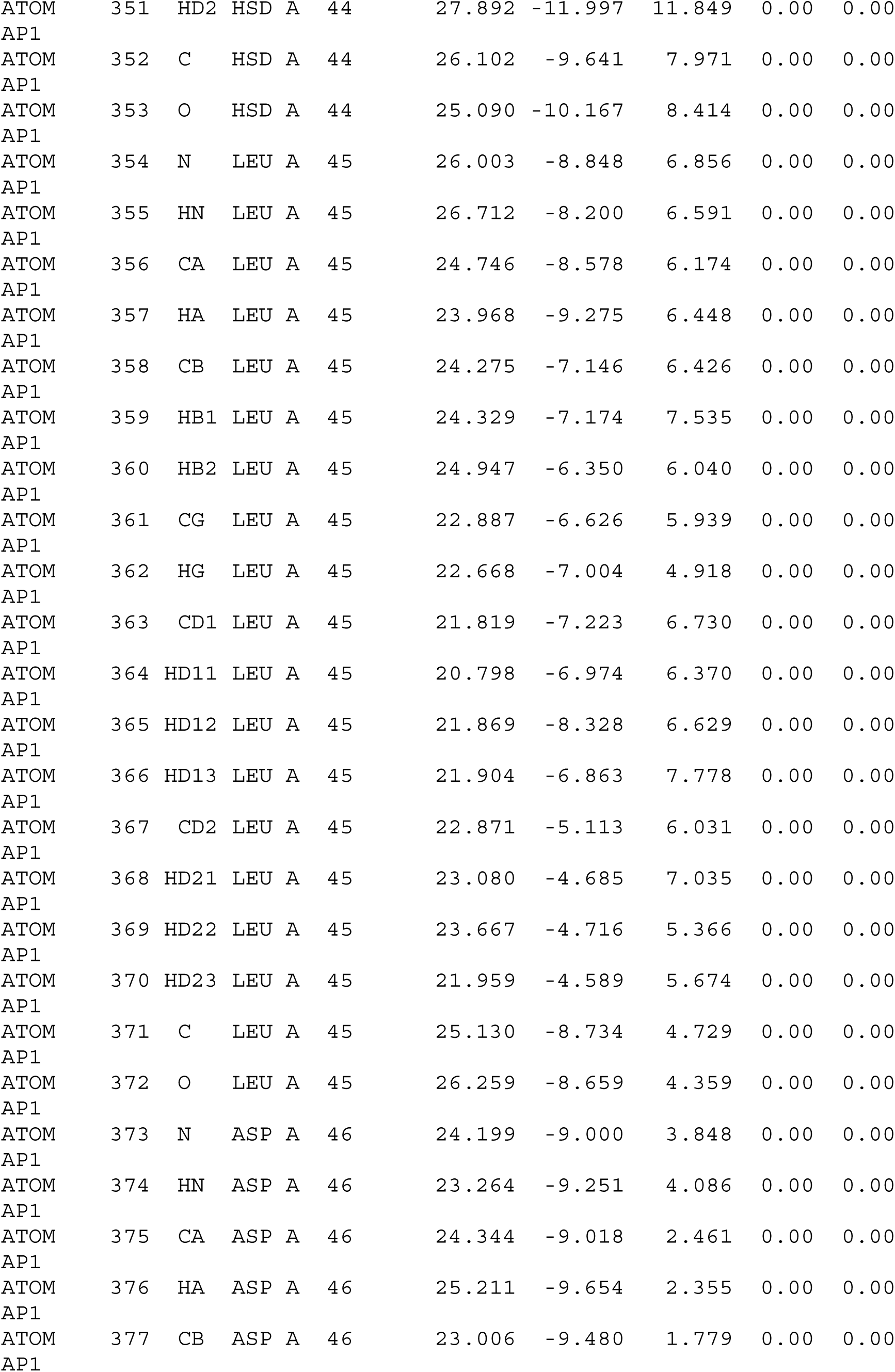

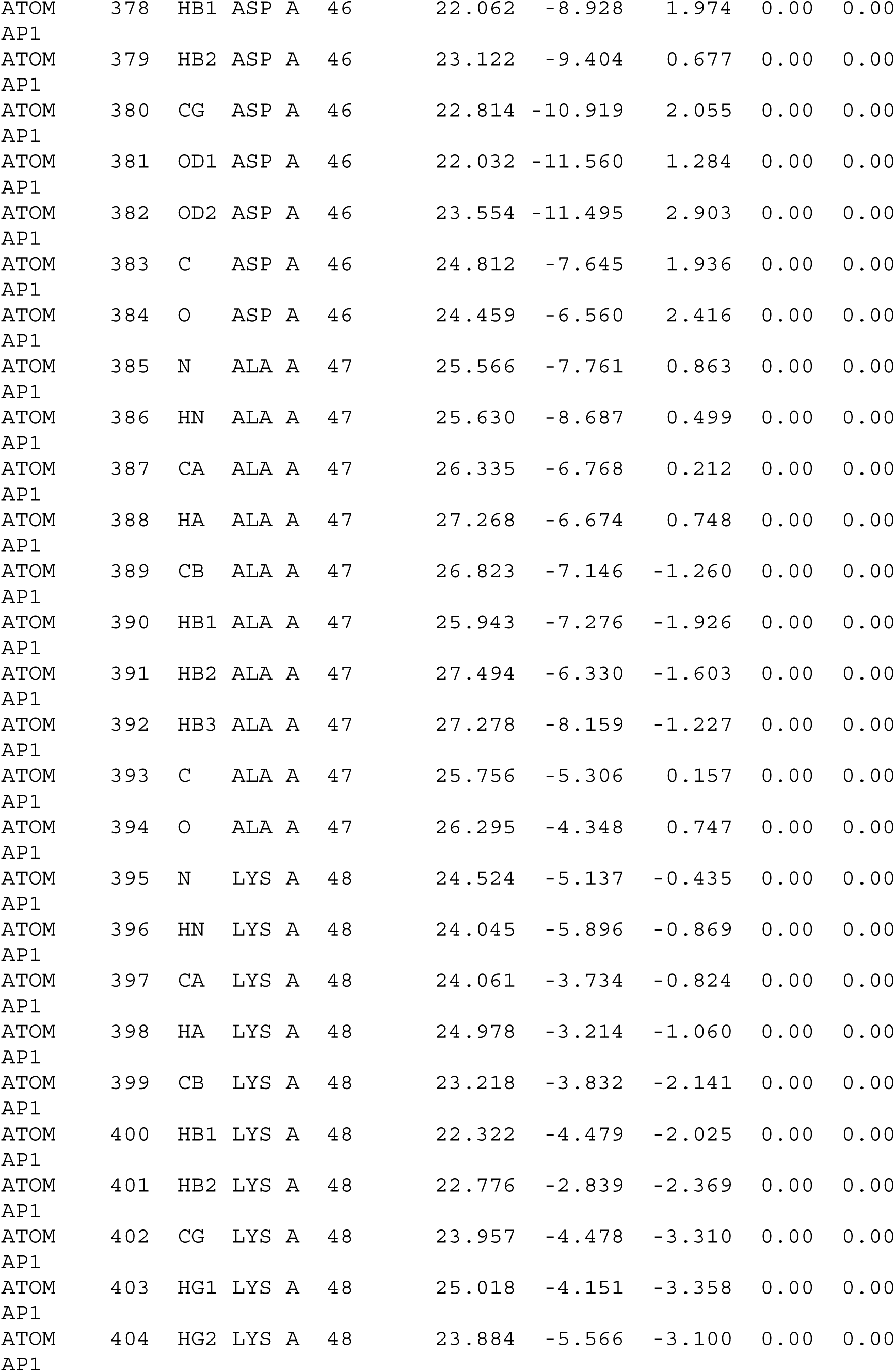

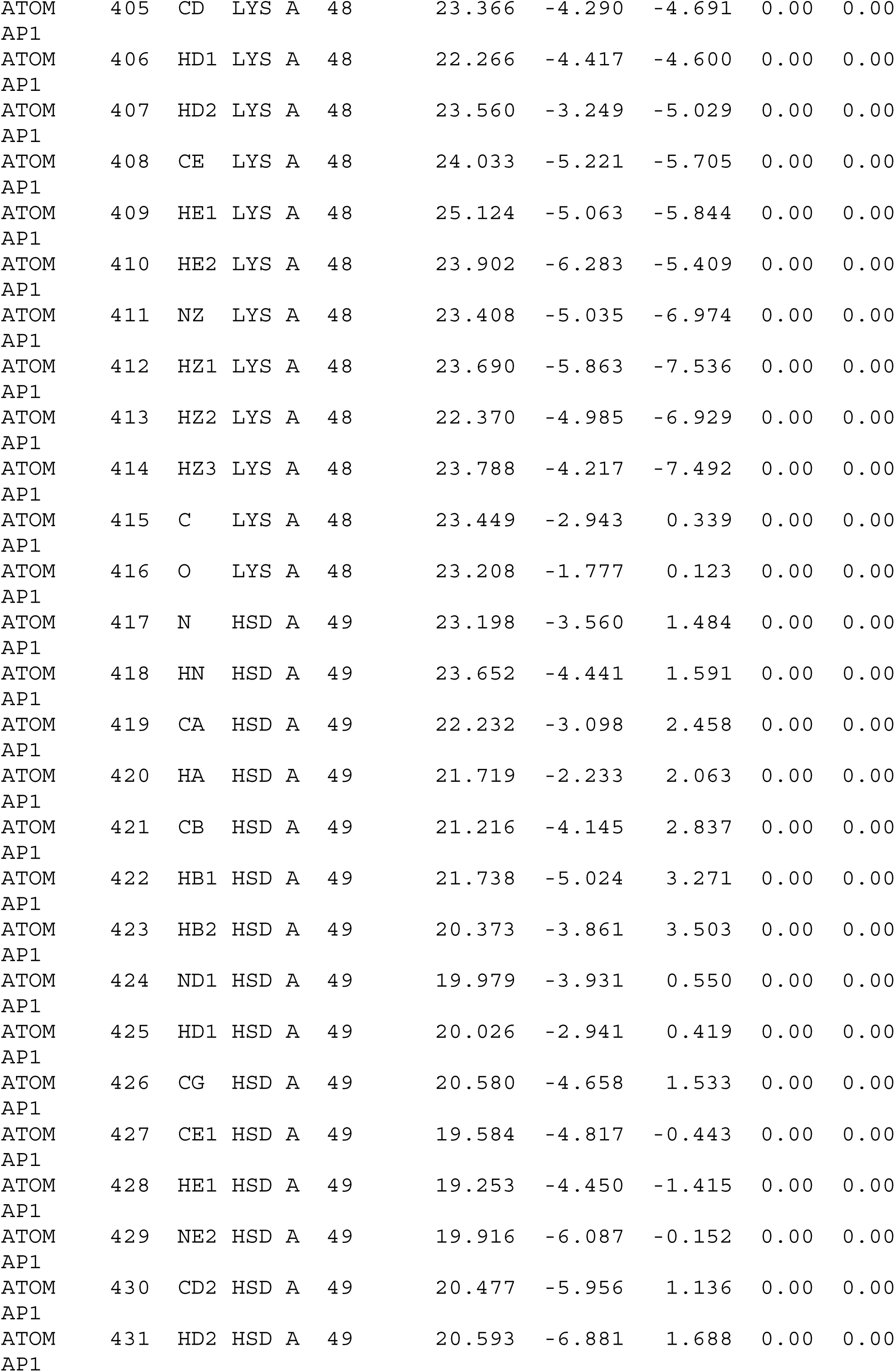

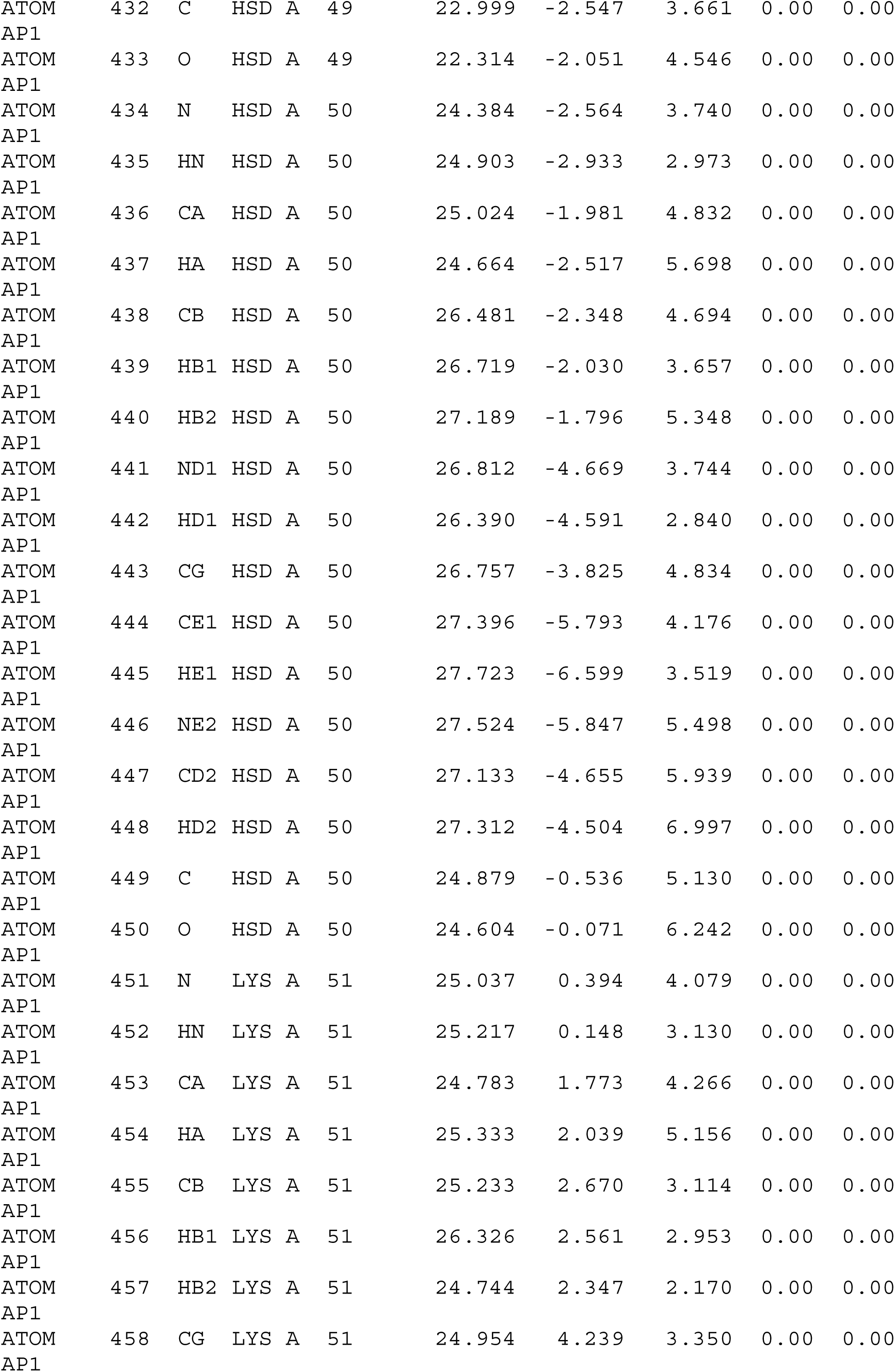

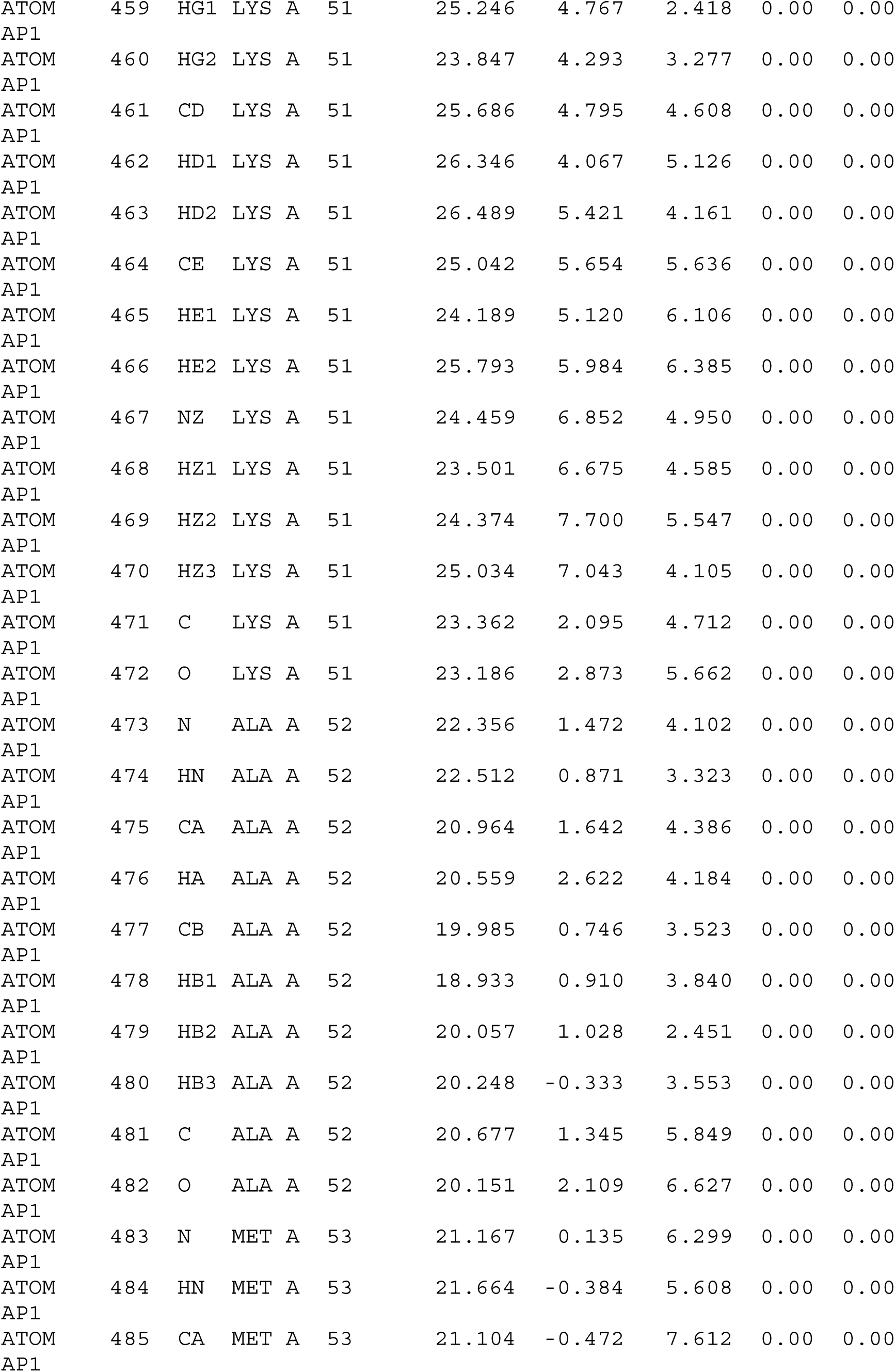

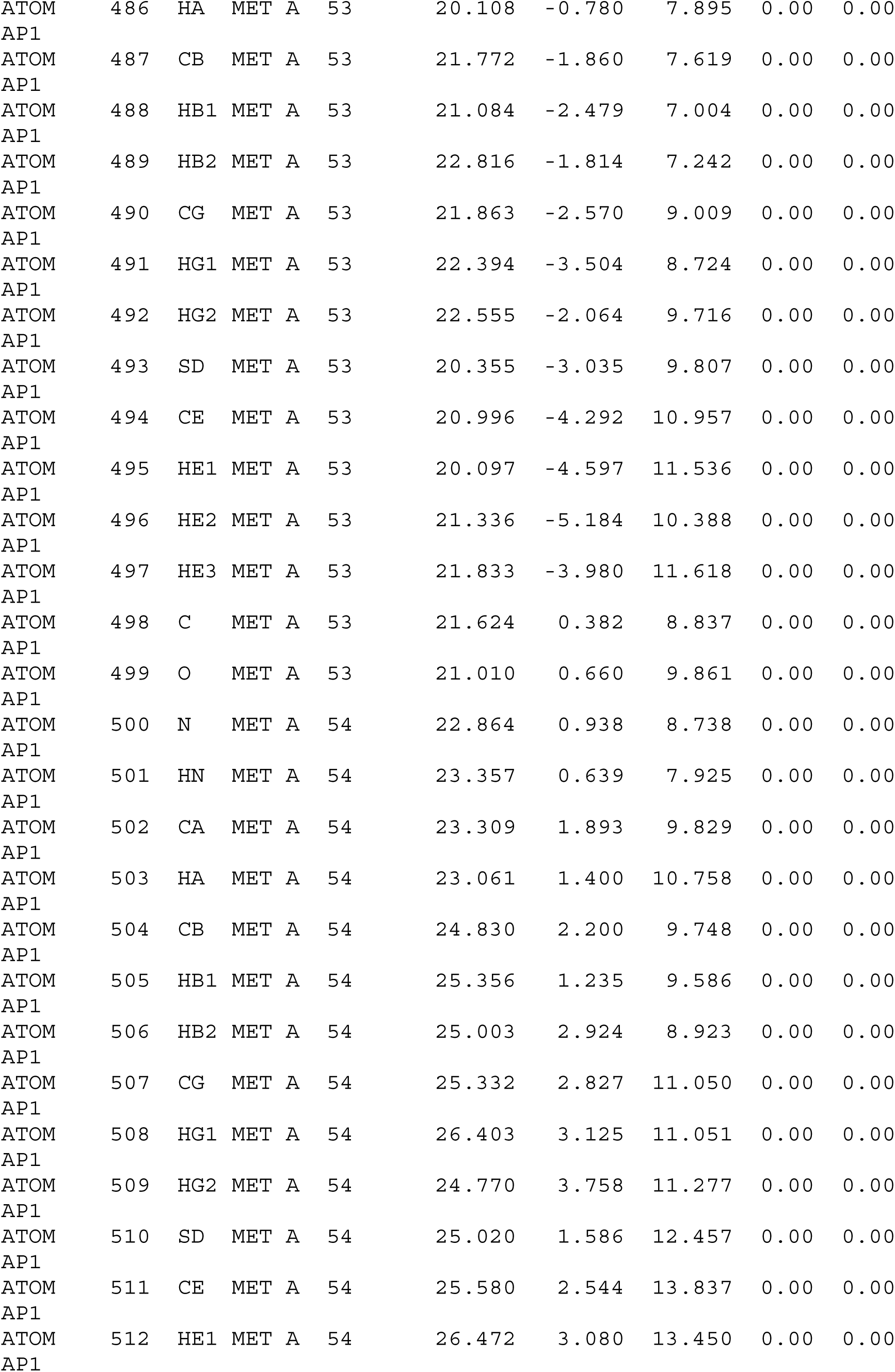

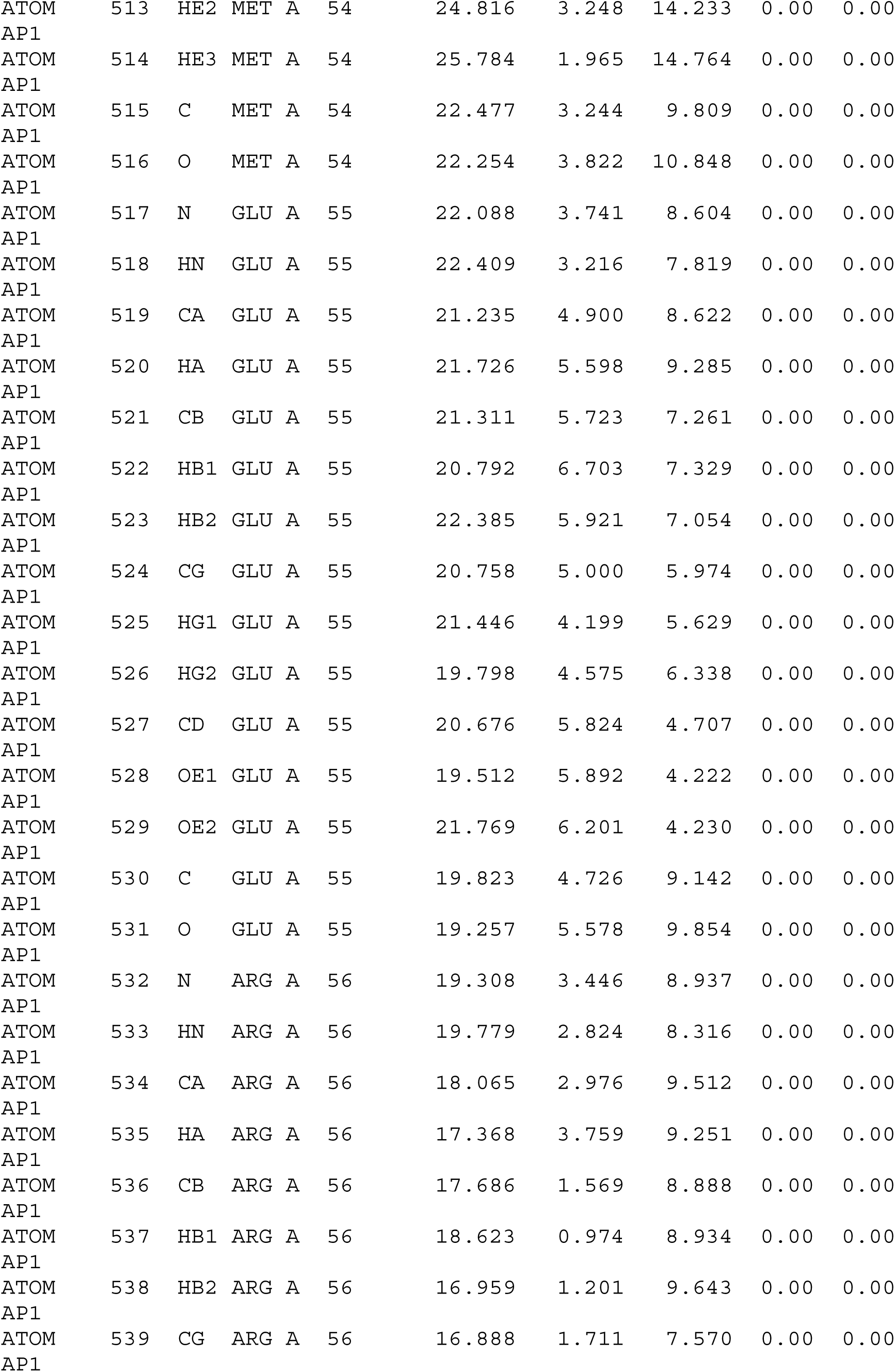

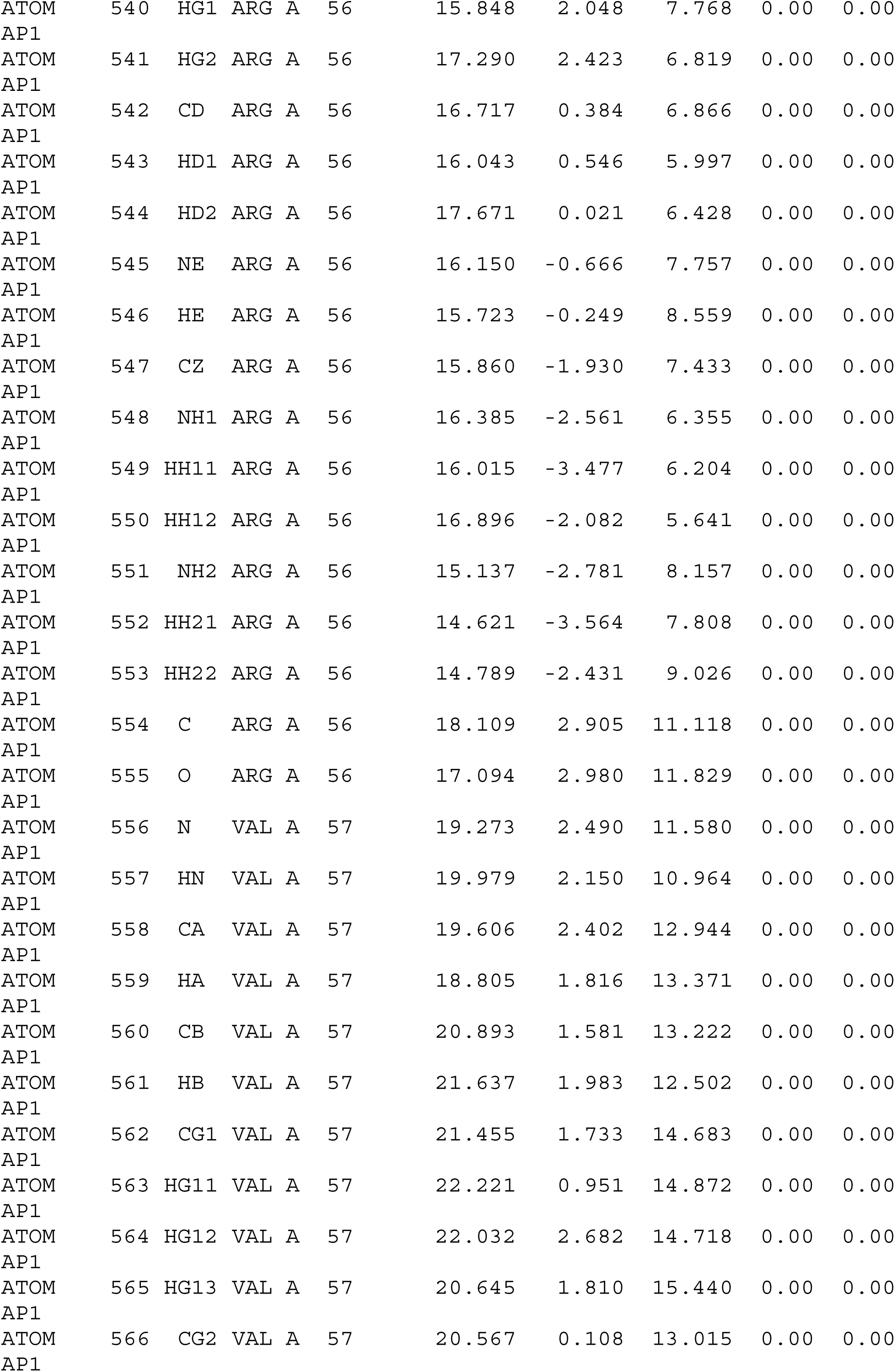

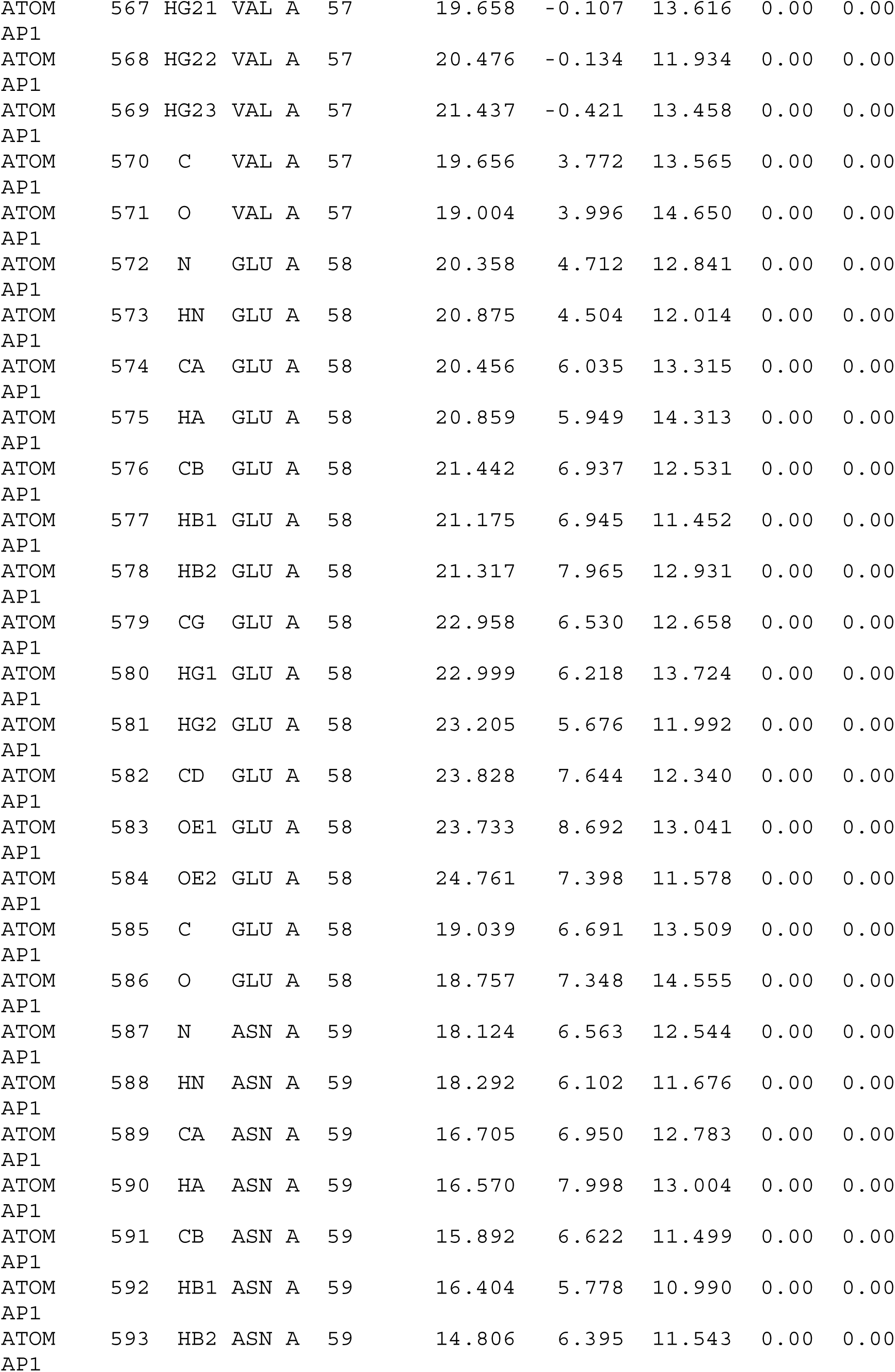

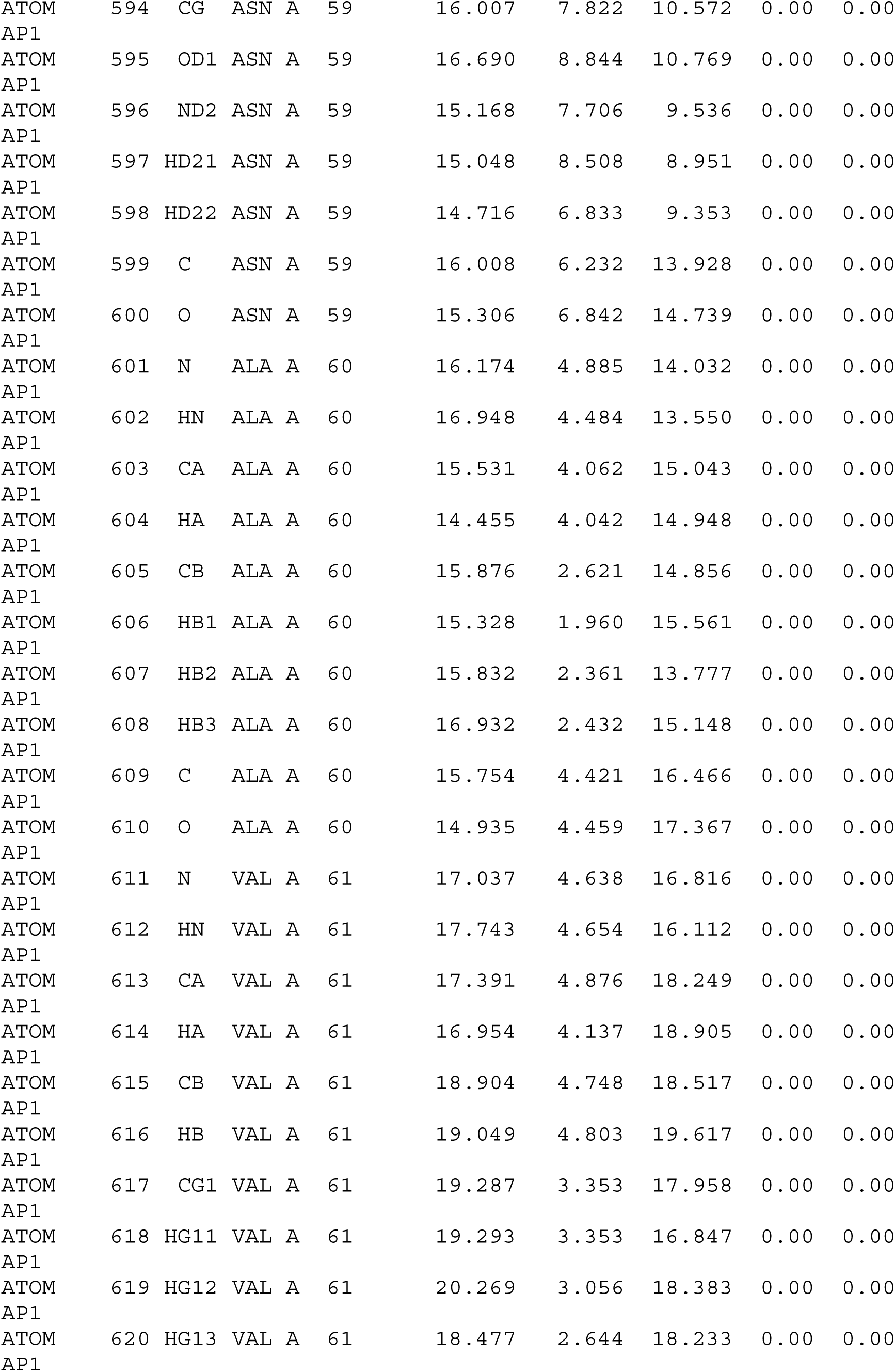

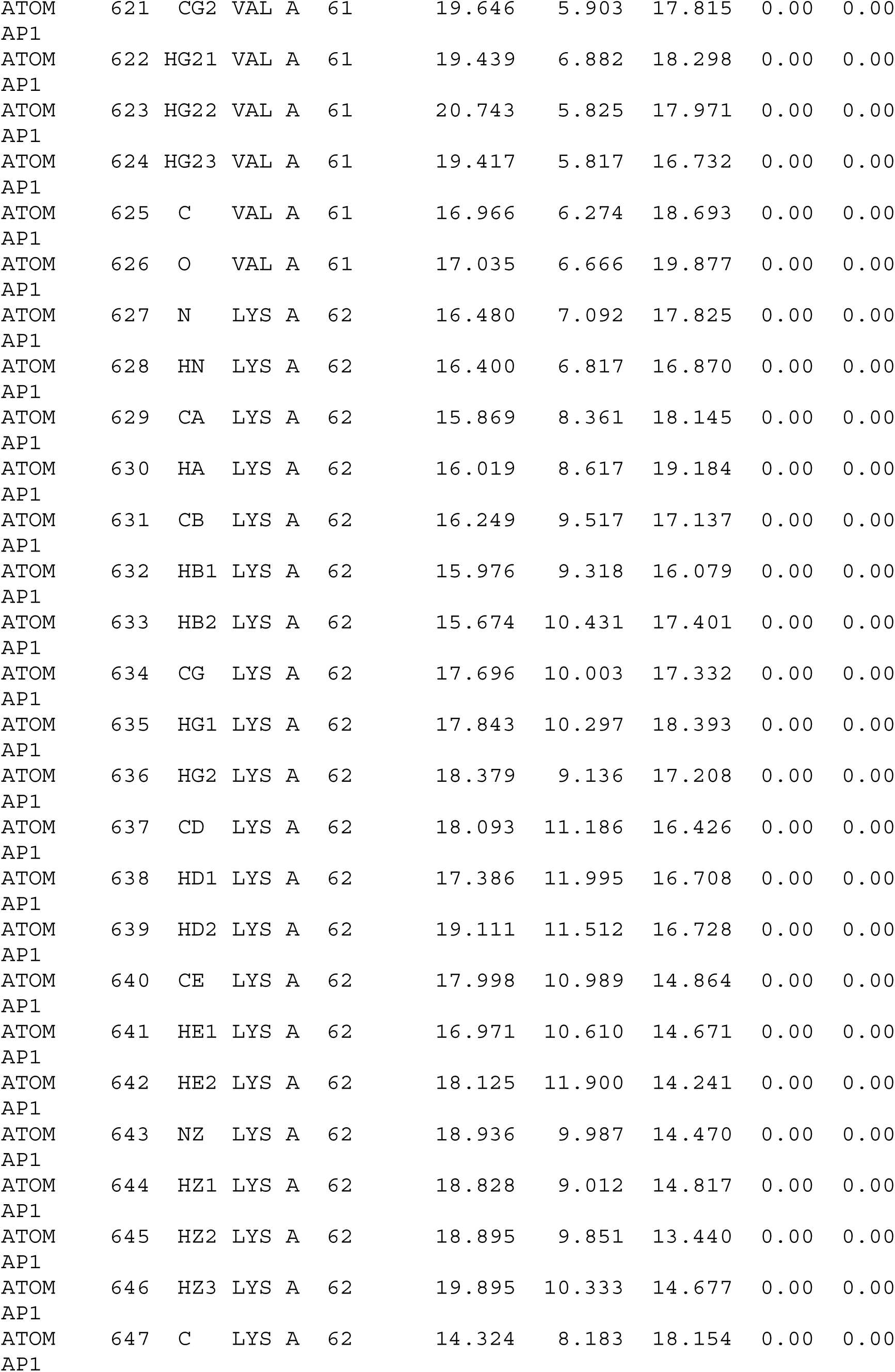

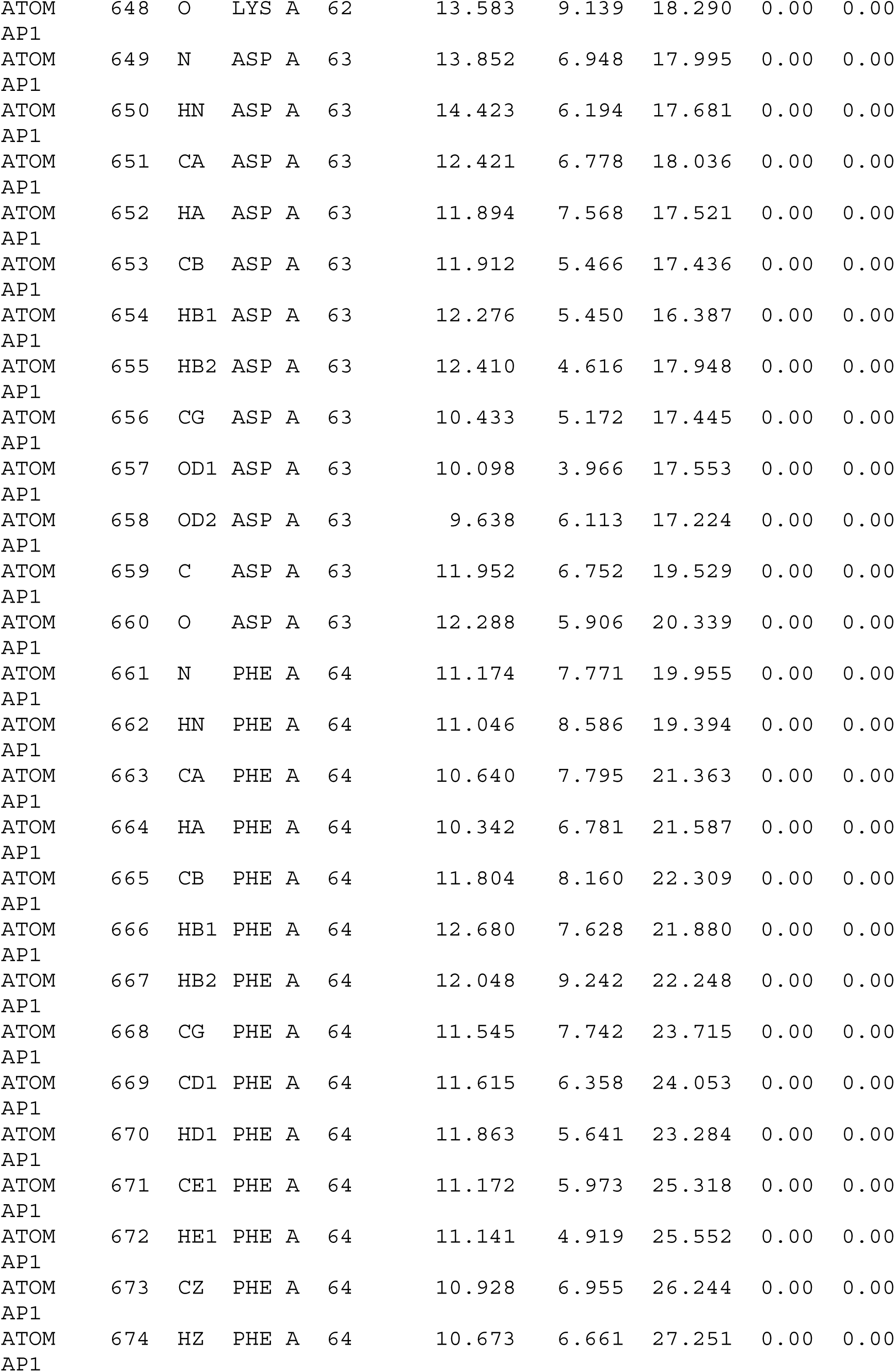

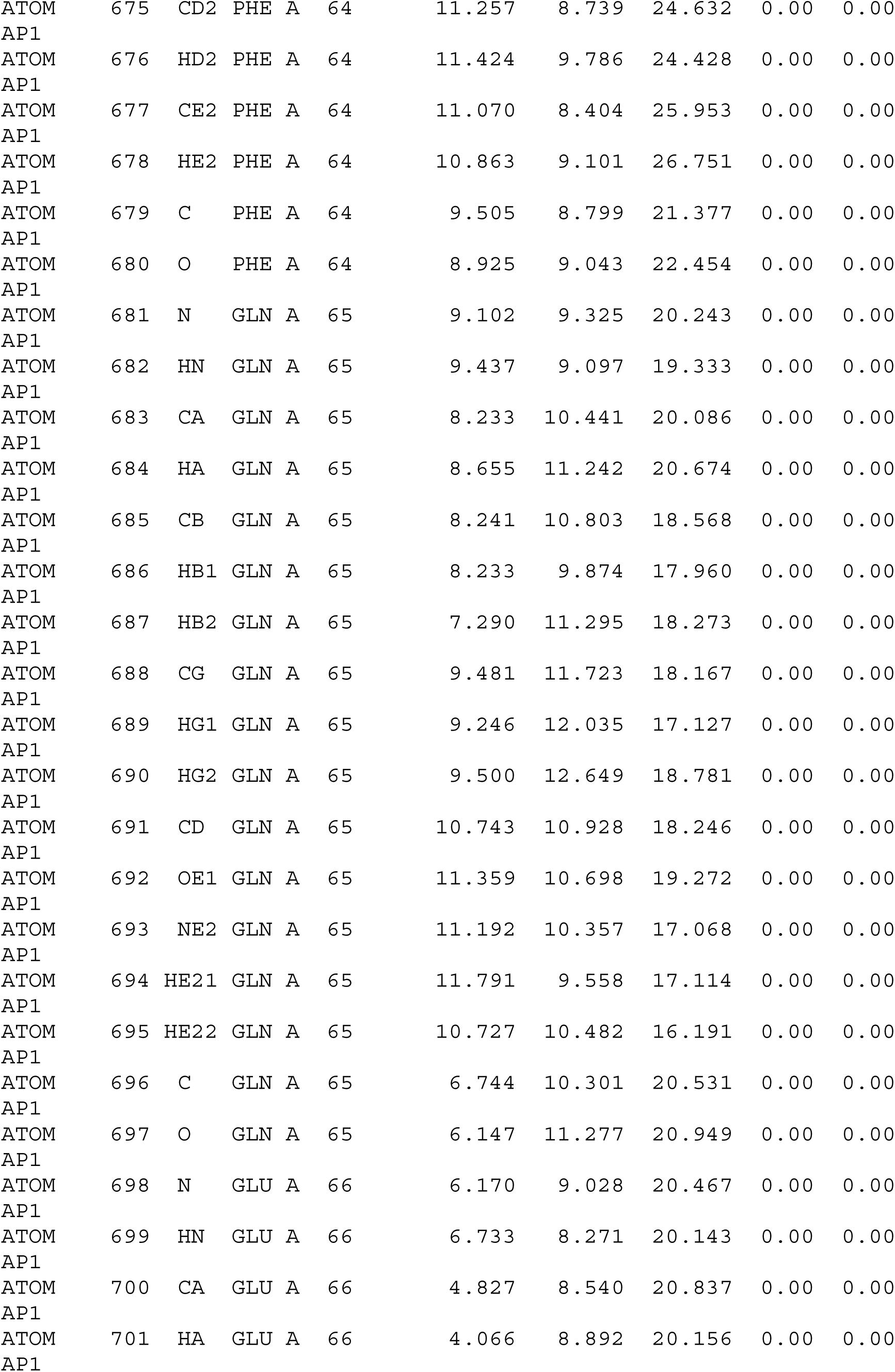

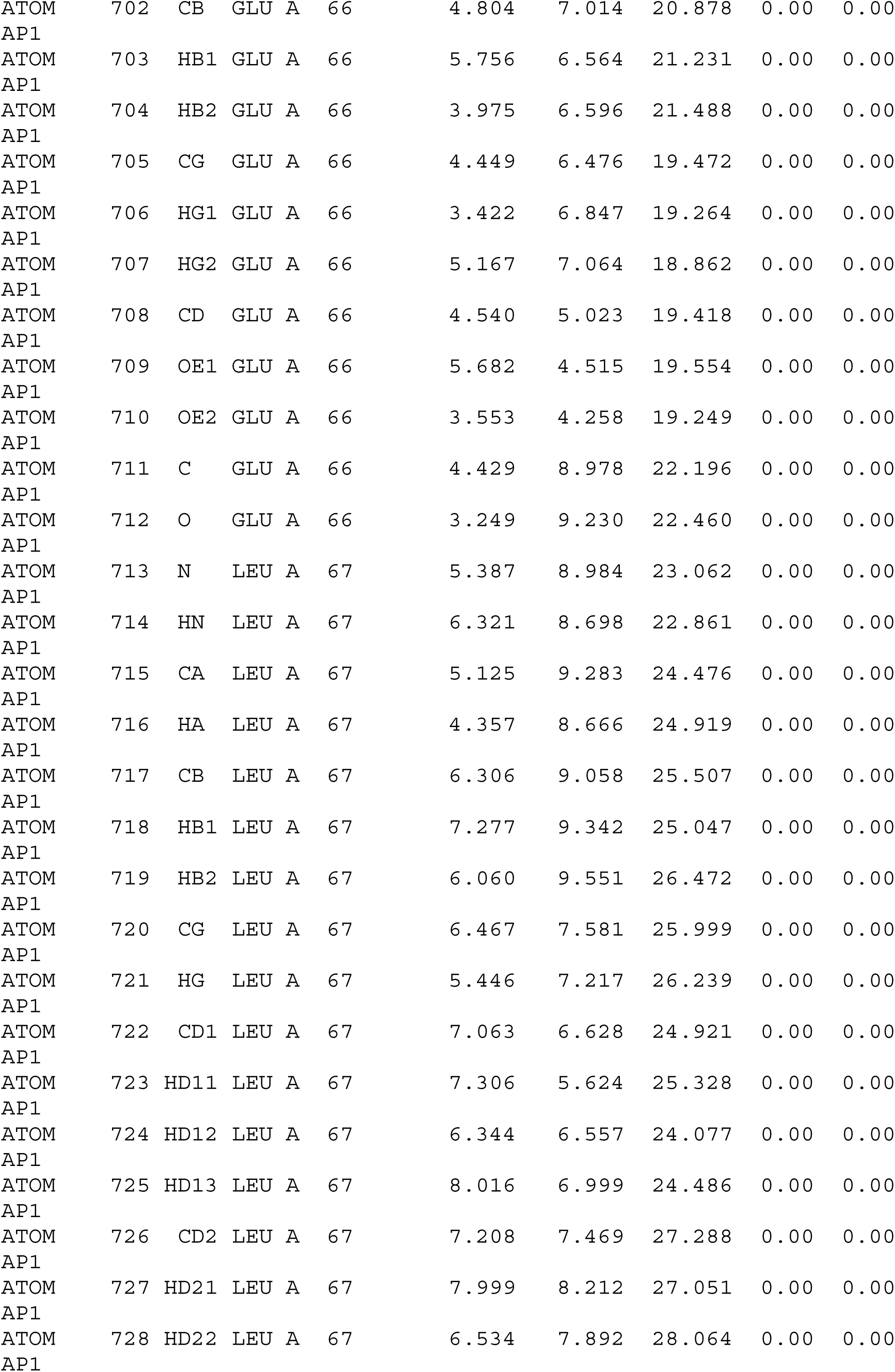

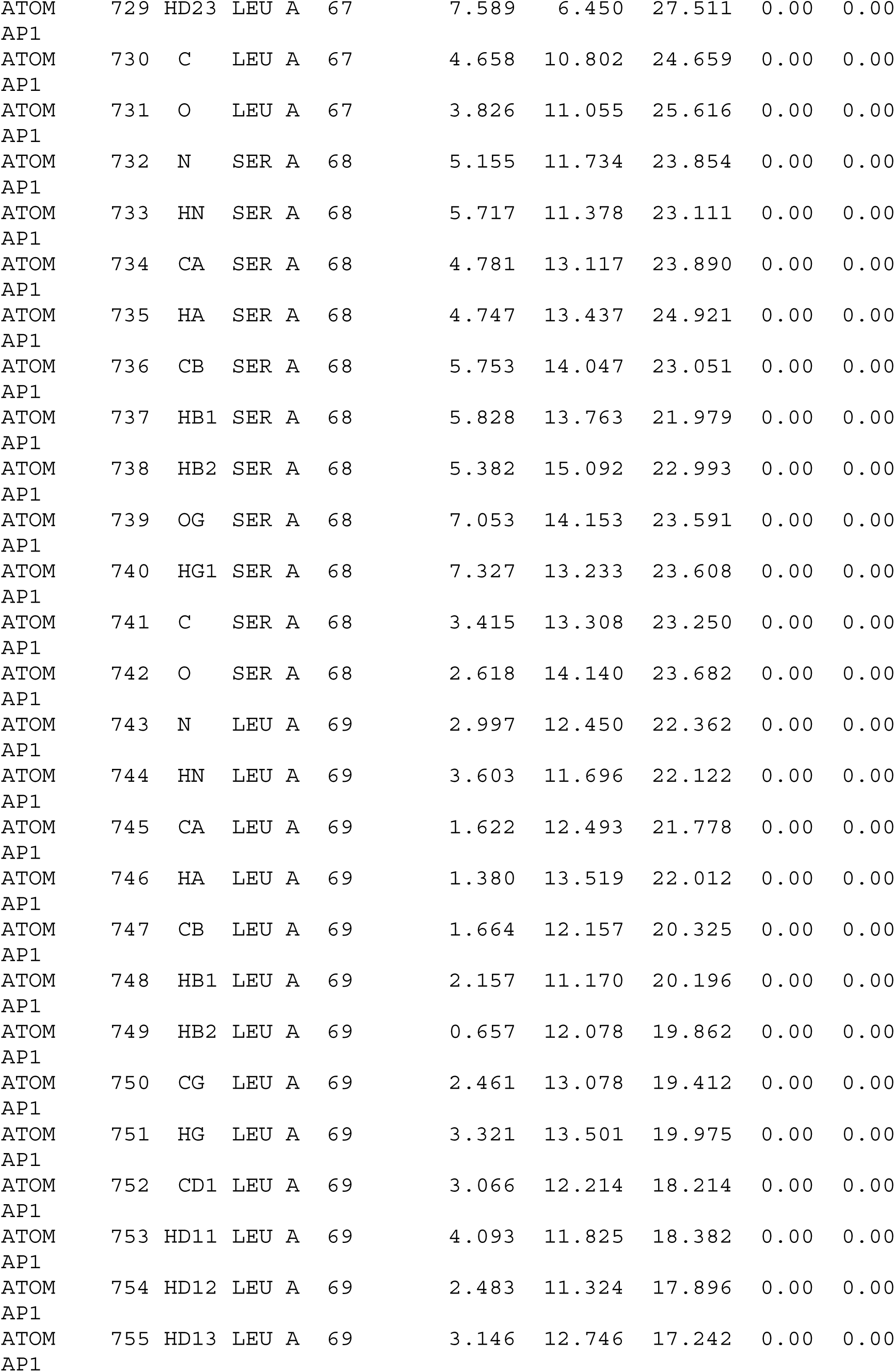

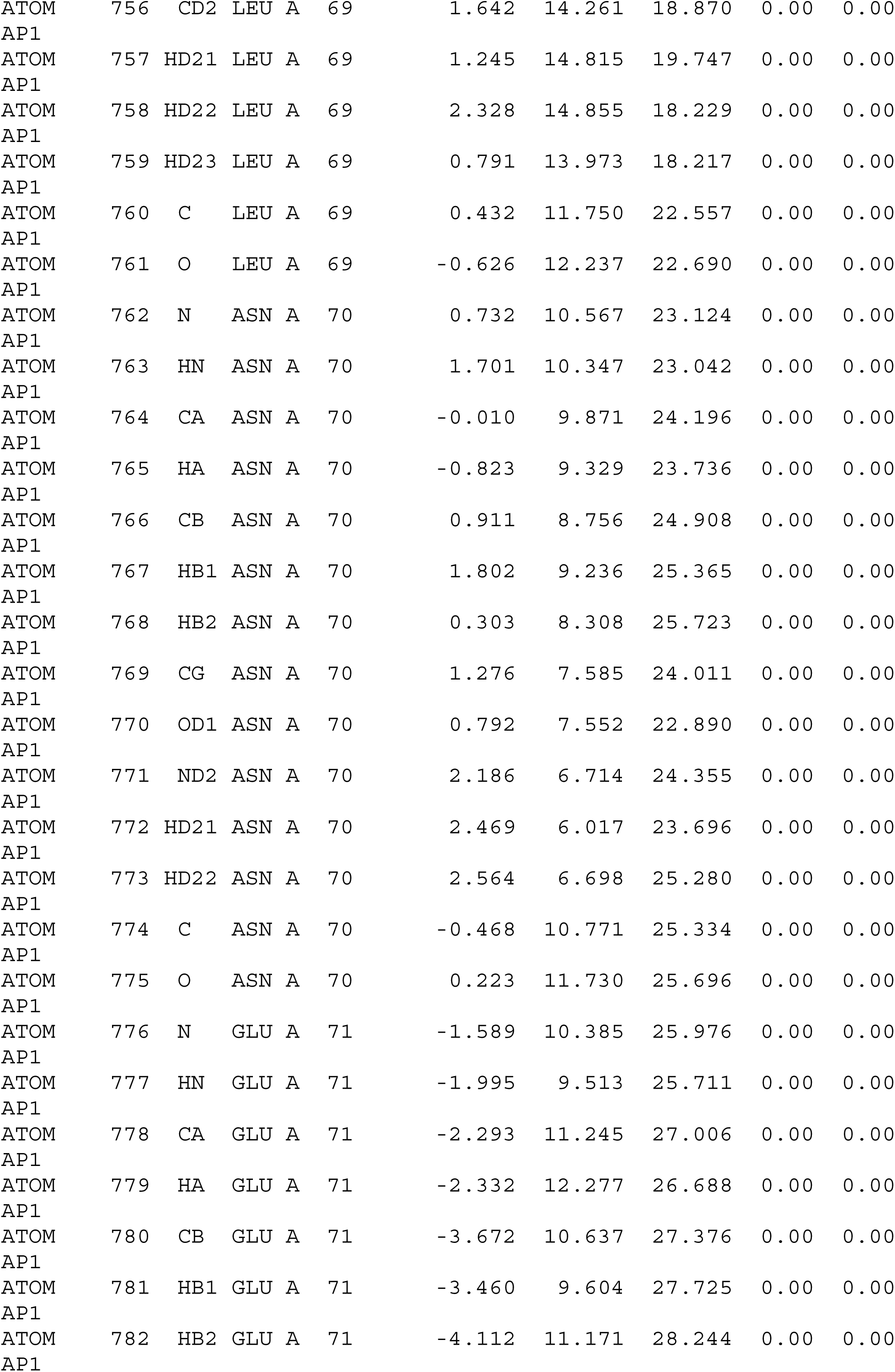

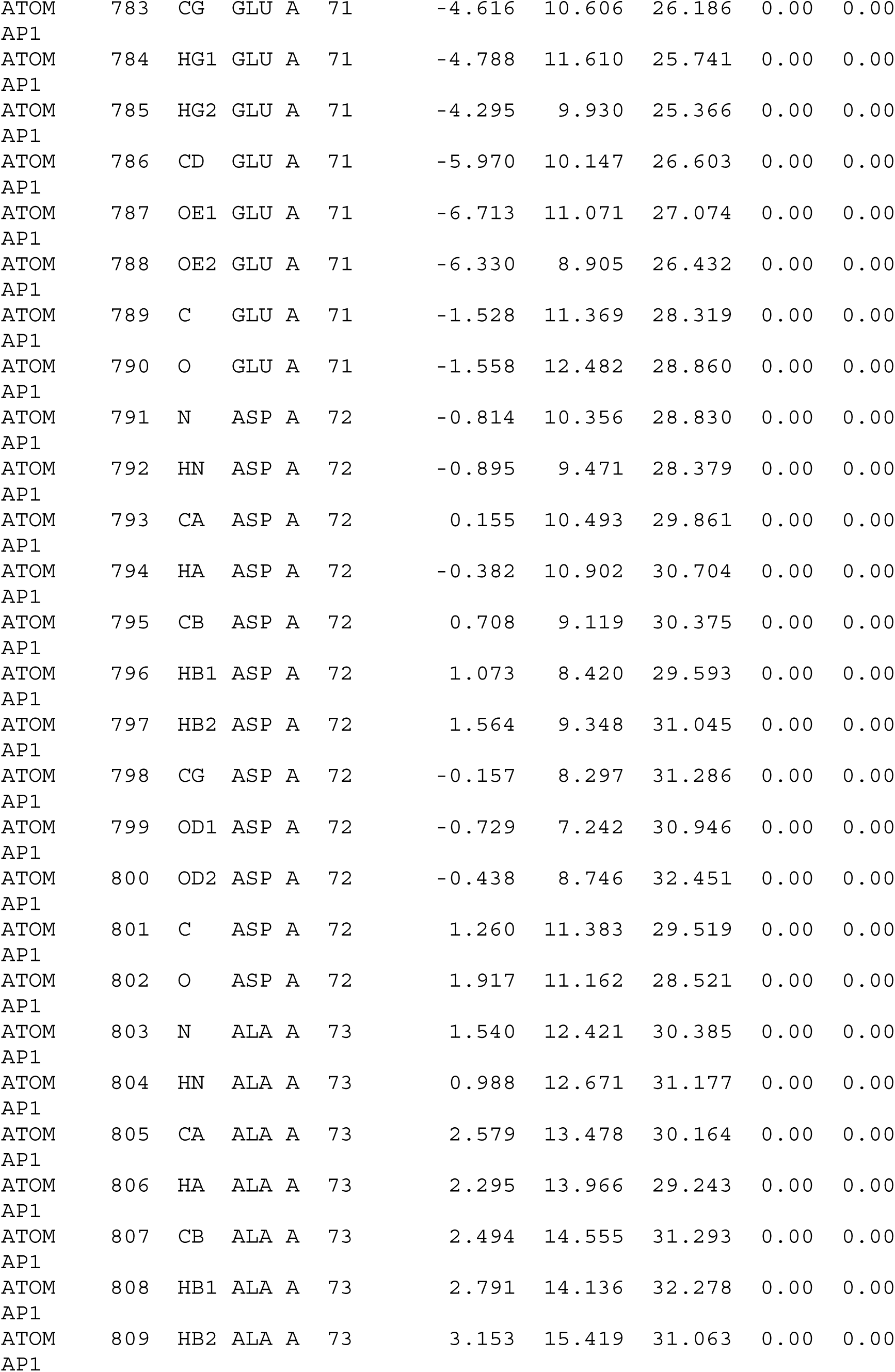

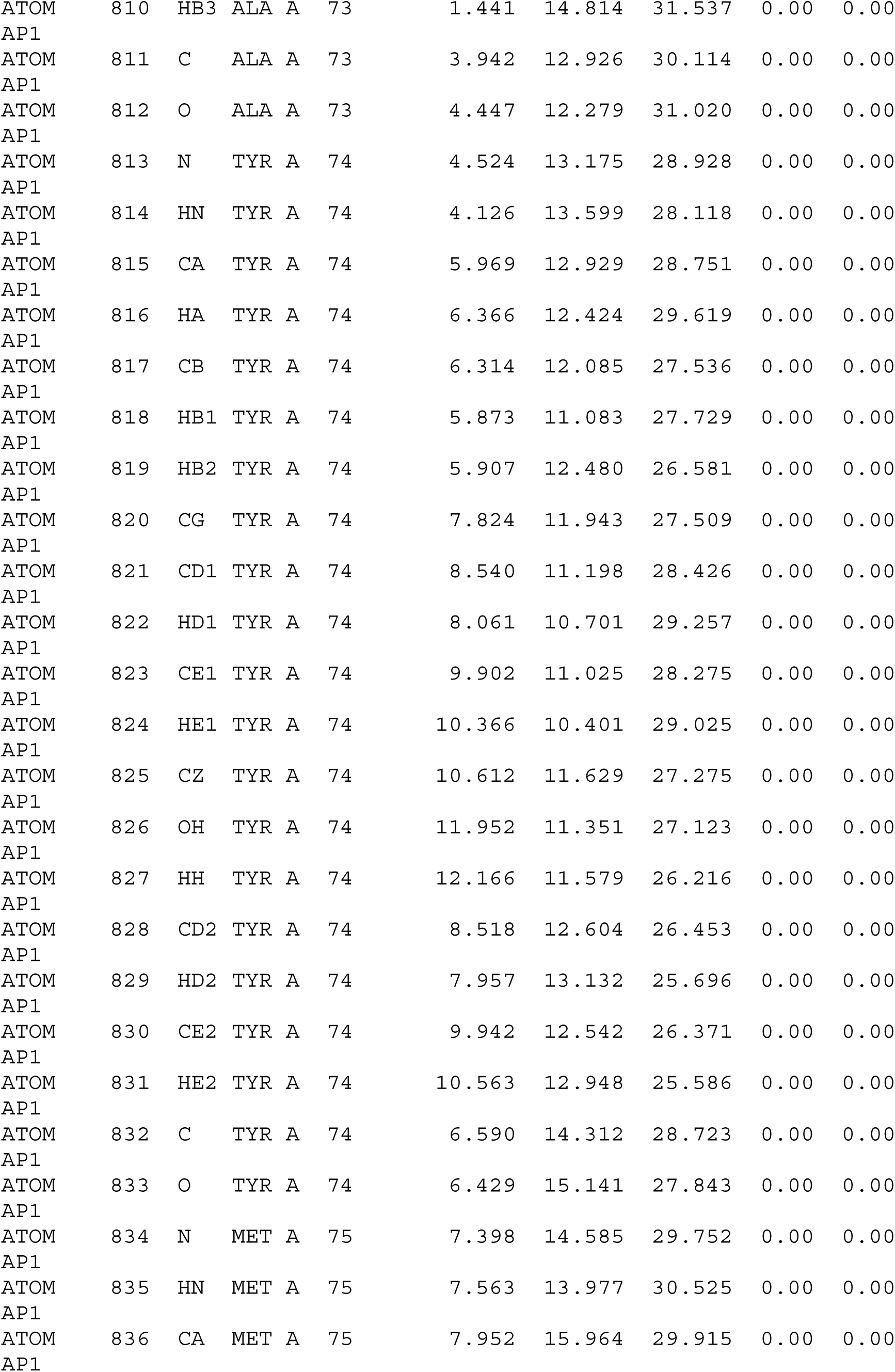

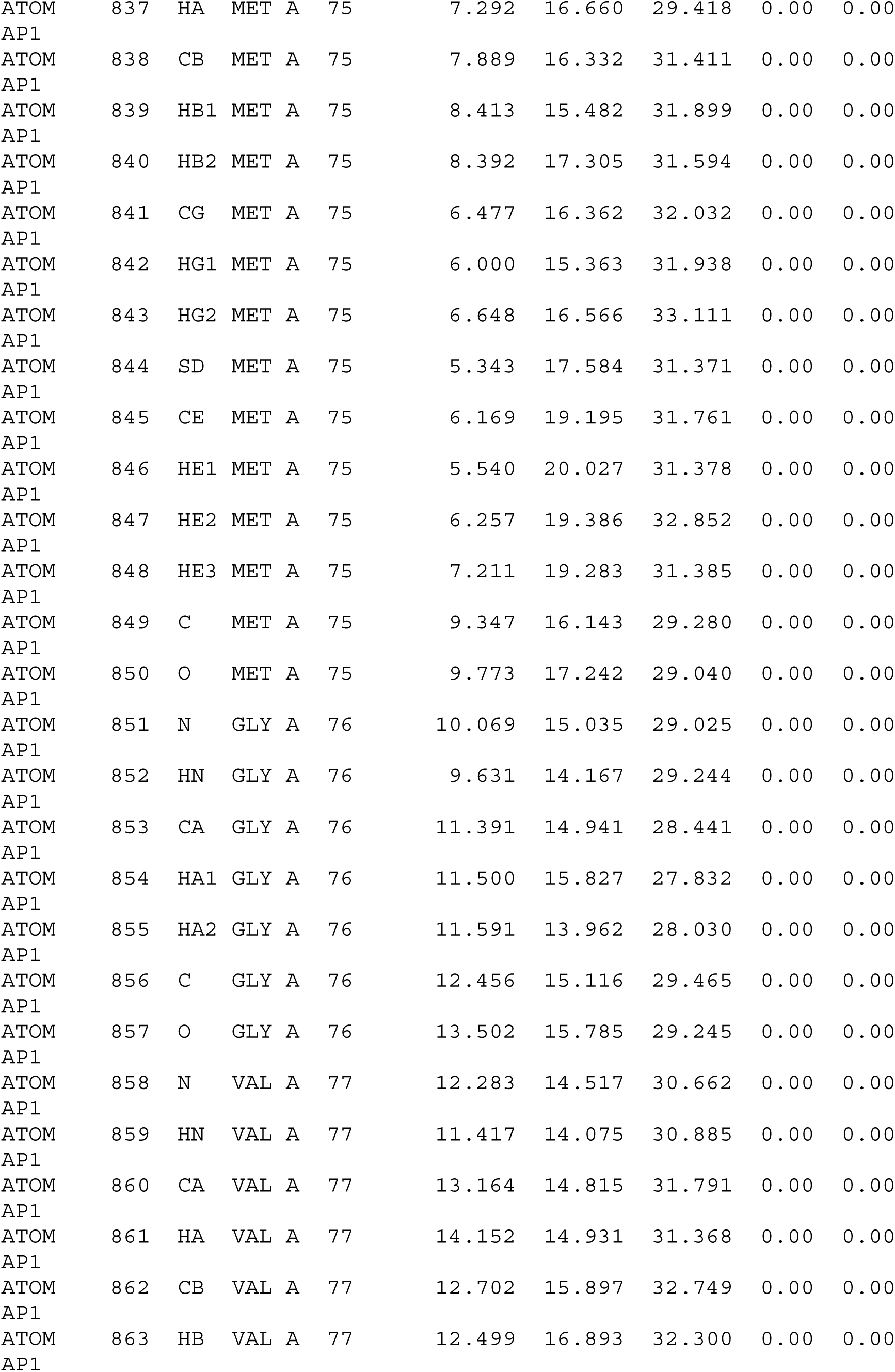

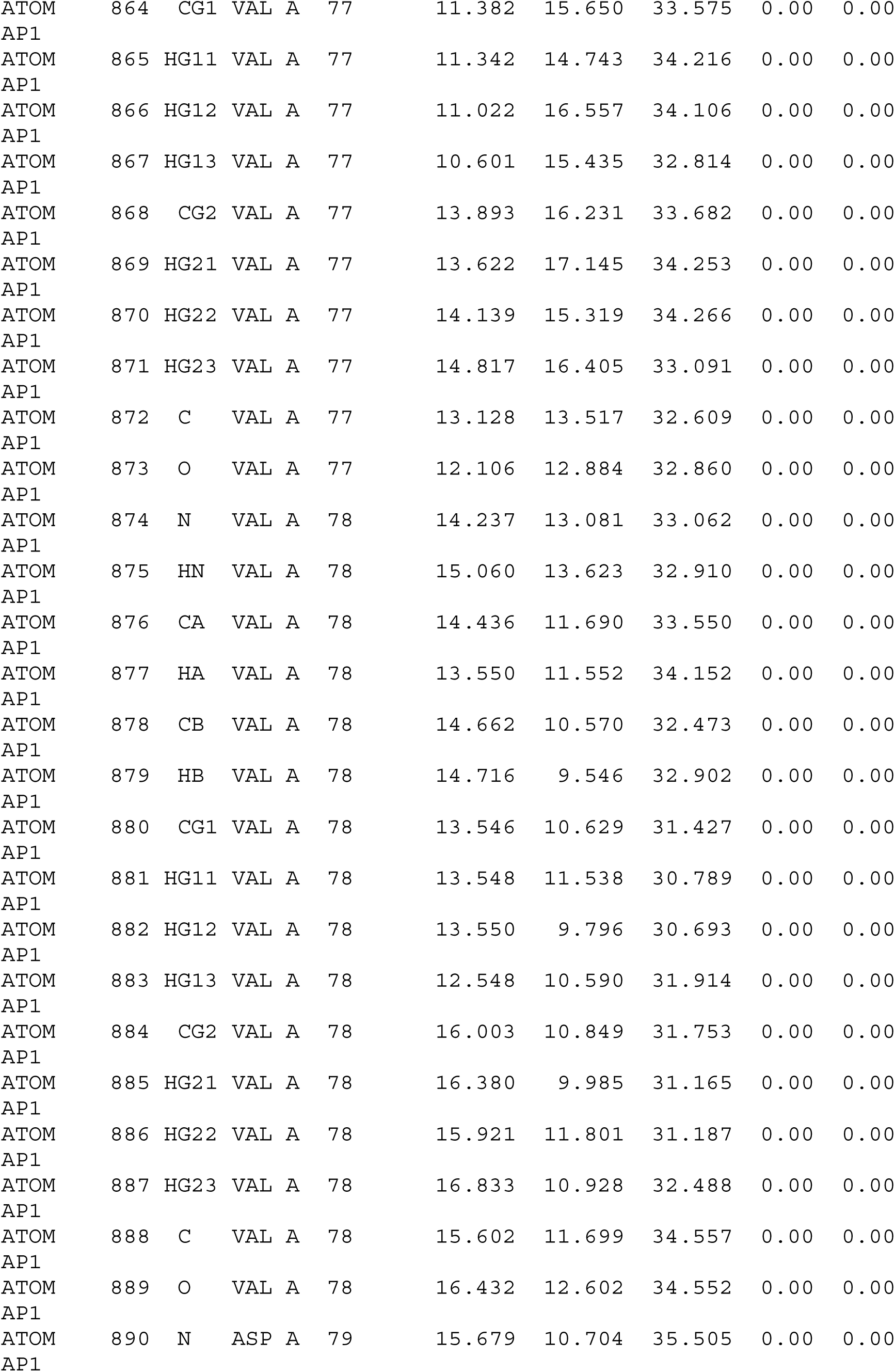

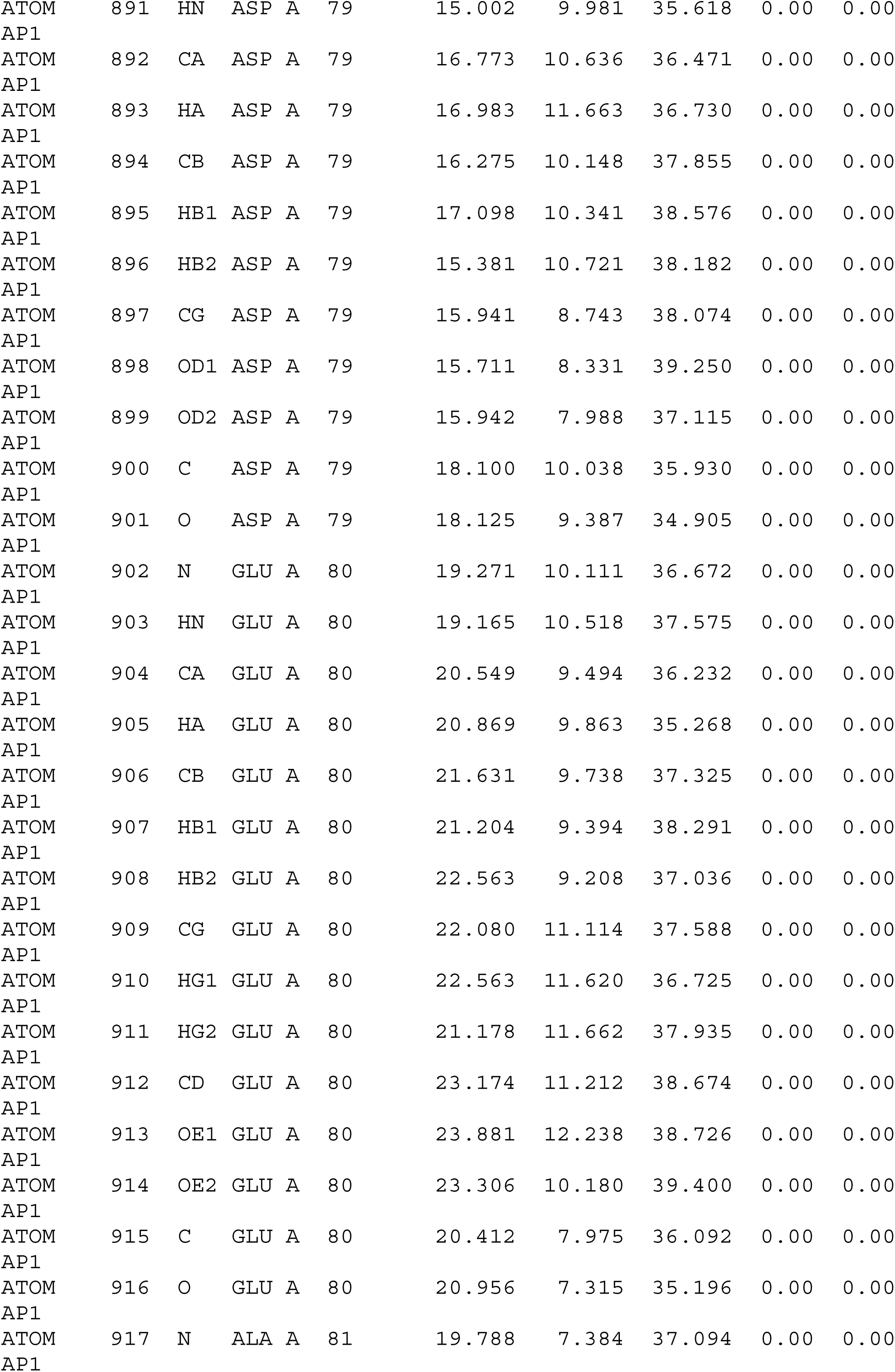

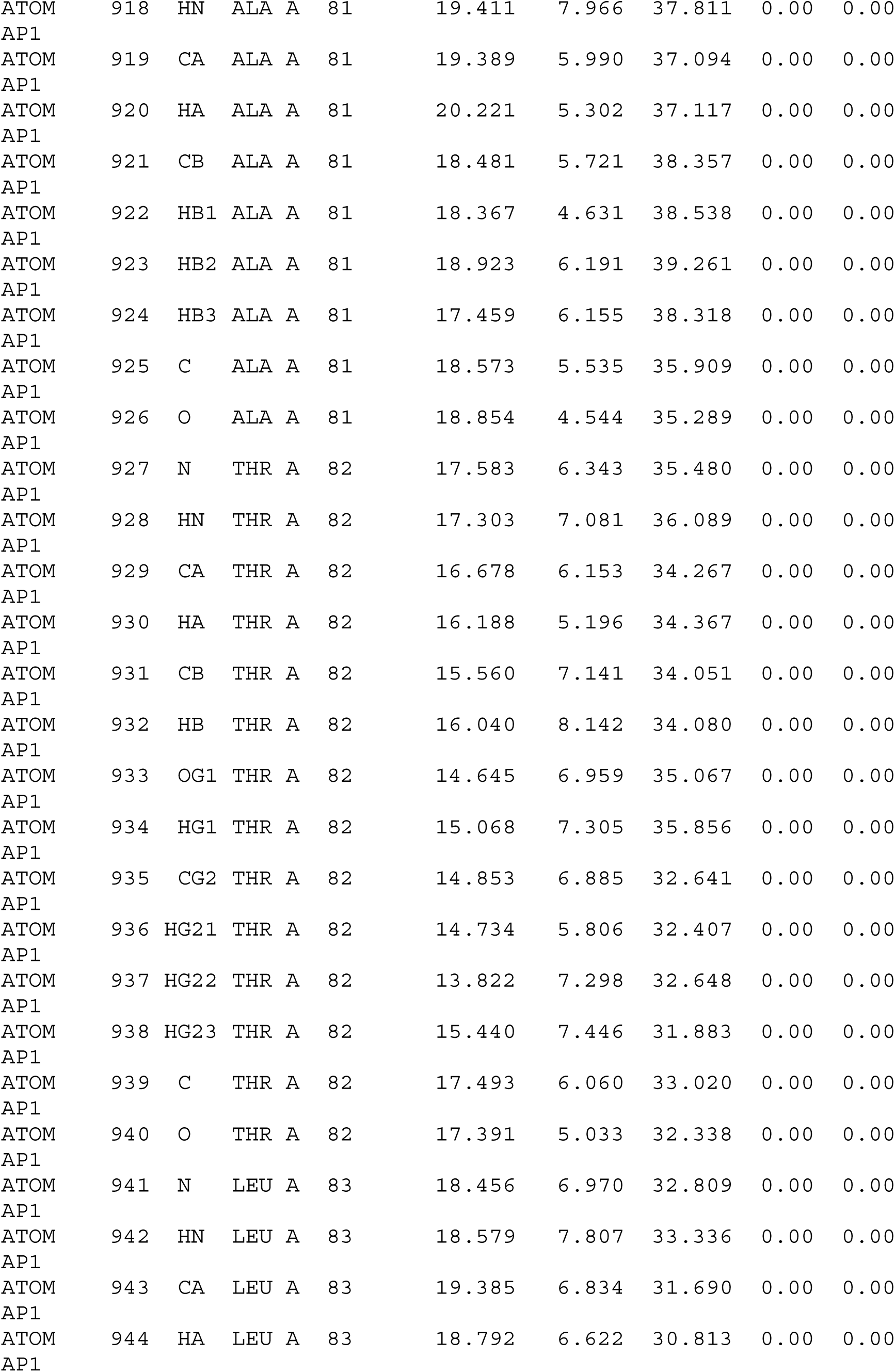

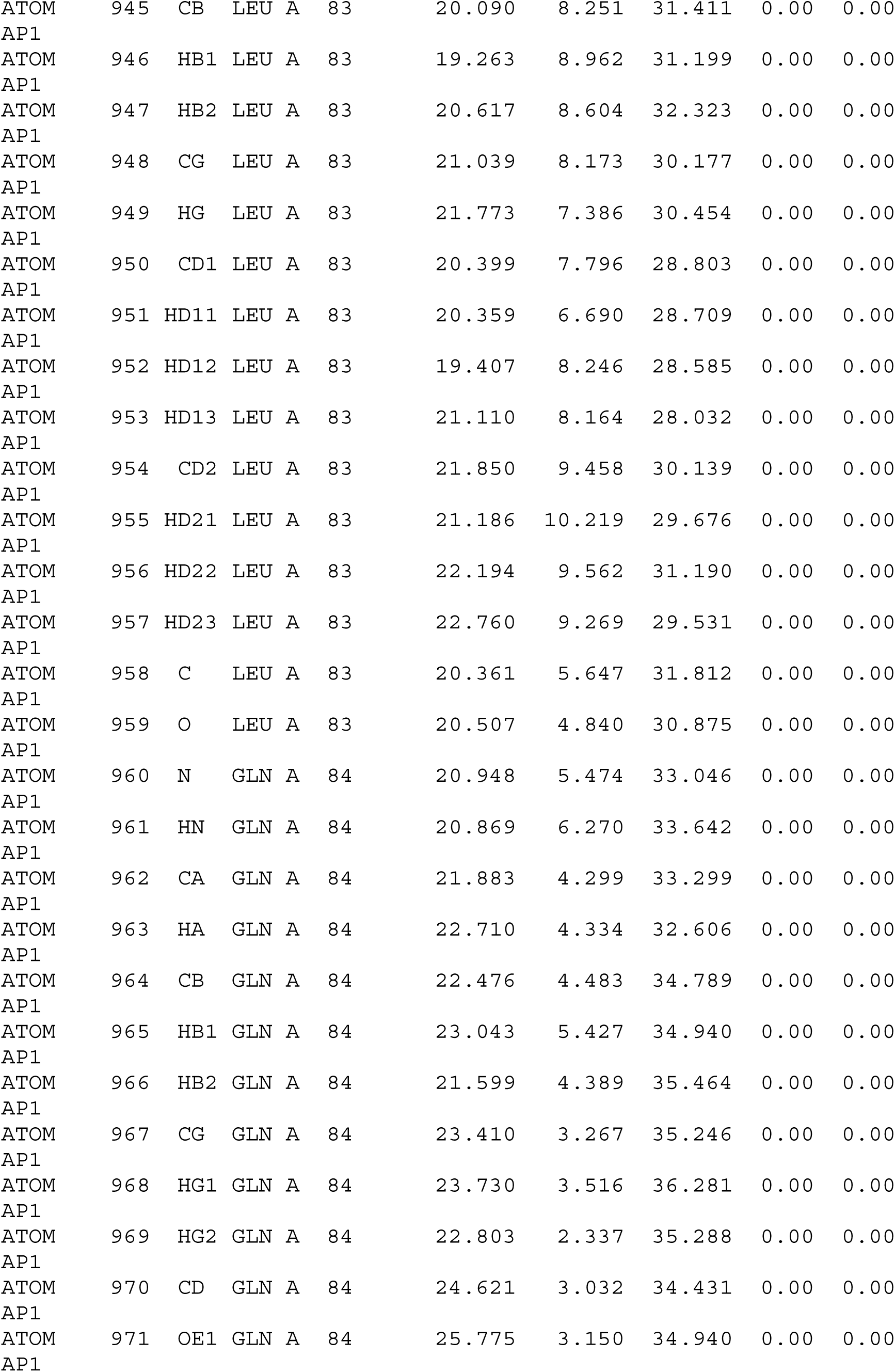

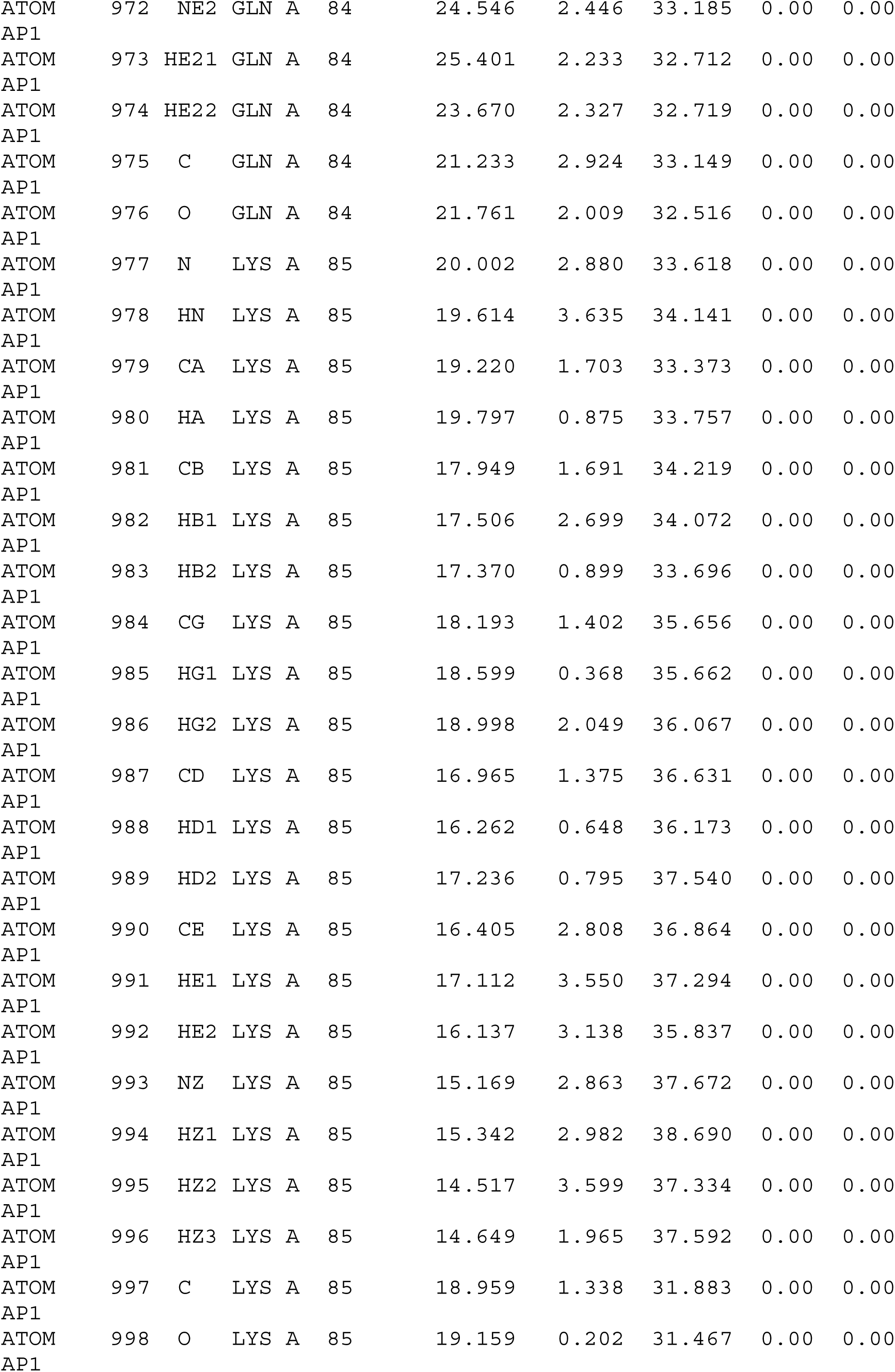

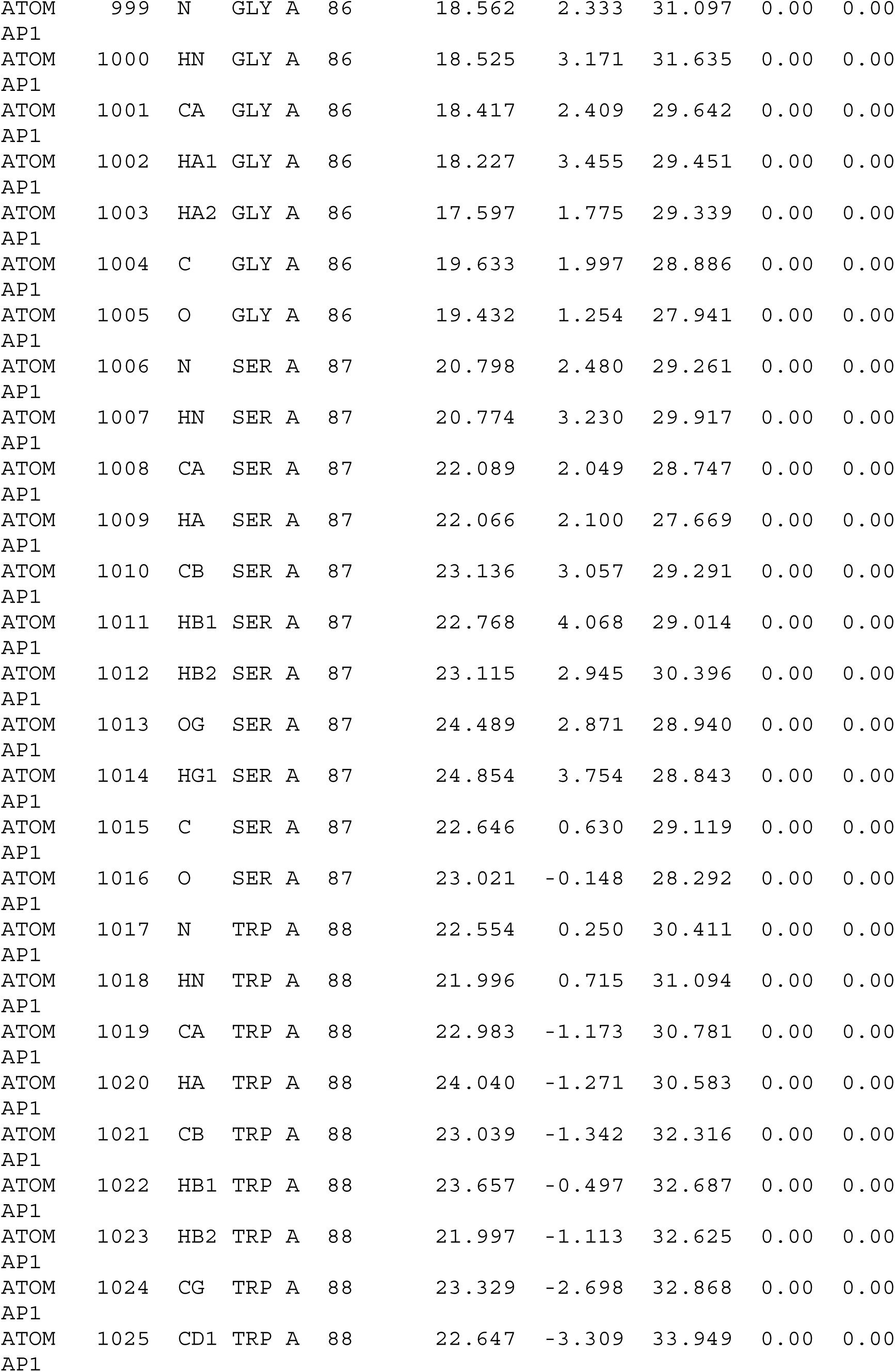

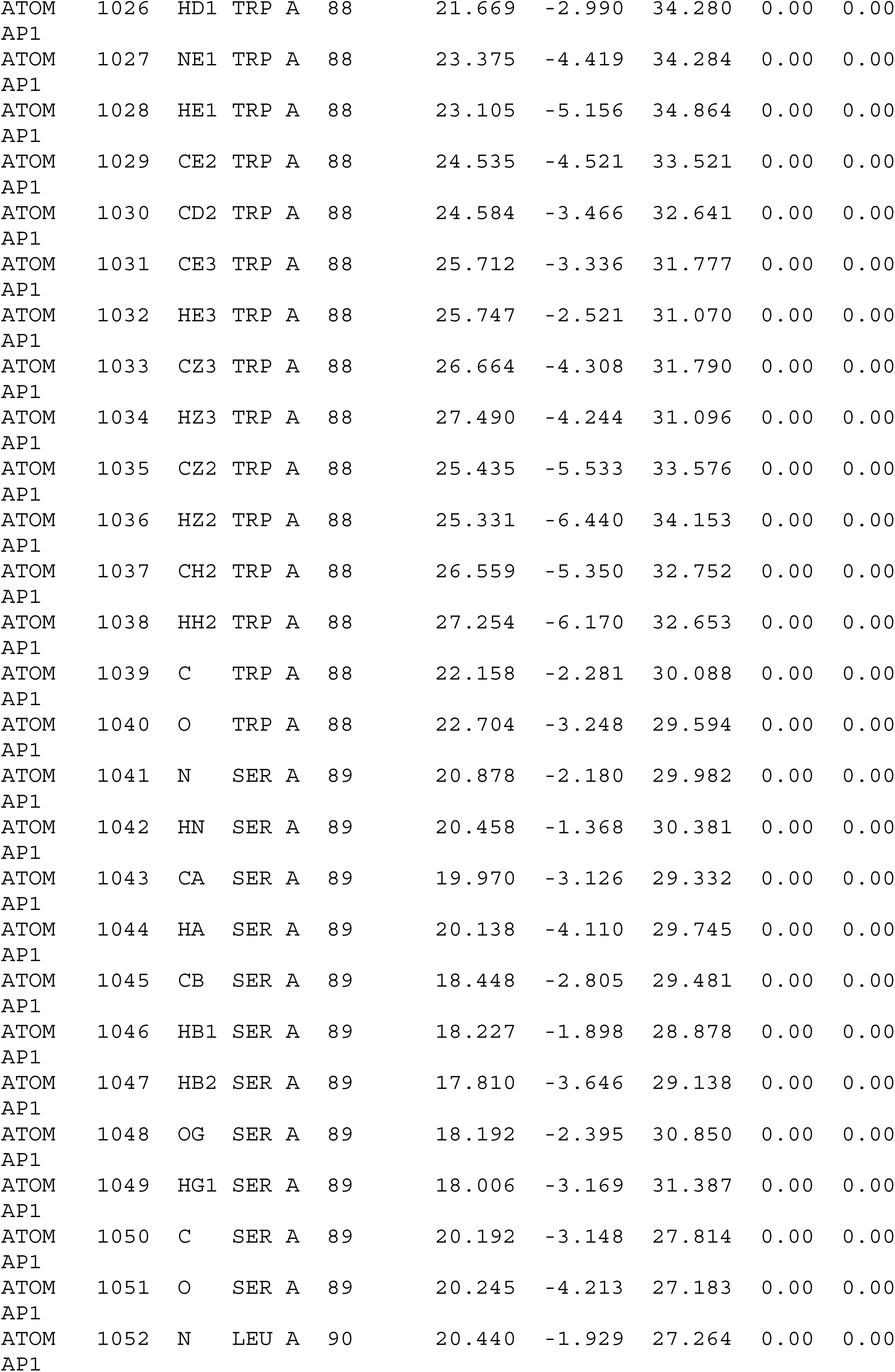

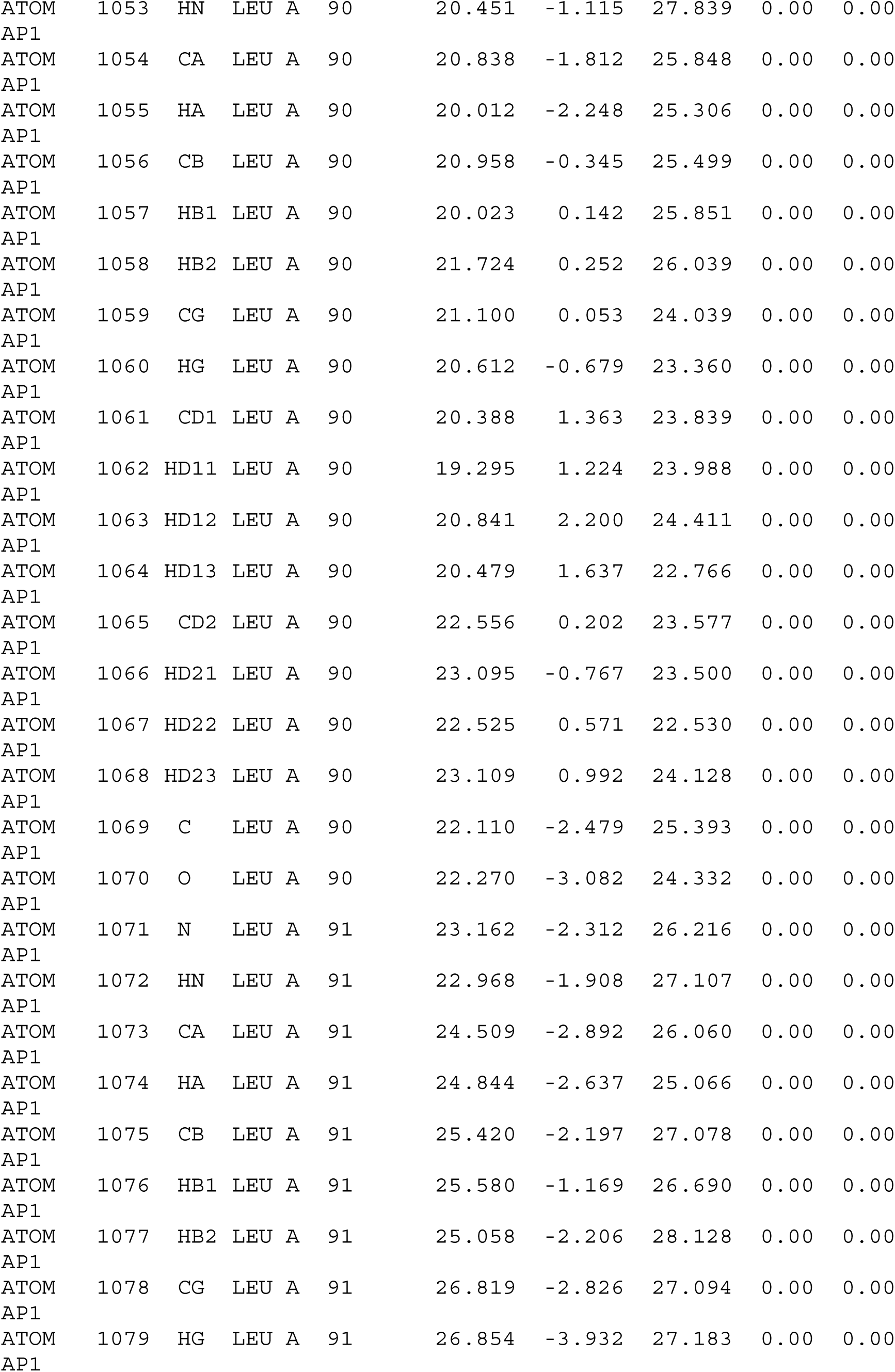

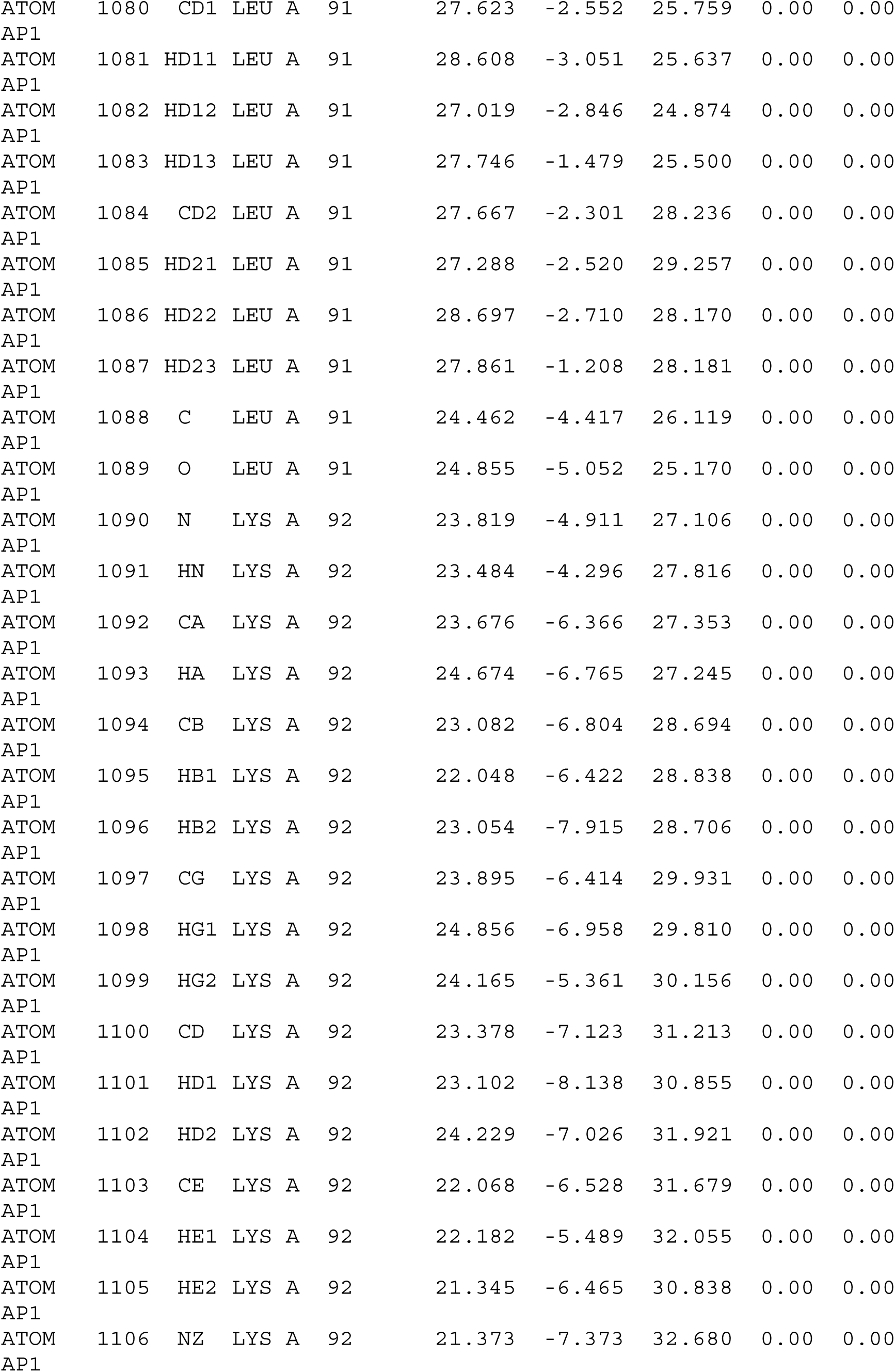

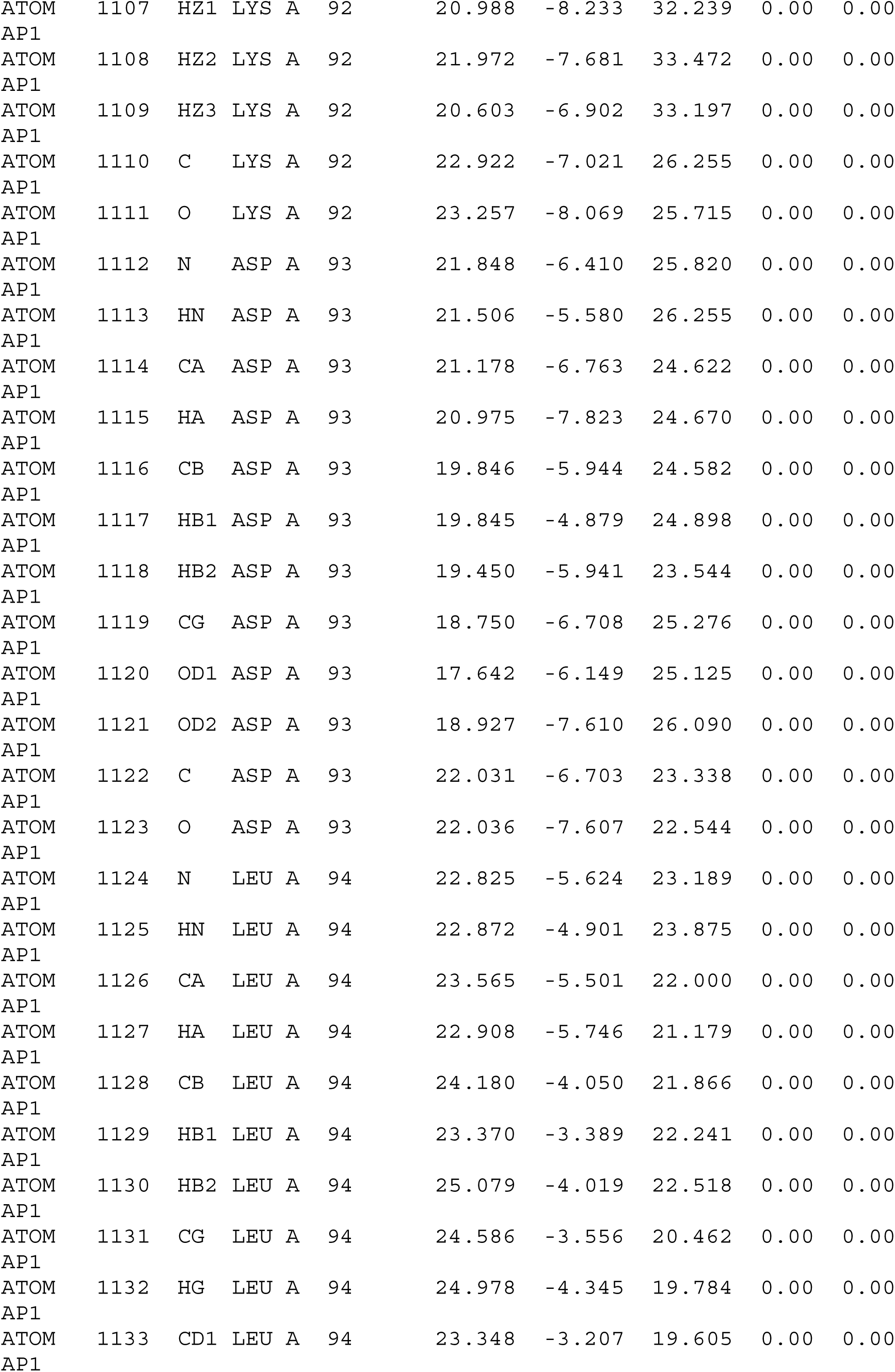

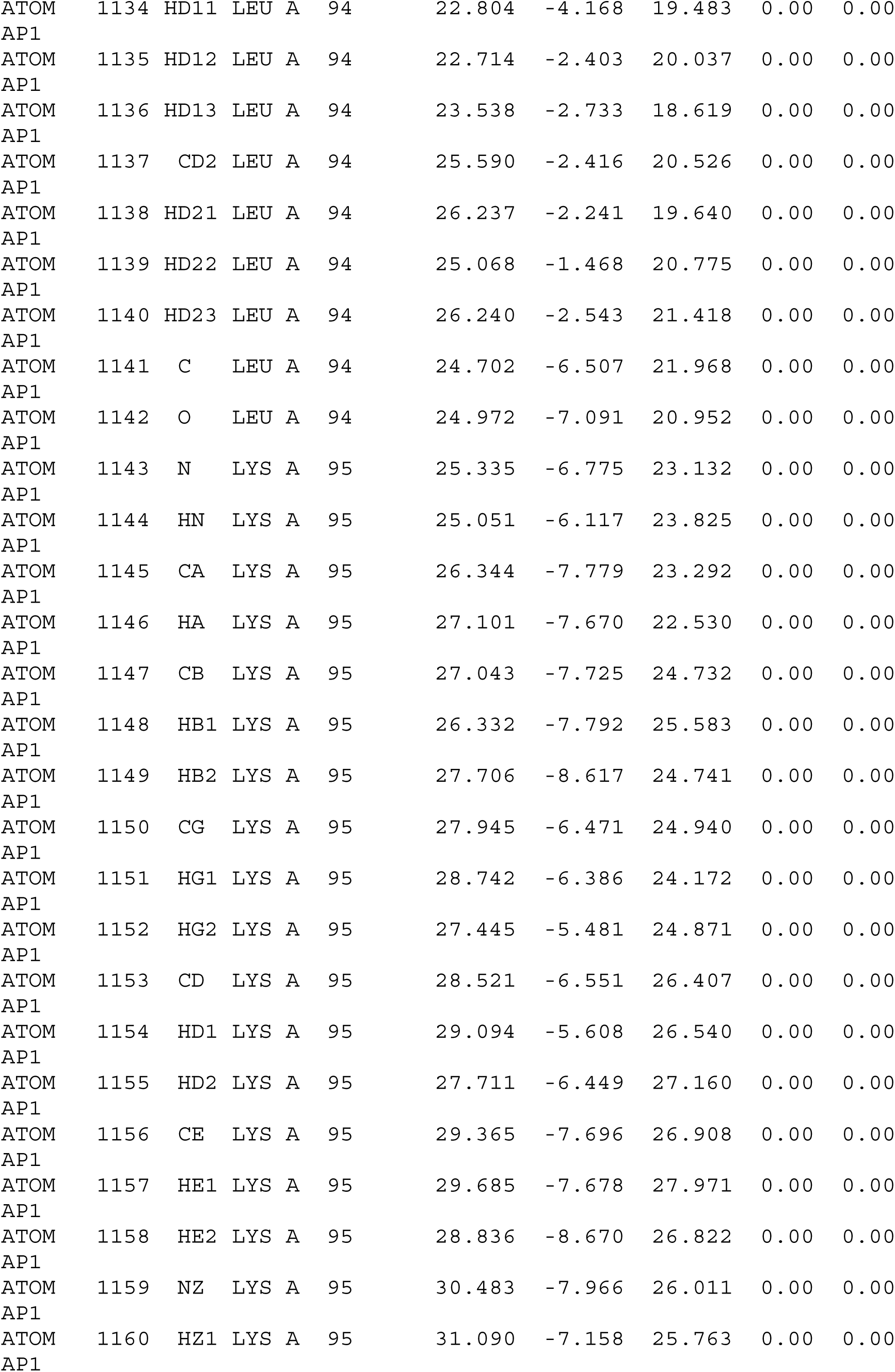

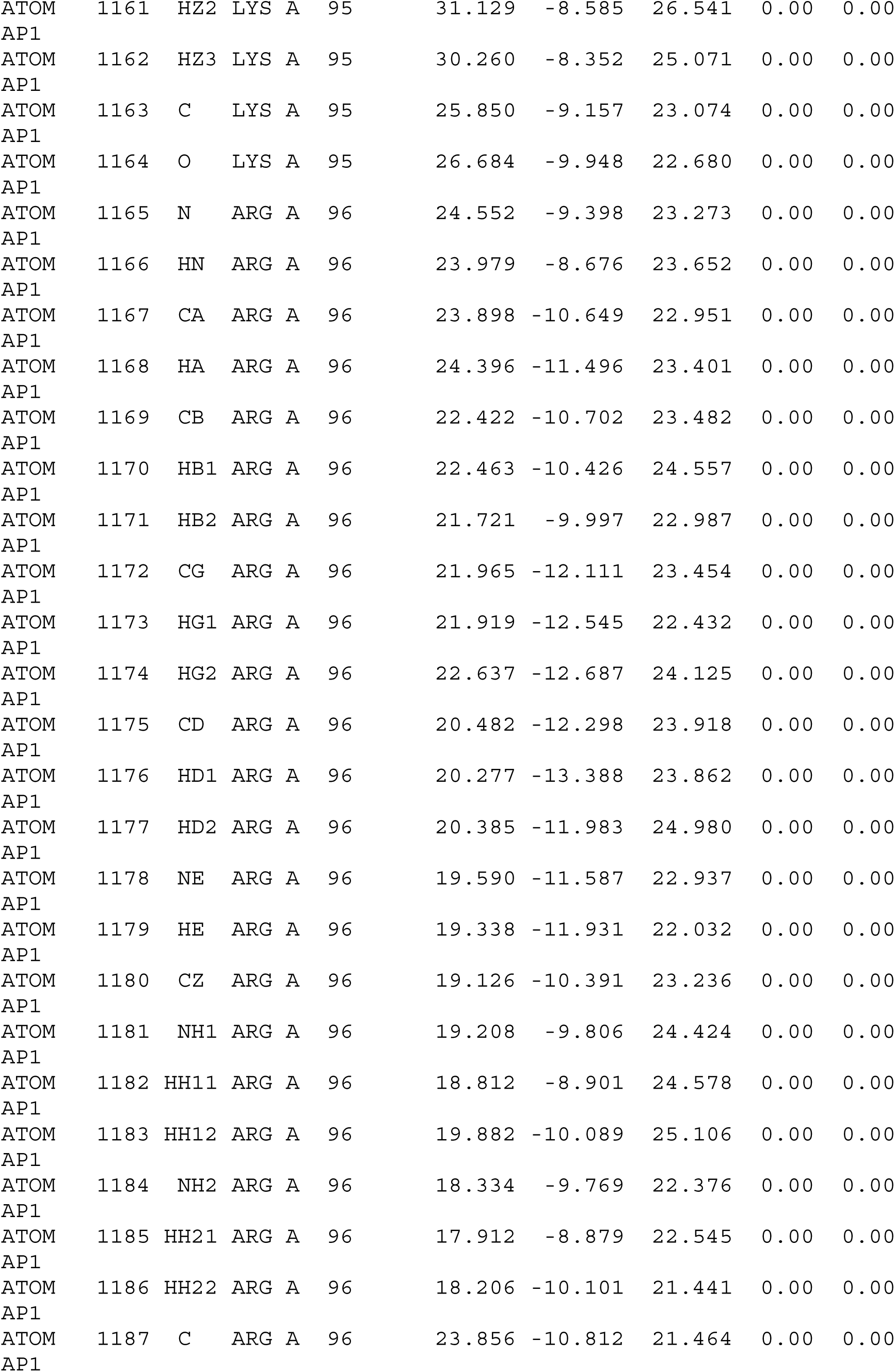

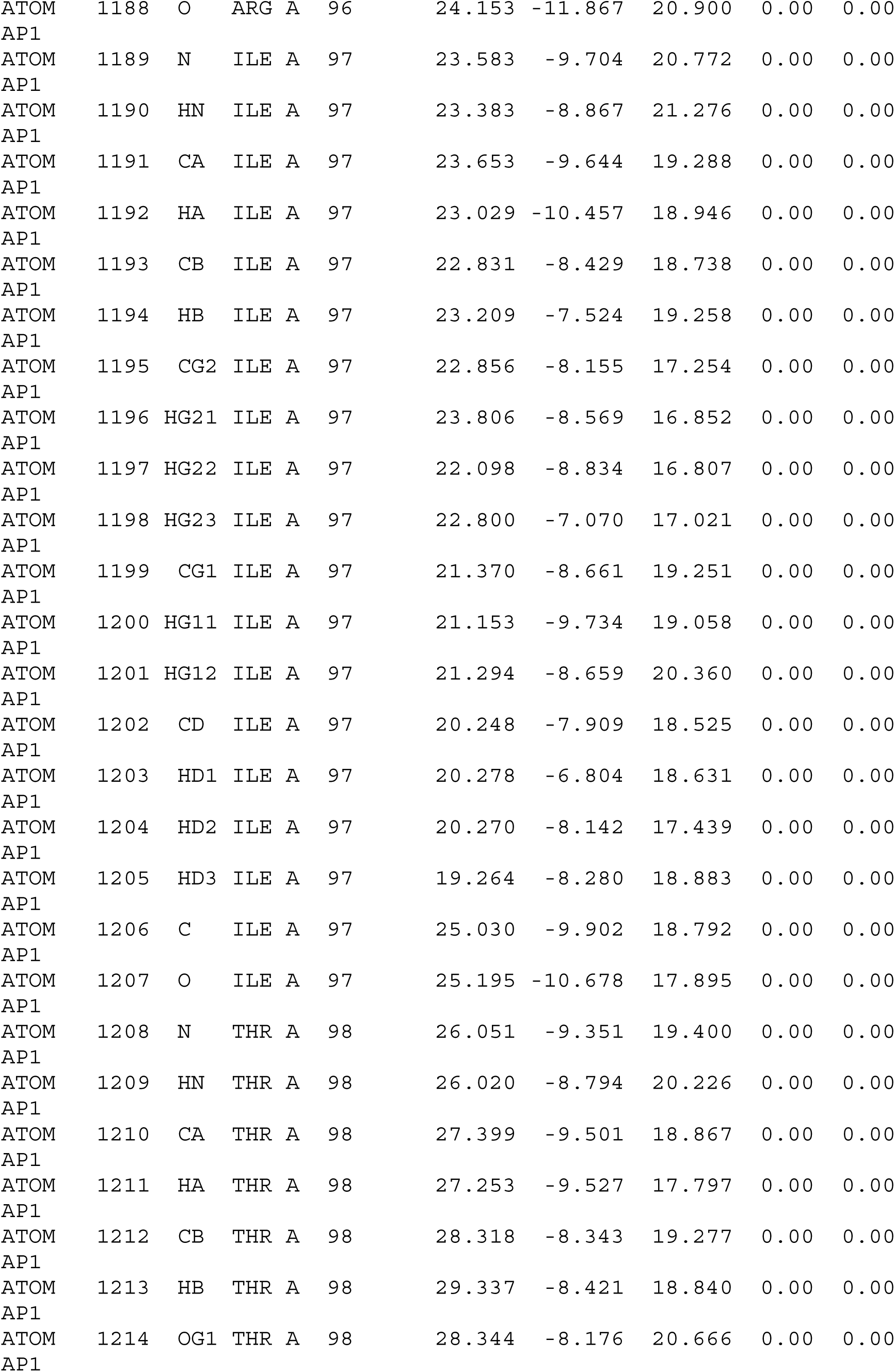

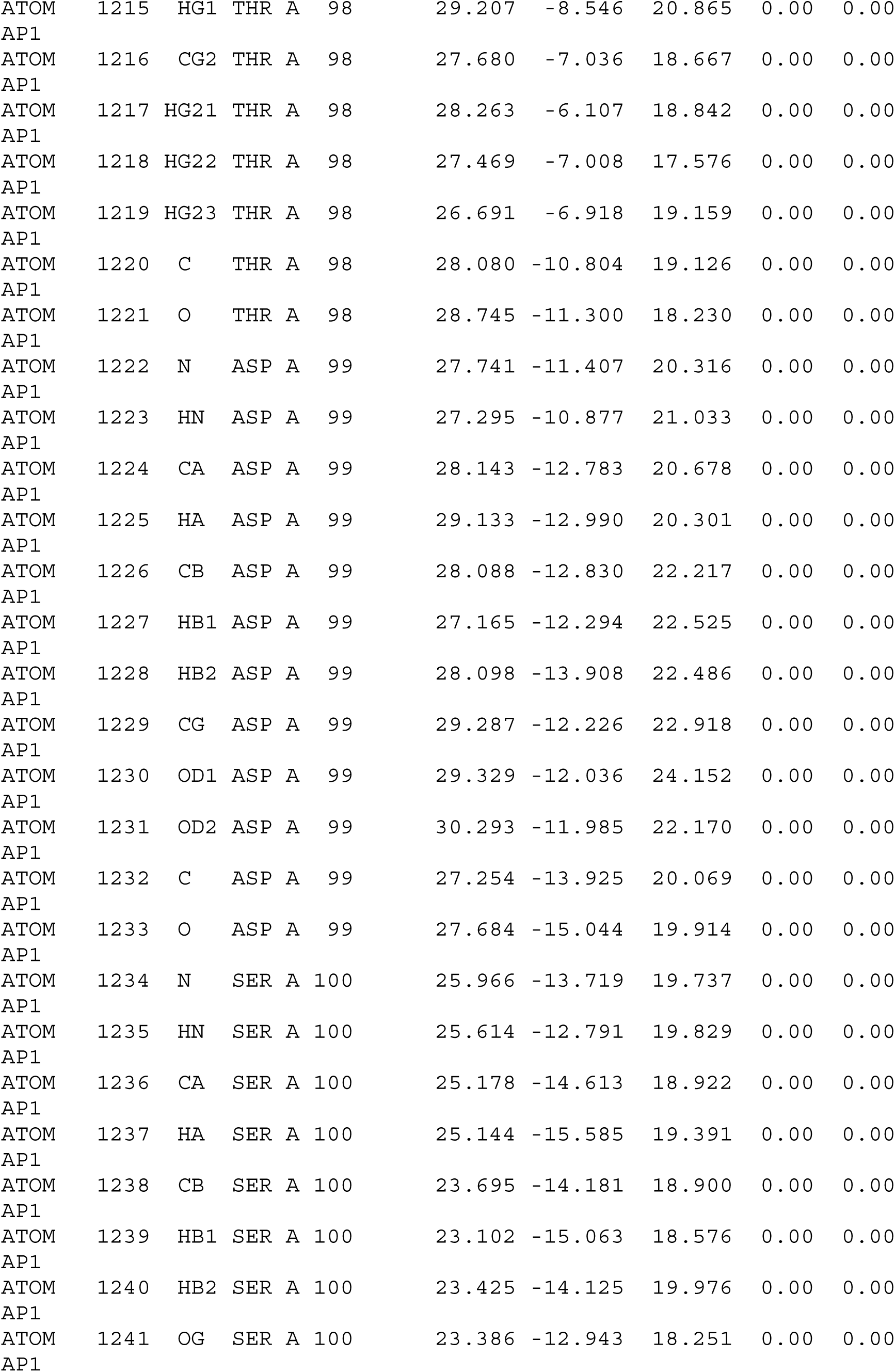

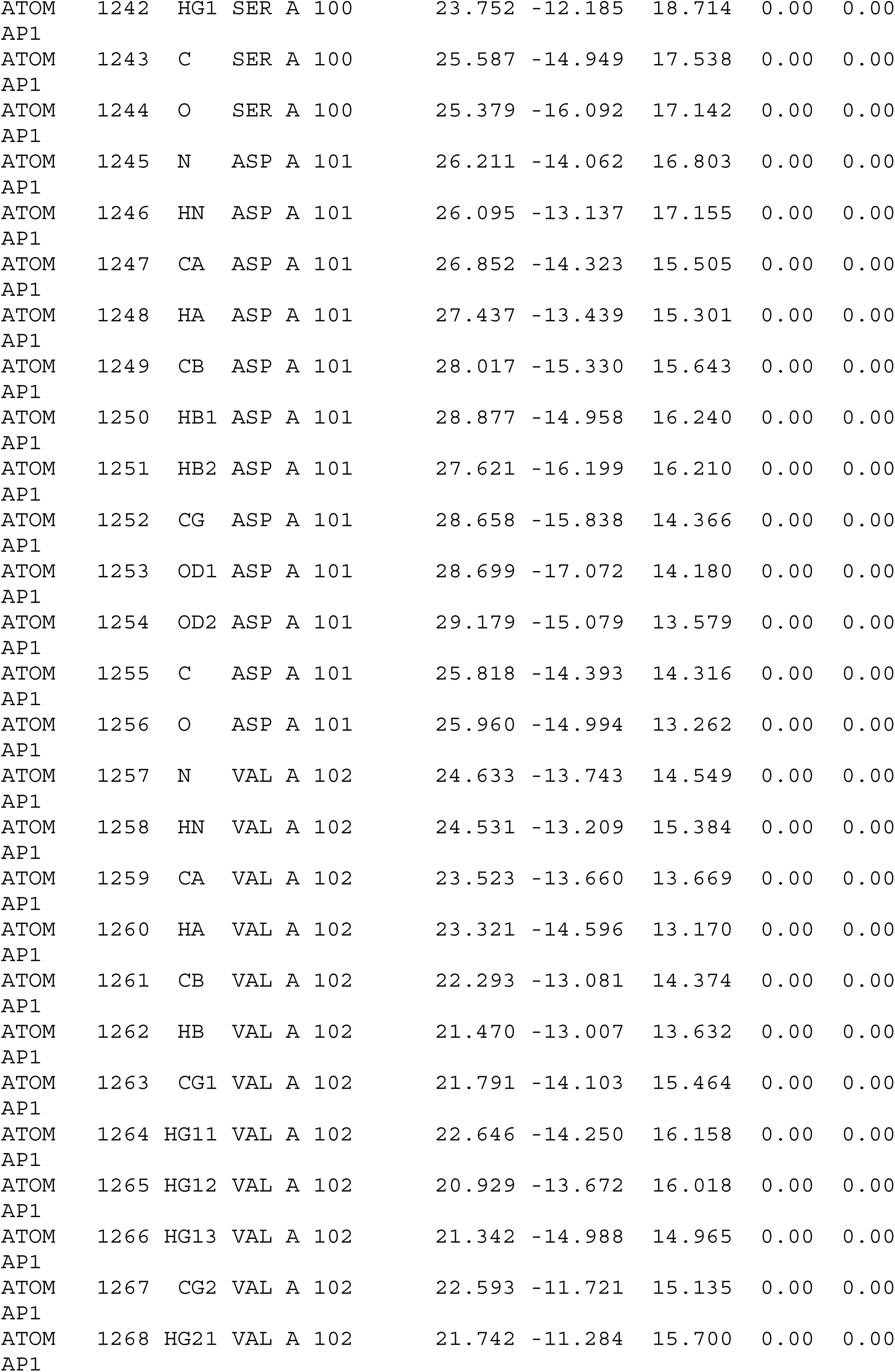

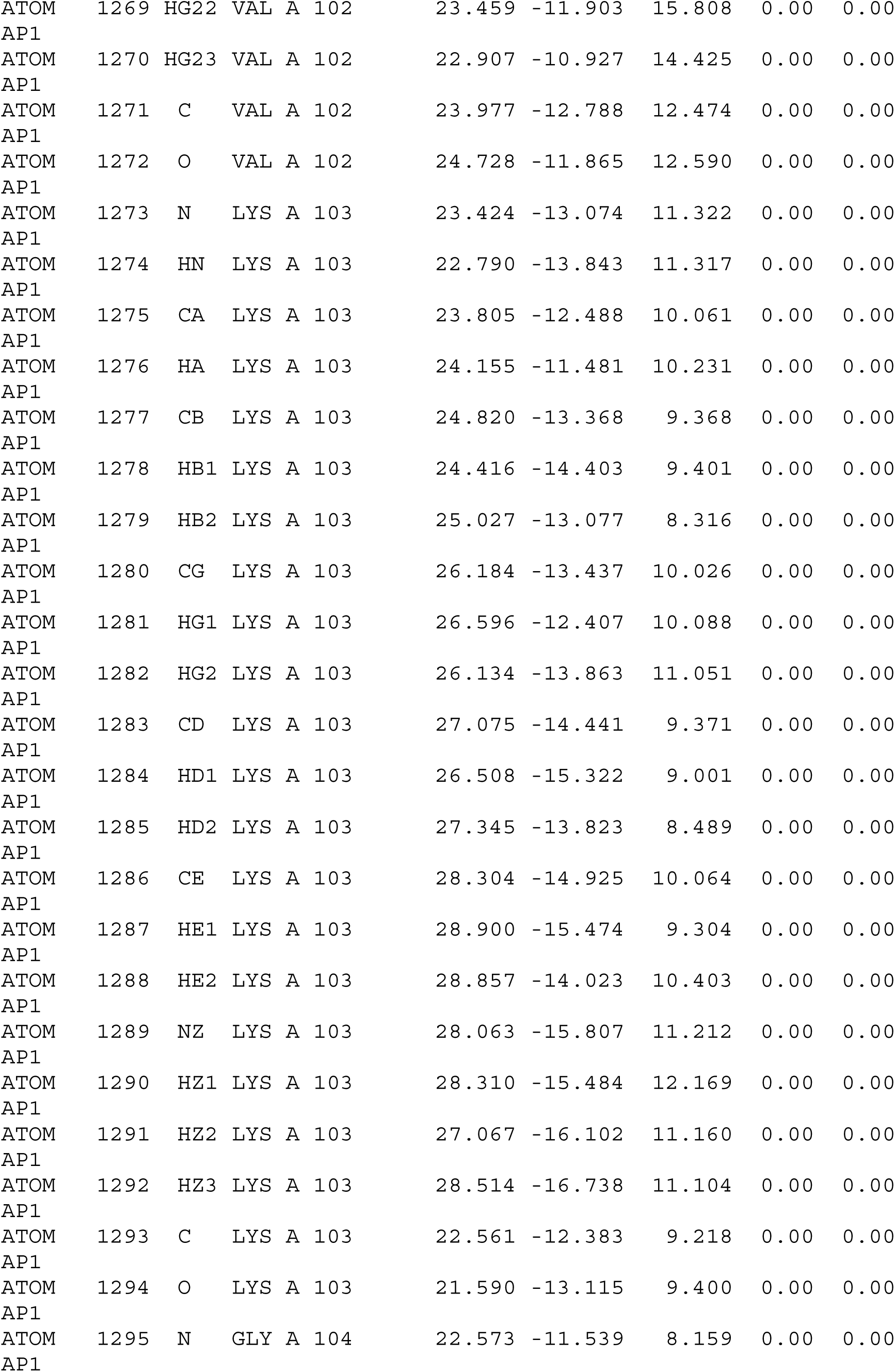

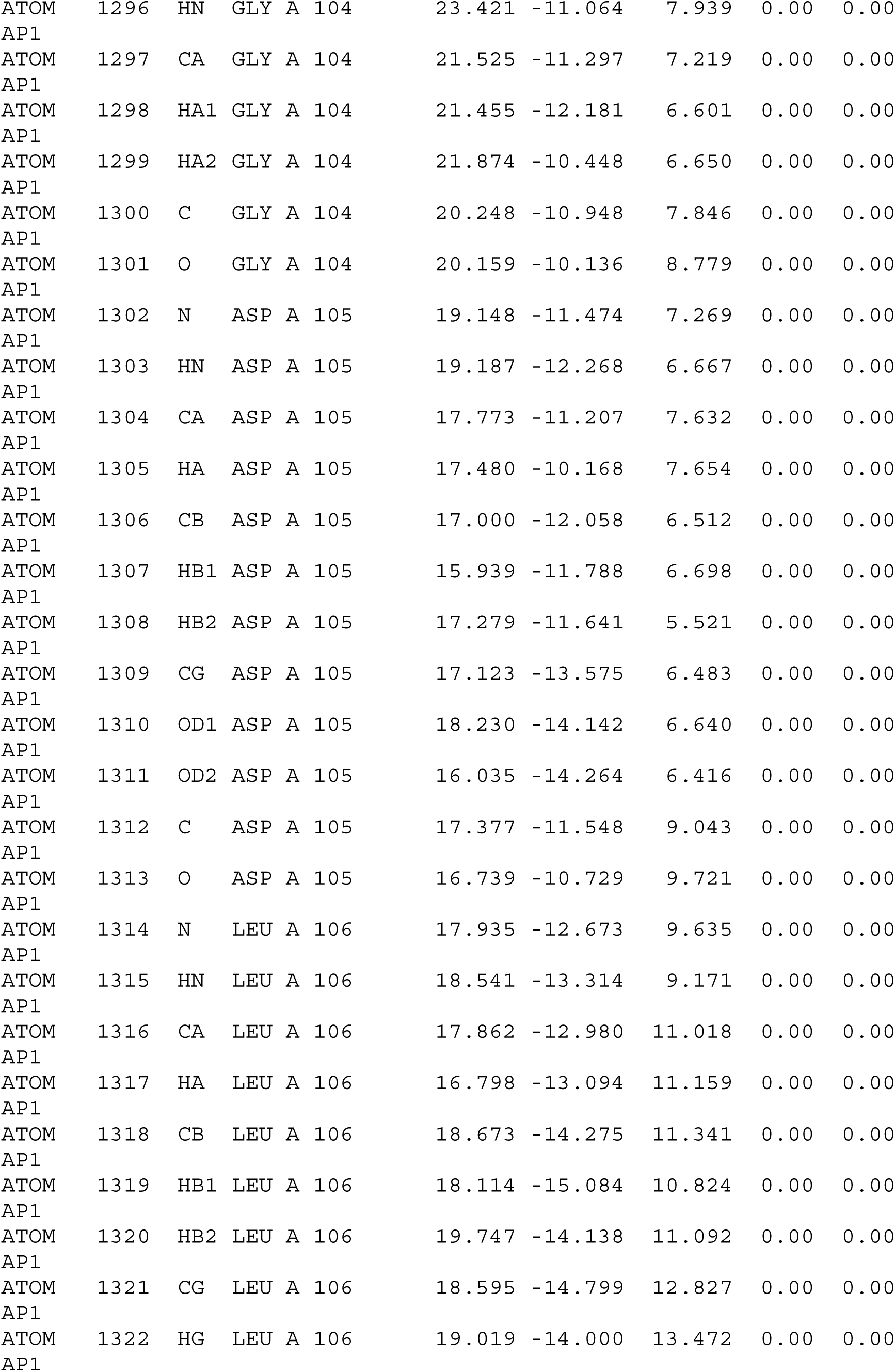

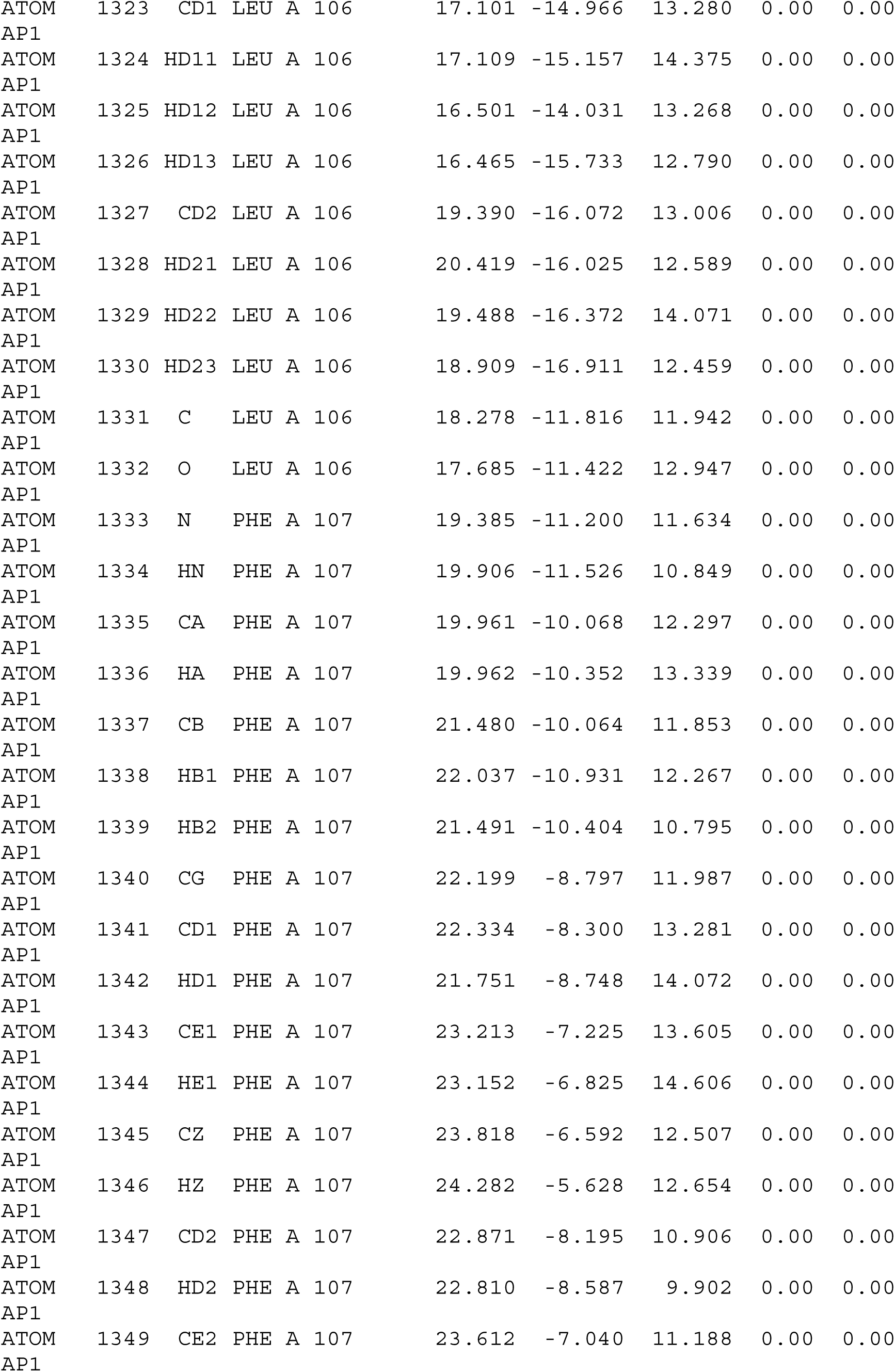

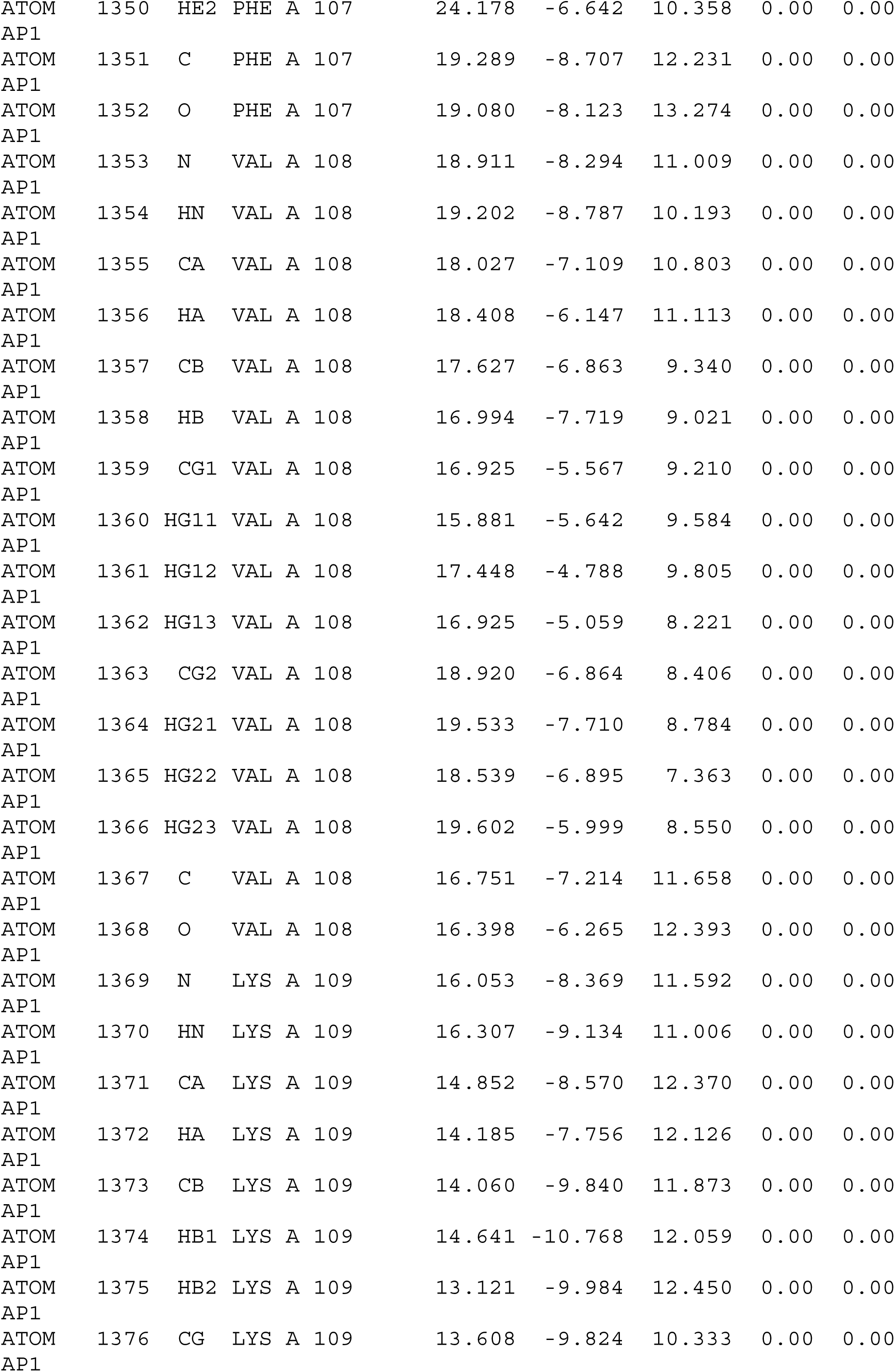

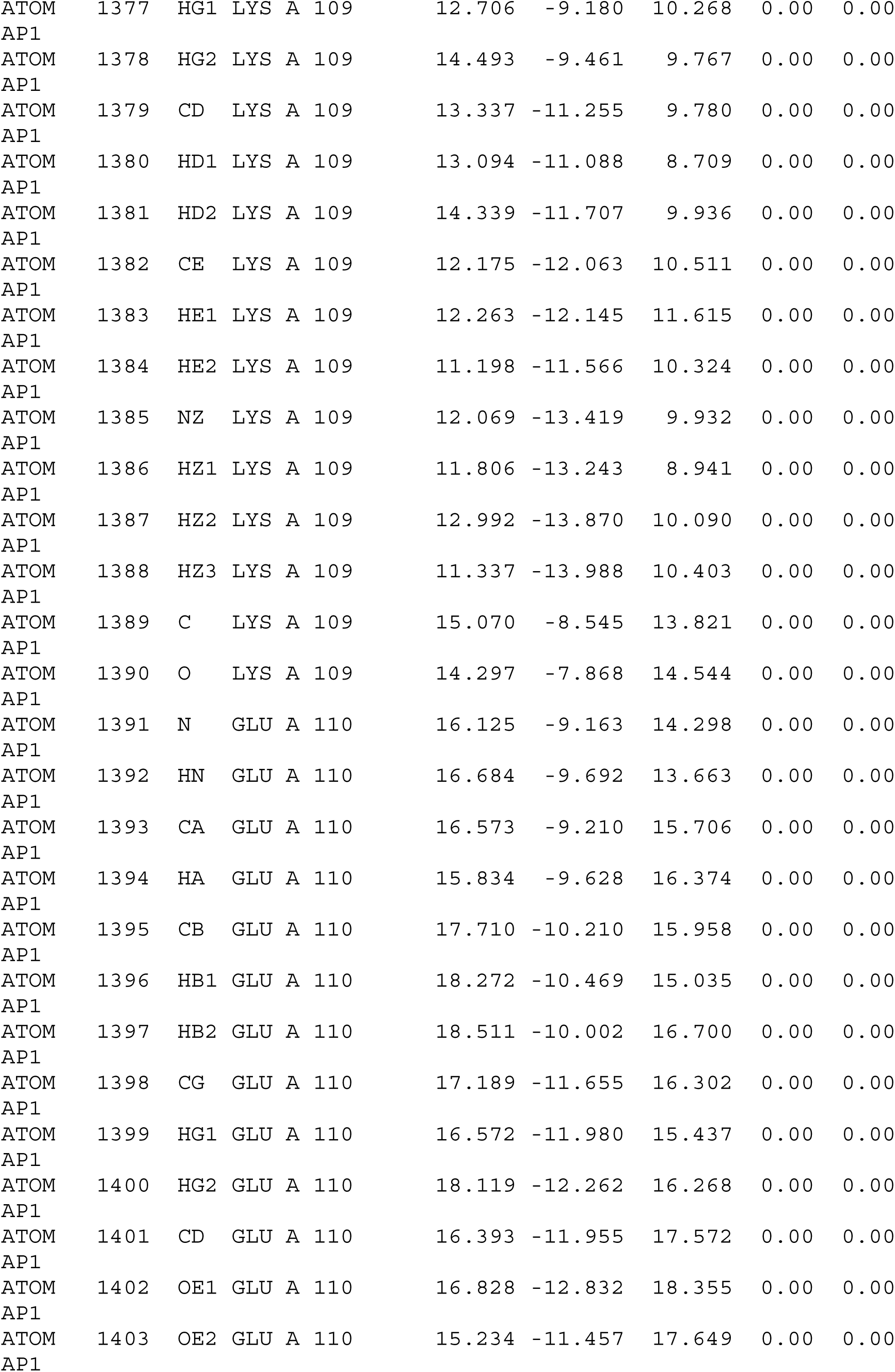

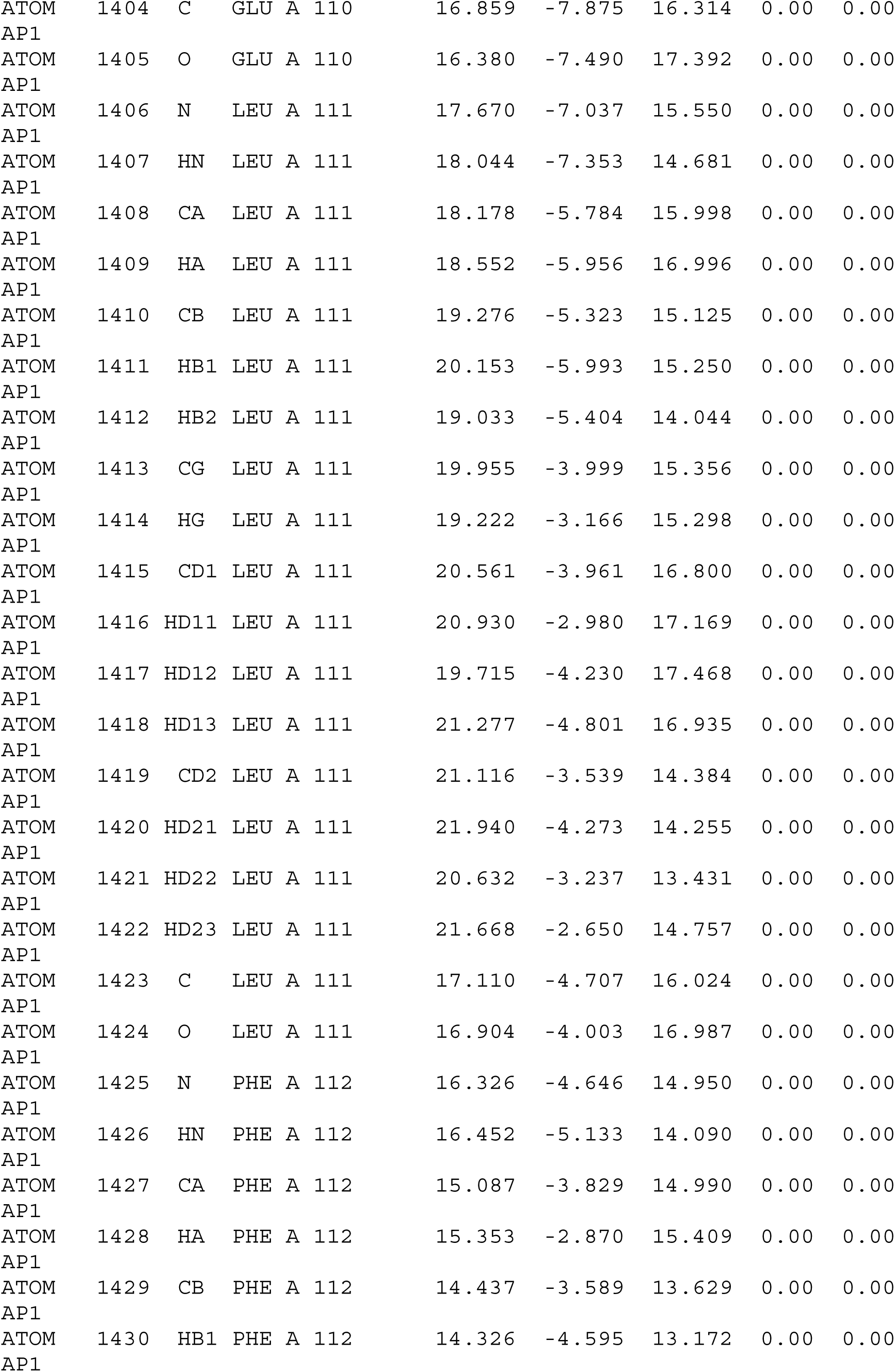

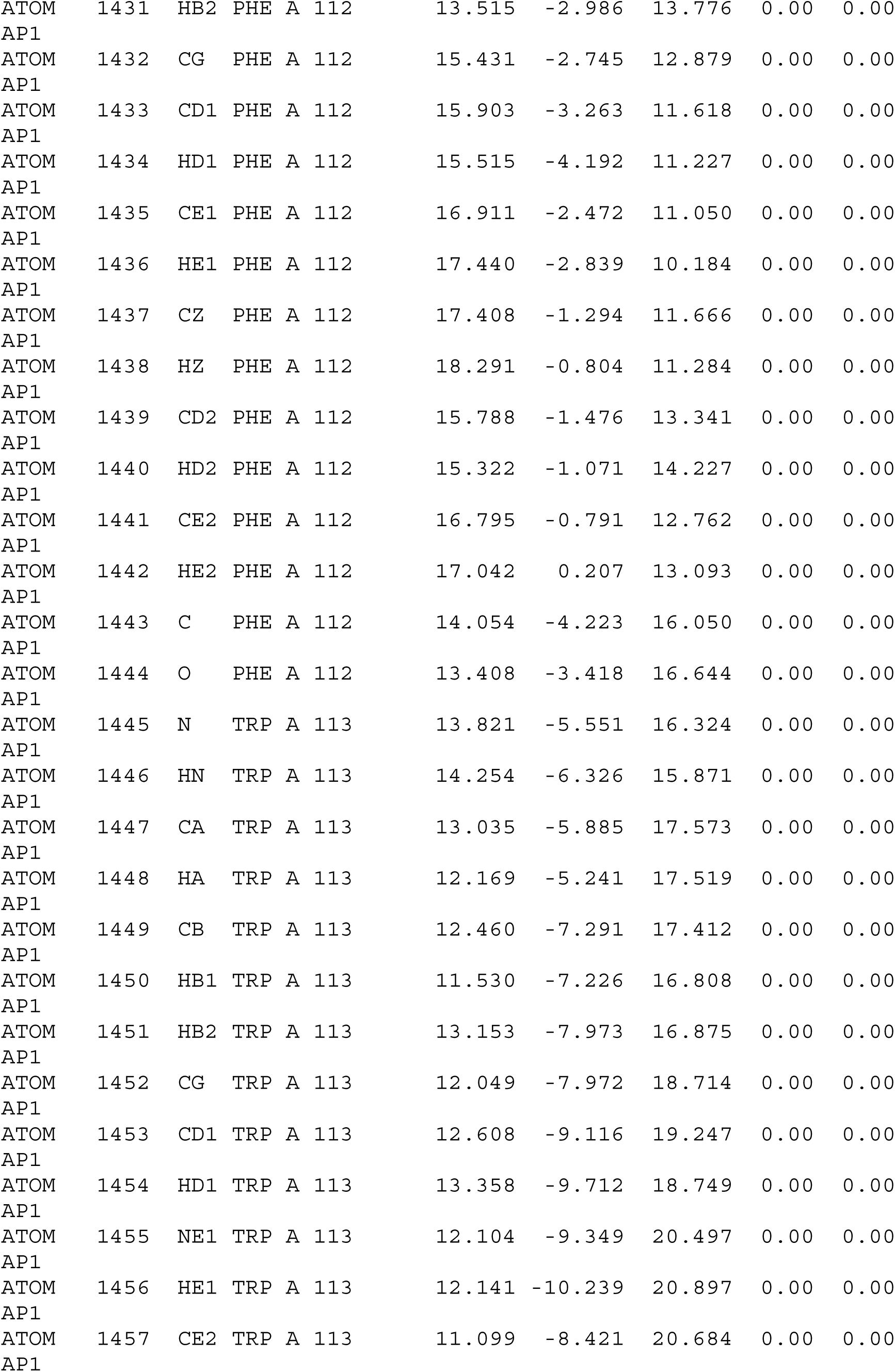

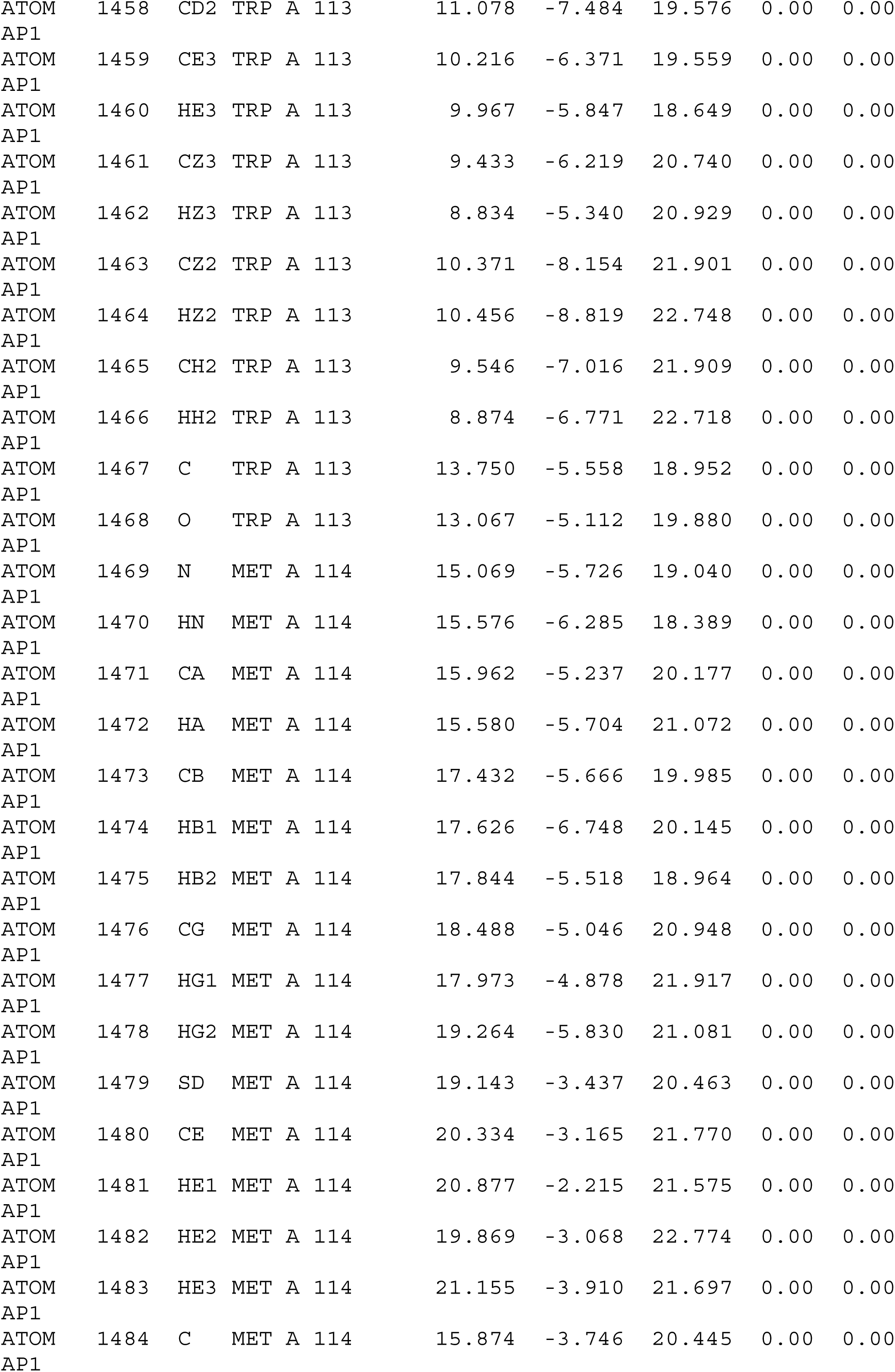

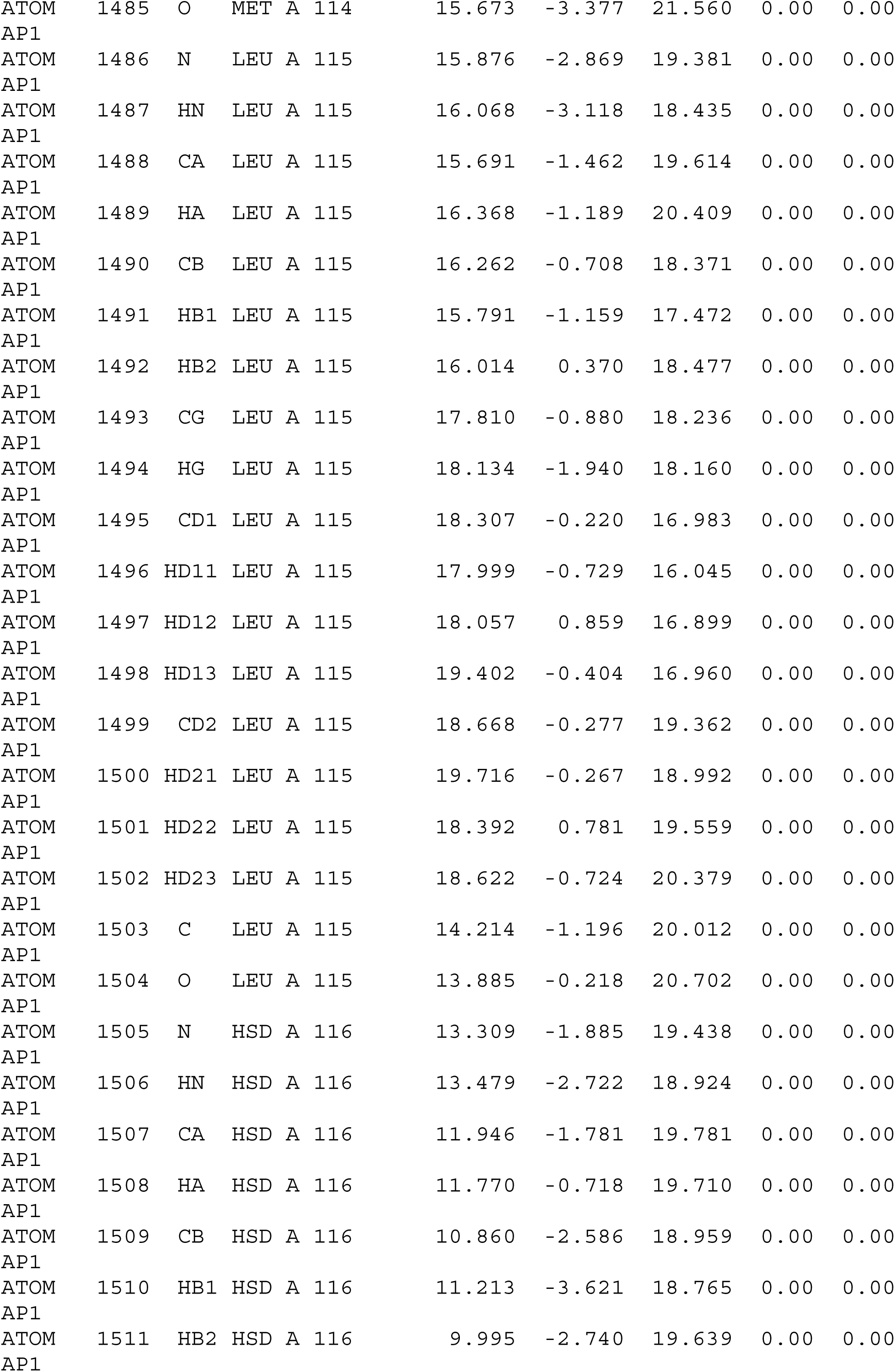

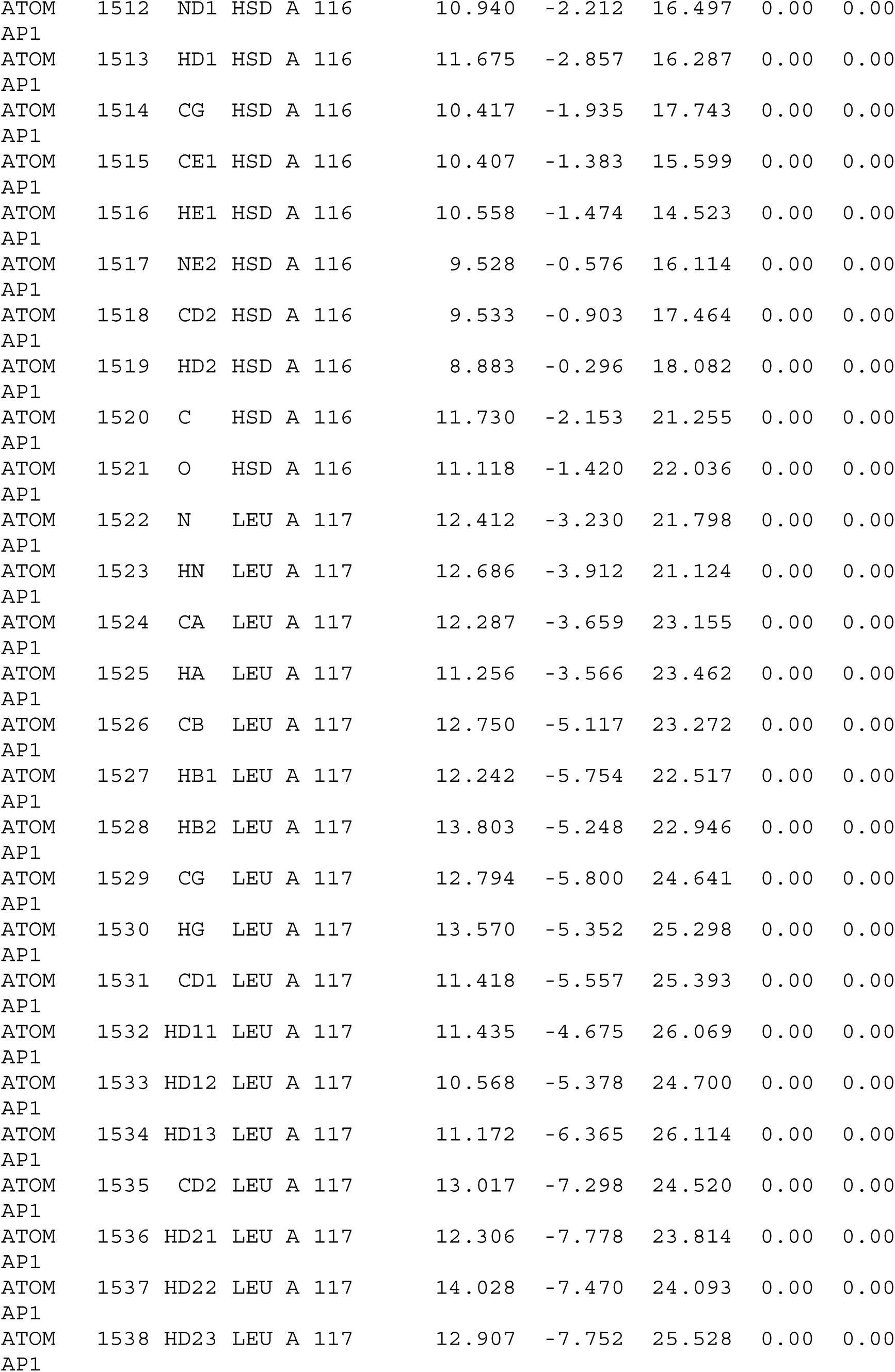

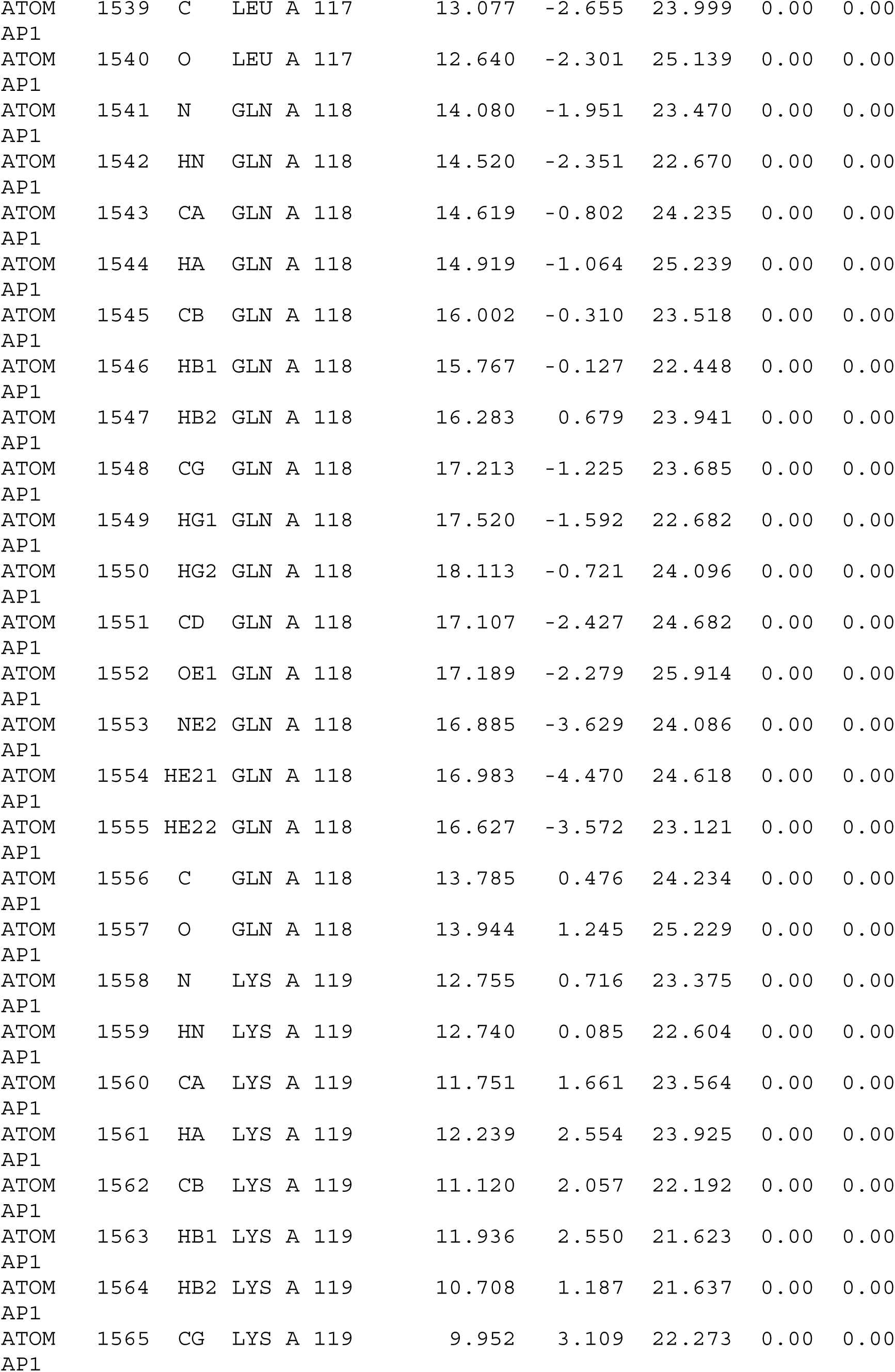

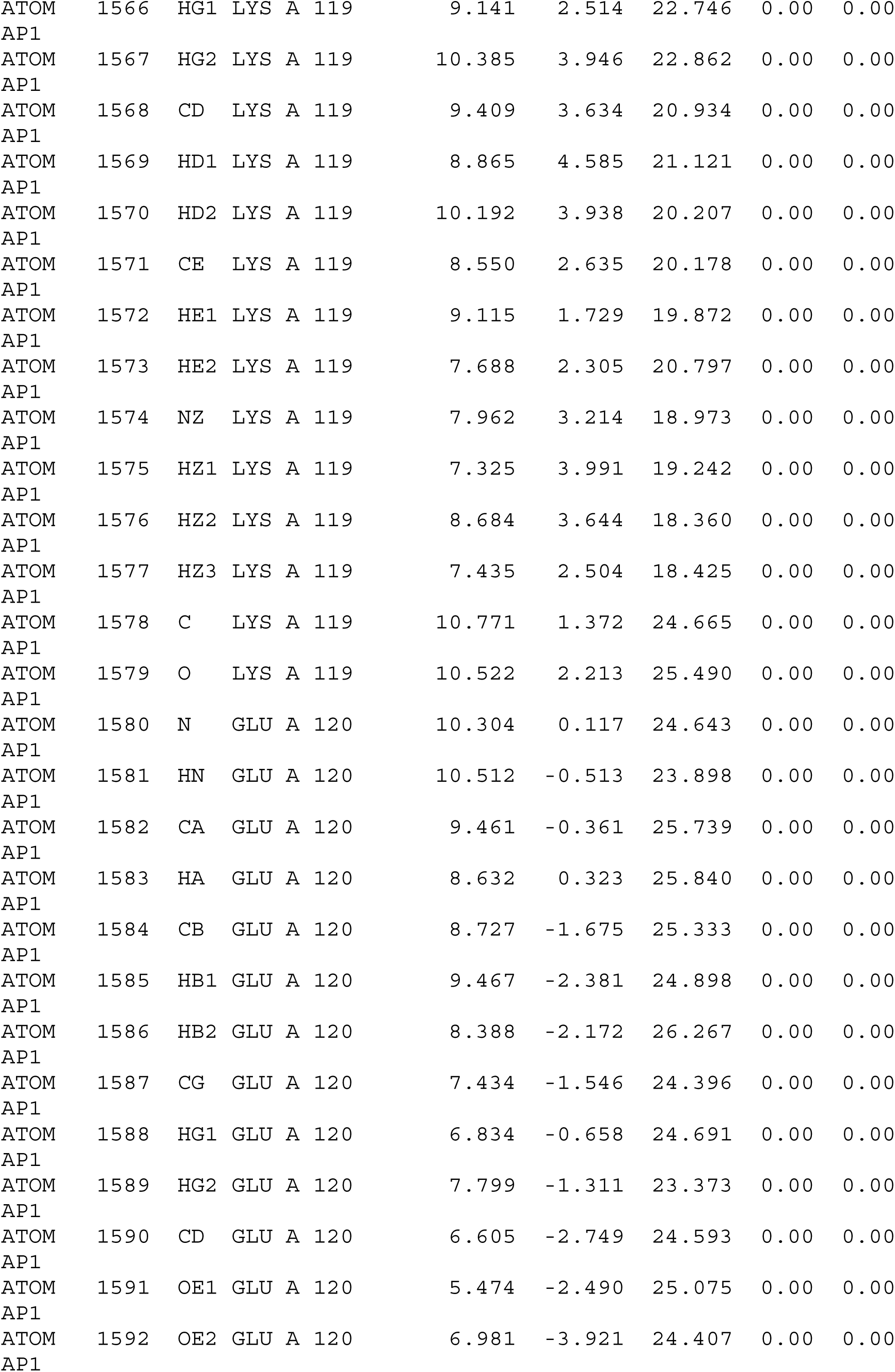

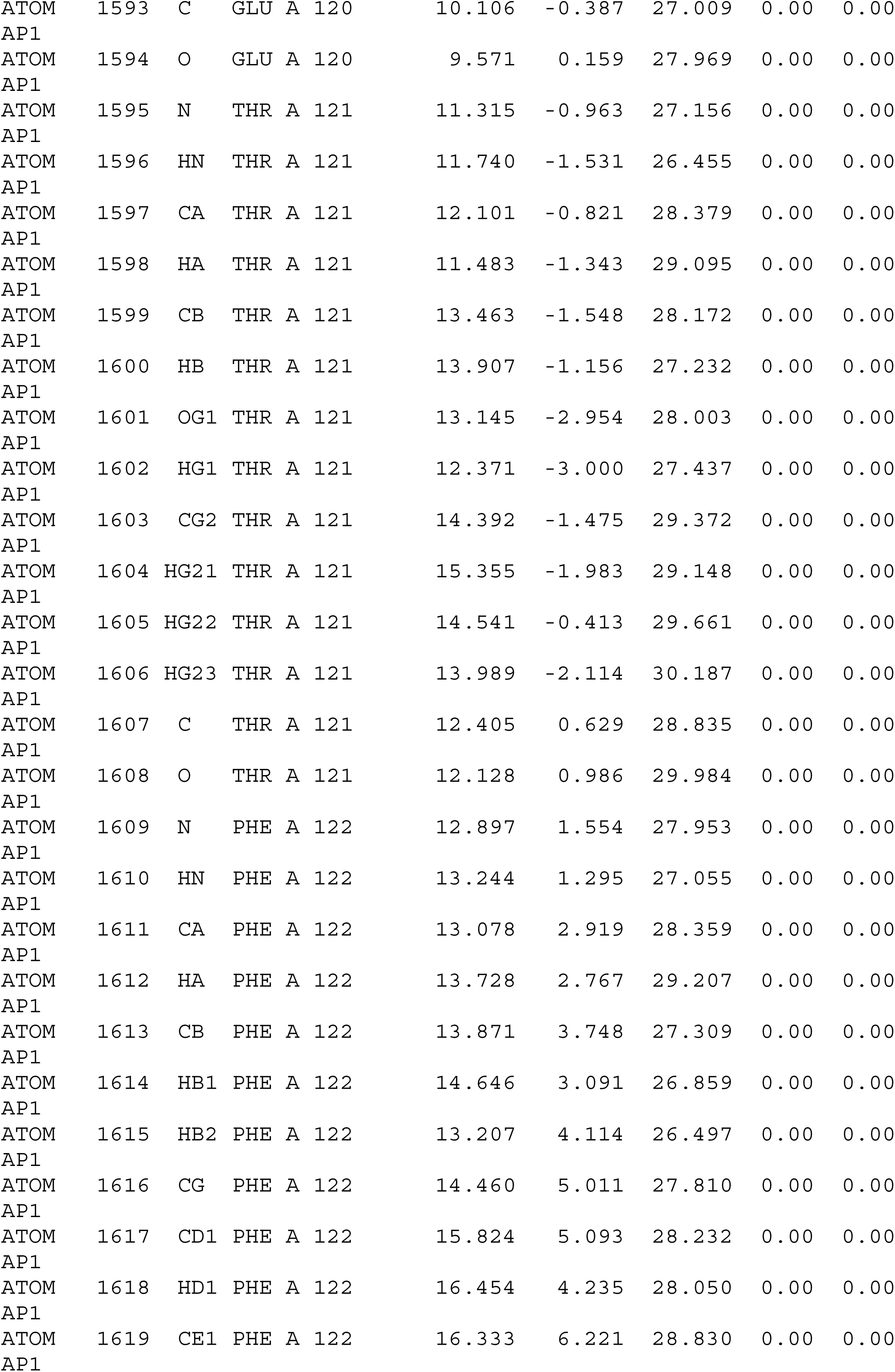

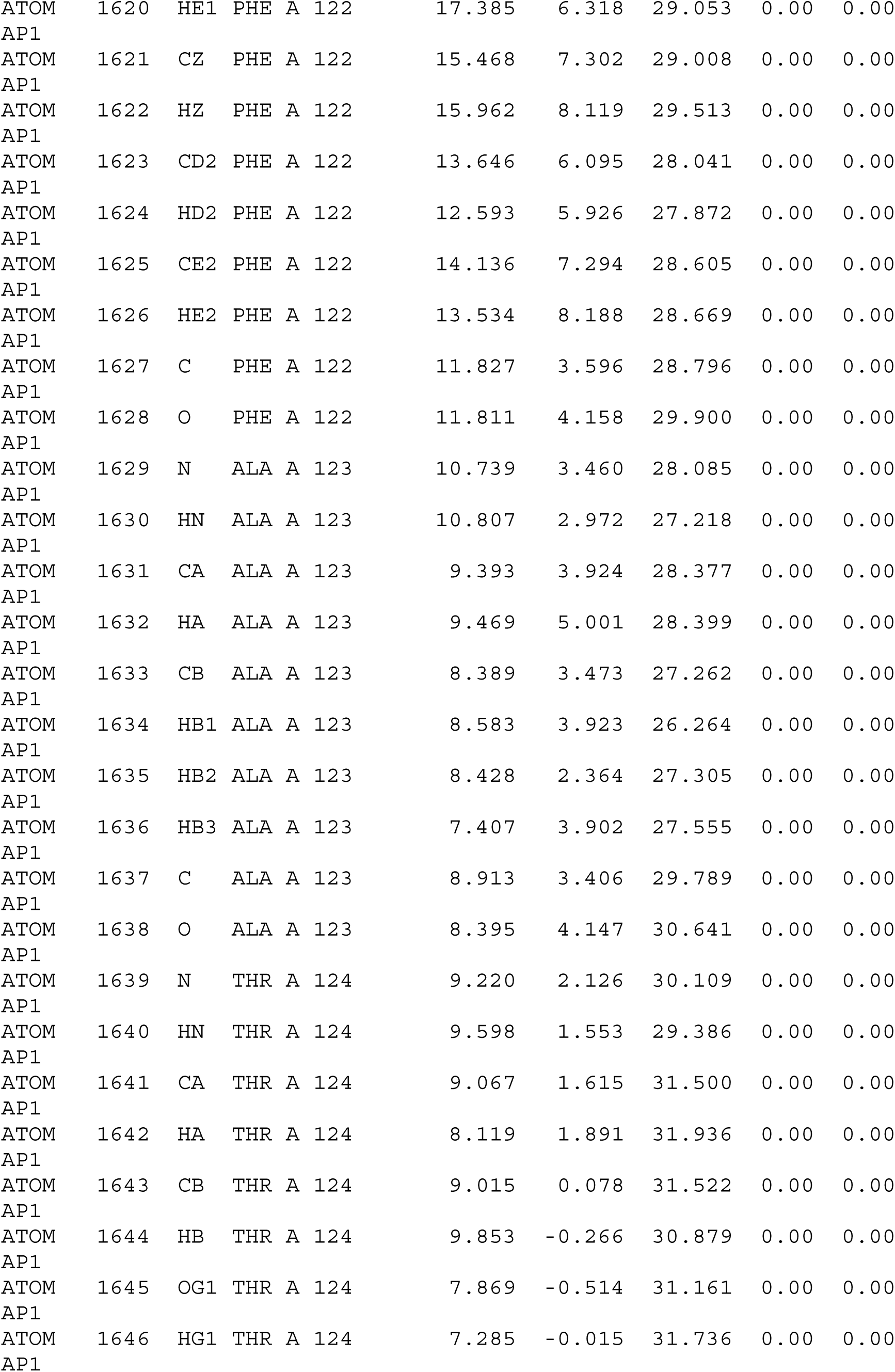

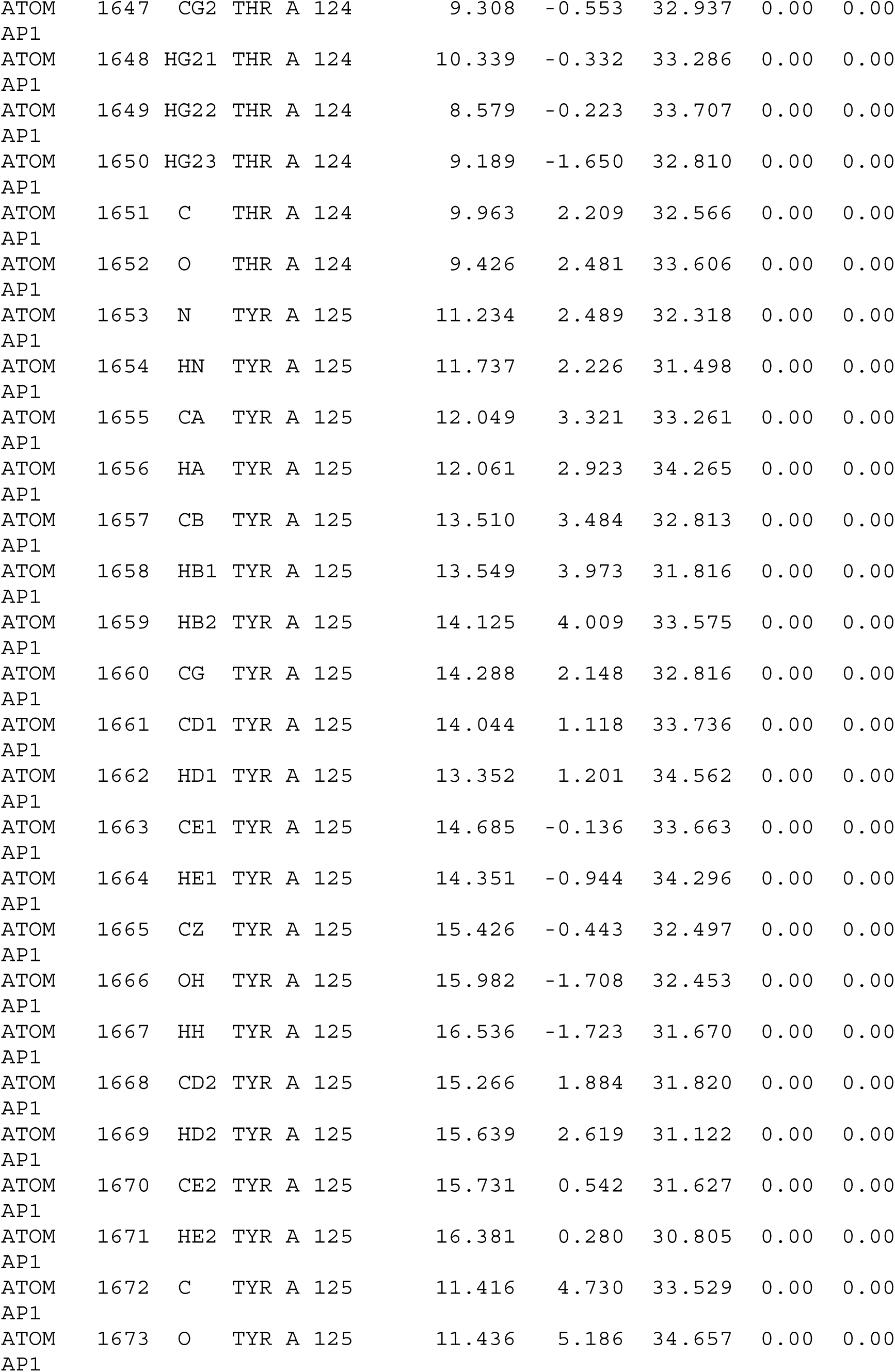

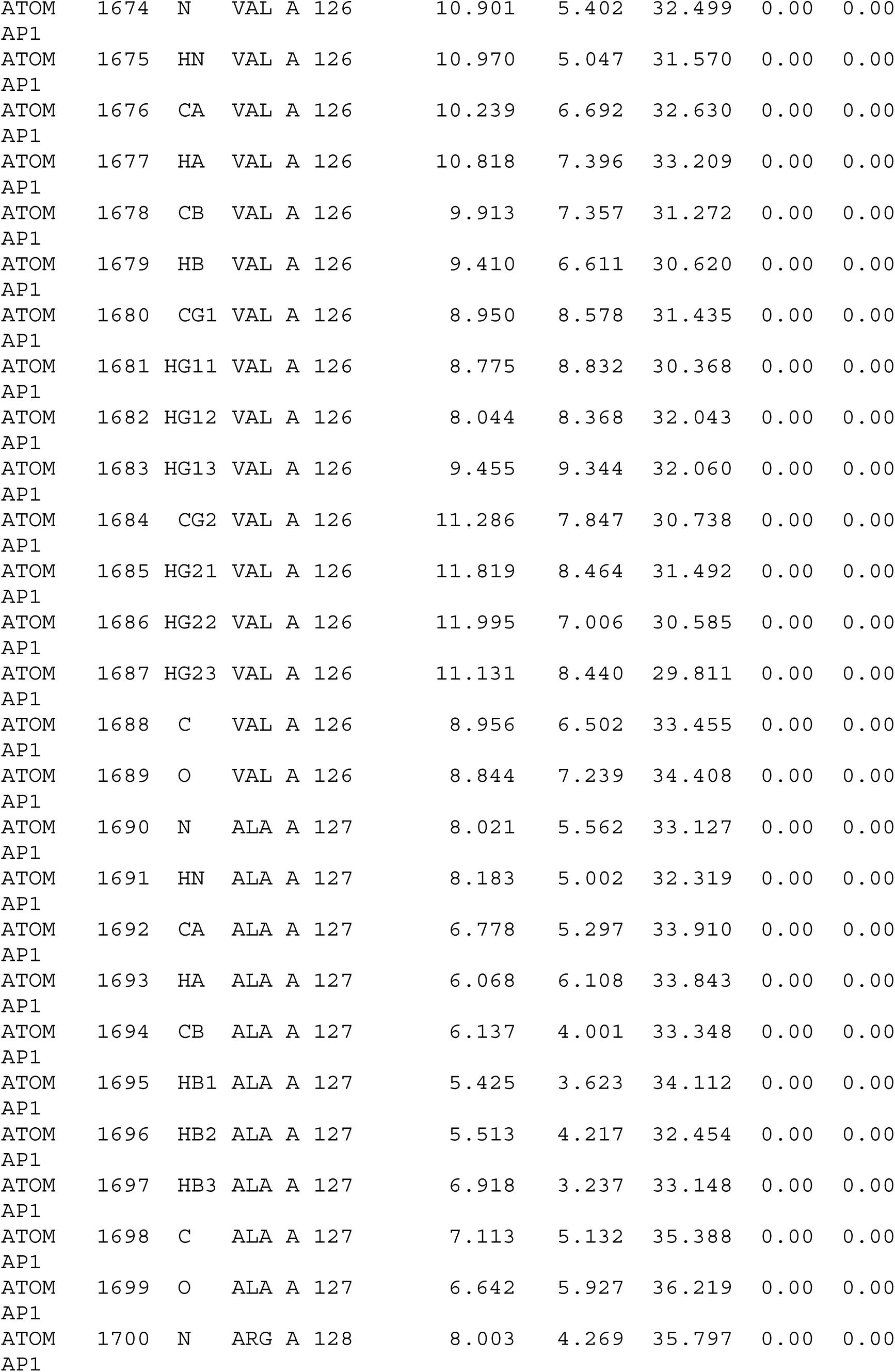

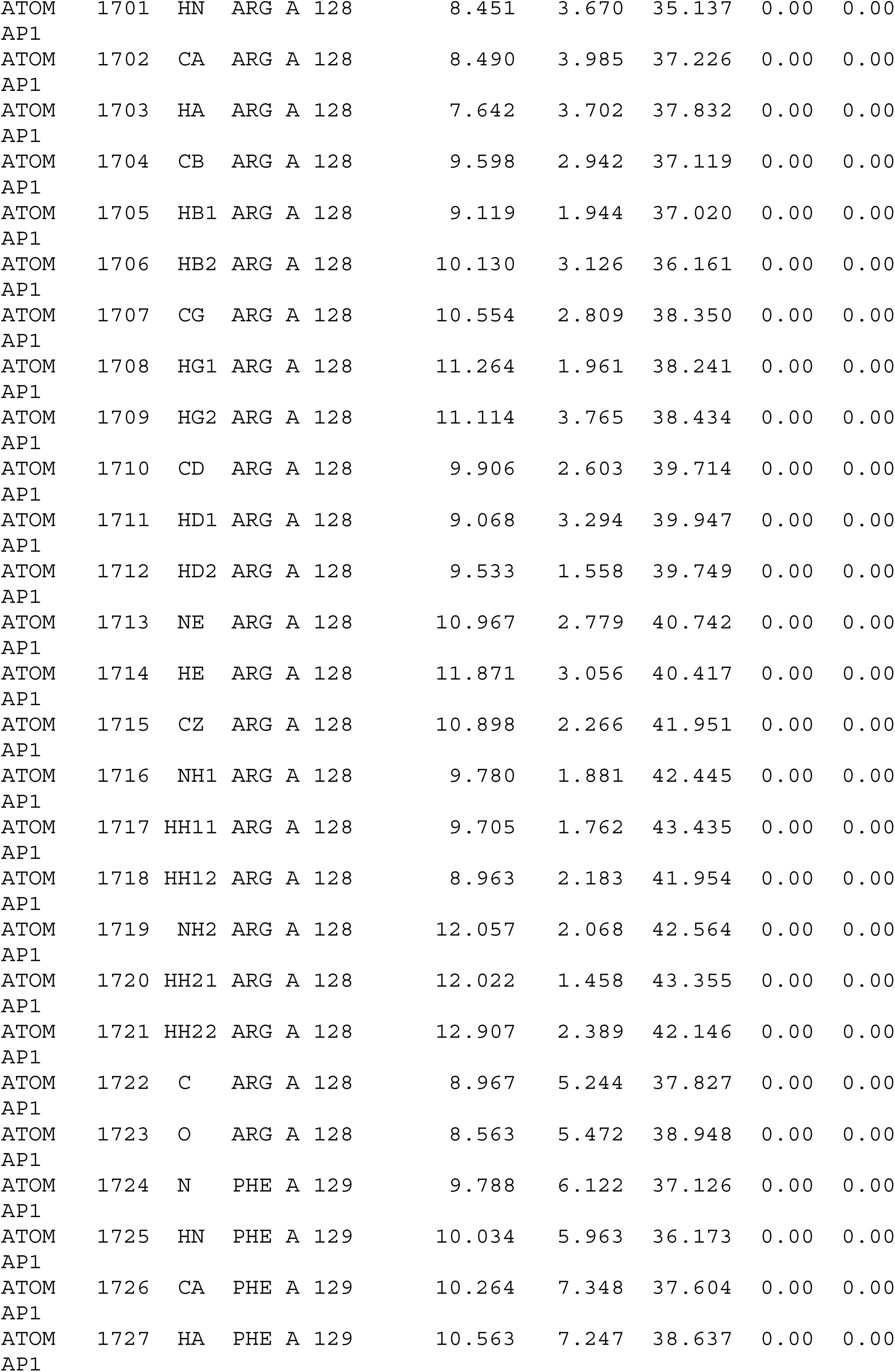

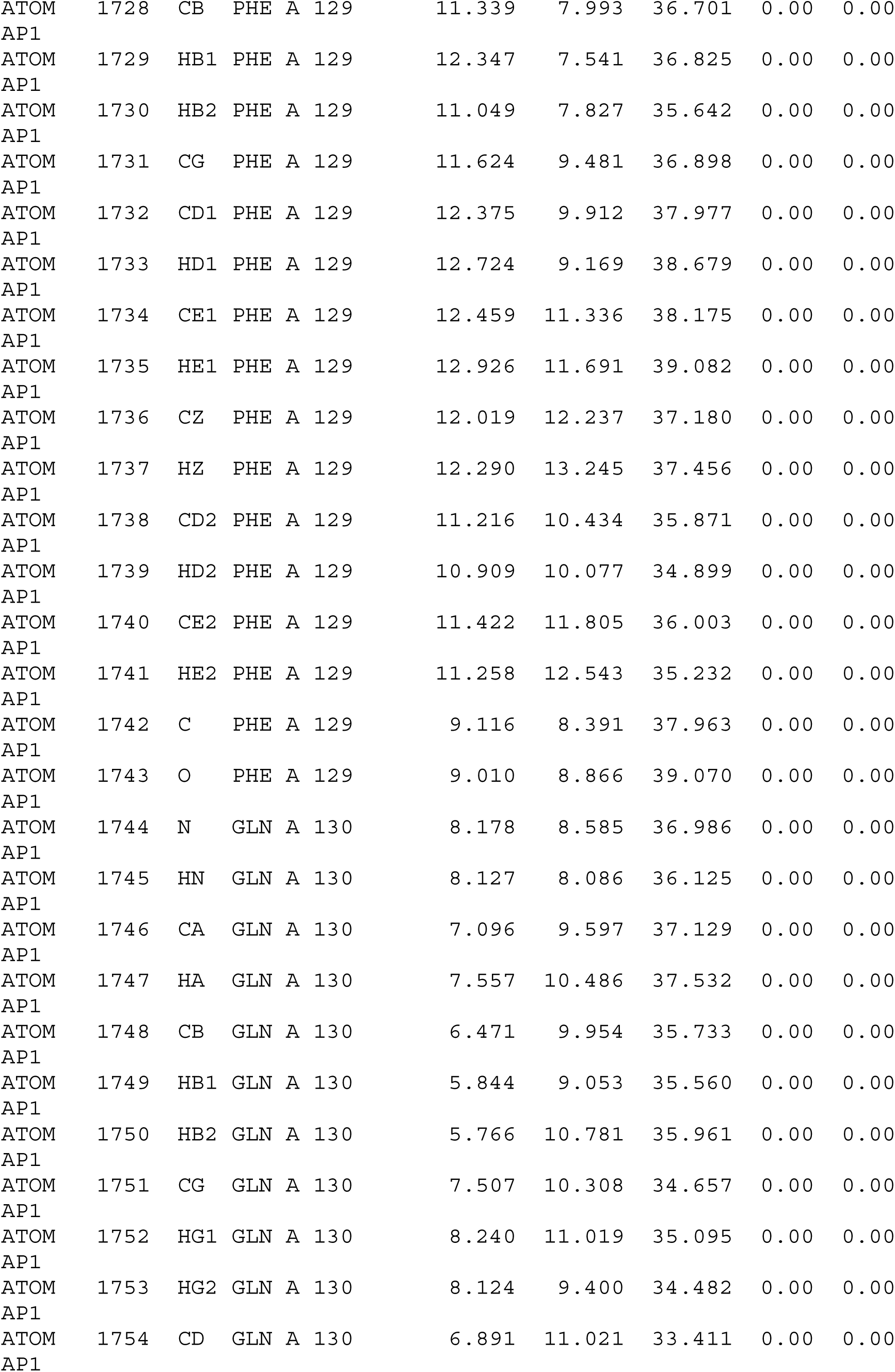

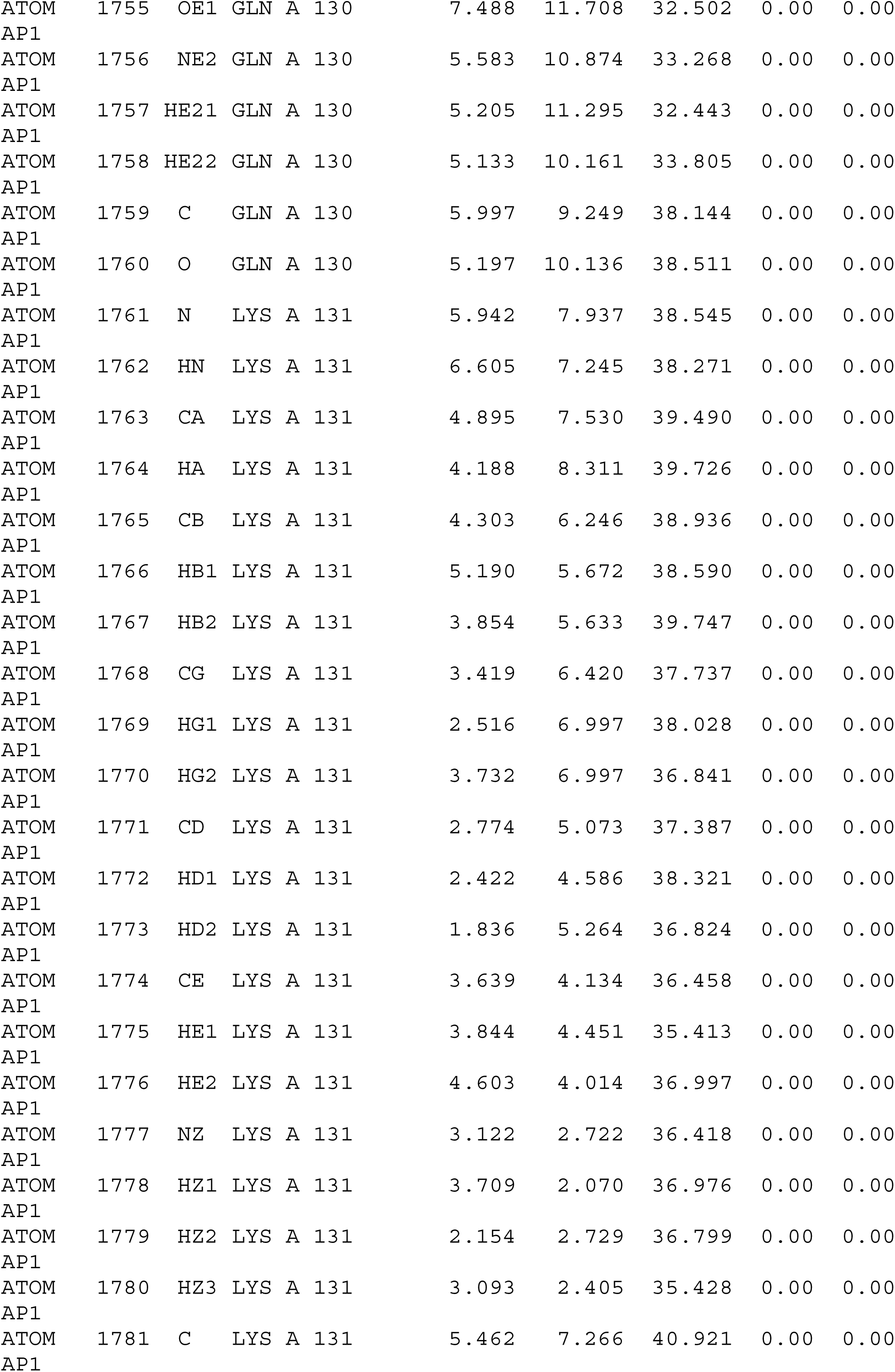

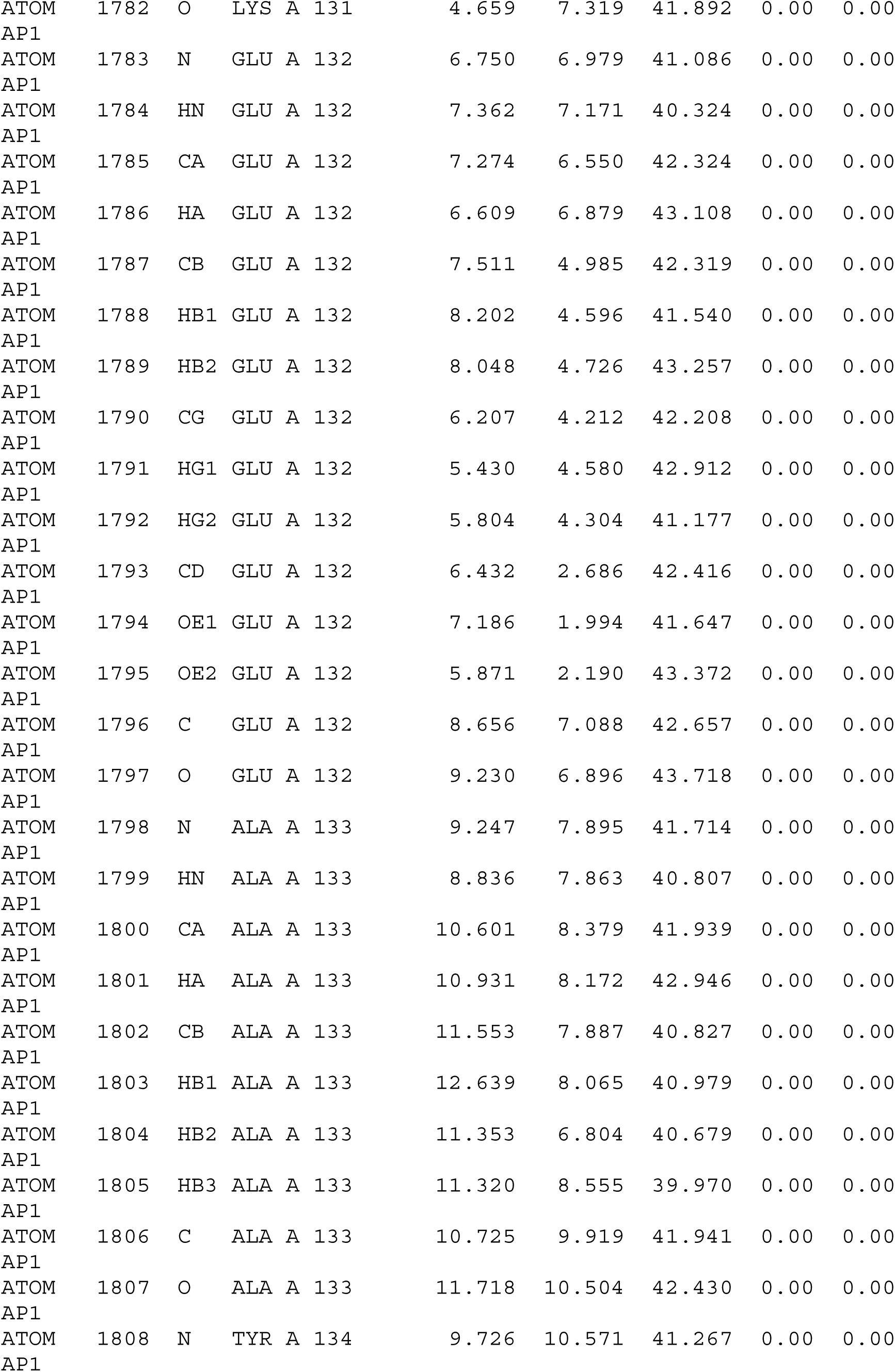

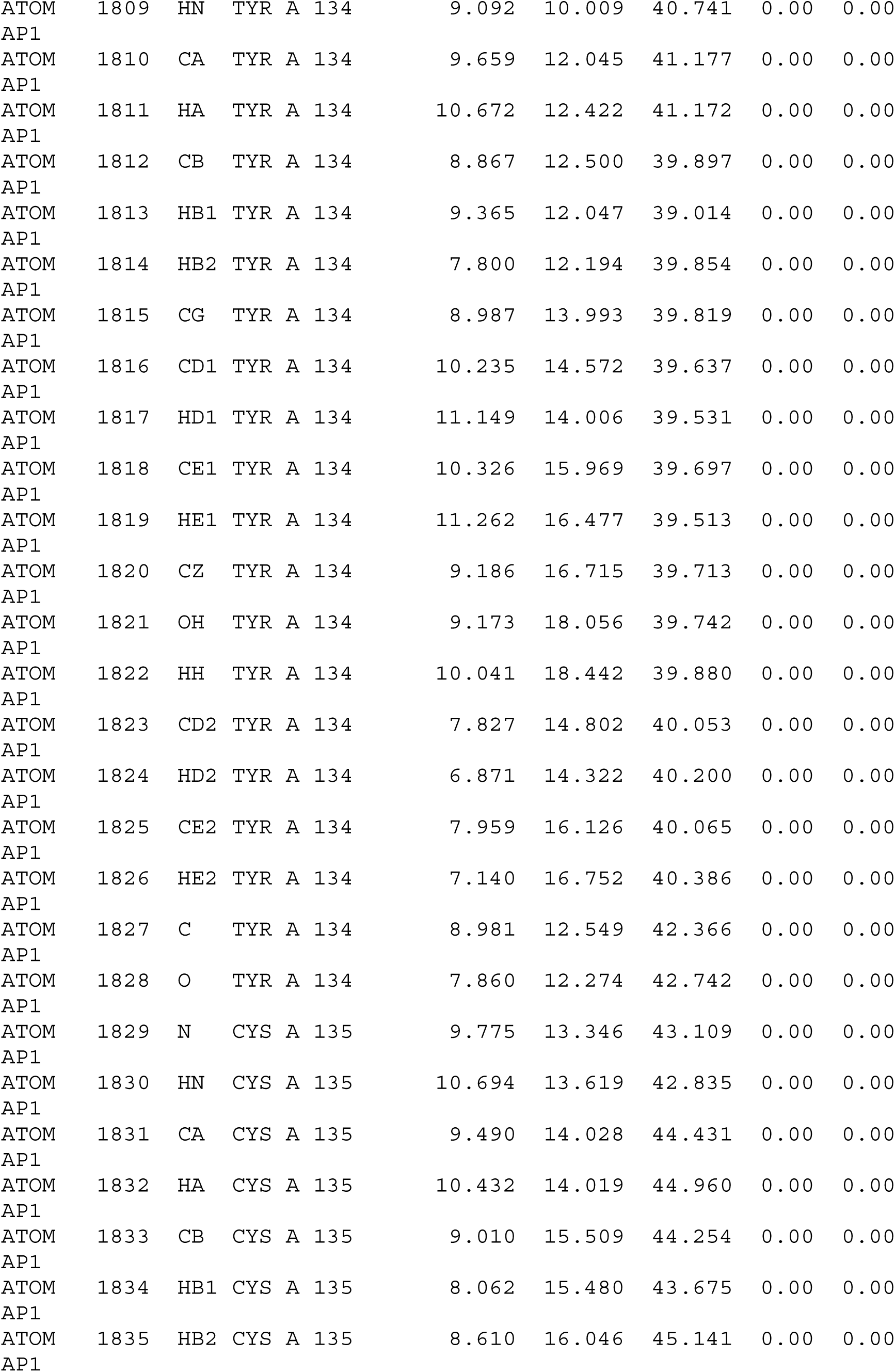

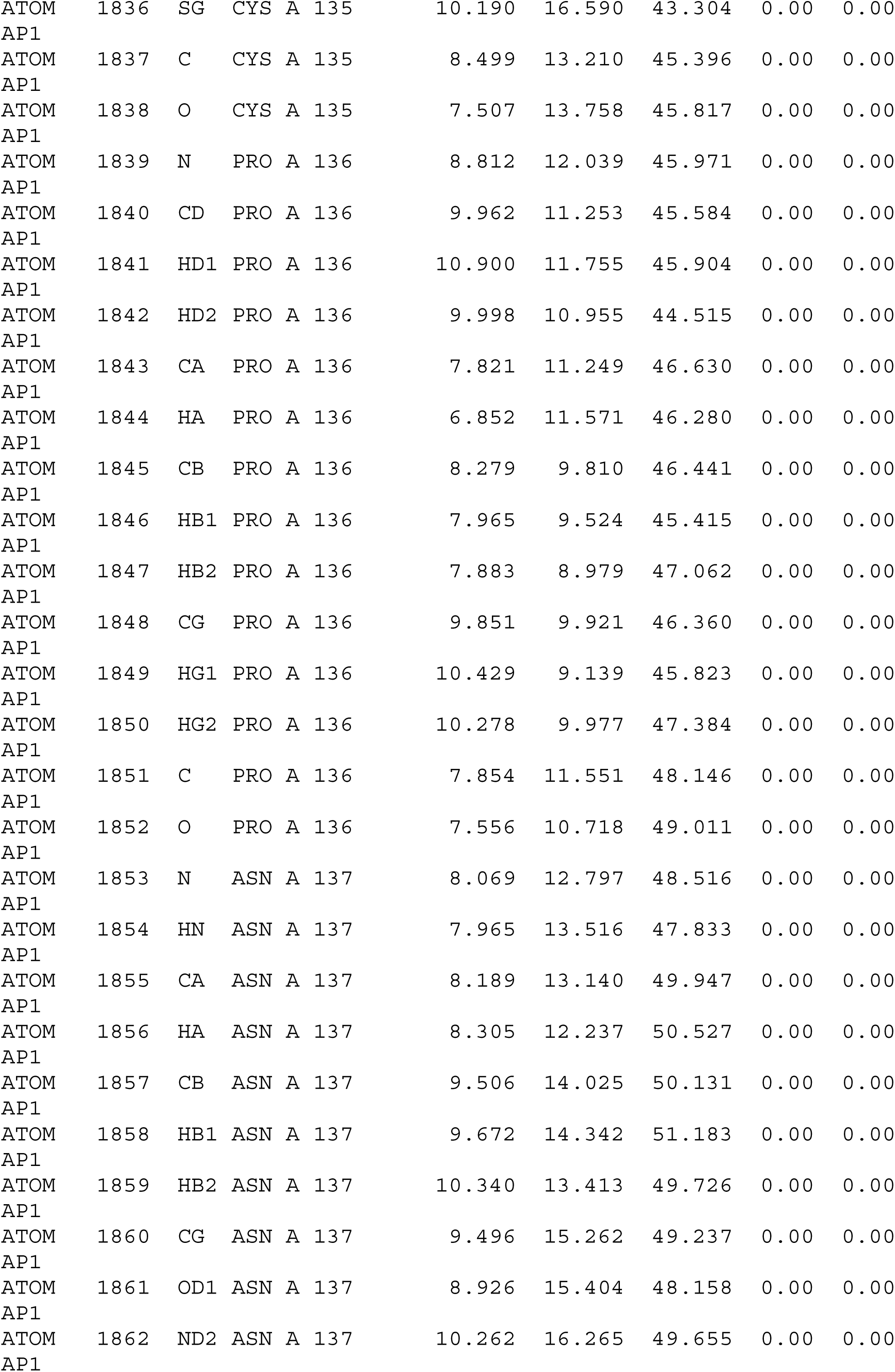

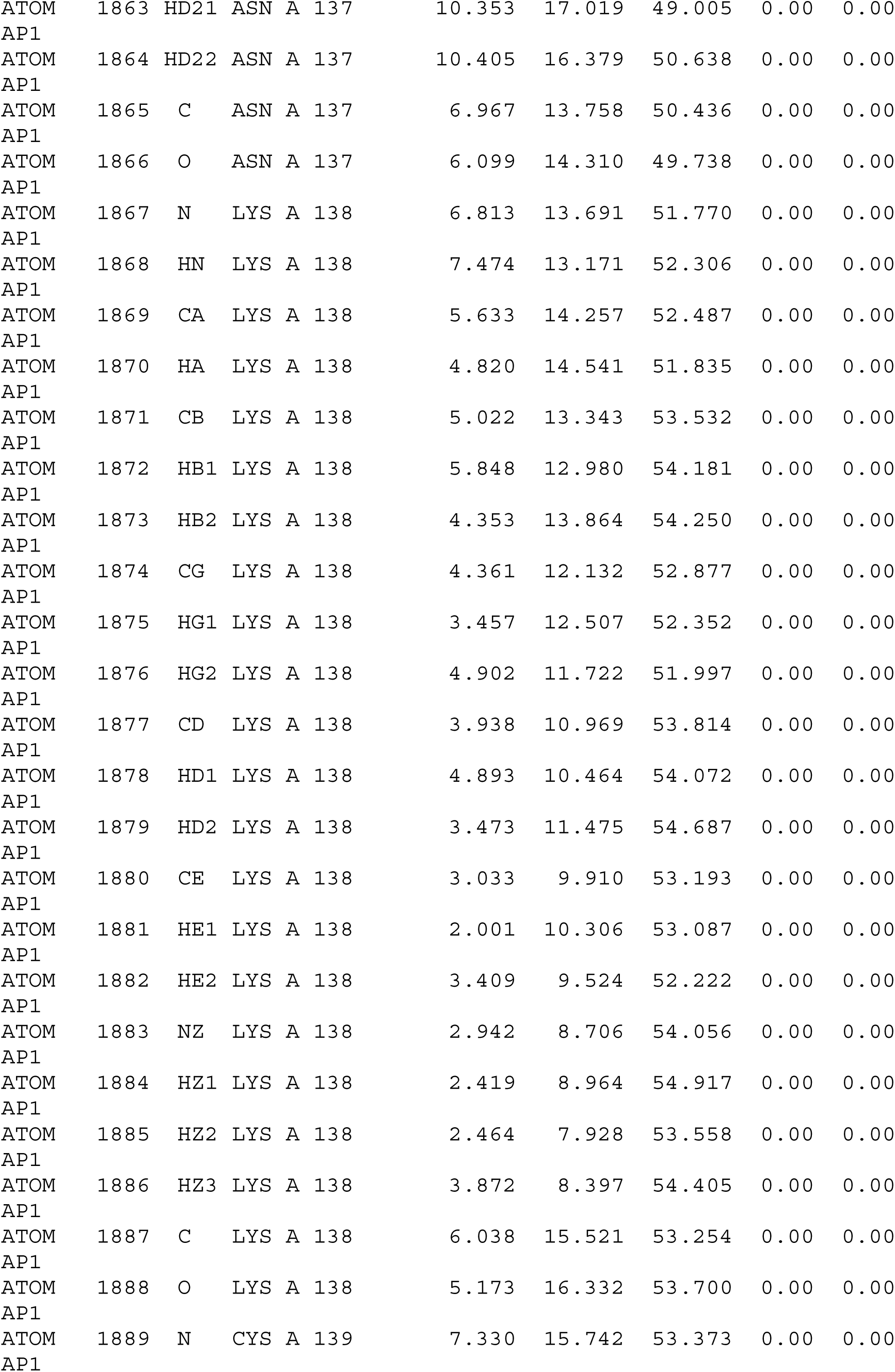

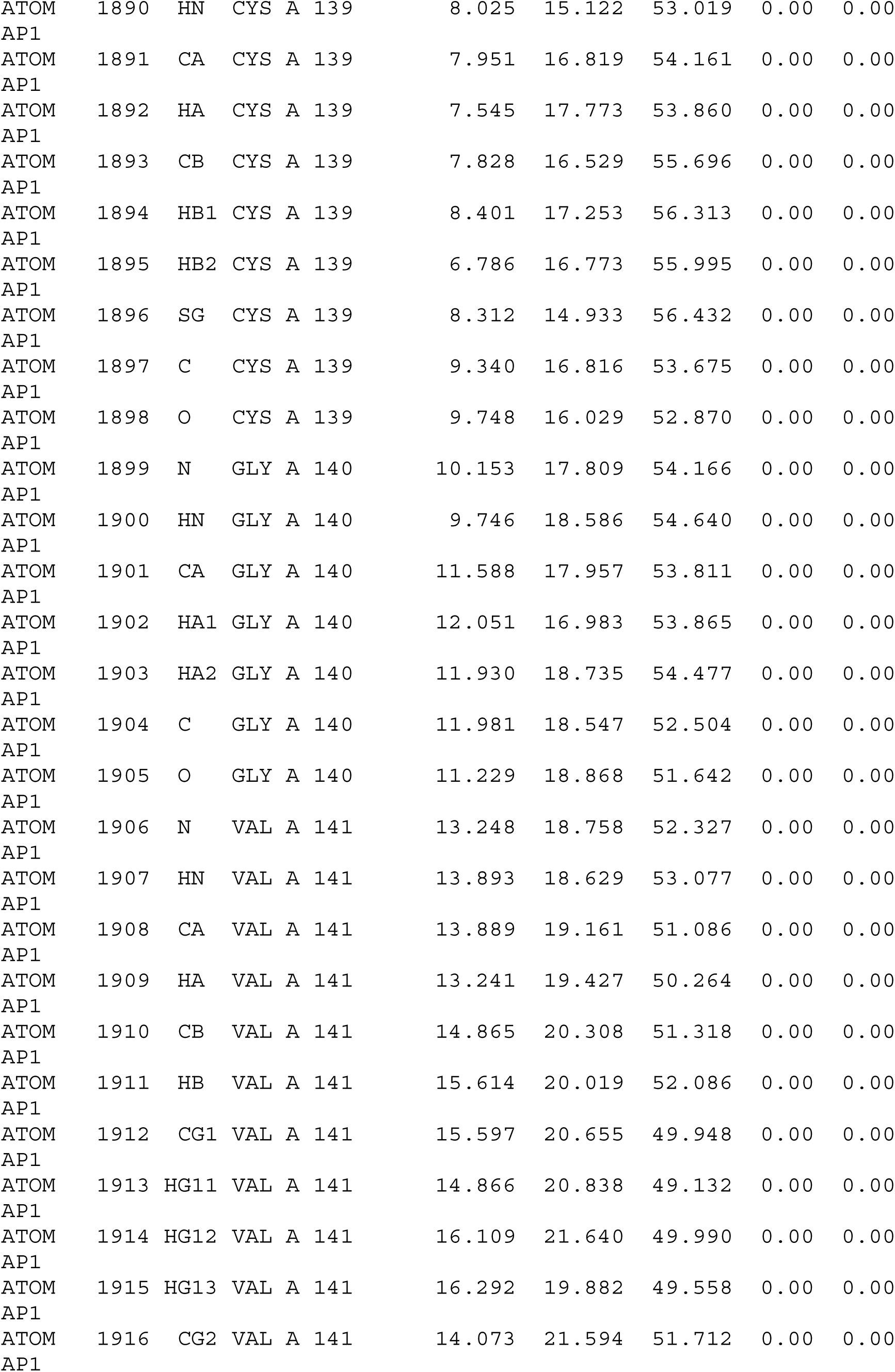

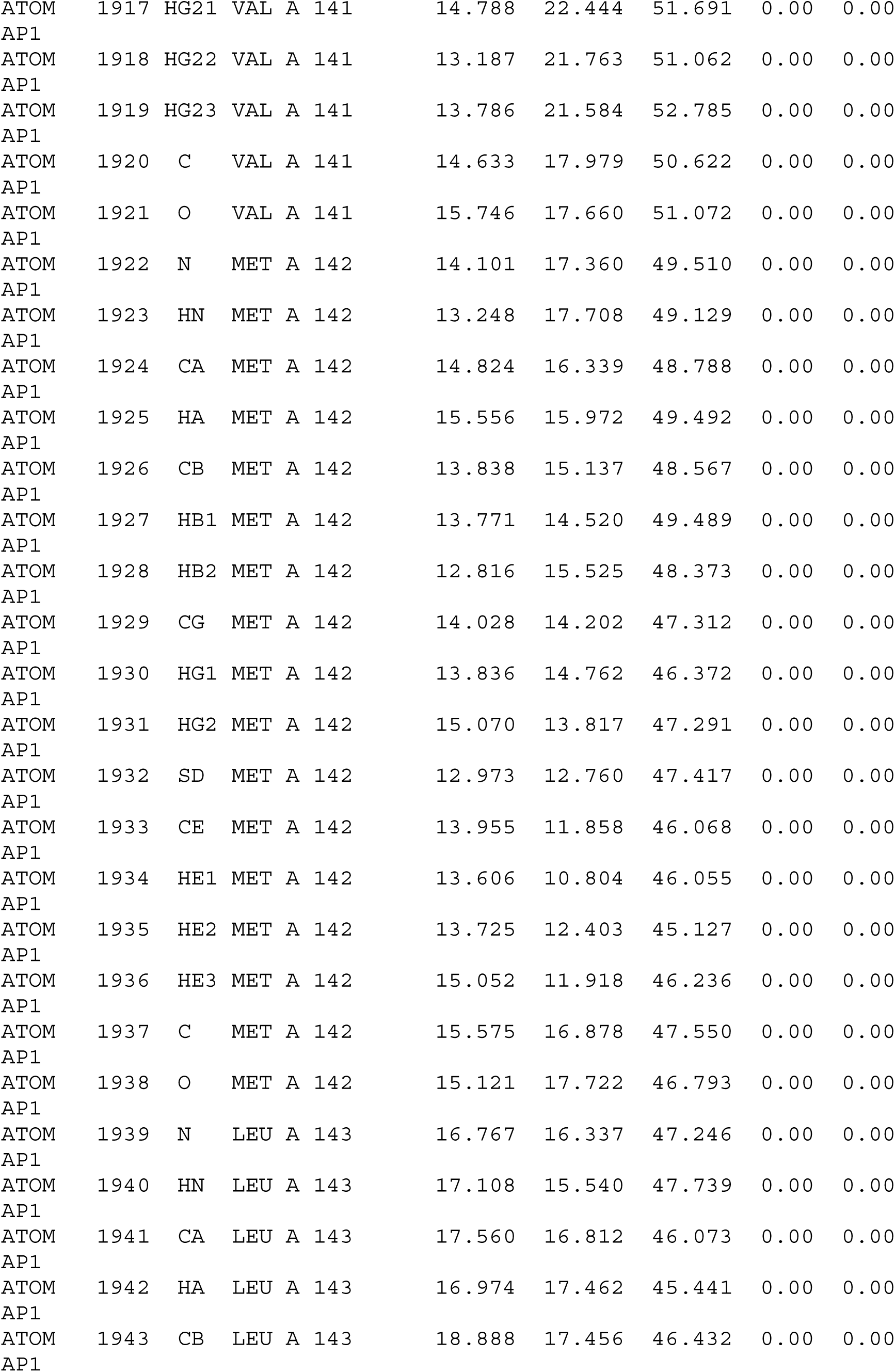

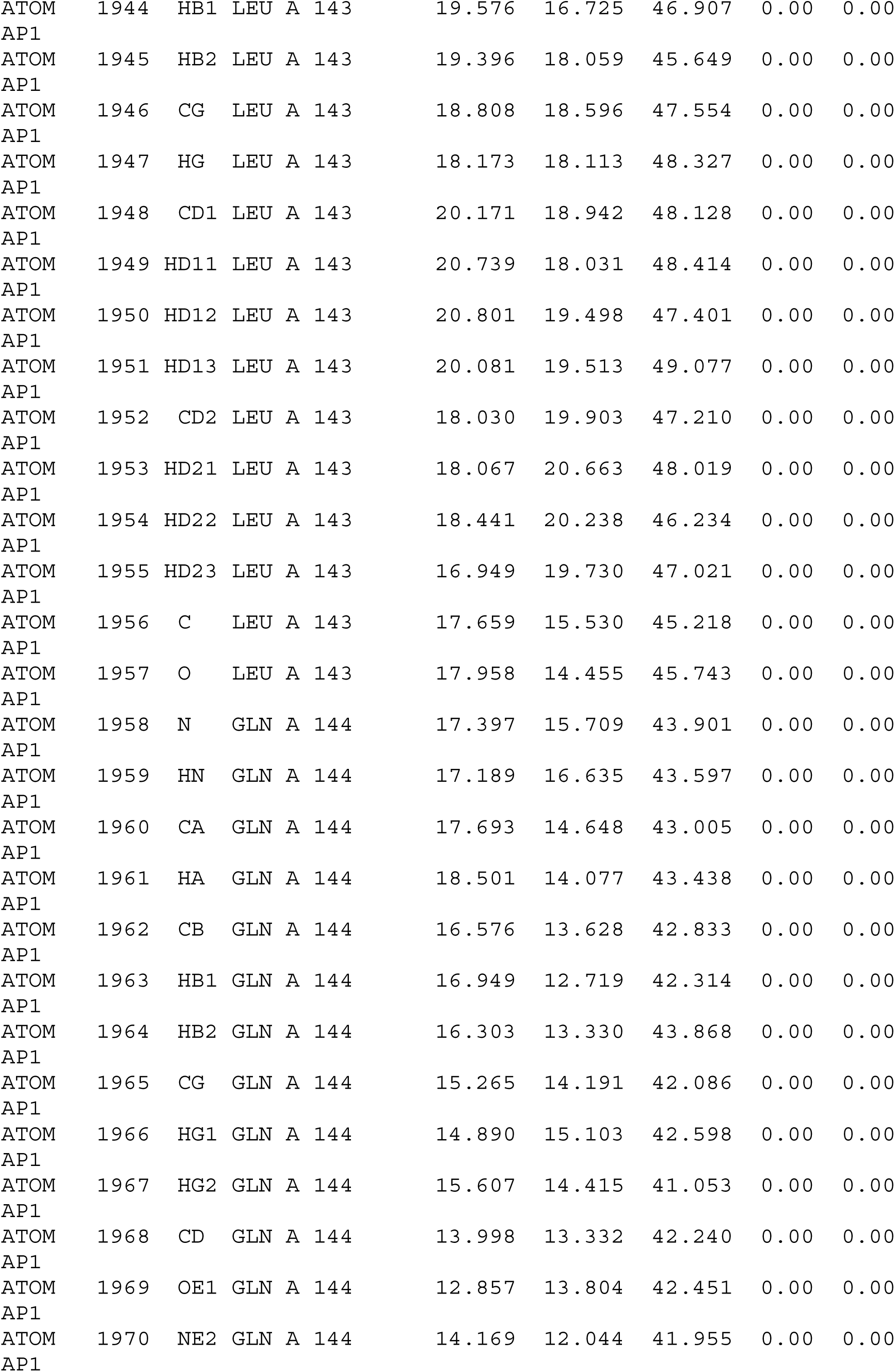

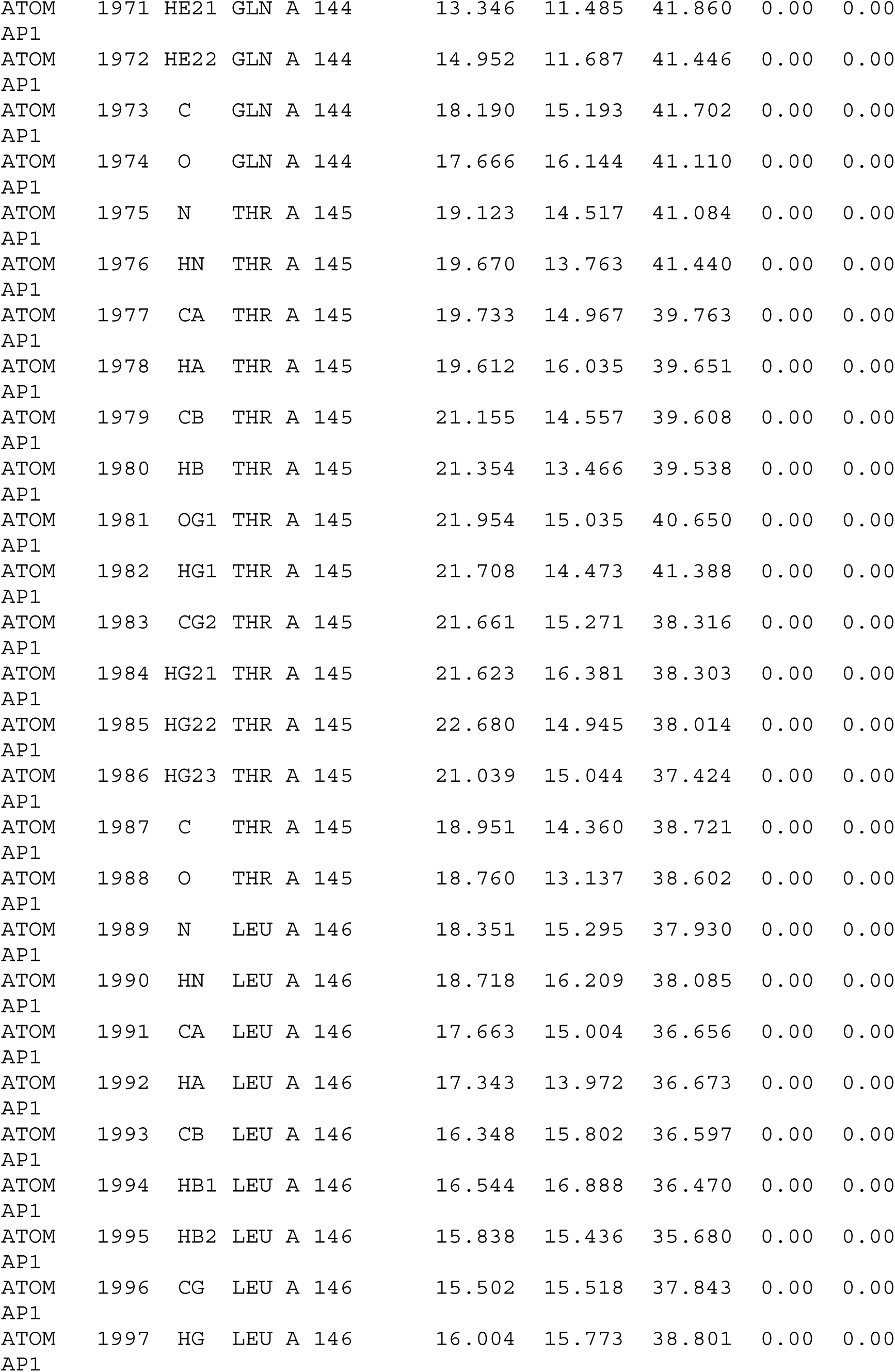

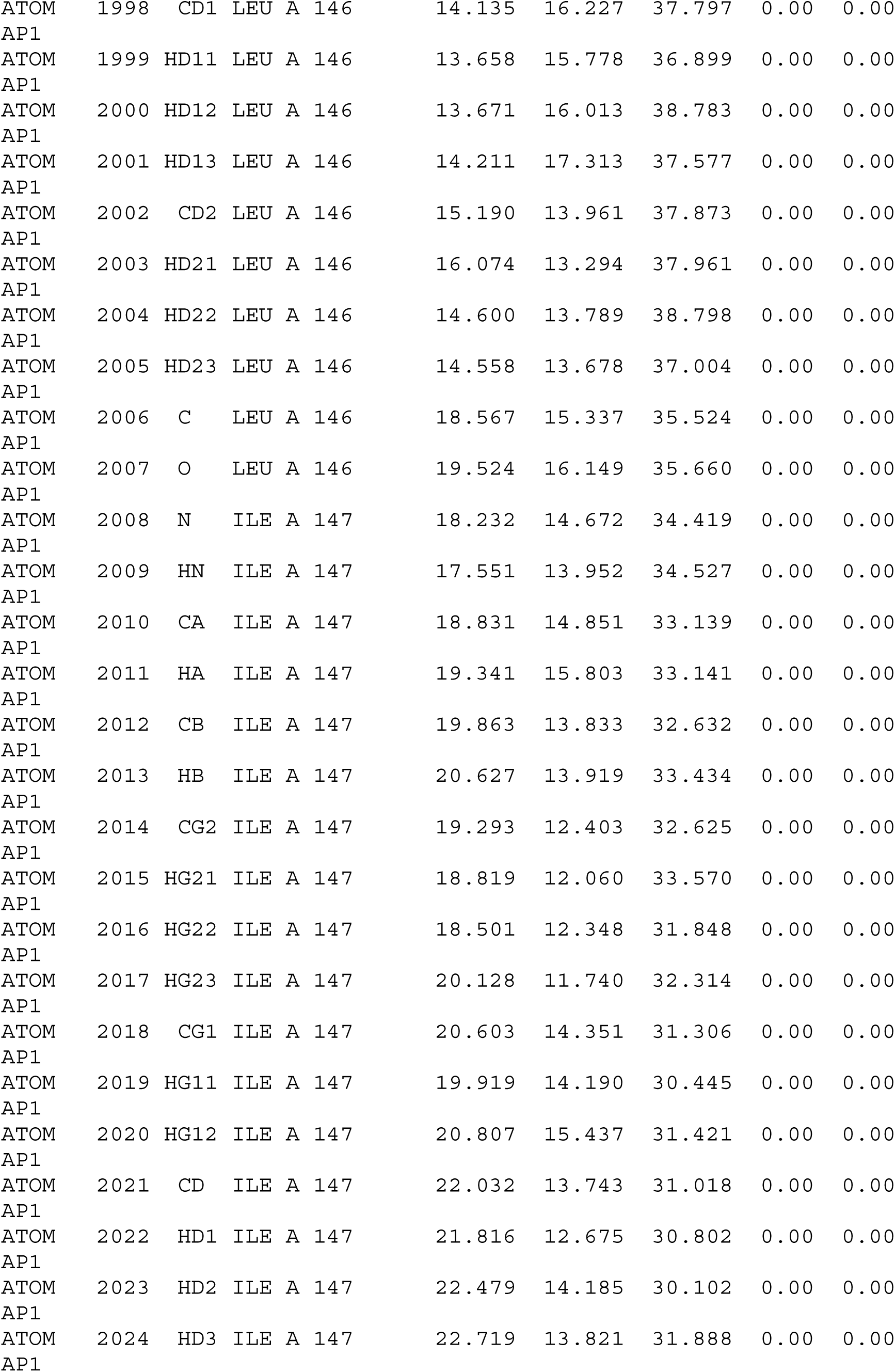

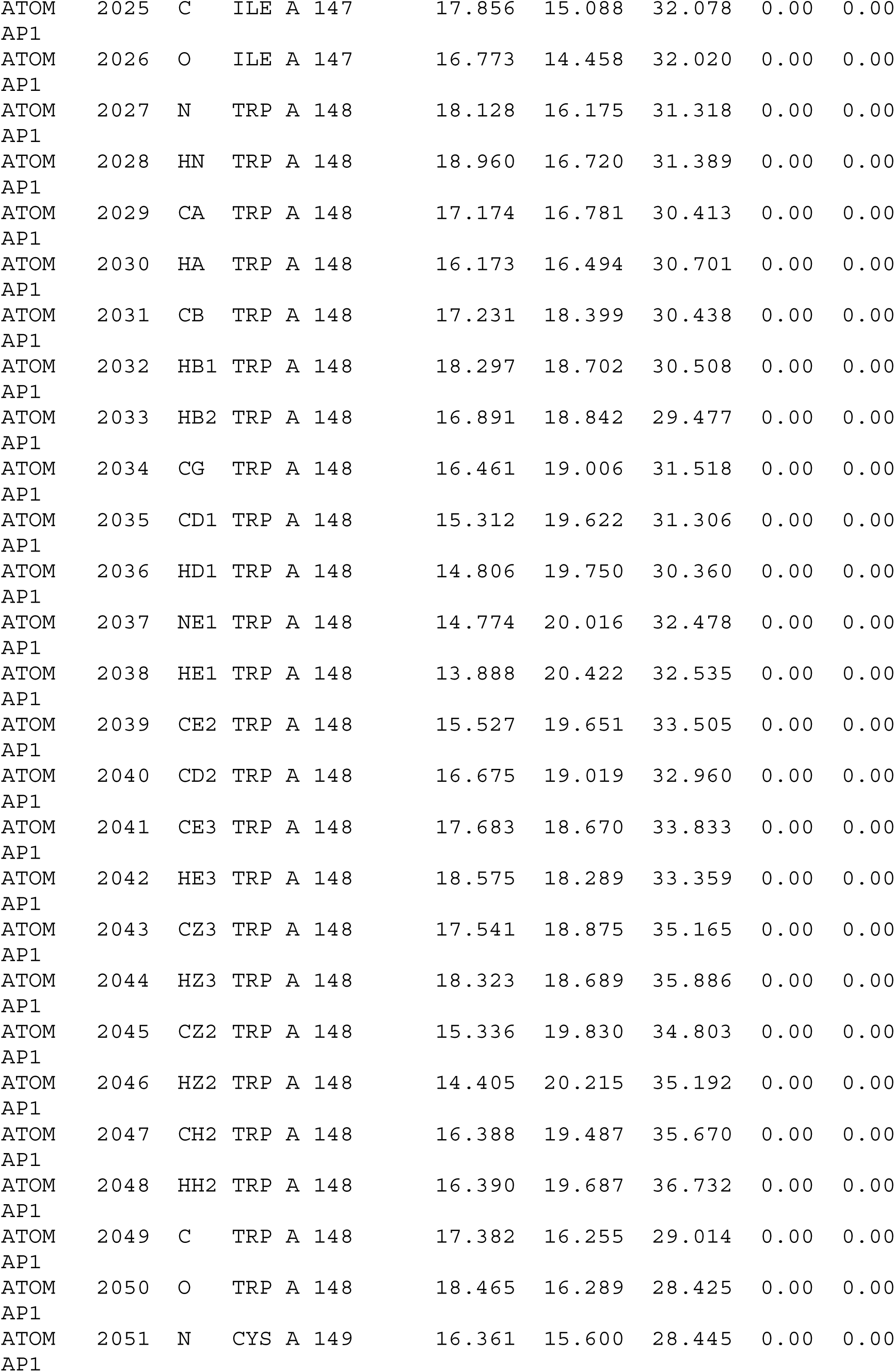

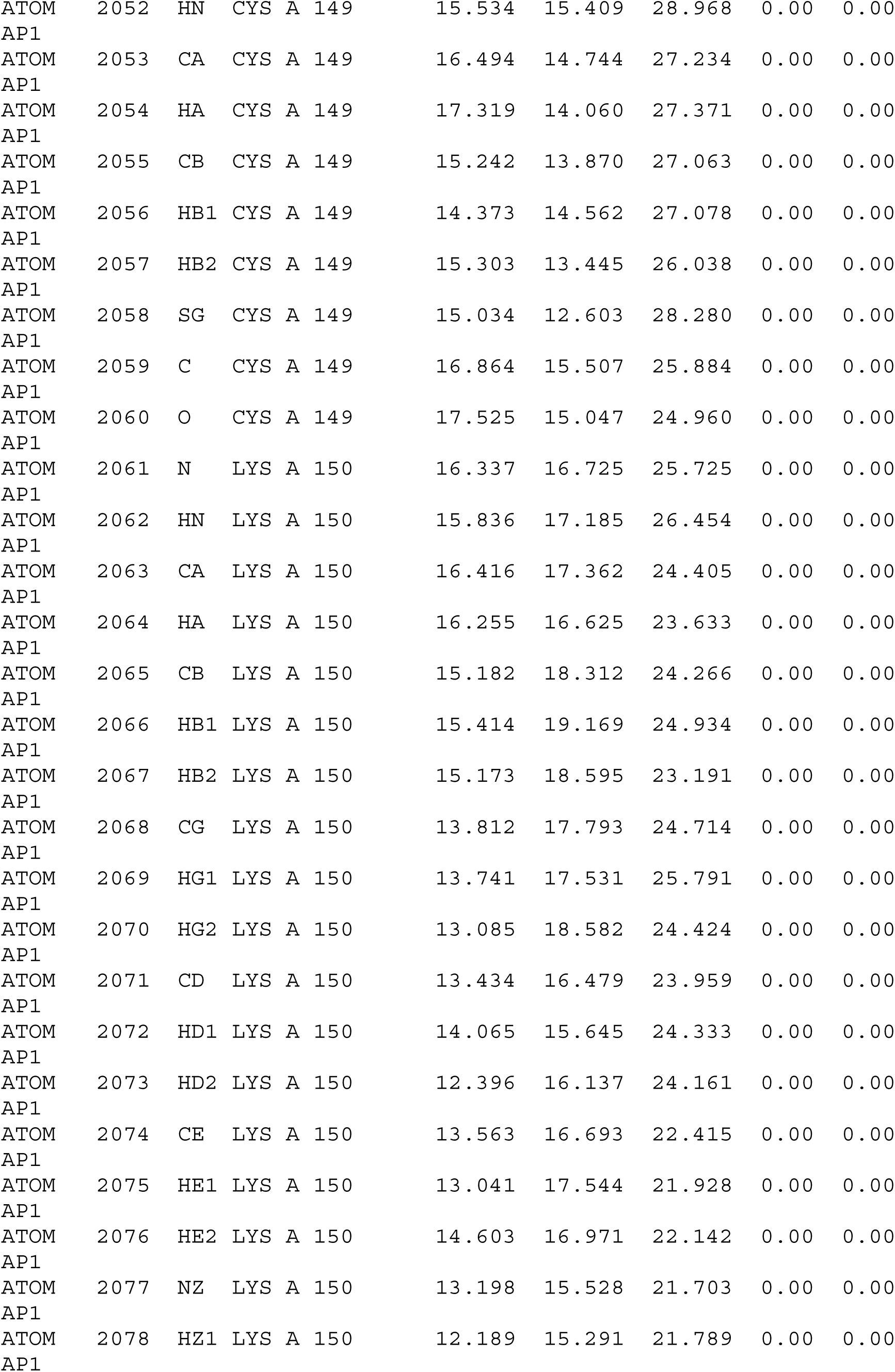

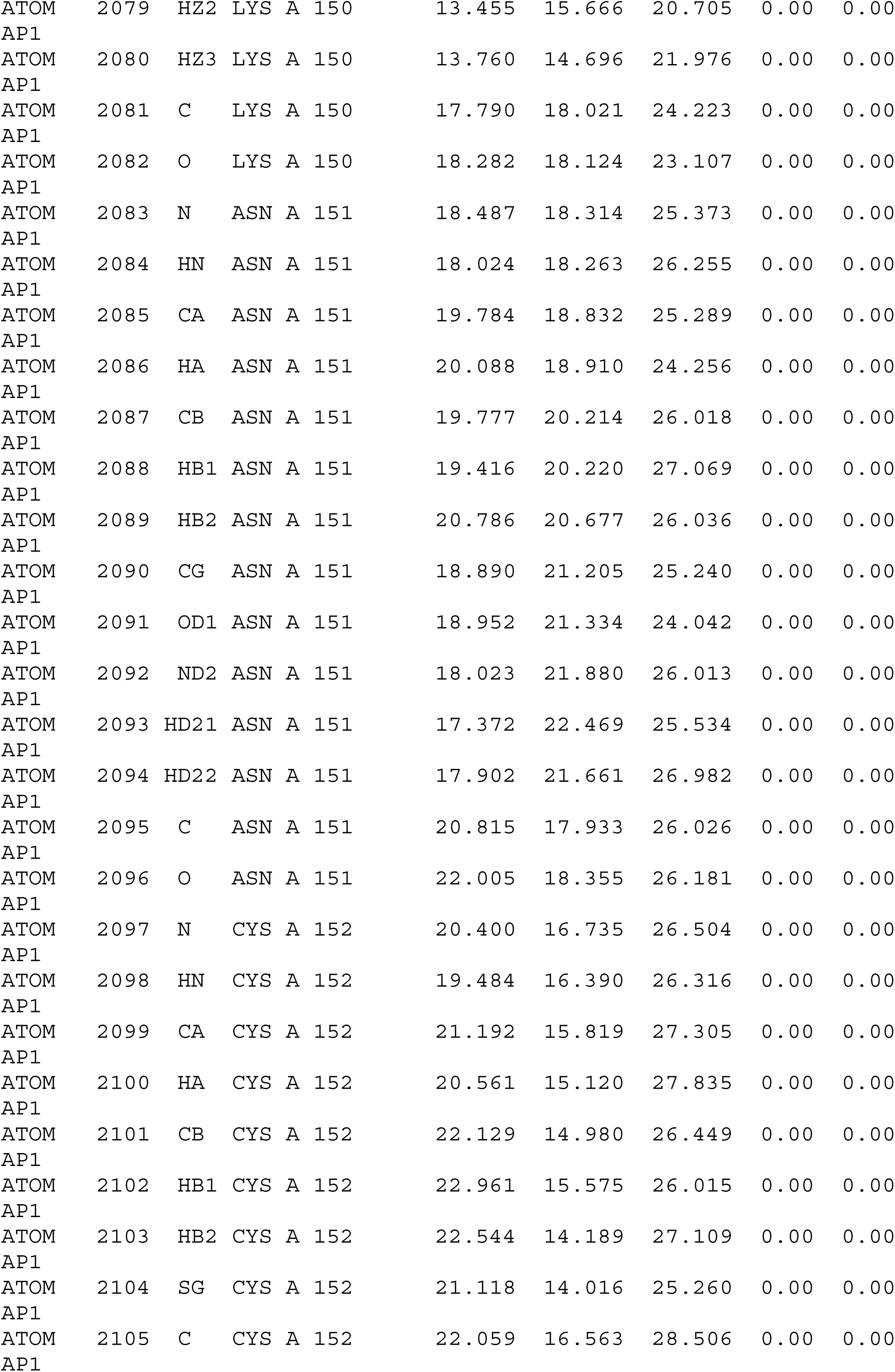

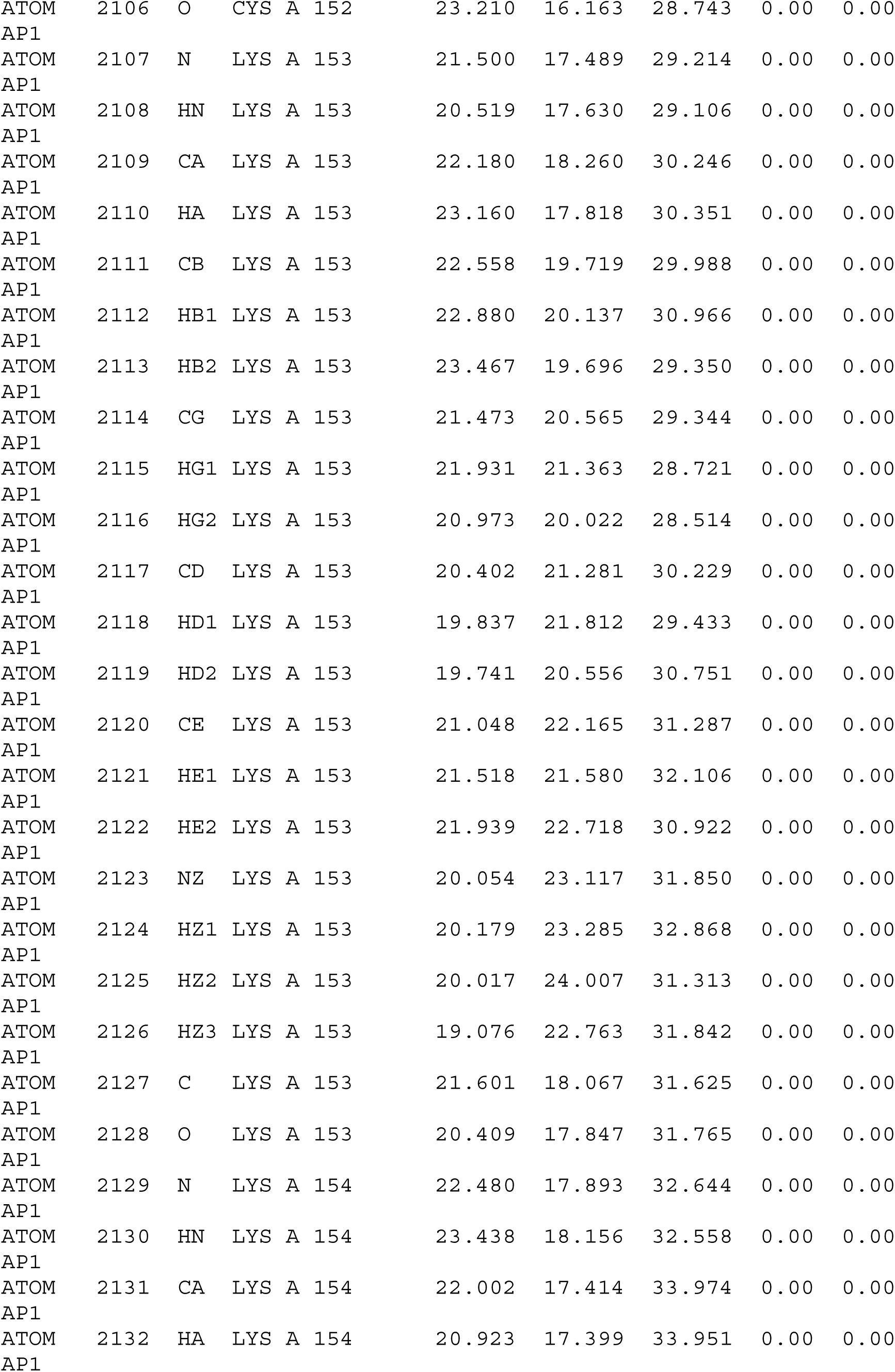

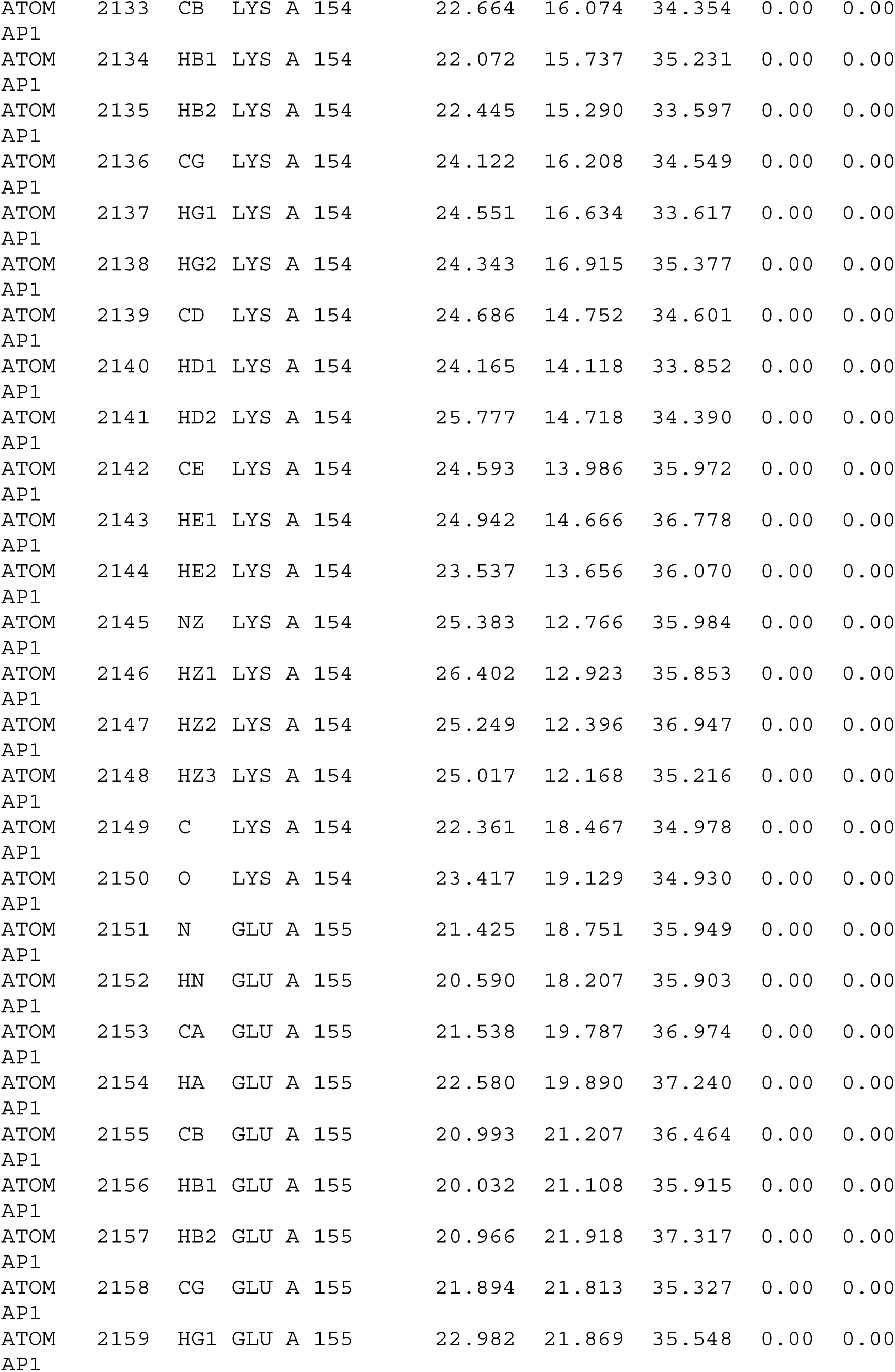

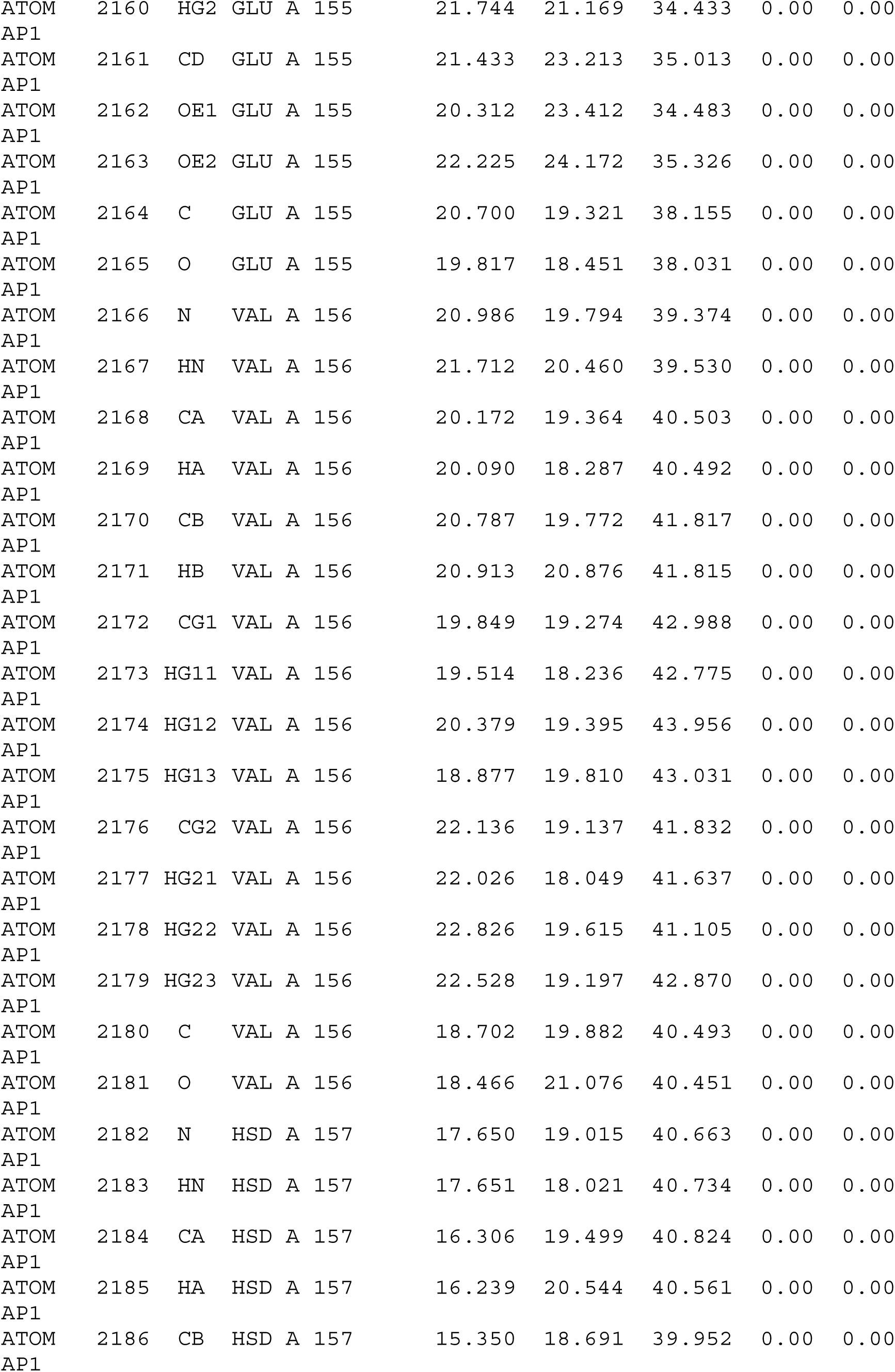

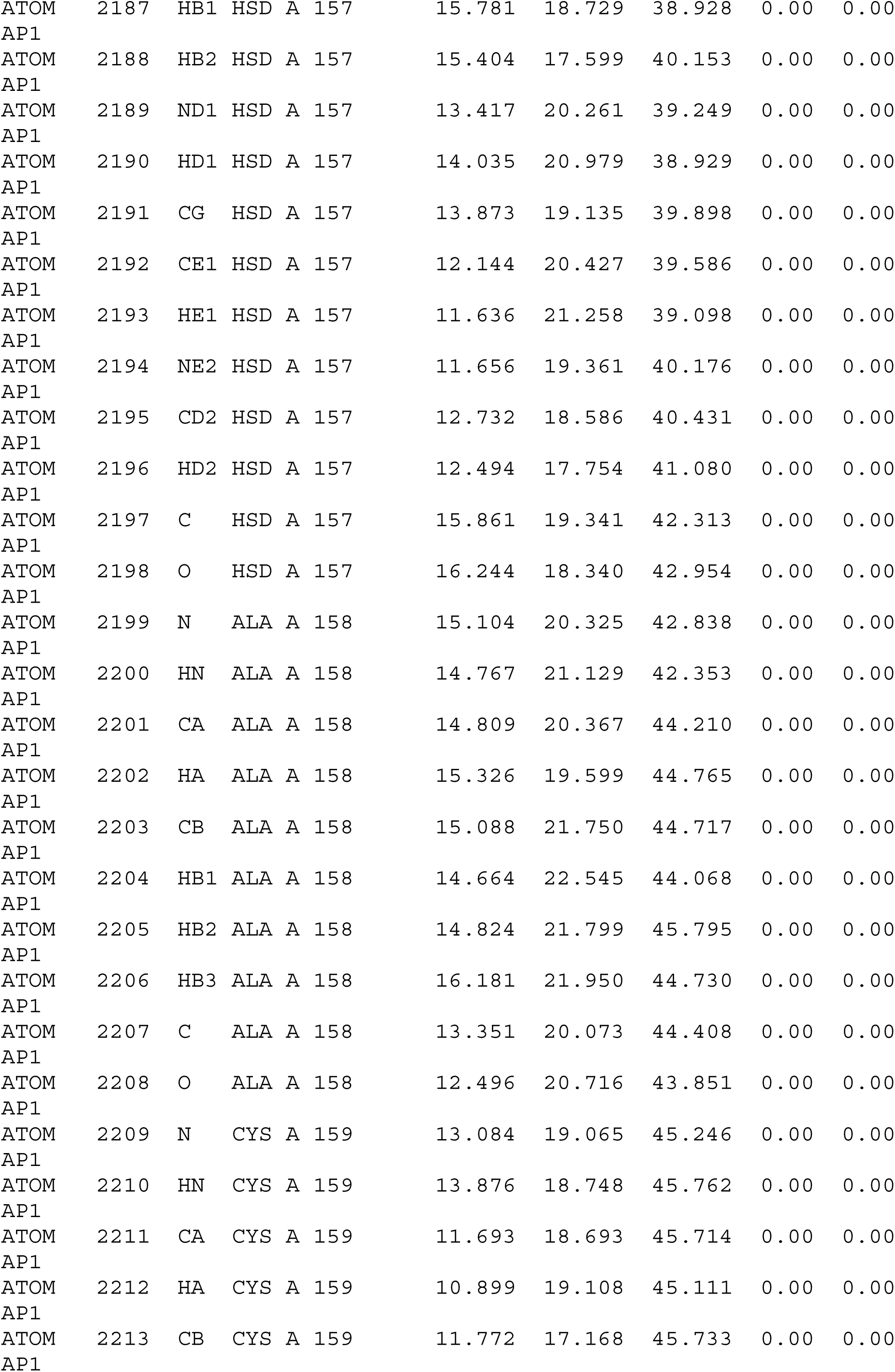

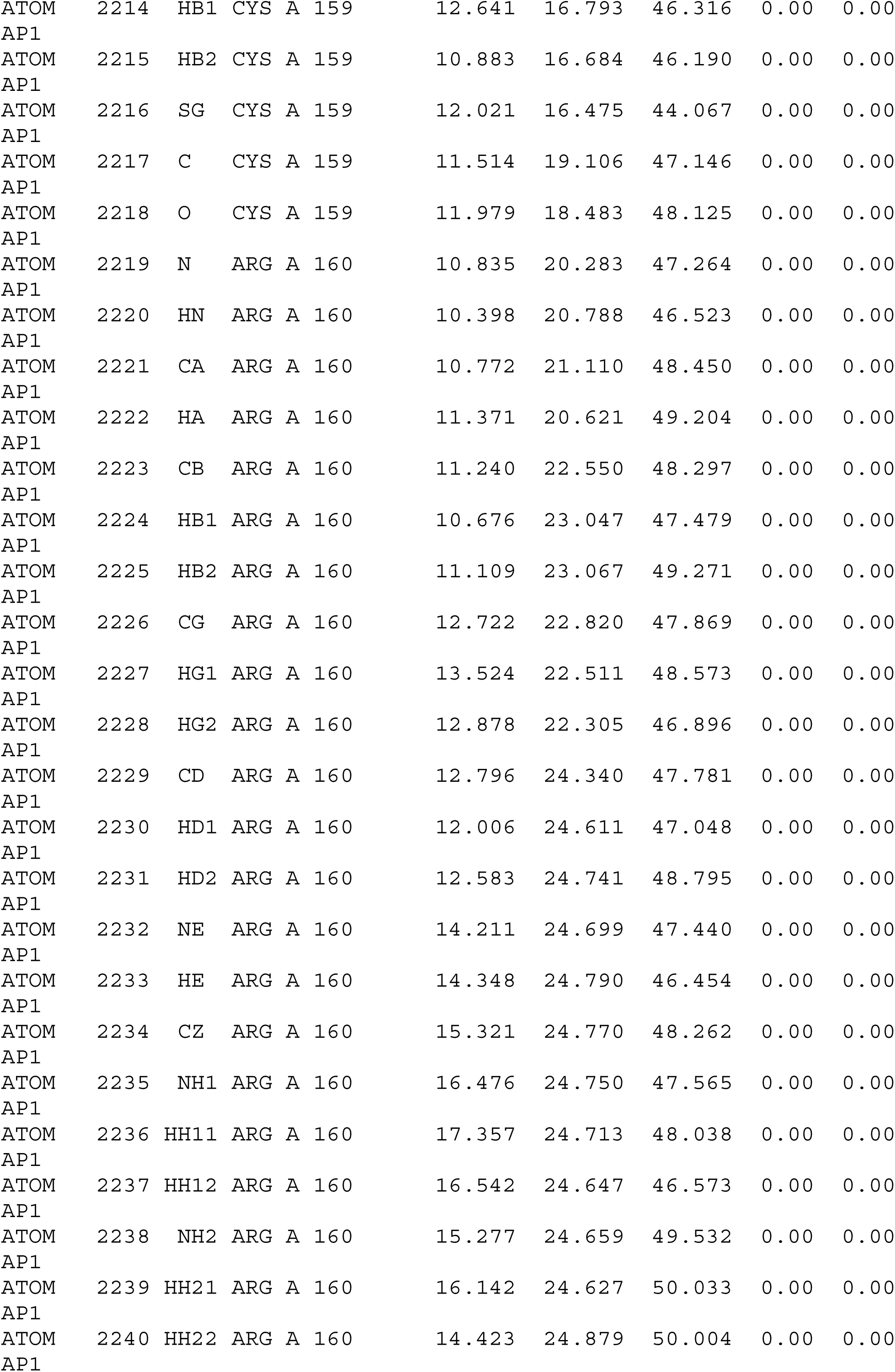

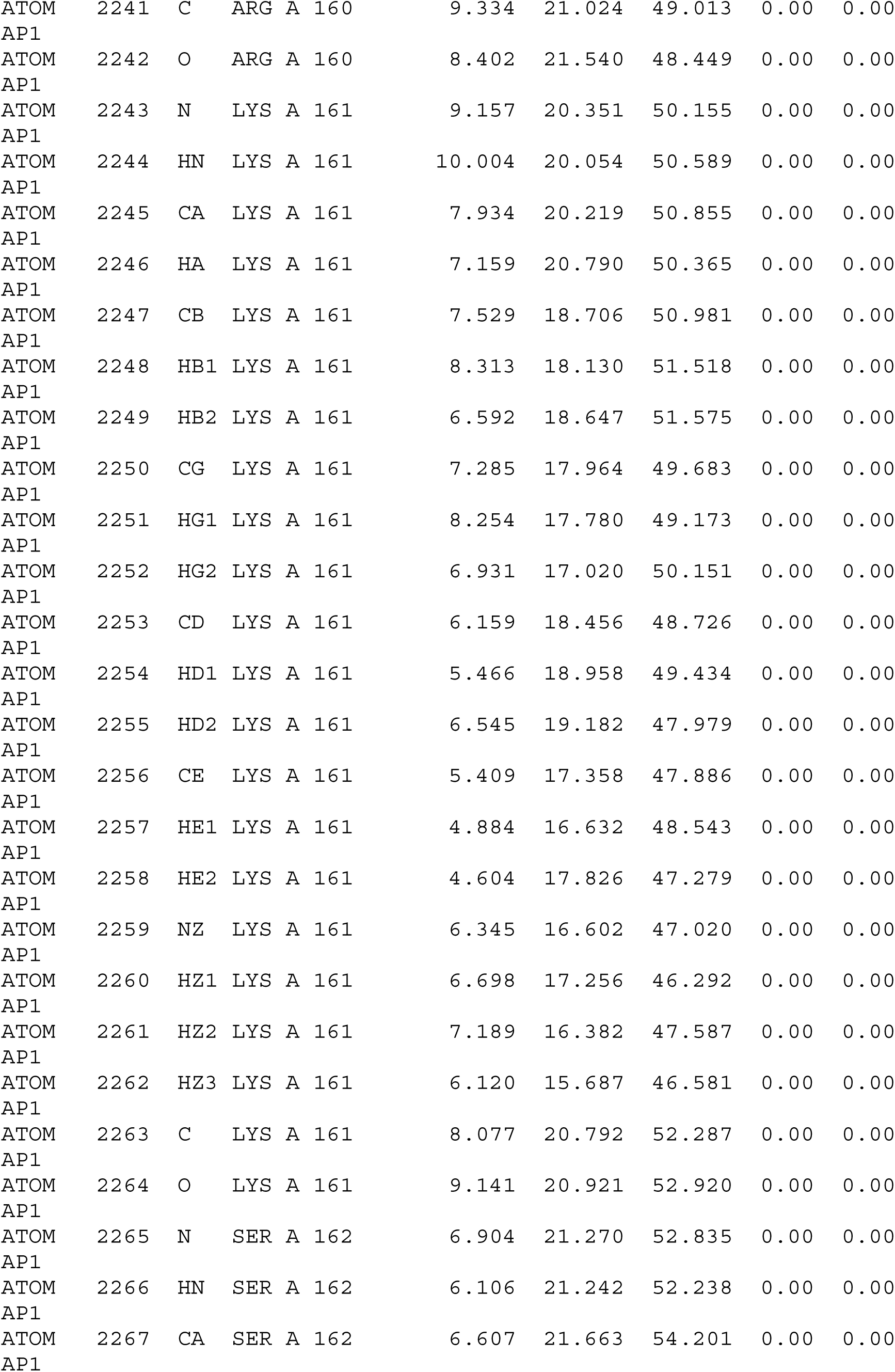

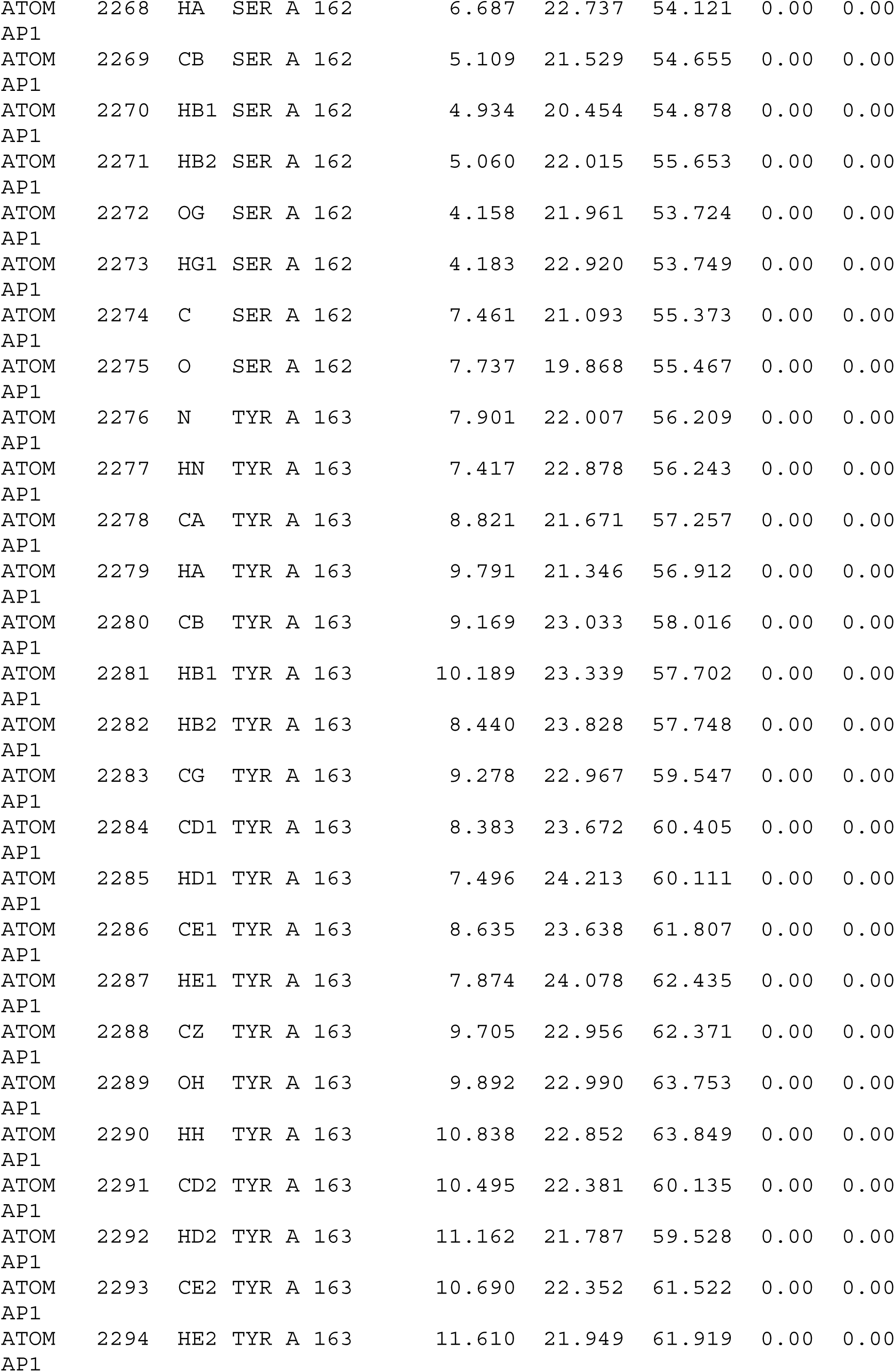

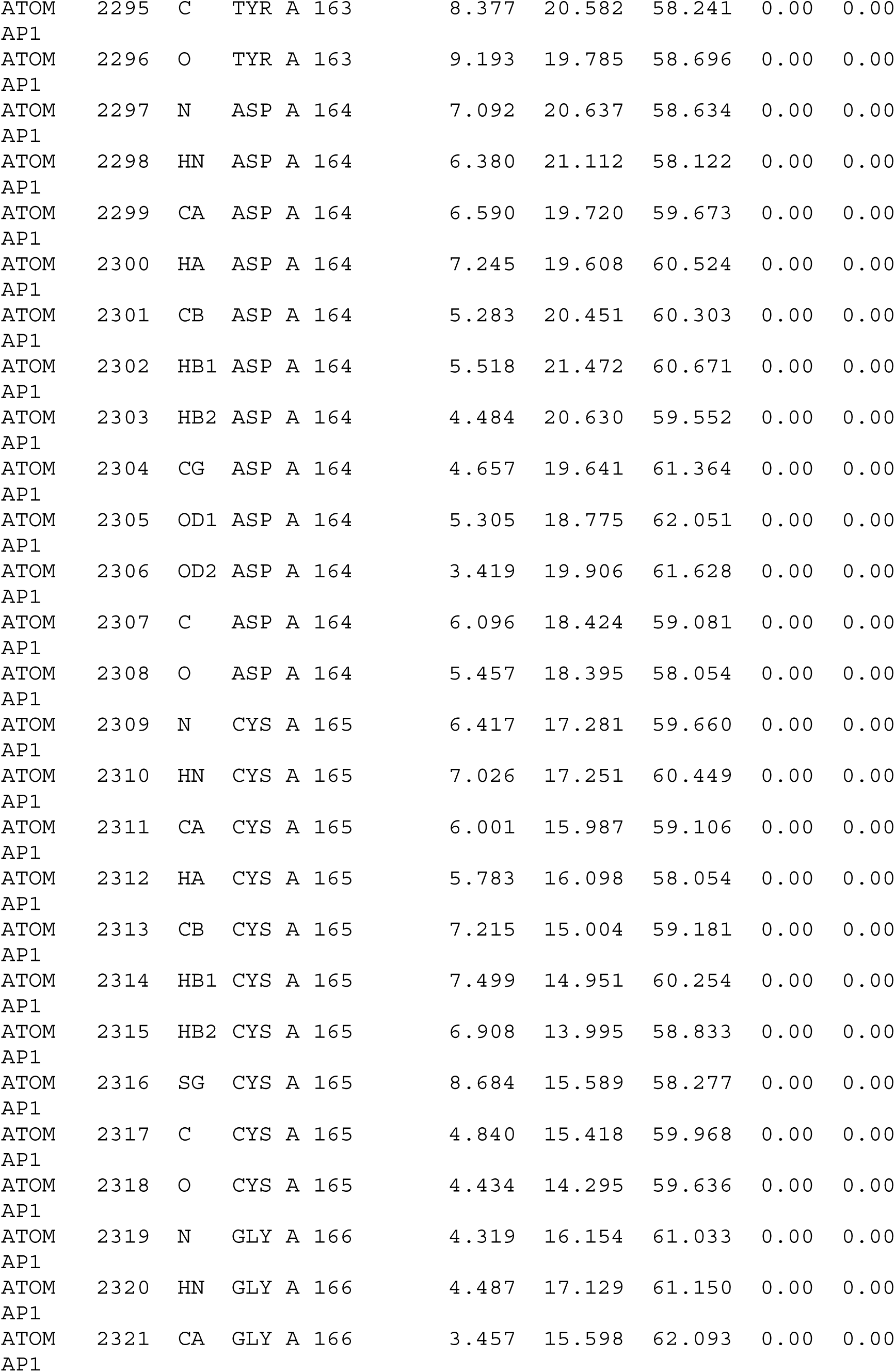

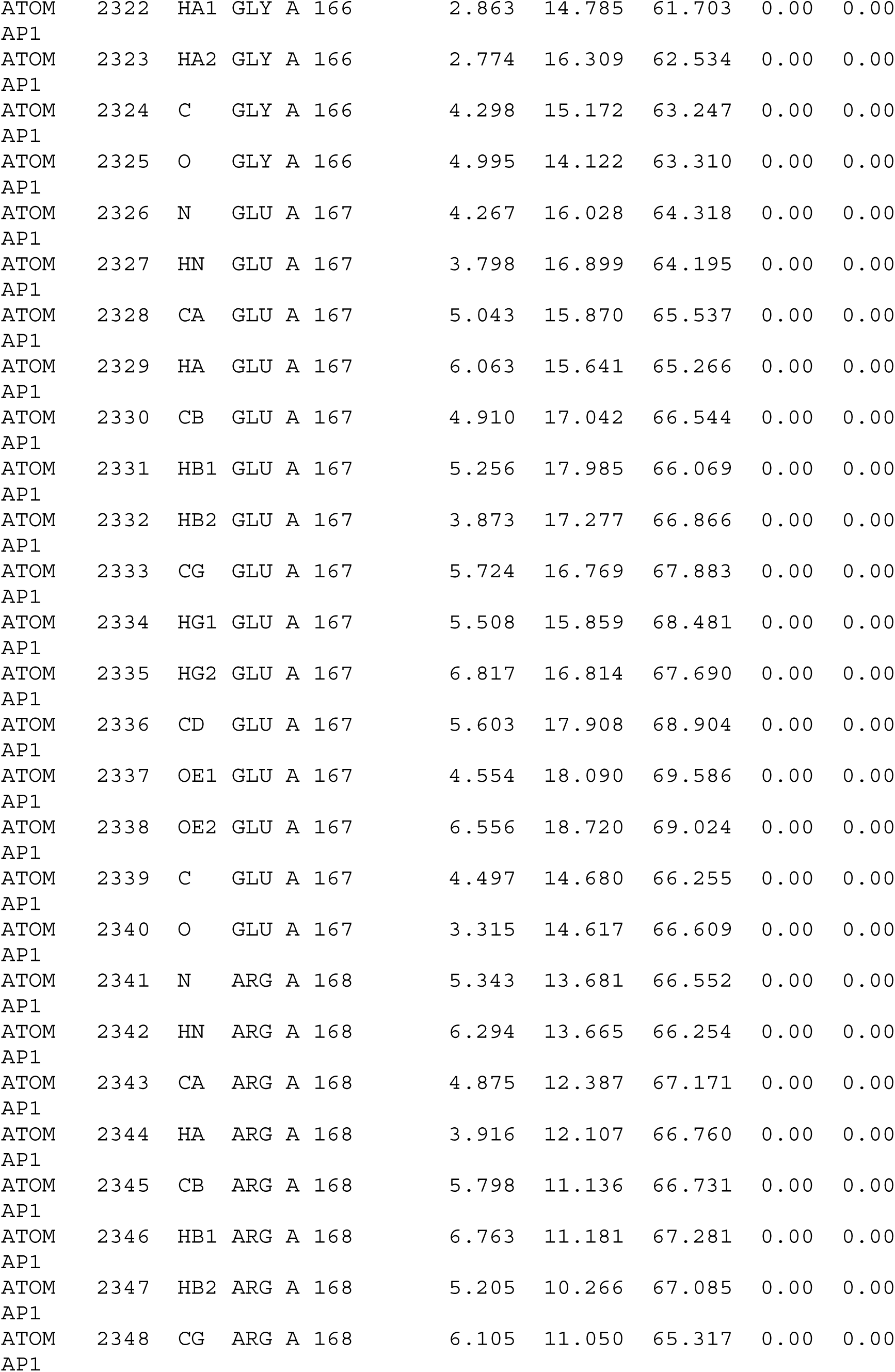

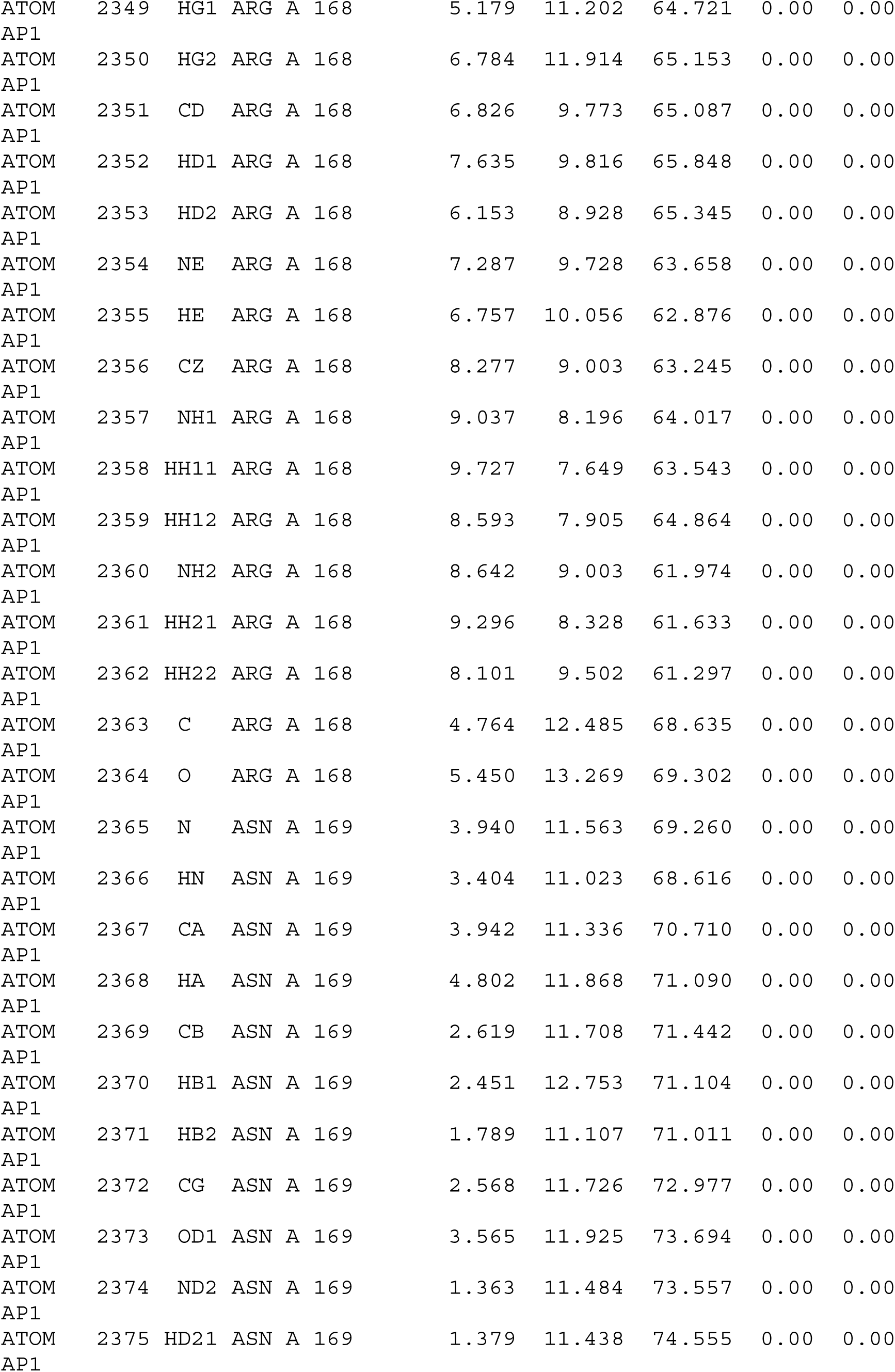

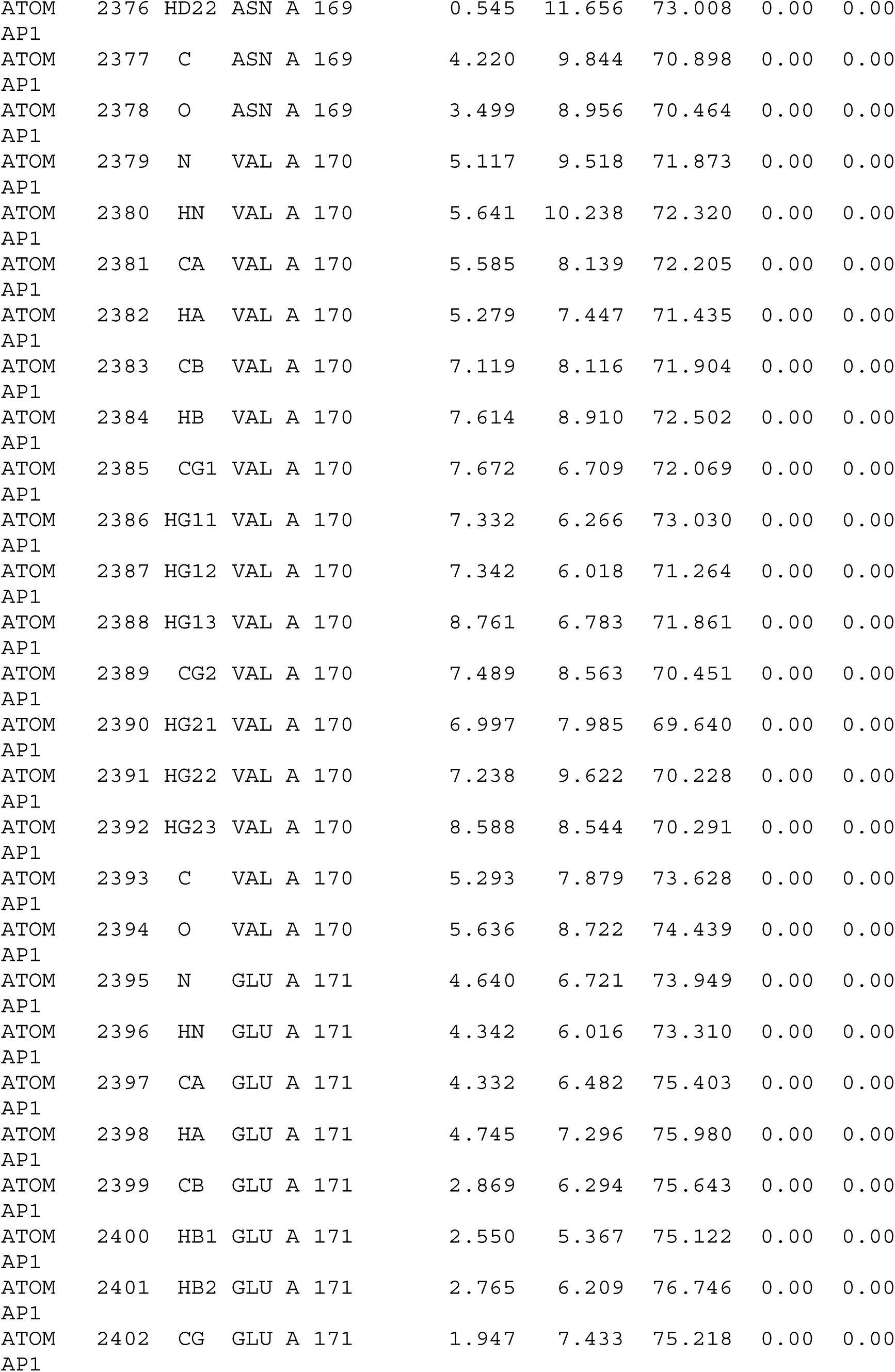

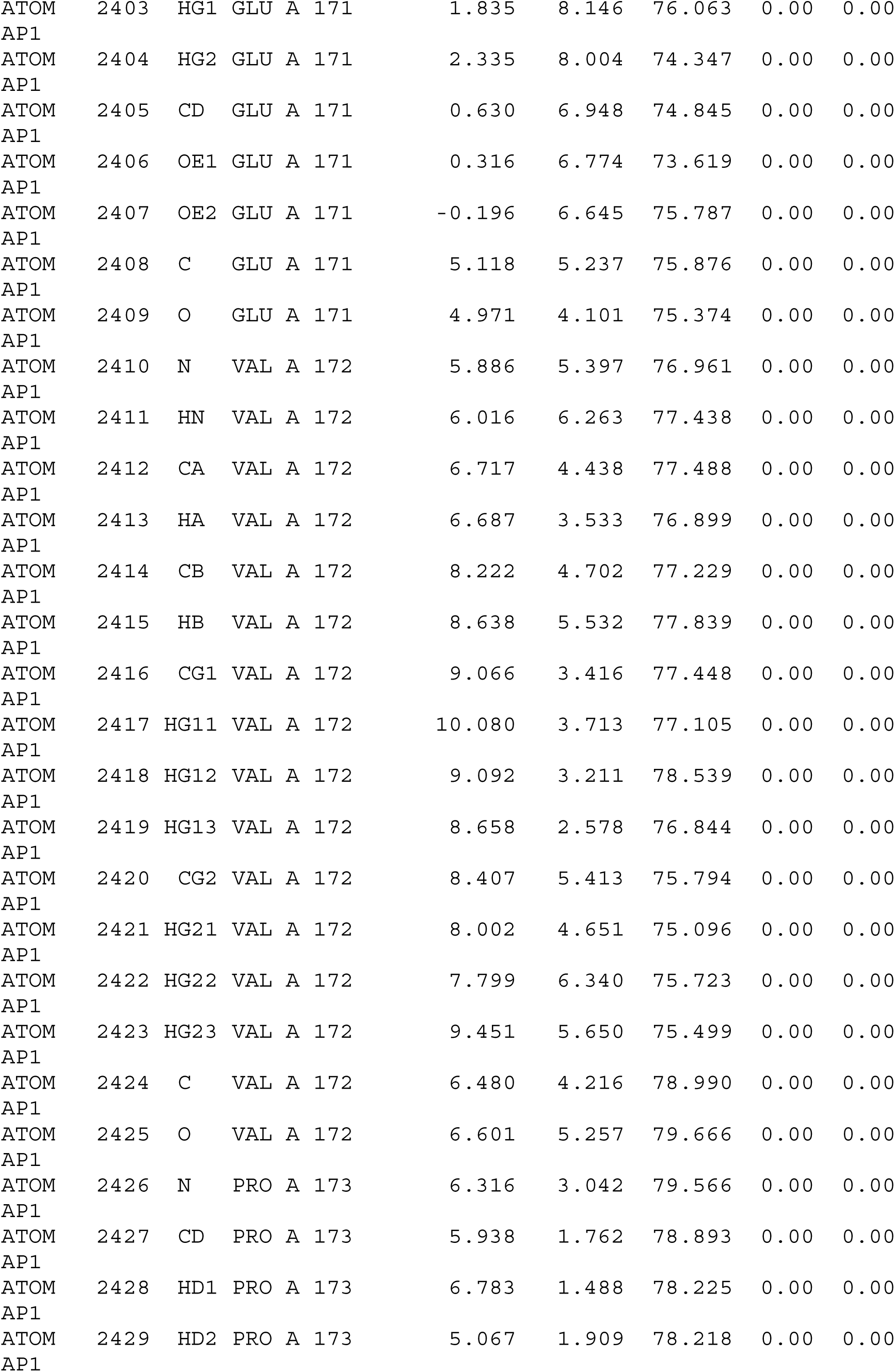

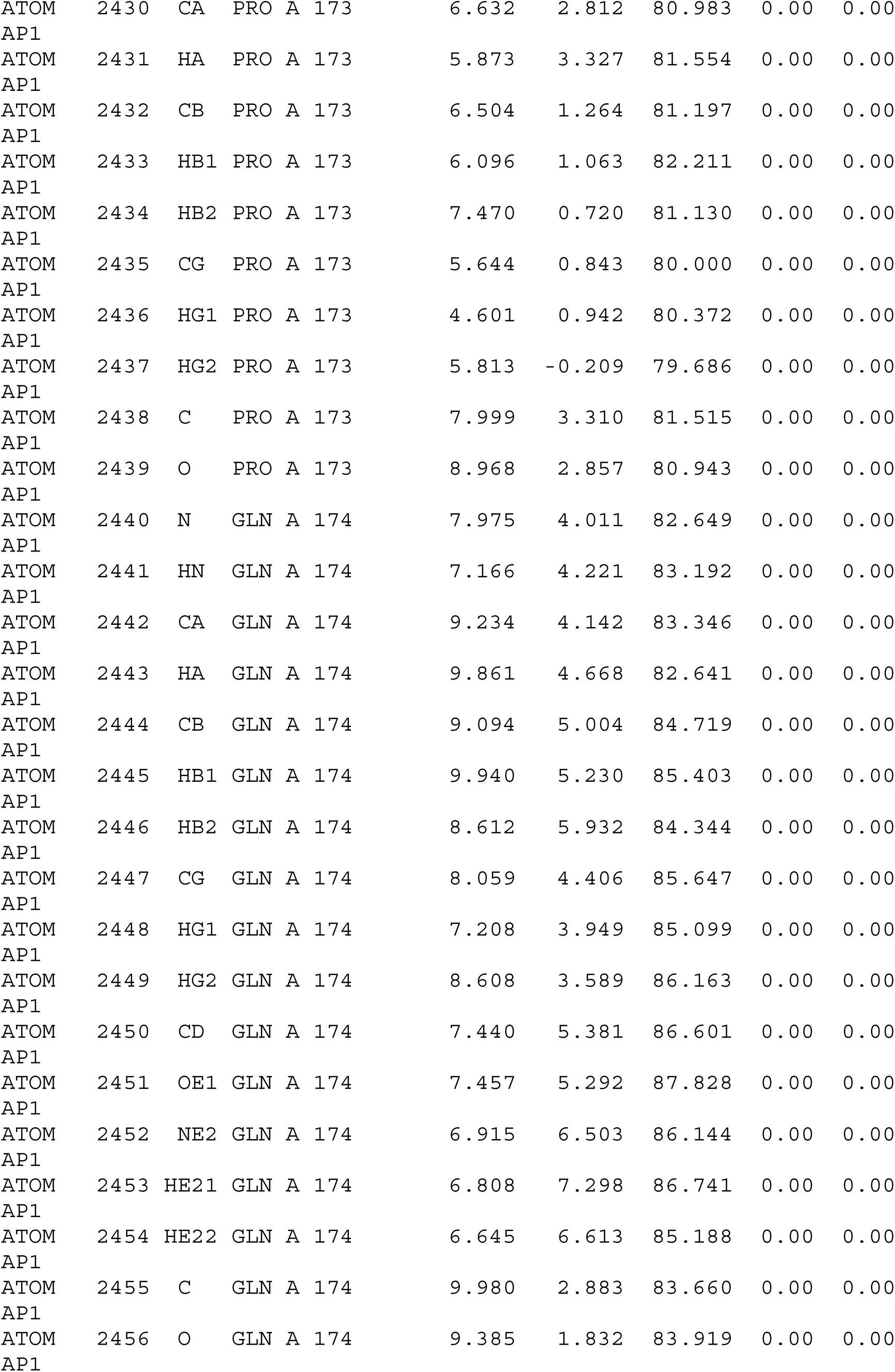

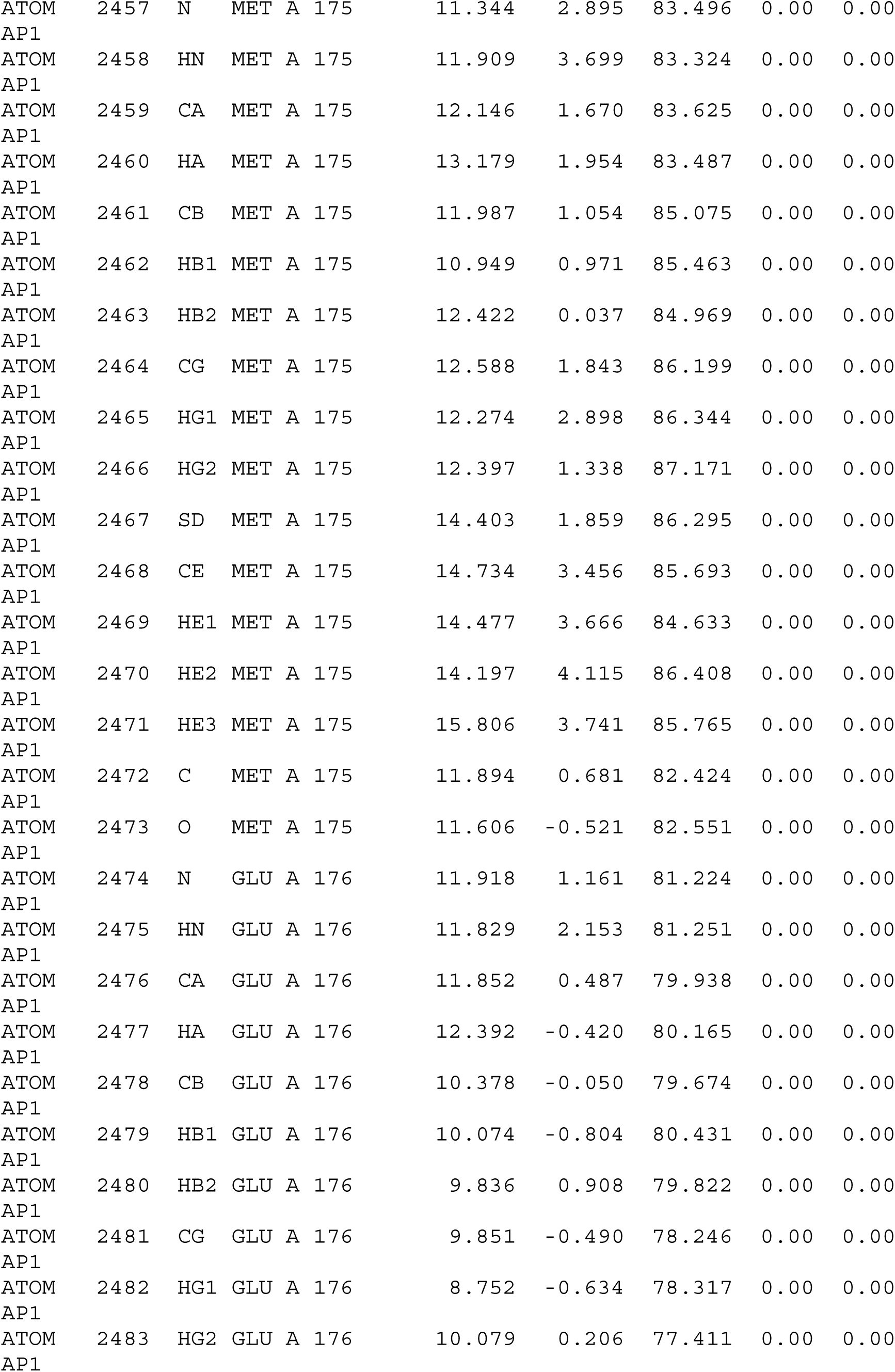

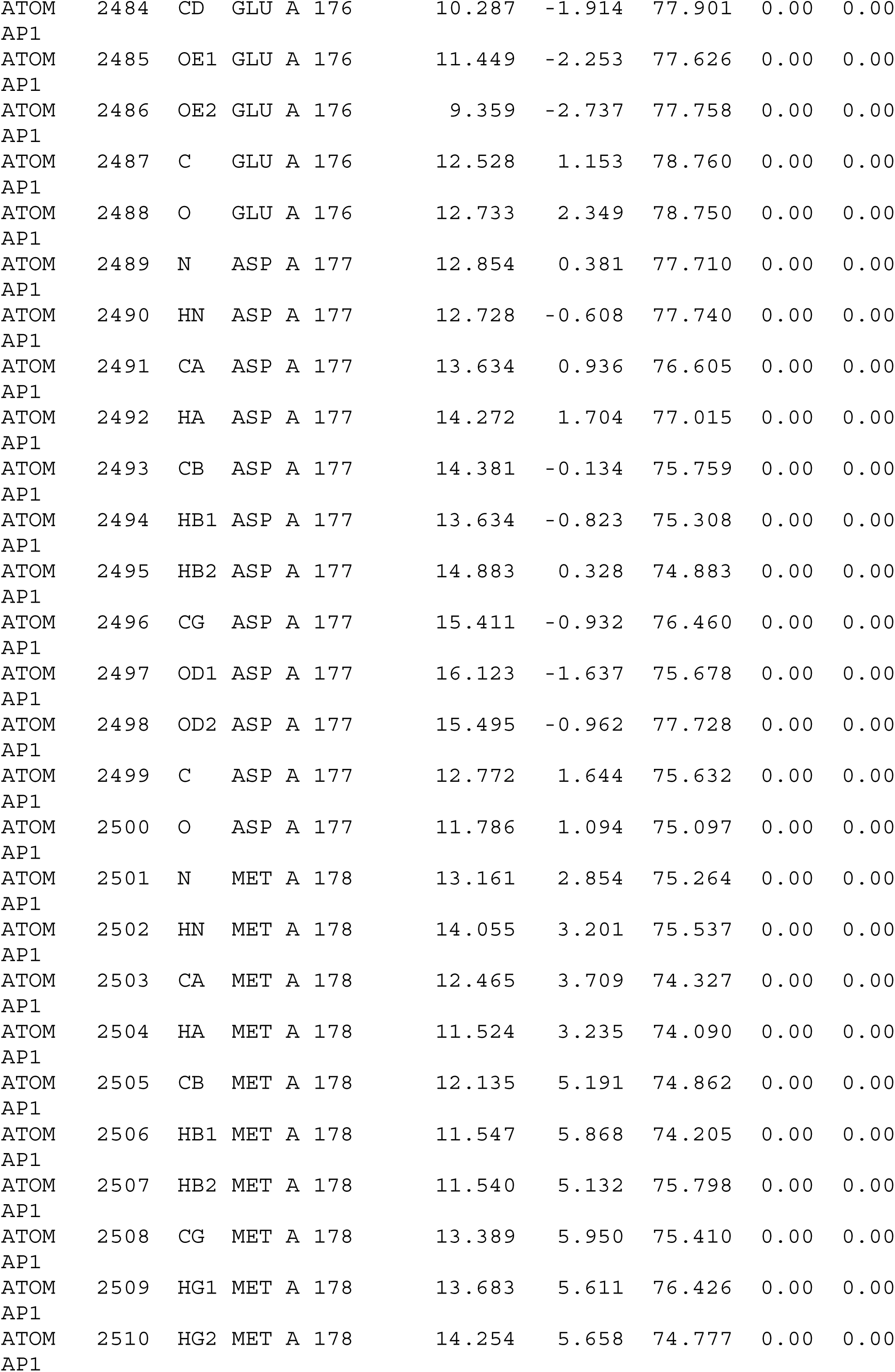

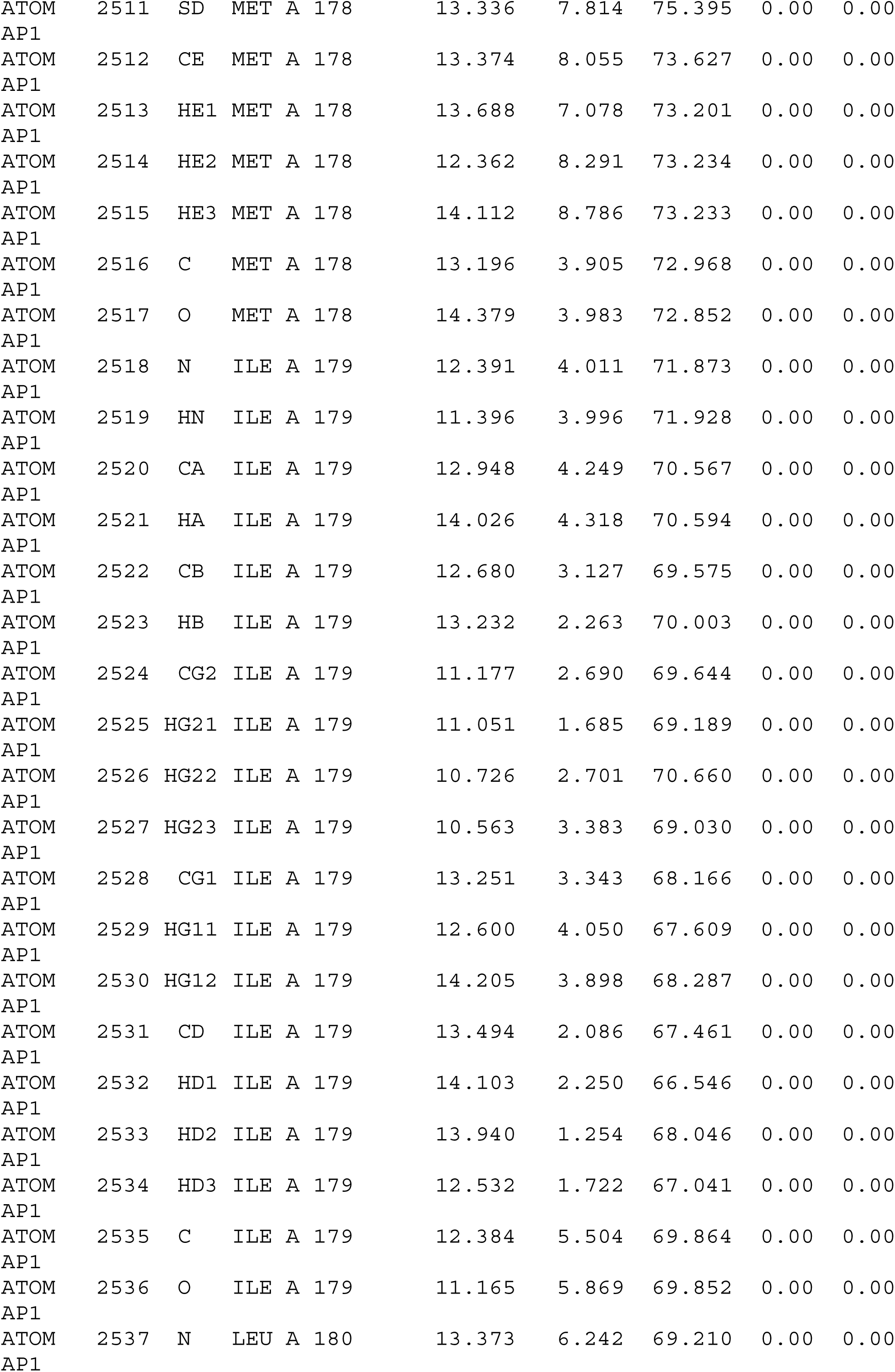

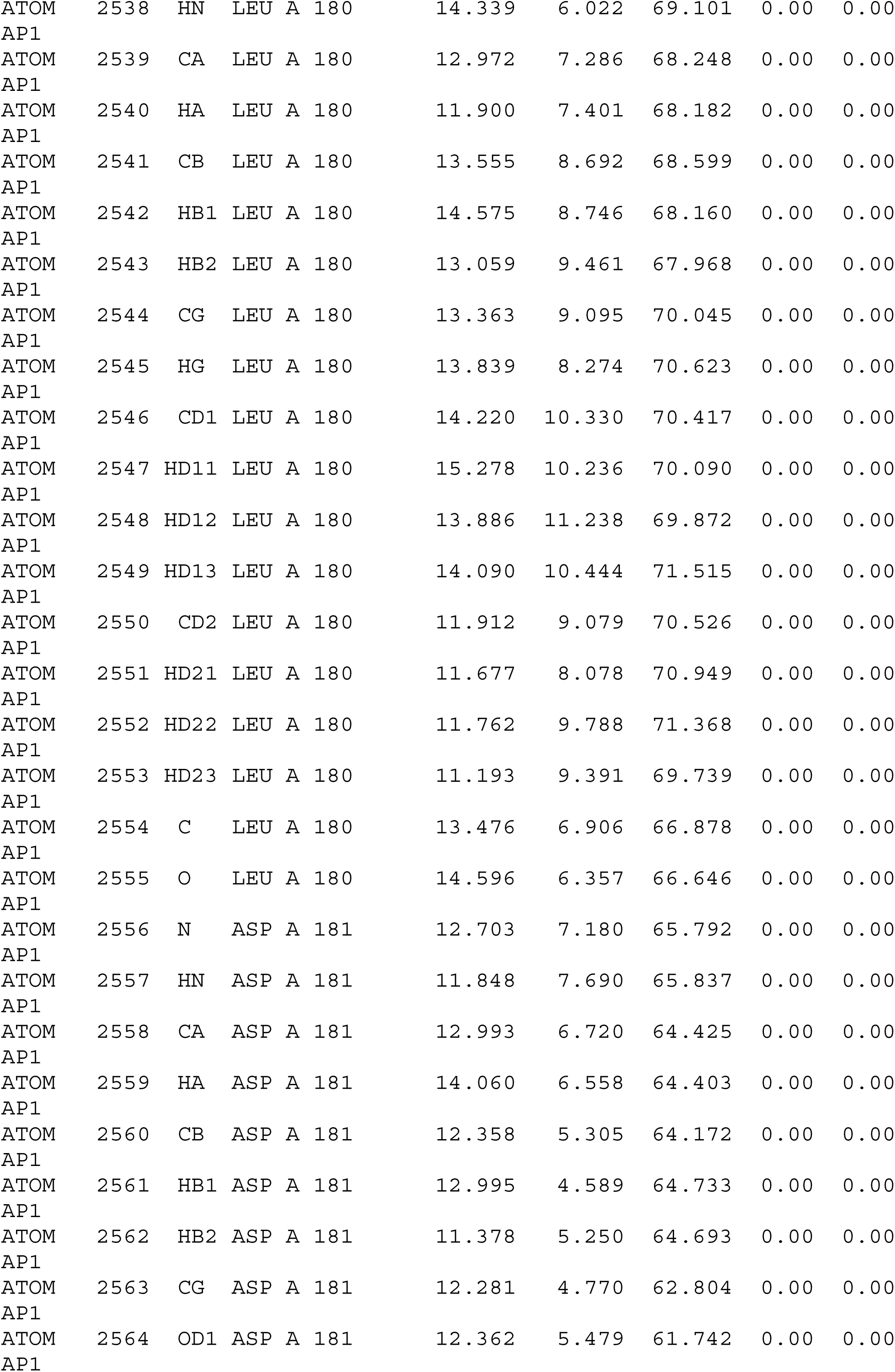

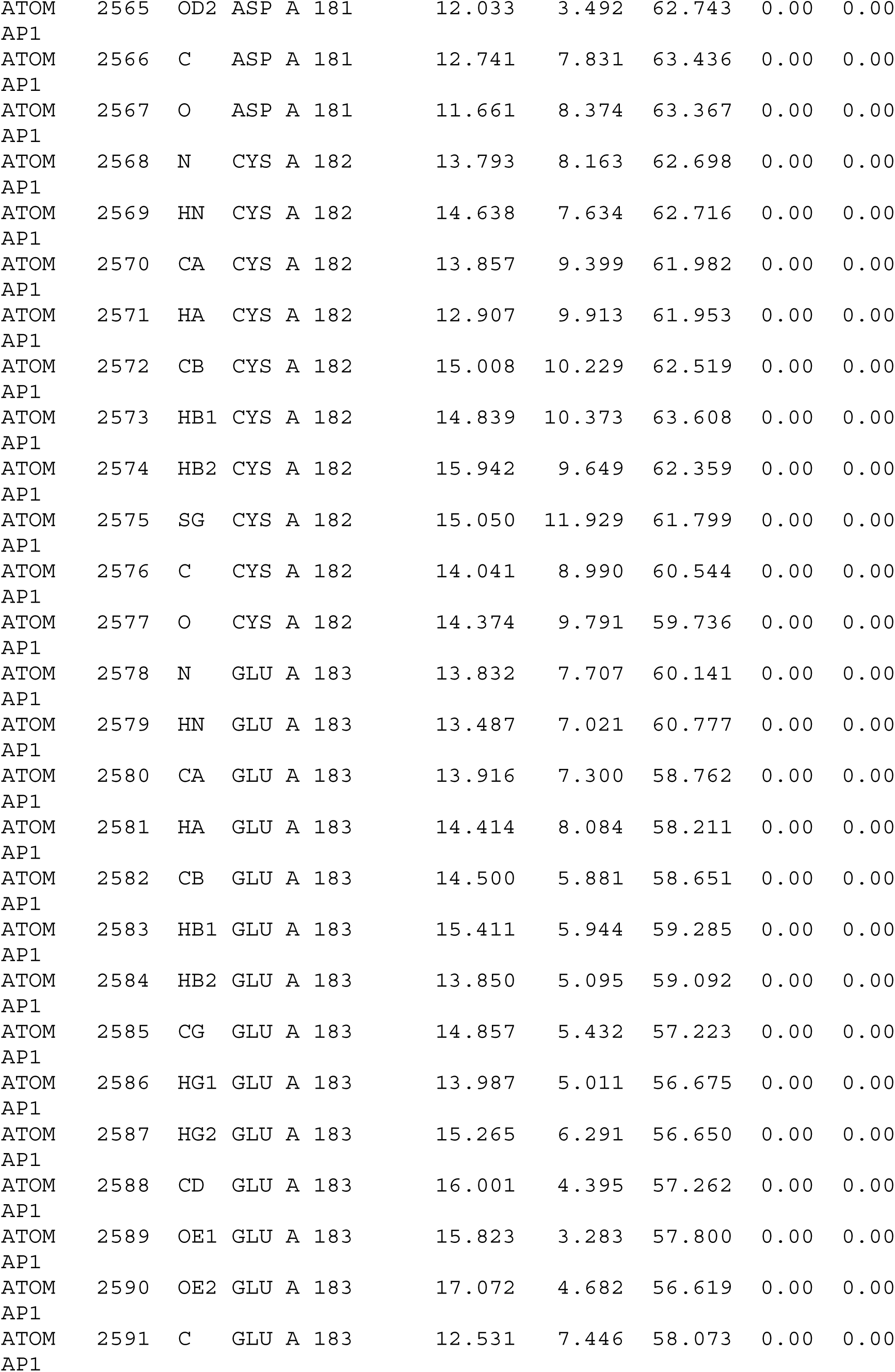

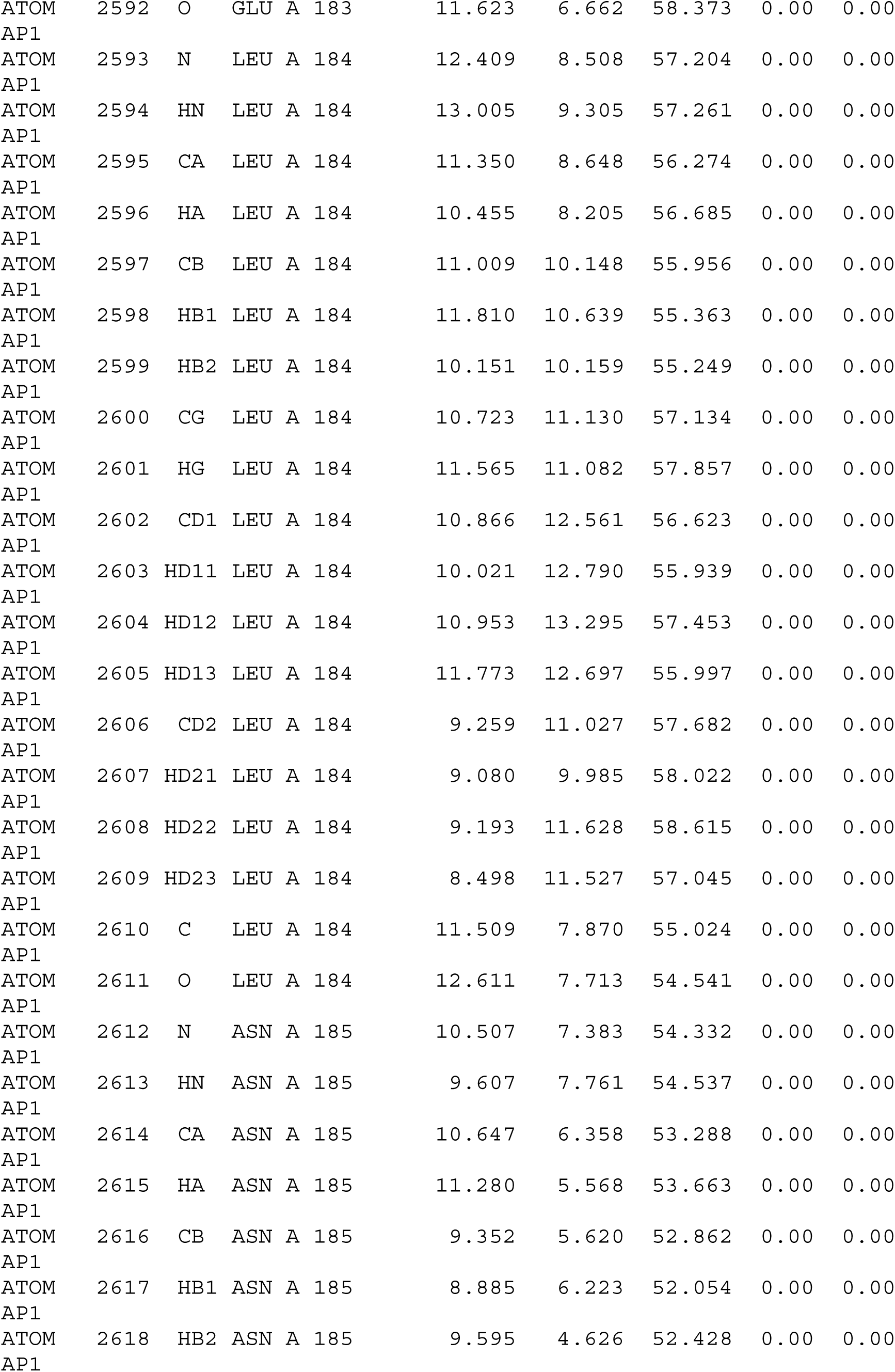

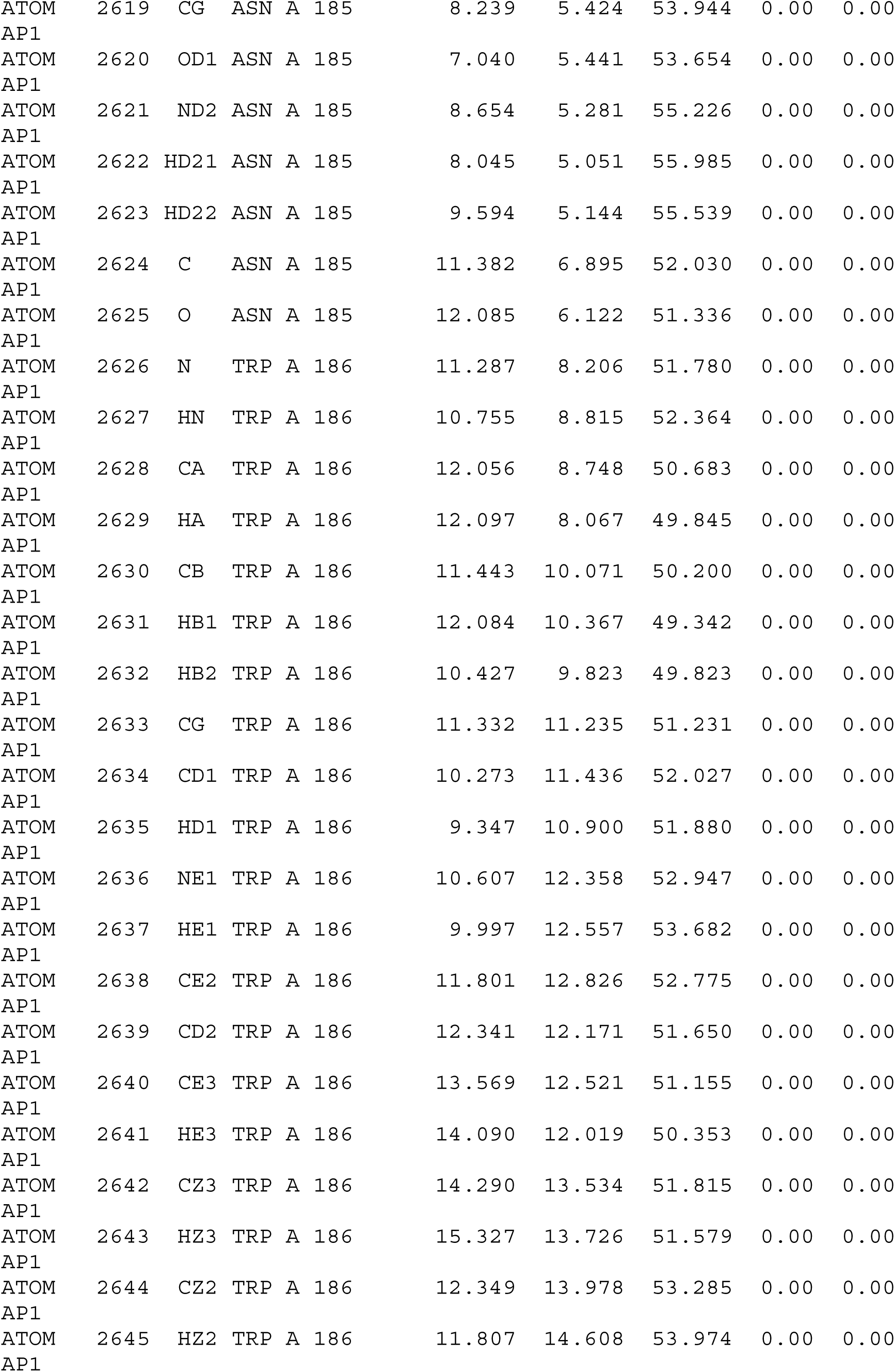

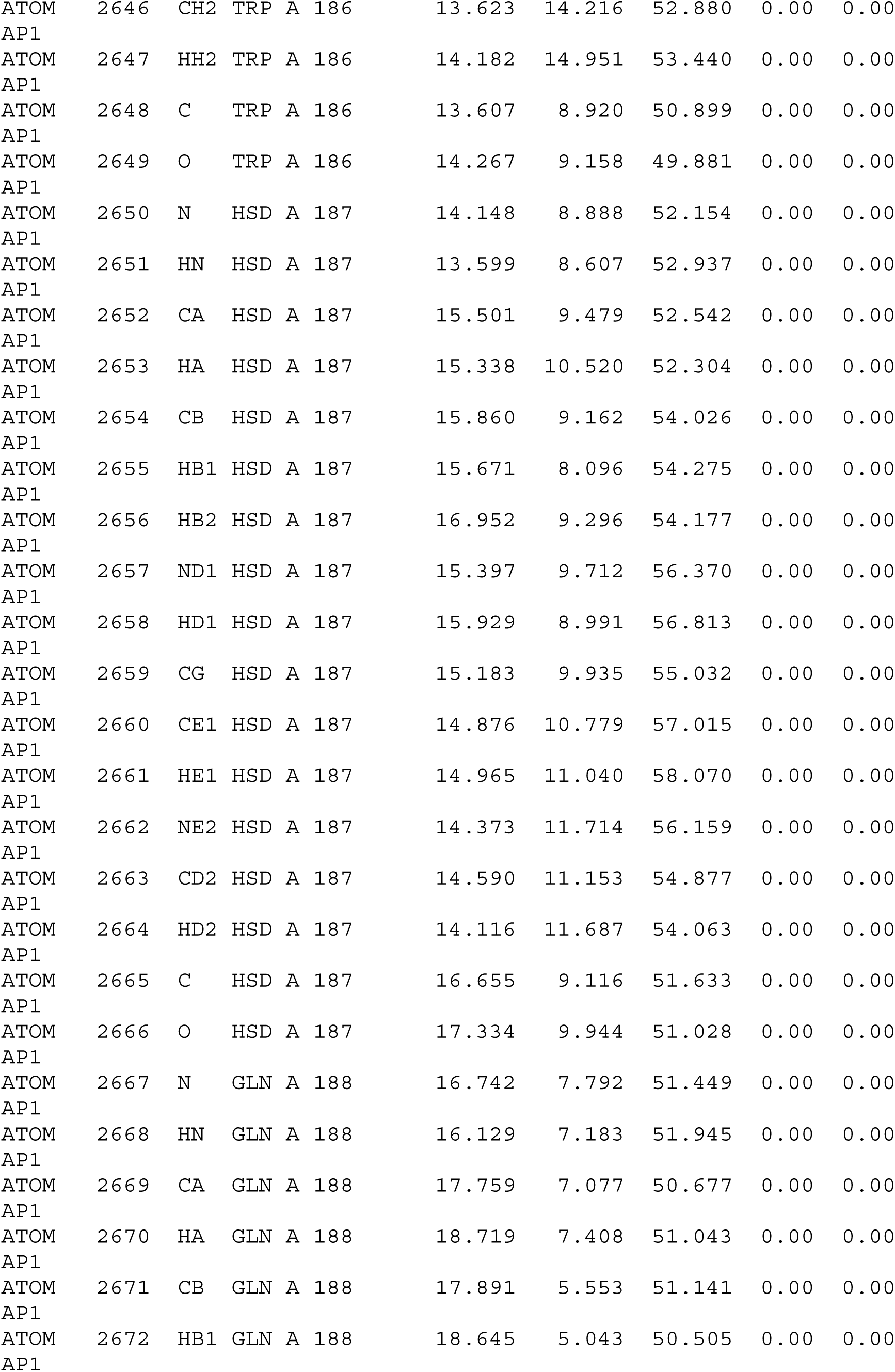

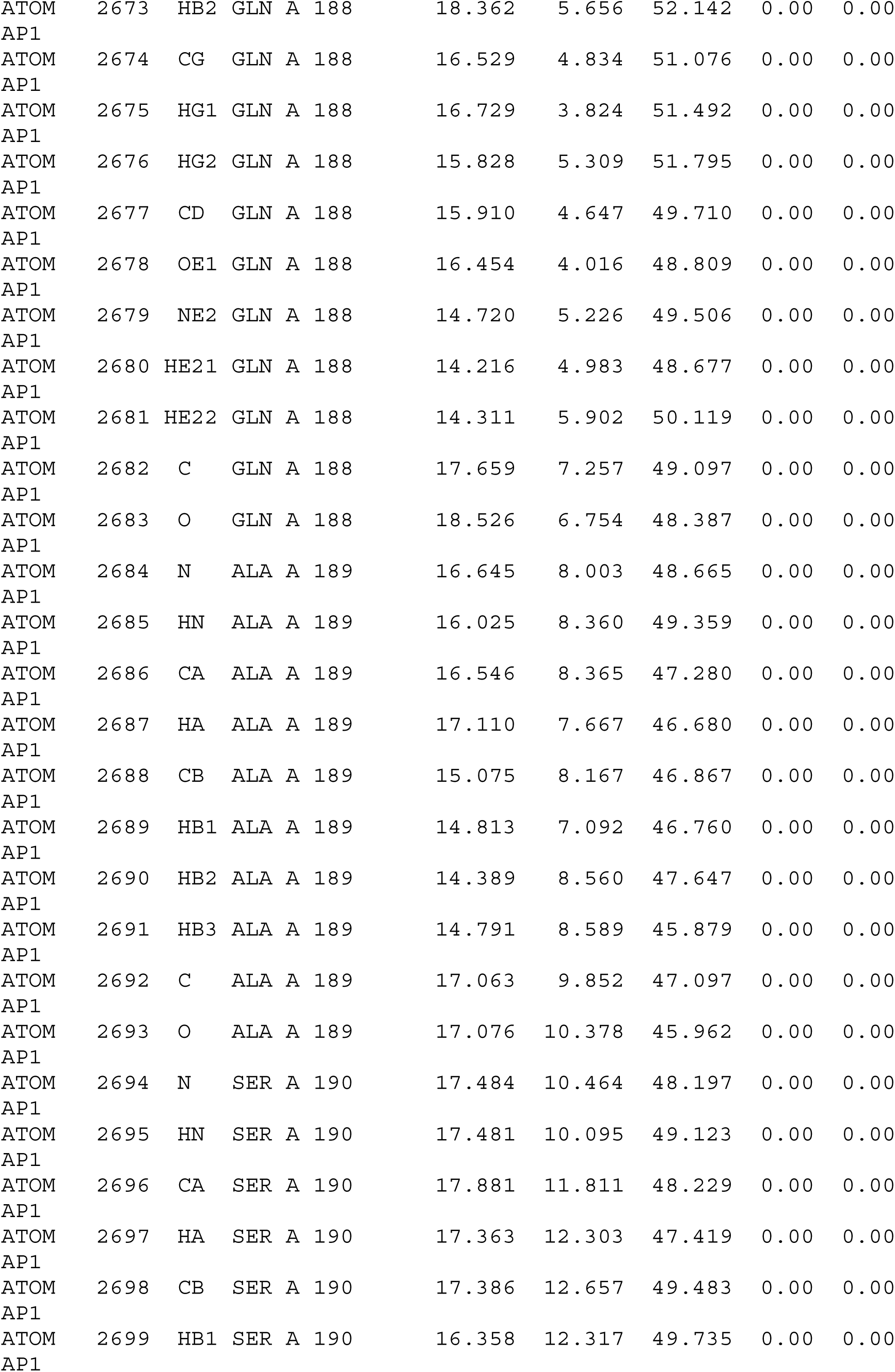

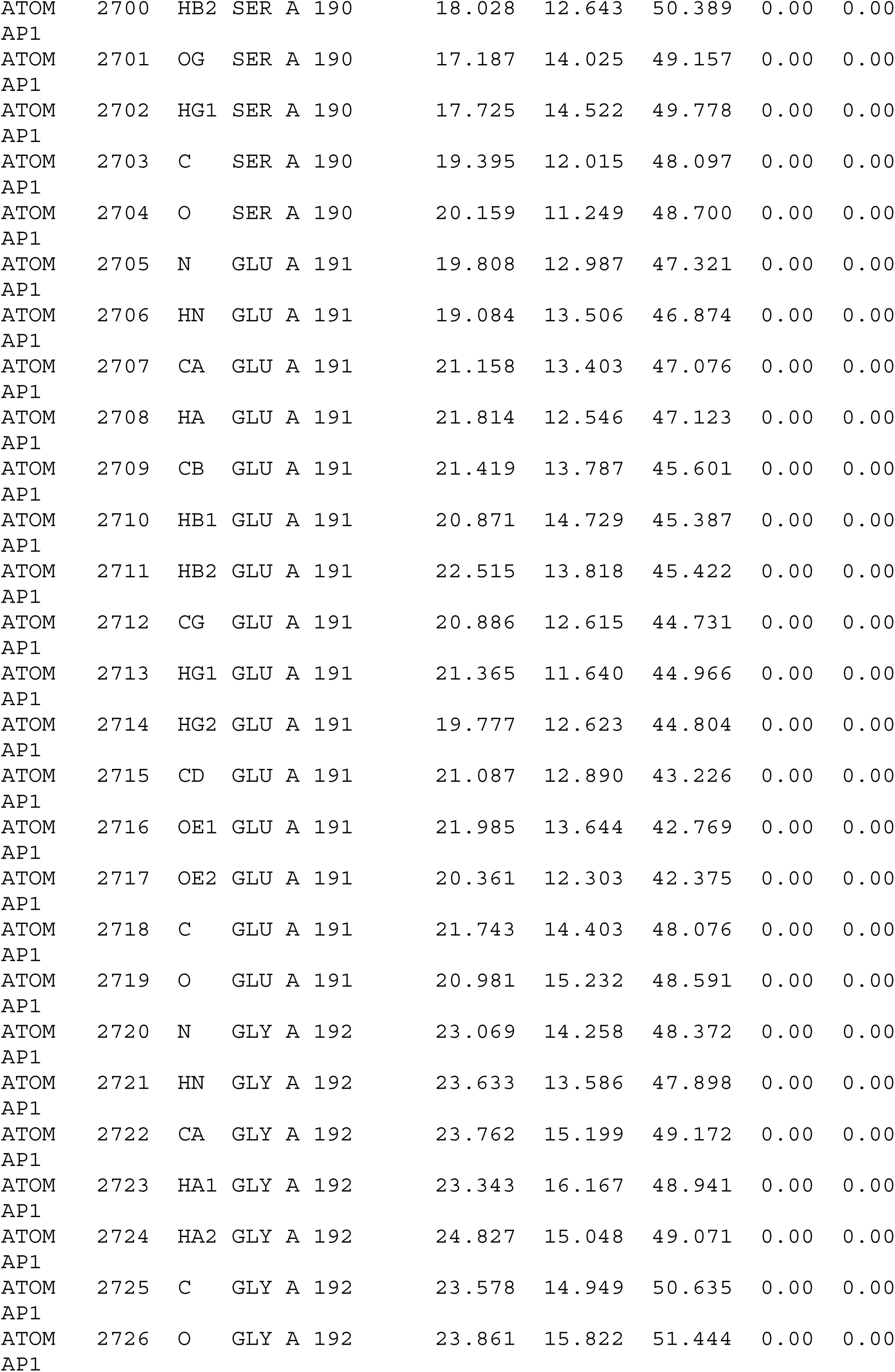

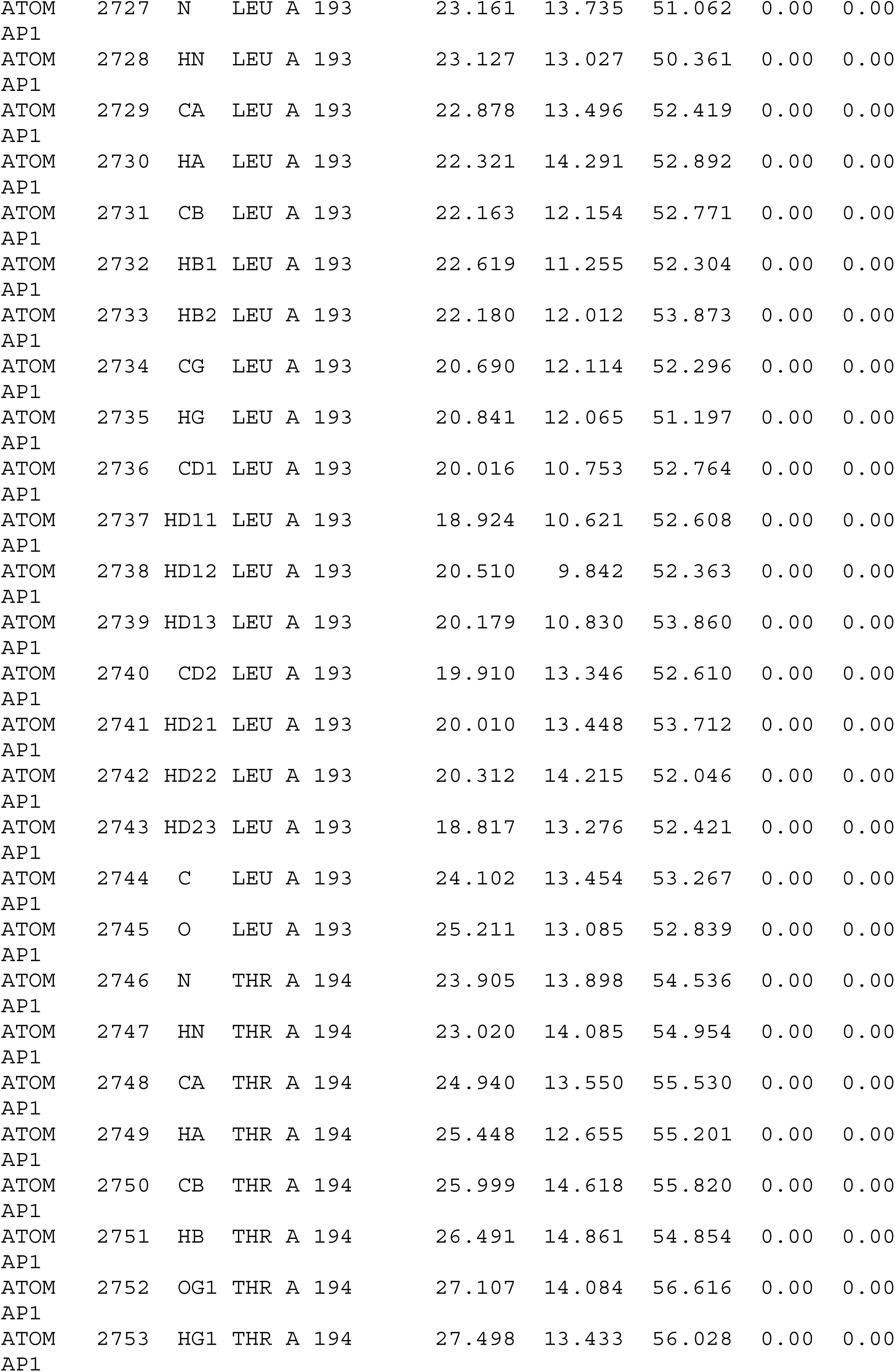

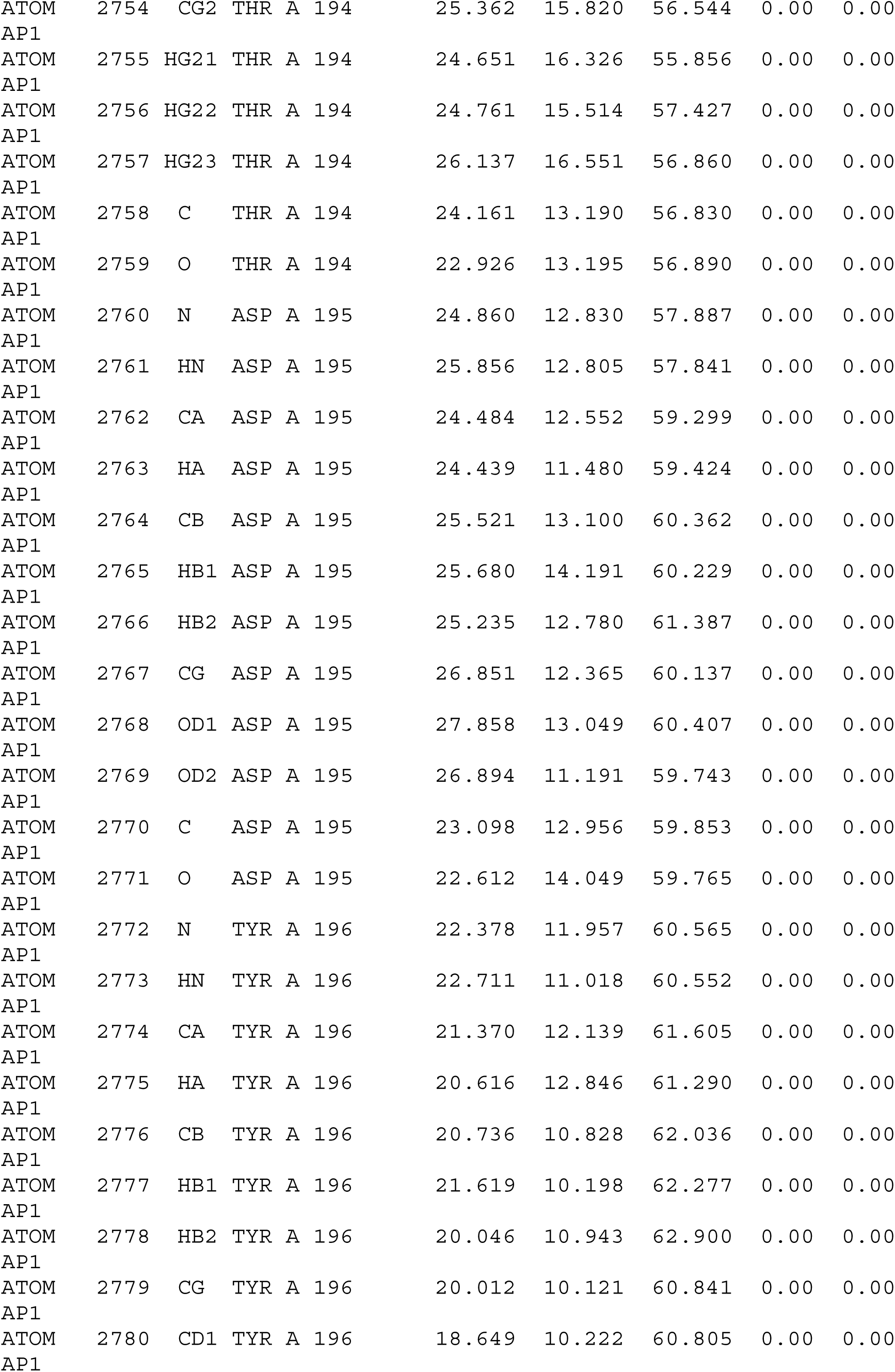

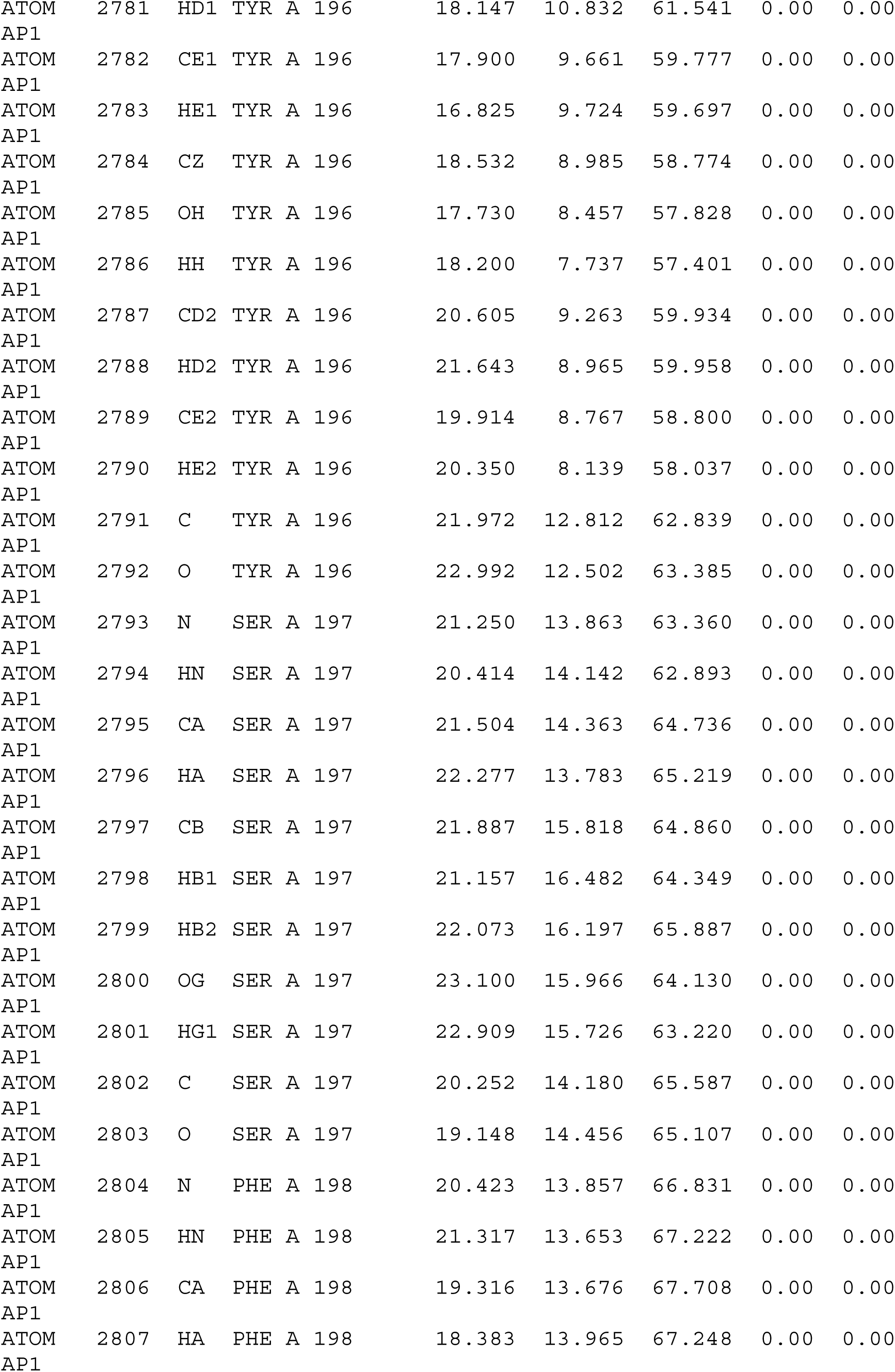

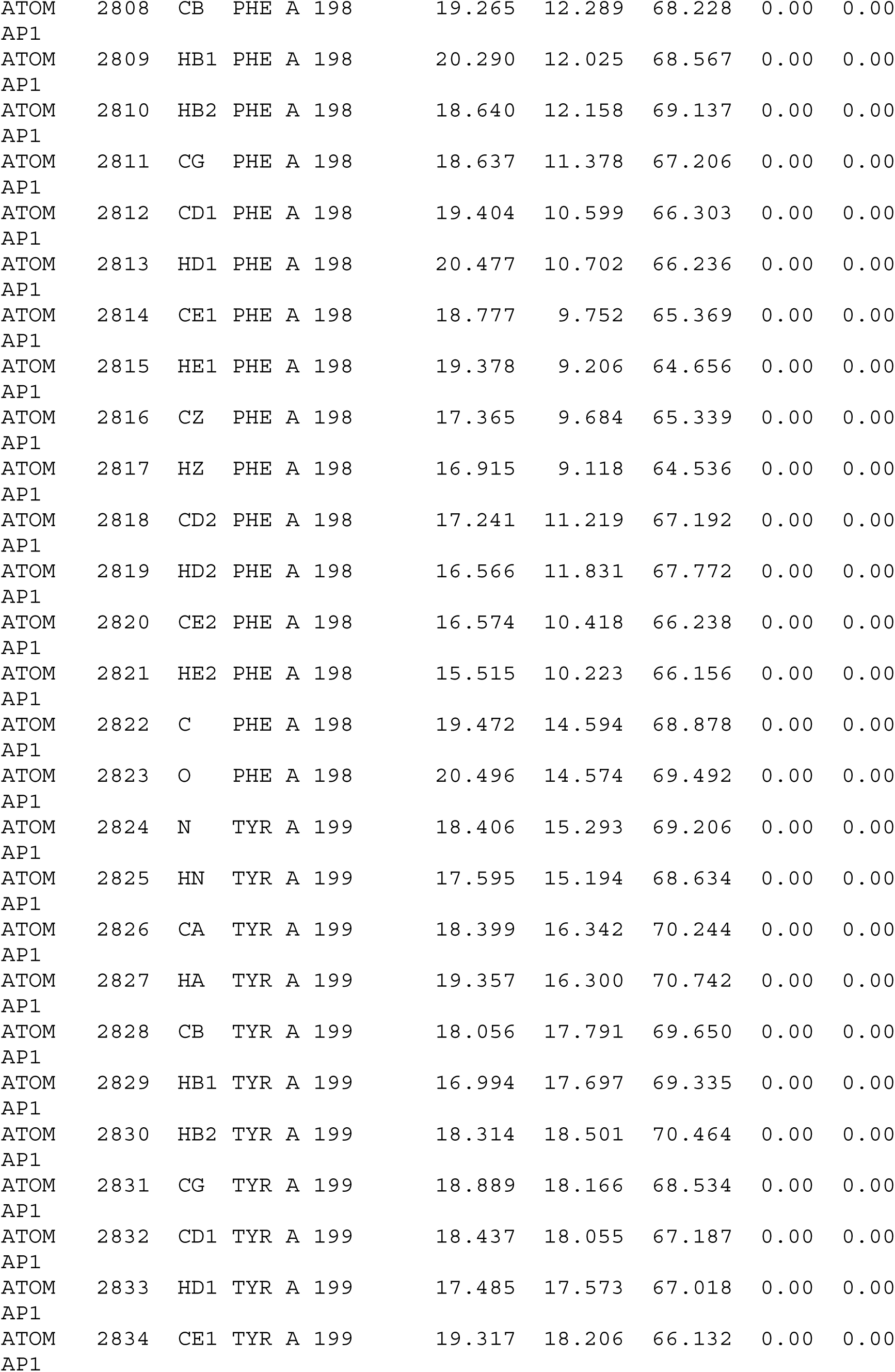

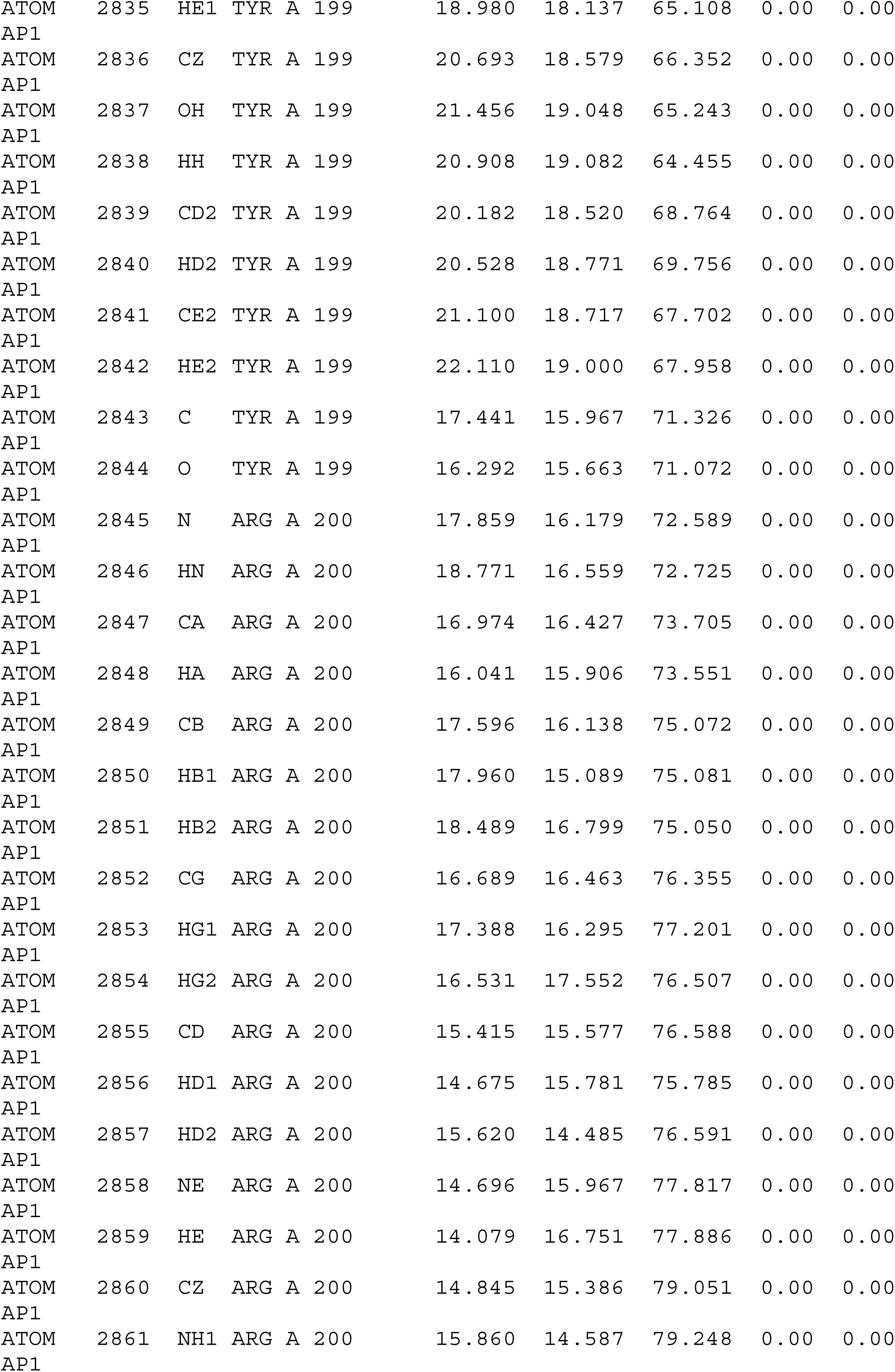

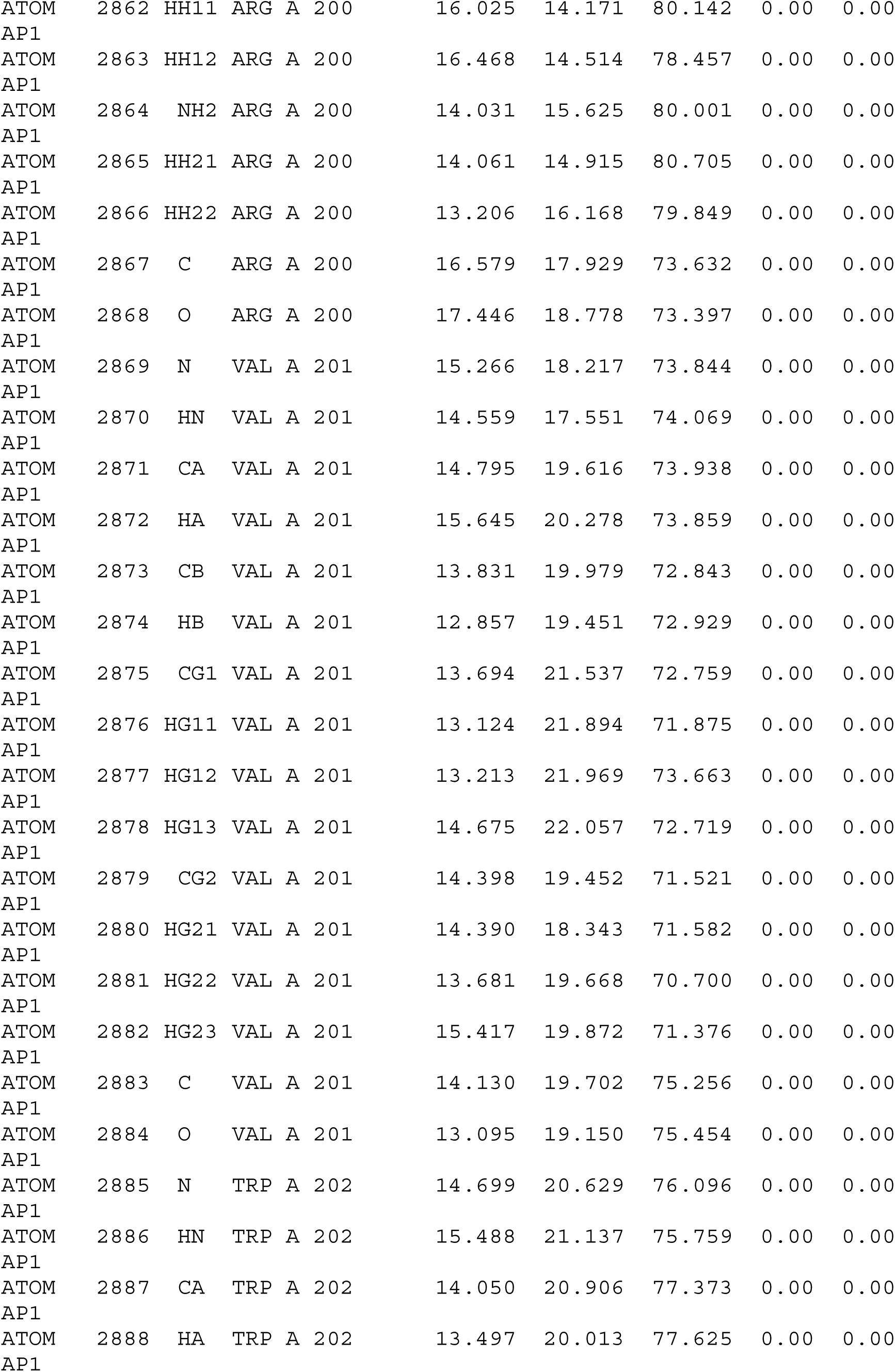

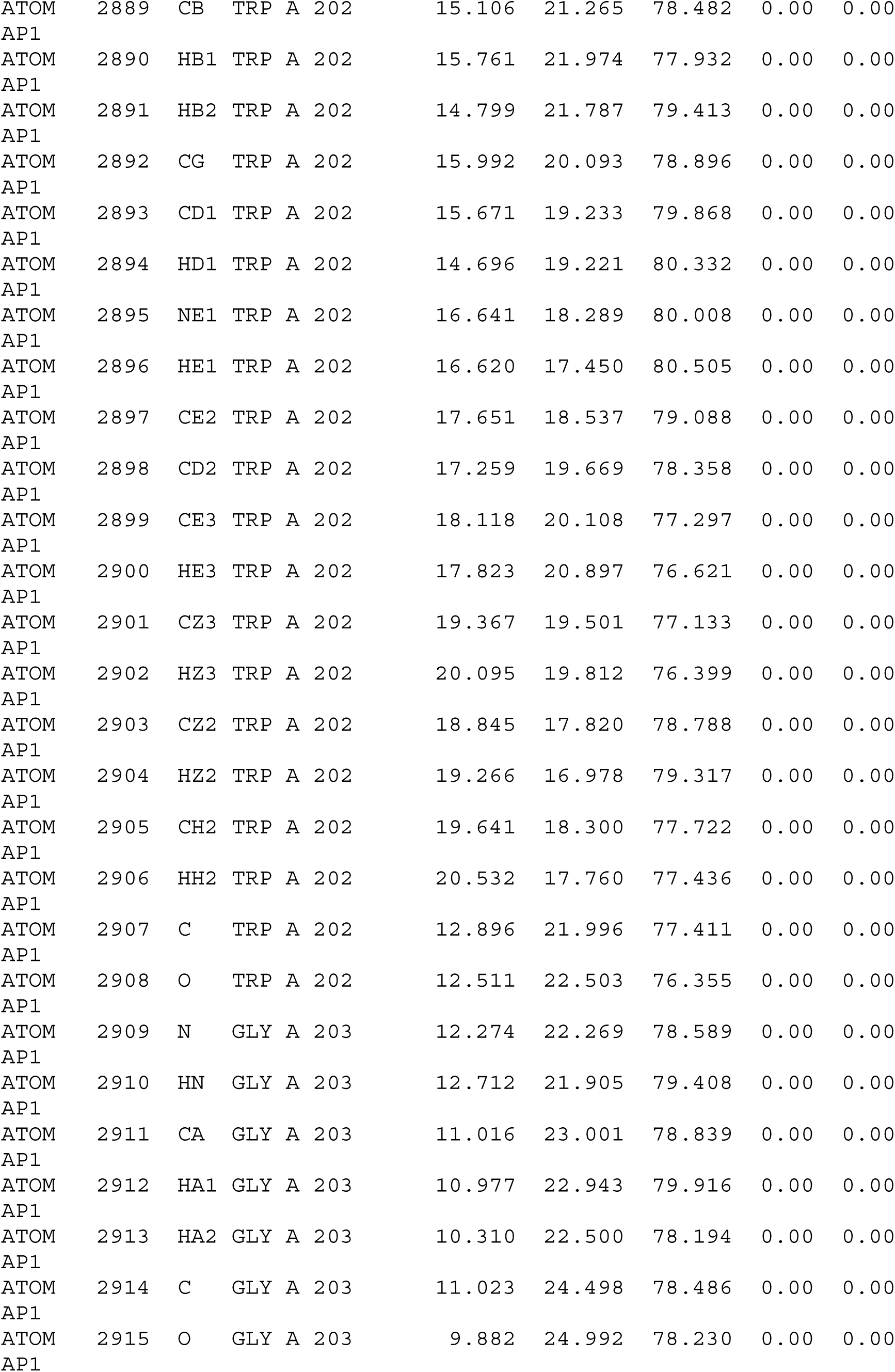

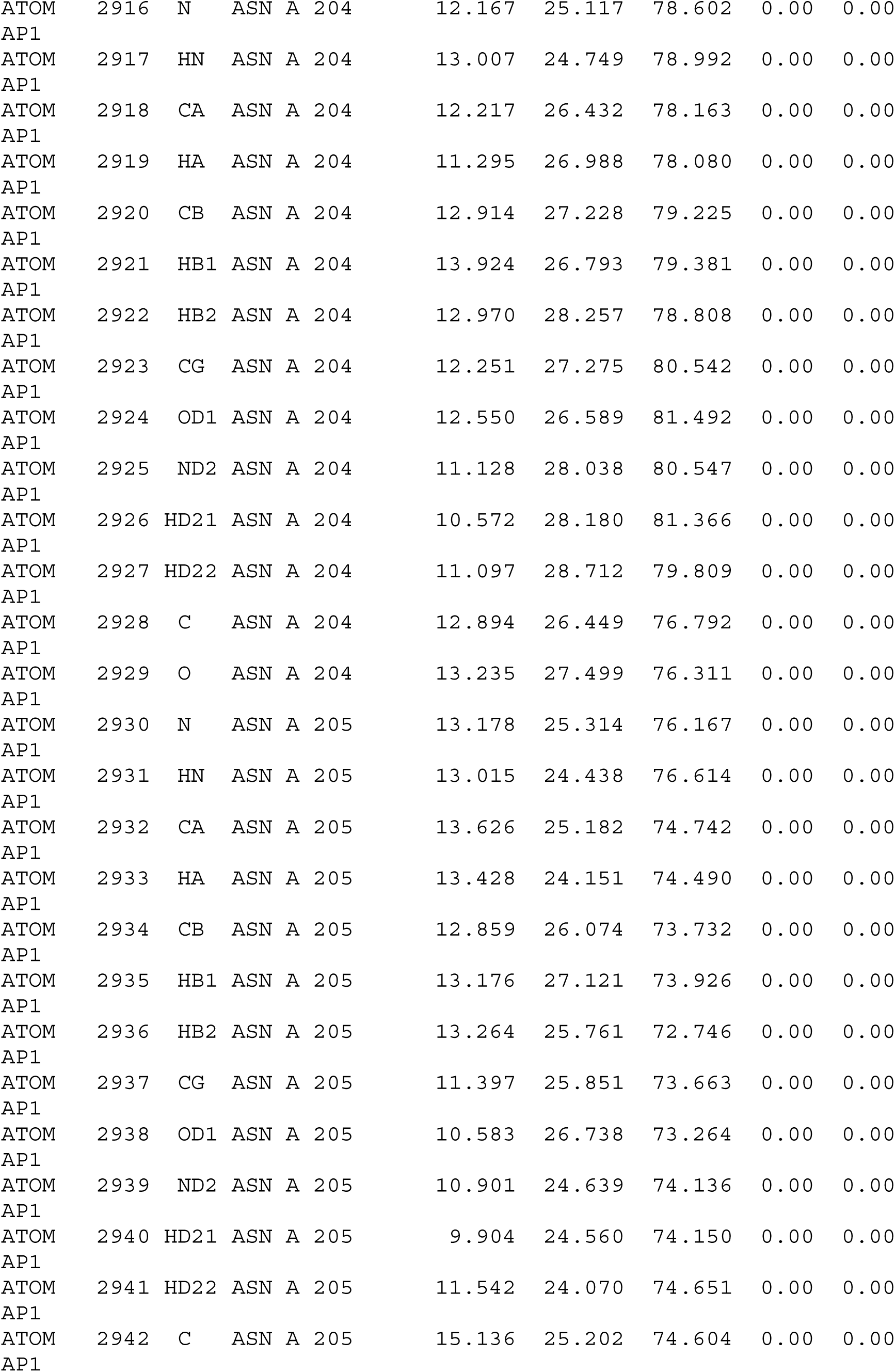

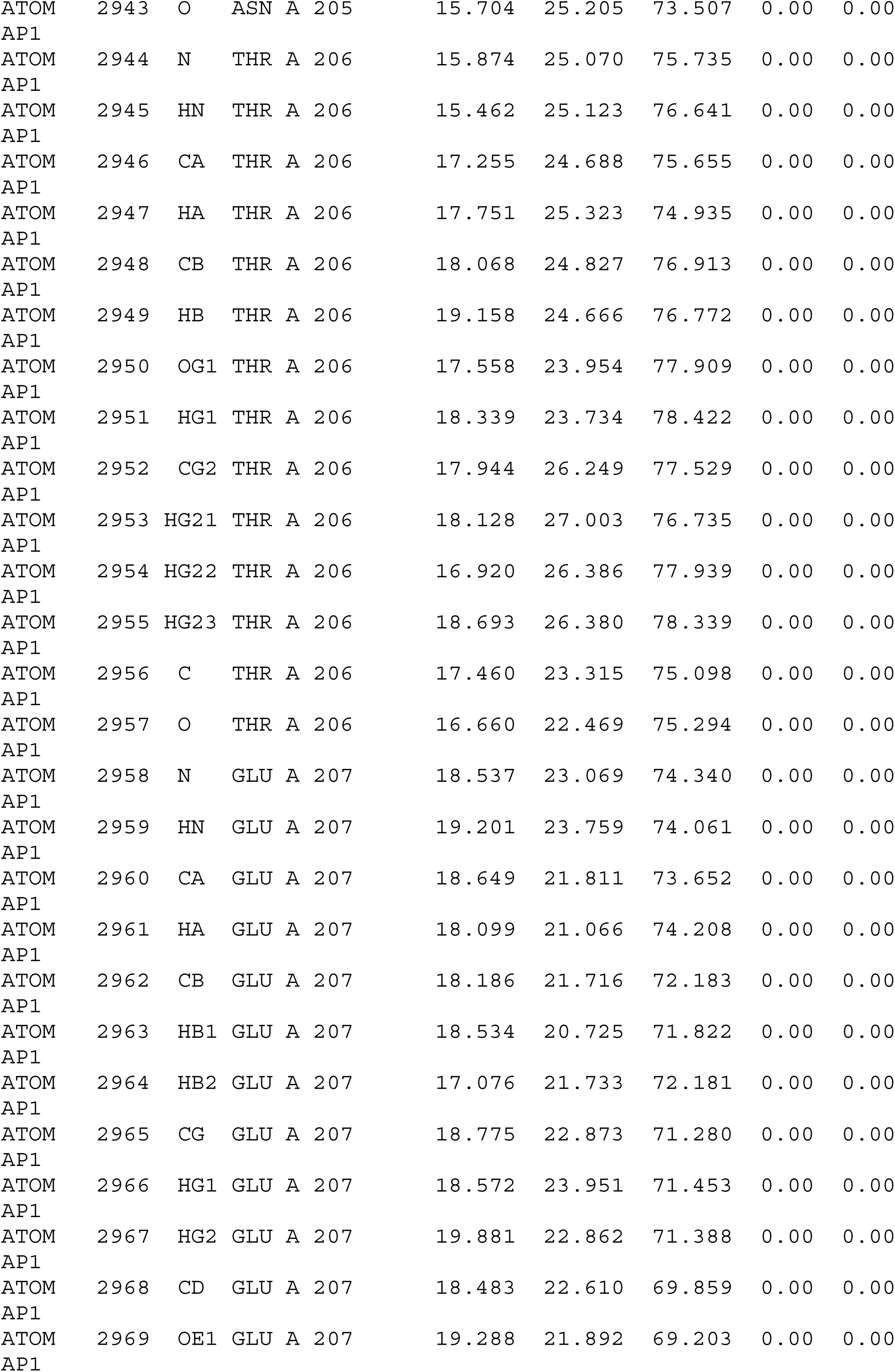

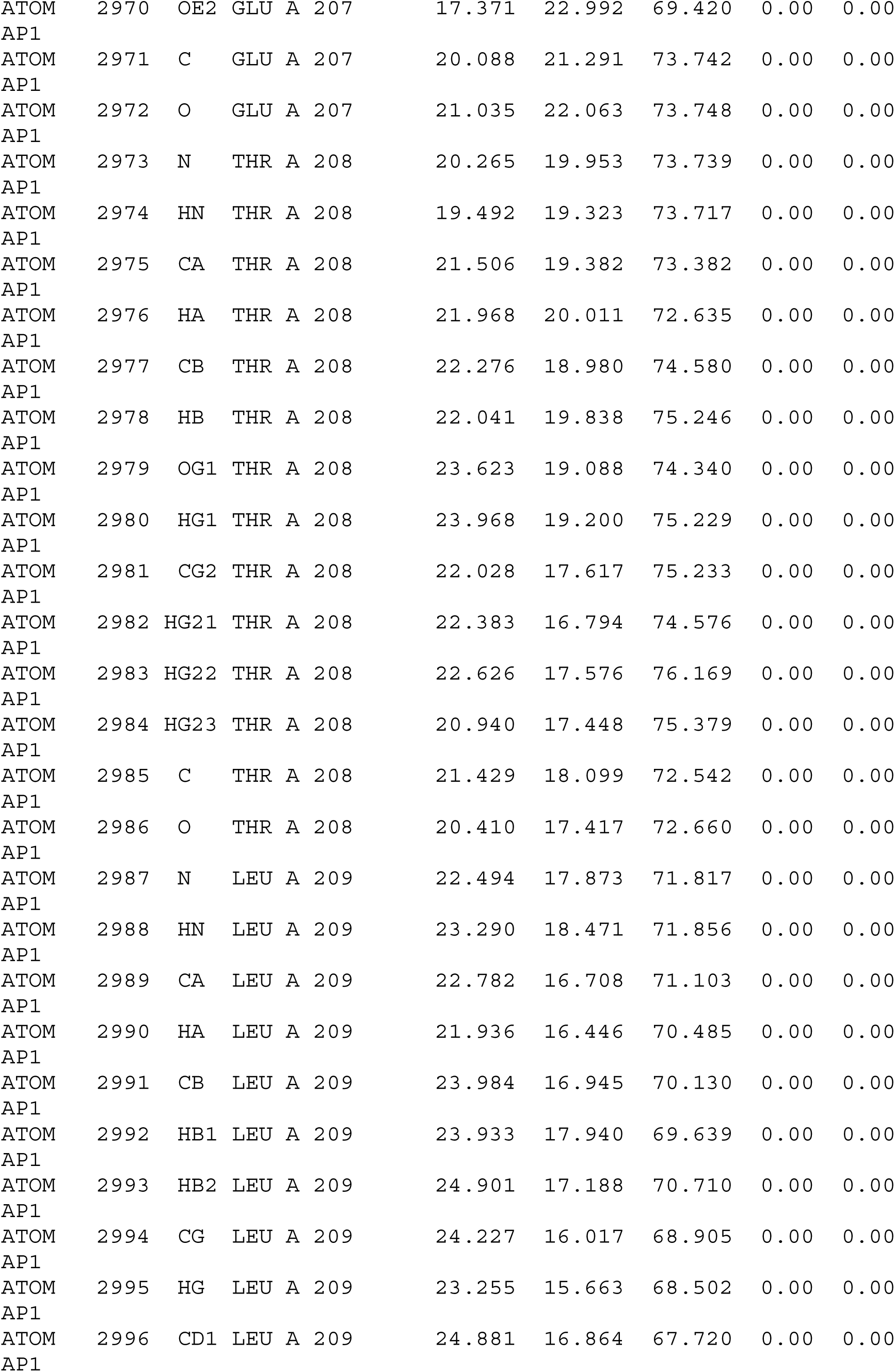

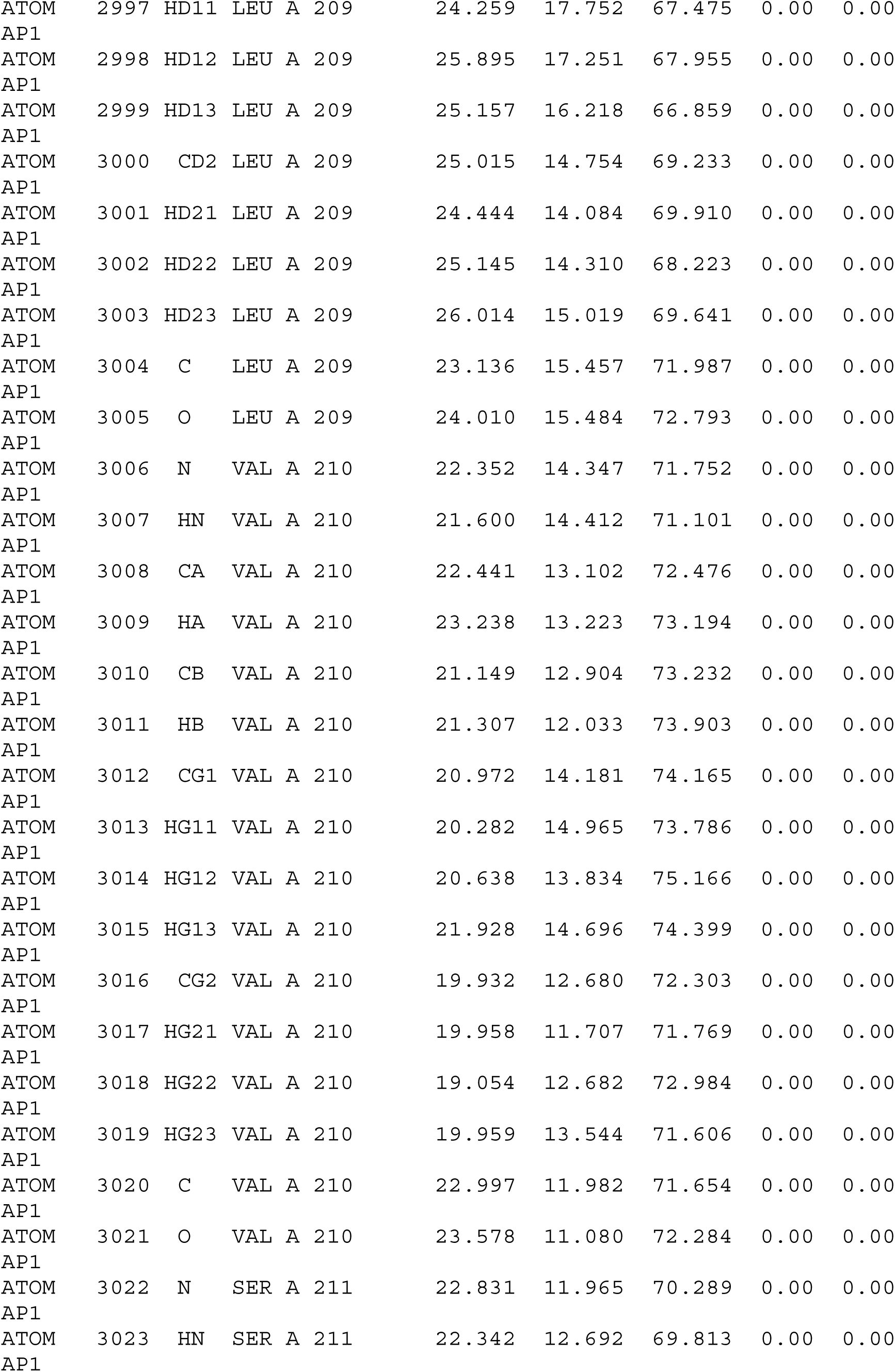

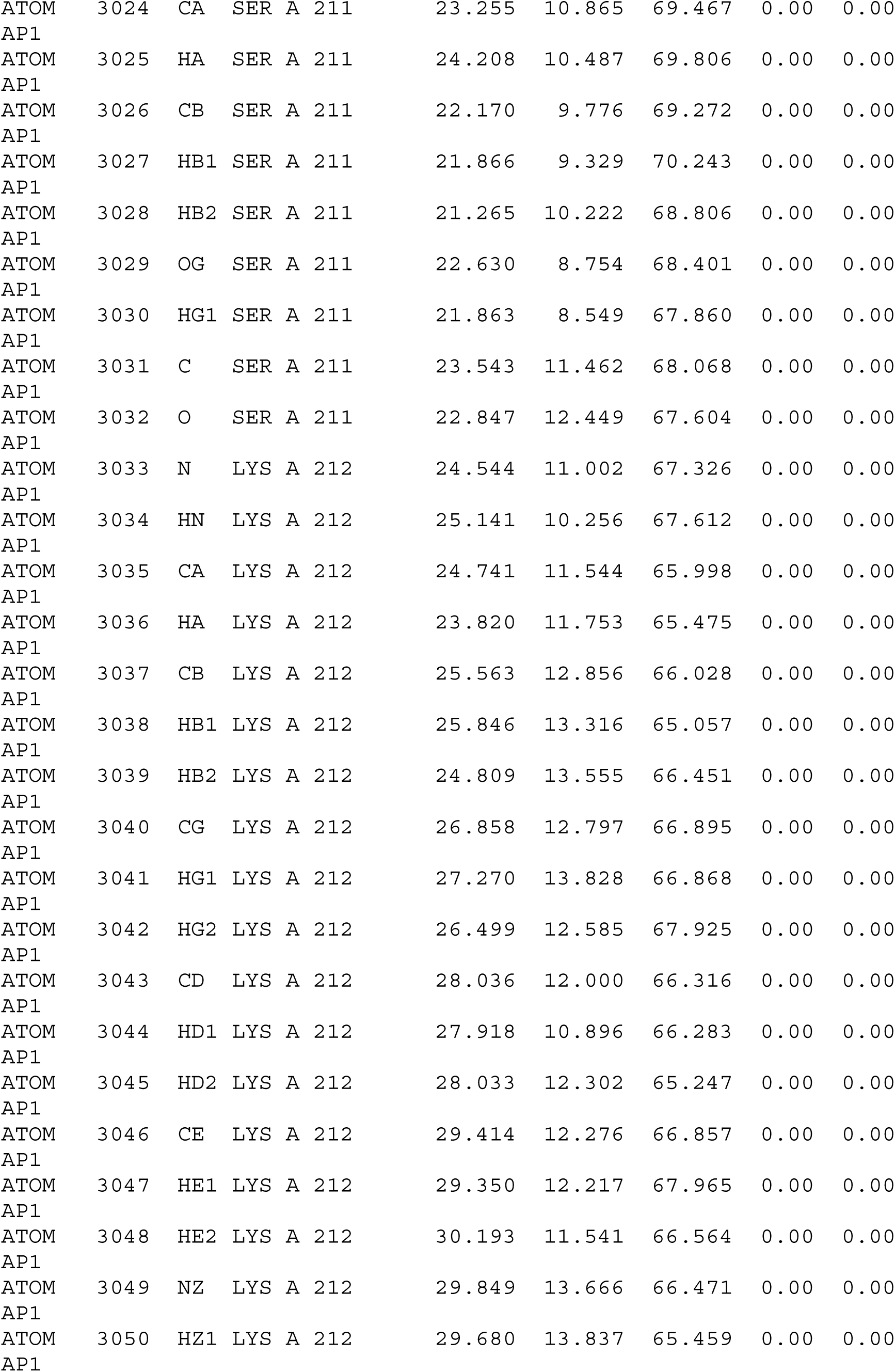

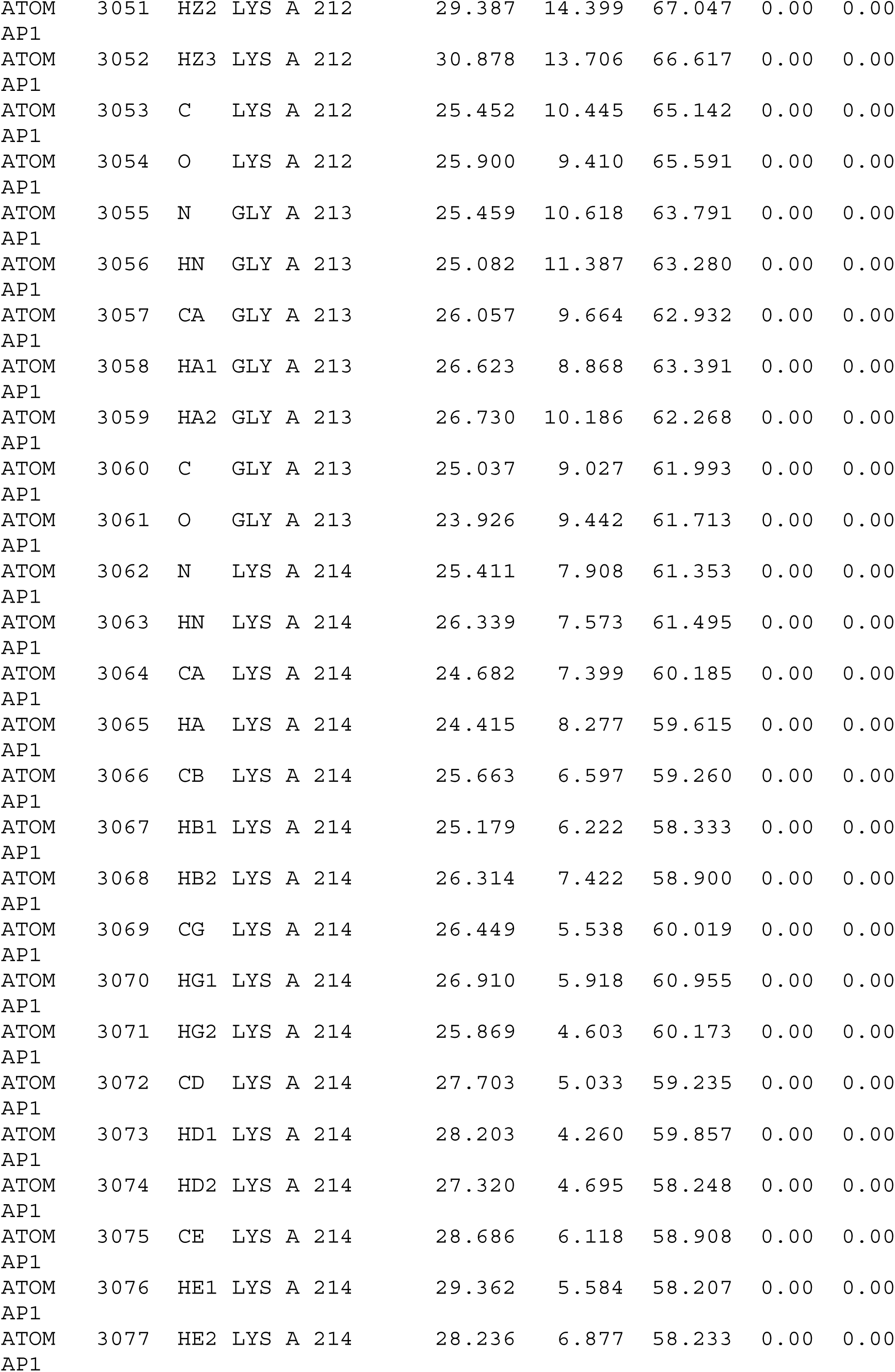

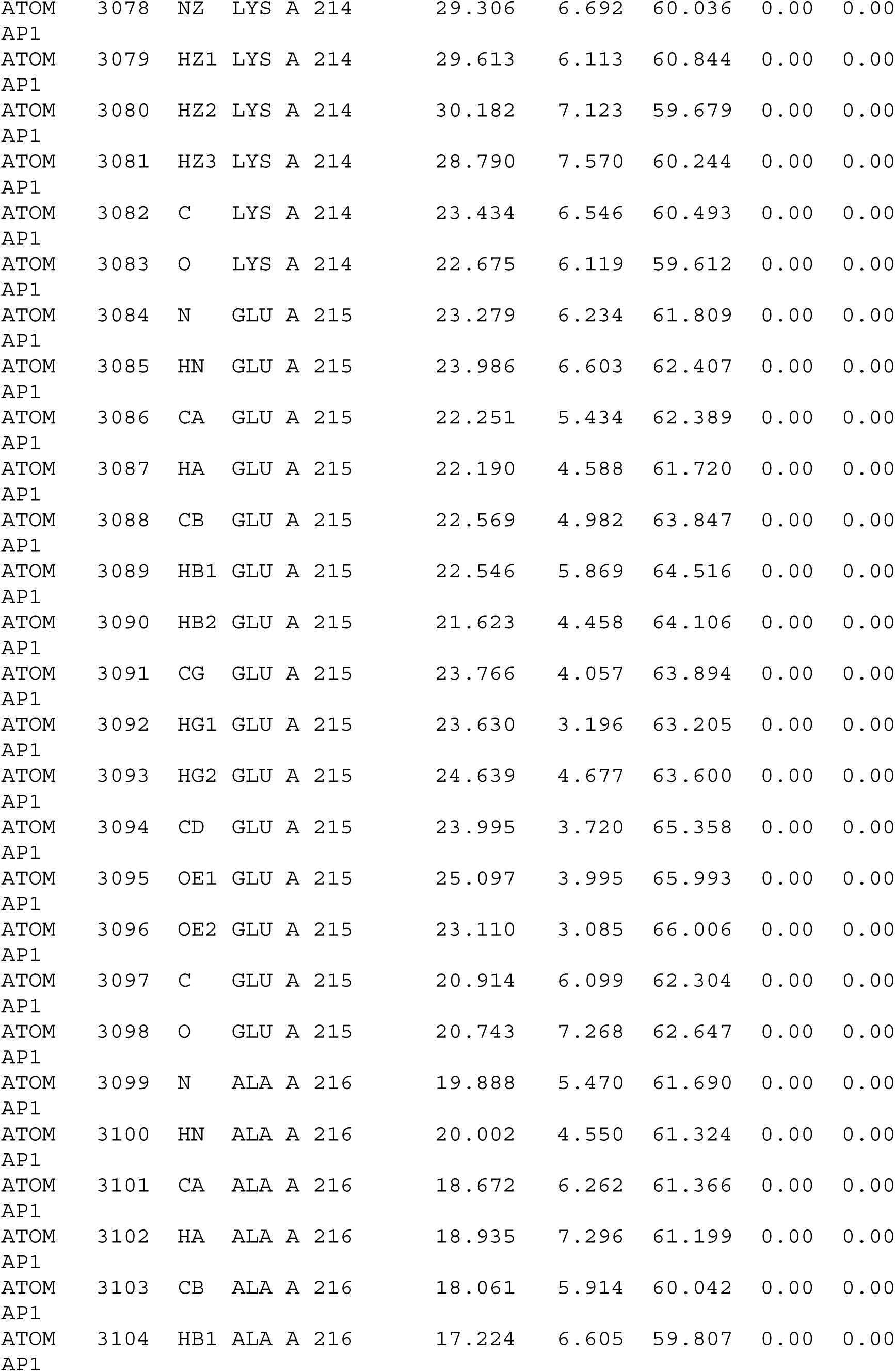

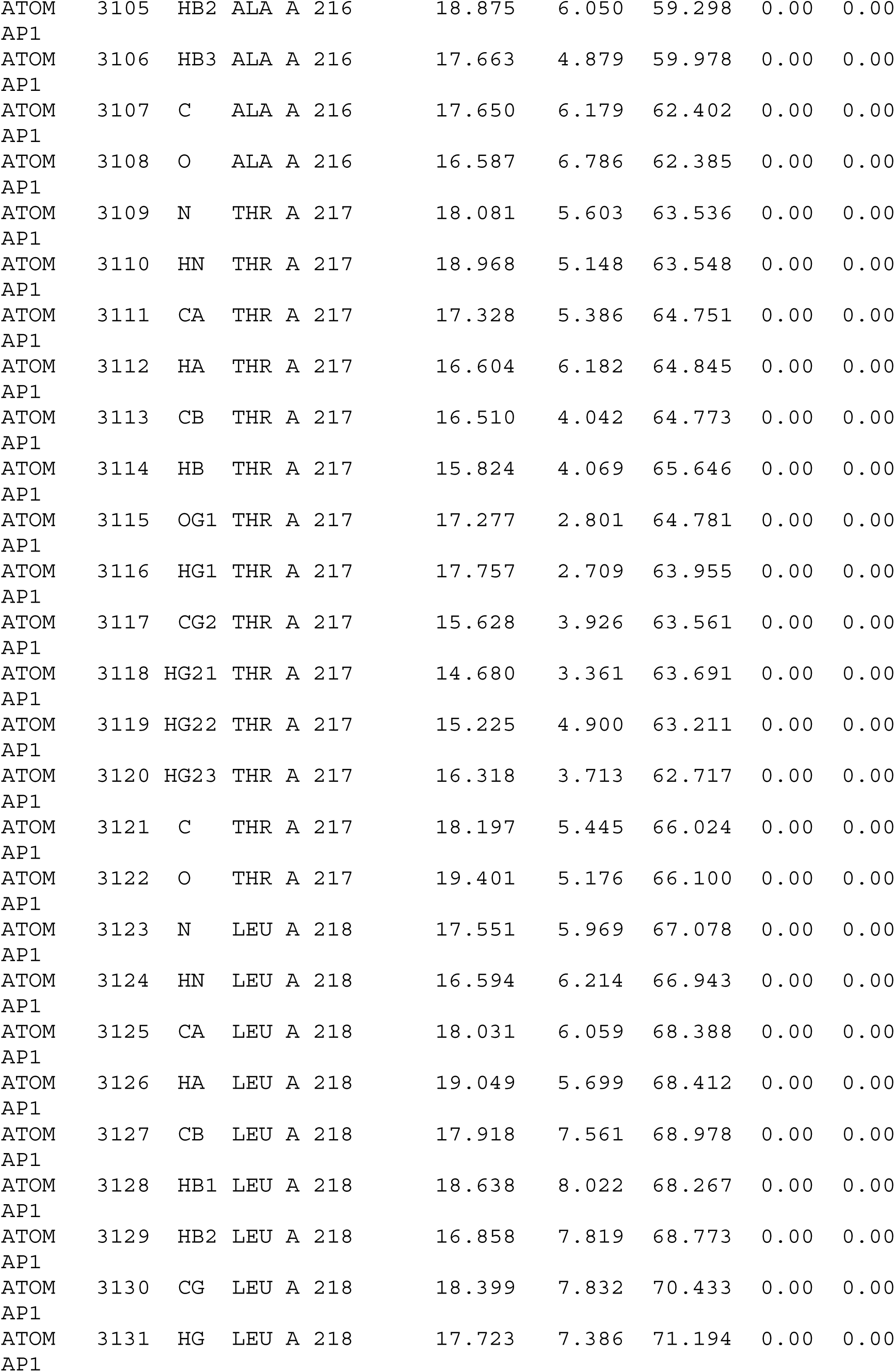

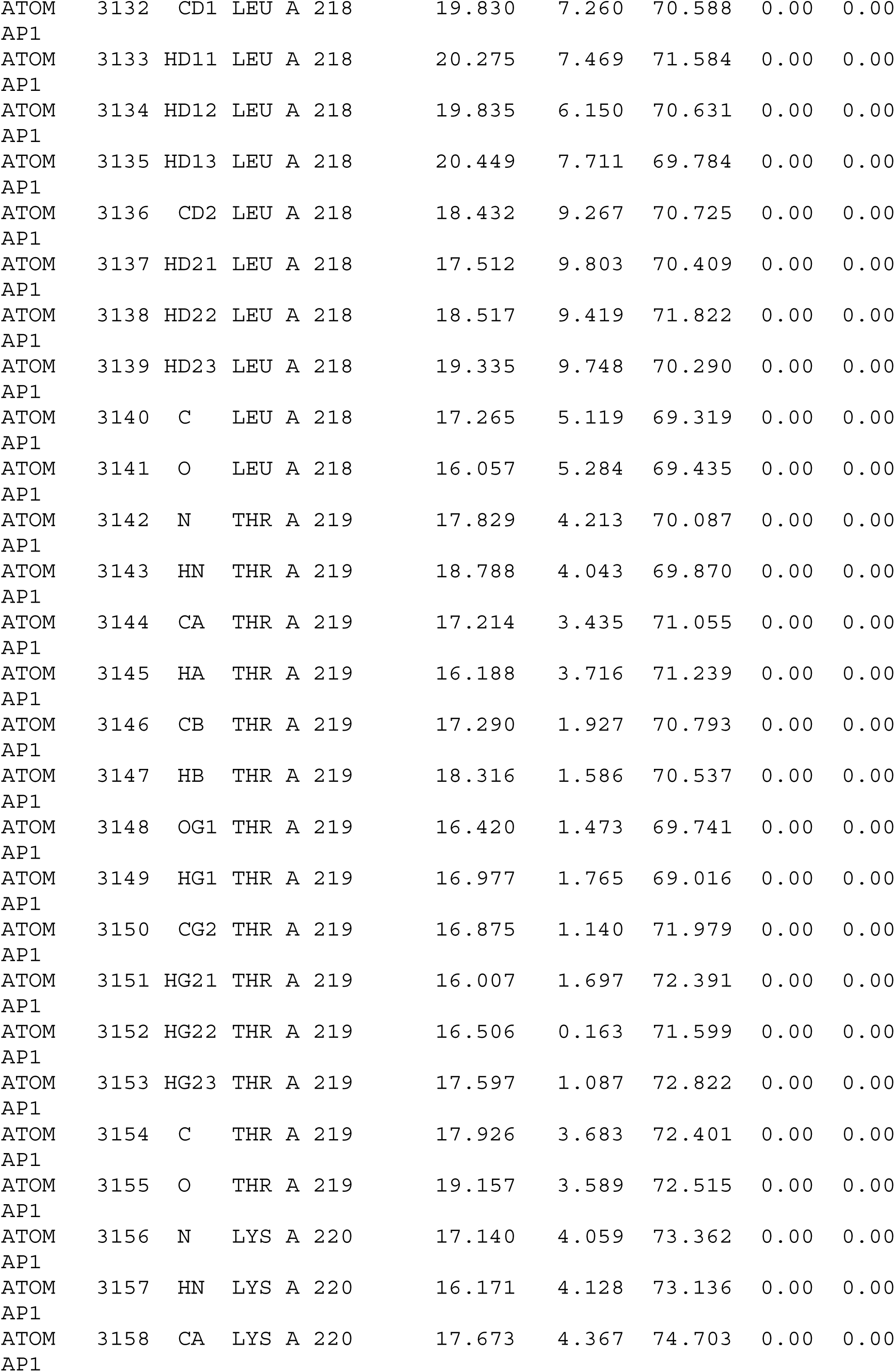

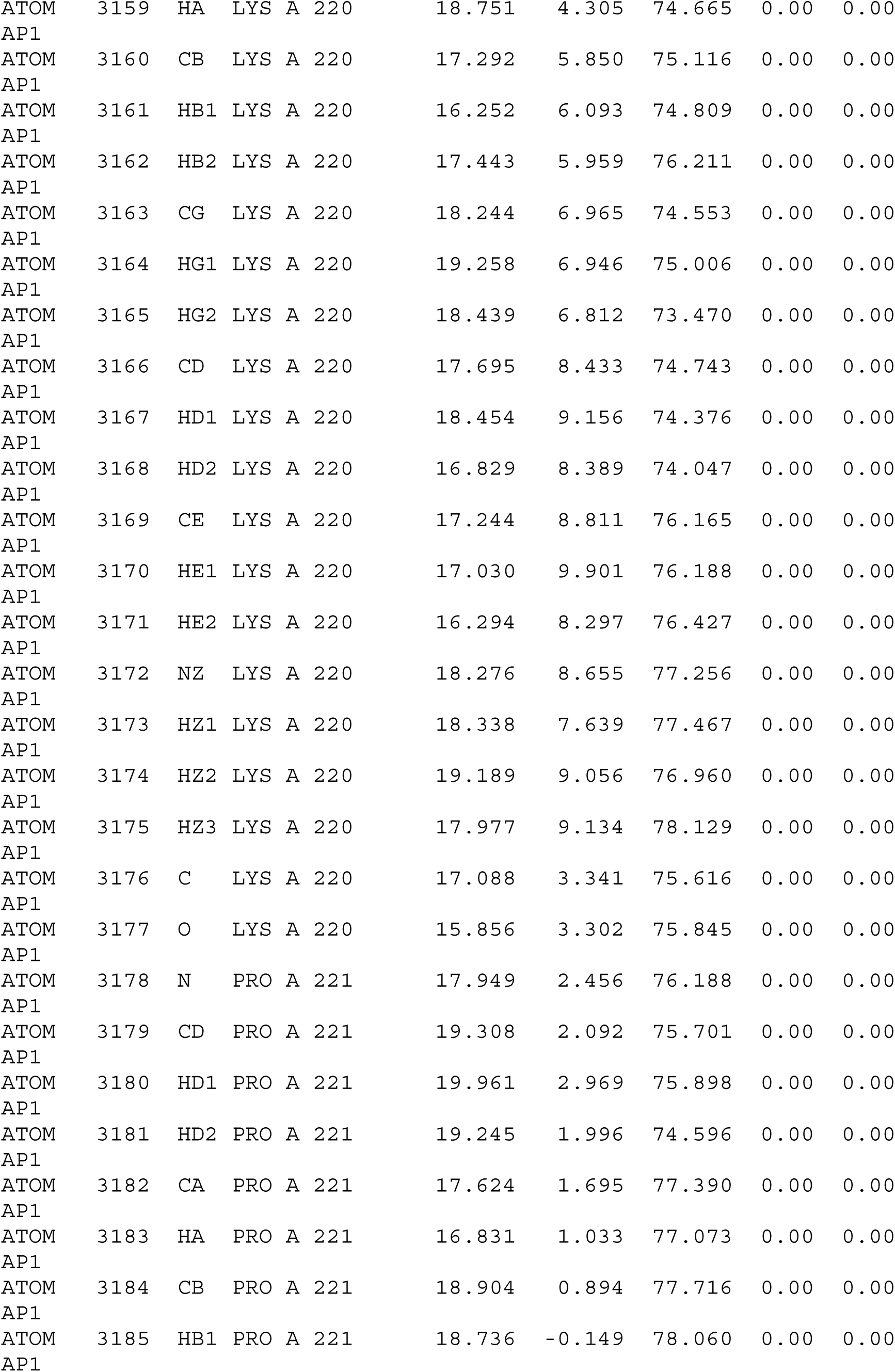

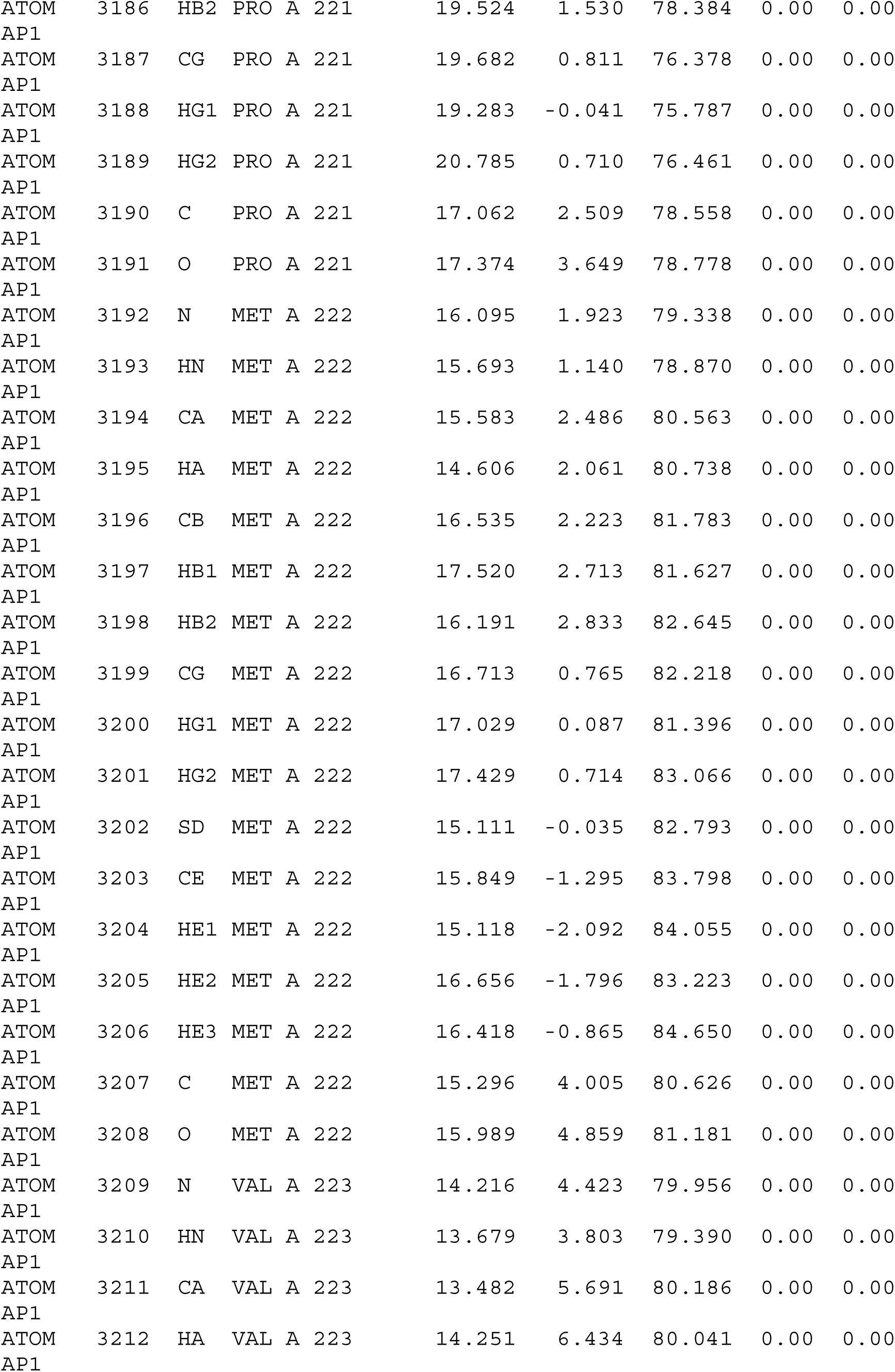

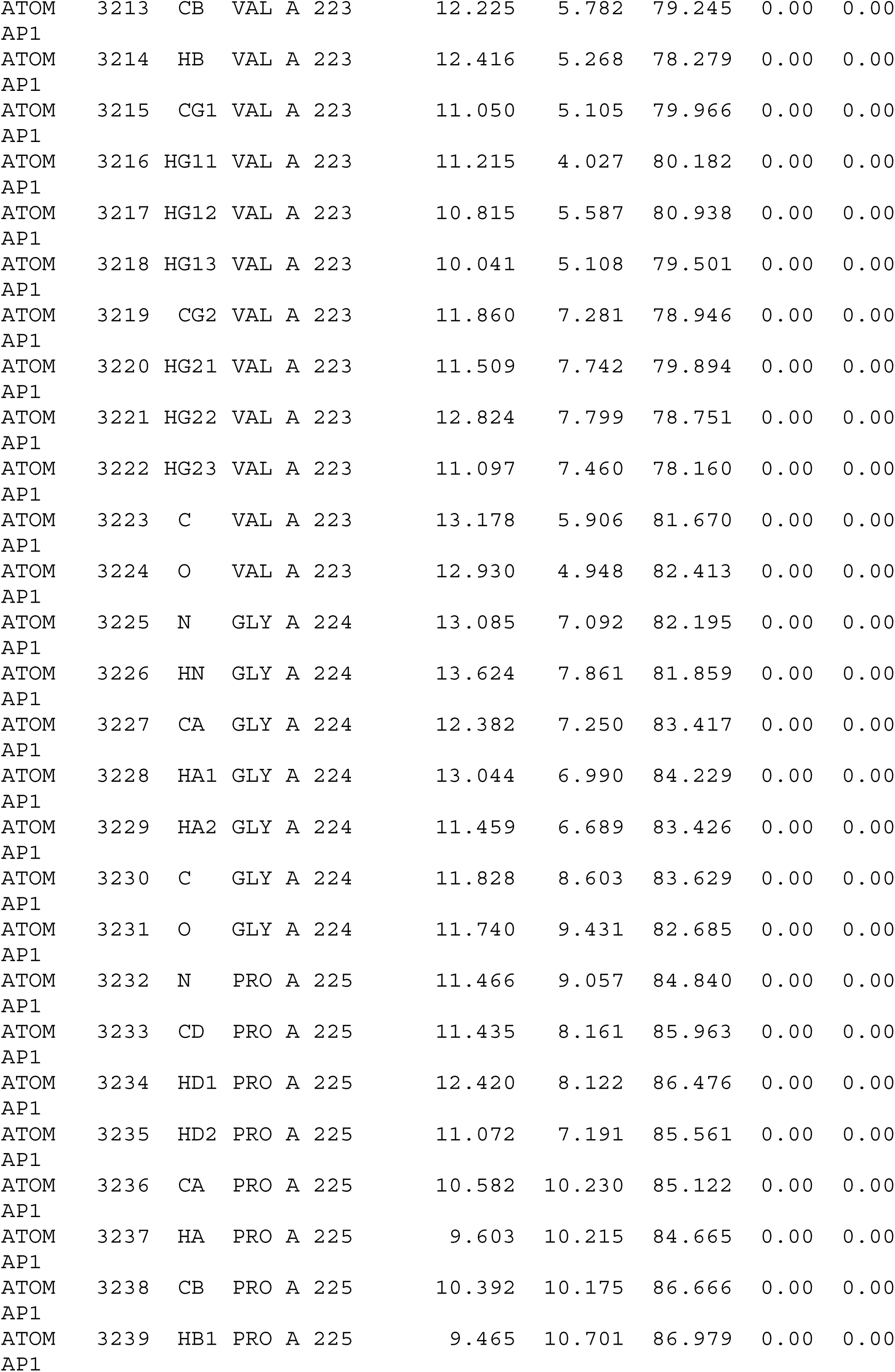

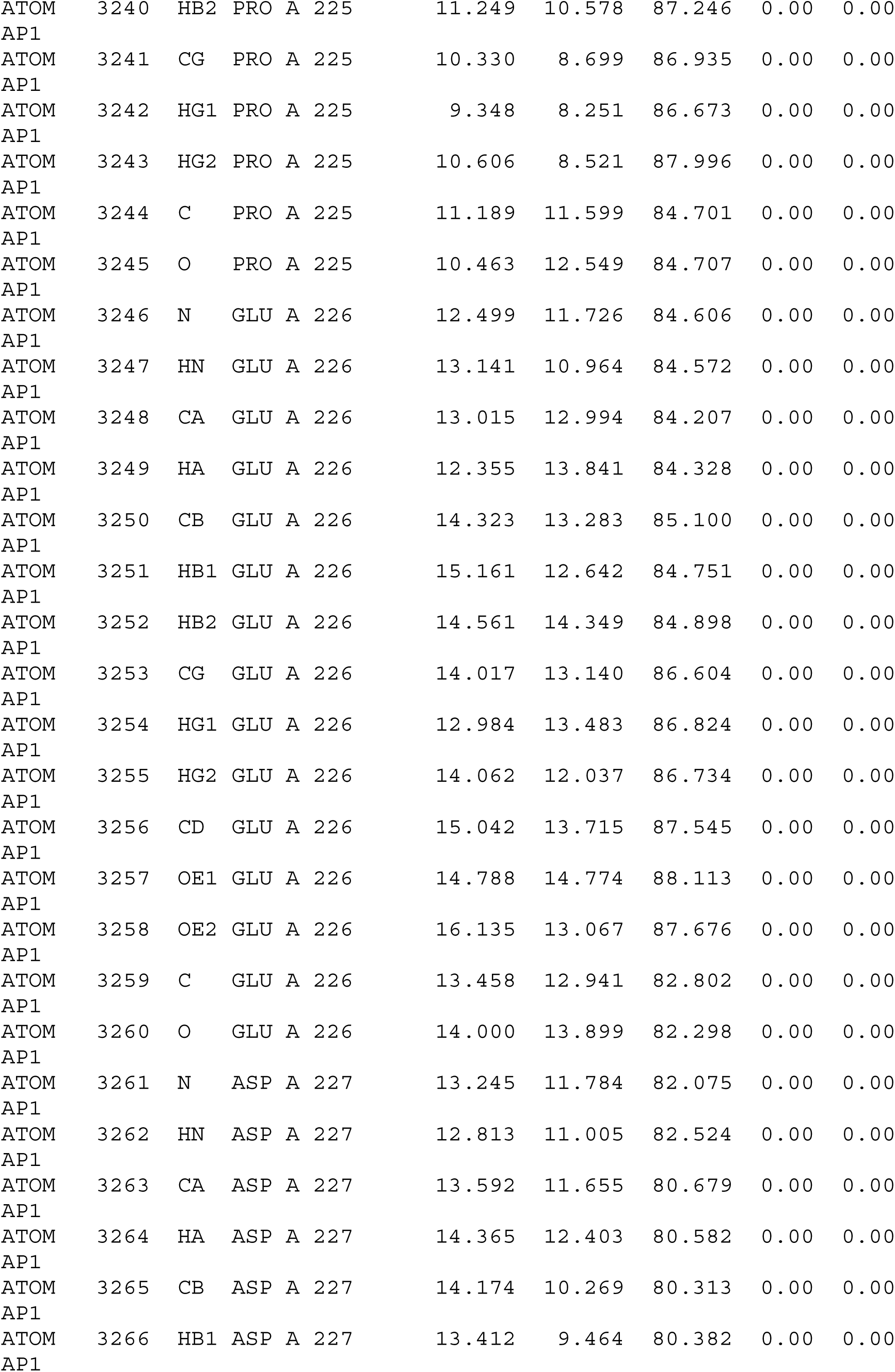

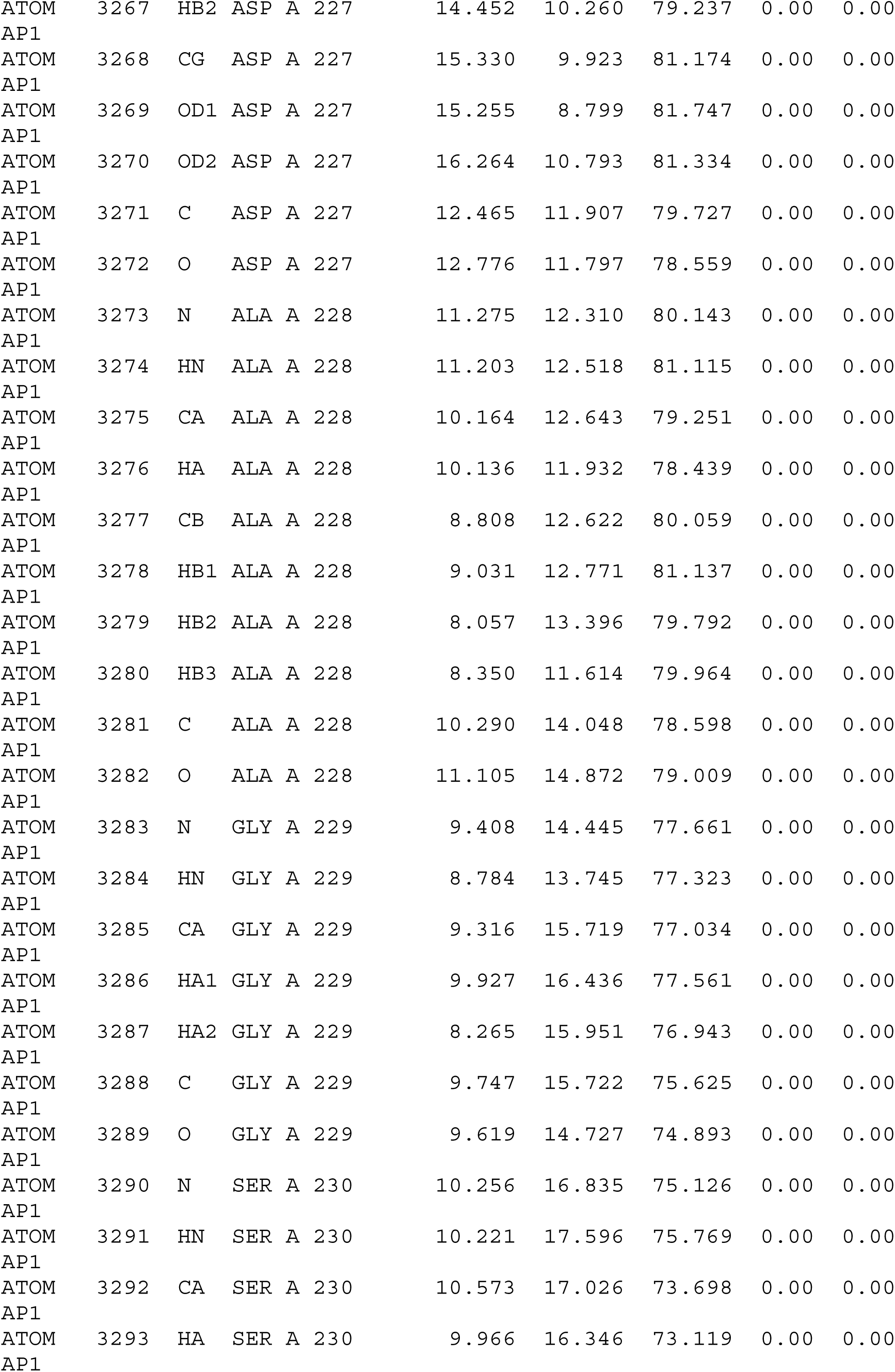

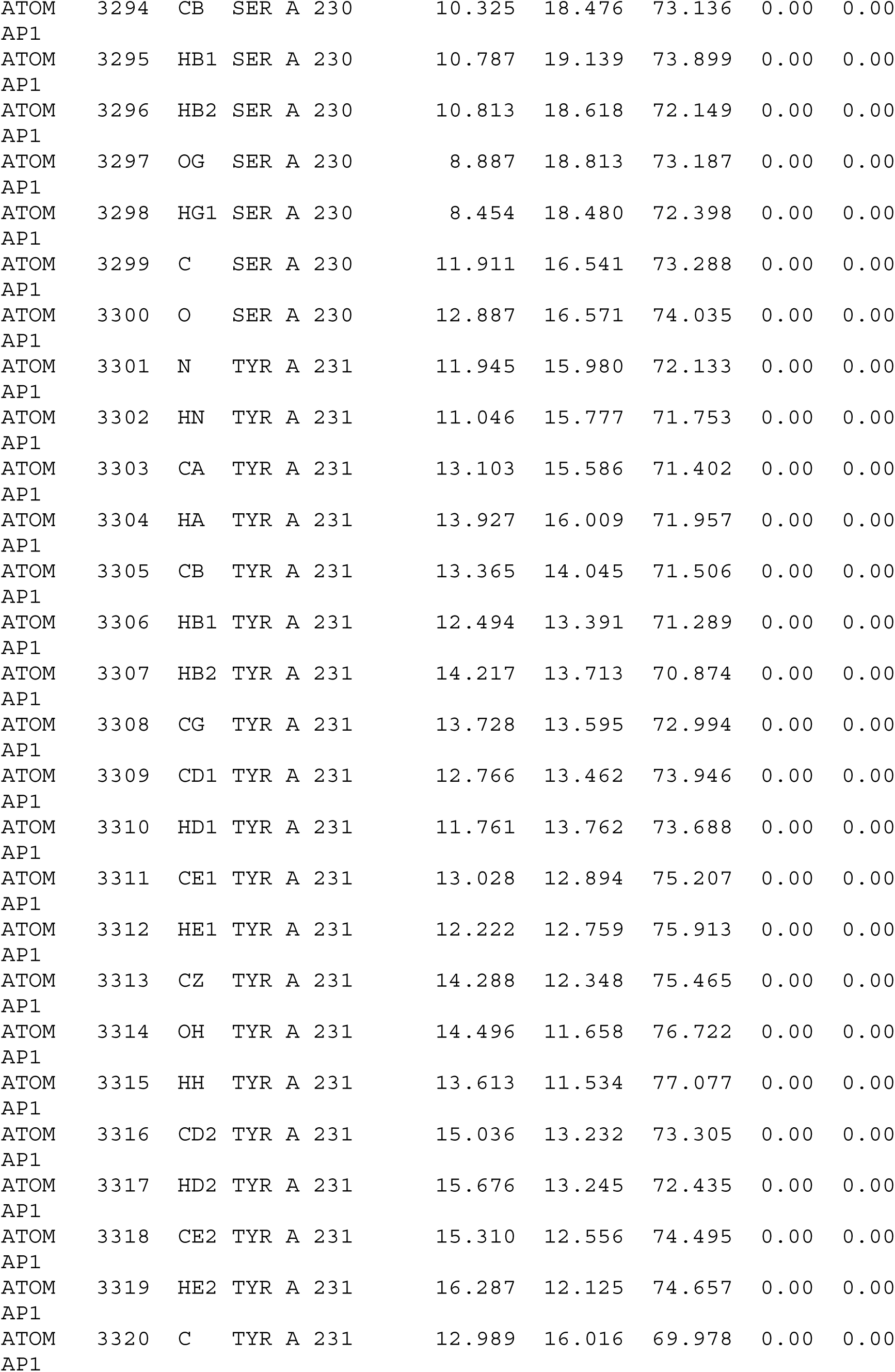

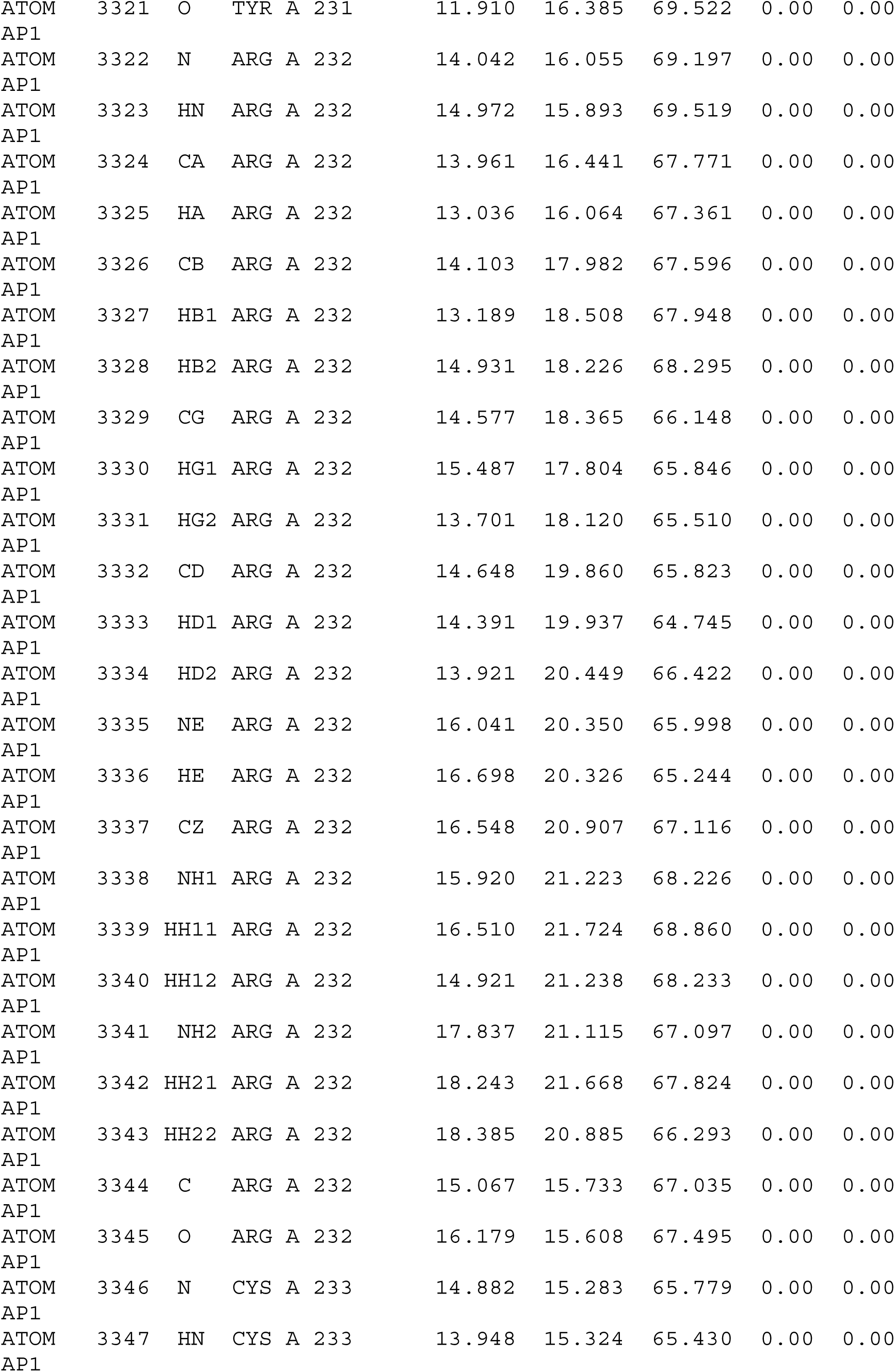

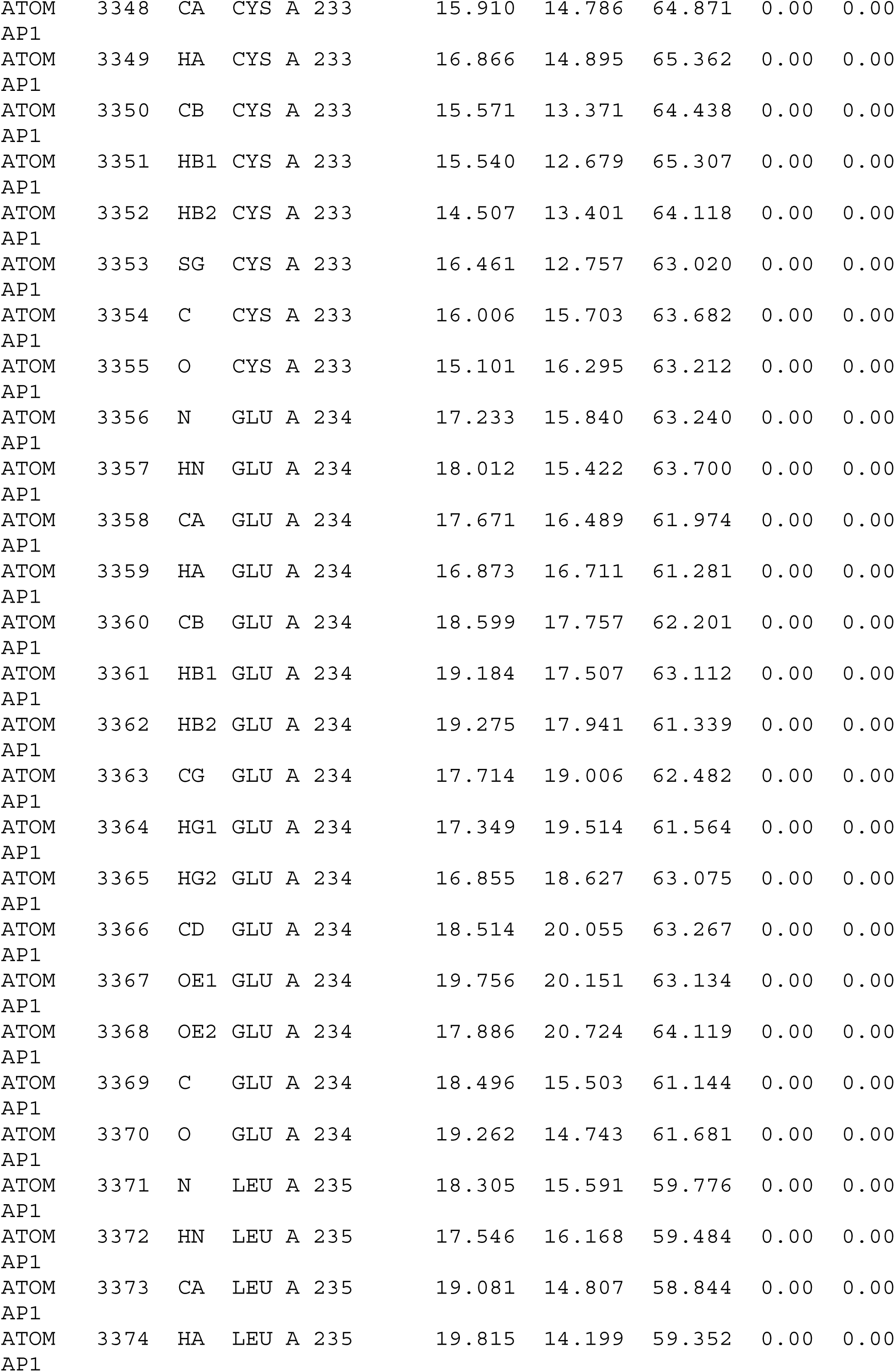

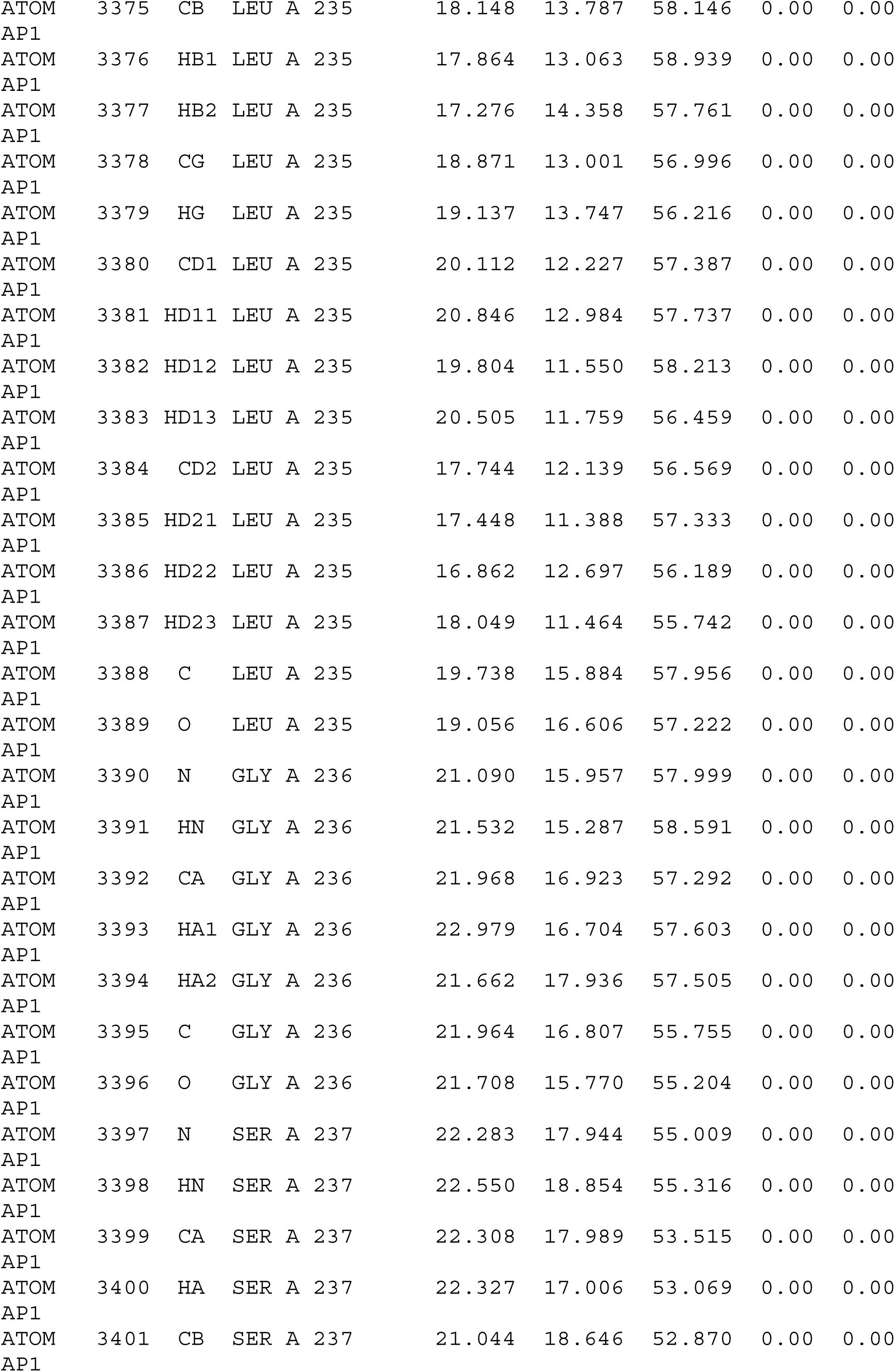

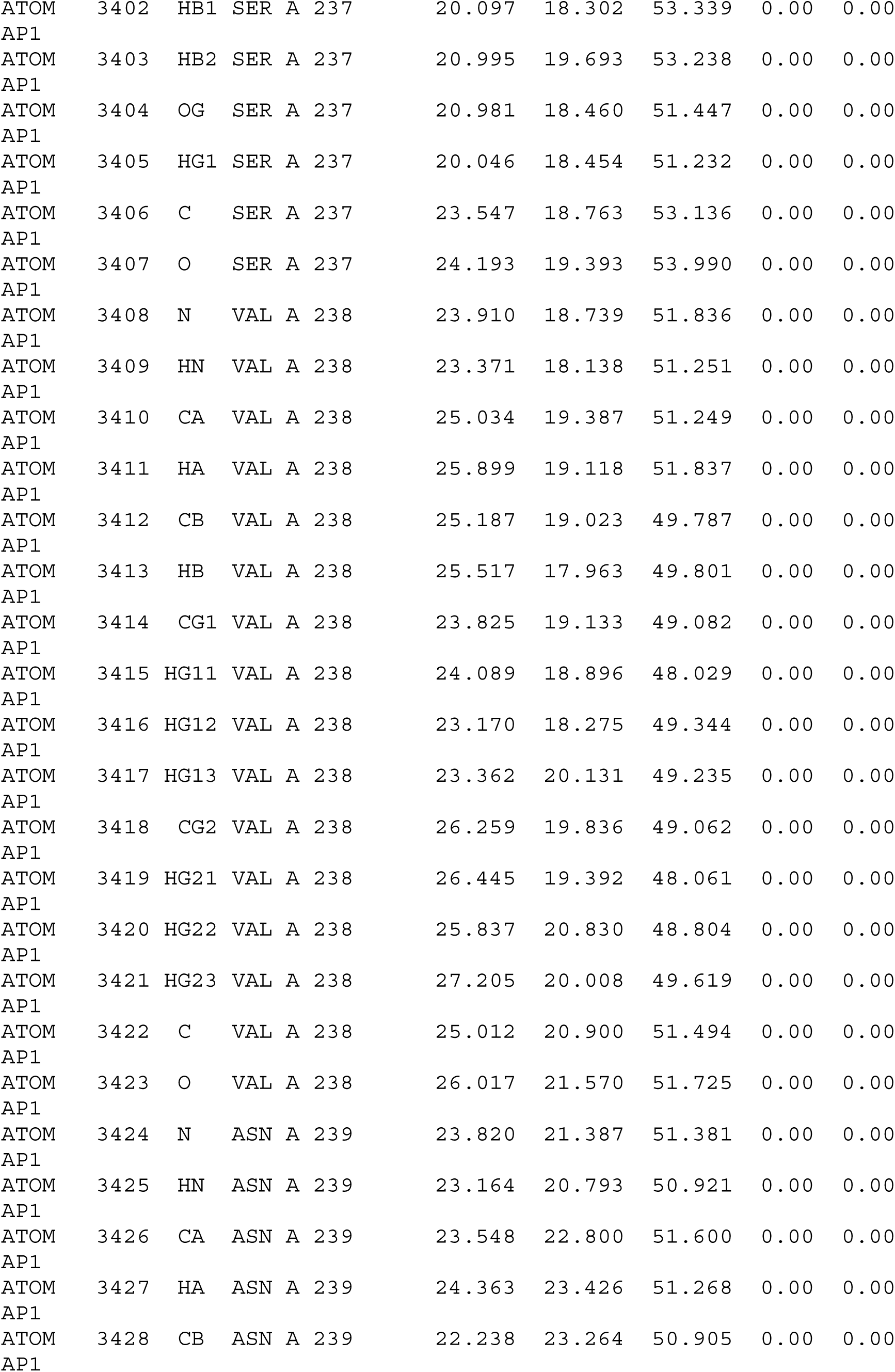

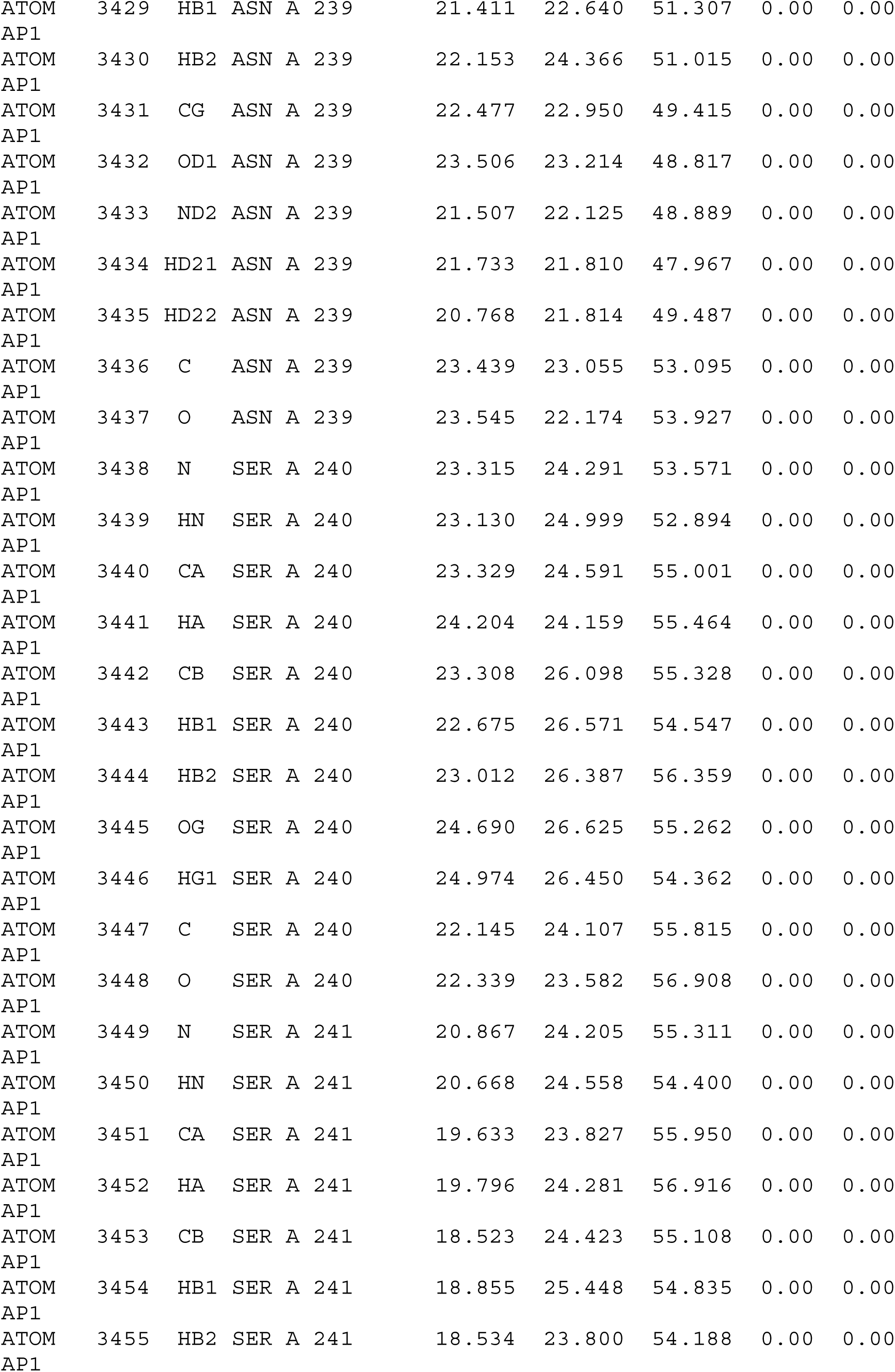

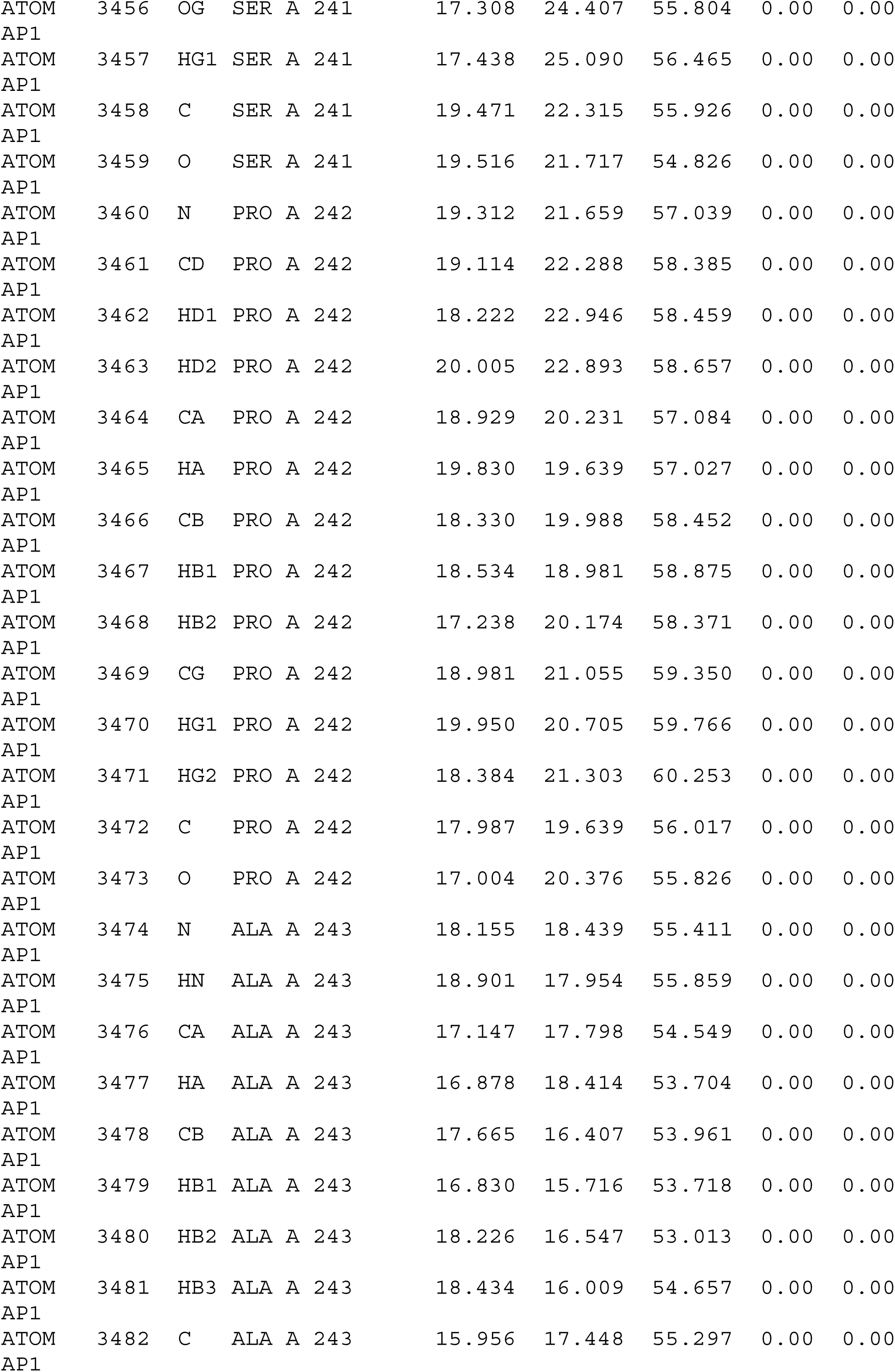

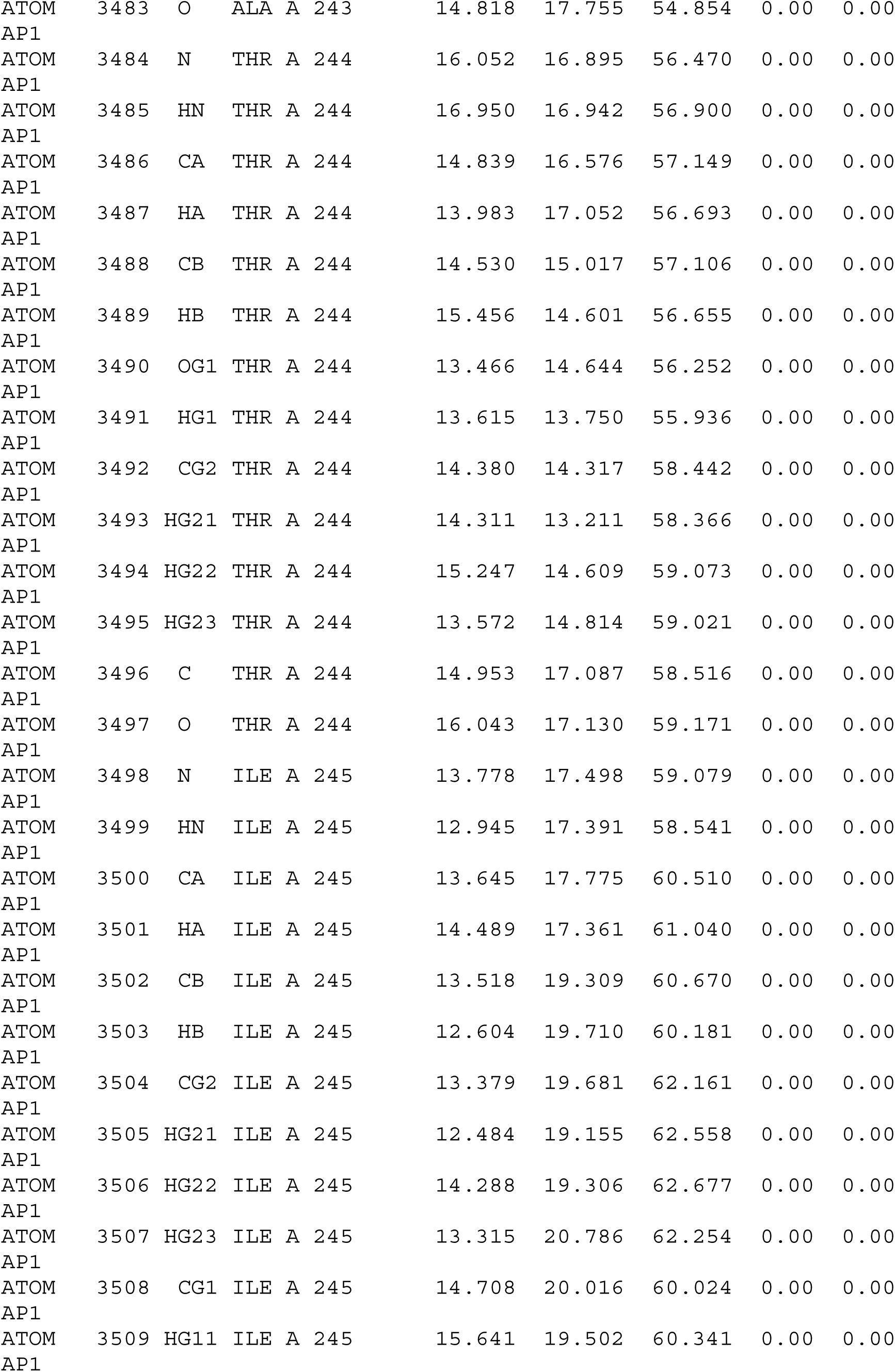

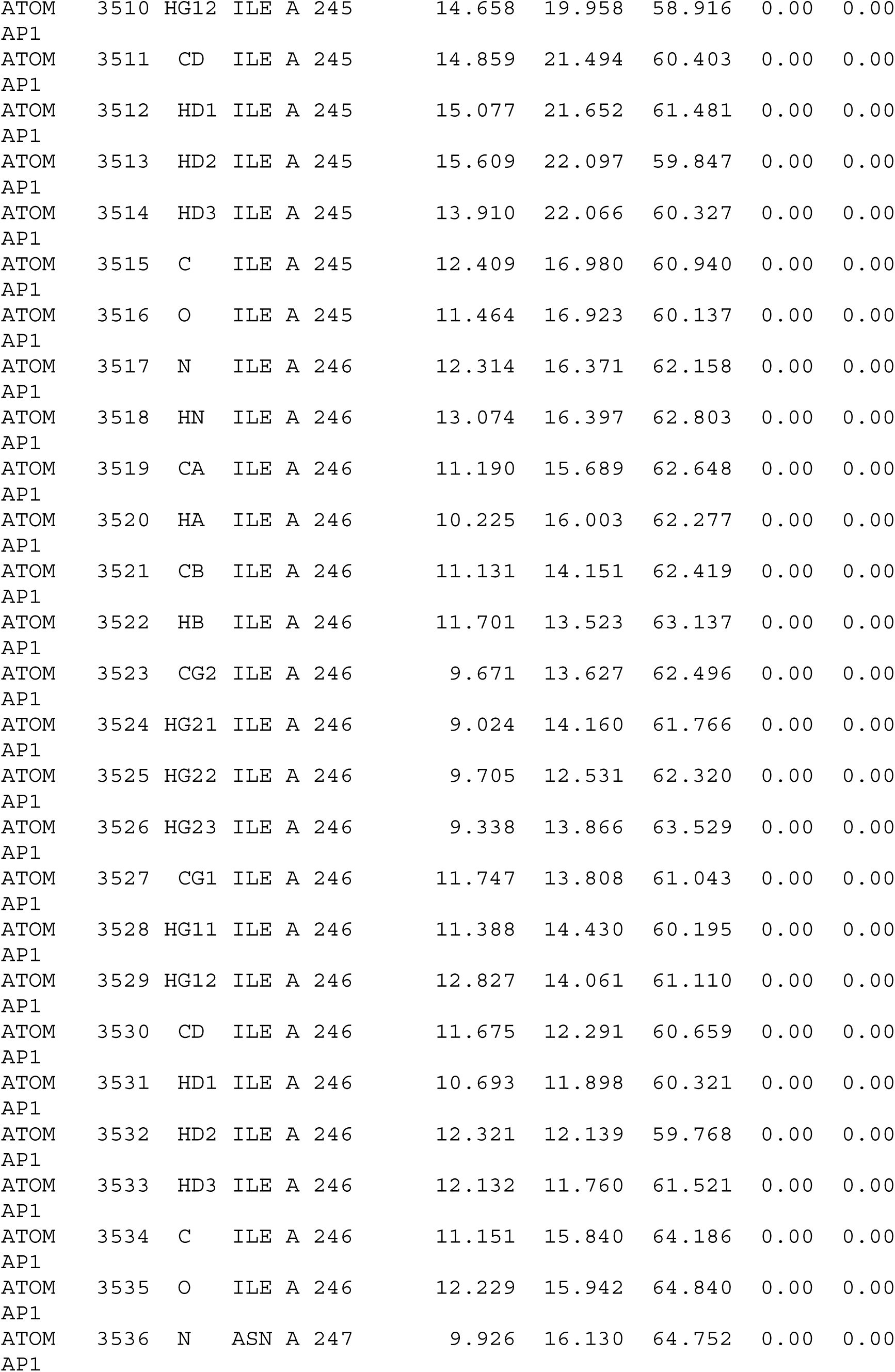

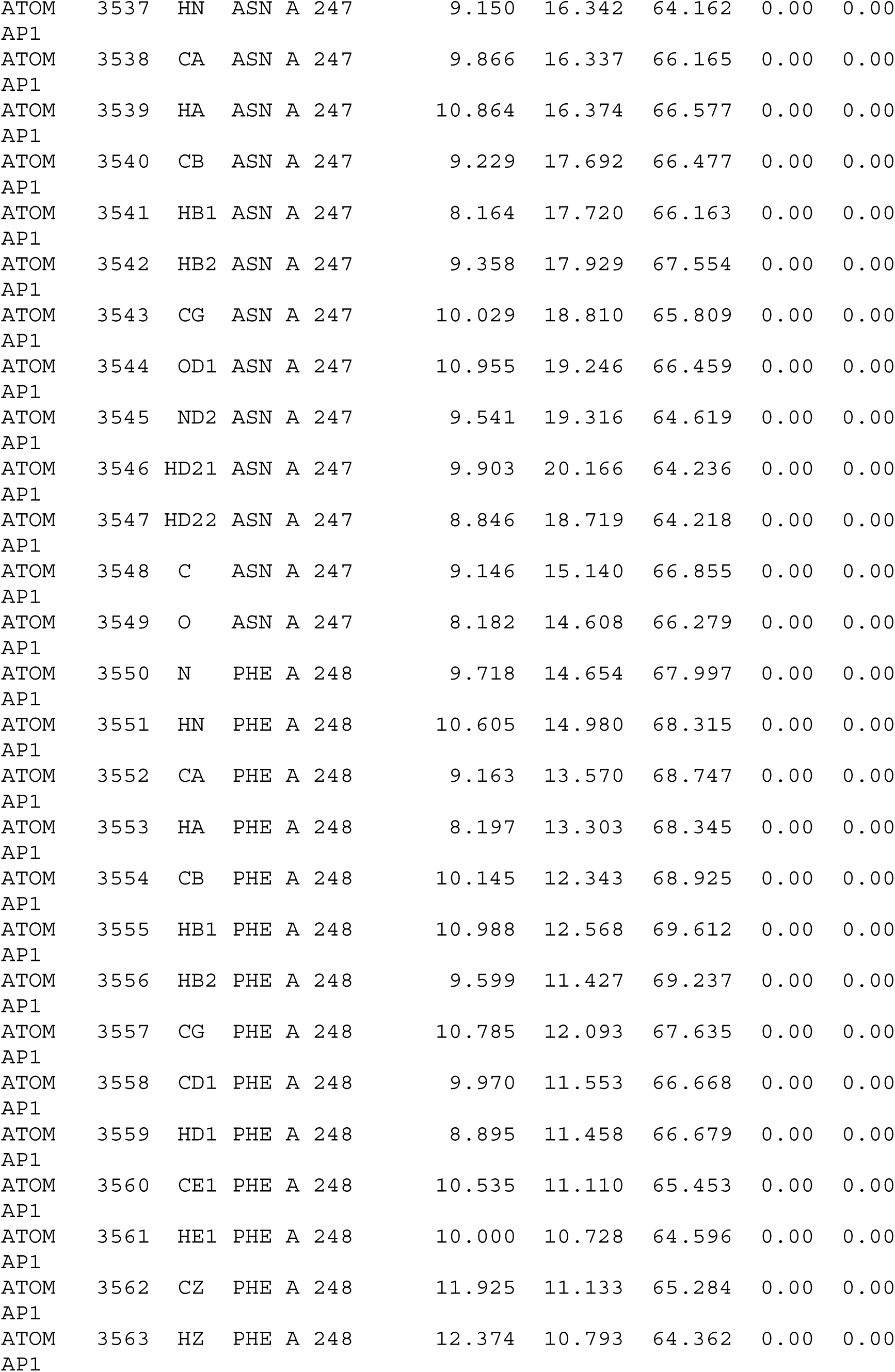

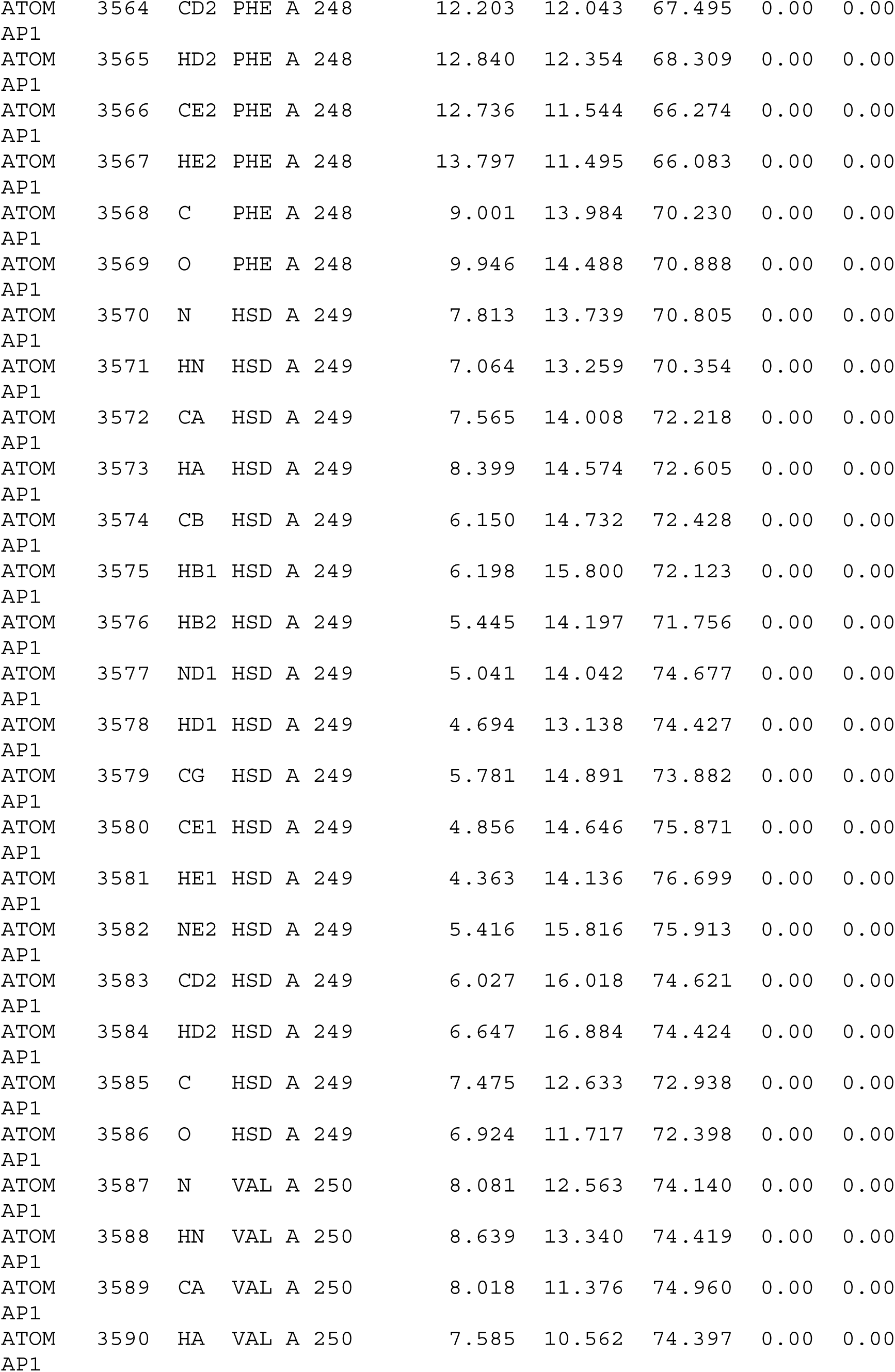

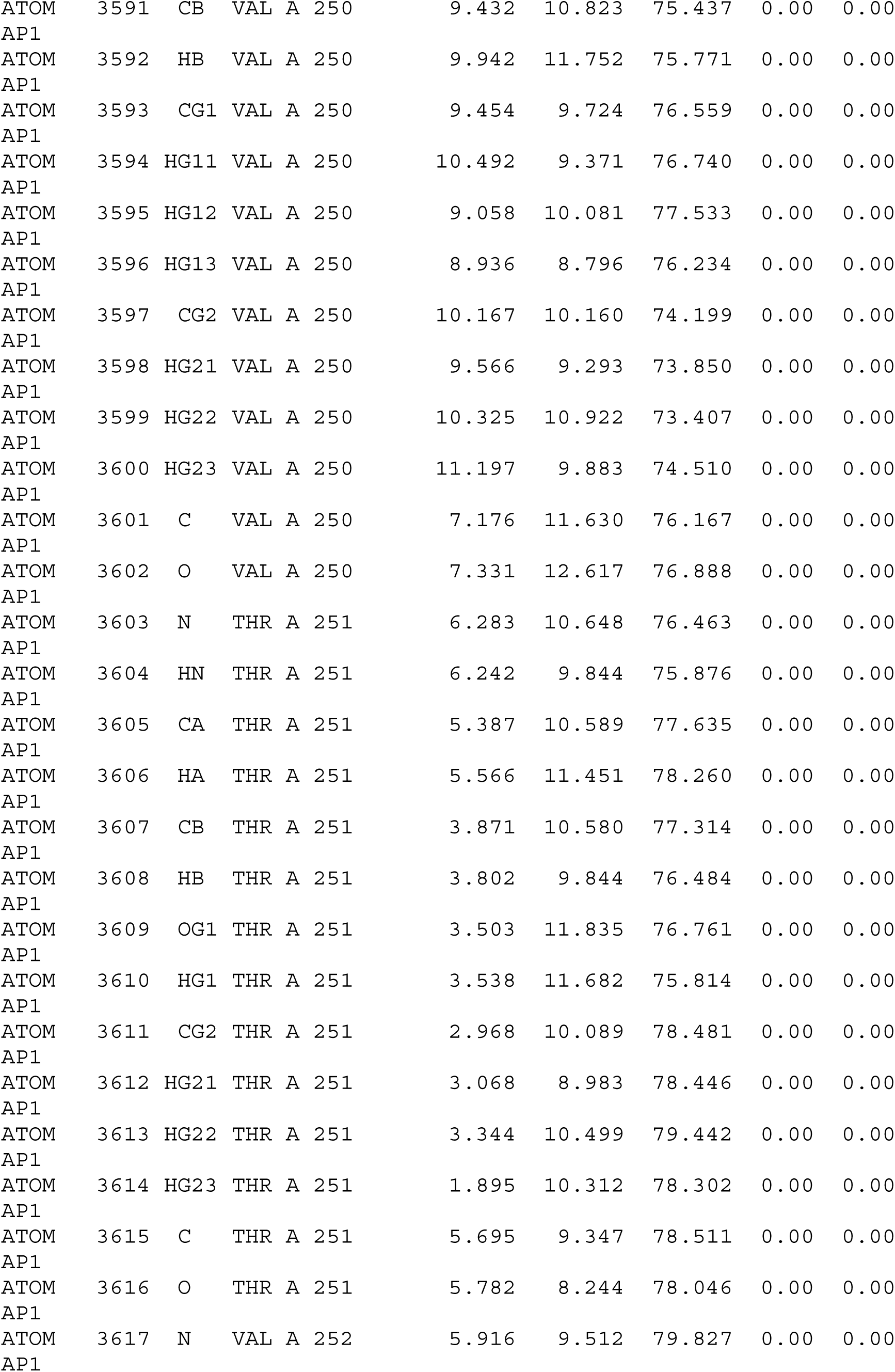

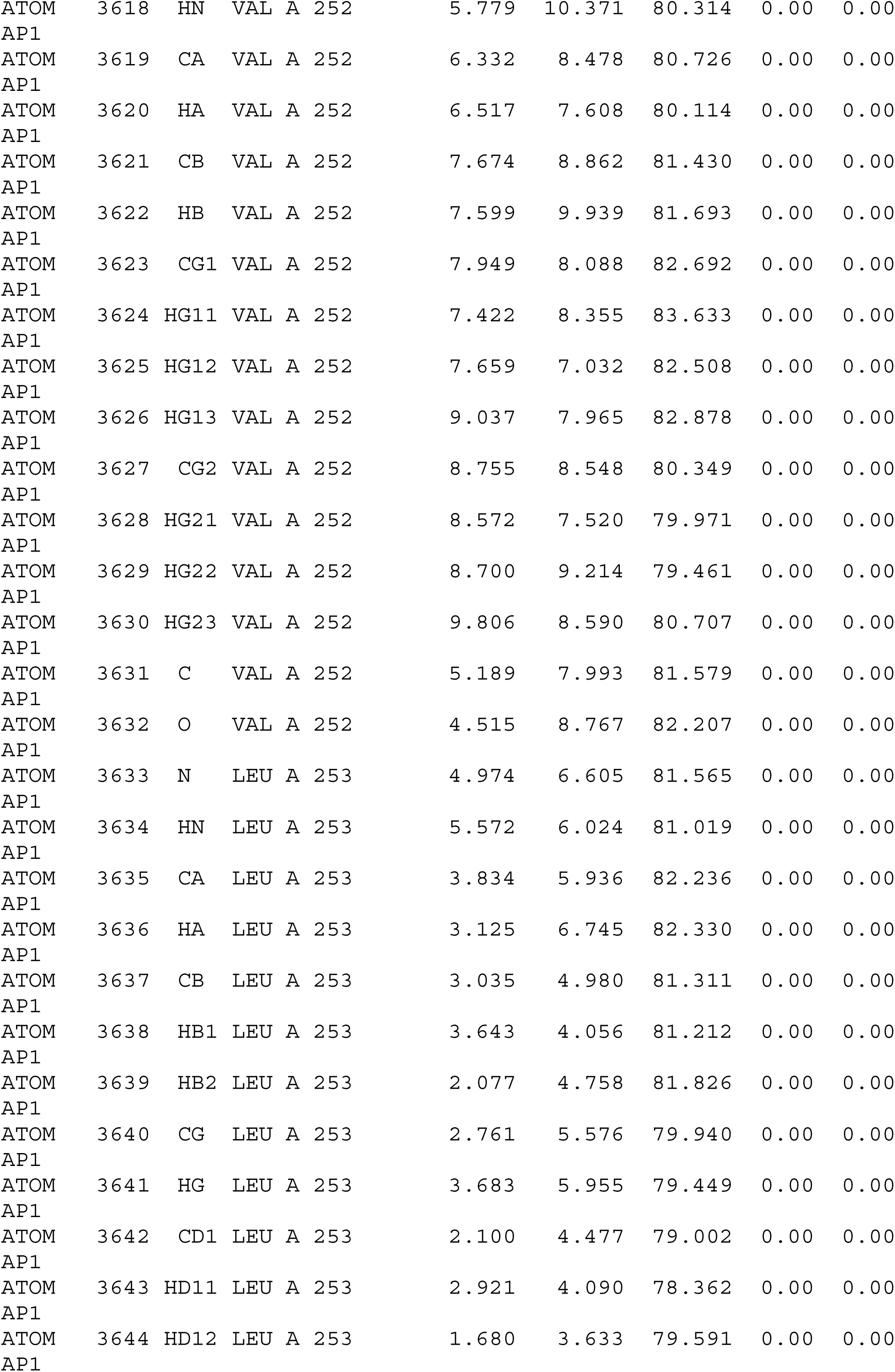

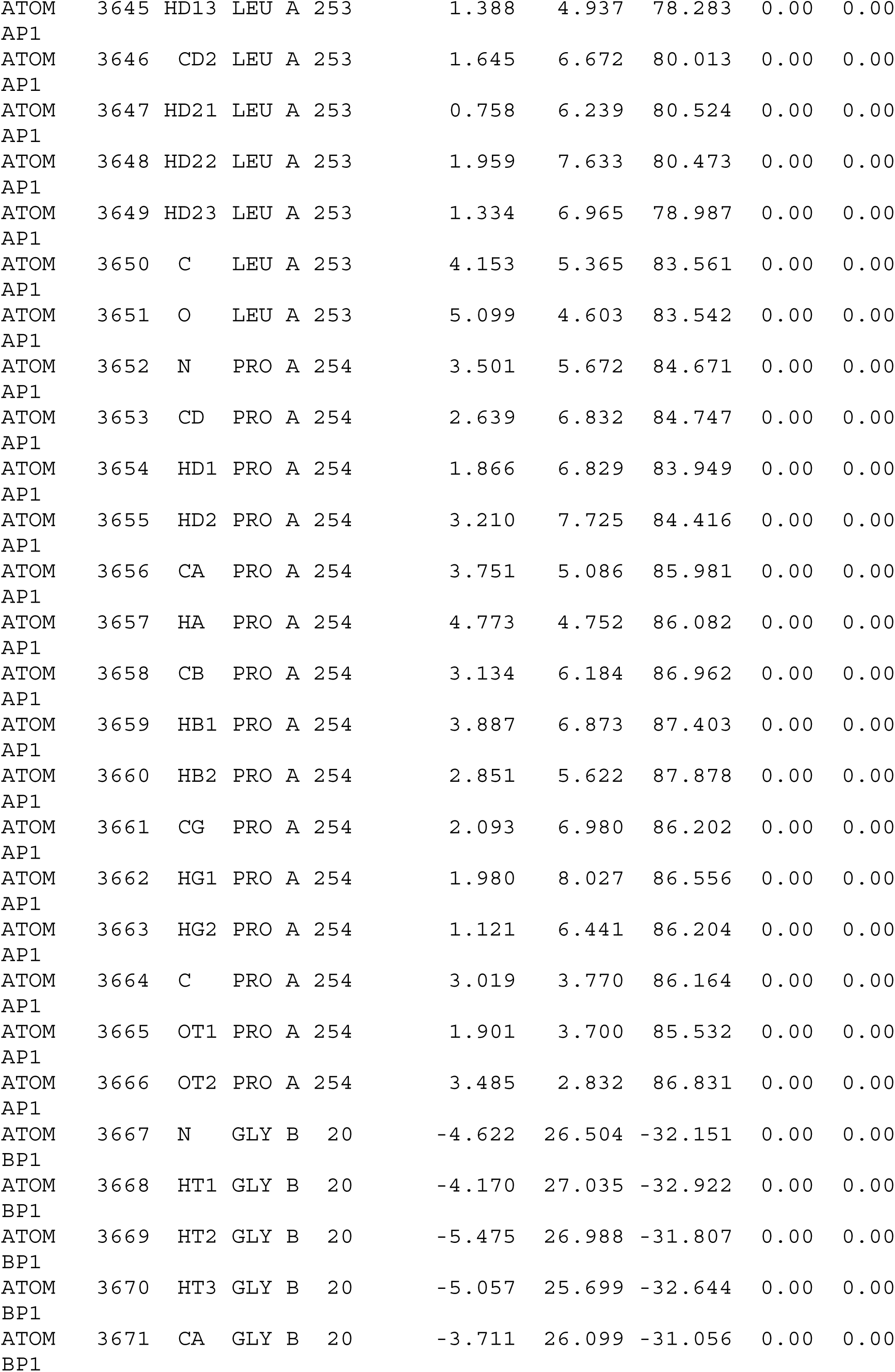

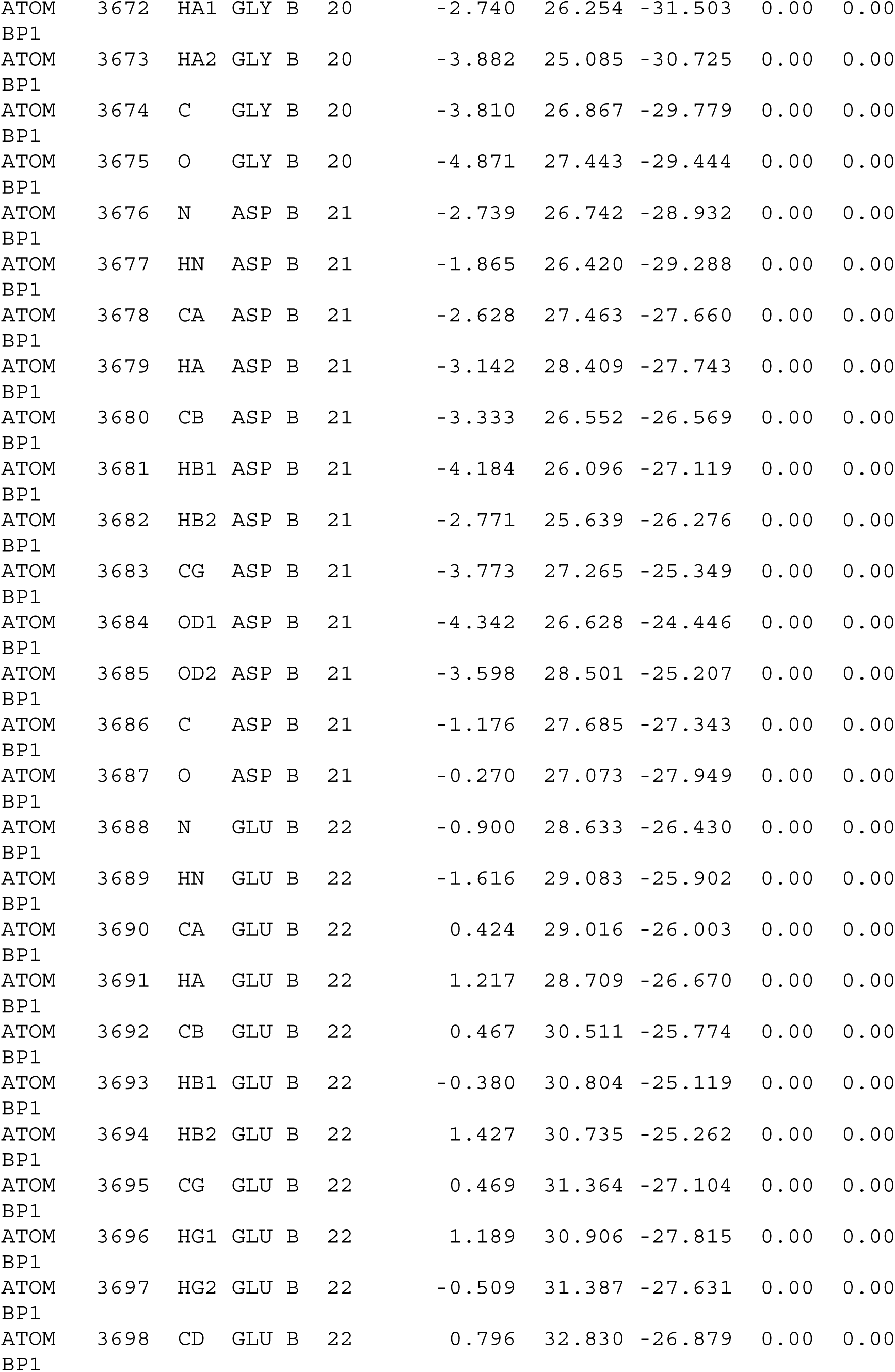

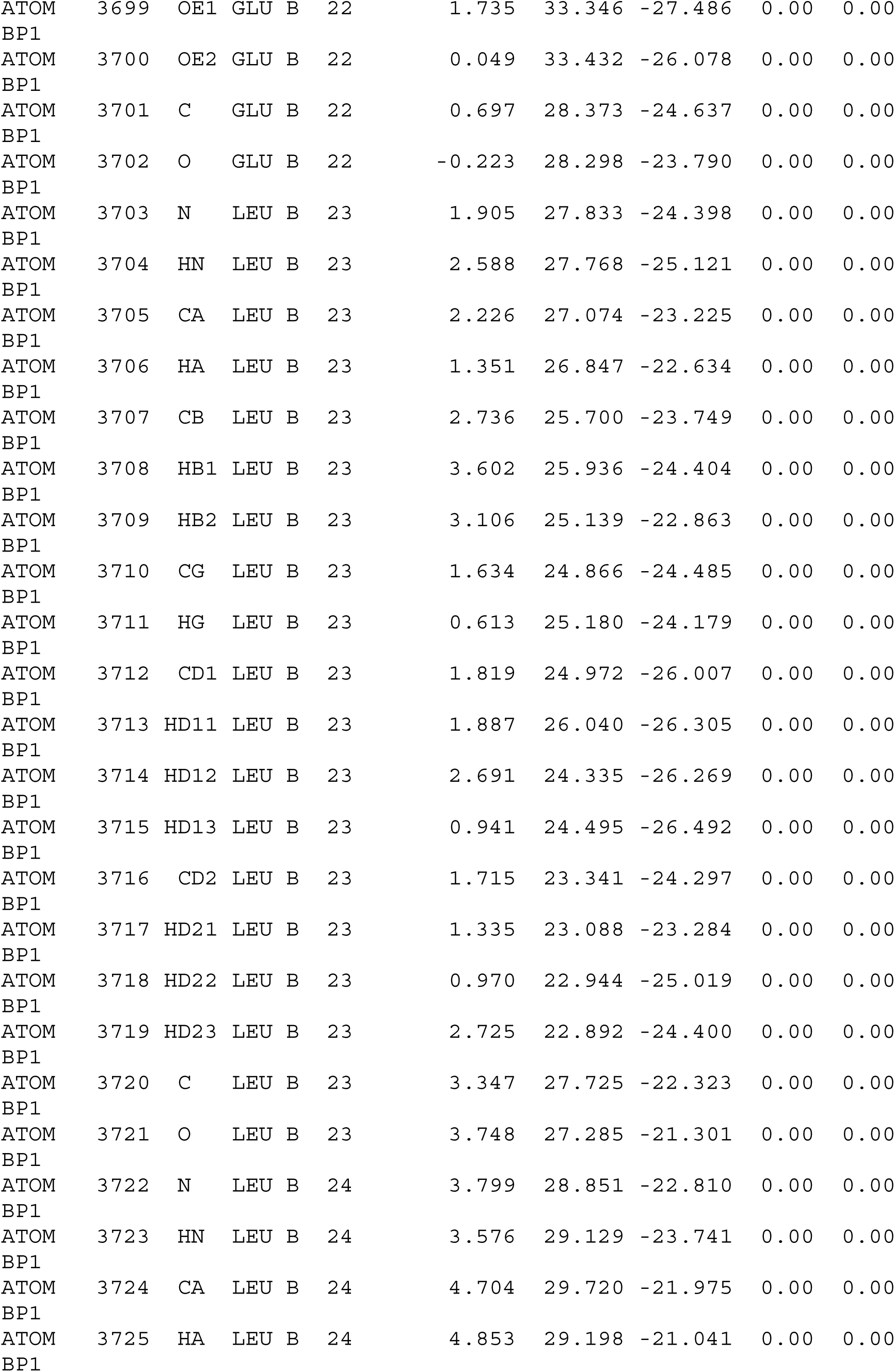

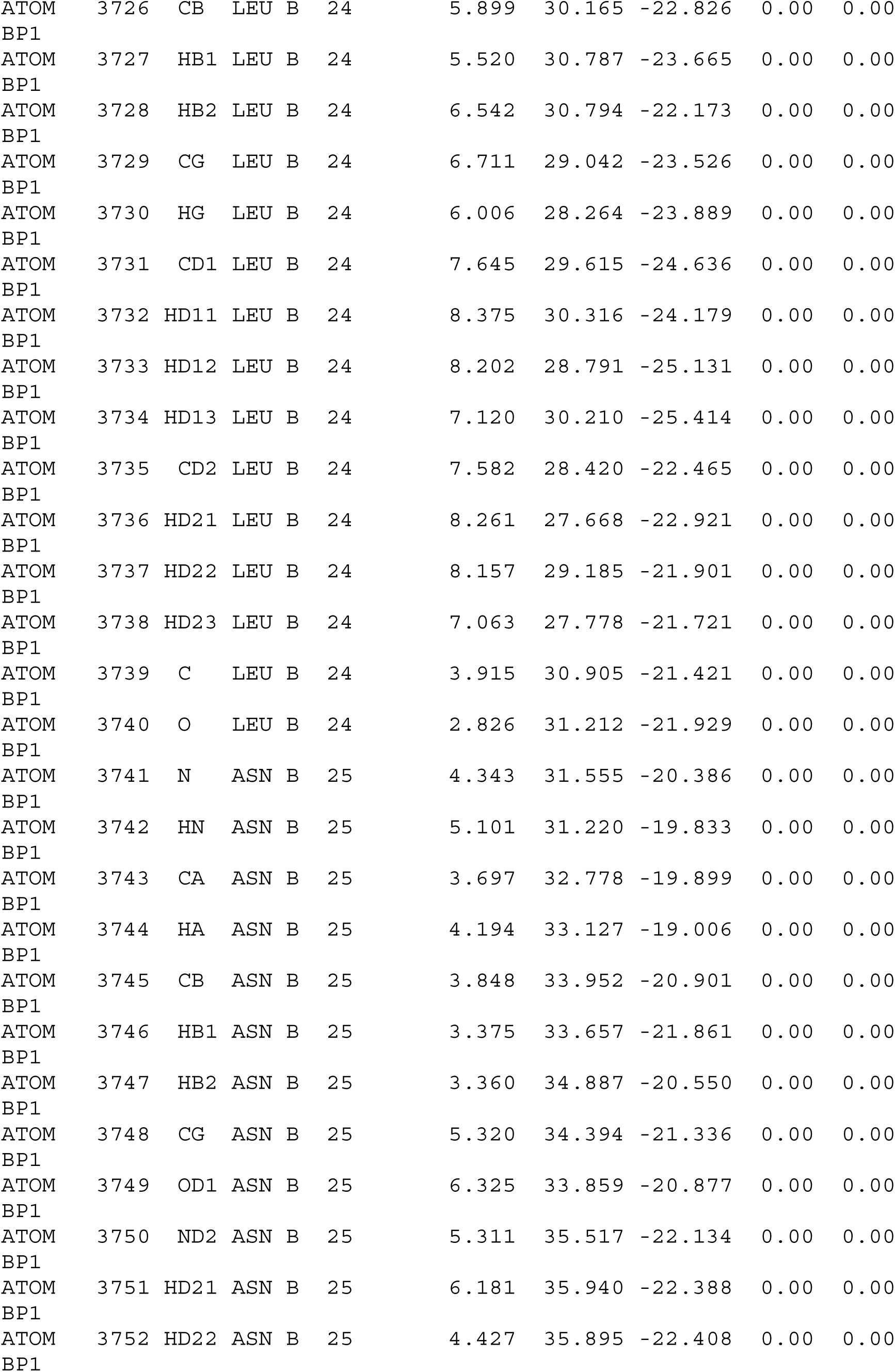

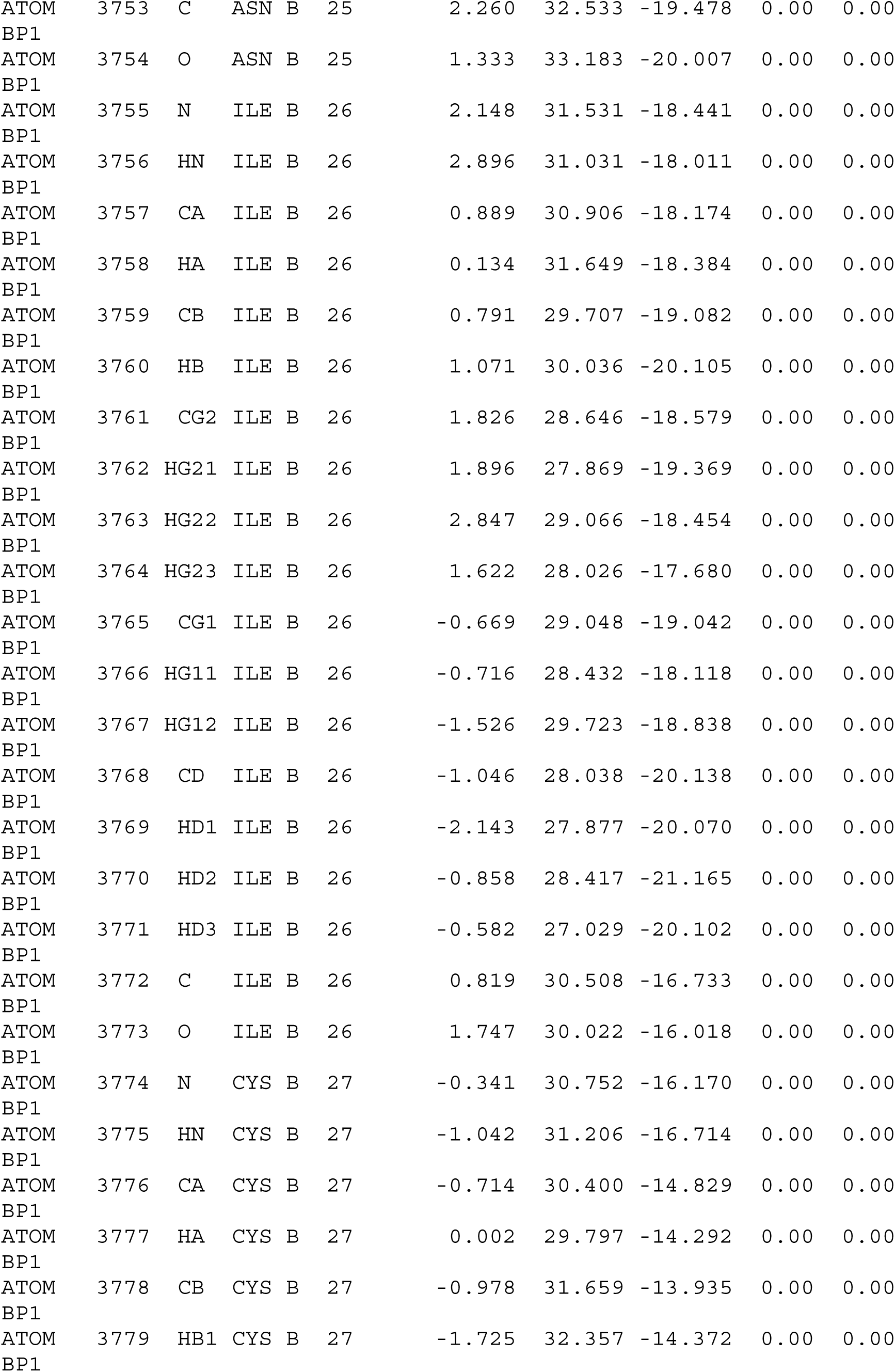

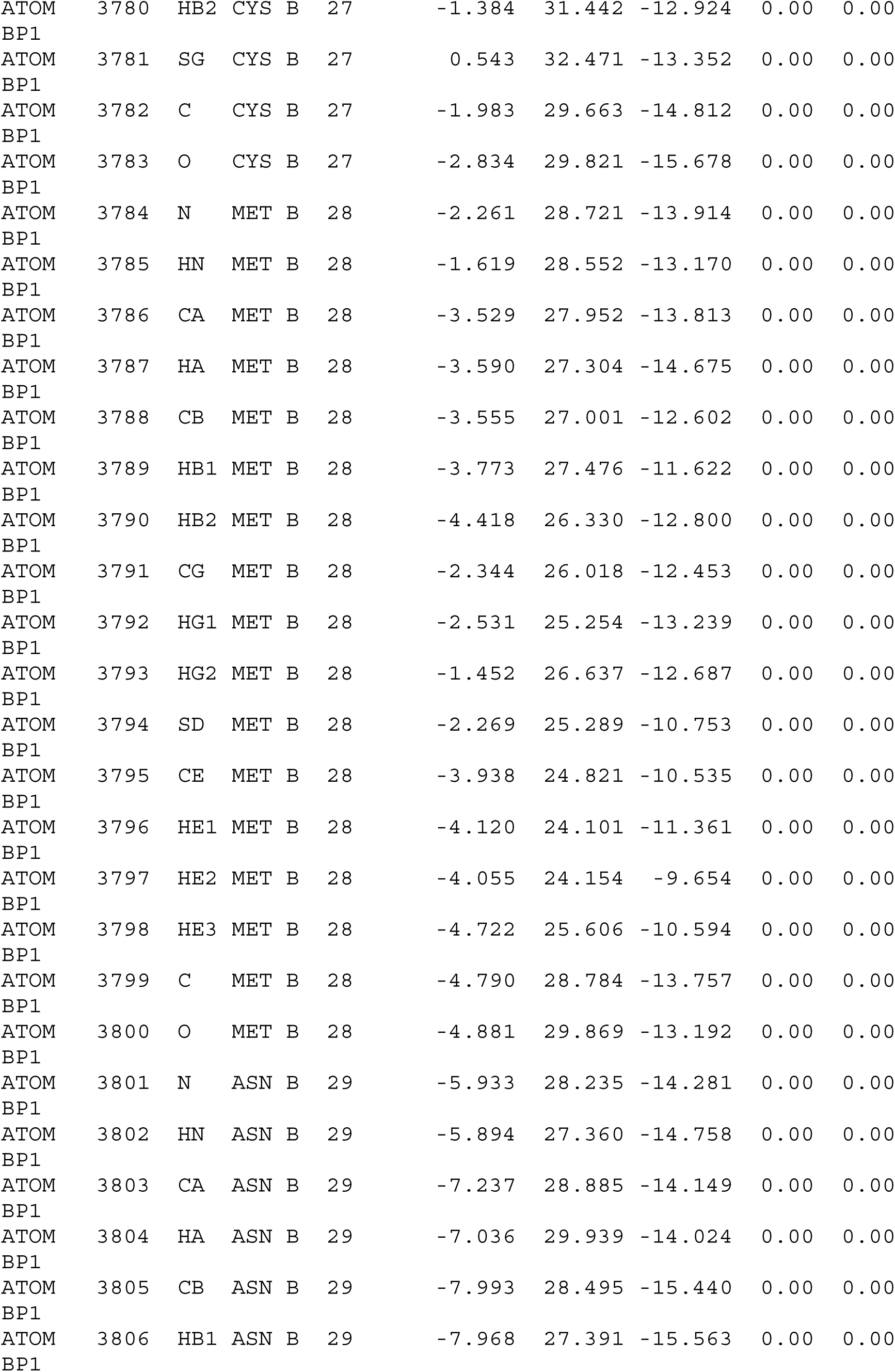

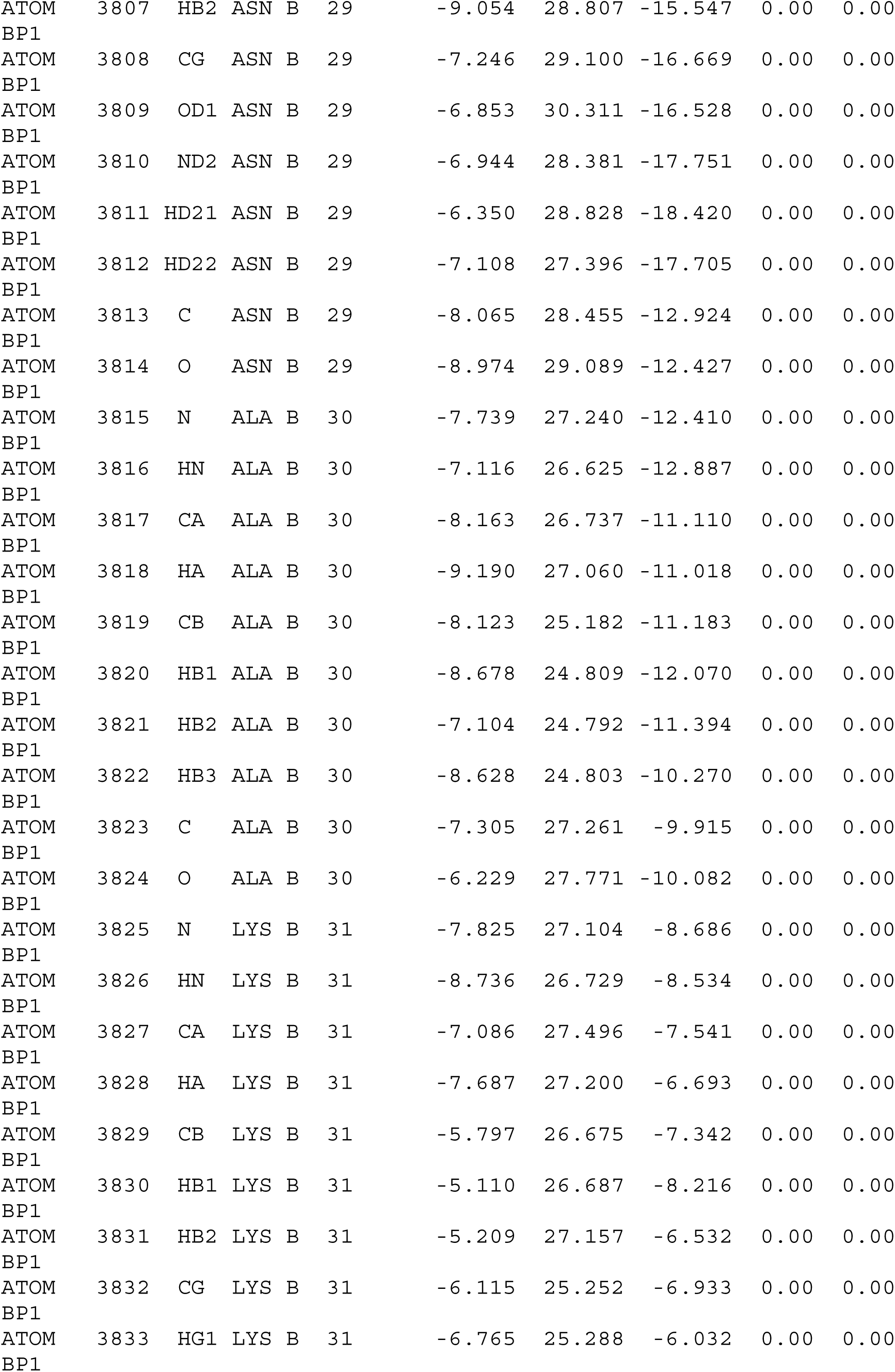

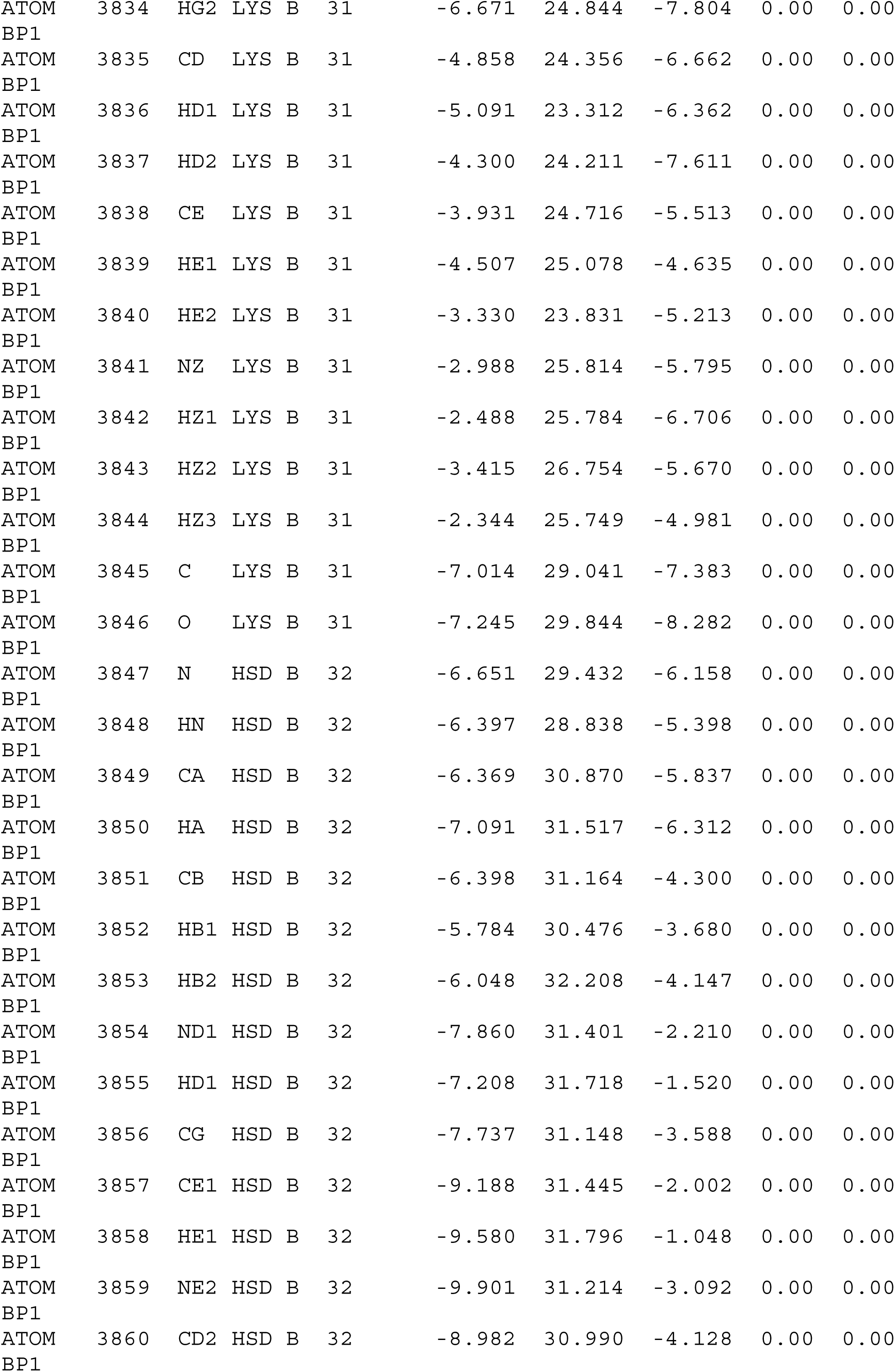

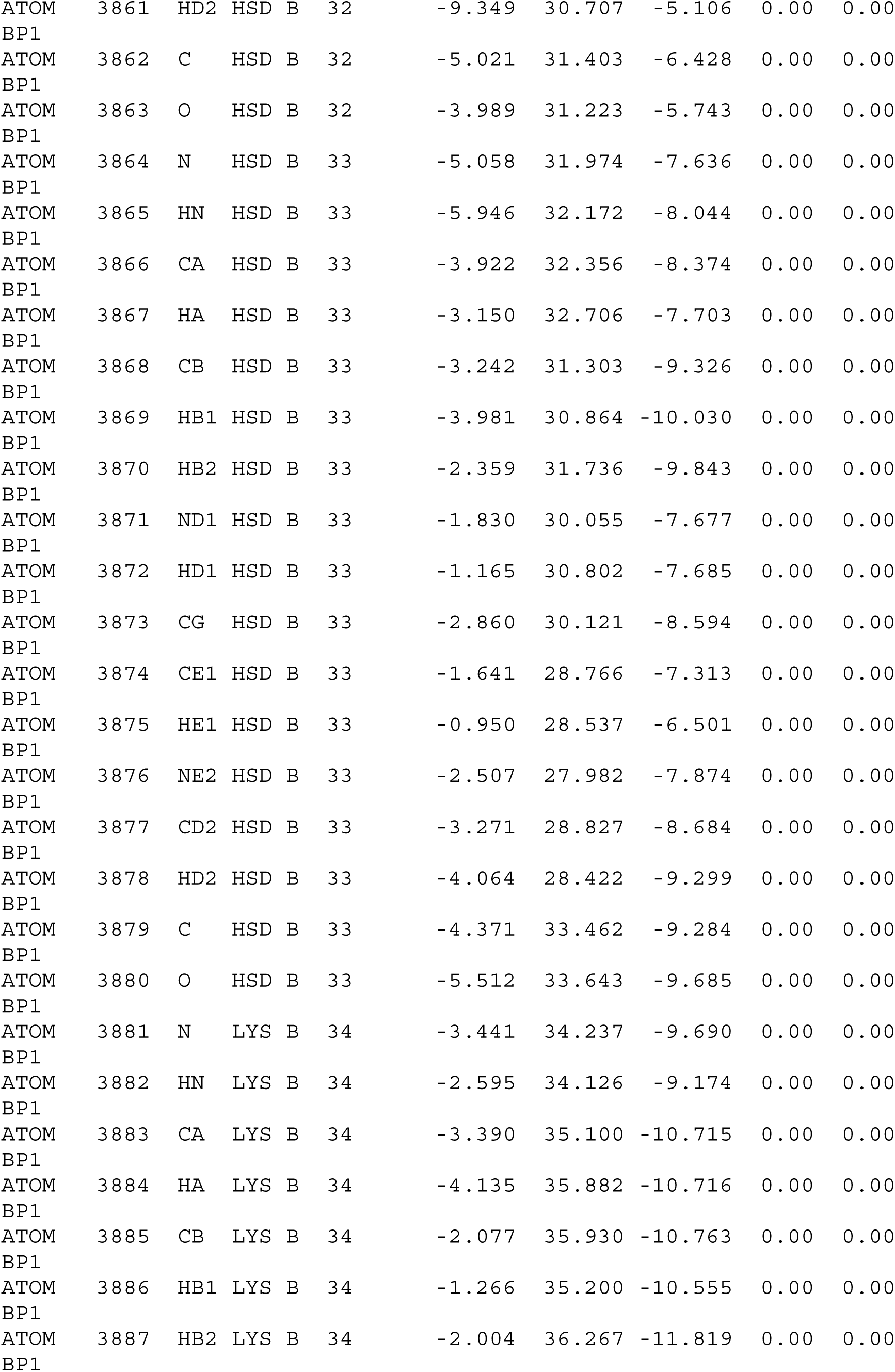

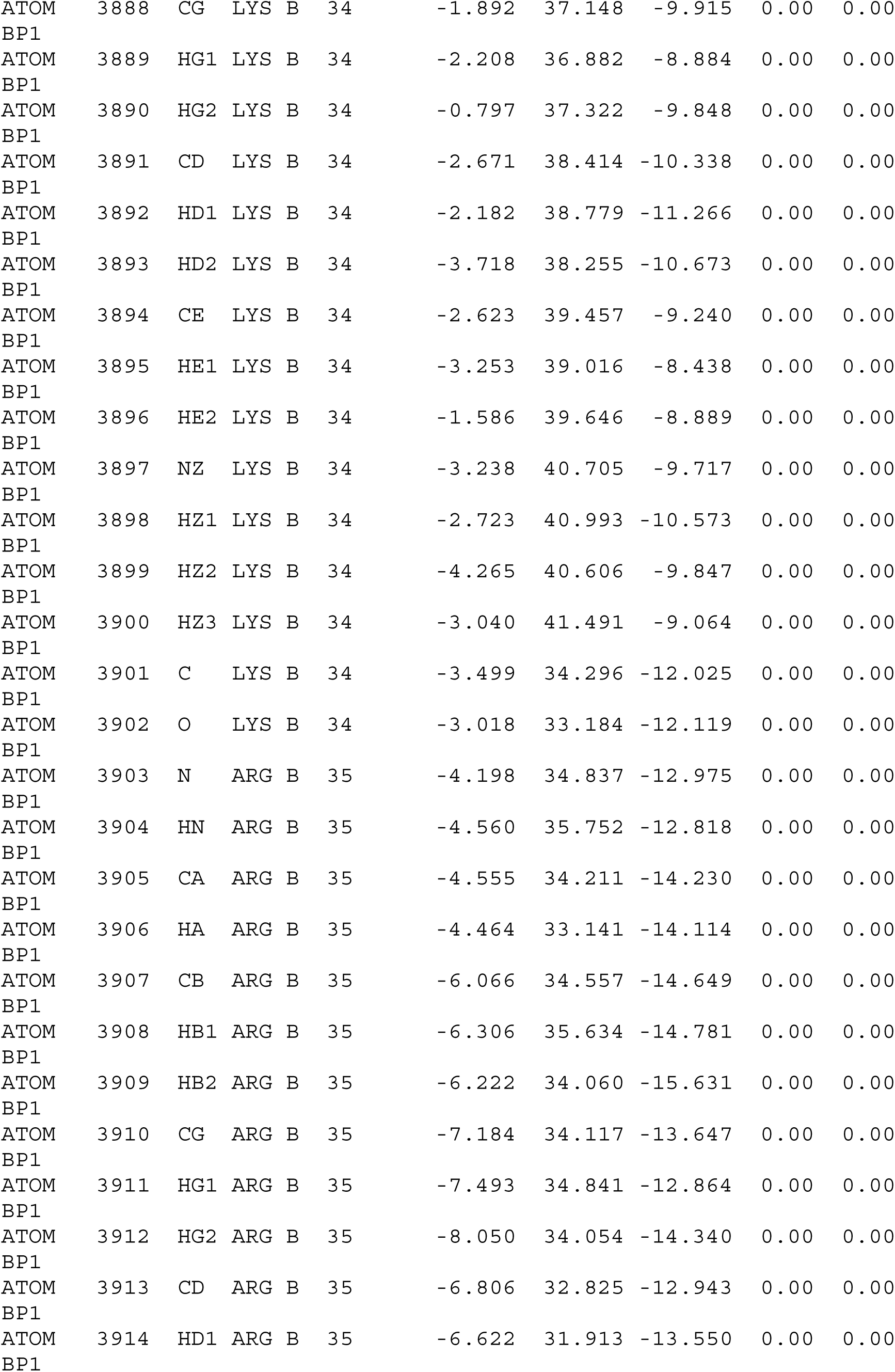

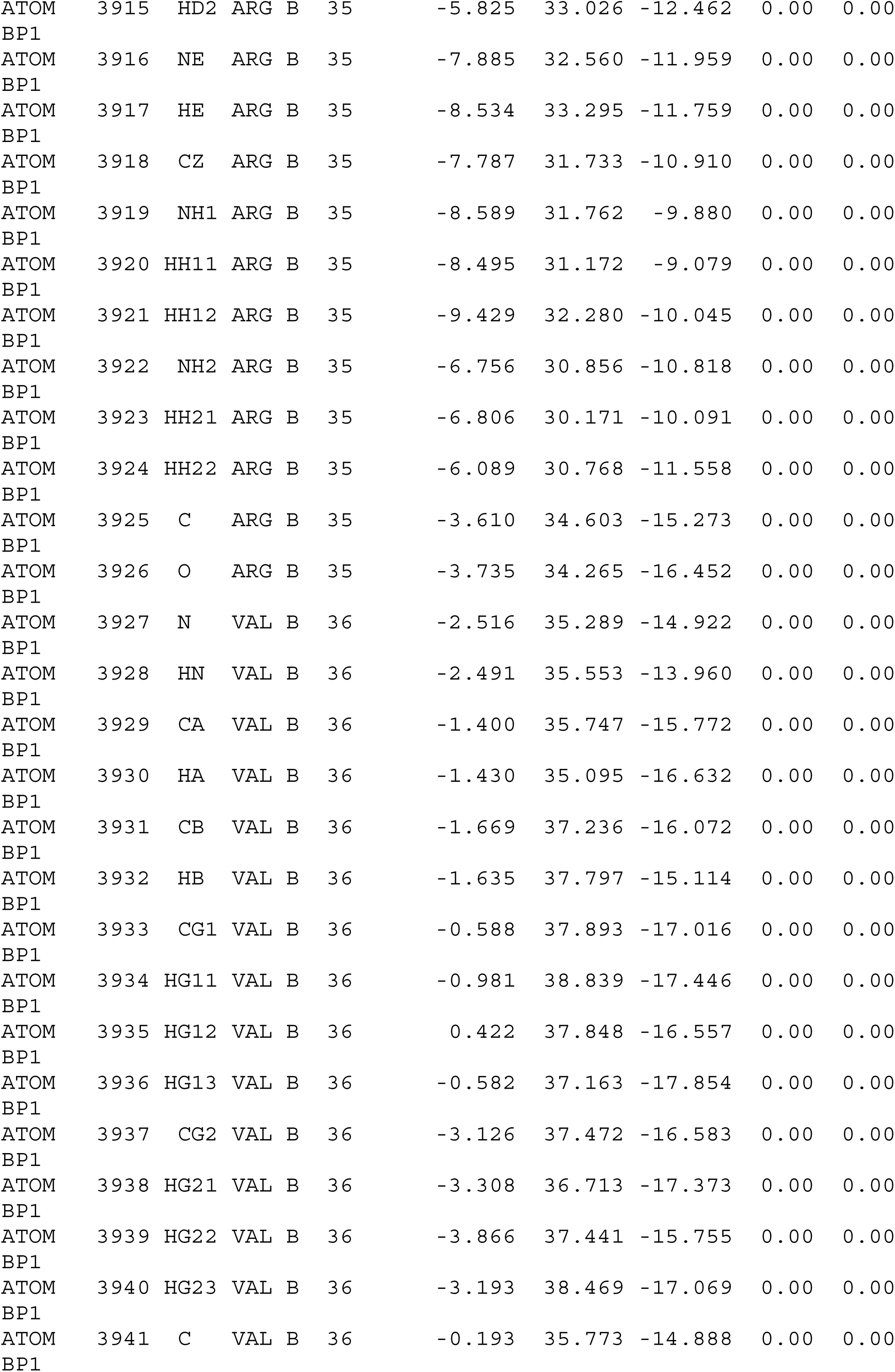

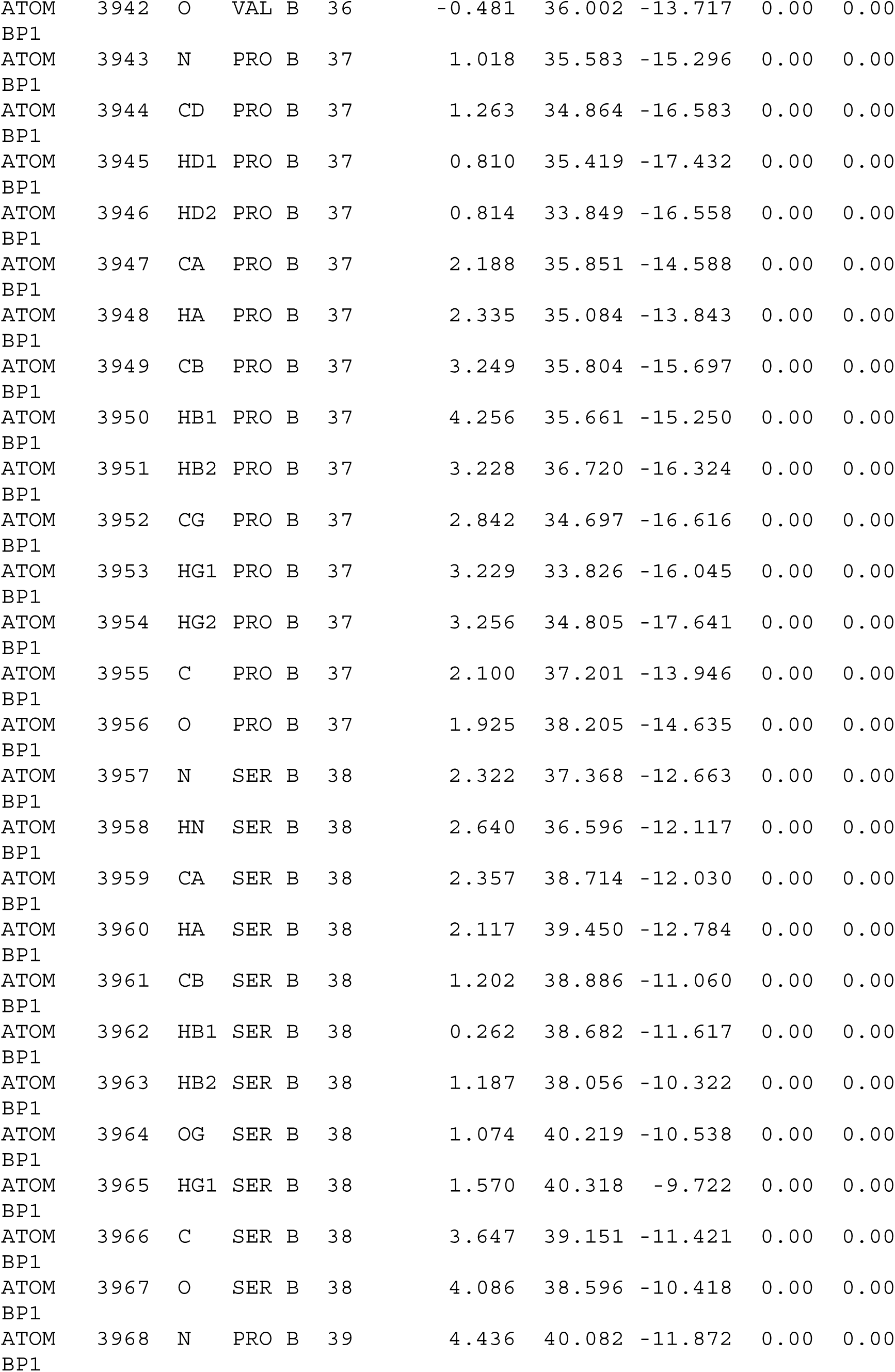

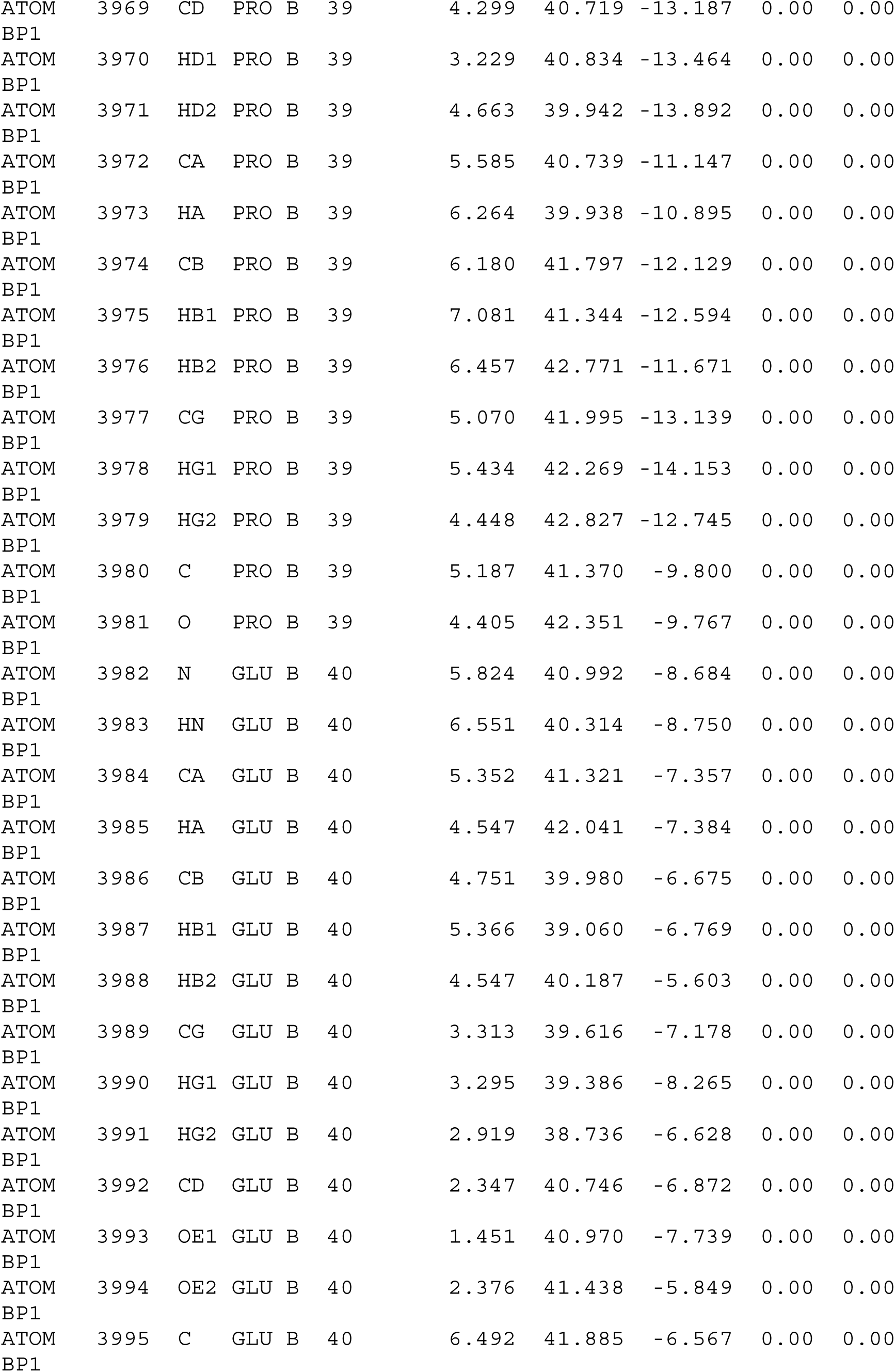

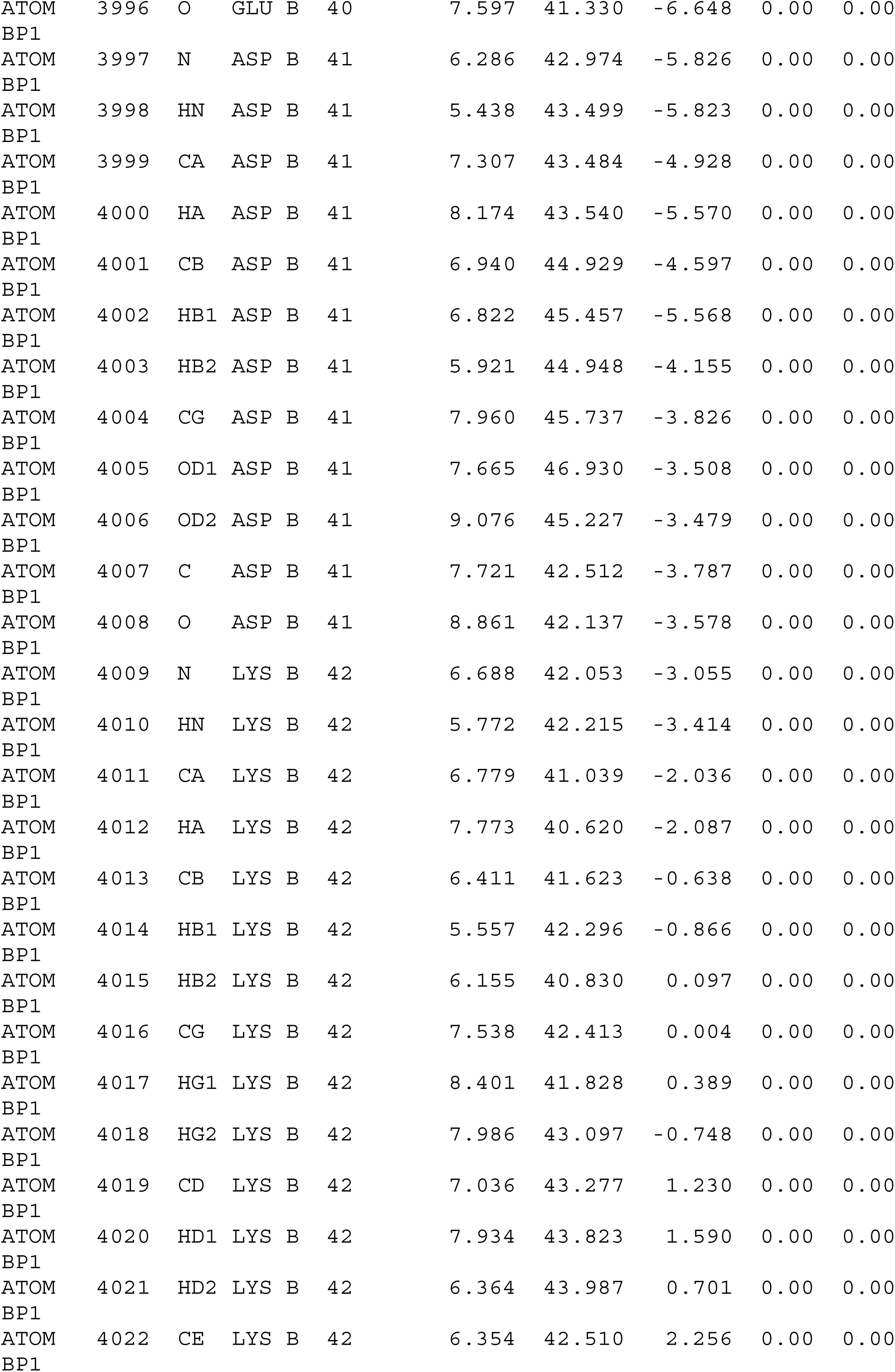

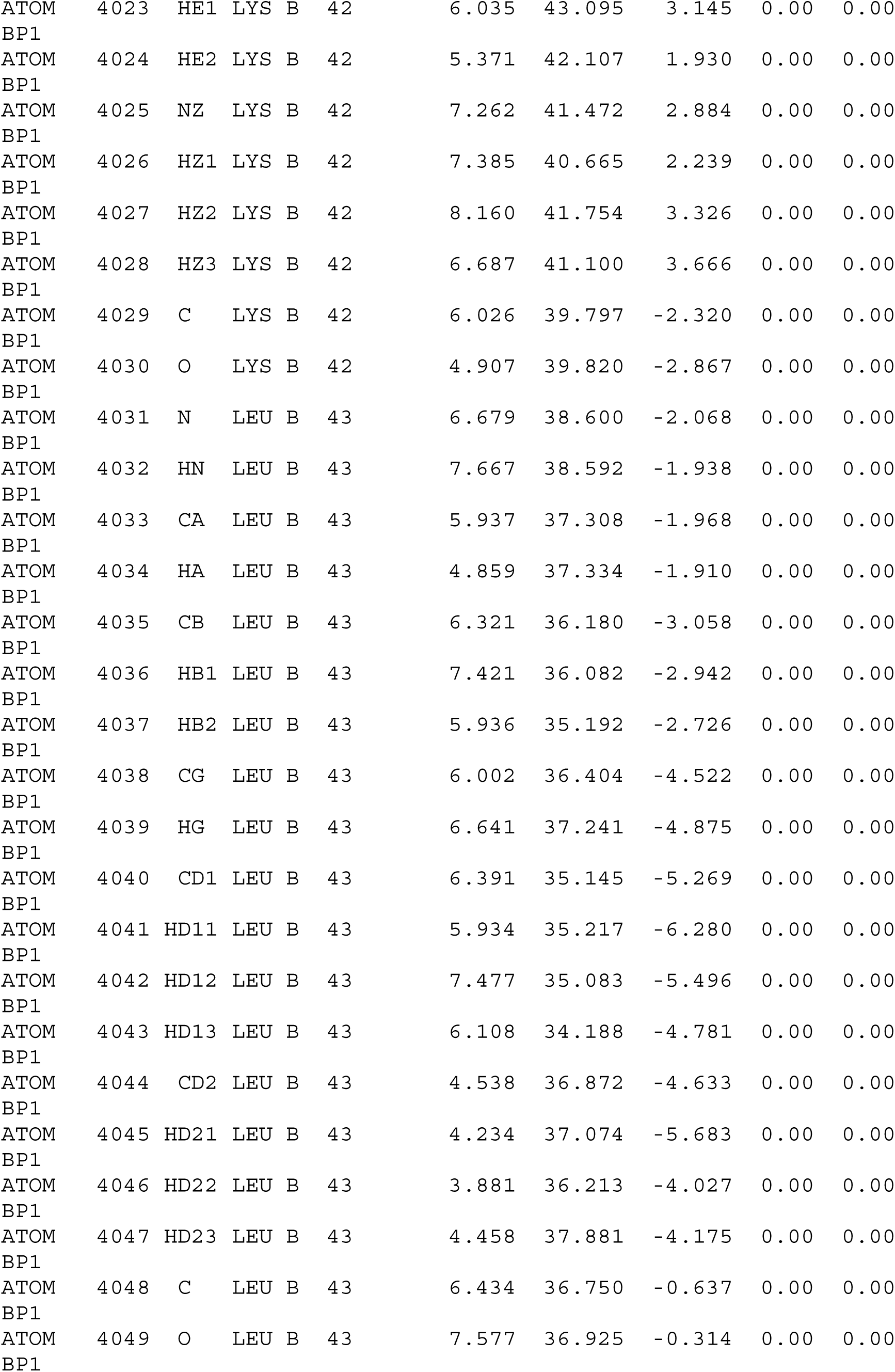

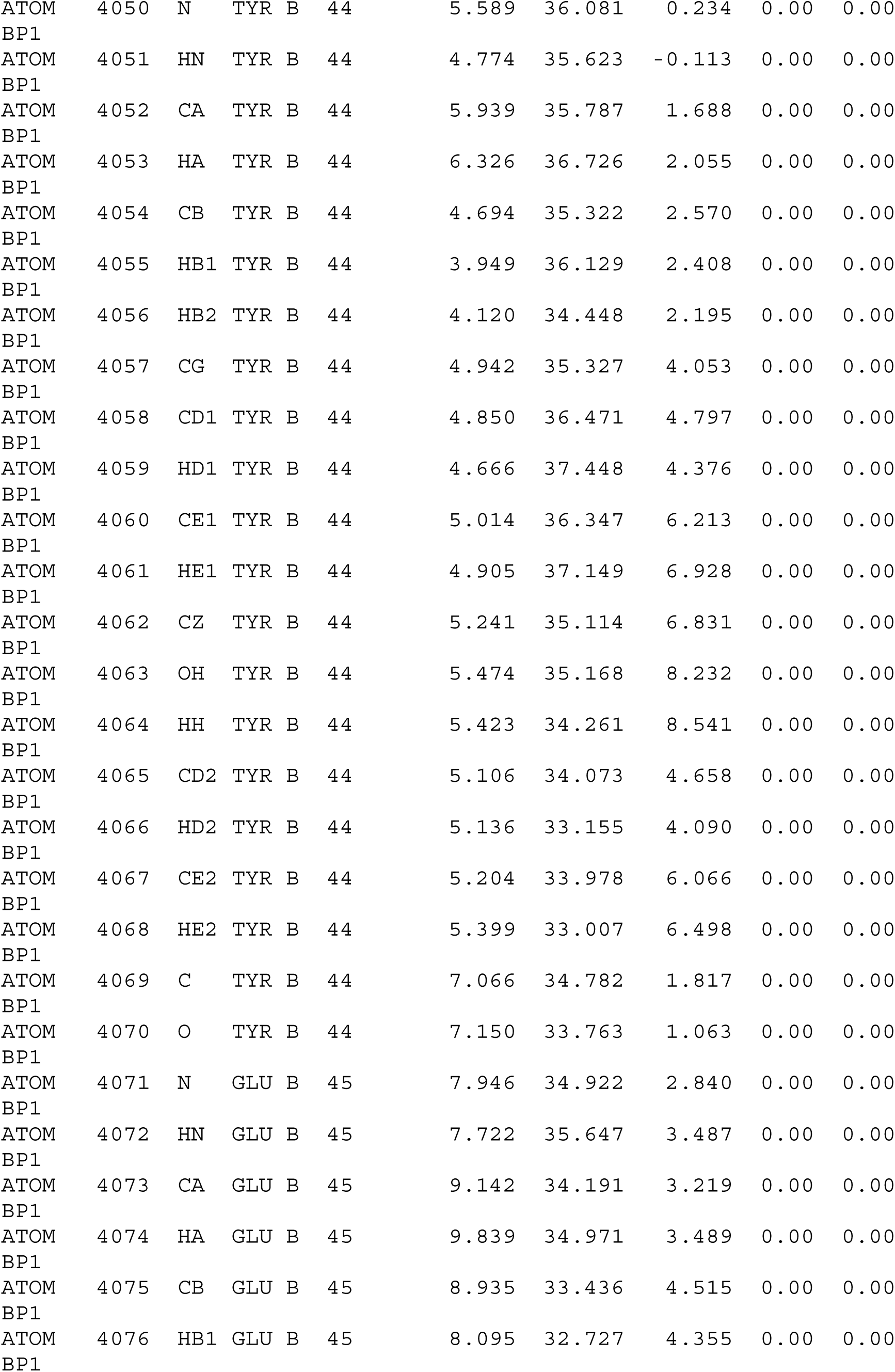

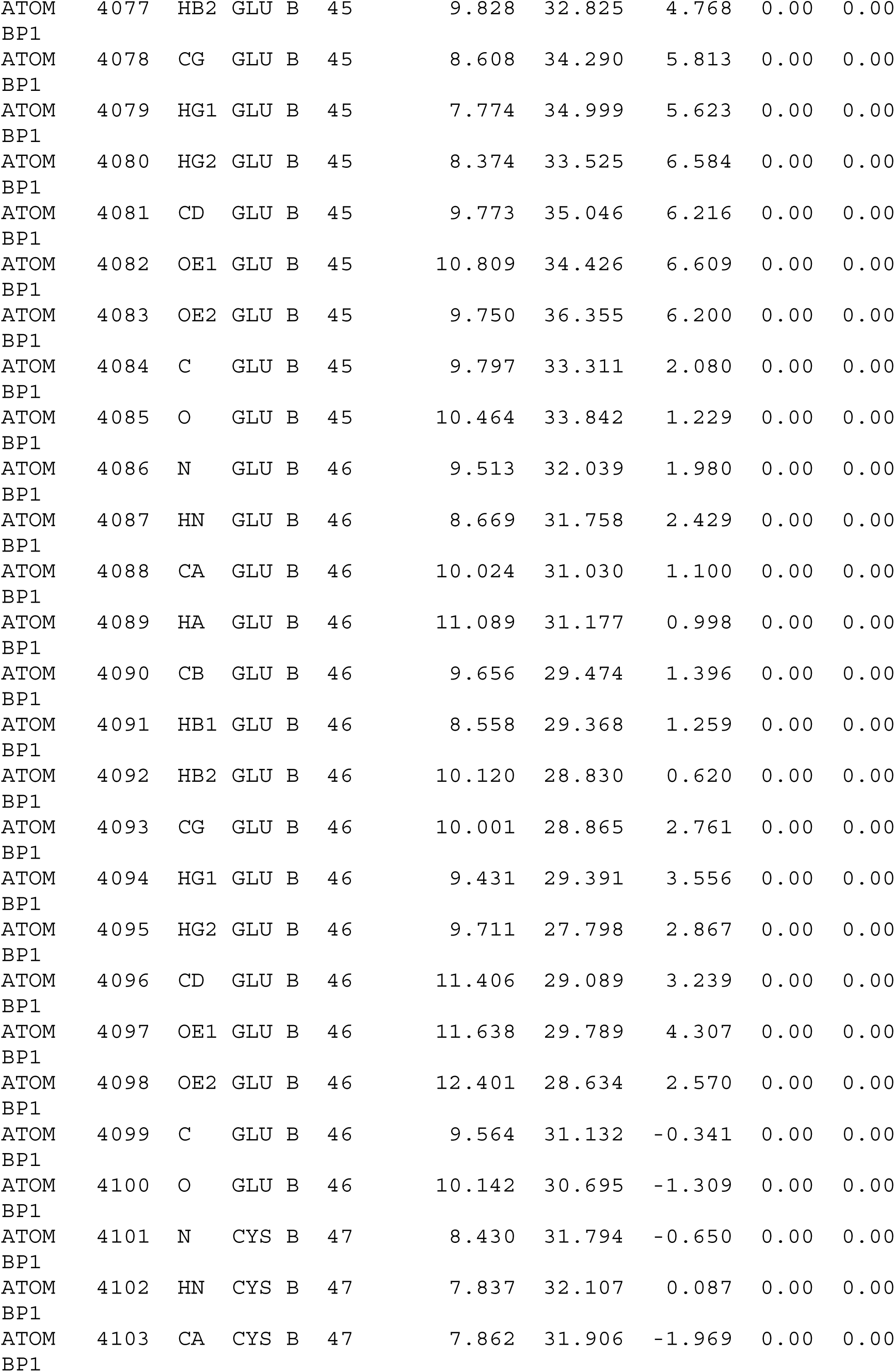

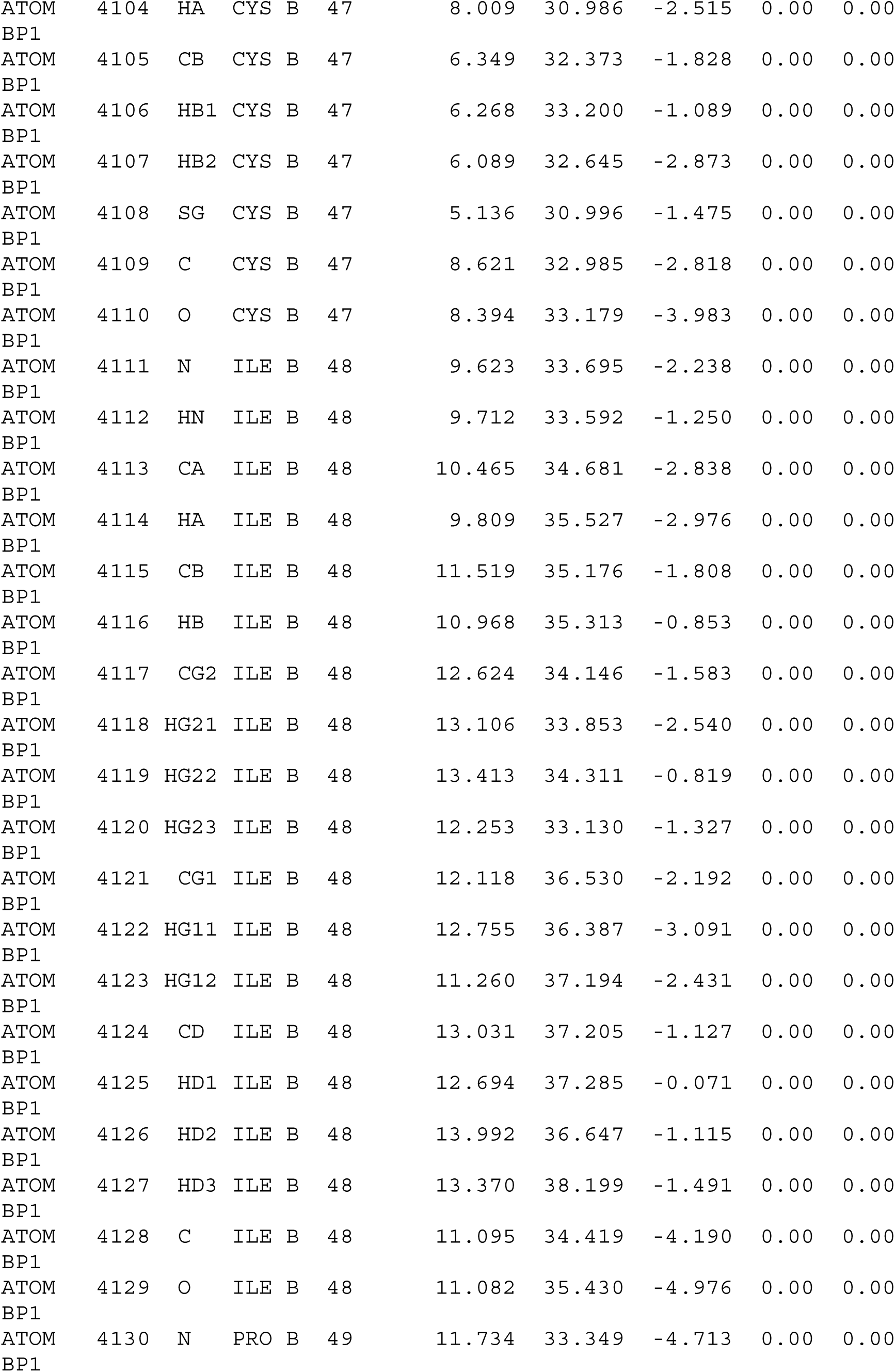

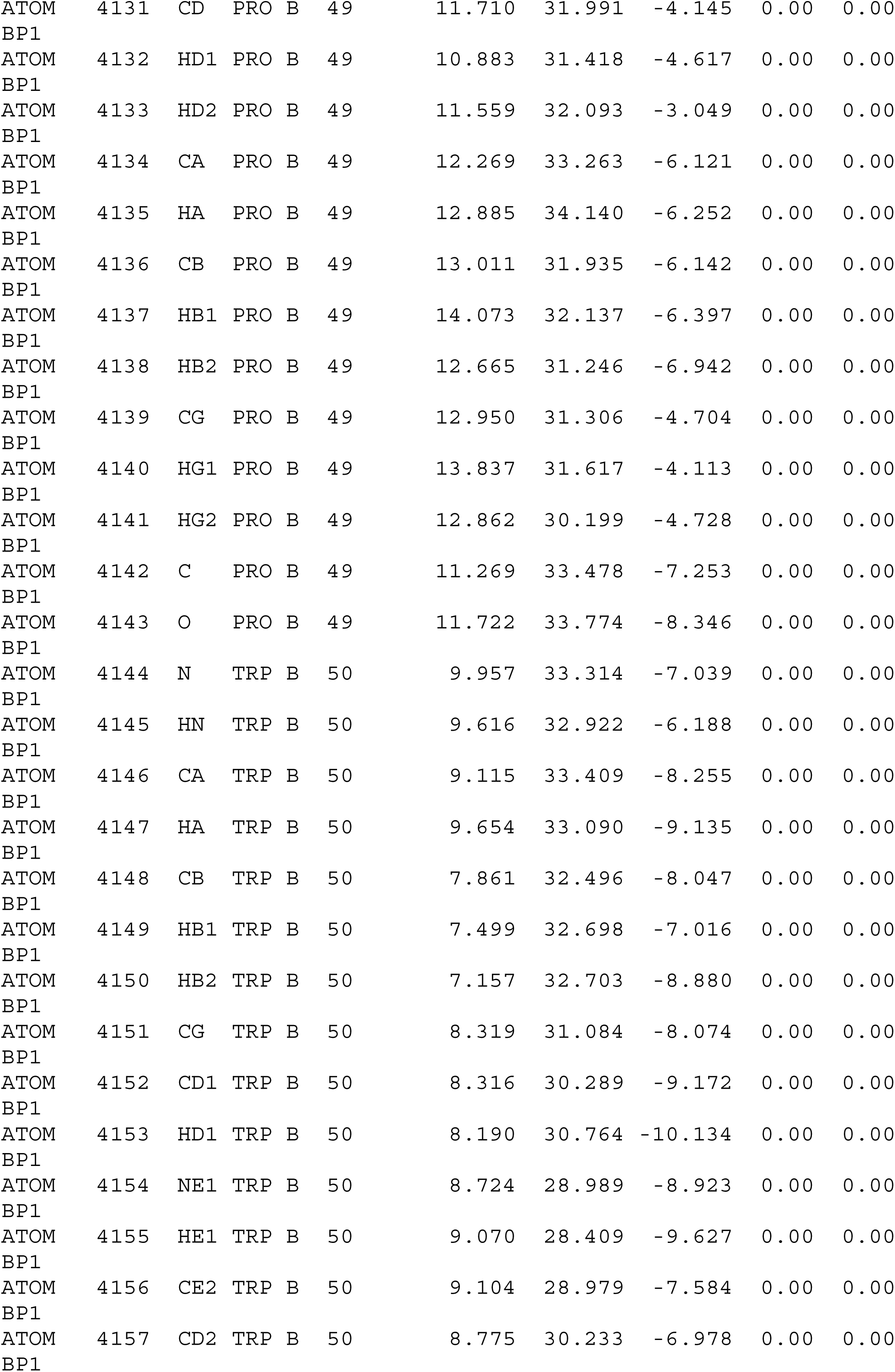

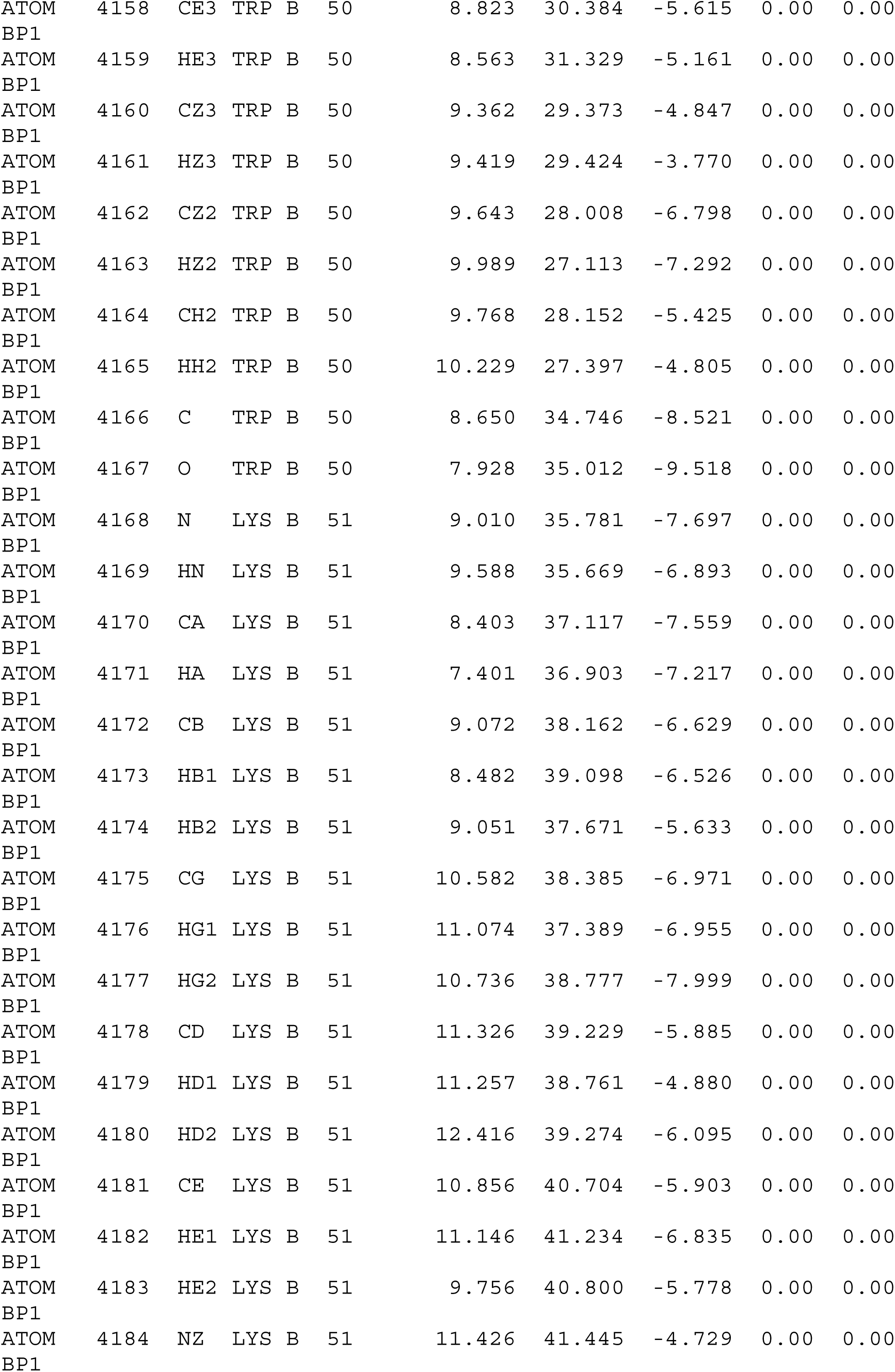

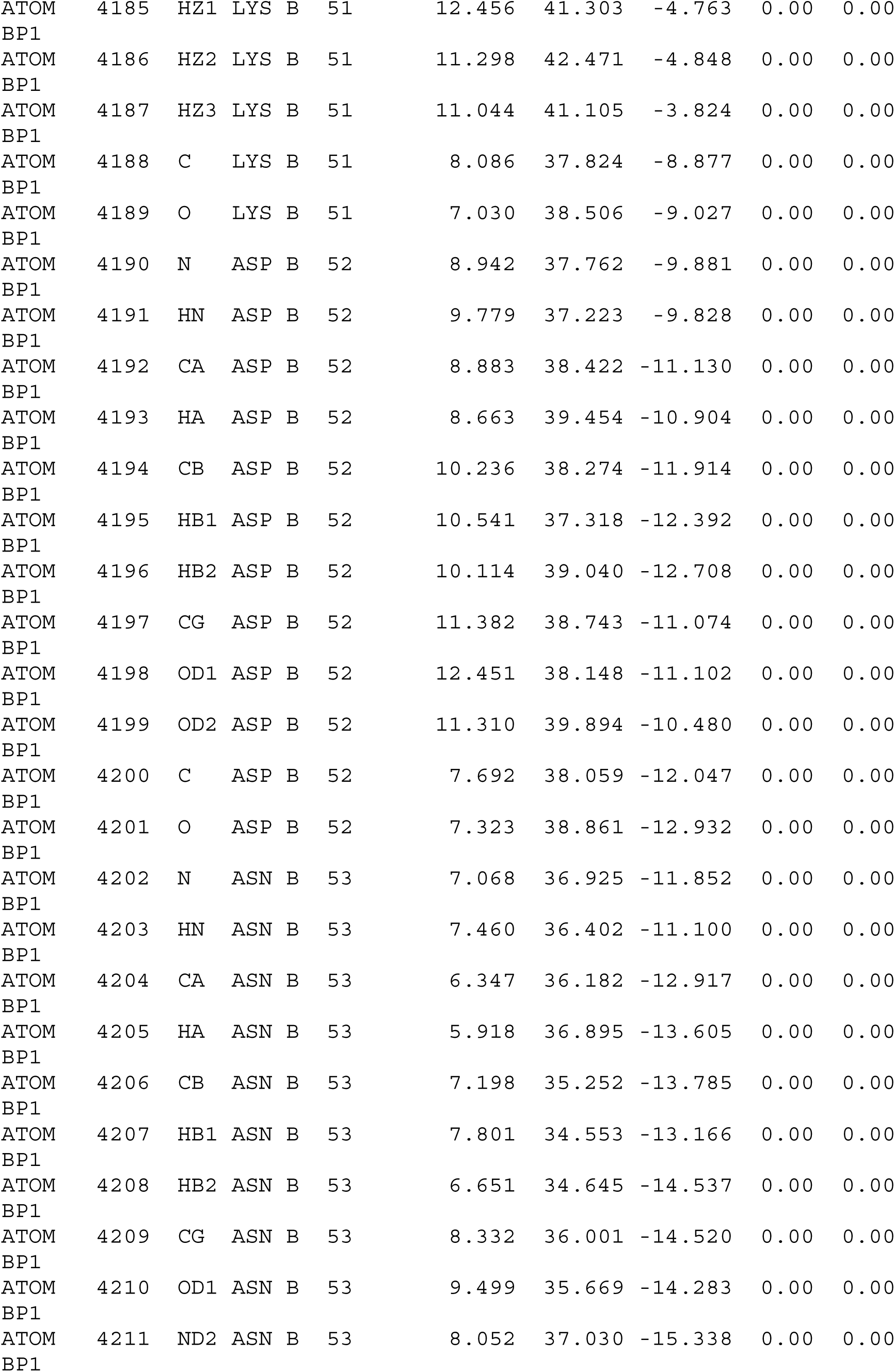

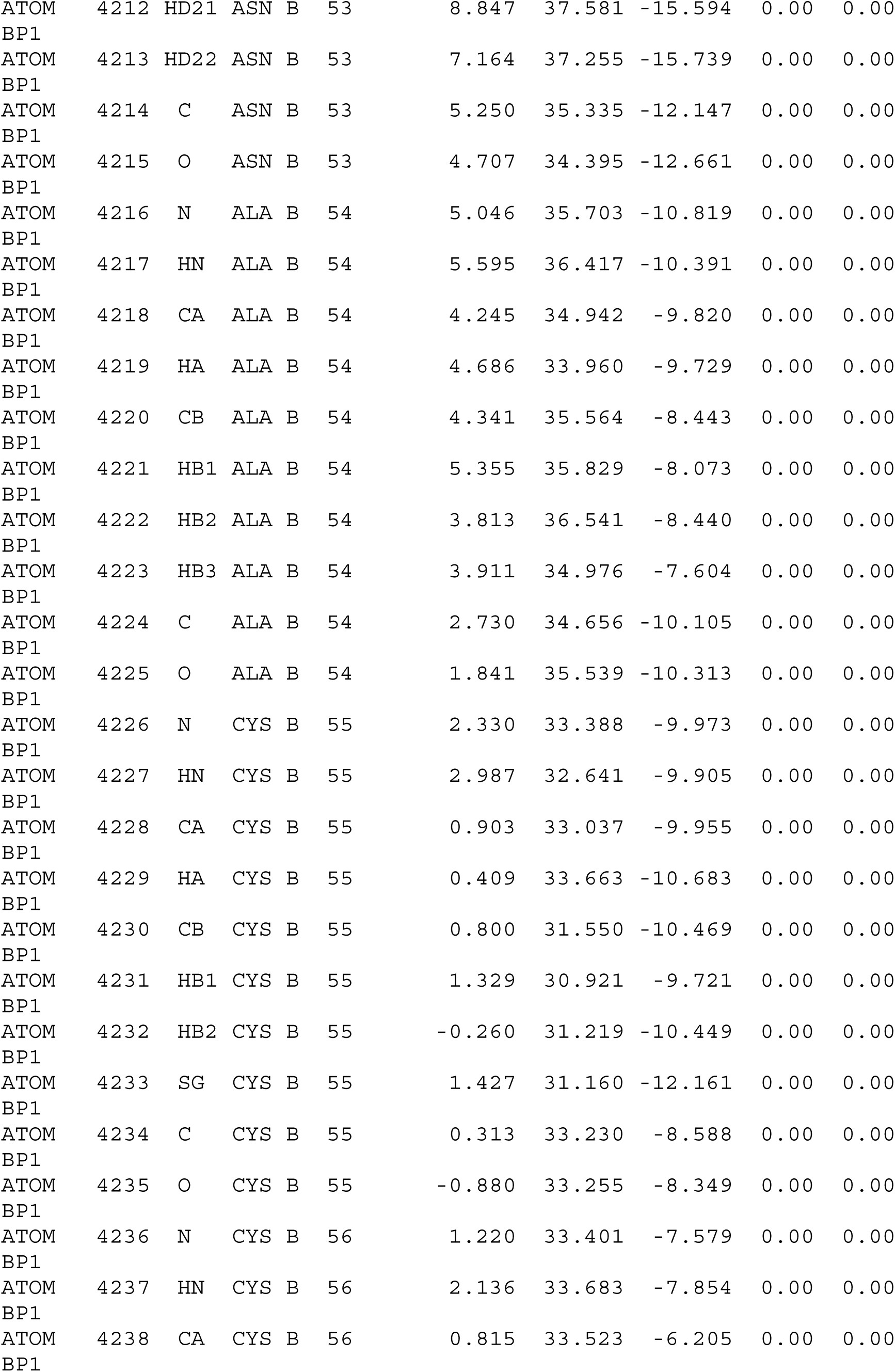

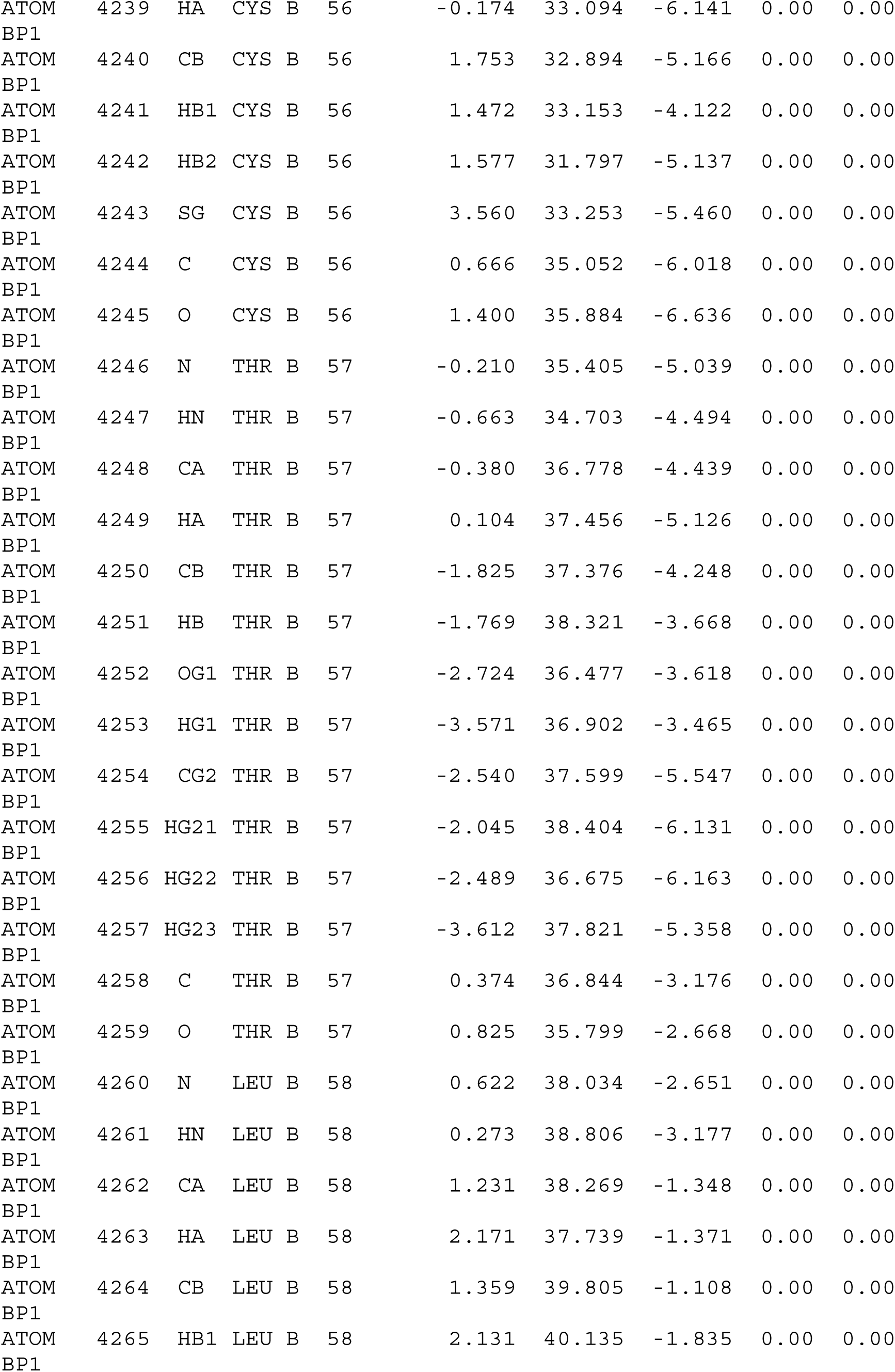

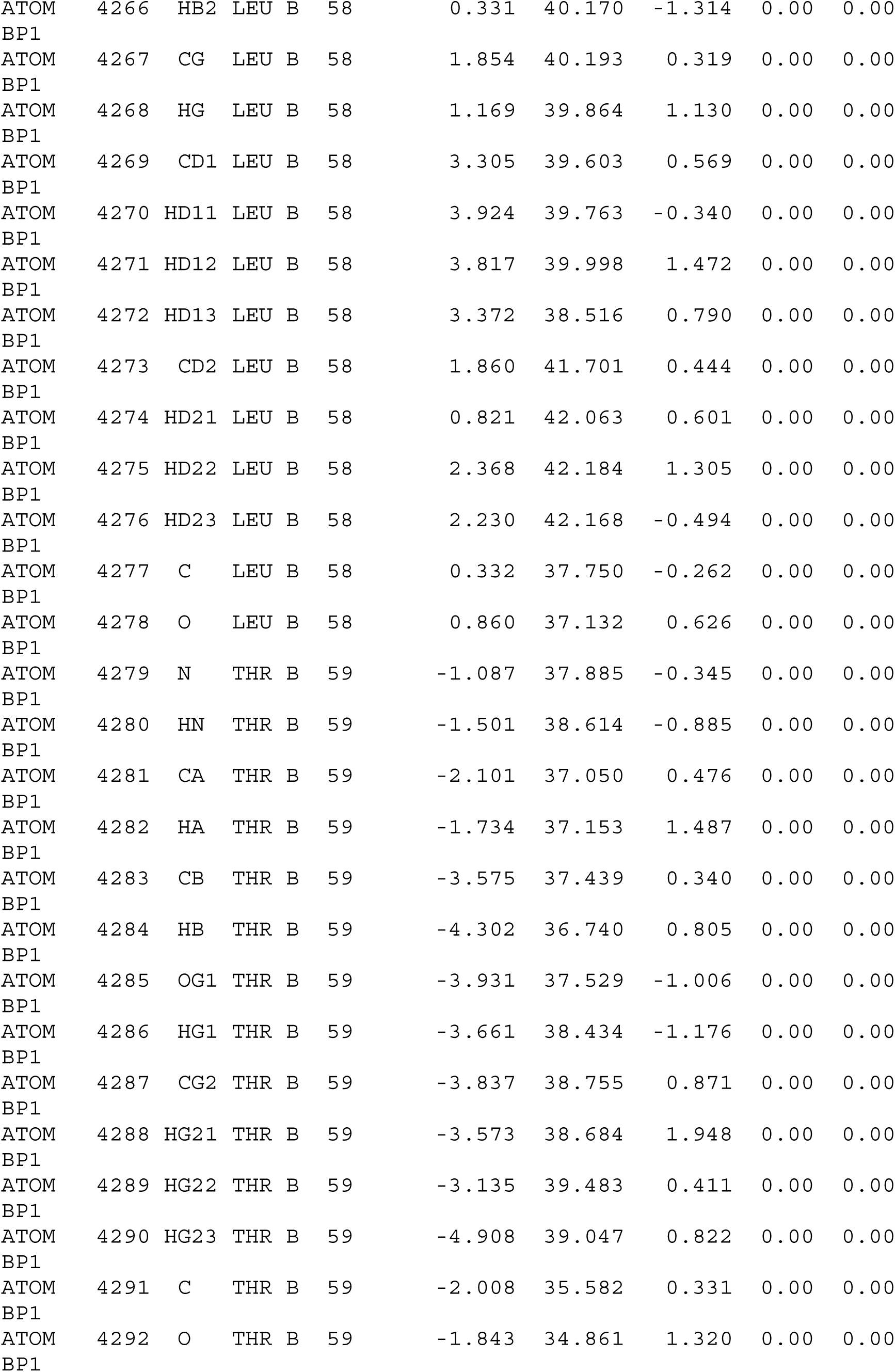

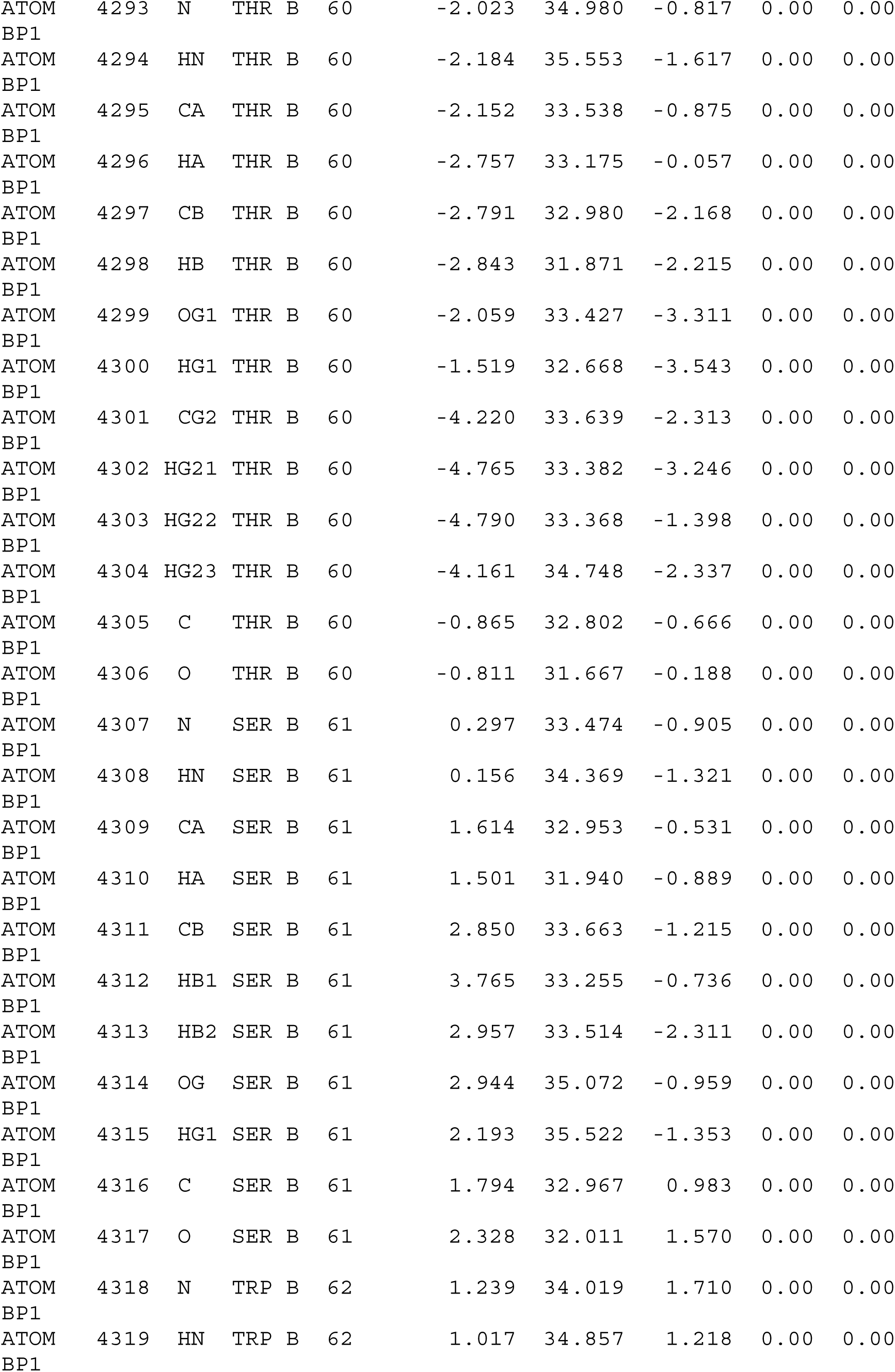

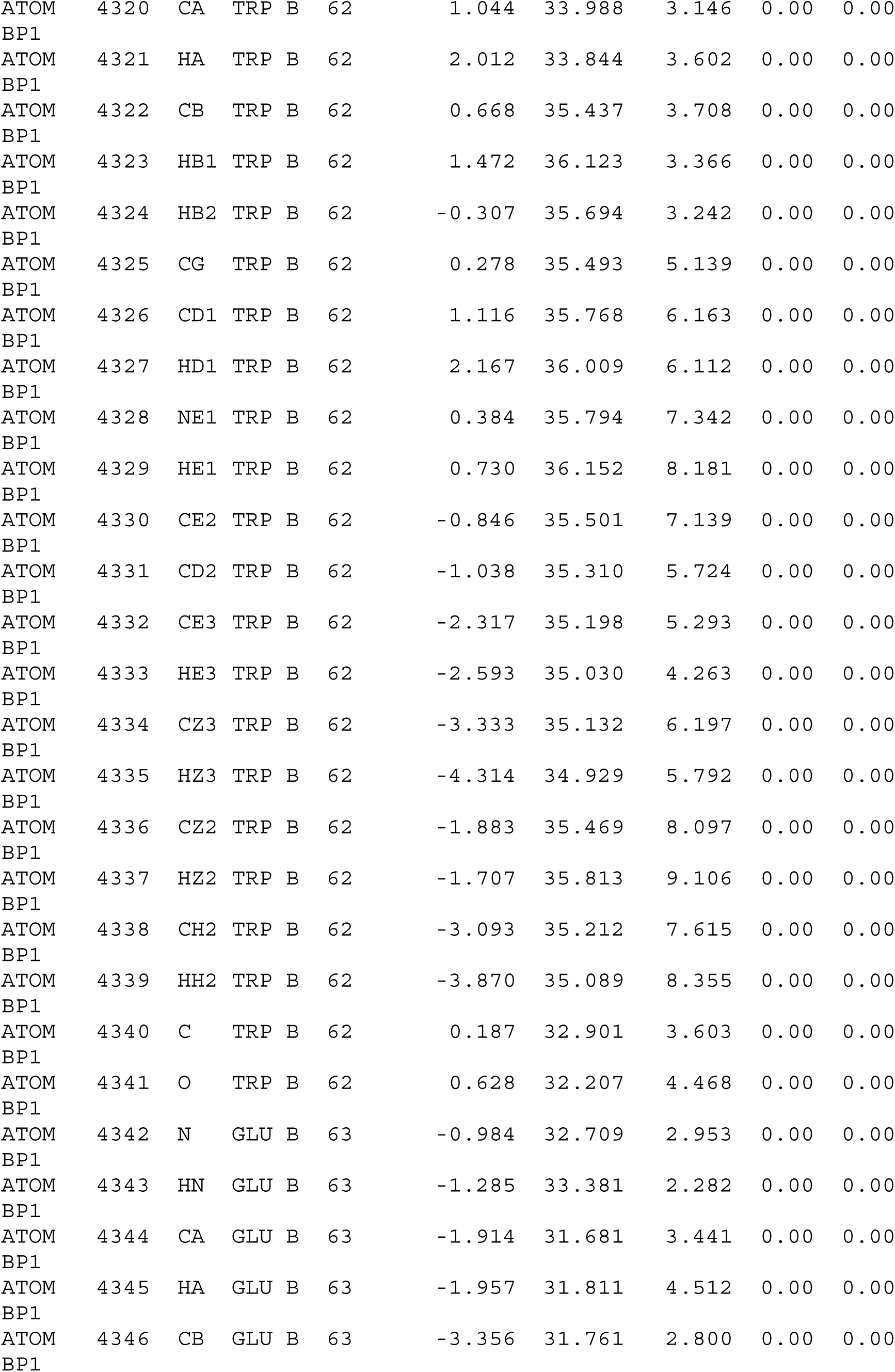

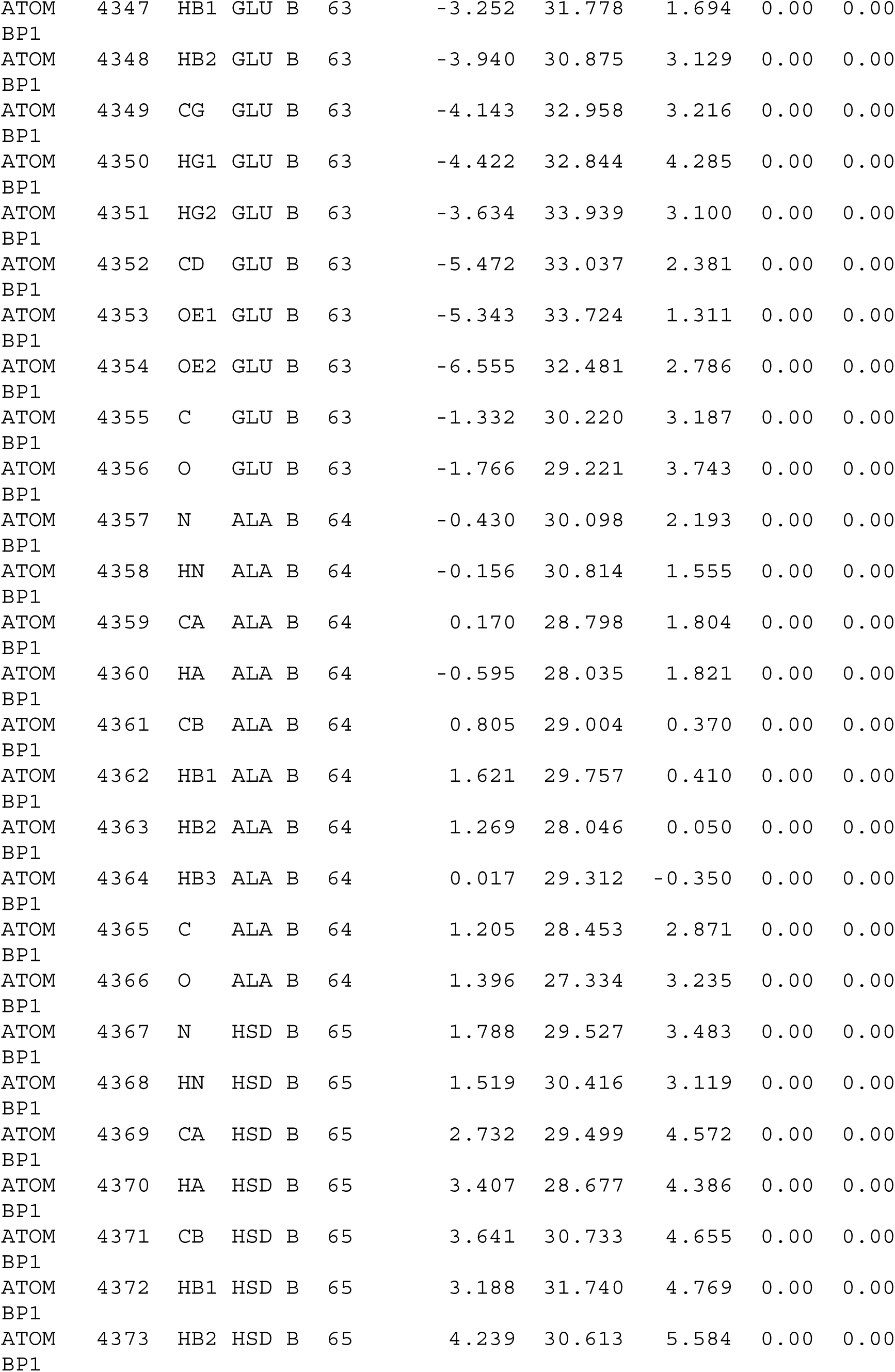

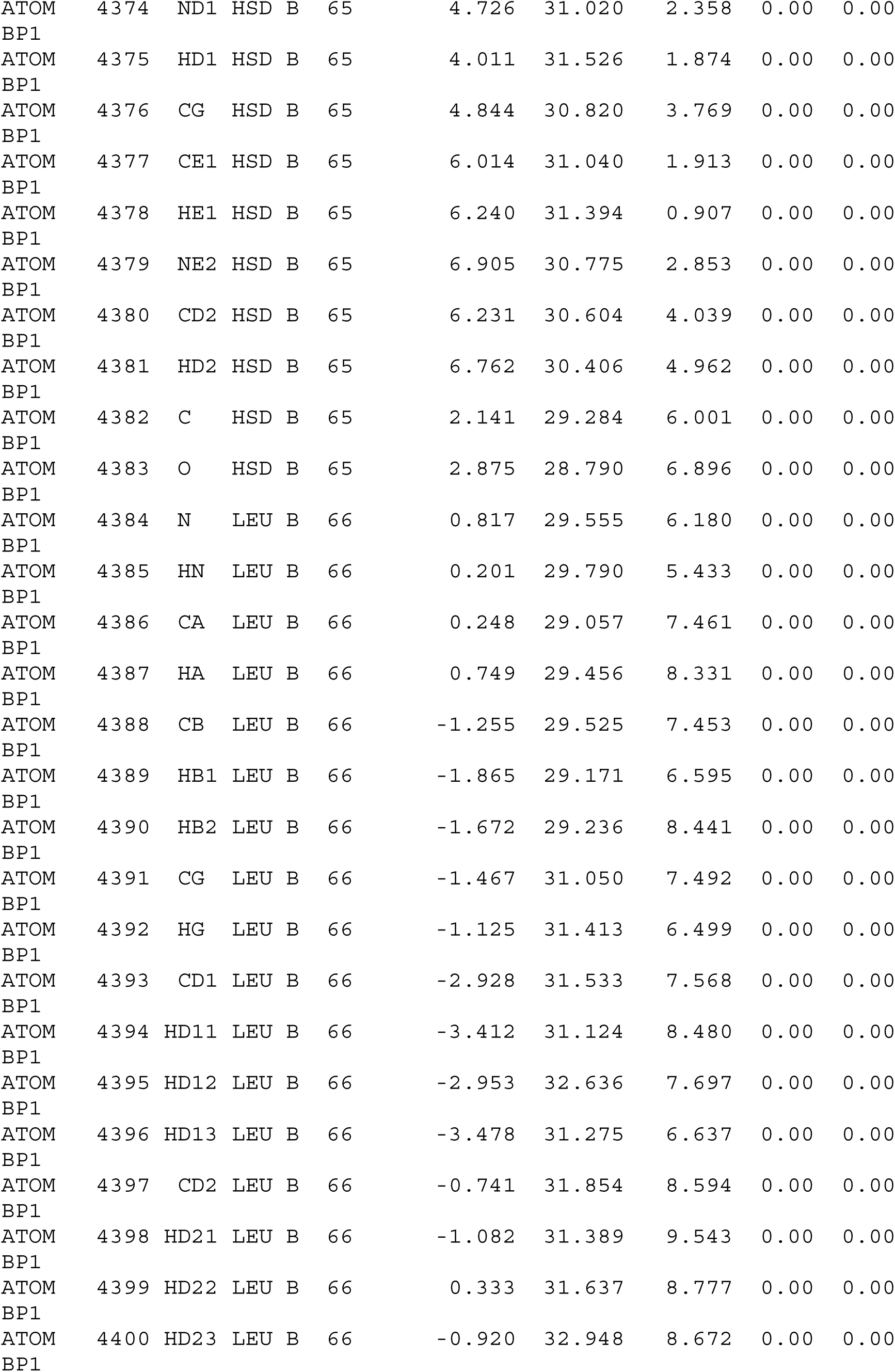

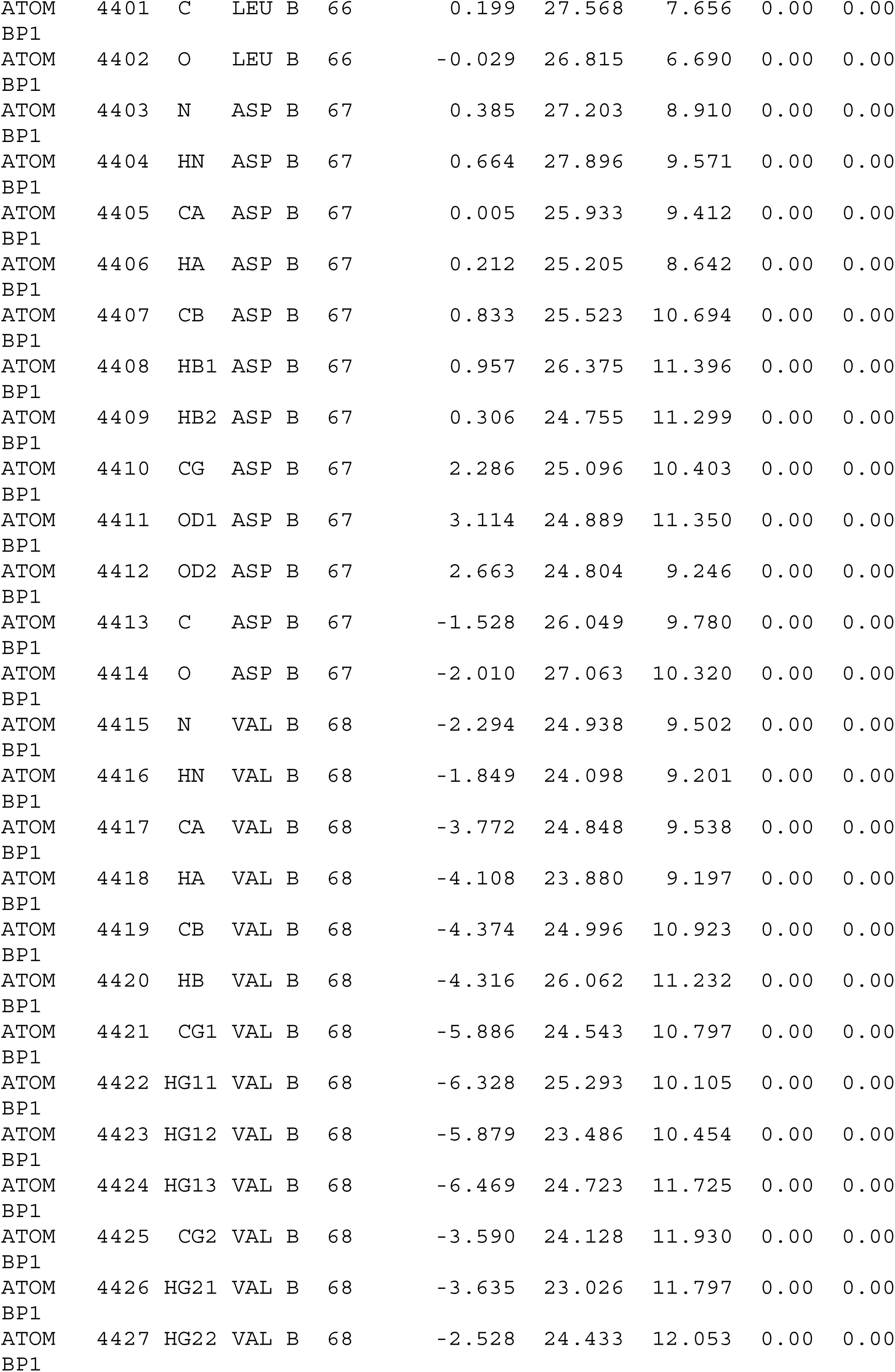

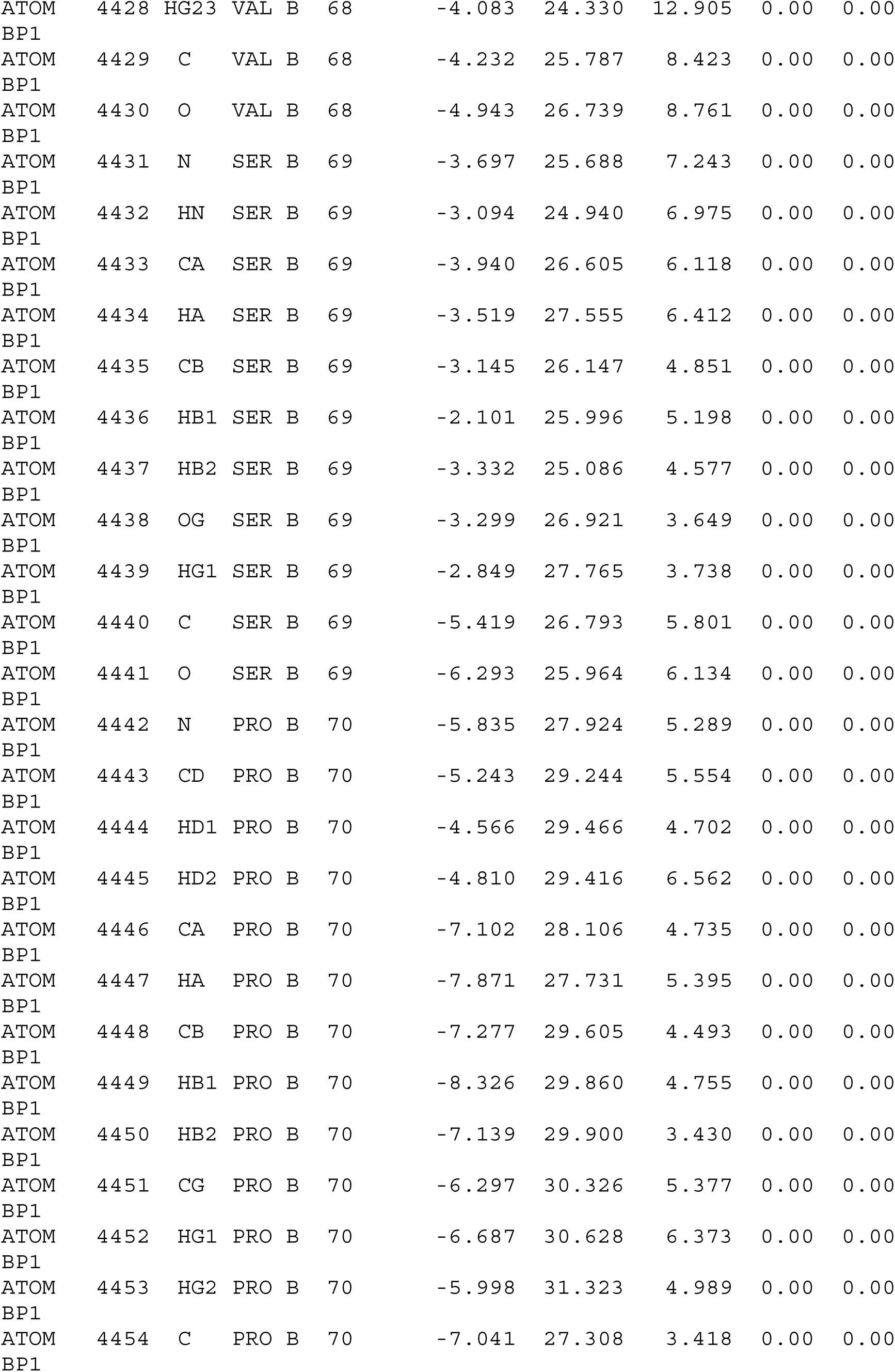

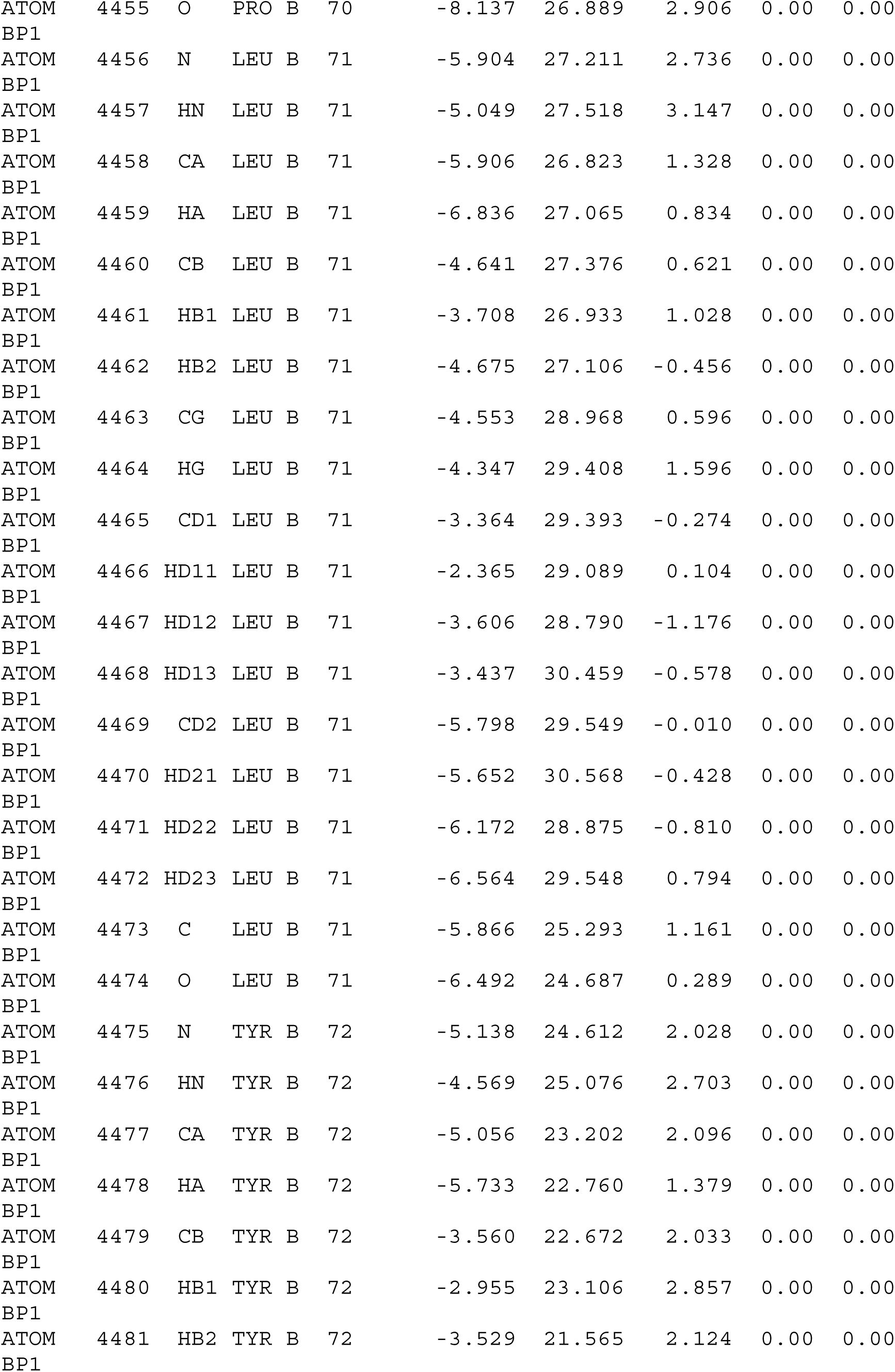

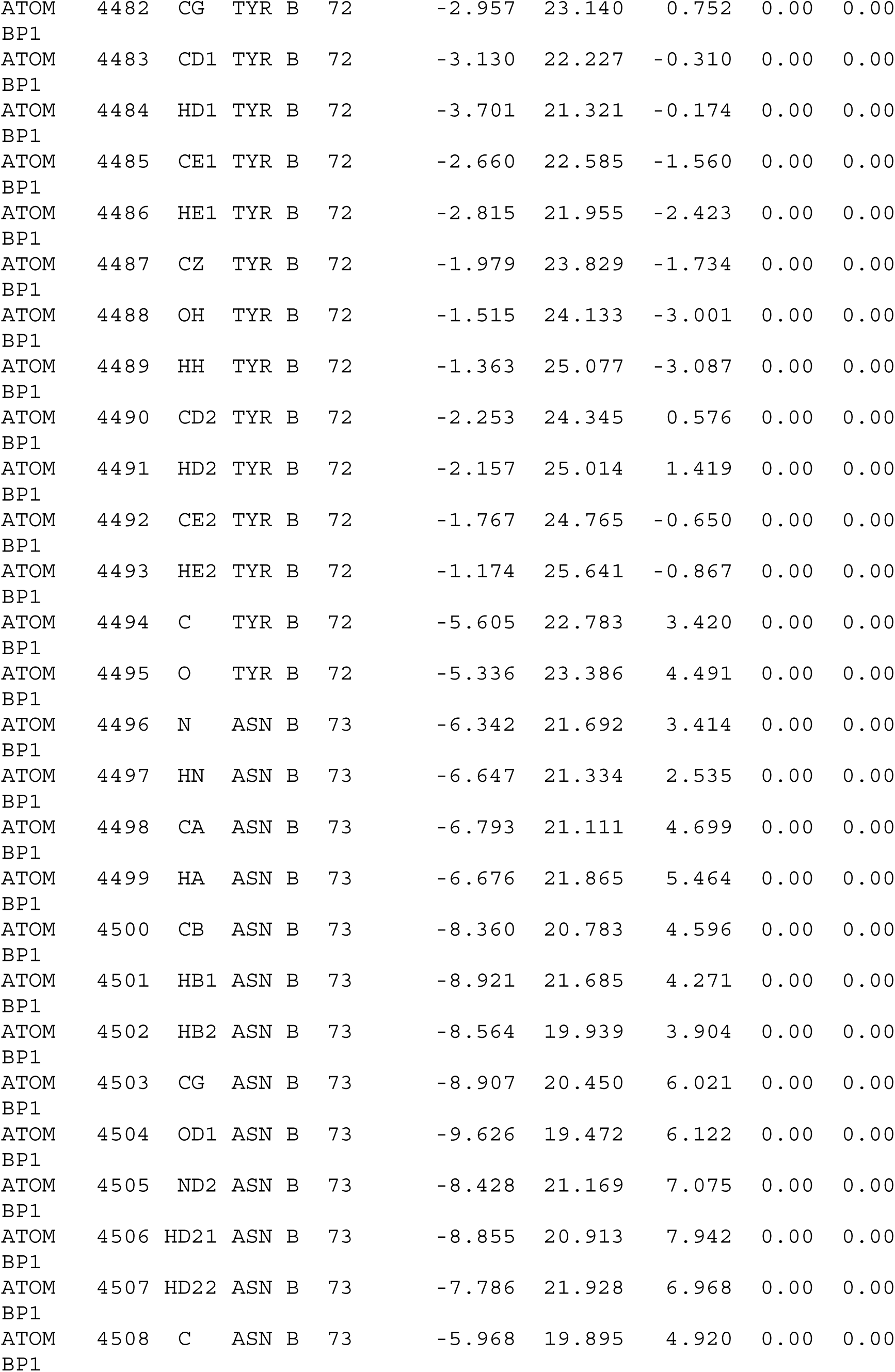

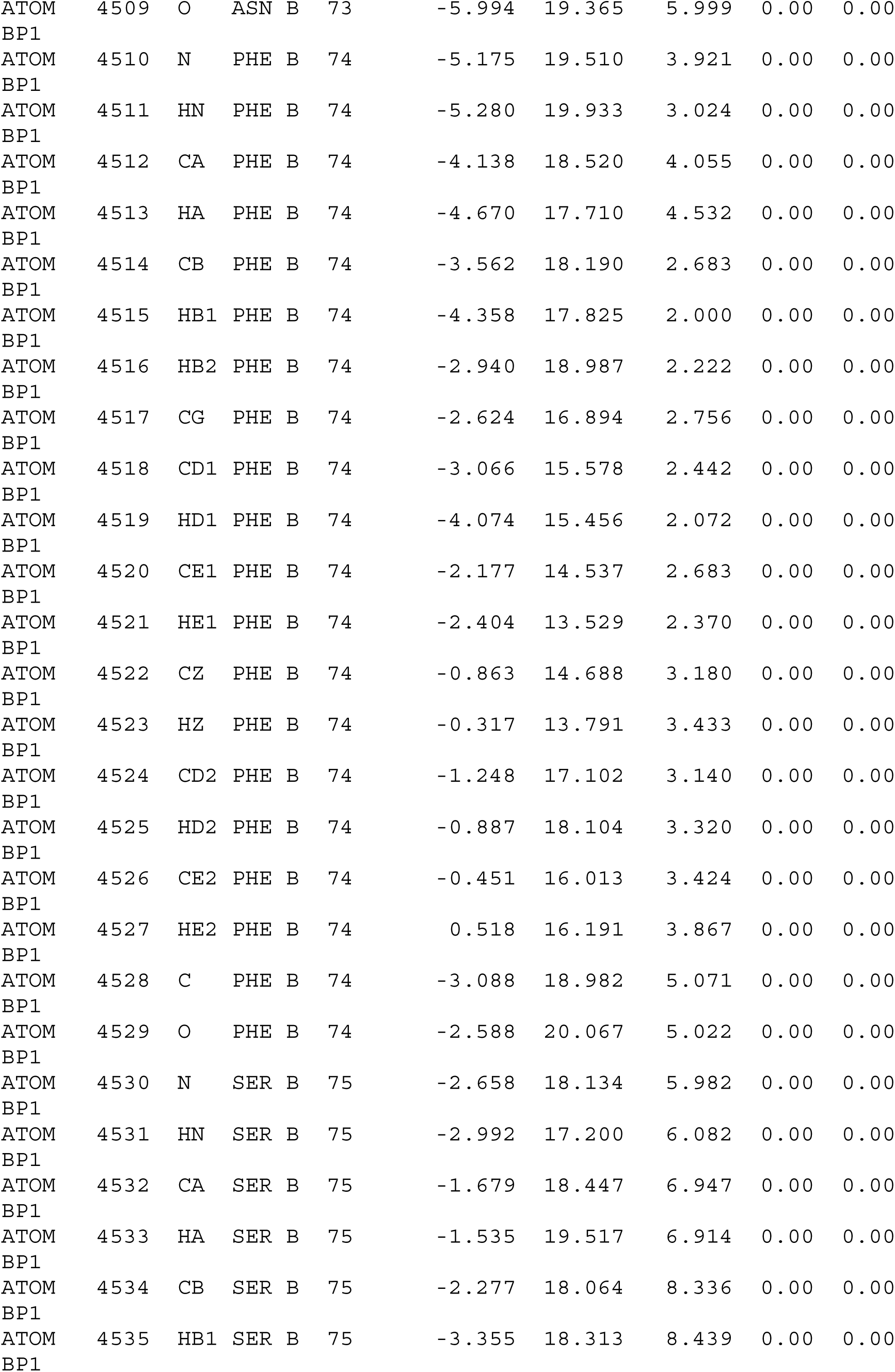

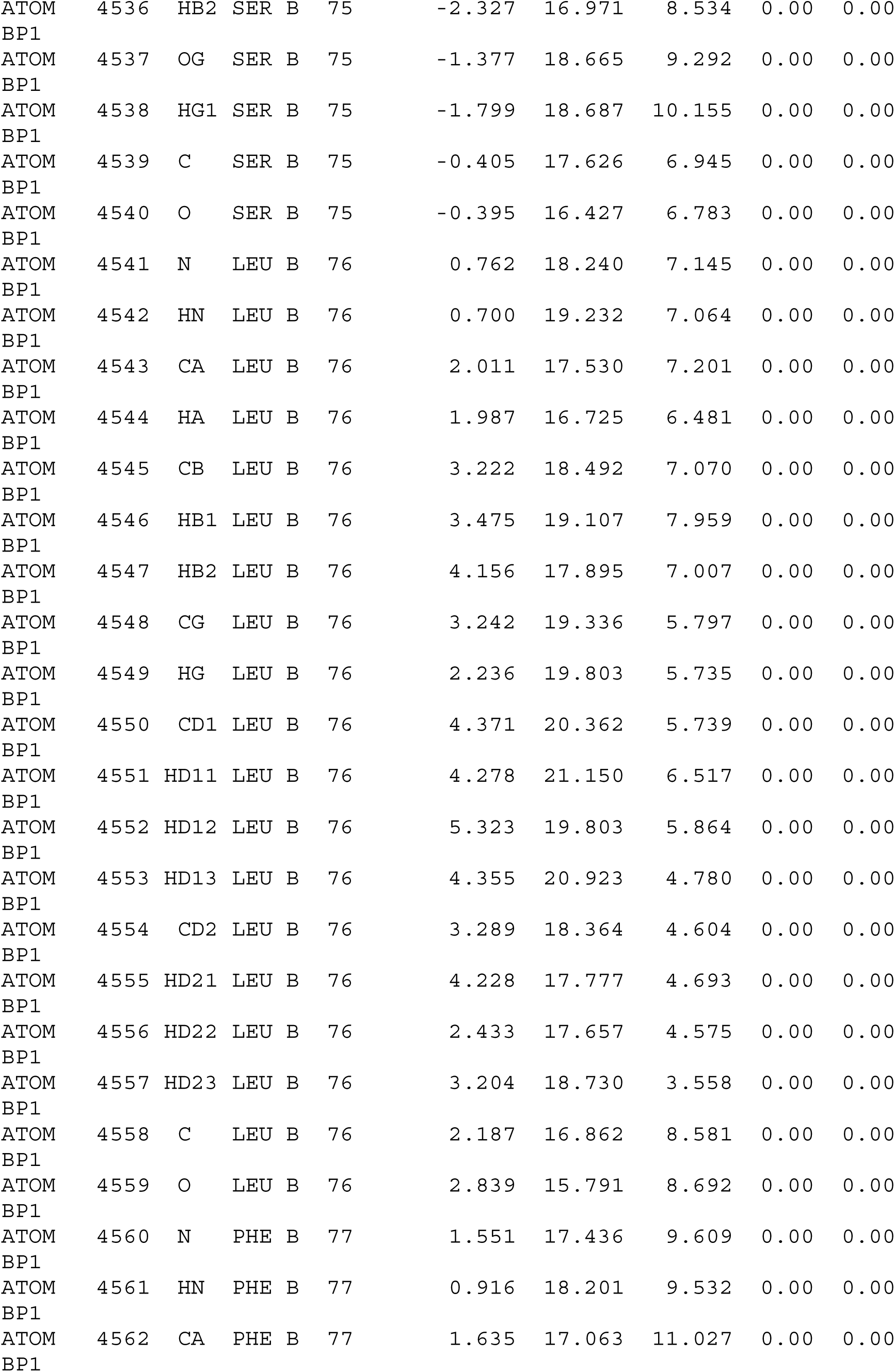

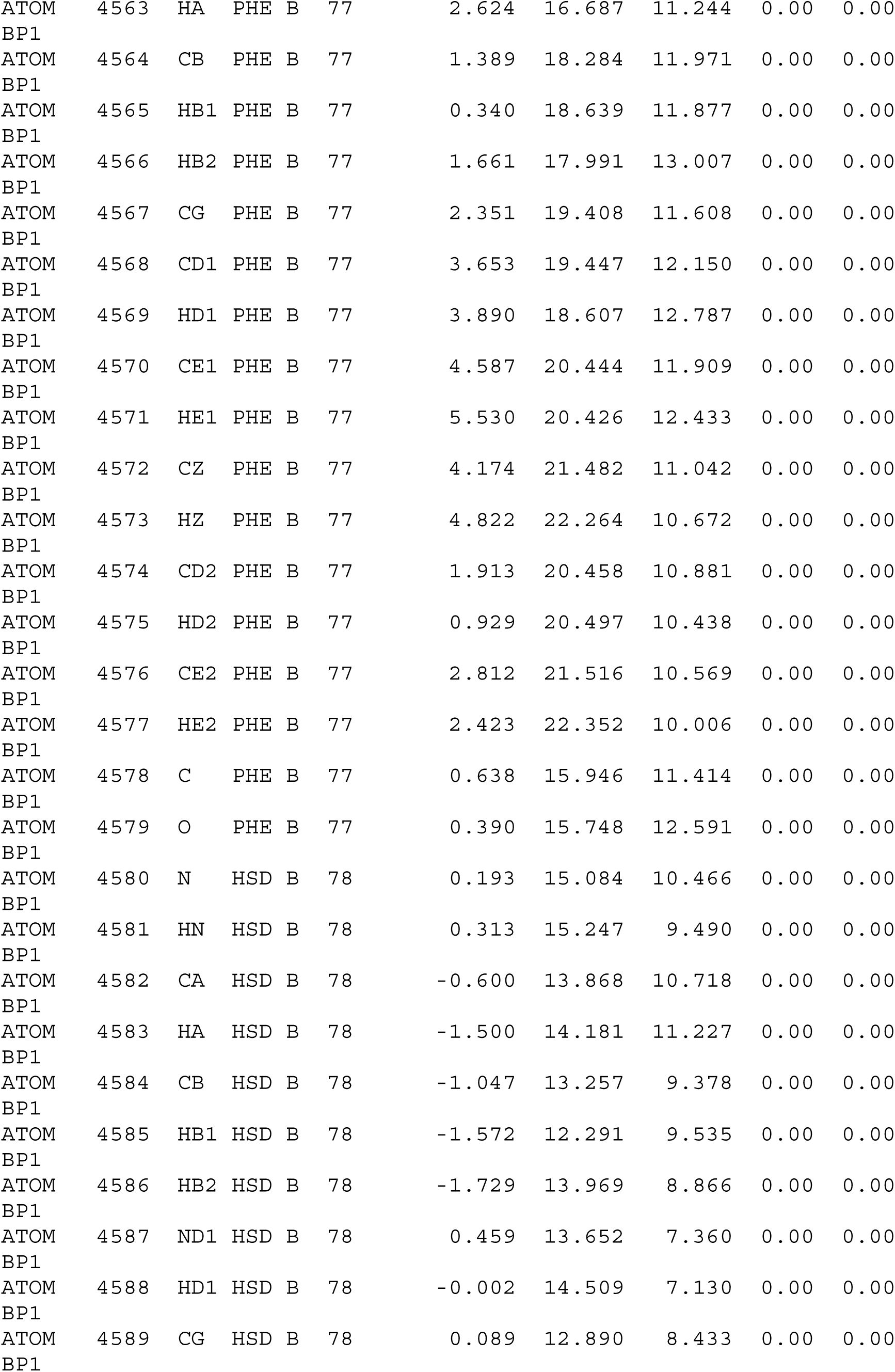

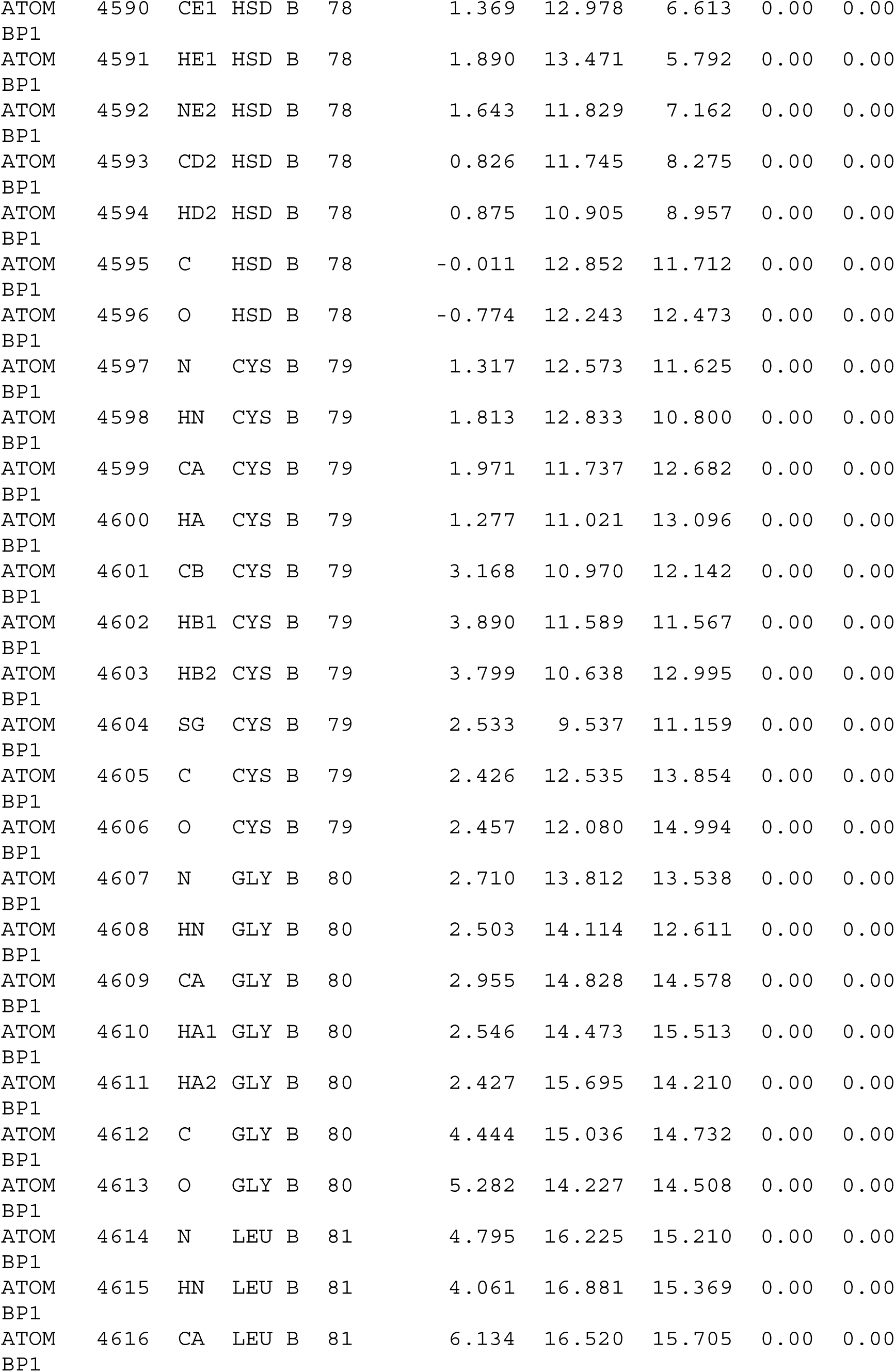

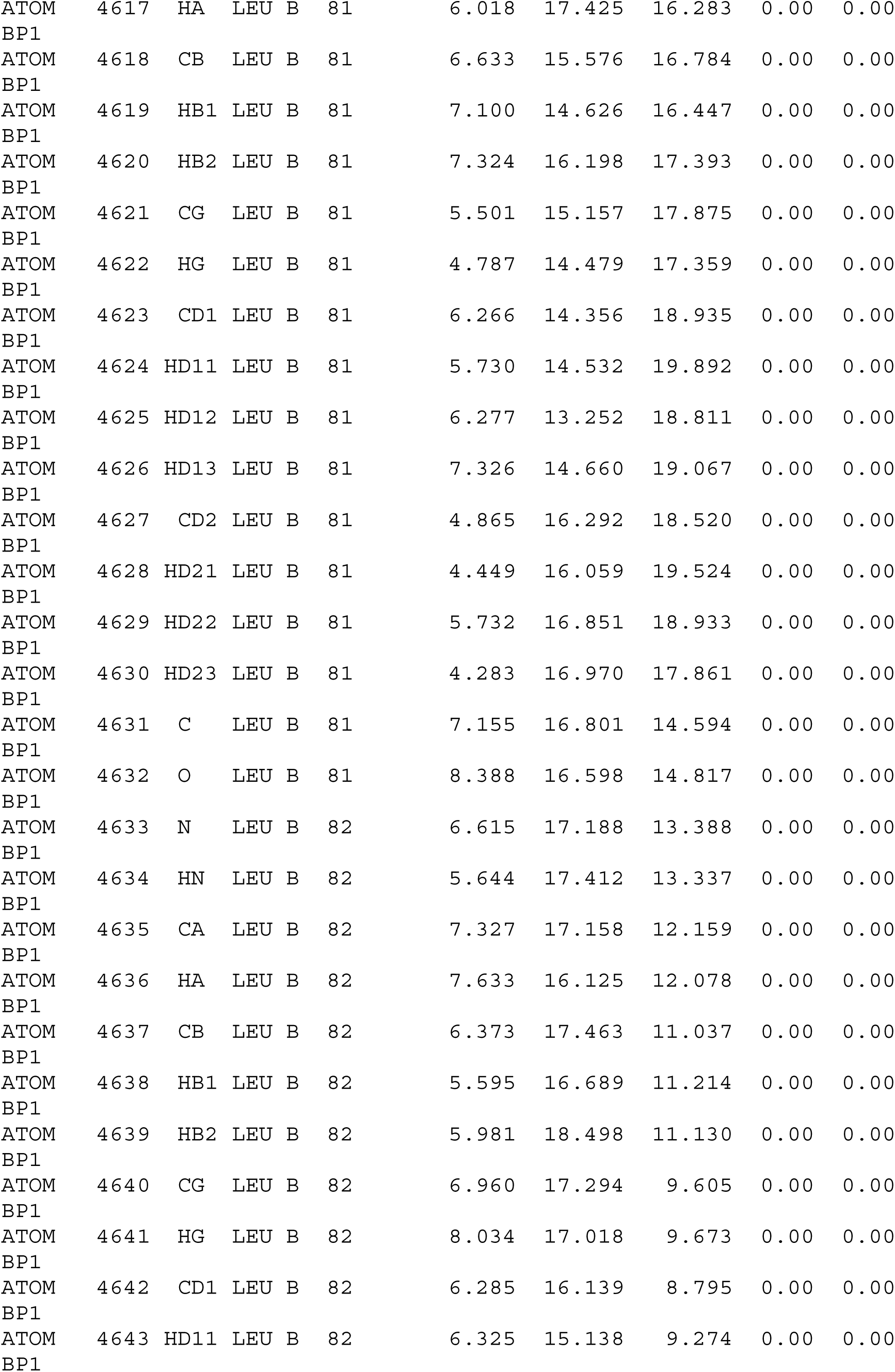

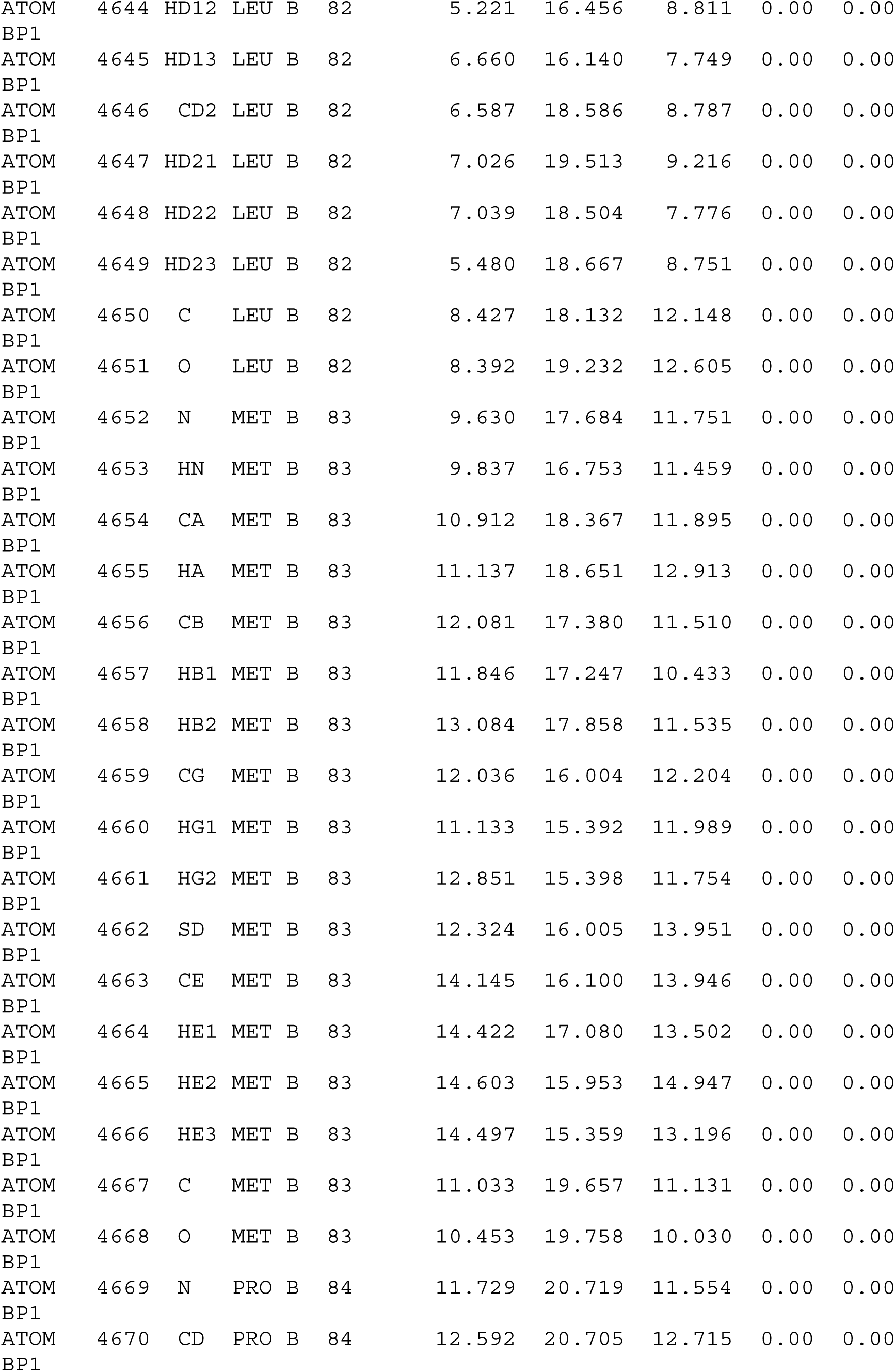

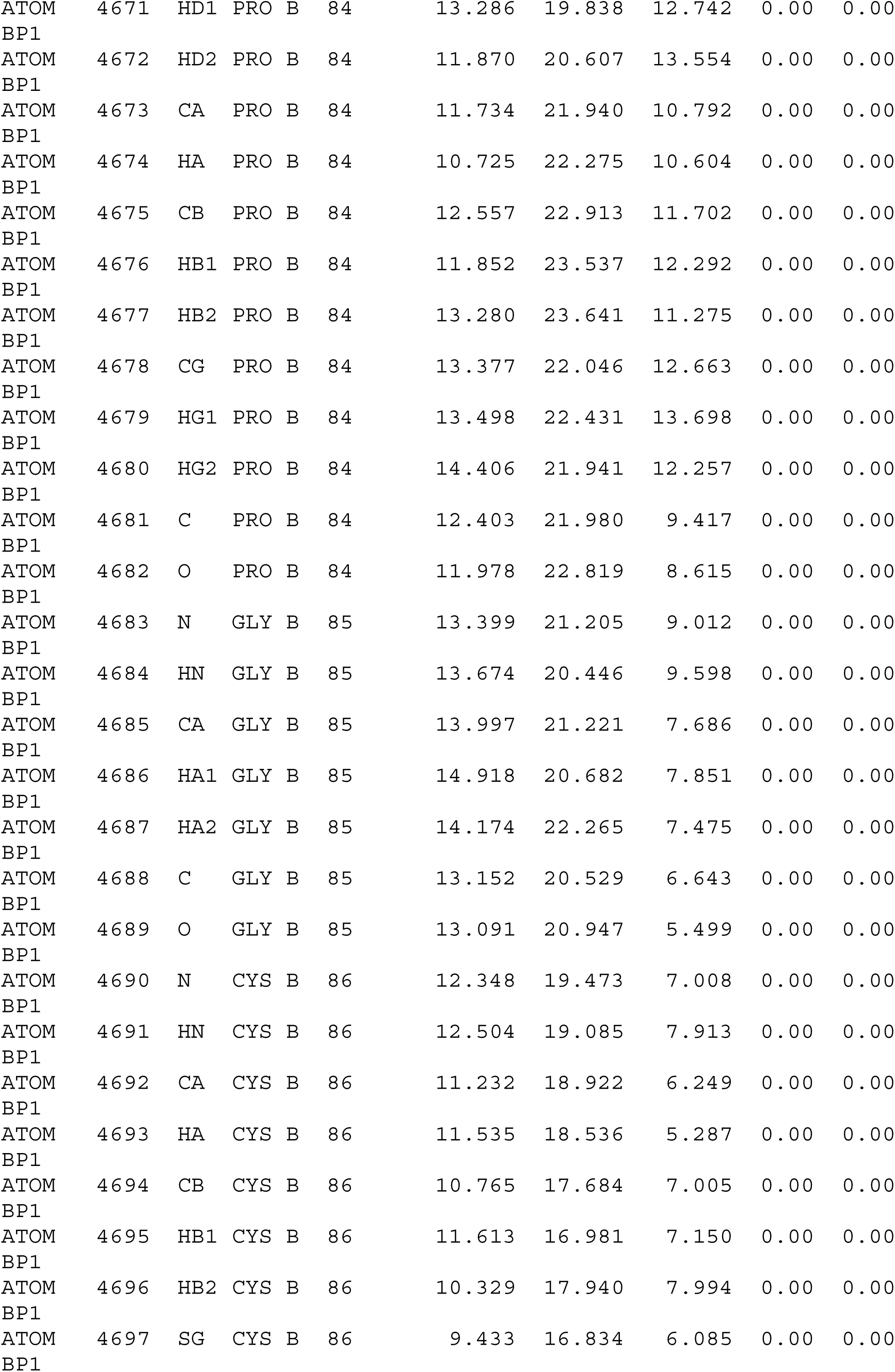

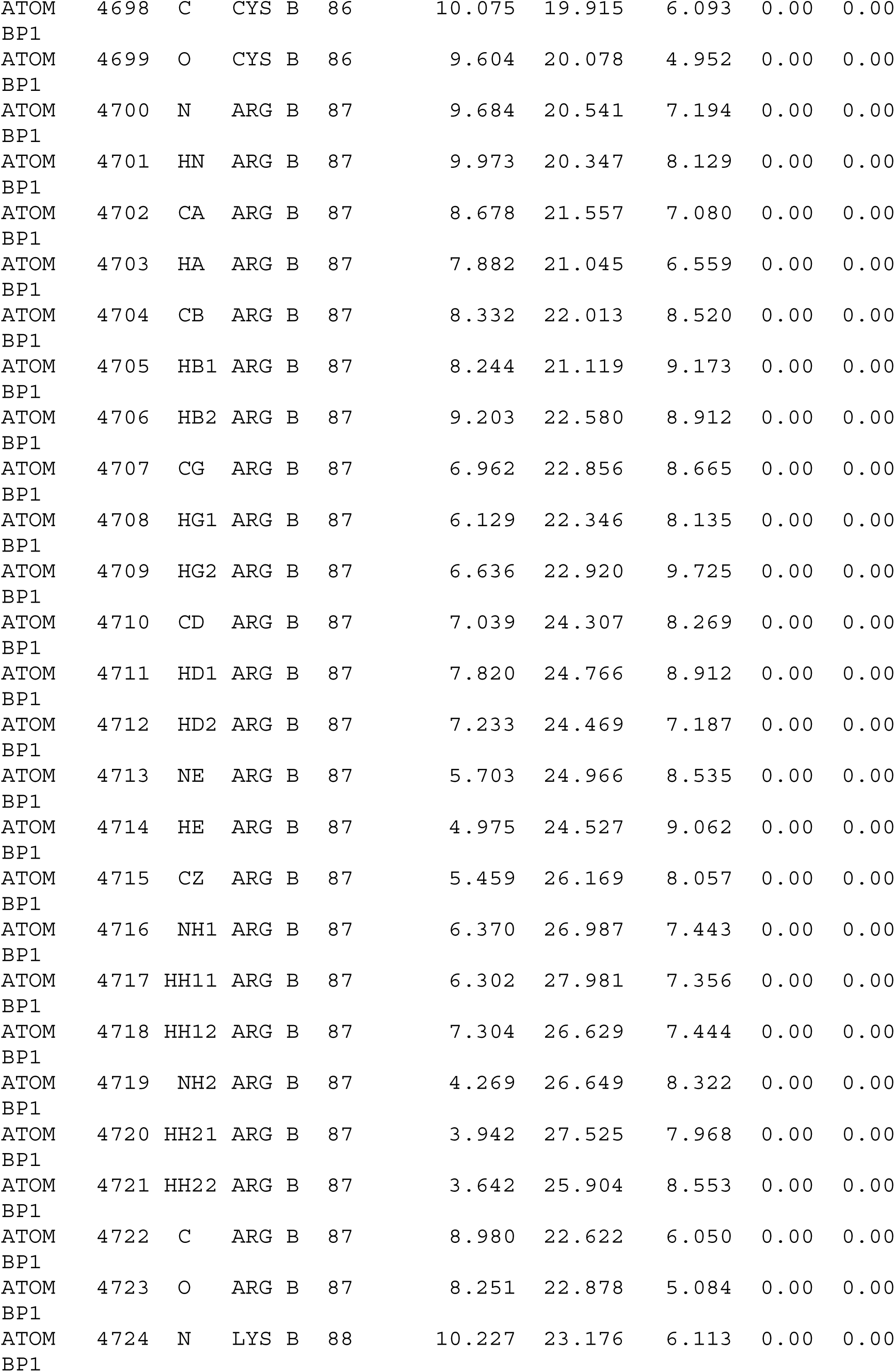

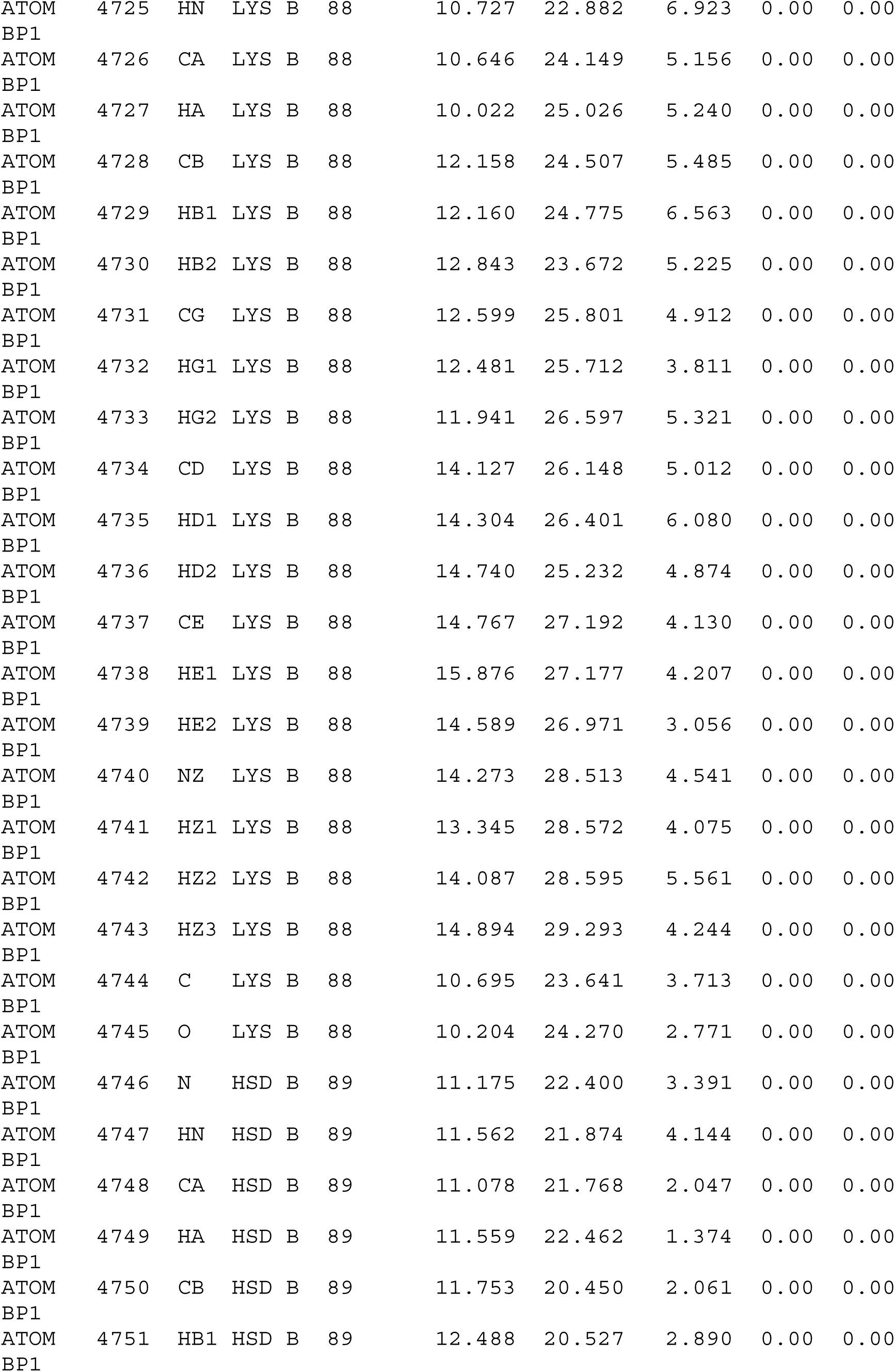

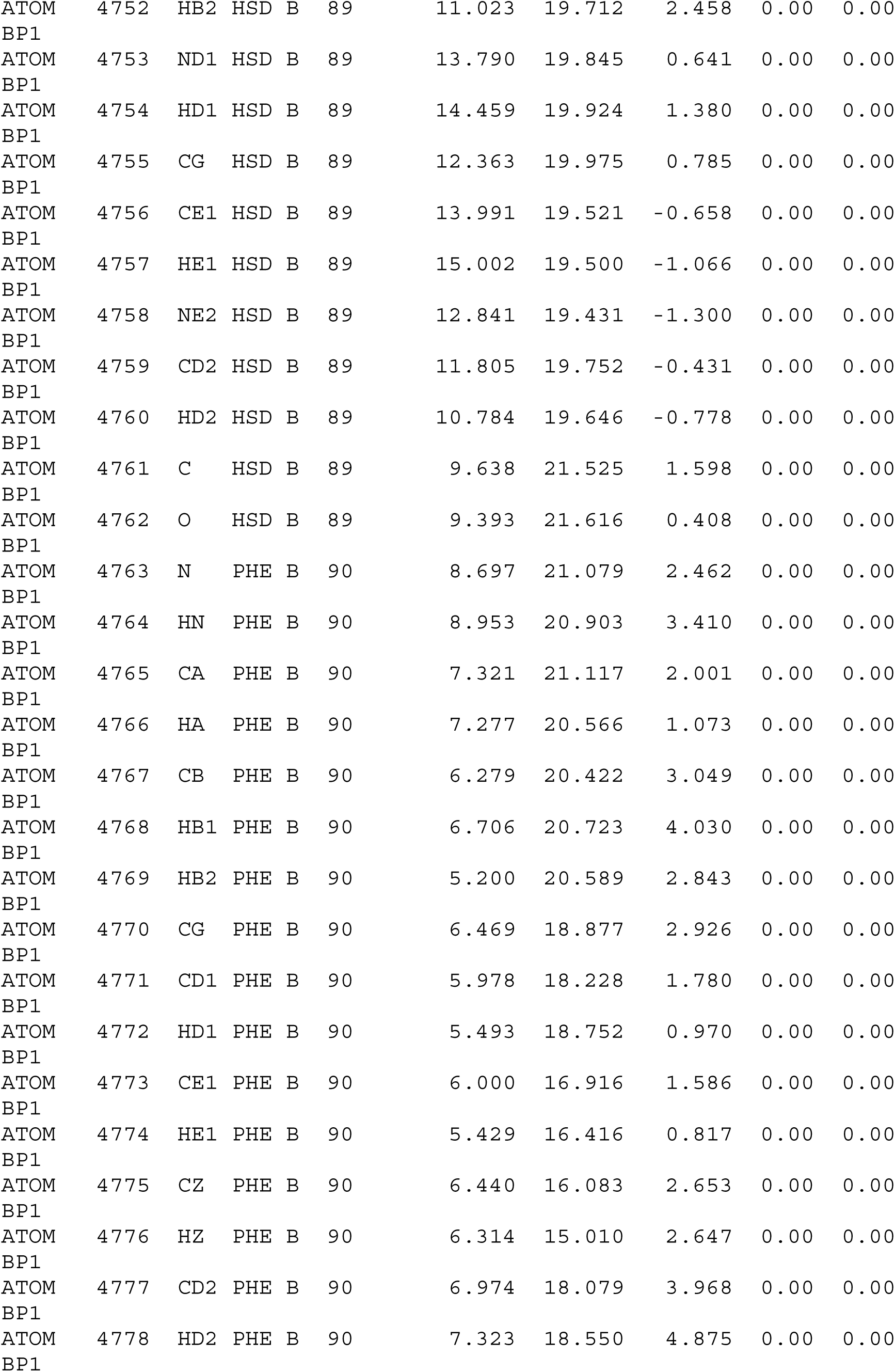

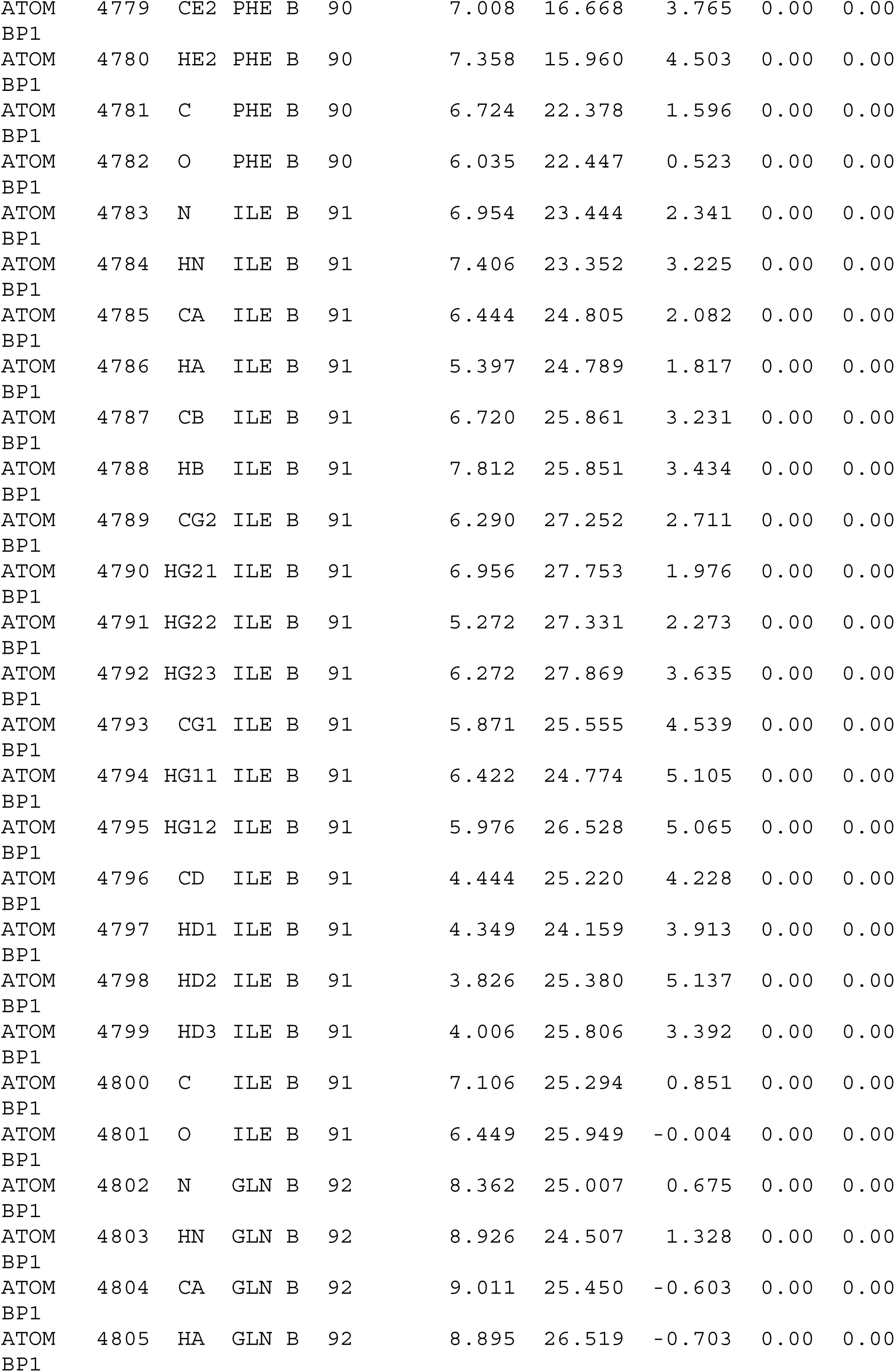

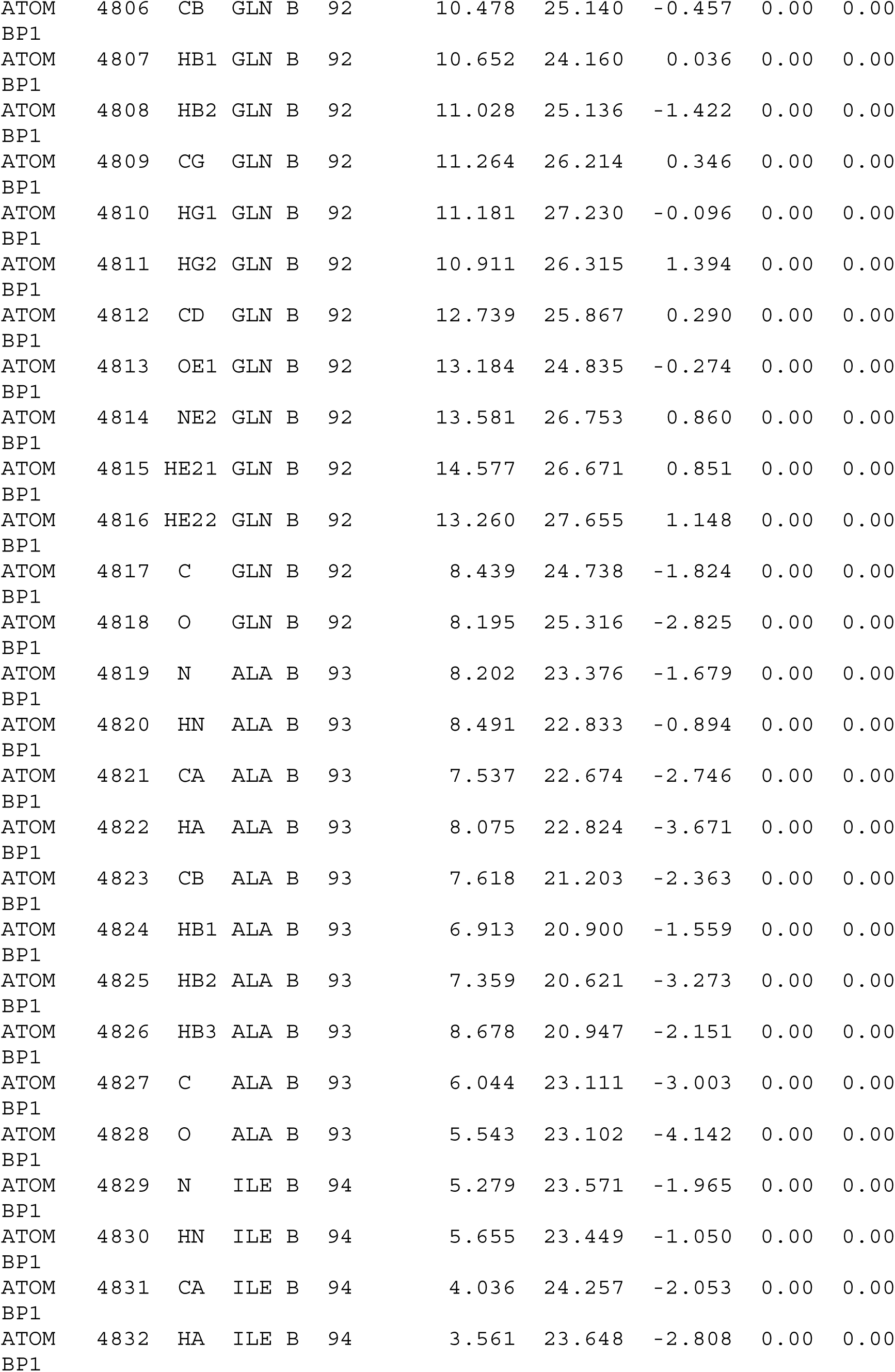

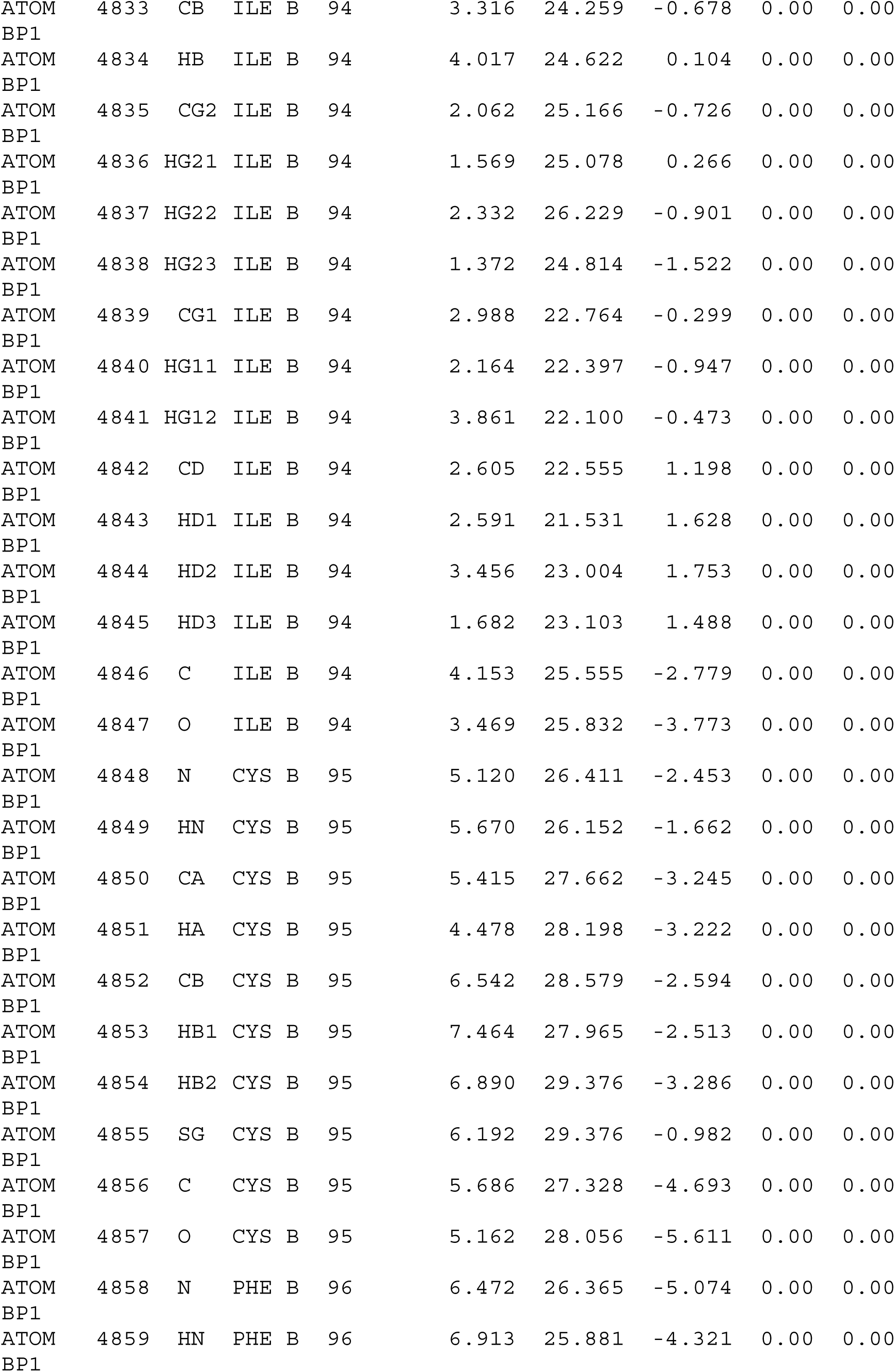

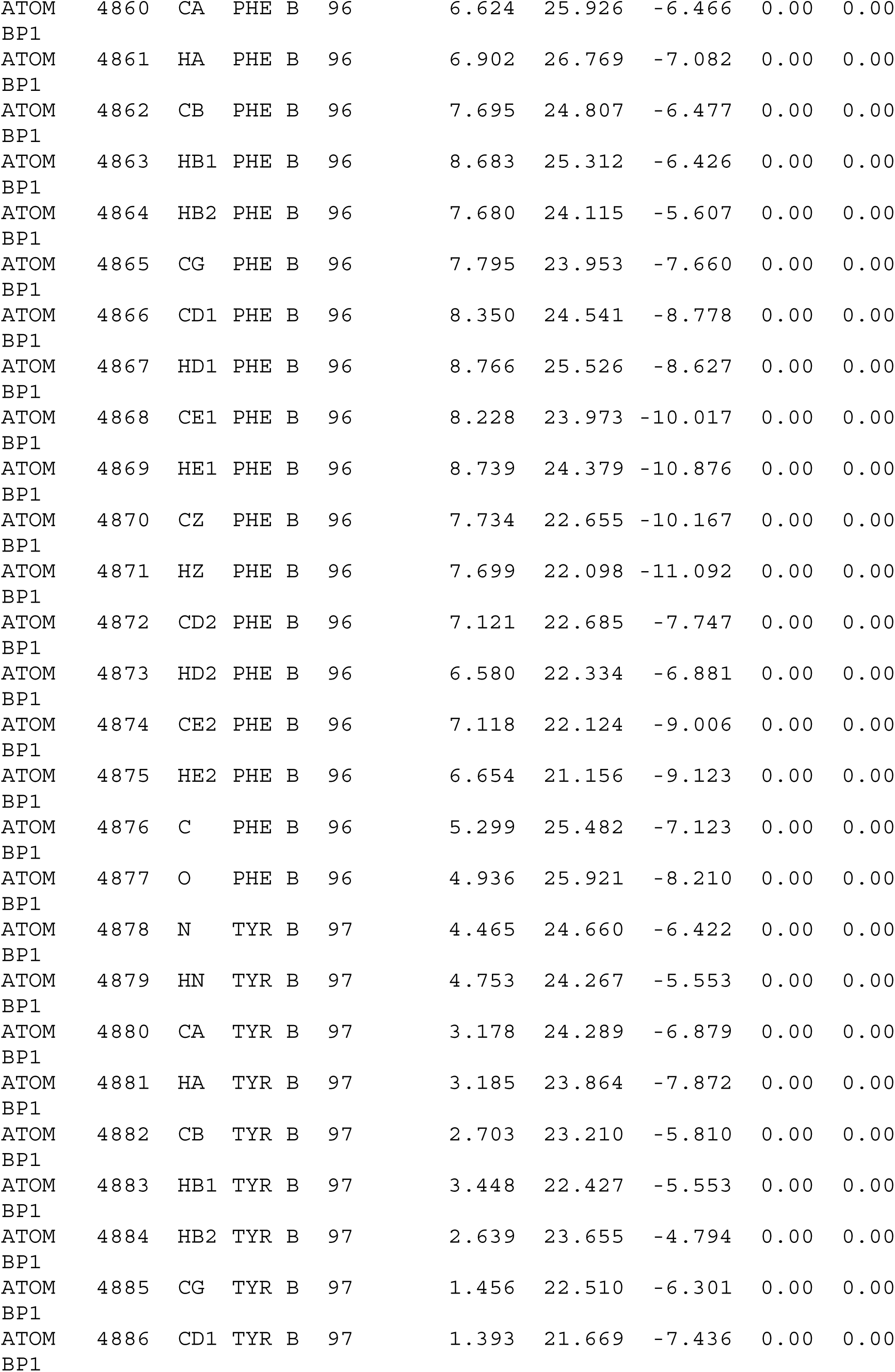

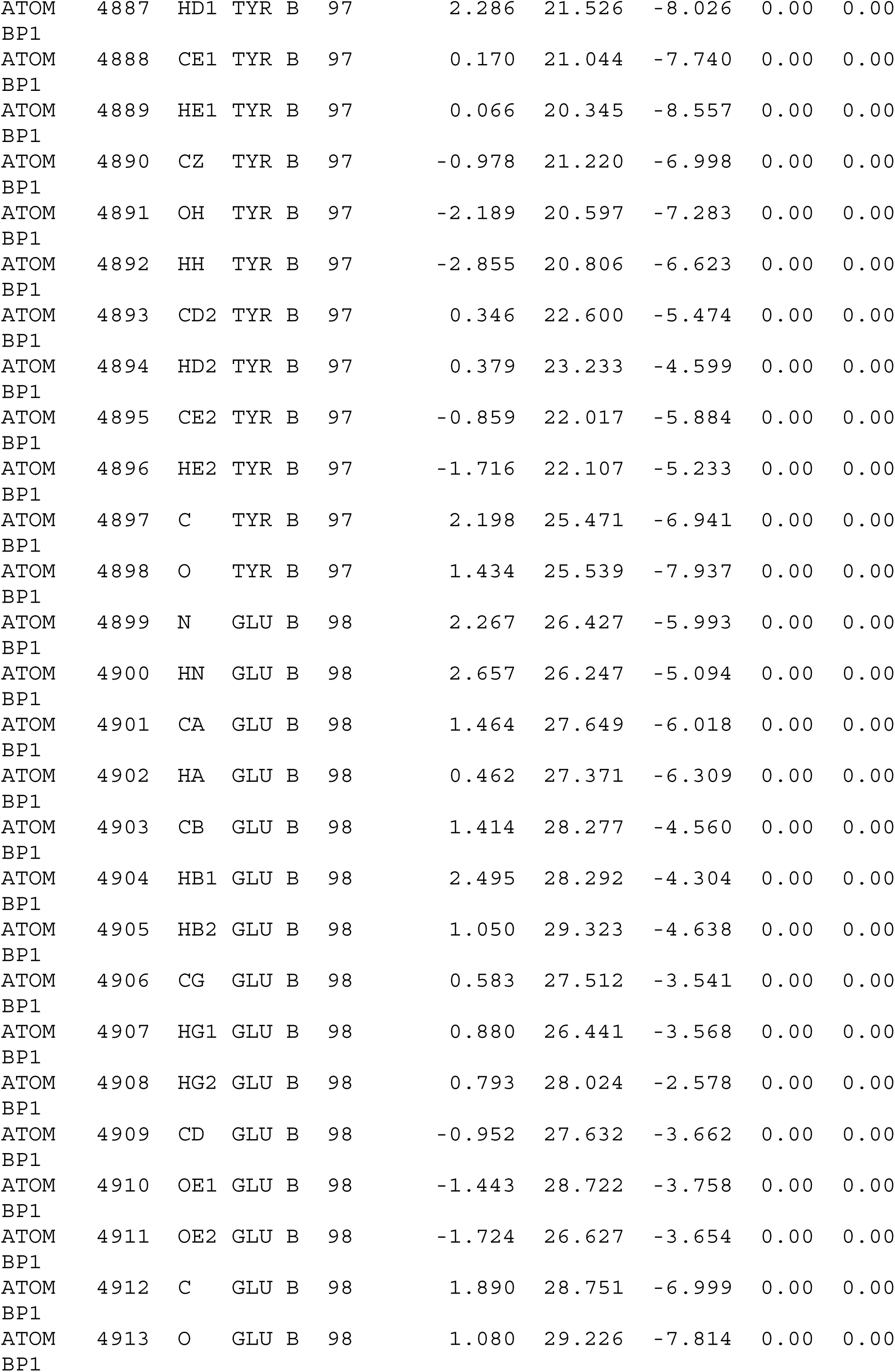

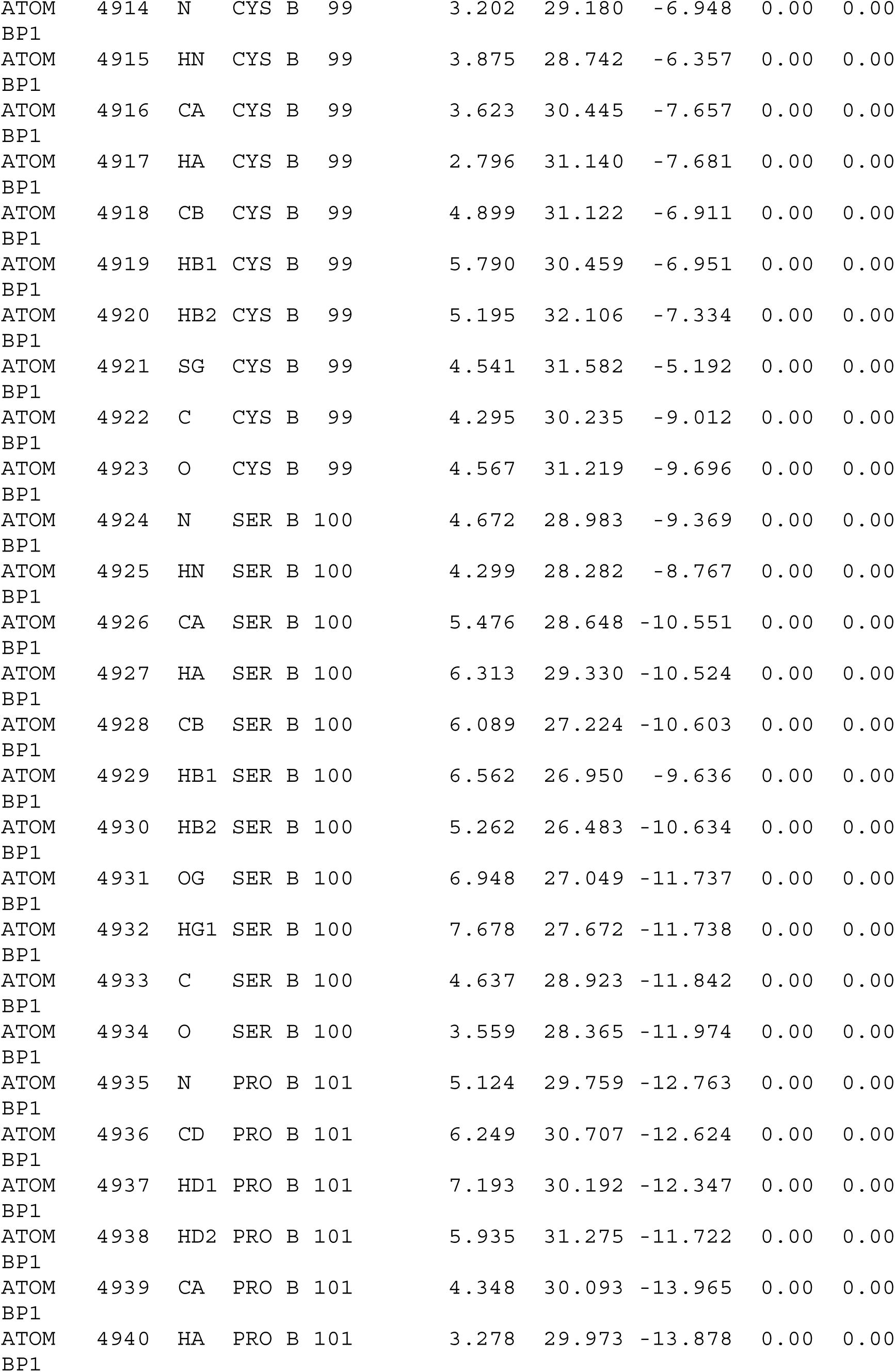

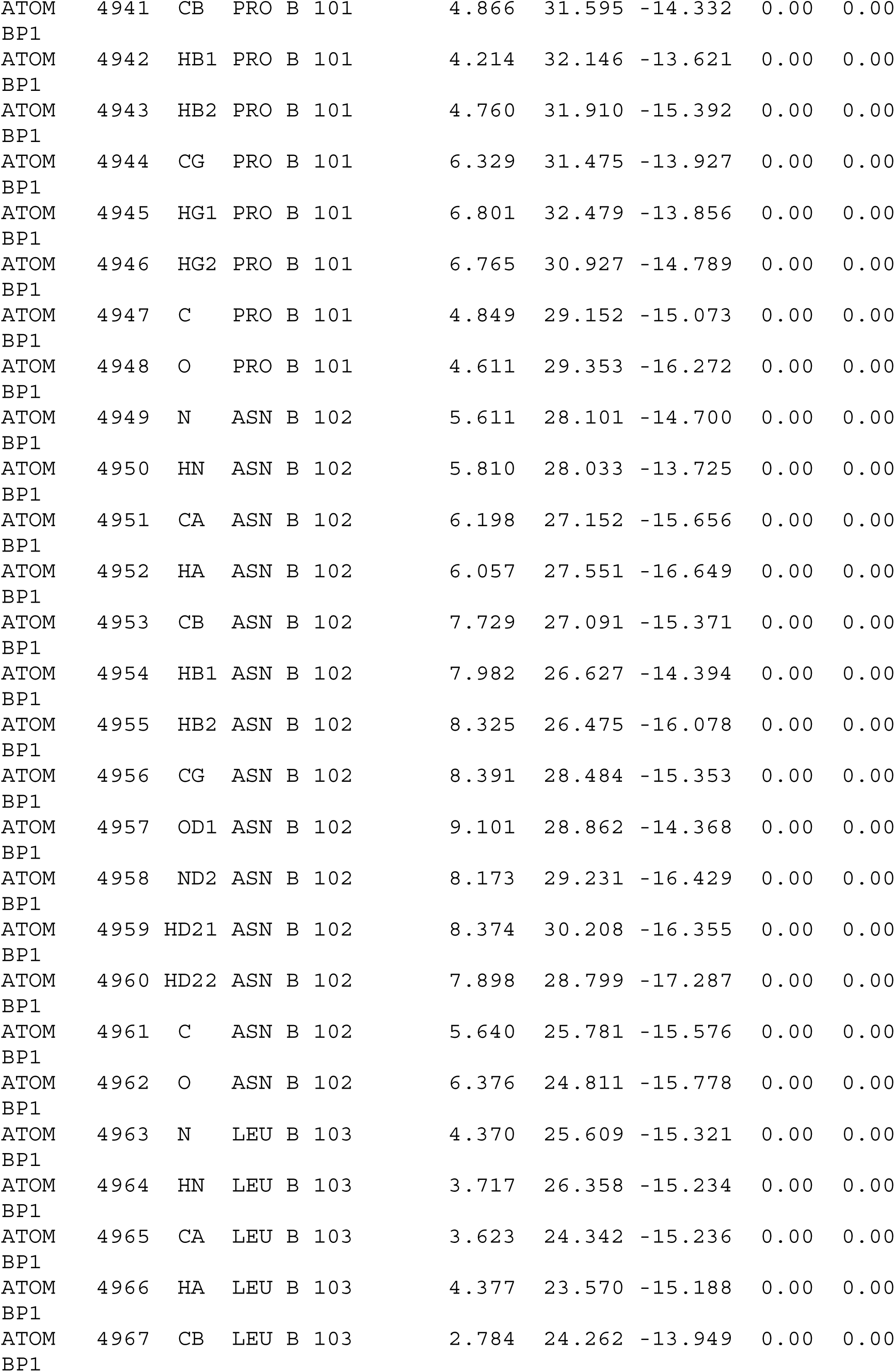

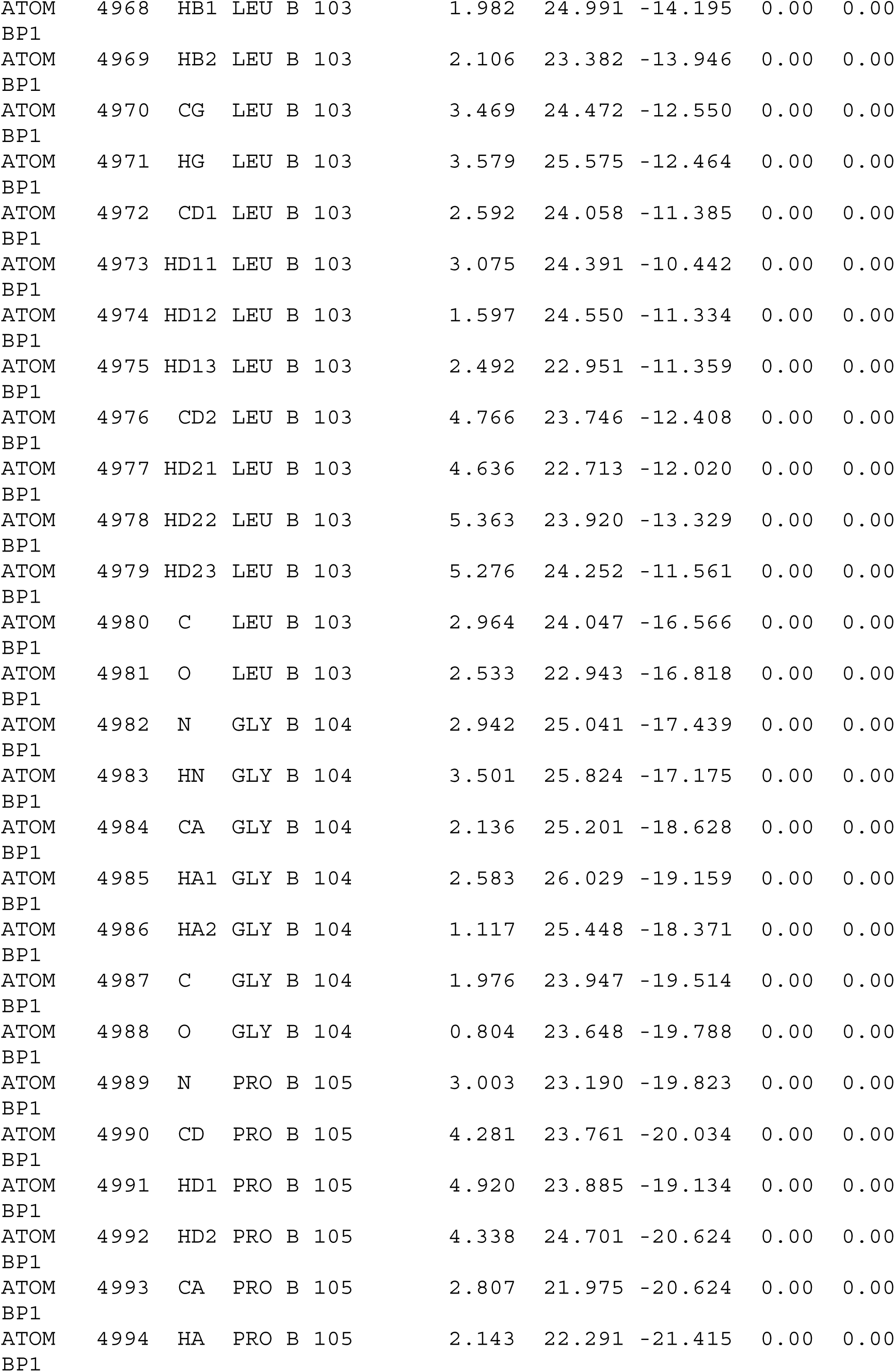

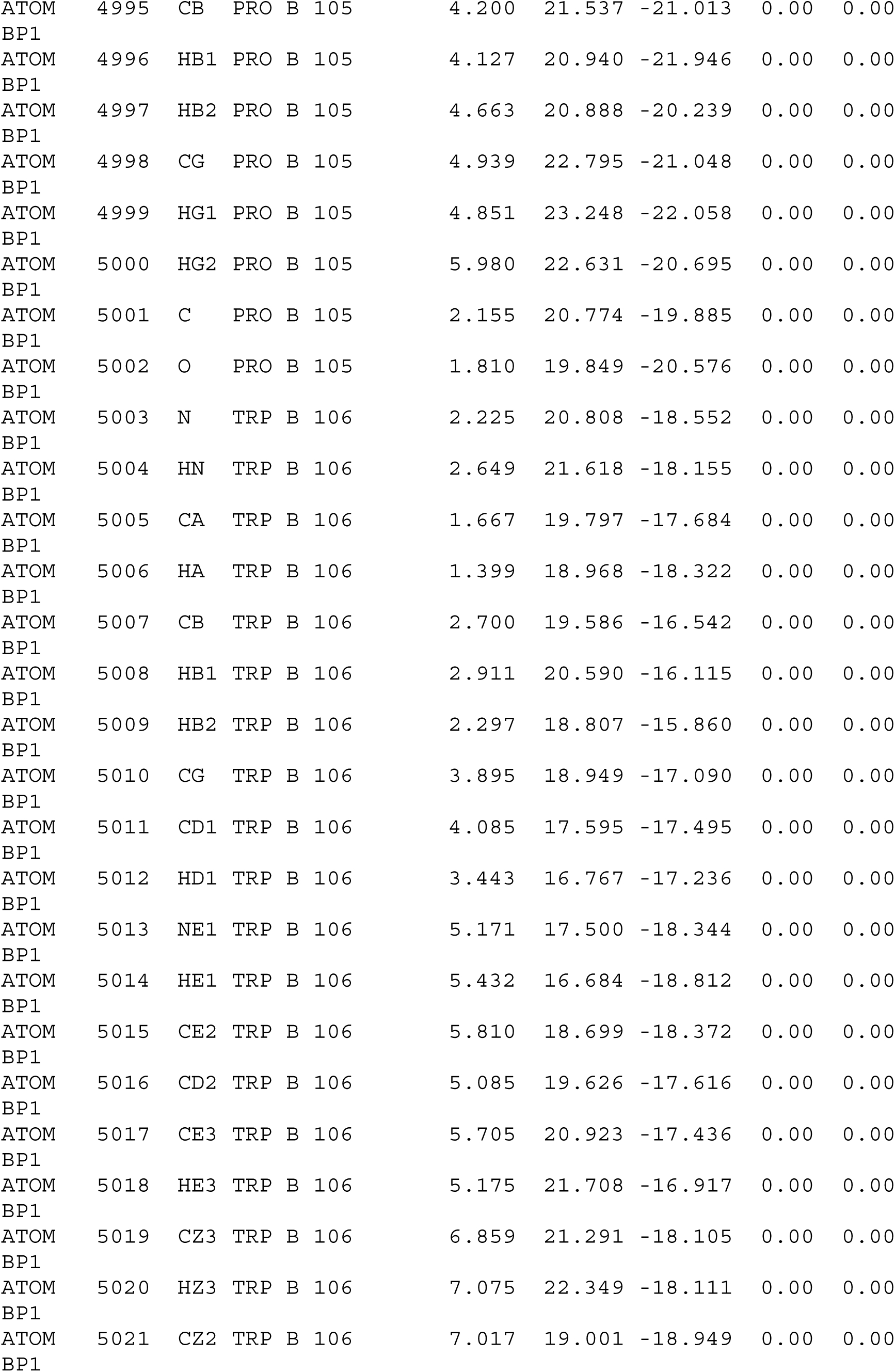

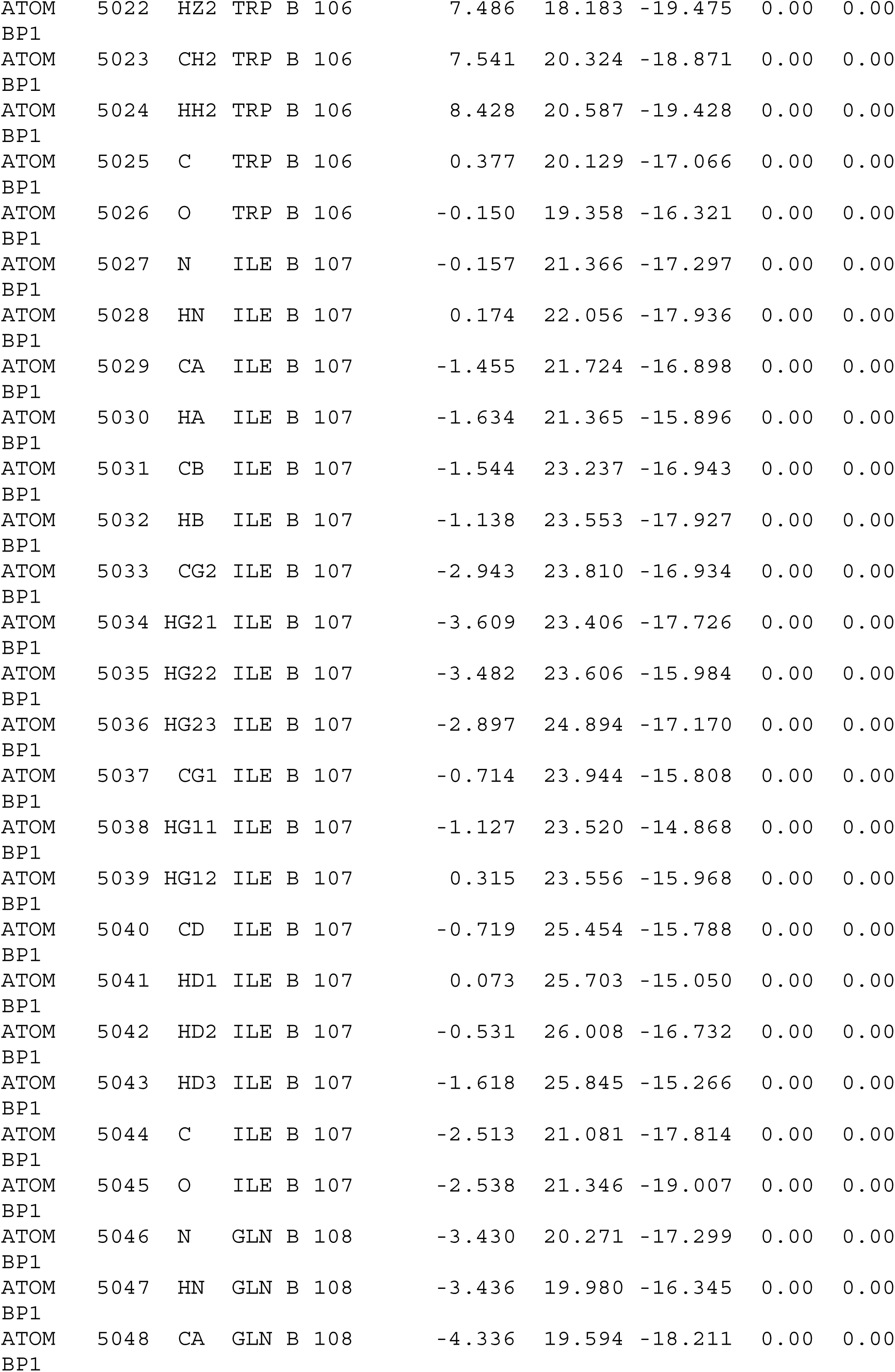

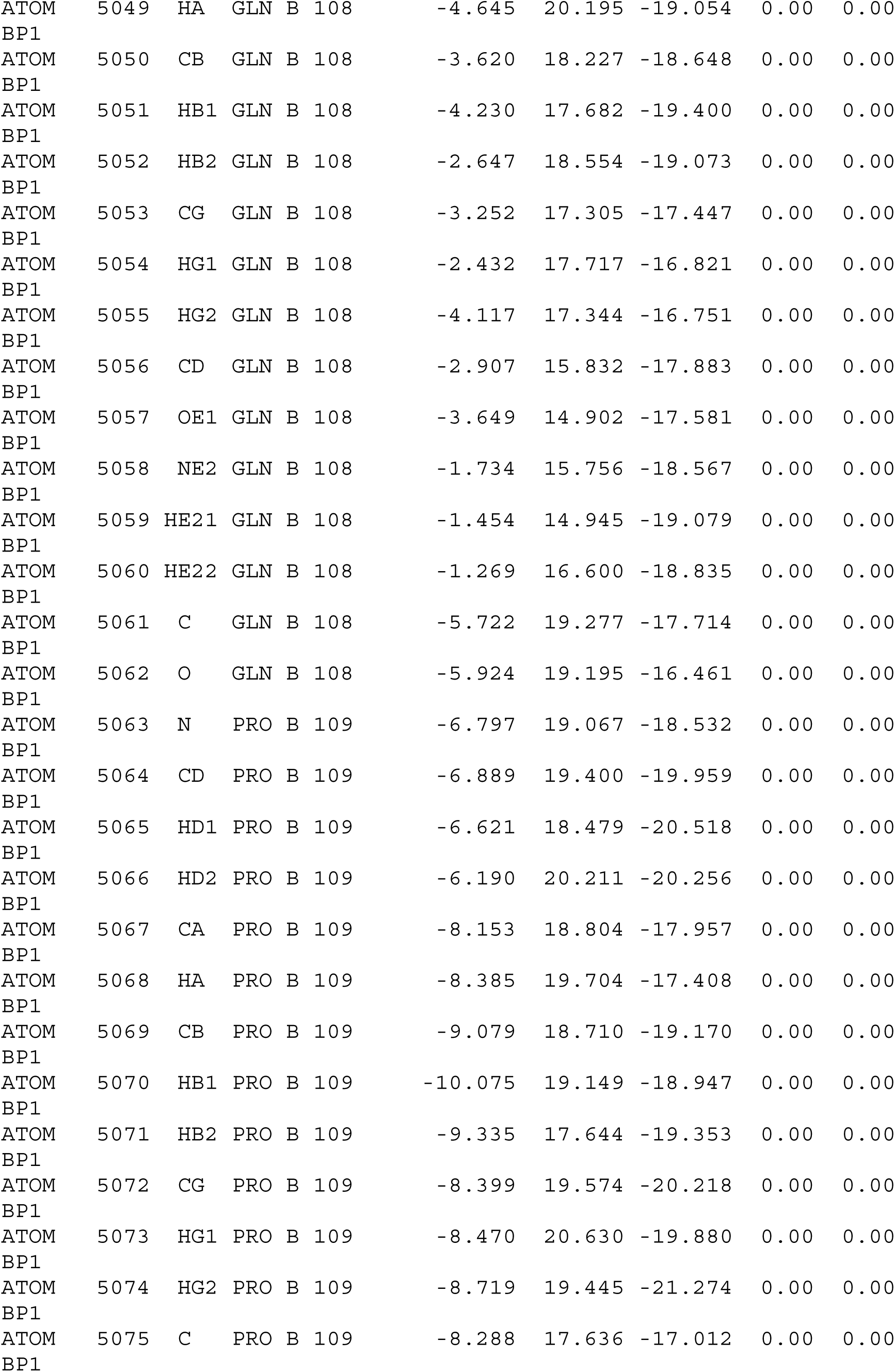

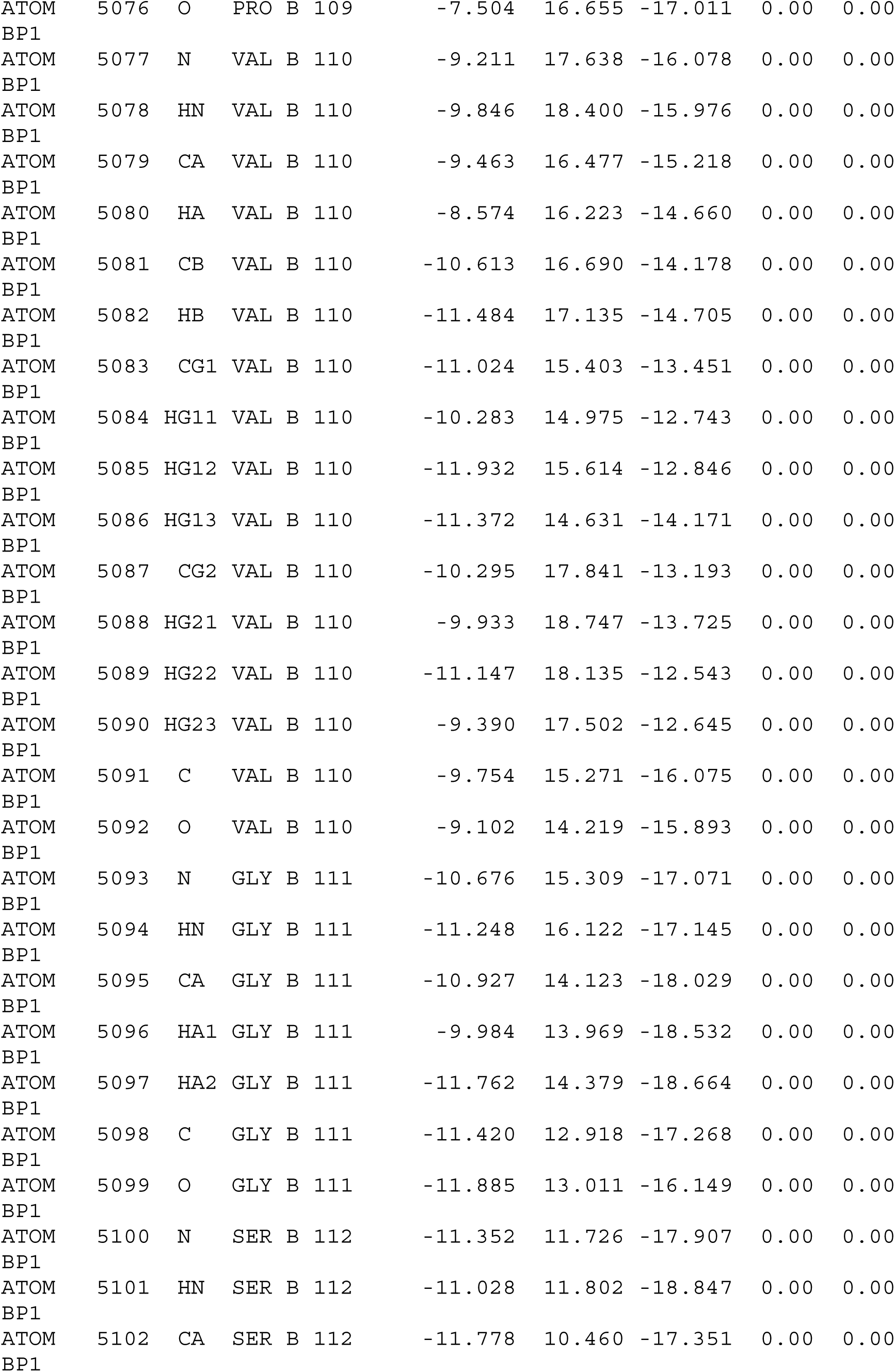

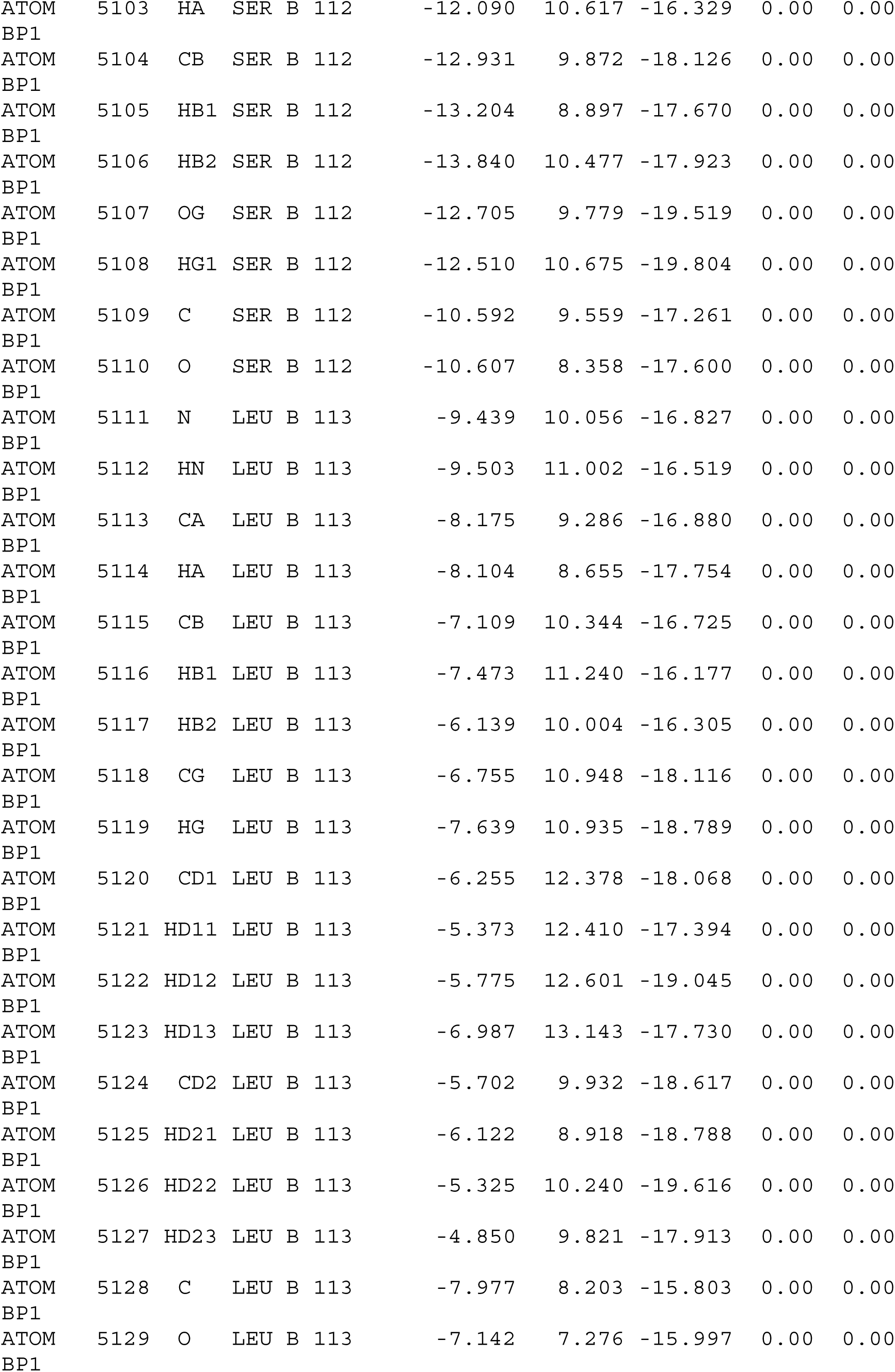

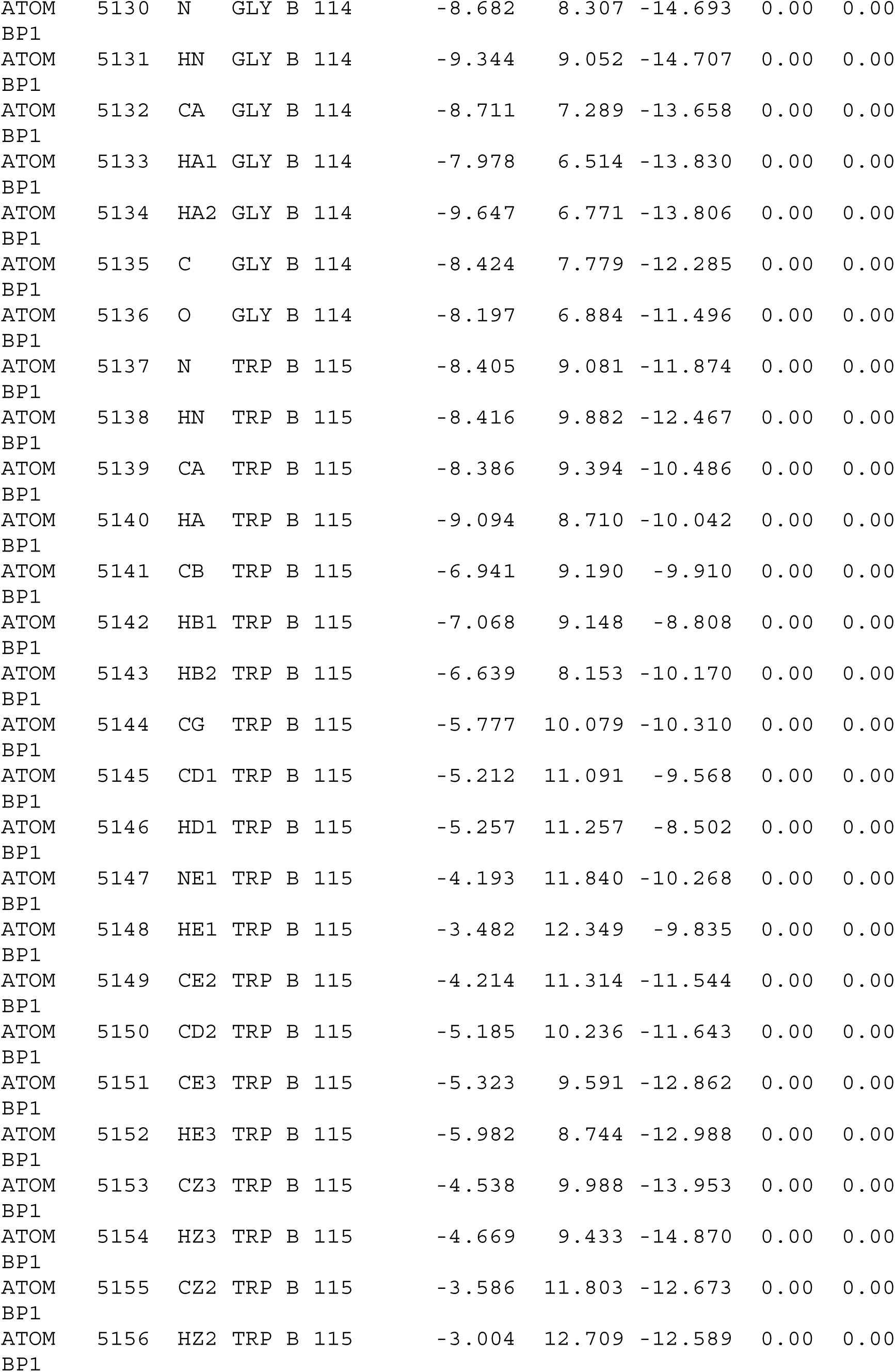

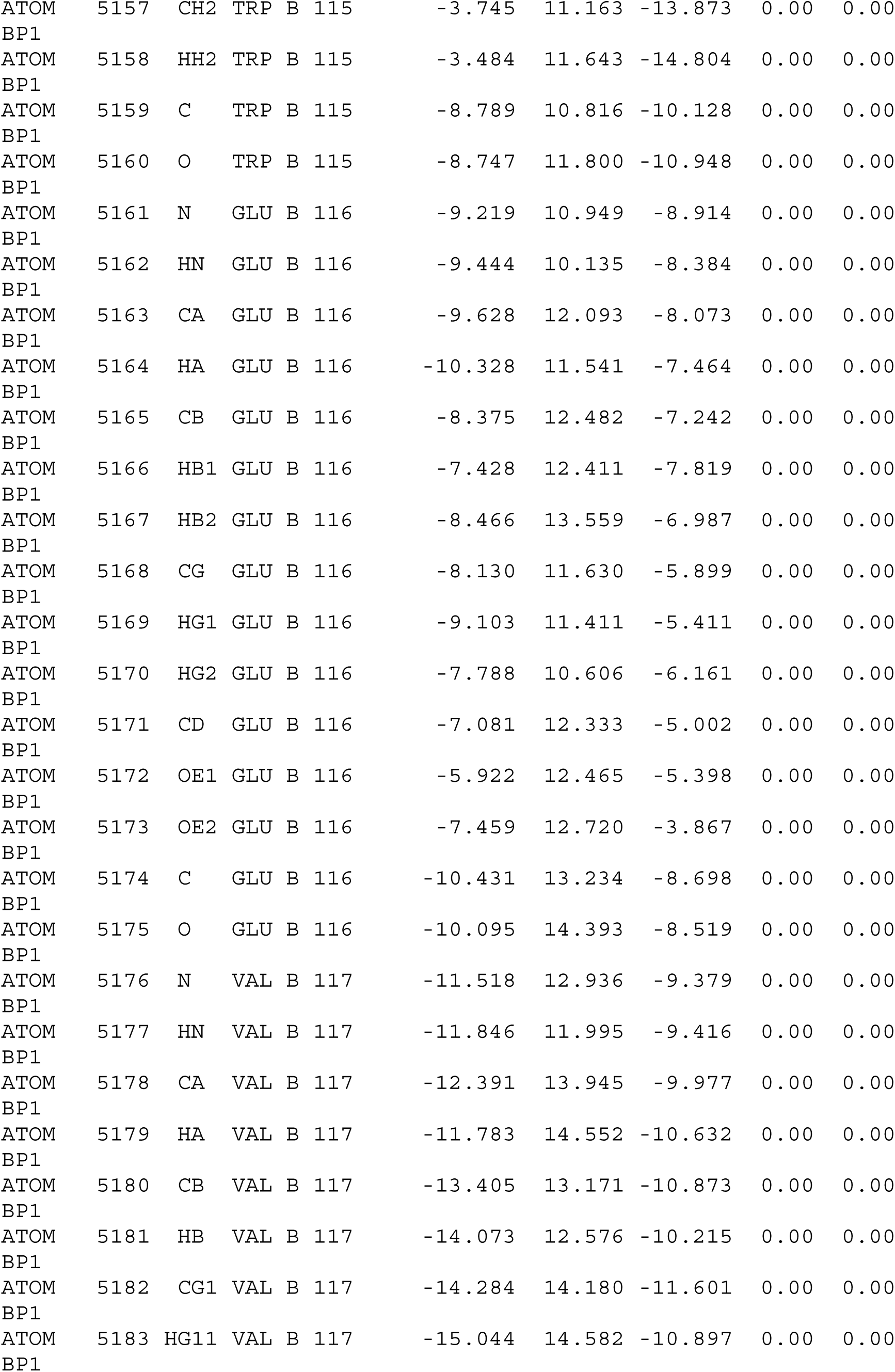

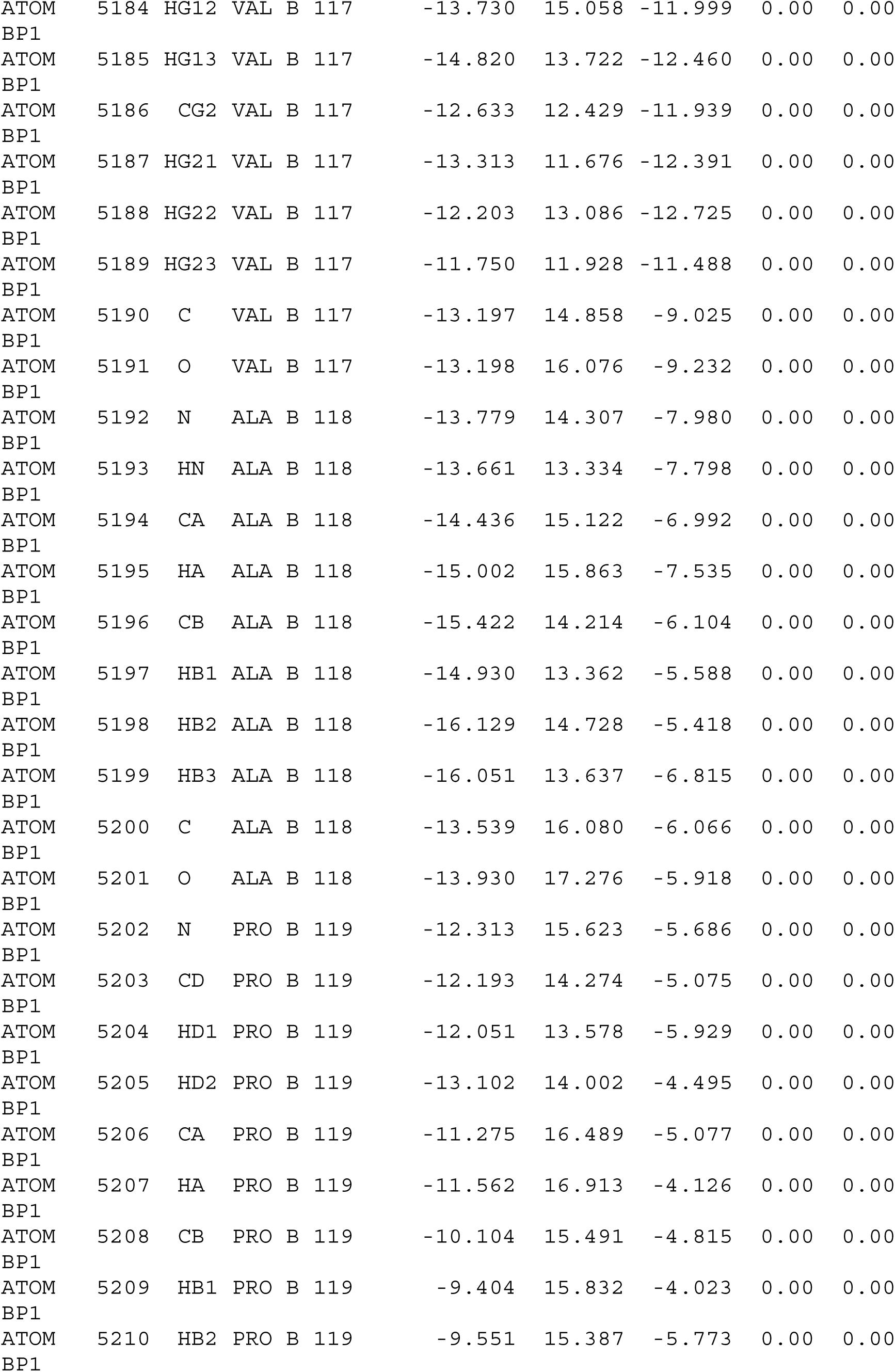

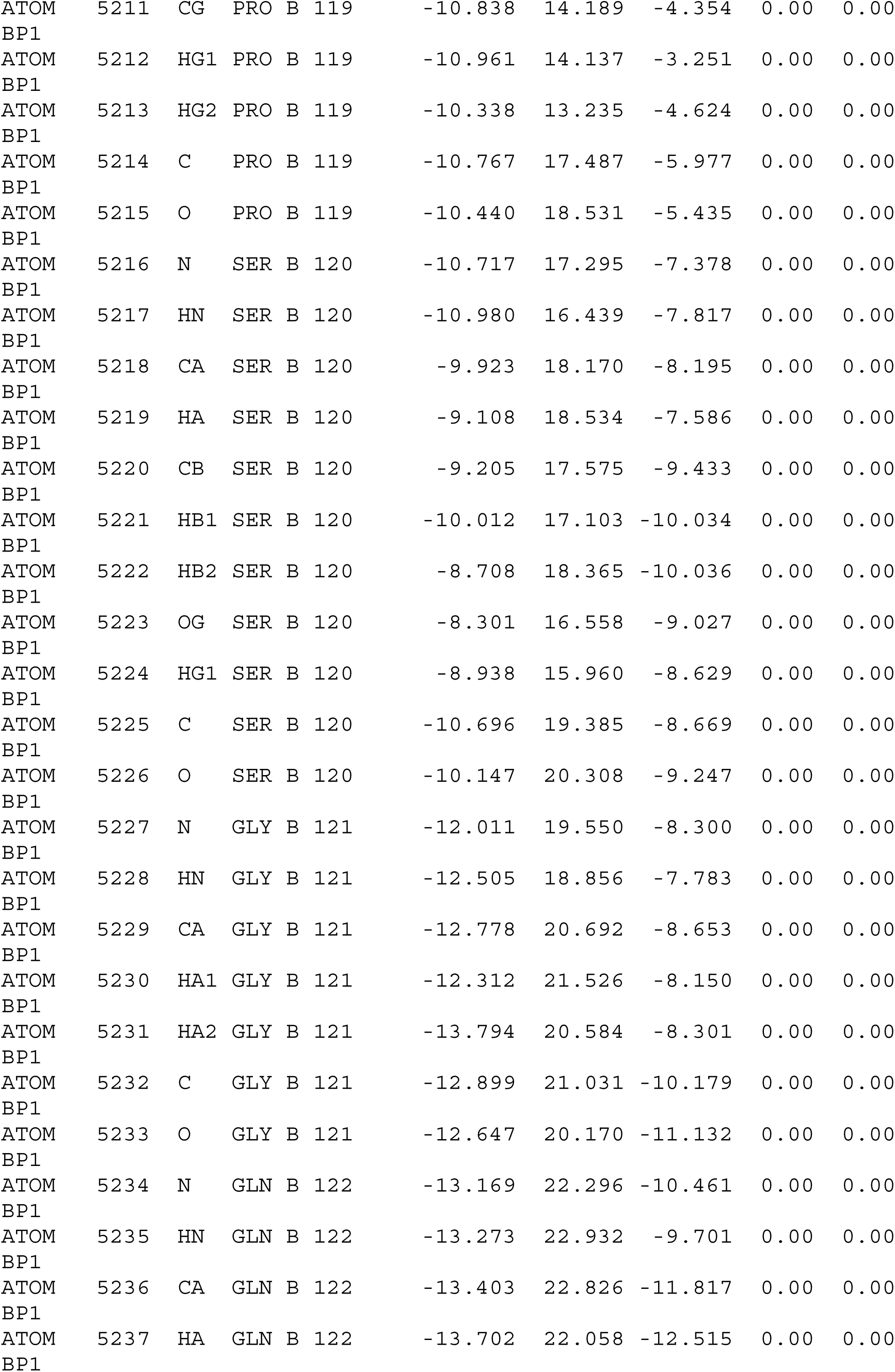

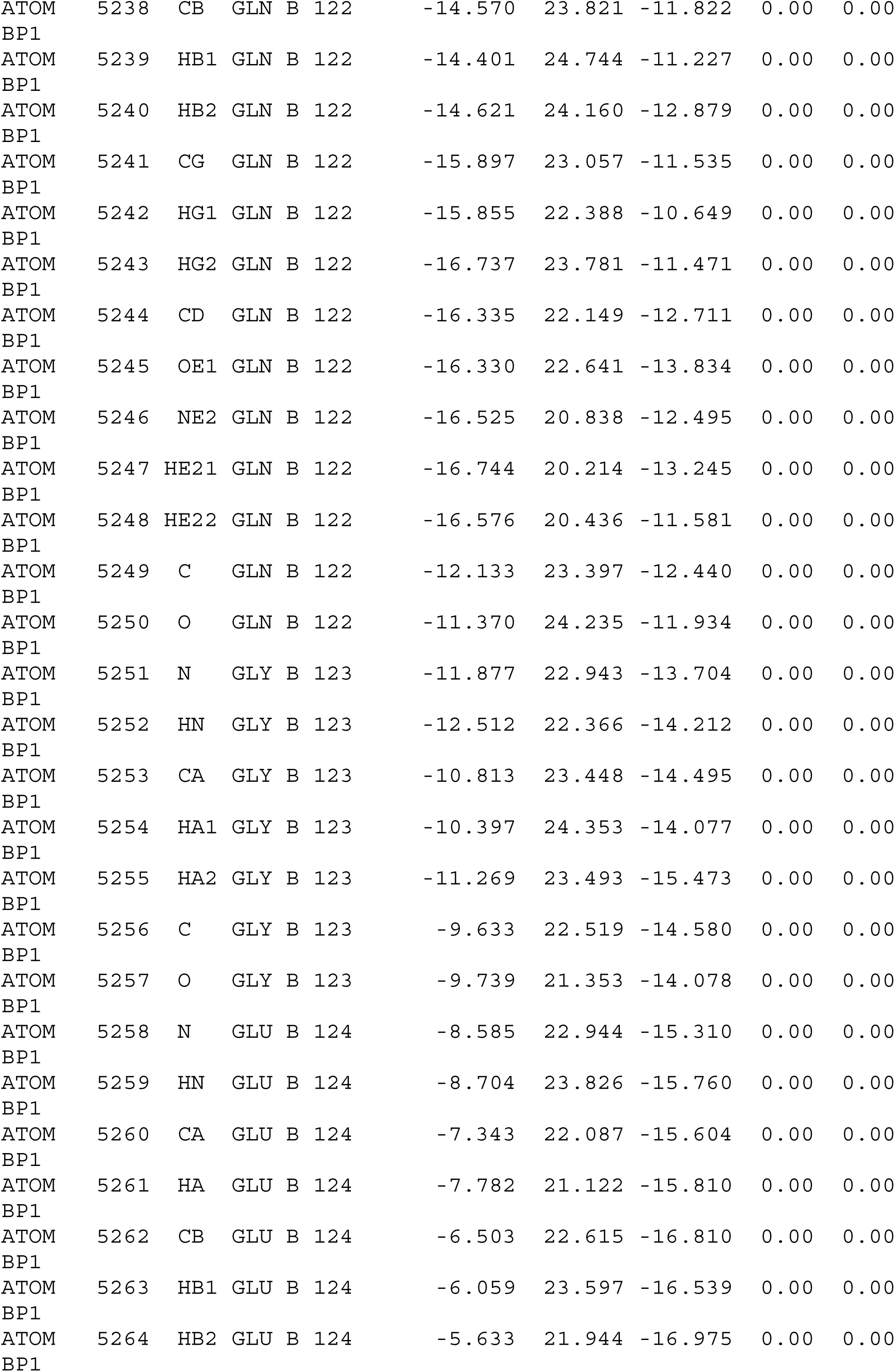

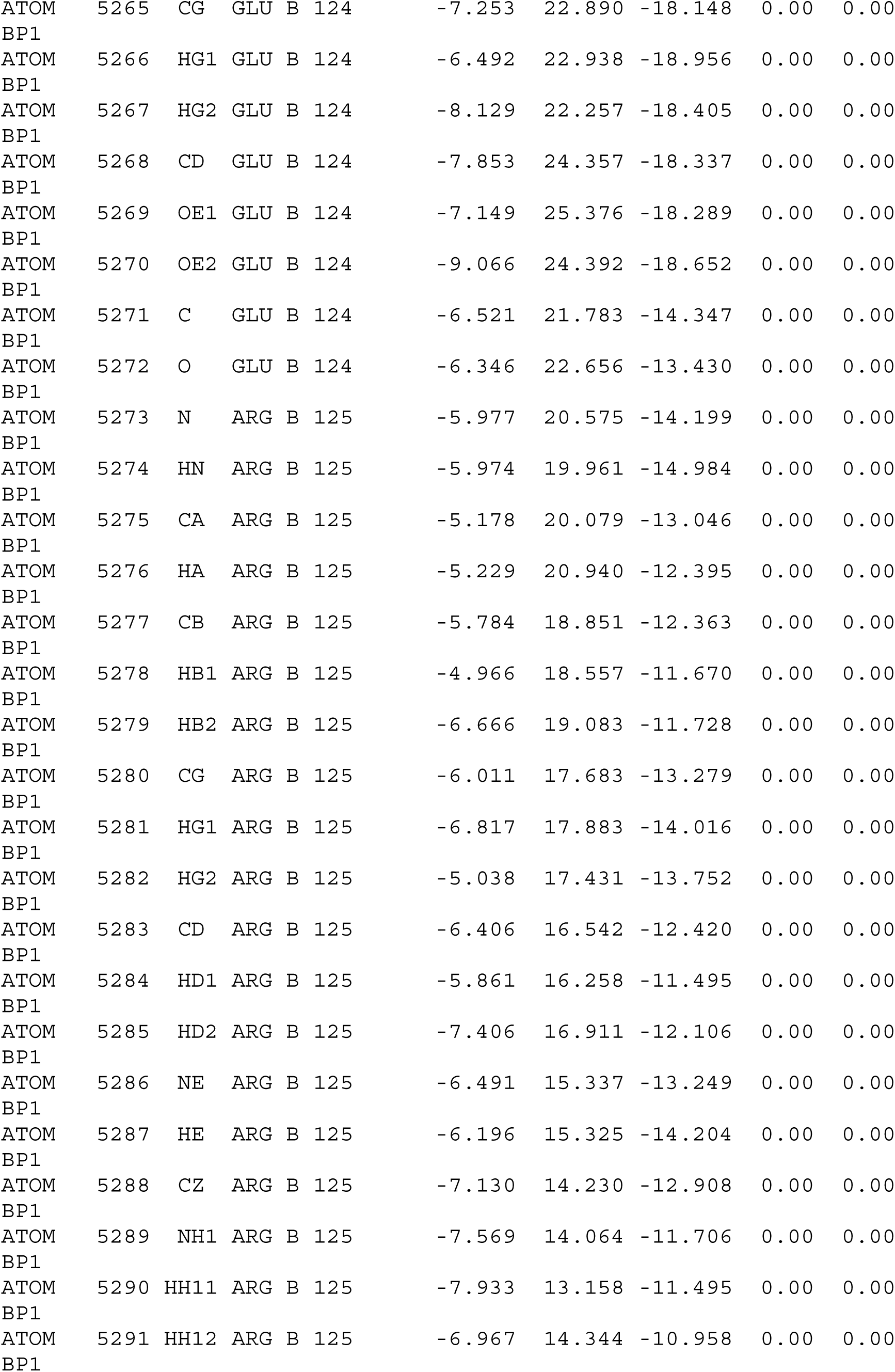

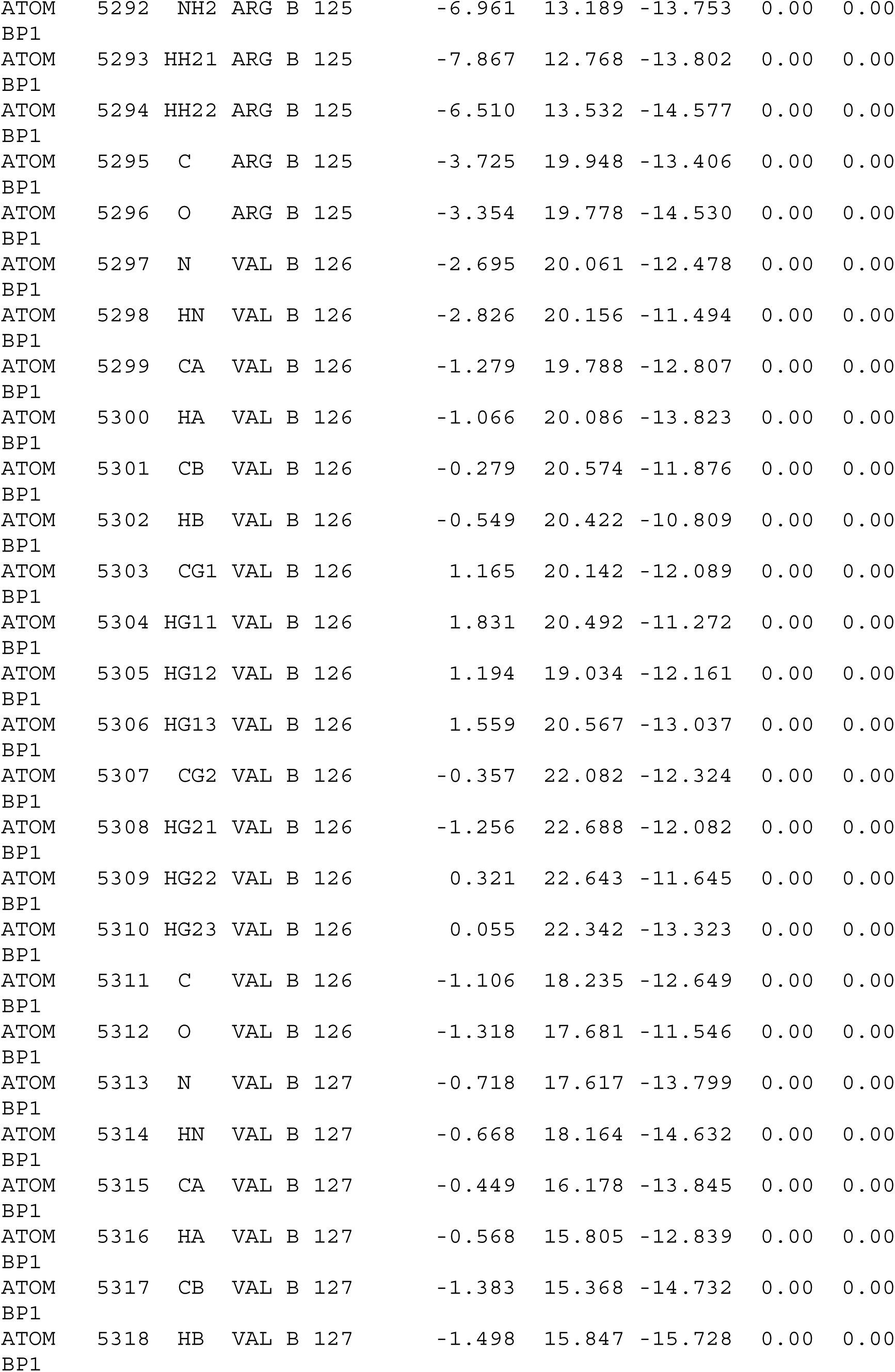

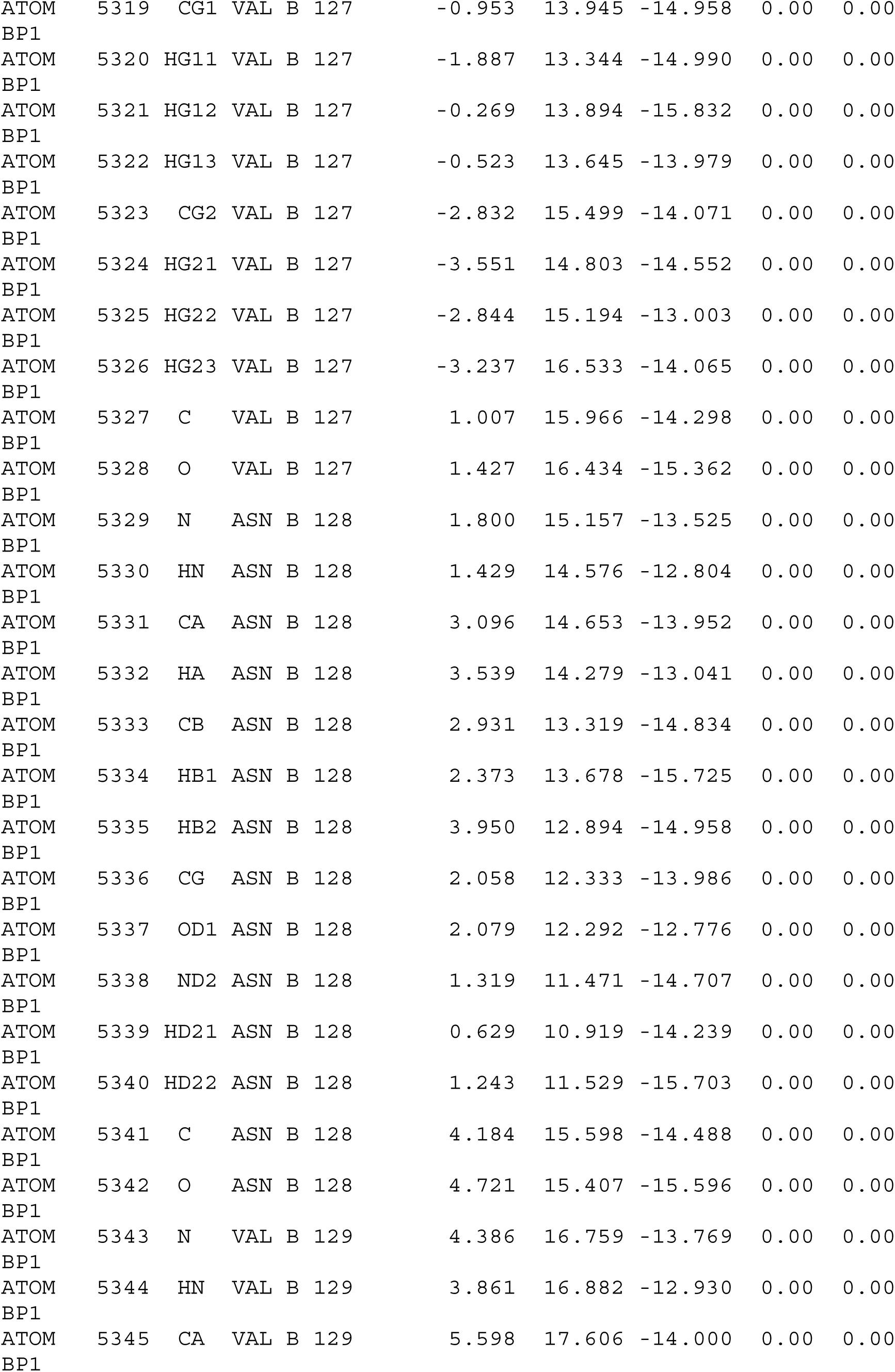

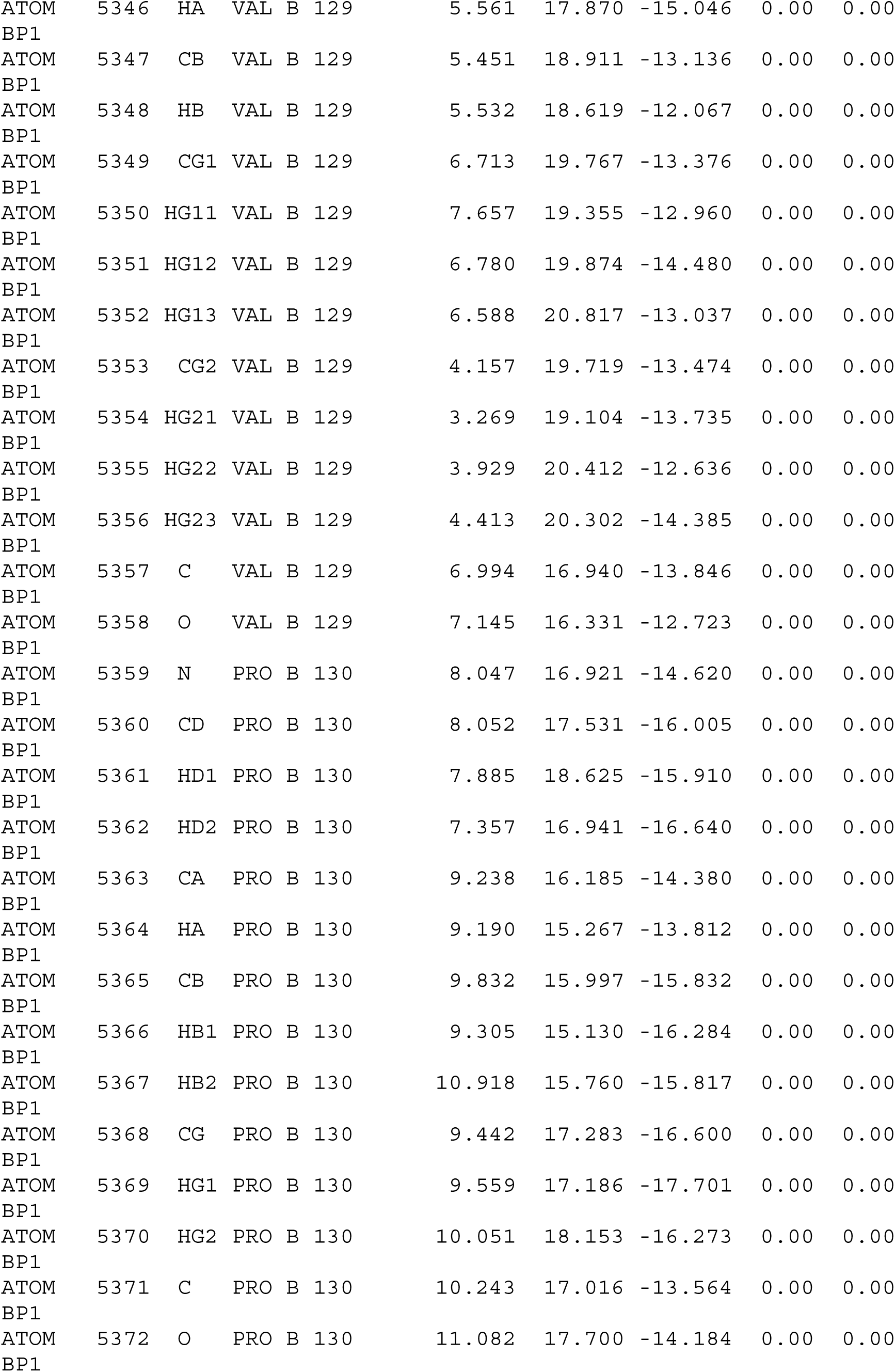

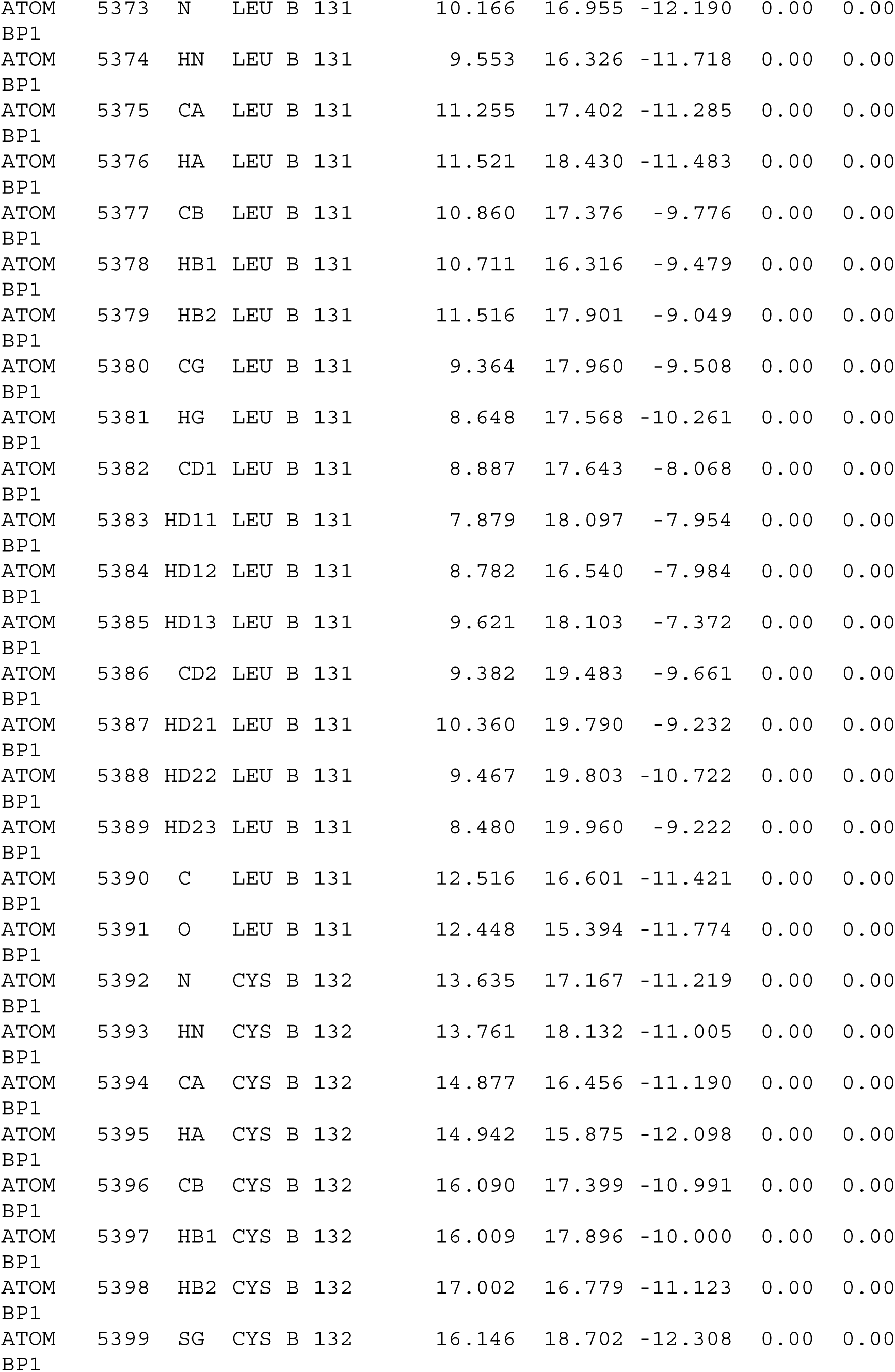

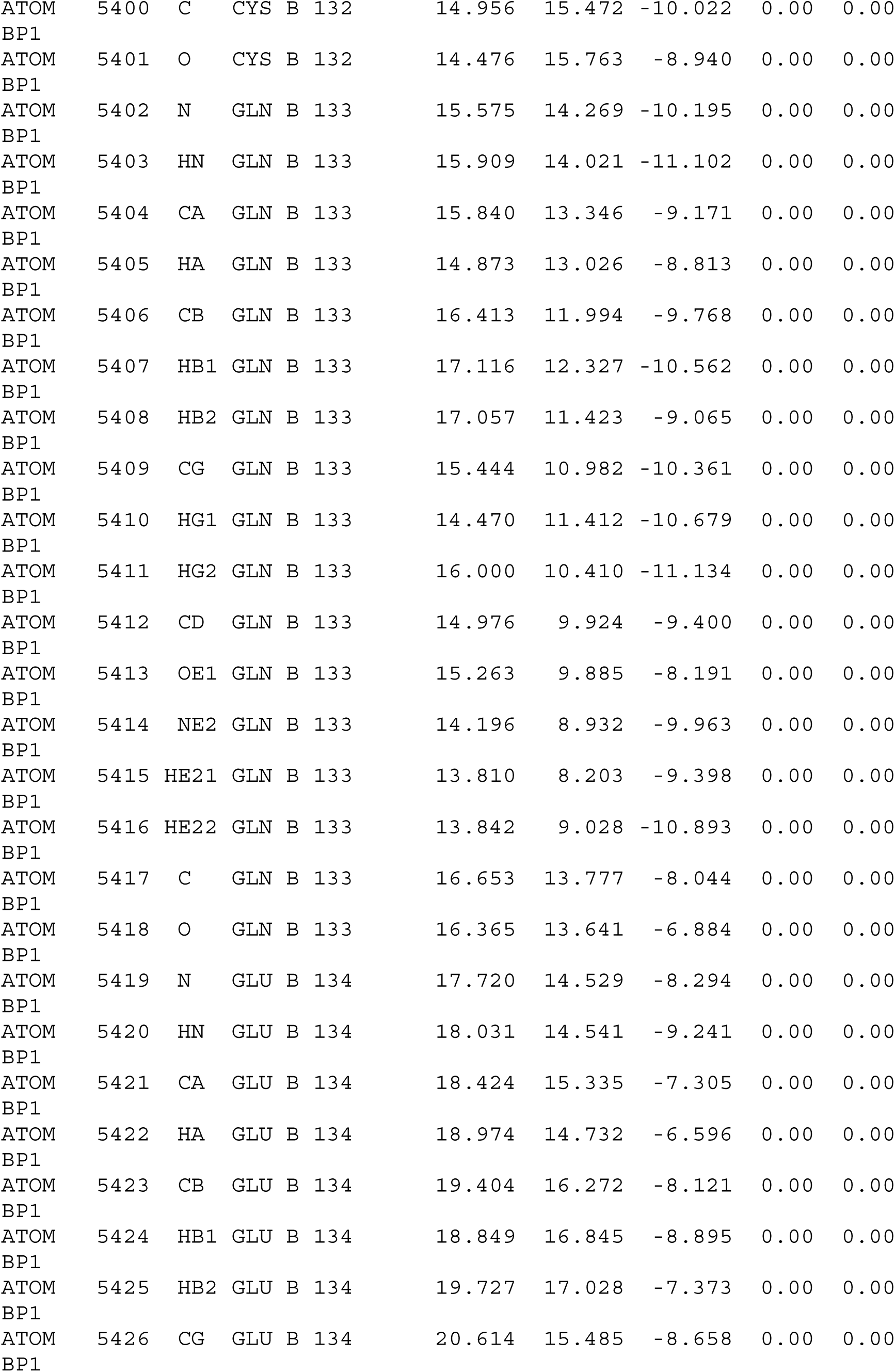

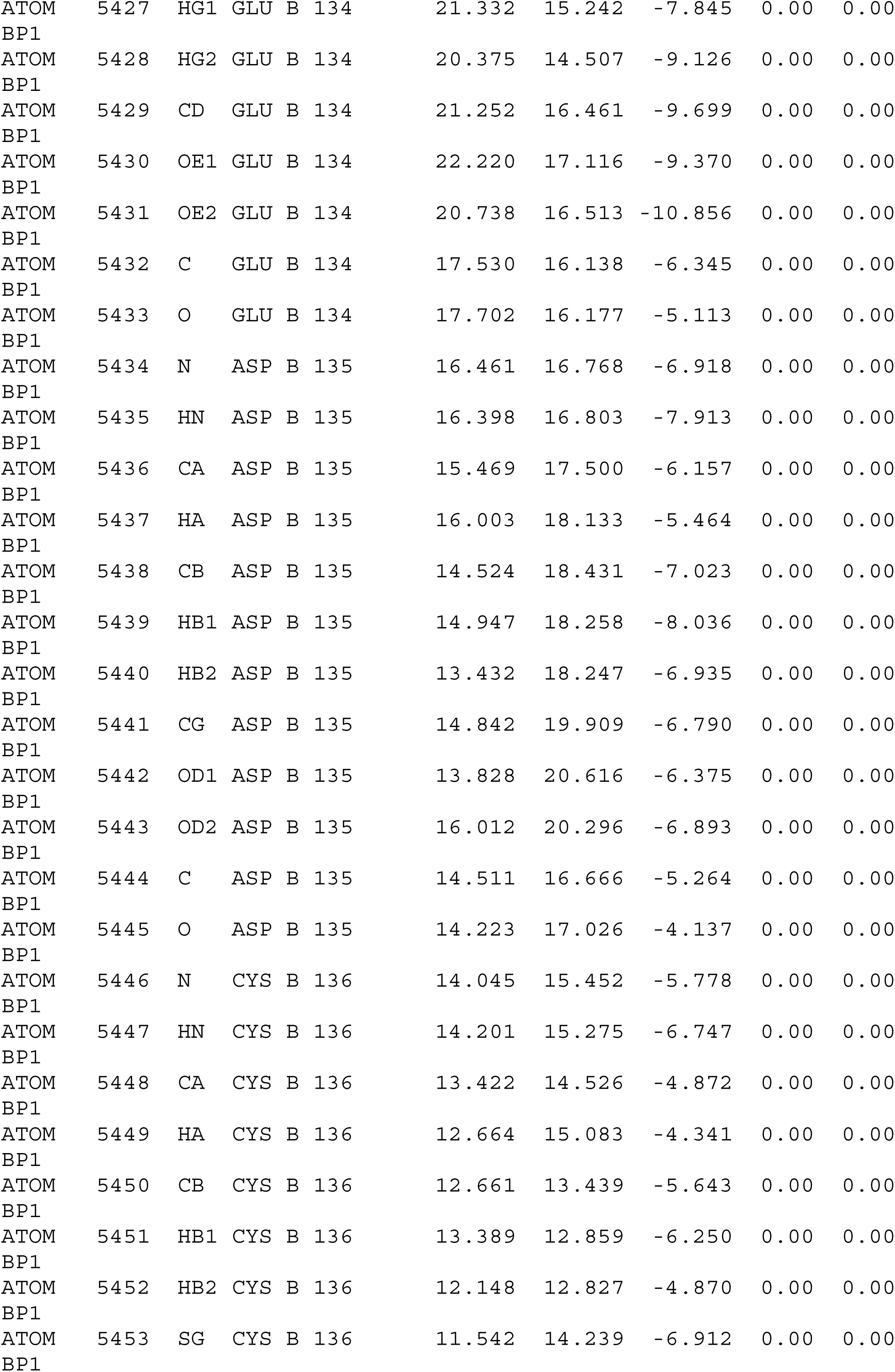

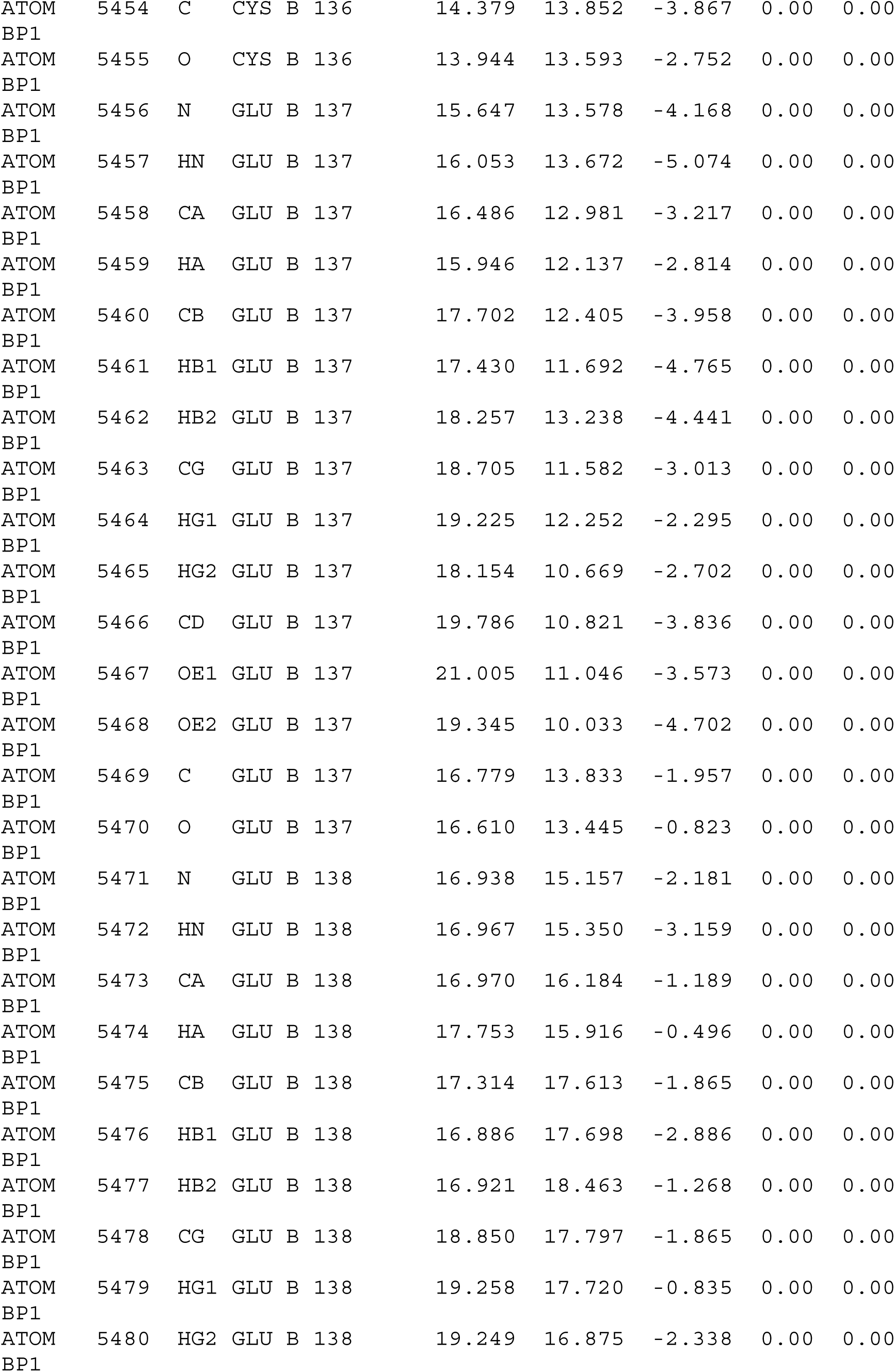

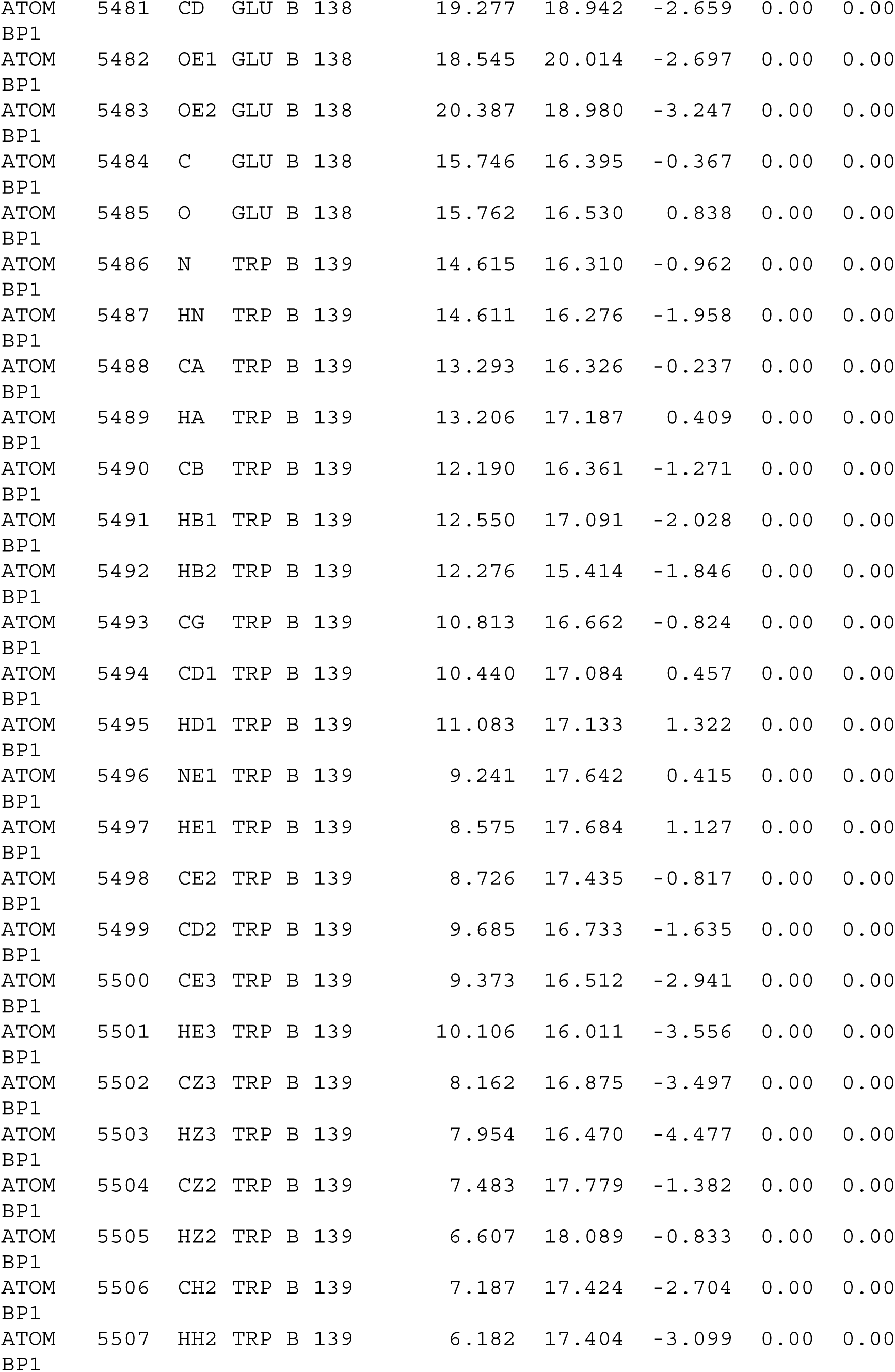

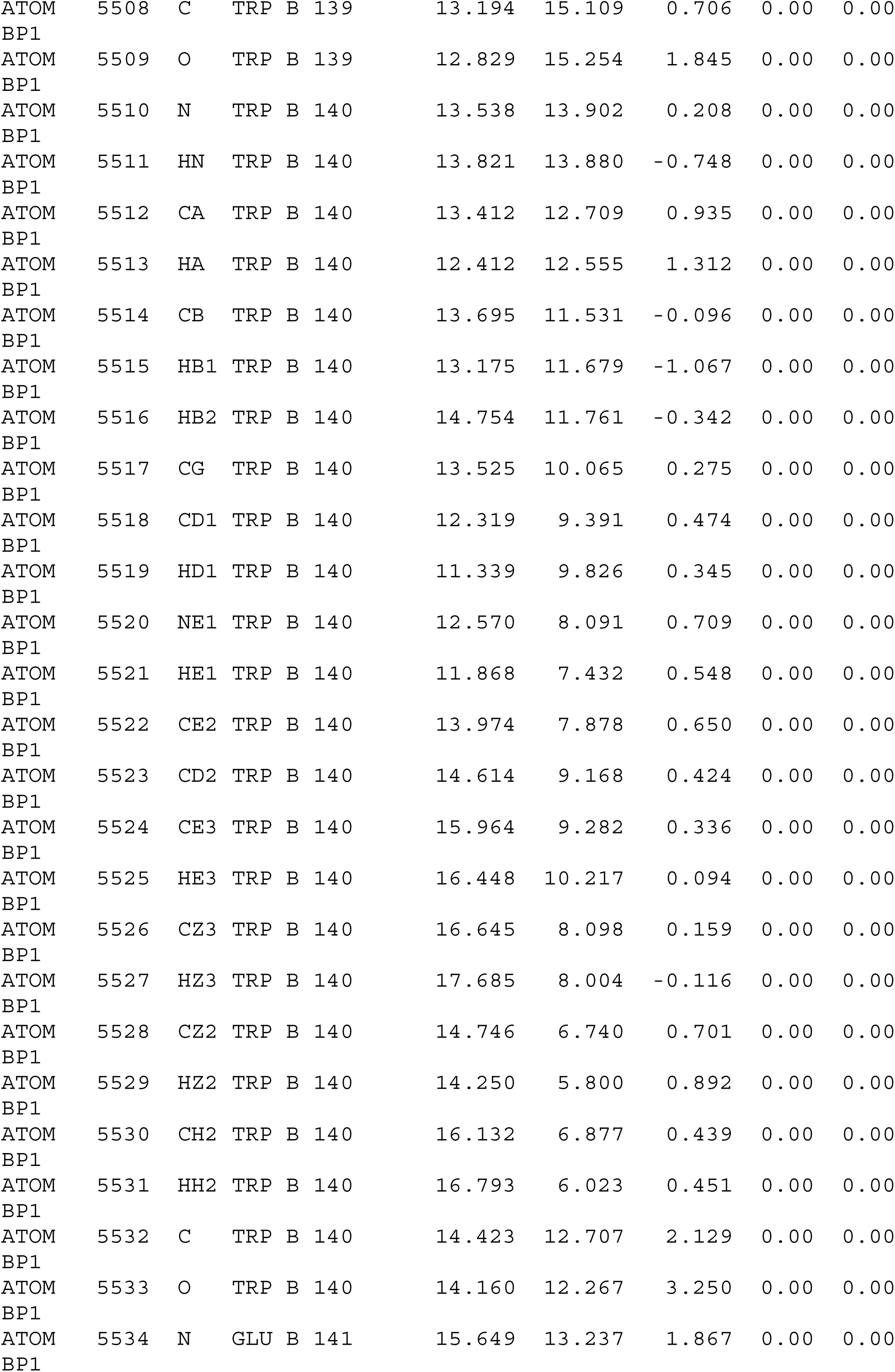

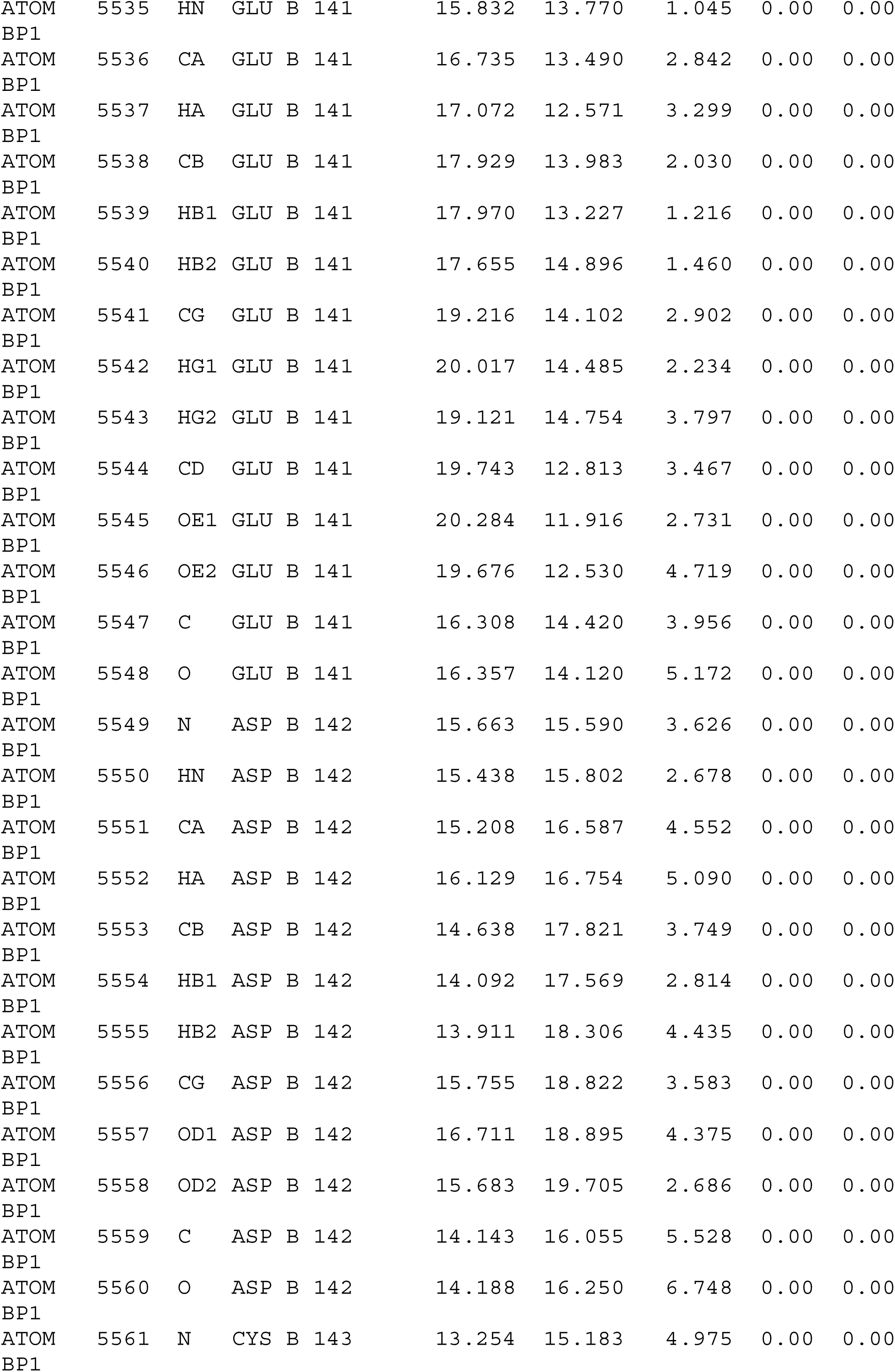

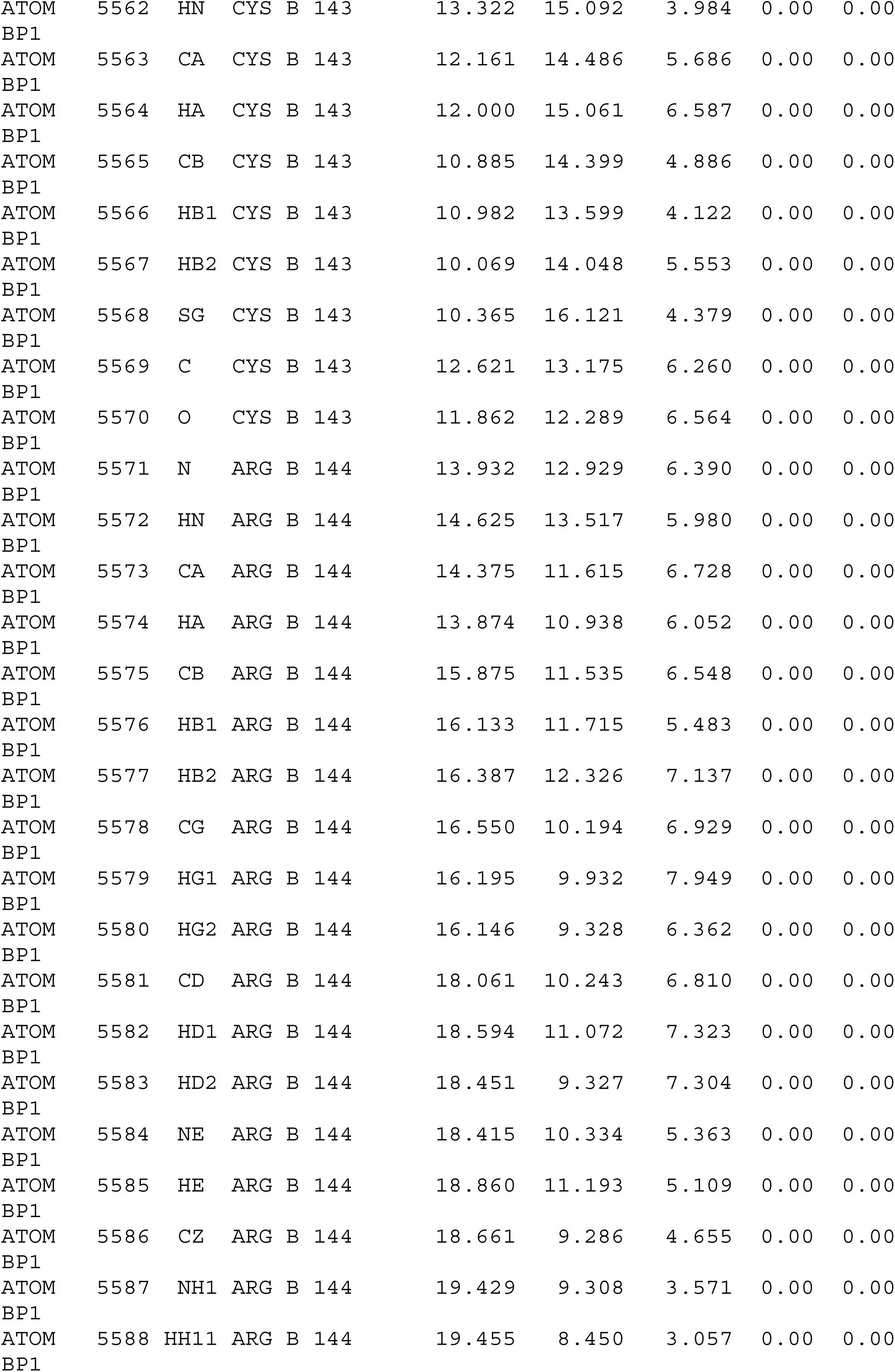

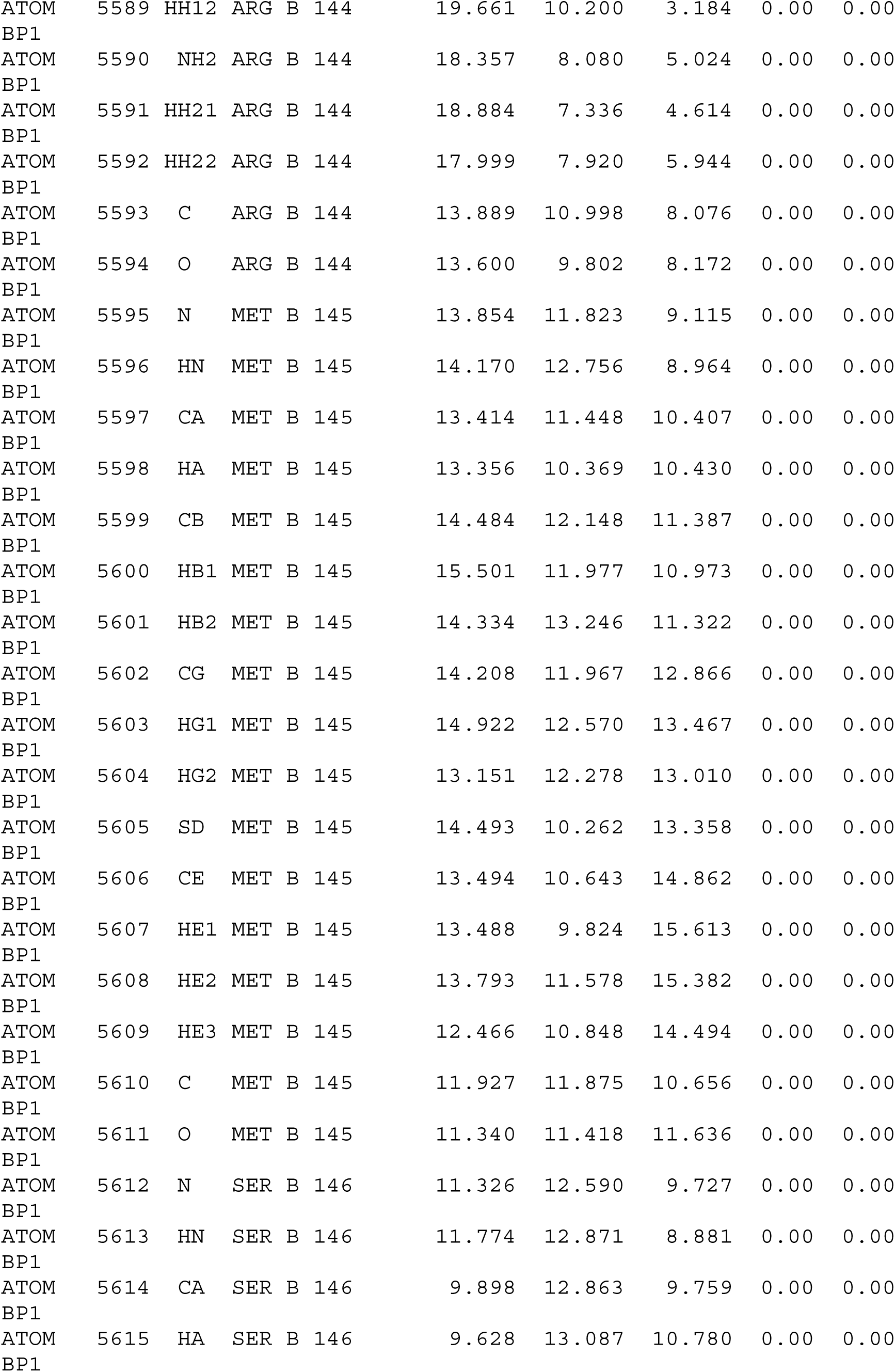

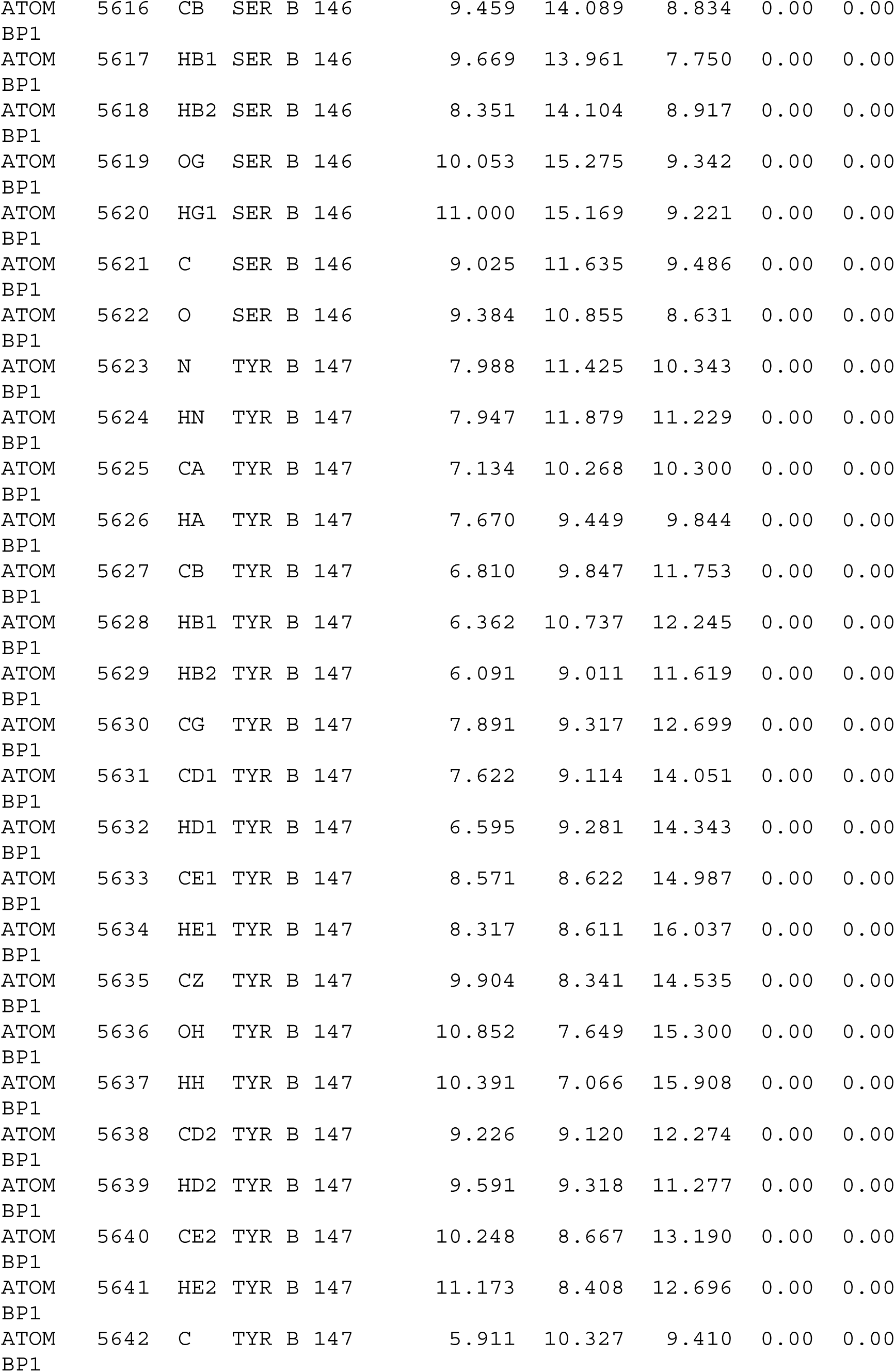

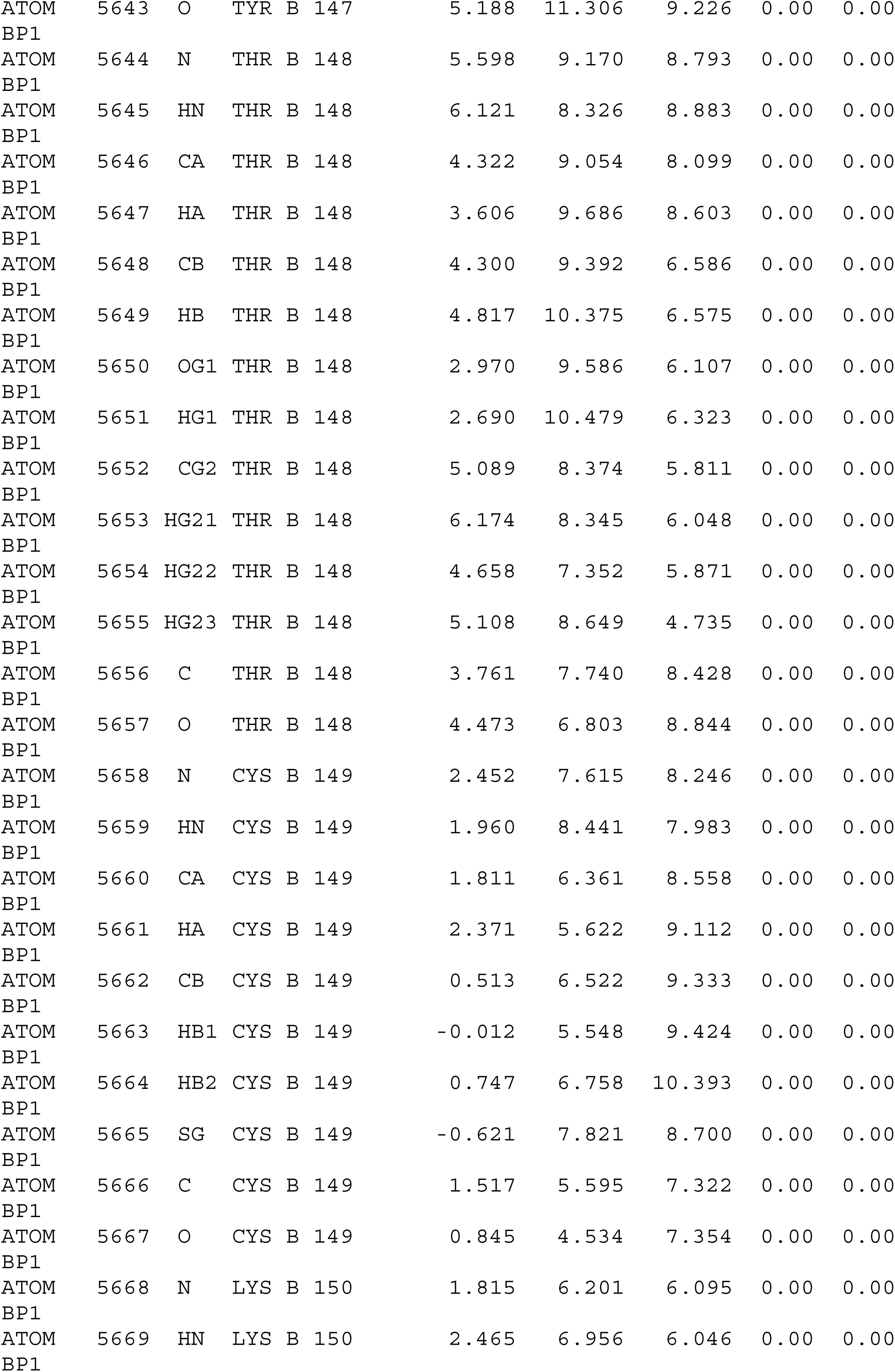

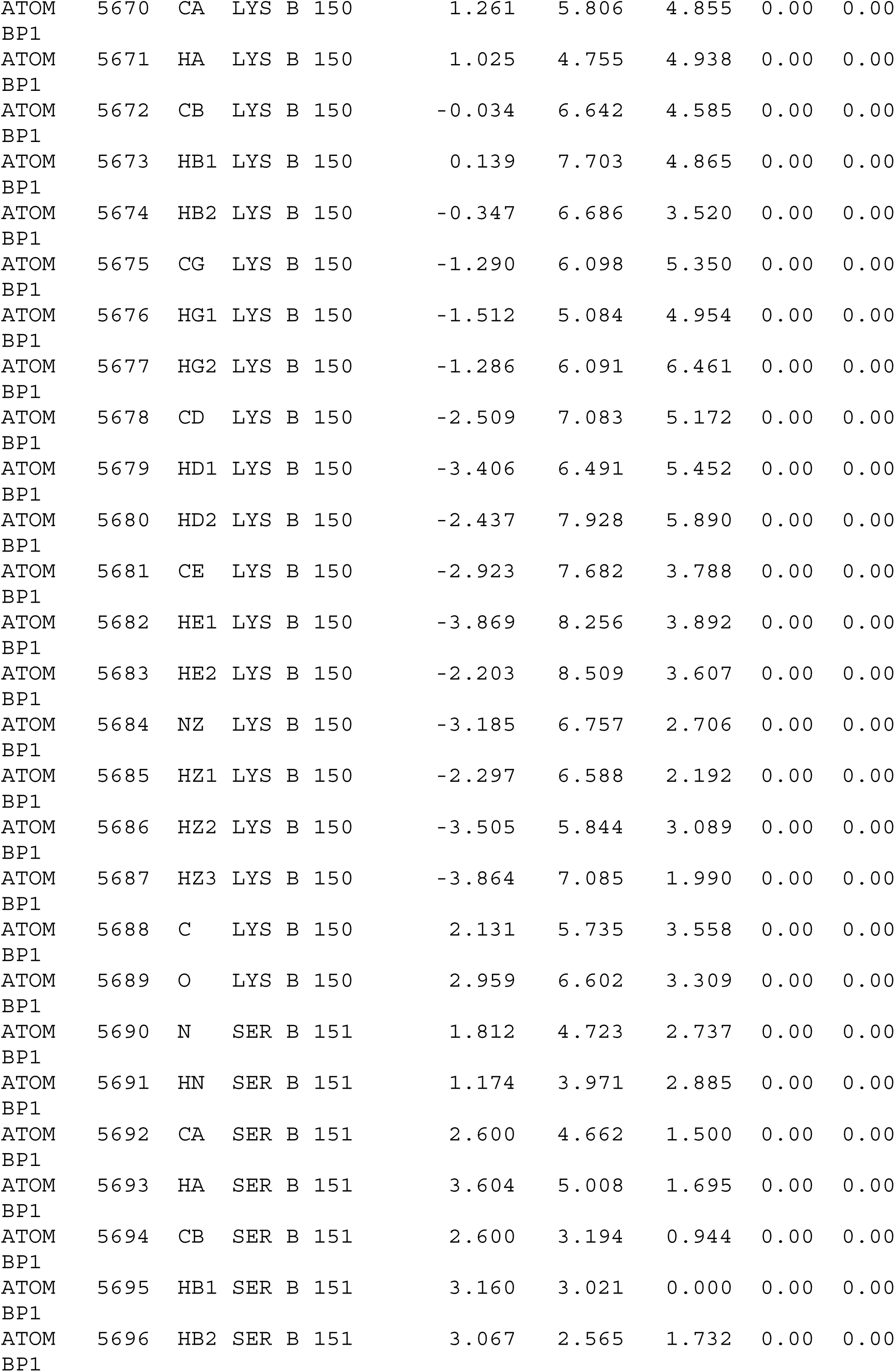

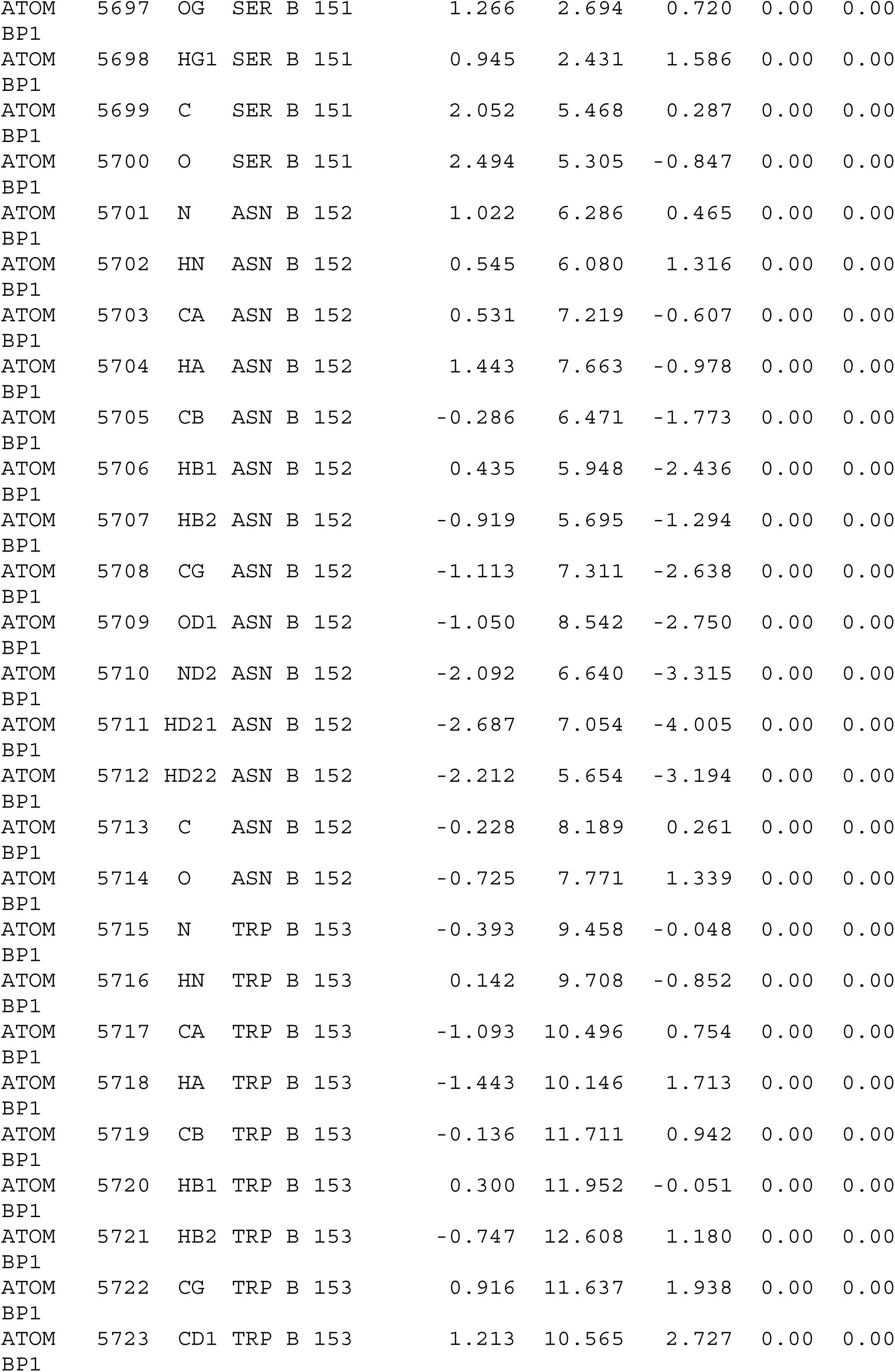

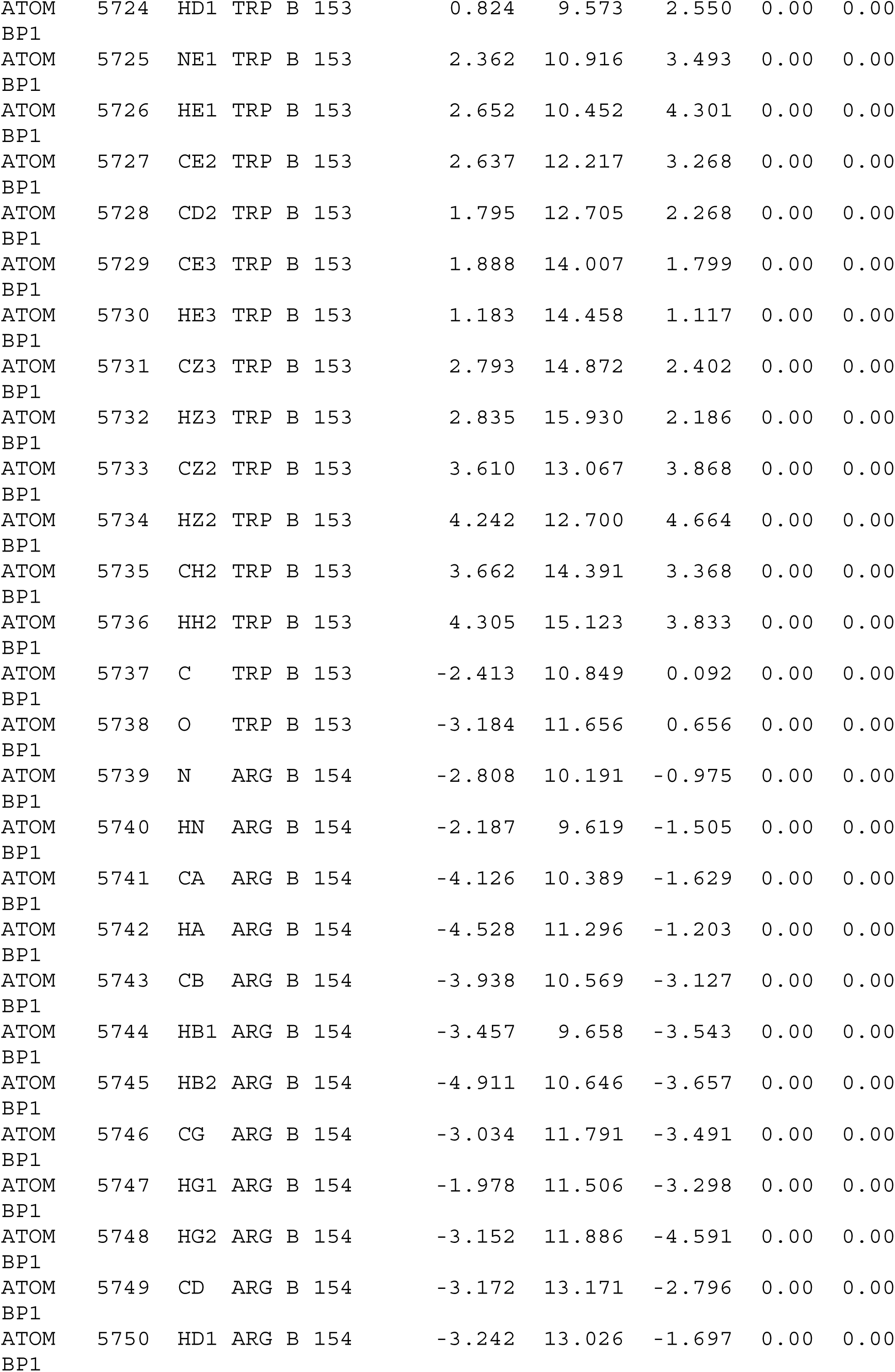

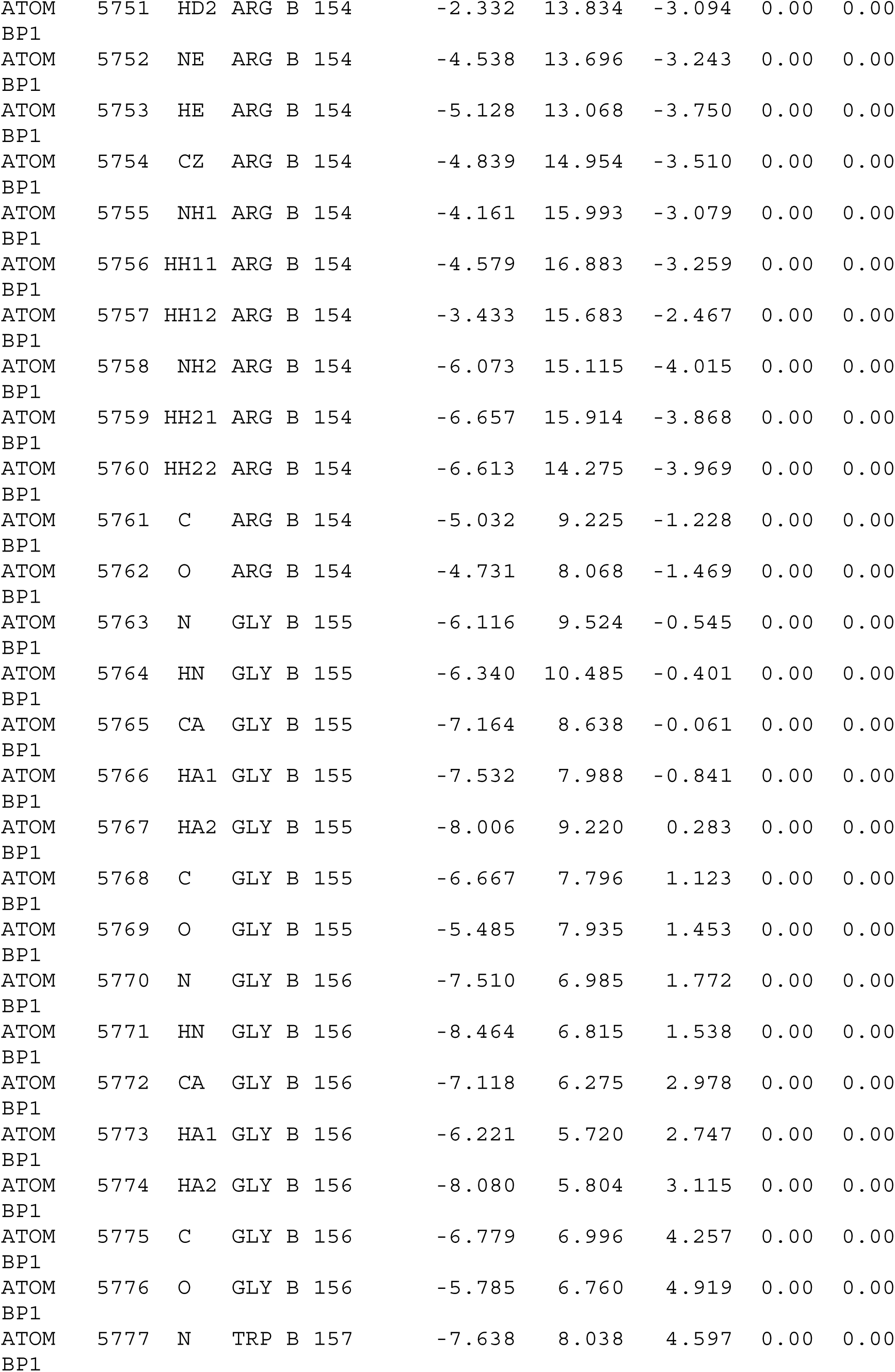

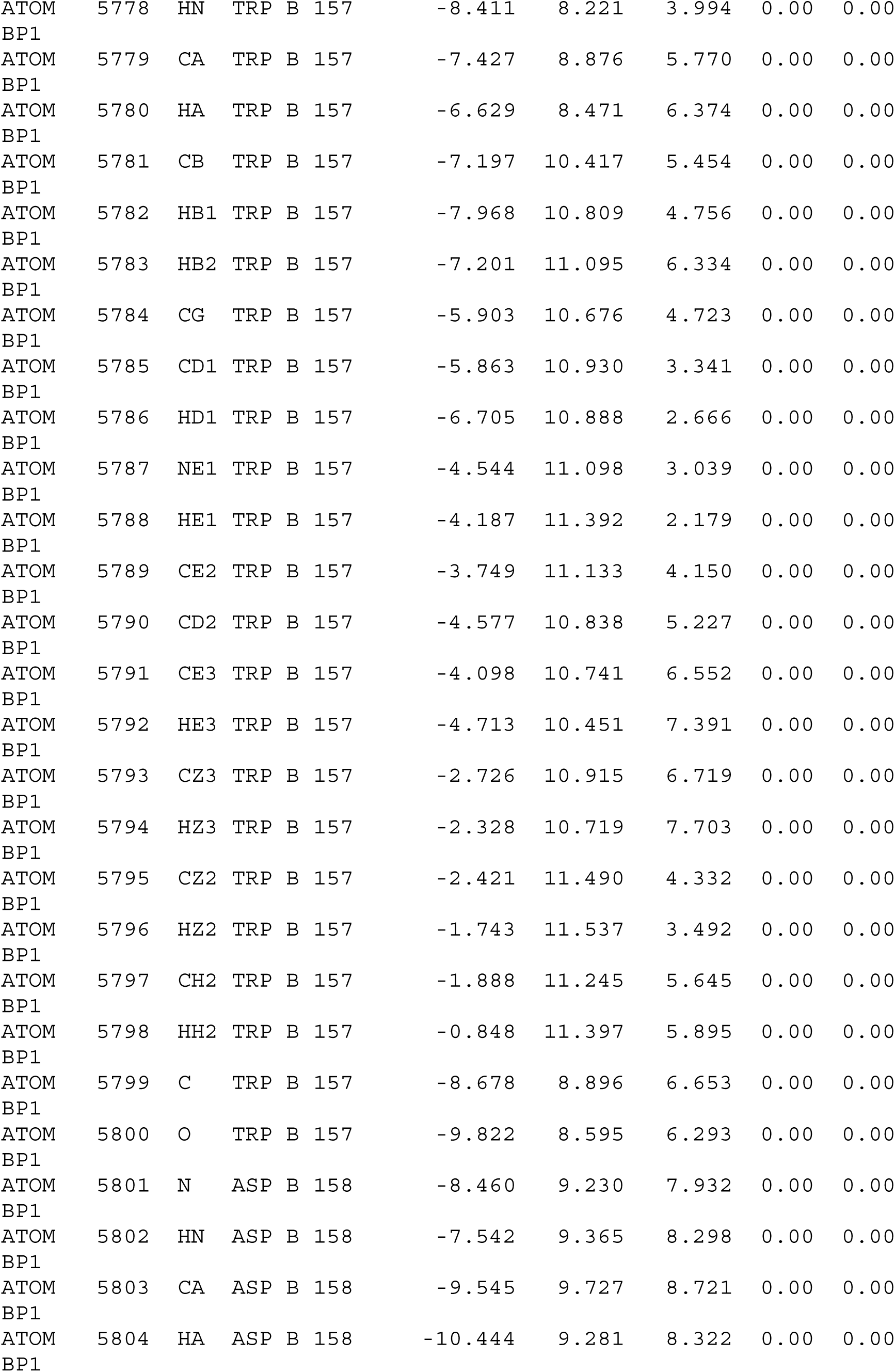

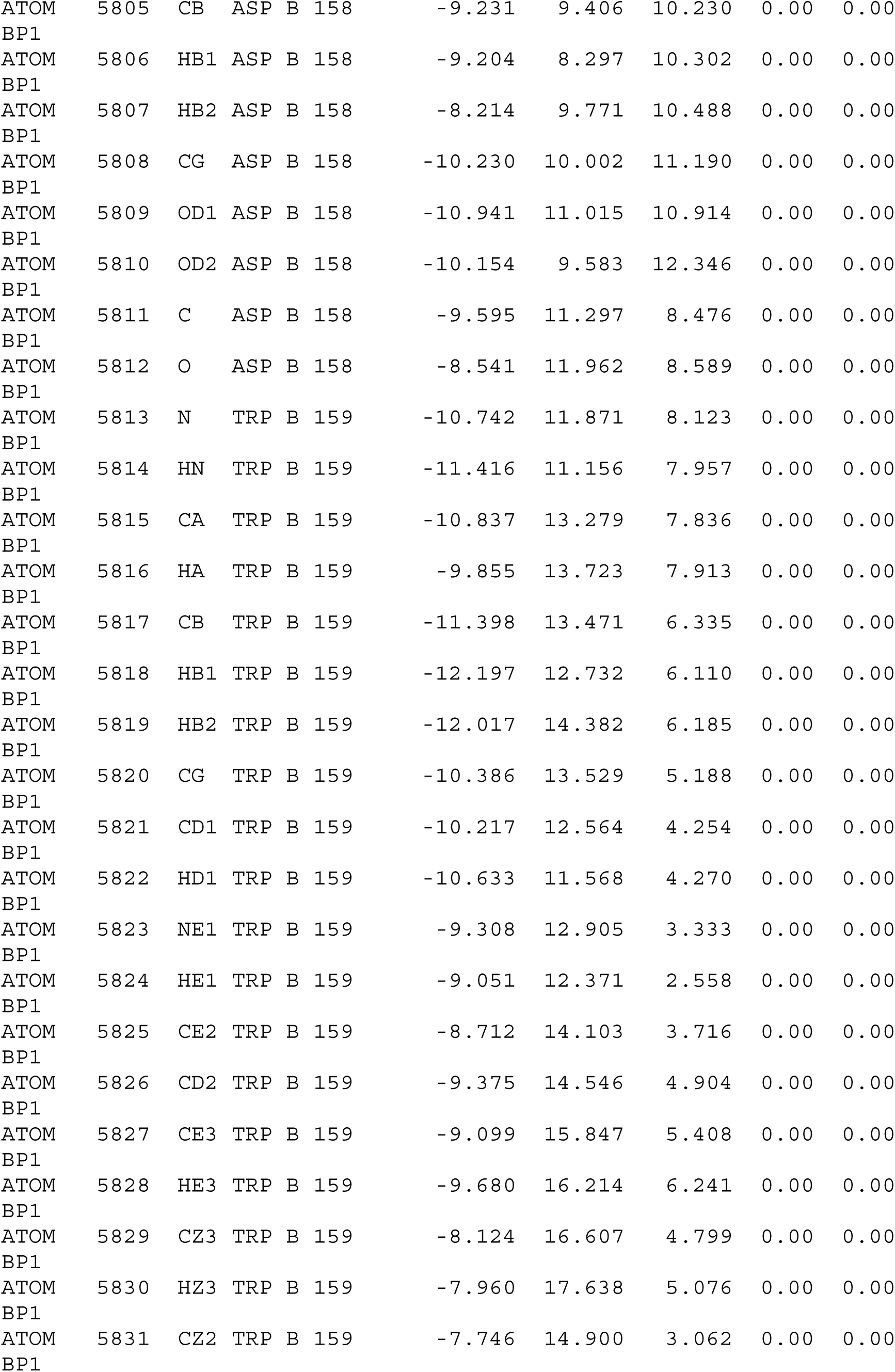

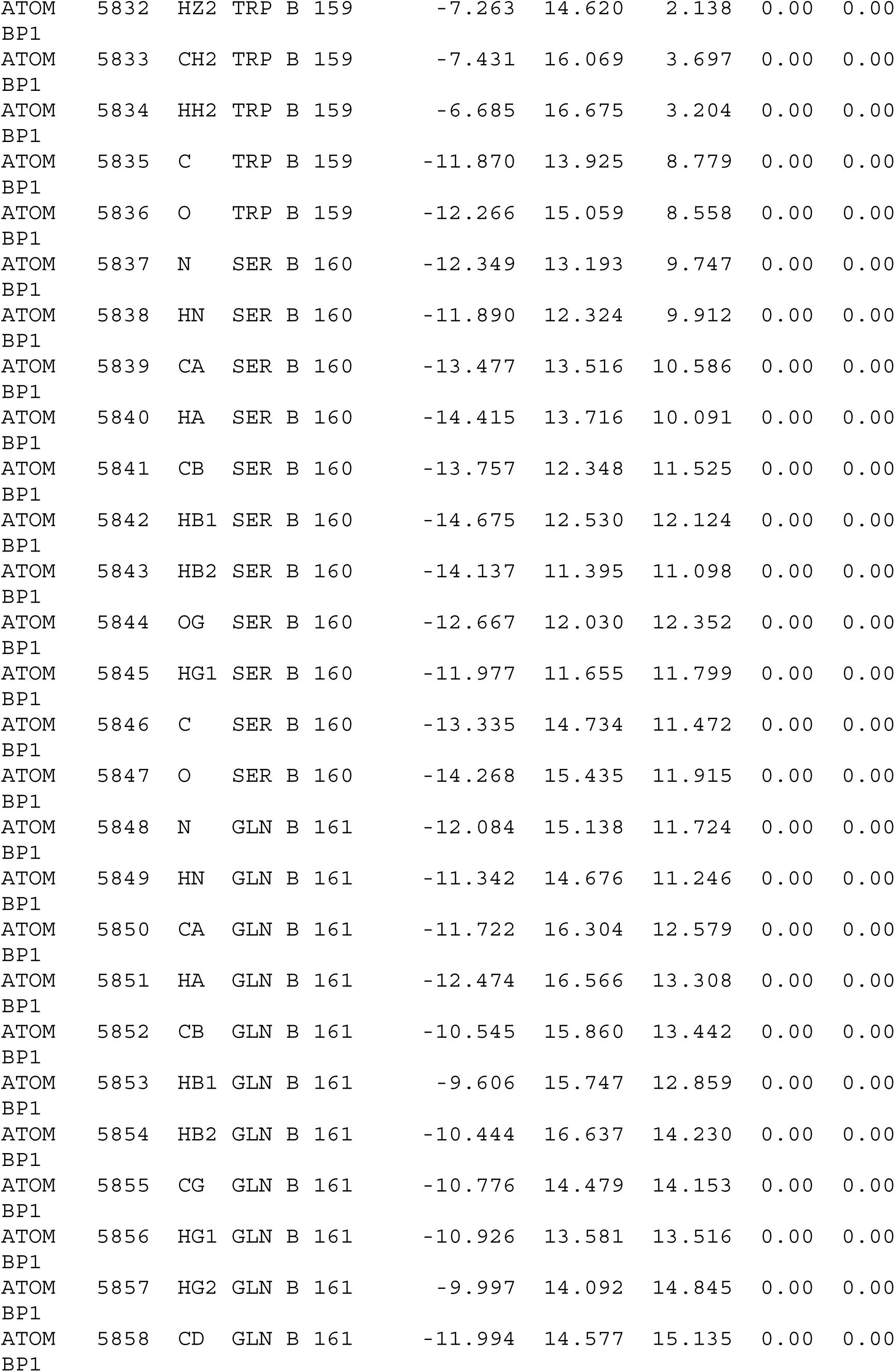

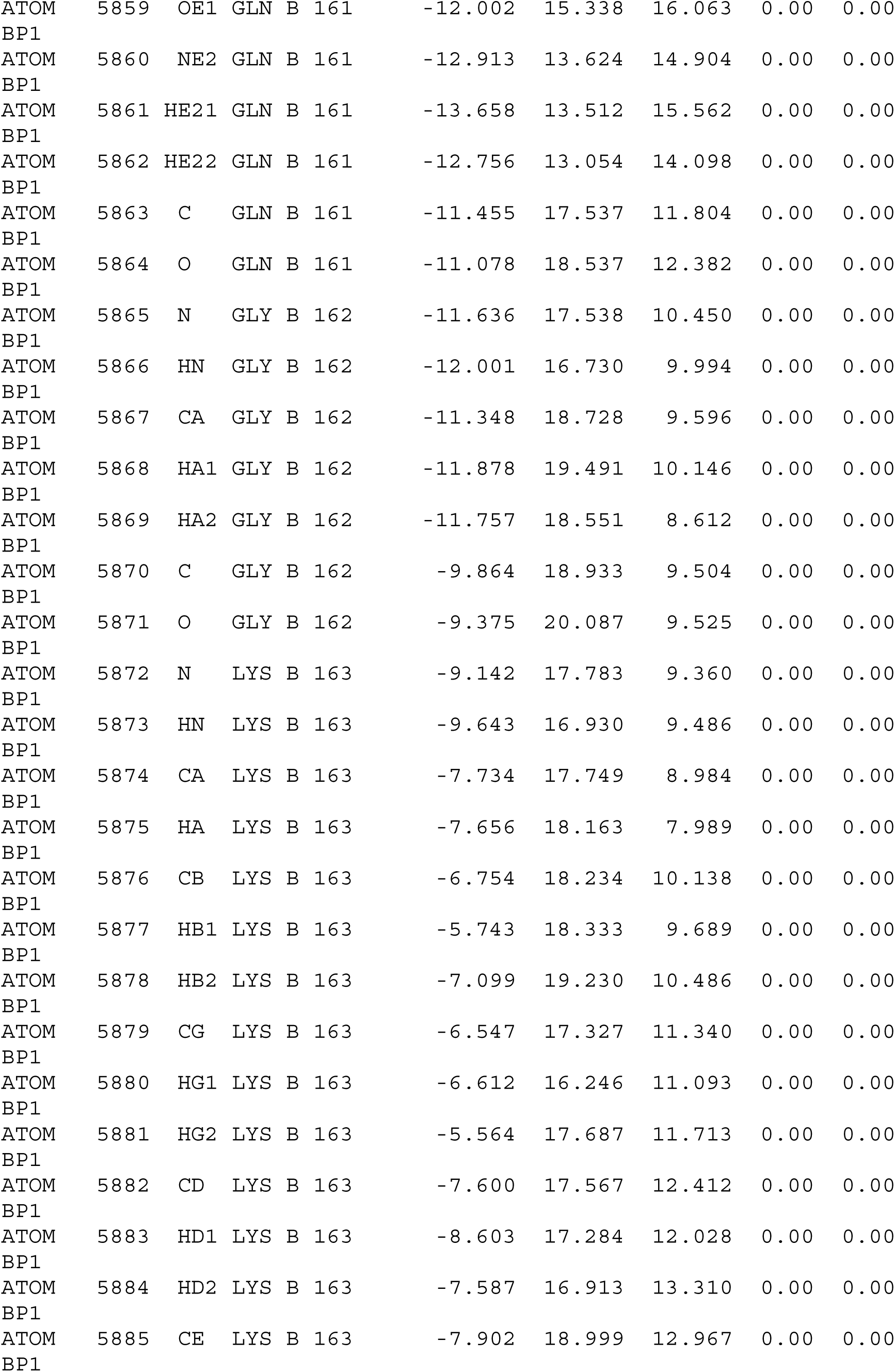

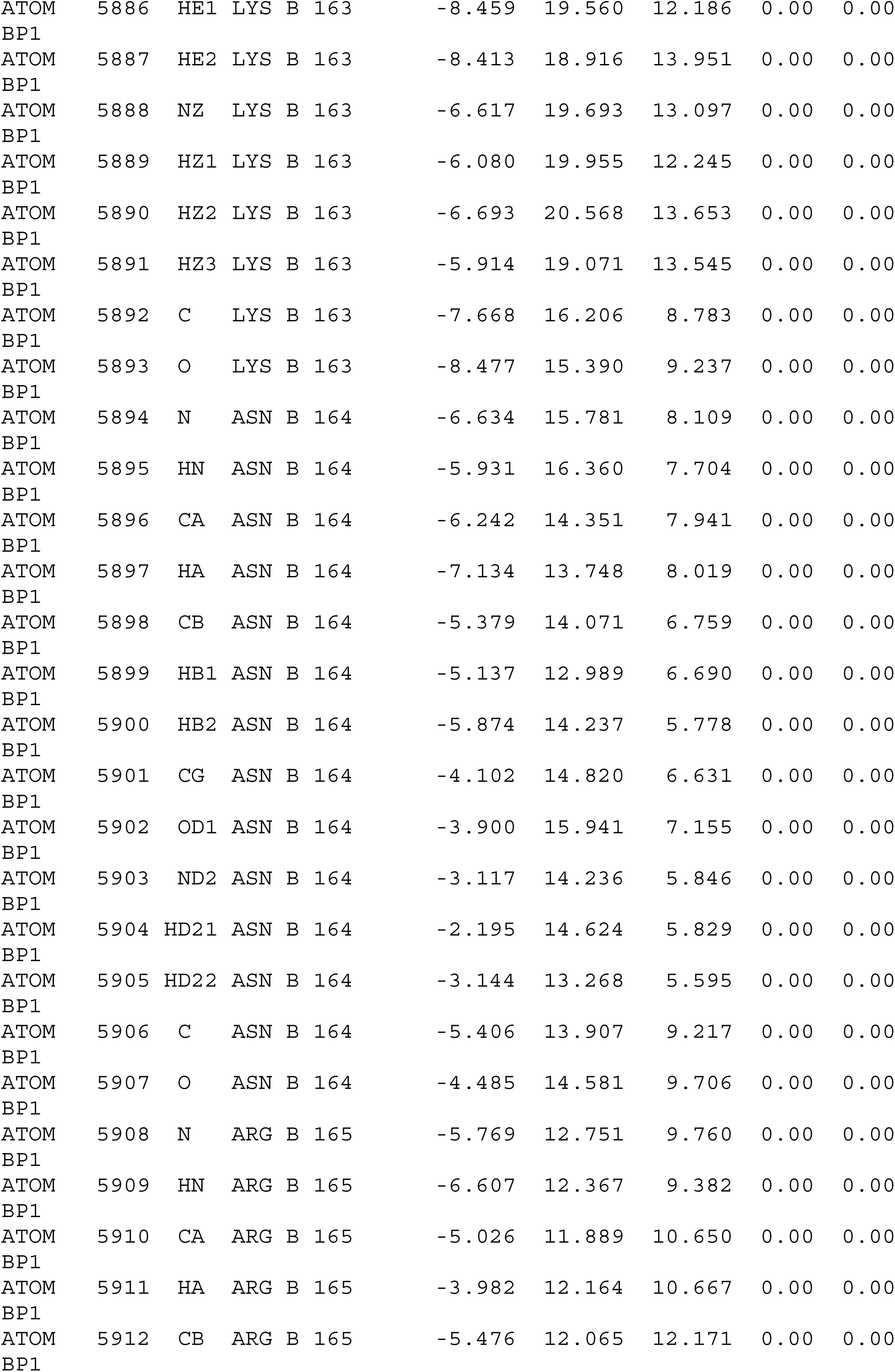

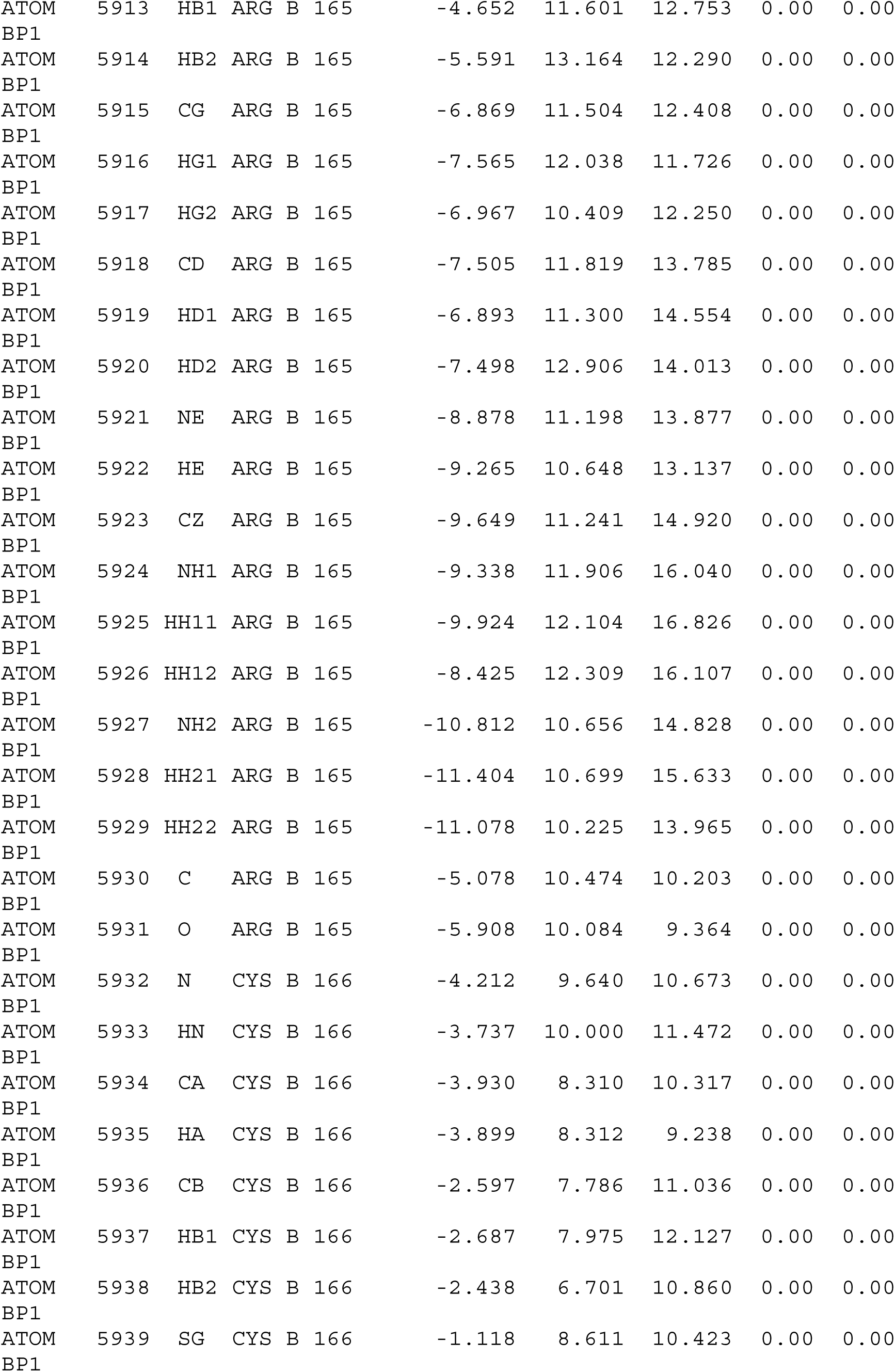

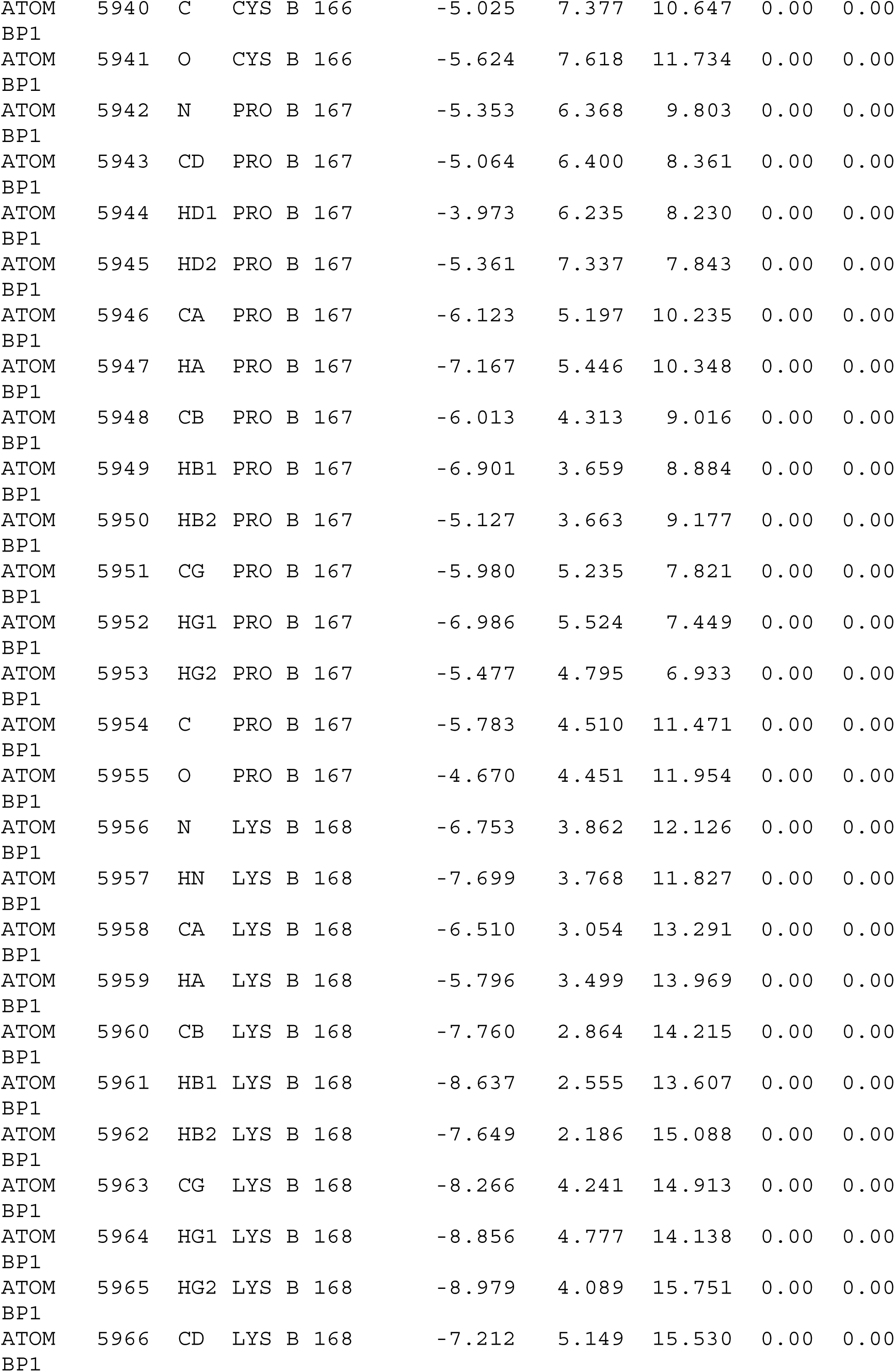

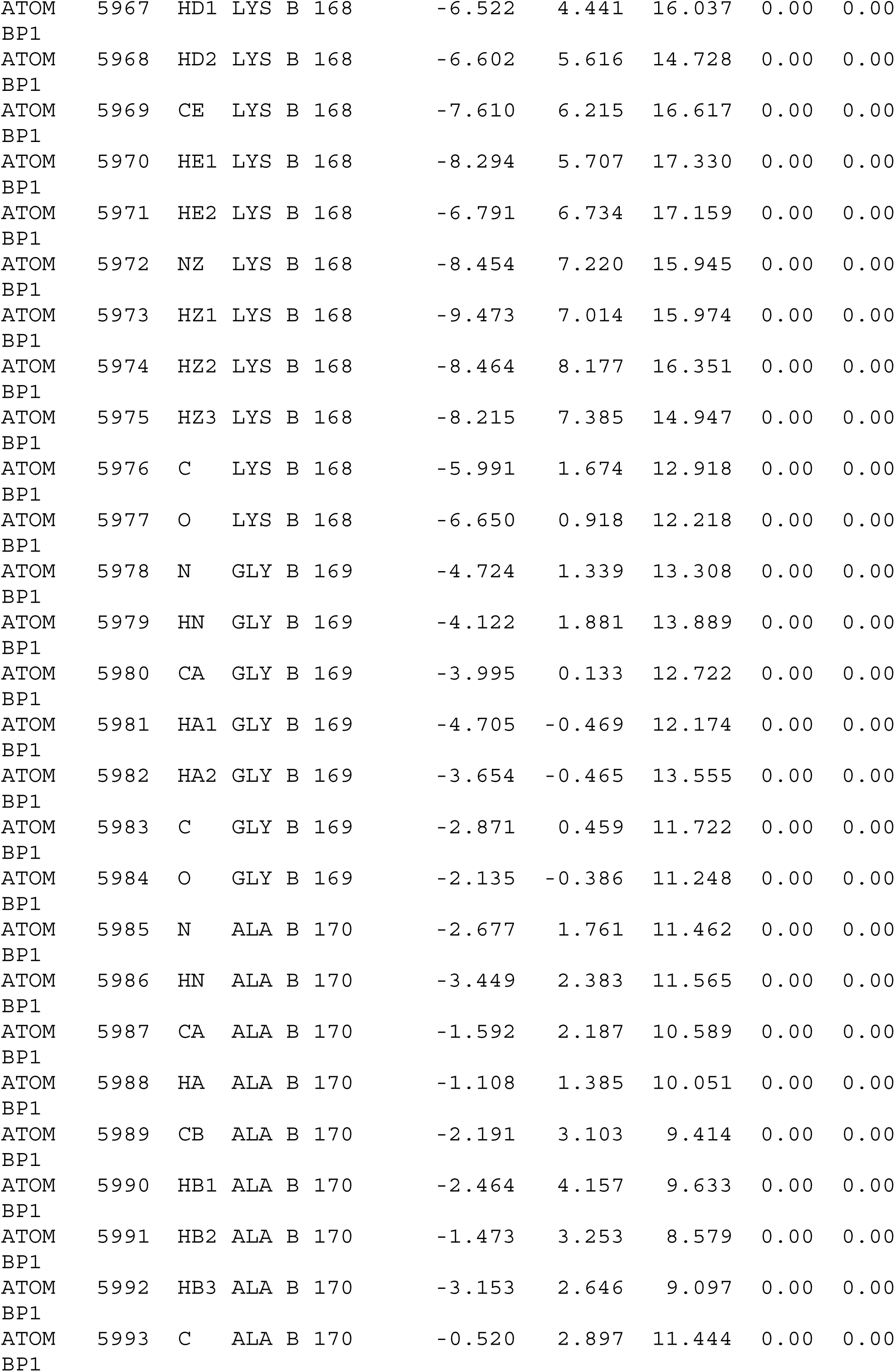

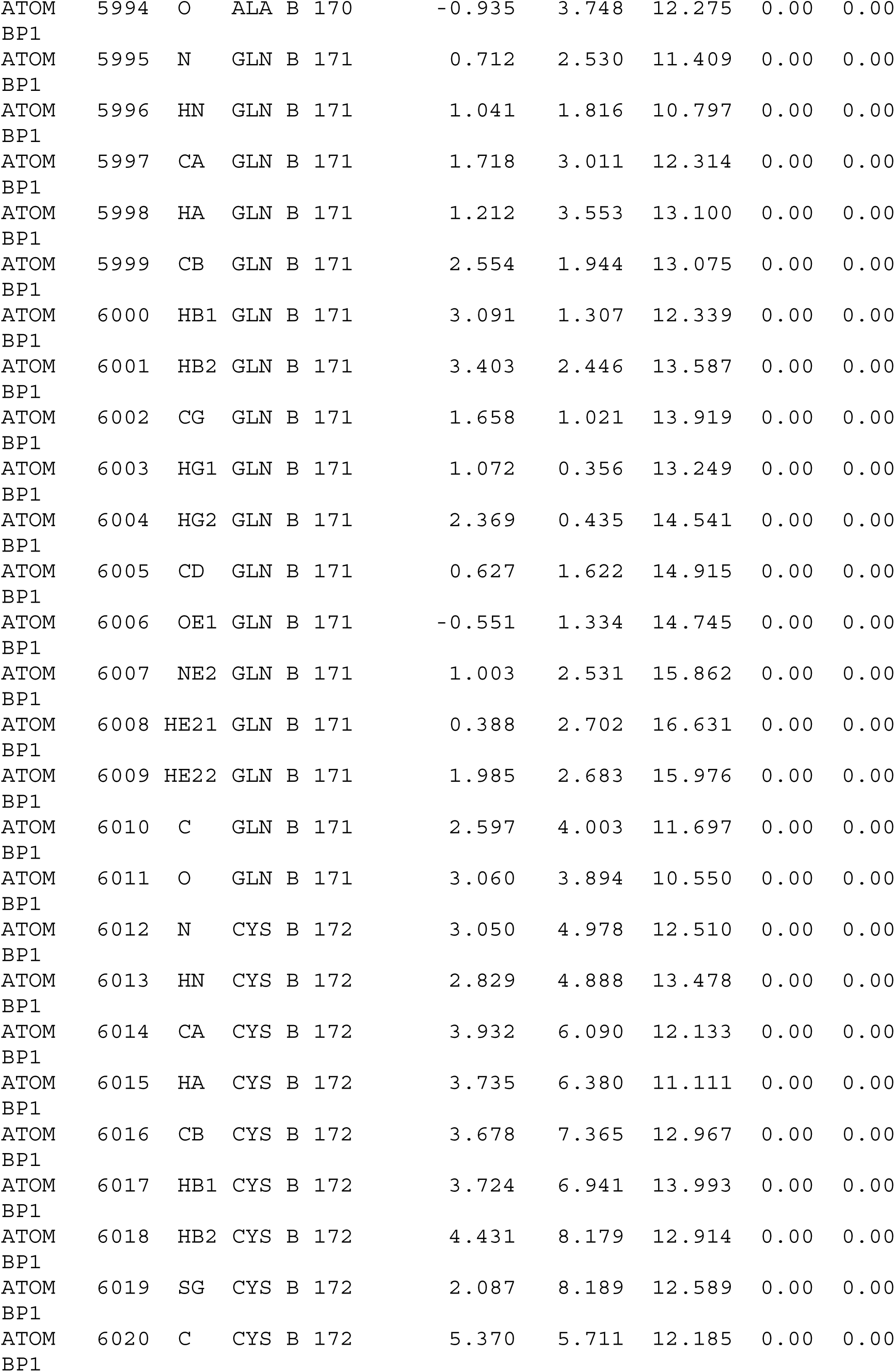

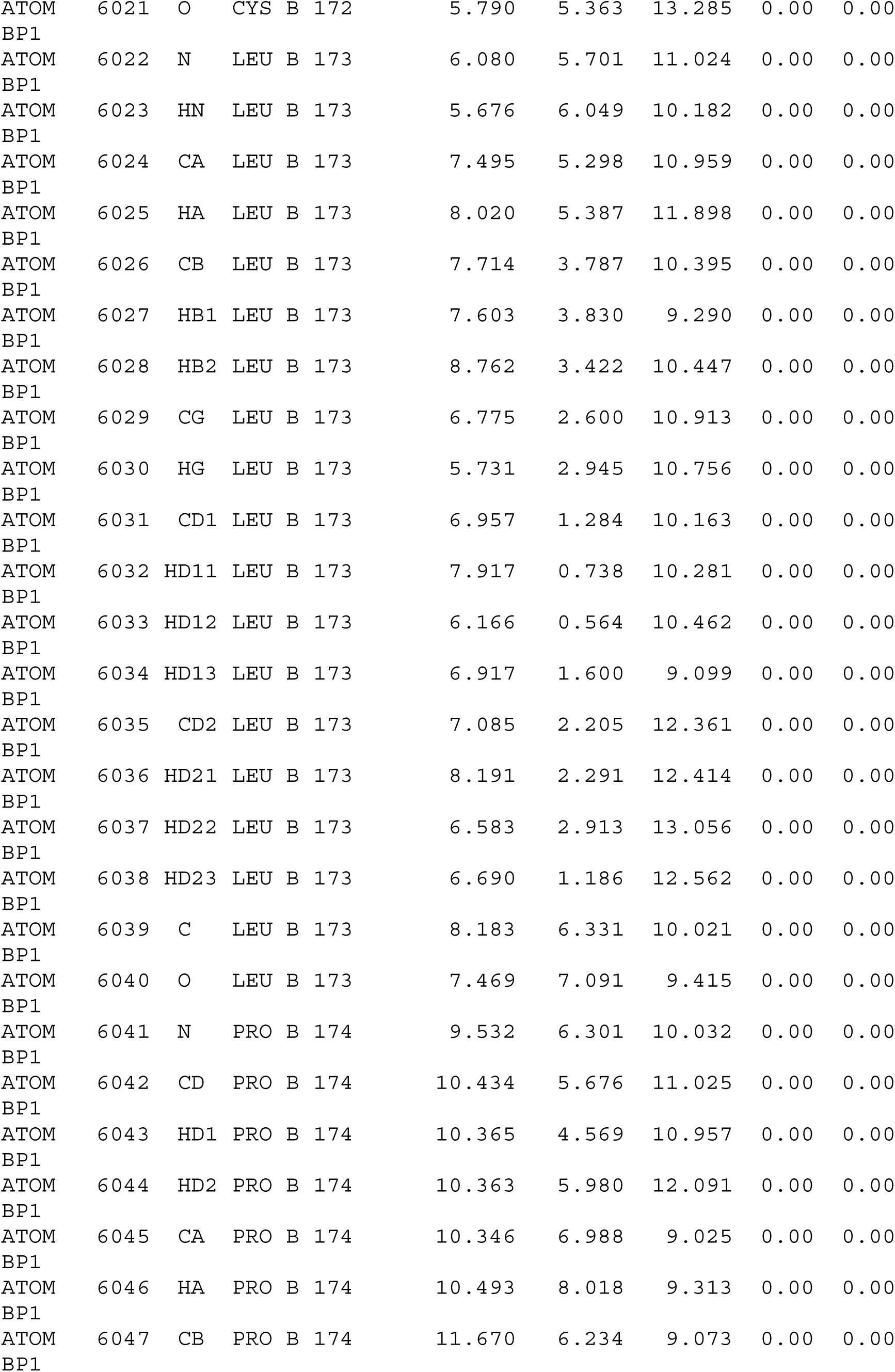

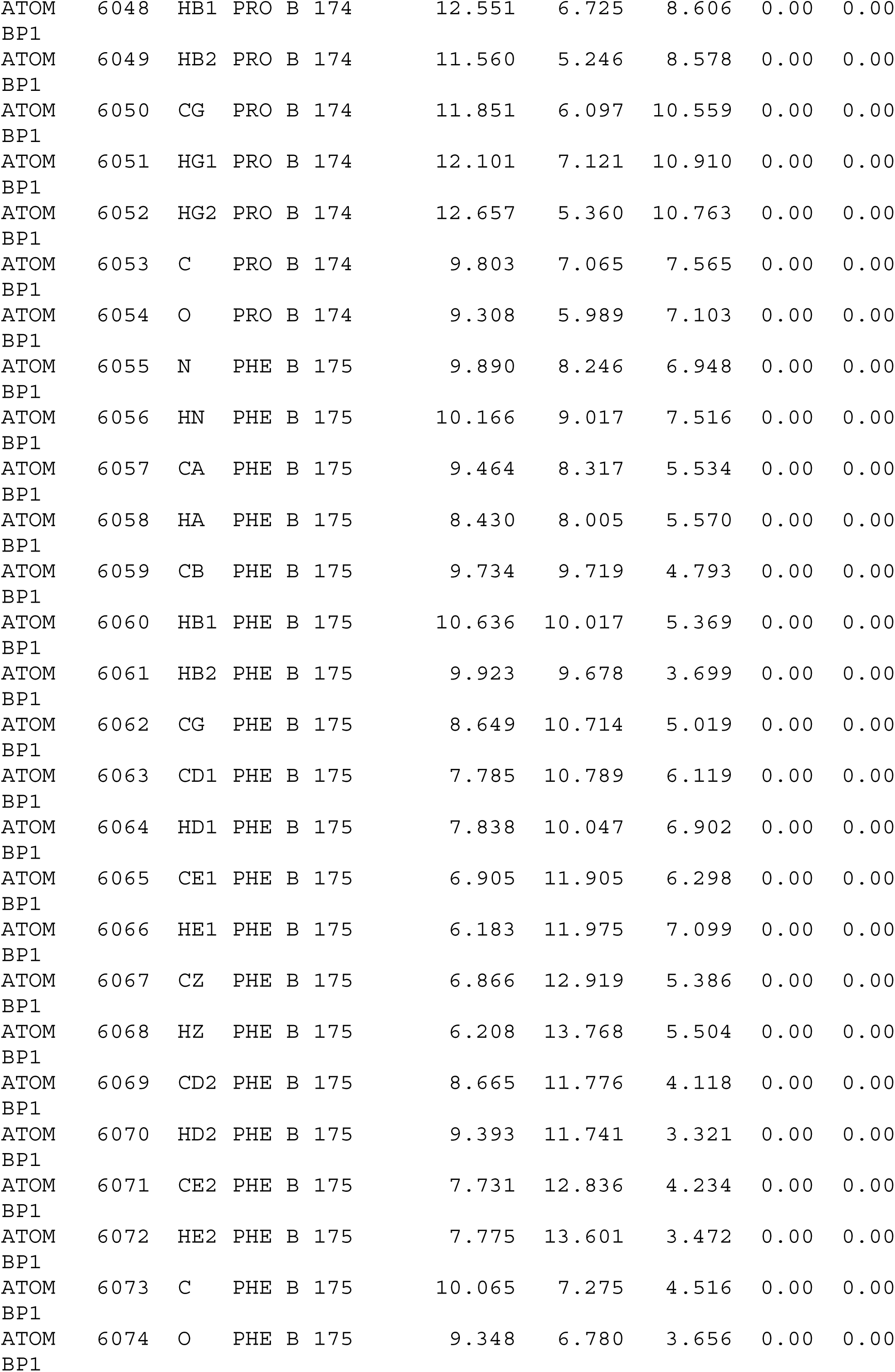

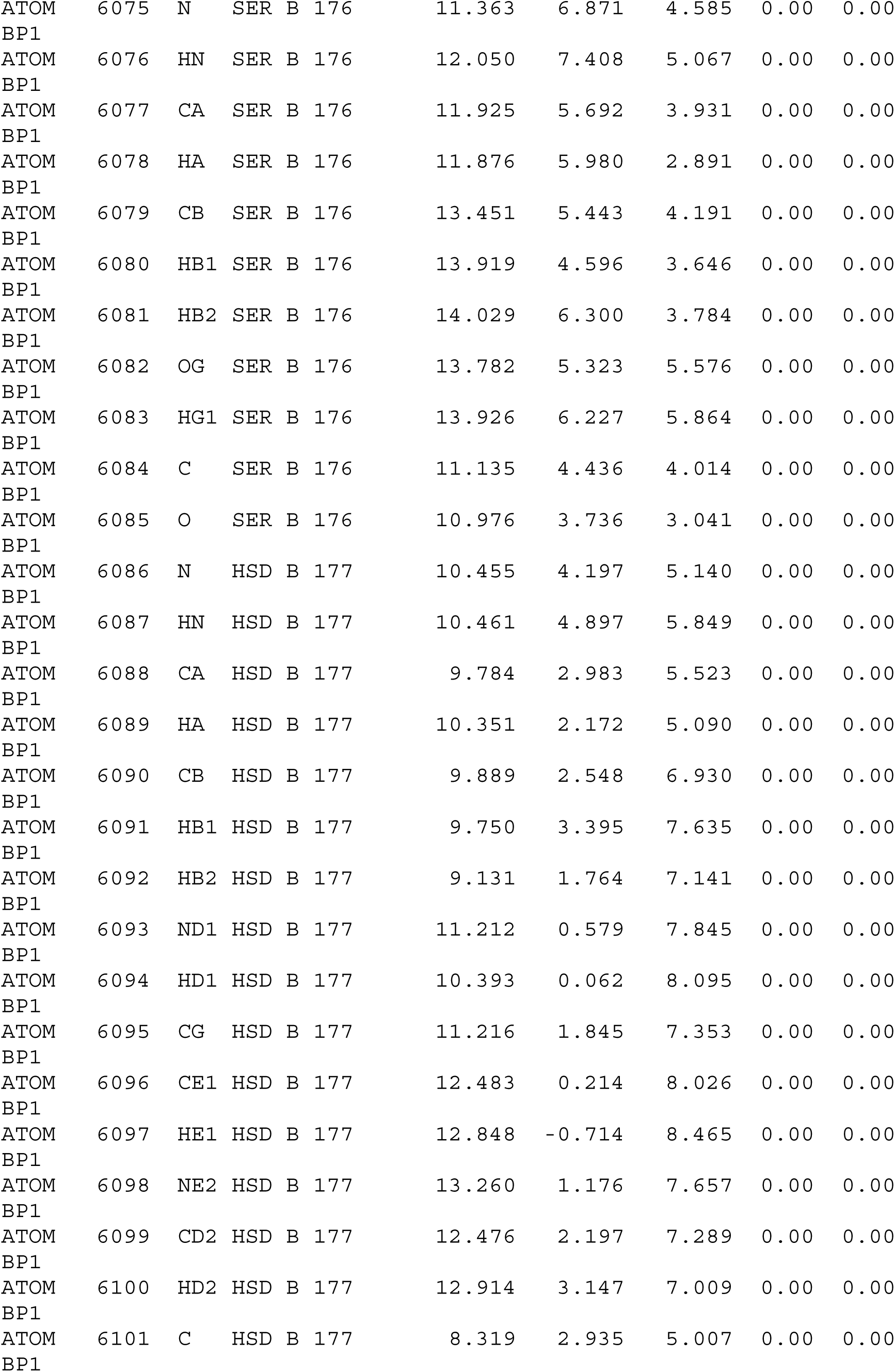

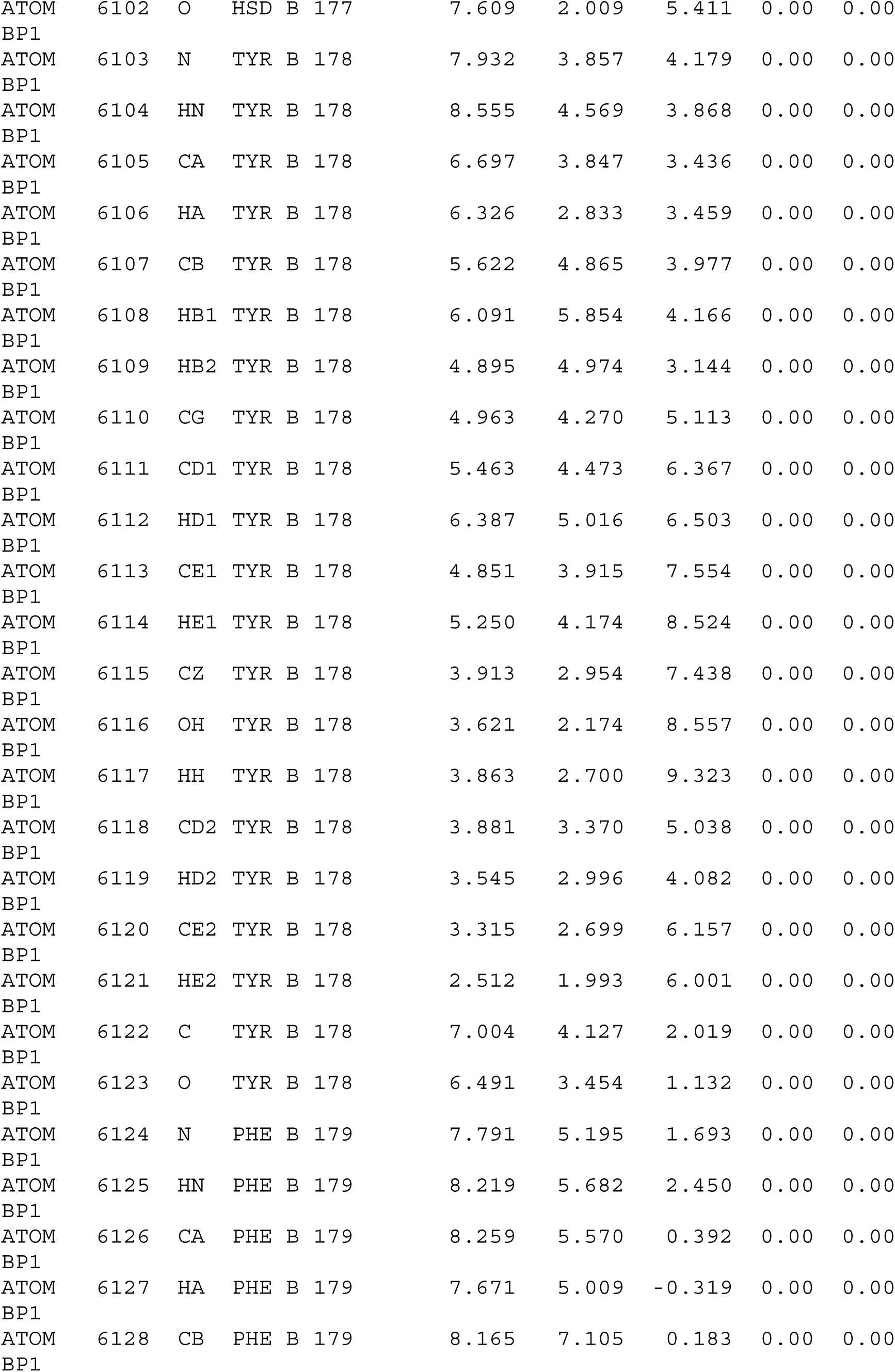

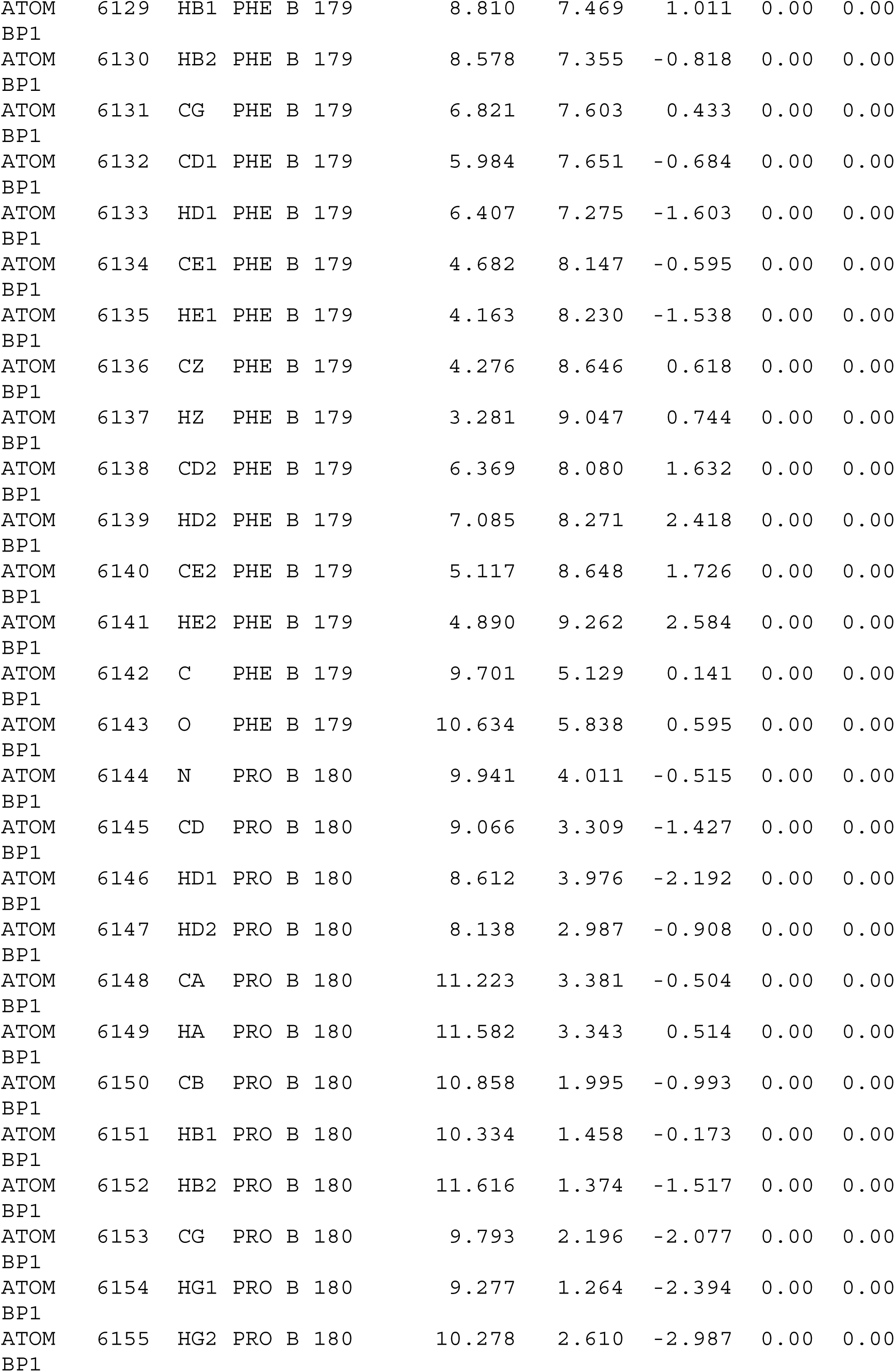

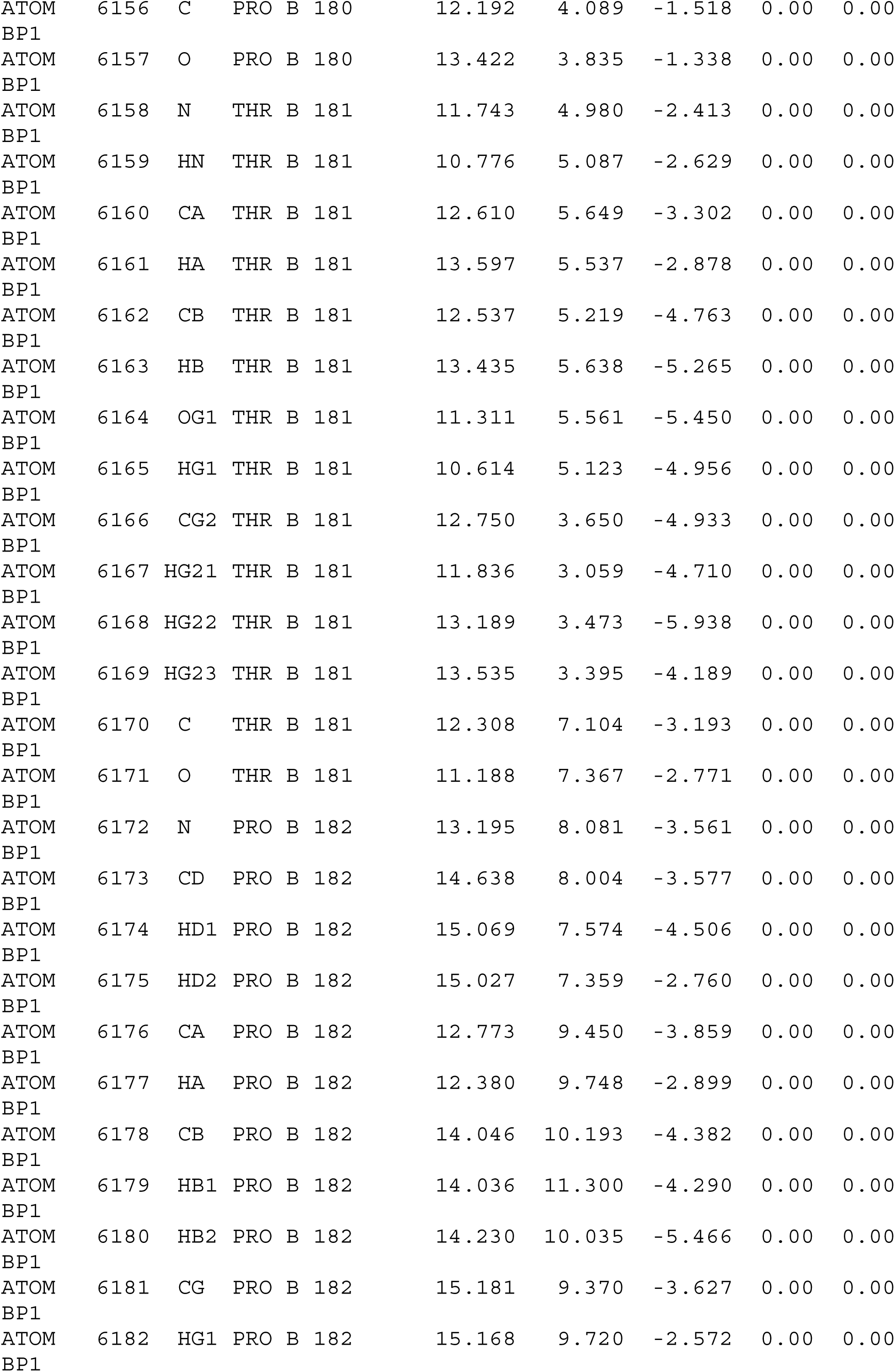

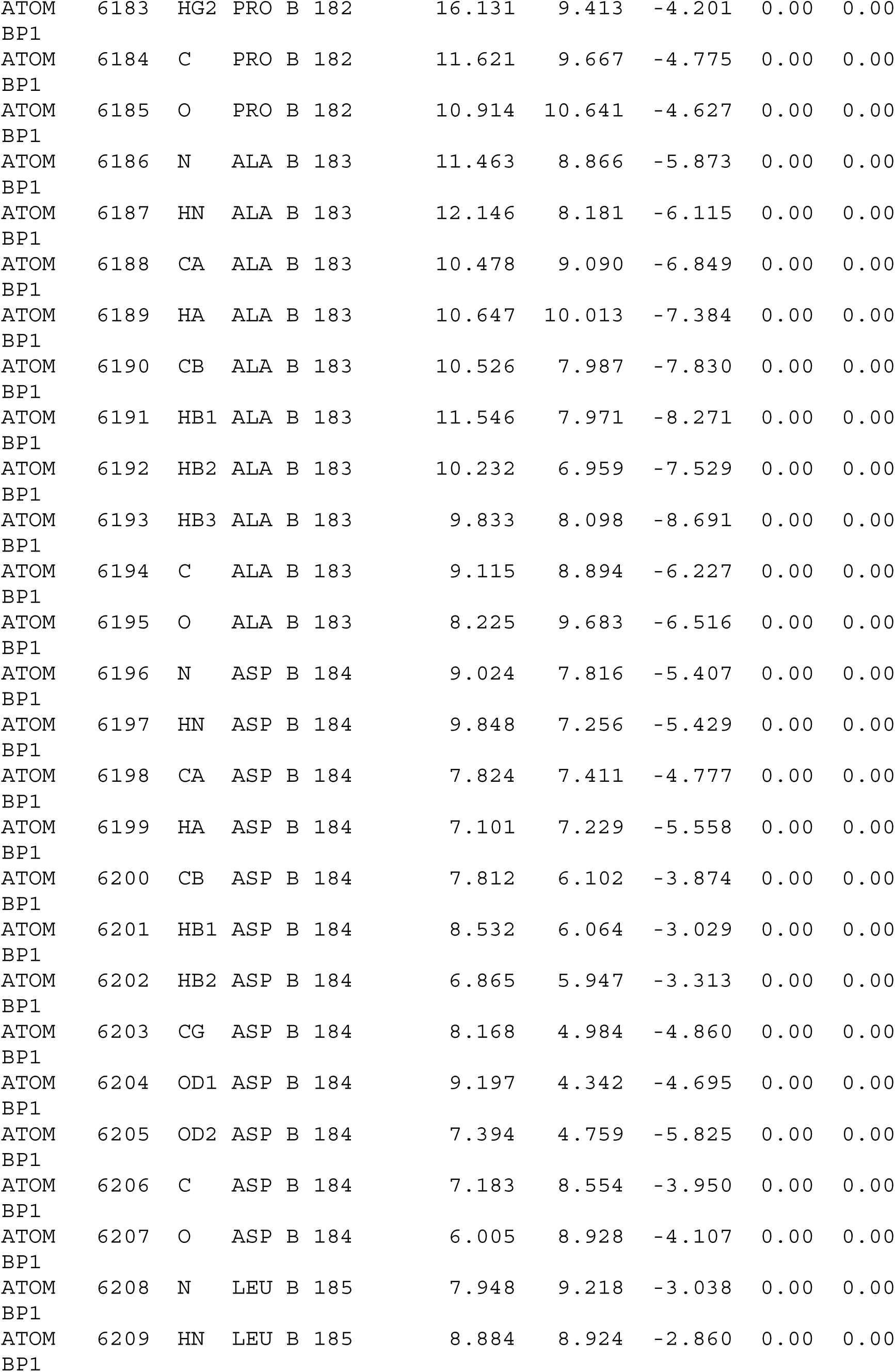

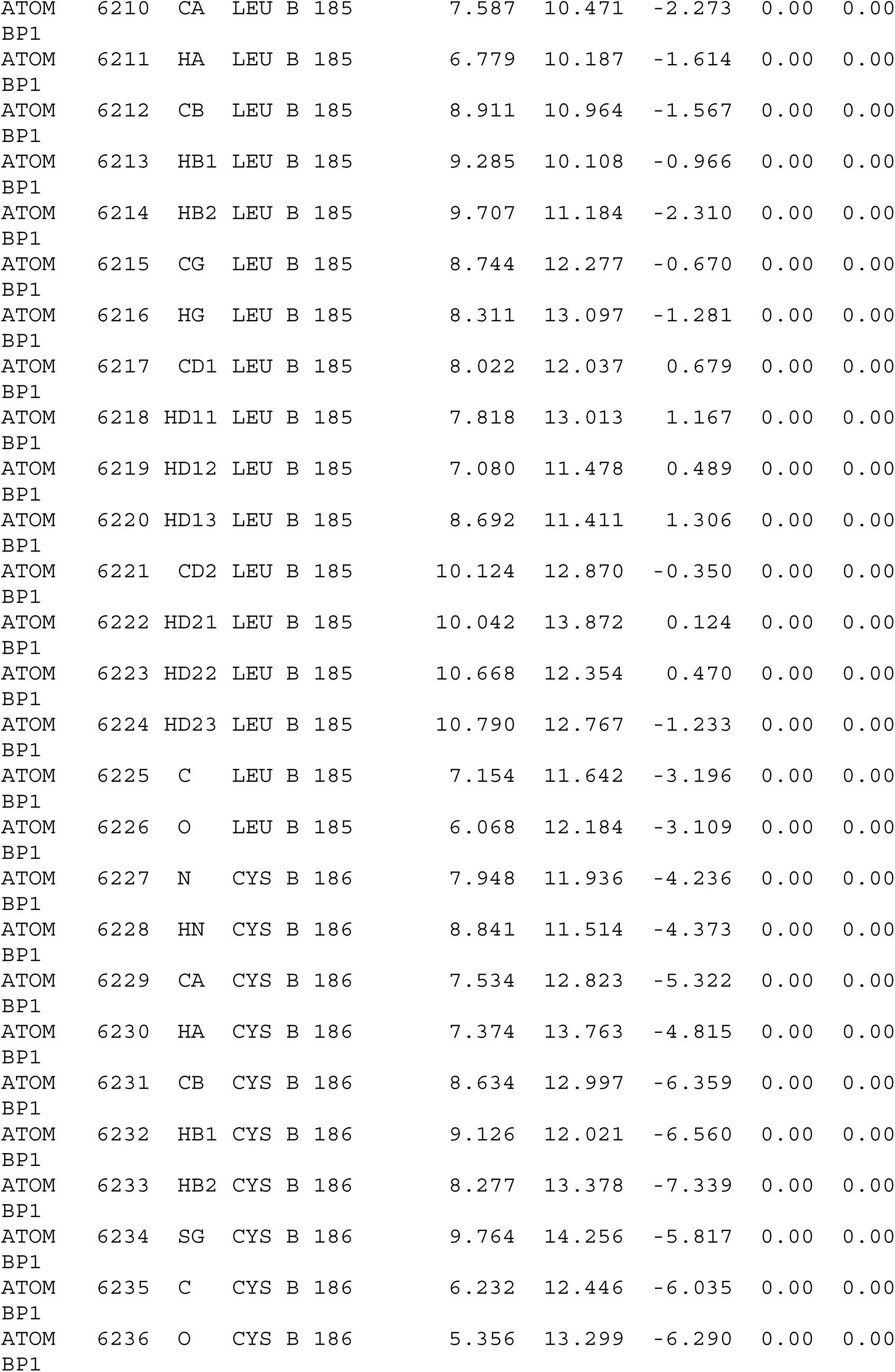

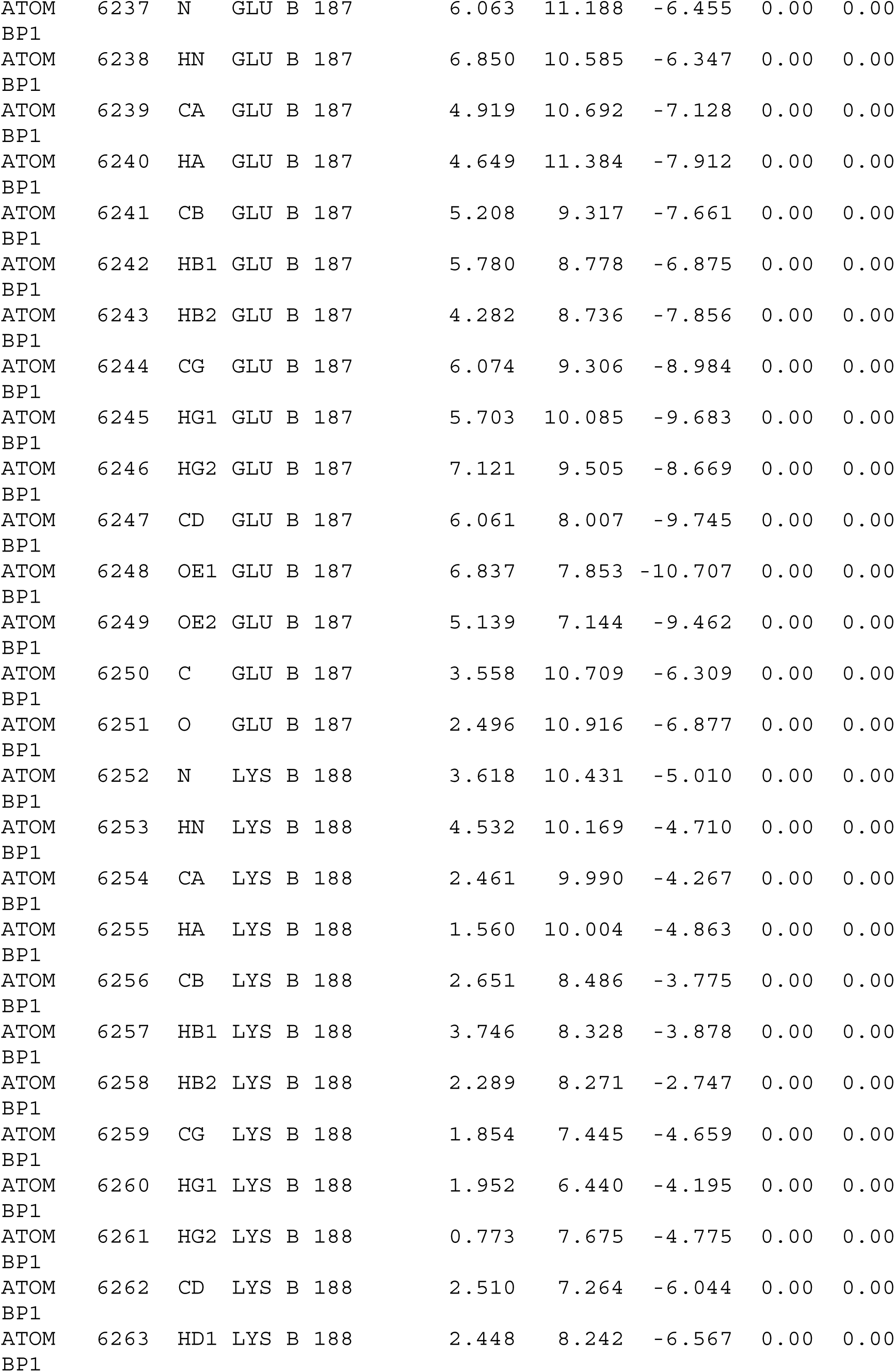

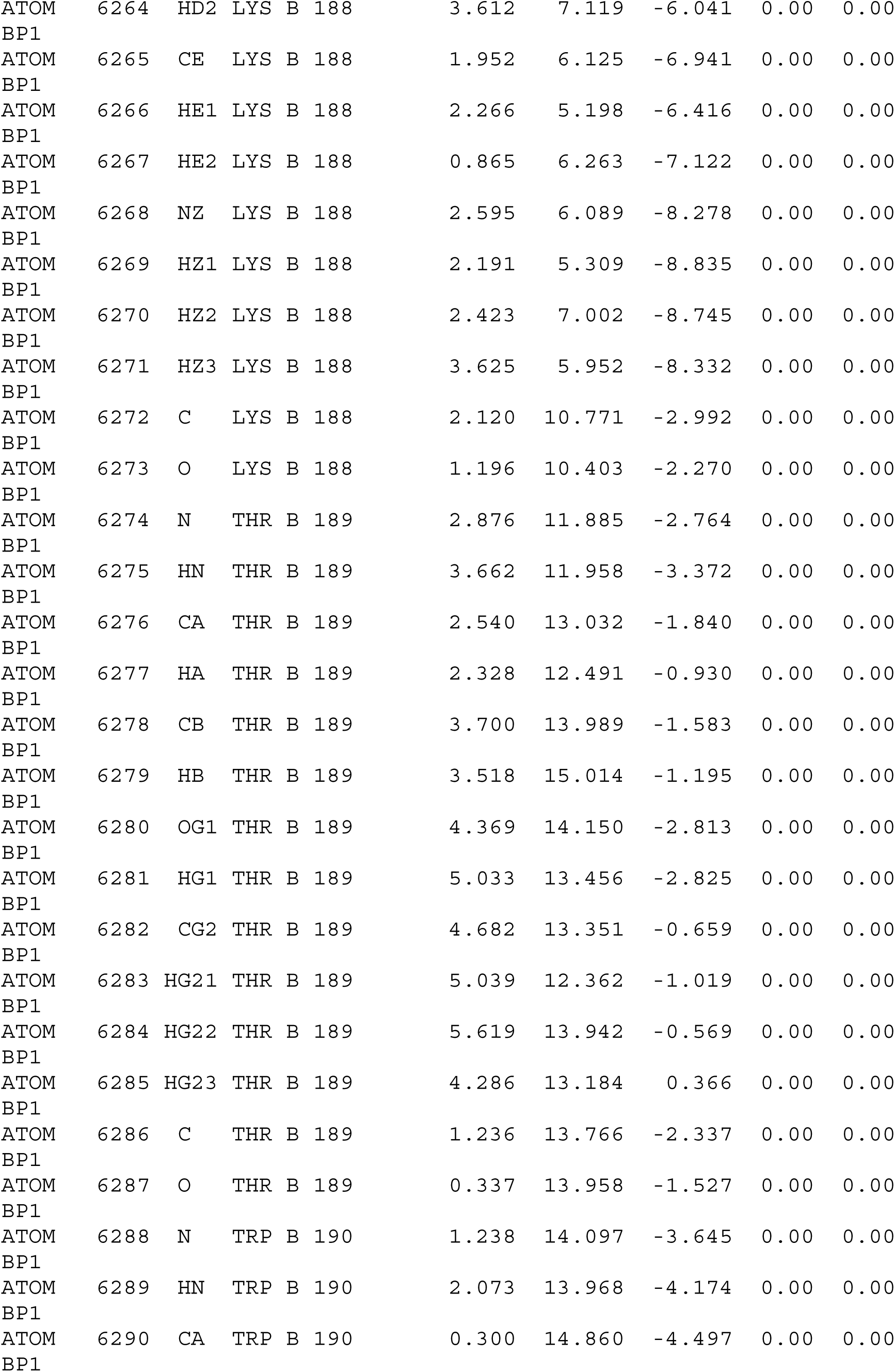

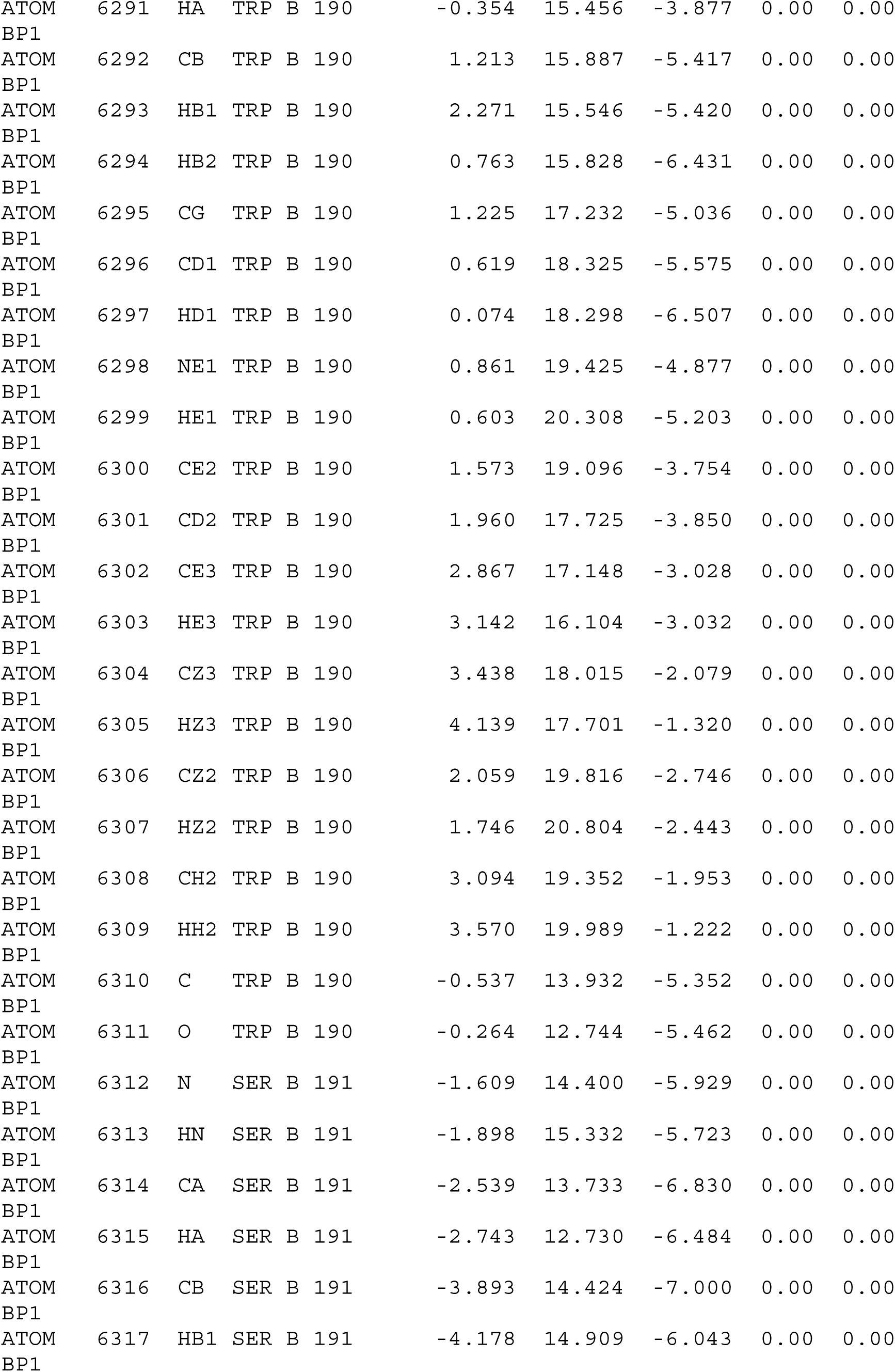

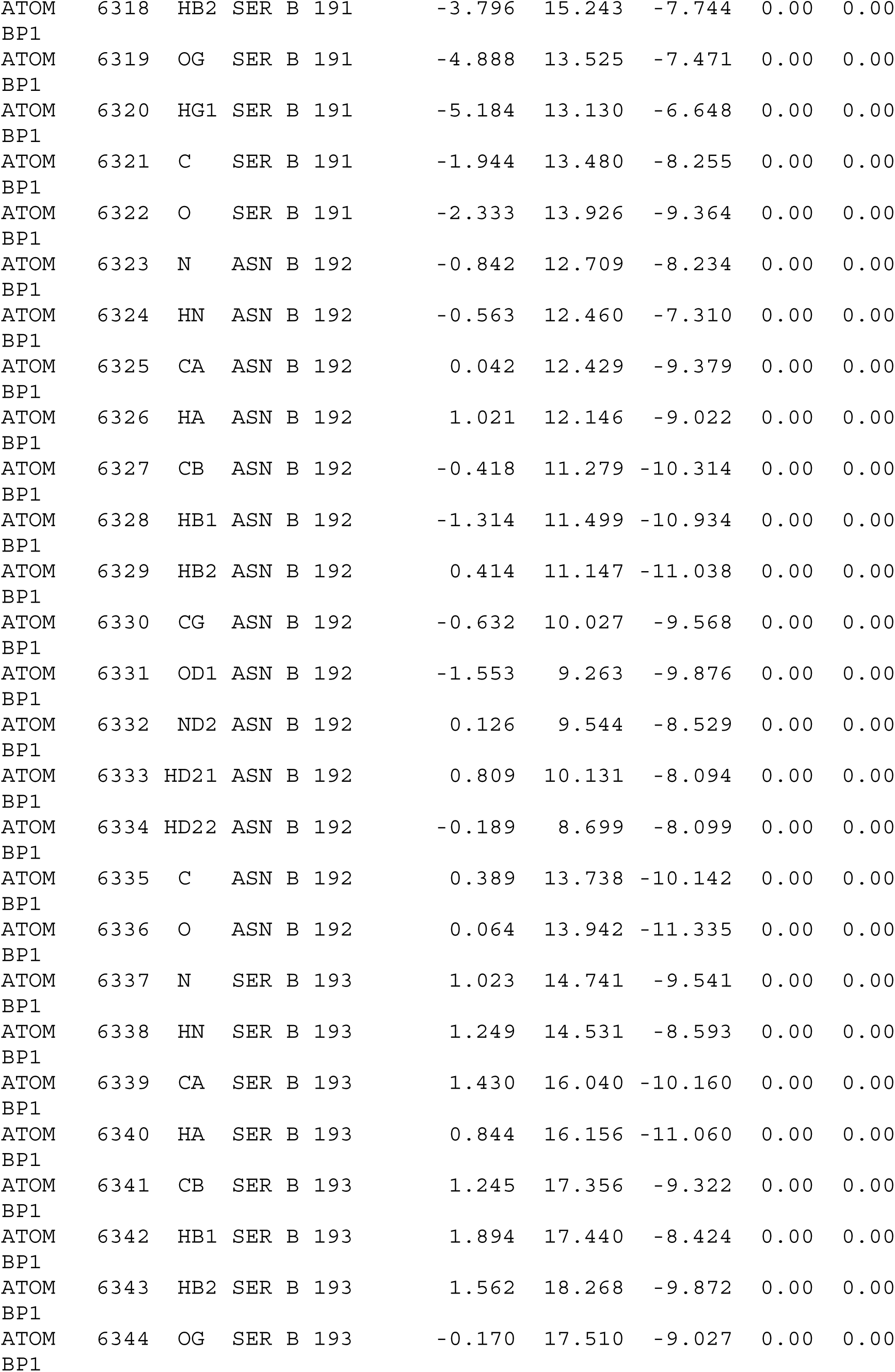

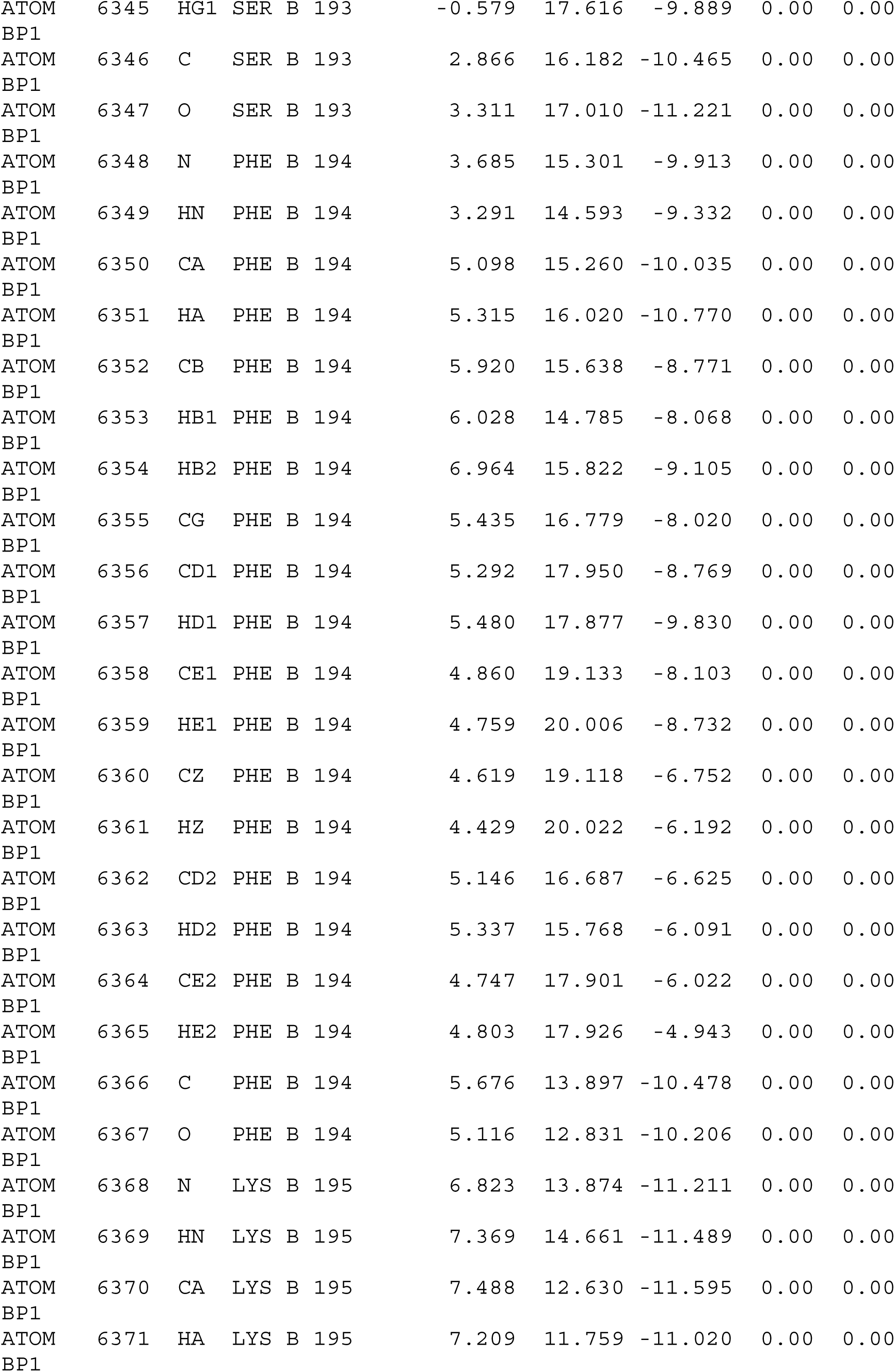

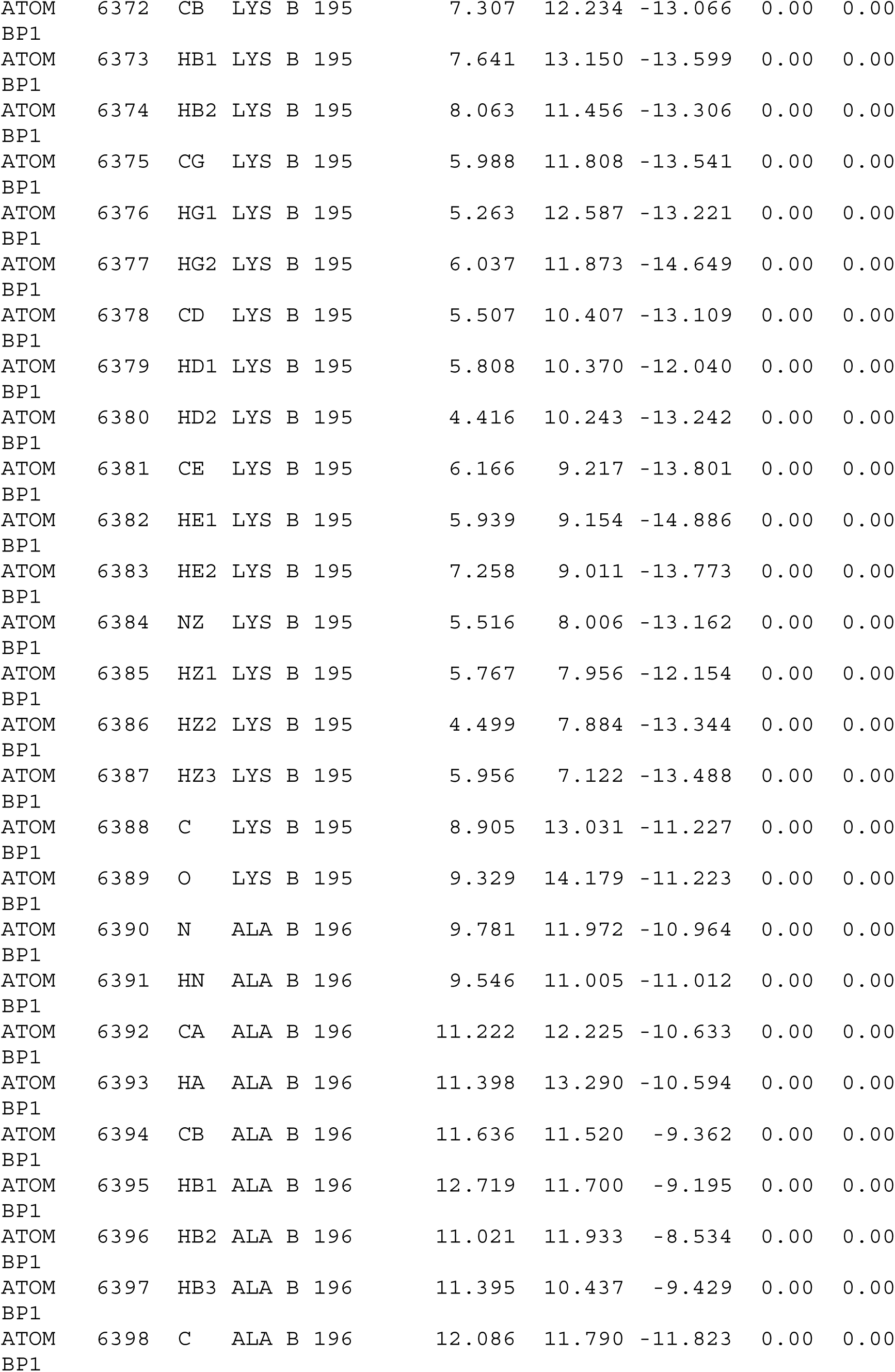

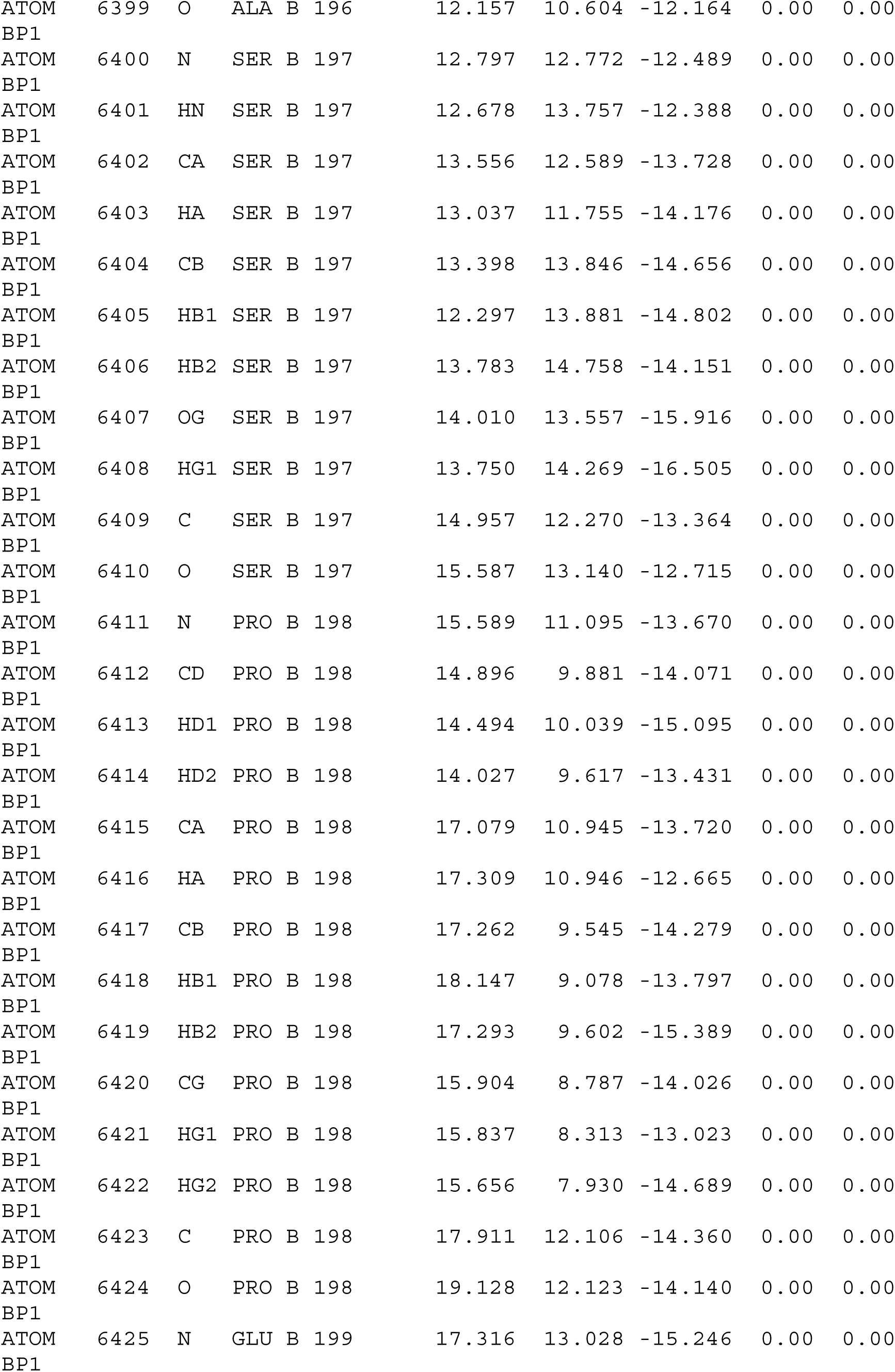

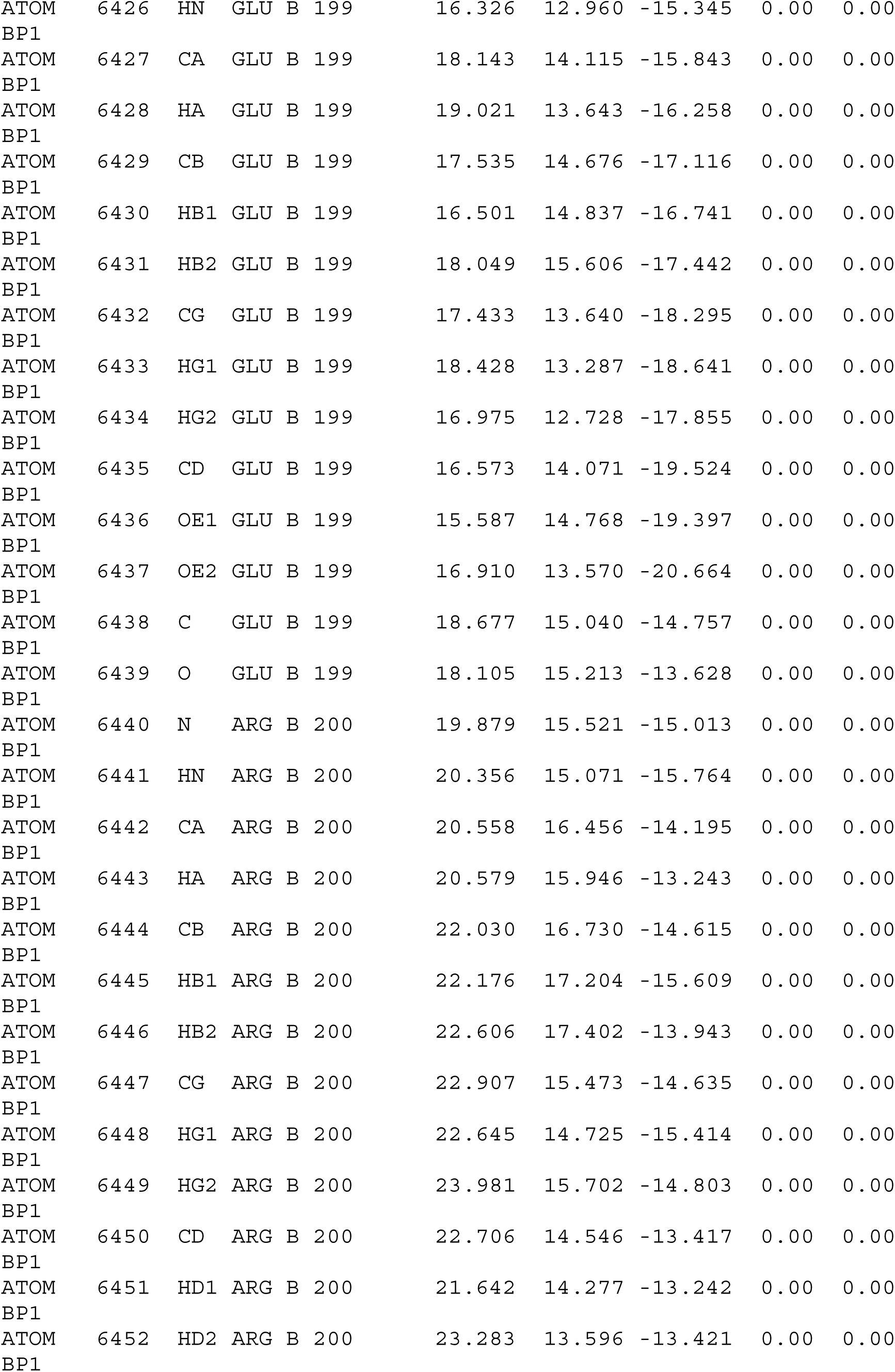

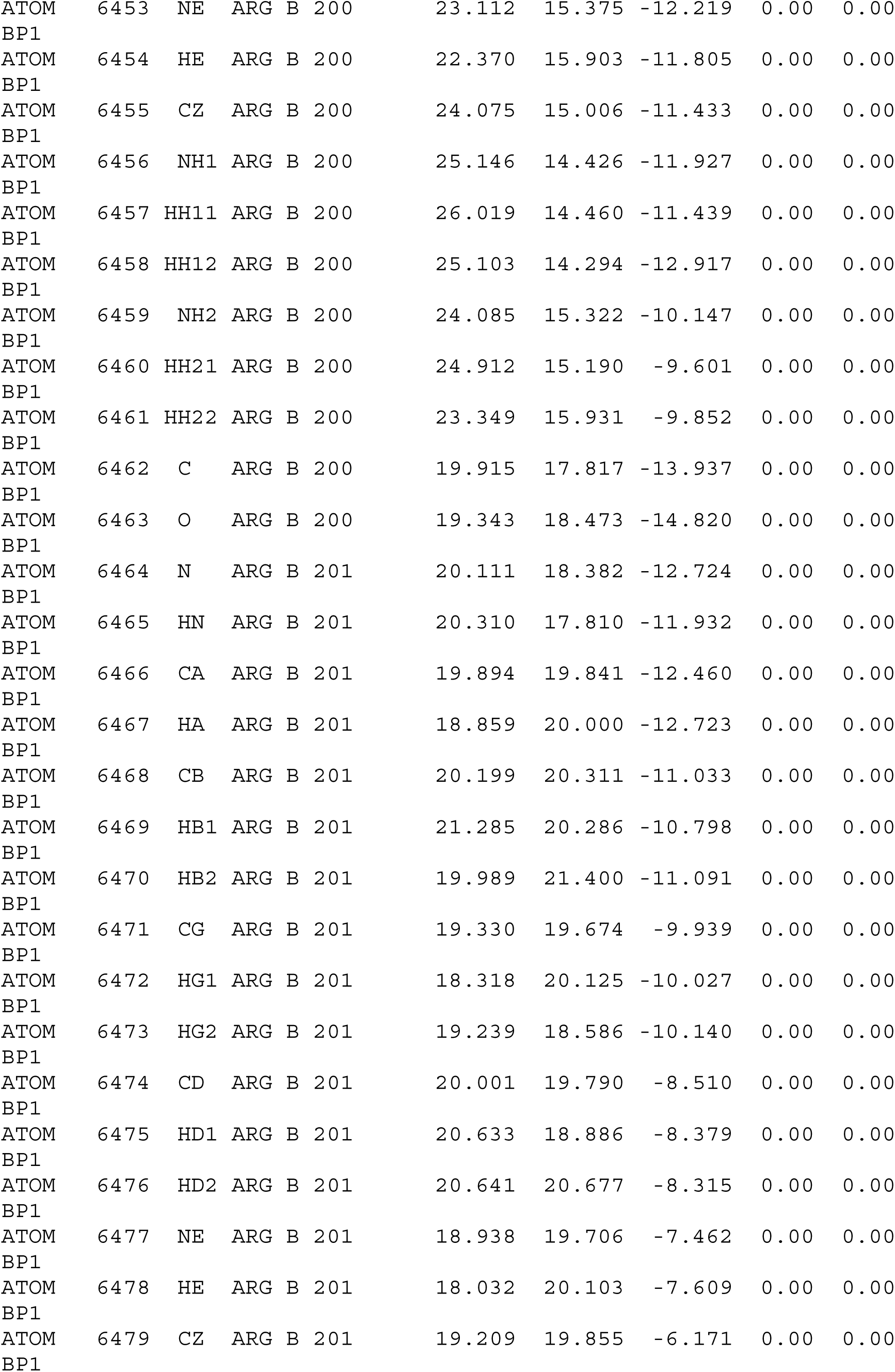

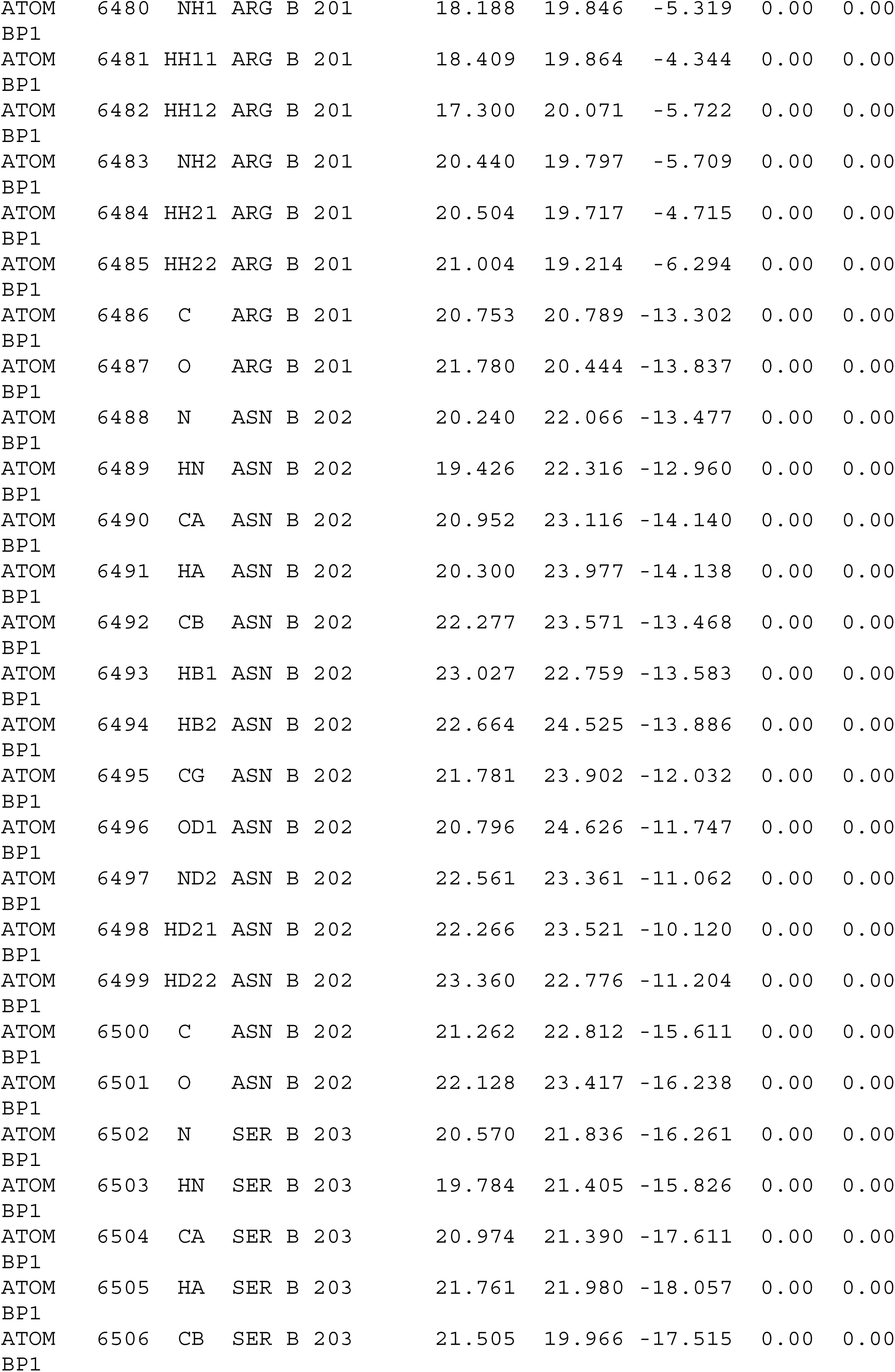

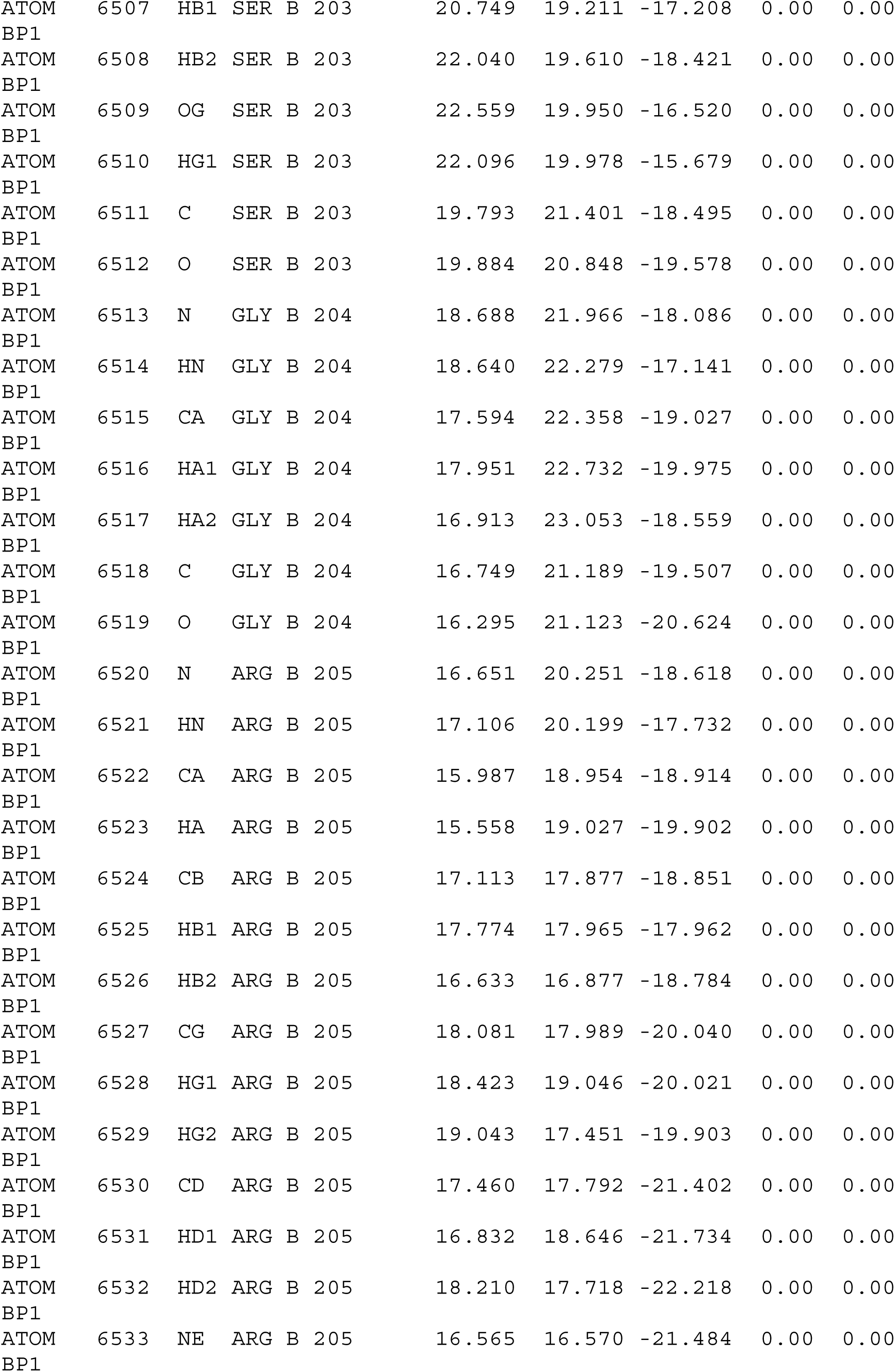

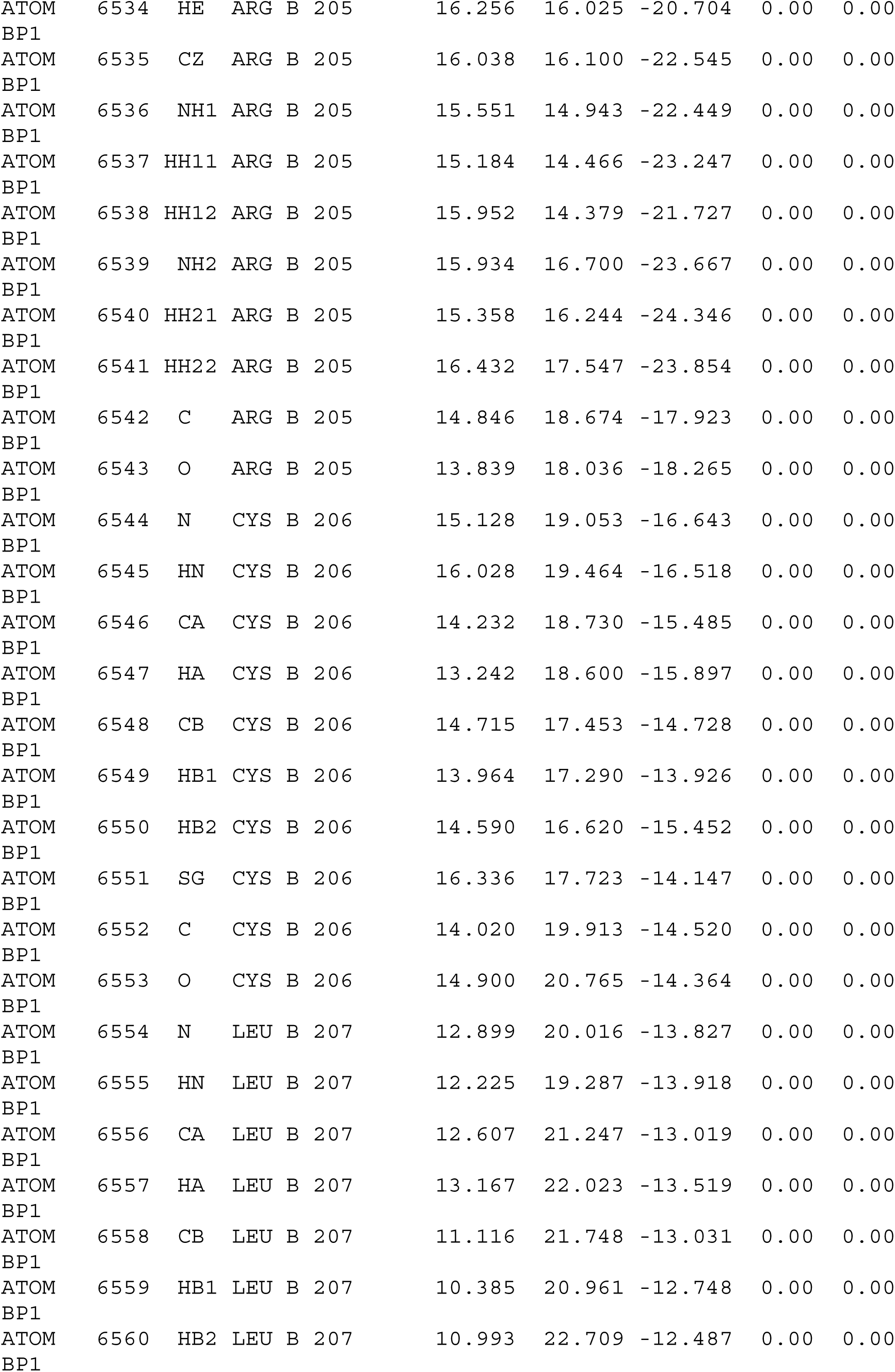

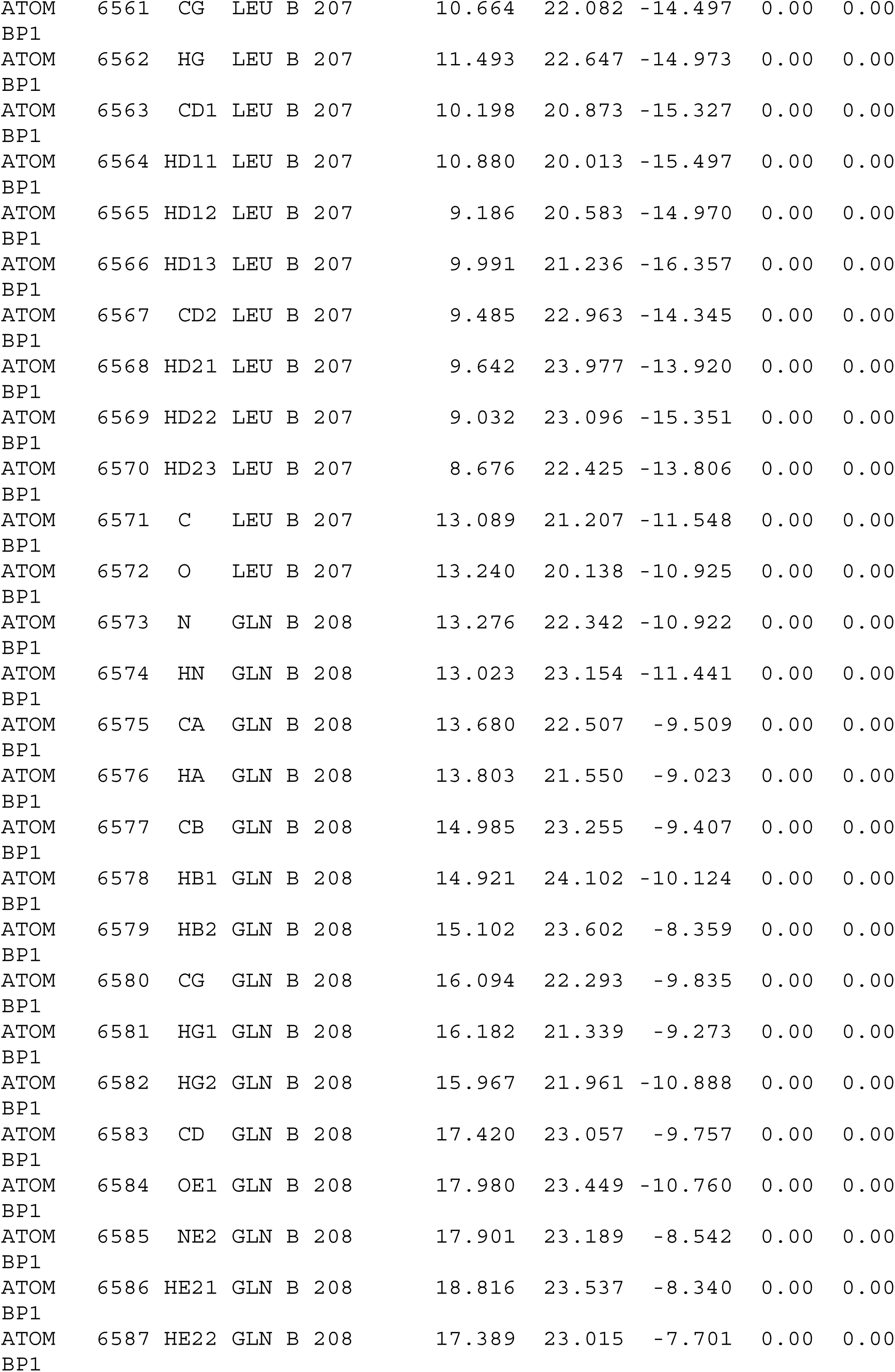

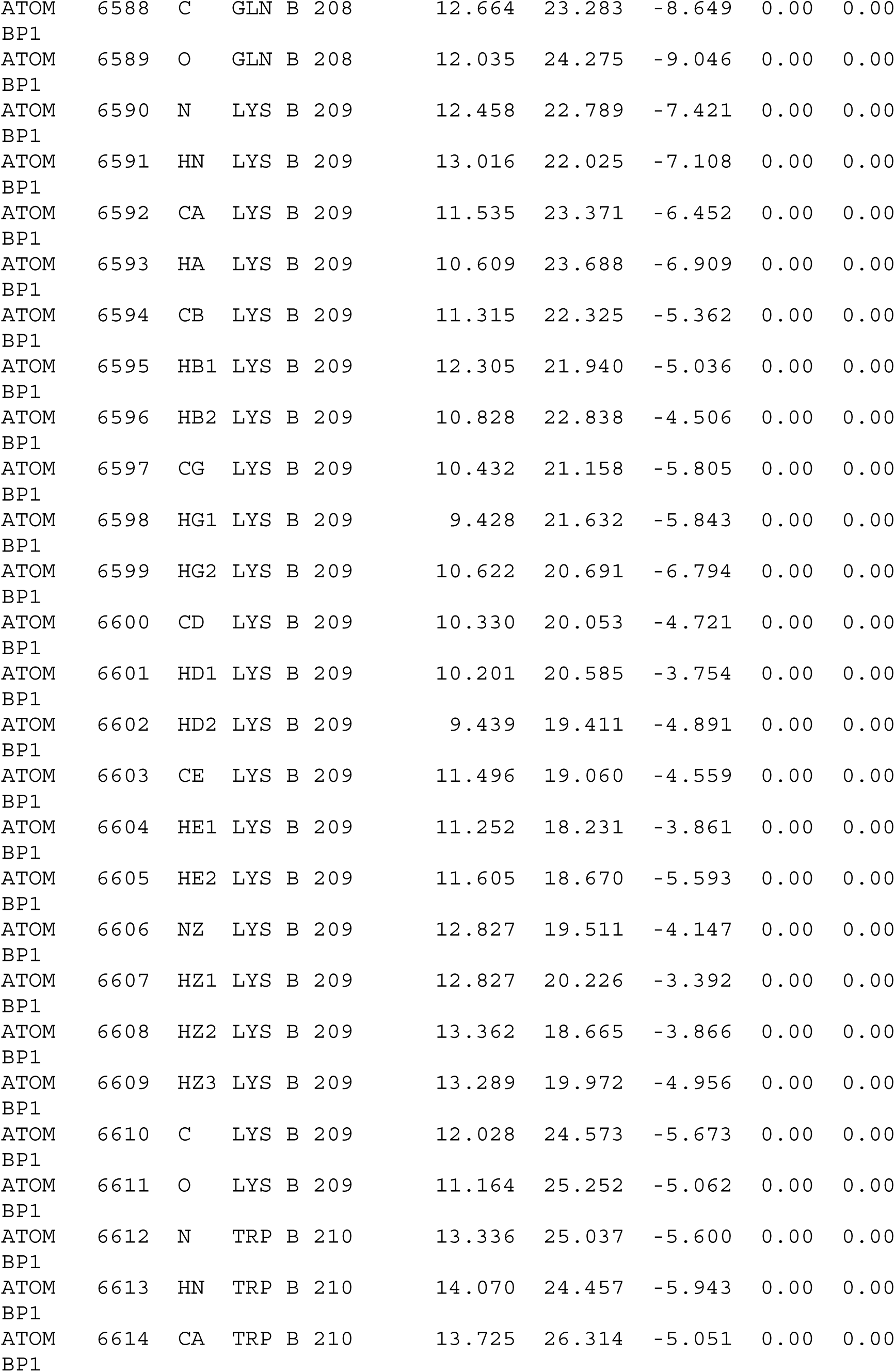

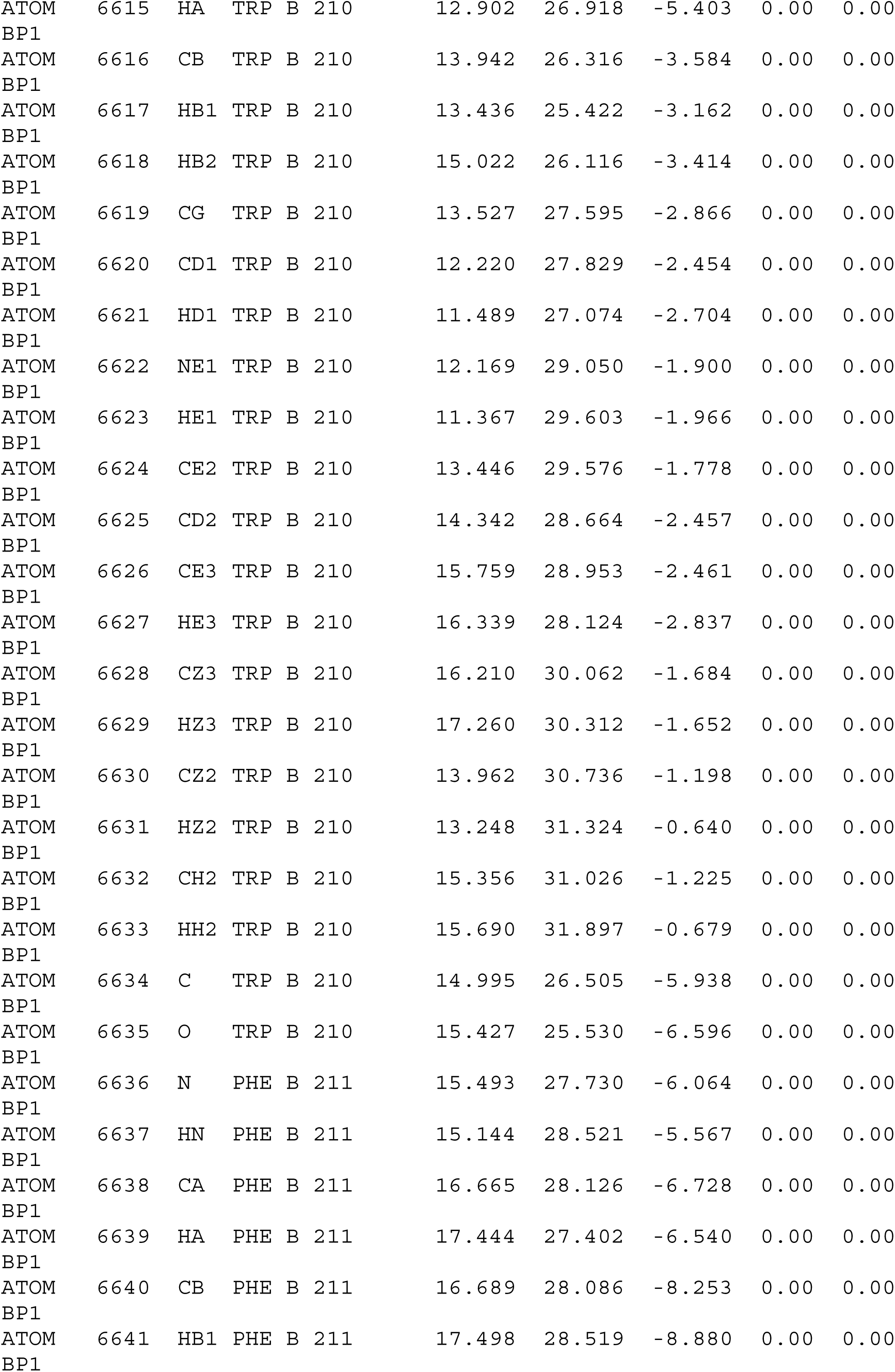

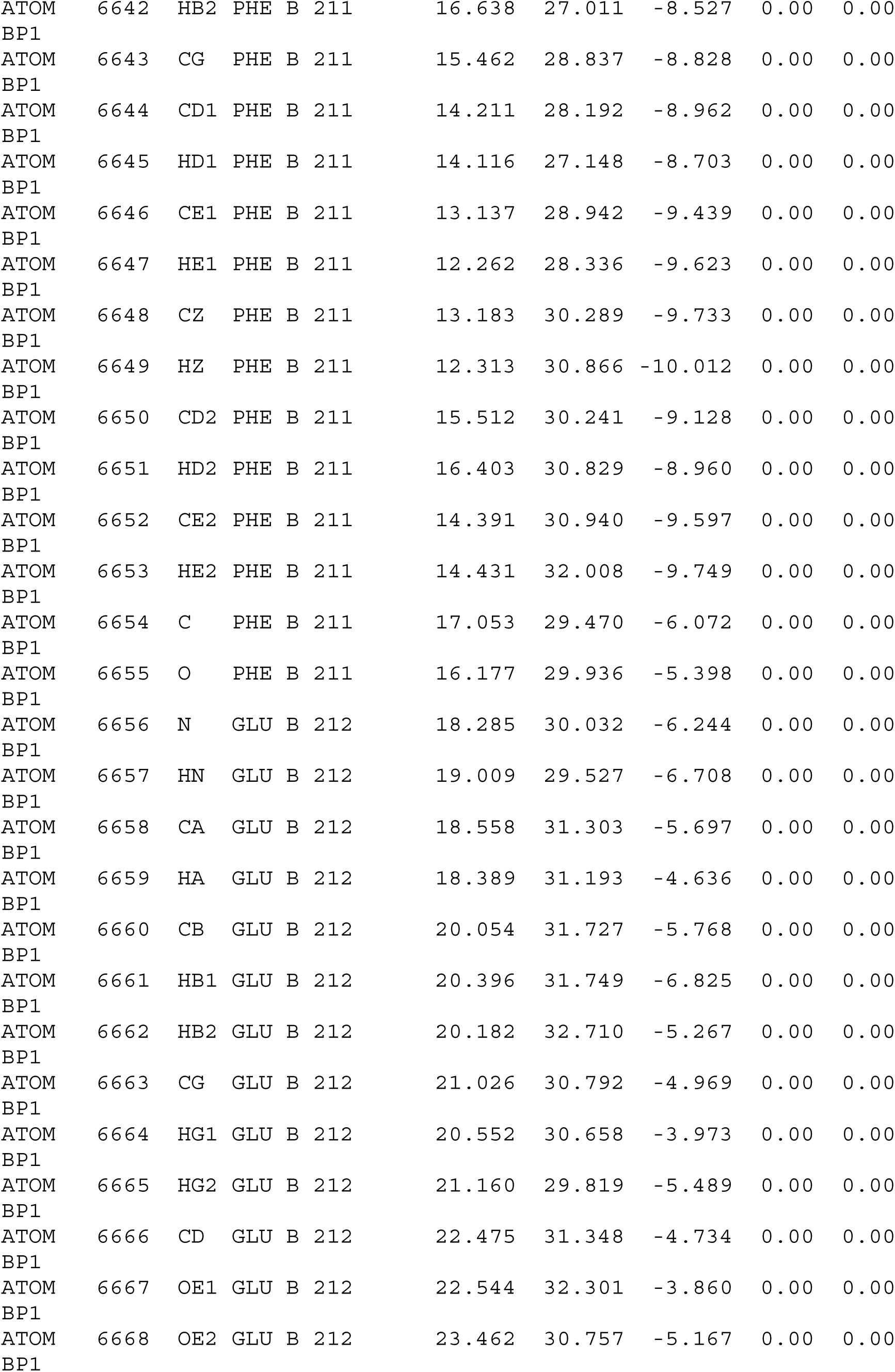

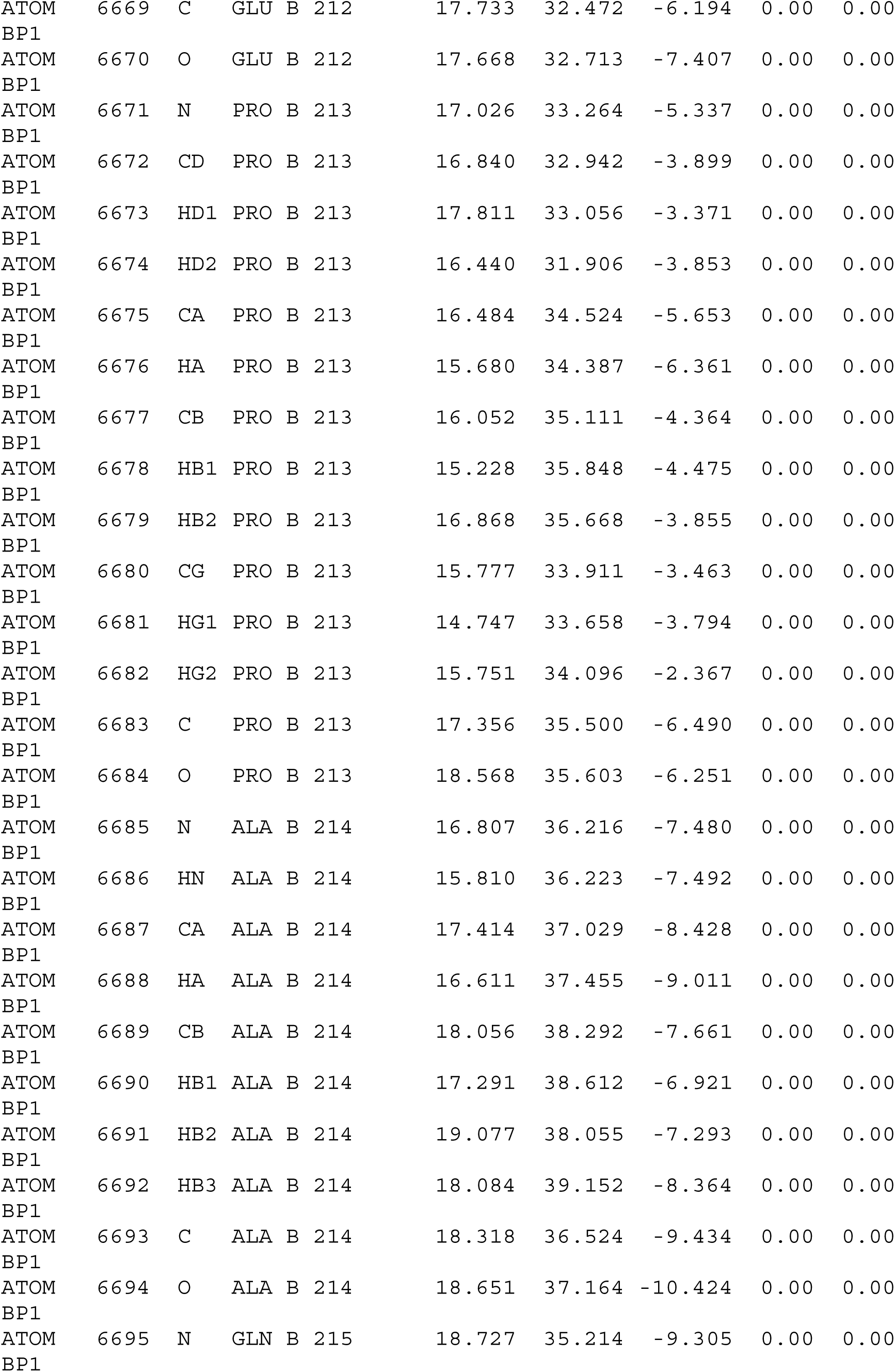

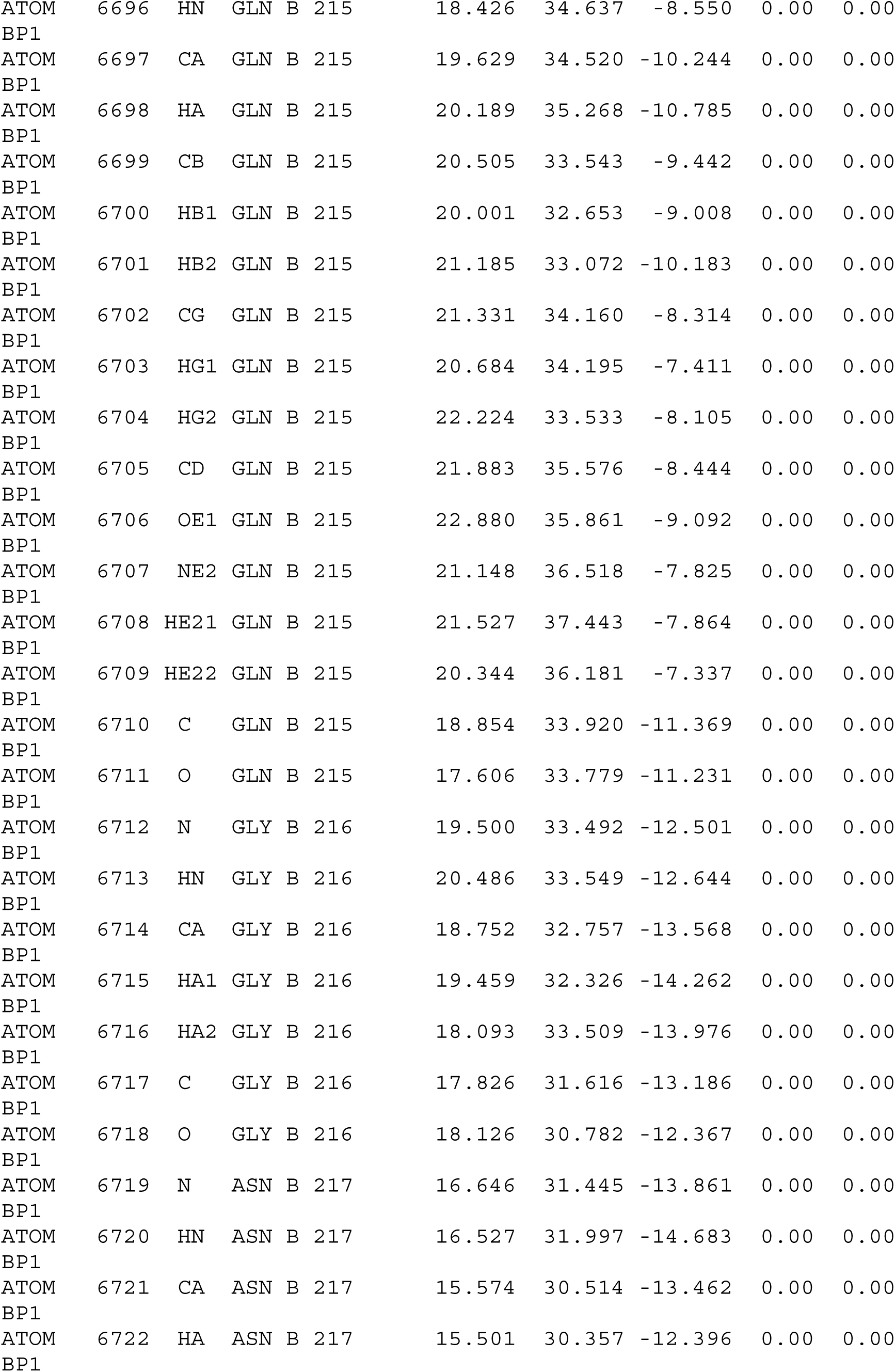

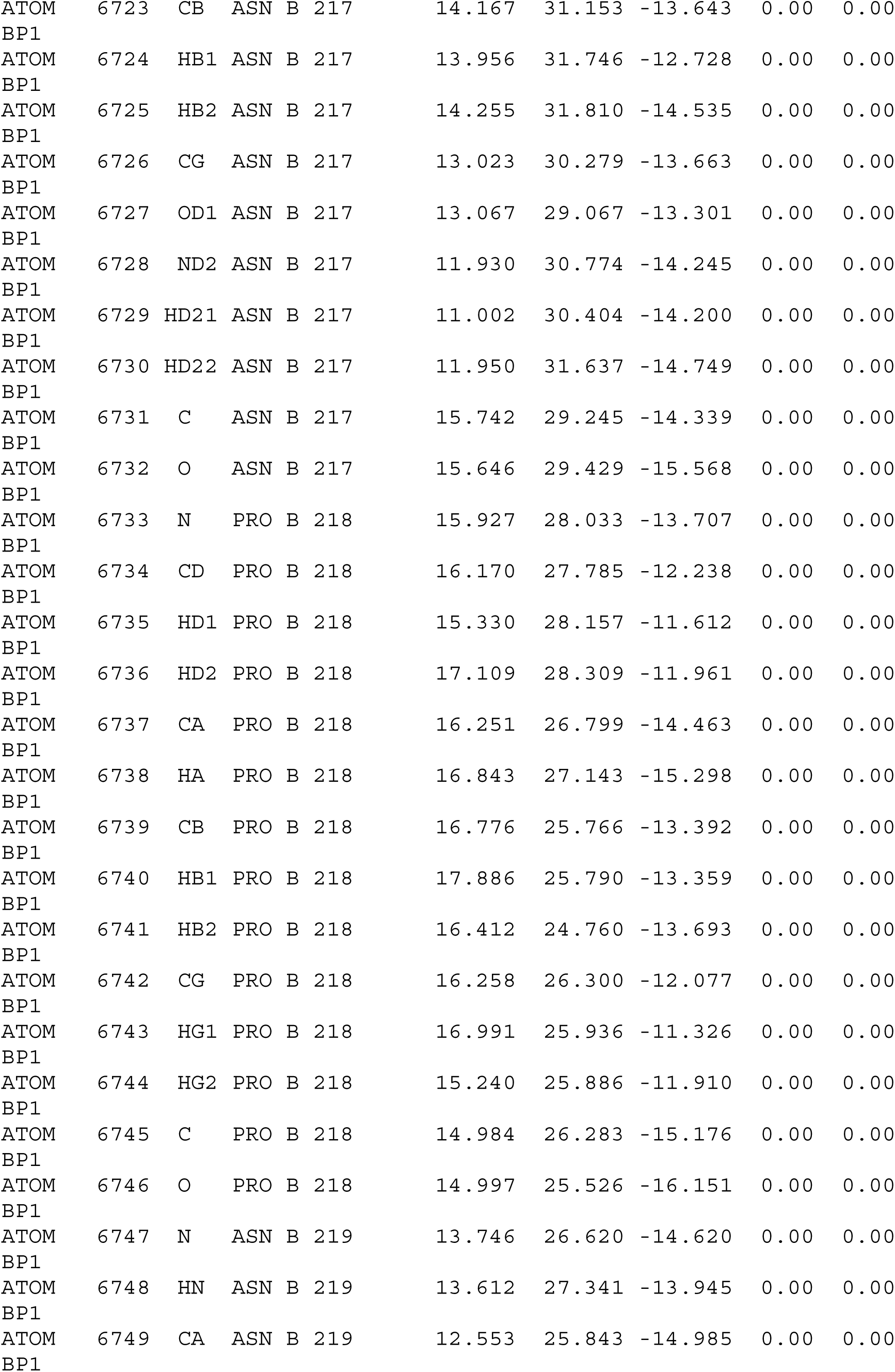

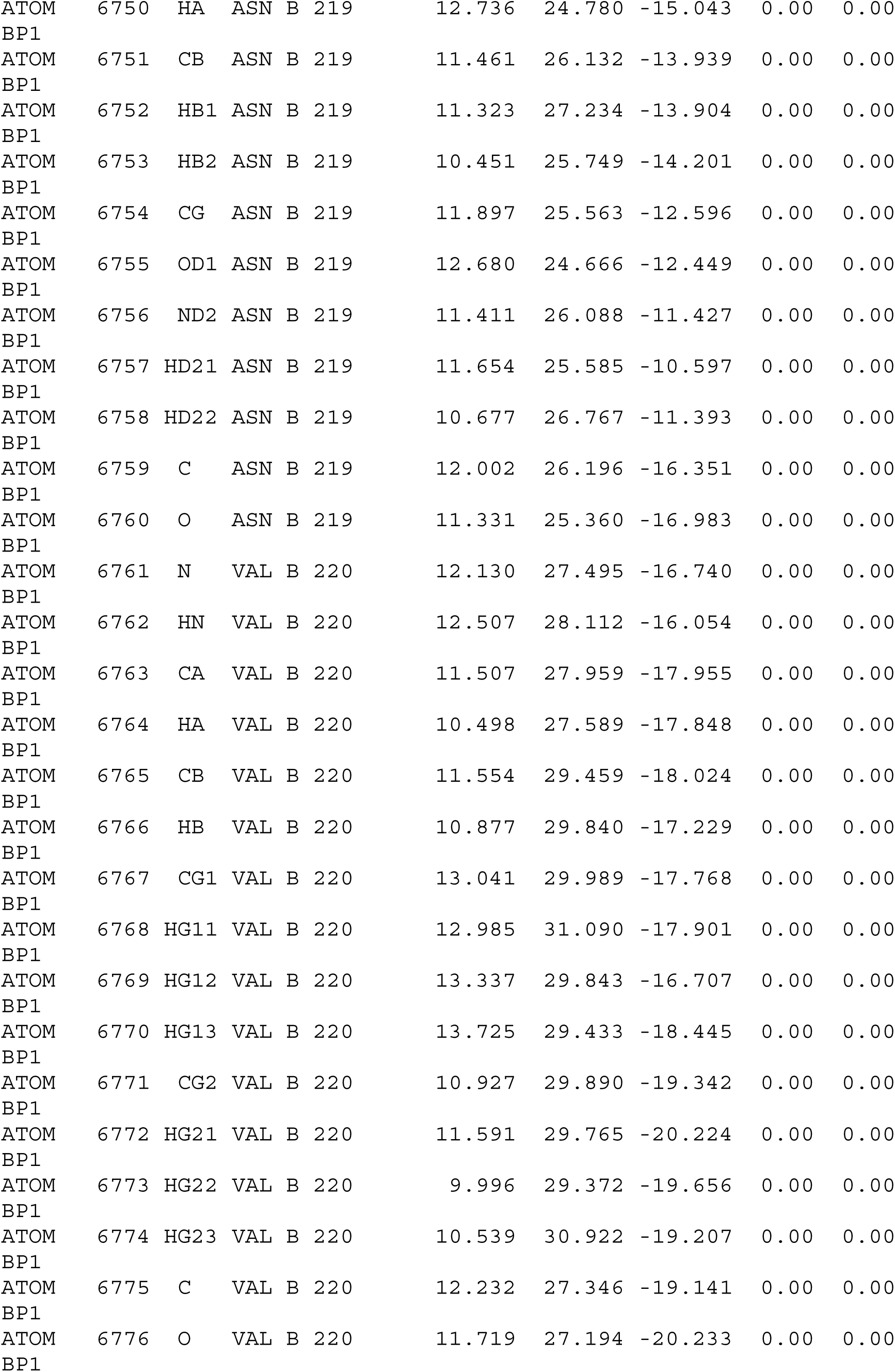

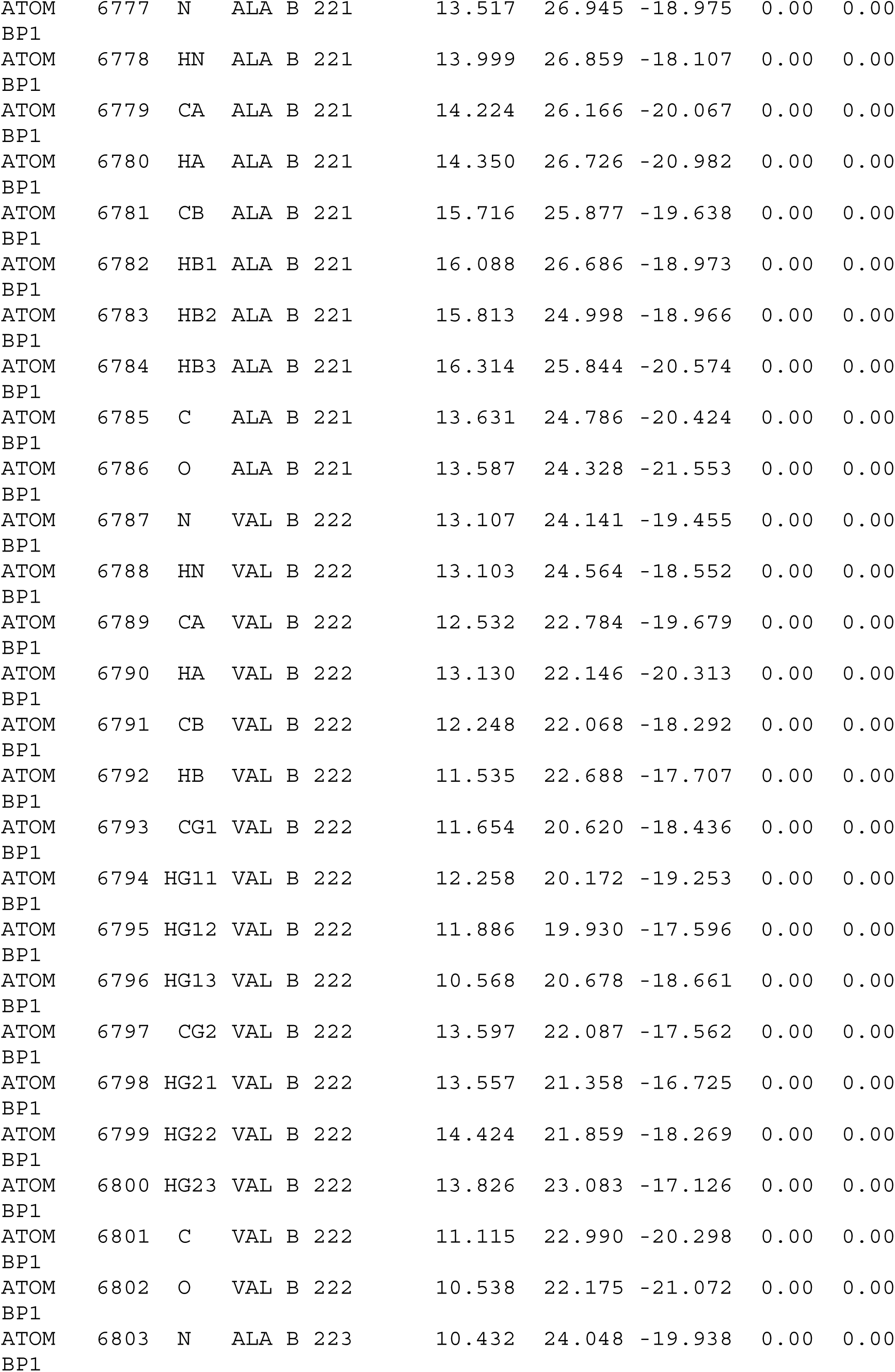

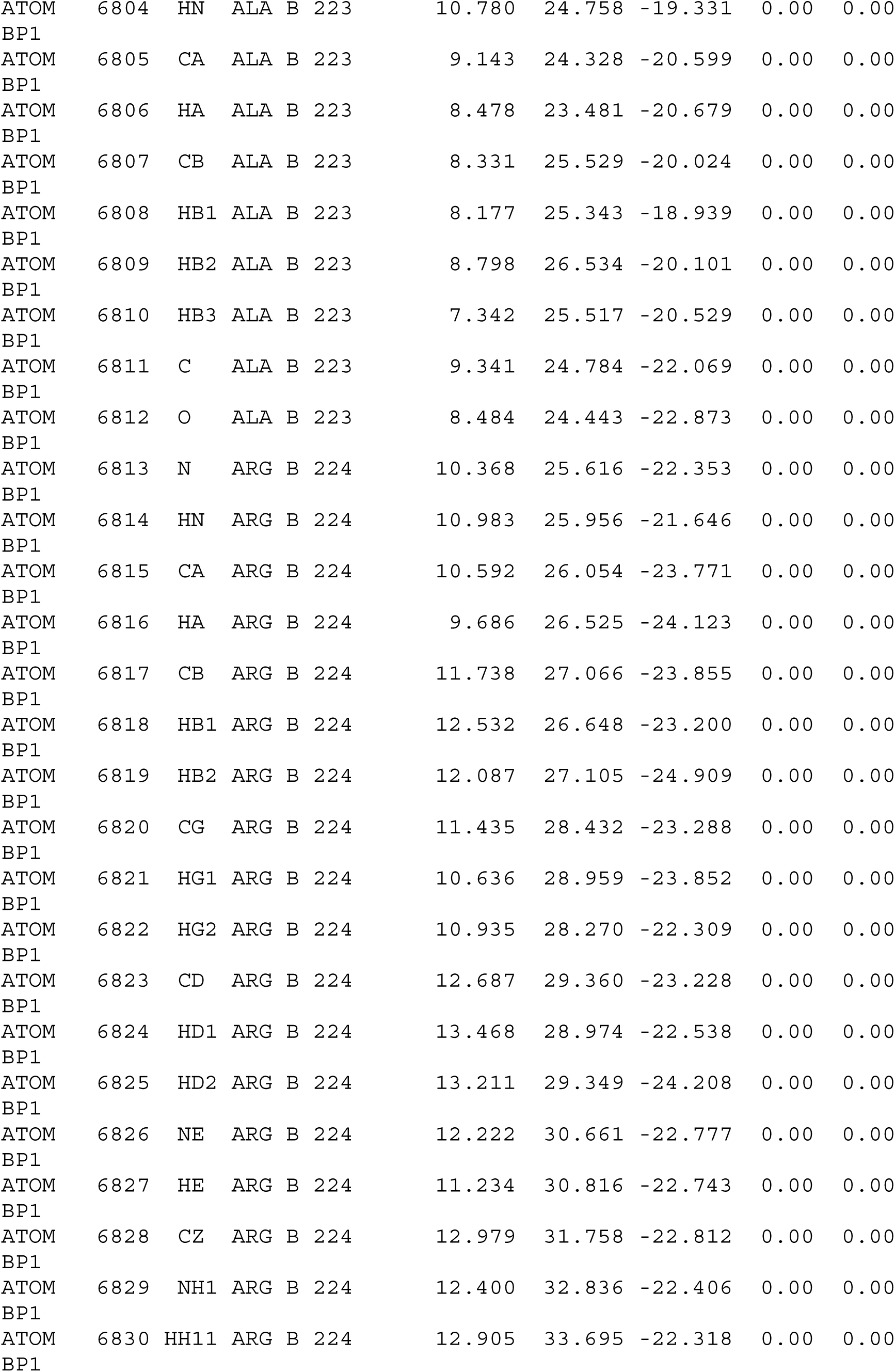

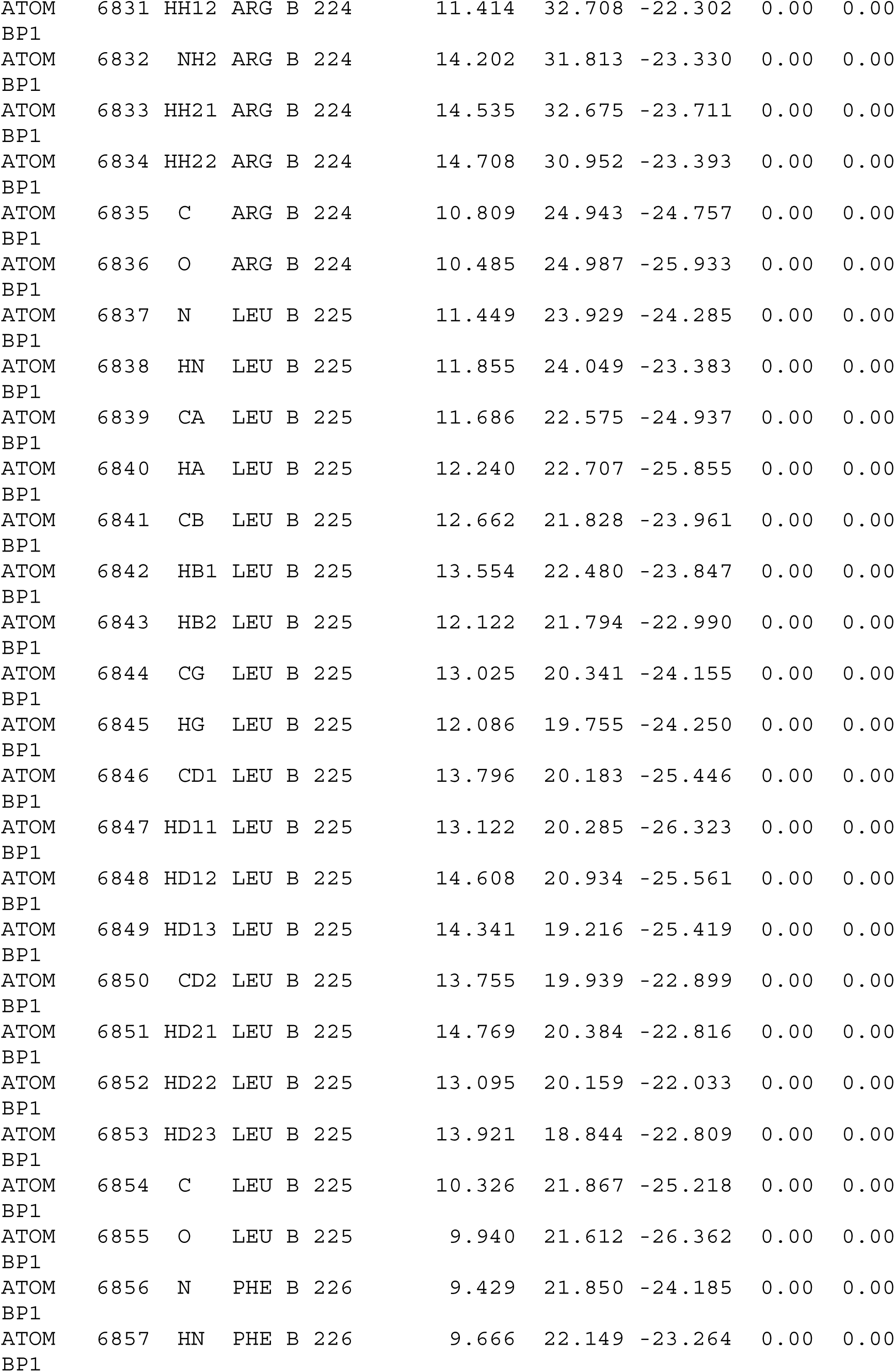

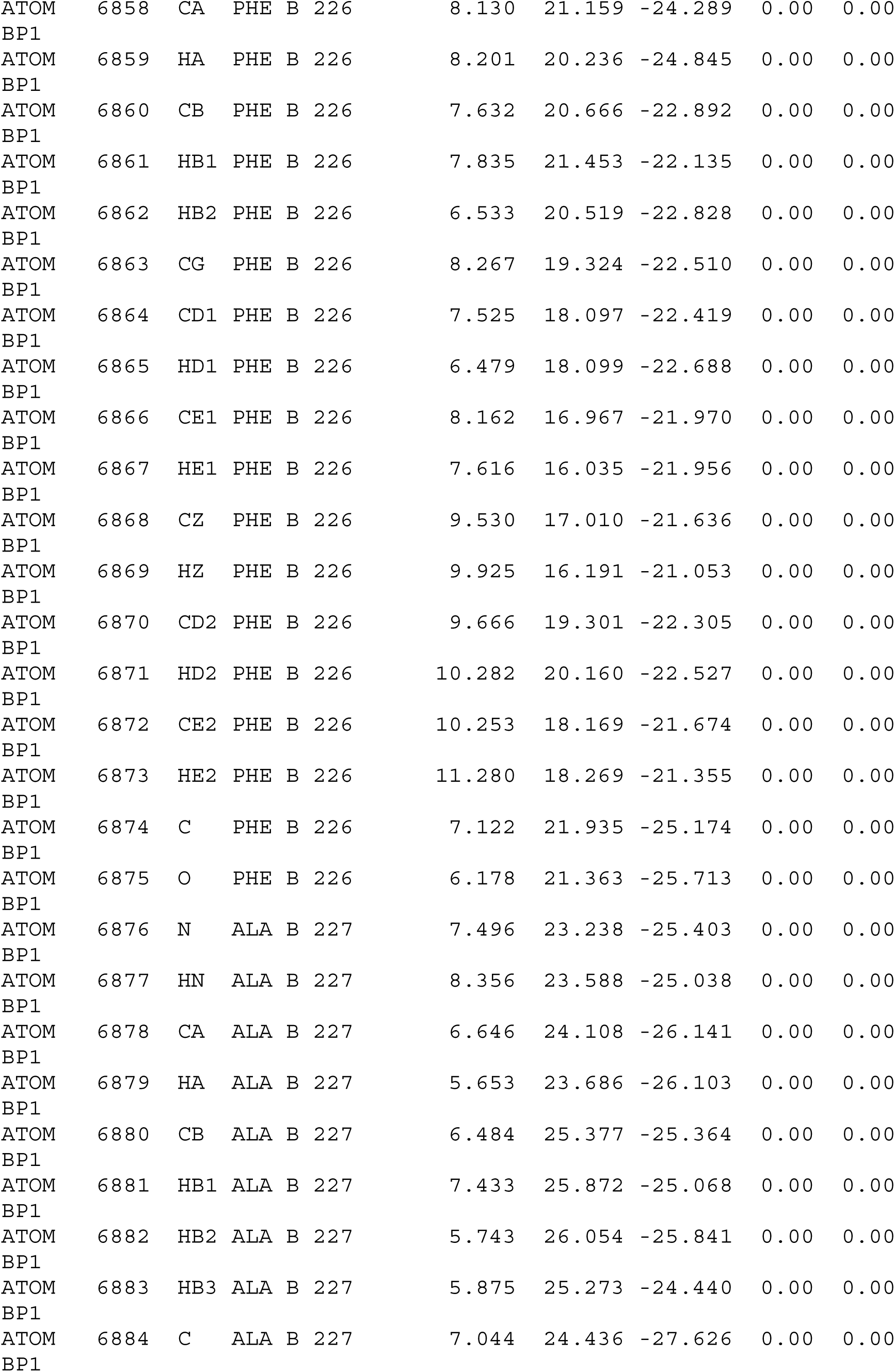

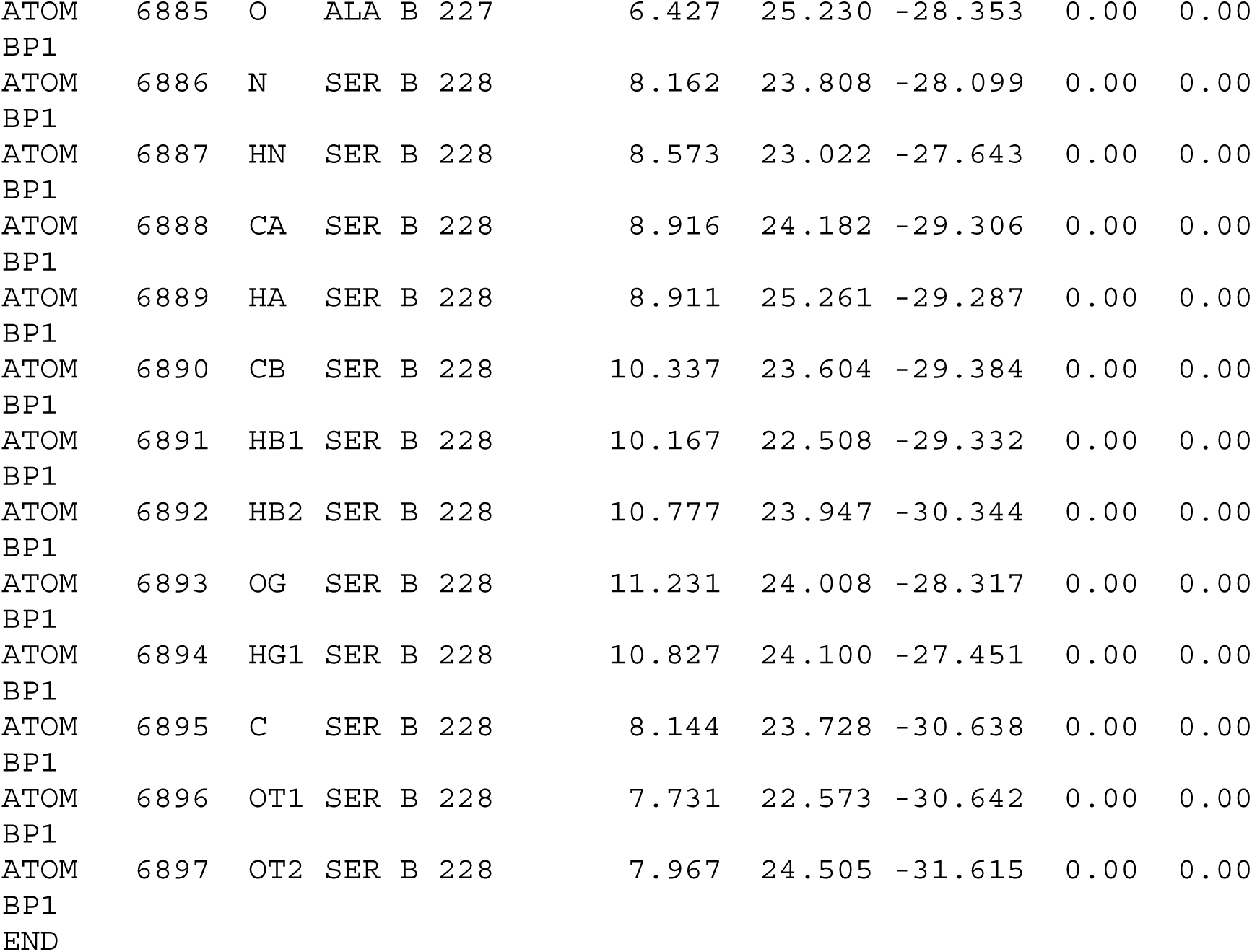

