## Supplementary Information for "Force-dependent Reorganization and Mechanostability of the Izumo1:Juno Complex Involved in Human Fertilization"

#### Materials and Methods

Construct design and site-directed mutagenesis

**Constructs for use in insect cell expression systems.** The ectodomains of Izumo1 and Juno were cloned with a C-terminal $His_{10}$ and SpyTag into pFastBac1 vectors for use in the Bac-to-Bac$^{TM}$ Baculovirus expression system (Thermo Fisher Scientific, Massachusetts, United States). To this end, wild-type Izumo1-$His_{10}$-SpyTag-LPETGG and Juno-$His_{10}$-SpyTag-LPETGG were synthesized as codon-optimized linear DNA fragments (GeneArt – ThermoFisher Scientific, Regensburg, Germany) for Gibson Assembly using primers designed in SnapGene software. Izumo1 and Juno point mutations were generated as required using the Q5® Site-Directed Mutagenesis Kit and NEBuilder v1 software for primer design (New England Biolabs, Massachusetts, United States).

**Constructs for use in bacterial expression systems.** For biophysical characterization, the SpyTag/SpyCatcher system facilitated the covalent attachment of the polyproteins required for AFM-SMFS and flow cytometry binding studies. All constructs containing polyproteins required for the AFM-SMFS and flow cytometry binding studies were cloned in pET28a vectors for expression in *E. coli* NiCo21(DE3) (New England Biolabs, Massachusetts, United States) using Gibson Assembly and primers designed in SnapGene software. The construct (N-to C-terminus) used for surface functionalization in the constant speed AFM-SMFS and flow cytometry binding studies was ybbR-$His_6$-ELP-$(MV7E2)_3$-ddFLN4-SpyCatcher (*47*). The construct (N-to C- terminus) used for surface functionalization in force clamp AFM-SMFS was ybbR-$His_6$-ELP-$(MV7E2)_3$-SpyCatcher. The construct (N-to-C-terminus) used for the fluorescent tagging in the flow cytometry binding studies was ybbR-HRV3C-$His_6$-GFP-SpyCatcher. In each instance, the terminal ybbR-tag allowed for the site-specific immobilization of target proteins to surfaces coated in coenzyme A using surfactin phosphopantetheinyl transferase (SFP) (*48*).

All constructs and point mutations were subsequently validated using Sanger sequencing (MicroSynth, Wein, Austria).

**Full protein construct sequences.** Where human rhinovirus (HRV) 3C Protease cleavage sites (LEVLFQGP) or sortase motifs (LPETGG) are present in the constructs, they were not used in this study. Further, the cysteine residue at position 18 in wild-type *Dictyostelium discoideum* 4$^{th}$ filamin domain (ddFLN4, UniProt: P13466, residues 549 - 649) has previously been mutated to C18S to avoid a potential cross-reaction to maleimide groups present from surface functionalization (*47*). For human Izumo1 constructs, an N-terminal Hemolin (HEM) secretion signal peptide was added. For human Juno constructs, an N-terminal Honeybee Melittin (HBM) secretion signal peptide was added. For mouse Izumo1 and Juno constructs, an N-terminal Binding Immunoglobulin Protein (BiP) secretion signal peptide was added. Secretion signal peptides are required to direct recombinant proteins in *Spodoptera frugiperda* (Sf9) based cell lines to the secretory pathways. HEM and HBM include sites for proteolytic cleavage such that they are not present in the mature proteins. For both Izumo1 and Juno point mutants, the numbering is based on the consensus sequence provided by UniProt (Izumo1: Q8IYV9, Juno: A6ND01).

HEM-Izumo1-His$_{10}$-SpyTag-LPETGG

MAFKSIAVLSACIIVGSACVICDPSVVLALKSLEKDYLPGHLDAKHHKAMMERVENAVKDFQELSLNEDAYM
GVVDEATLQKGSWSLLKDLKRITDSDVKGDLFVKELFWMLHLQKETFATYVARFQKEAYCPNKCGVMLQTLI
WCKNCKKEVHACRKSYDCGERNVEVPQMEDMILDCELNWHQASEGLTDYSFYRVWGNNTETLVSKGKEA
TLTKPMVGPEDAGSYRCELGSVNSSPATIINFHVTVLPHHHHHHHHHHAHIVMVDAYKPTKLPETGG

HEM-Izumo1E71A-His$_{10}$-SpyTag-LPETGG

MAFKSIAVLSACIIVGSACVICDPSVVLALKSLEKDYLPGHLDAKHHKAMMERVENAVKDFQELSLNADAYM
GVVDEATLQKGSWSLLKDLKRITDSDVKGDLFVKELFWMLHLQKETFATYVARFQKEAYCPNKCGVMLQTLI
WCKNCKKEVHACRKSYDCGERNVEVPQMEDMILDCELNWHQASEGLTDYSFYRVWGNNTETLVSKGKEA
TLTKPMVGPEDAGSYRCELGSVNSSPATIINFHVTVLPHHHHHHHHHHAHIVMVDAYKPTKLPETGG

HEM-Izumo1K150A-His$_{10}$-SpyTag-LPETGG

MAFKSIAVLSACIIVGSACVICDPSVVLALKSLEKDYLPGHLDAKHHKAMMERVENAVKDFQELSLNEDAYM
GVVDEATLQKGSWSLLKDLKRITDSDVKGDLFVKELFWMLHLQKETFATYVARFQKEAYCPNKCGVMLQTLI
WCANCKKEVHACRKSYDCGERNVEVPQMEDMILDCELNWHQASEGLTDYSFYRVWGNNTETLVSKGKEA
TLTKPMVGPEDAGSYRCELGSVNSSPATIINFHVTVLPHHHHHHHHHHAHIVMVDAYKPTKLPETGG

HEM-Izumo1H157A-His$_{10}$-SpyTag-LPETGG

MAFKSIAVLSACIIVGSACVICDPSVVLALKSLEKDYLPGHLDAKHHKAMMERVENAVKDFQELSLNEDAYM
GVVDEATLQKGSWSLLKDLKRITDSDVKGDLFVKELFWMLHLQKETFATYVARFQKEAYCPNKCGVMLQTLI
WCKNCKKEVAACRKSYDCGERNVEVPQMEDMILDCELNWHQASEGLTDYSFYRVWGNNTETLVSKGKEA
TLTKPMVGPEDAGSYRCELGSVNSSPATIINFHVTVLPHHHHHHHHHHAHIVMVDAYKPTKLPETGG

HEM-Izumo1R160A-His$_{10}$-SpyTag-LPETGG

MAFKSIAVLSACIIVGSACVICDPSVVLALKSLEKDYLPGHLDAKHHKAMMERVENAVKDFQELSLNEDAYM
GVVDEATLQKGSWSLLKDLKRITDSDVKGDLFVKELFWMLHLQKETFATYVARFQKEAYCPNKCGVMLQTLI
WCKNCKKEVHACAKSYDCGERNVEVPQMEDMILDCELNWHQASEGLTDYSFYRVWGNNTETLVSKGKEA
TLTKPMVGPEDAGSYRCELGSVNSSPATIINFHVTVLPHHHHHHHHHHAHIVMVDAYKPTKLPETGG

HBM-Juno-$His_{10}$-SpyTag-LPETGG

MKFLVNVALVFMVVYISYIYADGDELLNICMNAKHHKRVPSPEDKLYEECIPWKDNACCTLTTSWEAHLDVS
PLYNFSLFHCGLLMPGCRKHFIQAICFYECSPNLGPWIQPVGSLGWEVAPSGQGERVVNVPLCQEDCEEW
WEDCRMSYTCKSNWRGGWDWSQGKNRCPKGAQCLPFSHYFPTPADLCEKTWSNSFKASPERRNSGRCL
QKWFEPAQGNPNVAVARLFASHHHHHHHHHHAHIVMVDAYKPTKLPETGG

HBM-JunoH177Q-$His_{10}$-SpyTag-LPETGG

MKFLVNVALVFMVVYISYIYADGDELLNICMNAKHHKRVPSPEDKLYEECIPWKDNACCTLTTSWEAHLDVS
PLYNFSLFHCGLLMPGCRKHFIQAICFYECSPNLGPWIQPVGSLGWEVAPSGQGERVVNVPLCQEDCEEW
WEDCRMSYTCKSNWRGGWDWSQGKNRCPKGAQCLPFSQYFPTPADLCEKTWSNSFKASPERRNSGRCL
QKWFEPAQGNPNVAVARLFASHHHHHHHHHHAHIVMVDAYKPTKLPETGG

BiP-mIzumo1-$His_{10}$-SpyTag-LPETGG

MKLCILLAVVAFVGLSLGCVICDPSVVLALKSLEKDYLPGHLDAKHHKAMMERVENAVKDFQELSLNEDAYM
GVVDEATLQKGSWSLLKDLKRITDSDVKGDLFVKELFWMLHLQKETFATYVARFQKEAYCPNKCGVMLQTLI
WCKNCKKEVHACRKSYDCGERNVEVPQMEDMILDCELNWHQASEGLTDYSFYRVWGNNTETLVSKGKEA
TLTKPMVGPEDAGSYRCELGSVNSSPATIINFHVTVLPGSGSGSGSHHHHHHHHHHGSGGLLDAHIVMVDA
YKPTKLPETGG

BiP-mJuno-$His_{10}$-SpyTag-LPETGG

MKLCILLAVVAFVGLSLGGDELLNICMNAKHHKRVPSPEDKLYEECIPWKDNACCTLTTSWEAHLDVSPLYNF
SLFHCGLLMPGCRKHFIQAICFYECSPNLGPWIQPVGSLGWEVAPSGQGERVVNVPLCQEDCEEWWEDCR
MSYTCKSNWRGGWDWSQGKNRCPKGAQCLPFSHYFPTPADLCEKTWSNSFKASPERRNSGRCLQKWFEP
AQGNPNVAVARLFASGSGSGSGSHHHHHHHHHHGSGGLLDAHIVMVDAYKPTKLPETGG

ybbR-$His_6$-ELP-$(MV7E2)_3$-ddFLN4-SpyCatcher

MGTDSLEFIASKLAHHHHHHWGSGHGVGVPGMGVPGVGVPGVGVPGVGVPGVGVPGVGVPGVGVPGV
GVPGEGVPGEGVPGVGVPGMGVPGVGVPGVGVPGVGVPGVGVPGVGVPGVGVPGVGVPGEGVPGEG
VPGVGVPGMGVPGVGVPGVGVPGVGVPGVGVPGVGVPGVGVPGVGVPGEGVPGEGVPGWPSGSADP
EKSYAEGPGLDGGECFQPSKFKIHAVDPDGVHRTDGGDGFVVTIEGPAPVDPVMVDNGDGTYDVEFEPKE
AGDYVINLTLDGDNVNGFPKTVTVKPAPGSGSGSGSVDTLSGLSSEQGQSGDMTIEEDSATHIKFSKRDEDG
KELAGATMELRDSSGKTISTWISDGQVKDFYLYPGKYTFVETAAPDGYEVATAITFTVNEQGQVTVNGKATK
GDAHI

ybbR-His$_6$-ELP-(MV7E2)$_3$-SpyCatcher

MGTDSLEFIASKLAHHHHHHWGSGHGVGVPGMGVPGVGVPGVGVPGVGVPGVGVPGVGVPGVGVPGV
GVPGEGVPGEGVPGVGVPGMGVPGVGVPGVGVPGVGVPGVGVPGVGVPGVGVPGVGVPGEGVPGEG
VPGVGVPGMGVPGVGVPGVGVPGVGVPGVGVPGVGVPGVGVPGVGVPGEGVPGEGVPGWPSGSVDTL
SGLSSEQGQSGDMTIEEDSATHIKFSKRDEDGKELAGATMELRDSSGKTISTWISDGQVKDFYLYPGKYTFVE
TAAPDGYEVATAITFTVNEQGQVTVNGKATKGDAHI

ybbR-HRV3C-His$_6$-GFP-SpyCatcher
MGTDSLEFIASKLALEVLFQGPLQHHHHHHPWTSASSGGEELFAGIVPVLIELDGDVHGHKFSVRGEGEGDA
DYGKLEIKFICTTGKLPVPWPTLVTTLCYGIQCFARYPEHMKMNDFFKSAMPEGYIQERTIQFQDDGKYKTRG
EVKFEGDTLVNRIELKGKDFKEDGNILGHKLEYSFNSHNVYIRPDKANNGLEANFKTRHNIEGGGVQLADHY
QTNVPLGDGPVLIPINHYLSTQTKISKDRNEARDHMVLLESFSACCHTHGMDELYRGSGSGSGSVDTLSGLSS
EQGQSGDMTIEEDSATHIKFSKRDEDGKELAGATMELRDSSGKTISTWISDGQVKDFYLYPGKYTFVETAAPD
GYEVATAITFTVNEQGQVTVNGKATKGDAHI

Protein expression and purification

**Bacterial protein expression and purification.** All pET28a constructs were transformed into *E. coli* NiCo21(DE3) for subsequent expression and purification. Cultures were grown in terrific broth media (TB, 12 g/L tryptone, 24 g/L yeast extract, 4 mL/L glycerol, 170 mM M KH2PO4, 720 mM K2HPO4, pH 7.0) at 37 °C with shaking until the O.D.600 of culture reached 0.6-0.7 where they were induced using 1 mM isopropyl β-d-1-thiogalactopyranoside (IPTG) at 16 °C, with shaking. After an induction period of 16 h, cultures were harvested using centrifugation and stored at -80 °C until needed. Cell pellets were resuspended at 4 °C in lysis buffer (50 mM Tris-HCl, 50 mM NaCl, 5 mM $MgCl_2$, 0.1% (v/v) TritonX-100, Glycerol, pH 8.0) for subsequent sonication (Digital Sonifier 450, Branson Ultrasonics Corp, Connecticut, United States). The cell lysate then underwent centrifugation at 30,000 ×g for 30 min and was passed through a 0.45 µM filter. The clarified lysate was then incubated with Ni-NTA Resin (HisPur™, Thermo Fischer Scientific, Massachusetts, United States) for 1 h at 4 °C with shaking. Ni-NTA Resin was reconstituted by centrifugation 800 ×g for 5 mins and transferred to gravity flow columns (Pierce™, Thermo Fischer Scientific, Massachusetts, United States). The resin was then washed extensively in 10 mM imidazole-supplemented phosphate-buffered saline (PBS, 137 mM NaCl, 2.7 mM KCl, 10 mM $Na_2HPO_4$, and 1.8 mM $KH_2PO_4$, pH 7.4) and eluted in 250 mM imidazole supplemented PBS. Protein-containing elution fractions were concentrated in centrifugal filters (Amicon®, Merck, Darmstadt, Germany) and further processed using size-exclusion chromatography (SEC, Superdex 200 Increase 10/300 GL column, Cytiva, Marlborough, United States) into PBS. SEC elution fractions containing the protein of interest were again concentrated using centrifugal filters and stored in glycerol 33% (v/v) at -20 °C until used in experiments. Protein concentrations were measured by spectrophotometry at 280 nm (NanoDrop 1000,

Thermo Scientific, DE, USA). For each construct, final concentrations typically ranged from 100 - 500 µM.

**Baculovirus generation.** The Izumo1 and Juno genes were transposed into bacmids by the transformation of pFastBac1 parent vectors into *E. coli* DH10Bac cells and plating onto 5-Bromo-4-chloro-3-indolyl β-D-galactopyranoside (X-gal) selection plates (Luria-Bertani agar with 40 µg/mL X-gal, 0.5 mM IPTG, 30 µg/mL kanamycin, 30 µg/mL tetracycline, 30 µg/mL gentamicin, pH 7.4). Bacmids were isolated from white colonies using a generic isopropanol extraction protocol for DNA. To confirm the identity of each bacmid, the transposed regions underwent PCR amplification using standard M13 sequencing primers followed by sanger sequencing. GenJet™ (Thermo Fisher Scientific, Massachusetts, United States) DNA In Vitro Transfection Reagent were then used to transfect ExpiSF™ Sf9 cells (Thermo Fischer Scientific, Massachusetts, United States) with the purified bacmids as per the manufacturer's instructions. P0 virus stocks were harvested by centrifugation and passing of the supernatant through a 0.45 µM filter. P1 and P2 virus stocks were generated by infecting ExpiSF™ cells $1\times10^6$ cells/mL (Countess™ 3 automated cell counter, Thermo Fischer Scientific, Massachusetts, United States) with the previous generation of viral stock for 48 h at 27 °C after which the viral stocks were harvested as described above. Finally, the identity of recombinantly expressed Izumo1 and Juno in P2 viral stocks was verified using the C-terminal SpyTags, as described previously (*49*).

**Insect cell protein expression and purification.** ExpiSF™ cells were cultured in ExpiSF™ chemically defined medium (Thermo Fischer Scientific, Massachusetts, United States) at 27 °C with shaking. As directed by the manufacturer, cultures were seeded in fresh medium at a concentration of $5\times10^6$ cells/mL and mixed with ExpiSf™ Enhancer (Thermo Fischer Scientific, Massachusetts, United States) 18 h before transfection. Transfection with P2-derived viral stocks was monitored via GFP fluorescence and increased cell size 24 h post-infection. Successfully transfected cultures were then incubated at 27 °C, with shaking, for a further 48-72 h or until cell viability dropped below 70%. ExpiSF™ cells were removed via centrifugation and the culture medium passed through a 0.45 µM filter. Fastback Ni Advance Resin (Protein Ark, Cambridge, United Kingdom) was added to the clarified media and recombinant proteins were eluted into tris-buffered saline with glycerol (TBS-G, 20 mM Tris-HCl, 300 mM NaCl, and 10% glycerol (w/v), pH 8.0), supplemented with 0.1% (w/v) tween, using imidazole gradients across gravity-flow columns. Fractions containing the recombinant proteins were further purified using SEC into TBS-G (20 mM Tris-HCl, 300 mM NaCl, and 10% glycerol (w/v), pH 8.0) storage buffer and concentrated using centrifugal concentrators. Concentrated proteins were processed with detergent removal spin columns (Pierce®, Thermo Fischer Scientific, Massachusetts, United States) and stored at -20 °C. Directly prior to use in experiments, Izumo1-$His_{10}$-SpyTag-LPETGG and Juno-$His_{10}$-SpyTag-LPETGG aliquots were incubated over night at 4 °C with equimolar amounts of ybbR-$His_6$-ELP-$(MV7E2)_3$-ddFLN4-SpyCatcher or ybbR-$His_6$-ELP-$(MV7E2)_3$-SpyCatcher as required. Protein concentrations were measured by spectrophotometry at 280 nm. For each construct, final concentrations typically ranged from 30-150 µM.

Atomic force microscopy single-molecule force spectroscopy

**AFM sample preparation.** Site-specific immobilization of samples for atomic force microscopy-based single molecule force spectroscopy (AFM-SMFS) was performed as described previously (*50*, *51*). In brief, AFM cantilevers (Biolever Mini AC40TS, Olympus, Tokyo, Japan) were treated with an ultraviolet ozone cleaner (Novascan, Iowa, United States) for 40 minutes and 25 mm diameter round cover glass surfaces (Menzel Gläser, Braunschweig, Germany) were incubated in piranha solution (1:1 (v/v) 30% $H_2O_2$: concentrated $H_2SO_4$) for 30 minutes. Irradiated cantilevers and piranha etched cover glasses were then incubated with 50% (v/v) (3-Aminopropyl)dimethylethoxysilane (ABCR, Karlsruhe, Germany), diluted in isopropanol, with water then added to 0.5 % (v/v), for 5 minutes and 30 minutes, respectively. Silanated cantilevers were then washed successively in toluene, isopropanol, and water. Silanated cover glasses were thoroughly washed in water three times. The silanized cantilevers and cover glasses were cured at 60 °C for 40 minutes. Next, cantilevers and cover glasses were incubated in a solution containing 20 mg/mL of sulfosuccinimidyl 4-(N-maleimidomethyl)cyclohexane-1-carboxylate (Thermo Fisher Scientific, Massachusetts, United States) in 4-(2-Hydroxyethyl)piperazine-1-ethanesulfonic acid buffer (HEPES, 50 mM HEPES, 150 mM NaCl, pH 7.5) for 1 hour. The exposed maleimide groups on the cantilevers and cover glasses were reacted with 200 µM coenzyme A in in Mal-Coupling buffer (50mM $Na_2HPO_4$, 50mM NaCl, 10mM $C_10H_16N_2O_8$, pH 7.2) for 2 h at room temperature. In the final modification step, the ybbR-tags located at the C-terminus of the target proteins were incubated with cantilevers and cover glasses with 10 mM $MgCl_2$ and 1.5 µM SFP in PBS buffer for 4 h at room temperature. Between each modification step, following silanization and prior to the functionalization of proteins, cantilevers and cover glasses were washed three times with water. Following the functionalization of proteins, the cantilevers and cover glasses were thoroughly washed with PBS (pH 7.4).

**AFM-SMFS measurements, constant speed.** Constant speed SMFS measurements were performed on a ForceRobot® AFM (JPK instruments, Berlin, Germany). The contact-free method was used to calibrate cantilever spring constants, ranging from 0.06 to 0.14 $N \cdot m^{-1}$. The cantilever was brought into contact with the surface at a force setpoint of 0.18 nN for a period of 0.2 s and subsequently retracted at constant speeds of 800, 1600, 3200, and 6400 $nm \cdot s^{-1}$. Between measurements, the glass surface was moved horizontally by 100 nm. In the absence of surface-cantilever interactions over an interval of 24 approach attempts, the glass surface was moved 1000 nm vertically. Typically, a 10-15 h experimental run yielded 15,000-20,000 measurements. PBS, adjusted to a pH of 7.4, was utilized as the buffer solution for all AFM-SMFS measurements.

**Constant Speed SMFS data analysis and curve selection.** AFM data was analyzed using a combination of Python, MATLAB (MathWorks), and R scripts.

Analysis and selection of single molecule events followed transformation of baseline subtracted force *vs.* extension curves into contour length space. This transformation was performed by applying the assumptions of the freely rotating chain (FRC) model to each force *vs.* extension curve. In the FRC model, rigid freely rotating segments of length *b* are assumed to be connected by a fixed angle, *ϒ*. Here, the implementation given by Eq. (1) (*52*) was used to calculate contour length (*L*) as a function of force (*F*) and extension (*x*), giving force-contour length curves.

$$L = \begin{cases} \frac{3k_BT}{xFa} & \text{for } \frac{Fb}{k_BT} < \frac{l}{p} \\ \frac{x}{1-\left(\frac{4Fl}{xk_BT}\right)^{-0.5}} & \text{for } \frac{b}{l} < \frac{Fb}{k_BT} < \frac{l}{b} \\ \frac{x}{1-\left(\frac{2Fb}{k_BT}\right)^{-1}} & \text{for } \frac{Fb}{k_BT} < \frac{l}{b} \end{cases} \quad (1)$$

Where $k_B$ is the Boltzman constant, *T* is the temperature, *p* is persistence length, *b* is given by Eq. (2), and *l* is given by Eq. (3)

$$b = a\left(\frac{(1-\cos\gamma)\cos\frac{\gamma}{2}}{1+\cos\gamma}\right) \quad (2)$$

$$l = b\frac{\cos\frac{\gamma}{2}}{|\ln(\cos\gamma)|} \quad (3)$$

Following the transformation of force *vs.* extension curves into force *vs.* contour length curves, automated filtering and manual processing yielded ~70-150 force *vs.* contour length curves per pulling speed, per independent experiment. The automated filter was set to exclude curves showing forces and contour length increments indicative of non-specific cantilever-surface interactions, the absence of any cantilever-surface interaction, or the formation of multiple Izumo1:Juno complexes. Specifically, the automated filter excluded any force-extension curves that had sawtooth peaks within the first ~100 nm of extension, where no sawtooth peaks were observed, or more than 9 sawtooth peaks were observed. From the filtered dataset, single molecule interactions were then manually identified using two different selection criteria. Under the first selection criteria, single molecule interactions between Izumo1 and Juno were identified as individual rupture events < 100 pN at contour lengths of ~135 nm. This criterion was based on FRC-fitting of the expected contour length the polypeptide chains for this force range and the expectation that above these forces the ddFLN4 fingerprint domains would typically unfold. The remaining curves were then sorted for individual rupture events that followed the unfolding of two ddFLN4 fingerprint domains. The unfolding of each ddFLN4 domain contributed an additional 32 nm in contour length, producing final contour lengths of ~190 nm at rupture. The data sets were then further sorted based on the presence or absence of unfolding intermediates that could contribute on average an additional ~15-20 nm of contour length.

Next, rupture force distributions were fitted as gaussian mixed models (GMM) using the mclust package in R (*53*). This approach allowed for the systematic statistical classification of the majority of unbinding events as either P0, P1, or P2. Here, the fitting parameters were set to allow for the best possible fit of the pooled rupture force distributions from each independent experiment, using one, two, three, or four gaussians. To improve the robustness of the fitting approach across all of the tested variants, a square-root transformation was applied the pooled rupture force distributions, which were then normalized prior to GMM fitting. In instances where a fourth gaussian was detected in the GMM, they were limited to a single constant pulling speed for a non-wild type mutant and were largely overlapping with another Gaussian. Therefore, we could not attribute them to a distinct unbinding pathway. Instead they were attributed to minor experimental variance. Accordingly, where a fourth gaussian was fitted in as part of the GMM, these rupture events were reclassified into the overlapping parent gaussians. Nonetheless, we would like to highlight the possibility of future improvements to the resolution of AFM-SMFS leading to the isolation of additional unbinding pathways in wild-type Izumo1:Juno. Such pathways would, ofcourse, need to be accounted for in the fitting of future distributions.

To explore the theoretical energy landscape of Izumo1:Juno, the force-independent off-rate ($k_0$), the distance to the transition state along the reaction coordinate (Eq. (6-8) given as $\Delta x^‡$), and the force-independent free energy of activation ($\Delta G^‡$) for P0, P1 and P2 were first determined using the Dudko-Hummer-Szabo (DHS) model (*38*, *39*). For this purpose, the rupture force distributions of each pathway and each constant pulling speed were plotted as histograms with equal bin widths ($\Delta F$ = 10 pN). After converting the rupture force histograms into binned rupture force probability distributions, the force-dependent off-rates were calculated using the histogram transformation method outlined in Eq. (4).

$$k_{\text{off}}(F_k) = \frac{h_k r(F_k)}{\left(\frac{h_k}{2} + \sum_{i=k+1}^{N} h_i\right)\Delta F} \tag{4}$$

Where $k_{\text{off}}(F_k)$ is the force-dependent off-rate at the median rupture force of the k$^{\text{th}}$ bin, $r(F_k)$ is the median loading rate of the $k^{\text{th}}$ bin, and $h_k$ is the height of the $k^{\text{th}}$ bin as calculated using Eq. (5).

$$h_k = \frac{C_k}{C_{\text{tot}}\Delta F} \tag{5}$$

Where $C_k$ is the number of counts in the $k^{\text{th}}$ bin and $C_{\text{tot}}$ is the total number of counts in the histogram.

The resulting $k_{\text{off}}(F_k)$ values were plotted against the corresponding median rupture force of the bin and fitted using the DHS-model, outlined in Eq. (6). To ensure robustness against local minima, we employed a Monte Carlo-based numerical optimization approach for fitting the DHS-model. First, plausible ranges for each parameter, based on prior empirical evidence and theoretical considerations, were specified. For all Izumo1:Juno variants, the ranges were as follows: $k_0$ within [0.001, 20], $\Delta x^{\ddagger}$ within [0.001, 1.5], and $\Delta G^{\ddagger}$ within [0.01, 20]. For each iteration of the model fitting, initial parameter values for $k_0$, $\Delta x^{\ddagger}$, and $\Delta G^{\ddagger}$ were randomly sampled from uniform distributions defined by the above-mentioned parameter ranges. Each set of initial parameter values underwent optimization to minimize the residual sum of squares between the experimental data and the model predictions using the Levenberg-Marquardt nonlinear least squares algorithm available in base R ($n$ = 1000). This fitting process was repeated for both the cusp-like ($\upsilon$ = 1/2) and cubic linear ($\upsilon$ = 1/3) DHS-parameters that describe the shape of the energy barrier. To ensure robustness, the optimization process was subjected to multiple runs ($n$ = 3) with different random seeds, and the consistency of the results was verified.

$$k_{\text{off}}(F) = k_0 \left(1 - \frac{vF\Delta x^{\ddagger}}{\Delta G^{\ddagger}}\right)^{\frac{1}{v}-1} e^{\beta \Delta G^{\ddagger}\left[1-\left(1-\frac{vF\Delta x^{\ddagger}}{\Delta G^{\ddagger}}\right)^{\frac{1}{v}}\right]} \tag{6}$$

Where $\beta^{-1}=k_BT$, $k_B$ is the Boltzmann constant, and $T$ is the temperature. In addition, the shape of the energy barrier can be parameterized by choose between either a cusp-like or a cubic linear shape during DHS-fitting.

To then apply the DHS-fitted parameters to the fitting of the force *vs.* (ln) loading rate graphs, Eq. (7) was used.

$$F = \frac{\Delta G^{\ddagger}}{v\Delta x^{\ddagger}} \left[1 - \left(\frac{1}{\Delta G^{\ddagger}} \ln \frac{k_0 e^{-\beta \Delta G^{\ddagger}}}{\Delta x^{\ddagger} l}\right)^{v}\right] \tag{7}$$

Where $F$ is the rupture force and where $l$ is (ln) loading rate.

For comparison to DHS-fitting, the Bell-Evans (BE) model (*36*, *37*) was used to obtain $k_0$ and $\Delta x^{\ddagger}$, from fitting of the median rupture force against the logarithm of the median loading rate. In contrast to the DHS-model, in the BE-model, $\Delta x^{\ddagger}$ is assumed to be independent of force.

$$F = \frac{k_B T}{\Delta x^\ddagger} \ln \left[ \frac{r_f \Delta x^\ddagger}{k_0 k_B T} \right] \tag{8}$$

Where $F$ is the rupture force and $r_f$ is the loading rate at rupture.

**AFM-SMFS measurements, force clamp.** Force clamp SMFS measurements were performed on a ForceRobot® AFM. To measure the bond lifetime of protein complexes at different retraction force setpoints (20, 30, 40, 50, 60 70, 80, and 90 pN or 80, 90, 100, 110, 120, 130, 140, 150, 160, 170, and 180 pN), cantilevers were briefly brought into contact with cover glasses at a force setpoint < 40 pN for a period of 0.2 s and then retracted at 4000 $nm \cdot s^{-1}$. Where the surface-cantilever interactions formed, the measured force increased as the cantilevers were retracted until the target force setpoint was reached. At the target force setpoint, the cantilever tip-sample distance was continuously adjusted to maintain the force setpoint until the complex ruptured. Measurements were limited to a maximum clamp time of 10 s. To assist in the correct selection of force *vs.* time curves, additional controls were also performed without Juno functionalized to the cover glass. PBS (pH 7.4) was utilized as the buffer solution for all measurements.

**Force clamp SMFS curve selection and data analysis.** Over a typical 15 h measurement period, ~500-1,000 force *vs.* time curves were collected for subsequent analysis. Following a combination of automated and manual processing of force clamp data, each experiment yielded ~10-60 force *vs.* time curves for each force setpoint. In the absence of fingerprint domains on the AFM constructs, an automated filter was written to exclude the signature of non-specific adhesion events identified from the non-Juno functionalized cover glass controls. These controls directly followed the Izumo1:Juno force clamp measurements, using the same cantilevers. In brief, the automated filtering excluded force *vs.* time curves with the following features as non-specific force clamping events: more than three discrete AFM-head height steps, retraction distances less than 40 nm, retraction distances greater than 200 nm. Further, bond lifetimes less than 15 ms were excluded to avoid potential artifacts due to the regularization times of the feedback loop. The resulting plotted force *vs.* times curves were then inspected manually to ensure the absence of optical or mechanical artifacts and that the target force setpoint was properly maintained.

**Force clamp data analysis.** In practice, the measured force setpoint on the ForceRobot® 300 AFM using BioLevever mini cantilevers was only accurate to within approximately 5 pN of the target setpoint. For the plotting and binning of force *vs.* time curves for subsequent statistical analysis, the force setpoint was identified as the highest mean force across any given ~5 ms window during bond clamping (as determined by the minimum bond-lifetime cut-off limit). Lifetimes were calculated as the time at which the bond was clamped until the AFM resumed its retraction phase following bond rupture. Where the catch bond states of Izumo1:Juno represent the interplay of multiple unbinding pathways, we did not expect the assumption of normality to hold for all of the binned bond lifetimes within a force clamp experiment. To validate this assumption, the Shapiro-Wilk Test was used. Subsequently, the non-parametric Kruskal–Wallis test followed by Dunn's multiple-comparison test was used for hypothesis testing of changes to bond lifetime within the binned lifetime distributions (10 pN). In all instances, $p$-values < 0.05 were considered statistically significant.

Flow cytometry-based bead-binding assay

**Polystyrene bead preparation.** With the exception of the silanization step, site-specific immobilization of samples for bead-based flow cytometry binding assays was performed as described above for the AFM-SMFS cover glass preparation. For each binding assay, 30 million 3.75 µm amino-coated polystyrene breads (SPHERO™ Amino Polystyrene Particles, Sterotech, Illinois, United States). For each washing step outlined in the site-specific immobilization protocol, the beads were centrifuged at 14,000 ×g for 3 mins, removal of the supernatant, and then resuspended in required solution.

**Flow cytometry binding assay.** Flow cytometry binding assays were performed on an Attune NxT Flow Cytometer (Invitrogen, Massachusetts, United States). Izumo1-SpyTag fusions were labeled with SpyCatcher-GFP in 0.1% tween 20 (w/v) TBS-G, as described previously (*54*). Starting at 100× the expected $K_D$, 5-fold dilution series of Izumo1-GFP were prepared in 0.1% tween 20 (w/v) TBS-G, for a total of 7 dilutions. Approximately 1 µM Juno-SpyCatcher-ddFLN4-ELP-ybbR fusion was incubated with 30 million SpyCatcher-ddFLN4 functionalized polystyrene beads in TBS-G buffer. Each Izumo1-GFP dilution was then incubated with approximately 3.5 million Juno conjugated polystyrene beads for 3 h before measurement. Directly before measurement, the polystyrene beads were washed in 0.1% tween 20 (w/v) TBS-G and resuspended to a final volume of 500 µL. For each concentration and biological replicate, 10,000 events were recorded and the measurements repeated in triplicate. The datasets were analyzed using R scripts and the Langmuir binding model was fitted to extract the $K_D$. Due to protein limitations, for the measurement of mouse Izumo1:Juno, the GFP-labeling and bead functionalization of Izumo1 and Juno, respectively, were reversed.

Monte Carlo simulations

Monte Carlo simulations based on Kramer's theory used to validate and explore the proposed multi-state catch bond model for Izumo1:Juno. Specifically, we sought to generate estimates for the kinetic parameters that govern the transition between each unbinding pathway, to test if reversible or non-reversible transitions were more likely, and to evaluate forces at which catch bond behavior might emerge. For these purposes, two different Monte Carlo approaches were conceived: one simulating constant speed AFM-SMFS and one simulating force clamp AFM-SMFS.

To briefly clarify the goal of generating estimates for the kinetic parameters that govern the transition between each unbinding pathway: unlike the kinetic parameters for P0, P1, and P2 unbinding, the energy barriers that govern Izumo1:Juno pathway switching cannot be directly observed or measured using constant speed AFM-SMFS. Instead, as described below, we performed a non-linear least square fitting of Monte Carlo simulated rupture force histograms against the experimental rupture force histograms to evaluate and refine estimates of the pathway switching kinetic parameters (see below). Thus, the Monte Carlo rupture force histograms were generated using a combination of the experimental BE $k_0$ and $\Delta x$ values for P0, P1, and P2 unbinding and randomly generated $k_0$ and $\Delta x$ values for the unbinding pathway transitions. Between random generation and non-linear least squares optimization, millions of possible $k_0$ and $\Delta x$ combinations were considered for each pathway transition.

**Constant Speed Monte Carlo simulation.** In the constant speed Monte Carlo simulations, a worm-like chain model (WLC) (*55*) was used to generate force values $F(t_i)$ across an evenly distributed molecular extension axis $X(t_i)$. The values on the molecular extension axis $X(t_i)$ were then converted to the equivalent AFM head height $H(t_i)$ using a bending correction (Eq. (9)).

$$H(t_i) = X(t_i) + \frac{F(t_i)}{k} \tag{9}$$

Together, the above steps allow for the correlation between the time of the simulation that has passed and the predicted forces acting on the simulated receptor:ligand complex. More specifically, it gives the WLC-predicted force at the current AFM head height $H(t_i)$, which is itself calculated based on the retraction speed of the AFM head and time of the simulation that has elapsed.

$$t_{i+1} = t_i + \frac{H(t_{i+1}) - H(t_i)}{V} \tag{10}$$

Next, using the relevant kinetic parameters ($k_0$ and $\Delta x$), the force-dependent off-rates were calculated for each of the unbinding pathways and the transition barriers that separate them using Eq. (11).

$$k_{\text{off}}(F) = k_0 e^{\beta F \Delta x} \tag{11}$$

For the unbinding pathways P0, P1, and P2, the experimentally derived BE kinetic parameters ($k_0$ and $\Delta x$, see **table S1**) were used. As referenced above, the energy barriers that govern transition between the unbinding pathways were assumed to behave in a manner consistent with the BE model. Accordingly, for each iteration of the Monte Carlo, the kinetic parameters for the transition between unbinding pathways were randomly generated within a defined range as follows: $k_0$ within [0.00001, 100] and $\Delta x$ within [0.01, 10].

During constant speed Monte Carlo simulations, an array of time points ($\underline{t}$) were generated and the force-dependent off-rate for each energy barrier at that time point was calculated as described above. These off-rates were then converted into the dissociation probabilities for all possible events, using Eq. (12).

$$P(F) = 1 - e^{-k_{\text{off}}(F)\Delta t} \tag{12}$$

At each time point in the simulation, the dissociation probability $P(F)$ is compared to a random number between zero and unity. If the random number is smaller than $P(F)$ the receptor:ligand unbinding or the unbinding pathway transition event occurs.

In the Monte Carlo simulations describing our kinetic models, the receptor:ligand complex always begins in the Native state (N). From N, there are two possibilities, unbinding through P0, or transition to intermediate state 1 ($I_1$). From, $I_1$ the simulated receptor:ligand complex can unbind through P1, transition to intermediate state 2 ($I_2$), or depending on if reversible transition is allowed, return to N. from, $I_2$ the simulated receptor:ligand complex can unbind through P2, or depending on if reversible transition is allowed, return to $I_1$. At each time interval, all the possible events at the current state were randomly shuffled and the first event executed. If unbinding occurs, the corresponding force is recorded as the rupture force. If a transition state occurs, the simulation moves to the next time interval. To reflect the experimental results, constant speed Monte Carlo simulations repeated for 500 replicates for each constant pulling speed.

Using bin widths of 20 pN, the Monte Carlo rupture force distributions for each constant pulling speed were evaluated against the corresponding experimental rupture force distributions using the sum of squared residuals. Each set of initial parameter values then underwent optimization to minimize the residual sum of squares. This random numerical approach to fitting was then repeated 10,000 times. To ensure robustness and consistency, the optimization process was subjected to multiple runs with different random seeds. Across 100,000+ iterations for each batch of simulations, we retained the parameter set that resulted in the lowest sum of squared residuals, thus identifying the set of parameters that best represented the force-dependent kinetics for the switching between P0, P1, and P2.

**Force clamp Monte Carlo simulation.** In the force clamp Monte Carlo, an array of force setpoints were first generated. The parameter set from the constant speed Monte Carlo simulations that resulted in the lowest sum of squared residuals when compared to the experimental constant speed data histogram distributions was retained for the force clamp Monte Carlo simulations. For each force setpoint, the force-dependent off-rates were calculated for each unbinding pathway and transition barrier separating the pathways using Eq. (11). For each time interval, the force-dependent off-rate was multiplied by the total time elapsed to give the probability of bond rupture. From there, Monte Carlo simulations followed the same rules as the constant speed Monte Carlo simulations regarding starting state, transition between unbinding pathways, and complex rupture. If rupture occurs, the corresponding elapsed time was recorded as the bond lifetime. Simulations were repeated 100 times for each force setpoint.

Molecular Dynamics Simulation and Analysis

**Molecular dynamics simulations setup.** The human Izumo1:Juno complex was modeled based on existing crystal structures (*9*, *43*) and validated as described in detail in (*44*). Standardized protocols were used for Steer Molecular Dynamics (SMD) simulations, as they have proven successful in previous projects characterizing bacterial adhesins that bind to the Fibronectin (*56*) and Vinculin (*57*). Briefly, using the NAMD version 2.14 (*58*) software and CHARMM36 force field (*59*, *60*), our systems were simulated under constant velocity pulling in an all-atom approach in explicit TIP3 water solution with periodic boundary conditions and a time step of 2 fs. Systems were neutralized and simulated in physiological salt concentration (150 mM NaCl) and NPT (number of particles, pressure, and temperature) ensemble with constant pressure (1 bar) and constant temperature (310 K) using Langevin dynamics.

To define the starting geometry of the Izumo1:Juno complex, we considered the orientation imposed by two factors: their respective membrane contact sites and the behavior of their gamete carriers in fertilization. Izumo1, a transmembrane protein on motile sperm, is represented in this study only by its extracellular region, which is crucial for fertilization. The C-terminal residue P254, where Izumo1 traverses the sperm membrane, was chosen as the pulling point and the moving restraint with a force constant of 7 kcal $mol^{-1}·Å^{-2}$ was applied on it. Juno, located on the extracellular membrane of the egg, is attached via a glycosylphosphatidylinositol anchor at residue 228. This point was used as an anchor in simulations with a force constant of 2 kcal $mol^{-1}·Å^{-2}$. Constant velocity pulling was performed at 0.001 $Å·ps^{-1}$ until full unbinding or 90 ns. This strategy allowed us to closely mimic the experimental conditions of the AFM-SMFS setup (**Fig. 1C**). Despite requiring pulling speeds several orders of magnitude higher, it has been demonstrated that SMD can recapitulate key features of mechanically induced unbinding and provide valuable insights into experimental results (*25*, *26*, *56*, *57*). We simulated 50 independent trajectories for wild-type Izumo1:Juno, Izumo1:JunoH177Q, and Izumo1:JunoH177E. mutations were prepared using QwikMD (*61*). All the simulations were run on the Piz Daint supercomputer.

**SMD data analysis and visualization.** Trajectories were analyzed using previously published protocols (*56*, *57*) and in house VMD/tool command line (TCL) and Python scripts based on the PyContact (*62*) and MDAnalysis (*63*) packages. Simulations were visualized in VMD (*64*, *65*). Contacts between Izumo1 and Juno were calculated using the distance threshold of 4.5 Å (**fig. S17**). Rupture events were defined as frames where the total number of contacts between Izumo1 and Juno dropped to 0. Since multiple unbinding and rebinding events were observed along each Izumo1:Juno pulling trajectory, the peak force was defined as the single rupture event with the highest overall force. The force loading rate was then determined by linearly fitting the force *vs*. time data immediately prior (0.5%-2% of the total trajectory frames) to the rupture event with the peak force. Rebinding distance and rebinding time were measured between the first and the last time Izumo1:Juno lost all the contacts. Rebinding rate was calculated as the number of rupture events per $ns^{-1}$. C-terminal to C-terminal distances for Izumo1:Juno complexes were measured between the center of mass of anchoring residue on Juno (S228) and pulling residue on Izumo1 (P254). Contact surface area was calculated in VMD (*64*, *65*) using the TCL terminal for each trajectory at the initial production frame assuming it represents the unloaded state and at 2 ns assuming it is in a loaded state (**fig. S18**).

#### Supplementary Text

Flow cytometry-based bead-binding assay controls

Juno-coated polystyrene beads were functionalized as per the described method. These beads were then incubated with GFP at the concentrations shown in **fig. S1**. For the GFP control we observed minimal non-specific binding at concentrations up to 20 µM. As an additional control, to ensure that the upper dynamic range for our flow cytometry-based bead-binding assay covered all variants of interest, the binding affinity for mouse Izumo1:Juno (mIzumo1:Juno) was also measured at 6.05 ± 0.15 µM (± SE) (**fig. S1**). This result is several fold above the highest expected binding affinity for the human Izumo1:Juno (hIzumo1:Juno) mutants and within 2-fold of the previously reported surface plasmon resonance (SPR) measured value of mIzumo1:Juno, 12.3 ± 0.2 µM (*2*). As with the experimental measurements, both controls were completed for two biological replicates. In combination with the accuracy of the wild-type hIzumo1:Juno complex in our flow cytometry-based bead-binding assay, relative to SPR (*9*), which provides information on the lower dynamic range of our assay, we are confident in the accuracy of the equilibrium binding behavior described for the surface functionalized Izumo1:Juno variants used in this study.

Confirming the absence of intermediate states for P0

We sought to verify that the near complete absence of intermediate states in P0 was not simply the result of bias during force *vs.* extension curve selection or classification. A specific scenario where this might be problematic is the exclusion of two concurrent P0 unbinding events (two-to-from an intermediate followed by P0 unbinding. In both instances you would expect to see two peaks at ~20-60 pN. To address this concern, we plotted the force distributions relating to the initial appearance of the intermediates in P1 and P2 for each constant pulling speed (**fig. S4**, **B** and **D**). Here, we see that the force distributions for the initial appearance of the intermediate states do not overlap with the rupture force distribution for P0 shown in **Fig. 1F**. Thus, we conclude that it is more likely that the contour length increments measured relate to a specific conformational change linked to, or following, the state changes that result in P1 or P2 unbinding.

On possibility of Izumo1 Ig-like domain unfolding contributing to intermediate state formation

A more conventional explanation for the observation of intermediate states during Izumo1:Juno unbinding is the unfolding of Izumo1 residues located outside of the interconnected disulfide network. Such a scenario would be largely limited to the partial unfolding of the Izumo1 Ig-like domain (**fig. S4E**). While the unfolding of Izumo1 residues 166 to 181 and 234 to 255 would contribute a contour length increment of 13.1 nm (15 aa × 0.365 nm = 5.5 nm and 21 aa × 0.365

nm = 7.7 nm, respectively), we see no evidence of the Ig-like domain unfolding prior to rupture in the SMD simulations (**Supplementary Movie 1, Supplementary Movie 2**). Furthermore, across the different experimental constant pulling speeds, we often observe secondary peaks in the plotted contour length distributions (**fig. S4**). The ~25-40 nm final contour lengths of these secondary peaks are best explained by some combination of rare independent unfolding events (*i.e.* the unfolding of the Ig-like domain) in addition to the force-dependent re-organization of the Izumo1:Juno interface. The possibility of Izumo1 unfolding is further discussed in relation to Supplementary Text sections for the constant speed AFM-SMFS of the Izumo1 point mutations, in particular for Izumo1K150A:Juno and Izumo1R160A:Juno.

An important consideration for any scenario involving fertilization is that any unfolding of Izumo1 or Juno that increases the membrane-to-membrane distance > 10 nm would actively inhibit membrane fusion (*66*). Thus, unfolding of the Ig-like domain at the experimentally observed rates for the appearance of intermediate states during Izumo1:Juno unbinding would seem counterintuitive to sperm-egg cell fusion. We do not believe this assumption holds true in the force-dependent reorganization of the Izumo1:Juno interface purported by our SMD simulations. Here, Juno acts like a fulcrum to drive the N-terminal fusion associated residues of Izumo1 (F28, W88, and W113 (*11*)) towards the egg cell membrane (**fig. S21**). Additionally, contact between Izumo1 and the egg-cell membrane would alleviate tension from the C-terminal Juno α-helix, prevents unfolding. Therefore, the increased distance between the C-termini of Izumo1 and Juno in our unbinding model would not necessarily inhibit membrane fusion in the same way that Ig-like domain unfolding might. Instead, the observed intermediate states may in fact represent key molecular events in overcoming the energetic barriers of membrane fusion. As discussed in the main text, such an unbinding mechanism also provides a possible explanation for the unexpectedly low rates of Izumo1 fusogenic activity observed under equilibrium conditions (*11*).

<u>Rationale for the selection of Izumo1 mutants based on sequence conservation, equilibrium binding behavior, and SMD simulations</u>

**Izumo1E71A.** Izumo1 E71 is a divergent residue that forms a salt bridge with Juno K163 at the equilibrium binding interface (*9*) (**fig. S5B**). An alanine substitution at Izumo1 E71, forming the complex Izumo1E71A:Juno, has previously been shown to have minimal impact on the equilibrium binding behavior (*9*). Similarly, we observed a $K_D$ for Izumo1E71A:Juno of 43 ± 5 nM (± SE) (**fig. S1**). To reiterate, in our wild-type Izumo1:Juno SMD simulations, when force is applied to through the pulling of the C-termini residues at a constant speed, Cluster 1 unbinds first (**Fig. 2, G** and **H**). The resulting second-state Izumo1:Juno interface then consists primarily of Cluster 2 amino acid residues, which includes Izumo1 E71. During unbinding from the second state, Izumo E71 typically forms one of the terminal interactions with Juno (**fig. S5B, Supplementary Movie 1**). Stochastically, the Cluster 2 amino acids stabilize Izumo1:Juno long enough for the

Cluster 3 interactions to form, thereby transitioning the system to a third state and therefore the three-state unbinding pathway (**fig. S5C, Supplementary Movie 2**). Once in the three-state unbinding pathway, Izumo1 E71 is removed from any interaction with Juno. Based on the analysis of these SMD simulations we reasoned that where Izumo1 E71A has minimal impact on the $K_D$, Izumo1E71A might impact the mechanostability of P1 and, or, the P1 → P2 transition rate. Based on this analysis and to further build upon previous characterization efforts of Izumo1E71A, we selected Izumo1E71A as one of the divergent Cluster 2 mutations used in this study.

**Izumo1K150A.** Izumo1 K150 is a divergent residue that forms a hydrogen bond with Juno C79 at the equilibrium binding interface (*9*) (**fig. S6B**). Previously uncharacterized, our equilibrium binding affinity experiments in **fig. S1** demonstrates a modest decrease in $K_D$ for Izumo1K150A:Juno to 110 ± 27 nM (± SE). At equilibrium, Izumo1 K150 represents the N-terminal-most Cluster 2 amino acid that is involved in the initial binding interface. With the application of force, additional contacts are made that extend the binding interface beyond Izumo1 K150, thereby increasing the contact surface area (**Fig. 2E**; **fig. S6, B** and **C**). This increase in contact surface area under load is linked to the catch bond behavior we report for Izumo1:Juno. Similar to Izumo E71, as the system transitions from the two-state to the three-state unbinding pathway, Izumo1 K150 forms one of the terminal interaction points with Juno. Stochastically, these terminal interactions in our SMD simulations seem to support the formation of the Cluster 3 contacts necessary for transition to the three-state pathway. Furthermore, whilst Izumo1 K150 is a divergent residue, it is part of a highly-conserved short two amino acid loop boarded by two different disulfide bonds (*9*). Of note, one of these cysteines, Izumo1 C152, forms a disulfide bond with the N-terminal most residue of Izumo1, C22 (**fig. S4E**). We therefore had to consider the possibility that a mutation in this region could alter the disulfide network and cause large intermediate unfolding events that might affect long term stability of AFM-SMFS measurements. Overall, we were interested in any possible effects of an alanine substitution Izumo1 K150 on the function of the short Izumo1 loop, in particular on the P0 → P1 and P1 → P2 transition rates.

**Izumo1H157A.** Izumo1 H157 is a highly conserved residue forming no clear non-covalent interactions with Juno in the crystal structure (*9*). However, as previously shown (*44*), Izumo1 H157 forms a hydrogen with Juno R87 following equilibration in MD structures (**fig. S7B**). Experimentally, a single alanine substitution, forming the complex Izumo1H157A:Juno, has previously been shown to cause large changes to the equilibrium binding behavior of Izumo1:Juno (*9*). In this study, we observed a $K_D$ for Izumo1H157A:Juno of 243 ± 23 nM (± SE). As a Cluster 1 residue, we observed the involvement of Izumo1 H157 in the initial contacts between Izumo1:Juno before unbinding during the transition to the second-state in our SMD simulations (**fig. S7B**). Following, unbinding of Izumo1 H157 and transition to the second-state, Izumo1 H157 formed no additional interactions (**fig. S7, B** and **C**). Therefore, we expected that any resulting changes to unbinding the behavior would be localized to the interaction between Izumo1 H157-Juno R87. Our rationale to select Izumo1 H157 was based on the high degree of sequence conservation across mammalian species (*9*) and as Cluster 1 with an established negative impact on equilibrium binding affinity (*9*). From a mechanical perspective, we expected it to impact primarily the P0 → P1 transition rate.

**Izumo1R160A.** Izumo1 R160 is a conserved residue that forms hydrogen bonds with Juno E45 and Juno H65 in the equilibrium crystal structure (*9*) (**fig. S8B**). A single alanine substitution, forming the complex Izumo1R160A:Juno, has previously been shown to impact the equilibrium binding behavior of Izumo1:Juno (*9*). Similarly, here, we observed a $K_D$ for Izumo1R160A:Juno of 157 ± 14 nM (± SE) (**fig. S1**). A Cluster 2 residue, our SMD simulations predict that Izumo1R160 unbinds early following the transition to the second-state. Following this initial unbinding, Izumo1R160 forms no subsequent interactions and the system can evolve as either a two-or-three state unbinding trajectory, depending on if cluster the three amino acids form stable interactions (**fig. S8, B** and **C**). Finally, as a Cluster 1 residue, Izumo1 R160 is located close to the Ig-like domain and it was hypothesized that disruption of the force-propagation networks in this area might provide additional insight into the origin of the contour length intermediates.

<u>Analysis of fitted DHS and BE parameters for Izumo1 point mutations</u>

We performed constant speed AFM-SMFS on the Izumo1 point mutations in triplicate, using the same experimental approach described for wild-type Izumo1:Juno. Similarly, to obtain insights into the theoretical energy landscapes for the unbinding pathways, we used the same force *vs.* extension curve sorting and model fitting approaches as wild-type Izumo1:Juno (**table S2**). Where broad agreements were observed in the DHS and BE derived $k_0$ values, they were limited to the increase in P2 $k_0$ for Izumo1E71A:Juno, the decrease in P1 $k_0$ for Izumo1K150A:Juno, and the decrease in P2 $k_0$ for Izumo1R160A:Juno (**table S2**). For the DHS and BE derived $\Delta x$ values, agreement was found for decreased P0, increased P1, and decreased P2 in Izumo1E71A:Juno; and increased P0 and decreased P2 in Izumo1K150A:Juno (**table S2**). However, in no instances were these differences sufficient to result in more than subtle changes to mechanostability of the force activated pathways. Indeed, with the exception of P1 for Izumo1K150A:Juno, we see similar or greater median rupture forces for each of the Izumo1 mutants in P1 and P2 for all pulling speeds (**fig. S5-8**). Thus, as explained in the main text, it becomes clear that where alanine substitutions of single Izumo1 residues are sufficient to disrupt the equilibrium binding behavior of Izumo1:Juno (**fig. S1**) (*9*), the mechanical behavior of Izumo1:Juno offers a far greater level of functional redundancy. In the context of natural selection and the necessity of successful fertilization, this suggests that the mechanical stability of Izumo1:Juno is of greater import than the equilibrium binding affinity.

Analysis of unbinding pathway distribution and intermediate states for Izumo1 point mutations

**Izumo1E71A:Juno.** Interestingly, despite Izumo1 E71 forming terminal interactions during the two-state unbinding SMD simulations of the wild-type complex (**fig. S5B**); experimentally, we observed the ratio of P0 to P1 and P1 to P2 transitions increasing for Izumo1E71A:Juno (**fig. S5, D** and **E**). This behavior, observed at all constant pulling speeds, suggests that the terminal interactions observed for Izumo1 E71 during SMD are not directly relevant for the mechanostablility of each unbinding pathway. Instead, we propose they regulate the formation of Cluster 3 interactions between Izumo1 and Juno (**fig. S5, B** and **C**). Unique amongst the Izumo1:Juno variants used in this study, for Izumo1E71A:Juno we also observe the increased presence of long intermediate states in P0 at pulling speeds of 3200 and 6400 $nms^{-1}$ (**fig. S5F**). Here, the median contour length changes of 30.1 ± 9.8 nm (± MAD) were observed. Given the established relationship between the higher order unbinding pathways and the observation of intermediate states in wild-type Izumo1:Juno, these results indicate the presence of unique intermediate states for Izumo1E71A:Juno in P0. We also see a tailing in the contour length distribution of P1 and P2 intermediates for Izumo1E71A:Juno that is a further indication that the additional intermediate states observed in P0 are also present in P1 and P2 (**fig. S5, G** and **H**). However, the resulting median P1 (15.6 ± 7.2 nm, MAD) and P2 (16.2 ± 9.2 nm, MAD) contour length values remain similar to the wild-type Izumo1:Juno, suggesting that any such intermediates are of low abundance relative to the most prominent intermediate states. Overall, our constant speed AFM-SMFS experiments and SMD simulation implicate Izumo1 E71 primarily in the transition between the unbinding pathways.

**Izumo1K150A:Juno.** Like Izumo1E71A, Izumo1K150A is a Cluster 2 single alanine substitution**.** Unlike Izumo1E71A:Juno, the ratios for P0:P1:P2 in Izumo1K150A:Juno remain largely unchanged (**fig. S6, D** and **E**). However, similar to Izumo1E71A:Juno, for Izumo1K150A:Juno the median P1 (20.7 ± 13.1 nm, MAD) and P2 (21.4 ± 15.5 nm, MAD) contour length values increase relative to the wild-type Izumo1:Juno at 3200 $nms^{-1}$ (**fig. S6, G** and **H**). We note that for Izumo1K150A:Juno, the AFM-SMFS measurements displayed poor stability, with cantilevers regularly losing activity after only 1-2 h. As such, only one replicate remained active for the entire measurement duration of 12 h and is shown in **fig. S6**. We posit that the increased contour lengths and broader distribution of the intermediate states for this mutant may indicate the presence of some additional states unique to Izumo1K150A:Juno (**fig. S6, G** and **H**). It is possible that these additional intermediates impact long-term functionality following mechanical stresses. As previously discussed in the rationale for the selection of K150A as an Izumo1 mutant, this loss of mechanostability could be related to the bordering disulfide bonds (C22-C149). Disruption of this particular disulfide bond could lead to unfolding of parts of the Izumo1 alpha helical bundle (**fig. S4E**) and early failure of the cantilever tips through loss of Juno recognition.

**Izumo1H157A:Juno.** For the remaining Cluster 2 alanine substitution, Izumo1H157A, we observe inhibition of transition to the P2 pathway at all pulling speeds. Of particular note, only 7.9 ± 6.8% (SEM) of Izumo1H157A:Juno complexes unbind through P2 at 6400 $nms^{-1}$ (**fig. S7, D** and **E**). Furthermore, the median P0 and P1 rupture forces for Izumo1H157A:Juno are between 25.1 to 36.8 pN for loading rates of 1.2 to 14.6 $nNs^{-1}$ and 89.2 to 144.9 pN for loading rates of 2.6 to 39.6 $nNs^{-1}$, respectively (**fig. S7D**). Thus, the possibility that the decrease in P2 unbinding is due to the premature rupture of Izumo1H157A:Juno can be excluded. Where the transition from P1 to P2 in Izumo1H157A:Juno is inhibited, a reverse transition to P0 increases in likelihood, thereby increasing the abundance of P0 at all pulling speeds except 800 $nms^{-1}$. Finally, for Izumo1H157A:Juno we observe no change to the relative rates for intermediate transition (**fig. S7F**) or the median contour lengths of the intermediate states for P1 or P2 (**fig. S7, G** and **H**). Overall, it seems likely the conserved Cluster 2 residue Izumo1 H157 is involved in orienting the complex in the second-state for subsequent transition into the P2 unbinding pathway.

**Izumo1R160A:Juno.** Several interesting changes were observed for the Cluster 1 mutant Izumo1R160A (**fig. S8**). First, unique amongst all of the Izumo1:Juno variants tested, the GMM classifier identified a small proportion of Izumo1R160A:Juno complexes, 3.6 ± 2.8% (SEM), unbinding through P2 at pulling speeds of 800 $nms^{-1}$ (**fig. S8, D** and **E**). Yet, the percentage of P2 unbinding events for Izumo1R160A:Juno is lower for all other pulling speeds (**fig. S8, D** and **E**). This effect is particularly evident at pulling speeds of 6400 $nms^{-1}$ , where a ~20% reduction relative to wild-type Izumo1:Juno is observed (**fig. S8, D** and **E**). Second, Izumo1R160A:Juno demonstrates an increase in P1 and P2 intermediate transitions events at all pulling speeds (**fig. S8F**). Third, we note that the median change in contour length following transition through these intermediate states increases for Izumo1R160A:Juno relative to wild-type Izumo1:Juno (**fig. S8G**). Here, the median contour length change is 22.4 ± 13.8 and 28.4 ± 14.9 nm (MAD), for P1 and P2 at 3200 $nms^{-1}$, respectively, nearly double that of the wild-type Izumo1:Juno. We predict that due to the proximity of Izumo1 Cluster 1 residues to the Ig-like domain, the disruption of the force propagation pathway in Izumo1R160A:Juno (*i.e.* the hydrogen bonds between Izumo1 R160 and Juno E45 & H65) results in the unfolding of the Izumo1 Ig-like domain (amino acids 165-182 and 233-255, 17 + 22 × 0.365 = 14.2 nm) (**fig. S8G**). Given the possible sources of contour length increments for Izumo1 and Juno (shown in **fig. S4, E** and **F**, respectively), the additional increases in contour length observed in Izumo1R160A:Juno support our initial conclusions from the wild-type Izumo1:Juno SMD simulations. Specifically, that the contour length increments that are observed in wild-type Izumo1:Juno are largely the result of force induced reorientation of the Izumo1 and Juno binding interface rather than direct unfolding events.

Support for the reversibility of unbinding pathway transitions from constant speed AFM-SMFS

Despite the overall preservation of mechanostability for each unbinding pathway in the studied Izumo1 mutants, the alanine substitution of the well-conserved Cluster 2 residues had clear impacts on the P0, P1, and P2 unbinding ratios (**fig. S5-8**). This effect was particularly prevalent for the probability of unbinding through P2 at higher pulling speeds. At 6400 $nms^{-1}$, we observe a ~20% increase in P2 unbinding for Izumo1E71A:Juno and a ~20% decrease in P2 unbinding for Izumo1H157A:Juno (**fig. S5** and **fig. S7**). For reference, the well-conserved Cluster 1 mutant, Izumo1R160A:Juno, showed a ~10% decrease in P2 unbinding at 6400 $nms^{-1}$ (**fig. S8**) and the divergent Cluster 2 mutant, Izumo1K150A:Juno, showed no change in P2 unbinding at 6400 $nms^{-1}$ (**fig. S6**). This dichotomy of Cluster 2 alanine substitutions increasing or decreasing unbinding through higher order unbinding pathways, whilst subtly enhancing overall mechanostability in both instances, suggests the possibility of a dynamic equilibrium between the unbinding pathways. In particular it brings into focus a mechanistic question of whether the alanine substitutions are simply affecting the forward transition to higher order unbinding pathways (*i.e.* P0 → P1 → P2) or if reversible transitions are indeed possible on the experimental timescale (*i.e*. P0 ⇌ P1 ⇌ P2) (*34*). In scenarios or mutants that favor the transition to higher order pathways or inhibit the transition back to lower order pathways, the mechanostability of the eventual unbinding pathway would be enhanced as less time is spent in lower mechanostability states (*34*). Thus, we hypothesize that the small increases in mechanostability of Izumo1E71A:Juno and Izumo1H157A:Juno are the result of less time spent in lower order states during unbinding.

Unfolding events observed in SMD simulations

We observed two types of secondary structure adjustments across the SMD simulations. The first was the previously discussed unfurling of the C-terminal Juno α-helix, where the occurrence increased from 4% in wild-type to 14% in both JunoH177Q and JunoH177E (**Fig. S19**). The second major adjustment involved the stretching of the Izumo1 between residues E66 and V78, and subsequent unfurling of the α2-helix of Izumo1 (**Fig. S20**). This structural change was most prevalent in the Izumo1:JunoH177Q and Izumo1:JunoH177E simulations, but was also observed to a lesser degree in the wild-type simulations.

#### Supplementary Figures

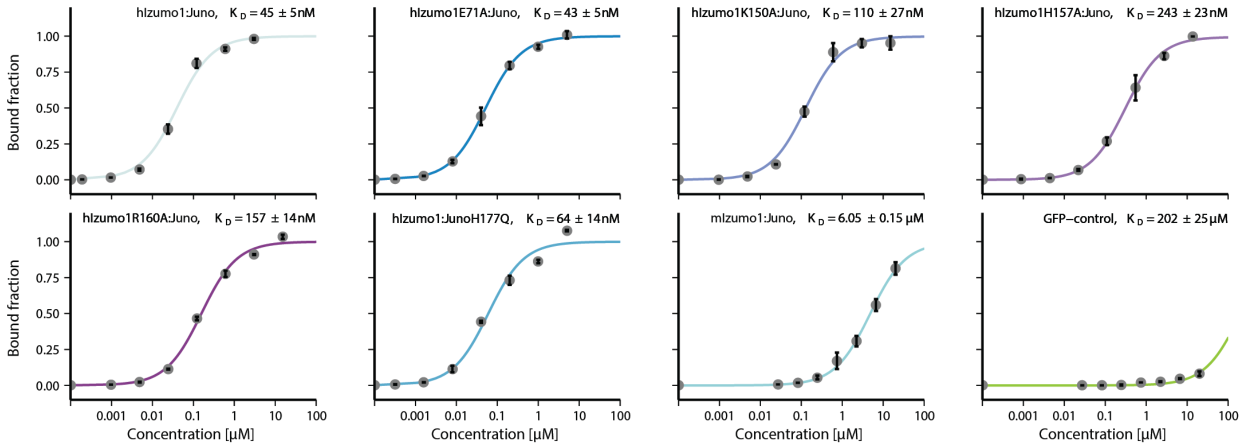


**Figure S1: Flow cytometry-based bead-binding assay.** Binding assays were performed for wild-type human (h) Izumo1:Juno, Izumo1E71A:Juno, Izumo1K150A, Izumo1H15A:Juno, Izumo1R160A:Juno, and Izumo1:JunoH177Q. Also included are mouse (m) Izumo1:Juno as an upper dynamic range control and GFP-only as a negative control. The bound fraction for each biological replicate represents the average of three measurements. Dots represent the average of two independent biological replicates. Error bars represent the range of the two independent biological replicates. Lines represent non-linear least squares fitting with a single binding site. Binding affinities reported as $K_D$ ± SE of the fitted parameters.

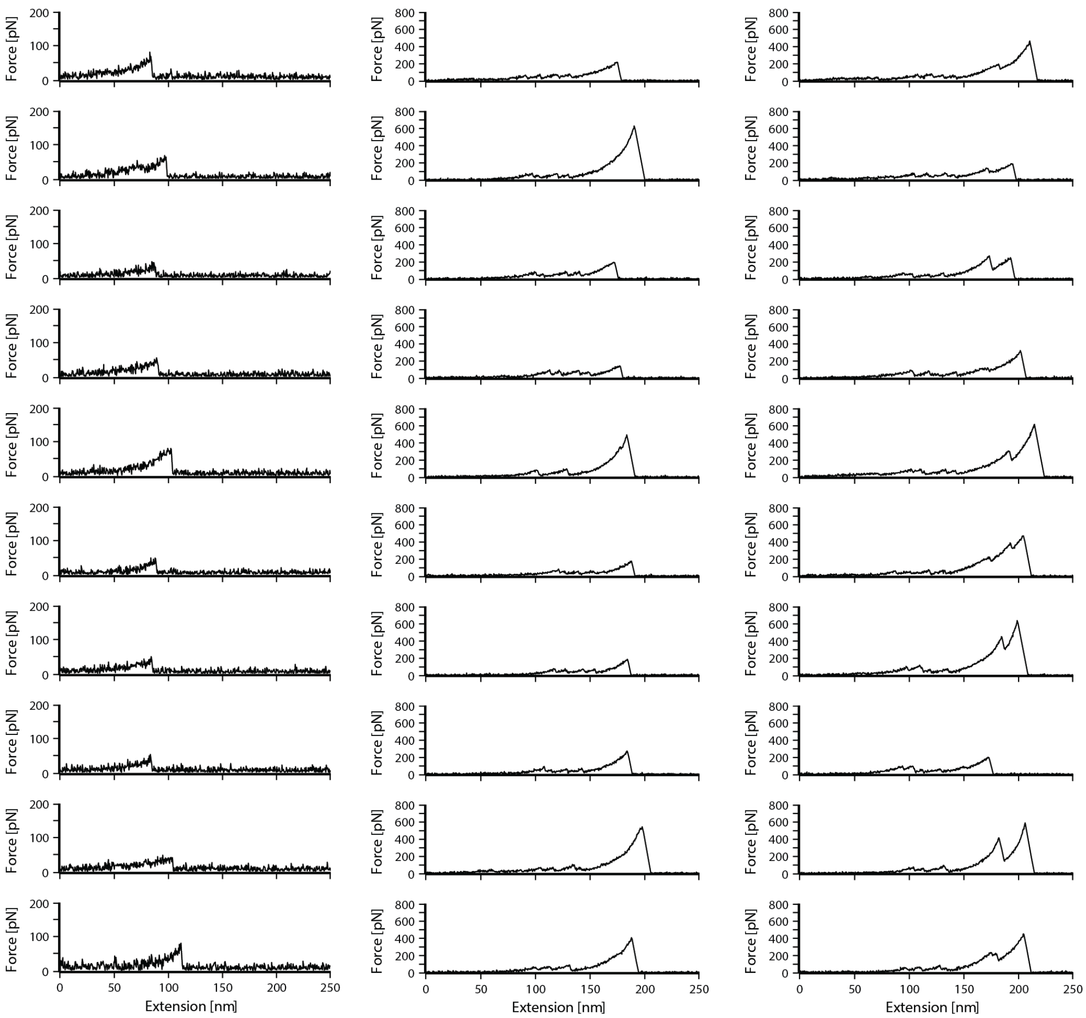


**Figure S2: Representative constant speed SMFS force *vs.* extension curves from this study following automated and manual sorting.** Each curve represents a distinct single Izumo1:Juno rupture event corresponding to each unbinding pathway subgroup a, b, or c. In the first column are force *vs.* extension curves showing Izumo1:Juno rupture at low force and in the absence of ddFLN4 fingerprint unbinding, classified as a. In the second column are force *vs.* extension curves showing Izumo1:Juno rupture following ddFLN4 fingerprint unfolding, classified as b. In the third column are force *vs.* extension curves showing Izumo1:Juno rupture following ddFLN4 fingerprint unfolding and an additional conformational rearrangement within the Izumo1:Juno complex, classified as c.

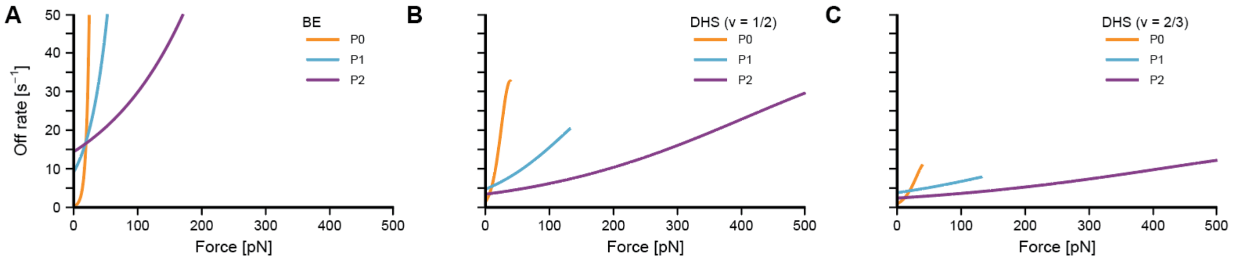


**Figure S3: Force-dependent off-rates for Izumo1:Juno.** Force-dependent off-rates for Izumo1:Juno calculated from the fitted Bell-Evans (BE) and Dudko-Hummer-Szabo (DHS) kinetic parameters, $k_0$ and $\Delta x$. (**A**) BE model fitted kinetic parameters (Eq. 11). (**B**) DHS model fitted kinetic parameters where $v$ = 1/2. (**C**) DHS model fitted kinetic parameters where $v$ = 2/3. All fitted kinetic parameters are available in **table S1**.

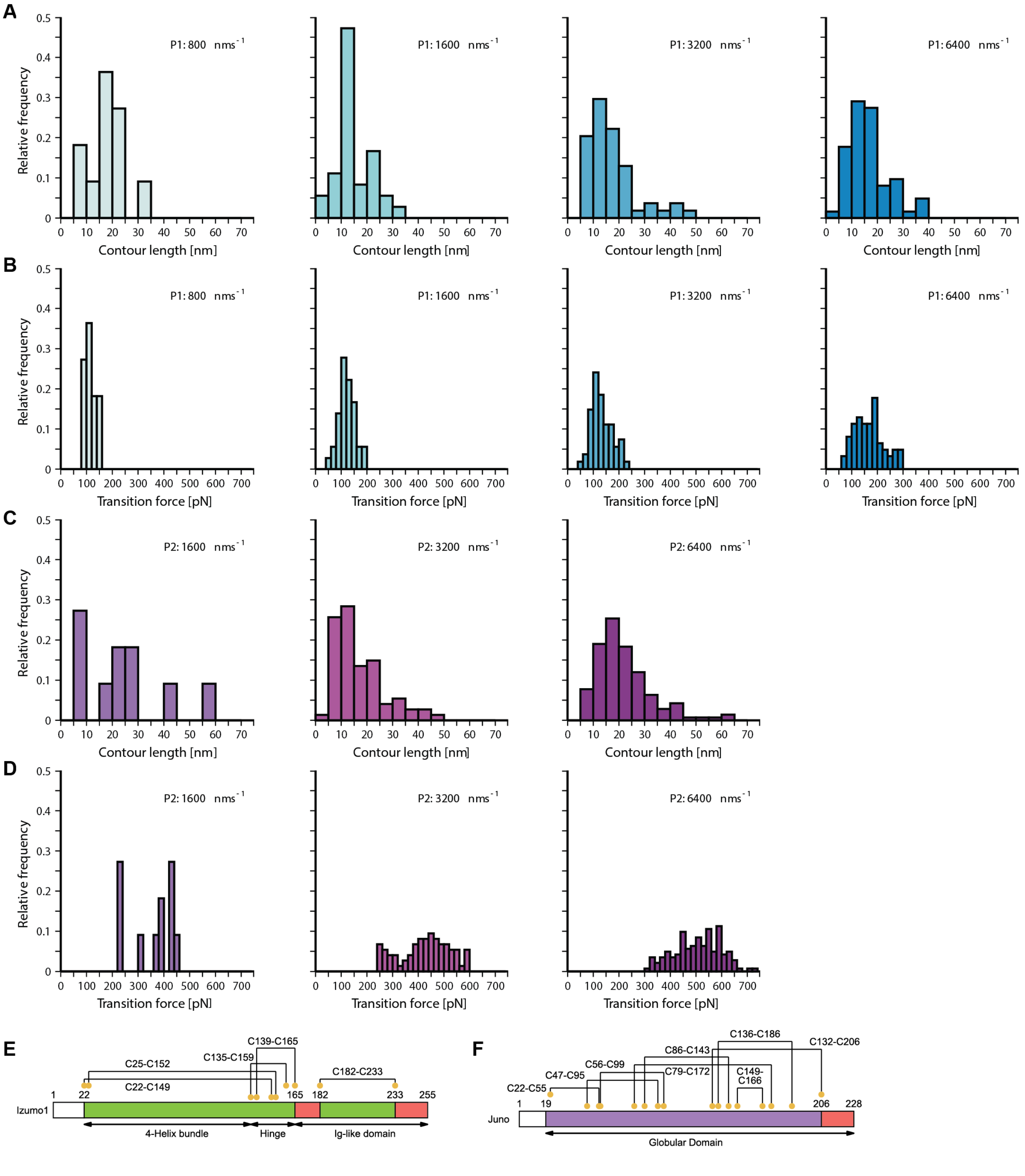
A
B
C
D
E
F
Relative frequency
Contour length [nm]
Transition force [pN]
P1: 800 nms⁻¹
P1: 1600 nms⁻¹
P1: 3200 nms⁻¹
P1: 6400 nms⁻¹
P2: 1600 nms⁻¹
P2: 3200 nms⁻¹
P2: 6400 nms⁻¹
Izumo1
C25-C152
C22-C149
C135-C159
C139-C165
C182-C233
1 22 165 182 233 255
4-Helix bundle
Hinge
Ig-like domain
Juno
C22-C55
C47-C95
C56-C99
C86-C143
C79-C172
C136-C186
C149-C166
C132-C206
1 19 206 228
Globular Domain

**Figure S4: Contour length distributions for Izumo1:Juno folded intermediate states.** (**A**) Contour length distributions for Izumo1:Juno rearrangements occurring prior to rupture through P1. (**B**) Force distributions for Izumo1:Juno rearrangements occurring prior to rupture through P1. (**C**) Contour length distributions for Izumo1:Juno rearrangements occurring prior to rupture through P2. (**D**) Force distributions for Izumo1:Juno rearrangements occurring prior to rupture through P2. Contour lengths show the total contour length change of Izumo1:Juno complexes following ddFLN4 unfolding until final Izumo1:Juno rupture. Transition forces show the force at which the first non-ddFLN4 contour length change occurs. For P1 800, 1600, 3200, and 6400 $nms^{-1}$, $n$ = 122-322. For P2 1600, 3200, and 6400 $nms^{-1}$, $n$ = 23-207. (**E**) Structural topology for Izumo1, including the conserved disulfide network. (**F**) Structural topology for Juno, including the conserved disulfide network. For both Izumo1 and Juno topologies, red indicates regions available for mechanically induced unfolding. Ig-like = Immunoglobulin-like.

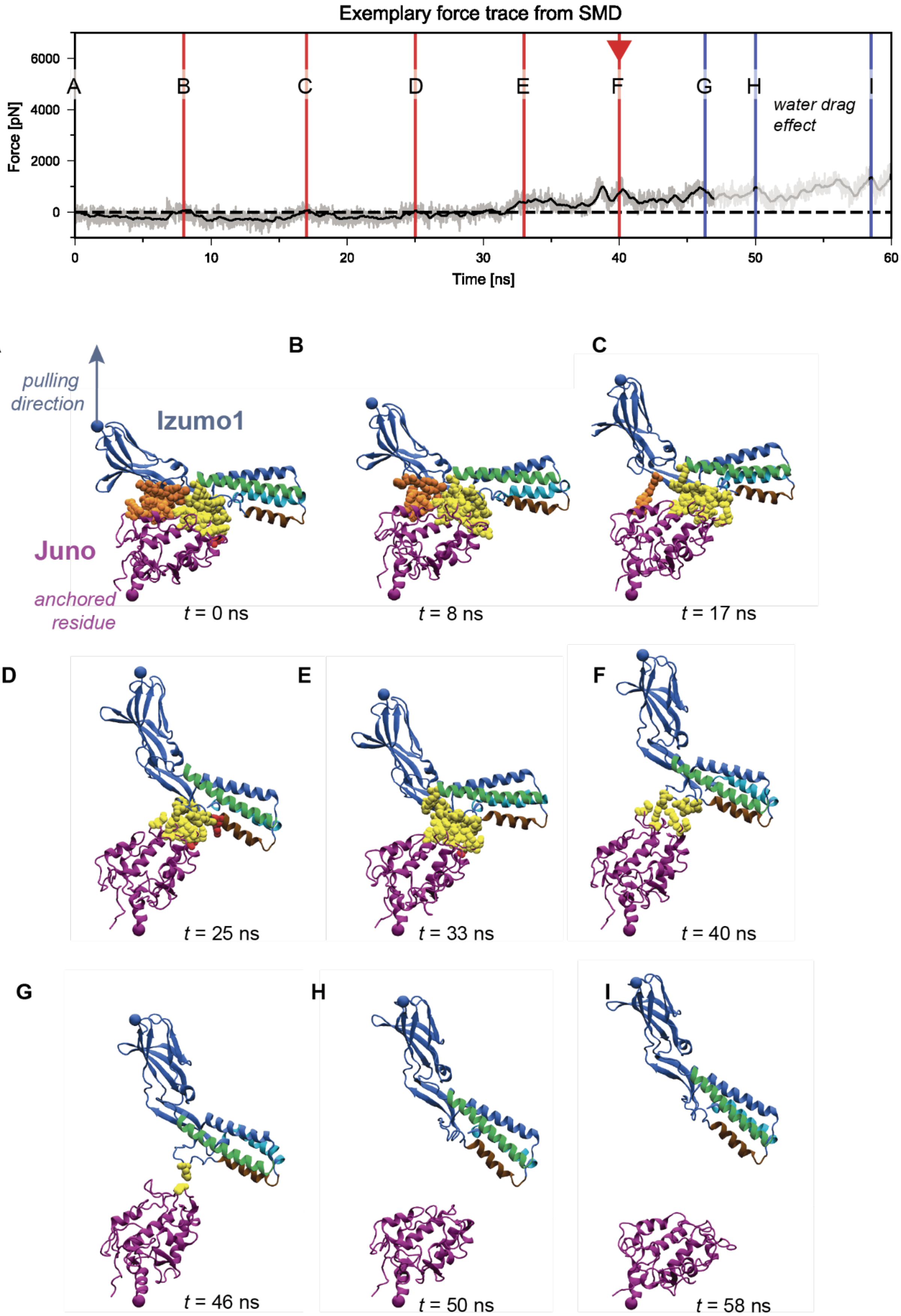

Exemplary force trace from SMD
Force [pN]
6000
4000
2000
0
A
B
C
D
E
F
G
H
I
water drag effect
0
10
20
30
40
50
60
Time [ns]
A
pulling direction
Izumo1
Juno
anchored residue
t = 0 ns
B
t = 8 ns
C
t = 17 ns
D
t = 25 ns
E
t = 33 ns
F
t = 40 ns
G
t = 46 ns
H
t = 50 ns
I
t = 58 ns

**Figure S5: Force *vs*. time trace with structure snapshots representative of the two-state wild-type Izumo1:Juno unbinding observed in SMD simulations.** Letters and lines mark timepoints at which snapshots were taken and the red triangle marks the first Izumo1:Juno rupture event. Gray traces represent force for individual frames and black lines are the rolling average force across the 50 frames. The number of contacts between Izumo1 and Juno as a function of time can be found in **fig. S9A** and the details of the corresponding interactions are shown in **fig. S6**. Preview of the unbinding process can be observed in **Supplementary Movie 1**.

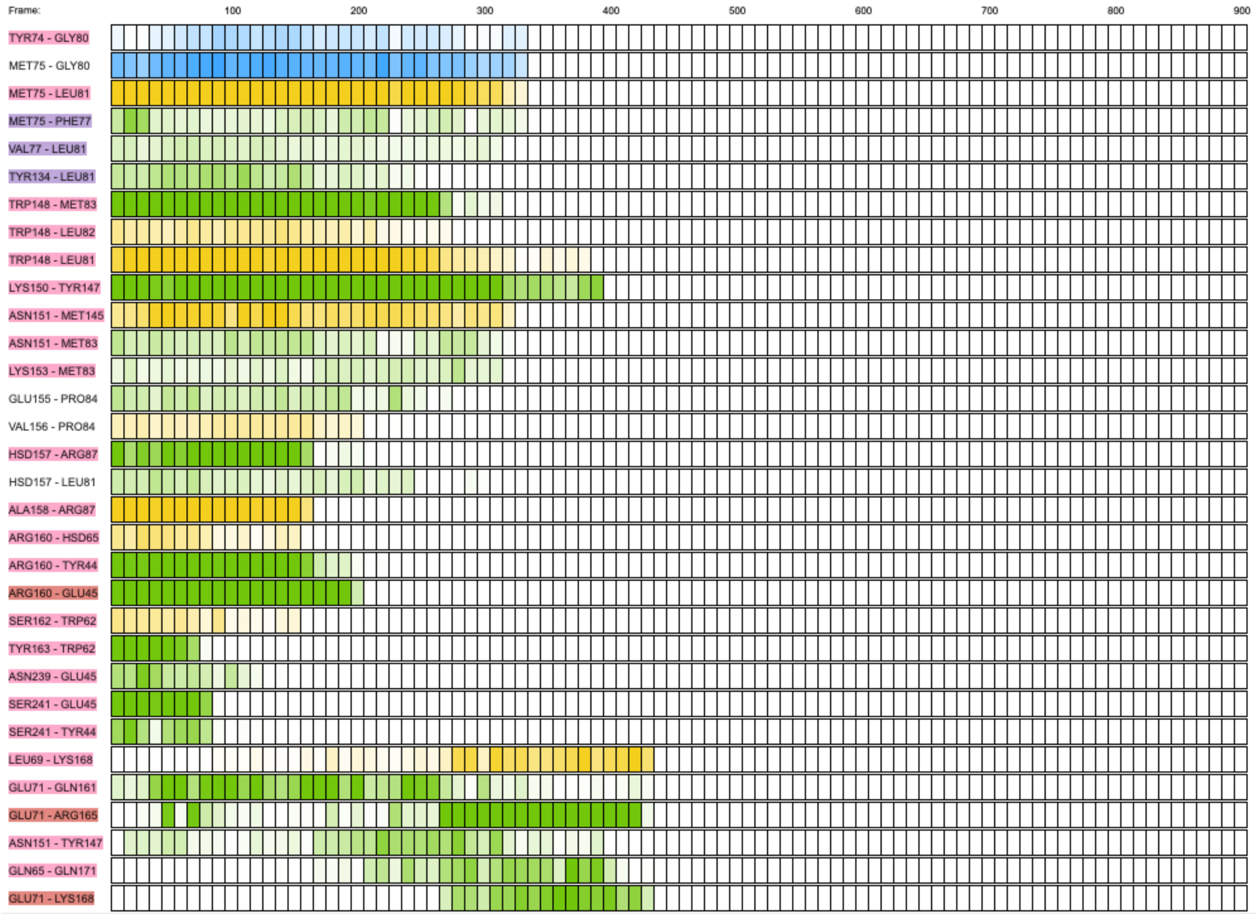


**Figure S6: Exemplary time series of the non-covalent interactions as observed in the two-state unbinding pathway of the wild-type Izumo1:Juno complex in SMD simulations.** Known binding interface of Izumo1:Juno complex gradually breaks within 50 ns, forming only few transient interactions in the process. **Supplementary Movie 1** shows the preview of the corresponding simulation. The first residue in the label belongs to Izumo1, and the second residue belongs to Juno. The color of the label is determined by interaction type and geometry: red, salt bridge; blue, hydrophobic; and pink, hydrogen bond. Each box corresponds to the equal trajectory segment, their color signifies interaction type and the intensity shows the strength of the interaction: green, side chain/side chain; yellow, side chain/backbone; blue, backbone/backbone. PyContact (*62*) was used to extract this data with a filter of mean score value > 0.25 for trajectory of the total simulation time of 90 ns (900 frames extracted).

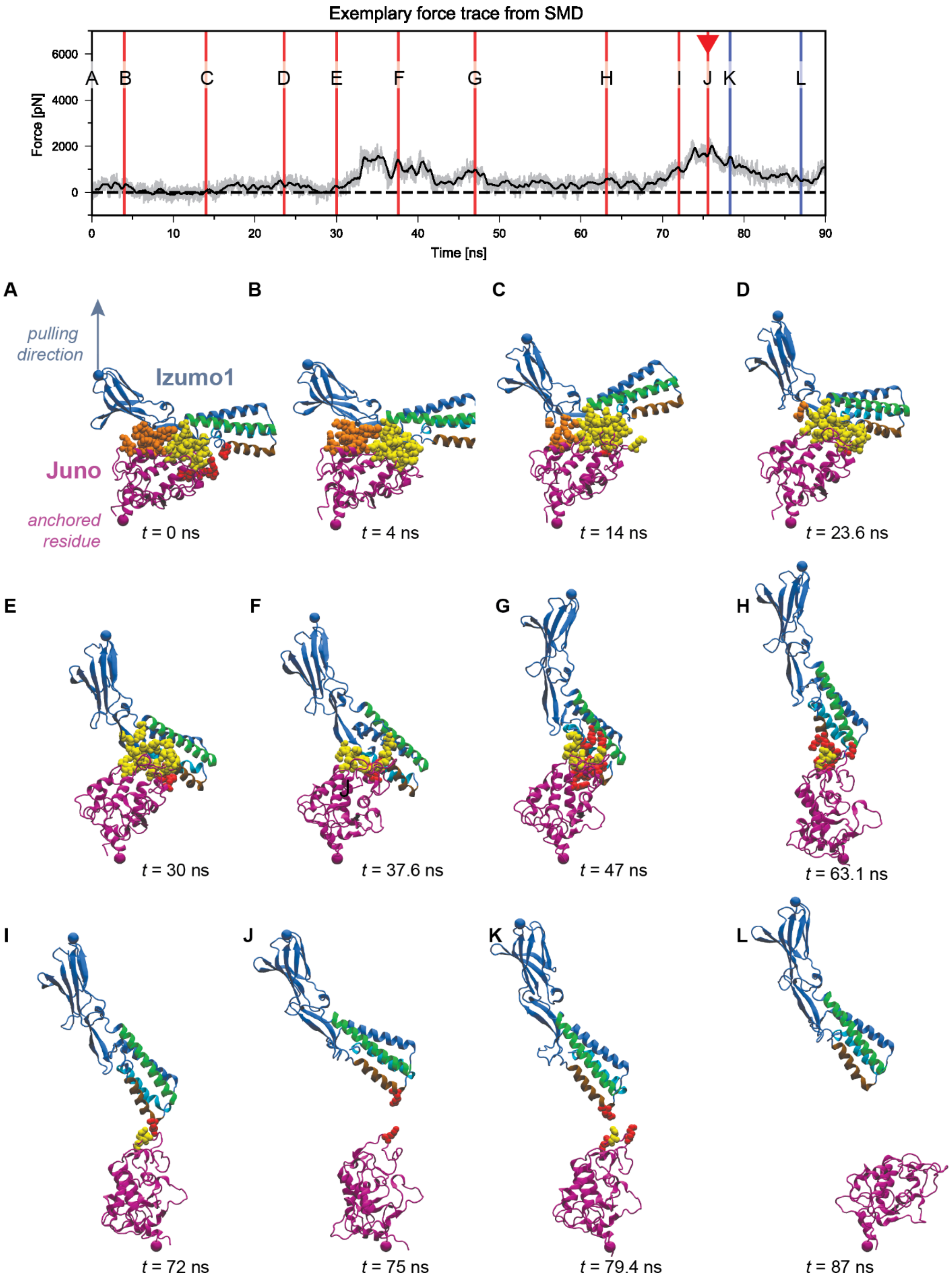

Exemplary force trace from SMD
Force [pN]
Time [ns]
A B C D E F G H I J K L
pulling direction
Izumo1
Juno
anchored residue
t = 0 ns
t = 4 ns
t = 14 ns
t = 23.6 ns
t = 30 ns
t = 37.6 ns
t = 47 ns
t = 63.1 ns
t = 72 ns
t = 75 ns
t = 79.4 ns
t = 87 ns

**Figure S7: Force *vs*. time trace with structure snapshots representative of the three-state wild-type Izumo1:Juno unbinding observed in SMD simulations.** Letters and lines mark timepoints at which snapshots were taken and the red triangle marks the first Izumo1:Juno rupture event. Gray traces represent force for individual frames and black lines are the rolling average force across the 50 frames. The number of contacts between Izumo1 and Juno as a function of time can be found in **fig. S9B** and the details of the corresponding interactions are shown in **fig. S8**. Preview of the unbinding process can be observed in **Supplementary Movie 2**.

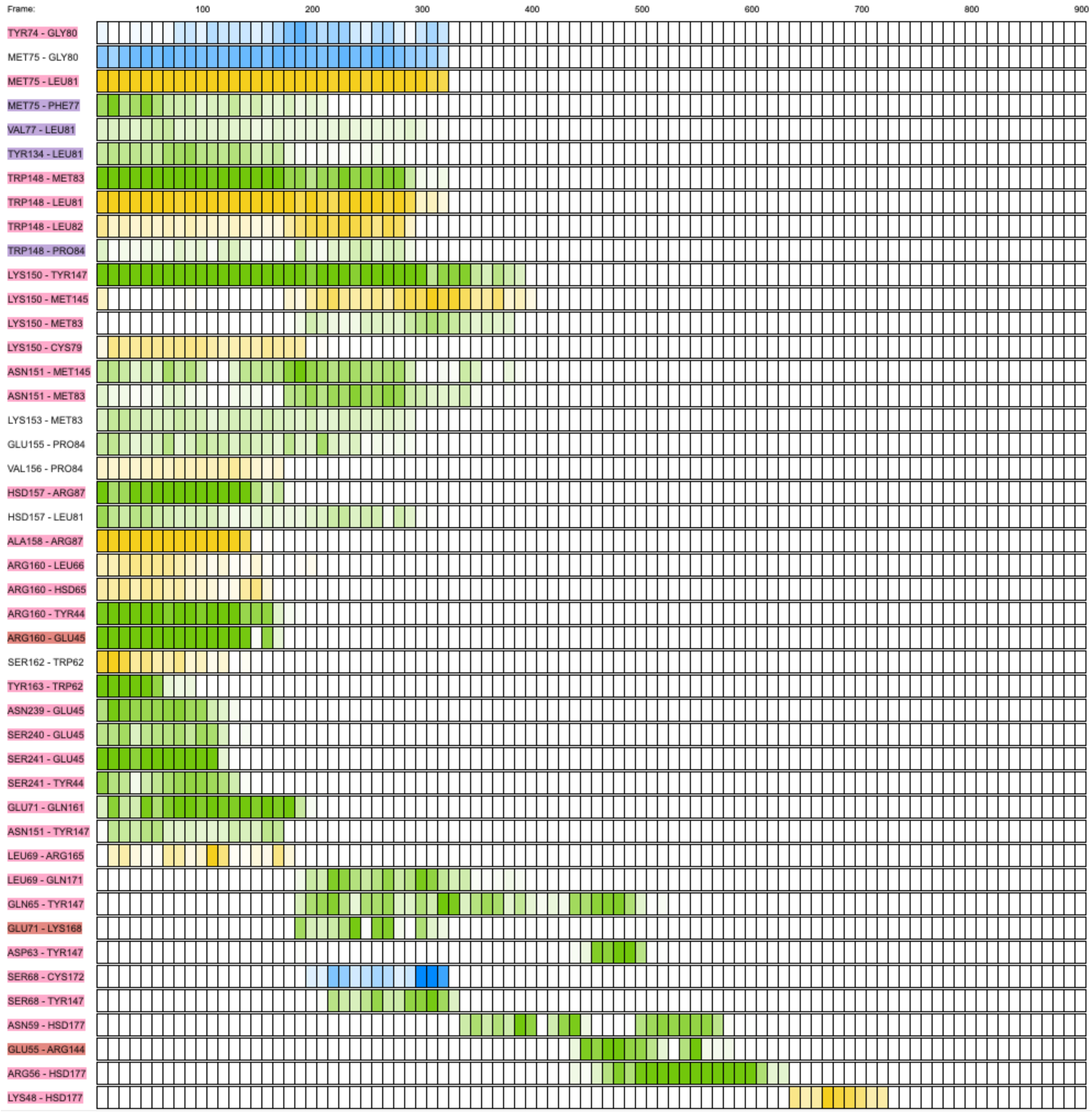
Frame:
100
200
300
400
500
600
700
800
900
TYR74 - GLY80
MET75 - GLY80
MET75 - LEU81
MET75 - PHE77
VAL77 - LEU81
TYR134 - LEU81
TRP148 - MET83
TRP148 - LEU81
TRP148 - LEU82
TRP148 - PRO84
LYS150 - TYR147
LYS150 - MET145
LYS150 - MET83
LYS150 - CYS79
ASN151 - MET145
ASN151 - MET83
LYS153 - MET83
GLU155 - PRO84
VAL156 - PRO84
HSD157 - ARG87
HSD157 - LEU81
ALA158 - ARG87
ARG160 - LEU66
ARG160 - HSD65
ARG160 - TYR44
ARG160 - GLU45
SER162 - TRP62
TYR163 - TRP62
ASN239 - GLU45
SER240 - GLU45
SER241 - GLU45
SER241 - TYR44
GLU71 - GLN161
ASN151 - TYR147
LEU69 - ARG165
LEU69 - GLN171
GLN65 - TYR147
GLU71 - LYS168
ASP63 - TYR147
SER68 - CYS172
SER68 - TYR147
ASN59 - HSD177
GLU55 - ARG144
ARG56 - HSD177
LYS48 - HSD177

**Figure S8: Exemplary time series of the non-covalent interactions as observed in the three-state unbinding pathway of the wild-type Izumo1:Juno complex in SMD simulations.** In this trajectory the strongest late stage interactions were mediated by H177 of Juno forming three hydrogen bonds with N59, R56 and K48 of Izumo1**.** Total unbinding time was prolonged to over 70 ns. The strongest late stage interactions are mediated by H177 of Juno**. Supplementary Movie 2** shows the preview of the corresponding simulation. The first residue in the label belongs to Izumo1, and the second residue belongs to Juno. The color of the label is determined by interaction type and geometry: red, salt bridge; blue, hydrophobic; and pink, hydrogen bond. Each box corresponds to the equal trajectory segment, their color signifies interaction type and the intensity shows the strength of the interaction: green, side chain/side chain; yellow, side chain/backbone; blue, backbone/backbone. PyContact (*62*) was used to extract this data with a filter of mean score value > 0.25 for trajectory of the total simulation time of 90 ns (900 frames extracted).

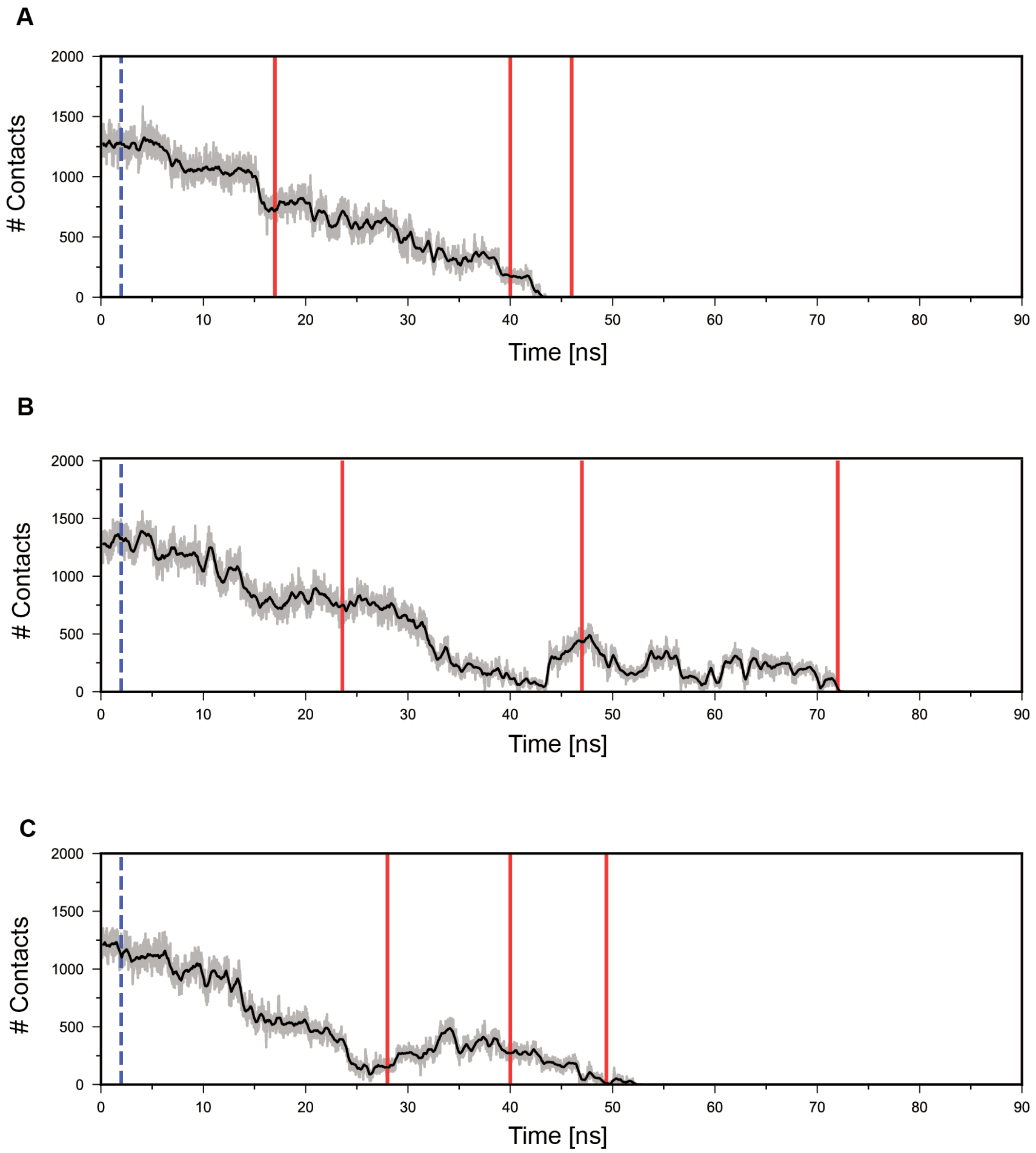
A
B
C
### Contacts
Time [ns]
2000
1500
1000
500
0
0
10
20
30
40
50
60
70
80
90

**Figure S9: Number of contacts between Izumo1 and Juno traced as a function of time for representative unbinding trajectories of each type. (A)** During two-state unbinding, the number of contacts between Izumo1 and Juno steadily decreases over time. **(B)** During three-state unbinding, the number of contacts increases during the first 2 ns. This is followed by an extended transition phase from ~15-30 ns, during which the number of contacts between Izumo1 and Juno plateaus at ~750. During the detachment of the second binding site at ~43 ns, the rotation of Izumo1 and Juno allows new contacts amongst residues that were previously unconnected. **(C)** Number of contacts for the three-state unbinding subtype is consistent with the number of contacts during three-state unbinding shown in panel **B**. While the initial binding interface is lost earlier at ~25 ns, new contacts are formed and increase from 0 to ~500. Gray traces represent number of contacts for individual frames and black lines are the rolling average number of contacts across the 50 frames. Red lines mark frames as shown in **Fig. 2**. Dashed blue line marks frame from which contact surface area was extracted to represent the loaded state.

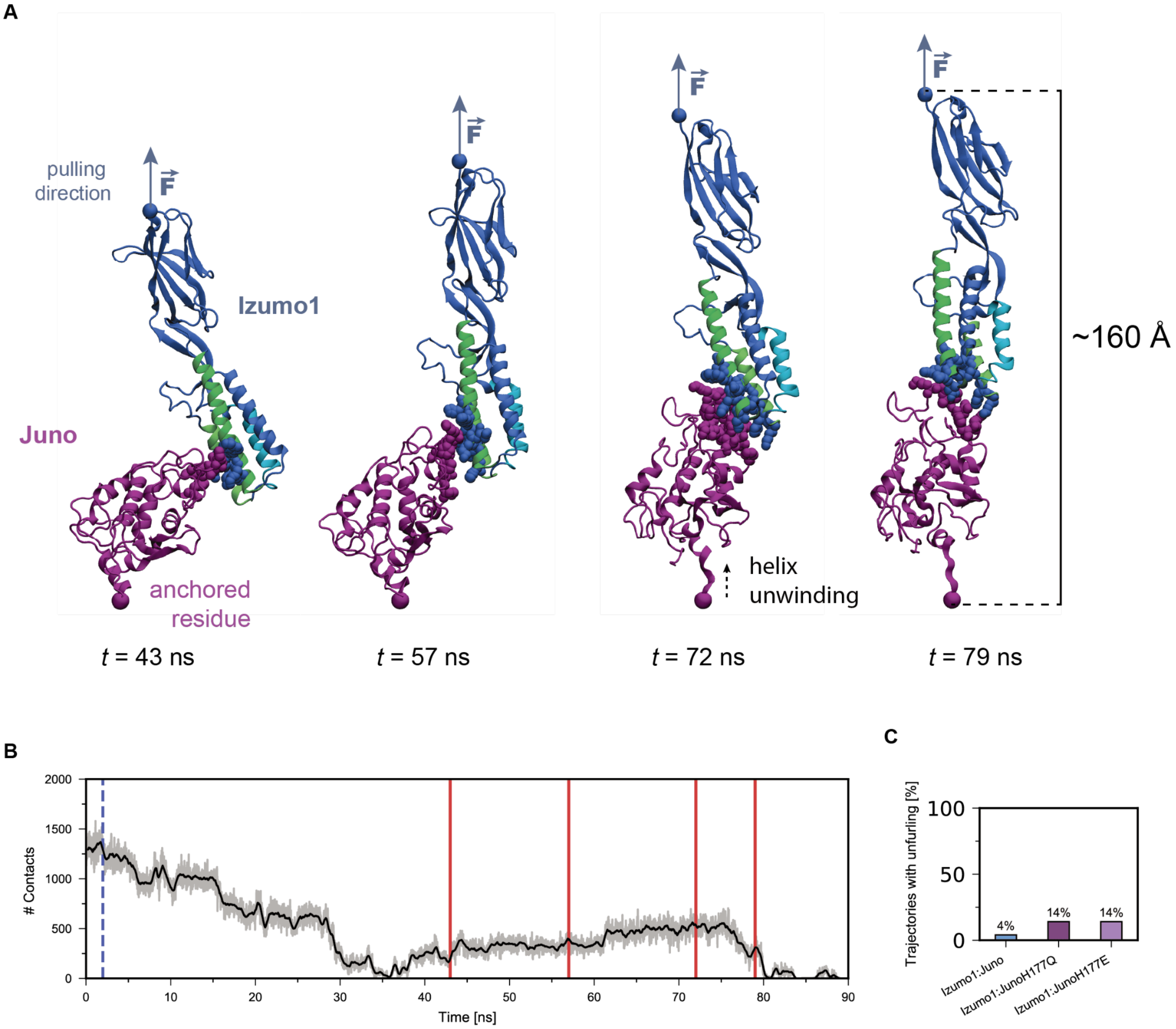


**Figure S10: The unwinding of the C-terminal helix of Juno as observed in some SMD simulations of Izumo1:Juno complex under constant velocity pulling. (A)** Representative snapshots of structural deformations in the secondary structure of Juno during unbinding. As the known binding interface is lost, new interactions are gradually formed, prolonging the interaction between Izumo1 and Juno until the C-terminal of Juno unwinds. **(B)** Number of contacts between Izumo1 and Juno traced as a function of time. Gray traces represent number of contacts for individual frames and the black line represents the rolling average number of contacts across the 50 frames. Red lines mark frames shown in **A**. **(C)** Percentage of trajectories that show unwinding of the Juno C-terminal α-helix in wild-type Izumo1:Juno. Izumo1:JunoH177Q, and Izumo1:JunoH177E.

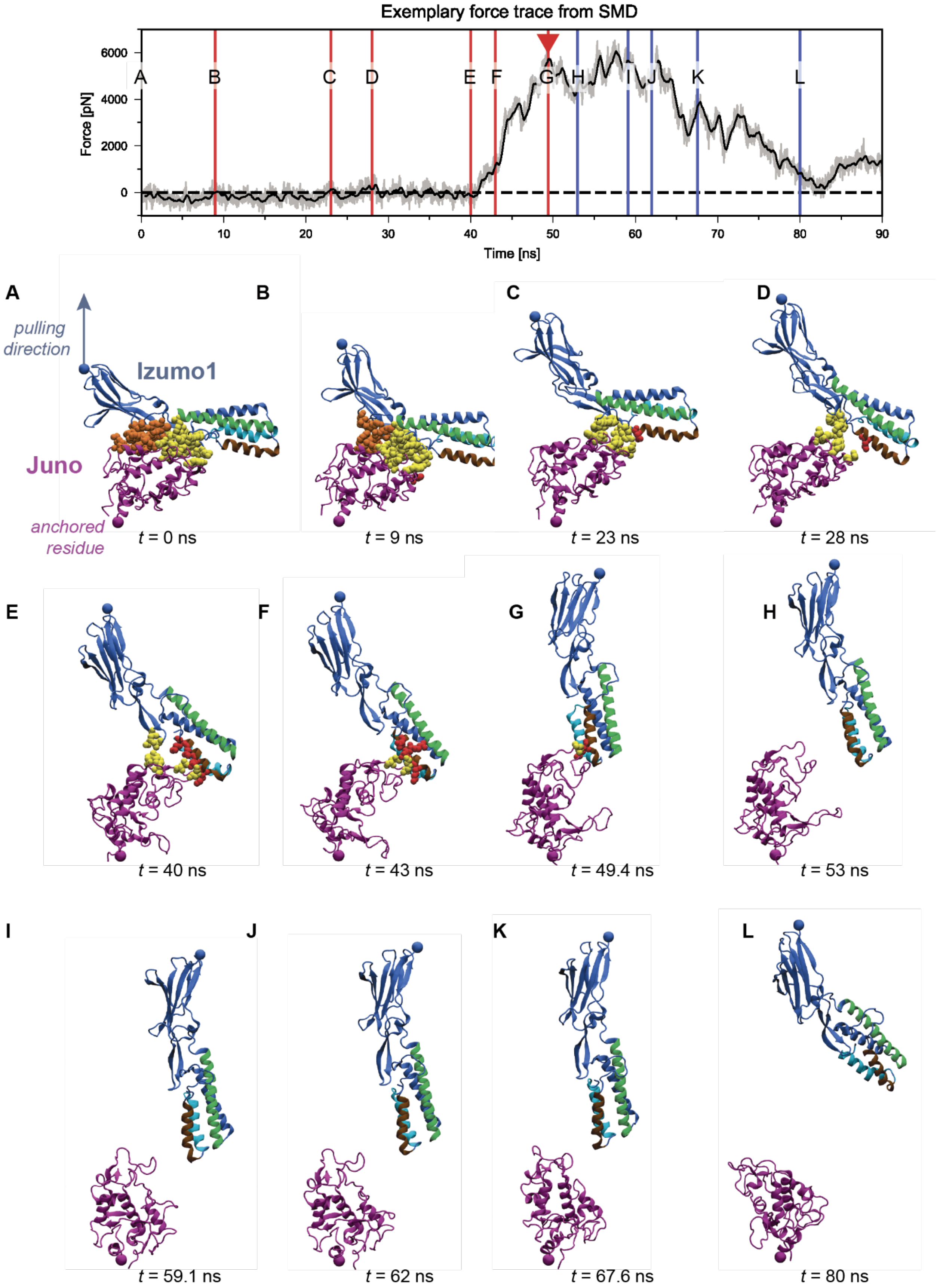

Exemplary force trace from SMD
Force [pN]
6000
4000
2000
0
Time [ns]
0
10
20
30
40
50
60
70
80
90
A
B
C
D
E
F
G
H
I
J
K
L
pulling direction
Izumo1
Juno
anchored residue
t = 0 ns
t = 9 ns
t = 23 ns
t = 28 ns
t = 40 ns
t = 43 ns
t = 49.4 ns
t = 53 ns
t = 59.1 ns
t = 62 ns
t = 67.6 ns
t = 80 ns

**Figure S11: Force *vs.* time trace with structural snapshots representative of the three-state wild-type Izumo1:Juno unbinding subtype observed in SMD simulations.** Different residues are involved in the Cluster 3 (red) formation in three-state subtype trajectories when compared to the majority of the three-state unbinding trajectories. Letters and lines mark timepoints at which snapshots were taken and the red triangle marks the first rupture event. Gray traces represent force for individual frames and black lines are the rolling average force across the 50 frames. The number of contacts between Izumo1 and Juno as a function of time can be found on **fig. S9C**. Preview of the unbinding process can be observed in **Supplementary Movie 3**.

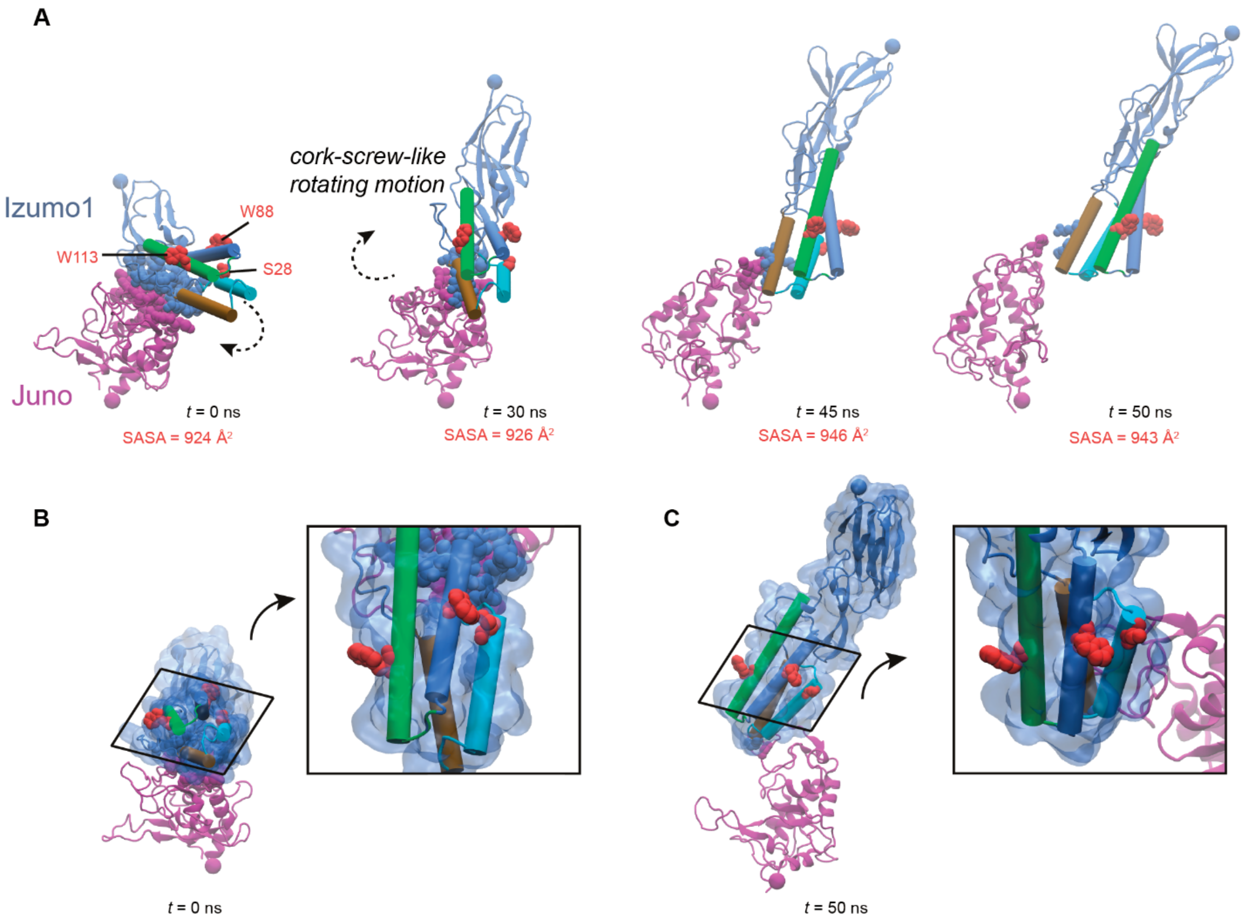


**Figure S12: Reorganization of membrane fusion-essential residues in the three-state wild-type Izumo1:Juno unbinding subtype observed during SMD simulations**. **(A)** Upwards pulling of Izumo1 results in a cork-screw-like rotation of Izumo1 with respect to Juno. During this process, the membrane fusion-essential residues; W113, W88 and S28 (*11*); partially buried in the 4-helix bundle of Izumo1 (shown in red), become increasingly solvent exposed (SASA calculated using water probe radius of 1.4 Å reported for each frame) and aligned across same interaction plane. Such conformational changes would allow for larger hydrophobic interaction paths between Izumo1 and the egg cell membrane, supporting the membrane fusion activity of Izumo1 expressing cells. **(B)** Initial orientation of W113, W88 and S28 in the 4-helix bundle of Izumo1. **(C)** Final orientation of W113, W88 and S28, now aligned to form a large hydrophobic patch on the Izumo1 surface. See also **Supplementary Movie S4**.

**A**

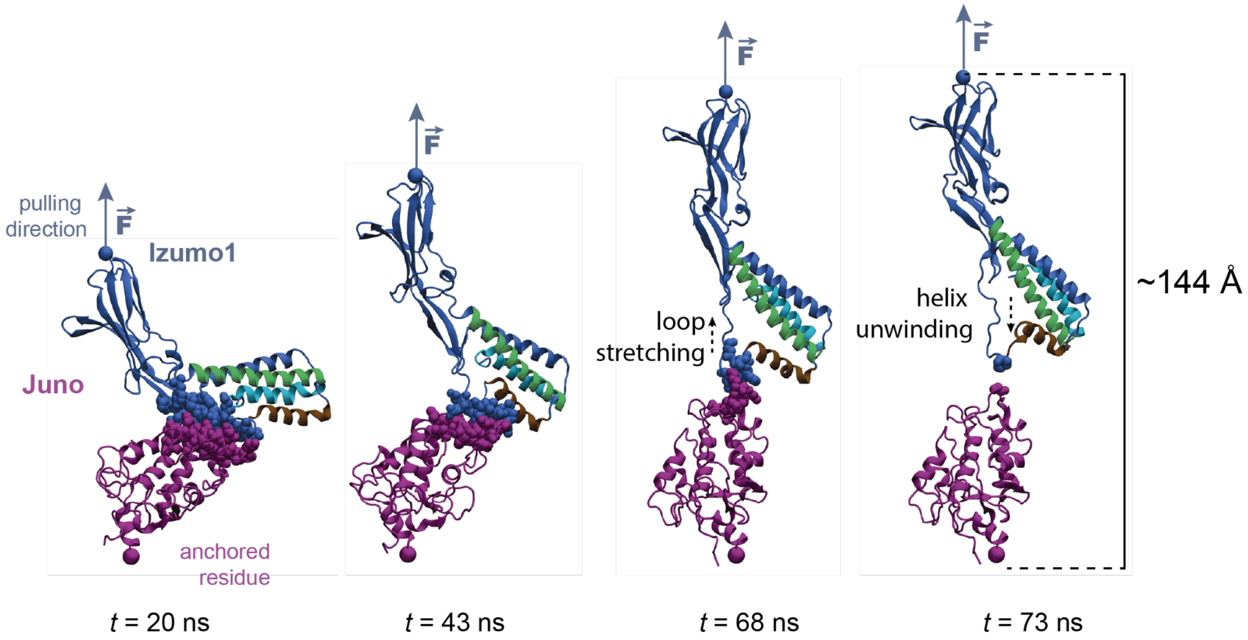


**B**

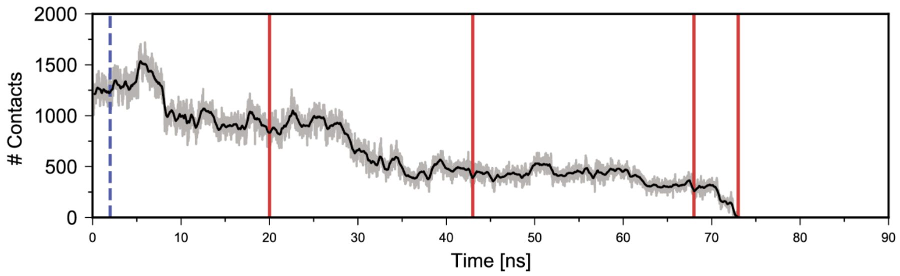


**Figure S13: The partial unwinding of Izumo1 4-helix bundle stochastically observed in SMD simulations of the Izumo1:JunoH177Q complex. (A)** Representative snapshots of structural deformations of the Izumo1 secondary structure during unbinding. The Cluster 2 interactions formed a strong association between Juno and the deformed region of Izumo1 around the residue E71. Where SMD simulations are performed at 0.001 Å·ps$^{-1}$, it is possible that the 4-helix bundle of Izumo1 undergoes further rearrangements at physiological speeds. **(B)** Number of contacts between Izumo1 and Juno traced as a function of time. Gray traces represent number of contacts for individual frames and the black line represents the rolling average number of contacts across the selected frames. Red lines mark frames shown in **A**.

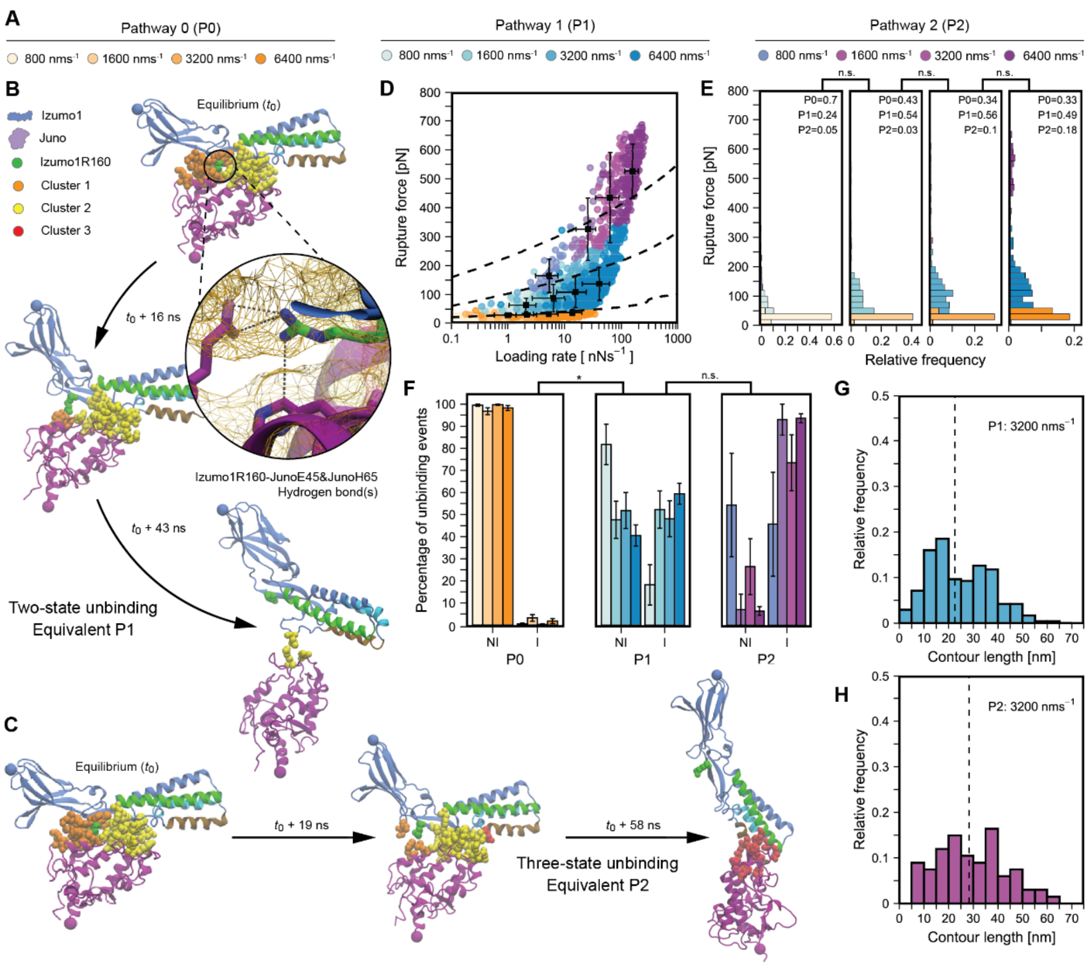


**Figure S14: Supplementary constant speed AFM-SMFS data for Izumo1R160A:Juno.** (**A**) Legend for P0, P1, and P2 unbinding events at each constant pulling speed. (**B**) Equilibrated starting structure ($t = t_0$) for Izumo1:Juno showing Izumo1R160 (green) and amino acids clusters 1 (orange), 2 (yellow) and 3 (red). Izumo1:Juno complex unbinding though the two-state pathway. (**C**) Equilibrated starting structure ($t = t_0$), color coded as per panel **B**, with Izumo1:Juno unbinding through the third-state pathway. (**D**) Izumo1R160A:Juno rupture force *vs.* loading rates (median ± MAD). Dashed lines show the DHS model fits (Eq. (6), Eq. (7), **table S1**). (**E**) Izumo1R160A:Juno rupture force histograms (bins = 20 pN). (**F**) Direct Izumo1R160A:Juno unbinding (NI) compared to Izumo1R160A:Juno unbinding through intermediate folded states (I) (mean ± SEM). (**G**) Contour length distributions for Izumo1R160A:Juno rearrangements occurring prior to rupture through P1 at 3200 nms$^{-1}$ ($n$ = 238). (**H**) Contour length distributions for Izumo1R160A:Juno rearrangements occurring prior to rupture through P2 at 3200 nms$^{-1}$ ($n$ = 67). Data in **D**, **E**, and **F** represent four pooled independent experiments ($n$ = 691-947) with significance (*$p < 0.05$) determined using Kurskal-Wallis tests followed by pairwise Wilcoxon rank-sum tests.

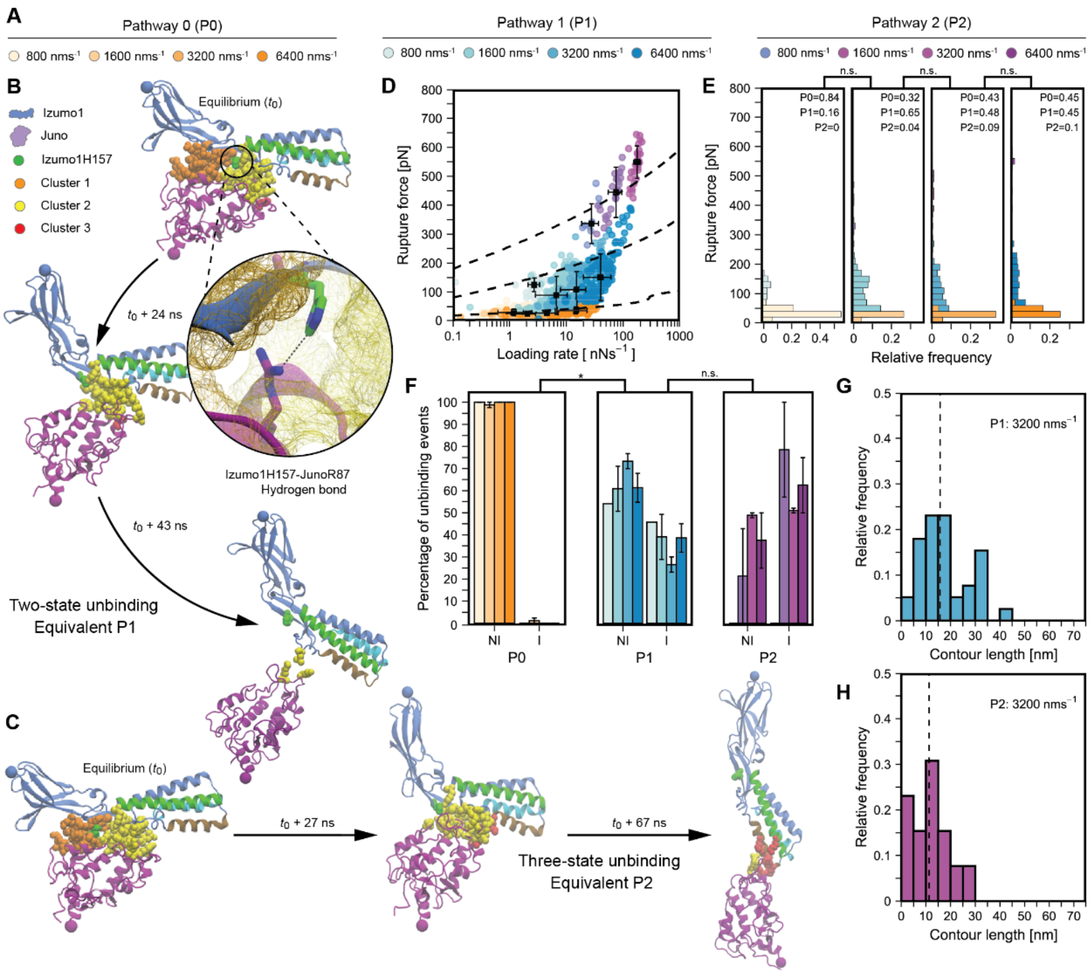


**Figure S15: Supplementary constant speed AFM-SMFS data for Izumo1H157A:Juno.** (**A**) Legend for P0, P1, and P2 unbinding events at each constant pulling speed. (**B**) Equilibrated starting structure ($t = t_0$) for Izumo1:Juno showing Izumo1H157 (green) and amino acids clusters 1 (orange), 2 (yellow) and 3 (red). Izumo1:Juno complex unbinding though the two-state pathway. (**C**) Equilibrated starting structure ($t = t_0$), color coded as per panel **B**, with Izumo1:Juno unbinding through the three-state pathway. (**D**) Izumo1H157A:Juno rupture force *vs.* loading rates (median ± MAD). Dashed lines show the DHS model fits (Eq. (6), Eq. (7), **table S1**). (**E**) Izumo1H157A:Juno rupture force histograms (bins = 20 pN). (**F**) Direct Izumo1H157A:Juno unbinding (NI) compared to Izumo1H157A:Juno unbinding through intermediate folded states (I) (mean ± SEM). (**G**) Contour length distributions for Izumo1H157A:Juno rearrangements occurring prior to rupture through P1 at 3200 nms$^{-1}$ ($n$ = 39). (**H**) Contour length distributions for Izumo1H157A:Juno rearrangements occurring prior to rupture through P2 at 3200 nms$^{-1}$ ($n$ = 13). Data in **D**, **E**, and **F** represent four pooled independent experiments ($n$ = 153-294) with significance (*$p$ < 0.05) determined using Kurskal-Wallis tests followed by pairwise Wilcoxon rank-sum tests.

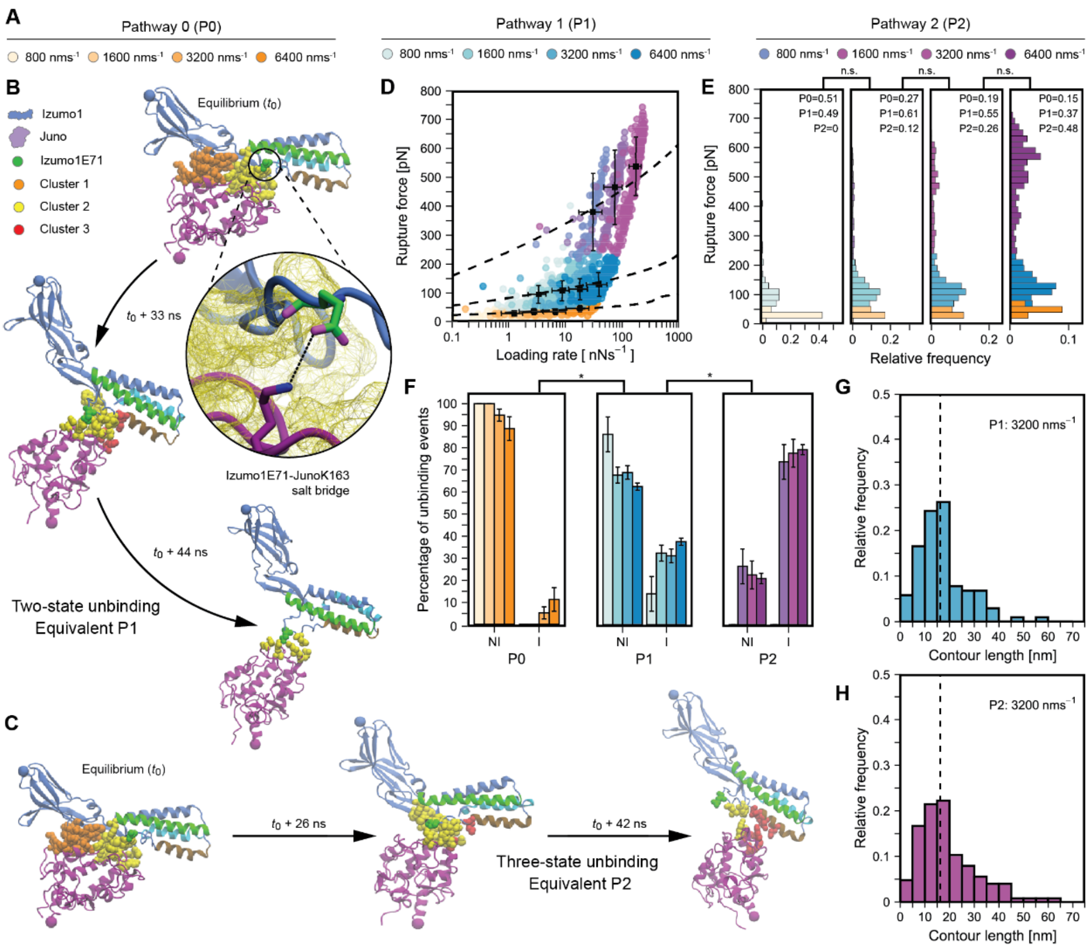


**Figure S16: Supplementary constant speed AFM-SMFS data for Izumo1E71A:Juno.** (**A**) Legend for P0, P1, and P2 unbinding events at each constant pulling speed. (**B**) Equilibrated starting structure ($t = t_0$) for Izumo1:Juno showing Izumo1E71 (green) and amino acids clusters 1 (orange), 2 (yellow) and 3 (red). Izumo1:Juno complex unbinding though the two-state pathway. (**C**) Equilibrated starting structure ($t = t_0$), color coded as per panel **B**, with Izumo1:Juno unbinding through the three-state pathway. (**D**) Izumo1E71A:Juno rupture force *vs.* loading rates (median ± MAD). Dashed lines show the DHS model fits (Eq. (6), Eq. (7), **table S1**). (**E**) Izumo1E71A:Juno rupture force histograms (bins = 20 pN). (**F**) Direct Izumo1E71A:Juno unbinding (NI) compared to Izumo1E71A:Juno unbinding through intermediate folded states (I) (mean ± SEM). (**G**) Contour length distributions for Izumo1E71A:Juno rearrangements occurring prior to rupture through P1 at 3200 $nms^{-1}$ ($n$ = 103). (**H**) Contour length distributions for Izumo1E71A:Juno rearrangements occurring prior to rupture through P2 at 3200 $nms^{-1}$ ($n$ = 126). Data in **D**, **E**, and **F** represent four pooled independent experiments ($n$ = 420-617) with significance (*$p < 0.05$) determined using Kurskal-Wallis tests followed by pairwise Wilcoxon rank-sum tests.

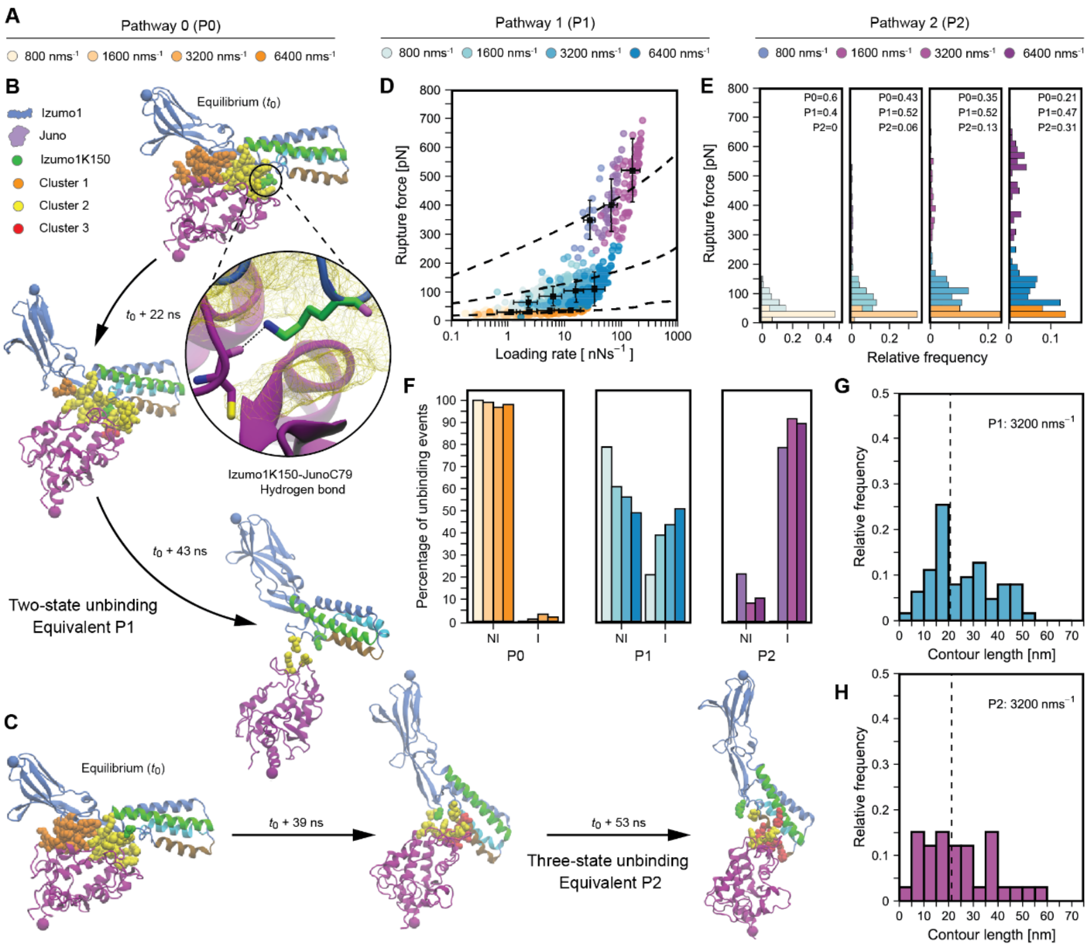


**Figure S17: Supplementary constant speed AFM-SMFS data for Izumo1K150:Juno.** (**A**) Legend for P0, P1, and P2 unbinding events at each constant pulling speed. (**B**) Equilibrated starting structure ($t = t_0$) for Izumo1:Juno showing Izumo1K150 (green) and amino acids clusters 1 (orange), 2 (yellow) and 3 (red). Izumo1:Juno complex unbinding though the two-state pathway. (**C**) Equilibrated starting structure ($t = t_0$), color coded as per panel **B**, with Izumo1:Juno unbinding through the three-state pathway. (**D**) Izumo1K150A:Juno rupture force *vs.* loading rates (median ± MAD). Dashed lines show the DHS model fits (Eq. (6), Eq. (7), **table S1**). (**E**) Izumo1K150A:Juno rupture force histograms (bins = 20 pN). (**F**) Direct Izumo1K150A:Juno unbinding (NI) compared to Izumo1K150A:Juno unbinding through intermediate folded states (I) (mean ± SEM). (**G**) Contour length distributions for Izumo1K150A:Juno rearrangements occurring prior to rupture through P1 at 3200 nms$^{-1}$ ($n$ = 63). (**H**) Contour length distributions for Izumo1K150A:Juno rearrangements occurring prior to rupture through P2 at 3200 nms$^{-1}$ ($n$ =33). Data in **D**, **E**, and **F** represent a single experiment stable for over 12 h ($n$ = 225-275). Hypothesis testing was not performed for this variant due to long-term instability of replicates.

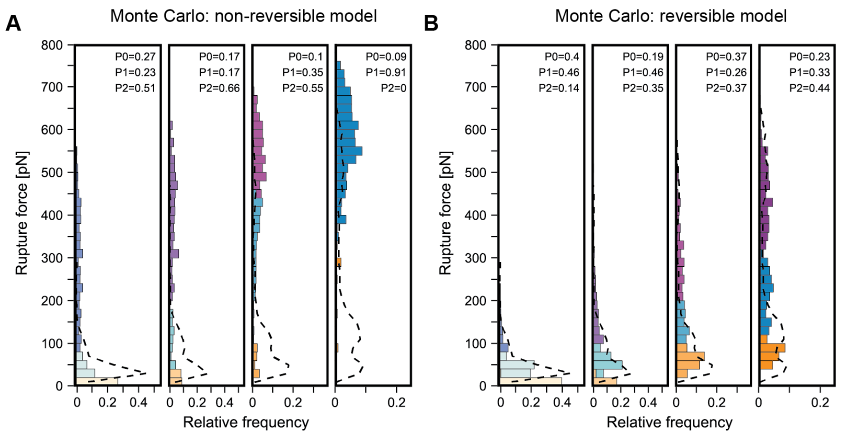


**Figure S18: Constant speed Monte Carlo simulations of multi-pathway Izumo1:Juno unbinding through intermediate states.** (**A**) Non-reversible transitions between intermediate states that give rise to P0, P1, and P2 unbinding pathways. Residual sum of squares fit to the binned experimental and Monte Carlo simulated rupture force distributions (bins = 20 pN) is 0.189. (**B**) Reversible transitions between intermediate states that give rise to P0, P1, and P2 unbinding pathways. Residual sum of squares fit to the binned experimental and Monte Carlo simulated rupture force distributions (bins = 20 pN) is 0.173. For all constant speed Monte Carlo simulations, time step = 1000, *n* = 500, and dashed lines represent a smoothed approximation of the experimental rupture force histogram distribution (fitted using a cubic spline).

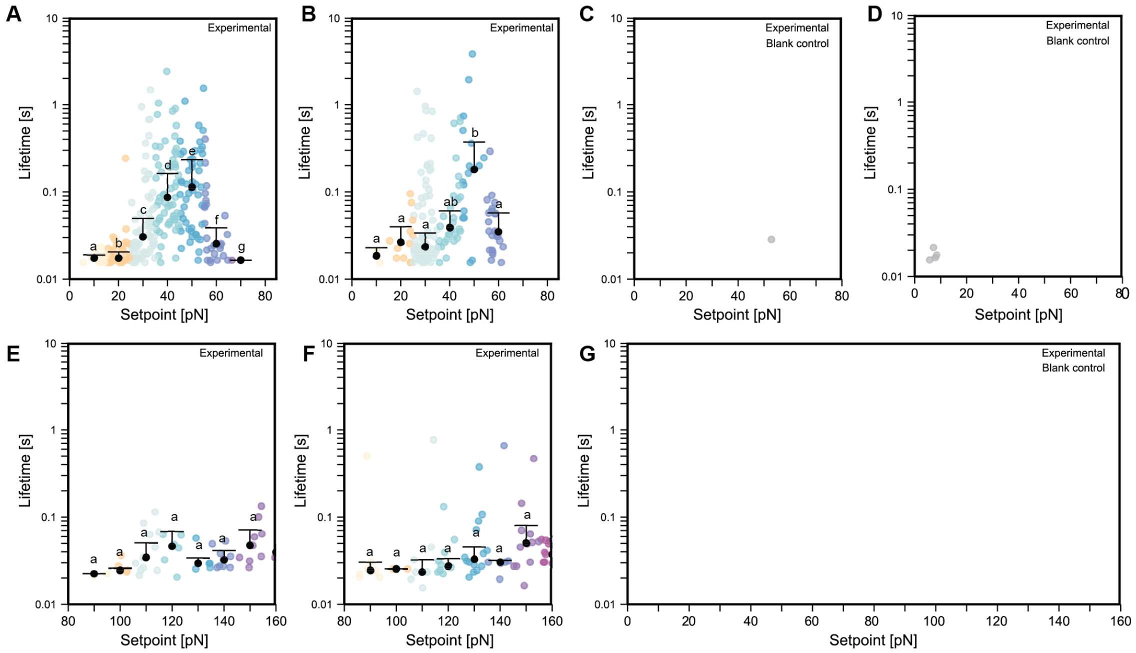


**Figure S19: Force clamp AFM-SMFS Izumo1:Juno biological replicates and blank controls.** (**A-B**) Biological replicate 2 and 3 for Izumo1:Juno force clamp AFM-SMFS performed in the 10-80 pN setpoint range. (**C-D**) Representative blank controls for Izumo1:Juno force clamp AFM-SMFS performed in the 10-80 pN setpoint range. (**E**) Biological replicate 1 for Izumo1:Juno force clamp AFM-SMFS performed in the 80-160 pN setpoint range. (**F**) Biological replicate 2 for Izumo1:Juno force clamp AFM-SMFS performed in the 80-160 pN setpoint range. (**G**) Representative blank controls for Izumo1:Juno force clamp AFM-SMFS performed in the 10-160 pN setpoint range. No force clamping events were observed that passed the automated filtering. Colored dots represent individual bond lifetime measurements. Dark gray dots represent non-specific force clamping events. Black dots represent median bond lifetime (± MAD) across 10 pN bins. Shared letters indicate the absence of statistically significant differences ($p > 0.05$) using Kruskal–Wallis tests followed by Dunn's multiple-comparison tests. Blank control consists of the Izumo1 functionalized tips, with activity established using constant speed SMFS, probed against cover glasses with only the AFM-handle functionalized (*i.e*. no Juno) directly following their measurement.

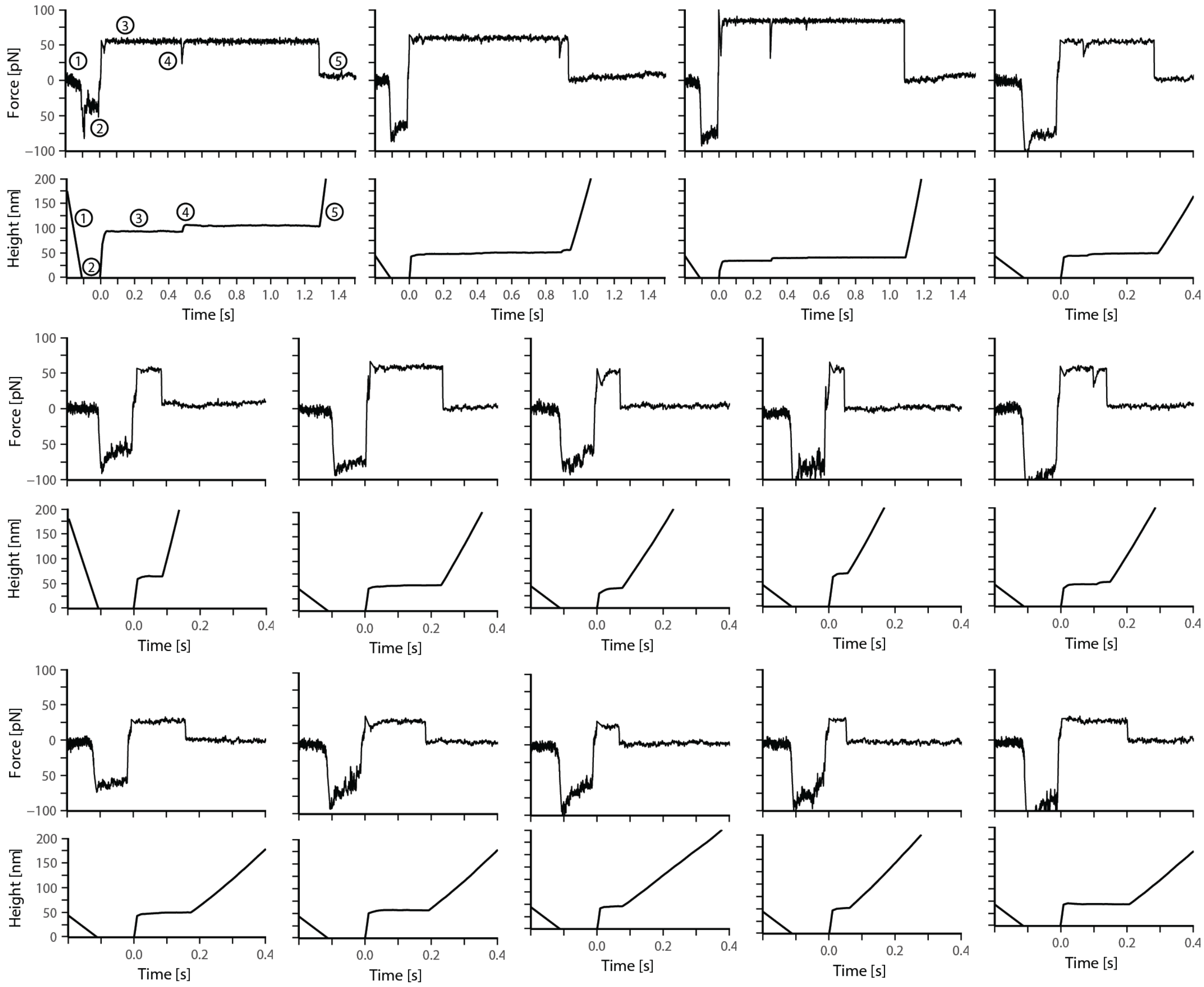


**Figure S20: Force clamp AFM-SMFS representative curves.** Top panels show force *vs.* time curves. Bottom panels show height *vs.* time curves. (1) Shows the approach of the cantilever to the surface. (2) Shows the pause of the cantilever tip at the surface for 0.2 s. (3) Shows the retraction of the cantilever to the target setpoint. (4) Where present in force *vs.* time and height *vs.* time curves this feature shows the presence of a contour length change in the Izumo1:Juno complex. (5) Shows complex rupture and resumes retraction of the cantilever.

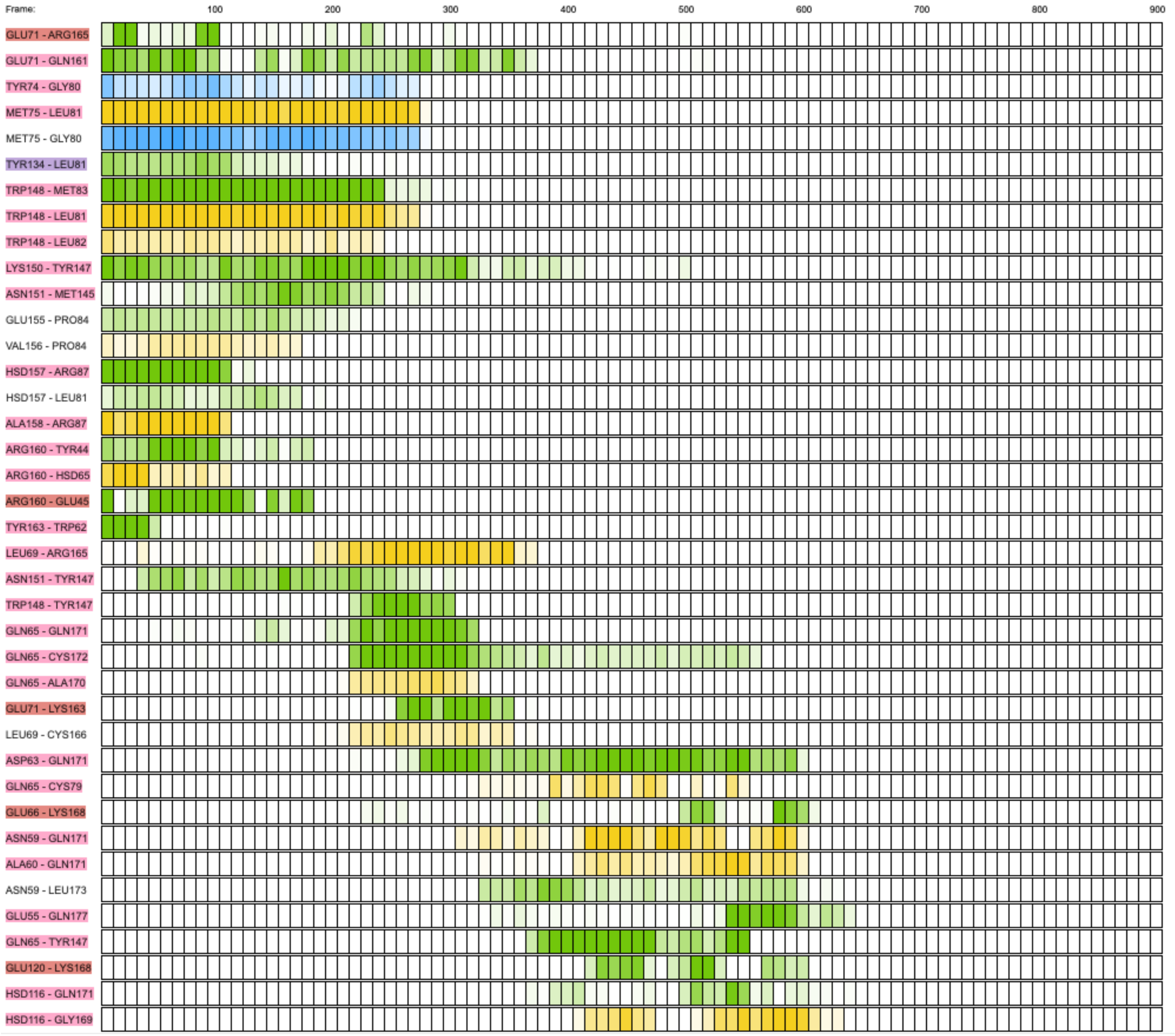


**Figure S21: Exemplary time series of the non-covalent interactions as observed in the three-state unbinding pathway of the Izumo1:JunoH177Q mutant complex in SMD simulations.** In simulations of Izumo1:JunoH177Q, fewer trajectories showed entry into the third unbinding pathway. Overall, the residues on Izumo1 that formed prolonged interactions with JunoH177Q were similar to those interacting with Juno in the wild-type Izumo1:Juno complex. However, where Juno H177 was unavailable to form strong interactions, Juno Q177 formed only one hydrogen bond with Izumo1 E55. **Supplementary Movie 5** shows the preview of the corresponding simulation. The first residue in the label belongs to Izumo1, and the second residue belongs to Juno. The color of the label is determined by interaction type and geometry: red, salt bridge; blue, hydrophobic; and pink, hydrogen bond. Each box corresponds to the equal trajectory segment, their color signifies interaction type and the intensity shows the strength of the interaction: green, side chain/side chain; yellow, side chain/backbone; blue, backbone/backbone. PyContact (*62*) was used to extract this data with a filter of mean score value > 0.25 for trajectory of the total simulation time of 90 ns (900 frames extracted).

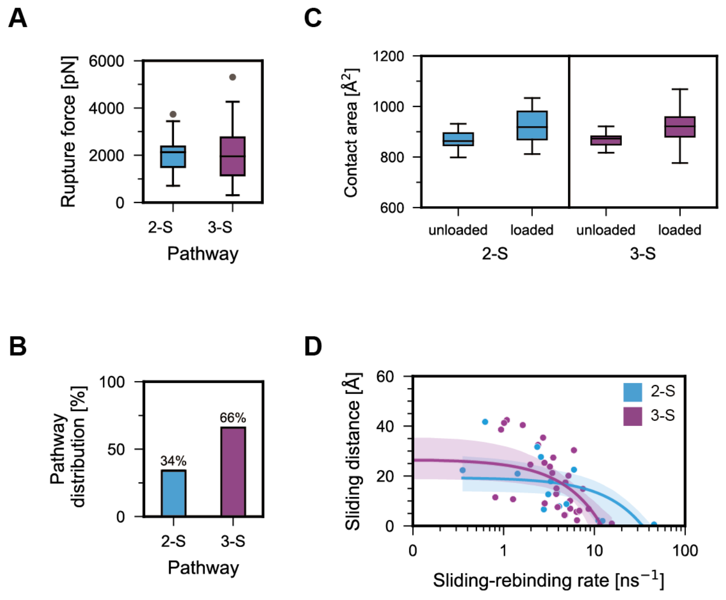


**Figure S22: SMD simulations of Izumo1:JunoH177E unbinding.** (**A**) Peak forces extracted from two-state (2-S) and three-state (3-S) SMD simulations of Izumo1:JunoH177E complexes (median ± MAD). (**B**) Two-state and three-state unbinding pathway distribution for Izumo1:H177E across 50 SMD trajectories. (**C**) Observed increases in contact area between Izumo1 and JunoH177E in unloaded ($t$ = 0 ns) and loaded ($t$ = 0 ns) complexes for the two-state and three-state pathways. (**D**) Sliding-rebinding distance *vs.* sliding-rebinding rate for the force-dependent rearrangement of Izumo1:JunoH177E complexes in the two-state and three-state pathways. Boxplots show the median, interquartile range (IQR), and whiskers extending to 1.5 times the IQR. Points beyond the whiskers are outliers.

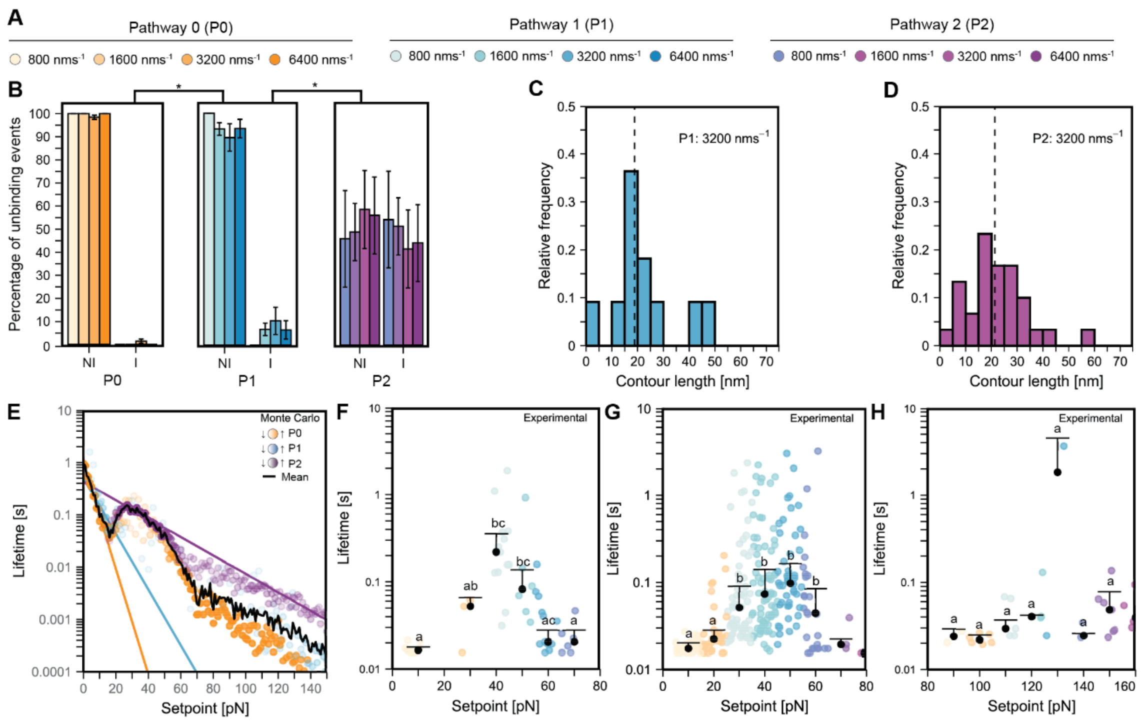

**Figure S23: Monte Carlo force clamp and supplemental AFM-SMFS data for Izumo1:JunoH177Q.** (**A**) Legend for P0, P1, and P2 unbinding events at each constant pulling speed. (**B**) Direct Izumo1:JunoH177Q unbinding (NI) compared to unbinding through intermediate folded states (I) (mean ± SEM) with significance (*$p < 0.05$) determined using Kurskal-Wallis tests followed by pairwise Wilcoxon rank-sum tests. (**C**) Contour length distributions for rearrangements during P1 at 3200 nms$^{-1}$ ($n = 11$). (**D**) Contour length distributions for rearrangements during P2 at 3200 nms$^{-1}$ ($n = 30$). (**E**) Monte Carlo simulated force clamp AFM-SMFS for Izumo1:JunoH177Q. Colored dots represent mean bond lifetime for each unbinding pathway at each setpoint with intensity representing the relative abundance of an unbinding pathway. Colored lines represent lifetimes calculated directly from BE-derived kinetic parameters (**table S1,** Eq. (11)). Black line represents the mean bond lifetime across all three pathways. (**F-G**) Biological replicates for force clamp AFM-SMFS performed between 10-80 pN. (**H**) Representative force clamp AFM-SMFS experiment performed between 90-160 pN. Colored dots represent individual bond lifetime measurements. Black dots represent median bond lifetime (± MAD) across 10 pN bins. Shared letters indicate the absence of significance ($p > 0.05$) determined using Kruskal–Wallis tests followed by Dunn's multiple-comparison tests.

**Table S1: Non-equilibrium kinetic parameters determined using constant speed AFM-SMFS.** Error reported as standard error.

| Protein Complex | Pathway | BE model | | DHS model ($\upsilon$ = 1/2) | | | DHS model ($\upsilon$ = 2/3) | | |
|---|---|---|---|---|---|---|---|---|---|
| | | $k_0$ [$s^{-1}$] | $\Delta x$ [nm] | $k_0$ [$s^{-1}$] | $\Delta x$ [nm] | $\Delta G^{\ddagger}$ [$k_BT$] | $k_0$ [$s^{-1}$] | $\Delta x$ [nm] | $\Delta G^{\ddagger}$ [$k_BT$] |
| Izumo1:Juno | 0 | 1.28± 2.56 | 0.779± 0.264 | 1.46± 0.95 | 0.681± 0.191 | 19.52± 4.38 | 0.92± 0.68 | 0.405± 0.136 | 15.14± 4.29 |
| | 1 | 3.62± 3.22 | 0.240± 0.039 | 4.66± 0.25 | 0.072± 0.006 | 13.89± 0.96 | 3.78± 0.24 | 0.032± 0.004 | 9.49± 2.22 |
| | 2 | 4.20± 0.37 | 0.046± 0.001 | 3.41± 0.19 | 0.039± 0.001 | 16.12± 0.37 | 2.36± 0.15 | 0.022± 0.001 | 11.75± 0.35 |
| Izumo1E71A:Juno | 0 | 1.27± 1.79 | 0.726± 0.178 | 2.43± 0.86 | 0.447± 0.082 | 17.28± 2.43 | 1.69± 0.64 | 0.252± 0.056 | 12.70± 2.26 |
| | 1 | 4.10± 4.49 | 0.244± 0.047 | 3.34± 0.46 | 0.116± 0.012 | 15.13± 1.00 | 2.49± 0.38 | 0.061± 0.008 | 10.44± 0.92 |
| | 2 | 4.30± 0.67 | 0.045± 0.002 | 3.88± 0.18 | 0.035± 0.001 | 15.94± 0.38 | 2.76± 0.14 | 0.018± 0.001 | 11.35± 0.34 |
| Izumo1K150A:Juno | 0 | 1.24± 1.98 | 0.809± 0.226 | 1.84± 0.75 | 0.663± 0.123 | 18.07± 2.75 | 1.20± 0.52 | 0.387± 0.085 | 13.72± 2.53 |
| | 1 | 1.39± 0.57 | 0.227± 0.022 | 3.78± 0.58 | 0.097± 0.014 | 14.78± 1.23 | 2.00± 0.51 | 0.049± 0.010 | 9.78± 1.08 |
| | 2 | 2.55± 2.11 | 0.041± 0.01 | 3.66± 0.26 | 0.038± 0.002 | 16.16± 0.70 | 2.65± 0.27 | 0.020± 0.002 | 12.14± 1.61 |
| Izumo1H157A:Juno | 0 | 1.07± 3.5 | 0.937± 0.525 | 3.46± 0.68 | 0.401± 0.047 | 14.69± 1.53 | 2.57± 0.51 | 0.221± 0.029 | 10.07± 1.31 |
| | 1 | 4.34± 20.09 | 0.23± 0.192 | 4.01± 0.34 | 0.063± 0.006 | 14.66± 0.90 | 3.34± 0.32 | 0.030± 0.004 | 9.51± 1.08 |
| | 2 | 2.14± 0.20 | 0.036± 0.001 | 3.43± 0.23 | 0.040± 0.002 | 17.44± 0.62 | 2.30± 0.18 | 0.022± 0.001 | 12.85± 0.62 |
| Izumo1R160A:Juno | 0 | 1.14± 1.68 | 0.875± 0.222 | 3.06± 0.83 | 0.376± 0.074 | 14.74± 2.15 | 2.28± 0.66 | 0.198± 0.048 | 10.10± 1.94 |
| | 1 | 0.91± 0.21 | 0.169± 0.011 | 4.57± 0.32 | 0.067± 0.006 | 13.53± 0.63 | 3.70± 0.30 | 0.032± 0.004 | 8.64± 0.59 |
| | 2 | 2.54± 0.13 | 0.039± 0.001 | 3.03± 0.20 | 0.043± 0.002 | 16.72± 0.41 | 2.06± 0.17 | 0.024± 0.001 | 12.42± 0.39 |

| | | | | | | | | | |
|---|---|---|---|---|---|---|---|---|---|
| Izumo1:JunoH177Q* | 0 | 1.05± 1.66 | 0.956± 0.261 | 1.98± 0.91 | 0.752± 0.186 | 17.31± 3.15 | 1.30± 0.65 | 0.434± 0.127 | 13.01± 2.95 |
| | 1 | 1.98± 8.10 | 0.506± 0.361 | 2.21± 1.08 | 0.329± 0.076 | 17.77± 3.31 | 1.59± 0.82 | 0.183± 0.051 | 12.95± 3.06 |
| | 2 | 2.34± 3.46 | 0.167± 0.051 | 5.03± 0.28 | 0.048± 0.006 | 10.89± 0.62 | 4.18± 0.31 | 0.020± 0.004 | 6.12± 0.60 |

***** During global fitting of the DHS model, forces greater than 400 pN were excluded as outliers that impacted convergence of the fitting algorithm.

Table S2: Summary of the SMD simulations performed.

| Protein complex | Number of trajectories | Time [ns] |
|---|---|---|
| Izumo1:Juno | 50 | 90 |
| Izumo1:JunoH177Q | 50 | 90 |
| Izumo1:JunoH177E | 50 | 90 |

**Table S3: Distribution of Izumo1:Juno residues between different interaction clusters. Positions of bolded residues can be seen in Fig. 2.**

| Cluster | Izumo1 residues | Juno residues |
|---|---|---|
| 1 | **A158, C159, R160, K161, S162, Y163, D164**<br>**N239,** S241, **P242, A243** | **K42, Y44, E45**<br>**L58, T59, W62, H65, L66** |
| 2 | **Q65, S68, L69, E71, D72, A73, Y74, M75,** P84<br>**Y134**<br>**L146,** W148**, K150, N151, E155, H157** | **D67**<br>**F77, L81, M83, P84, R87**<br>**M145**<br>**Q161, R165, L168, Q171, L173, P174** |
| 3 | **K34, E37, K38**<br>**K48, H49, K51, M54, E55, R56, N59, K62, N63**<br>**D105, K109, F112, W113, H116, K119** | **Q133**<br>**W140, E141, R144, Y147**<br>**H177**<br>**T181** |

**Table S4: Kinetic parameters for unbinding pathway switching determined using Monte Carlo simulation of non-reversible transitions.**

| Protein complex | Pathway | BE model | |
|---|---|---|---|
| | | $k_0$ [$s^{-1}$] | $\Delta x$ [nm] |
| Izumo1:Juno | 0 → 1 | 0.0002 | 5.28 |
| | 1 → 2 | 0.0008 | 3.05 |

Table S5: Kinetic parameters for unbinding pathway switching determined using Monte Carlo simulation of reversible transitions.

| **Protein complex** | **Pathway** | **BE model** | |
|---|---|---|---|
| | | **$k_0$ [s$^{-1}$]** | **$\Delta x$ [nm]** |
| Izumo1:Juno | 0 → 1 | 0.0008 | 3.46 |
| | 1 → 0 | 41.0662 | 0.28 |
| | 1 → 2 | 0.8874 | 1.34 |
| | 2 → 1 | 129.2846 | 0.03 |

#### Supplementary Movies Captions

**Movie 1: Exemplary trajectory of Izumo1:Juno unbinding along the two-state pathway in Steered Molecular Dynamics simulations of constant velocity at a pulling speed of 0.001 Å·ps$^{-1}$.** Contact area between proteins is marked by van der Waals representation. Spheres mark the harmonically restrained residue of Juno (S228) and the pulling residue of Izumo1 (P245). Simulation time shown is 90 ns. Water and ions were removed for clarity. Rendered in VMD (*64*, *65*).

**Movie 2: Exemplary trajectory Izumo1:Juno unbinding along the three-state pathway Steered Molecular Dynamics simulations of constant velocity at a pulling speed of 0.001 Å· ps$^{-1}$.** Contact area between proteins is marked by van der Waals representation. Spheres mark the harmonically restrained anchored residue of Juno (S228) and the pulling residue of Izumo1 (P245). Simulation time shown is 90 ns. Water and ions were removed for clarity. Rendered in VMD(*64*, *65*).

**Movie 3: Exemplary trajectory Izumo1:Juno unbinding along the subtype of the three-state Steered Molecular Dynamics simulations of constant velocity at a pulling speed of 0.001 Å·ps$^{-1}$.** Contact area between proteins is marked by van der Waals representation. Spheres mark the harmonically restrained anchored residue of Juno (S228) and the pulling residue of Izumo1 (P245). Simulation time shown is 90 ns. Water and ions were removed for clarity. Rendered in VMD (*64*, *65*).

**Movie 4: Exemplary trajectory of the Izumo1:Juno unbinding where the reorganization of the fusion-related residues on the Izumo1 helical bundle is observed.** Contact area between proteins is marked by van der Waals representation. Spheres mark the anchored residue of Juno (S228) and the pulling residue of Izumo1 (P245). Red residues mark W113, W88 and S28, fusion-essential residues identified in (*11*) . Simulation time is 90 ns. Water and ions were removed for clarity. Rendered in VMD (*64*, *65*).

**Movie 5: Exemplary trajectory Izumo1:Juno unbinding of the Izumo1:JunoH177Q as observed in Steered Molecular Dynamics simulations of constant velocity at a pulling speed of 0.001 Å·ps$^{-1}$.** JunoH177 often creates the last observed contact in the slowed pathways and without it the new stable interaction interface cannot be fully formed. Contact area between proteins is marked by van der Waals representation. Spheres mark the anchored residue of Juno (S228) and the pulling residue of Izumo1 (P245). Simulation time is 90 ns. Water and ions were removed for clarity. Rendered in VMD (*64*, *65*).

#### Supplementary PDB Files

**File F1: PDB coordinate file of the intermediate unbinding structure of the Izumo1:Juno complex extracted from the two-state pathway**. This orientation represents the suggested state $I_1$ **(Fig. 3A)**.

**File F2: PDB coordinate file of the intermediate unbinding structure of the Izumo1:Juno complex extracted from the three-state pathway**. This orientation represents the suggested state $I_2$ **(Fig. 3A).**

**File F3: PDB coordinate file of the intermediate unbinding structure of the Izumo1:Juno complex extracted from the subset of the three-state pathway**. This orientation represents the variation of the suggested state $I_2$ **(Fig. 3A** and **fig. S12).**
